# The kinetics and basal levels of the transcriptome response during Effector-Triggered Immunity in Arabidopsis are mainly controlled by four immune signaling sectors

**DOI:** 10.1101/2023.05.10.540266

**Authors:** Rachel A. Hillmer, Daisuke Igarashi, Thomas Stoddard, You Lu, Xiaotong Liu, Kenichi Tsuda, Fumiaki Katagiri

**Affiliations:** Department of Plant and Microbial Biology, University of Minnesota - Twin Cities, St Paul, MN, 55108; Ajinomoto Co Inc, Tokyo, Japan; Bioinformatics and Computational Biology Graduate Program, University of Minnesota - Twin Cities, Minneapolis, MN, 55455; State Key Laboratory of Agricultural Microbiology, Hubei Hongshan Laboratory, Hubei Key Lab of Plant Pathology, College of Plant Science and Technology, Huazhong Agricultural University, Wuhan 430070, China

**Keywords:** Plant immunity, effector-triggered immunity, transcriptome, network reconstitution, time-series analysis, gamma distribution model, averaging model, AvrRpt2, Arabidopsis

## Abstract

To observe the transcriptome response during Effector-Triggered Immunity (ETI) without complications from any other pathogen factors or heterogeneously responding cell populations, we transgenically and conditionally expressed the *Pseudomonas syringae* effector AvrRpt2 in Arabidopsis leaves. We studied this ETI-specific, cell-autonomous transcriptome response in 16 exhaustively combinatorial genetic backgrounds for the jasmonate (JA), ethylene (ET), PAD4, and salicylate (SA) immune signaling sectors. Removal of some or all four sectors had relatively small impacts on the intensity of the overall ETI transcriptome response (1972 upregulated and 1290 downregulated genes). Yet, we found that the four signaling sectors strongly affect the kinetics of the ETI transcriptome response based on analysis of individual genes via time-course modeling and of the collective behaviors of the genes via a PCA-based method: the PAD4 sector alone and the JA;SA sector interaction (defined by the averaging model) accelerated the response, while the ET;SA sector interaction delayed it. The response acceleration by the PAD4 sector or the JA;SA sector interaction was consistent with their positive contributions to ETI measured by pathogen growth inhibition. The responsive genes overlapping between ETI and Pattern-Triggered Immunity (PTI) had distinct regulatory trends regarding the four sectors, indicating different regulatory circuits in upstream parts of ETI and PTI signaling. The basal mRNA levels of most ETI-upregulated genes, but not downregulated genes, were predominantly positively regulated by the PAD4;SA sector interaction. This detailed mechanistic decomposition of the roles of four signaling sectors allowed us to propose a potential regulatory network involved in ETI signaling.

## INTRODUCTION

Regulatory mechanisms of plant immunity during interactions with pathogens are complex. This is because plant immune systems must have a high level of resilience to withstand assaults from fast-evolving pathogens (Katagiri, 2018). A common approach to a complex problem is reductionism: reducing a complex problem to a combination of a small number of simpler problems. In plant immunity, a common reduced conceptual framework is a sequence of Pattern-Triggered Immunity (PTI), Effector-Triggered Susceptibility (ETS), and Effector-Triggered Immunity (ETI) (Dodds & Rathjen, 2010; Jones & Dangl, 2006). PTI is elicited when pattern recognition receptors (PRRs) on the plant cell membrane recognize the cognate molecular patterns, which are conserved among phylogenetically similar microbes or are plant molecules associated with damage. For example, a part of bacterial flagellin, flg22, is recognized as a microbe-associated molecular pattern (MAMP) by the Arabidopsis PRR FLS2 (Gómez-Gómez *et al*, 1999; Zipfel *et al*, 2004). Pathogens well-adapted to the host plant deliver effector molecules into the plant cell and compromise PTI signaling, which is the state of ETS. Plants have intracellular ETI receptors called resistance (R) proteins that sometimes could directly or indirectly recognize the cognate effectors delivered. For example, the *Pseudomonas syringae* effector AvrRpt2 is recognized by the Arabidopsis ETI receptor RPS2 (Bent *et al*, 1994; Mindrinos *et al*, 1994). This recognition leads to elicitation of ETI.

To apply reductionism to ETI according to this framework, the state of ETI needs to be studied in isolation from PTI and ETS. If an ETI-eliciting effector is delivered from a pathogen, the pathogen necessarily presents MAMPs, and consequently PTI is elicited as well. Since ETI and PTI interact in a highly context dependent manner (Hatsugai *et al*, 2017; Ngou *et al*, 2021; Wang *et al*, 2023; Yuan *et al*, 2021), subtraction of the quantitative measure of plant immunity with a pathogen strain (a mixture of PTI and ETS states) from that with the congenic pathogen strain carrying an ETI-eliciting effector (a mixture of PTI, ETS, and ETI states) does not provide a measure of the ETI-only state. Another complexity for studying plant immunity is heterogenous states among plant cells. During ETI, the plant cells that directly recognize an ETI-eliciting effector typically undergo the hypersensitive response (HR), which is a cell-autonomous programmed cell death (Coll *et al*, 2011). The surrounding plant cells activate a later immune response based on some secondary signal(s) from the autonomously responding plant cells and/or diffusible pathogen factors (Liu *et al*, 2022; Lu & Tsuda, 2021). Spatially separating the responses of these two cell populations experimentally via single cell RNA-seq with good time resolution and adequate replication is technically and financially challenging. Artificial expression of an ETI-eliciting effector in plant cells circumvents these issues (Hatsugai *et al.*, 2017; Ngou *et al.*, 2021; Yuan *et al.*, 2021): the ETI-only, cell-autonomous-response-only state can be achieved. In a whole-plant application, use of a chemical-inducible promoter controlling expression of an effector is a practical method, e.g., (McNellis *et al*, 1998; Tsuda *et al*, 2012).

Another area in which reductionism can be applied to the study of ETI is dissection of the ETI signaling network. Signaling machineries involved in the network interact with each other, making the behavior of the signaling system complex: particularly, backup mechanisms (buffering interactions) in the network provide resilience against disabling attack on some of the machineries (Hillmer *et al*, 2017; Katagiri, 2018; Tsuda *et al*, 2009). We have reduced the signaling network to a network of five major immune signaling sectors, the jasmonate (JA), ethylene (ET), PAD4, and salicylate (SA) sectors and the ETI-Mediating, PTI-Inhibited sector (EMPIS), in Arabidopsis (Hatsugai *et al.*, 2017; Hillmer *et al.*, 2017; Tsuda *et al.*, 2009). Disabling all five sectors mostly abolished ETI elicited by AvrRpt2 (AvrRpt2-ETI). ETI can be largely restored from the ETI-abolished state in three independent ways (i.e., through three backup mechanisms): (i) restoring EMPIS only; (ii) restoring the PAD4 sector only; or (iii) restoring both the JA and SA sectors (Hatsugai *et al.*, 2017; Katagiri, 2022; Tsuda *et al.*, 2009). Thus, these three machineries were largely sufficient for ETI signaling in the five-sector context.

An important step in reductionism is to synthesize understanding of the original complex problem by combining what we learned from several simpler problems. However, when no generic rules about how to combine them are known, it is necessary to measure what happens when they are combined in every combination, i.e., study their interactions in an exhaustive manner. Regarding four signaling sectors, the JA, ET, PAD4, and SA sectors, we have been using 16 exhaustively combinatorial genotypes among the hub genes of the four sectors to estimate all possible interactions among them (Hillmer *et al.*, 2017; Tsuda *et al.*, 2009). This essentially reconstitutes the intact network sector by sector, so we call this analytical approach network reconstitution (formerly, signaling allocation) (Katagiri, 2017). We improved a general linear model describing network reconstitution so that it allows an intuitive and consistent interpretation of higher order interactions among sectors (Katagiri, 2022). The new model is called the averaging model, and we call the analysis Network Reconstitution via Averaging Model (NRAM). The definition of interactions in the averaging model is different from that in an additive model, so a “;” is used to indicate an averaging model interaction, e.g., JA;SA interaction.

Here, we collected Arabidopsis RNA-seq data in response to Estradiol (Ed) -inducible *in planta* expression of AvrRpt2 in 16 exhaustively combinatorial genetic backgrounds for the four major immune signaling sectors, the JA, ET, PAD4, and SA sectors. We applied NRAM with the four sectors to the cell-autonomous ETI transcriptome response. The transcriptome response was characterized by the kinetic parameters of the peak amplitude and time of the response time-course for each of 1972 ETI-upregulated and 1290 ETI-downregulated genes. The four sectors did not strongly affect the overall response intensity, consistent with the fact that EMPIS (Hatsugai *et al.*, 2017), which could be inhibited by PTI, was not disabled. However, the PAD4 sector alone or the JA;SA sector interaction, which positively contribute to AvrRpt2-ETI, accelerated the responses of both upregulated and downregulated genes. This observation suggests the importance of a fast response during AvrRpt2-ETI for strong immunity. In contrast, the ET;SA sector interaction delayed these responses. We applied similar analyses to a previously published flg22-PTI transcriptome response data set (Hillmer *et al.*, 2017). Although the responsive genes were highly overlapping between AvrRpt2-ETI and flg22-PTI, regulation of the transcriptome response via the four signaling sectors was distinct between AvrRpt2-ETI and flg22-PTI. We also found that the basal mRNA levels of ETI-upregulated, but not ETI-downregulated, genes were predominantly and positively regulated by the PAD4;SA sector interaction. We propose a regulatory network model that can explain our observations regarding ETI transcriptome response regulation across the four signaling sectors.

## RESULTS

### Genotype nomenclatures

For simplicity, we use the following genotype nomenclature. The names of transgenes for Ed-induced expression are preceded by *Ed*-, such as *Ed-AvrRpt2*. All genotypes carried the *Ed-AvrRpt2* transgene, except the *GUS* genotype carrying *Ed-GUS* in a wild-type genetic background. The wild-type alleles regarding of the hub genes of the four signaling sectors, the JA, ET, PAD4, and SA sectors, are shown by single letters, *J, E, P*, and *S*, respectively. The corresponding mutant alleles are shown by the letter *x* for the corresponding position. For example, *JEPS* represents the wild type alleles for all four genes, and *JxPx* represents the wild-type alleles for the hub genes of the JA and PAD4 sectors and mutant alleles for those of the ET and SA sectors. This genotype nomenclature for the four sectors, using “*x*”, makes it clear which sectors remain functional in a particular genotype, which is important in mechanistic interpretations of the sectors (Katagiri, 2022; Tsuda *et al.*, 2009). The *r2r1* genotype carried mutant alleles of the *R* genes, *rps2* and *rpm1*, while its four sector hub genes had wild-type alleles.

### The transcriptome data set

We used 18 Arabidopsis genotypes, *JEPS, xEPS, JxPS, JExS, JEPx, xxPS, xExS, xEPx, JxxS, JxPx, JExx, xxxS, xxPx, xExx, Jxxx, xxxx, r2r1*, and *GUS*. For each genotype, leaf tissue was collected at 0, 2, and 5 hours post treatment (hpt) of mock and 0, 1, 2, 3, 4, 5, and 6 hpt of 50 µM Ed. The 6 hpt of Ed was chosen as the last time point since macroscopic HR-associated turgor pressure loss was evident by 9 hpt in *JEPS*. Three biological replicates were made for each of 18 genotypes * (3 time points with mock + 7 time points with Ed) * 3 biological replicates = 540 RNA-tag-seq libraries.

### Overview of the RNA-seq data

The RNA-seq data for each genotype * treatment * time combination were consistent across biological replicates by visual inspection, indicating high reproducibility of the data (Figure S1). The mean estimates for four genotypes, *JEPS*, *xxxx*, *r2r1*, and *GUS* across the treatment and the time for 18970 well-expressed genes were subjected to PCA. The results are shown in a PC1 x PC2 plot (Figure 1A). While the PC1 values of *JEPS* and *xxxx* with Ed showed large variation in a time-dependent manner, the PC1 values among the negative controls (*r2r1*, and *GUS* with Ed and all mocks) showed only small variation. Therefore, PC1, which captures 45.1% of the data variance, mostly represents the ETI-specific transcriptome response. On the other hand, PC2, which represents 17.3% of the variance, mainly captures circadian/diurnal responses (Yang *et al*, 2020) in the negative controls (Text S1 for details). Although the ETI transcriptome response in *xxxx* was very similar to the response in *JEPS* as the trajectories along their time points were very close in the PC1 x PC2 space, the *xxxx* response was generally slower than the *JEPS* response (compare “E3” to “E6” between the genotypes). These overview observations regarding the ETI-response in *JEPS* and *xxxx* are in agreement with previous observations when AvrRpt2 was delivered from *P. syringae* (Mine *et al*, 2018).

**Figure 1.**
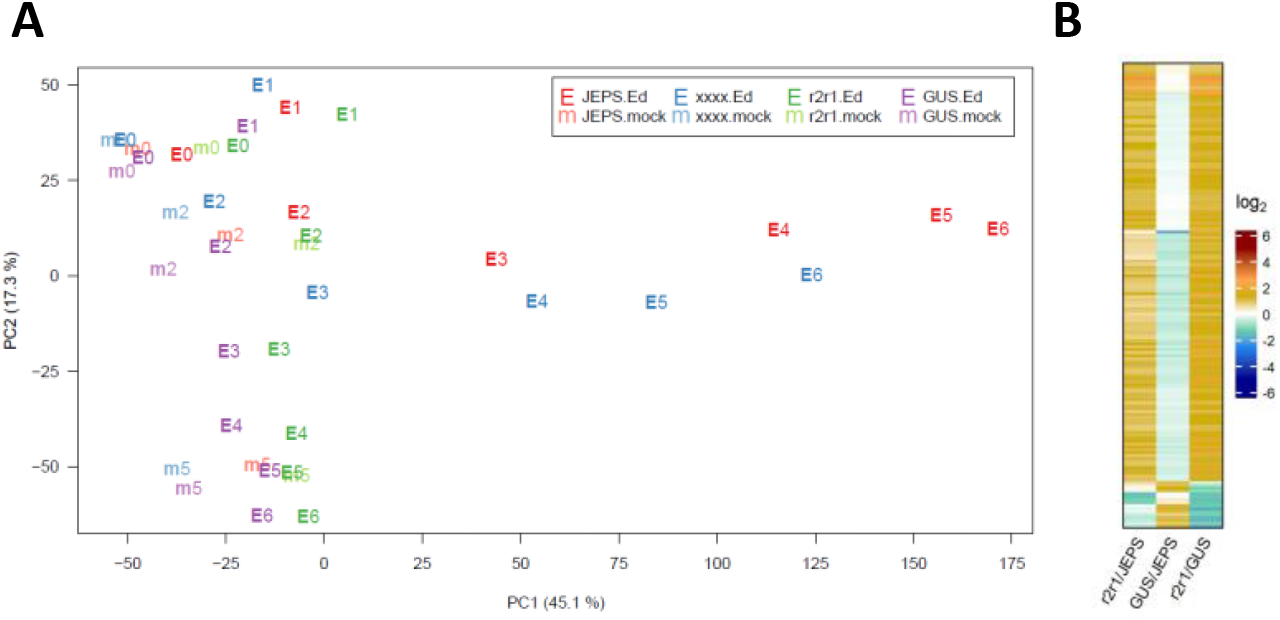
PC1 and PC2 capture characteristic biological responses in the transcriptome data. A. PC1 and PC2 values resulting from PCA of the transcriptome level values for *JEPS*, *xxxx*, *r2r1*, and *GUS* lines (point colors, red, blue, green, and purple, respectively) at 0, 2, and 5 hpt of mock (point characters, “m” plus hpt values, such as “m2”, with fainter colors) and 0, 1, 2, 3, 4, 5, 6 hpt of Ed (point characters, “E” plus hpt values, such as “E4”, with solid colors). B. The mRNA levels of 158 genes that have significant differences in at least one of the pairwise comparisons among *JEPS*, *r2r1*, and *GUS* at 0 hpt (of both mock and Ed) are shown in a heatmap. All the mRNA level-related values are on a log_2_ scale in this and the subsequent figures. The color scale bar is shown at the right in B. This scale is used for all the mRNA level ratio-related values in the main figures with heatmaps for easy comparisons across the figures.

### *AvrRpt2* mRNA level variation did not cause the ETI transcriptome response variation across the combinatorial genotypes

Since AvrRpt2 is the signal that initiates an ETI response in our experimental system, if AvrRpt2 expression was significantly leaky at 0 hpt or affected by the combinatorial genotypes, it could complicate interpretations of the results. Thus, before further analyzing the ETI transcriptome response across the combinatorial genotypes, we examined these possibilities.

First, uninduced, leaky expression of AvrRpt2 was investigated because leaky AvrRpt2 expression in the *Ed-AvrRpt2*-carrying lines could elicit a constitutive, weak ETI response. Although our RNA-seq read depth per library was generally limited (1.2 to 8.4 million aligned reads per library; median 3.4 million), 96 out of 102 libraries for the genotypes carrying *Ed-AvrRpt2* at 0 hpt had no read counts for *AvrRpt2*, indicating very low leaky expression of *AvrRpt2* in general. Then, we compared mRNA level differences gene by gene between every pair of *JEPS*, *r2r1*, and *GUS* genotypes at 0 hpt. Figure 1B shows a heatmap for three pairwise comparisons with 158 genes that showed significant differences in at least one of the pairwise comparisons. If leaky AvrRpt2 expression caused a weak ETI response, the genes for a weak ETI response should show little difference in *r2r1*/*GUS* and significant and similar differences in *r2r1*/*JEPS* and *GUS*/*JEPS* since a weak ETI response would be expected only in *JEPS*. There was no gene that fit this profile of a potential weak ETI response. All genotypes except *GUS* shared the same *Ed-AvrRpt2* transgene locus introduced by genetic crosses from a single transgenic line (see Methods). Thus, we conclude that there was no appreciable effect of leaky AvrRpt2 expression in our lines.

Second, we investigated induced *AvrRpt2* mRNA levels across the combinatorial genotypes. The induced *AvrRpt2* mRNA levels (the log_2_(Ed/mock) mean estimates) indeed varied substantially across the genotypes (Figure 2A). However, the induced *AvrRpt2* mRNA level was generally strongly anticorrelated with those of ETI-responsive genes across the genotypes (Figure 2B), indicating that variation in the *AvrRpt2* mRNA level was not a major cause of the ETI transcriptome response variation we observed across the genotypes.

**Figure 2.**
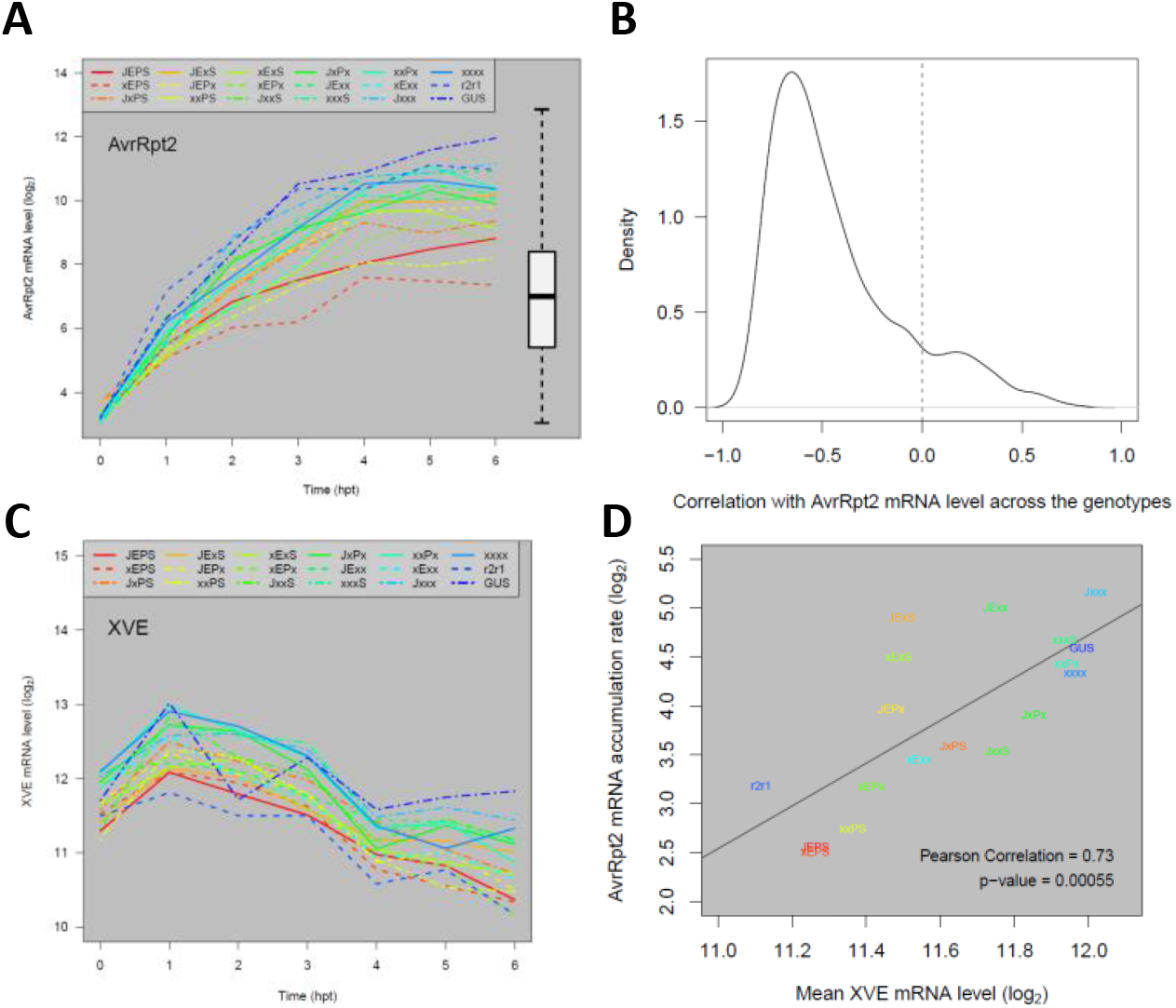
The major trends of how the four signaling sectors affect the ETI transcriptome response are not results of the *AvrRpt2* mRNA level affected by the four sectors. A. The mRNA level time courses of *AvrRpt2* in all the Arabidopsis genotypes. For the *GUS* line, the *GUS* mRNA level is shown instead. The box-and-whiskers on the right shows the distribution of the mRNA levels of the entire transcriptome for comparison. B. The distribution of the Pearson correlation coefficient for the mRNA levels across the 16 combinatorial genotypes between each of 1972 ETI-upregulated genes and *AvrRpt2*: the time point at which the mRNA level is highest in *JEPS* was chosen as a representative time point for each gene for calculation of the correlation. The correlation zero is indicated by a dashed vertical line. C. The mRNA level time courses of *XVE* in all genotypes. D. A significantly positive correlation between the mean XVE mRNA levels and the accumulation rate ratios from 1 hpt to 4 hpt of the *AvrRpt2* mRNA across the genotypes. The regression line, the correlation coefficient, and the significance of the correlation are also shown. The color codes used to indicate the genotypes in A, C, and D are the same.

### The four signaling sectors mainly affect the response kinetics

Figure 3A shows a heatmap of the log_2_(Ed/mock) mean estimates for 16 combinatorial genotypes at 7 time points (0, 1, 2, 3, 4, 5, and 6 hpt). It contains 1972 ETI-upregulated and 1290 downregulated genes, which were well-modeled for the time course and ordered according to the model-fit peak times in *JEPS* (see below for the time-course model). The combinatorial genotypes from *JEPS* to *xxxx* did not show large differences in the log_2_(Ed/mock) time courses across the genes. This was not surprising because EMPIS can mediate relatively intact ETI when it is not inhibited by PTI signaling (i.e., the four signaling sectors are not absolutely required for an ETI response when EMPIS is functional) (Hatsugai *et al.*, 2017). However, when the log_2_(Ed/mock) value for *xxxx* were subtracted from those for *JEPS* at each time point, the peaks of the differences were generally earlier than the peaks of the values in *xxxx* (Figure 3B), indicating that the peak times were earlier in *JEPS* than *xxxx* for most of the upregulated and downregulated genes.

**Figure 3.**
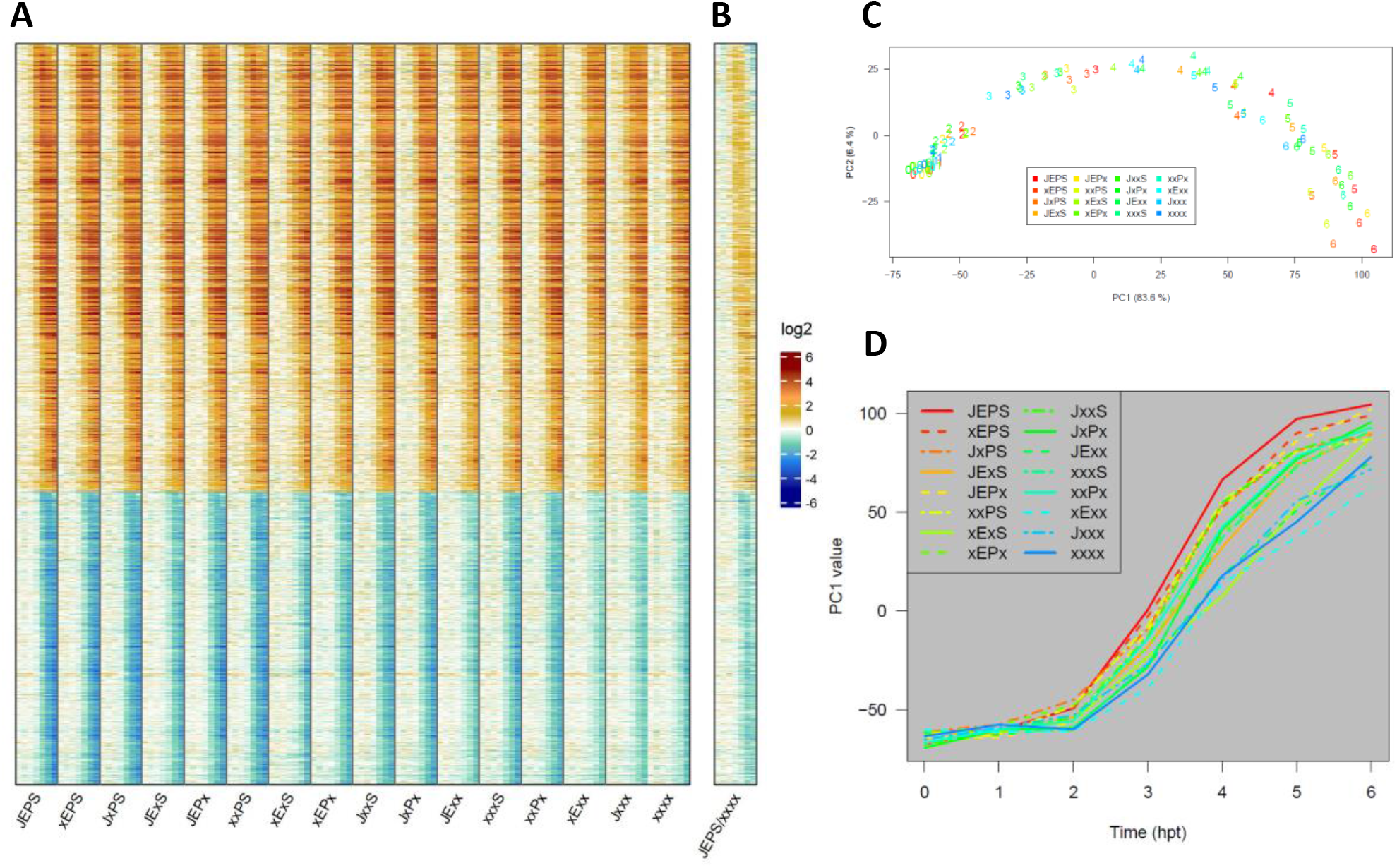
The fold-change of the transcriptome response during ETI is not strongly affected by the four signaling sectors. A. A heat map showing the mRNA level fold-change compared to 0 hpt in 1972 upregulated and 1290 downregulated genes during ETI in 16 combinatorial genotypes regarding the four signaling sectors. For each genotype, the fold-change values for 0, 1, 2, 3, 4, 5, and 6 hpt with 50 µM estradiol are shown in this order. B. The ratio of the mRNA level fold-changes between the genotypes *JEPS* (WT) and *xxxx* (quadruple mutant) at each time point is shown for the same gene set and order as A. All the mRNA level ratio values are in the log_2_ scale. The color scale bar is shown at the right of B. C. PC1 and PC2 values resulting from PCA of the ETI-responsive transcriptomes shown in A. The point character numbers show the hpt. D. The time course of PC1 (ETI-PC1) values in C are shown for each combinatorial genotype background.

Figure 3C shows the PC1 x PC2 plot of genotype * time data resulting from PCA of the values used in Figure 3A. PC1 captured most of the ETI response across the genotypes (83.6% variance). In the PC1 x PC2 plot (together, 90% variance), the response trajectories of the time points for the genotypes were all very similar, monotonously increasing along PC1, and differed in how fast the PC1 value increased. This observation confirms that the ETI-transcriptome response was similar across the genotypes and that the major difference across the genotypes was response kinetics. Subsequently, the PC1 value in Figure 3C is used as the representative value for the ETI-response of each genotype (designated the “ETI-PC1” value). Figure 3D shows the time courses of ETI-PC1. Roughly speaking, the more sectors that were functional, the faster the ETI response was. Note that the time courses of fast-responding genotypes, such as *JEPS*, show signs of saturating responses in this time range, while slow-responding genotypes, such as *xxxx*, do not.

### The PAD4 sector effect and the JA;SA sector interaction accelerated the peak time while the ET;SA sector interaction delayed it

To characterize the dynamics of the ETI transcriptome response gene by gene, we fit a gamma probability density function (gamma-pdf) time course model to the response of each gene in each genotype (Figure 4A for a schematic of the model; Figures S2 and S3 show the model fit by heatmaps and time courses, respectively; Table S1 shows the fitted parameter values and the fitted log_2_(Ed/mock) values at each time point). The model parameters were the peak amplitude (positive and negative for upregulated and downregulated), the peak time, the peak shape, and the time lag. The log_2_(Ed/mock) time courses were well-modeled for 1972 upregulated and 1290 downregulated genes. However, since the peak characteristics of the models with peak times well after the last data time point (6 hpt) are not reliable, 866 upregulated and 234 downregulated genes with estimated peak times < 7.5 hpt in at least 13 out of 16 genotypes were selected for NRAM analysis of the peak amplitude and time (Figures 4B and 4C). The left-most lane in the NRAM heatmap for the peak amplitude shows the log_2_-ratio of the peak amplitudes between *JEPS* and *xxxx* for each gene (“PA,JEPS/xxxx” in Figure 4B). The general trend was that the absolute value of the peak amplitude was larger in *JEPS* than *xxxx*: log_2_(*JEPS*/*xxxx*) medians 0.51 and -0.74 for the upregulated and downregulated genes, respectively. In the case of the peak time differences between *JEPS* and *xxxx* (“PT,JEPS-xxxx” in Figure 4C), in agreement with the observations in “JEPS/xxxx” in Figure 3B (the same lanes are also shown in Figure 4B as a reference), the general trend was delays in the peak time in *xxxx* compared to *JEPS*: medians 0.92- and 0.77-hour delays for the upregulated and downregulated genes, respectively. These values, “PA,JEPS/xxxx” and “PT,JEPS-xxxx”, were subjected to NRAM-decomposition for the peak amplitude and time, respectively. Since the number of downregulated genes analyzed was small, it is not clear from the figures whether there are any common NRAM trends across the downregulated genes in either the peak amplitude or time. On the other hand, common NRAM trends are evident among modeled upregulated genes in both the peak amplitude and time as similar color bands dominate in some of the sector effects and interactions, e.g.: the PAD4 and SA sectors positively and the PAD4;SA sector interaction negatively affected the peak amplitude; the PAD4 sector and the JA;SA sector interactions accelerated and the ET;SA sector interactions delayed the peak time. (Here we consider up to two-sector interactions because a mechanistic interpretation of each sector influence could be complicated with interactions among three or more sectors.) These visual observations were confirmed by the means across the 866 upregulated genes for each of the sector effects and interactions (Figures 4D and 4E).

**Figure 4.**
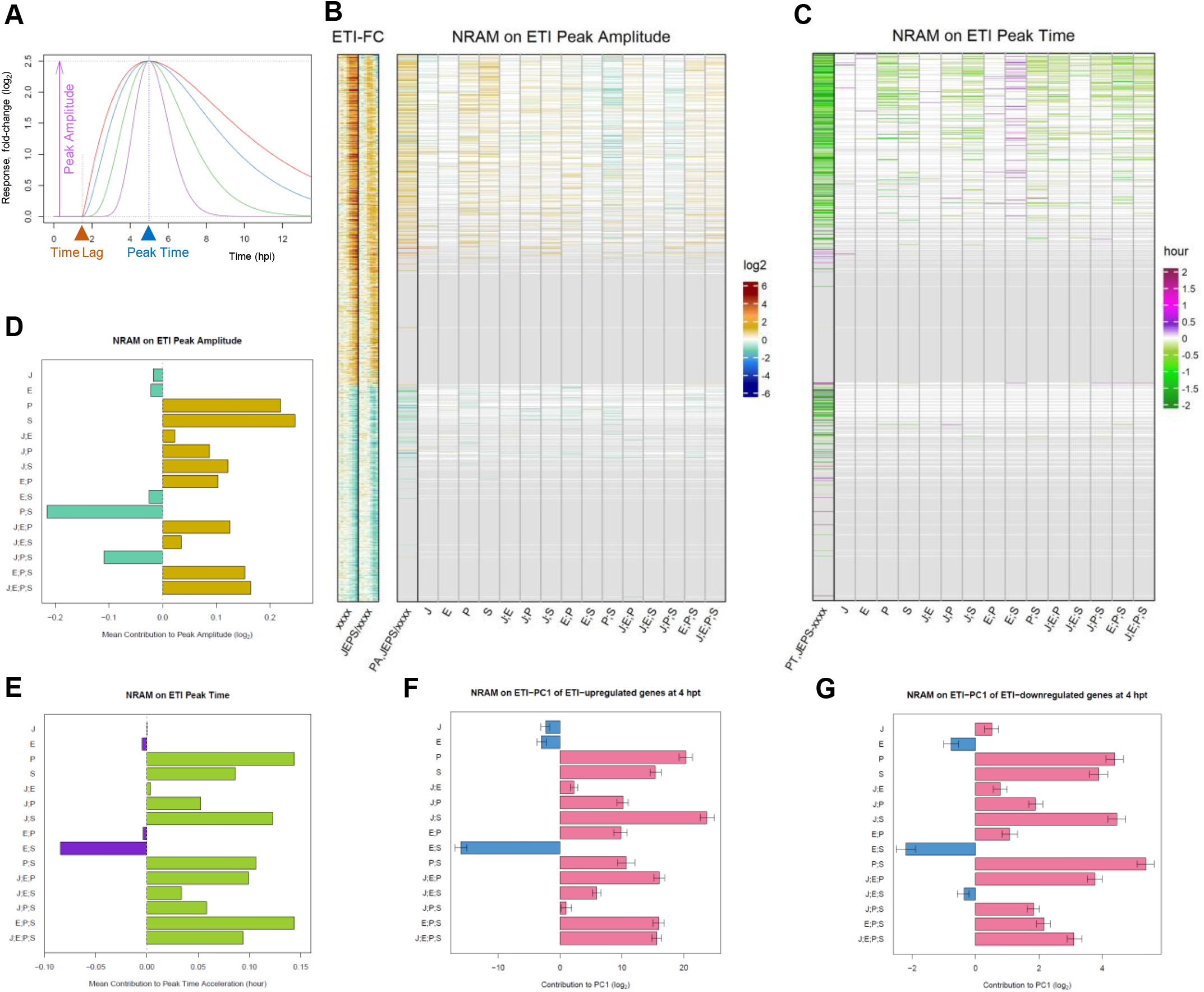
The peak times of many ETI-responsive genes are accelerated by the PAD4 sector and the JA;SA sector interaction but delayed by the ET;SA sector interaction. A. A schematic representation of the gamma-pdf time course model. Single-peak time course curves in different colors (red, blue, green, and purple) are time course models with different shape parameter values. In addition to the shape parameter, the model is parameterized by the peak amplitude (purple arrow), the time lag (brow arrowhead), and the peak time (blue arrowhead) parameters. The values for the latter three parameters used for these example time course models are 2.5, 1.5 hpt, and 5 hpt, respectively. The value for the baseline parameter, *B*, is considered subtracted, and therefore its value is zero and not shown. B. NRAM analysis regarding the four sectors was applied to the peak amplitudes estimated by gamma-pdf time course models for each gene during ETI. The left column, indicated as “PA,JEPS/xxxx”, shows the ratio of the peak amplitudes between *JEPS* and *xxxx*. The set and order of the genes are the same as those in Figure 3A (1972 ETI-upregulated and 1290 ETI-downregulated genes), and the *xxxx* fold-change values and the *JEPS*/*xxxx* fold-change ratios from Figures 3A and 3B are included on the left (“ETI-FC”) as references. The color scale bar on a log_2_ scale shown on the right is the same as that in Figure 3B. C. NRAM analysis regarding the four sectors was applied to the peak time estimated by gamma-pdf time course models for each gene during ETI. The left column, indicated as “PT,JEPS-xxxx”, shows the difference of the peak times of *JEPS* from that of *xxxx*. Gray horizontal lines in B and C indicate genes for which the peaks were not estimated with confidence due to their late peak times: the numbers of modeled genes that were subjected to NRAM analysis (genes not grayed out) were 866 and 234 for ETI-upregulated and - downregulated genes, respectively. D. The mean value across the genes of each term in NRAM of the peak amplitudes of the 866 upregulated genes. E. The mean value across the genes of each term in NRAM of the peak times of the 866 upregulated genes. The signs of the values were flipped to show the acceleration of the peak time. F. NRAM analysis of the ETI-PC1 values at 4 hpt for ETI-upregulated genes. The mean and its 95% confidence interval (error bar) were estimated based on 500 bootstrapped sets of the ETI-upregulated genes. The projection of each bootstrapped set on the ETI-PC1 was subjected to NRAM analysis. G. NRAM analysis of the ETI-PC1 values at 4 hpt for ETI-downregulated genes. The mean and its 95% confidence interval (error bar) were estimated based on 500 bootstrapped sets of the ETI-downregulated genes. The projection of each bootstrapped set on the ETI-PC1 was subjected to NRAM analysis.

These trends observed in NRAM analysis were only for the well-modeled upregulated, early-peak genes, which are about half of the well-modeled upregulated genes, and no trends were evident for the small number of well-modeled downregulated, early-peak genes. Since the ETI-PC1 captured most of the ETI response and its timing (Figure 3C), we subjected the ETI-PC1 values at 4 hpt to NRAM analysis. The time of 4 hpt was chosen because most genes started responding by this time, and most genes had not reached their peaks by this time, so that this time point likely captures the response kinetics of the gene sets with little bias. In fact, the obtained ETI-PC1 NRAM profiles at 4 hpt for 1972 ETI-upregulated genes (Figure 4F) and 1290 downregulated genes (Figure 4G) were very similar to the peak time NRAM mean profiles of 866 upregulated genes (Figure 4E). The similar mean trends of the NRAM profiles between the upregulated and downregulated genes indicate that the peak times of the upregulated and downregulated genes were on average regulated essentially in the same manner by the four signaling sectors. Using bootstrapping, the 95% confidence intervals of the NRAM terms were estimated for the upregulated and downregulated genes. The small 95% confidence intervals relative to the mean values suggest that the observed NRAM profile trends are very common across the genes in each of the upregulated and downregulated gene sets, i.e., the peak times of most of the upregulated and downregulated genes were regulated essentially in the same manner by the four signaling sectors.

### The roles of the four sectors in regulation of PTI-responsive genes were clearly different from those in regulation of ETI-responsive genes

A similar analysis was conducted with the flg22-PTI transcriptome response (data from (Hillmer *et al.*, 2017)) for the genes in common between this response and the 3262 ETI-responsive well-modeled genes (602 ETI-upregulated and 594 downregulated genes were common; Figure 5). The gamma-pdf time course model (with no time lag parameter) was fit to the estimated log_2_(flg22/mock) data at 2, 3, 5, and 9 hpt for each of the 16 combinatorial genotypes for each gene (Figures S4 and S5 show the model fit by heatmaps and time courses, respectively; Table S2 shows the fitted parameter values and the fitted log_2_(flg22/mock) values at each time point). The data at 1 hpt were not used because the response at 1 hpt likely represents a general stress response rather than an immune-specific response (Bjornson *et al*, 2021). The data at 18 hpt were not used because we wanted to compare the responses between ETI and PTI over similar time ranges. The genes upregulated or downregulated in ETI and PTI were generally consistent across the common genes (“xxxx” in Figure 5A). The collective effect of the four sectors on the peak amplitude was limited for most genes (faint colors in “PA,JEPS/xxxx” compared to “xxxx” in Figure 5A), and the collective effect was not consistent across the upregulated or downregulated genes (reddish and blueish color bands are mixed in “PA,JEPS/xxx”). Furthermore, no consistent pattern was evident in the NRAM results for the PTI peak amplitude. On the other hand, consistent patterns were evident in the NRAM results for the PTI peak time: The ET sector accelerated and the JA;SA sector interaction delayed the PTI transcriptome response (Figure 5B), which was confirmed by the mean values of the NRAM terms (Figure 5C). When PCA was applied to the PTI transcriptome response data, PC1 appeared not to capture the timing of the response (the genotype-specific trajectories were not highly overlapping; Figure 5D). Therefore, the PC1 values cannot be used for analysis of the peak time in the PTI response. In short, control of the peak amplitude and time by the four signaling sectors is distinct between the ETI and PTI transcriptome responses.

**Figure 5.**
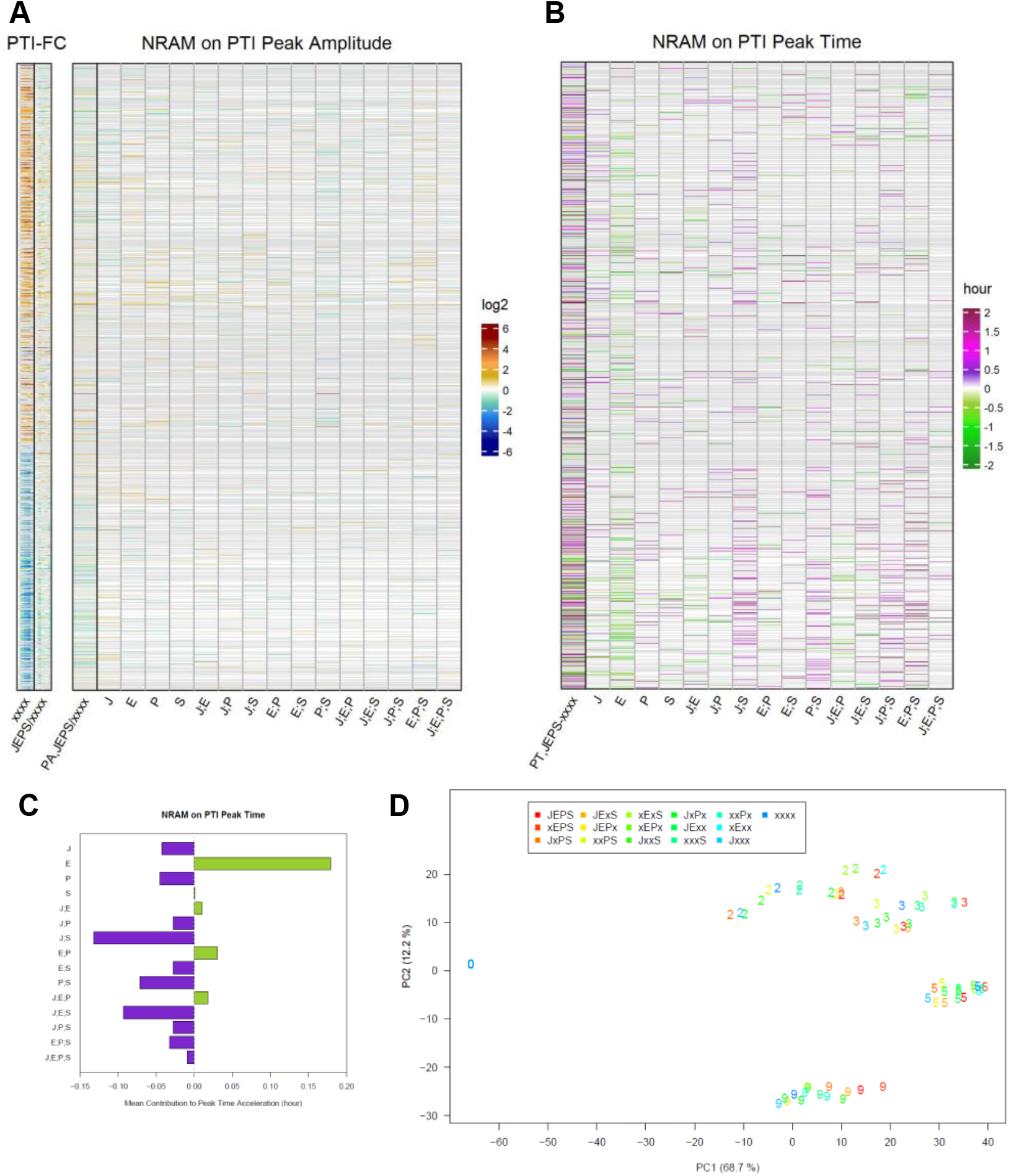
The influence of four sectors on the peak times of PTI responses is distinct from that on the peak times of ETI responses for the ETI-responsive genes. A. NRAM analysis regarding the four sectors was applied to the peak amplitudes estimated by gamma-pdf time course models for each gene during PTI elicited by flg22. The left column, indicated as “PA,JEPS/xxxx”, shows the ratio of the peak amplitudes between *JEPS* and *xxxx*. In this figure, the references on the left are also based on the PTI data (“PTI-FC”). The gene set and order are the same as those in Figure 3A (1972 ETI-upregulated and 1290 ETI-downregulated genes). The log_2_ color scale bar shown on the right is the same as that in Figure 3B. B. NRAM analysis regarding the four sectors was applied to the peak time estimated by gamma-pdf time course models for each gene during PTI. The left column, indicated as “PT,JEPS-xxxx”, shows the difference of the peak times of *JEPS* from that of *xxxx*. Gray horizontal lines in A and B indicate genes that were not significantly responsive in PTI or that were not well modeled by the gamma-pdf time course model (602 upregulated and 594 downregulated genes are not grayed out in the NRAM heatmaps in A and B). C. The mean value across the genes of each term in NRAM on the peak time in B. The signs of the values were flipped to show the acceleration of the peak time. E. PC1 and PC2 values resulting from PCA of the PTI response transcriptomes. The color coding of the genotypes is the same as that in Figure 3C. The point character numbers, 0, 2, 3, 5, and 9, indicate the hpt. The data for 1 and 18 hpt were excluded from the analysis. The values for 0 hpt are the same for all genotypes and consequently only the point with the *xxxx* color is visible in the figure.

### The basal mRNA levels of most ETI-upregulated genes were largely positively controlled by the PAD4;SA sector interaction

We also investigated the influence of the four signaling sectors on the uninduced, basal mRNA levels of the ETI-responsive genes (Figure 6). The basal mRNA levels of most ETI-upregulated genes but not the downregulated genes were positively regulated by the PAD4;SA sector interaction (Figure 6A). Thus, the basal levels of the ETI-upregulated genes are likely positively regulated by the PAD4;SA sector interactions. We examined the converse of this observation, whether the genes whose basal mRNA levels are positively regulated by the PAD4;SA sector interactions are likely to be ETI-upregulated genes. When the 4562 genes whose basal mRNA levels were affected in some way by the four signaling sectors were subjected to NRAM, the genes with positive PAD4;SA sector interaction values were strongly associated with the ETI-upregulated genes (Figure 6B). Thus, the gene set for the ETI-upregulated genes and the set of genes whose basal mRNA levels are positively regulated by the PAD4;SA sector interactions are largely overlapping.

**Figure 6.**
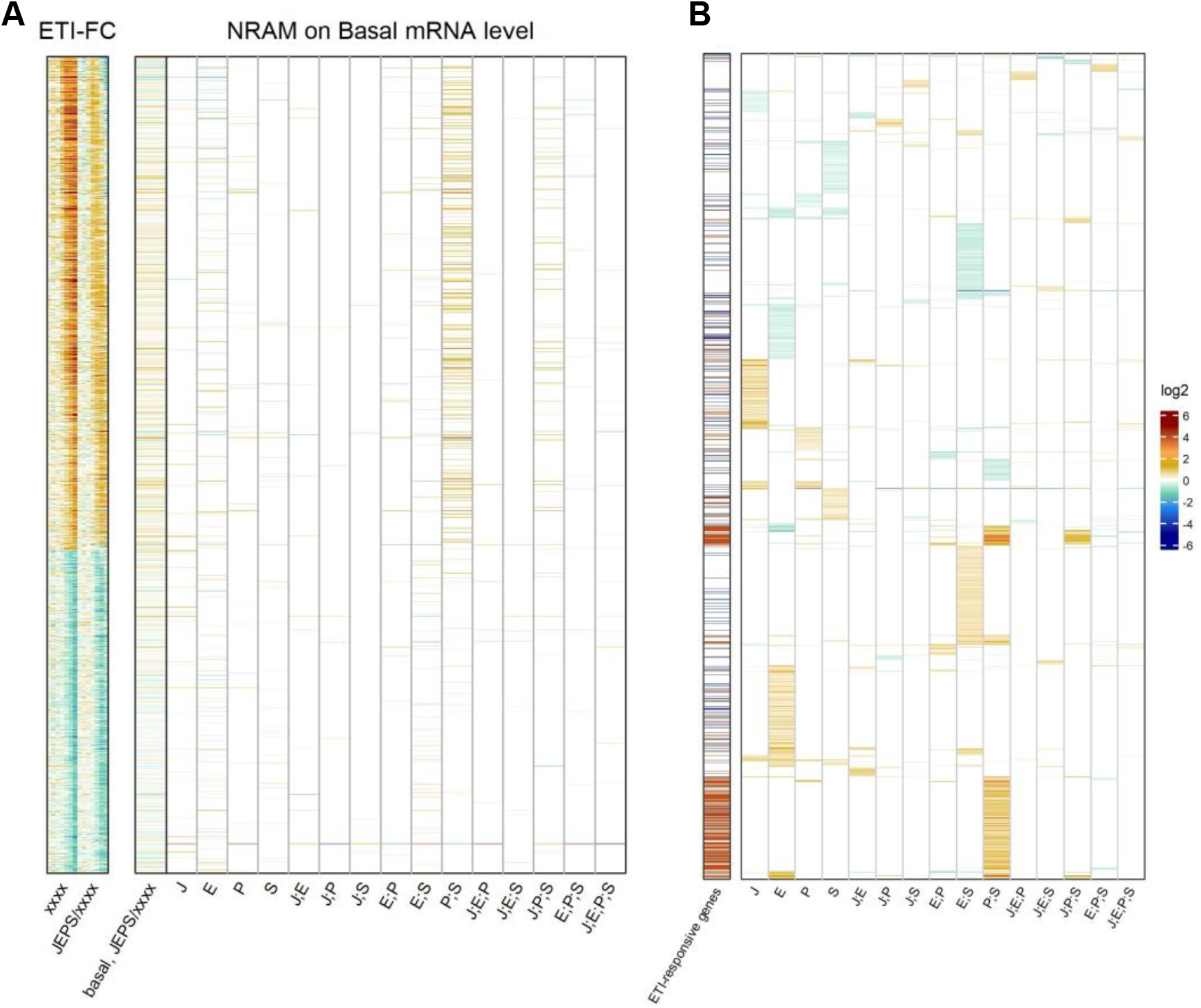
The genes whose basal mRNA level is primarily positively regulated by the PAD4;SA sector interaction are largely overlapping with ETI-upregulated genes. A. NRAM analysis regarding the four sectors was applied to the basal mRNA level of each ETI-upregulated or downregulated gene. The gene set and order are the same as those in Figures 3A. The left column, indicated as “basal,JEPS/xxxx”, shows the ratio of the basal levels between *JEPS* and *xxxx*. The references on the left, “ETI-FC”, are the same as those in Figure 4B. B. NRAM analysis results of 4562 genes that had at least one significant NRAM term when the analysis was applied to their basal mRNA levels. The genes were hierarchically clustered with the NRAM values for better visualization. The left column indicates the ETI-upregulated (red) and downregulated (blue) genes.

## DISCUSSION

In this study, we combined simplification of an ETI experimental system by *in planta* expression of the ETI-eliciting effector AvrRpt2, dynamical analysis of the ETI transcriptome response, and decomposition of the signaling sector contributions to the transcriptome response by NRAM. This multifaceted analytical design allowed us to reveal how the dynamical characteristics of the ETI-transcriptome responses are regulated through specific signaling sectors and their interactions. Similar analysis of the PTI transcriptome response indicated that dynamical characteristics of the overlapping gene set were regulated differently from those in ETI. Furthermore, the ETI upregulated genes were characterized by positive regulation of their basal mRNA levels via the PAD4;SA sector interaction.

The wild type alleles of *RPS2* and/or *RPM1* altered the transcriptome without ETI elicitation (Figure 1B). The genes downregulated by *RPS2* and/or *RPM1* (R2R1_down) overlap with ETI-upregulated genes (ETI_up; Figure 7A). This tendency is also detectable in Figure 1A, which shows that the data points for the *r2r1* genotype have slightly higher PC1 values than the other data points for no ETI elicitation in the genotypes containing the *R2R1* wild-type alleles. Since PC1 mainly represents the ETI response, the slightly higher PC1 values in *r2r1* correlate with downregulation of the ETI response by *RPS2* and/or *RPM1*. Since it has been reported that functional alleles of *RPS2* downregulate the mRNA levels of some defense-related genes (MacQueen *et al*, 2016), this observation could be explained solely by the *RPS2* effect. We only saw 12 genes significantly upregulated by *RPS2* and/or *RPM1,* and they have virtually no overlap with ETI-responsive genes (Figures 7A and 7B). The *R2R1* downregulated gene set (142 genes) was mostly included in a larger set of genes whose basal mRNA levels were positively regulated by the PAD4;SA interaction (738 genes; Figure 7C), suggesting that *RPS2* and/or *RPM1* lowers the basal mRNA levels of the *R2R1* downregulated genes by weakly interfering with the interaction between the PAD4 and SA sectors.

**Figure 7.**
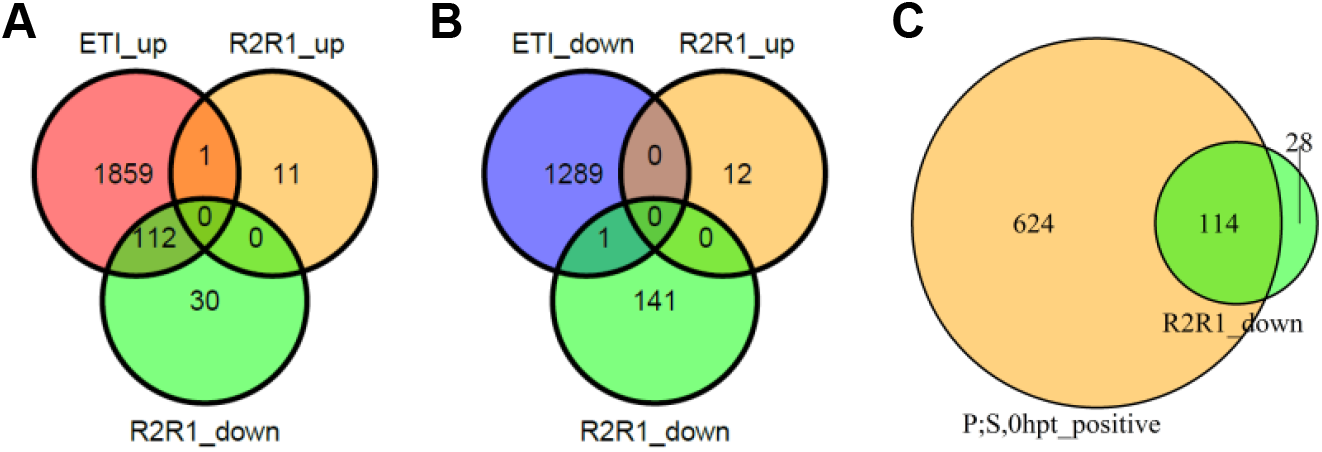
The basal mRNA levels of most genes downregulated in the lines carrying wild-type *RPS2* and *RPM1* genes are primarily positively regulated by the PAD4;SA sector interaction. A. A Venn diagram showing the number of genes overlapping among the ETI-upregulated genes (ETI_up) and *RPS2 RPM1*-upregulated (R2R1_up) and downregulated (R2R1_down) genes. B. A Venn diagram showing the number of genes overlapping among the ETI-downregulated genes (ETI_down) and *RPS2 RPM1*-upregulated (R2R1_up) and downregulated (R2R1_down) genes. C. A Venn diagram showing the number of genes overlapping between the genes whose basal mRNA level is upregulated by the PAD4;SA sector interaction (P;S,0hpt_positive) and *RPS2 RPM1*-downregulated genes (R2R1_down).

The four-sector influence on *AvrRpt2* mRNA level upregulation is completely different from those in most ETI-upregulated genes (Figure 2B). The genotype difference in the *AvrRpt2* mRNA level was already clear at 1 hpt (Figure 2A) while not much ETI transcriptome response was observed before 2 hpt (e.g., Figures 1A, 3A, and 3C). These two observations suggest that the genotype difference in the *AvrRpt2* mRNA level is not a consequence of a difference in ETI response across the genotypes. The difference in the XVE protein level (the artificial TF for the Ed-inducible promoter activation; (Zuo *et al*, 2000)) likely explains it as the mean *XVE* mRNA level (Figure 2C) positively correlates with the *AvrRpt2* mRNA accumulation rate (Figure 2D).

The ETI signaling system is built in a way that allows three signaling mechanisms, the EMPIS only, the PAD4 sector only, and the JA and SA sectors together, to back up each other – none of the three is essential, but any one of the three is sufficient (Hatsugai *et al.*, 2017; Katagiri, 2022; Tsuda *et al.*, 2009). In our experimental system, EMPIS was not disabled, and thus the influence of the four signaling sectors we observed was limited to modification of the EMPIS-mediated ETI response. Currently the only known way to disable EMPIS is to elicit PTI signaling (Hatsugai *et al.*, 2017). However, this is not compatible with our objective of studying ETI in isolation from PTI. In addition, PTI signaling interferes with inducible promoter systems (Igarashi *et al*, 2013), so it would be experimentally difficult to exclude PTI influence on *Ed-AvrRpt2* expression. If we knew how to disable EMPIS independent of PTI, such as by a genetic mutation, it would be interesting to investigate how the ETI response is restored from a completely disabled state through each of the three ways.

The peak amplitude and time in the ETI transcriptome response were regulated differently by the four signaling sectors. We estimated the four signaling sector influence on the peak amplitude and time for 866 upregulated genes with relatively early peak times (Figures 4B-E). For most of these genes, the peak amplitude was regulated by the PAD4 and SA sectors positively and by the PAD4;SA sector interaction negatively. For the same genes, the peak time was accelerated by the PAD4 sector and the JA;SA sector interaction. Using the ETI-PC1 value, we extended the observation of the four signaling sector influence on the peak time to most of the upregulated and downregulated genes. The PAD4 sector and the JA;SA sector interaction significantly and positively contribute to ETI-mediated resistance against *P. syringae* (Katagiri, 2022; Tsuda *et al.*, 2009): The influences of the two mechanisms were obvious in the case of AvrRpt2-ETI, in which EMPIS was inhibited by PTI signaling. They were also evident even when EMPIS was active in the case of AvrRpm1-ETI (Figures 7A and 7B in (Katagiri, 2022). The latter suggests that acceleration of the ETI response is important to enhance ETI-mediated resistance against *P. syringae*.

One simple signaling circuit that allows regulation of the peak amplitude and time separately is a feedforward loop with a linear combination at the signal convergence point (Figure 8A). Using a multi-compartment model (Godfrey, 1983; Kermack & McKendrick, 1927; Liu *et al.*, 2022; Prabakaran *et al*, 2021; Rescigno, 1960) for a feedforward loop, the peak time can be changed by changing the linear coefficients and the decay rate of the converging node (Figure 8B). The peak amplitude can be changed by these three model parameters and the input amplification ratio parameter before the loop. Four model parameters can be used to determine two model characteristics of the peak amplitude and time to change, and thus the feedforward model is highly flexible in determining the peak amplitude and time.

**Figure 8.**
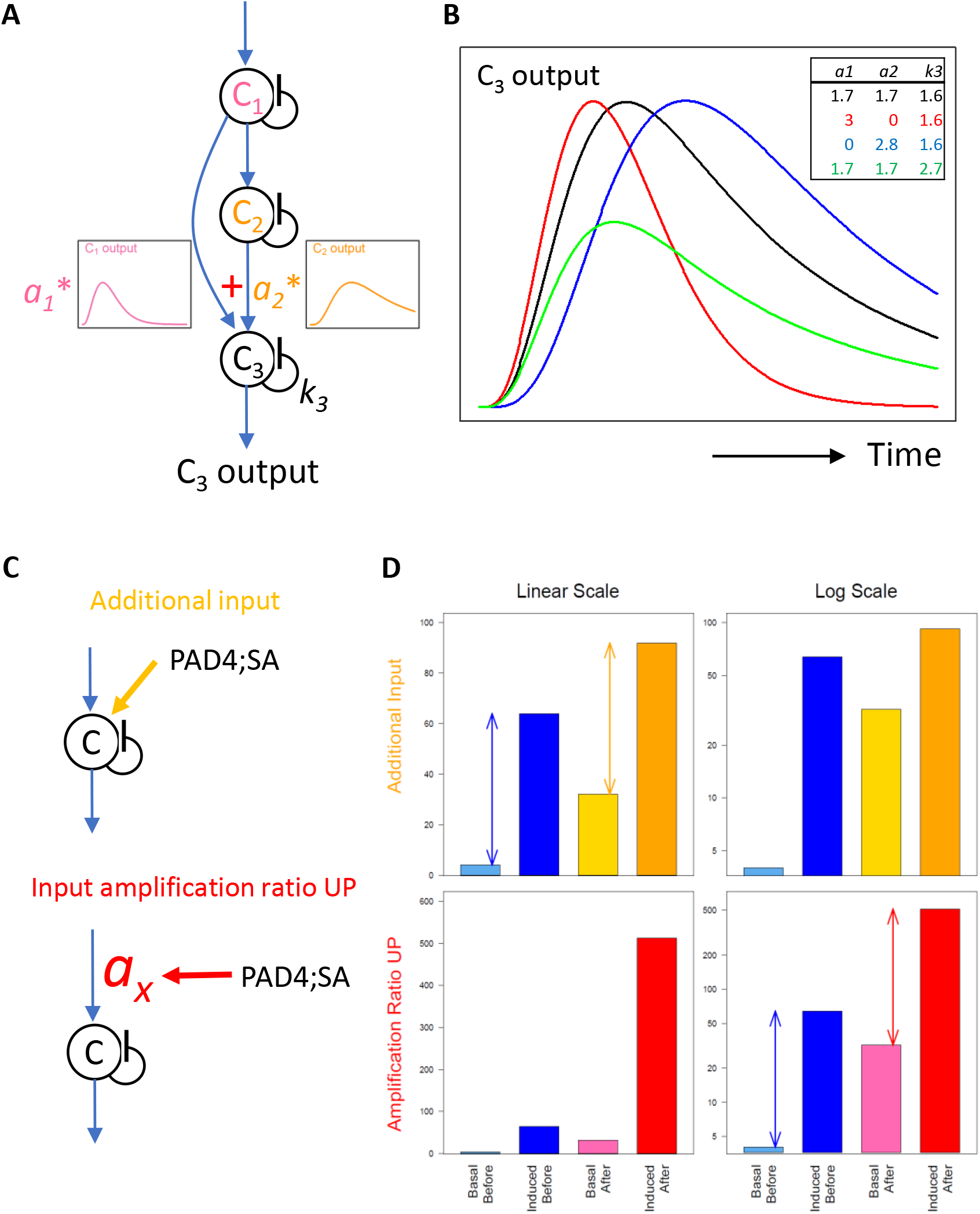
Possible network motifs for the transcriptome response regulation: a feedforward loop for the peak time and amplitude regulation (A and B); an additional input case and an input amplification ratio upregulation case for the basal level regulation (C and D). A. A schematic representation of a feedforward loop using a multicompartment model with three compartments. The input to compartment C_3_ is a linear combination of the outputs from compartments C_1_ and C_2_. The linear coefficients for the combination are *a_1_* and *a_2_*. The first-order self-decay rate for C_3_ is *k_3_*. B. Example time courses generated by a feedback loop shown in A. The parameter values corresponding to each time-course curve are indicated in the table at the top right of the plot with the corresponding color. C. In two network motifs, a motif with an additional input (top, orange) and a motif with upregulation of the input amplification ratio parameter (bottom, red), the PAD4;SA sector interactions can upregulate the basal level of the signal. Note that the input signal to compartment C shown by a blue arrow is induced by the ETI signal initiated by AvrRpt2 recognition. D. The signal outputs of the two motifs shown in C. For each of the additional input case (top) and the input amplification ratio upregulation case (bottom), the “basal” and “induced” output signal levels “before” and “after” a positive influence of the PAD4;SA sector interaction are shown on a linear scale (left) and a log scale (right). The example parameter values used are: the “before” basal and induced levels, 4 and 64 (common for both cases, blueish bars); the additional input (top), 28; the input amplification ratio (top) and that for “before” (bottom), 1; the input amplification ratio for “after” (bottom), 8. The arrows in the top left and bottom right plots indicate that the difference between the induced and the basal levels does not change on a linear scale in the additional input case and on a log scale in the input amplification ratio upregulation case, respectively.

We observed a highly common characteristic that the basal mRNA levels of ETI-upregulated genes were positively regulated by the PAD4;SA sector interactions (Figure 6). This characteristic was not shared by ETI-downregulated genes, strongly suggesting that this basal level regulation acts after signals for upregulation and downregulation are split. Such a basal level regulation could be explained by an additional non-ETI-responding input or upregulation of the input amplification ratio parameter shared among the upregulated genes (Figure 8C, top and bottom, respectively). The additional input does not change the difference between the peak level and the basal level on a linear scale (Figure 8D, top left). The input amplification ratio can multiply the basal level and the peak level equally. In the latter scenario, the difference between the induced and basal levels is unchanged on a log scale (Figure 8D, bottom right), which indicates that the fold change (induced / basal) is not changed when the input amplification ratio is changed. The data in this study does not have sufficient statistical power to test these two possible scenarios.

In defense priming situations in Arabidopsis, the mRNA levels of defense-related genes after mock or pathogen inoculation in secondary leaf tissue were compared between prior mock and a defense primer (*Pto* DC3000 or pipecolic acid) treatments of primary leaf tissue (Bernsdorff *et al*, 2016). Defense primer treatments induce the systemic acquired resistance (SAR) state in the secondary tissue. Generally, SAR induction changed the basal level but did not strongly change the fold change of pathogen-upregulated genes in the secondary tissue, which is consistent with basal level regulation by the input amplification ratio (Figures 6 and 8 in (Bernsdorff *et al.*, 2016)): defense priming substantially increases the input amplification ratio. Since the genes upregulated in different types of immunity overlap, it is likely that the basal level regulation we observed with the ETI-upregulated genes is also mediated by regulation of the input amplification ratio.

The transcriptome response regulation by the four signaling sectors was very different between ETI and PTI while the overlap of upregulated gene sets was large. We proposed a WRKY network activation hypothesis (Liu *et al.*, 2022). In this hypothesis, PTI and ETI signaling have separate upstream parts so that they can be regulated separately, and then the signals are fed into different entry points of the transcriptionally interconnected WRKY network. These inputs through different entry points would activate highly overlapping WRKY subgroups although the specific WRKY members that are activated are not the same, explaining the overlapping but not identical upregulated gene sets.

We combined two feedforward loops for ETI and PTI for both upregulated and downregulated genes, basal level regulation by the input amplification ratio for upregulated genes, and WRKY network activation for upregulated genes, to propose one possible regulatory network model for ETI and PTI transcriptome responses (Figure 9). Although it was not clear how extendable was the peak amplitude regulation we observed with ETI-upregulated genes with relatively early peak times, we tentatively assumed that this regulation is common among both upregulated and downregulated genes in this model. One ambiguity we left in the model is whether the input amplification ratio-based basal level regulation is part of the WRKY network or upstream of the WRKY network. Since many genes whose basal mRNA levels are positively regulated by the PAD4;SA sector interaction are upregulated in both ETI and PTI, it is likely part of the WRKY network. If this is the case, probably the input amplification ratios of multiple WRKYs are positively regulated by the PAD4;SA sector interactions.

**Figure 9.**
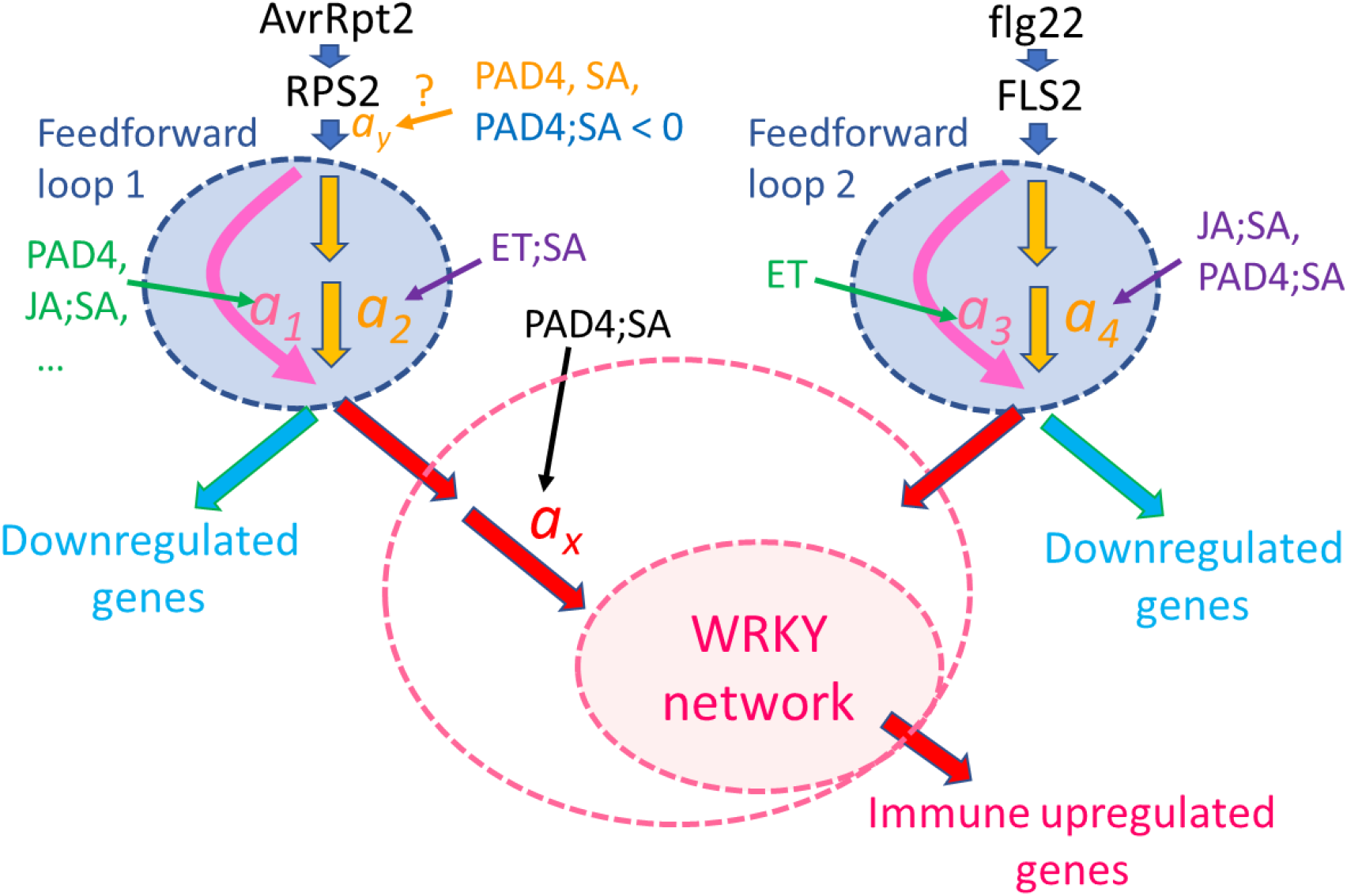
A possible regulatory model for the immune transcriptome response. A signaling event for the ETI response (left side) is initiated by recognition of AvrRpt2 by RPS2 (top left). The generated signal has a single-peak time course. The signal is processed through Feedforward loop 1, which can regulate the signal kinetics through modulation of the linear coefficients *a_1_* and *a_2_*.: accelerated by the PAD4 sector, the JA;SA sector interaction, and others; delayed by the ET;SA interaction. Although the peak amplitude positive regulation by the PAD4 and SA sectors and the negative regulation by the PAD4;SA sector interaction are tentatively placed before Feedforward loop 1, the current data cannot exclude the possibilities that this peak amplitude regulation occurs at later steps (thus, “?” is attached). The output of Feedforward loop 1 is processed separately for upregulated genes and downregulated genes. The amplification ratio *a_x_* for the upregulated gene regulation is positively regulated by the PAD4;SA sector interaction, which was observed as upregulation of the basal mRNA level of ETI-upregulated genes. Then the signal is fed into a transcriptionally interconnected WRKY network (Liu *et al.*, 2022), which upregulates the ETI-upregulated genes. The larger pink dashed ellipse around “WRKY network” indicates the possibility that the PAD4;SA sector interaction control of the amplification ratio *a_x_* is included within the function of the WRKY network. A signaling event for the PTI response (right side) is initiated by recognition of flg22 by FLS2 (top right). The generated signal is processed through a separate feedforward loop, Feedforward loop 2, which can regulate the signal kinetics through modulation of coefficients *a_3_* and *a_4_*. The signal for regulation of PTI-upregulated genes is fed into the WRKY network through different entry-point WRKYs. With different entry points, the WRKY network can upregulate similar but not identical gene sets (Liu *et al.*, 2022).

We emphasize that Figure 9 represents just one simple model that can explain all the observations discussed and that there are other possible models. However, we also emphasize that this is a highly testable mechanistic model as it is characterized by a well-defined network architecture and mechanistic model parameters. This level of mechanistic modeling was enabled by simplification of an ETI system by *in planta* effector expression, dynamical analysis that characterized the response with highly interpretable, quantitative dynamical parameters, and NRAM analysis that allowed proper estimation of interaction interpretations. This approach, comprised of a simplified experimental system, dynamical analysis, and NRAM on dynamical parameters, forms a platform highly applicable to mechanistic studies of other transient response systems that involve interactions of multiple components for their regulation.

## METHODS

### Plant materials and growth conditions

An Arabidopsis Col-0 line carrying *Ed-AvrRpt2* as a single copy insertion (*JEPS*; (Tsuda *et al.*, 2012)) was crossed to a *dde2-2 ein2-1 pad4-1 sid2-2* quadruple mutant line with Col-0 background (Tsuda *et al.*, 2012). *DDE2*, *EIN2*, *PAD4*, and *SID2* are the hub genes of the four immune signaling sectors, the JA, ET, PAD4, and SA sectors, respectively (Alonso *et al*, 1999; Jirage *et al*, 1999; von Malek *et al*, 2002; Wildermuth *et al*, 2001). Approximately 3500 F_2_ plants from the cross were progressively screened by PCR-based genotyping (Tsuda *et al.*, 2012; Tsuda *et al.*, 2009) for homozygosity of *Ed-AvrRpt2* and either wild-type- or mutant-allele homozygosity for *SID2*, *PAD4*, *DDE2*, and *EIN2*. For three of the desired genotypes, we only obtained plants with one gene that was heterozygous, so F_3_ plants from such F_2_ plants were screened for homozygosity. In this way, all 16 combinatorial genotype lines with the *Ed-AvrRpt2* transgene were prepared (*JEPS* to *xxxx*). *GUS* (Tsuda *et al.*, 2012) and *r1r2* (Hatsugai *et al.*, 2017) plant lines with Col-0 background were also used. Arabidopsis plants were grown in a controlled environment at 22°C with a 12-hour/12-hour light/dark cycle and 75% relative humidity.

### Experimental design of plant treatments

A single 18-genotype set was grown in two 15 cm * 15 cm pots (9 plants per pot), and four 18-genotype sets were grown in a single flat of 2 * 4 = 8 pots. In a single biological replicate, four and two flats of plants (16 and 8 18-genotype sets) were grown for Ed and mock treatments, respectively. The flats for Ed and mock treatments were separated to avoid cross-contamination of the treatments. For each of the genotype * treatment * time combinations (180 biological samples), two plants from two different flats were pooled. Thus, in a single biological replicate, 14 and 6 18-genotype sets for Ed and mock treatments (7 and 3 time points), respectively, were used. We used the remaining plants (2 18-genoteype sets for each treatment) as backups in case some plants did not grow well. Plants for three biological replicates were grown one week apart. In addition, the relative growing positions of 18 genotypes across two pots were randomized across the biological replicates.

### Plant treatment and collection

Twenty-four-day old plants were sprayed with 50 µM Ed or mock (both contained 0.25% ethanol in water) three hours after dawn (the time the lights switched on in the growth chamber). The aerial parts of plants were harvested at the indicated time (1, 2, 3, 4, 5, or 6 hpt). Unsprayed plants were used for 0 hpt. The aerial parts of two plants were pooled for each genotype * treatment * time * replicate sample. The harvested plant tissue was immediately flash frozen in liquid N_2_.

### RNA-tag-seq and preprocessing of the RNA-tag-seq data

Each of 540 tissue samples was pulverized and RNA was extracted using TRIzol (Thermo Fisher) and RNeasy (Qiagen) according to the manufacturers’ instructions. The RNA-tag-seq libraries were generated according to a 3’-tag sequencing method (Rallapalli *et al*, 2014). Thirty-three 16-plexed and one 12-plexed libraries were sequenced by the University of Minnesota Genomics Center using an Illumina HiSeq 2000 sequencer. The raw sequence data were preprocessed to obtain read-counts-per-gene data for each library according to (Hillmer *et al.*, 2017). The preprocessed data set included 33603 genes that had at least one read count in at least one library. The raw sequence data and the preprocessed read-counts-per-gene data have been deposited in NCBI GEO (accession: GSE196892).

### Estimation of the mean and the continuous mRNA level data

From the ETI-response preprocessed data, genes for which the read count was 0 in more than half of the libraries or for which the 15^th^ highest read count value across the libraries was fewer than 25 were removed, resulting in 18978 well-expressed genes. Pseudocounts were added to each library proportional to the 90^th^ percentile of the library (One pseudocount was added to the library with 55 read count as the 90^th^ percentile value) to have at least one read count in every gene in each library. GLM-NB (negative binomial generalized linear model) with the fixed effect with 18 genotypes * (7 Ed-treated times + 3 mock-treated times) = 180 levels was fit to the pseudocount-added data for each gene with the log-ratio of the 90^th^-percentile read count of the libraries over 500 as the offset, to obtain 180 mean estimates, their associated standard errors, and their deviance residuals on a log_2_ scale. Subsequently all the mRNA level values are on a log_2_ scale. The deviance residuals were added to their mean estimates to generate approximated continuous log_2_-transformed mRNA level data (the “continuous mRNA level” data). Tne deviance residuals were used for this purpose because they normally distribute in a log-scale. The inverse of the square of the standard errors were used as the weights in linear models when they were fit to the continuous mRNA level data since the continuous data do not preserve the count data errors of the sequence read data.

Using the mean estimates and standard errors from GLM-NB, dynamically responding genes were selected that had significantly different mRNA level values (*q* < 0.05 (by Storey FDR (Storey *et al*, 2005)) and |mean difference| > 1 (2-fold)) from 0 hpt at two or more consecutive time points later than 1 hpt in at least one of 18 genotypes, resulting in 10833 dynamical genes.

The experimentally manipulated genes, four genes for combinatorial genotypes, *DDE2*, *EIN2*, *PAD4*, and *SID2*, two *R* genes, *RPS2* and *RPM1*, and two transgenes, *Ed-AvrRpt2* and *XEV*, were removed from the figures that show the effects of the Ed-treatment and 18 genotypes unless specifically stated. When 16 combinatorial genotypes were compared, the two *R* genes, *RPS2* and *RPM1* were kept in the gene set as they were not manipulated within the 16 genotypes.

### Modeling mock time courses using the time course of *GUS* with Ed for each gene and the mean estimates of log_2_(Ed/mock) at every time point in every genotype

For each of 10833 dynamical genes in each combinatorial genotype, the mock treatment gives no-ETI response control values. However, we only had three time points for the mock treatment. We modeled mock mRNA levels for each gene at all seven time points using the time course model for *GUS* treated with Ed. In Figure 1B, the most prominent pattern was little difference in *GUS*/*JEPS* and similar and large differences in *r2r1*/*JEPS* and *r2r1*/*GUS*, indicating substantial effects of the *RPS2* and/or *RPM1* genes regardless of the transgene. Therefore, we used the data from *GUS* instead of *r2r1* to model the mock mRNA level values at 7 time points. First, a 4^th^-order time polynomial regression model was fit to the *GUS* continuous mRNA level data. Second the best polynomial model was selected using the step function and was used as the *GUS* time course model for the gene. Third, a linear model, ∼genotype/time, was fit to the mock continuous mRNA level data for 18 genotypes * 3 time points. Fourth, the best model among ∼genotype/time, ∼genotype, and ∼1 was selected using the step function, and the mean value for 0 hpt of mock was estimated using the best model. Fifth, for each genotype, the estimated 0 hpt of mock was used for the intercept of the time course model of *GUS* with Ed to obtain the “mock time-course model” (Figure S6). The mock time-course model was used to obtain the mean estimates of the mock values and their standard errors for all genotypes at seven time points. Then, the mock mean estimates were subtracted from the Ed mean estimates from the GLM-NB model to calculate the mean estimates for log_2_(Ed/mock) for each gene at every time point in every genotype. The associated standard errors were also calculated. ETI-responsive genes were selected using the criteria that the log_2_(Ed/mock) values for *q* < 0.05 and |mean| > log_2_3 (i.e., 3-fold) at two or more consecutive time points after 2 hpt in at least one genotype, resulting in 3499 genes.

### Modeling the dynamics of the ETI-responsive genes

From 3499 ETI-responsive genes, three non-overlapping gene sets were selected, 404 early.peak genes, 972 early.peak2 genes, and 1912 late.peak genes. For the early.peak genes, genes were selected that had the maximum |log_2_(Ed/mock)| value among 7 time points later than 2 hpt and earlier than 5 hpt in at least one genotype. For the early.peak2 genes, genes were selected that had the maximum |log_2_(Ed/mock)| value among 7 time points later than 2 hpt and earlier than 6 hpt in at least one genotype, and then the genes overlapping with early.peak genes were removed. The late.peak genes had the maximum |log_2_(Ed/mock)| value among 7 time points later than 2 hpt in at least one genotype, and then the genes overlapping with early.peak or early.peak2 genes were removed. Based on visual inspections of the time courses, AT3G03190, AT3G14870, AT3G62950, and ATCG00190 were removed from early.peak genes, and AT1G21160, AT2G14610, AT4G23130, AT5G02320, AT5G10140, AT5G41761, and ATCG01120 were removed from early.peak2 genes, as the time course shapes of some genotypes of these genes did not seem fit the gamma-pdf shape (see the next paragraph). The *DDE2*, *PAD4,* and *SID2* experimentally manipulated genes and 13 plastid-encoded genes were removed from these gene sets. In the end, we had 399 early.peak, 963 early.peak2, and 1900 late.peak genes (total 3262 “modeled” genes; 1972 upregulated and 1290 downregulated genes) selected for the time-course model fitting described below.

We assumed that the shape of the log_2_(Ed/mock) time course was approximated by the shape of gamma-pdf. Thus, the following function *f*(*t*) was fit to each of the early.peak, early.peak2, and late.peak genes.

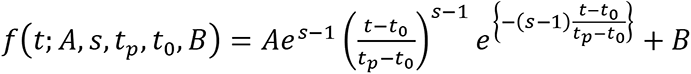

where *t* is the time in hpt, *A* is the peak amplitude on a log_2_ scale (*A* > 0 and *A* < 0 for upregulated and downregulated), *s* is the shape parameter (*s* > 1; the larger the *s*, the sharper the peak shape), *t*_*p*_ is the peak time in hpt; *t*_0_ is the time lag in hpt (*f*(*t* < *t*_0_) = 0); *B* is the baseline to make *f*(0)∼0 and *f*(∞)∼0. This model is called a gamma-pdf time course model. Example time course models are shown in Figure 4A. The model was fit to the approximated continuous log_2_(Ed/mock) mRNA ratio data using least square. Among the model parameters, *A*, *s* (log_2_- scaled for a better distribution), and *t*_*p*_ were fit for a time course of each genotype for each gene, while *t*_0_ and *B* were made common across the genotypes for each gene. As the statistical power of the data for constraining the parameter values was often limited, the parameter ranges in model fitting were *ad hoc* constrained. This statistical power issue was particularly severe with the genes and the genotypes with peak times later than 6 hpt, which was the last time point in the data. This was the reason that the genes were first divided into three gene sets according to the peak time, and then the model with different parameter ranges was fit separately for different gene sets. See the R script in Dataset S1 for details.

### Network reconstitution via averaging model (NRAM) analysis

For NRAM of the peak amplitudes and times of each gene for the four signaling sectors, the genes with peak times before 7.5 hpt in more than 13 out of 16 combinatorial genotypes were selected from the early.peak and early.peak2 genes (866 “early-modeled” genes). This was because late peak time estimates are not reliable. NRAM analysis was conducted according to (Katagiri, 2022) for the peak amplitudes, the peak times, the basal levels of genes, and the ETI-PC1 values. Briefly, NRAM is a general linear model approach to decompose the phenotype values across the 16 exhaustively combinatorial genotypes. They are decomposed into four signaling sector effects and their interactions. In the averaging model, an interaction (which is denoted using a semicolon, “;”) was defined as the difference between the phenotype of the genotype with all sectors in the interaction functional and the average of phenotypes of all genotypes with one of the sectors dysfunctional. This interaction definition makes the averaging model interaction mathematically stable, consistent, and interpretable for any number of sectors involved in the interaction.

### Reanalysis of the RNA-seq data for PTI response

The previously published RNA-seq data set for the flg22-PTI response (NCBI GEO accession, GSE78735; (Hillmer *et al.*, 2017)) was reanalyzed similarly to the RNA-seq data set for the ETI response in this study. Details of the analysis are described in Text S2. Briefly, 17423 well-expressed genes, 12537 dynamical genes, and 2946 PTI-responsive genes were progressively selected, and 2484 genes well-modeled by the gamma-pdf time-course model were obtained. See the R script in Dataset S1 for further details.

## ACKNOWLEDGEMENTS

We thank the University of Minnesota Genomics Center for sequencing. This work was supported by grants from the National Science Foundation to FK (grant nos. MCB-0918908, MCB-1518058, and IOS-1645460), a grant from the United States Department of Agriculture-National Institute of Food and Agriculture to FK (grant no. 2020-67013-31187), and by a grant from Ajinomoto Co., Inc. to FK.

## SUPPLEMENTAL INFORMATION

Text S1. Detailed interpretations of the PCA results shown in Figure 1A.

Text S2. Descriptions of reanalysis of the PTI transcriptome response data.

**Figure S1.**
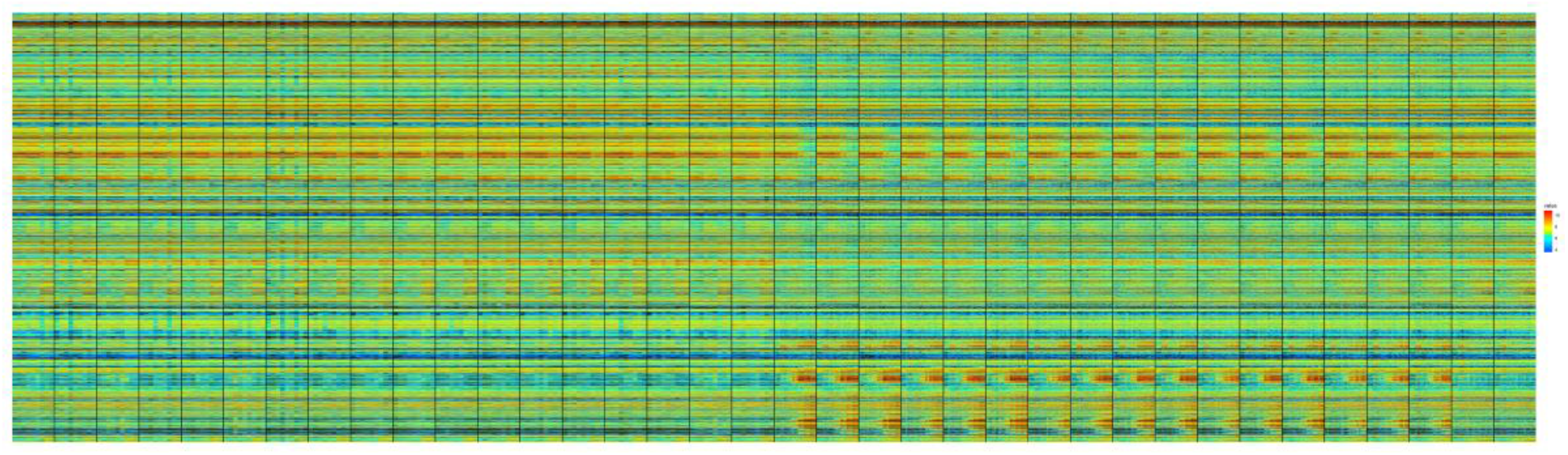
Heatmap of the mRNA levels of all 18978 well-expressed genes x 540 libraries. Major column divisions by gray vertical lines (36 major columns) divide each genotype * treatment. The major columns 1 to 18 are mock treated and the major columns 19 to 36 are Ed treated. For each treatment, the genotype order is *JEPS, xEPS, JxPS, JExS, JEPx, xxPS, xExS, xEPx, JxxS, JxPx, JExx, xxxS, xxPx, xExx, Jxxx, xxxx, r2r1*, and *GUS*. Within a mock major column, the major column is first divided into 3 time points (0, 2, and 5 hpt) and then the time points are divided into three biological replicates. Within an Ed major column, the major column is first divided into 7 time points (0, 1, 2, 3, 4, 5, and 6 hpt) and then the time points are divided into three biological replicates. Note that since the mock major columns have fewer time points, the width of a minor column (each replicate) is wider than that for the Ed major columns. In this way, the major column widths are the same, and the approximately corresponding times across the mock and Ed major columns can be intuitively compared. The color scale for the log_2_- transformed value is shown at the right. Rainbow colors are used so that relatively small differences in the heatmap are readily visible.

**Figure S2.**
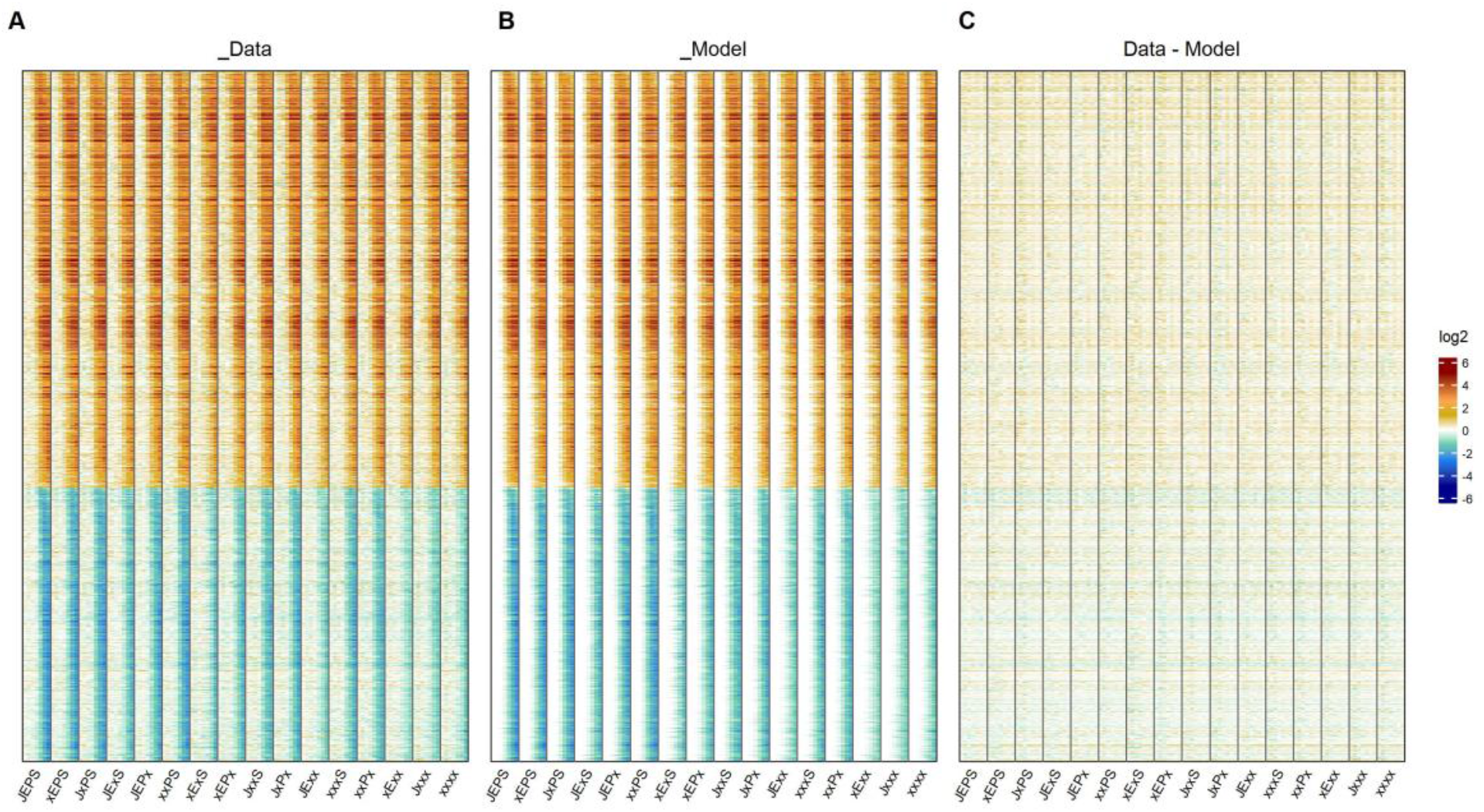
The fit of the ETI-response gamma-pdf time-course models by heatmaps. In all three heatmaps, the gene set (1972 upregulated and 1290 downregulated genes) and order for the rows and the genotype/time columns are the same as those in Figure 3A. The color code for the log_2_ ratio is the same as that in Figure 3B. A. A heatmap of the data for the log_2_ ratio of Ed/mock, which is the same as Figure 3A. The Ed values are the mean estimates of the GLM-NB. The mock values are from the mock time course estimated based on the *GUS* Ed time course and the three time point mock data of the genotype for each gene. B. A heatmap of the modeled values for the log_2_ ratio of Ed/mock. C. A heatmap of the discrepancies between the data and the model.

**Figure S3.** The fit of the ETI-response gamma-pdf time-course models by a time course plot for each of 1972 ETI-upregulated and 1290 downregulated genes. For each gene, the data and the model plots are shown side by side (plots for two genes per line). The data and the model values are the same as those used in Figures 3A and S2A and Figure S2B. The gene order is the same as that in these figures, top to bottom. The genotype color code and line type code are the same as those used in Figure 3D. Note that the 1-hpt data points were not used in model fitting although some genes show a transient response at 1 hpt. This is because such very early transient responses likely represent general stress responses and are unlikely to represent immune-specific responses (Bjornson *et al.*, 2021). This figure is provided as a separate file since it is very large.

**Figure S4.**
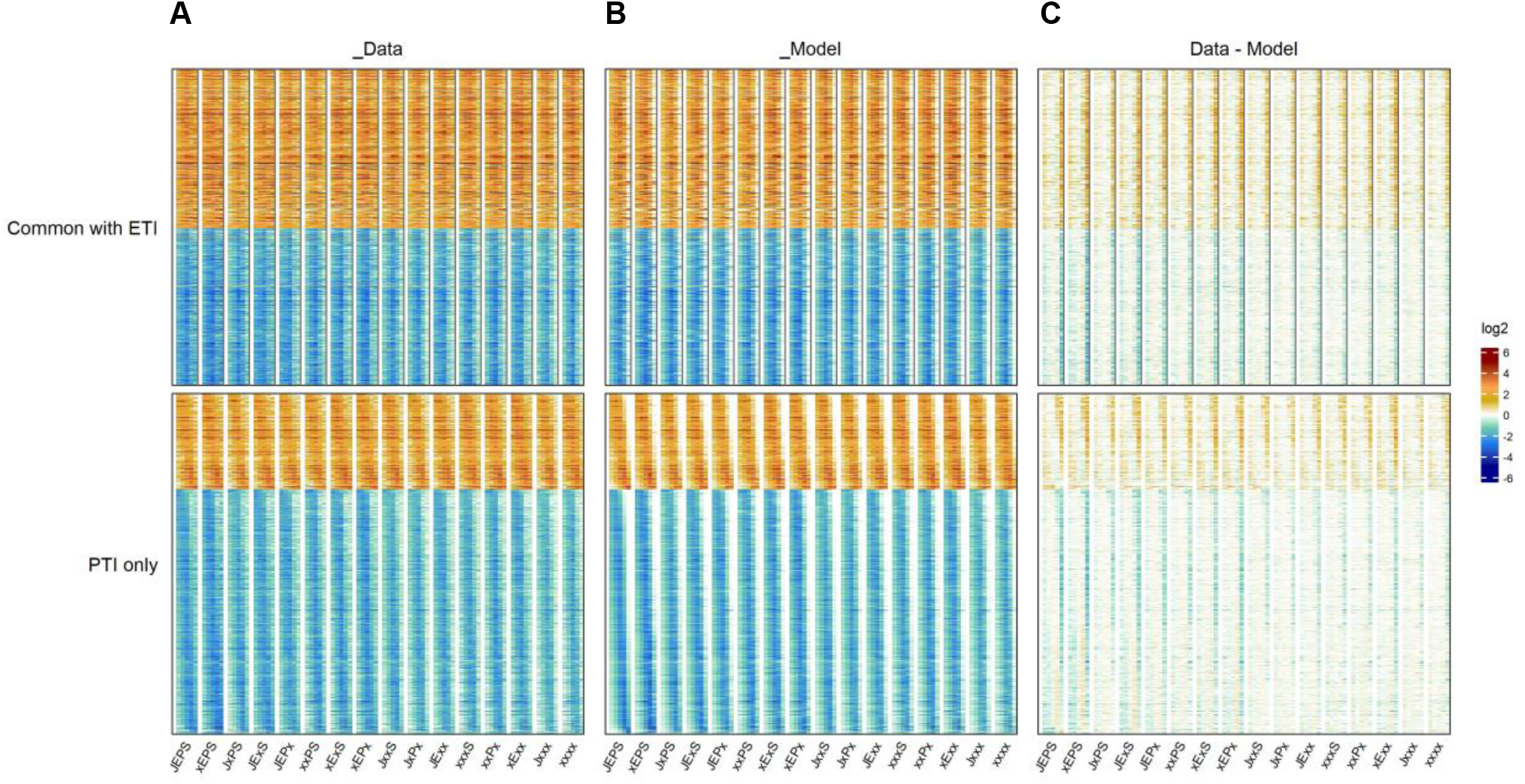
The fit of the PTI-response gamma-pdf time-course models for 2484 well-modeled genes by heatmaps. The heatmaps are divided into two for each panel: top, “Common with ETI”, the heatmap of the genes overlapping with ETI-response models (602 ETI-upregulated and 594 downregulated genes; Figure 5A) in the same order but with non-overlapping genes removed; bottom, “PTI only”, the rest of the genes for the PTI response models (357 upregulated and 931 downregulated genes). The genes were ordered according to the JEPS peak time for each set of the upregulated and the downregulated genes. Each genotype column (a main column of each panel) is divided into six columns for time points, 0, 2, 3, 5, 9, and 18 hpt. A. Heatmaps of the data. It is the log_2_ ratio of flg22/mock for each time point in each genotype for each gene. The flg22 values are the mean estimates of the GLM-NB. The mock values are from the mock time course estimated based on the *fls2* time course. B. A heatmap of the gamma-pdf time-course modeled values of the log_2_ ratio of flg22/mock. C. A heatmap of the discrepancies between the data and the model. The color code for the log_2_ ratio is the same as that in Figure 3B. Note that the 1-hpt data points were not used in model fitting although many genes show a transient response at 1 hpt. This is because such very early transient responses likely represent general stress responses and are unlikely to represent immune-specific responses (Bjornson *et al.*, 2021). The 18-hpt data points were not used in model fitting either, as we tried to compare PTI and ETI responses over a comparable time range. However, a substantial number of genes showed late second-peak responses while the time-course model used can only model single-peak responses. This is the reason for non-negligible discrepancies at 18 hpt in C.

**Figure S5.** The fit of the PTI-response gamma-pdf time-course models by a time course plot for each of 2484 well-modeled genes. For each gene the data and the model plots are shown side by side (plots for two genes per line). The data and the model values are the same as those used in Figure S4A and Figure S4B. The time axis (the *x*-axis) of each plot is squire-root-scaled to have the peak shape close to symmetric. The gene order is the same as that in Figure S4A, top to bottom. The genotype color code and the line type code are the same as those used in Figure 3D. The 1- and 18-hpt data were not used for modeling as explained in the Figure S4 legend. This figure is provided as a separate file since it is very large.

**Figure S6.**
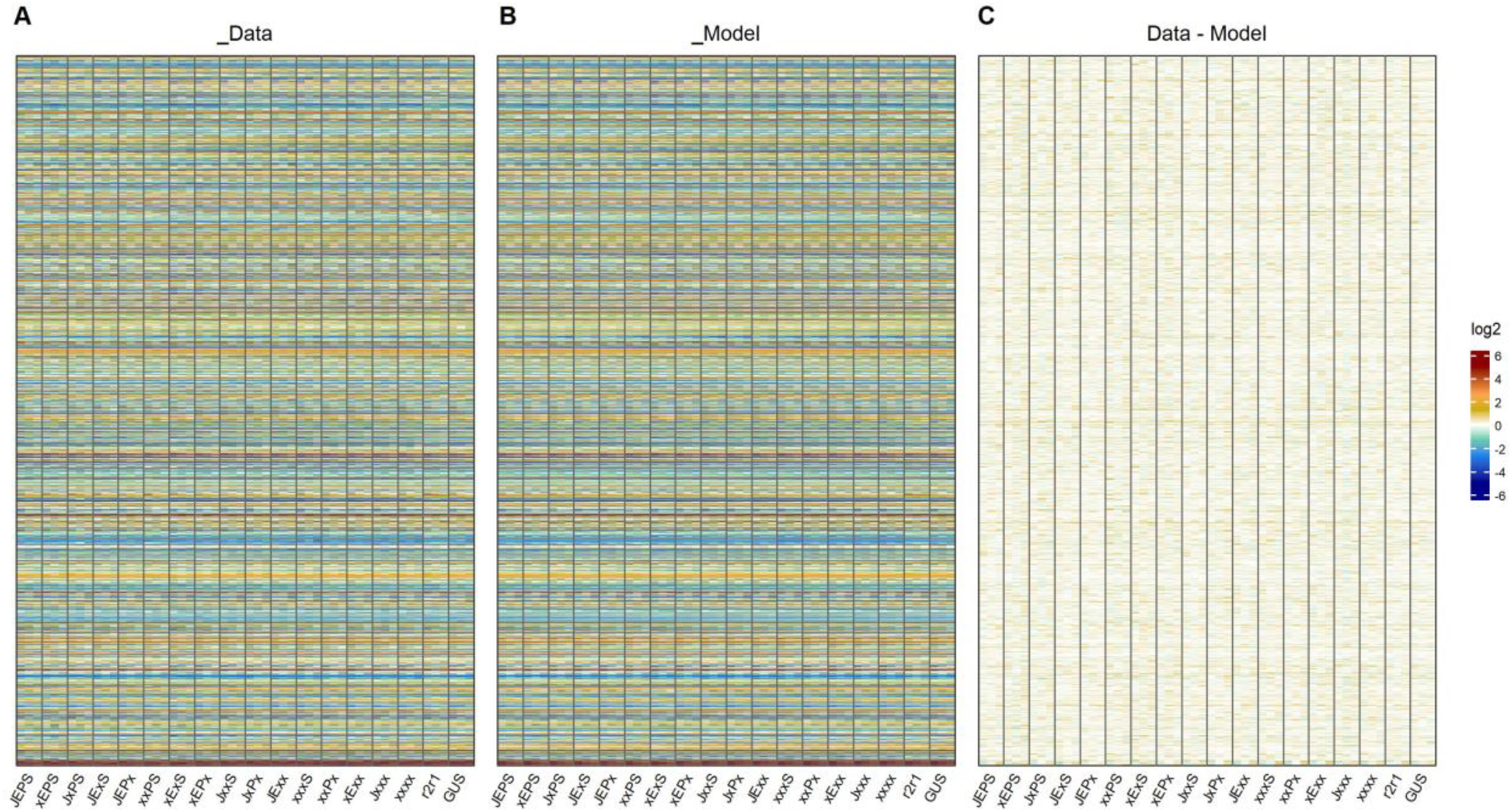
The fit of the mock time-course polynomial models for the ETI-response transcriptome data by heatmaps. A. A heatmap of the data. The data values are the mean estimates of GLM-NB at three time points (0, 2, and 5 hpt) after mock treatment for 18 genotypes for 10833 dynamical genes. B. A heatmap of the model. The mock time-course modeled values were used. C. A heatmap of the discrepancies between the data and the model. The genes are ordered according to the AGI gene code. The genotypes for each major column are indicated. Each major column has three minor columns for 0, 2, and 5 hpt. The row and column names are the same across three heatmaps for easy comparisons. The grand mean of the values in each heatmap was subtracted for better visualization in A and B, while the values in C were calculated from the data and model values before subtraction of the grand means.

Table S1. The fitted parameter values of the ETI-response gamma-pdf time-course models and the fitted values of the model at 0, 1, 2, 3, 4, 5, and 6 hpt for each genotype for each of 3262 well-modeled genes. The gene order in the row is the same as the gene order in Figure 3A. Generally, the column names are the genotype name followed by the parameter name or the time point, separated by “_”, except the parameters *t*_0_ and *B*, which have single values across the genotypes for each gene. The parameters are indicated by “A”, “log2s”, “pt”, “t0”, and “B” for *A*, log_2_ *s*, *p*_*t*_, *t*_0_, and *B* in the model, respectively. For example, “JxPS_log2s” indicates the log_2_ *s* value for the genotype *JxPS*, and “xExx_2h” indicates the 2 hpt modeled value for the genotype *xExx*.

Table S2. The fitted parameter values of the PTI-response gamma-pdf time-course models and the fitted values of the model at 0, 2, 3, 5, 9, and 18 hpt for each genotype for each of 2484 well-modeled genes. The gene order in the row is the same as the gene order in Figure S4. Generally, the column names are the genotype name followed by the parameter name or the time point, separated by “_”, except the parameters *t*_0_ and *B*, which have single values across the genotype for each gene. The parameters are indicated by “A”, “log2s”, “pt”, “t0”, and “B” for *A*, log_2_ *s*, *p*_*t*_, *t*_0_, and *B* in the model, respectively. For example, “JxPS_log2s” indicates the log_2_ *s* value for the genotype *JxPS*, and “xExx_2h” indicates the 2 hpt modeled value for the genotype *xExx*.

Dataset S1. R scripts and the data sets used in the study. Dataset S1 is available from https://github.com/fumikatagiri/Arabidopsis_ETI_transcriptome_response. The dataset contains an R script, “Script_Ed-AvrRpt2_ms_230503f.r”, which was used to generate Figures 1, 2, 3, 4, 6, S1, S2, S3, and S6 and Table S1, and another R script, “Script_flg22.230503f.r”, which was used to generate Figures 5, S4, and S5, and Table S2. Two input RNA-seq read count data files used in the scripts (from NCBI GEO accessions GSE196892 (this work) and GSE78735 (Hillmer *et al.*, 2017)) are in the “data” subfolder. Files for the circadian and diurnal gene data from (Yang *et al.*, 2020) are also in the “data” subfolder, which are required for generating figures in Text S1. To run these R scripts, an empty “outputs” subfolder is required. Run “Script_Ed-AvrRpt2_ms_230503f.r” first, as “Script_flg22.230503f.r” requires some output files generated by the former.

