## Supplementary material for "The kinetics and basal levels of the transcriptome response during Effector-Triggered Immunity in Arabidopsis are mainly controlled by four immune signaling sectors": Text S1

Text S1. Details of the PC1 x PC2 plot in Fig. 1A

This supplemental text describes detailed interpretations of the PCA results shown in Fig. 1A.

### **PC2 failed to capture major biological variation.**

In Fig. TS1.1 "eigengenes" for PC1 and PC2 are plotted for the four genotypes, *JEPS*, *xxxx*, *r1r2*, and *GUS* ("PC1 gene" and "PC2 gene", respectively; red, Ed-treated; blue, mock-treated).

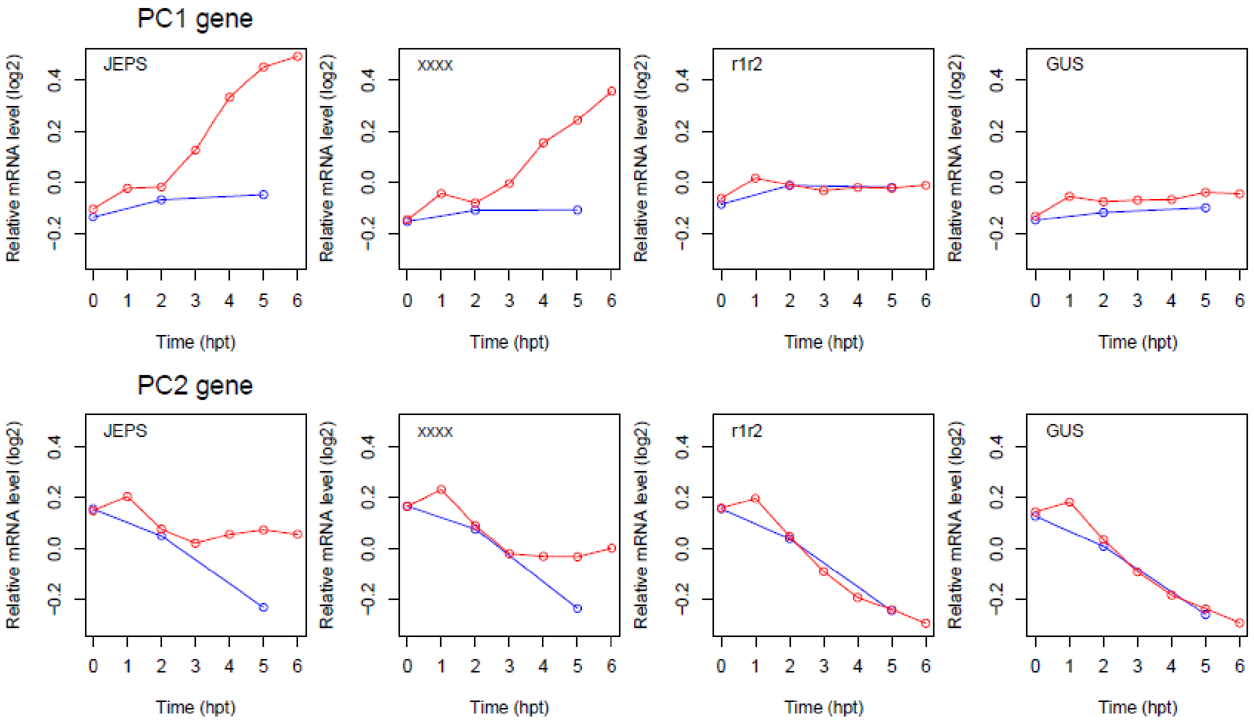


Fig. TS1.1. The time courses of the PC1 (top) and PC2 (bottom) "eigengenes" in the *JEPS*, *xxxx*, *r1r2*, and *GUS* genotypes. The Ed-treated (red) had 7 time points, 0, 1, …, 6 hpt, and the mock-treated (blue) had 3 time points, 0, 2, and 5 hpt. Note that the values were centered for each gene, so the absolute value of the basal levels (at 0 hpt) do not have any meaning. Also, values were scaled to vector size 1 in each gene. The positive and negative signs of PC1 and PC2 genes are arbitrary according to the definition of PCA.

Since we define the ETI response as $\log_2(\text{Ed/mock}) = \log_2(\text{Ed}) - \log_2(\text{mock})$, not only PC1 but also PC2 represent the ETI response (in the background of changing mock-treated) in *JEPS* and *xxxx*. This ETI response in PC2 gene occurred due to the orthogonality requirement between PC1 and PC2: in PC2 gene, Ed-treated stays fairly stable around 0 in *JEPS* and *xxxx* when the corresponding time course changes substantially in PC1 gene ( > 2 hpt in *JEPS* and > 3 hpt in *xxxx*); all the other genotype:treatment combinations are fairly stable in PC1 gene and change substantially in PC2 gene.

Thus, a gene with the time-course pattern we expect for a gene whose mRNA level monotonously changes along the time course irrespective of the treatment or the genotype can be well-approximated by a linear combination of $-a$ * "PC1 gene" + $a$ * "PC2 gene" ($a = 1/\sqrt{2}$ for the unit vector size), which is designated as "PC2 gene – PC1 gene" (Fig. TS1.2).

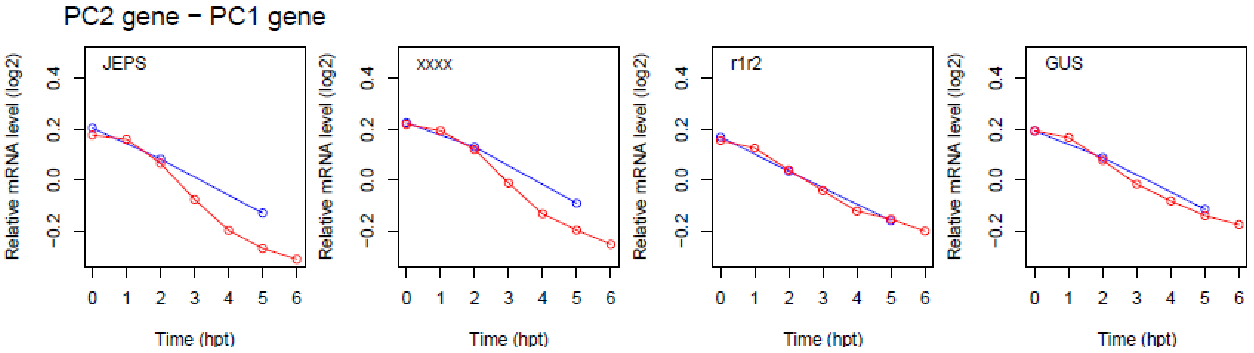


Fig. TS1.2. The time courses of the "PC2 gene – PC1 gene" eigengene in the *JEPS*, *xxxx*, *r1r2*, and *GUS* genotypes. Red, Ed-treated; blue, mock-treated.

When the 18970 genes used in Fig. 1A were plotted with the PC1 gene and PC2 gene coordinates (Fig. TS1.3), there were more genes along the "PC2 gene – PC1 gene" axis (green line) than along the PC2 gene axis, particularly "PC2 gene" > 0.

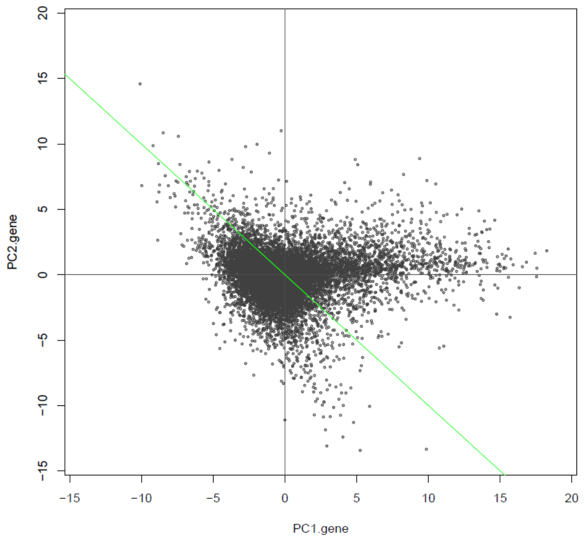


Fig. TS1.3. The distribution of the mRNA level patterns of 18970 genes in the PC1 gene x PC2 gene space. The green line shows "PC2 gene – PC1 gene" axis.

This figure indicates that the major biological variation captured in the PC1 x PC2 space (62.4% of the variance) was along PC1 (particularly on the positive side) and PC2-PC1. PC2 itself failed to capture major biological variation due to the orthogonal requirement among PCs. Actually, PC2 captured a minor type of ETI-specific mRNA level response while the baseline mRNA level was monotonously changing along hpt (Fig. TS1.1, bottom).

In fact, when the time-course modeled ETI-responsive genes (1251 up- and 835 down-regulated genes, Fig. 3A) were overlayed in the same plot (Fig. TS1.4), it was clear that the both PC1 gene and PC2 gene are ETI-responsive and that the "PC2 gene – PC1 gene" axis well represents the borderline between up- and down-regulated.

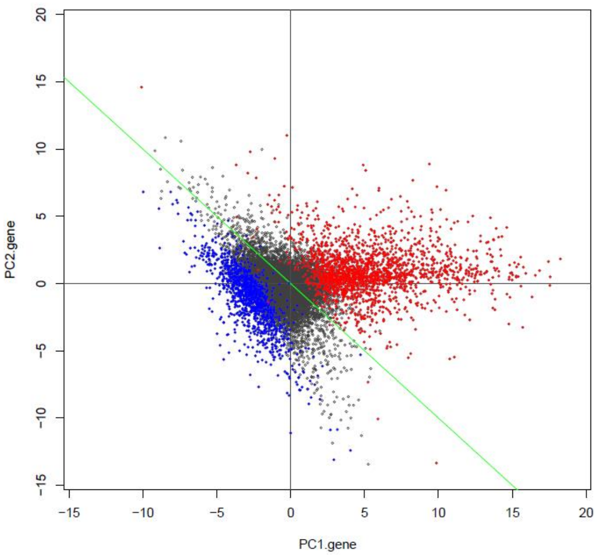


Fig. TS1.4. Time-course modeled ETI upregulated (red) and downregulated (blue) genes among the 18970 genes in the PC1 gene x PC2 gene space.

### The gene types associated with PC2-PC1

The observation of the "PC2 gene – PC1 gene" line as the border for ETI up- and down-regulated genes suggests that the values projected onto the "PC1 gene + PC2 gene" line (diagonal to "PC2 gene – PC1 gene") would well represent the ETI responsiveness, particularly for upregulated genes. Based on the spatial distributions of the modeled up- and down-regulated genes (red and blue in Fig. TS1.4), the genes with the values projected onto the "PC1 gene + PC2 gene" axis > 2 and < -2.5 were designated PC1+PC2.up genes (orange in Fig. TS1.5) and PC1+PC2.down genes (turquoise), and these two gene sets were subjected to GO term enrichment analysis.

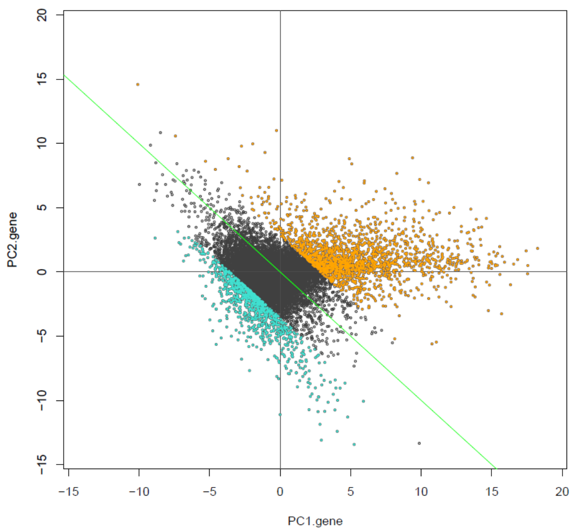


Fig. TS1.5. The PC1+PC2.up genes (orange, 1877 genes) and the PC1+PC2.down genes (turquoise, 967 genes) in the PC1 gene x PC2 gene plot.

As expected, PC1+PC2.up genes (1877 genes) were extremely highly enriched with immunity-related biological process terms, such as "response to biotic stimulus" (GO:0009607) for 799 genes (Bonferroni-corrected $p$ = 7.71E-296). However, no terms containing "light" or "circadian" were significantly enriched (i.e., such terms had Bonferroni-corrected $p > 0.05$). PC1+2.down genes (967 genes) were highly enriched with biological process terms related to light response and photosynthesis, such as "response to light stimulus" (GO:0009416) for 227 genes (Bonferroni-corrected $p$ = 3.70E-52) and "tetrapyrrole metabolic process" (GO:0033013) for 46 genes (Bonferroni-corrected $p$ = 4.26E-14).

We also applied the GO term enrichment analysis to the genes along the "PC2 gene – PC1 gene" (green) line. PC2-PC1.up genes (magenta in Fig. TS1.6) and PC2-PC1.down.genes (gold) were defined as the genes with values projected onto the "PC2 gene – PC1 gene" axis > 2.5 and < -2.5, respectively, and not overlapping with PC1+PC2.up genes or PC1+PC2.down genes.

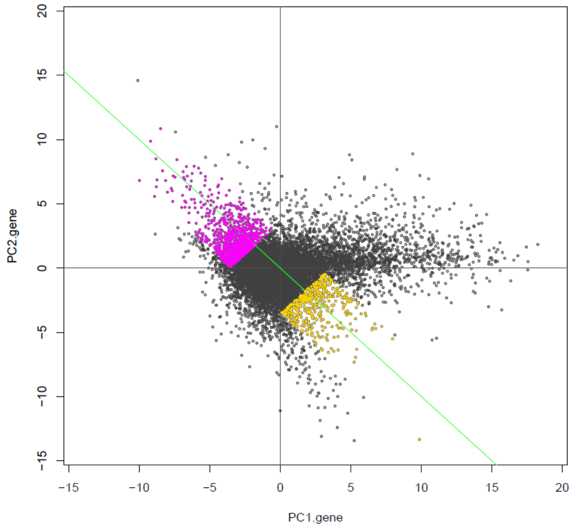


Fig. TS1.6. The PC2-PC1.up genes (magenta, 1071 genes) and the PC2-PC1.down genes (gold, 458 genes) in the PC1 gene x PC2 gene plot.

The PC2-PC1.up genes (1071 genes) were highly significantly enriched with biological process terms related to response to light, such as "response to radiation" (GO:0009314) for 266 genes (Bonferroni-corrected $p$ = 9.52E-66). In addition, the biological process term "circadian rhythm" (GO:0007623) was significantly enriched (Bonferroni-corrected $p$ = 6.24E-7). The PC2-PC1.down genes (458 genes) were highly significantly enriched with biological process terms like "response to stimulus" (GO:0050896, Bonferroni-corrected $p$ = 4.70E-41). The PC2-PC1.down genes appear to generally contain responsive genes but do not appear to strongly represent specific types of responses.

Since we conducted the infiltration procedures two to four hours after the light in the growth chambers turned on, we thought that the "PC2 gene – PC1 gene" axis was associated with circadian/diurnal responses, which is consistent with the GO term enrichment with the PC2-PC1.up genes. We selected with high confidence 1465 genes regulated by the circadian clock (circadian genes, magenta in Fig. TS1.7) and 917 diurnally regulated genes (diurnal genes, green) from Yang et al. (Yang *et al*, 2020), by applying the filter of meta2d_BH.Q < 0.0001 ("meta2d_BH.Q" is the false discovery rate based on the integrated $p$-values according to Yang et al.). 370 genes overlapped between the circadian and diurnal genes. It indeed appeared that circadian and diurnal genes largely distributed along the "PC2 gene – PC1 gene" axis (Fig. TS1.7).

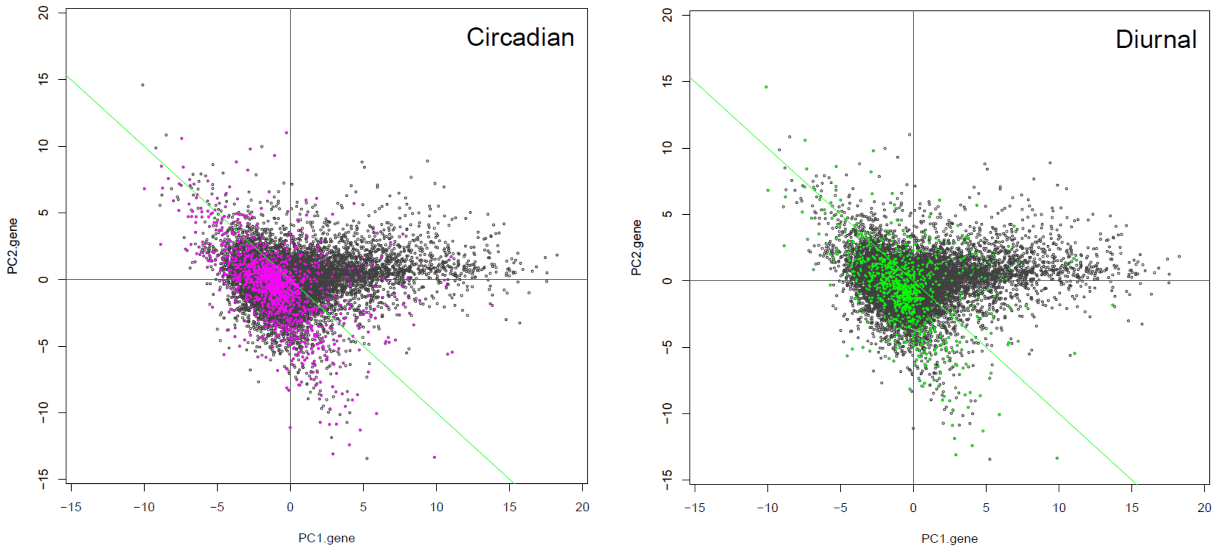


Fig. TS1.7. The circadian genes (magenta, 1465 genes; left panel) and the diurnal genes (green, 917 genes; right panel) in the PC1 gene x PC2 gene plot.

If the “PC2 gene – PC1 gene” axis mainly represents circadian and diurnal responses during our experimental time period, it is expected that genes with particular phases should be enriched along the “PC2 gene – PC1 gene” axis. When we color-coded the circadian and diurnal genes according to their phases (“ARS_adjphase” by Yang et al. (Yang *et al.*, 2020)) (Fig. TS1.8), indeed the genes with phases around 0 hours (purple to red) tended to have high “PC2 gene – PC1 gene” values and genes with opposite phases around 12 hours (green to cyan) tended to have low “PC2 gene – PC1 gene” values. Thus, we conclude that the PC2-PC1 mainly represents circadian/diurnal responses during our experimental time period.

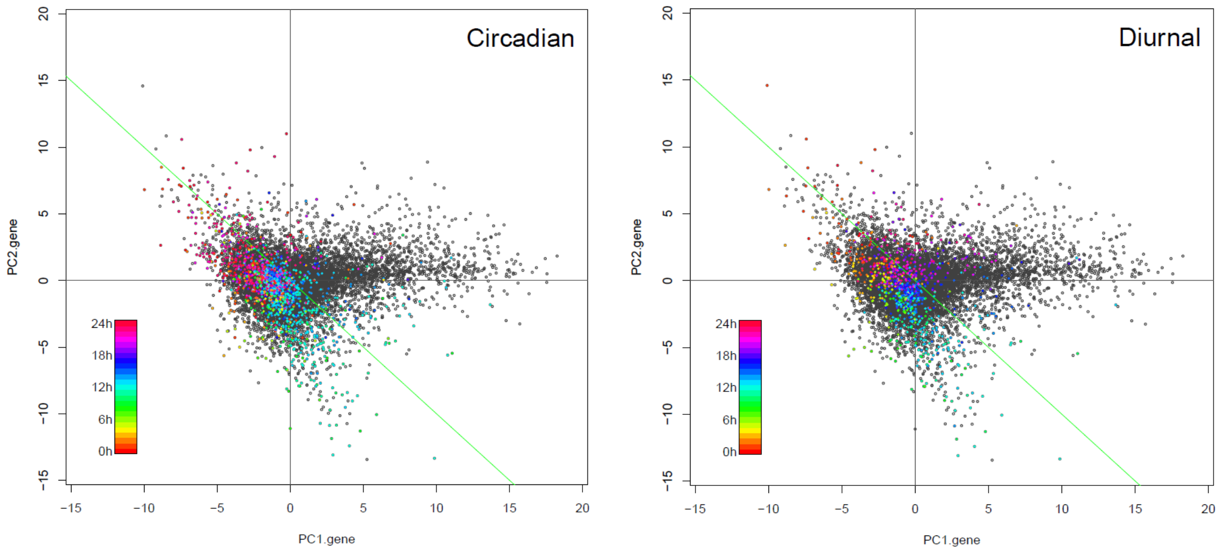


Fig. TS1.8. The phases of the circadian (left) and diurnal (right) genes were color-coded in the PC1 gene x PC2 gene plot.

Whereas PC2 does not capture the circadian and diurnal response across all treatment * genotype combinations, if focused on the non-ETI-response, Ed-treated *r2r1* and *GUS* and mock-treated all genotypes, the time courses of "PC2 gene" and "PC2 gene – PC1 gene" are similar (Fig. TS1.1, bottom and Fig. TS1.2). This indicates that PC2 mainly captures circadian/diurnal responses for Ed-treated *r2r1* and *GUS* and mock-treated all genotypes while this statement is not true for ETI-responding, Ed-treated *JEPS* and *xxxx*.

### METHODS

#### PCA

The R function prcomp (centred and non-scaled) was applied to the GLM-NB-estimated 40 genotype * time interaction values for the 18970 genes (a 18970-row, 40-column matrix) as in Fig. 1A. The unit vector along the PC1 gene axis was obtained by scaling the PC1 coordinates in the rotated data ($x) to the vector size 1. The inner product of a vector of the 40 values for a gene and the PC1 gene unit vector was the projected value of the gene on the PC1 gene axis. Similarly, the values projected onto the PC2 gene axis, "PC1 gene + PC2 gene" axis, and "PC2 gene – PC1 gene" axis were obtained for each gene.

#### GO term enrichment analysis

The Panther Classification System (http://go.pantherdb.org/) was used for the GO term enrichment analysis (Mi *et al*, 2019; Thomas *et al*, 2022).

#### The data for circadian and diurnal genes

The supplemental table files "159584_1_supp_509367_q8lbct.xlsx" and "159584_1_supp_509370_q8lbct.xlsx" for the circadian and diurnal genes, respectively, were obtained from Yang et al. (Yang *et al.*, 2020). After selecting $Type = "gene", the column value $meta2d_BH.Q < 0.0001 was used to select highly confident genes. The numbers of the genes in this document are those of the genes overlapping with the 18970 genes. The values in the $ARS_adjphase column were used as the phase values of the genes.
