## Supplementary material for "The kinetics and basal levels of the transcriptome response during Effector-Triggered Immunity in Arabidopsis are mainly controlled by four immune signaling sectors": Text S2

Text S2. Reanalysis of the RNA-seq data in response to flg22 in 16 combinatorial genotypes

**Estimation of the mean and the continuous mRNA level data**

From the preprocessed data (NCBI GEO accession GSE78735; (Hillmer *et al*, 2017), genes for which the read count was 0 in more than half of the libraries or for which the $10^{th}$ highest read count value across the libraries was fewer than 25 were removed, resulting in 17423 well-expressed genes. Pseudocounts were added to each library proportional to the $90^{th}$ percentile of the library (One pseudocount was added to the library with 27 read counts as the $90^{th}$ percentile value) to have at least one read count in every gene by library. GLM-NB with the fixed effect with 17 genotypes * 7 times = 119 levels was fit to the pseudocount-added data for each gene with the log-ratio of the $90^{th}$-percentile read count of the libraries over 500 as the offset, to obtain 119 mean estimates, their associated standard errors, and their deviance residuals on a $\log_2$ scale. Subsequently, all the mRNA level values are on a $\log_2$ scale. The deviance residuals were added to their mean estimates to generate approximated continuous $\log_2$-transformed mRNA level data (the "continuous mRNA level" data). The deviance residuals were used for this purpose because they normally distribute on a log scale. The inverse of the squares of the standard errors were used as the weights in linear models when they were fit to the continuous mRNA level data since the continuous data do not preserve the count data errors of the sequence read data.

Using the mean estimates and standard errors from GLM-NB, dynamically responding genes were selected that had significantly different mRNA level values ($q < 0.05$ (by Storey FDR (Storey *et al*, 2005)) and |mean difference| > 1 (2-fold)) from 0 hpt at two or more consecutive time points later than 1 hpt in at least one of 17 genotypes, resulting in 12537 dynamical genes.

**Modeling mock time courses using the time course of *fls2* with flg22 for each gene and the mean estimates of $\log_2$(flg22/mock) at every time point in every combinatorial genotype**

For each of 12537 dynamical genes in each combinatorial genotype, the mock treatment would give no-PTI response control values. However, we only had *fls2* as an equivalent of the mock treated *JEPS*. We modeled mock mRNA levels at all seven time points using the time course model for *fls2* in each gene. First, a $4^{th}$-order time polynomial regression model was fit to the *fls2* continuous mRNA level data. Second, the best polynomial model was selected using the step function and was used as the *fls2* time course model for the gene. Third, the $\log_2$(flg22/mock) mRNA level ratio was calculated at each time point of each of 16 combinatorial genotypes for the gene using the GLM-NB mean estimate values for the combinatorial genotypes. Fourth, the $\log_2$(flg22/mock) mRNA level ratio value at 0 hpt was subtracted to make the 0 hpt value zero in each genotype. The obtained $\log_2$(flg22/mock) mRNA level ratios are the mean estimates at each time point in each combinatorial genotype for the gene. The associated standard errors were also calculated.

PTI-responsive genes were selected with $\log_2$(flg22/mock) values for $q < 0.05$ and |mean| > $\log_2 3$ (i.e., 3-fold) at two or more consecutive time points after 1 hpt in at least one genotype, resulting in 2946 genes.

**Modeling the dynamics of the PTI-responsive genes**

The gamma-pdf time-course model was fit to the continuous $\log_2$(flg22/mock) values for each of 2946 PTI-responsive genes. First, two criteria were used to remove genes that were not appropriate for time-course modeling: (1) AT3G02480, as its time courses did not appear to have clear single-peak patterns; (2) Any gene whose maximum absolute values were at 1 hpt in more than one genotype; (3) Any gene whose sign of the maximum absolute values of the genotype across the genotypes was not consistent. Second, after fitting the model, any gene whose ratio of (the $2^{nd}$ latest peak time) / (the $2^{nd}$ earliest peak time) across the genotypes > 3 was removed as it was likely that some of the genotype time courses did not have clear single-peak patterns. After these quality controls, we obtained the gamma-pdf time-course models for 2484 genes. See the R script in Dataset S1 for further details.
