## Supplementary material for "The kinetics and basal levels of the transcriptome response during Effector-Triggered Immunity in Arabidopsis are mainly controlled by four immune signaling sectors": Figure S3

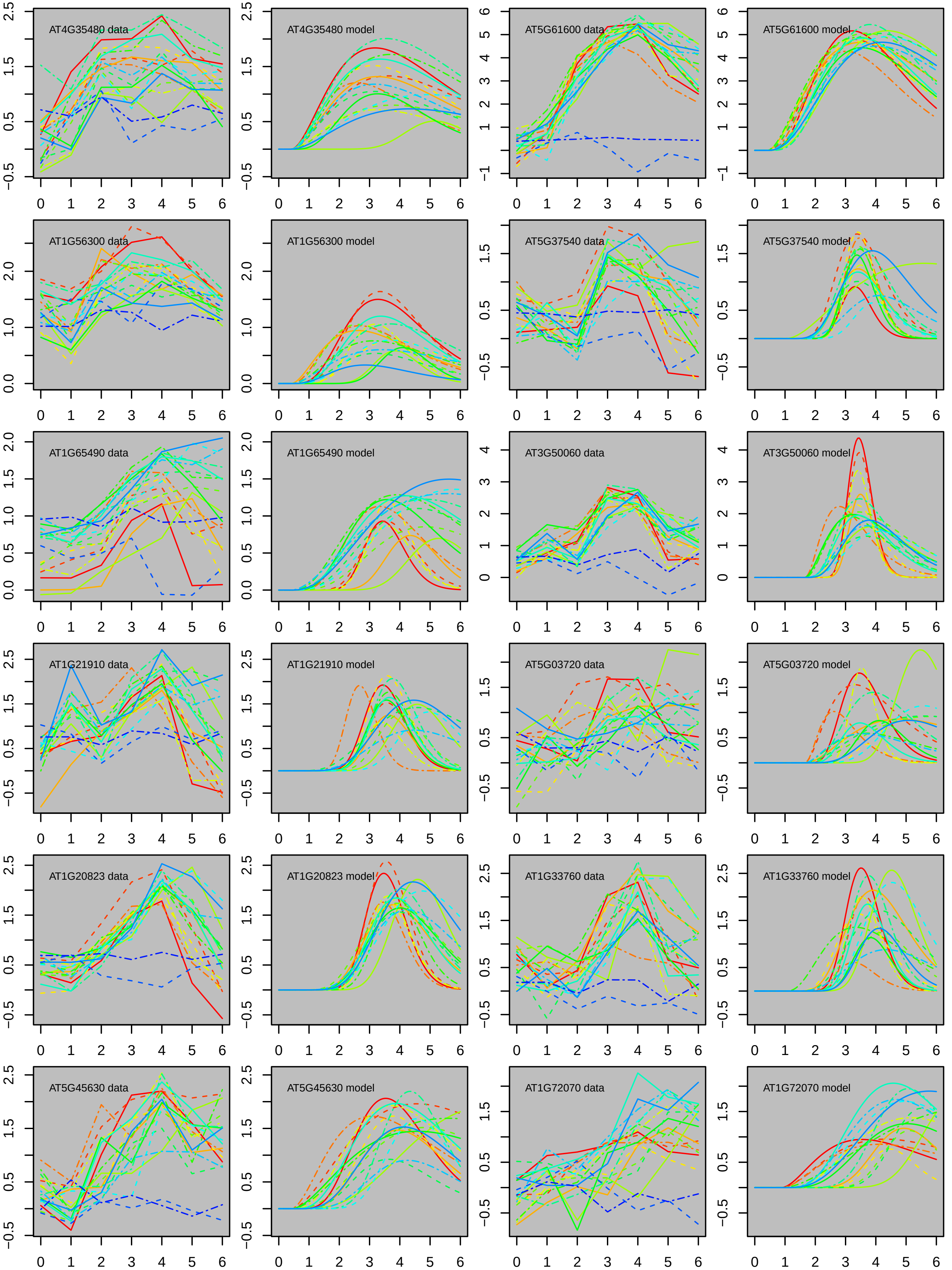
AT4G35480 data
AT4G35480 model
AT5G61600 data
AT5G61600 model
AT1G56300 data
AT1G56300 model
AT5G37540 data
AT5G37540 model
AT1G65490 data
AT1G65490 model
AT3G50060 data
AT3G50060 model
AT1G21910 data
AT1G21910 model
AT5G03720 data
AT5G03720 model
AT1G20823 data
AT1G20823 model
AT1G33760 data
AT1G33760 model
AT5G45630 data
AT5G45630 model
AT1G72070 data
AT1G72070 model

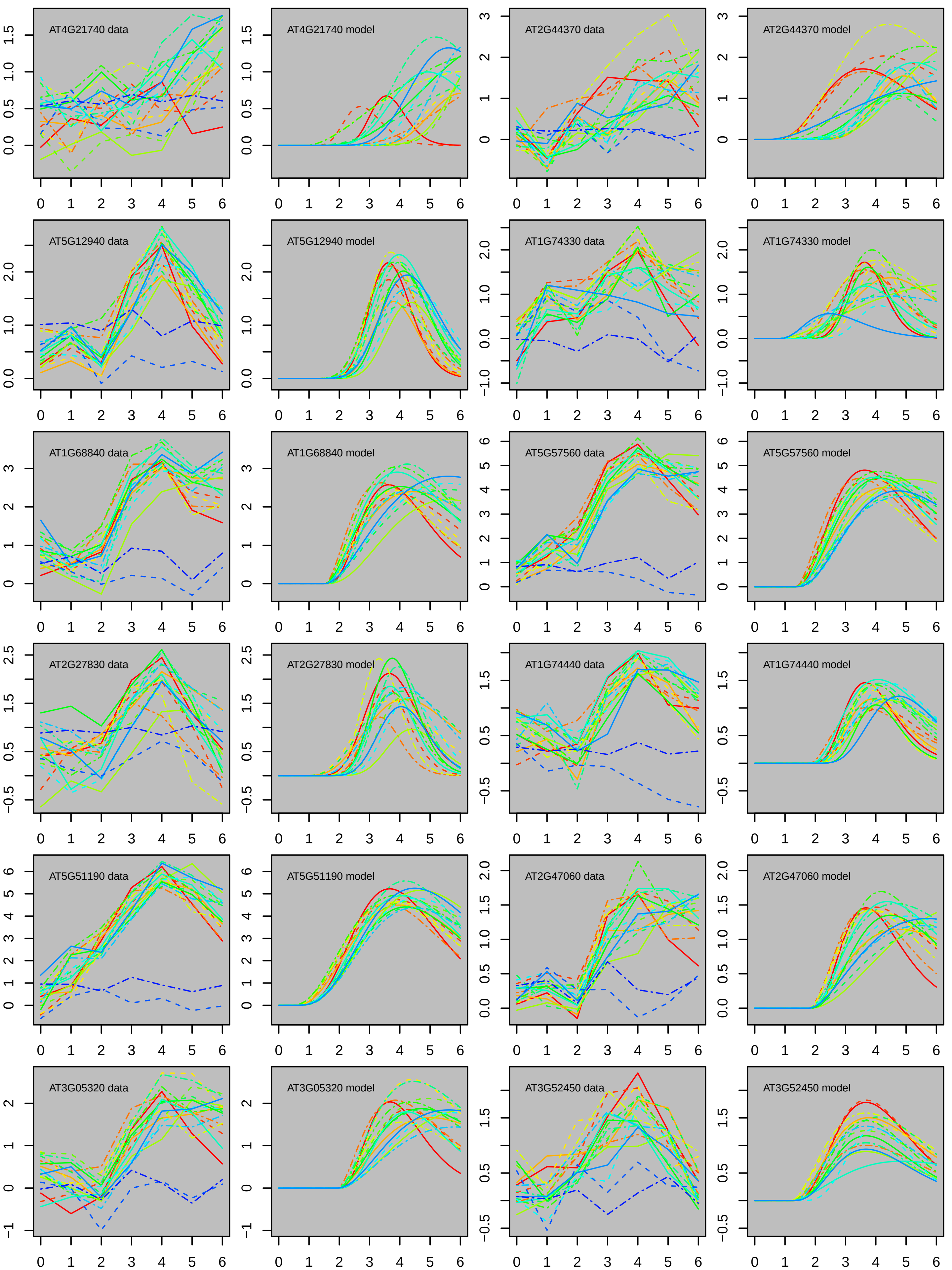
AT4G21740 data
AT4G21740 model
AT2G44370 data
AT2G44370 model
AT5G12940 data
AT5G12940 model
AT1G74330 data
AT1G74330 model
AT1G68840 data
AT1G68840 model
AT5G57560 data
AT5G57560 model
AT2G27830 data
AT2G27830 model
AT1G74440 data
AT1G74440 model
AT5G51190 data
AT5G51190 model
AT2G47060 data
AT2G47060 model
AT3G05320 data
AT3G05320 model
AT3G52450 data
AT3G52450 model

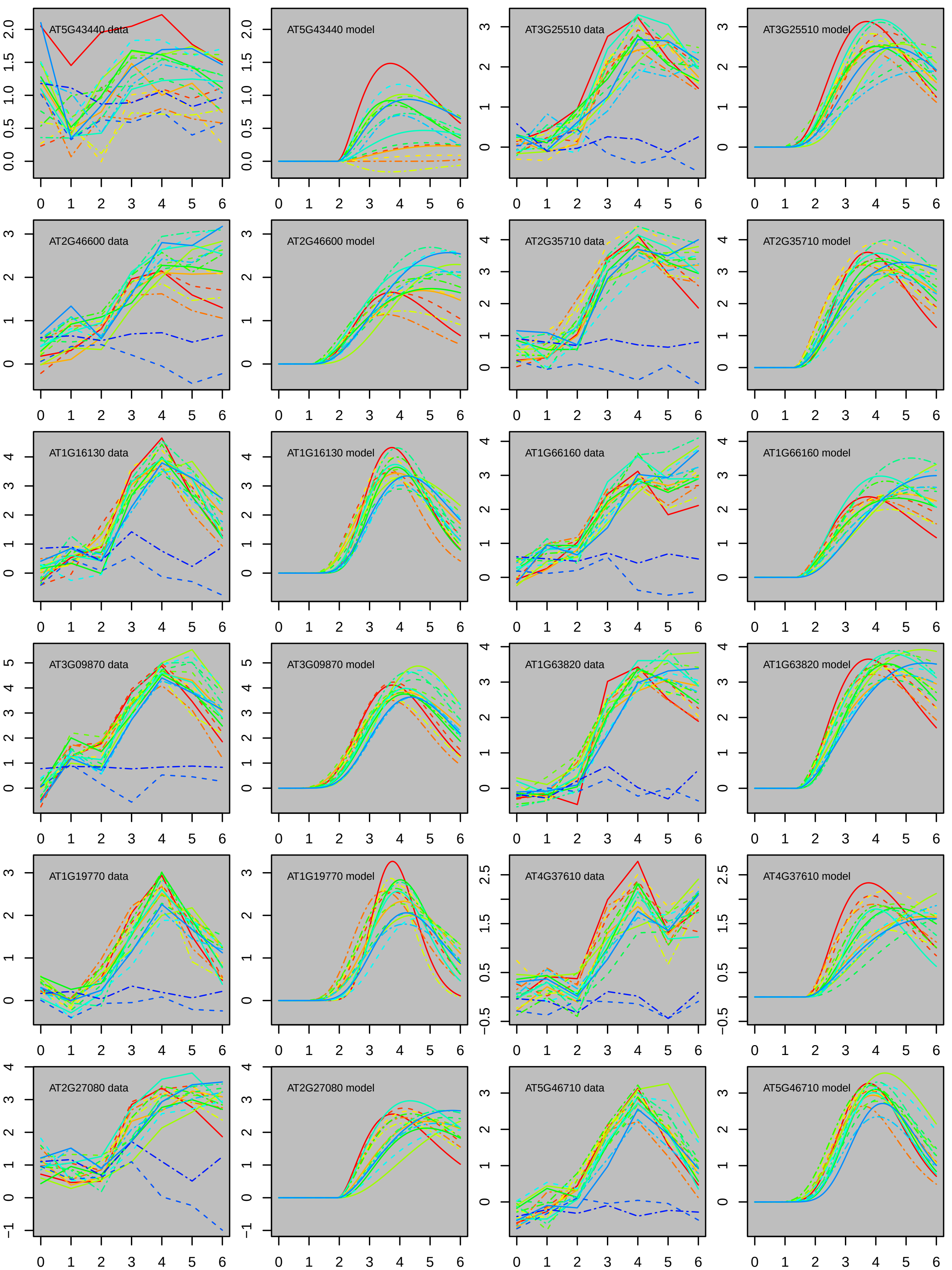
AT5G43440 data
AT5G43440 model
AT3G25510 data
AT3G25510 model
AT2G46600 data
AT2G46600 model
AT2G35710 data
AT2G35710 model
AT1G16130 data
AT1G16130 model
AT1G66160 data
AT1G66160 model
AT3G09870 data
AT3G09870 model
AT1G63820 data
AT1G63820 model
AT1G19770 data
AT1G19770 model
AT4G37610 data
AT4G37610 model
AT2G27080 data
AT2G27080 model
AT5G46710 data
AT5G46710 model

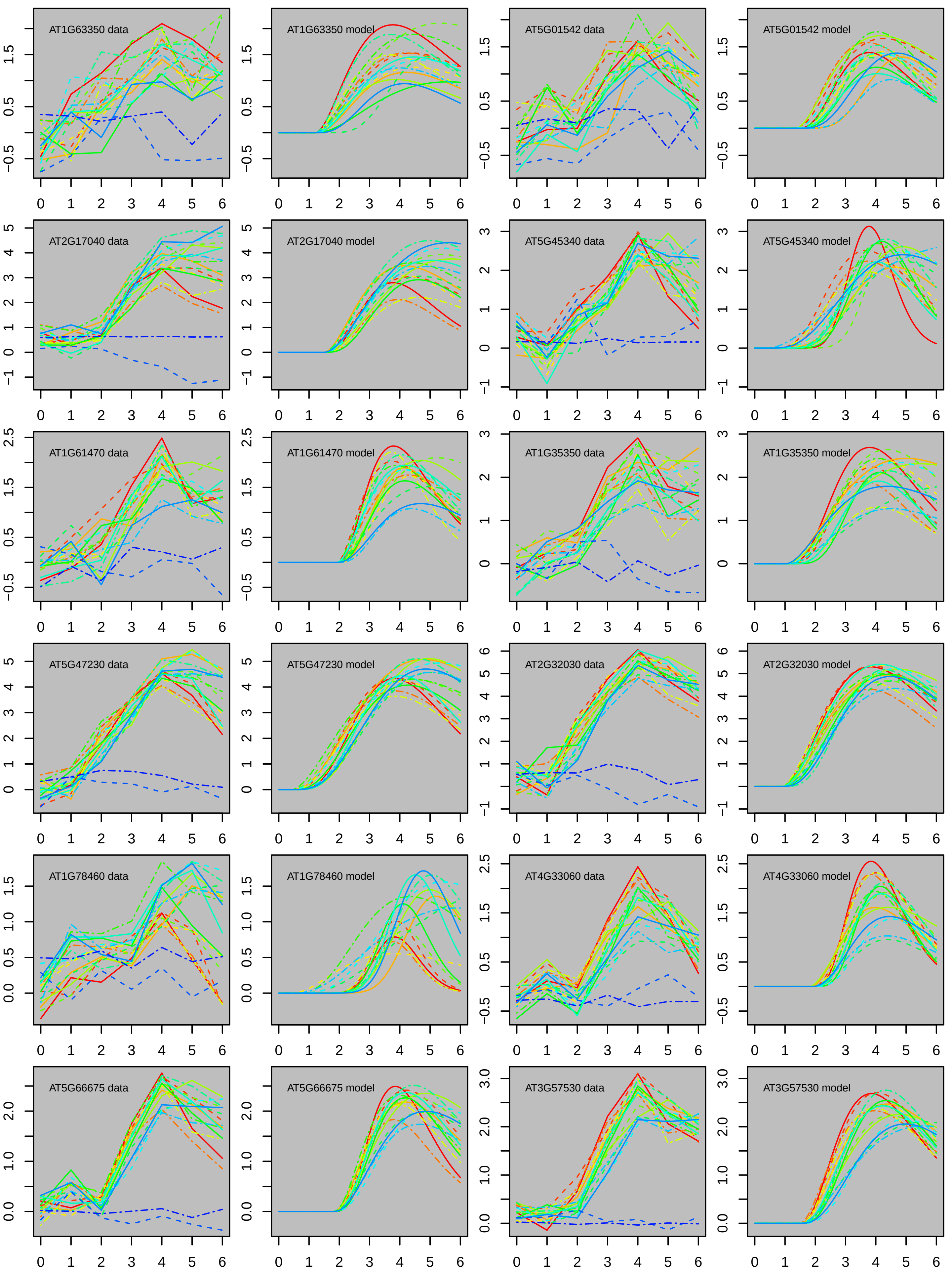
AT1G63350 data
AT1G63350 model
AT5G01542 data
AT5G01542 model
AT2G17040 data
AT2G17040 model
AT5G45340 data
AT5G45340 model
AT1G61470 data
AT1G61470 model
AT1G35350 data
AT1G35350 model
AT5G47230 data
AT5G47230 model
AT2G32030 data
AT2G32030 model
AT1G78460 data
AT1G78460 model
AT4G33060 data
AT4G33060 model
AT5G66675 data
AT5G66675 model
AT3G57530 data
AT3G57530 model

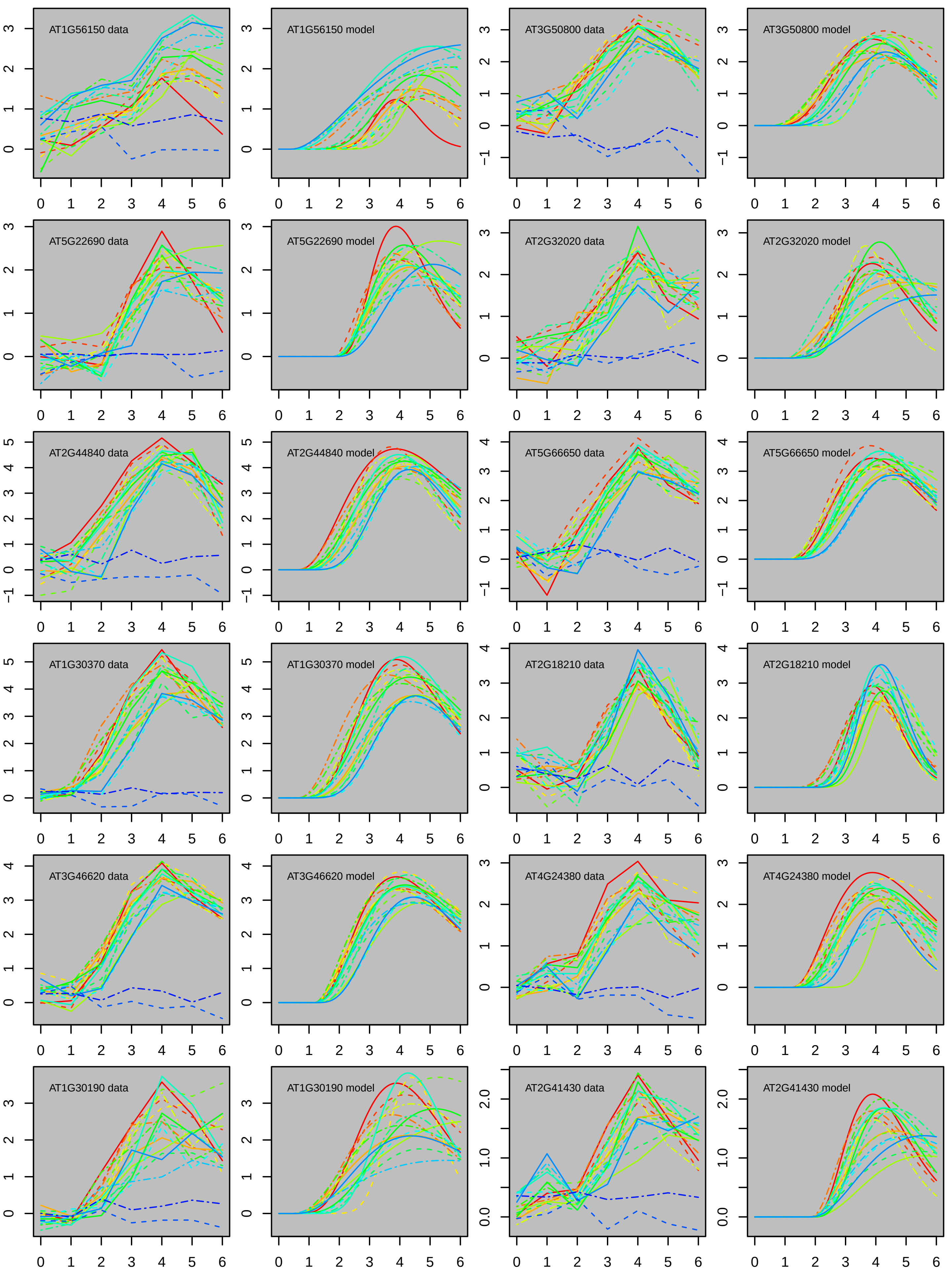
AT1G56150 data
AT1G56150 model
AT3G50800 data
AT3G50800 model
AT5G22690 data
AT5G22690 model
AT2G32020 data
AT2G32020 model
AT2G44840 data
AT2G44840 model
AT5G66650 data
AT5G66650 model
AT1G30370 data
AT1G30370 model
AT2G18210 data
AT2G18210 model
AT3G46620 data
AT3G46620 model
AT4G24380 data
AT4G24380 model
AT1G30190 data
AT1G30190 model
AT2G41430 data
AT2G41430 model

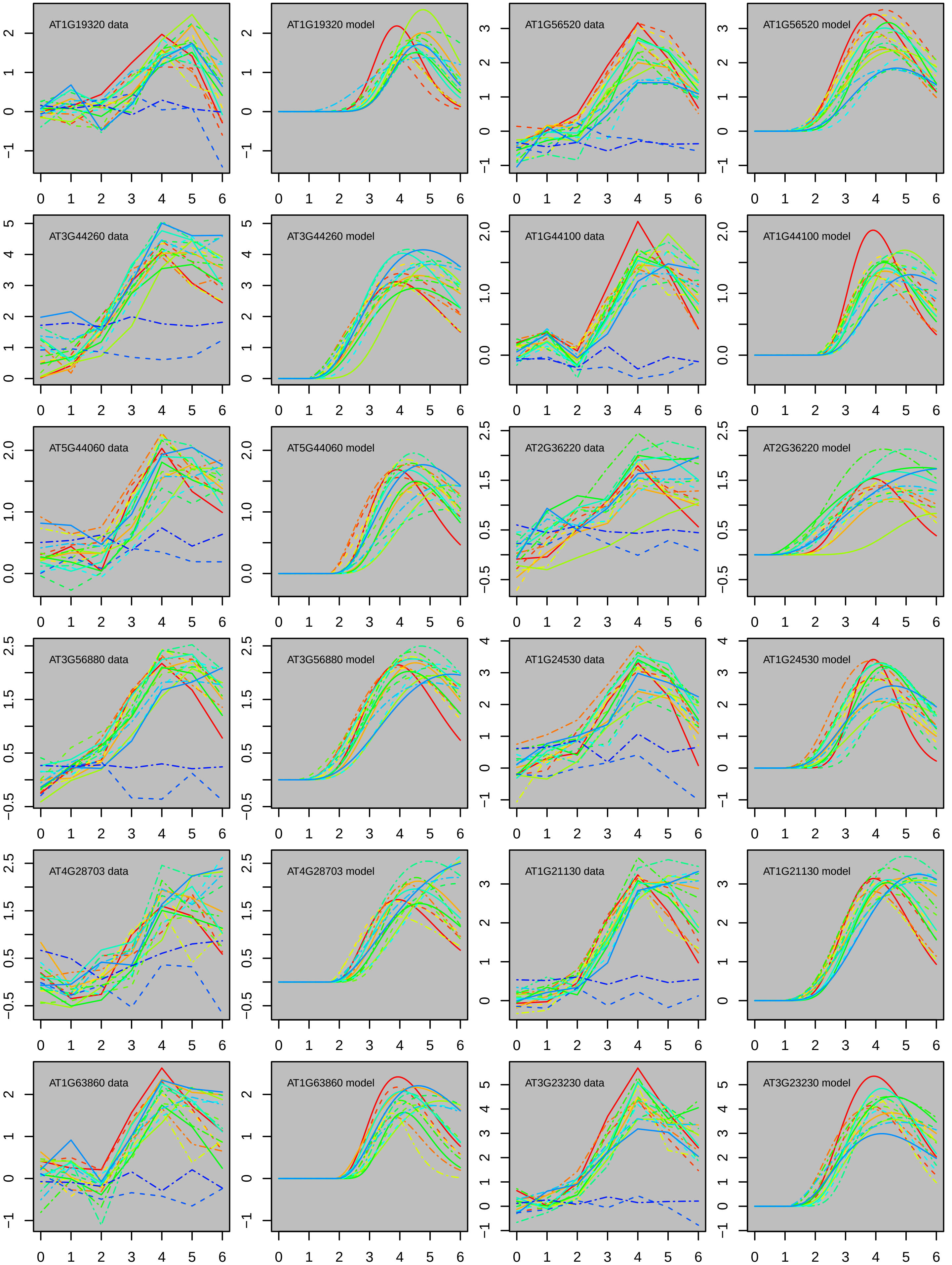
AT1G19320 data
AT1G19320 model
AT1G56520 data
AT1G56520 model
AT3G44260 data
AT3G44260 model
AT1G44100 data
AT1G44100 model
AT5G44060 data
AT5G44060 model
AT2G36220 data
AT2G36220 model
AT3G56880 data
AT3G56880 model
AT1G24530 data
AT1G24530 model
AT4G28703 data
AT4G28703 model
AT1G21130 data
AT1G21130 model
AT1G63860 data
AT1G63860 model
AT3G23230 data
AT3G23230 model

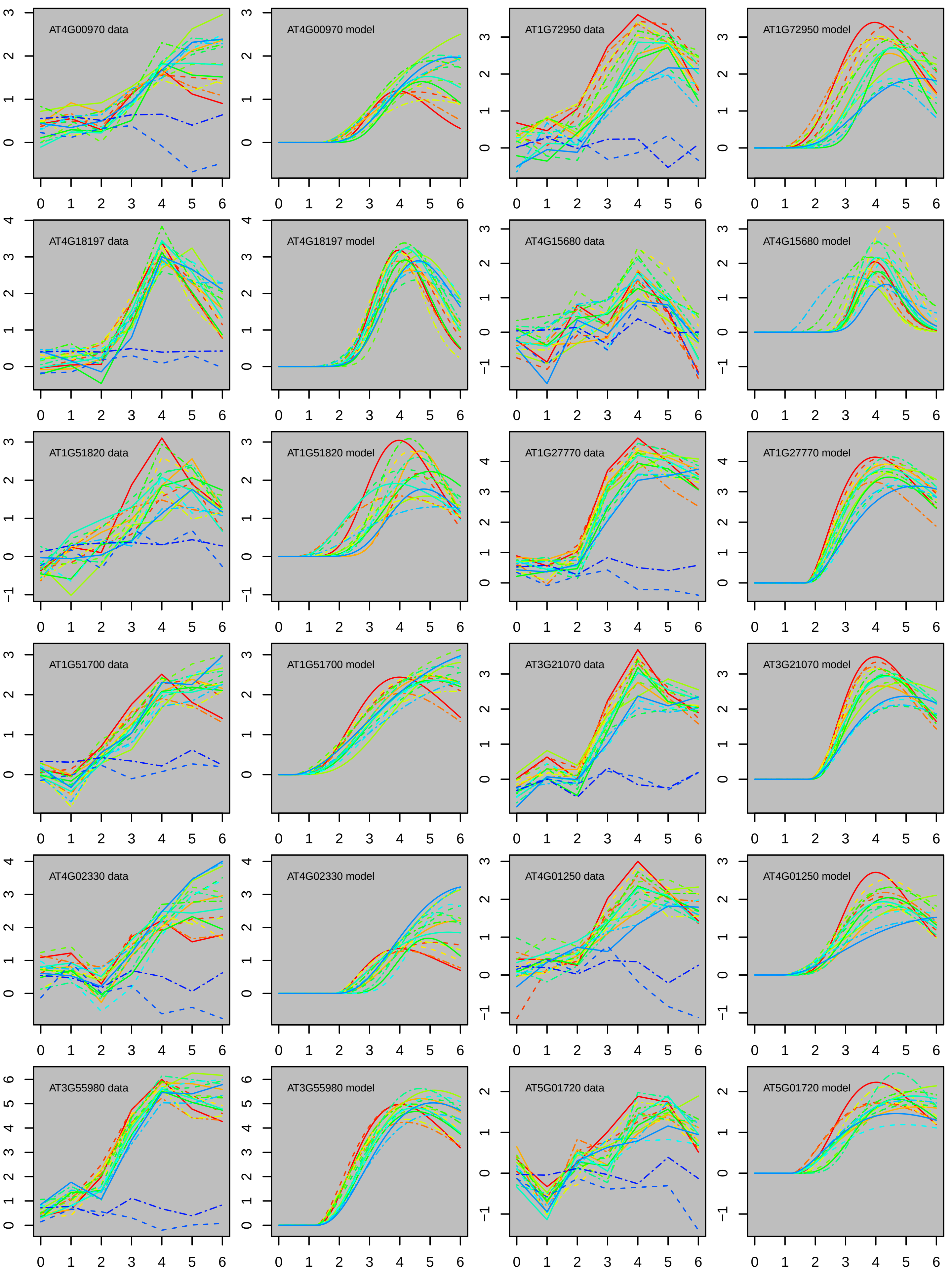
AT4G00970 data
AT4G00970 model
AT1G72950 data
AT1G72950 model
AT4G18197 data
AT4G18197 model
AT4G15680 data
AT4G15680 model
AT1G51820 data
AT1G51820 model
AT1G27770 data
AT1G27770 model
AT1G51700 data
AT1G51700 model
AT3G21070 data
AT3G21070 model
AT4G02330 data
AT4G02330 model
AT4G01250 data
AT4G01250 model
AT3G55980 data
AT3G55980 model
AT5G01720 data
AT5G01720 model

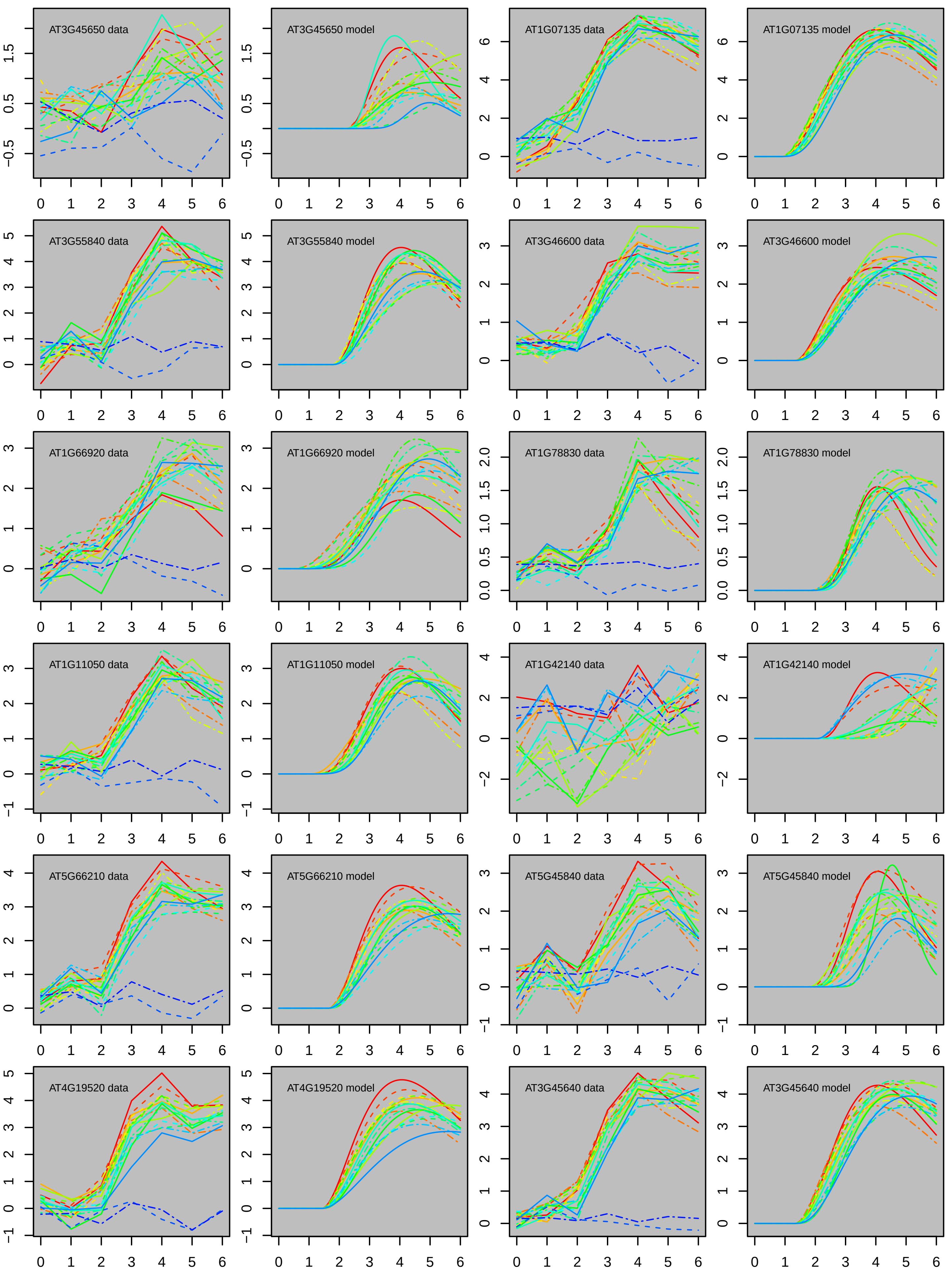
AT3G45650 data
AT3G45650 model
AT1G07135 data
AT1G07135 model
AT3G55840 data
AT3G55840 model
AT3G46600 data
AT3G46600 model
AT1G66920 data
AT1G66920 model
AT1G78830 data
AT1G78830 model
AT1G11050 data
AT1G11050 model
AT1G42140 data
AT1G42140 model
AT5G66210 data
AT5G66210 model
AT5G45840 data
AT5G45840 model
AT4G19520 data
AT4G19520 model
AT3G45640 data
AT3G45640 model

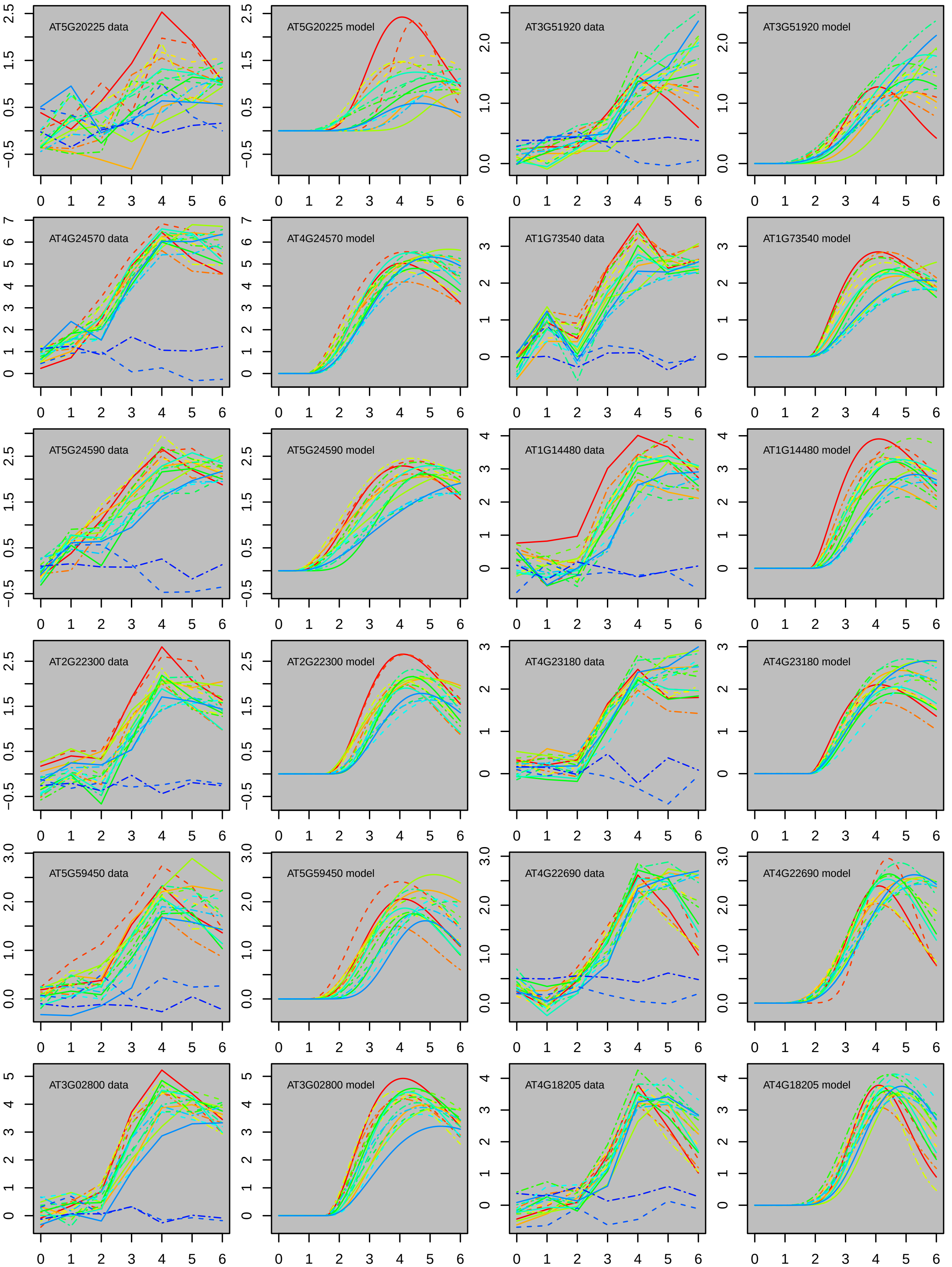
AT5G20225 data
AT5G20225 model
AT3G51920 data
AT3G51920 model
AT4G24570 data
AT4G24570 model
AT1G73540 data
AT1G73540 model
AT5G24590 data
AT5G24590 model
AT1G14480 data
AT1G14480 model
AT2G22300 data
AT2G22300 model
AT4G23180 data
AT4G23180 model
AT5G59450 data
AT5G59450 model
AT4G22690 data
AT4G22690 model
AT3G02800 data
AT3G02800 model
AT4G18205 data
AT4G18205 model

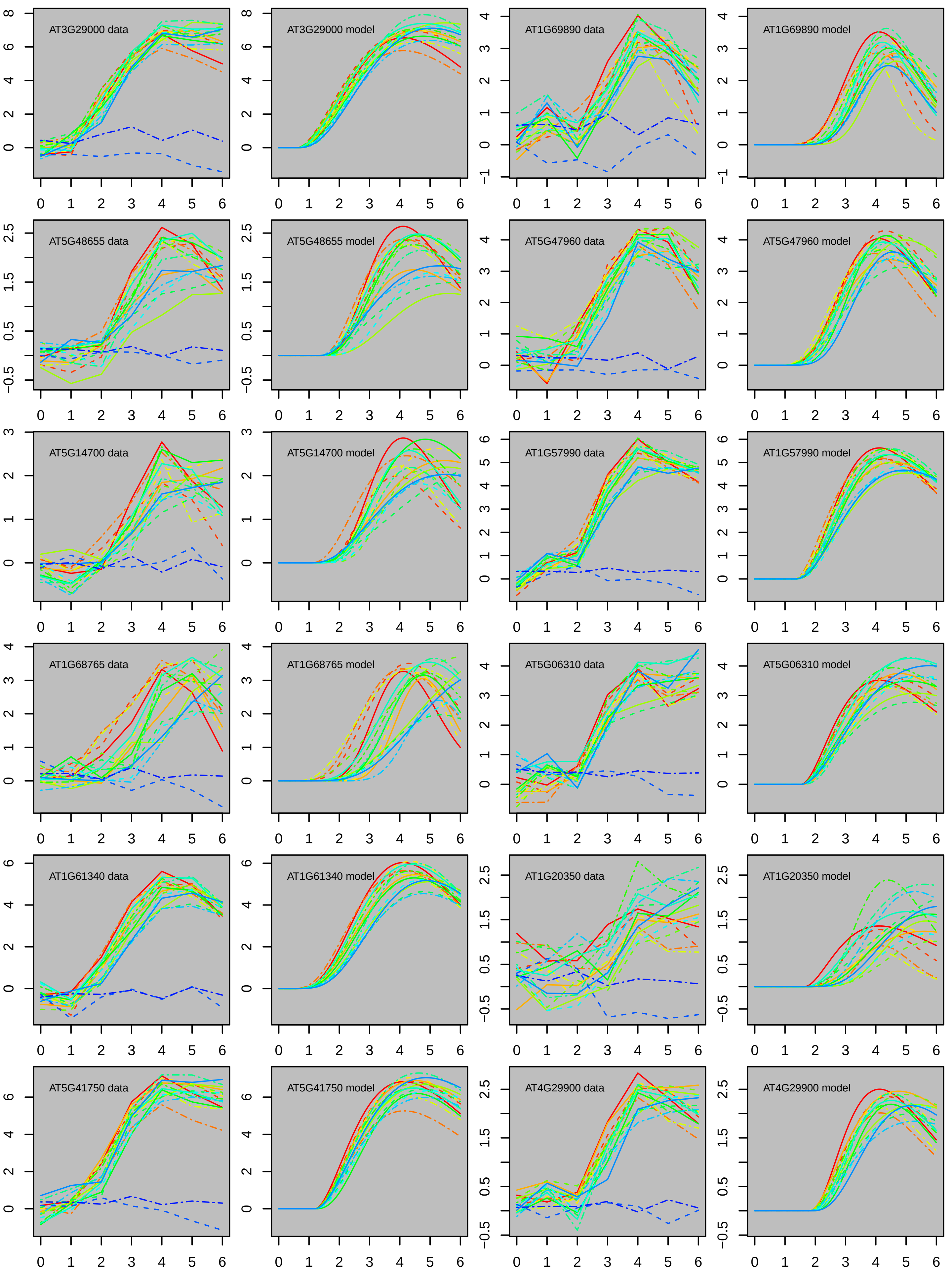
AT3G29000 data
AT3G29000 model
AT1G69890 data
AT1G69890 model
AT5G48655 data
AT5G48655 model
AT5G47960 data
AT5G47960 model
AT5G14700 data
AT5G14700 model
AT1G57990 data
AT1G57990 model
AT1G68765 data
AT1G68765 model
AT5G06310 data
AT5G06310 model
AT1G61340 data
AT1G61340 model
AT1G20350 data
AT1G20350 model
AT5G41750 data
AT5G41750 model
AT4G29900 data
AT4G29900 model

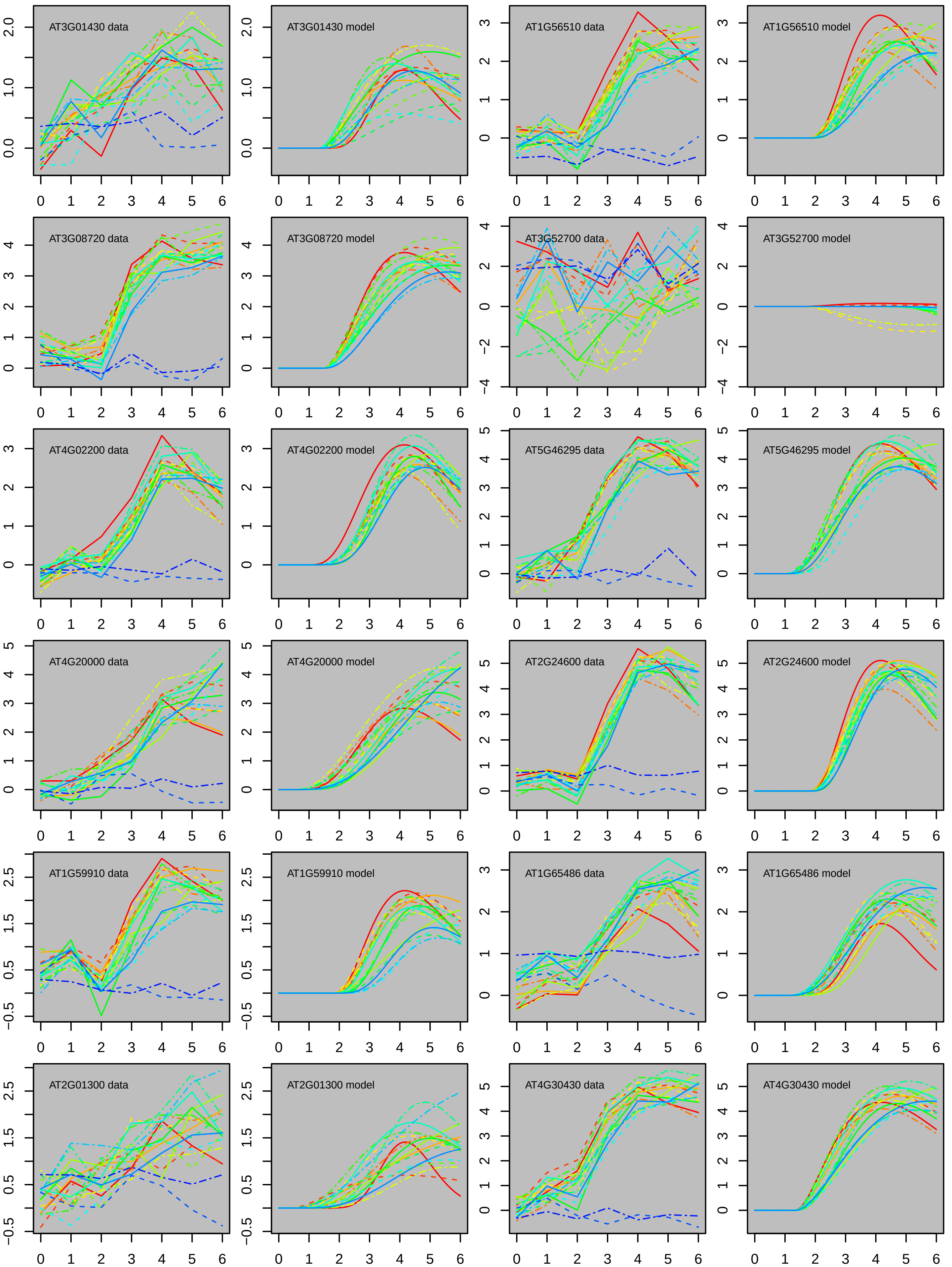
AT3G01430 data
AT3G01430 model
AT1G56510 data
AT1G56510 model
AT3G08720 data
AT3G08720 model
AT3G52700 data
AT3G52700 model
AT4G02200 data
AT4G02200 model
AT5G46295 data
AT5G46295 model
AT4G20000 data
AT4G20000 model
AT2G24600 data
AT2G24600 model
AT1G59910 data
AT1G59910 model
AT1G65486 data
AT1G65486 model
AT2G01300 data
AT2G01300 model
AT4G30430 data
AT4G30430 model

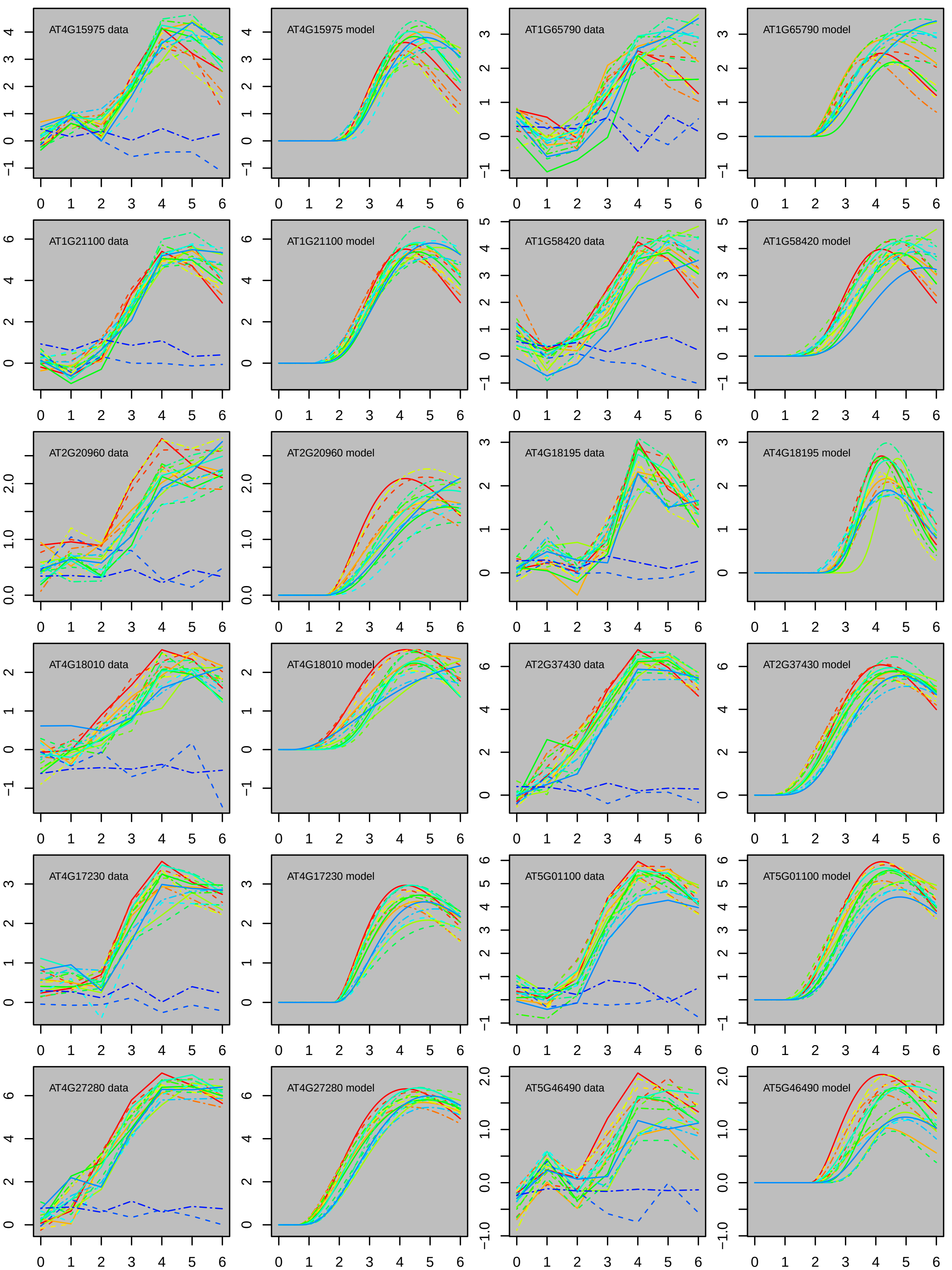
AT4G15975 data
AT4G15975 model
AT1G65790 data
AT1G65790 model
AT1G21100 data
AT1G21100 model
AT1G58420 data
AT1G58420 model
AT2G20960 data
AT2G20960 model
AT4G18195 data
AT4G18195 model
AT4G18010 data
AT4G18010 model
AT2G37430 data
AT2G37430 model
AT4G17230 data
AT4G17230 model
AT5G01100 data
AT5G01100 model
AT4G27280 data
AT4G27280 model
AT5G46490 data
AT5G46490 model

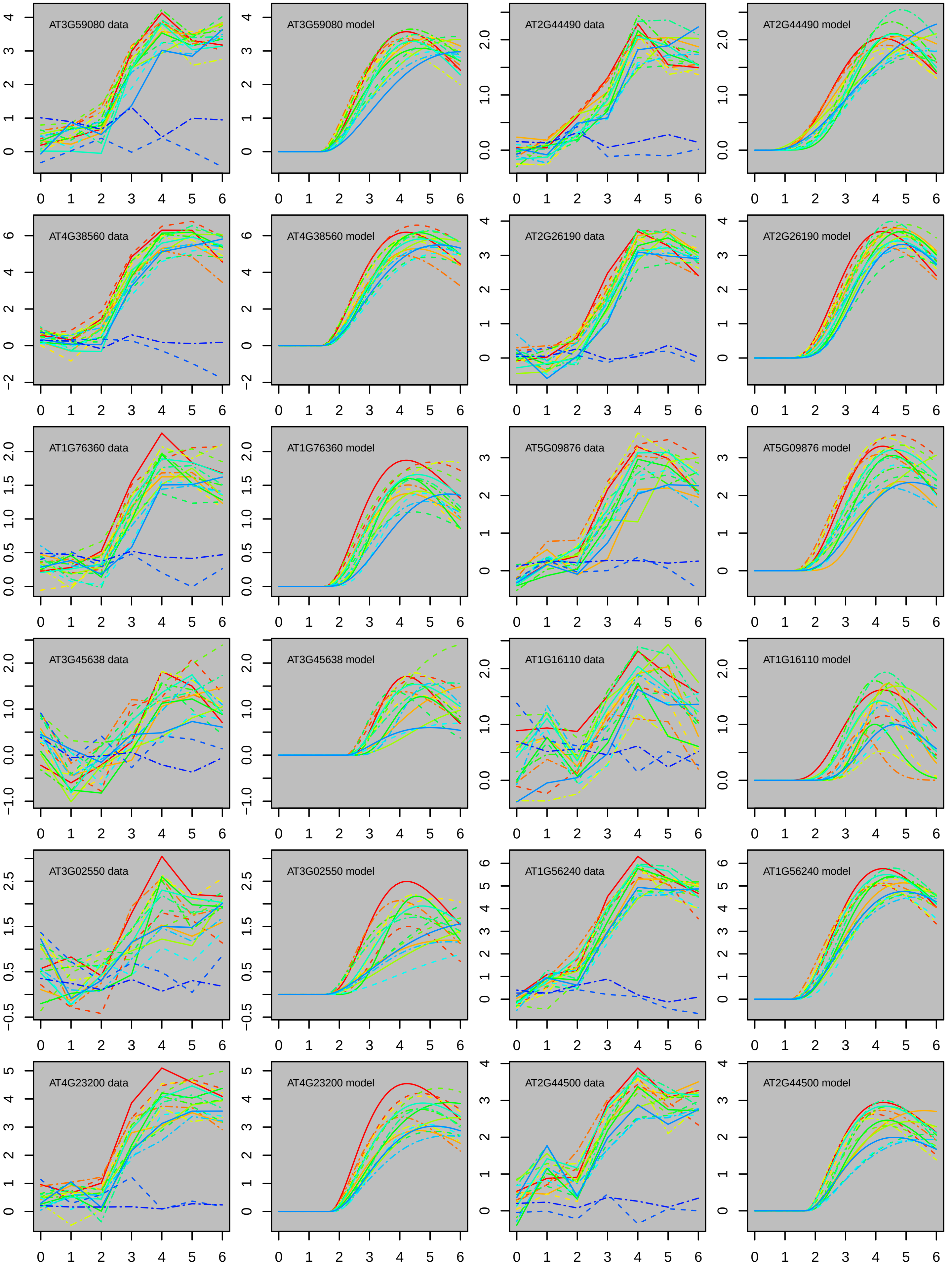
AT3G59080 data
AT3G59080 model
AT2G44490 data
AT2G44490 model
AT4G38560 data
AT4G38560 model
AT2G26190 data
AT2G26190 model
AT1G76360 data
AT1G76360 model
AT5G09876 data
AT5G09876 model
AT3G45638 data
AT3G45638 model
AT1G16110 data
AT1G16110 model
AT3G02550 data
AT3G02550 model
AT1G56240 data
AT1G56240 model
AT4G23200 data
AT4G23200 model
AT2G44500 data
AT2G44500 model

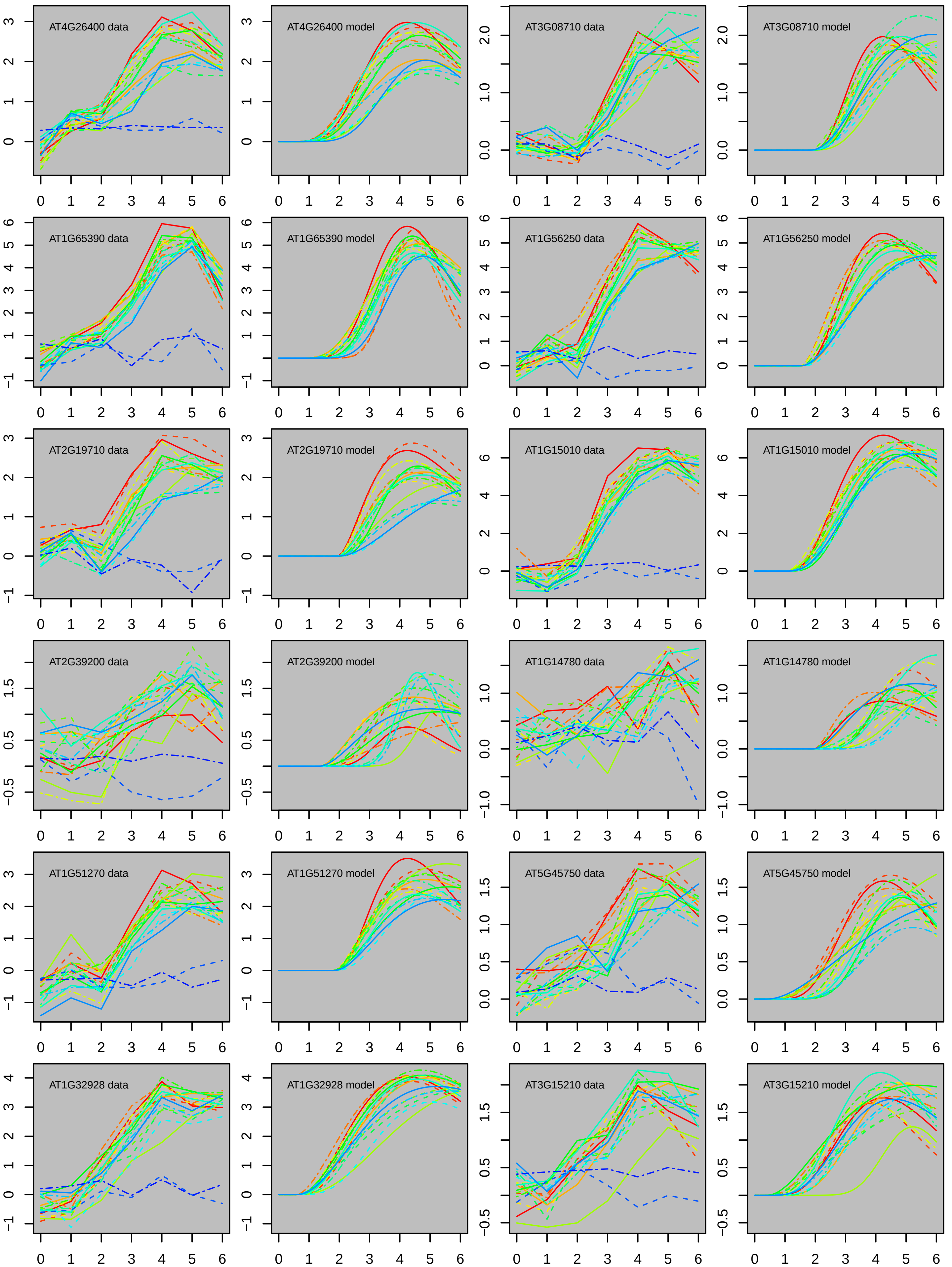
AT4G26400 data
AT4G26400 model
AT3G08710 data
AT3G08710 model
AT1G65390 data
AT1G65390 model
AT1G56250 data
AT1G56250 model
AT2G19710 data
AT2G19710 model
AT1G15010 data
AT1G15010 model
AT2G39200 data
AT2G39200 model
AT1G14780 data
AT1G14780 model
AT1G51270 data
AT1G51270 model
AT5G45750 data
AT5G45750 model
AT1G32928 data
AT1G32928 model
AT3G15210 data
AT3G15210 model

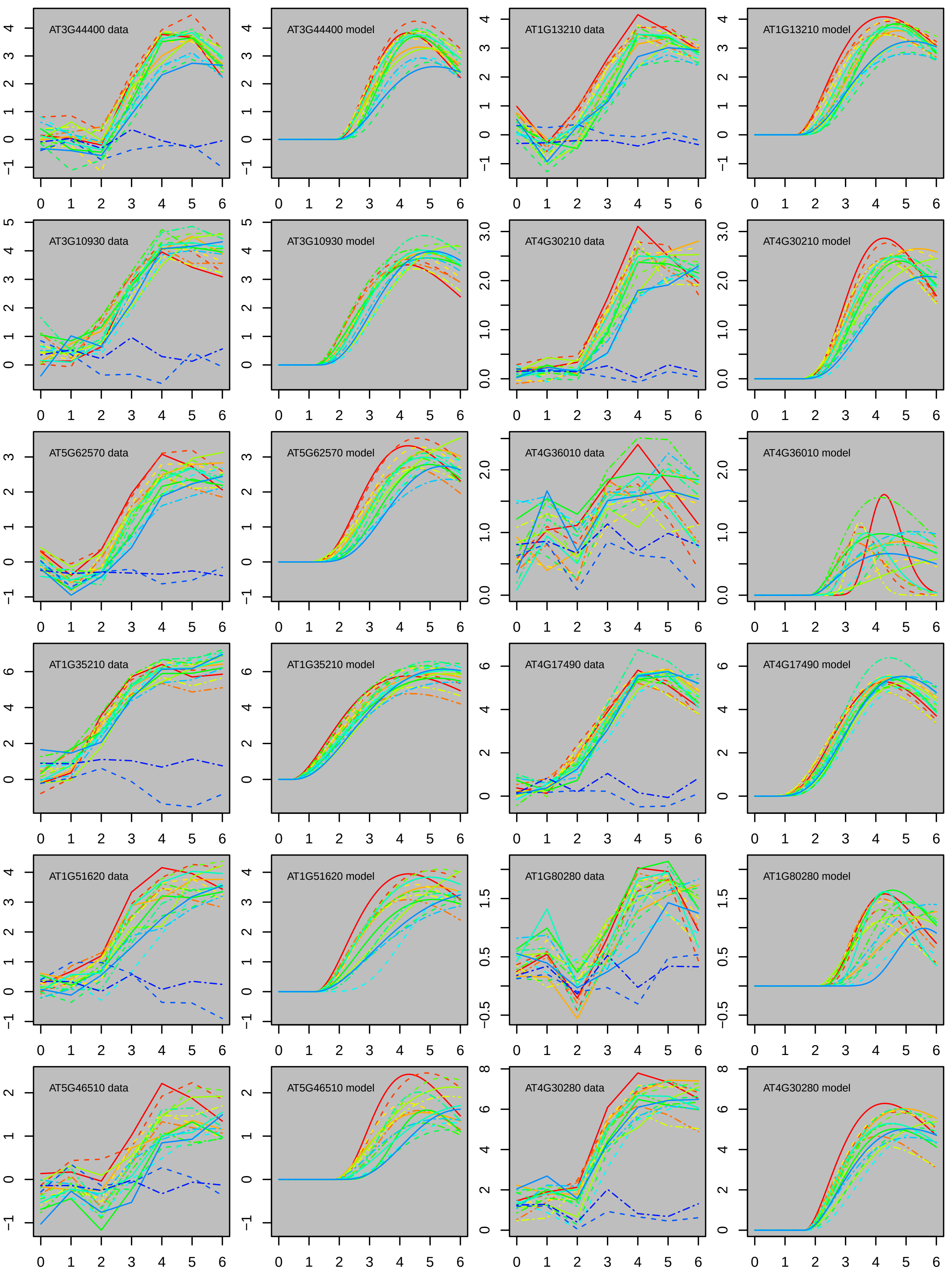
AT3G44400 data
AT3G44400 model
AT1G13210 data
AT1G13210 model
AT3G10930 data
AT3G10930 model
AT4G30210 data
AT4G30210 model
AT5G62570 data
AT5G62570 model
AT4G36010 data
AT4G36010 model
AT1G35210 data
AT1G35210 model
AT4G17490 data
AT4G17490 model
AT1G51620 data
AT1G51620 model
AT1G80280 data
AT1G80280 model
AT5G46510 data
AT5G46510 model
AT4G30280 data
AT4G30280 model

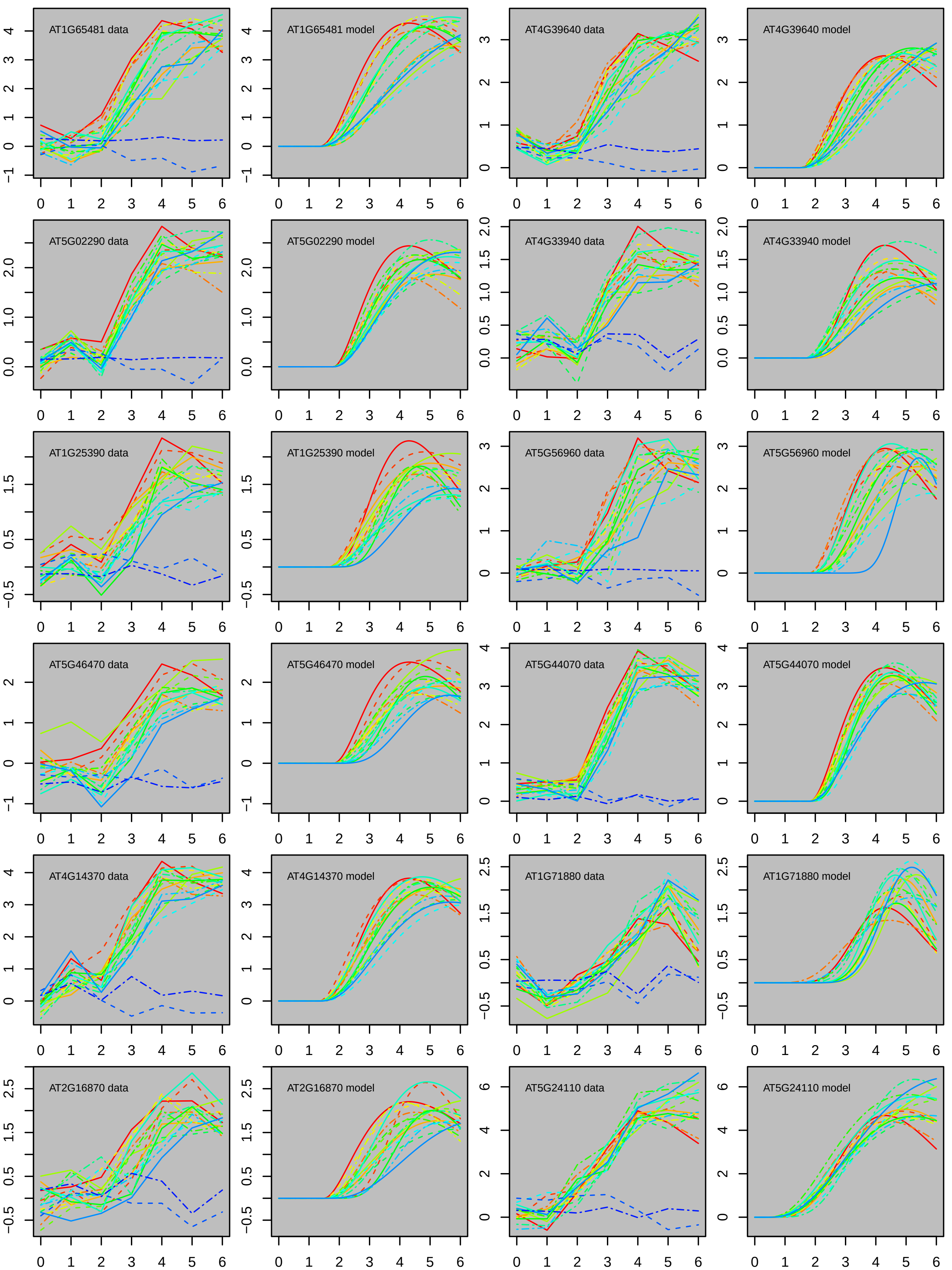
AT1G65481 data
AT1G65481 model
AT4G39640 data
AT4G39640 model
AT5G02290 data
AT5G02290 model
AT4G33940 data
AT4G33940 model
AT1G25390 data
AT1G25390 model
AT5G56960 data
AT5G56960 model
AT5G46470 data
AT5G46470 model
AT5G44070 data
AT5G44070 model
AT4G14370 data
AT4G14370 model
AT1G71880 data
AT1G71880 model
AT2G16870 data
AT2G16870 model
AT5G24110 data
AT5G24110 model

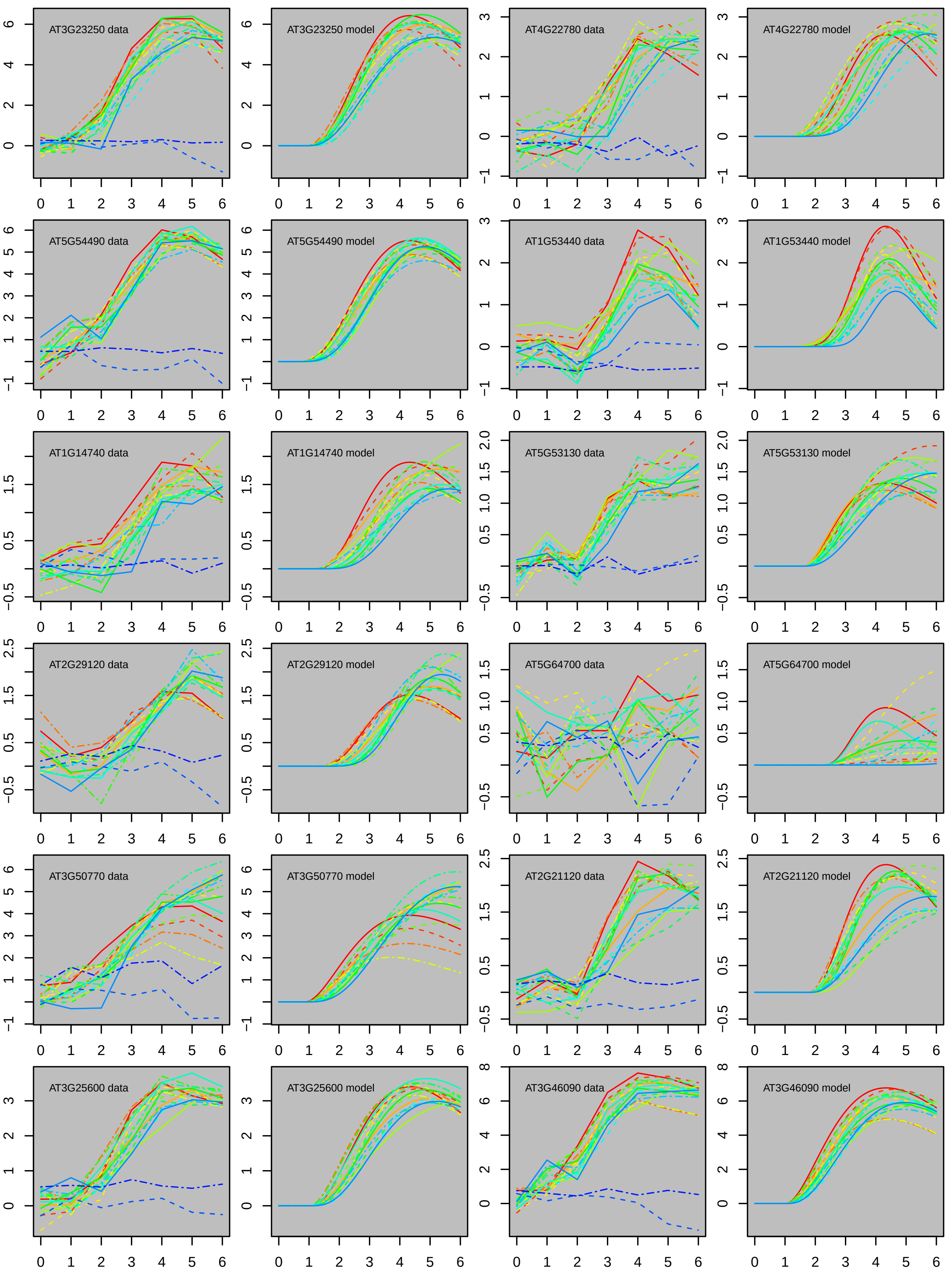

AT3G23250 data
AT3G23250 model
AT4G22780 data
AT4G22780 model
AT5G54490 data
AT5G54490 model
AT1G53440 data
AT1G53440 model
AT1G14740 data
AT1G14740 model
AT5G53130 data
AT5G53130 model
AT2G29120 data
AT2G29120 model
AT5G64700 data
AT5G64700 model
AT3G50770 data
AT3G50770 model
AT2G21120 data
AT2G21120 model
AT3G25600 data
AT3G25600 model
AT3G46090 data
AT3G46090 model

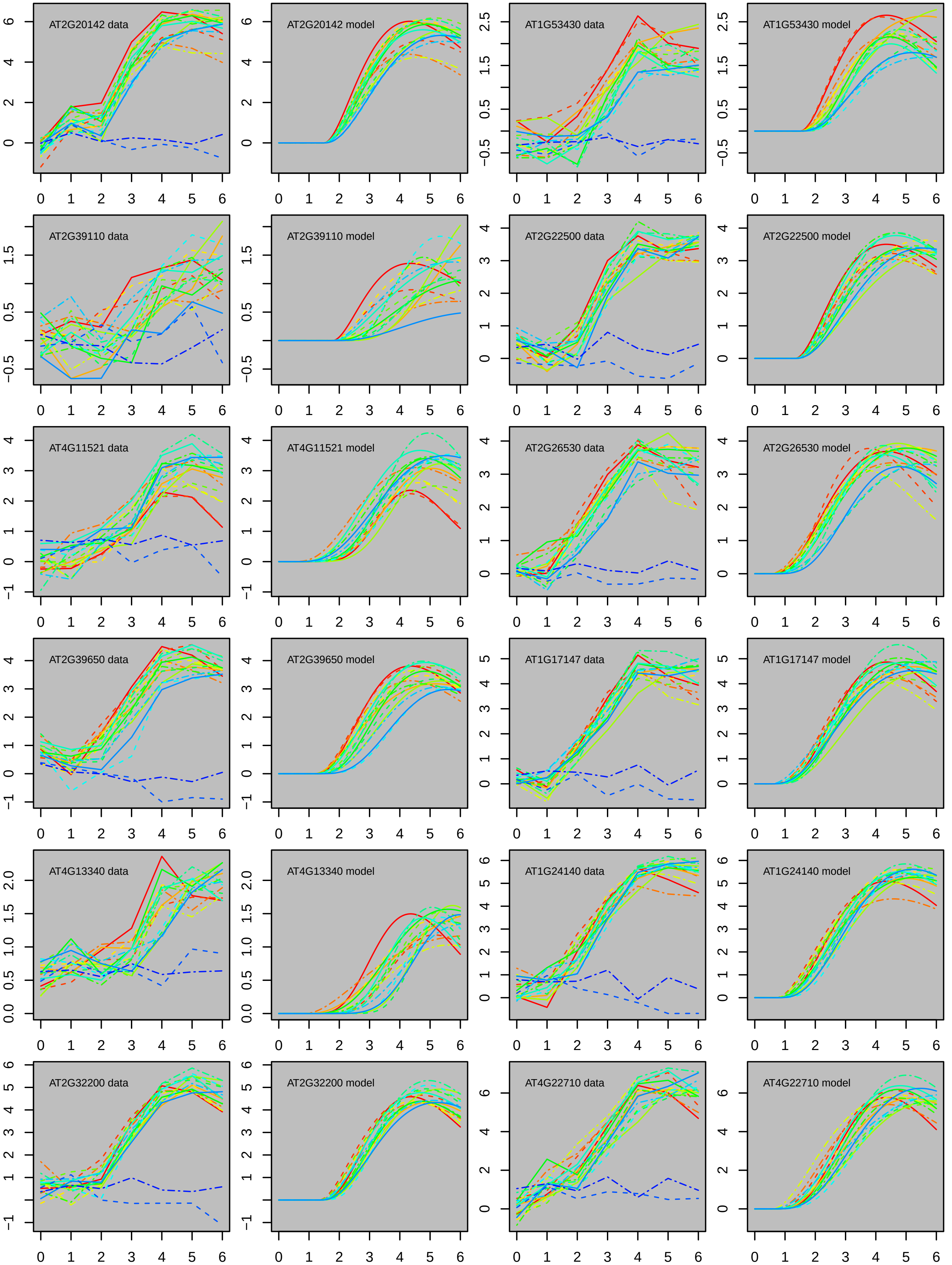

AT2G20142 data
AT2G20142 model
AT1G53430 data
AT1G53430 model
AT2G39110 data
AT2G39110 model
AT2G22500 data
AT2G22500 model
AT4G11521 data
AT4G11521 model
AT2G26530 data
AT2G26530 model
AT2G39650 data
AT2G39650 model
AT1G17147 data
AT1G17147 model
AT4G13340 data
AT4G13340 model
AT1G24140 data
AT1G24140 model
AT2G32200 data
AT2G32200 model
AT4G22710 data
AT4G22710 model

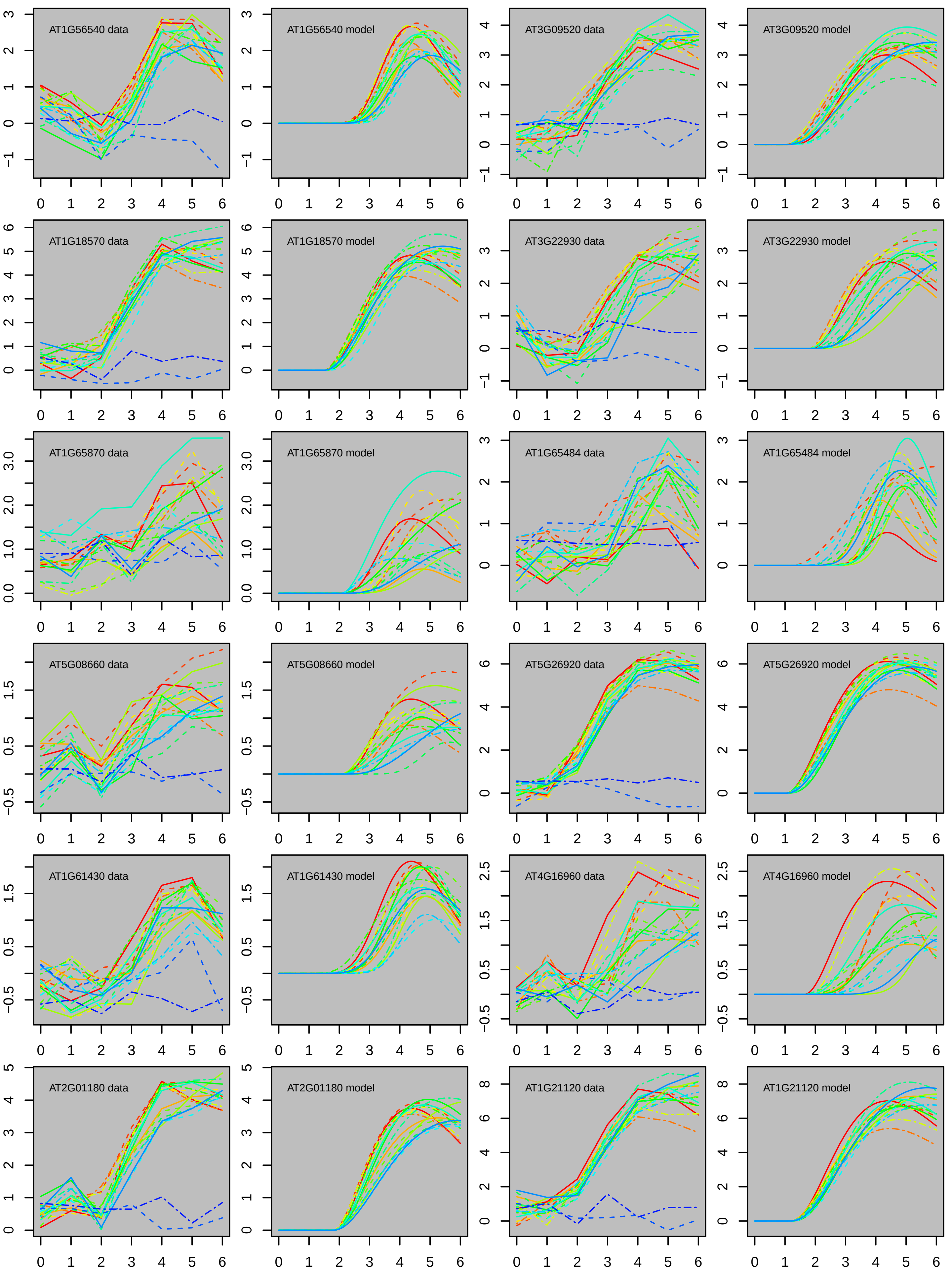
AT1G56540 data
AT1G56540 model
AT3G09520 data
AT3G09520 model
AT1G18570 data
AT1G18570 model
AT3G22930 data
AT3G22930 model
AT1G65870 data
AT1G65870 model
AT1G65484 data
AT1G65484 model
AT5G08660 data
AT5G08660 model
AT5G26920 data
AT5G26920 model
AT1G61430 data
AT1G61430 model
AT4G16960 data
AT4G16960 model
AT2G01180 data
AT2G01180 model
AT1G21120 data
AT1G21120 model

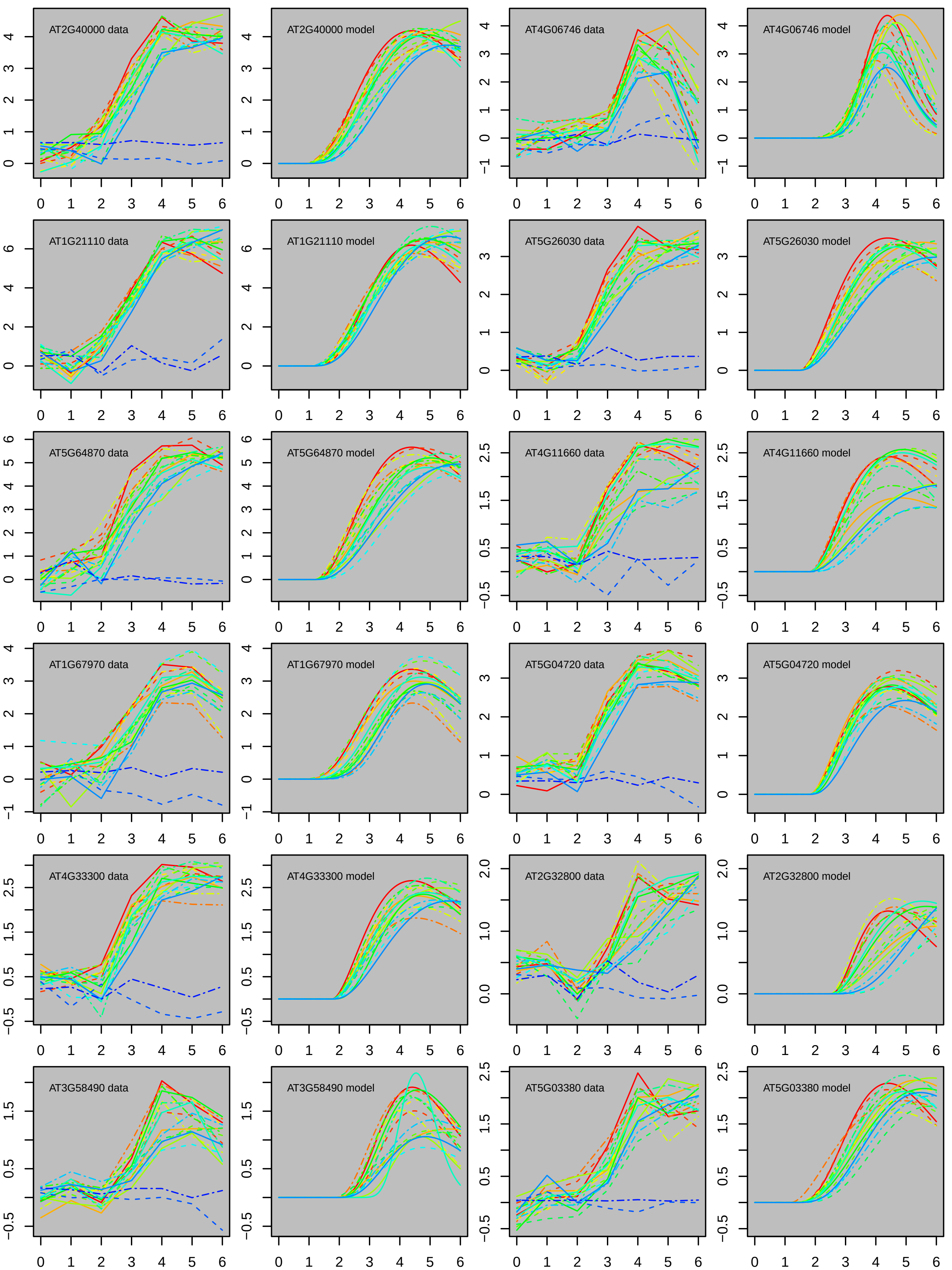
AT2G40000 data
AT2G40000 model
AT4G06746 data
AT4G06746 model
AT1G21110 data
AT1G21110 model
AT5G26030 data
AT5G26030 model
AT5G64870 data
AT5G64870 model
AT4G11660 data
AT4G11660 model
AT1G67970 data
AT1G67970 model
AT5G04720 data
AT5G04720 model
AT4G33300 data
AT4G33300 model
AT2G32800 data
AT2G32800 model
AT3G58490 data
AT3G58490 model
AT5G03380 data
AT5G03380 model

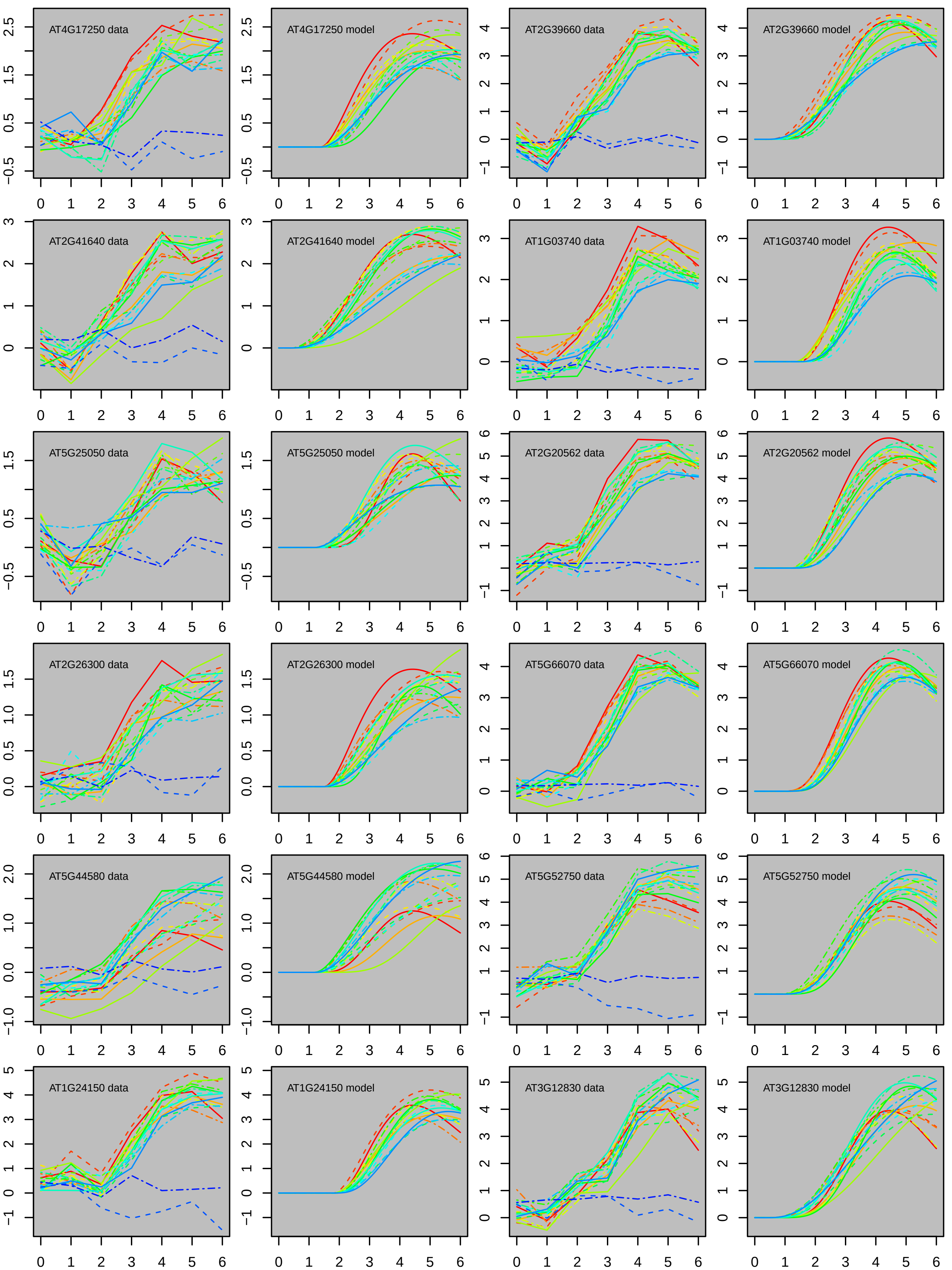

AT4G17250 data
AT4G17250 model
AT2G39660 data
AT2G39660 model
AT2G41640 data
AT2G41640 model
AT1G03740 data
AT1G03740 model
AT5G25050 data
AT5G25050 model
AT2G20562 data
AT2G20562 model
AT2G26300 data
AT2G26300 model
AT5G66070 data
AT5G66070 model
AT5G44580 data
AT5G44580 model
AT5G52750 data
AT5G52750 model
AT1G24150 data
AT1G24150 model
AT3G12830 data
AT3G12830 model

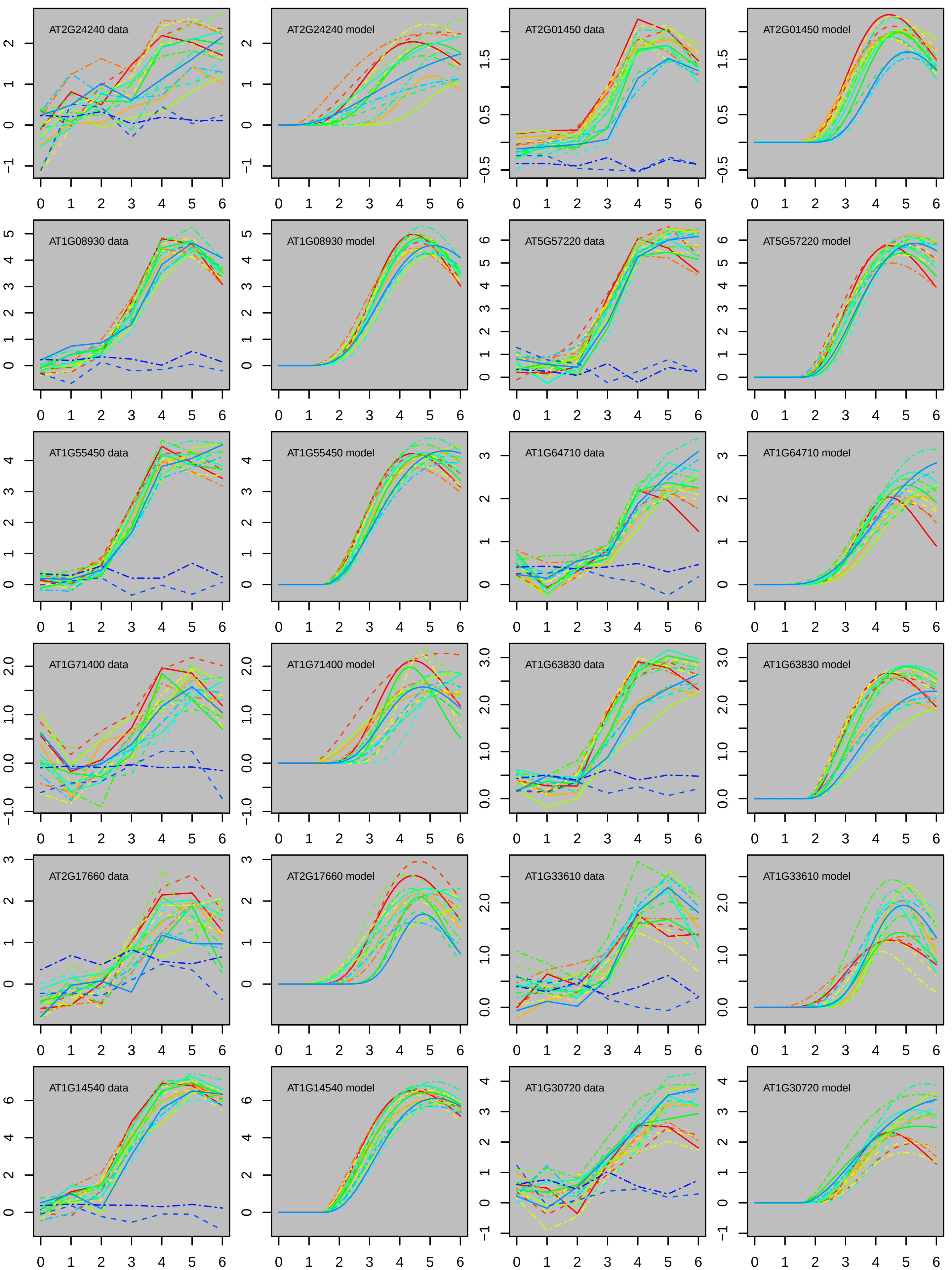

AT2G24240 data
AT2G24240 model
AT2G01450 data
AT2G01450 model
AT1G08930 data
AT1G08930 model
AT5G57220 data
AT5G57220 model
AT1G55450 data
AT1G55450 model
AT1G64710 data
AT1G64710 model
AT1G71400 data
AT1G71400 model
AT1G63830 data
AT1G63830 model
AT2G17660 data
AT2G17660 model
AT1G33610 data
AT1G33610 model
AT1G14540 data
AT1G14540 model
AT1G30720 data
AT1G30720 model

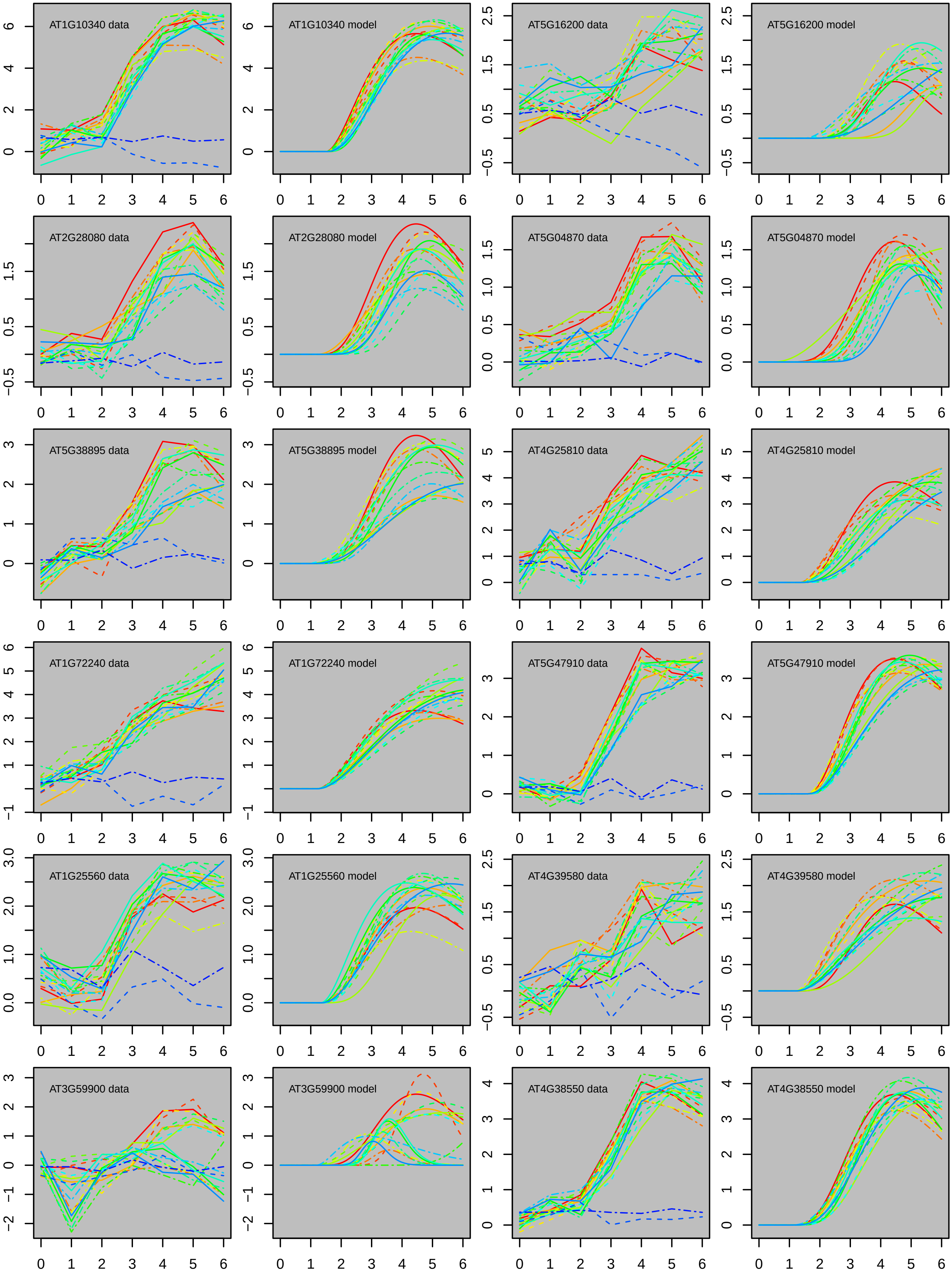

AT1G10340 data
AT1G10340 model
AT5G16200 data
AT5G16200 model
AT2G28080 data
AT2G28080 model
AT5G04870 data
AT5G04870 model
AT5G38895 data
AT5G38895 model
AT4G25810 data
AT4G25810 model
AT1G72240 data
AT1G72240 model
AT5G47910 data
AT5G47910 model
AT1G25560 data
AT1G25560 model
AT4G39580 data
AT4G39580 model
AT3G59900 data
AT3G59900 model
AT4G38550 data
AT4G38550 model

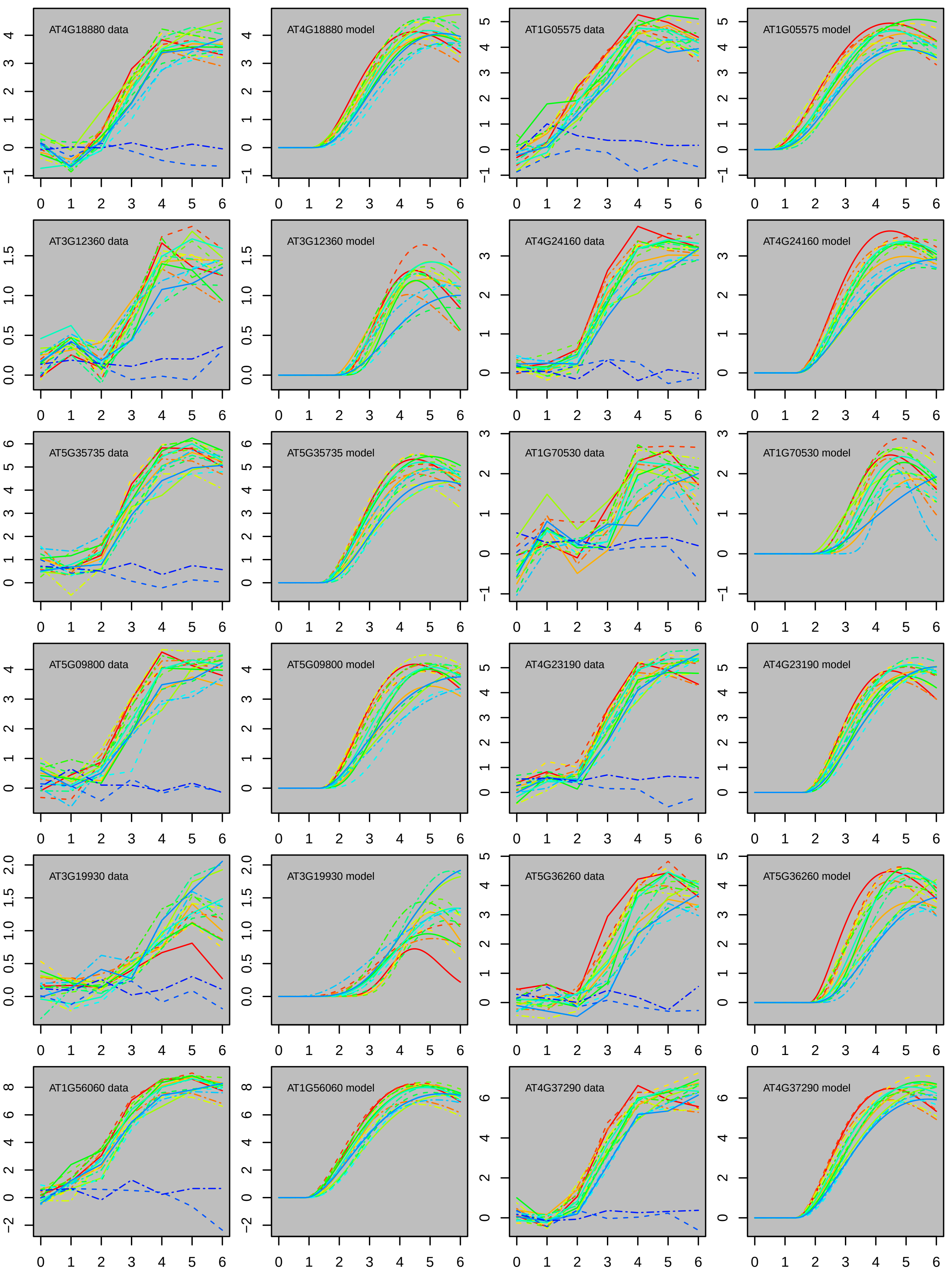
AT4G18880 data
AT4G18880 model
AT1G05575 data
AT1G05575 model
AT3G12360 data
AT3G12360 model
AT4G24160 data
AT4G24160 model
AT5G35735 data
AT5G35735 model
AT1G70530 data
AT1G70530 model
AT5G09800 data
AT5G09800 model
AT4G23190 data
AT4G23190 model
AT3G19930 data
AT3G19930 model
AT5G36260 data
AT5G36260 model
AT1G56060 data
AT1G56060 model
AT4G37290 data
AT4G37290 model

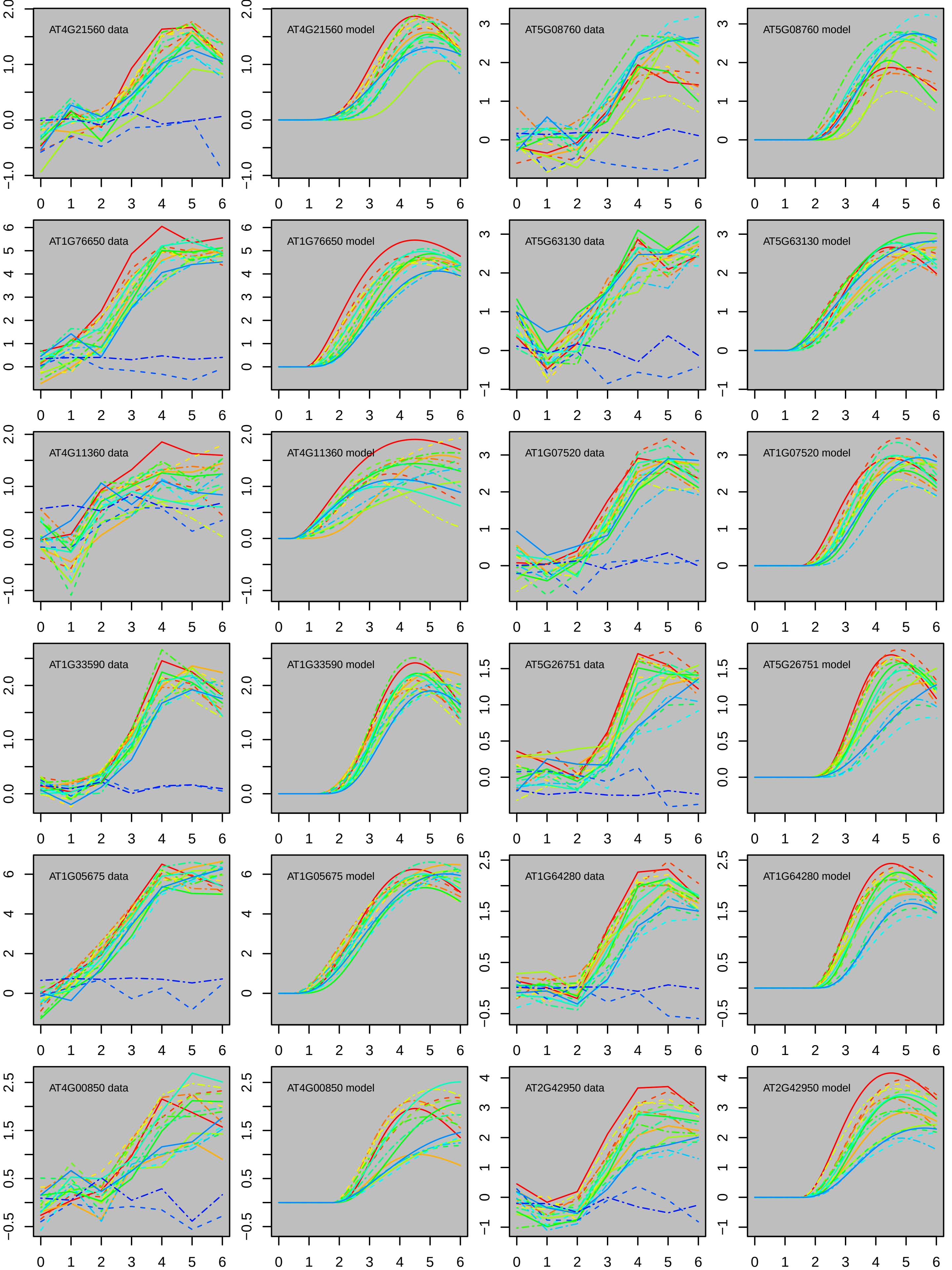

AT4G21560 data
AT4G21560 model
AT5G08760 data
AT5G08760 model
AT1G76650 data
AT1G76650 model
AT5G63130 data
AT5G63130 model
AT4G11360 data
AT4G11360 model
AT1G07520 data
AT1G07520 model
AT1G33590 data
AT1G33590 model
AT5G26751 data
AT5G26751 model
AT1G05675 data
AT1G05675 model
AT1G64280 data
AT1G64280 model
AT4G00850 data
AT4G00850 model
AT2G42950 data
AT2G42950 model

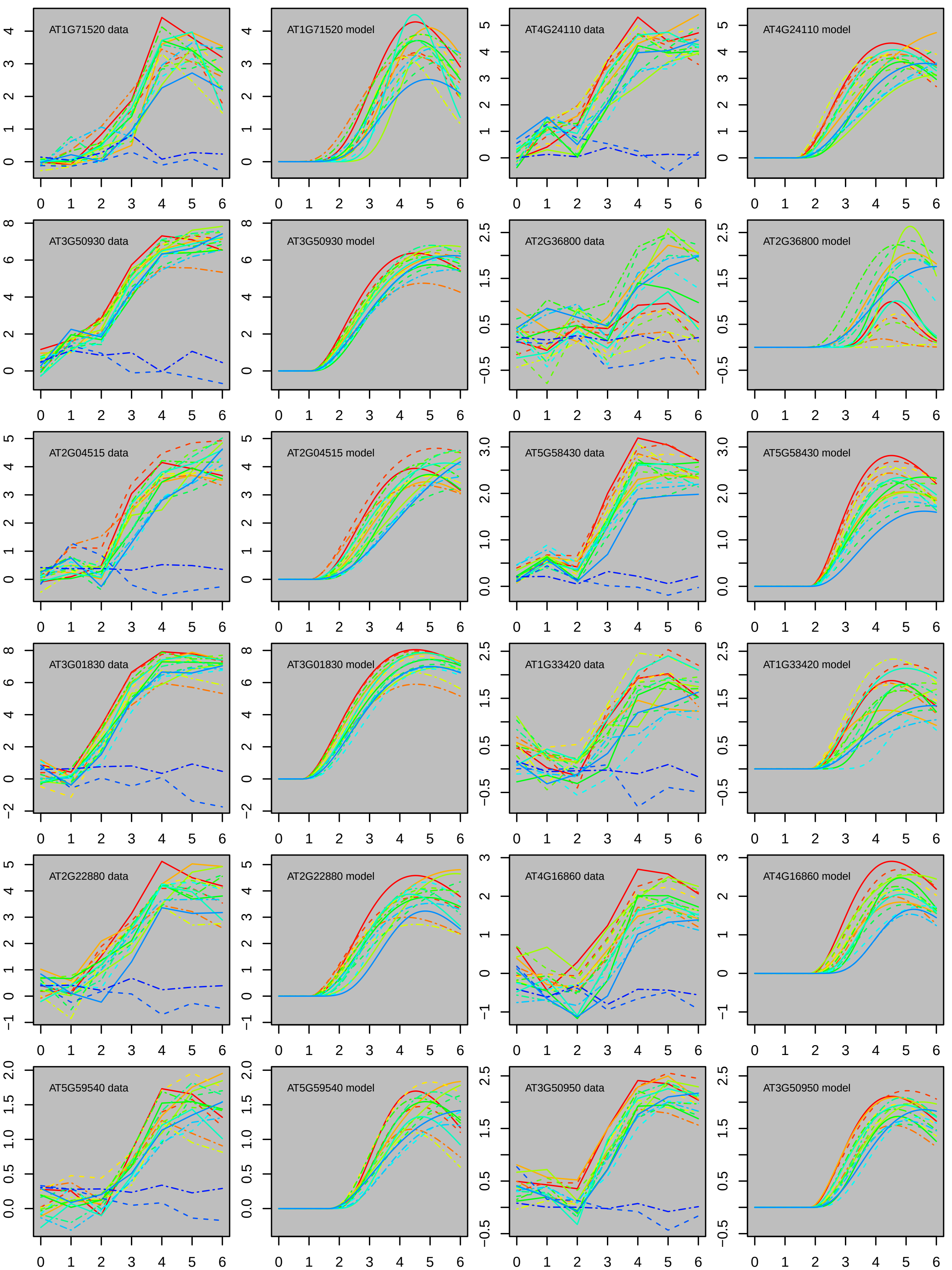
AT1G71520 data
AT1G71520 model
AT4G24110 data
AT4G24110 model
AT3G50930 data
AT3G50930 model
AT2G36800 data
AT2G36800 model
AT2G04515 data
AT2G04515 model
AT5G58430 data
AT5G58430 model
AT3G01830 data
AT3G01830 model
AT1G33420 data
AT1G33420 model
AT2G22880 data
AT2G22880 model
AT4G16860 data
AT4G16860 model
AT5G59540 data
AT5G59540 model
AT3G50950 data
AT3G50950 model

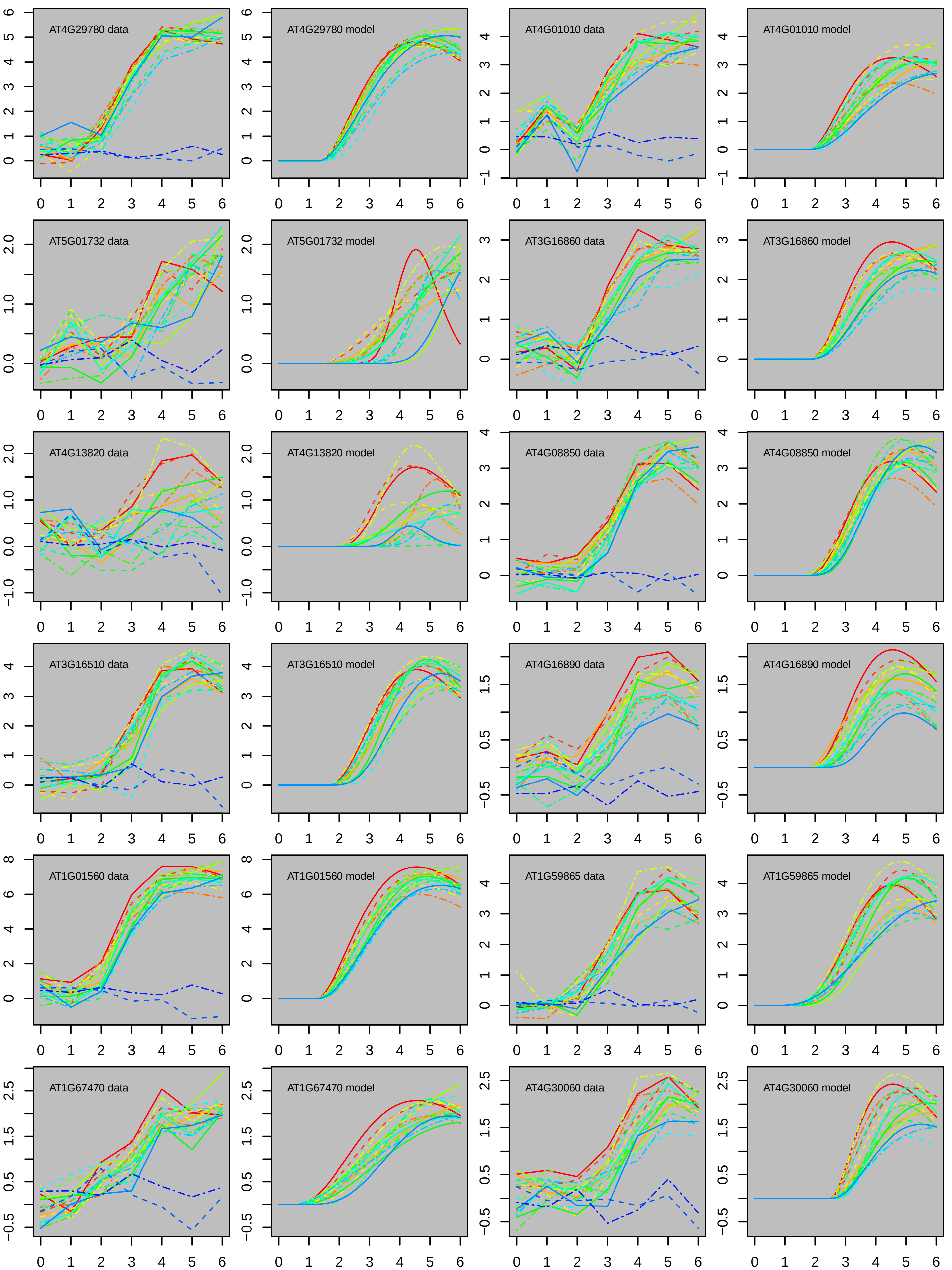
AT4G29780 data
AT4G29780 model
AT4G01010 data
AT4G01010 model
AT5G01732 data
AT5G01732 model
AT3G16860 data
AT3G16860 model
AT4G13820 data
AT4G13820 model
AT4G08850 data
AT4G08850 model
AT3G16510 data
AT3G16510 model
AT4G16890 data
AT4G16890 model
AT1G01560 data
AT1G01560 model
AT1G59865 data
AT1G59865 model
AT1G67470 data
AT1G67470 model
AT4G30060 data
AT4G30060 model

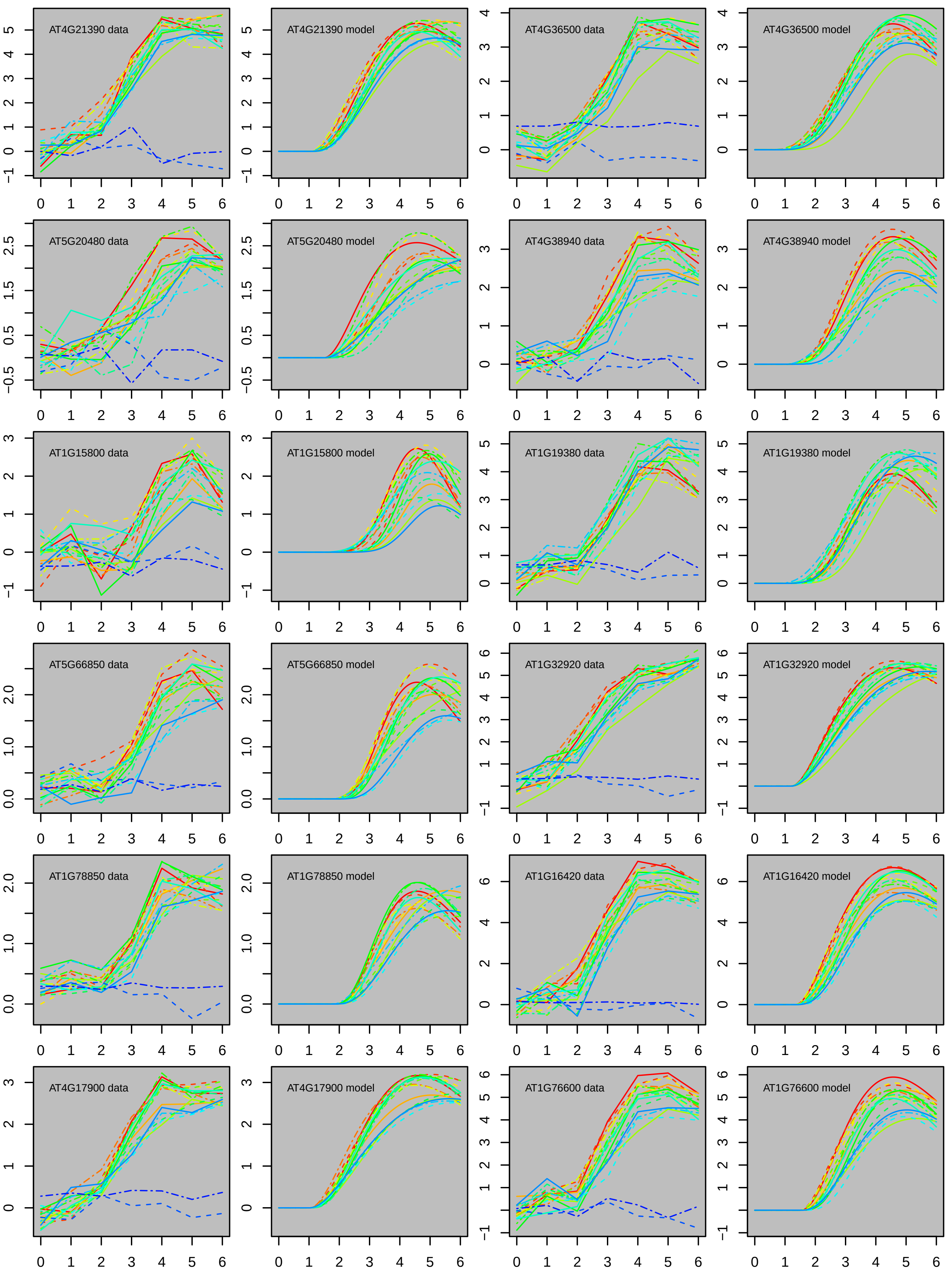
AT4G21390 data
AT4G21390 model
AT4G36500 data
AT4G36500 model
AT5G20480 data
AT5G20480 model
AT4G38940 data
AT4G38940 model
AT1G15800 data
AT1G15800 model
AT1G19380 data
AT1G19380 model
AT5G66850 data
AT5G66850 model
AT1G32920 data
AT1G32920 model
AT1G78850 data
AT1G78850 model
AT1G16420 data
AT1G16420 model
AT4G17900 data
AT4G17900 model
AT1G76600 data
AT1G76600 model

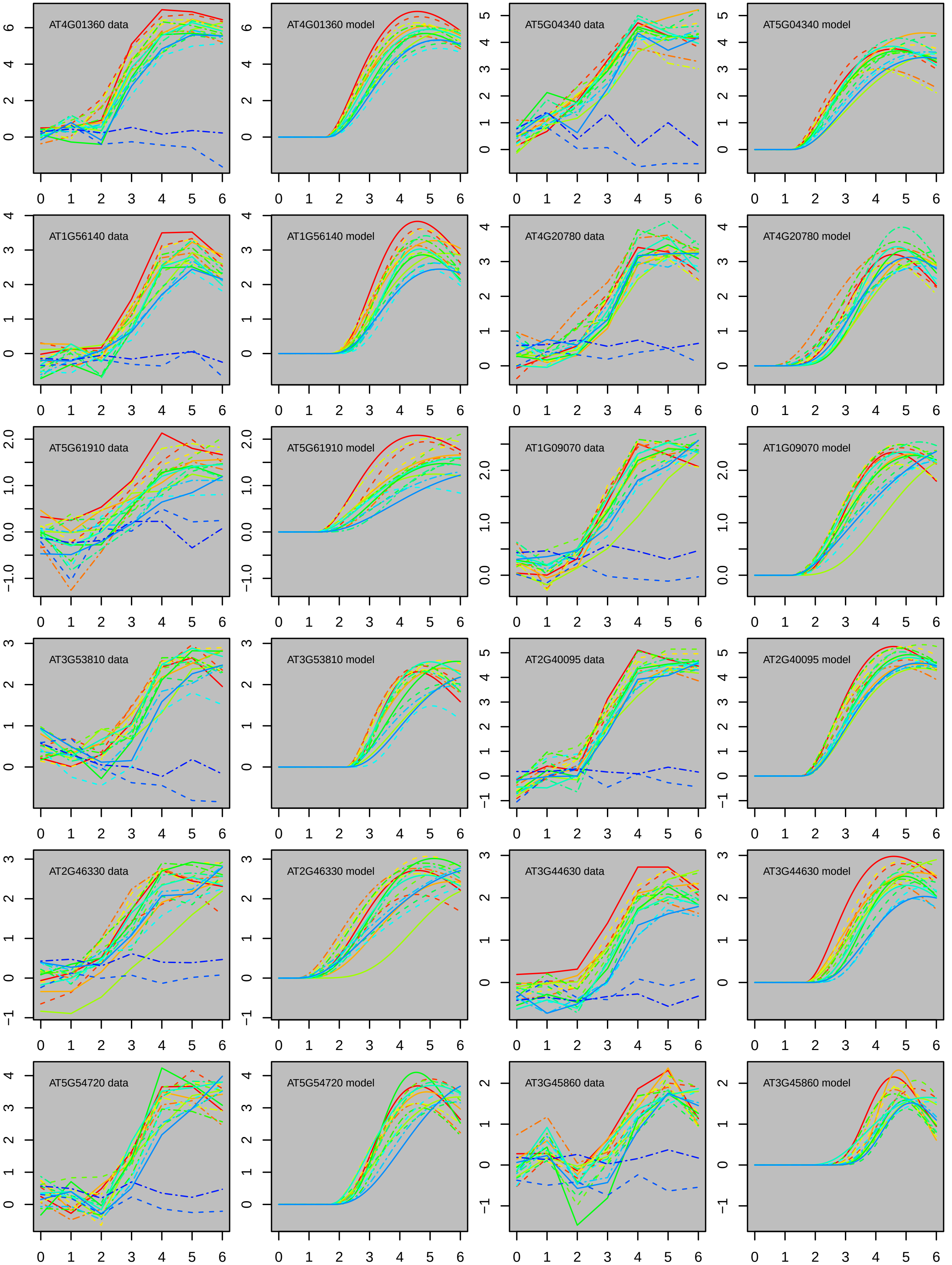
AT4G01360 data
AT4G01360 model
AT5G04340 data
AT5G04340 model
AT1G56140 data
AT1G56140 model
AT4G20780 data
AT4G20780 model
AT5G61910 data
AT5G61910 model
AT1G09070 data
AT1G09070 model
AT3G53810 data
AT3G53810 model
AT2G40095 data
AT2G40095 model
AT2G46330 data
AT2G46330 model
AT3G44630 data
AT3G44630 model
AT5G54720 data
AT5G54720 model
AT3G45860 data
AT3G45860 model

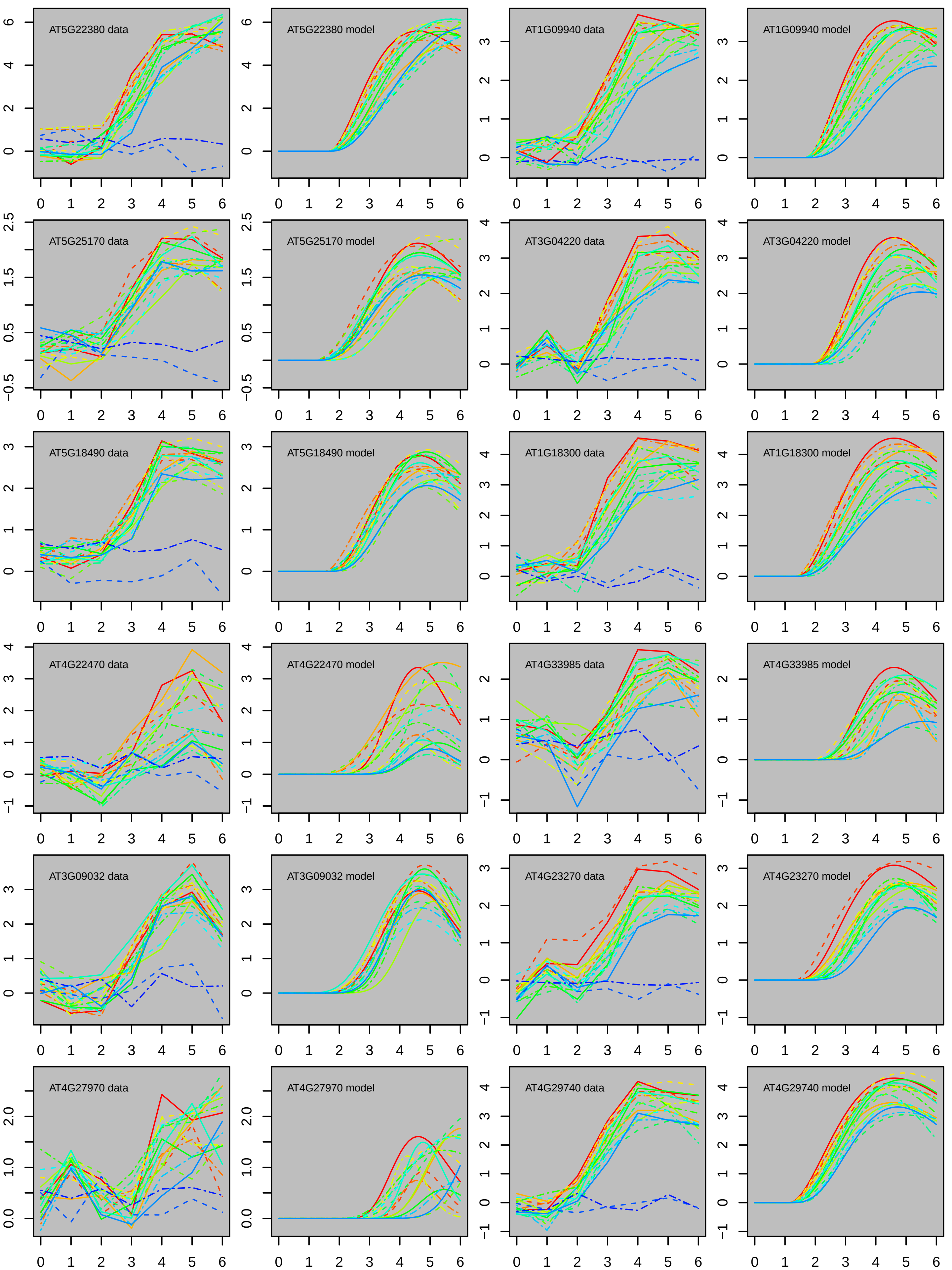
AT5G22380 data
AT5G22380 model
AT1G09940 data
AT1G09940 model
AT5G25170 data
AT5G25170 model
AT3G04220 data
AT3G04220 model
AT5G18490 data
AT5G18490 model
AT1G18300 data
AT1G18300 model
AT4G22470 data
AT4G22470 model
AT4G33985 data
AT4G33985 model
AT3G09032 data
AT3G09032 model
AT4G23270 data
AT4G23270 model
AT4G27970 data
AT4G27970 model
AT4G29740 data
AT4G29740 model

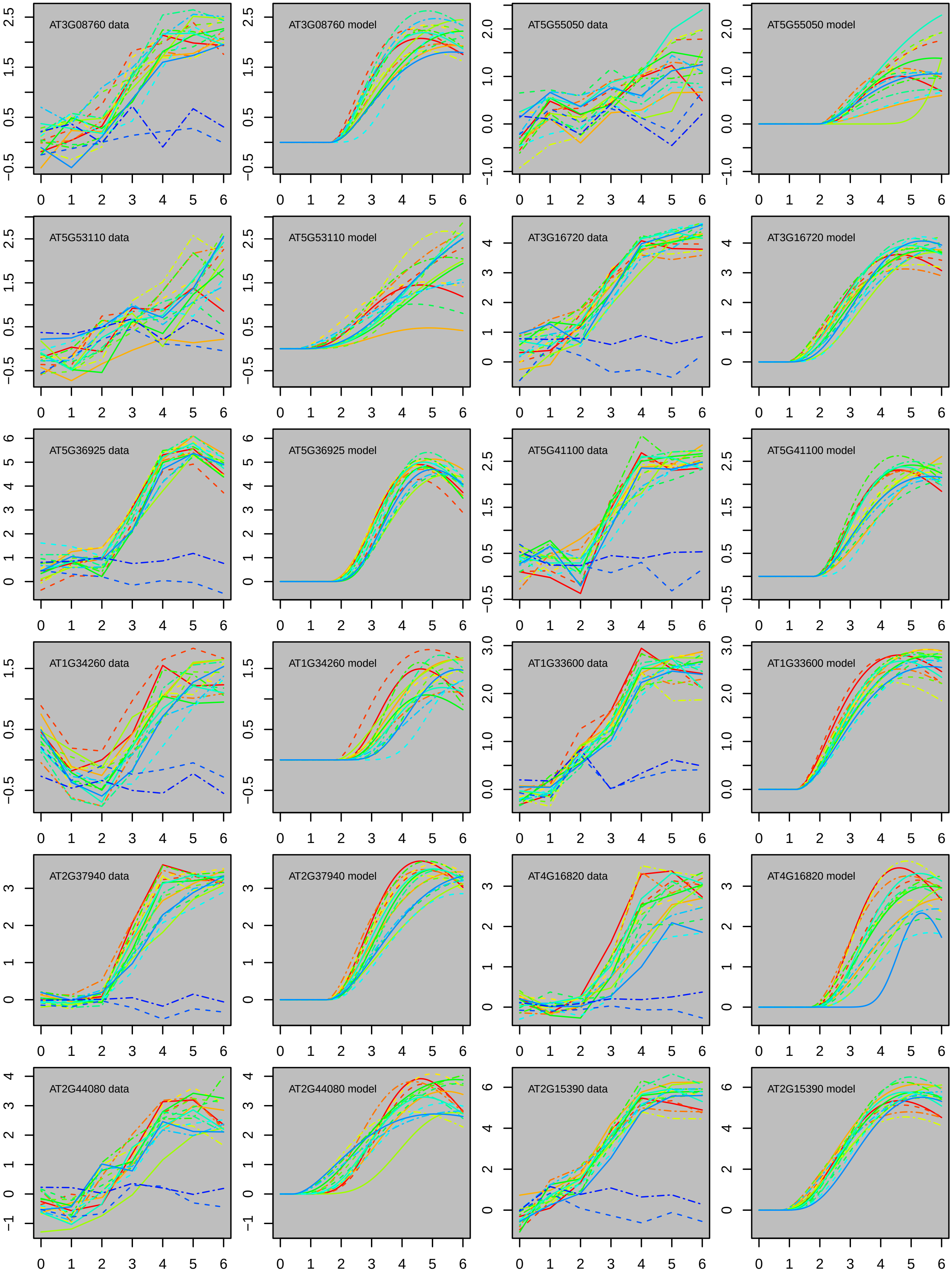
AT3G08760 data
AT3G08760 model
AT5G55050 data
AT5G55050 model
AT5G53110 data
AT5G53110 model
AT3G16720 data
AT3G16720 model
AT5G36925 data
AT5G36925 model
AT5G41100 data
AT5G41100 model
AT1G34260 data
AT1G34260 model
AT1G33600 data
AT1G33600 model
AT2G37940 data
AT2G37940 model
AT4G16820 data
AT4G16820 model
AT2G44080 data
AT2G44080 model
AT2G15390 data
AT2G15390 model

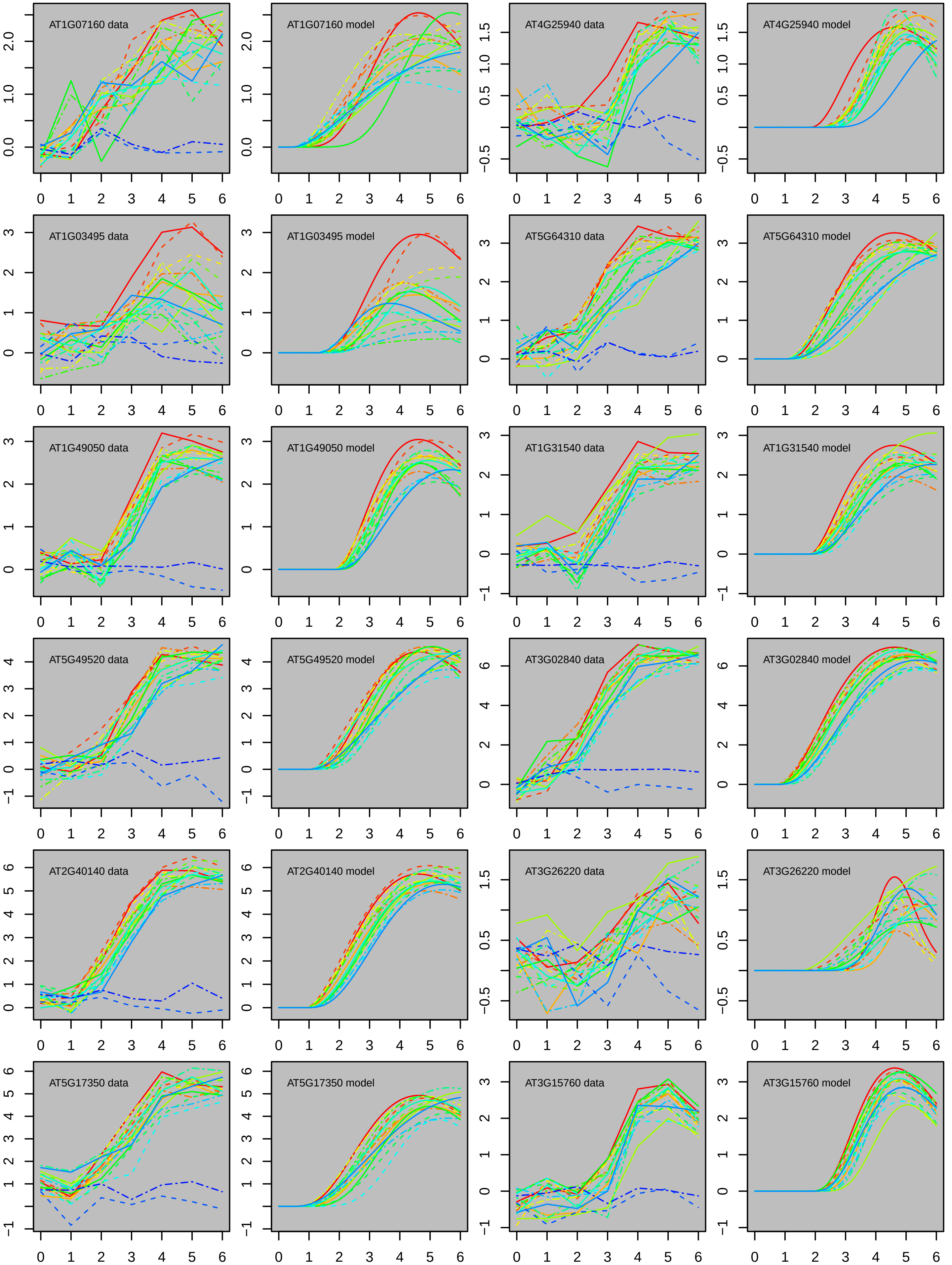

AT1G07160 data
AT1G07160 model
AT4G25940 data
AT4G25940 model
AT1G03495 data
AT1G03495 model
AT5G64310 data
AT5G64310 model
AT1G49050 data
AT1G49050 model
AT1G31540 data
AT1G31540 model
AT5G49520 data
AT5G49520 model
AT3G02840 data
AT3G02840 model
AT2G40140 data
AT2G40140 model
AT3G26220 data
AT3G26220 model
AT5G17350 data
AT5G17350 model
AT3G15760 data
AT3G15760 model

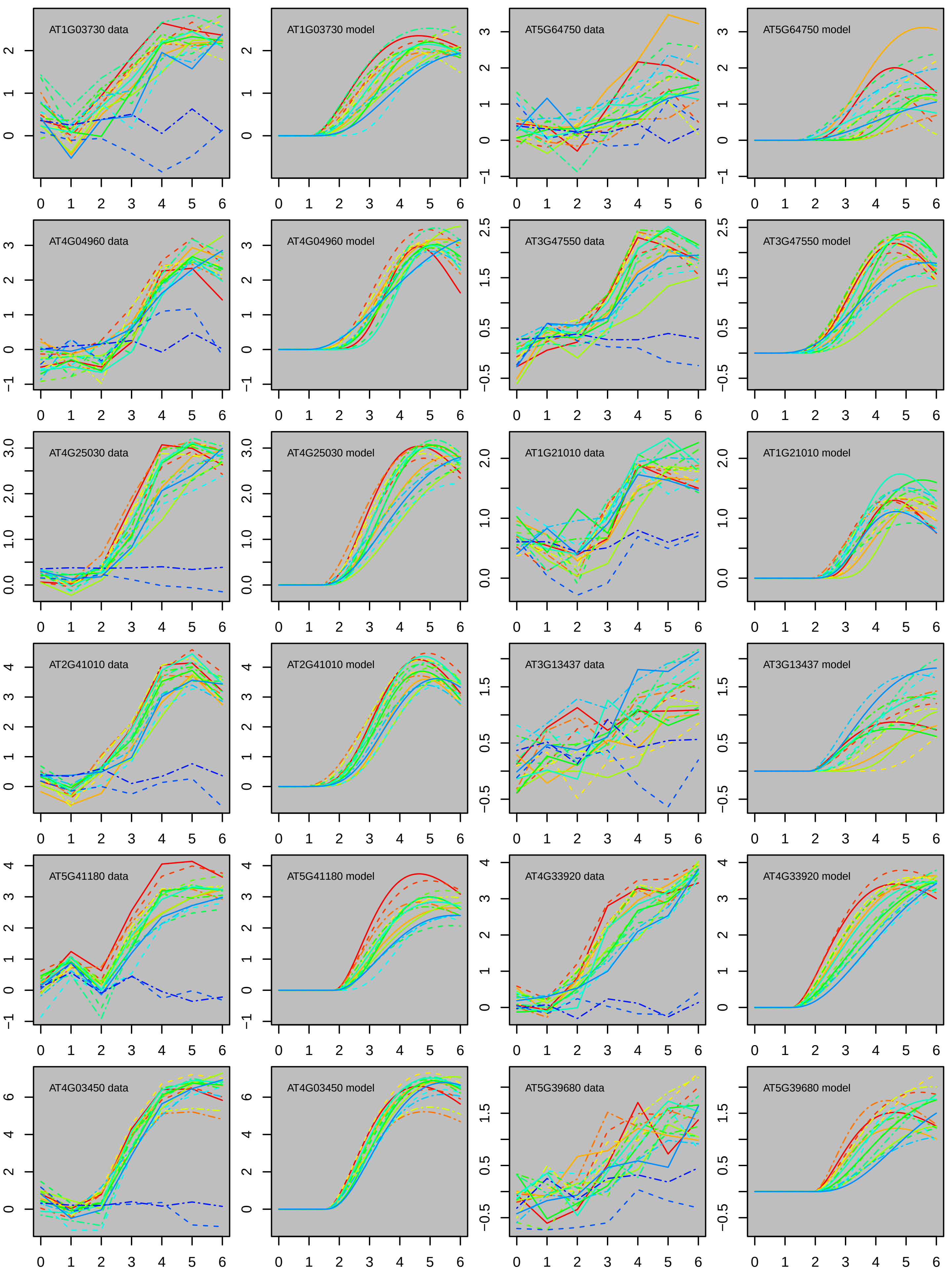
AT1G03730 data
AT1G03730 model
AT5G64750 data
AT5G64750 model
AT4G04960 data
AT4G04960 model
AT3G47550 data
AT3G47550 model
AT4G25030 data
AT4G25030 model
AT1G21010 data
AT1G21010 model
AT2G41010 data
AT2G41010 model
AT3G13437 data
AT3G13437 model
AT5G41180 data
AT5G41180 model
AT4G33920 data
AT4G33920 model
AT4G03450 data
AT4G03450 model
AT5G39680 data
AT5G39680 model

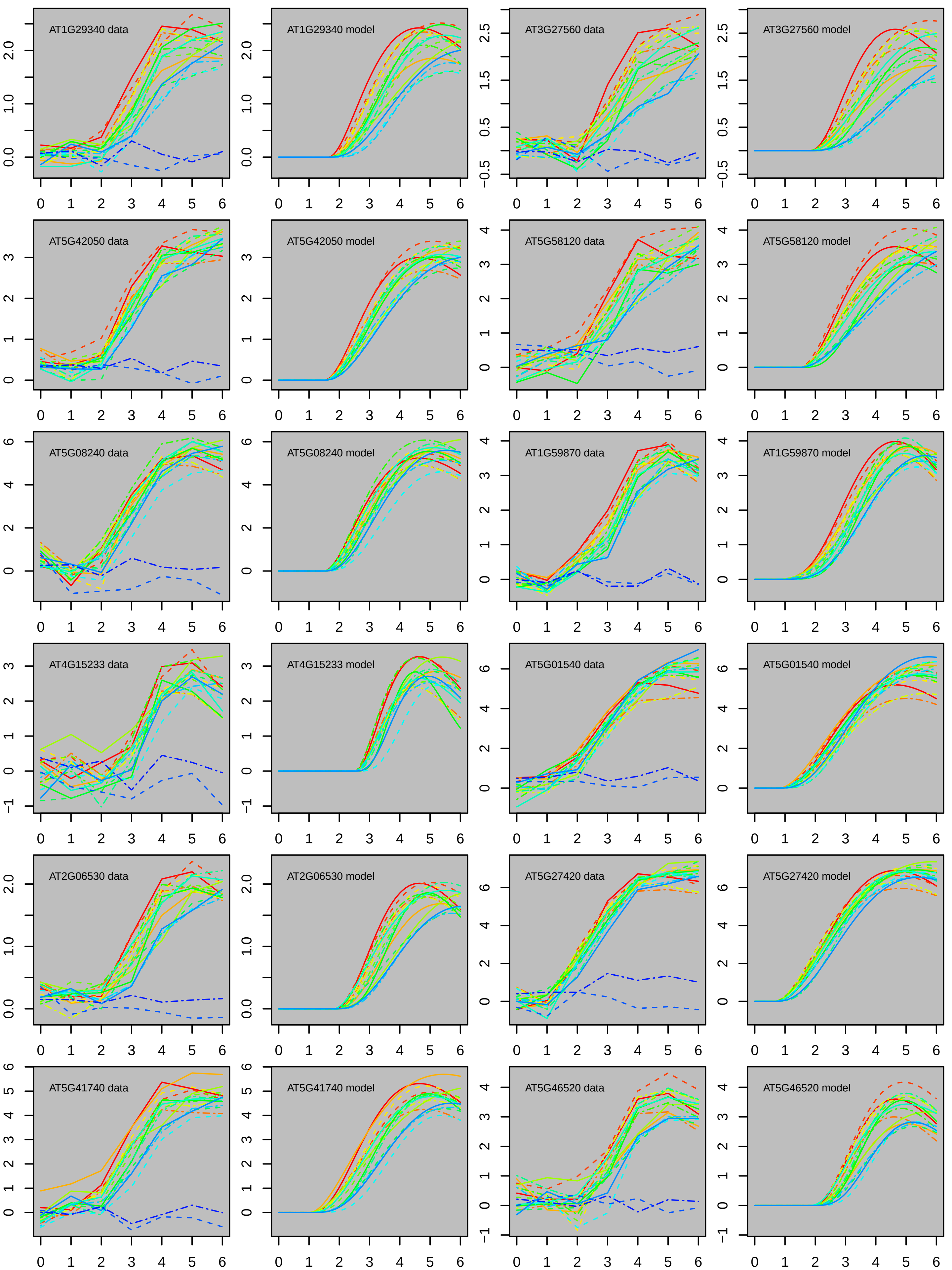

AT1G29340 data
AT1G29340 model
AT3G27560 data
AT3G27560 model
AT5G42050 data
AT5G42050 model
AT5G58120 data
AT5G58120 model
AT5G08240 data
AT5G08240 model
AT1G59870 data
AT1G59870 model
AT4G15233 data
AT4G15233 model
AT5G01540 data
AT5G01540 model
AT2G06530 data
AT2G06530 model
AT5G27420 data
AT5G27420 model
AT5G41740 data
AT5G41740 model
AT5G46520 data
AT5G46520 model

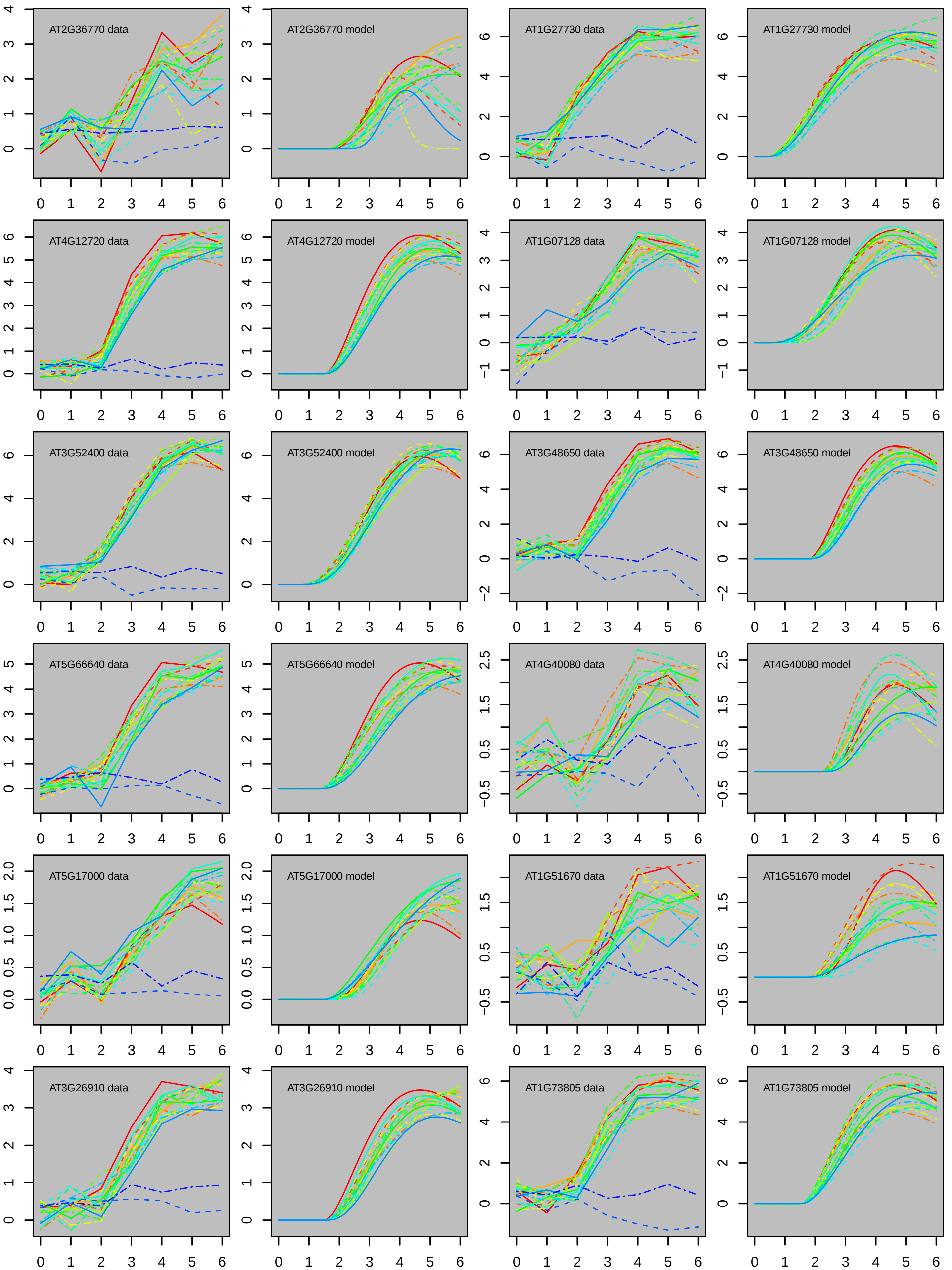
AT2G36770 data
AT2G36770 model
AT1G27730 data
AT1G27730 model
AT4G12720 data
AT4G12720 model
AT1G07128 data
AT1G07128 model
AT3G52400 data
AT3G52400 model
AT3G48650 data
AT3G48650 model
AT5G66640 data
AT5G66640 model
AT4G40080 data
AT4G40080 model
AT5G17000 data
AT5G17000 model
AT1G51670 data
AT1G51670 model
AT3G26910 data
AT3G26910 model
AT1G73805 data
AT1G73805 model

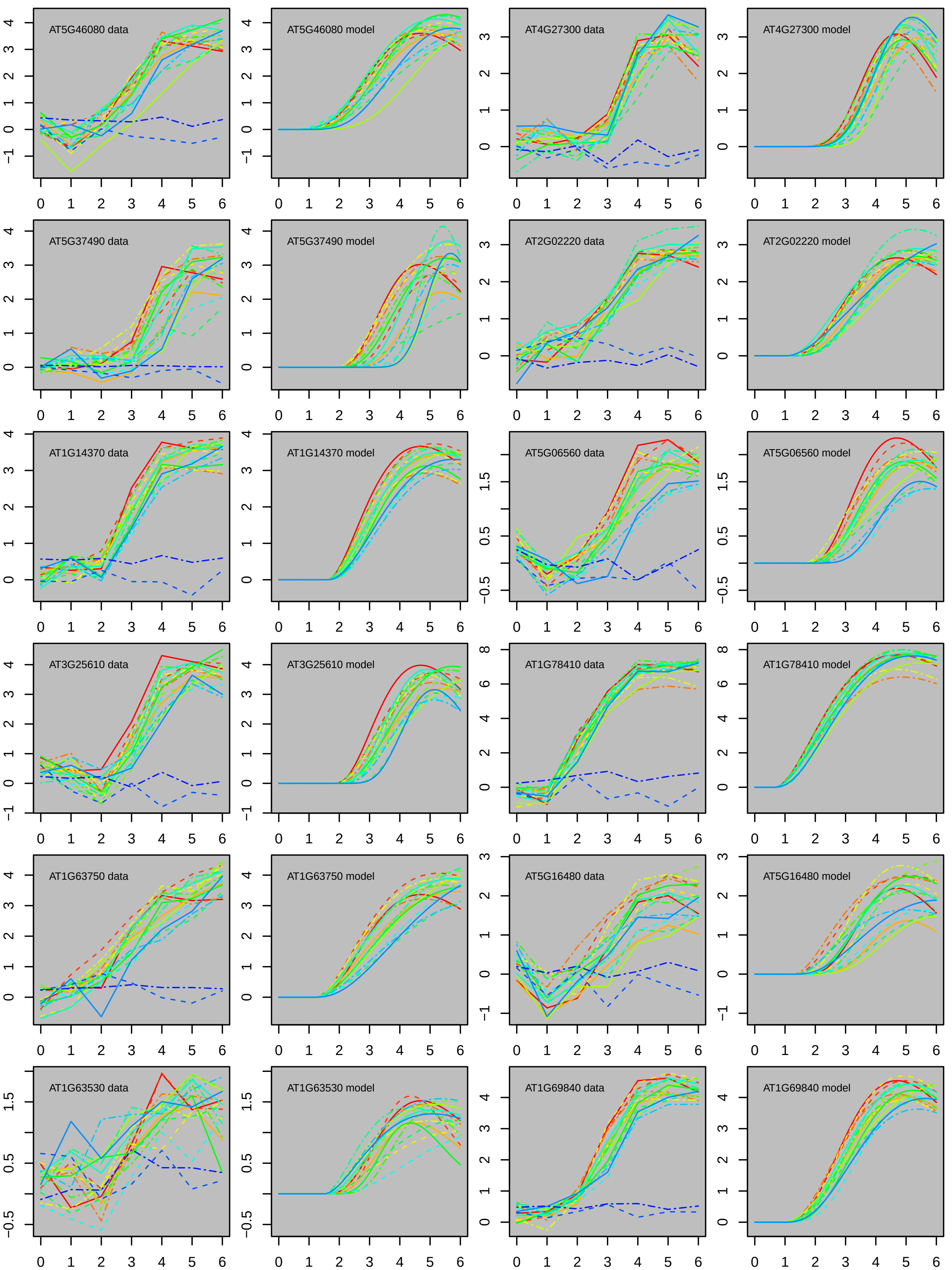

AT5G46080 data
AT5G46080 model
AT4G27300 data
AT4G27300 model
AT5G37490 data
AT5G37490 model
AT2G02220 data
AT2G02220 model
AT1G14370 data
AT1G14370 model
AT5G06560 data
AT5G06560 model
AT3G25610 data
AT3G25610 model
AT1G78410 data
AT1G78410 model
AT1G63750 data
AT1G63750 model
AT5G16480 data
AT5G16480 model
AT1G63530 data
AT1G63530 model
AT1G69840 data
AT1G69840 model

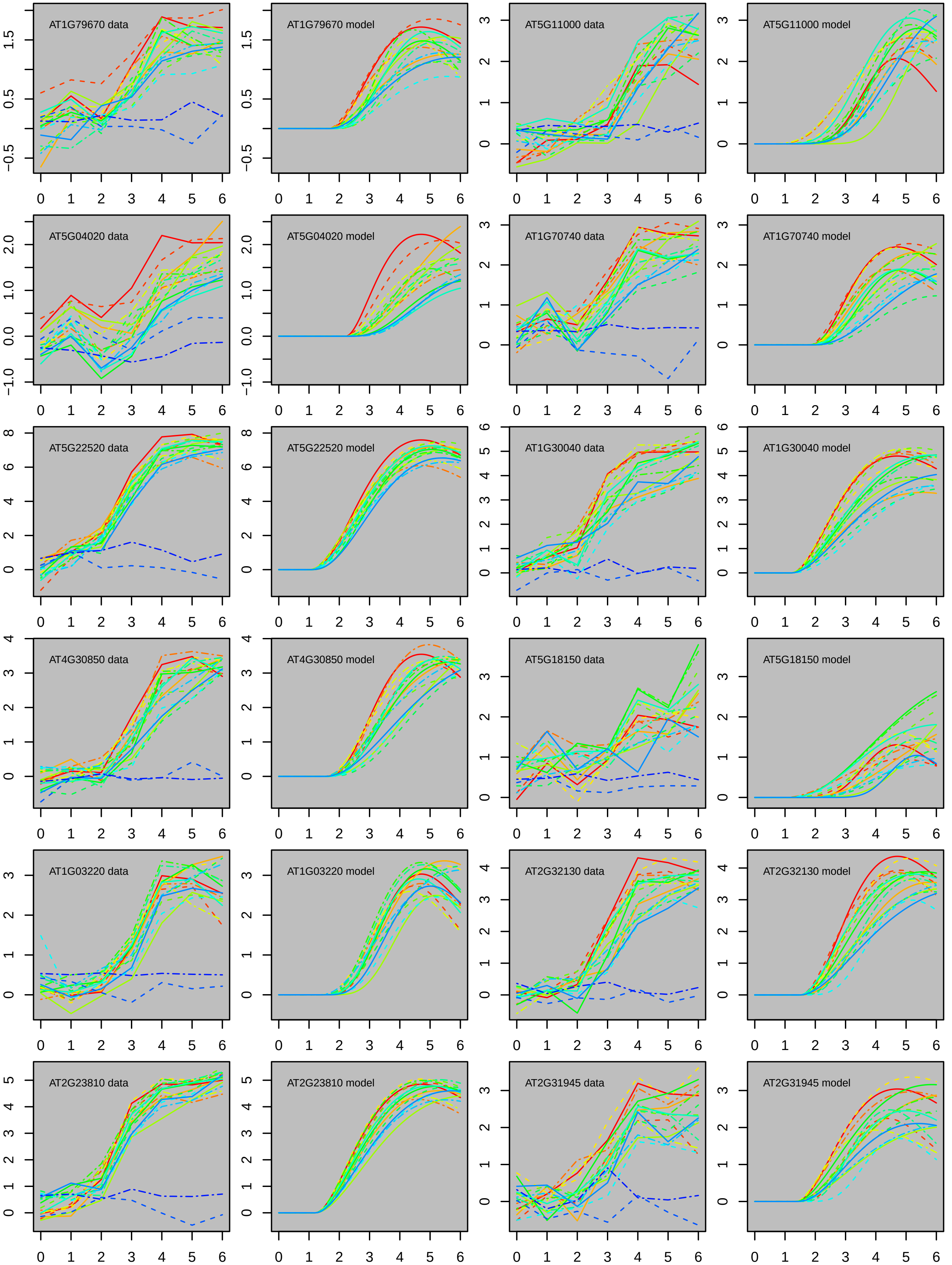

AT1G79670 data
AT1G79670 model
AT5G11000 data
AT5G11000 model
AT5G04020 data
AT5G04020 model
AT1G70740 data
AT1G70740 model
AT5G22520 data
AT5G22520 model
AT1G30040 data
AT1G30040 model
AT4G30850 data
AT4G30850 model
AT5G18150 data
AT5G18150 model
AT1G03220 data
AT1G03220 model
AT2G32130 data
AT2G32130 model
AT2G23810 data
AT2G23810 model
AT2G31945 data
AT2G31945 model

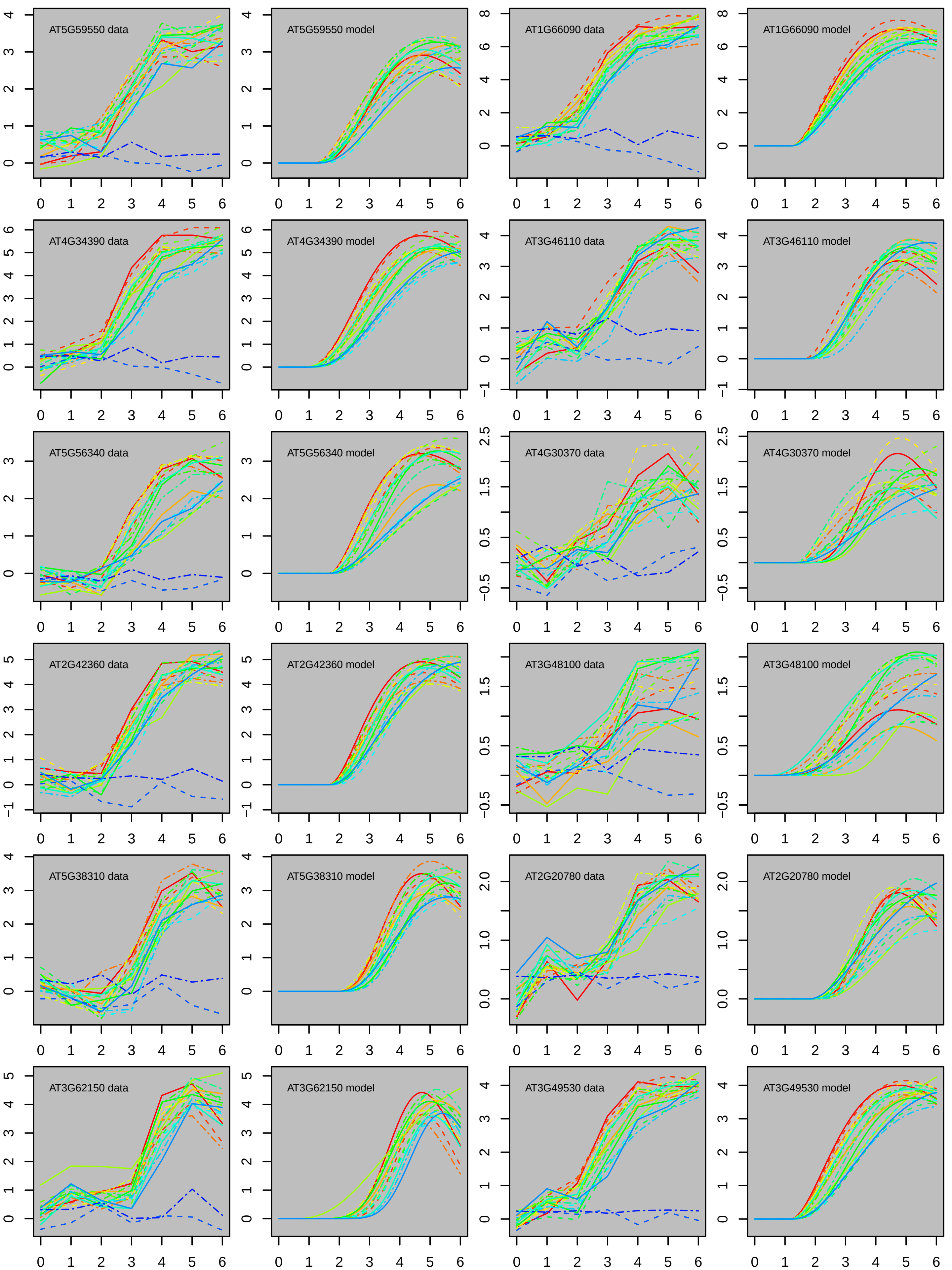
AT5G59550 data
AT5G59550 model
AT1G66090 data
AT1G66090 model
AT4G34390 data
AT4G34390 model
AT3G46110 data
AT3G46110 model
AT5G56340 data
AT5G56340 model
AT4G30370 data
AT4G30370 model
AT2G42360 data
AT2G42360 model
AT3G48100 data
AT3G48100 model
AT5G38310 data
AT5G38310 model
AT2G20780 data
AT2G20780 model
AT3G62150 data
AT3G62150 model
AT3G49530 data
AT3G49530 model

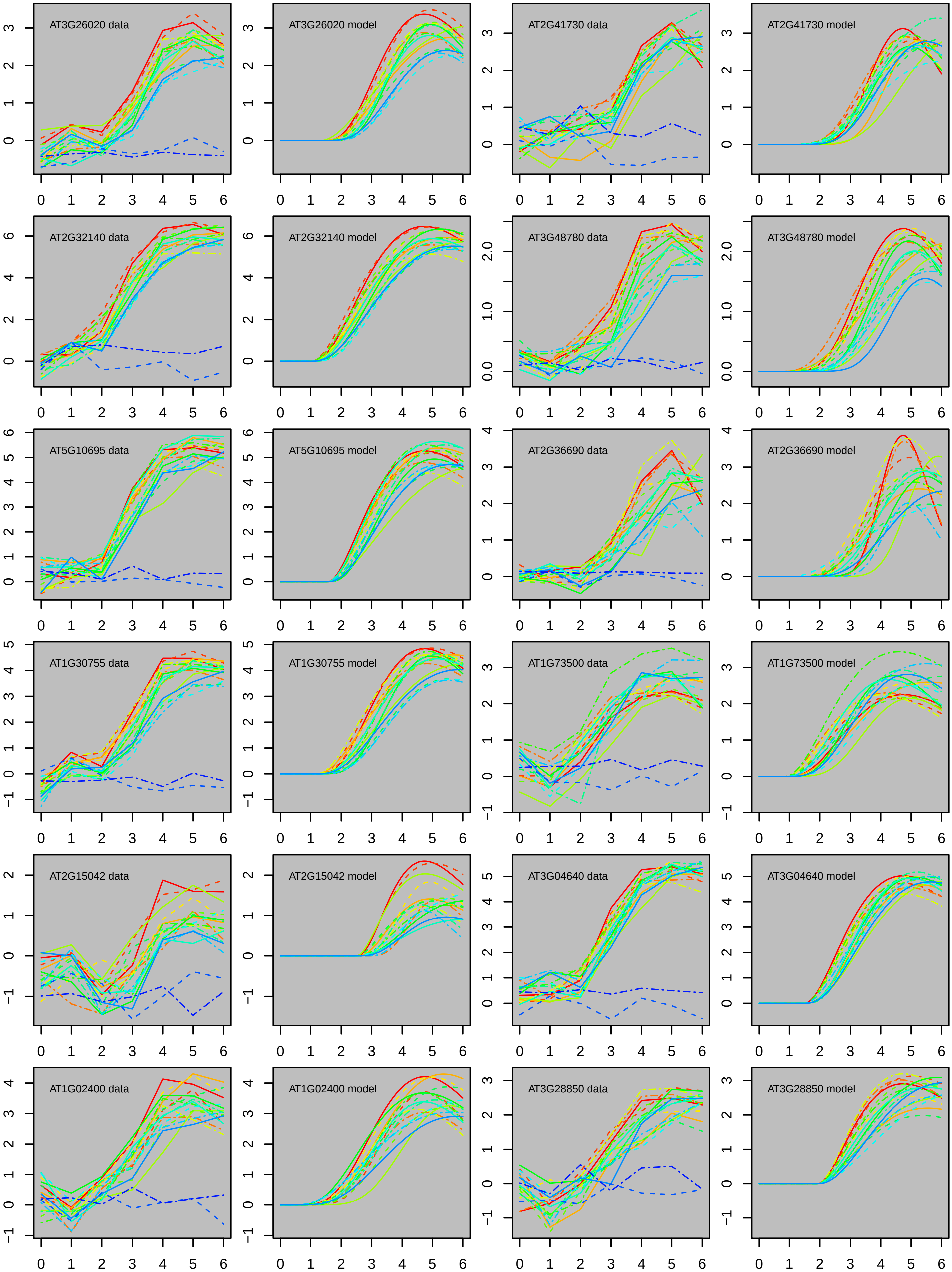

AT3G26020 data
AT3G26020 model
AT2G41730 data
AT2G41730 model
AT2G32140 data
AT2G32140 model
AT3G48780 data
AT3G48780 model
AT5G10695 data
AT5G10695 model
AT2G36690 data
AT2G36690 model
AT1G30755 data
AT1G30755 model
AT1G73500 data
AT1G73500 model
AT2G15042 data
AT2G15042 model
AT3G04640 data
AT3G04640 model
AT1G02400 data
AT1G02400 model
AT3G28850 data
AT3G28850 model

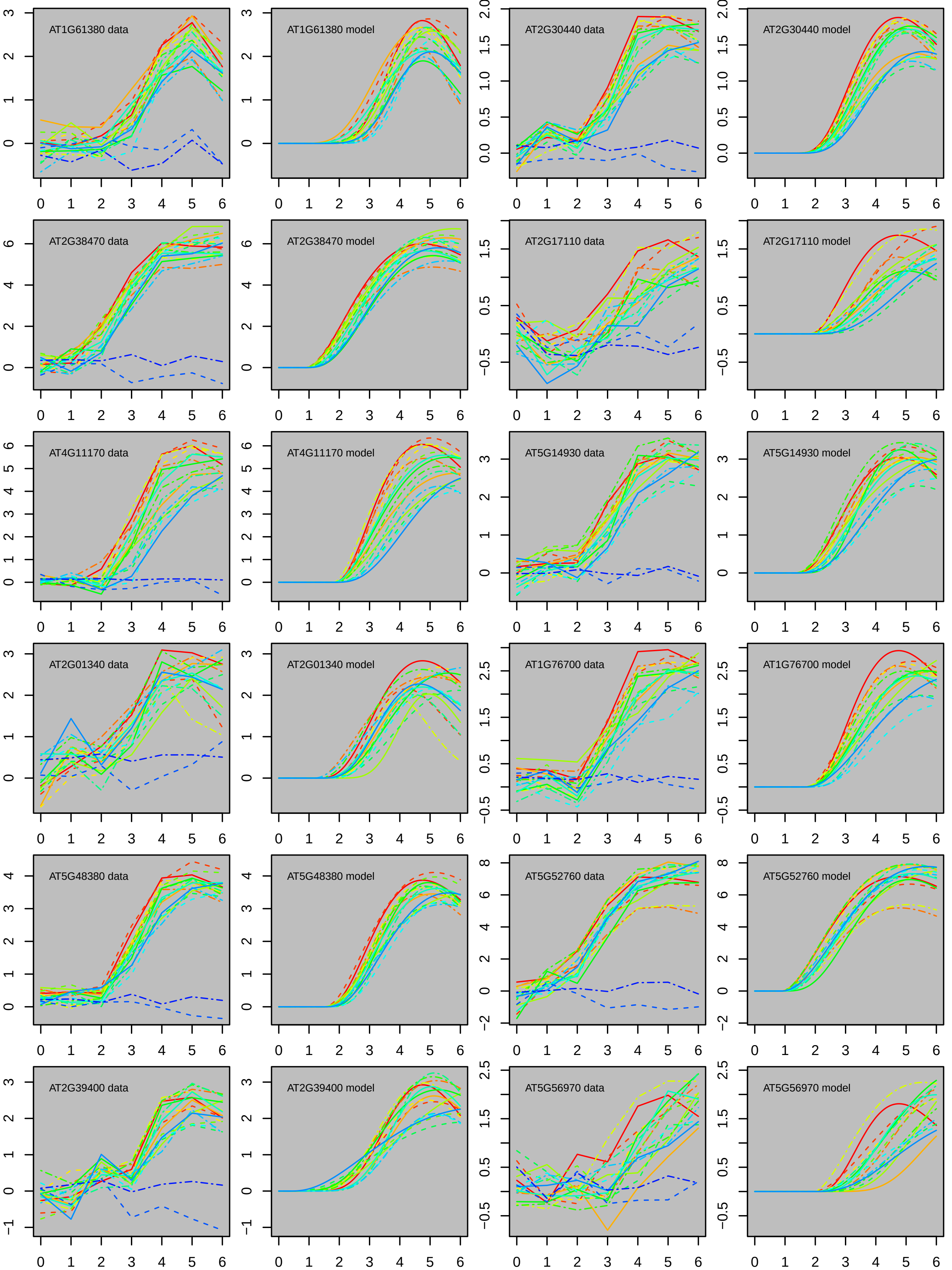
AT1G61380 data
AT1G61380 model
AT2G30440 data
AT2G30440 model
AT2G38470 data
AT2G38470 model
AT2G17110 data
AT2G17110 model
AT4G11170 data
AT4G11170 model
AT5G14930 data
AT5G14930 model
AT2G01340 data
AT2G01340 model
AT1G76700 data
AT1G76700 model
AT5G48380 data
AT5G48380 model
AT5G52760 data
AT5G52760 model
AT2G39400 data
AT2G39400 model
AT5G56970 data
AT5G56970 model

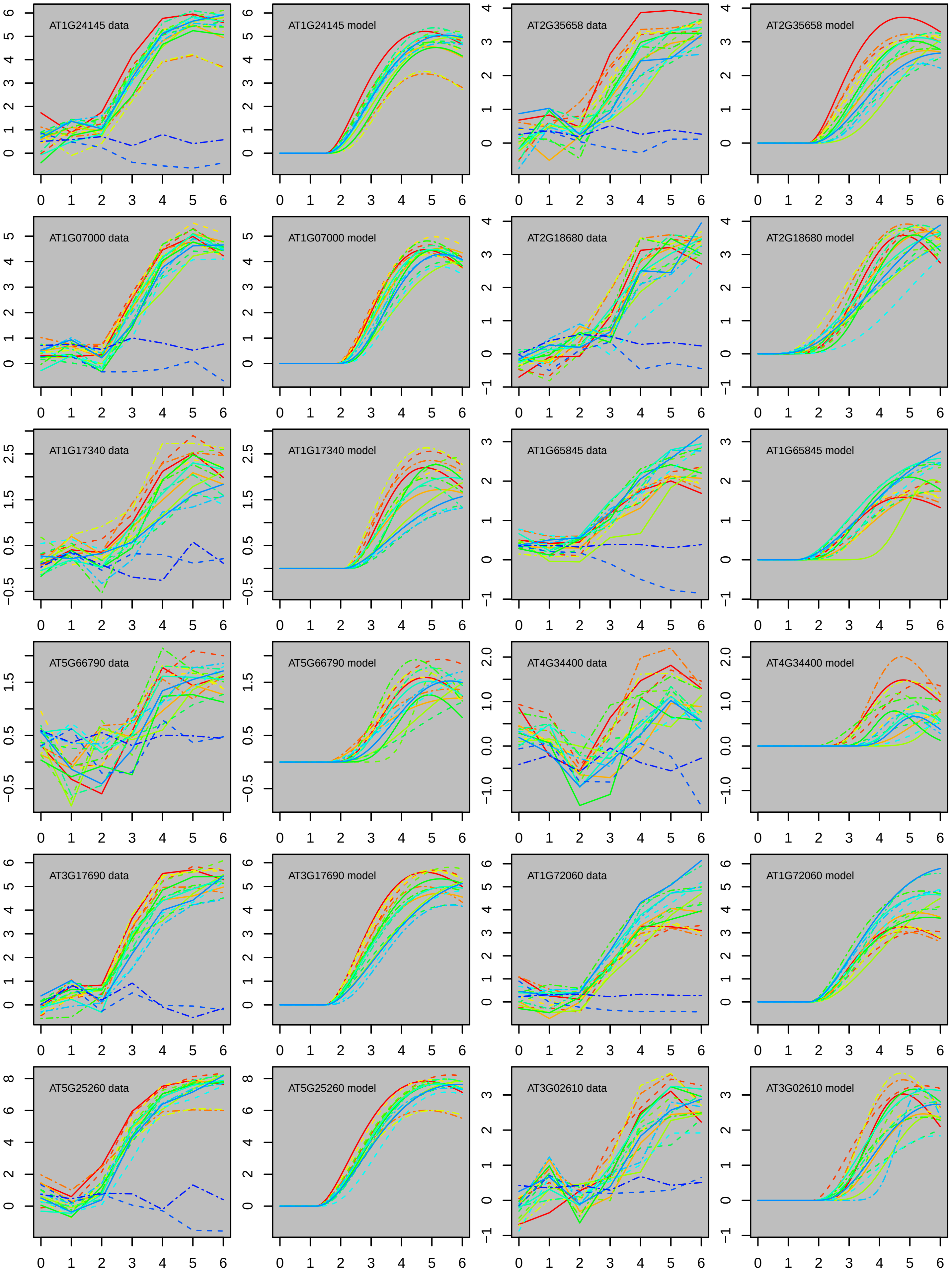

AT1G24145 data
AT1G24145 model
AT2G35658 data
AT2G35658 model
AT1G07000 data
AT1G07000 model
AT2G18680 data
AT2G18680 model
AT1G17340 data
AT1G17340 model
AT1G65845 data
AT1G65845 model
AT5G66790 data
AT5G66790 model
AT4G34400 data
AT4G34400 model
AT3G17690 data
AT3G17690 model
AT1G72060 data
AT1G72060 model
AT5G25260 data
AT5G25260 model
AT3G02610 data
AT3G02610 model

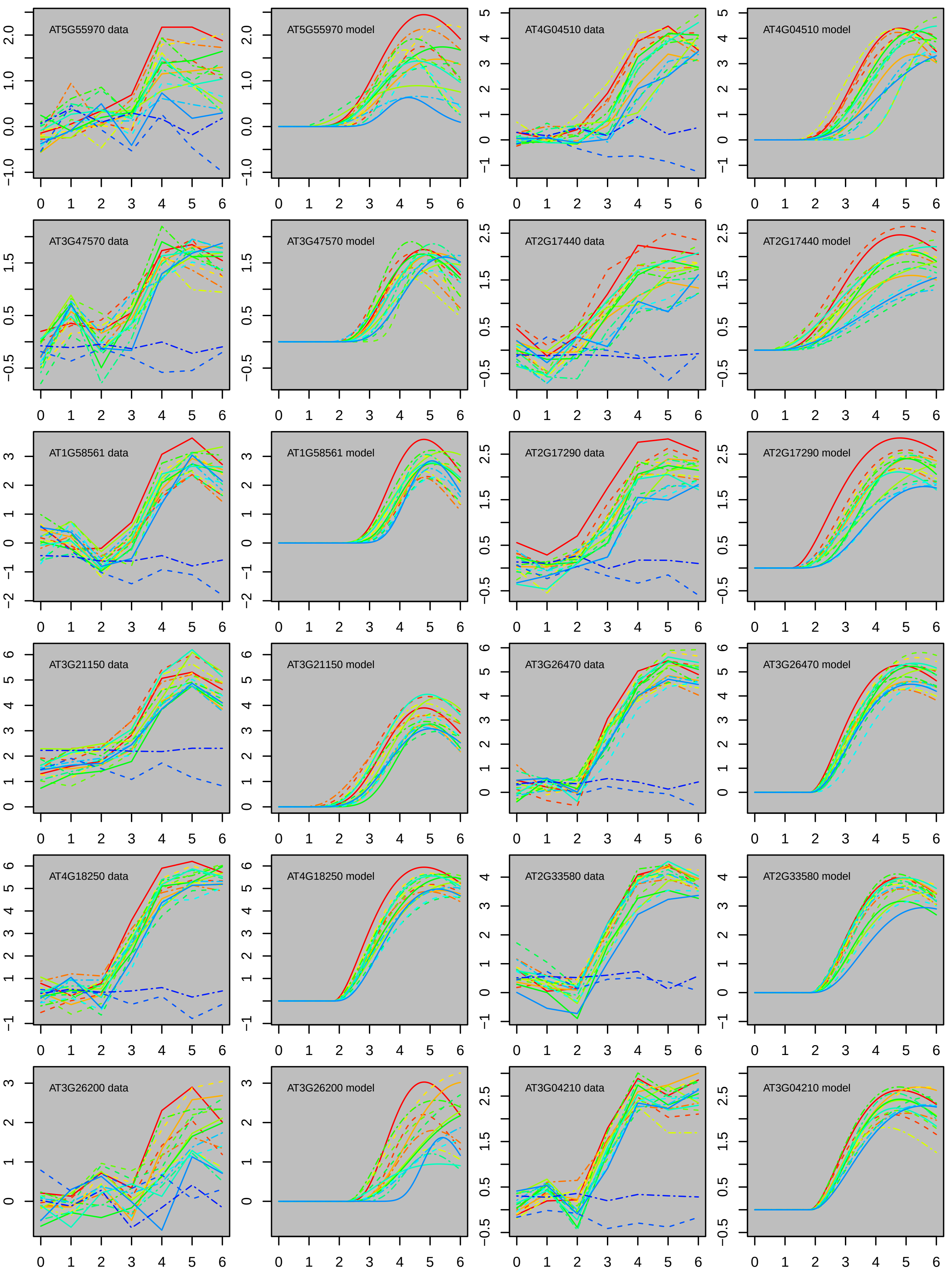
AT5G55970 data
AT5G55970 model
AT4G04510 data
AT4G04510 model
AT3G47570 data
AT3G47570 model
AT2G17440 data
AT2G17440 model
AT1G58561 data
AT1G58561 model
AT2G17290 data
AT2G17290 model
AT3G21150 data
AT3G21150 model
AT3G26470 data
AT3G26470 model
AT4G18250 data
AT4G18250 model
AT2G33580 data
AT2G33580 model
AT3G26200 data
AT3G26200 model
AT3G04210 data
AT3G04210 model

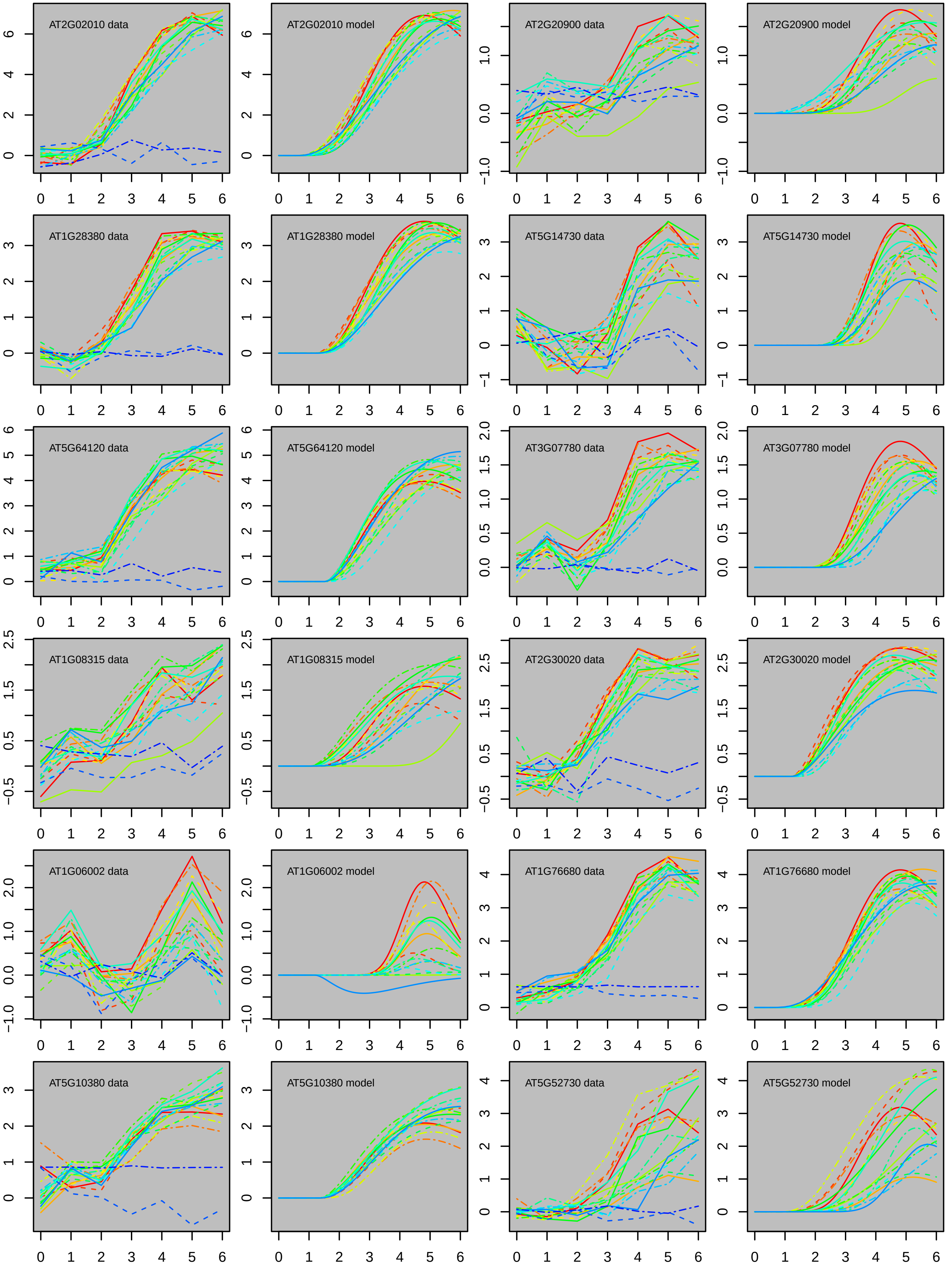

AT2G02010 data
AT2G02010 model
AT2G20900 data
AT2G20900 model
AT1G28380 data
AT1G28380 model
AT5G14730 data
AT5G14730 model
AT5G64120 data
AT5G64120 model
AT3G07780 data
AT3G07780 model
AT1G08315 data
AT1G08315 model
AT2G30020 data
AT2G30020 model
AT1G06002 data
AT1G06002 model
AT1G76680 data
AT1G76680 model
AT5G10380 data
AT5G10380 model
AT5G52730 data
AT5G52730 model

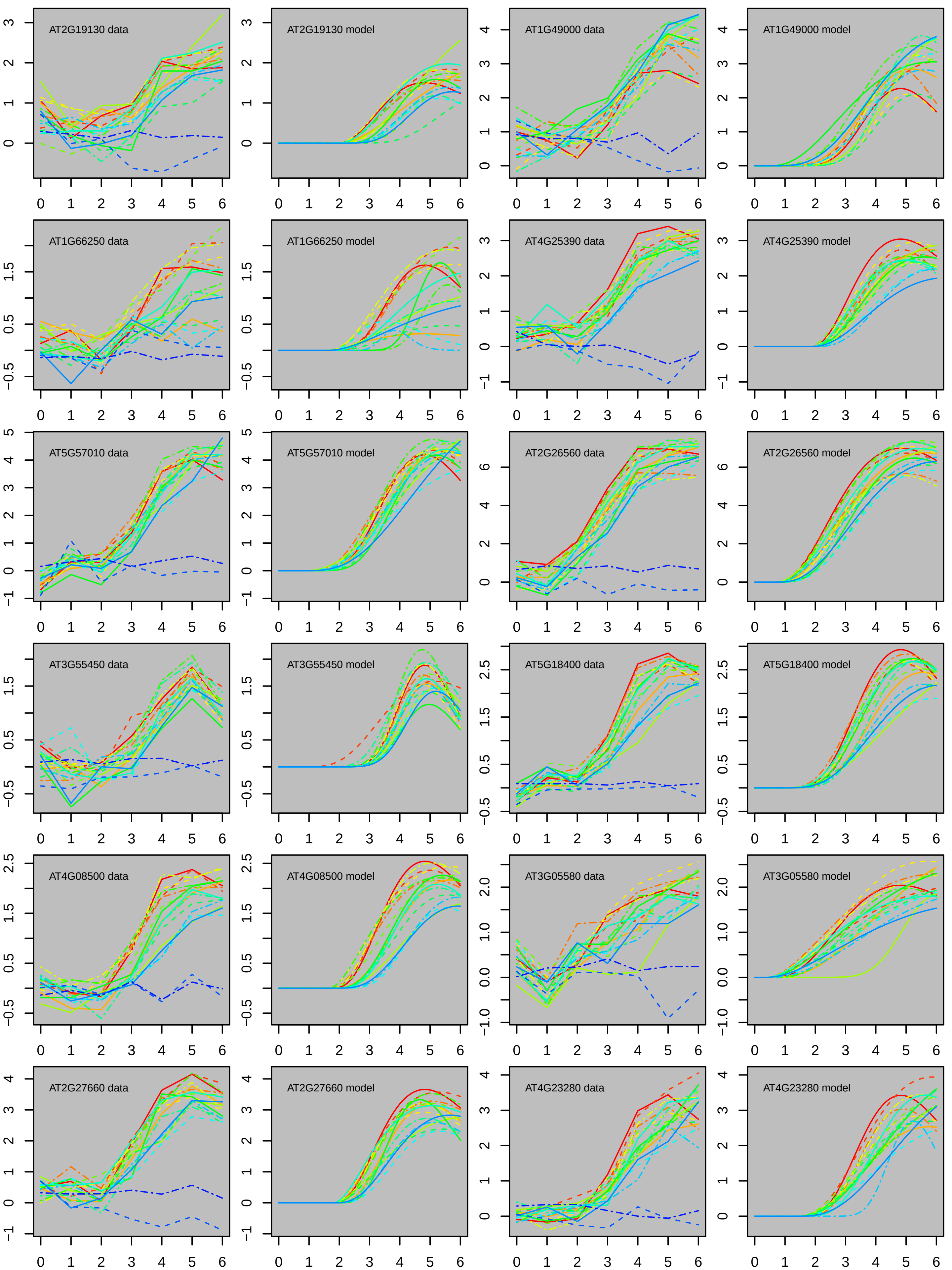
AT2G19130 data
AT2G19130 model
AT1G49000 data
AT1G49000 model
AT1G66250 data
AT1G66250 model
AT4G25390 data
AT4G25390 model
AT5G57010 data
AT5G57010 model
AT2G26560 data
AT2G26560 model
AT3G55450 data
AT3G55450 model
AT5G18400 data
AT5G18400 model
AT4G08500 data
AT4G08500 model
AT3G05580 data
AT3G05580 model
AT2G27660 data
AT2G27660 model
AT4G23280 data
AT4G23280 model

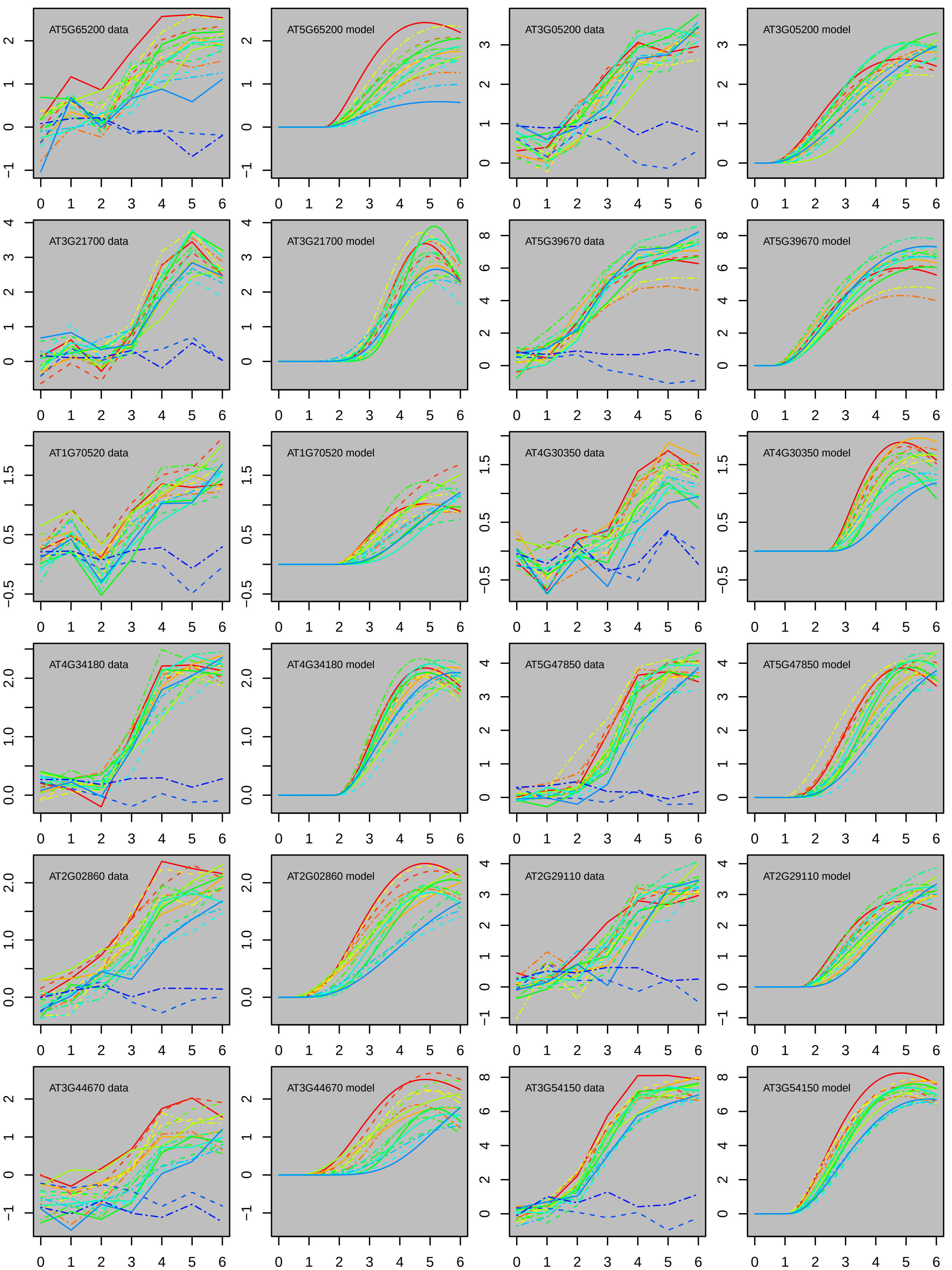
AT5G65200 data
AT5G65200 model
AT3G05200 data
AT3G05200 model
AT3G21700 data
AT3G21700 model
AT5G39670 data
AT5G39670 model
AT1G70520 data
AT1G70520 model
AT4G30350 data
AT4G30350 model
AT4G34180 data
AT4G34180 model
AT5G47850 data
AT5G47850 model
AT2G02860 data
AT2G02860 model
AT2G29110 data
AT2G29110 model
AT3G44670 data
AT3G44670 model
AT3G54150 data
AT3G54150 model

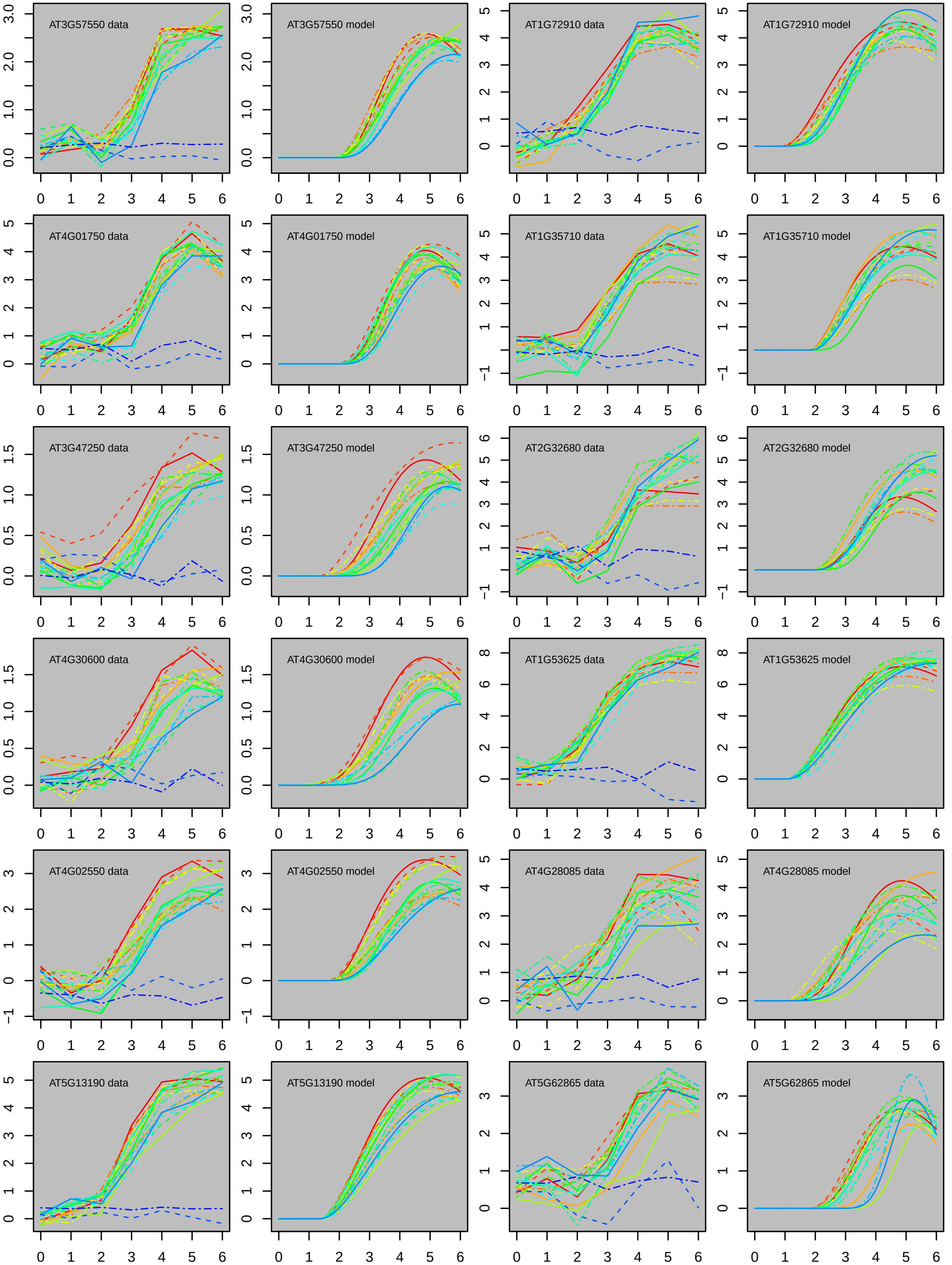

AT3G57550 data
AT3G57550 model
AT1G72910 data
AT1G72910 model
AT4G01750 data
AT4G01750 model
AT1G35710 data
AT1G35710 model
AT3G47250 data
AT3G47250 model
AT2G32680 data
AT2G32680 model
AT4G30600 data
AT4G30600 model
AT1G53625 data
AT1G53625 model
AT4G02550 data
AT4G02550 model
AT4G28085 data
AT4G28085 model
AT5G13190 data
AT5G13190 model
AT5G62865 data
AT5G62865 model

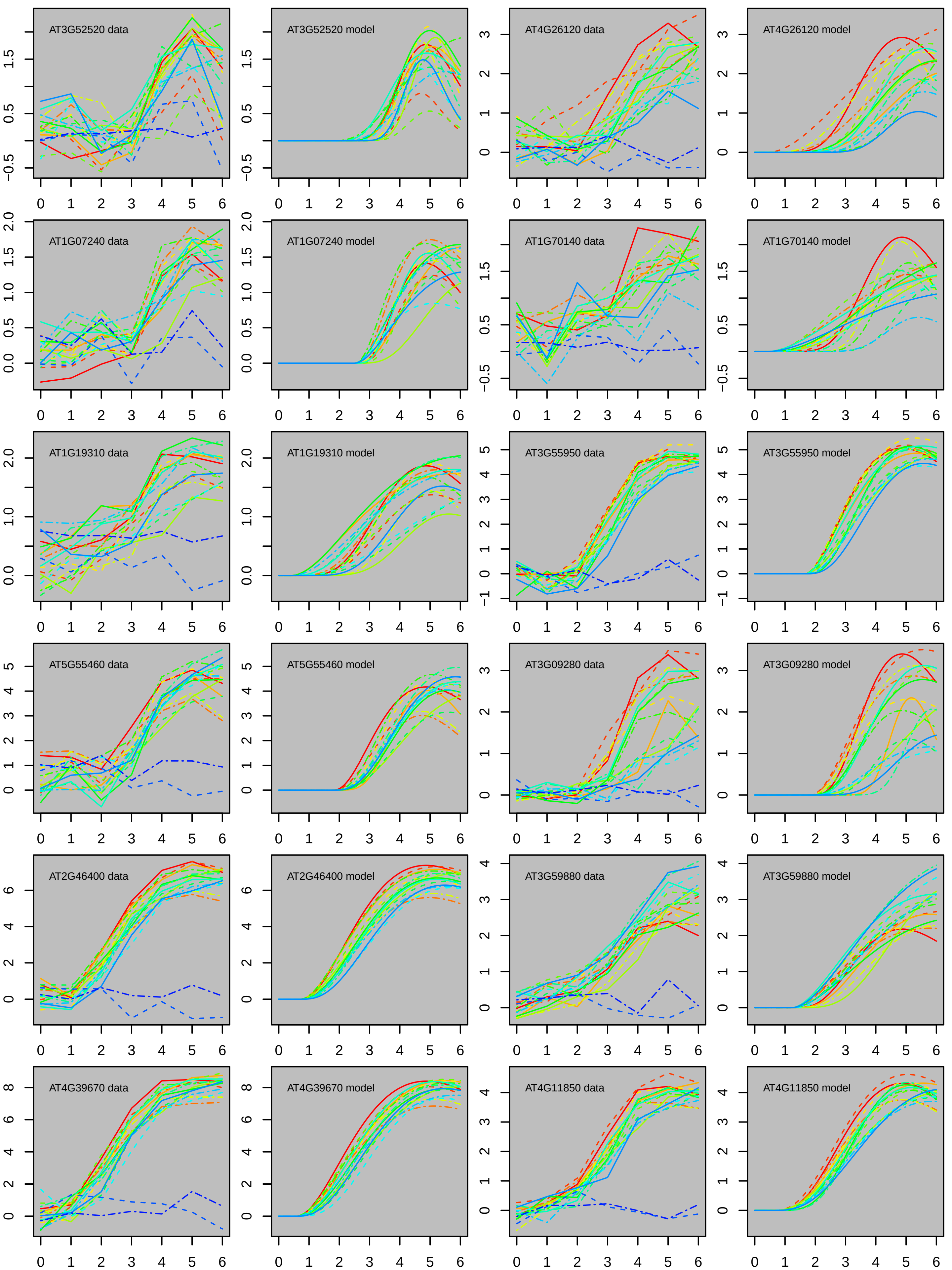
AT3G52520 data
AT3G52520 model
AT4G26120 data
AT4G26120 model
AT1G07240 data
AT1G07240 model
AT1G70140 data
AT1G70140 model
AT1G19310 data
AT1G19310 model
AT3G55950 data
AT3G55950 model
AT5G55460 data
AT5G55460 model
AT3G09280 data
AT3G09280 model
AT2G46400 data
AT2G46400 model
AT3G59880 data
AT3G59880 model
AT4G39670 data
AT4G39670 model
AT4G11850 data
AT4G11850 model

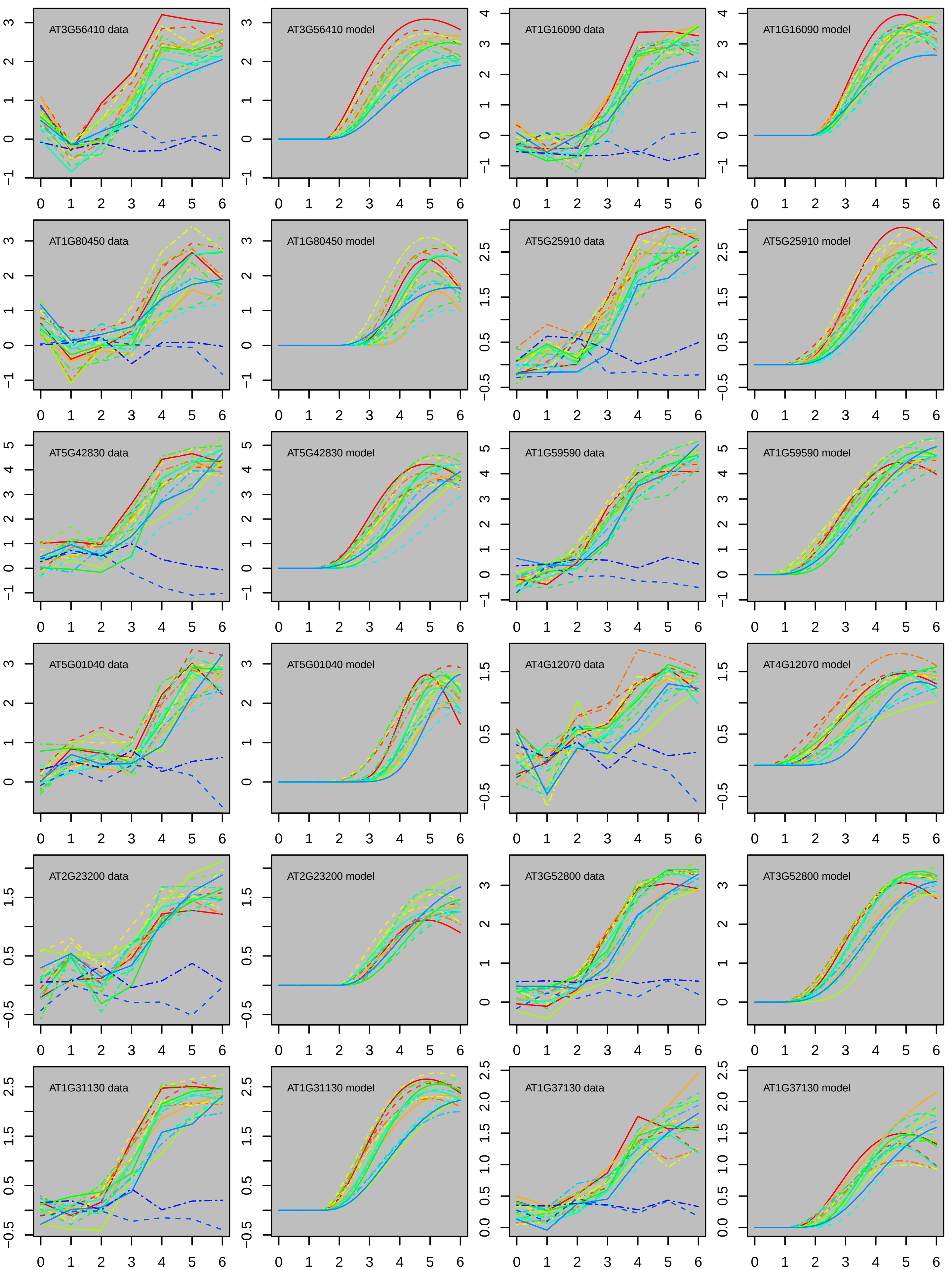

AT3G56410 data
AT3G56410 model
AT1G16090 data
AT1G16090 model
AT1G80450 data
AT1G80450 model
AT5G25910 data
AT5G25910 model
AT5G42830 data
AT5G42830 model
AT1G59590 data
AT1G59590 model
AT5G01040 data
AT5G01040 model
AT4G12070 data
AT4G12070 model
AT2G23200 data
AT2G23200 model
AT3G52800 data
AT3G52800 model
AT1G31130 data
AT1G31130 model
AT1G37130 data
AT1G37130 model

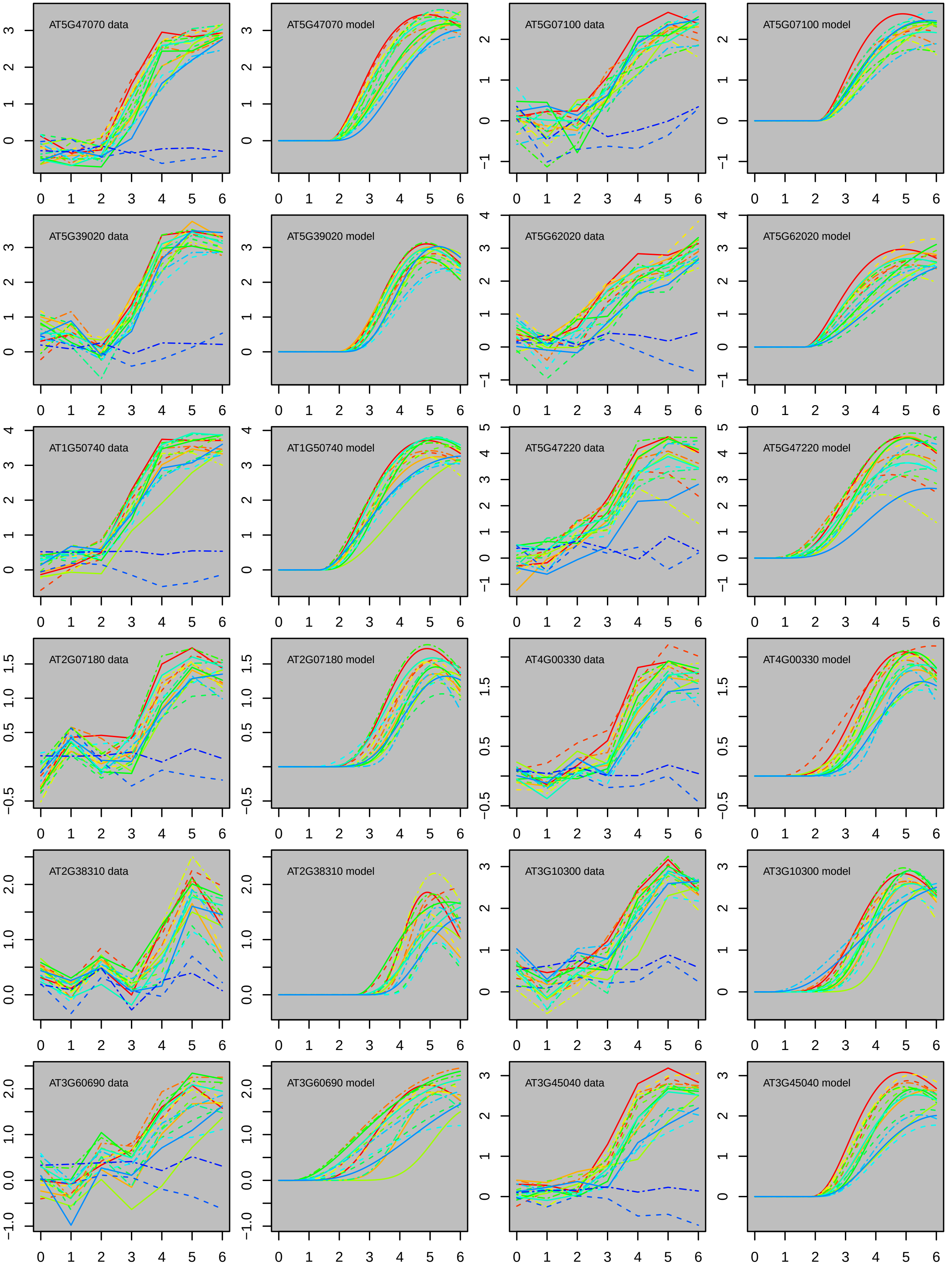
AT5G47070 data
AT5G47070 model
AT5G07100 data
AT5G07100 model
AT5G39020 data
AT5G39020 model
AT5G62020 data
AT5G62020 model
AT1G50740 data
AT1G50740 model
AT5G47220 data
AT5G47220 model
AT2G07180 data
AT2G07180 model
AT4G00330 data
AT4G00330 model
AT2G38310 data
AT2G38310 model
AT3G10300 data
AT3G10300 model
AT3G60690 data
AT3G60690 model
AT3G45040 data
AT3G45040 model

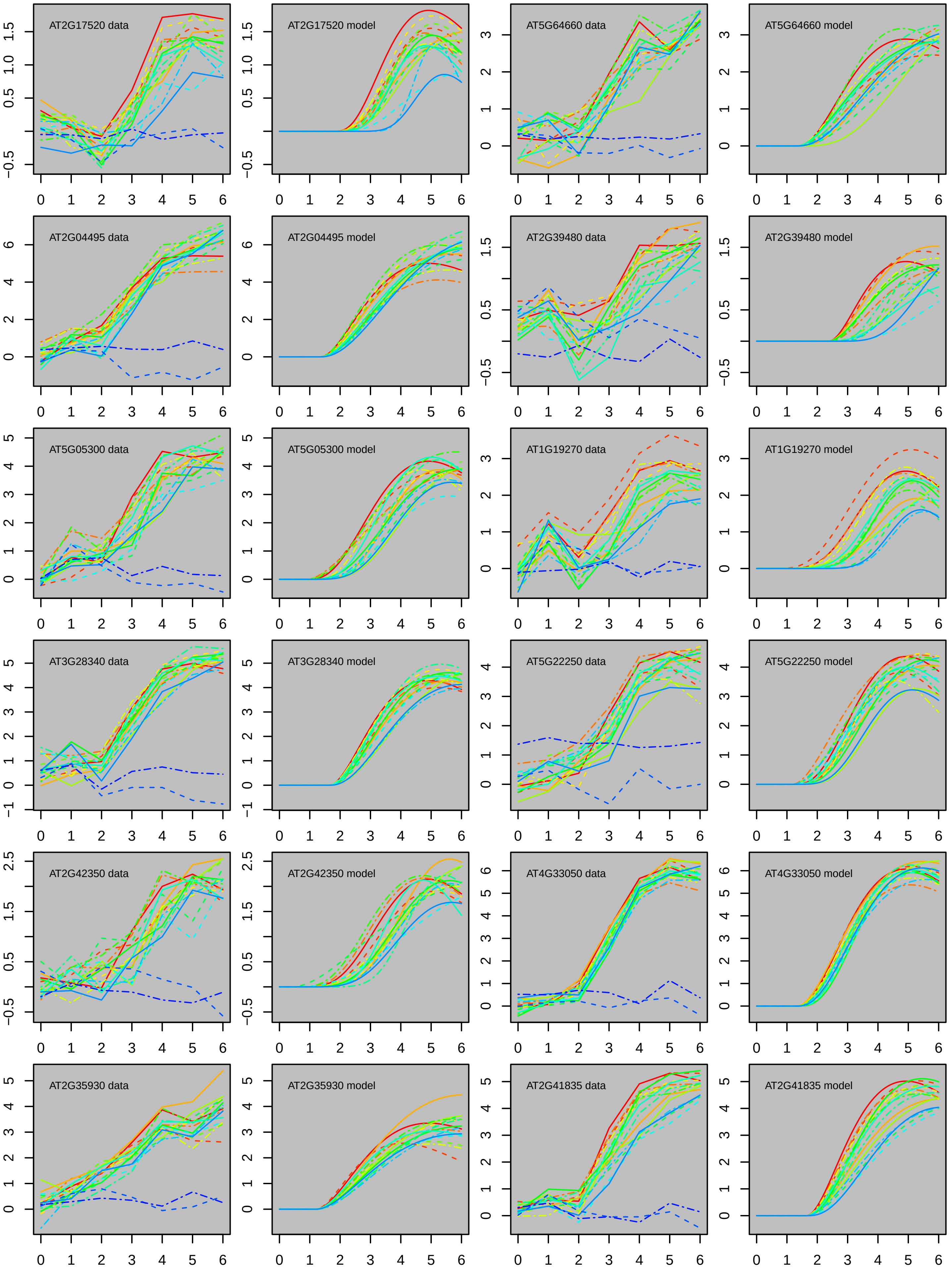
AT2G17520 data
AT2G17520 model
AT5G64660 data
AT5G64660 model
AT2G04495 data
AT2G04495 model
AT2G39480 data
AT2G39480 model
AT5G05300 data
AT5G05300 model
AT1G19270 data
AT1G19270 model
AT3G28340 data
AT3G28340 model
AT5G22250 data
AT5G22250 model
AT2G42350 data
AT2G42350 model
AT4G33050 data
AT4G33050 model
AT2G35930 data
AT2G35930 model
AT2G41835 data
AT2G41835 model

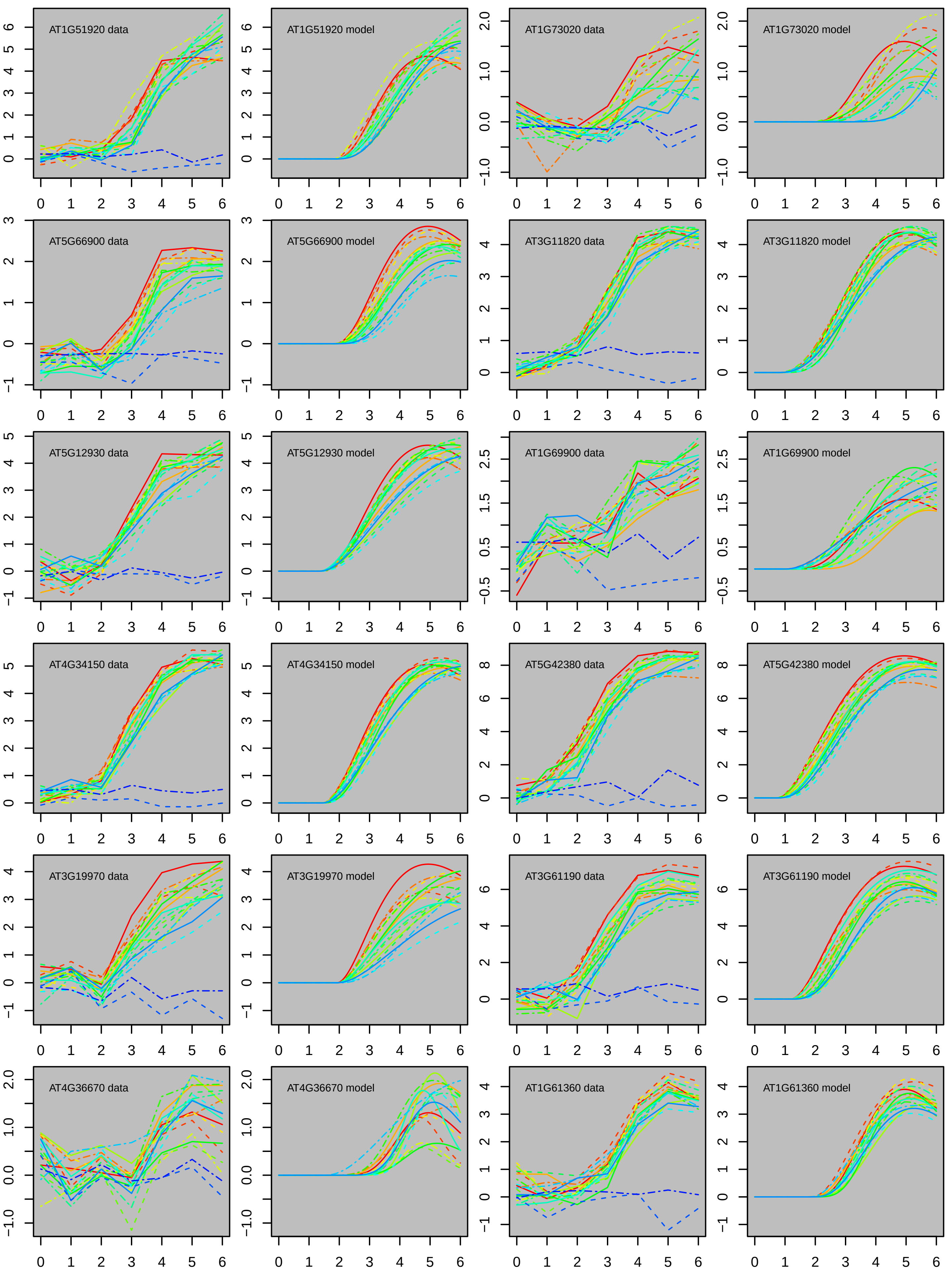

AT1G51920 data
AT1G51920 model
AT1G73020 data
AT1G73020 model
AT5G66900 data
AT5G66900 model
AT3G11820 data
AT3G11820 model
AT5G12930 data
AT5G12930 model
AT1G69900 data
AT1G69900 model
AT4G34150 data
AT4G34150 model
AT5G42380 data
AT5G42380 model
AT3G19970 data
AT3G19970 model
AT3G61190 data
AT3G61190 model
AT4G36670 data
AT4G36670 model
AT1G61360 data
AT1G61360 model

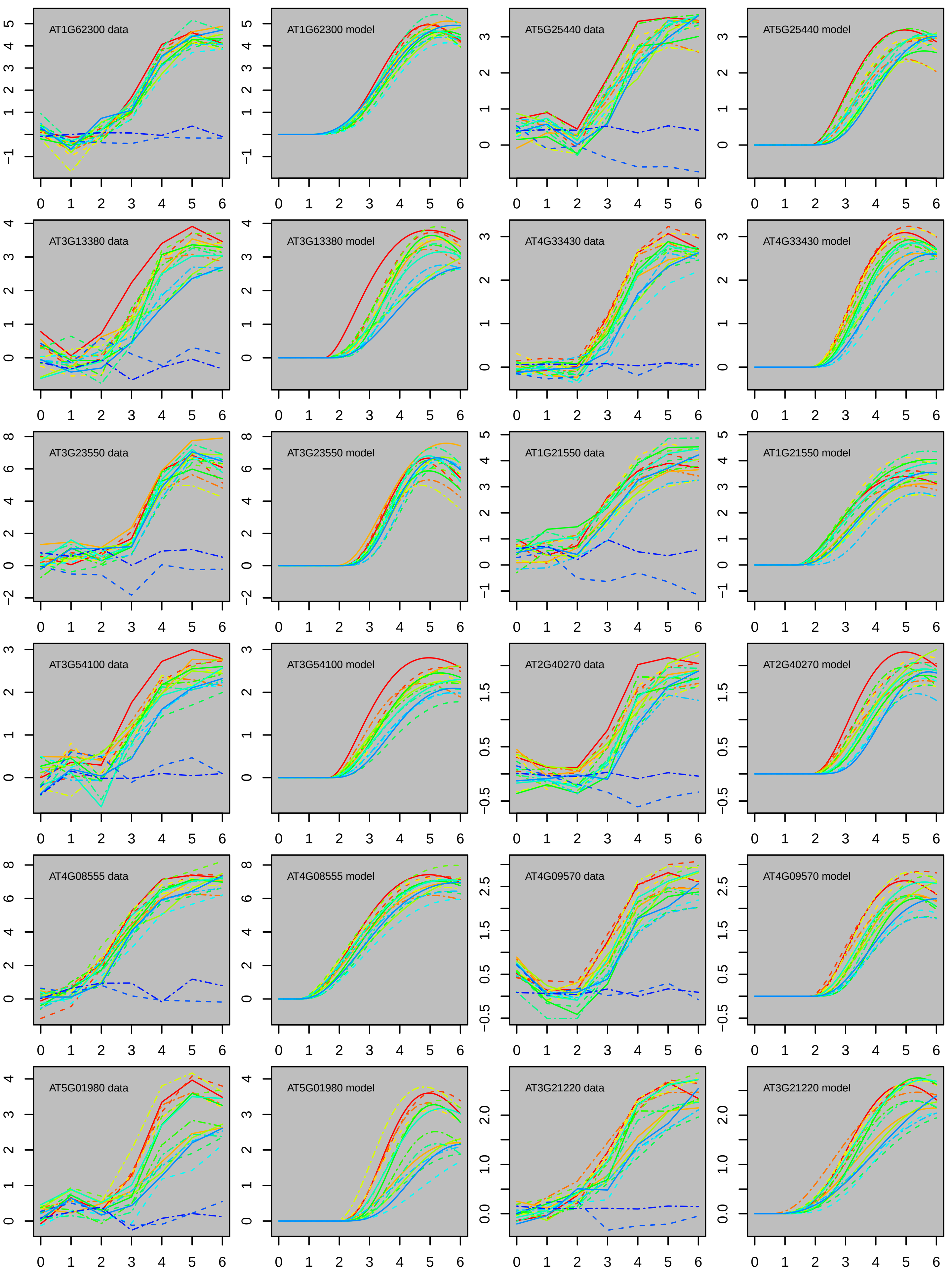
AT1G62300 data
AT1G62300 model
AT5G25440 data
AT5G25440 model
AT3G13380 data
AT3G13380 model
AT4G33430 data
AT4G33430 model
AT3G23550 data
AT3G23550 model
AT1G21550 data
AT1G21550 model
AT3G54100 data
AT3G54100 model
AT2G40270 data
AT2G40270 model
AT4G08555 data
AT4G08555 model
AT4G09570 data
AT4G09570 model
AT5G01980 data
AT5G01980 model
AT3G21220 data
AT3G21220 model

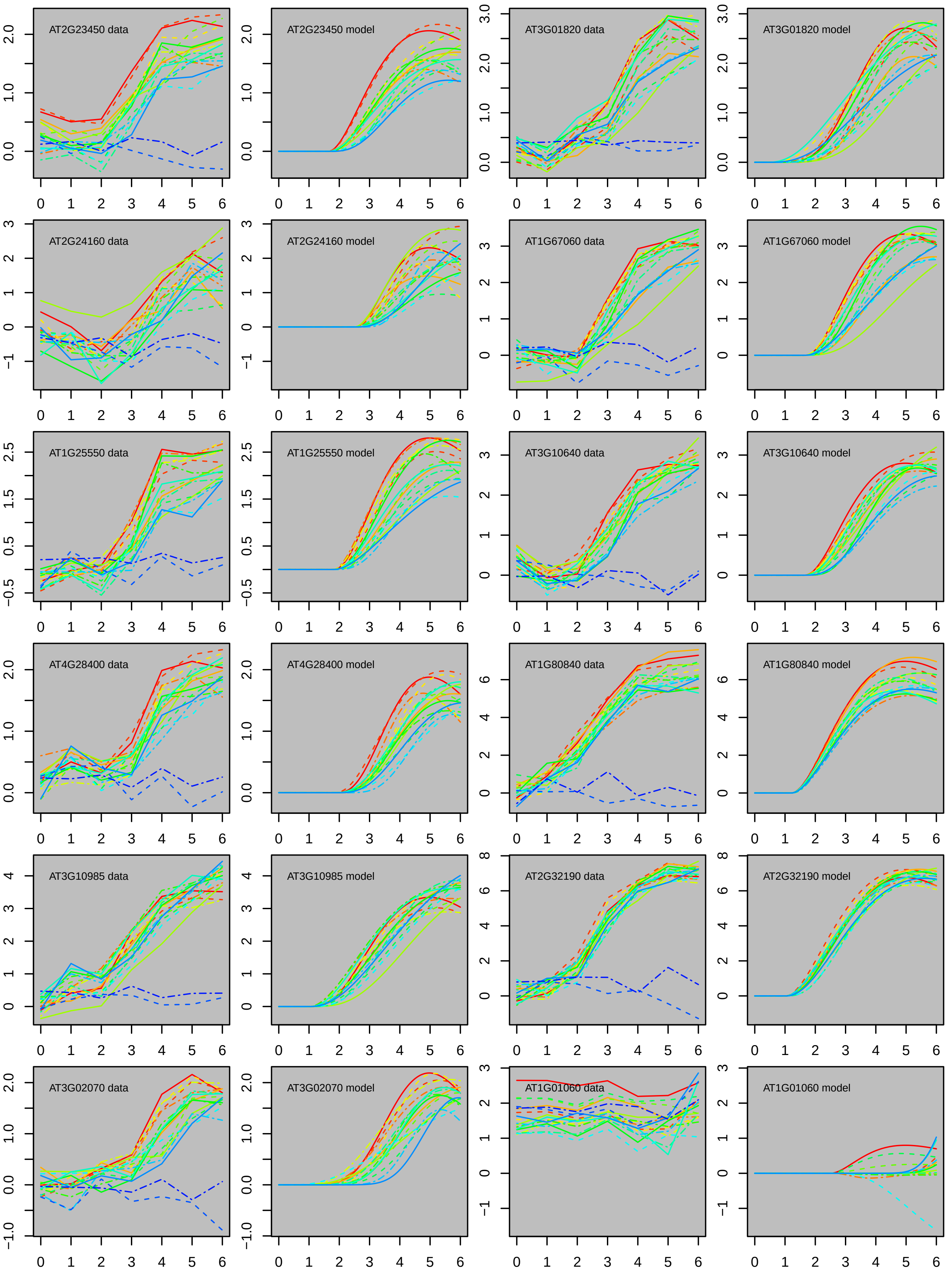
AT2G23450 data
AT2G23450 model
AT3G01820 data
AT3G01820 model
AT2G24160 data
AT2G24160 model
AT1G67060 data
AT1G67060 model
AT1G25550 data
AT1G25550 model
AT3G10640 data
AT3G10640 model
AT4G28400 data
AT4G28400 model
AT1G80840 data
AT1G80840 model
AT3G10985 data
AT3G10985 model
AT2G32190 data
AT2G32190 model
AT3G02070 data
AT3G02070 model
AT1G01060 data
AT1G01060 model

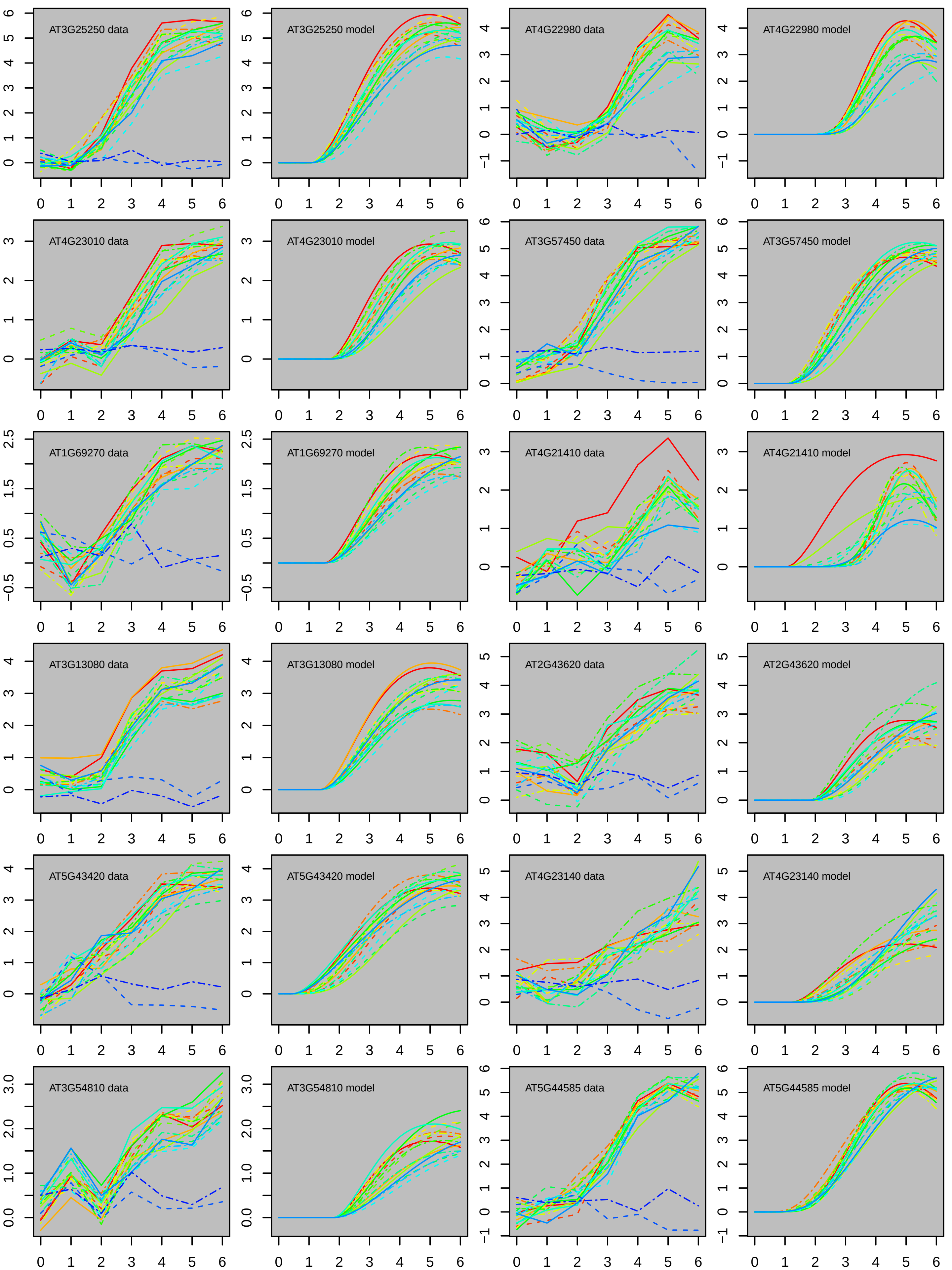
AT3G25250 data
AT3G25250 model
AT4G22980 data
AT4G22980 model
AT4G23010 data
AT4G23010 model
AT3G57450 data
AT3G57450 model
AT1G69270 data
AT1G69270 model
AT4G21410 data
AT4G21410 model
AT3G13080 data
AT3G13080 model
AT2G43620 data
AT2G43620 model
AT5G43420 data
AT5G43420 model
AT4G23140 data
AT4G23140 model
AT3G54810 data
AT3G54810 model
AT5G44585 data
AT5G44585 model

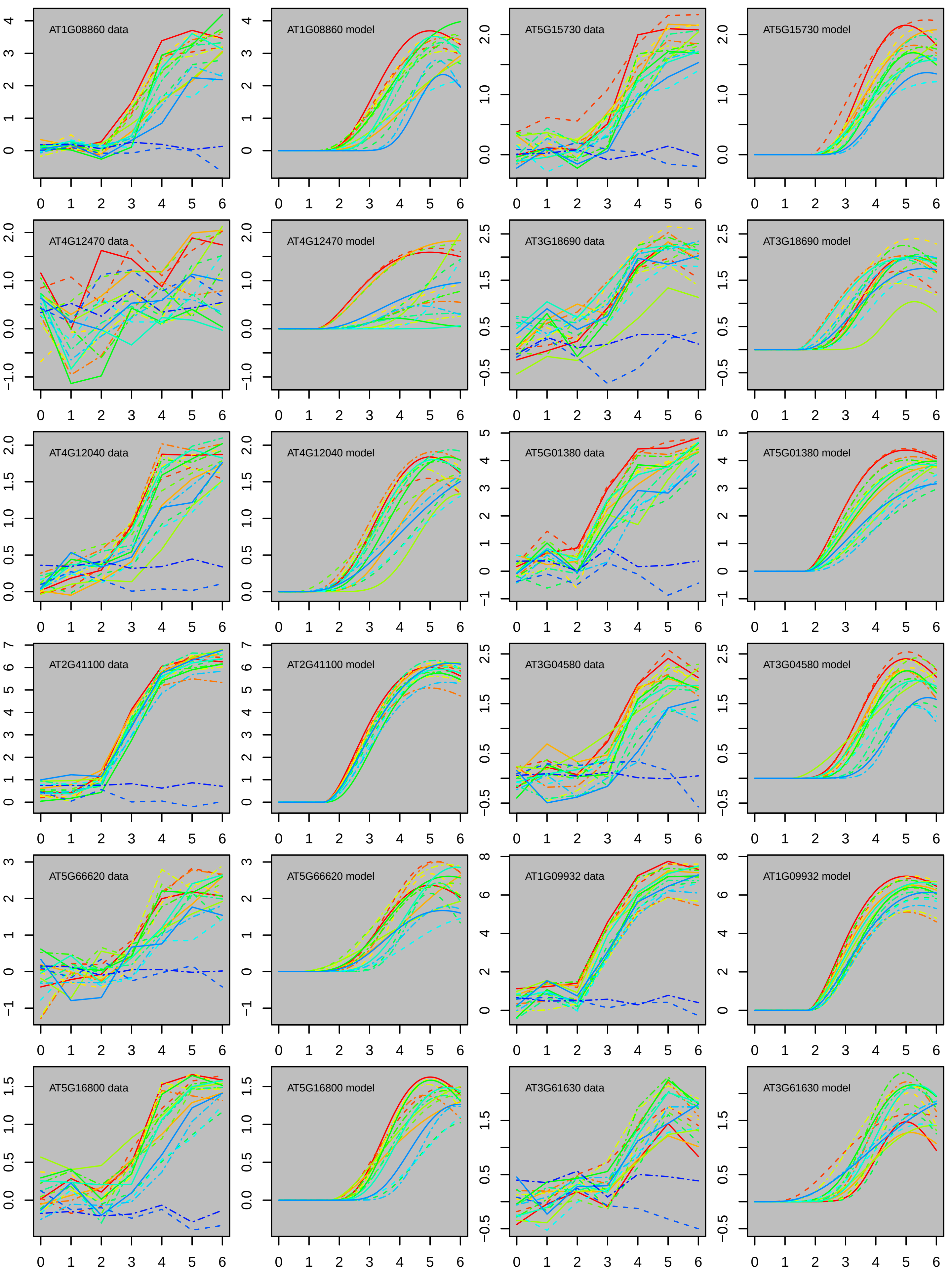
AT1G08860 data
AT1G08860 model
AT5G15730 data
AT5G15730 model
AT4G12470 data
AT4G12470 model
AT3G18690 data
AT3G18690 model
AT4G12040 data
AT4G12040 model
AT5G01380 data
AT5G01380 model
AT2G41100 data
AT2G41100 model
AT3G04580 data
AT3G04580 model
AT5G66620 data
AT5G66620 model
AT1G09932 data
AT1G09932 model
AT5G16800 data
AT5G16800 model
AT3G61630 data
AT3G61630 model

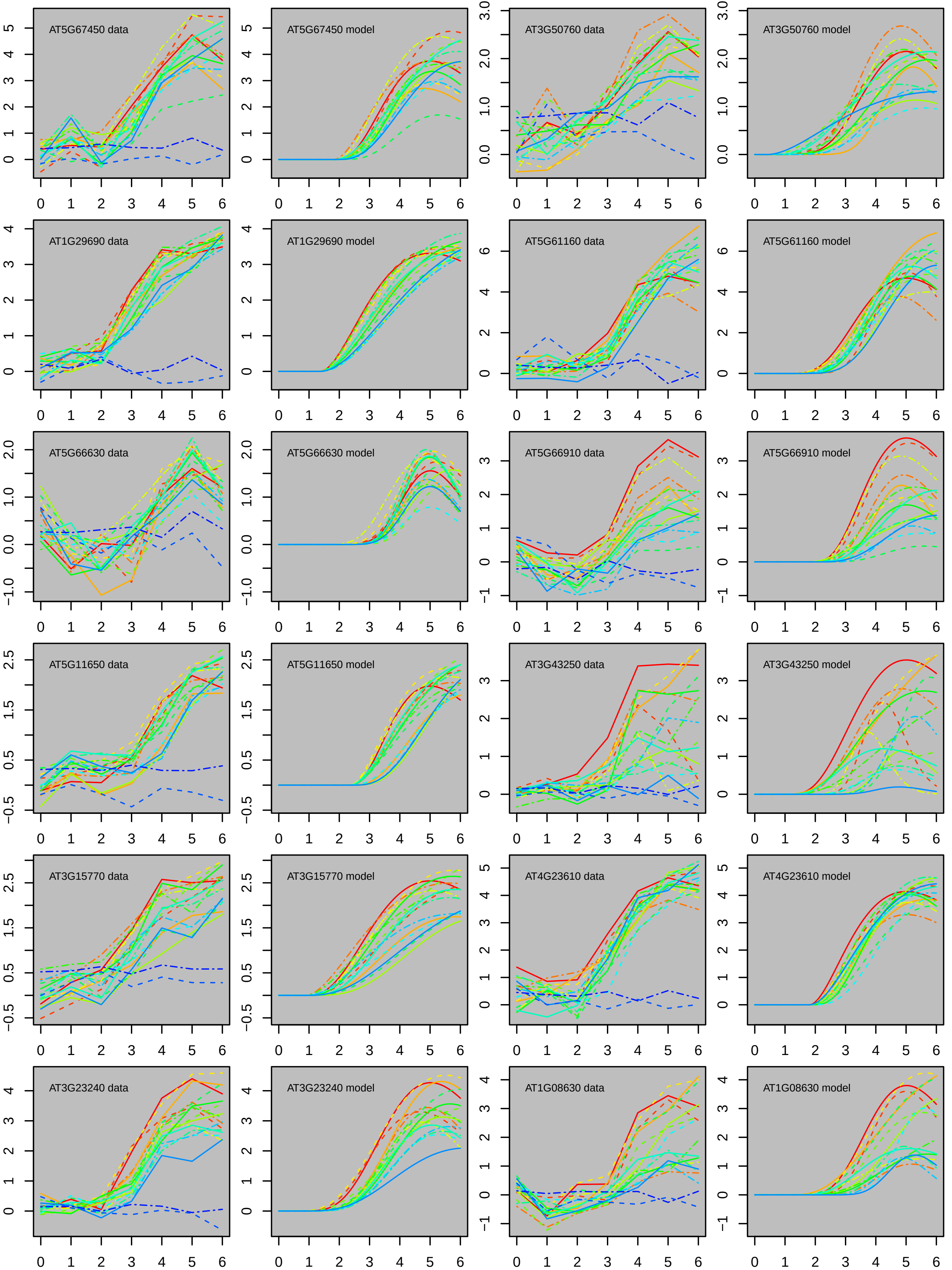

AT5G67450 data
AT5G67450 model
AT3G50760 data
AT3G50760 model
AT1G29690 data
AT1G29690 model
AT5G61160 data
AT5G61160 model
AT5G66630 data
AT5G66630 model
AT5G66910 data
AT5G66910 model
AT5G11650 data
AT5G11650 model
AT3G43250 data
AT3G43250 model
AT3G15770 data
AT3G15770 model
AT4G23610 data
AT4G23610 model
AT3G23240 data
AT3G23240 model
AT1G08630 data
AT1G08630 model

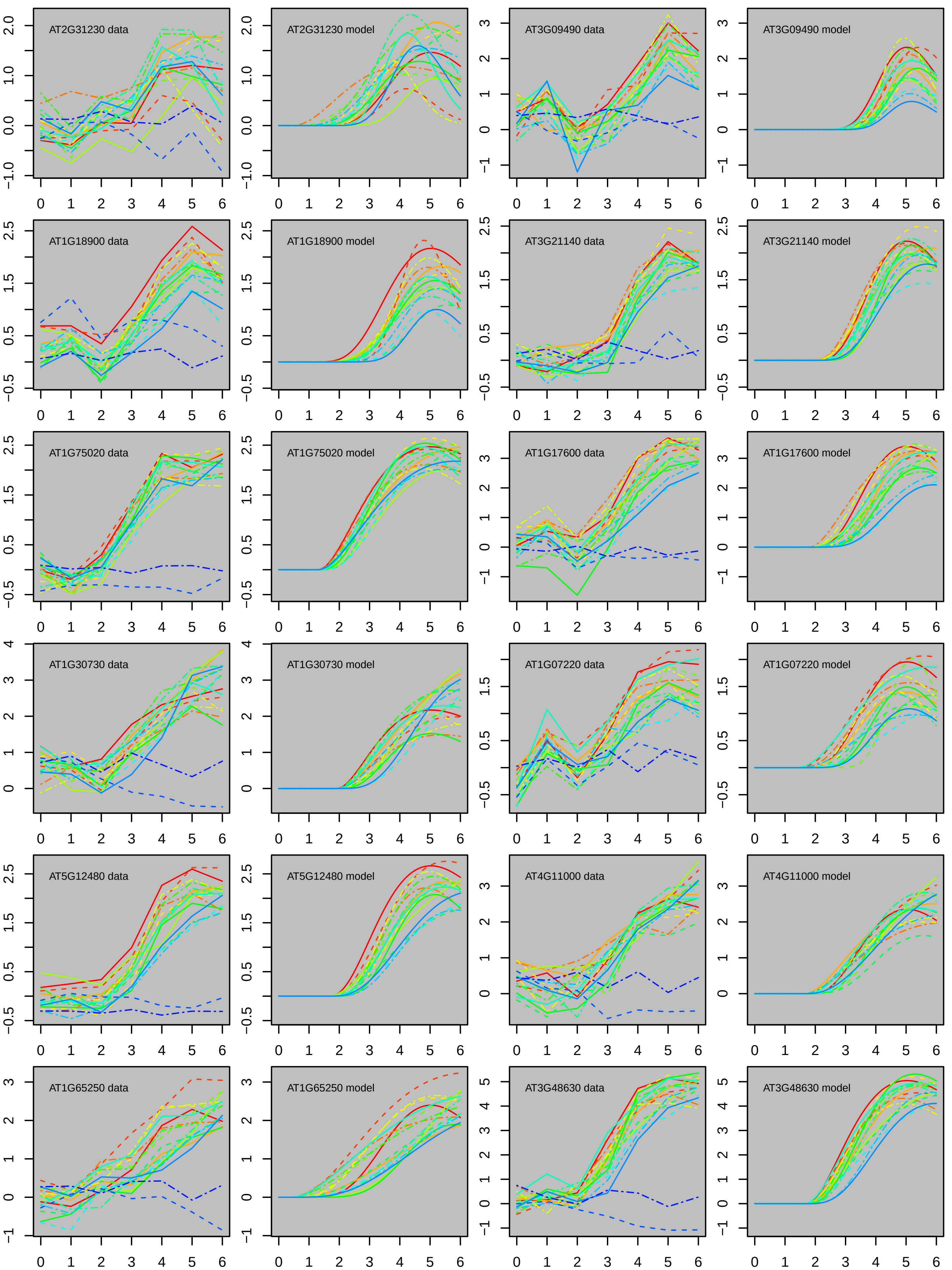
AT2G31230 data
AT2G31230 model
AT3G09490 data
AT3G09490 model
AT1G18900 data
AT1G18900 model
AT3G21140 data
AT3G21140 model
AT1G75020 data
AT1G75020 model
AT1G17600 data
AT1G17600 model
AT1G30730 data
AT1G30730 model
AT1G07220 data
AT1G07220 model
AT5G12480 data
AT5G12480 model
AT4G11000 data
AT4G11000 model
AT1G65250 data
AT1G65250 model
AT3G48630 data
AT3G48630 model

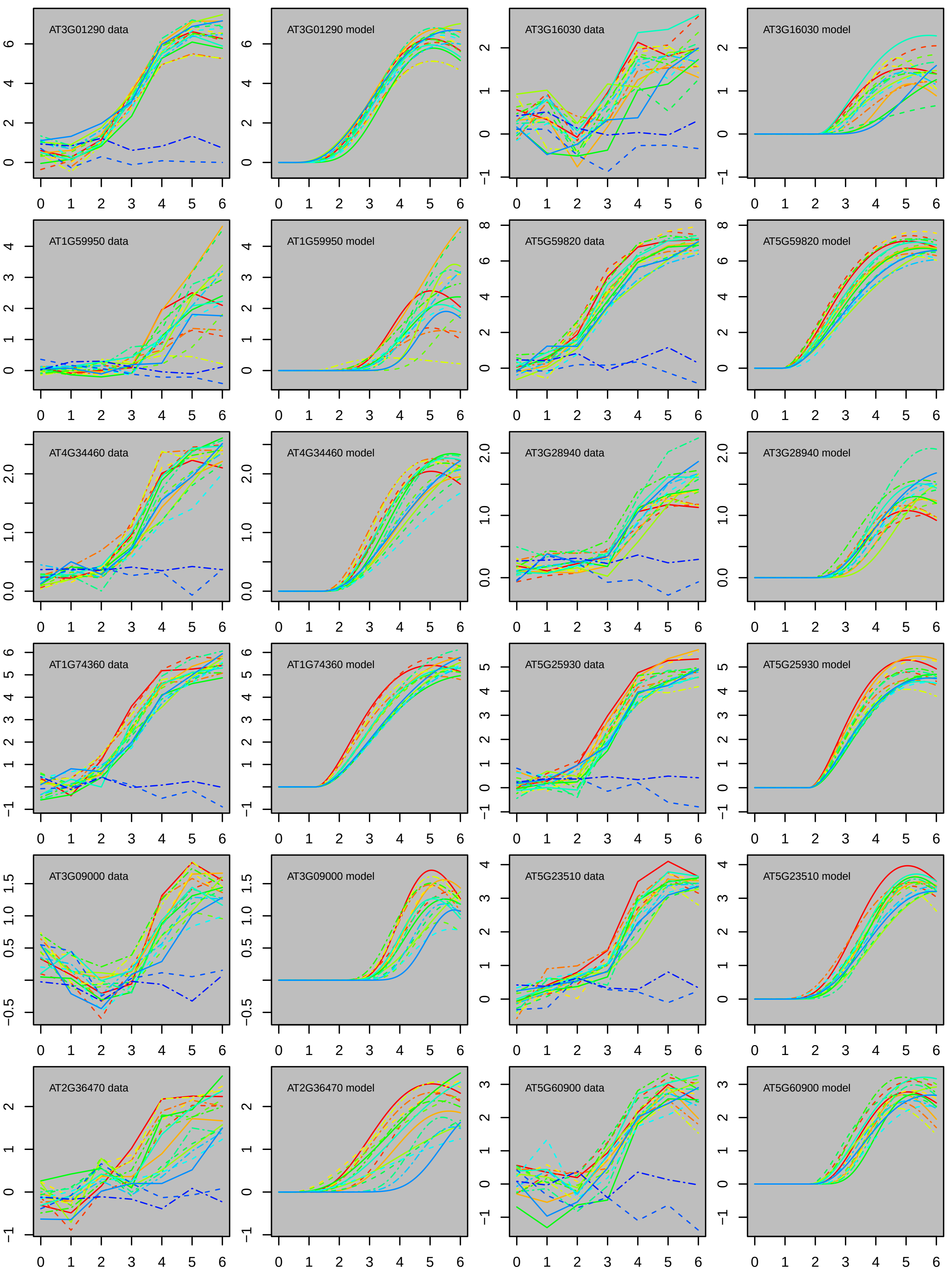
AT3G01290 data
AT3G01290 model
AT3G16030 data
AT3G16030 model
AT1G59950 data
AT1G59950 model
AT5G59820 data
AT5G59820 model
AT4G34460 data
AT4G34460 model
AT3G28940 data
AT3G28940 model
AT1G74360 data
AT1G74360 model
AT5G25930 data
AT5G25930 model
AT3G09000 data
AT3G09000 model
AT5G23510 data
AT5G23510 model
AT2G36470 data
AT2G36470 model
AT5G60900 data
AT5G60900 model

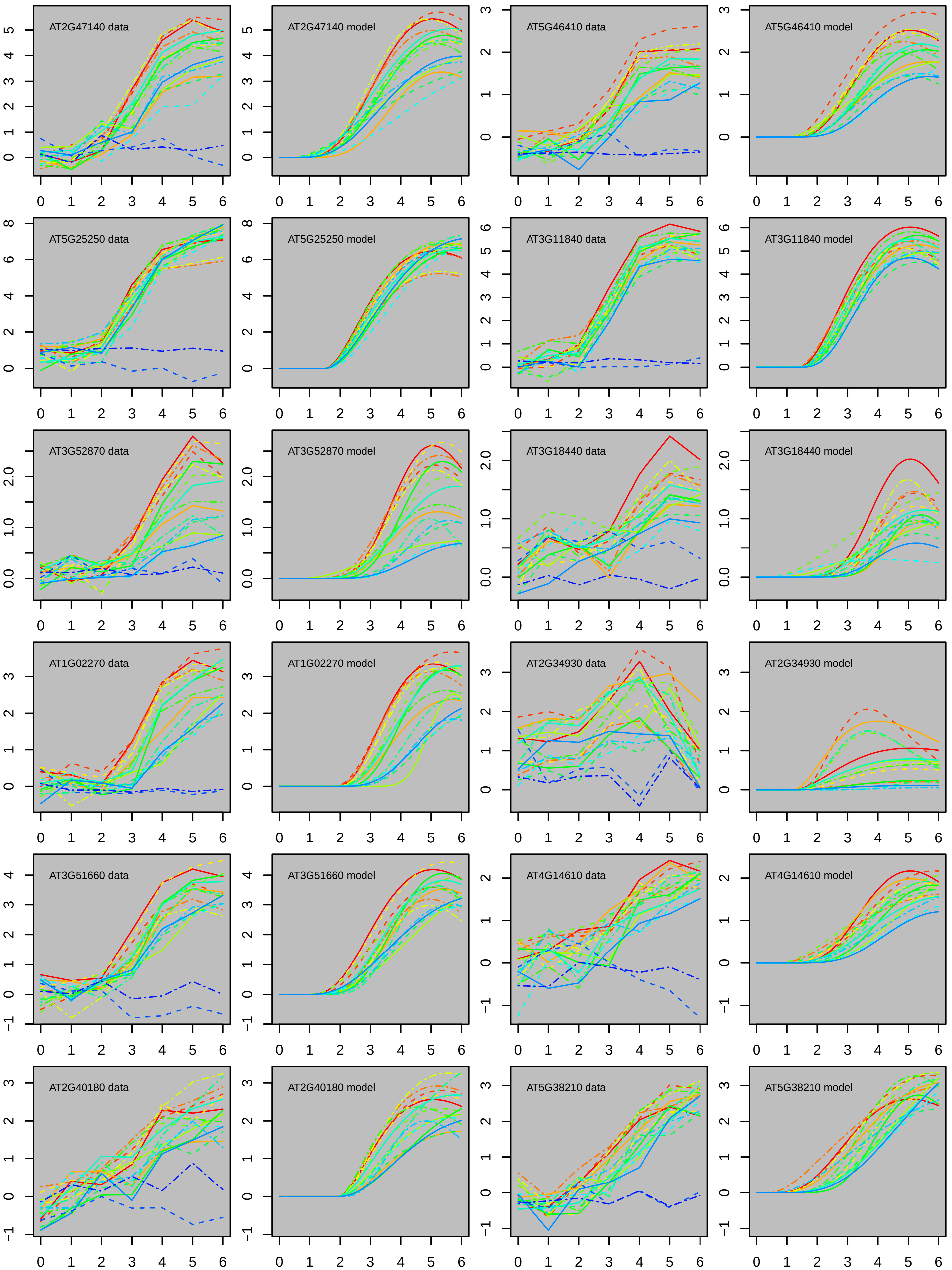
AT2G47140 data
AT2G47140 model
AT5G46410 data
AT5G46410 model
AT5G25250 data
AT5G25250 model
AT3G11840 data
AT3G11840 model
AT3G52870 data
AT3G52870 model
AT3G18440 data
AT3G18440 model
AT1G02270 data
AT1G02270 model
AT2G34930 data
AT2G34930 model
AT3G51660 data
AT3G51660 model
AT4G14610 data
AT4G14610 model
AT2G40180 data
AT2G40180 model
AT5G38210 data
AT5G38210 model

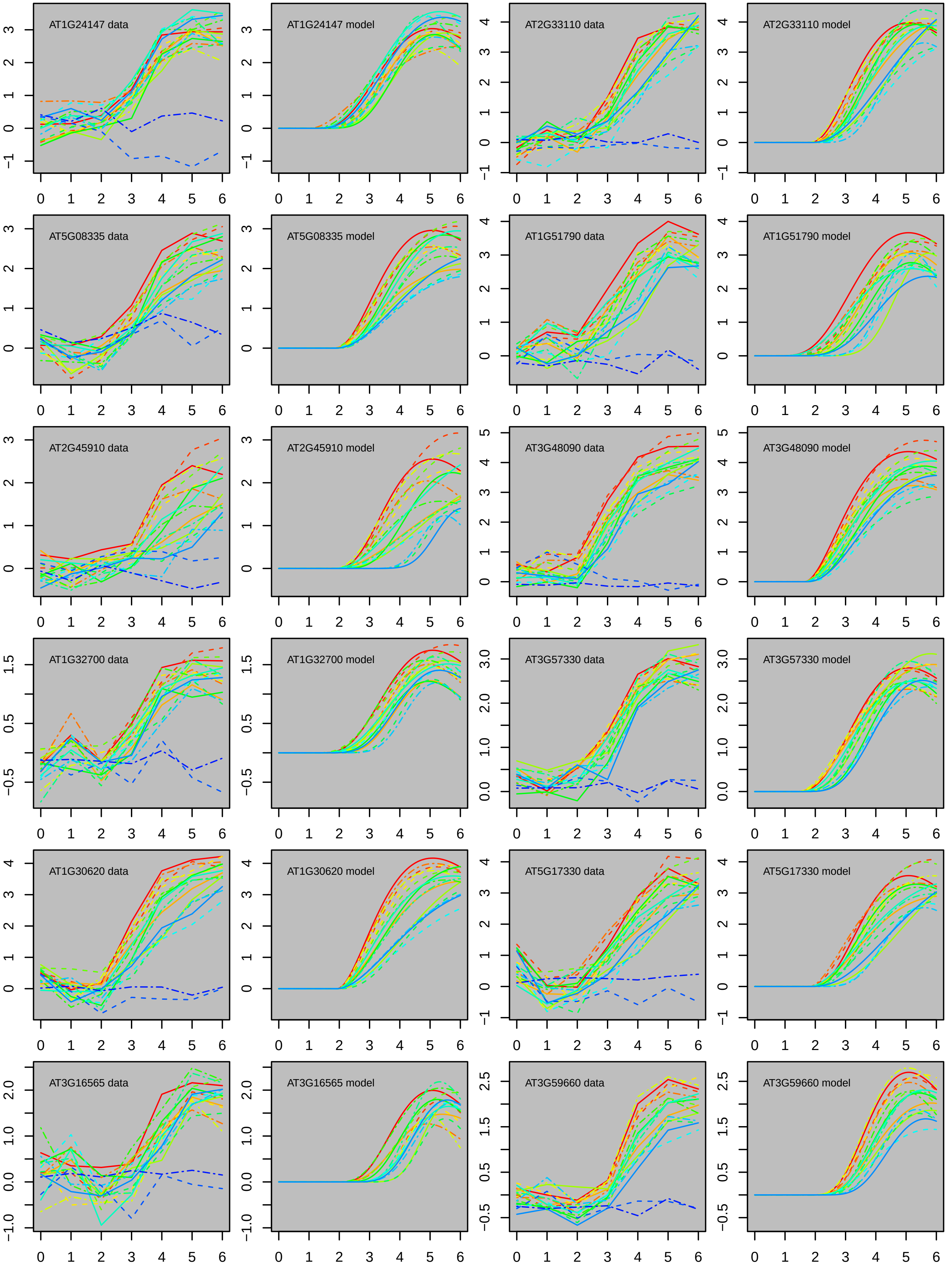
AT1G24147 data
AT1G24147 model
AT2G33110 data
AT2G33110 model
AT5G08335 data
AT5G08335 model
AT1G51790 data
AT1G51790 model
AT2G45910 data
AT2G45910 model
AT3G48090 data
AT3G48090 model
AT1G32700 data
AT1G32700 model
AT3G57330 data
AT3G57330 model
AT1G30620 data
AT1G30620 model
AT5G17330 data
AT5G17330 model
AT3G16565 data
AT3G16565 model
AT3G59660 data
AT3G59660 model

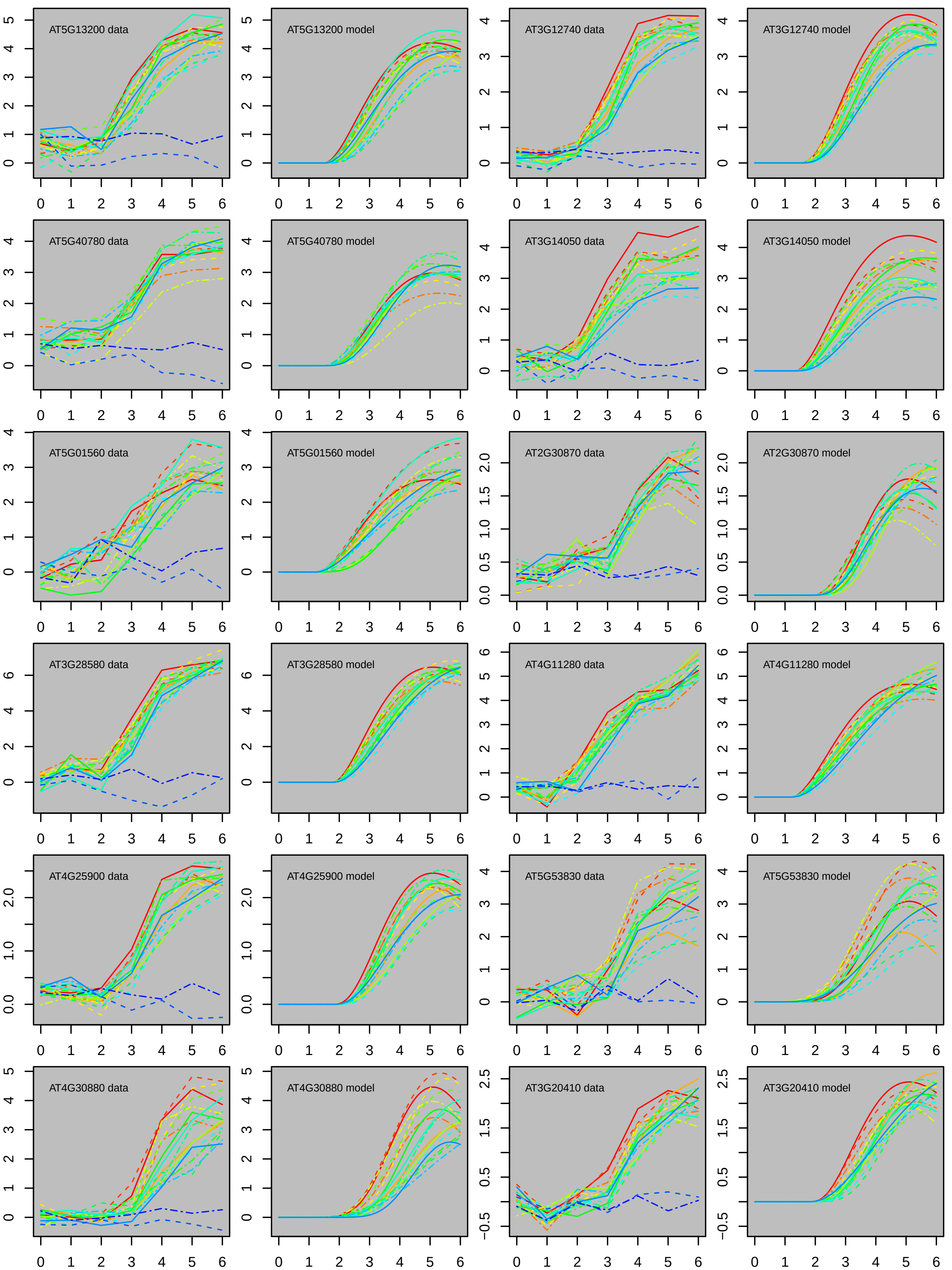
AT5G13200 data
AT5G13200 model
AT3G12740 data
AT3G12740 model
AT5G40780 data
AT5G40780 model
AT3G14050 data
AT3G14050 model
AT5G01560 data
AT5G01560 model
AT2G30870 data
AT2G30870 model
AT3G28580 data
AT3G28580 model
AT4G11280 data
AT4G11280 model
AT4G25900 data
AT4G25900 model
AT5G53830 data
AT5G53830 model
AT4G30880 data
AT4G30880 model
AT3G20410 data
AT3G20410 model

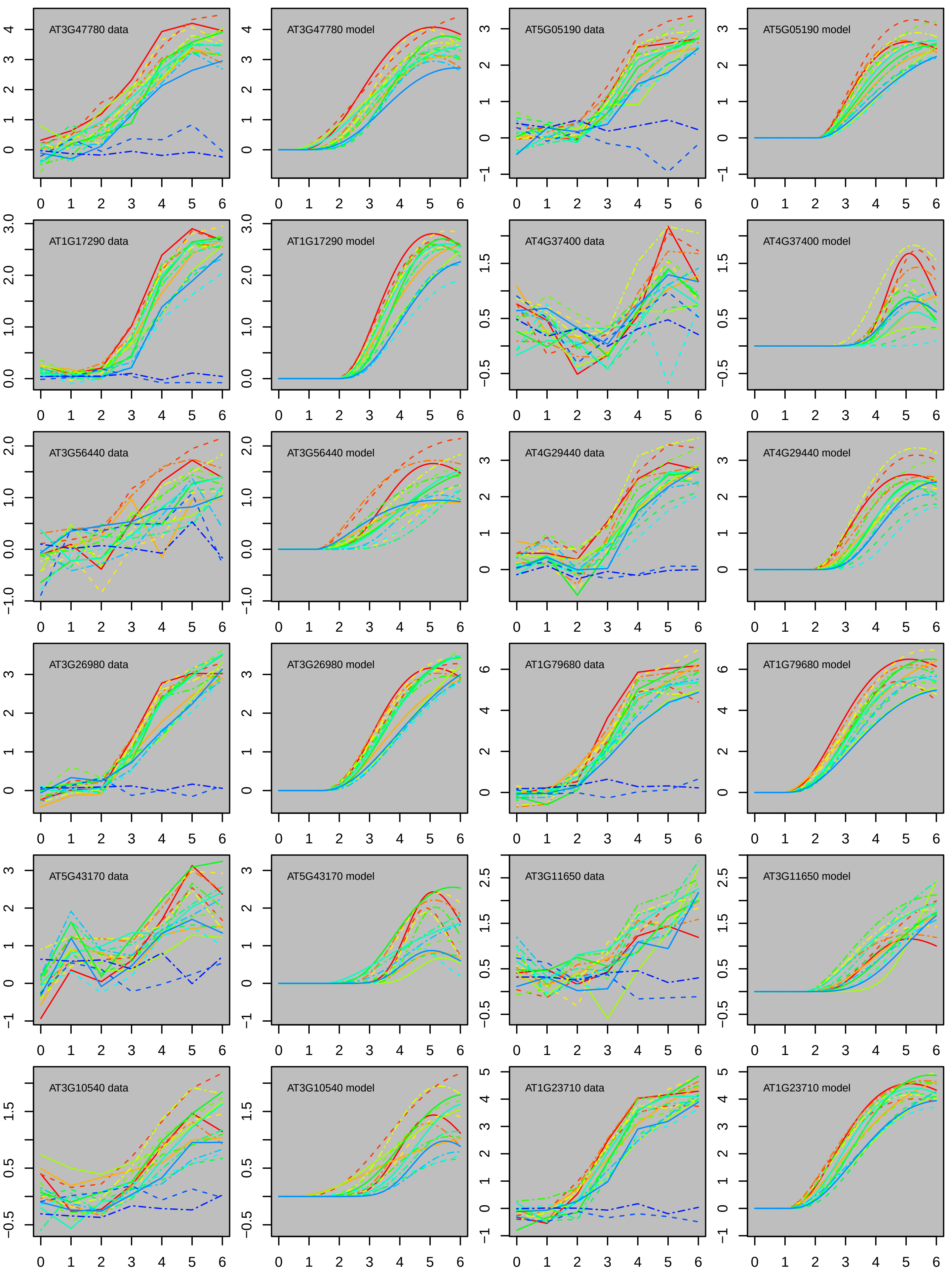

AT3G47780 data
AT3G47780 model
AT5G05190 data
AT5G05190 model
AT1G17290 data
AT1G17290 model
AT4G37400 data
AT4G37400 model
AT3G56440 data
AT3G56440 model
AT4G29440 data
AT4G29440 model
AT3G26980 data
AT3G26980 model
AT1G79680 data
AT1G79680 model
AT5G43170 data
AT5G43170 model
AT3G11650 data
AT3G11650 model
AT3G10540 data
AT3G10540 model
AT1G23710 data
AT1G23710 model

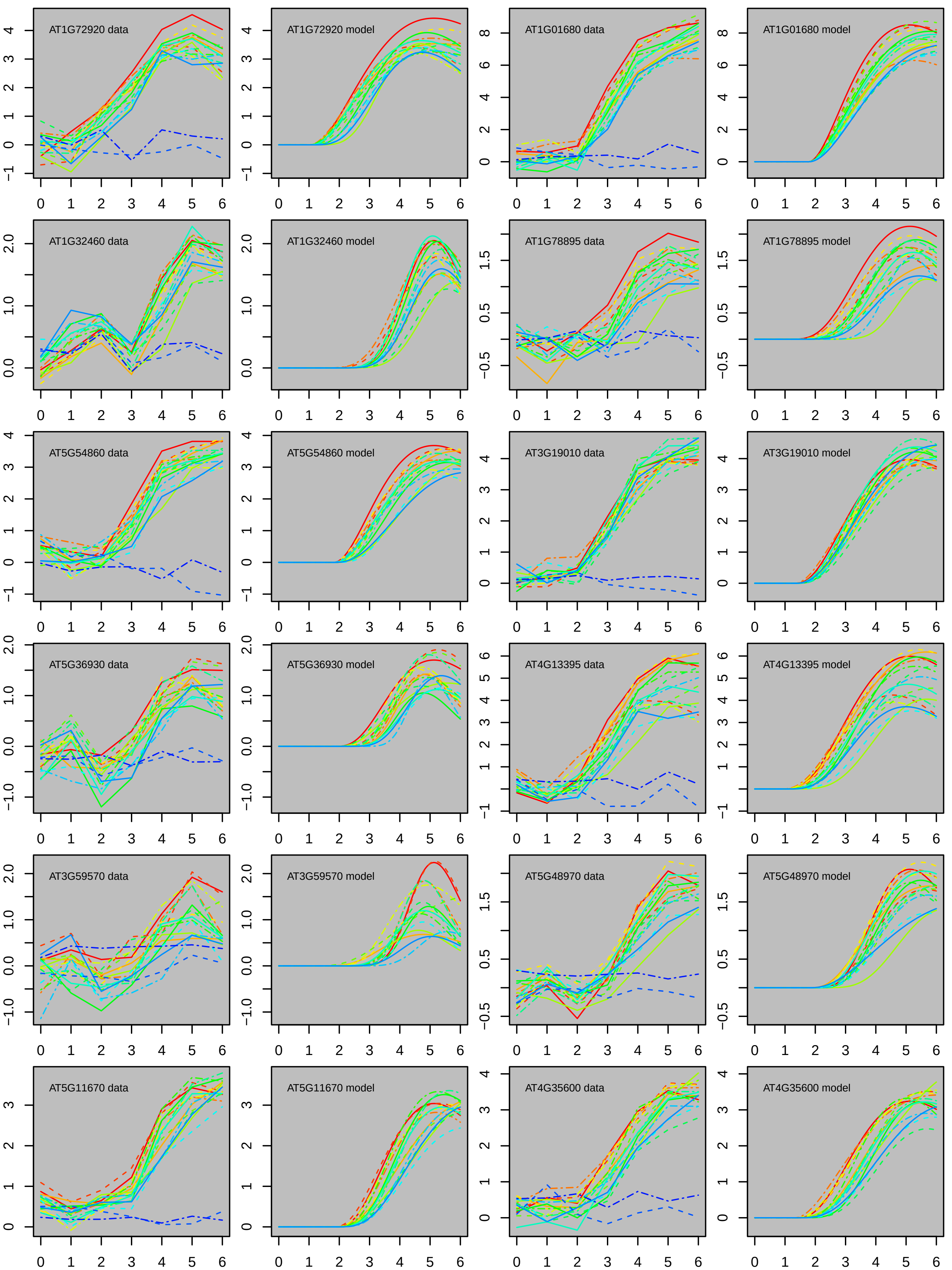
AT1G72920 data
AT1G72920 model
AT1G01680 data
AT1G01680 model
AT1G32460 data
AT1G32460 model
AT1G78895 data
AT1G78895 model
AT5G54860 data
AT5G54860 model
AT3G19010 data
AT3G19010 model
AT5G36930 data
AT5G36930 model
AT4G13395 data
AT4G13395 model
AT3G59570 data
AT3G59570 model
AT5G48970 data
AT5G48970 model
AT5G11670 data
AT5G11670 model
AT4G35600 data
AT4G35600 model

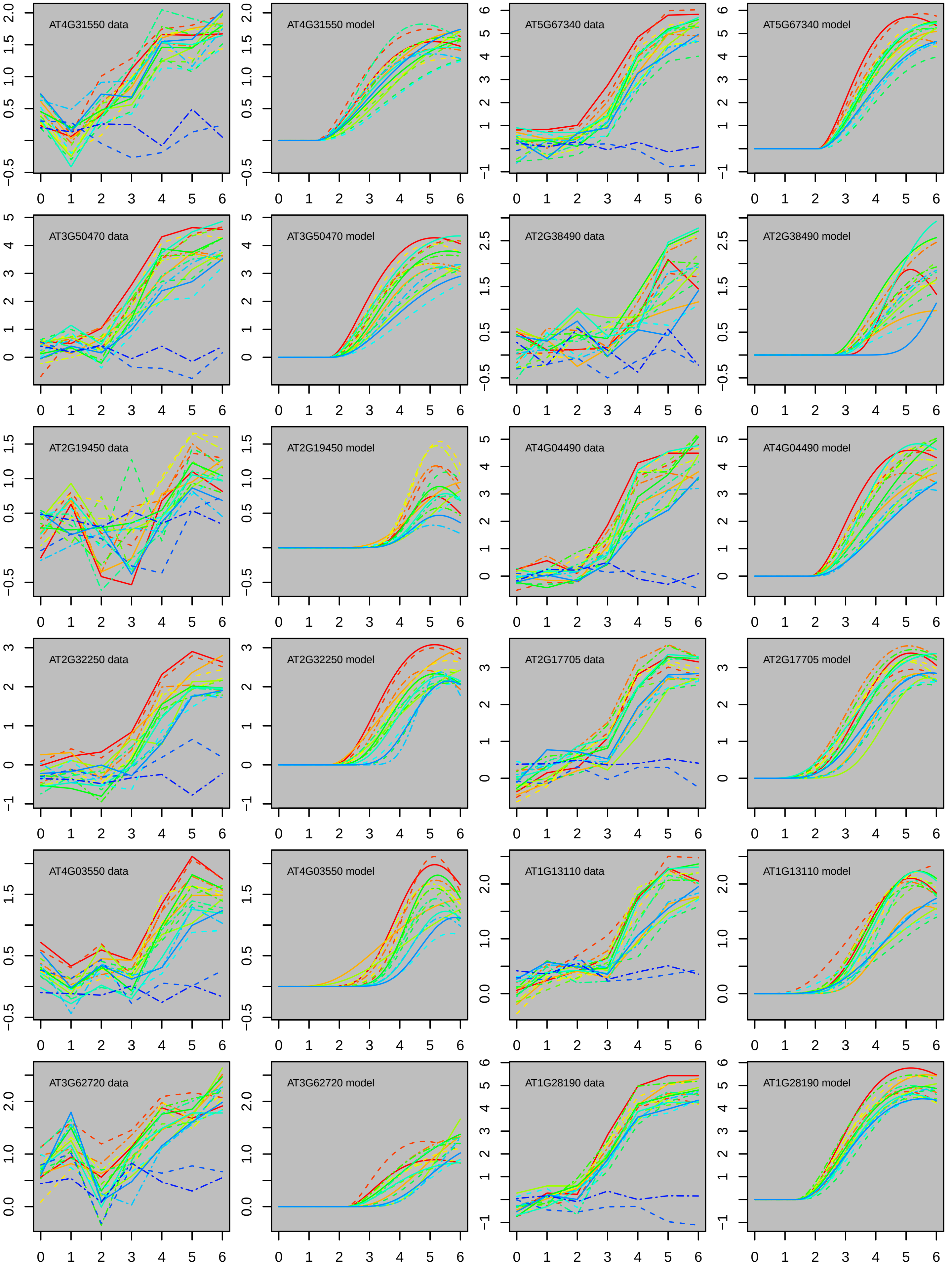

AT4G31550 data
AT4G31550 model
AT5G67340 data
AT5G67340 model
AT3G50470 data
AT3G50470 model
AT2G38490 data
AT2G38490 model
AT2G19450 data
AT2G19450 model
AT4G04490 data
AT4G04490 model
AT2G32250 data
AT2G32250 model
AT2G17705 data
AT2G17705 model
AT4G03550 data
AT4G03550 model
AT1G13110 data
AT1G13110 model
AT3G62720 data
AT3G62720 model
AT1G28190 data
AT1G28190 model
0 1 2 3 4 5 6

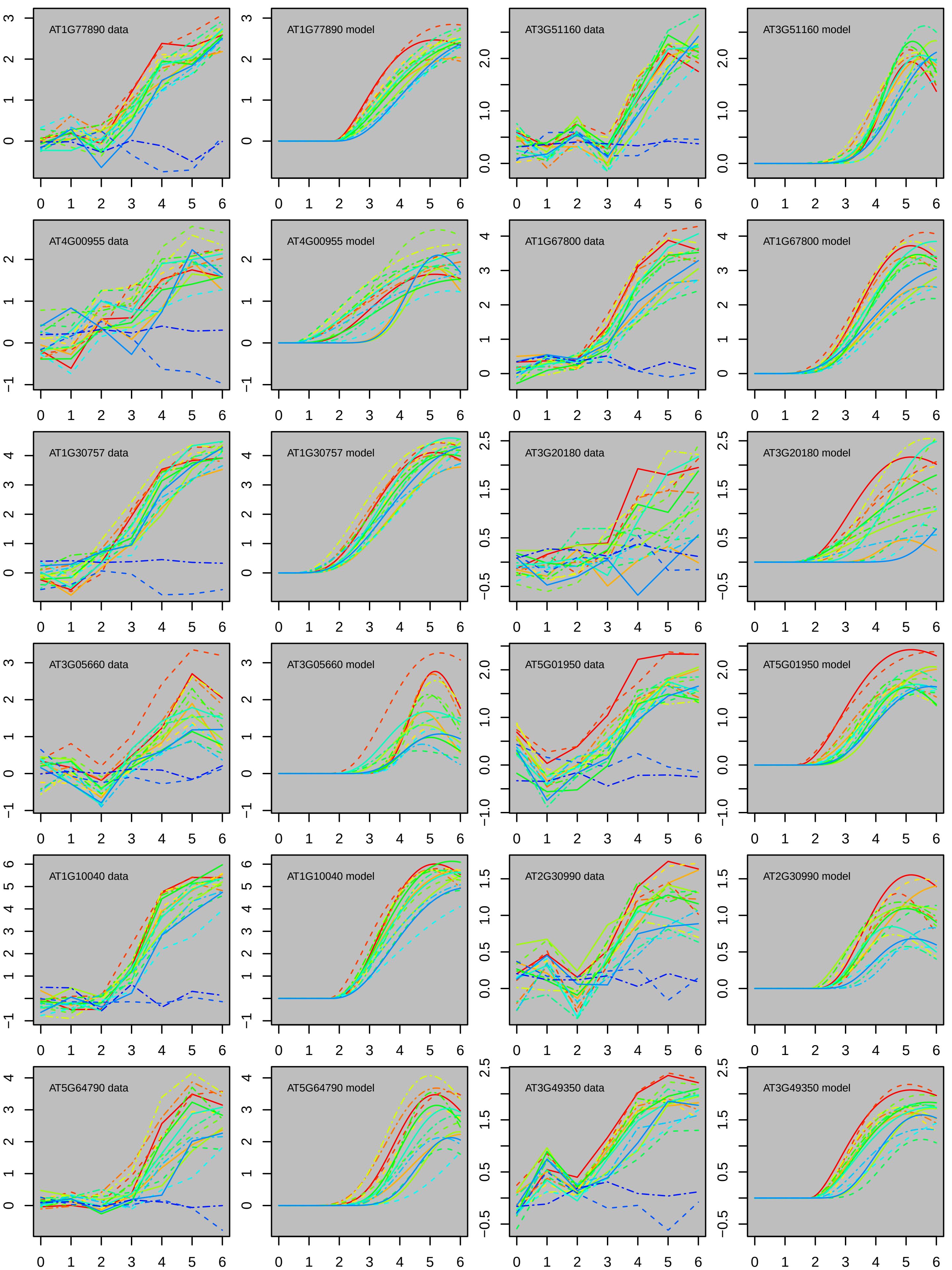
AT1G77890 data
AT1G77890 model
AT3G51160 data
AT3G51160 model
AT4G00955 data
AT4G00955 model
AT1G67800 data
AT1G67800 model
AT1G30757 data
AT1G30757 model
AT3G20180 data
AT3G20180 model
AT3G05660 data
AT3G05660 model
AT5G01950 data
AT5G01950 model
AT1G10040 data
AT1G10040 model
AT2G30990 data
AT2G30990 model
AT5G64790 data
AT5G64790 model
AT3G49350 data
AT3G49350 model

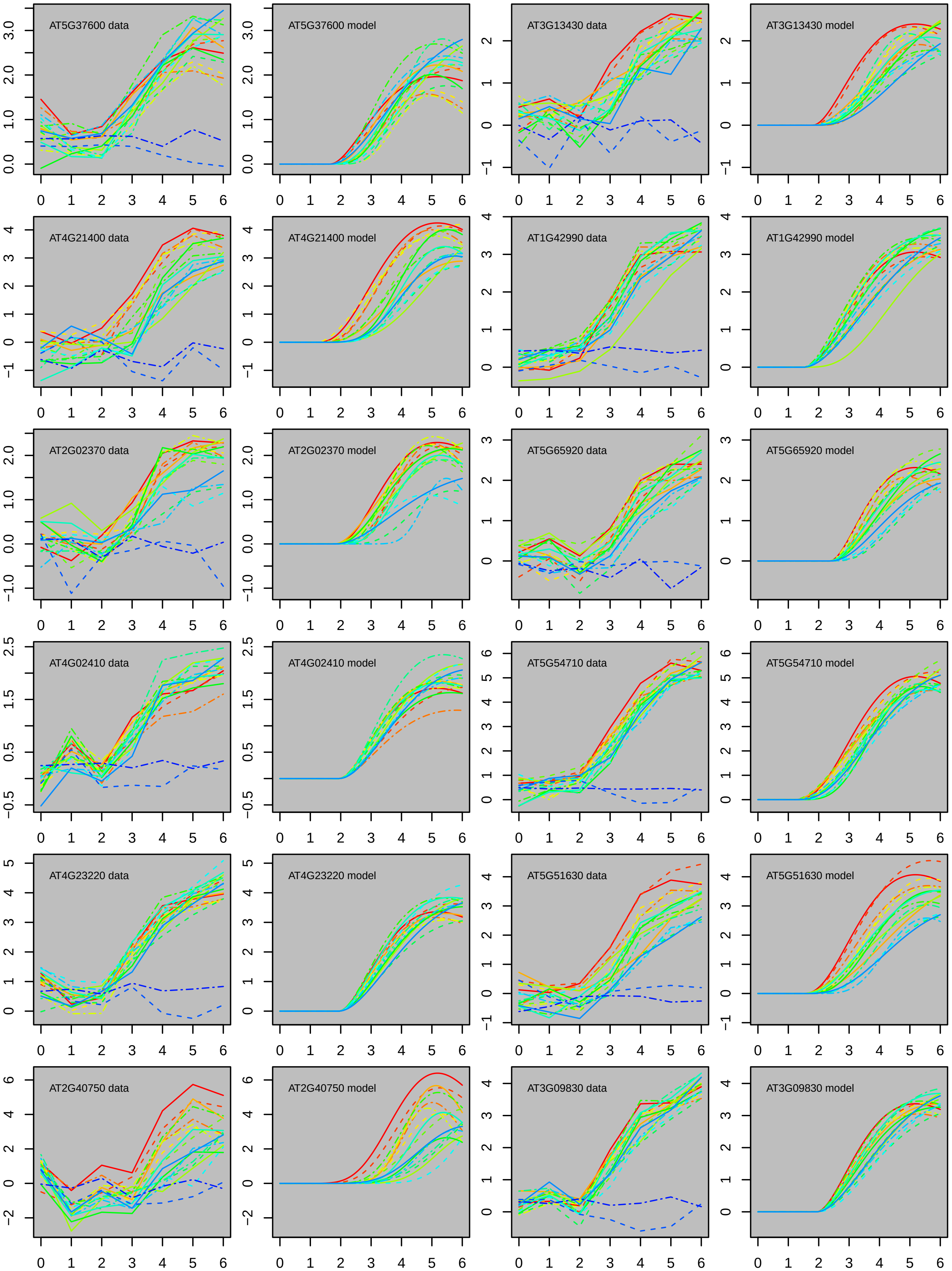
AT5G37600 data
AT5G37600 model
AT3G13430 data
AT3G13430 model
AT4G21400 data
AT4G21400 model
AT1G42990 data
AT1G42990 model
AT2G02370 data
AT2G02370 model
AT5G65920 data
AT5G65920 model
AT4G02410 data
AT4G02410 model
AT5G54710 data
AT5G54710 model
AT4G23220 data
AT4G23220 model
AT5G51630 data
AT5G51630 model
AT2G40750 data
AT2G40750 model
AT3G09830 data
AT3G09830 model

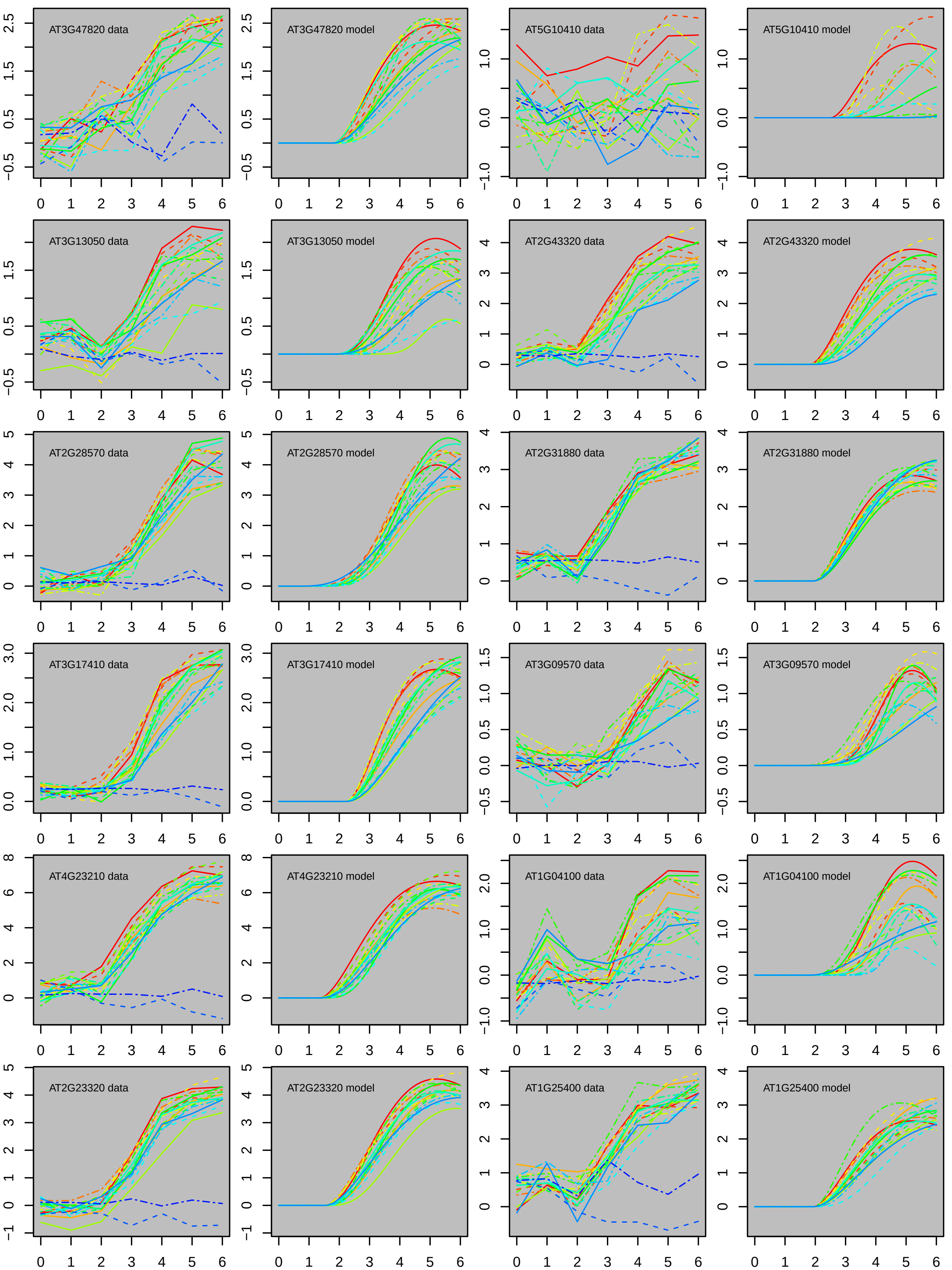
AT3G47820 data
AT3G47820 model
AT5G10410 data
AT5G10410 model
AT3G13050 data
AT3G13050 model
AT2G43320 data
AT2G43320 model
AT2G28570 data
AT2G28570 model
AT2G31880 data
AT2G31880 model
AT3G17410 data
AT3G17410 model
AT3G09570 data
AT3G09570 model
AT4G23210 data
AT4G23210 model
AT1G04100 data
AT1G04100 model
AT2G23320 data
AT2G23320 model
AT1G25400 data
AT1G25400 model

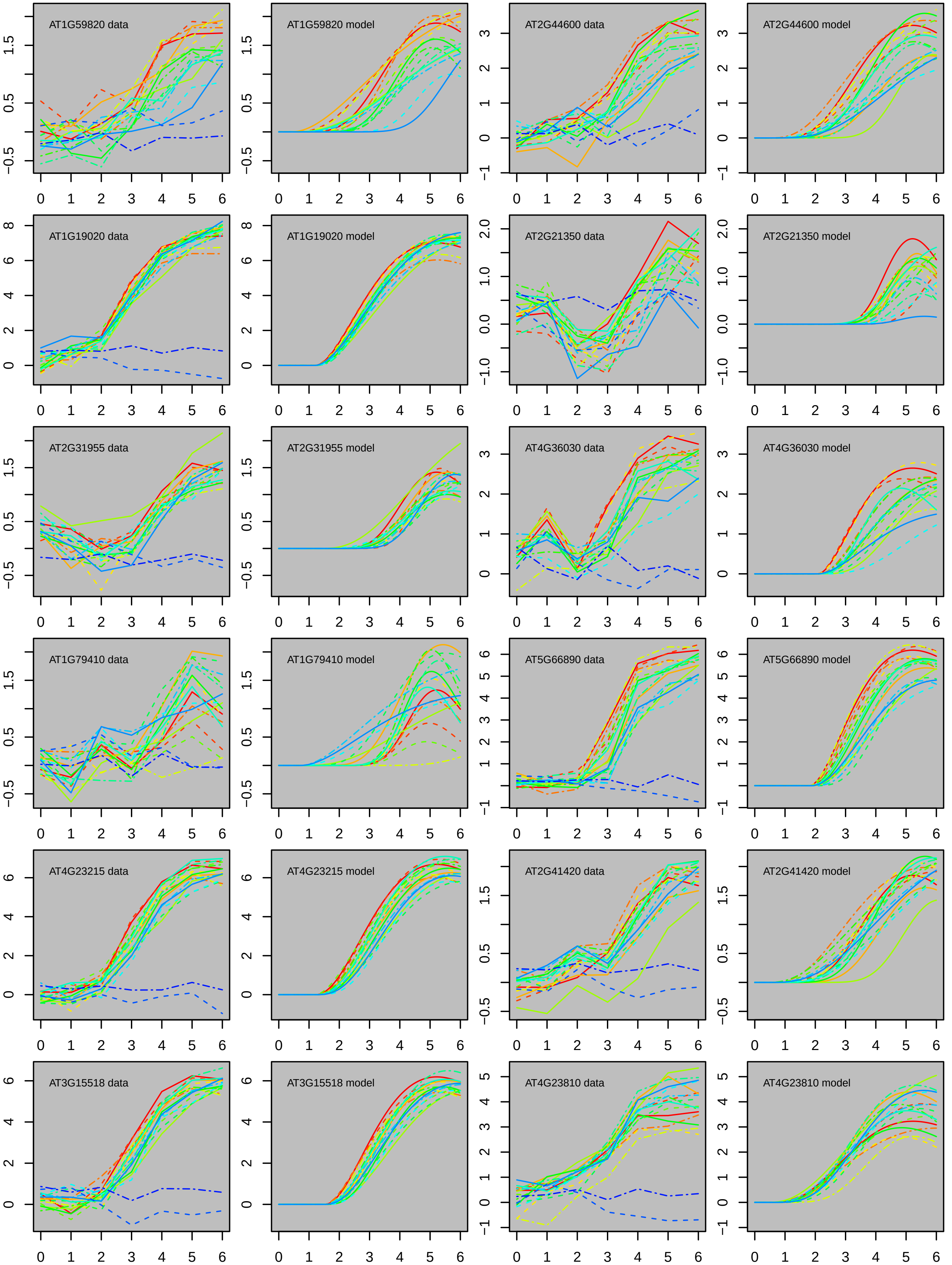
AT1G59820 data
AT1G59820 model
AT2G44600 data
AT2G44600 model
AT1G19020 data
AT1G19020 model
AT2G21350 data
AT2G21350 model
AT2G31955 data
AT2G31955 model
AT4G36030 data
AT4G36030 model
AT1G79410 data
AT1G79410 model
AT5G66890 data
AT5G66890 model
AT4G23215 data
AT4G23215 model
AT2G41420 data
AT2G41420 model
AT3G15518 data
AT3G15518 model
AT4G23810 data
AT4G23810 model

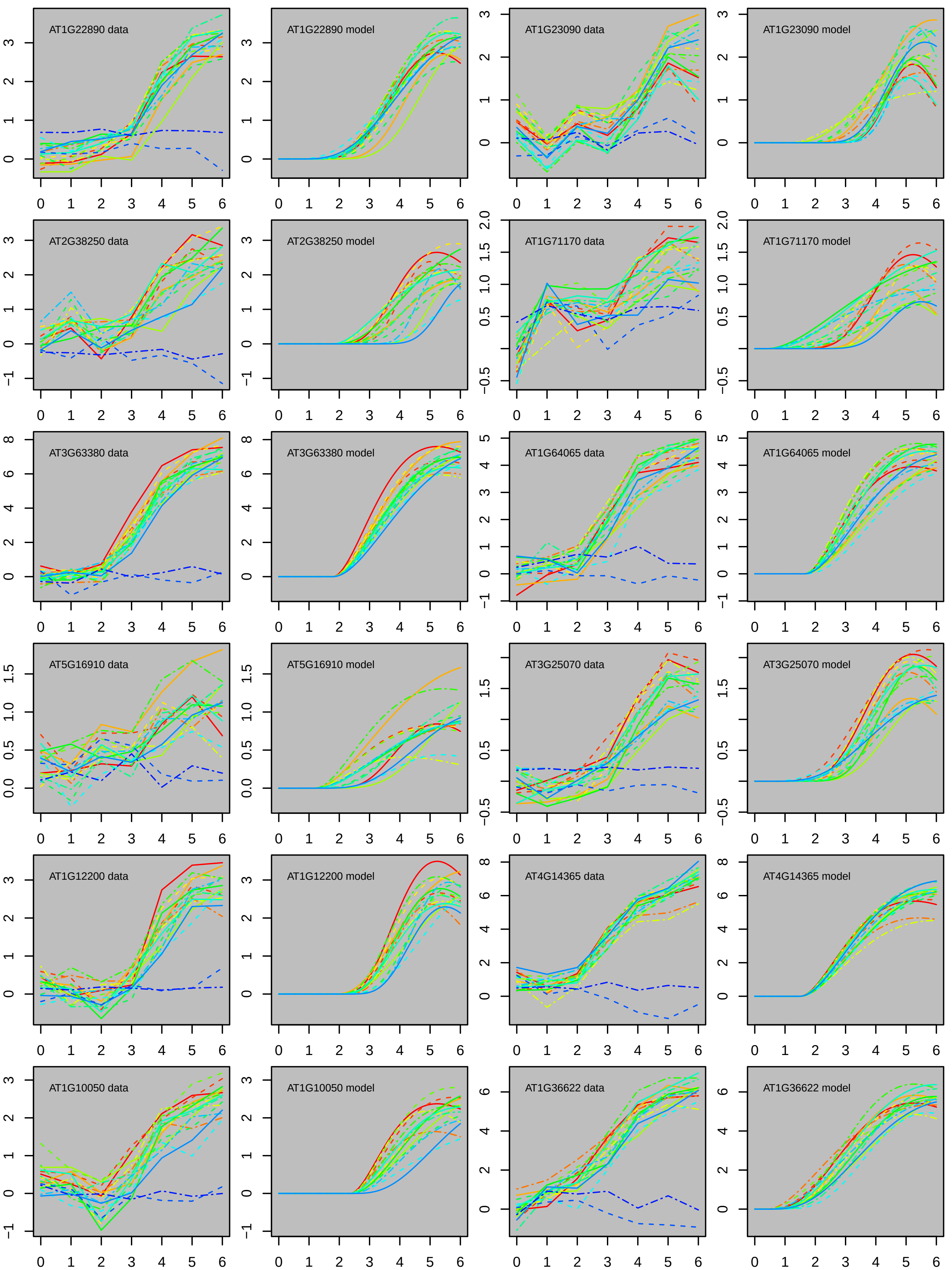

AT1G22890 data
AT1G22890 model
AT1G23090 data
AT1G23090 model
AT2G38250 data
AT2G38250 model
AT1G71170 data
AT1G71170 model
AT3G63380 data
AT3G63380 model
AT1G64065 data
AT1G64065 model
AT5G16910 data
AT5G16910 model
AT3G25070 data
AT3G25070 model
AT1G12200 data
AT1G12200 model
AT4G14365 data
AT4G14365 model
AT1G10050 data
AT1G10050 model
AT1G36622 data
AT1G36622 model

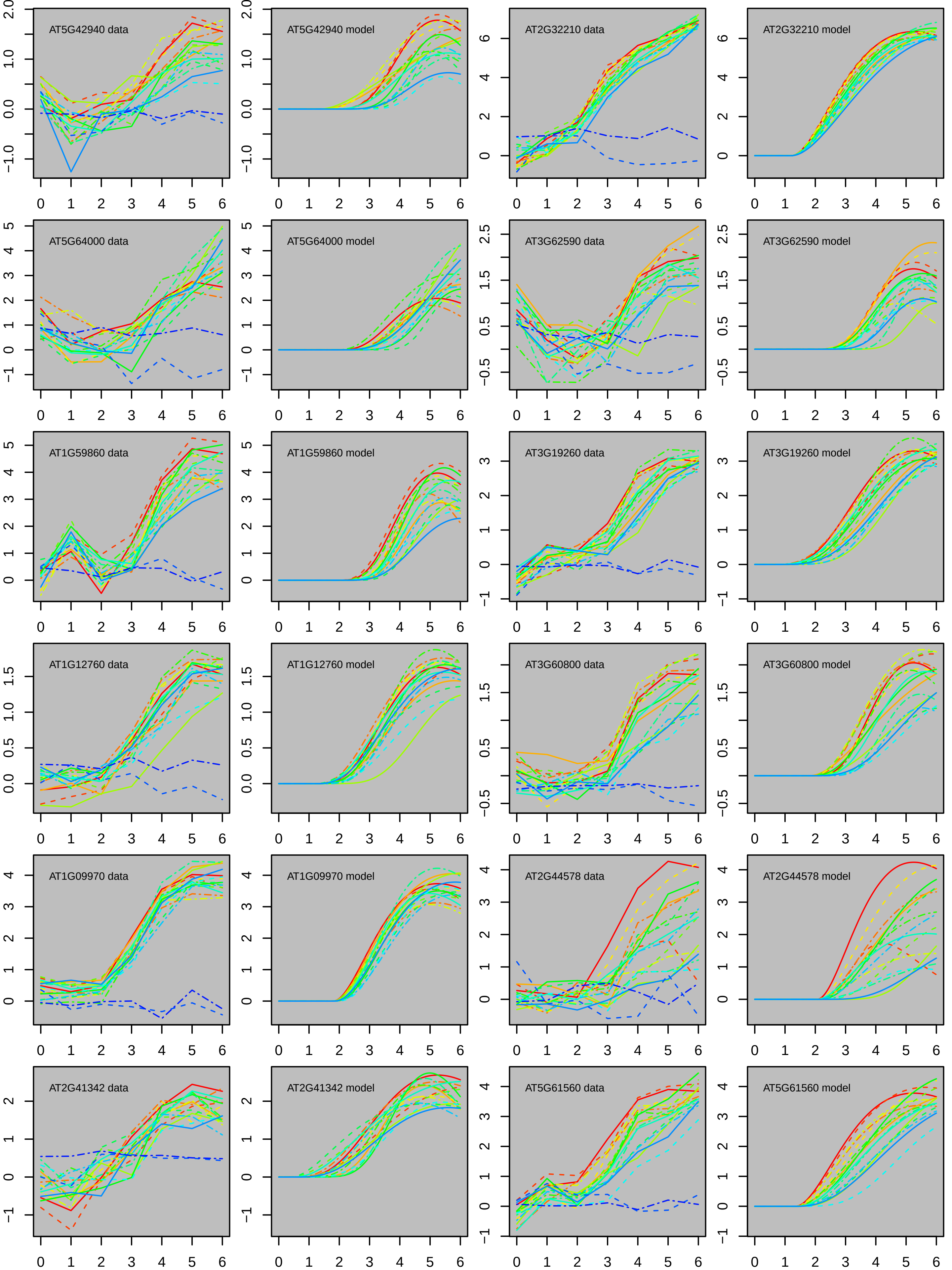
AT5G42940 data
AT5G42940 model
AT2G32210 data
AT2G32210 model
AT5G64000 data
AT5G64000 model
AT3G62590 data
AT3G62590 model
AT1G59860 data
AT1G59860 model
AT3G19260 data
AT3G19260 model
AT1G12760 data
AT1G12760 model
AT3G60800 data
AT3G60800 model
AT1G09970 data
AT1G09970 model
AT2G44578 data
AT2G44578 model
AT2G41342 data
AT2G41342 model
AT5G61560 data
AT5G61560 model

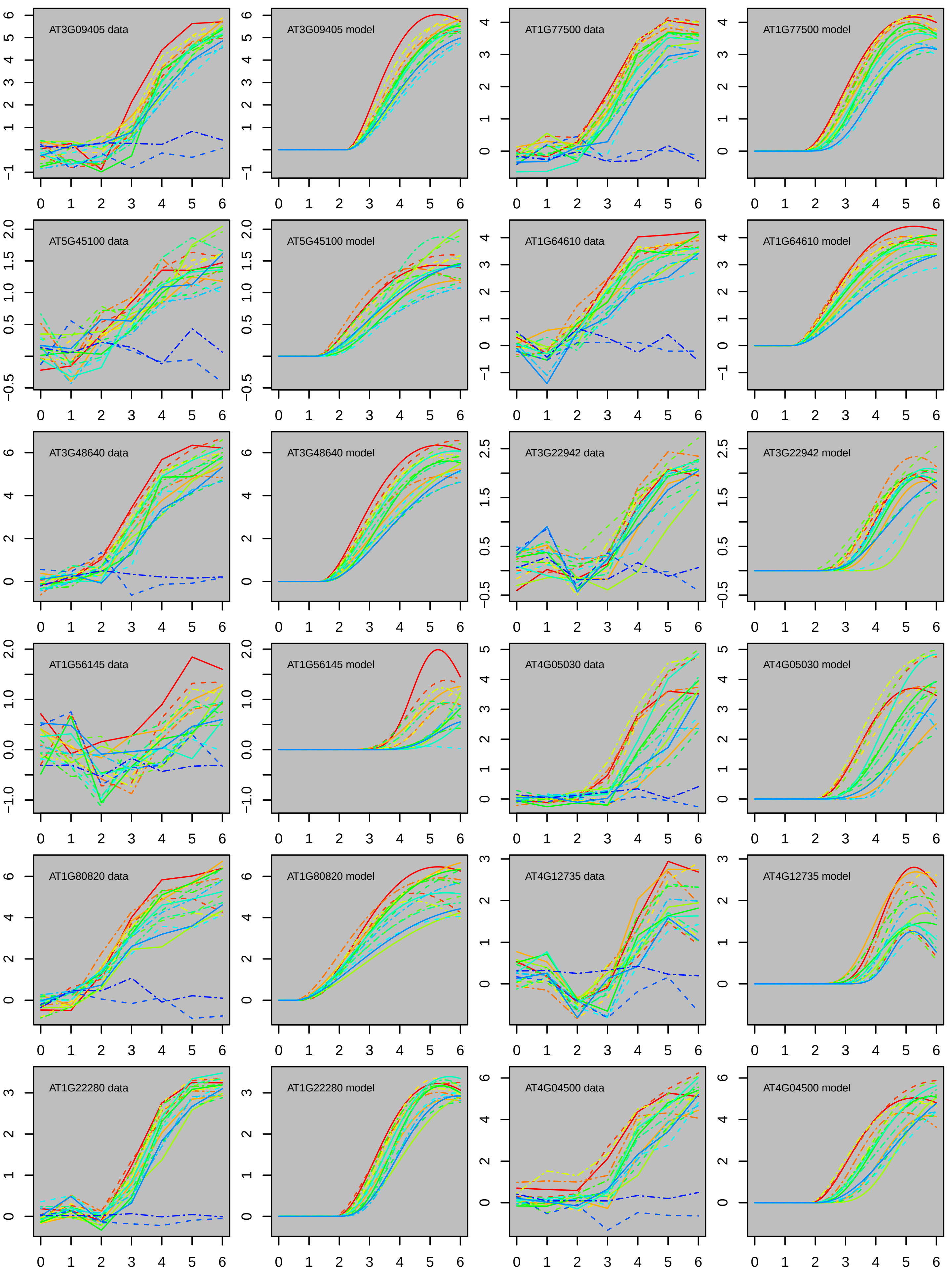
AT3G09405 data
AT3G09405 model
AT1G77500 data
AT1G77500 model
AT5G45100 data
AT5G45100 model
AT1G64610 data
AT1G64610 model
AT3G48640 data
AT3G48640 model
AT3G22942 data
AT3G22942 model
AT1G56145 data
AT1G56145 model
AT4G05030 data
AT4G05030 model
AT1G80820 data
AT1G80820 model
AT4G12735 data
AT4G12735 model
AT1G22280 data
AT1G22280 model
AT4G04500 data
AT4G04500 model

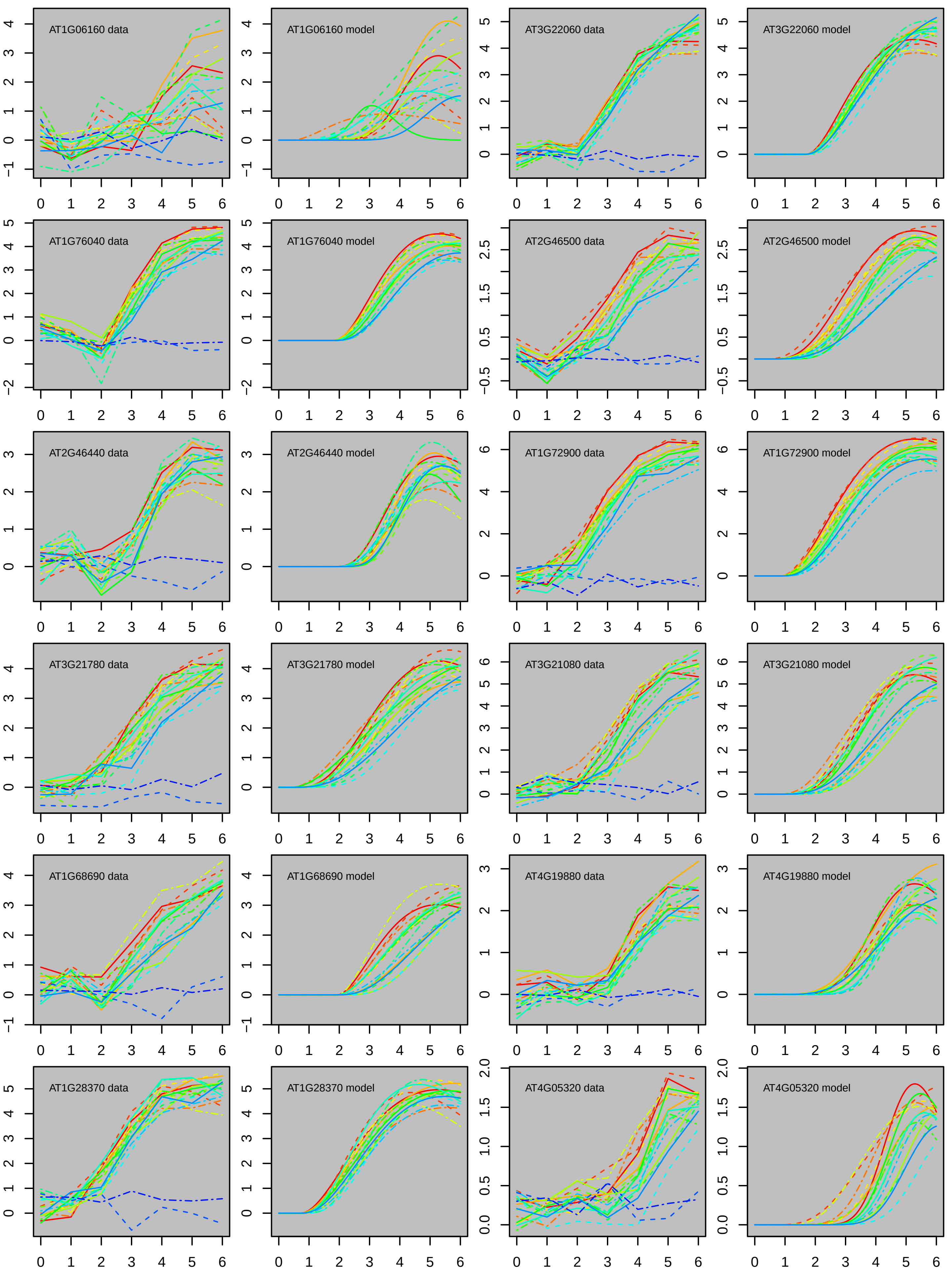
AT1G06160 data
AT1G06160 model
AT3G22060 data
AT3G22060 model
AT1G76040 data
AT1G76040 model
AT2G46500 data
AT2G46500 model
AT2G46440 data
AT2G46440 model
AT1G72900 data
AT1G72900 model
AT3G21780 data
AT3G21780 model
AT3G21080 data
AT3G21080 model
AT1G68690 data
AT1G68690 model
AT4G19880 data
AT4G19880 model
AT1G28370 data
AT1G28370 model
AT4G05320 data
AT4G05320 model

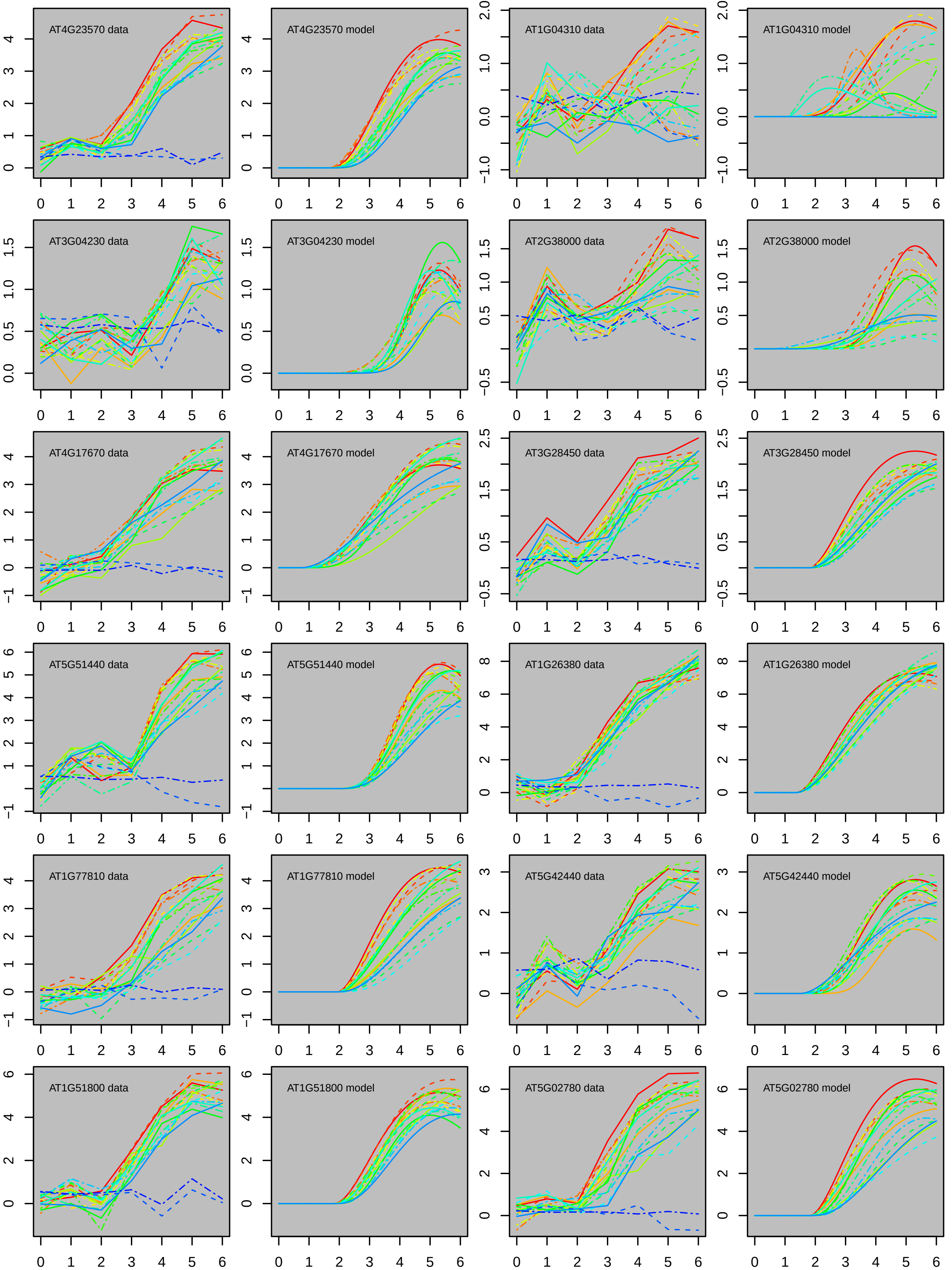

AT4G23570 data
AT4G23570 model
AT1G04310 data
AT1G04310 model
AT3G04230 data
AT3G04230 model
AT2G38000 data
AT2G38000 model
AT4G17670 data
AT4G17670 model
AT3G28450 data
AT3G28450 model
AT5G51440 data
AT5G51440 model
AT1G26380 data
AT1G26380 model
AT1G77810 data
AT1G77810 model
AT5G42440 data
AT5G42440 model
AT1G51800 data
AT1G51800 model
AT5G02780 data
AT5G02780 model

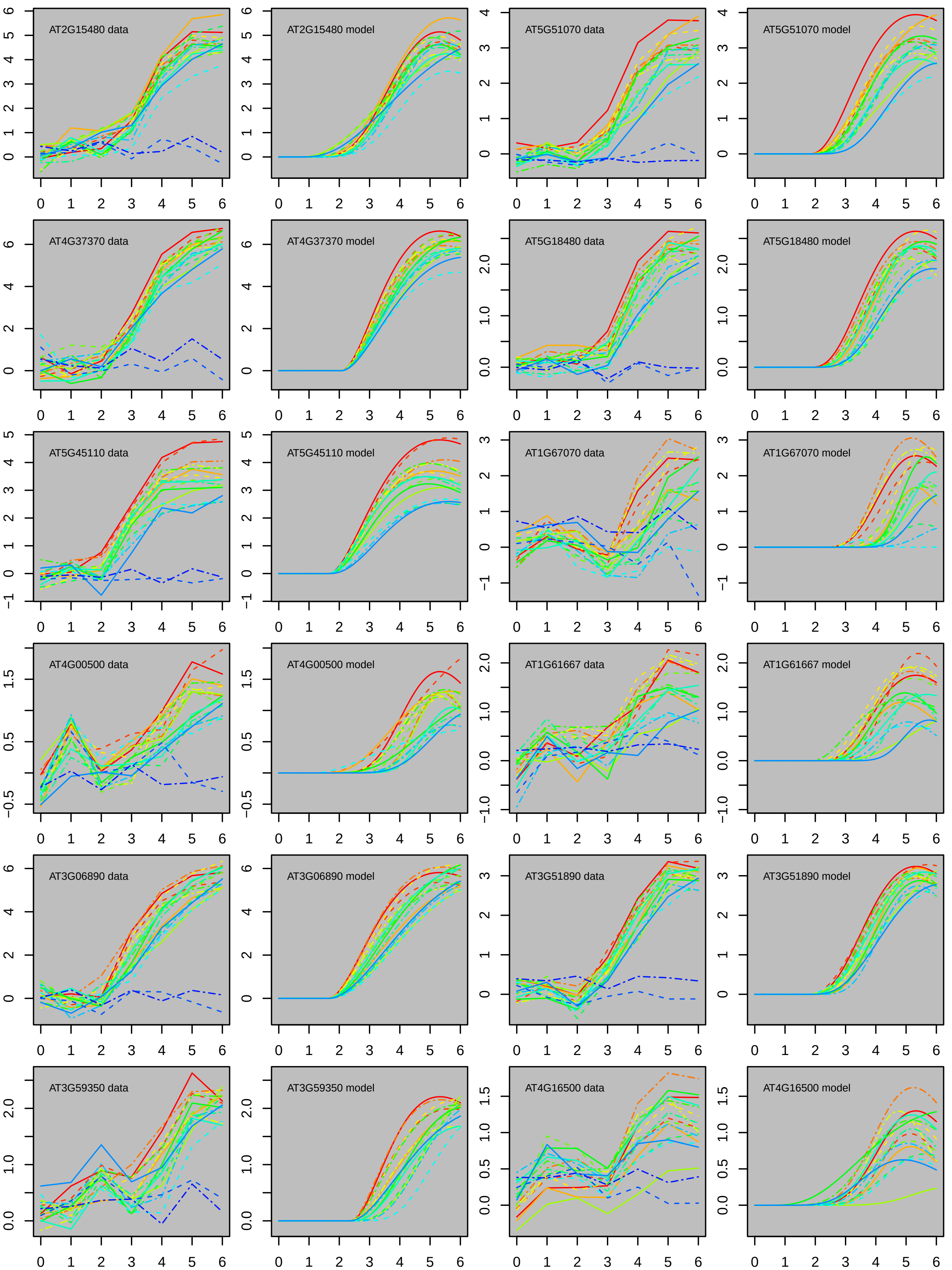
AT2G15480 data
AT2G15480 model
AT5G51070 data
AT5G51070 model
AT4G37370 data
AT4G37370 model
AT5G18480 data
AT5G18480 model
AT5G45110 data
AT5G45110 model
AT1G67070 data
AT1G67070 model
AT4G00500 data
AT4G00500 model
AT1G61667 data
AT1G61667 model
AT3G06890 data
AT3G06890 model
AT3G51890 data
AT3G51890 model
AT3G59350 data
AT3G59350 model
AT4G16500 data
AT4G16500 model

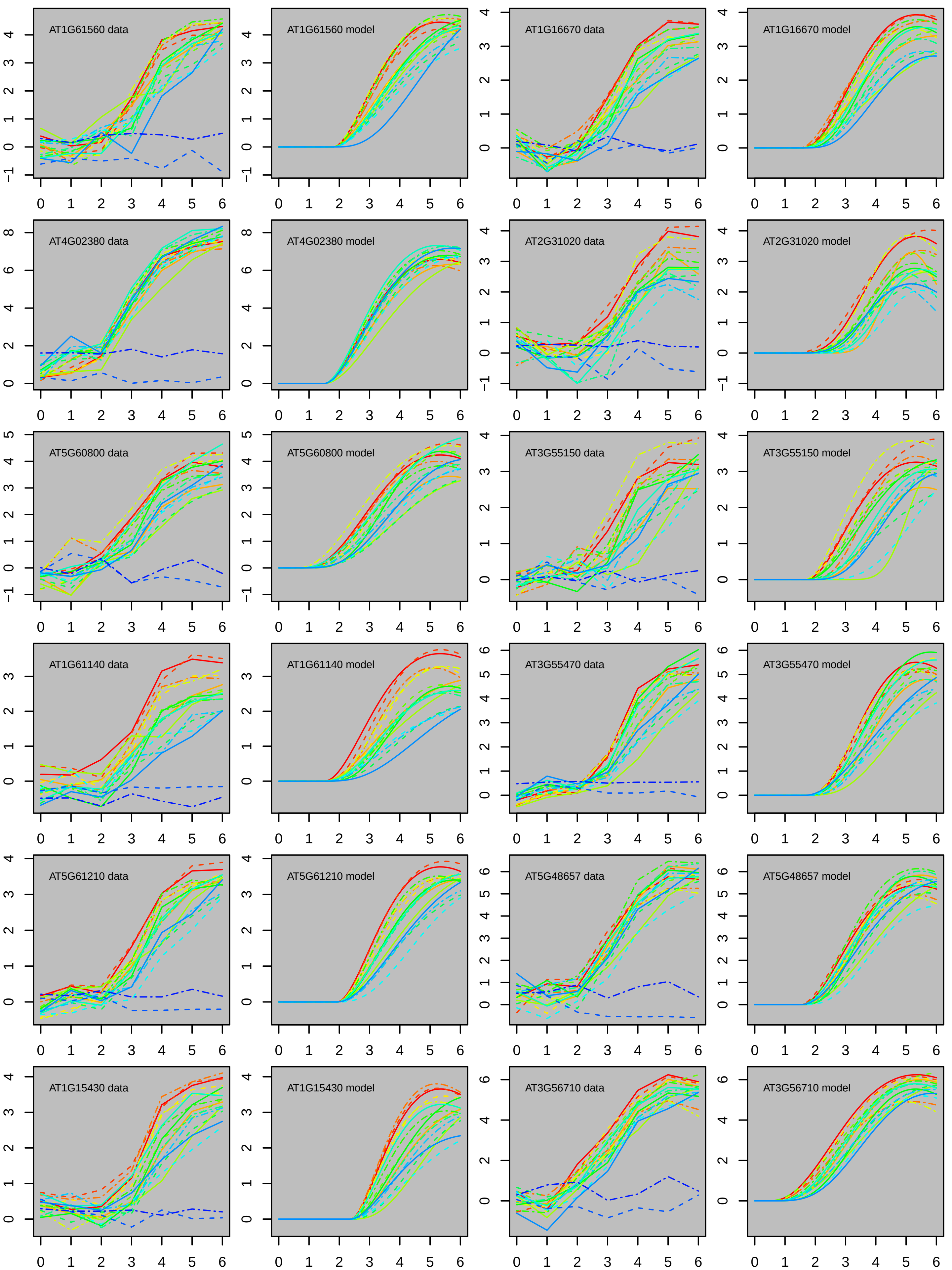
AT1G61560 data
AT1G61560 model
AT1G16670 data
AT1G16670 model
AT4G02380 data
AT4G02380 model
AT2G31020 data
AT2G31020 model
AT5G60800 data
AT5G60800 model
AT3G55150 data
AT3G55150 model
AT1G61140 data
AT1G61140 model
AT3G55470 data
AT3G55470 model
AT5G61210 data
AT5G61210 model
AT5G48657 data
AT5G48657 model
AT1G15430 data
AT1G15430 model
AT3G56710 data
AT3G56710 model

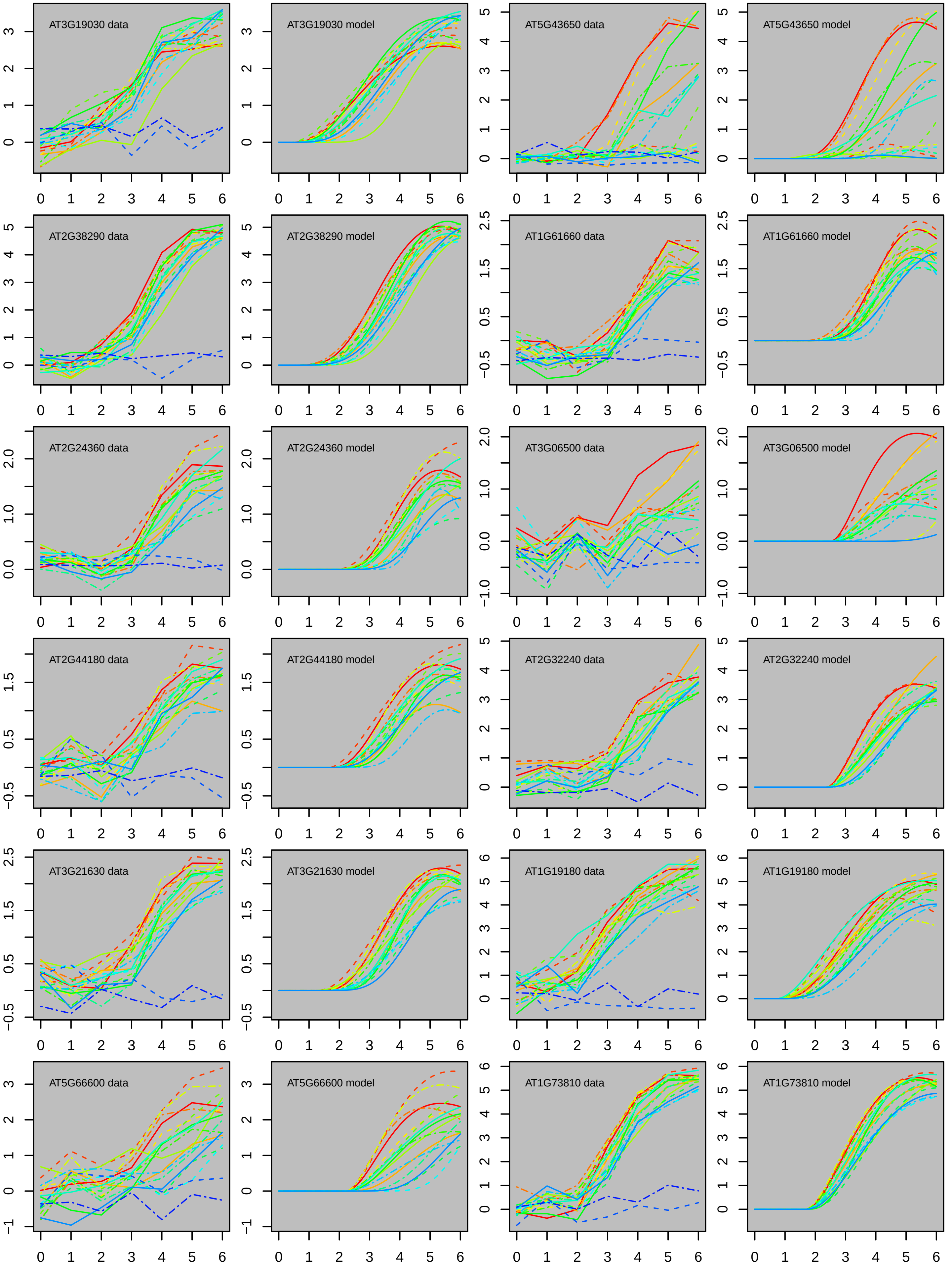
AT3G19030 data
AT3G19030 model
AT5G43650 data
AT5G43650 model
AT2G38290 data
AT2G38290 model
AT1G61660 data
AT1G61660 model
AT2G24360 data
AT2G24360 model
AT3G06500 data
AT3G06500 model
AT2G44180 data
AT2G44180 model
AT2G32240 data
AT2G32240 model
AT3G21630 data
AT3G21630 model
AT1G19180 data
AT1G19180 model
AT5G66600 data
AT5G66600 model
AT1G73810 data
AT1G73810 model

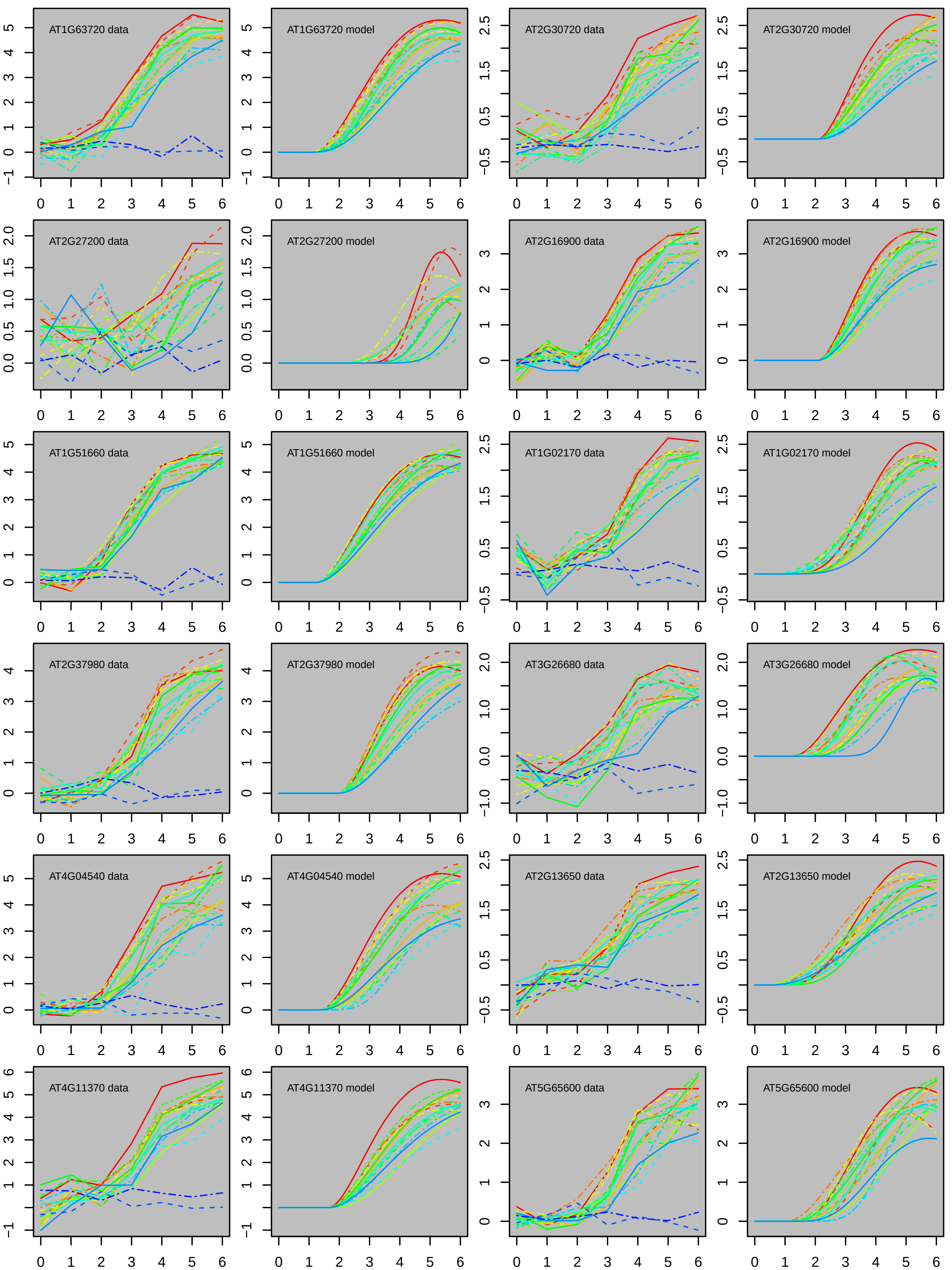

AT1G63720 data
AT1G63720 model
AT2G30720 data
AT2G30720 model
AT2G27200 data
AT2G27200 model
AT2G16900 data
AT2G16900 model
AT1G51660 data
AT1G51660 model
AT1G02170 data
AT1G02170 model
AT2G37980 data
AT2G37980 model
AT3G26680 data
AT3G26680 model
AT4G04540 data
AT4G04540 model
AT2G13650 data
AT2G13650 model
AT4G11370 data
AT4G11370 model
AT5G65600 data
AT5G65600 model

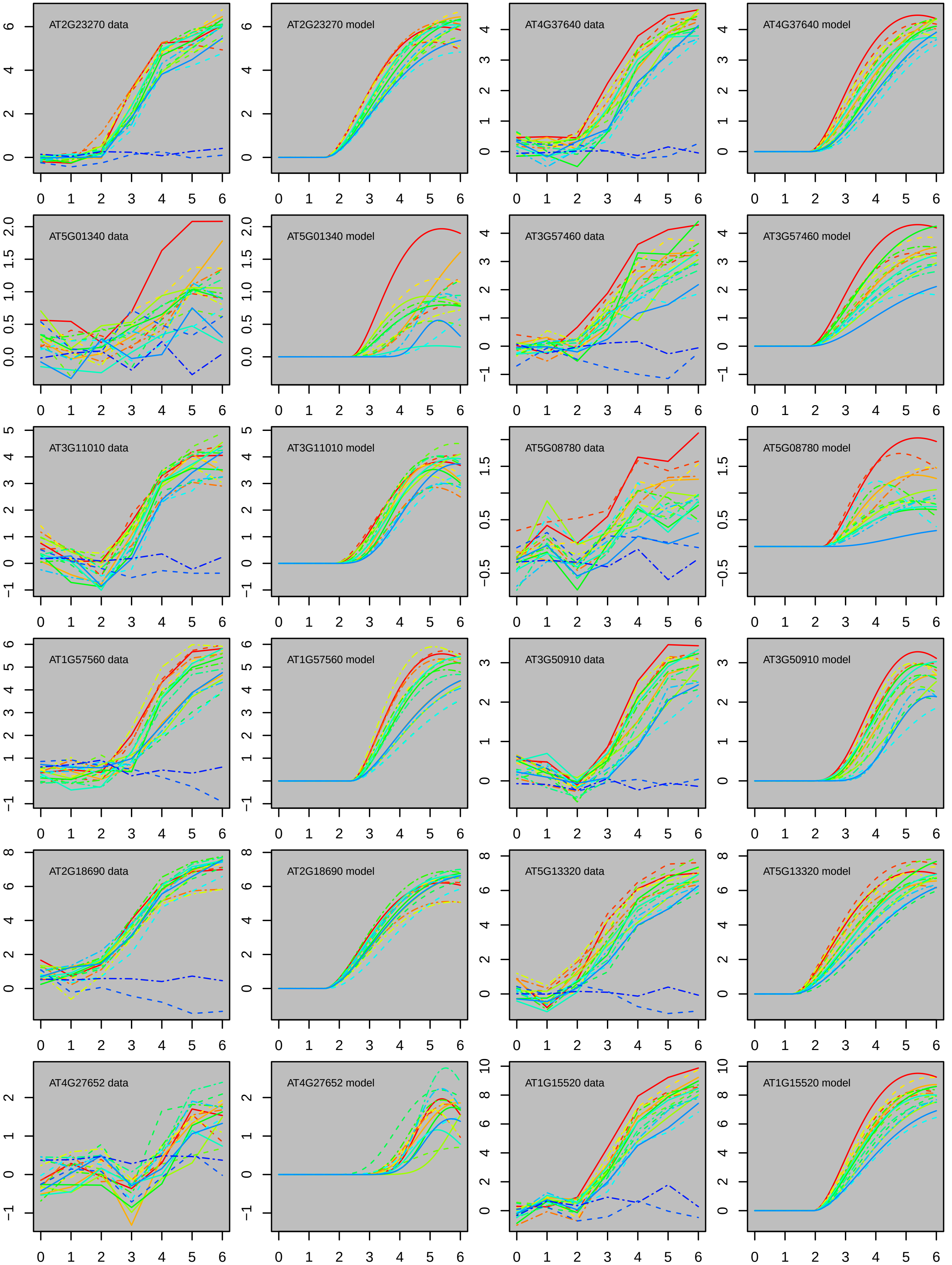
AT2G23270 data
AT2G23270 model
AT4G37640 data
AT4G37640 model
AT5G01340 data
AT5G01340 model
AT3G57460 data
AT3G57460 model
AT3G11010 data
AT3G11010 model
AT5G08780 data
AT5G08780 model
AT1G57560 data
AT1G57560 model
AT3G50910 data
AT3G50910 model
AT2G18690 data
AT2G18690 model
AT5G13320 data
AT5G13320 model
AT4G27652 data
AT4G27652 model
AT1G15520 data
AT1G15520 model

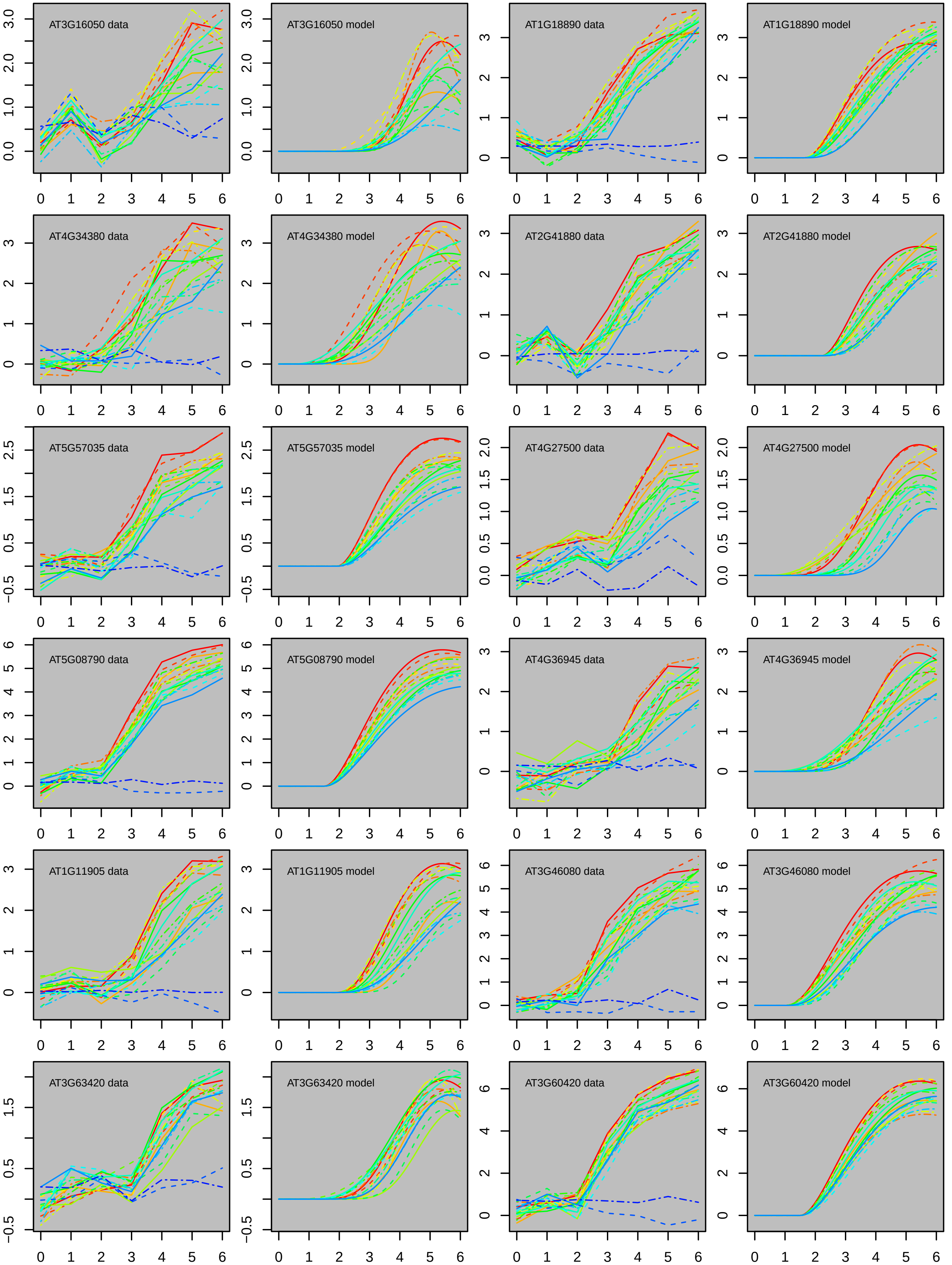
AT3G16050 data
AT3G16050 model
AT1G18890 data
AT1G18890 model
AT4G34380 data
AT4G34380 model
AT2G41880 data
AT2G41880 model
AT5G57035 data
AT5G57035 model
AT4G27500 data
AT4G27500 model
AT5G08790 data
AT5G08790 model
AT4G36945 data
AT4G36945 model
AT1G11905 data
AT1G11905 model
AT3G46080 data
AT3G46080 model
AT3G63420 data
AT3G63420 model
AT3G60420 data
AT3G60420 model

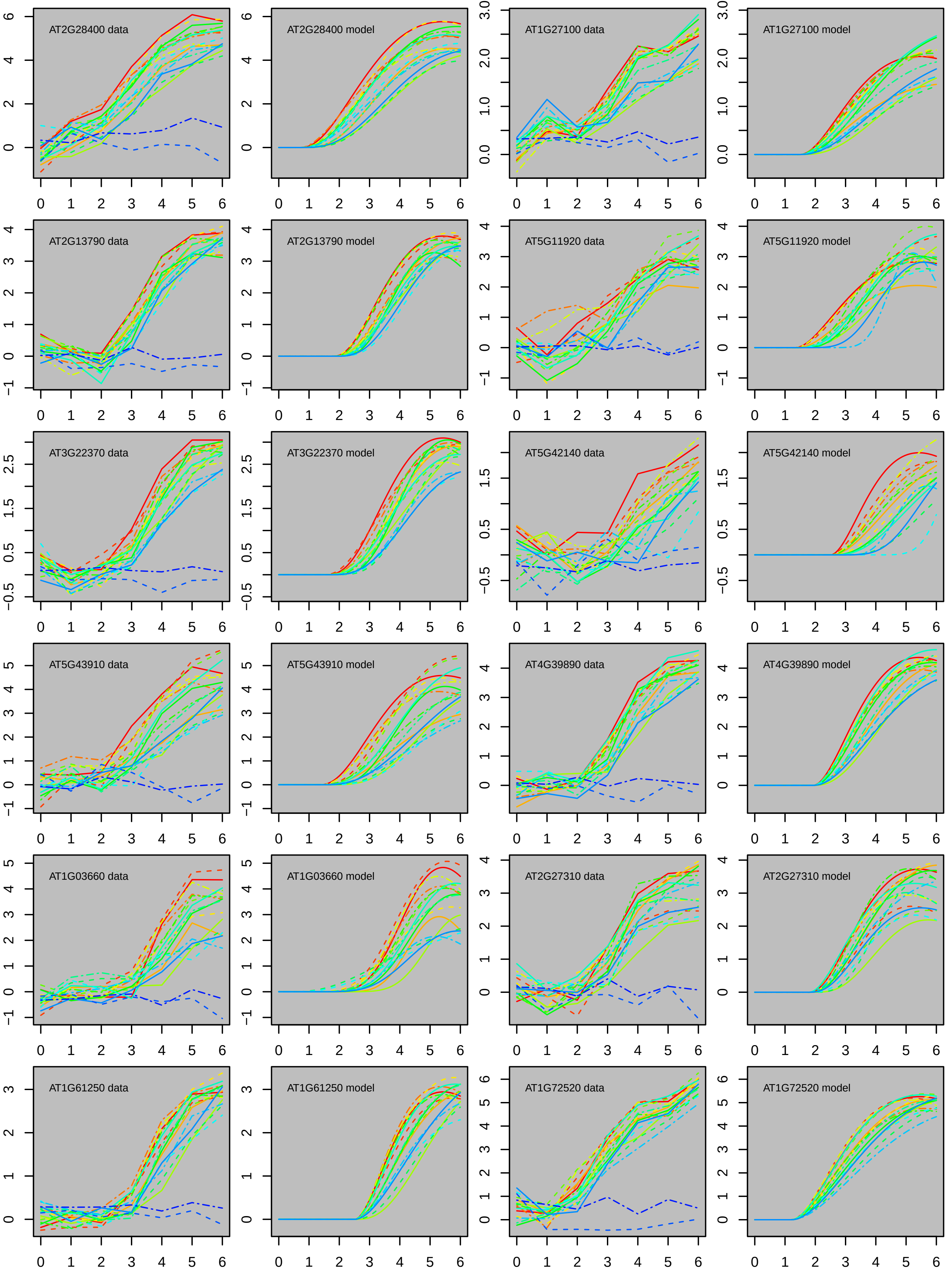
AT2G28400 data
AT2G28400 model
AT1G27100 data
AT1G27100 model
AT2G13790 data
AT2G13790 model
AT5G11920 data
AT5G11920 model
AT3G22370 data
AT3G22370 model
AT5G42140 data
AT5G42140 model
AT5G43910 data
AT5G43910 model
AT4G39890 data
AT4G39890 model
AT1G03660 data
AT1G03660 model
AT2G27310 data
AT2G27310 model
AT1G61250 data
AT1G61250 model
AT1G72520 data
AT1G72520 model

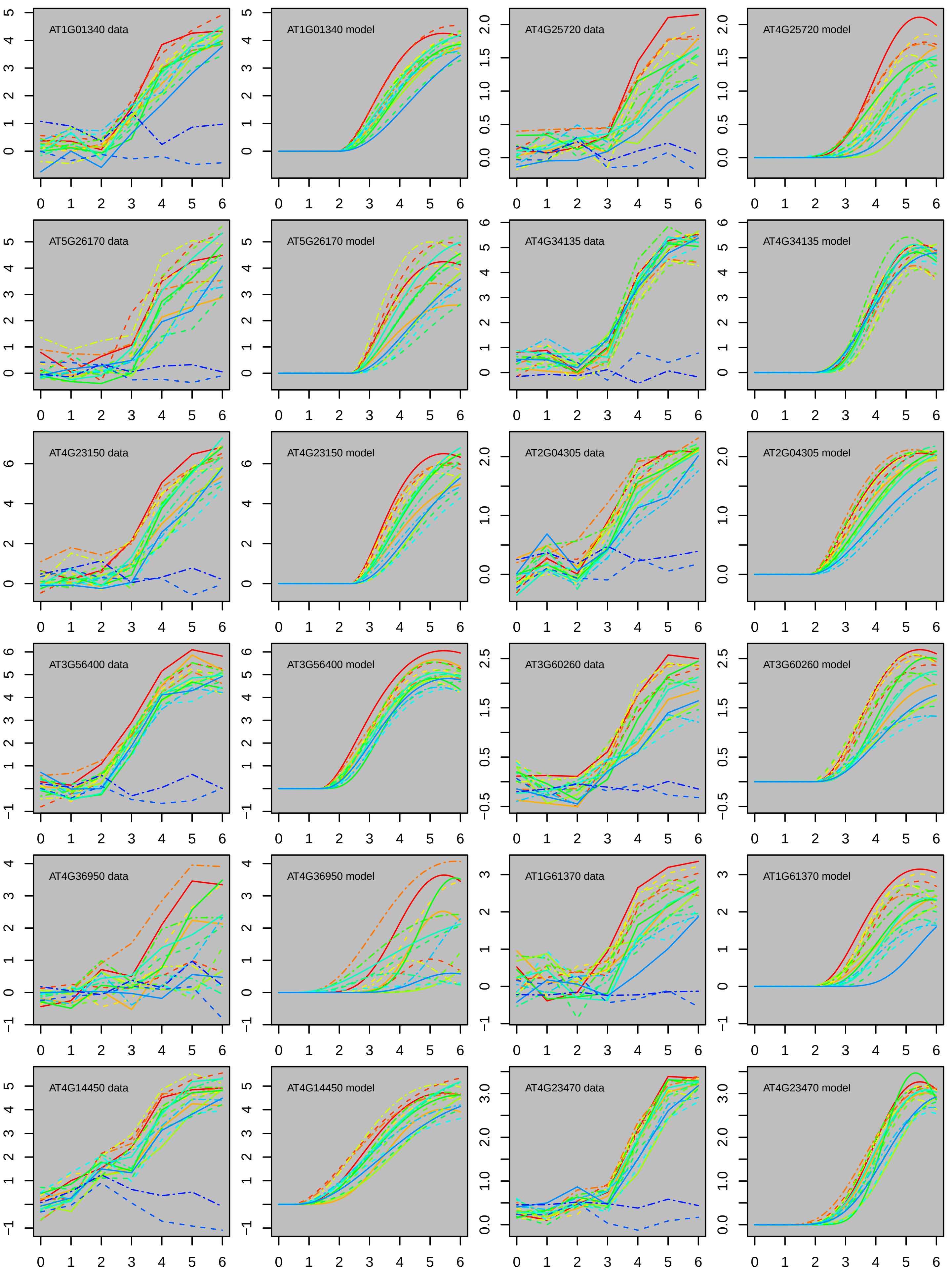
AT1G01340 data
AT1G01340 model
AT4G25720 data
AT4G25720 model
AT5G26170 data
AT5G26170 model
AT4G34135 data
AT4G34135 model
AT4G23150 data
AT4G23150 model
AT2G04305 data
AT2G04305 model
AT3G56400 data
AT3G56400 model
AT3G60260 data
AT3G60260 model
AT4G36950 data
AT4G36950 model
AT1G61370 data
AT1G61370 model
AT4G14450 data
AT4G14450 model
AT4G23470 data
AT4G23470 model

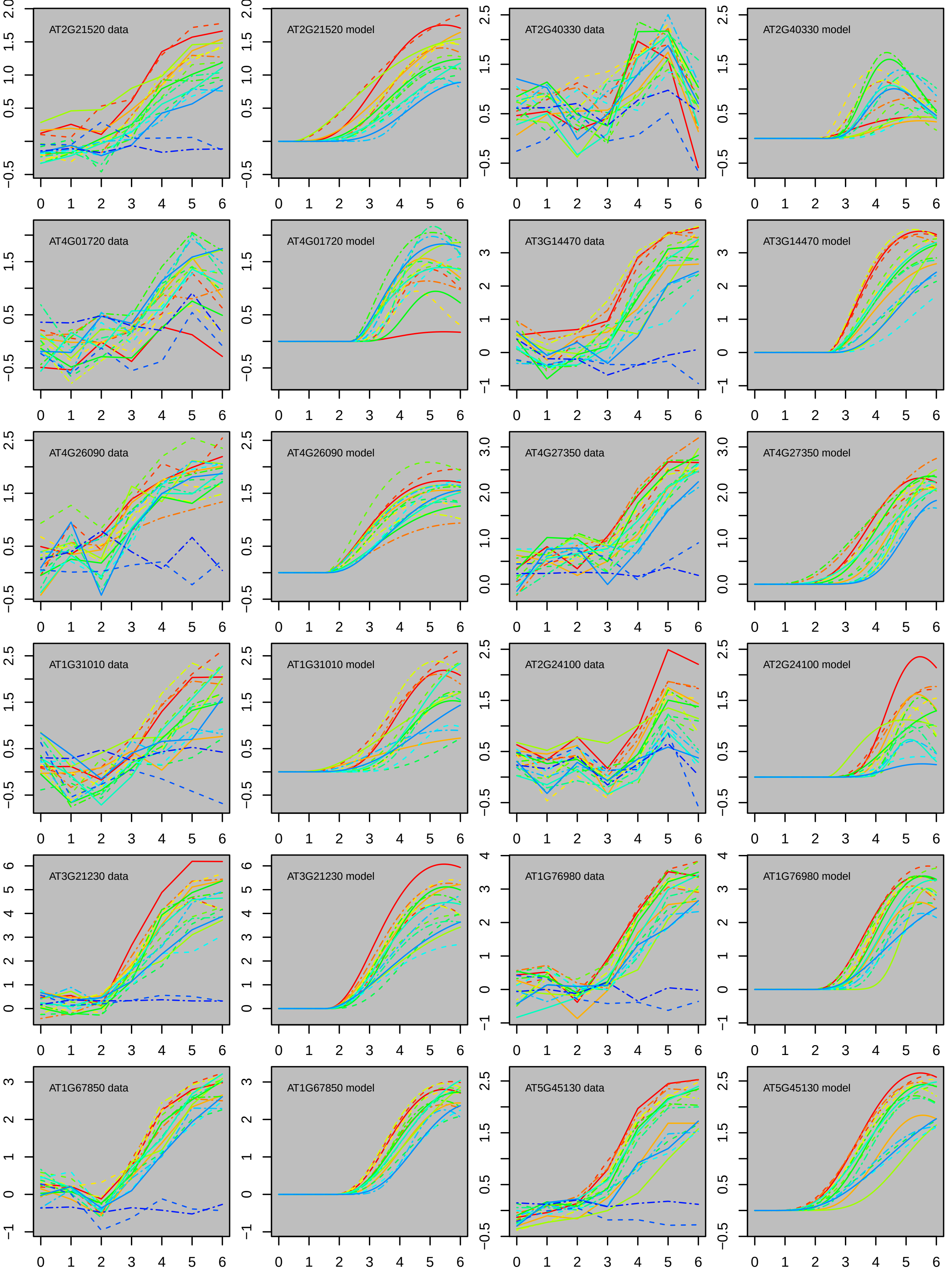
AT2G21520 data
AT2G21520 model
AT2G40330 data
AT2G40330 model
AT4G01720 data
AT4G01720 model
AT3G14470 data
AT3G14470 model
AT4G26090 data
AT4G26090 model
AT4G27350 data
AT4G27350 model
AT1G31010 data
AT1G31010 model
AT2G24100 data
AT2G24100 model
AT3G21230 data
AT3G21230 model
AT1G76980 data
AT1G76980 model
AT1G67850 data
AT1G67850 model
AT5G45130 data
AT5G45130 model

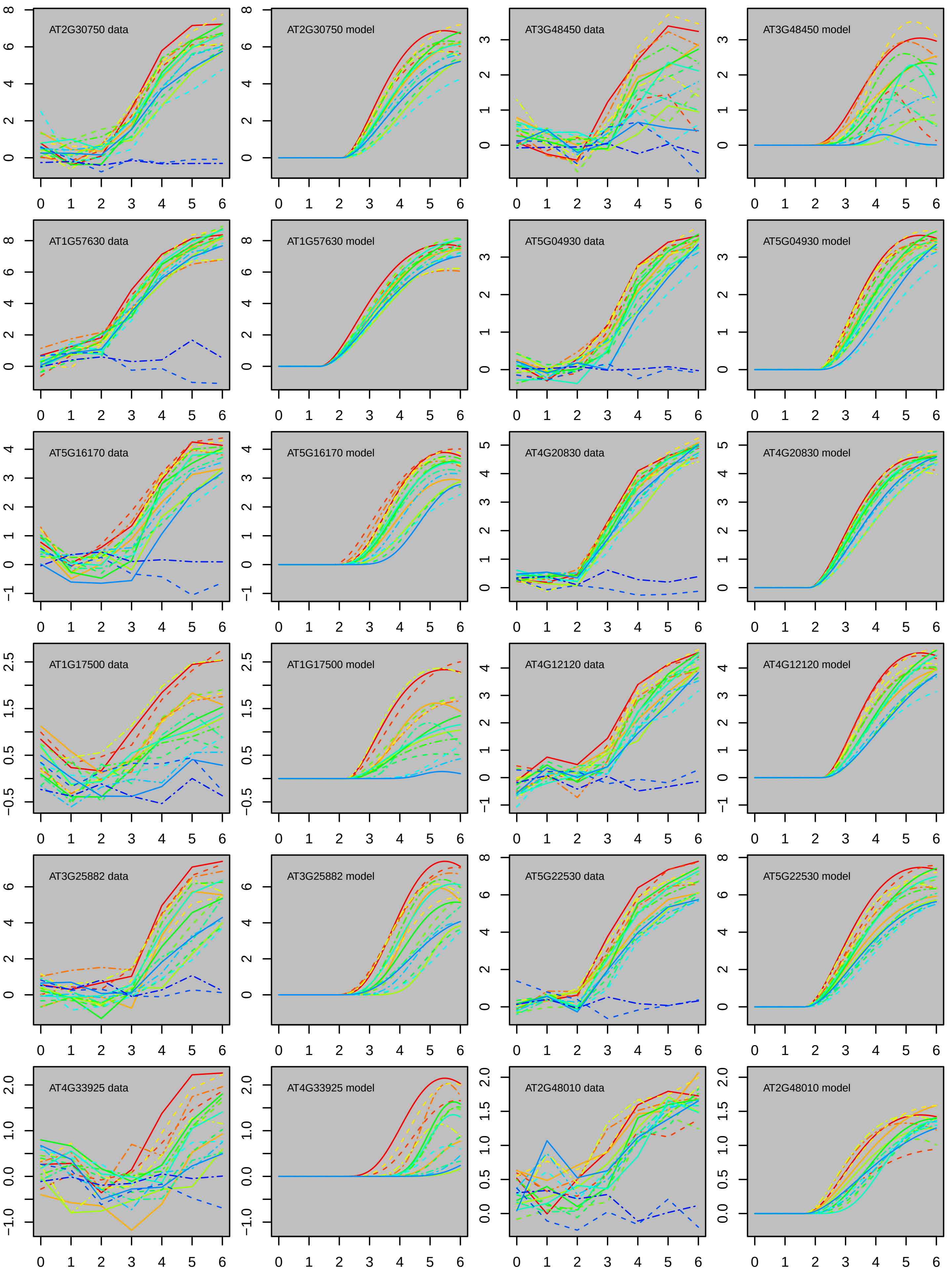

AT2G30750 data
AT2G30750 model
AT3G48450 data
AT3G48450 model
AT1G57630 data
AT1G57630 model
AT5G04930 data
AT5G04930 model
AT5G16170 data
AT5G16170 model
AT4G20830 data
AT4G20830 model
AT1G17500 data
AT1G17500 model
AT4G12120 data
AT4G12120 model
AT3G25882 data
AT3G25882 model
AT5G22530 data
AT5G22530 model
AT4G33925 data
AT4G33925 model
AT2G48010 data
AT2G48010 model

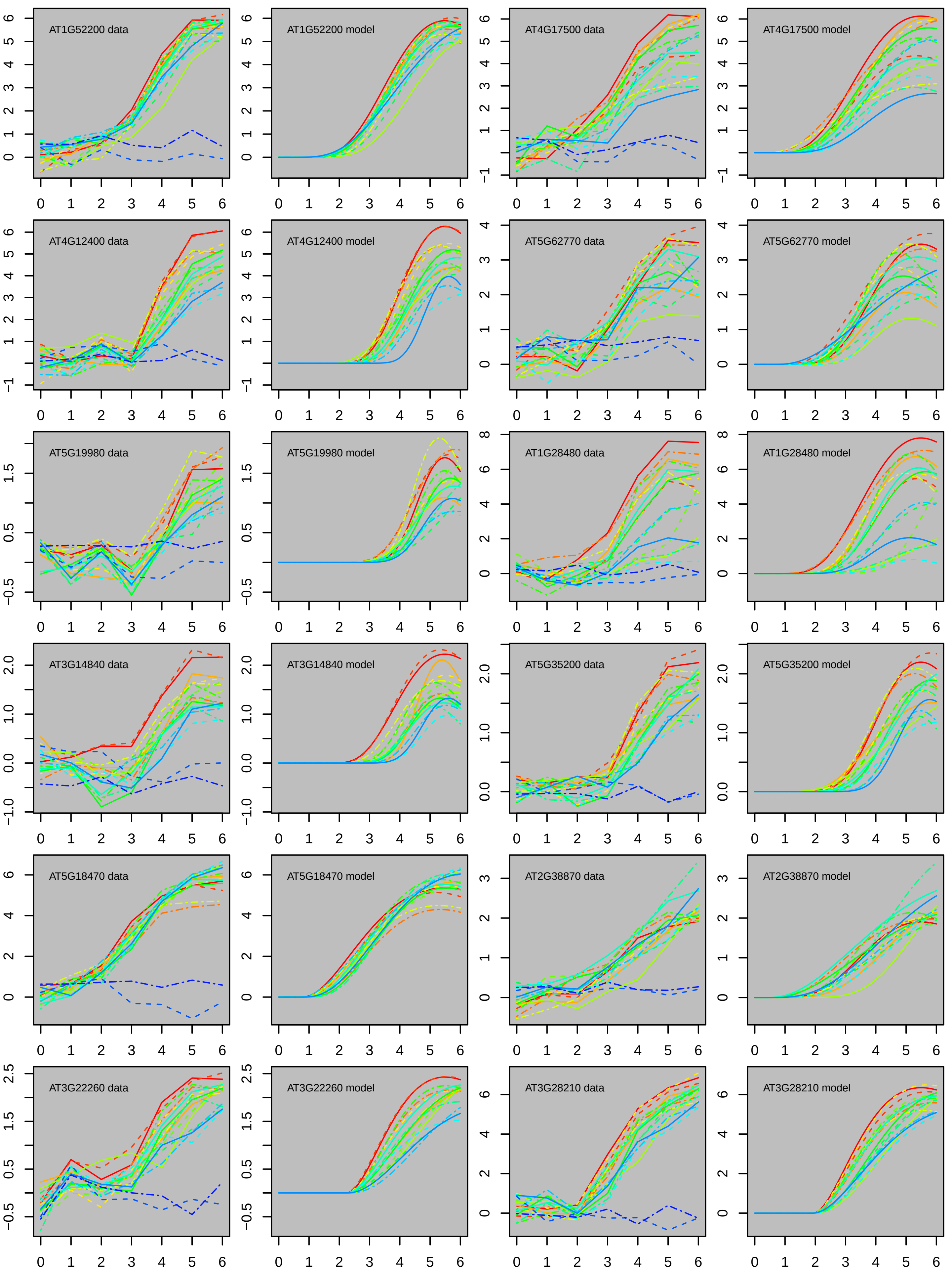

AT1G52200 data
AT1G52200 model
AT4G17500 data
AT4G17500 model
AT4G12400 data
AT4G12400 model
AT5G62770 data
AT5G62770 model
AT5G19980 data
AT5G19980 model
AT1G28480 data
AT1G28480 model
AT3G14840 data
AT3G14840 model
AT5G35200 data
AT5G35200 model
AT5G18470 data
AT5G18470 model
AT2G38870 data
AT2G38870 model
AT3G22260 data
AT3G22260 model
AT3G28210 data
AT3G28210 model

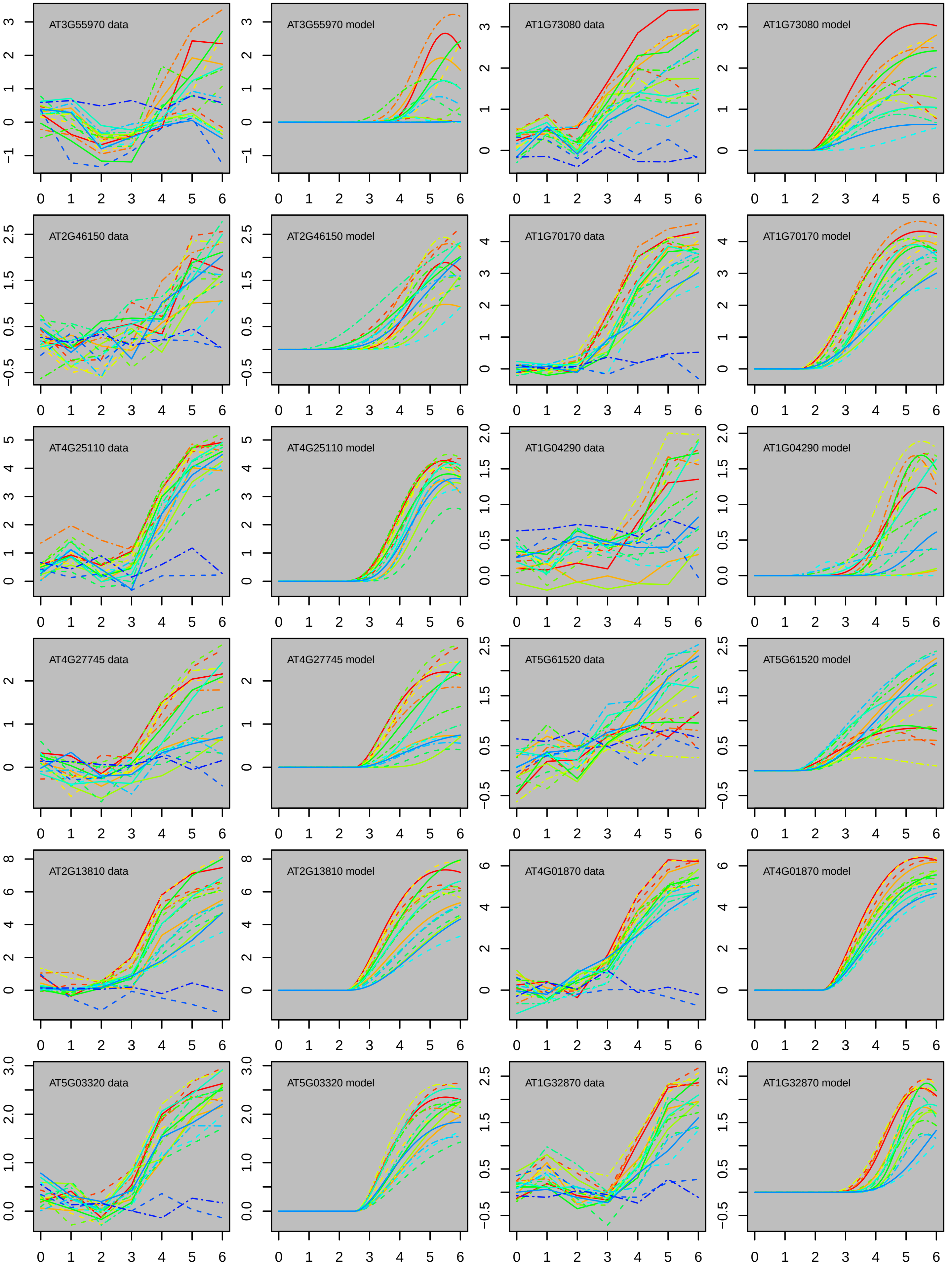
AT3G55970 data
AT3G55970 model
AT1G73080 data
AT1G73080 model
AT2G46150 data
AT2G46150 model
AT1G70170 data
AT1G70170 model
AT4G25110 data
AT4G25110 model
AT1G04290 data
AT1G04290 model
AT4G27745 data
AT4G27745 model
AT5G61520 data
AT5G61520 model
AT2G13810 data
AT2G13810 model
AT4G01870 data
AT4G01870 model
AT5G03320 data
AT5G03320 model
AT1G32870 data
AT1G32870 model

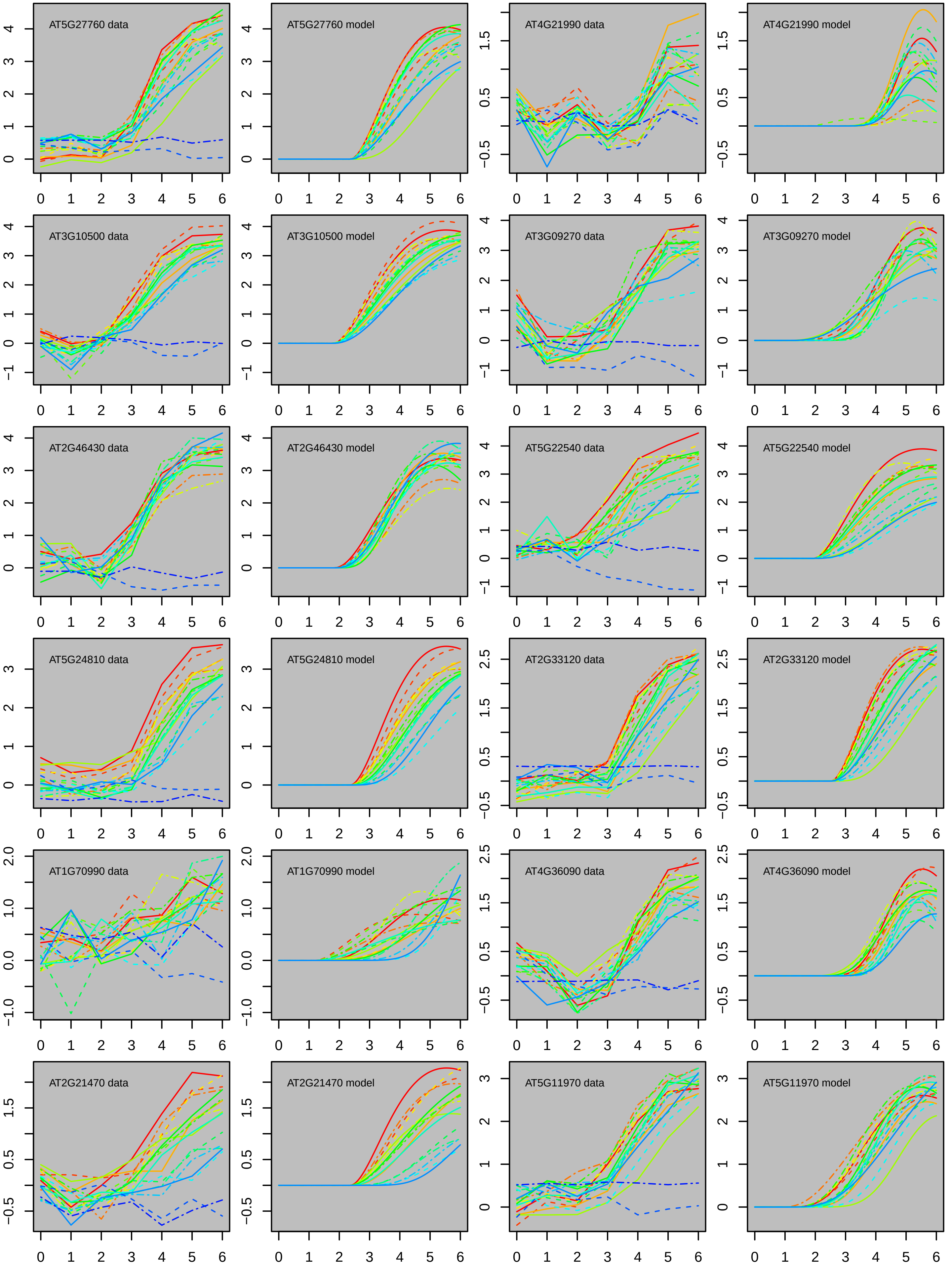
AT5G27760 data
AT5G27760 model
AT4G21990 data
AT4G21990 model
AT3G10500 data
AT3G10500 model
AT3G09270 data
AT3G09270 model
AT2G46430 data
AT2G46430 model
AT5G22540 data
AT5G22540 model
AT5G24810 data
AT5G24810 model
AT2G33120 data
AT2G33120 model
AT1G70990 data
AT1G70990 model
AT4G36090 data
AT4G36090 model
AT2G21470 data
AT2G21470 model
AT5G11970 data
AT5G11970 model

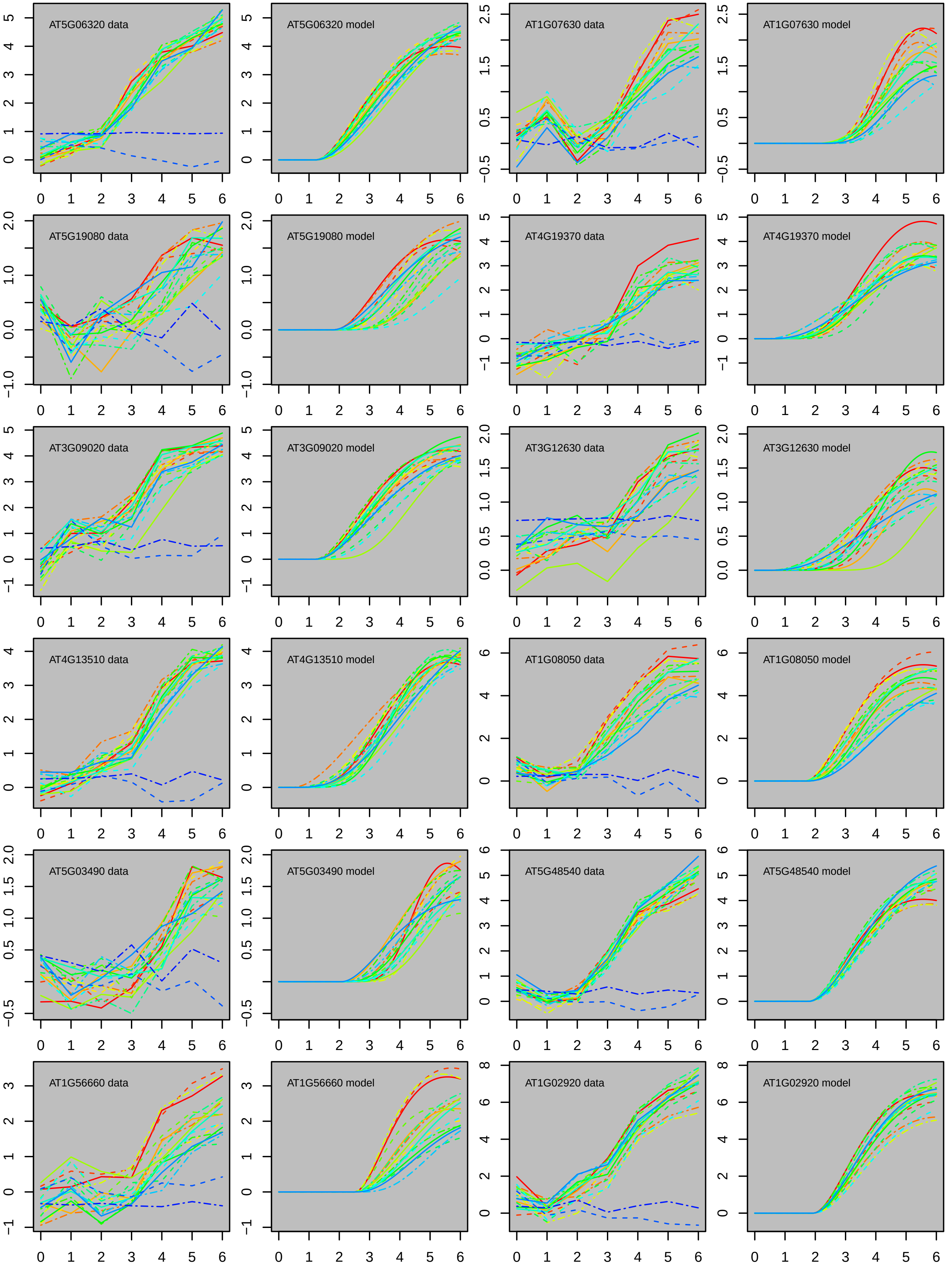

AT5G06320 data
AT5G06320 model
AT1G07630 data
AT1G07630 model
AT5G19080 data
AT5G19080 model
AT4G19370 data
AT4G19370 model
AT3G09020 data
AT3G09020 model
AT3G12630 data
AT3G12630 model
AT4G13510 data
AT4G13510 model
AT1G08050 data
AT1G08050 model
AT5G03490 data
AT5G03490 model
AT5G48540 data
AT5G48540 model
AT1G56660 data
AT1G56660 model
AT1G02920 data
AT1G02920 model

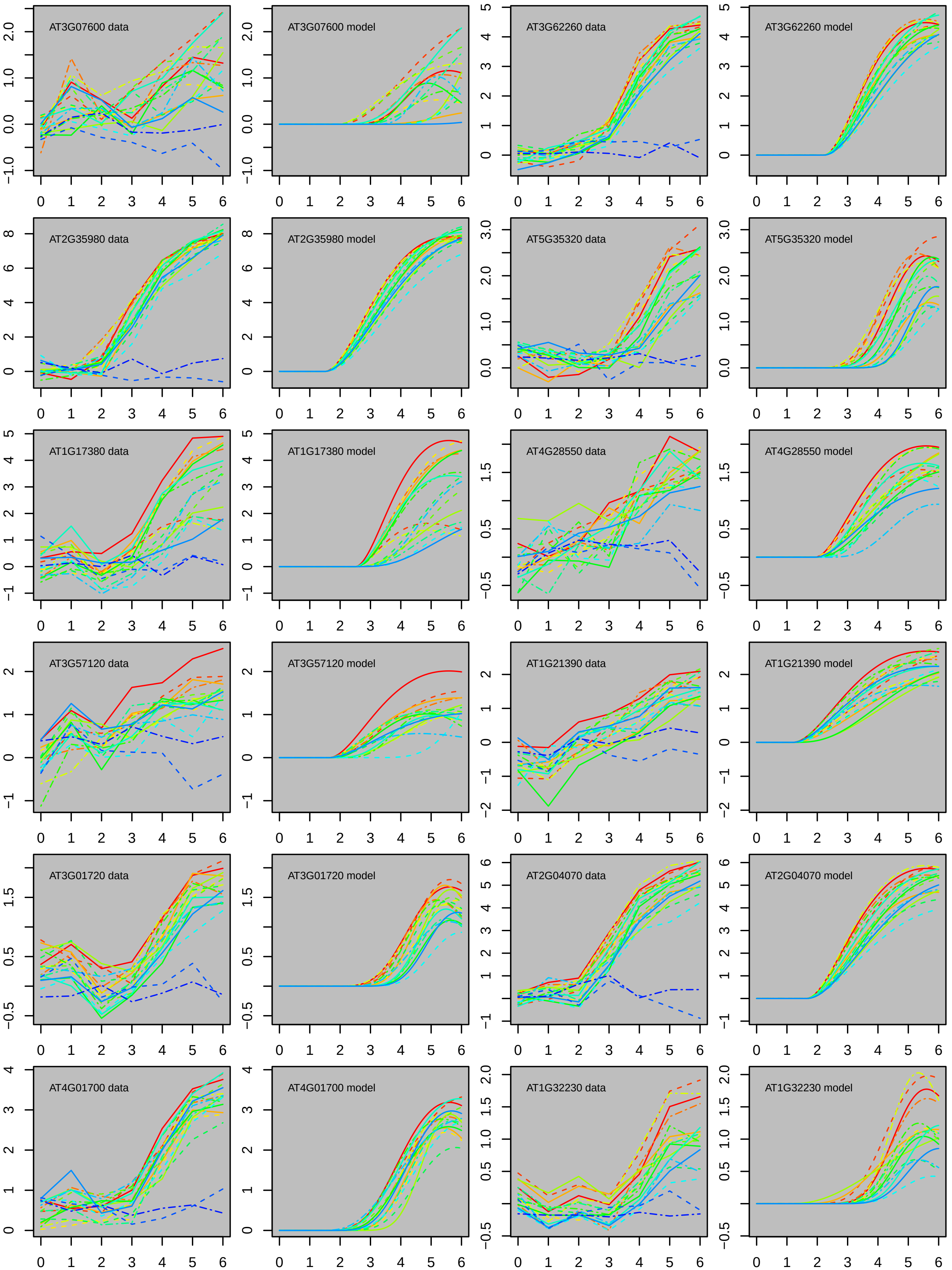
AT3G07600 data
AT3G07600 model
AT3G62260 data
AT3G62260 model
AT2G35980 data
AT2G35980 model
AT5G35320 data
AT5G35320 model
AT1G17380 data
AT1G17380 model
AT4G28550 data
AT4G28550 model
AT3G57120 data
AT3G57120 model
AT1G21390 data
AT1G21390 model
AT3G01720 data
AT3G01720 model
AT2G04070 data
AT2G04070 model
AT4G01700 data
AT4G01700 model
AT1G32230 data
AT1G32230 model

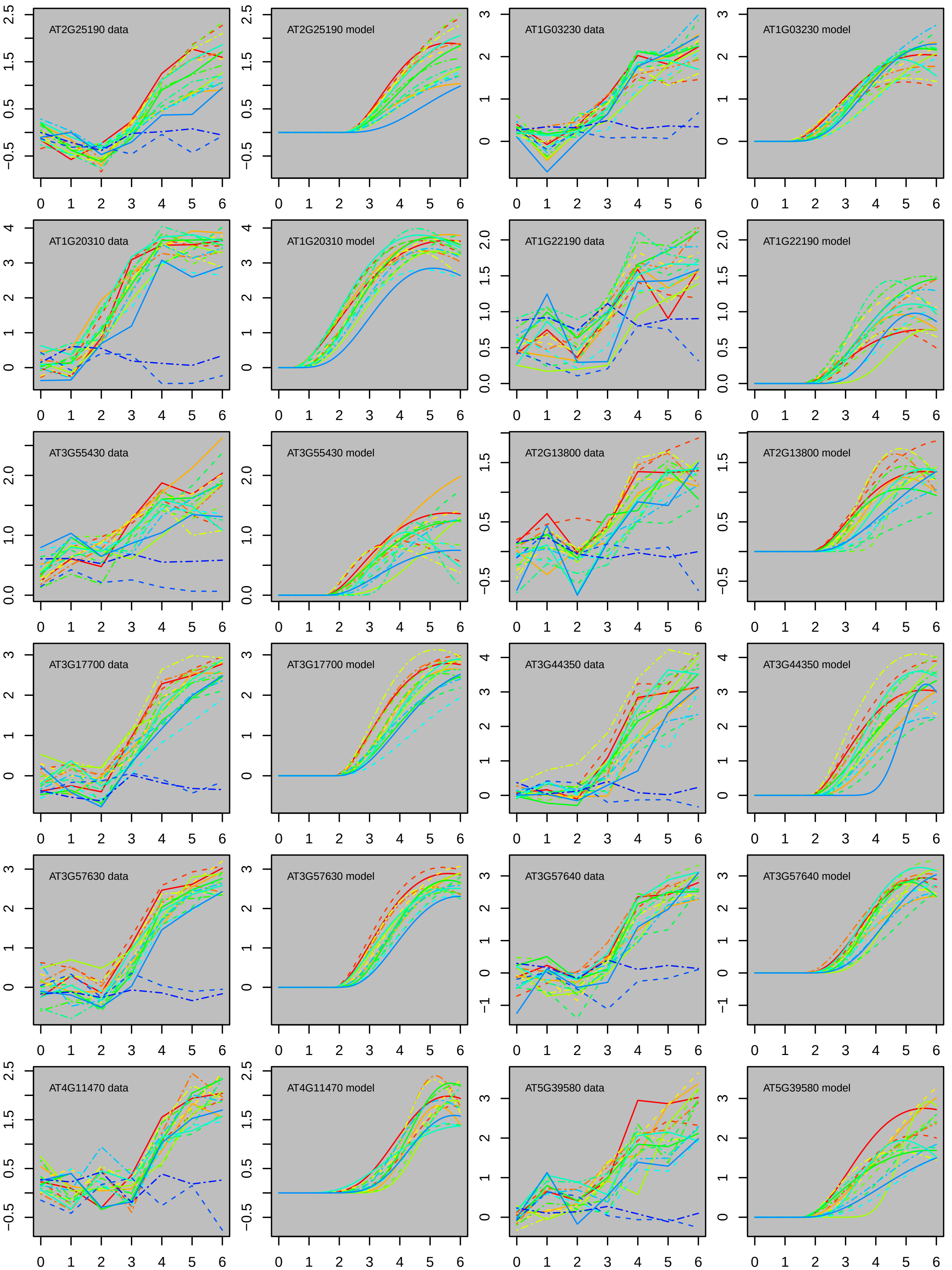
AT2G25190 data
AT2G25190 model
AT1G03230 data
AT1G03230 model
AT1G20310 data
AT1G20310 model
AT1G22190 data
AT1G22190 model
AT3G55430 data
AT3G55430 model
AT2G13800 data
AT2G13800 model
AT3G17700 data
AT3G17700 model
AT3G44350 data
AT3G44350 model
AT3G57630 data
AT3G57630 model
AT3G57640 data
AT3G57640 model
AT4G11470 data
AT4G11470 model
AT5G39580 data
AT5G39580 model

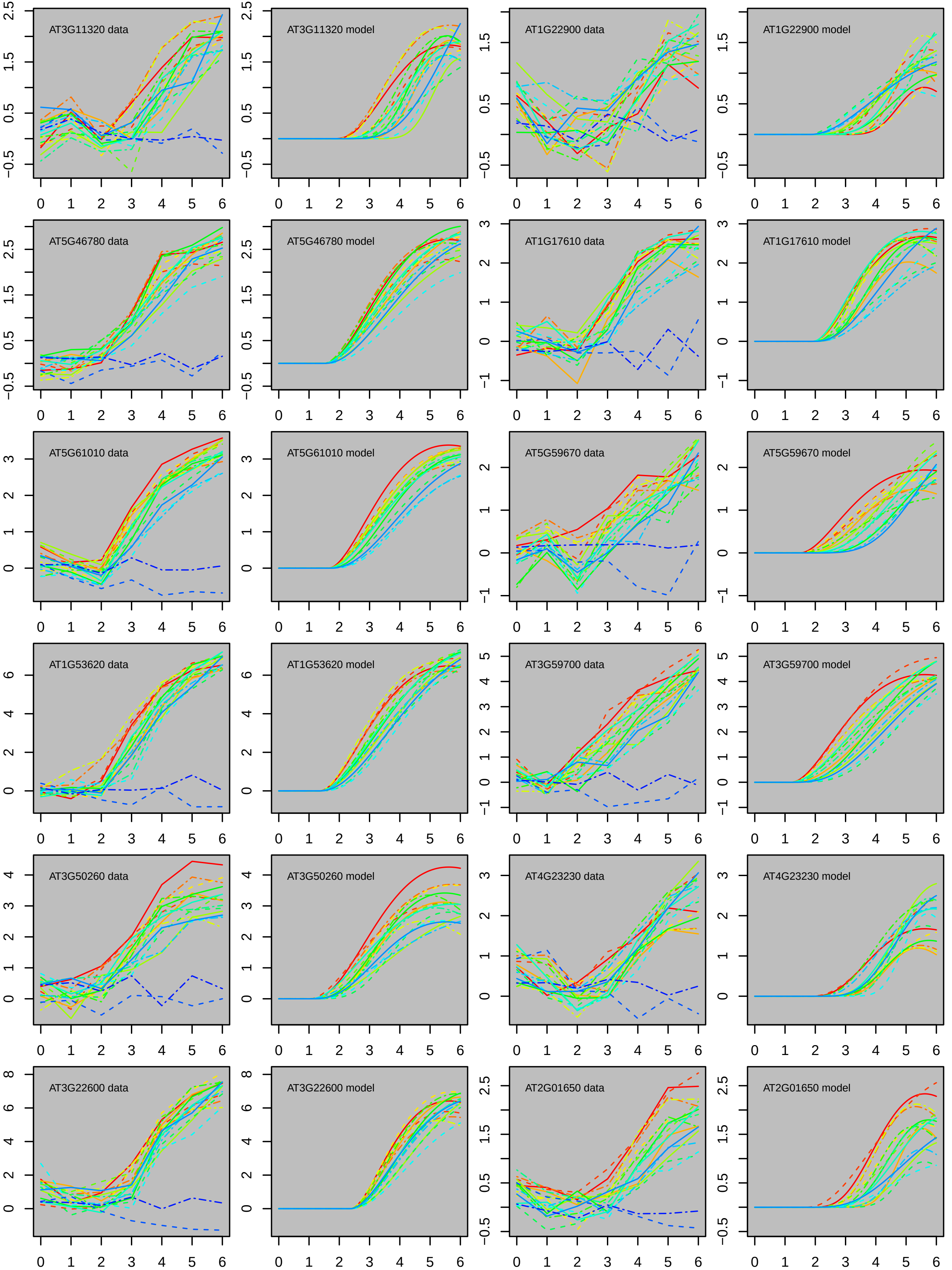

AT3G11320 data
AT3G11320 model
AT1G22900 data
AT1G22900 model
AT5G46780 data
AT5G46780 model
AT1G17610 data
AT1G17610 model
AT5G61010 data
AT5G61010 model
AT5G59670 data
AT5G59670 model
AT1G53620 data
AT1G53620 model
AT3G59700 data
AT3G59700 model
AT3G50260 data
AT3G50260 model
AT4G23230 data
AT4G23230 model
AT3G22600 data
AT3G22600 model
AT2G01650 data
AT2G01650 model

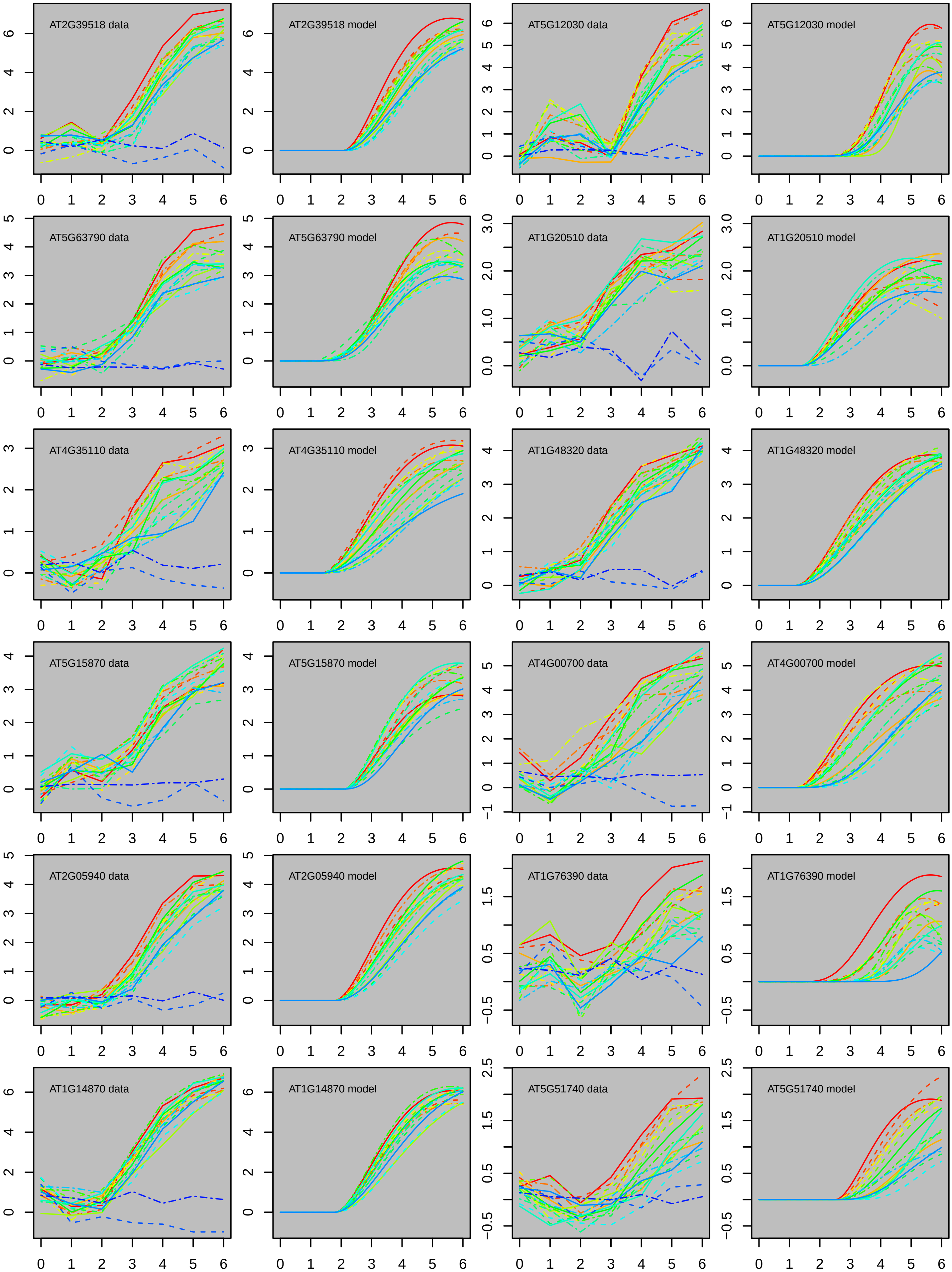
AT2G39518 data
AT2G39518 model
AT5G12030 data
AT5G12030 model
AT5G63790 data
AT5G63790 model
AT1G20510 data
AT1G20510 model
AT4G35110 data
AT4G35110 model
AT1G48320 data
AT1G48320 model
AT5G15870 data
AT5G15870 model
AT4G00700 data
AT4G00700 model
AT2G05940 data
AT2G05940 model
AT1G76390 data
AT1G76390 model
AT1G14870 data
AT1G14870 model
AT5G51740 data
AT5G51740 model

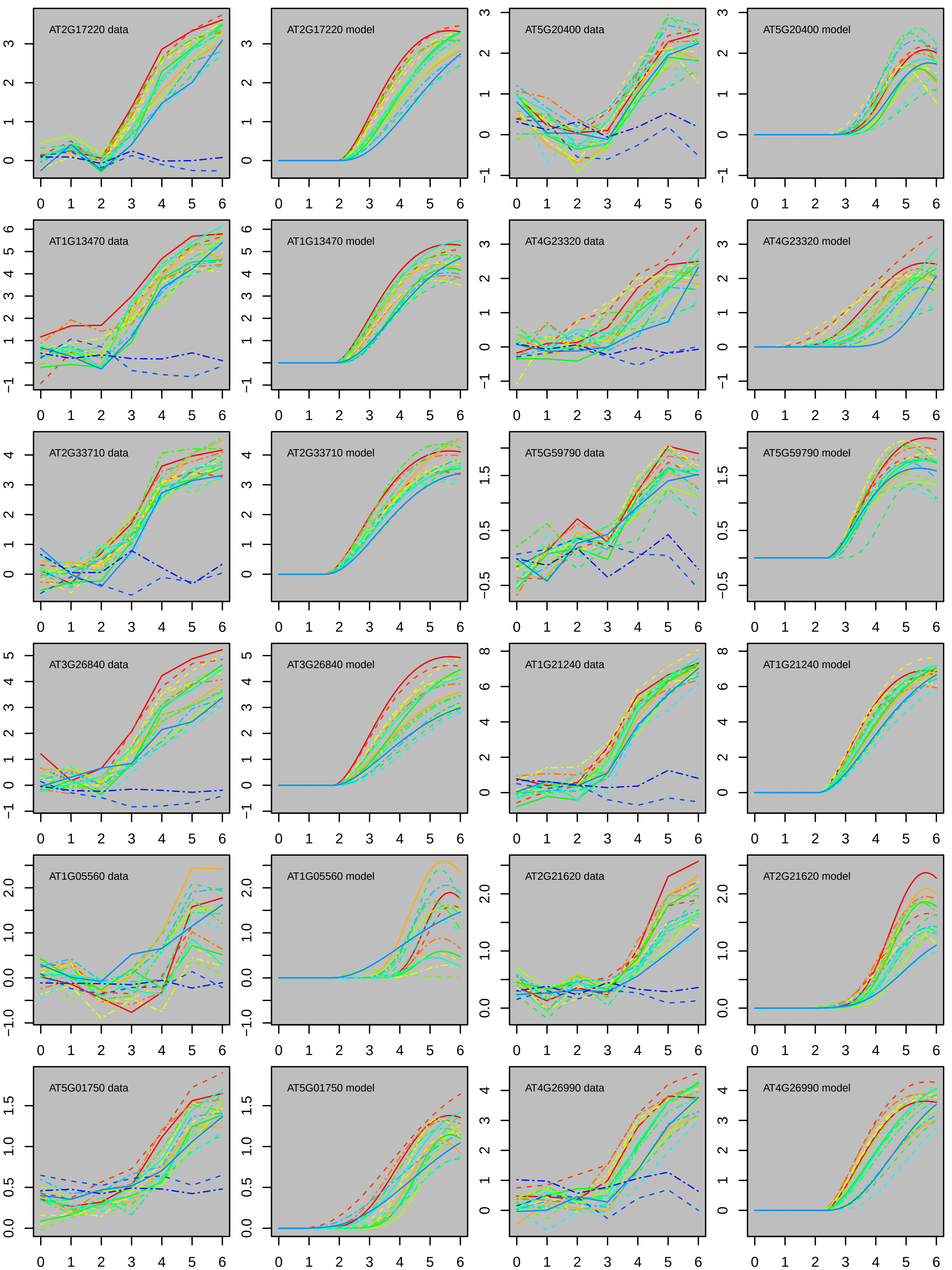

AT2G17220 data
AT2G17220 model
AT5G20400 data
AT5G20400 model
AT1G13470 data
AT1G13470 model
AT4G23320 data
AT4G23320 model
AT2G33710 data
AT2G33710 model
AT5G59790 data
AT5G59790 model
AT3G26840 data
AT3G26840 model
AT1G21240 data
AT1G21240 model
AT1G05560 data
AT1G05560 model
AT2G21620 data
AT2G21620 model
AT5G01750 data
AT5G01750 model
AT4G26990 data
AT4G26990 model

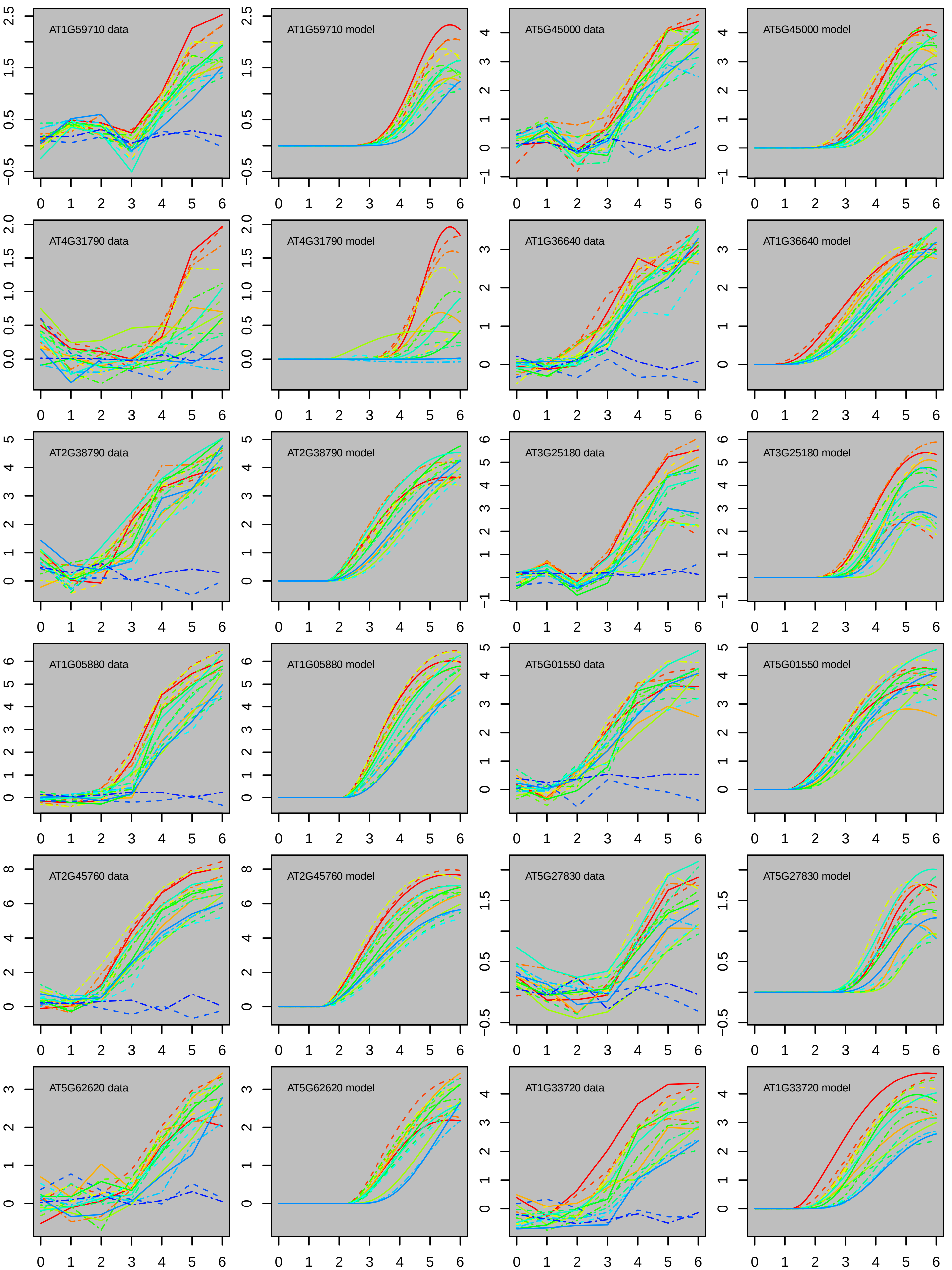
AT1G59710 data
AT1G59710 model
AT5G45000 data
AT5G45000 model
AT4G31790 data
AT4G31790 model
AT1G36640 data
AT1G36640 model
AT2G38790 data
AT2G38790 model
AT3G25180 data
AT3G25180 model
AT1G05880 data
AT1G05880 model
AT5G01550 data
AT5G01550 model
AT2G45760 data
AT2G45760 model
AT5G27830 data
AT5G27830 model
AT5G62620 data
AT5G62620 model
AT1G33720 data
AT1G33720 model

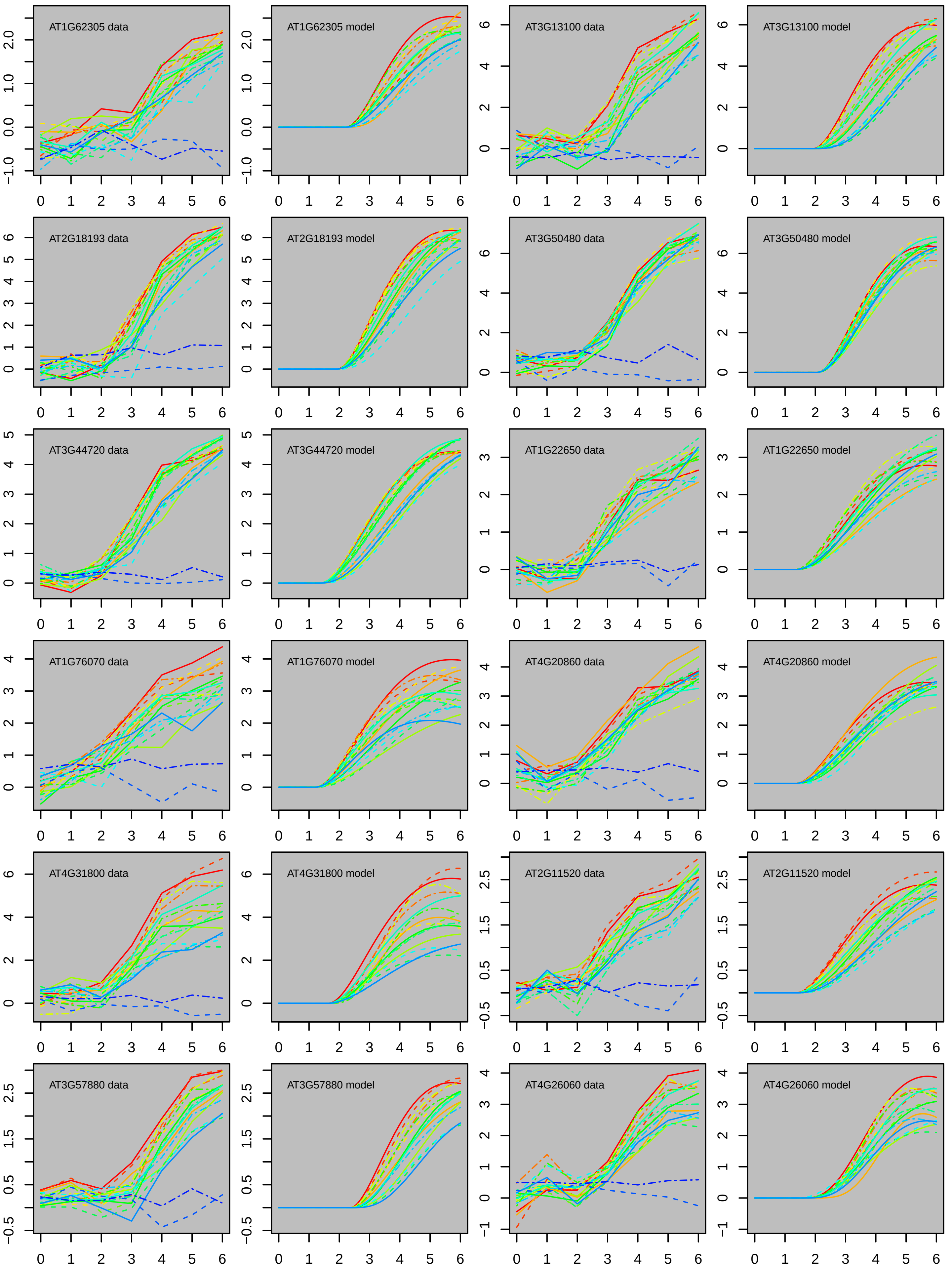
AT1G62305 data
AT1G62305 model
AT3G13100 data
AT3G13100 model
AT2G18193 data
AT2G18193 model
AT3G50480 data
AT3G50480 model
AT3G44720 data
AT3G44720 model
AT1G22650 data
AT1G22650 model
AT1G76070 data
AT1G76070 model
AT4G20860 data
AT4G20860 model
AT4G31800 data
AT4G31800 model
AT2G11520 data
AT2G11520 model
AT3G57880 data
AT3G57880 model
AT4G26060 data
AT4G26060 model

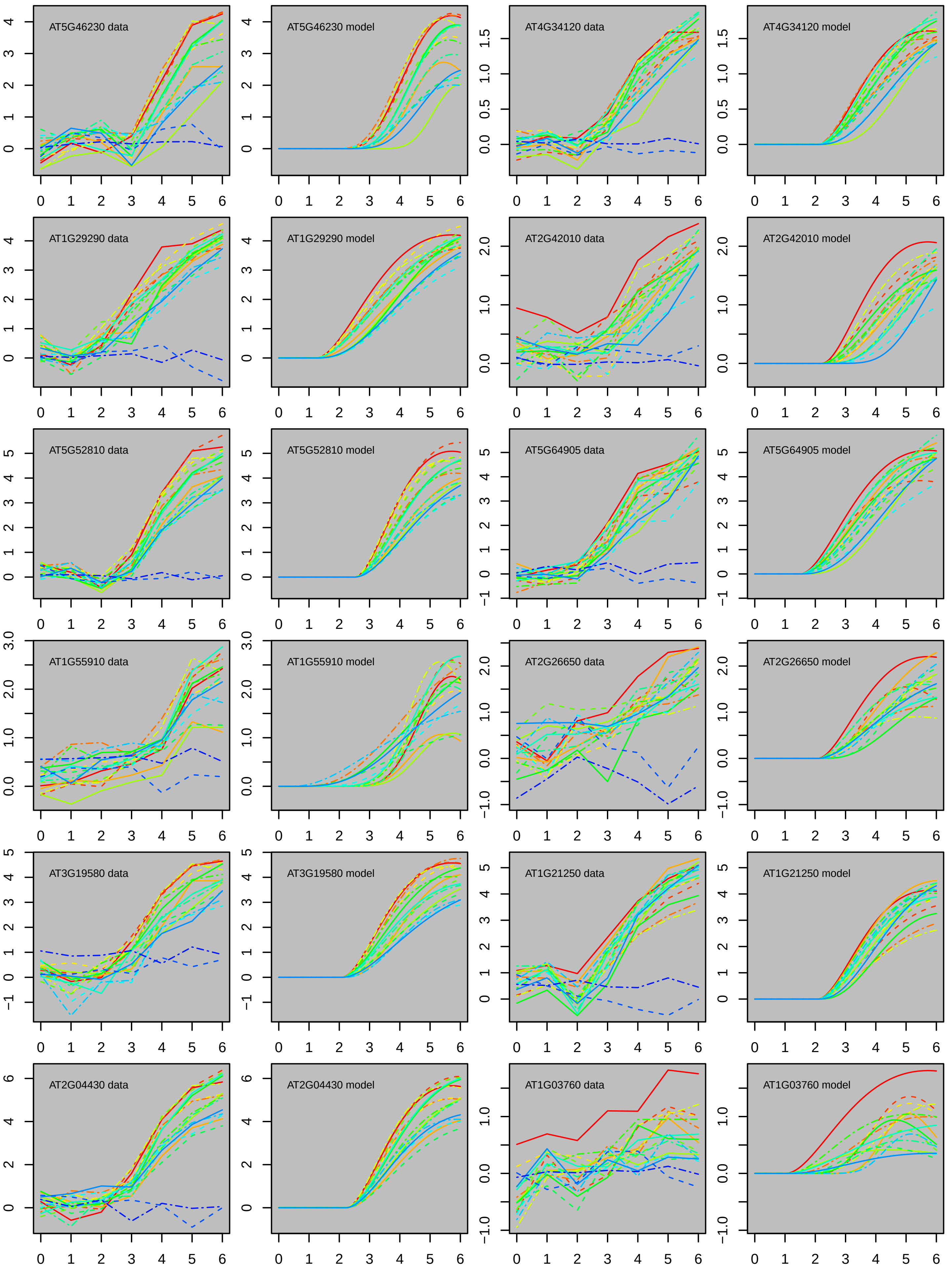
AT5G46230 data
AT5G46230 model
AT4G34120 data
AT4G34120 model
AT1G29290 data
AT1G29290 model
AT2G42010 data
AT2G42010 model
AT5G52810 data
AT5G52810 model
AT5G64905 data
AT5G64905 model
AT1G55910 data
AT1G55910 model
AT2G26650 data
AT2G26650 model
AT3G19580 data
AT3G19580 model
AT1G21250 data
AT1G21250 model
AT2G04430 data
AT2G04430 model
AT1G03760 data
AT1G03760 model

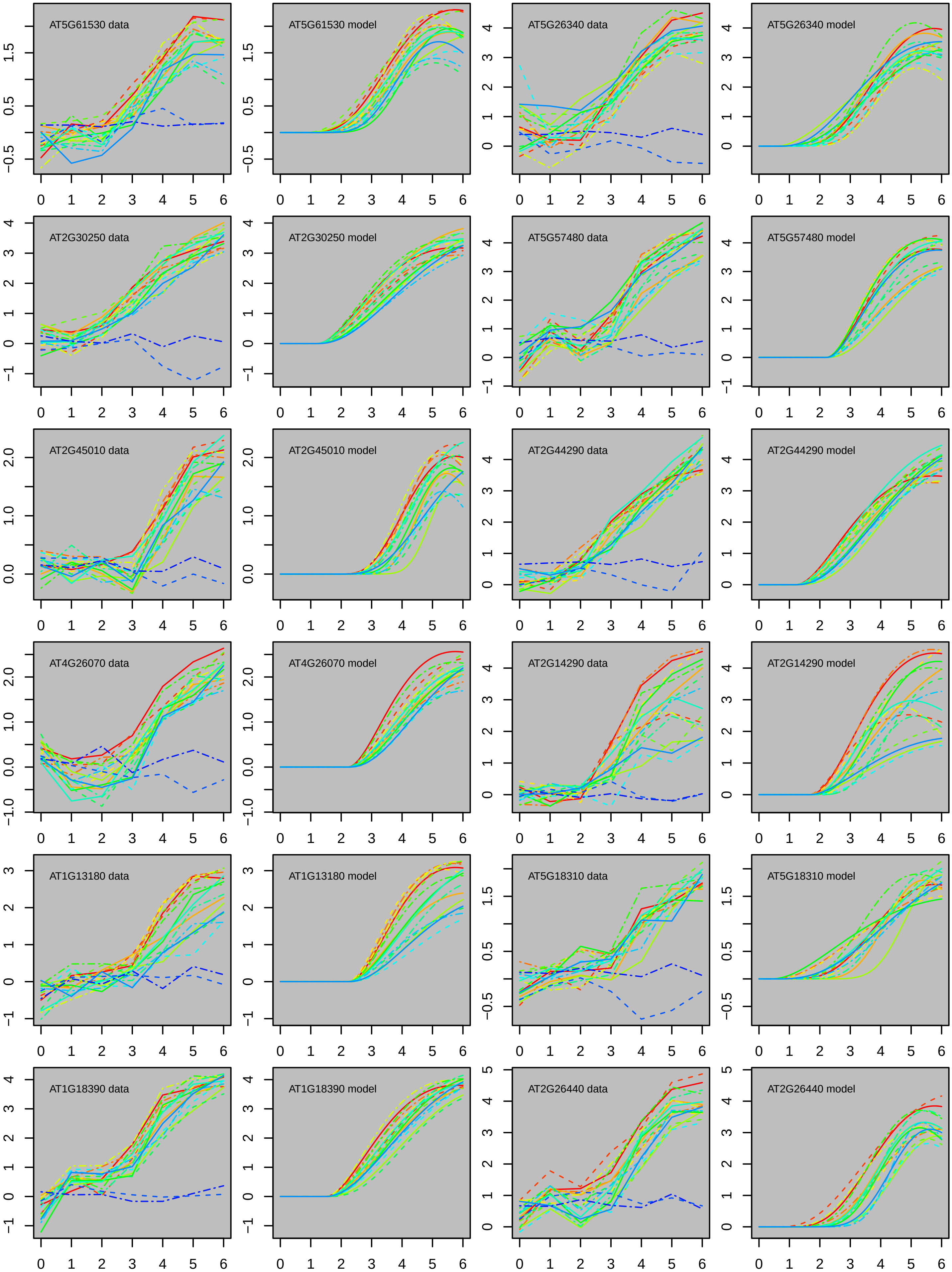
AT5G61530 data
AT5G61530 model
AT5G26340 data
AT5G26340 model
AT2G30250 data
AT2G30250 model
AT5G57480 data
AT5G57480 model
AT2G45010 data
AT2G45010 model
AT2G44290 data
AT2G44290 model
AT4G26070 data
AT4G26070 model
AT2G14290 data
AT2G14290 model
AT1G13180 data
AT1G13180 model
AT5G18310 data
AT5G18310 model
AT1G18390 data
AT1G18390 model
AT2G26440 data
AT2G26440 model

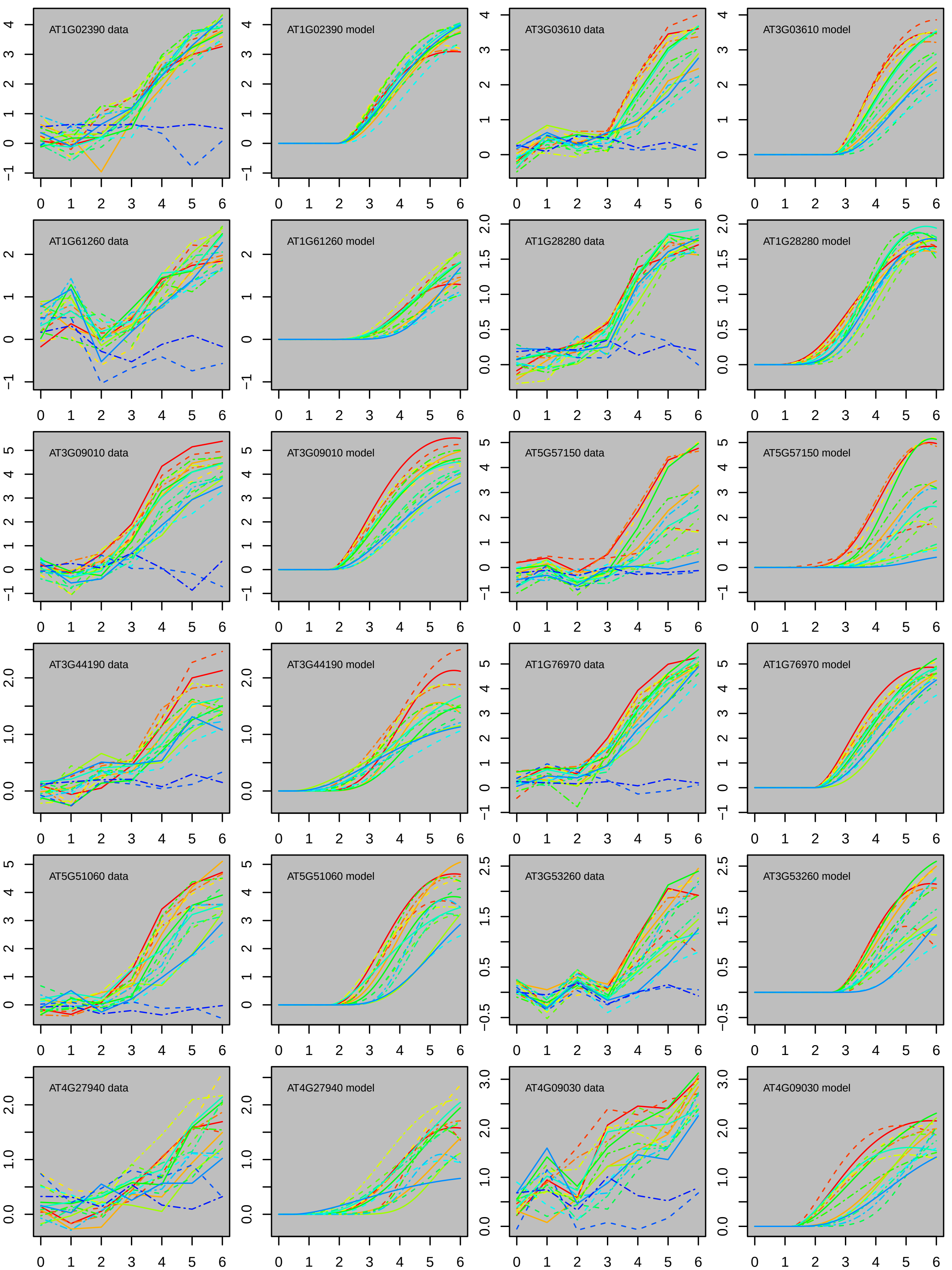
AT1G02390 data
AT1G02390 model
AT3G03610 data
AT3G03610 model
AT1G61260 data
AT1G61260 model
AT1G28280 data
AT1G28280 model
AT3G09010 data
AT3G09010 model
AT5G57150 data
AT5G57150 model
AT3G44190 data
AT3G44190 model
AT1G76970 data
AT1G76970 model
AT5G51060 data
AT5G51060 model
AT3G53260 data
AT3G53260 model
AT4G27940 data
AT4G27940 model
AT4G09030 data
AT4G09030 model

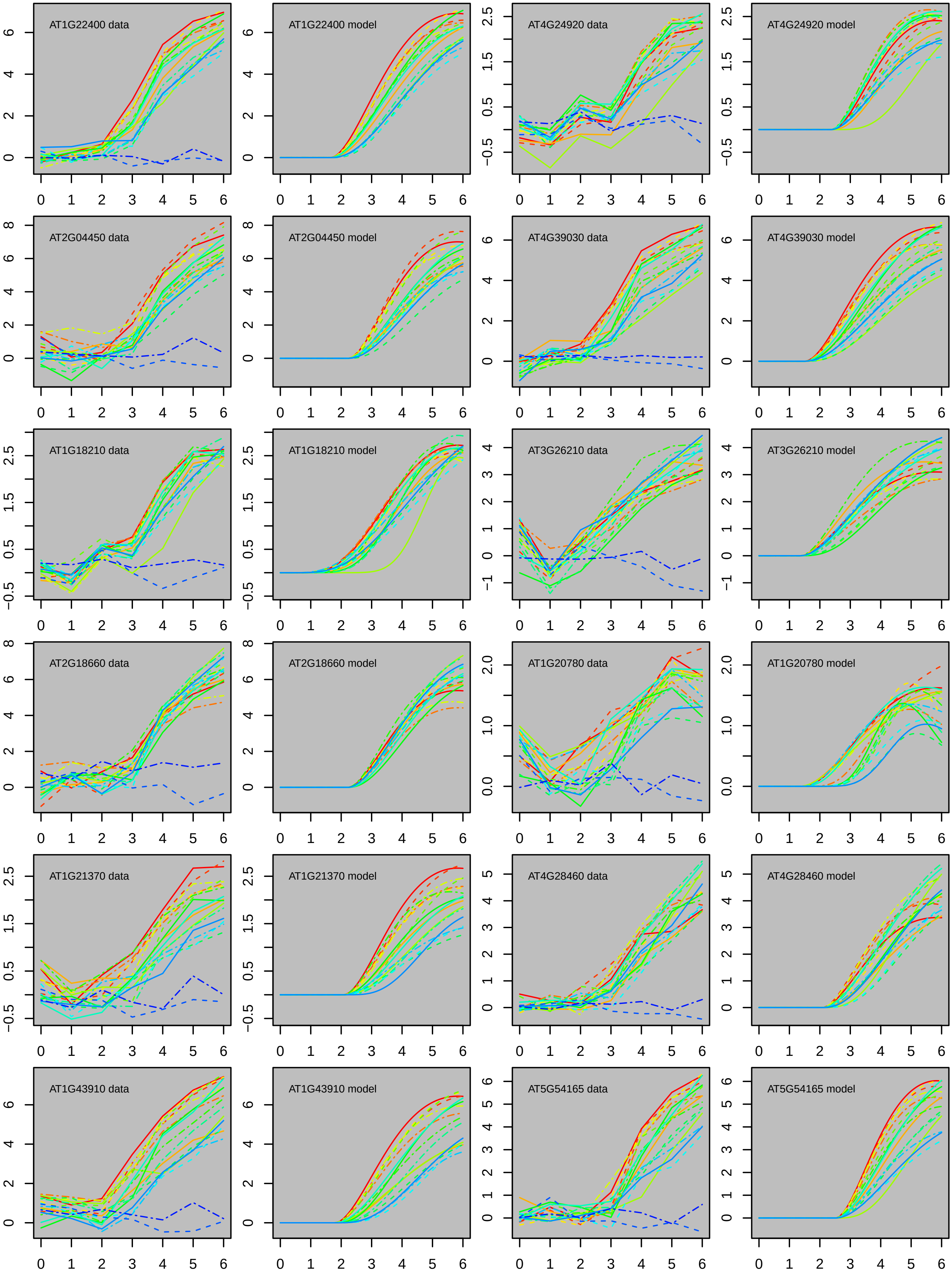
AT1G22400 data
AT1G22400 model
AT4G24920 data
AT4G24920 model
AT2G04450 data
AT2G04450 model
AT4G39030 data
AT4G39030 model
AT1G18210 data
AT1G18210 model
AT3G26210 data
AT3G26210 model
AT2G18660 data
AT2G18660 model
AT1G20780 data
AT1G20780 model
AT1G21370 data
AT1G21370 model
AT4G28460 data
AT4G28460 model
AT1G43910 data
AT1G43910 model
AT5G54165 data
AT5G54165 model

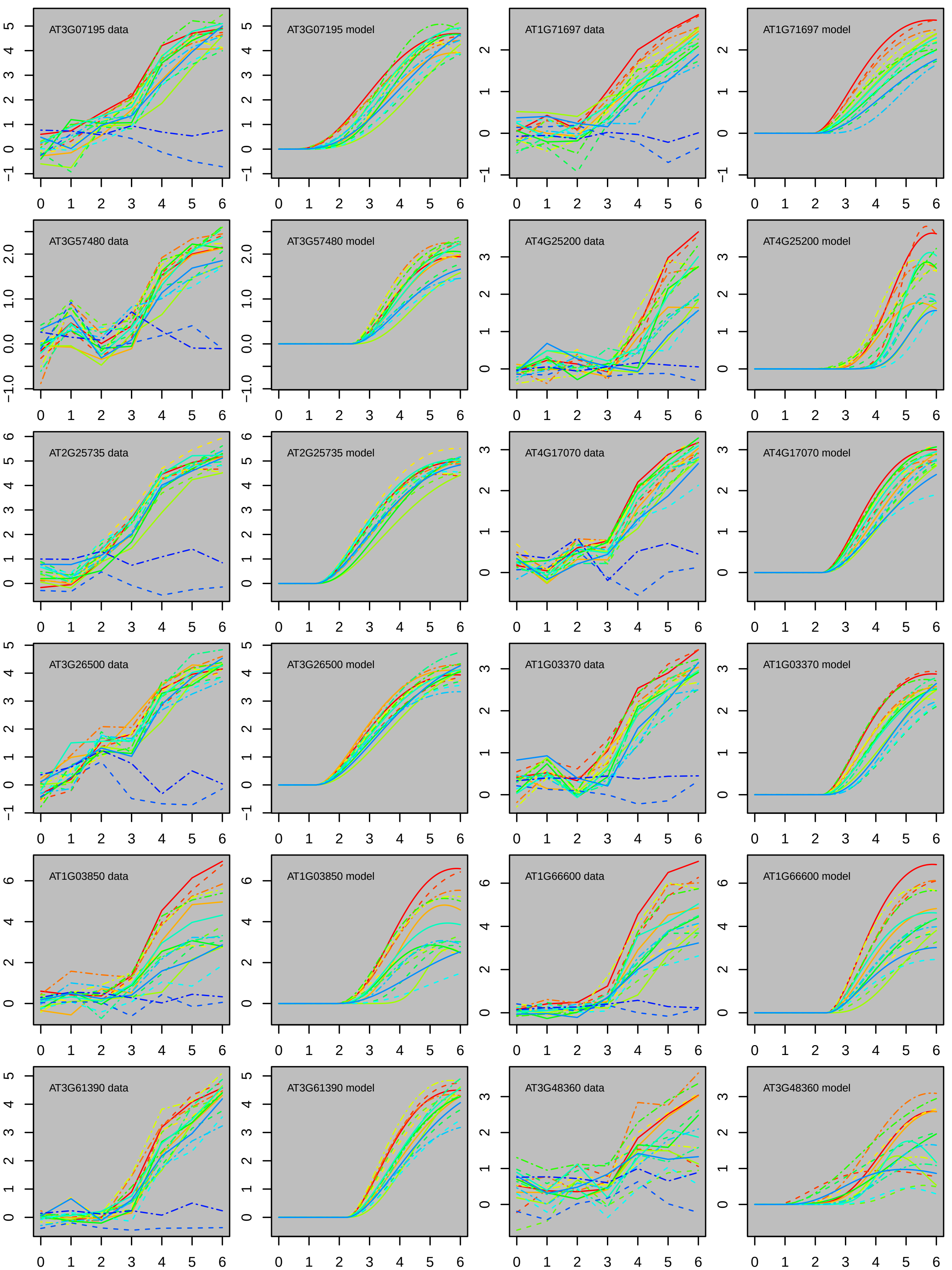
AT3G07195 data
AT3G07195 model
AT1G71697 data
AT1G71697 model
AT3G57480 data
AT3G57480 model
AT4G25200 data
AT4G25200 model
AT2G25735 data
AT2G25735 model
AT4G17070 data
AT4G17070 model
AT3G26500 data
AT3G26500 model
AT1G03370 data
AT1G03370 model
AT1G03850 data
AT1G03850 model
AT1G66600 data
AT1G66600 model
AT3G61390 data
AT3G61390 model
AT3G48360 data
AT3G48360 model

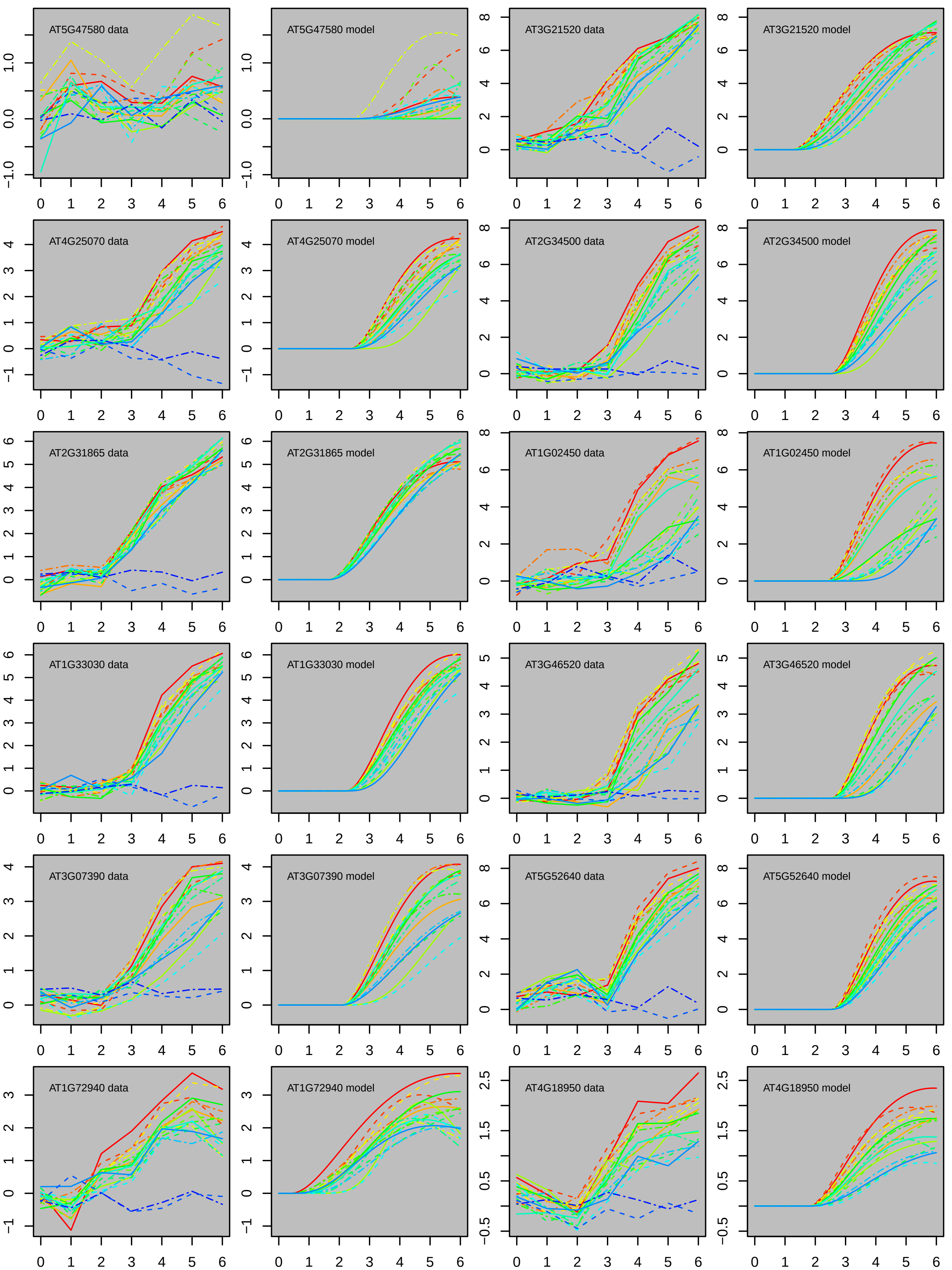
AT5G47580 data
AT5G47580 model
AT3G21520 data
AT3G21520 model
AT4G25070 data
AT4G25070 model
AT2G34500 data
AT2G34500 model
AT2G31865 data
AT2G31865 model
AT1G02450 data
AT1G02450 model
AT1G33030 data
AT1G33030 model
AT3G46520 data
AT3G46520 model
AT3G07390 data
AT3G07390 model
AT5G52640 data
AT5G52640 model
AT1G72940 data
AT1G72940 model
AT4G18950 data
AT4G18950 model

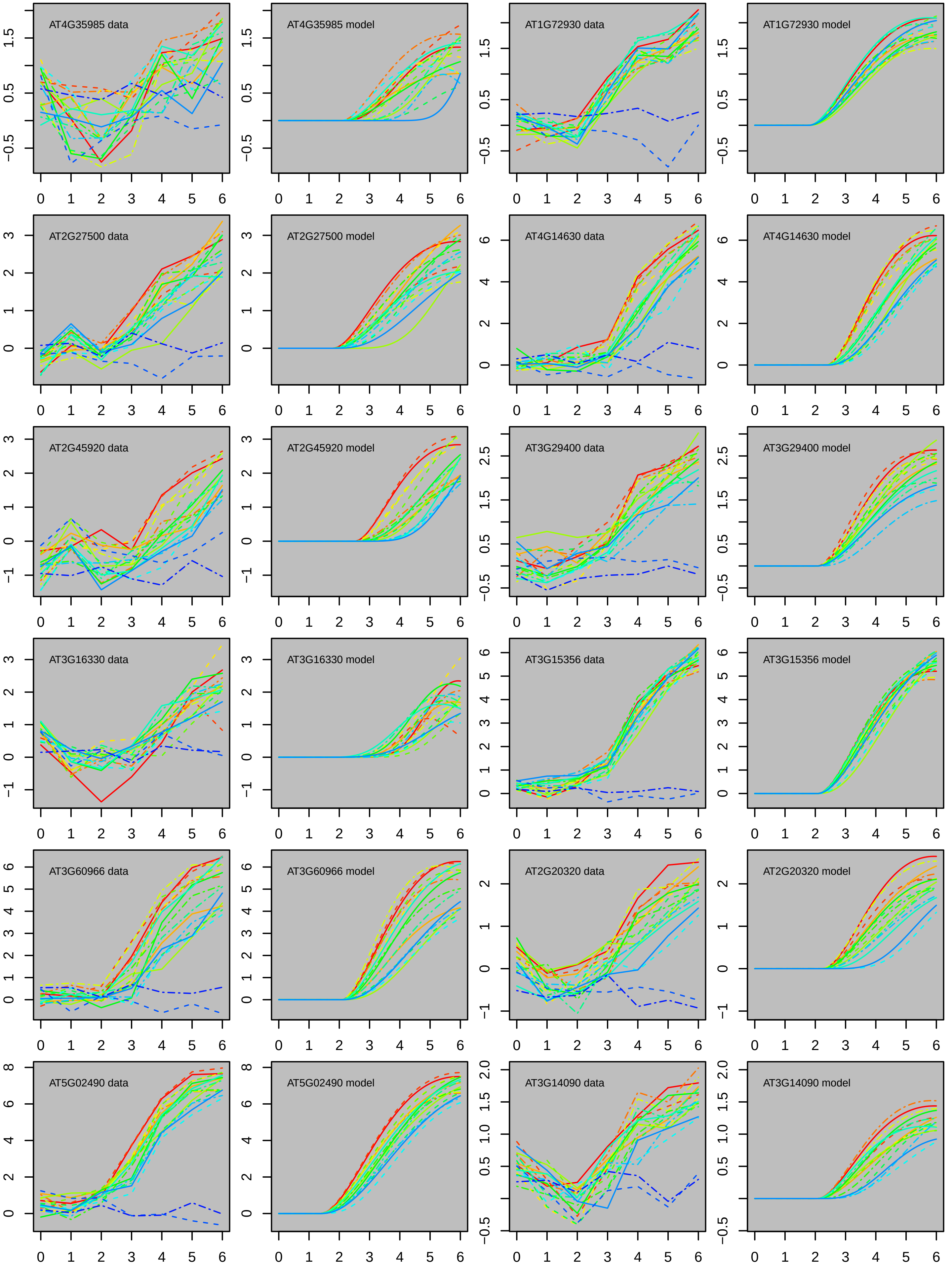
AT4G35985 data
AT4G35985 model
AT1G72930 data
AT1G72930 model
AT2G27500 data
AT2G27500 model
AT4G14630 data
AT4G14630 model
AT2G45920 data
AT2G45920 model
AT3G29400 data
AT3G29400 model
AT3G16330 data
AT3G16330 model
AT3G15356 data
AT3G15356 model
AT3G60966 data
AT3G60966 model
AT2G20320 data
AT2G20320 model
AT5G02490 data
AT5G02490 model
AT3G14090 data
AT3G14090 model

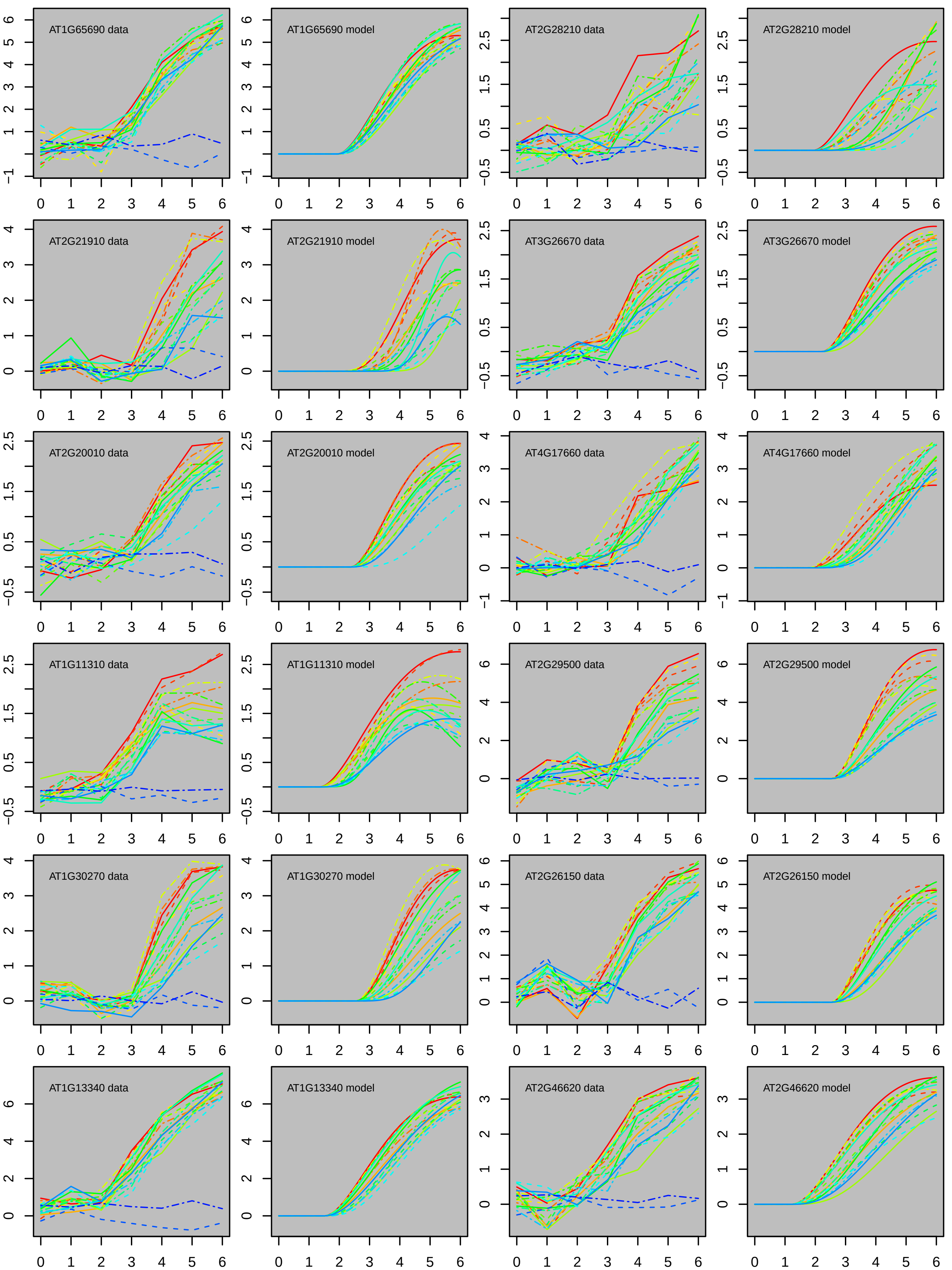
AT1G65690 data
AT1G65690 model
AT2G28210 data
AT2G28210 model
AT2G21910 data
AT2G21910 model
AT3G26670 data
AT3G26670 model
AT2G20010 data
AT2G20010 model
AT4G17660 data
AT4G17660 model
AT1G11310 data
AT1G11310 model
AT2G29500 data
AT2G29500 model
AT1G30270 data
AT1G30270 model
AT2G26150 data
AT2G26150 model
AT1G13340 data
AT1G13340 model
AT2G46620 data
AT2G46620 model
0 1 2 3 4 5 6

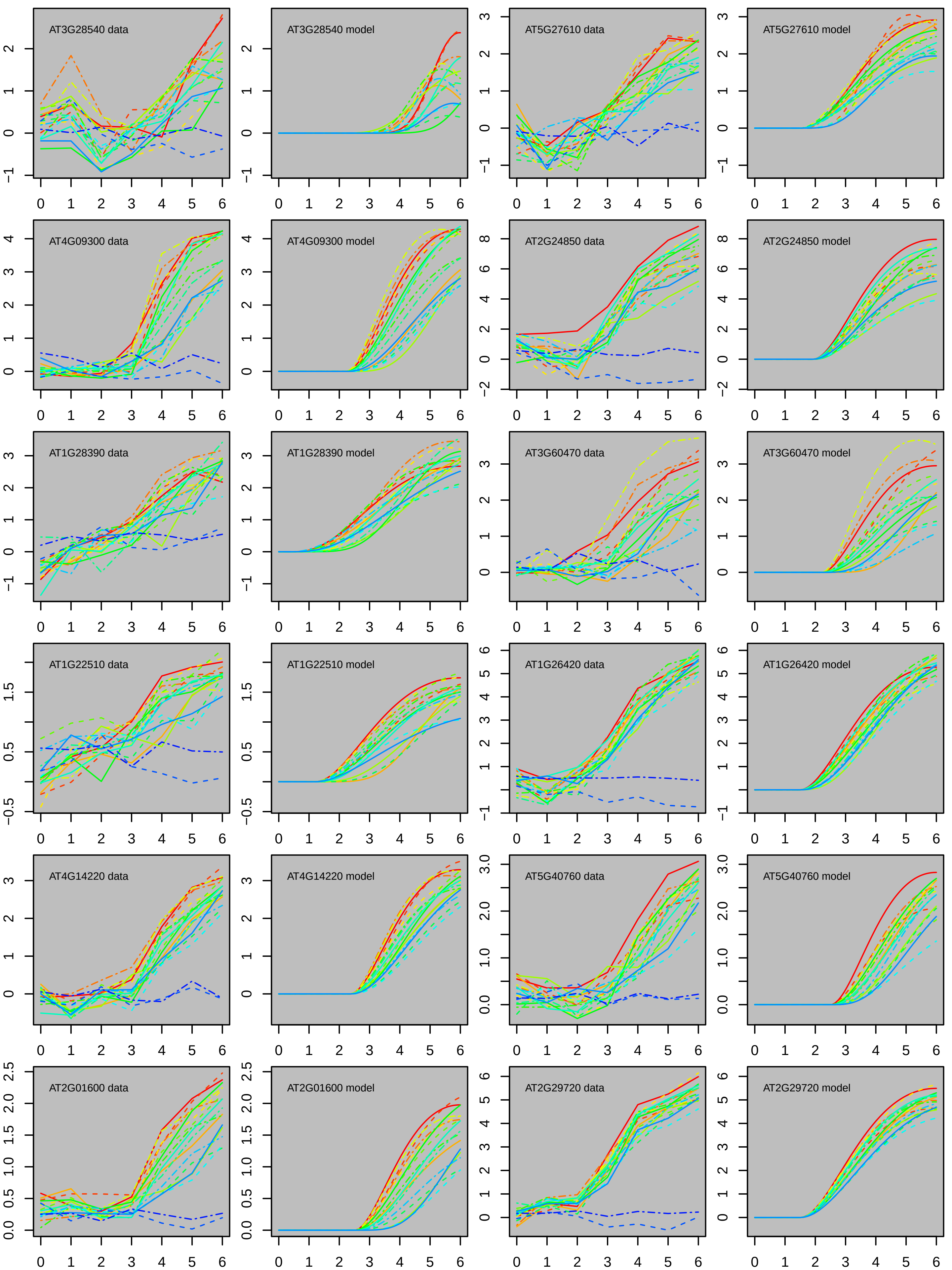
AT3G28540 data
AT3G28540 model
AT5G27610 data
AT5G27610 model
AT4G09300 data
AT4G09300 model
AT2G24850 data
AT2G24850 model
AT1G28390 data
AT1G28390 model
AT3G60470 data
AT3G60470 model
AT1G22510 data
AT1G22510 model
AT1G26420 data
AT1G26420 model
AT4G14220 data
AT4G14220 model
AT5G40760 data
AT5G40760 model
AT2G01600 data
AT2G01600 model
AT2G29720 data
AT2G29720 model

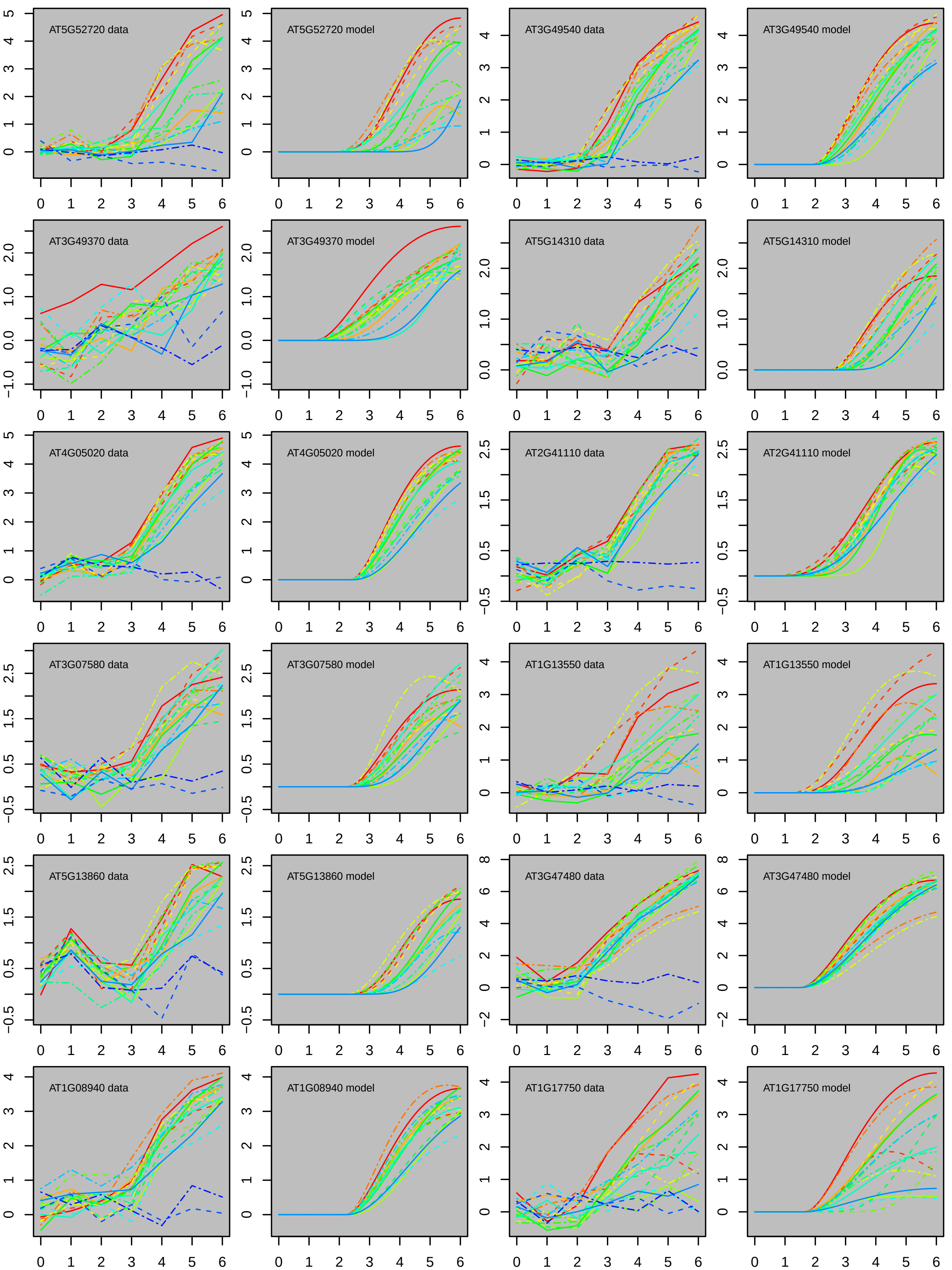

AT5G52720 data
AT5G52720 model
AT3G49540 data
AT3G49540 model
AT3G49370 data
AT3G49370 model
AT5G14310 data
AT5G14310 model
AT4G05020 data
AT4G05020 model
AT2G41110 data
AT2G41110 model
AT3G07580 data
AT3G07580 model
AT1G13550 data
AT1G13550 model
AT5G13860 data
AT5G13860 model
AT3G47480 data
AT3G47480 model
AT1G08940 data
AT1G08940 model
AT1G17750 data
AT1G17750 model

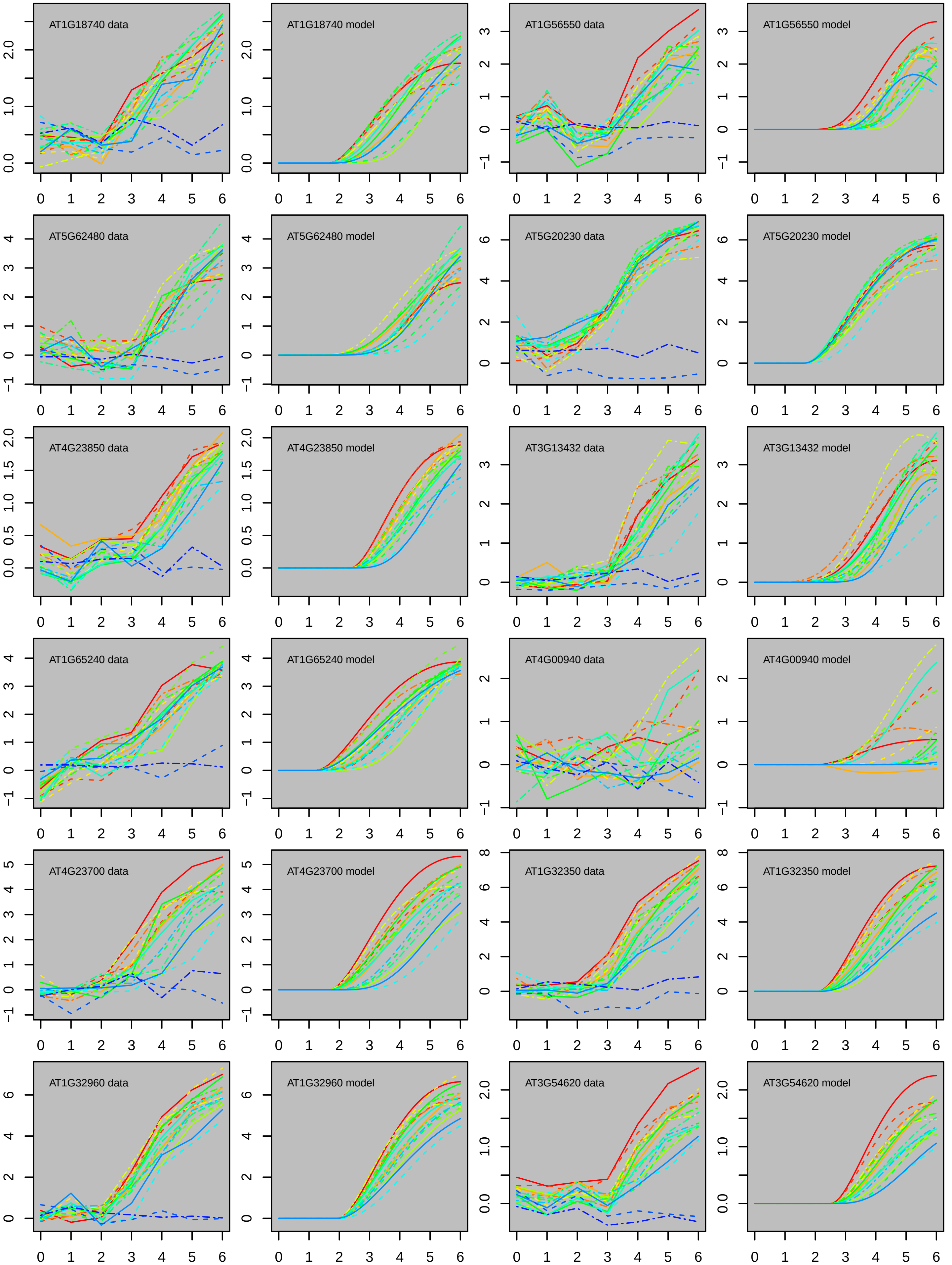
AT1G18740 data
AT1G18740 model
AT1G56550 data
AT1G56550 model
AT5G62480 data
AT5G62480 model
AT5G20230 data
AT5G20230 model
AT4G23850 data
AT4G23850 model
AT3G13432 data
AT3G13432 model
AT1G65240 data
AT1G65240 model
AT4G00940 data
AT4G00940 model
AT4G23700 data
AT4G23700 model
AT1G32350 data
AT1G32350 model
AT1G32960 data
AT1G32960 model
AT3G54620 data
AT3G54620 model

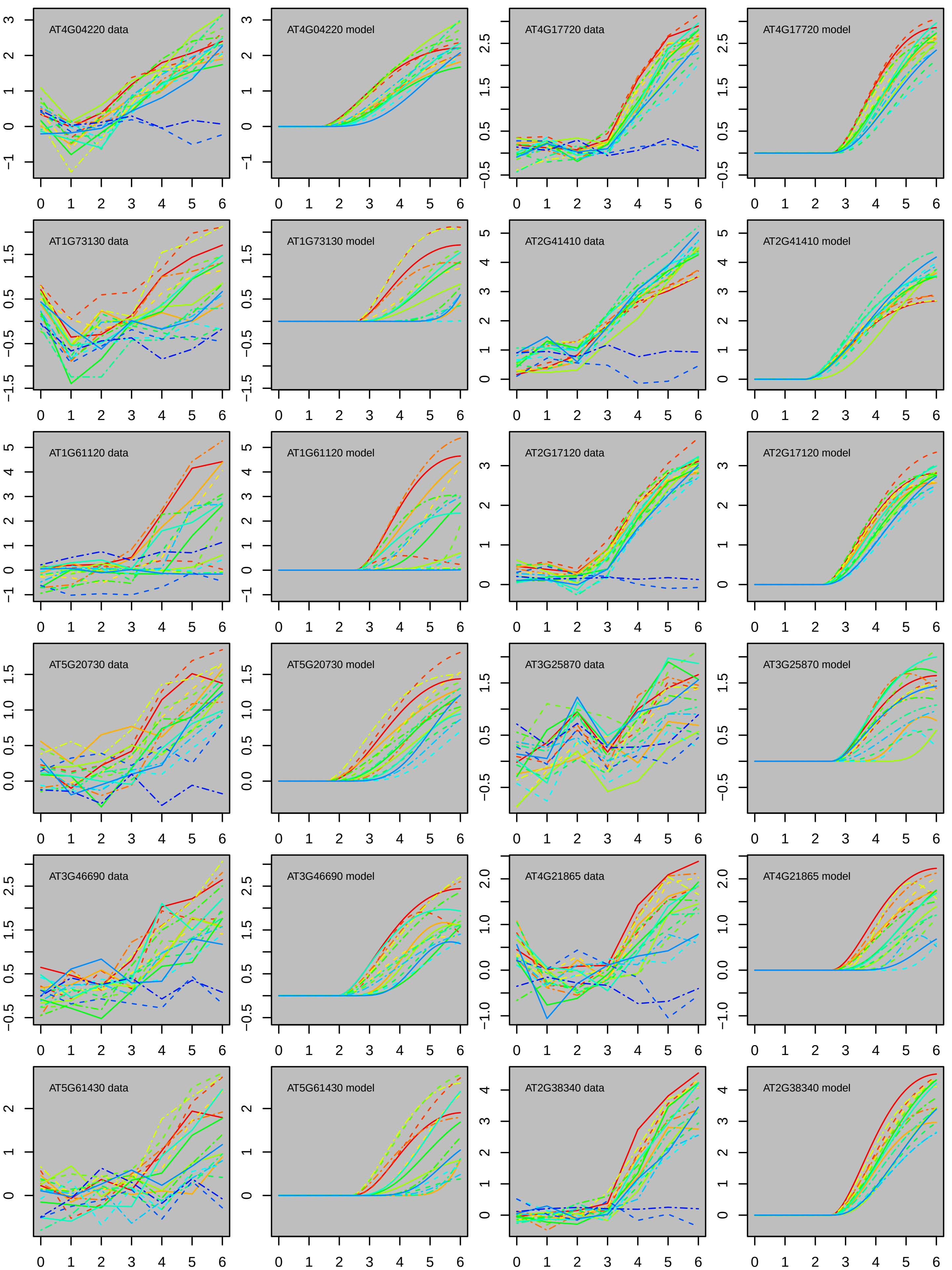

AT4G04220 data
AT4G04220 model
AT4G17720 data
AT4G17720 model
AT1G73130 data
AT1G73130 model
AT2G41410 data
AT2G41410 model
AT1G61120 data
AT1G61120 model
AT2G17120 data
AT2G17120 model
AT5G20730 data
AT5G20730 model
AT3G25870 data
AT3G25870 model
AT3G46690 data
AT3G46690 model
AT4G21865 data
AT4G21865 model
AT5G61430 data
AT5G61430 model
AT2G38340 data
AT2G38340 model

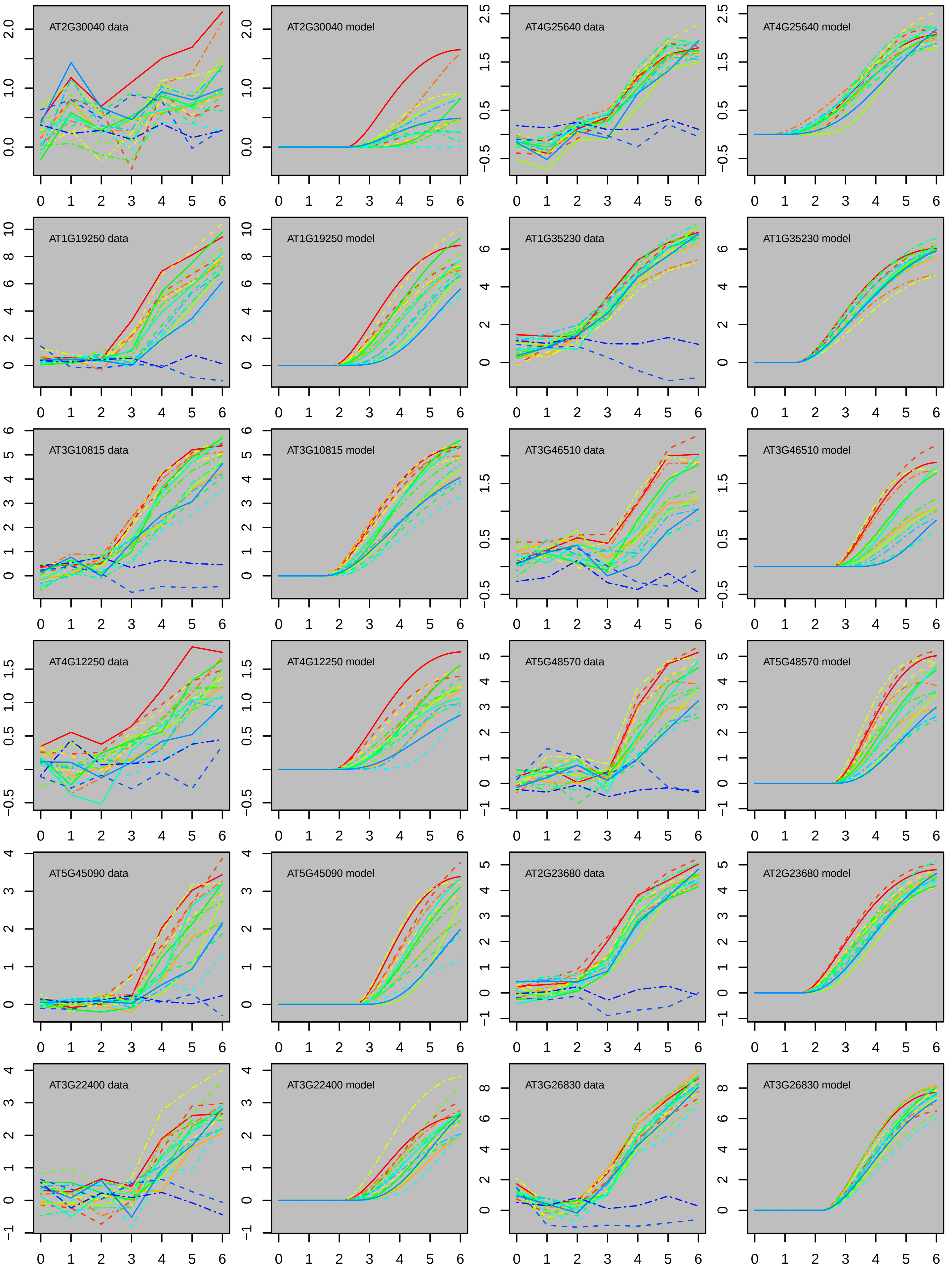

AT2G30040 data
AT2G30040 model
AT4G25640 data
AT4G25640 model
AT1G19250 data
AT1G19250 model
AT1G35230 data
AT1G35230 model
AT3G10815 data
AT3G10815 model
AT3G46510 data
AT3G46510 model
AT4G12250 data
AT4G12250 model
AT5G48570 data
AT5G48570 model
AT5G45090 data
AT5G45090 model
AT2G23680 data
AT2G23680 model
AT3G22400 data
AT3G22400 model
AT3G26830 data
AT3G26830 model
0 1 2 3 4 5 6

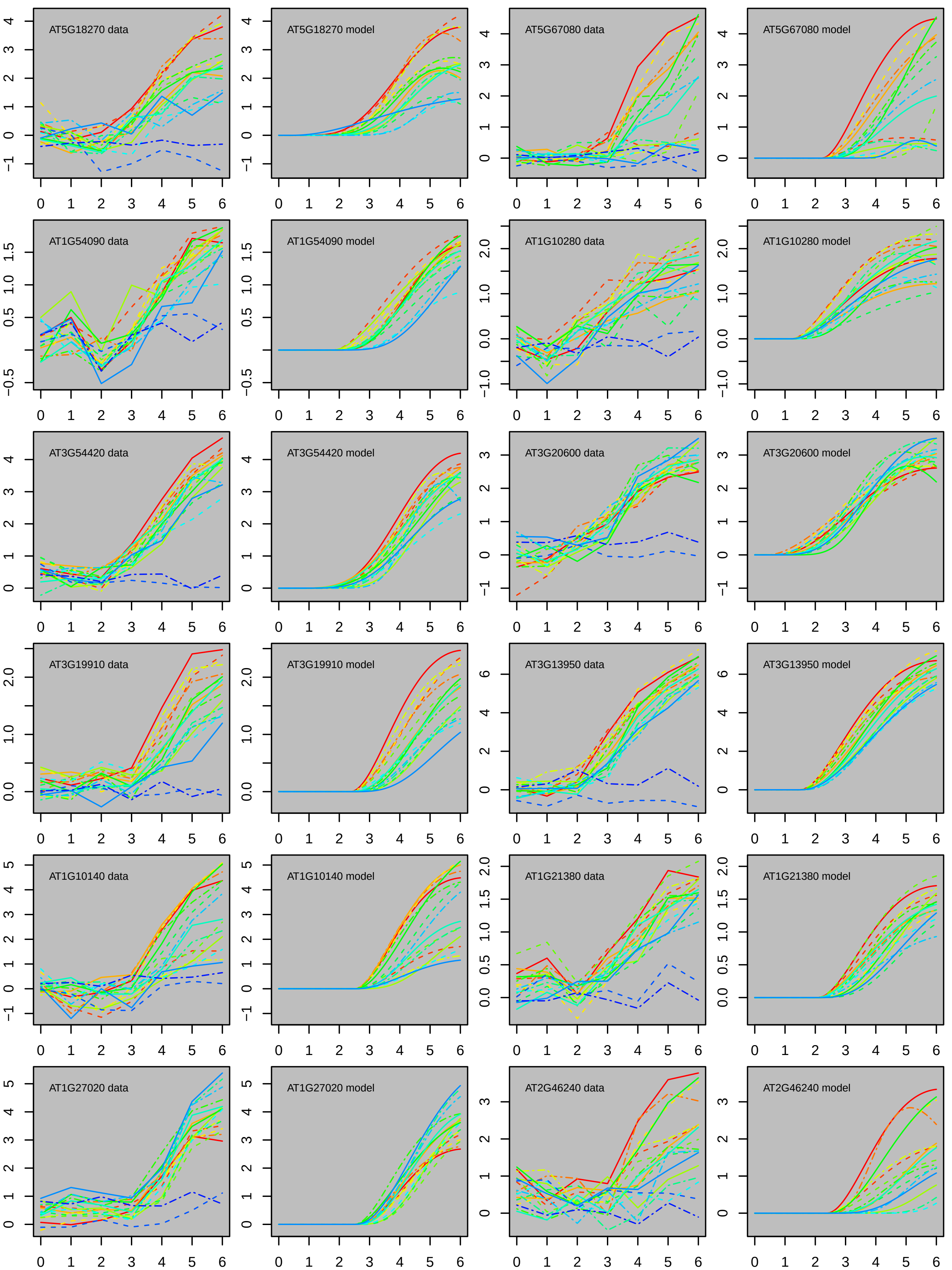
AT5G18270 data
AT5G18270 model
AT5G67080 data
AT5G67080 model
AT1G54090 data
AT1G54090 model
AT1G10280 data
AT1G10280 model
AT3G54420 data
AT3G54420 model
AT3G20600 data
AT3G20600 model
AT3G19910 data
AT3G19910 model
AT3G13950 data
AT3G13950 model
AT1G10140 data
AT1G10140 model
AT1G21380 data
AT1G21380 model
AT1G27020 data
AT1G27020 model
AT2G46240 data
AT2G46240 model

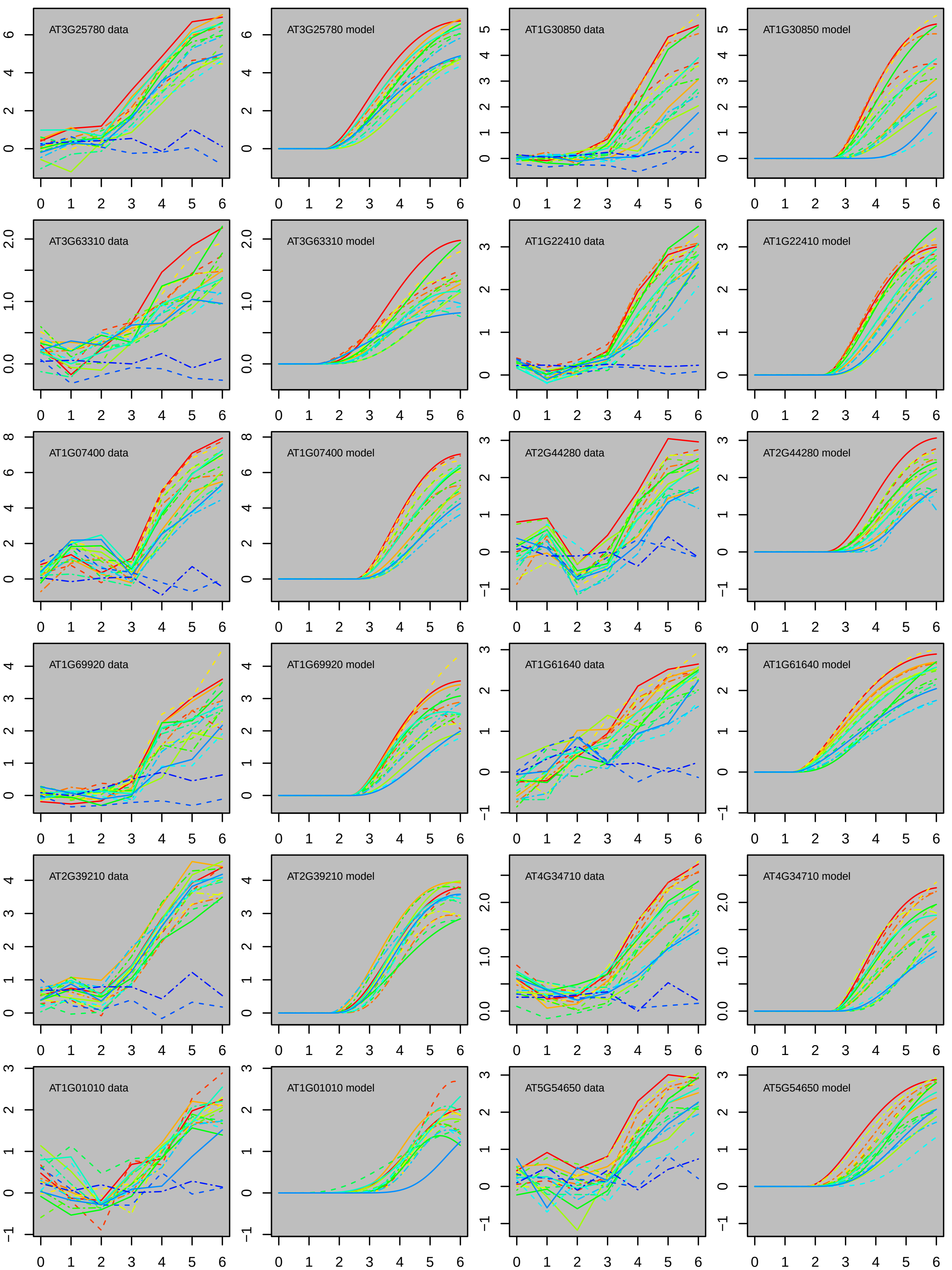
AT3G25780 data
AT3G25780 model
AT1G30850 data
AT1G30850 model
AT3G63310 data
AT3G63310 model
AT1G22410 data
AT1G22410 model
AT1G07400 data
AT1G07400 model
AT2G44280 data
AT2G44280 model
AT1G69920 data
AT1G69920 model
AT1G61640 data
AT1G61640 model
AT2G39210 data
AT2G39210 model
AT4G34710 data
AT4G34710 model
AT1G01010 data
AT1G01010 model
AT5G54650 data
AT5G54650 model

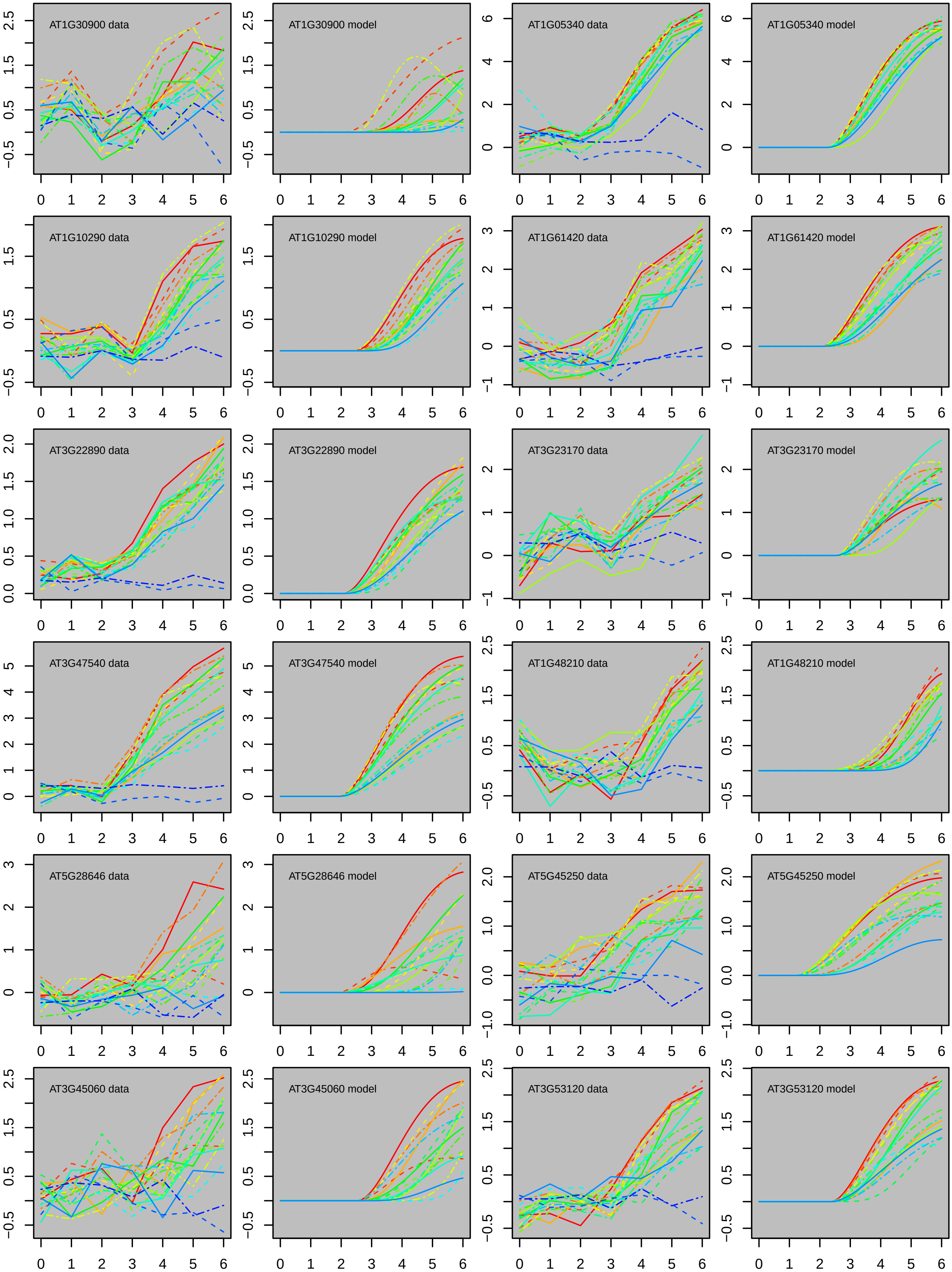
AT1G30900 data
AT1G30900 model
AT1G05340 data
AT1G05340 model
AT1G10290 data
AT1G10290 model
AT1G61420 data
AT1G61420 model
AT3G22890 data
AT3G22890 model
AT3G23170 data
AT3G23170 model
AT3G47540 data
AT3G47540 model
AT1G48210 data
AT1G48210 model
AT5G28646 data
AT5G28646 model
AT5G45250 data
AT5G45250 model
AT3G45060 data
AT3G45060 model
AT3G53120 data
AT3G53120 model

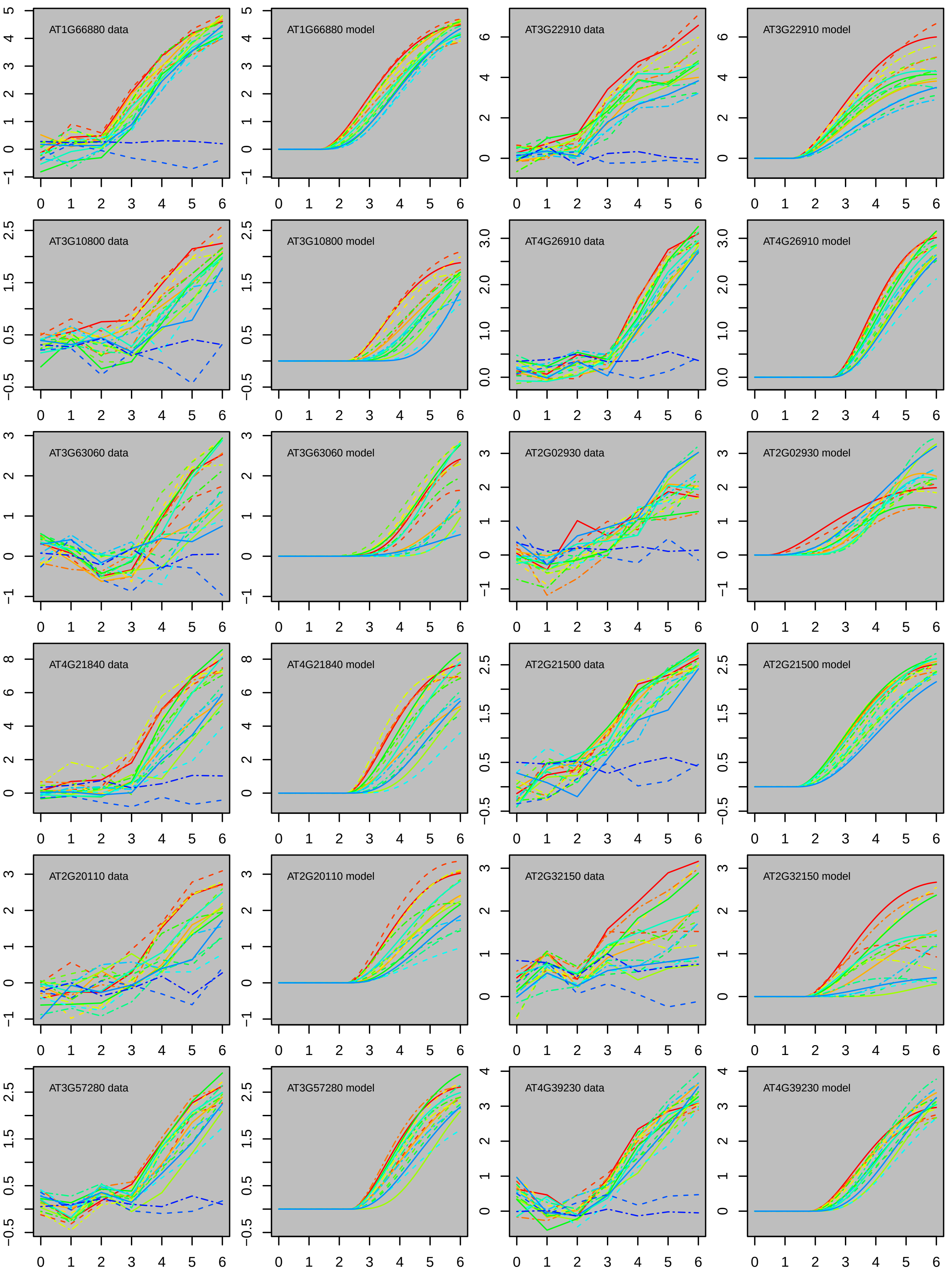

AT1G66880 data
AT1G66880 model
AT3G22910 data
AT3G22910 model
AT3G10800 data
AT3G10800 model
AT4G26910 data
AT4G26910 model
AT3G63060 data
AT3G63060 model
AT2G02930 data
AT2G02930 model
AT4G21840 data
AT4G21840 model
AT2G21500 data
AT2G21500 model
AT2G20110 data
AT2G20110 model
AT2G32150 data
AT2G32150 model
AT3G57280 data
AT3G57280 model
AT4G39230 data
AT4G39230 model

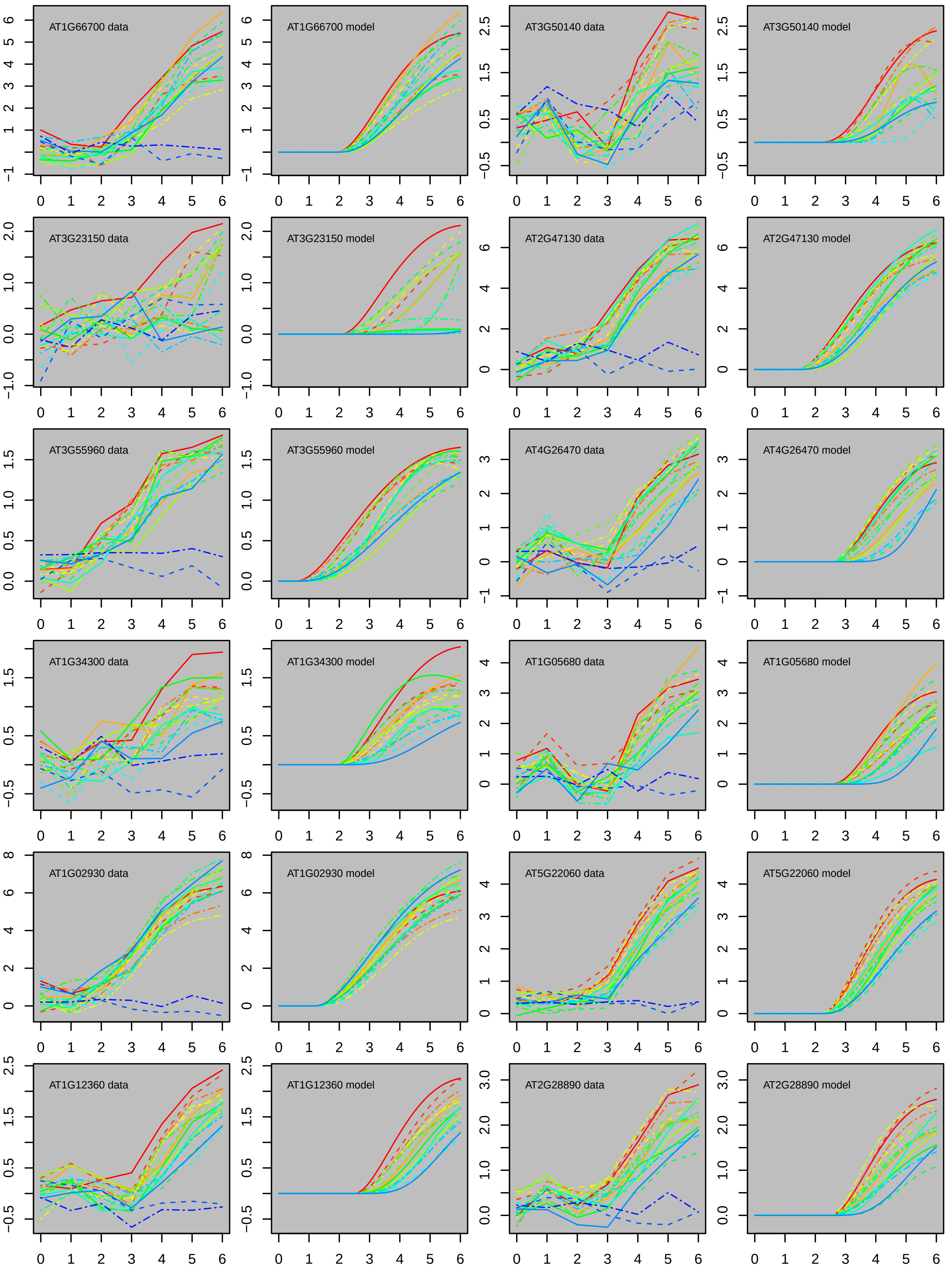
AT1G66700 data
AT1G66700 model
AT3G50140 data
AT3G50140 model
AT3G23150 data
AT3G23150 model
AT2G47130 data
AT2G47130 model
AT3G55960 data
AT3G55960 model
AT4G26470 data
AT4G26470 model
AT1G34300 data
AT1G34300 model
AT1G05680 data
AT1G05680 model
AT1G02930 data
AT1G02930 model
AT5G22060 data
AT5G22060 model
AT1G12360 data
AT1G12360 model
AT2G28890 data
AT2G28890 model

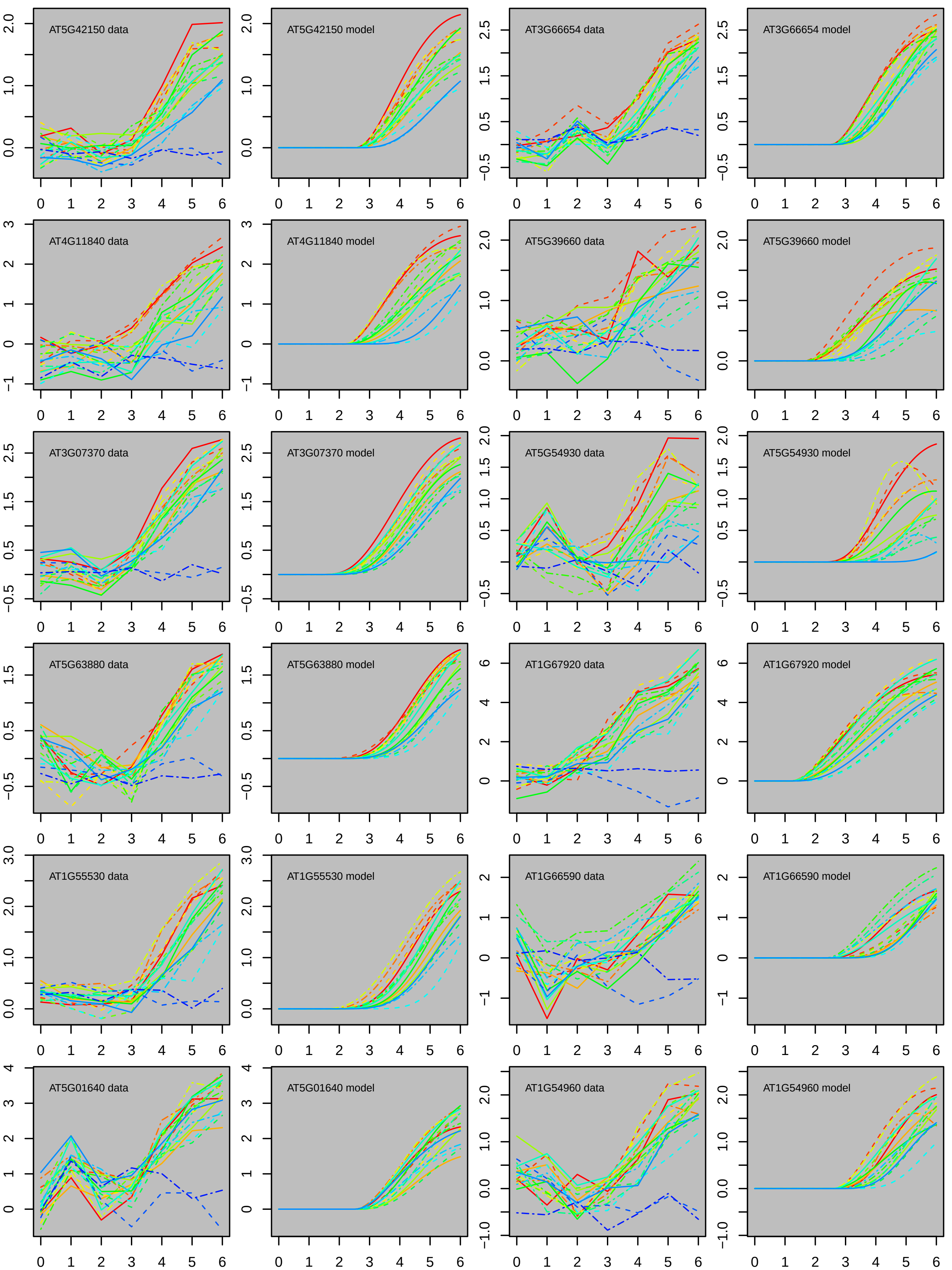
AT5G42150 data
AT5G42150 model
AT3G66654 data
AT3G66654 model
AT4G11840 data
AT4G11840 model
AT5G39660 data
AT5G39660 model
AT3G07370 data
AT3G07370 model
AT5G54930 data
AT5G54930 model
AT5G63880 data
AT5G63880 model
AT1G67920 data
AT1G67920 model
AT1G55530 data
AT1G55530 model
AT1G66590 data
AT1G66590 model
AT5G01640 data
AT5G01640 model
AT1G54960 data
AT1G54960 model

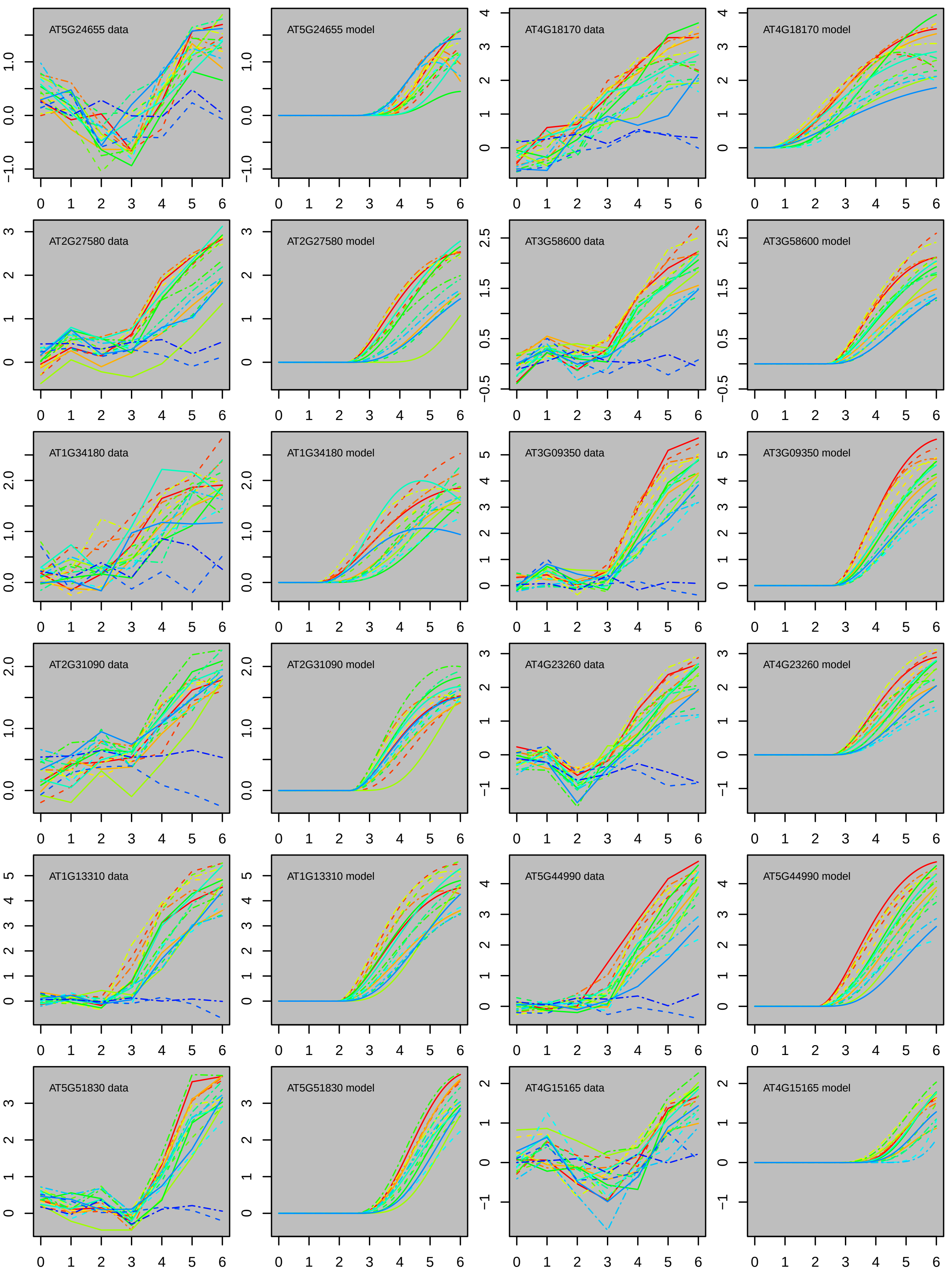
AT5G24655 data
AT5G24655 model
AT4G18170 data
AT4G18170 model
AT2G27580 data
AT2G27580 model
AT3G58600 data
AT3G58600 model
AT1G34180 data
AT1G34180 model
AT3G09350 data
AT3G09350 model
AT2G31090 data
AT2G31090 model
AT4G23260 data
AT4G23260 model
AT1G13310 data
AT1G13310 model
AT5G44990 data
AT5G44990 model
AT5G51830 data
AT5G51830 model
AT4G15165 data
AT4G15165 model

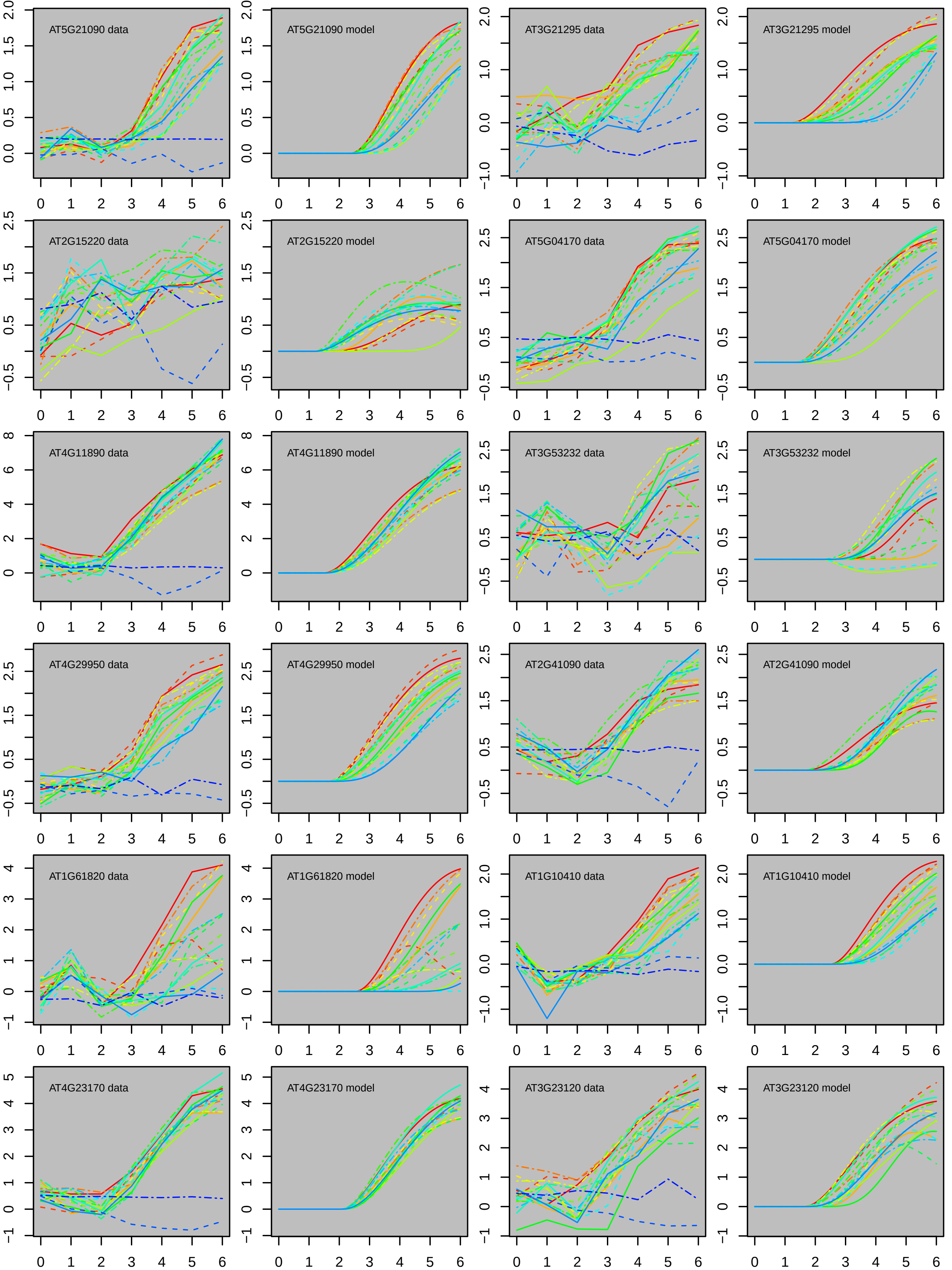

AT5G21090 data
AT5G21090 model
AT3G21295 data
AT3G21295 model
AT2G15220 data
AT2G15220 model
AT5G04170 data
AT5G04170 model
AT4G11890 data
AT4G11890 model
AT3G53232 data
AT3G53232 model
AT4G29950 data
AT4G29950 model
AT2G41090 data
AT2G41090 model
AT1G61820 data
AT1G61820 model
AT1G10410 data
AT1G10410 model
AT4G23170 data
AT4G23170 model
AT3G23120 data
AT3G23120 model

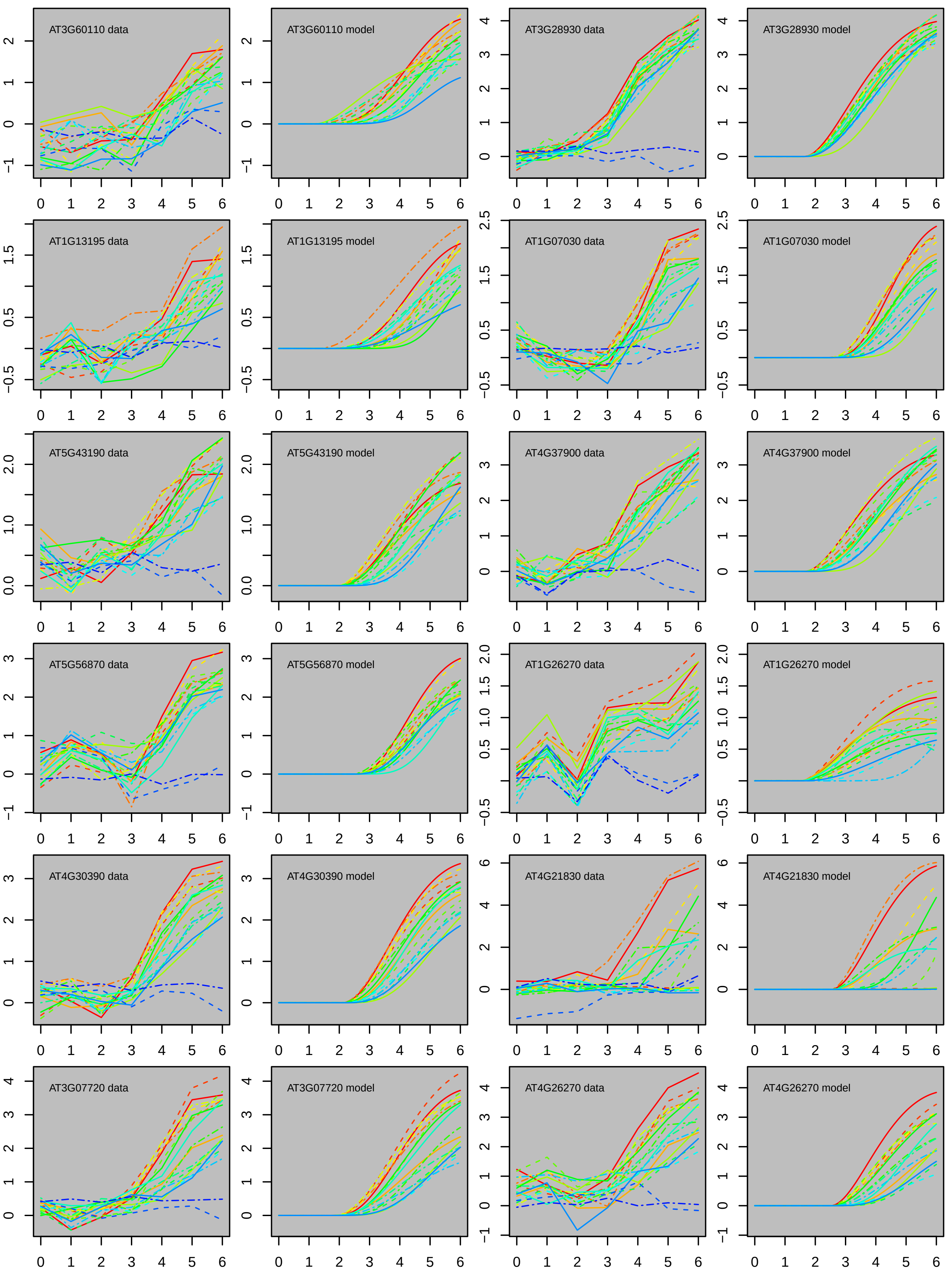
AT3G60110 data
AT3G60110 model
AT3G28930 data
AT3G28930 model
AT1G13195 data
AT1G13195 model
AT1G07030 data
AT1G07030 model
AT5G43190 data
AT5G43190 model
AT4G37900 data
AT4G37900 model
AT5G56870 data
AT5G56870 model
AT1G26270 data
AT1G26270 model
AT4G30390 data
AT4G30390 model
AT4G21830 data
AT4G21830 model
AT3G07720 data
AT3G07720 model
AT4G26270 data
AT4G26270 model

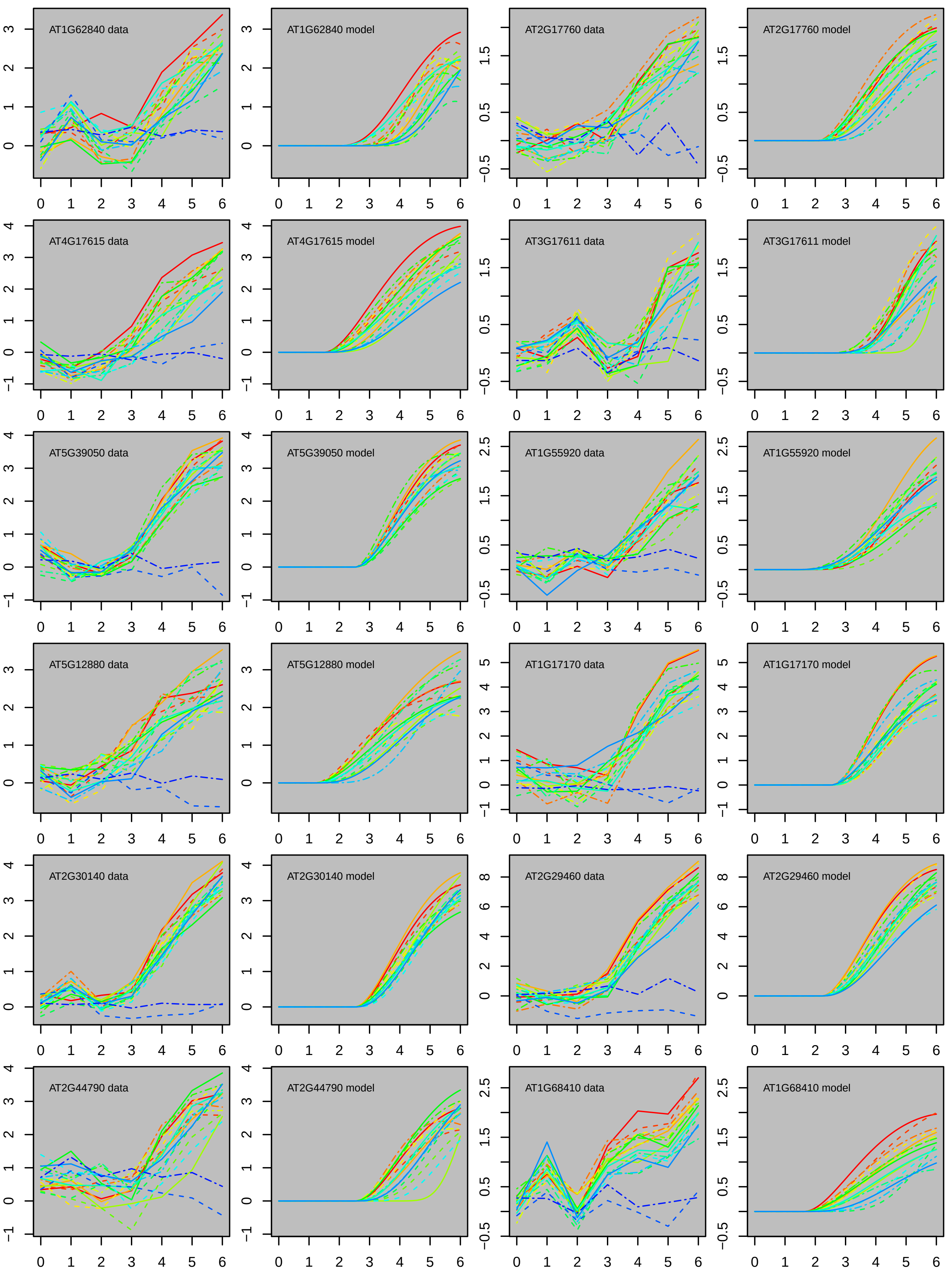
AT1G62840 data
AT1G62840 model
AT2G17760 data
AT2G17760 model
AT4G17615 data
AT4G17615 model
AT3G17611 data
AT3G17611 model
AT5G39050 data
AT5G39050 model
AT1G55920 data
AT1G55920 model
AT5G12880 data
AT5G12880 model
AT1G17170 data
AT1G17170 model
AT2G30140 data
AT2G30140 model
AT2G29460 data
AT2G29460 model
AT2G44790 data
AT2G44790 model
AT1G68410 data
AT1G68410 model

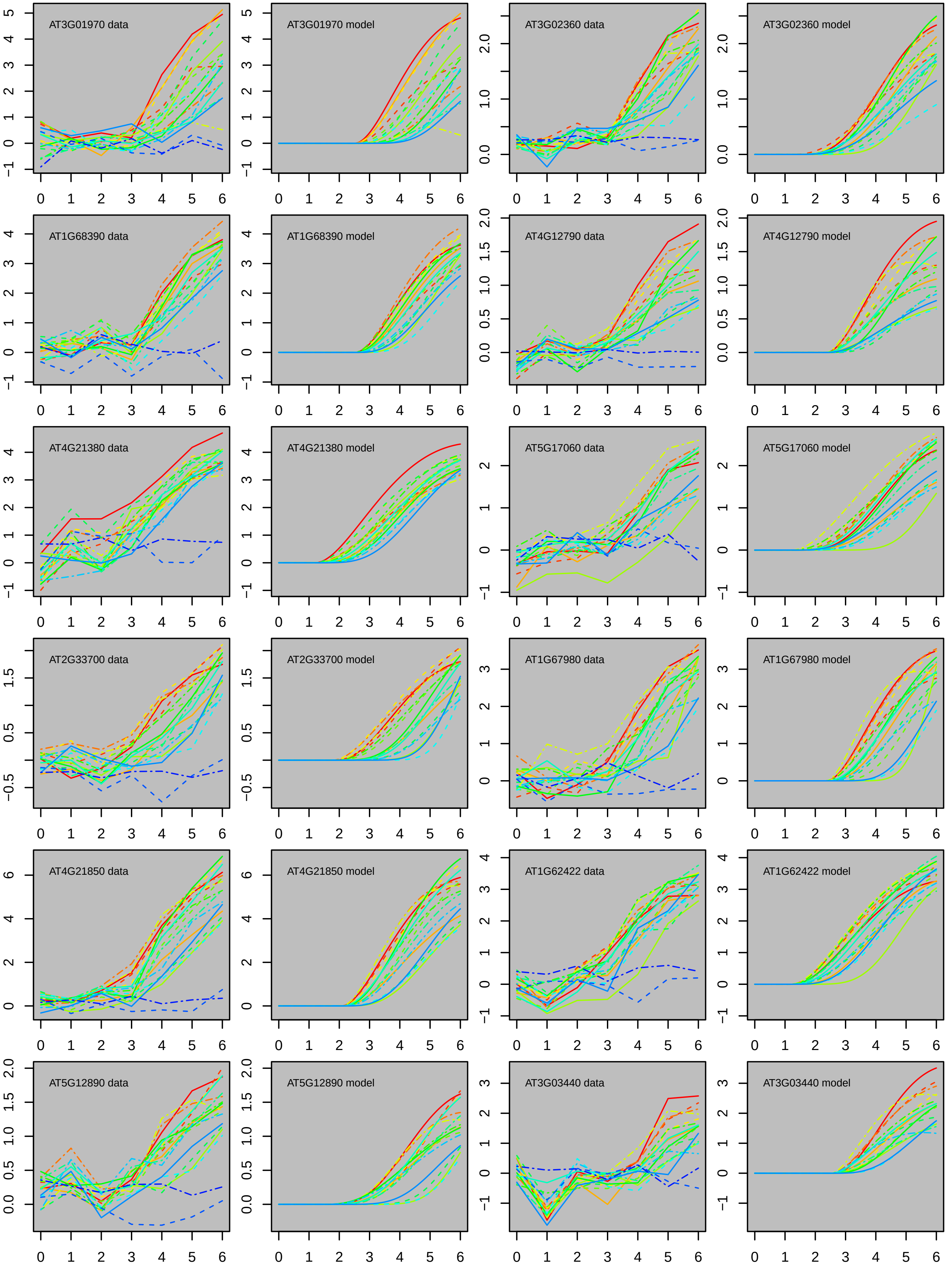

AT3G01970 data
AT3G01970 model
AT3G02360 data
AT3G02360 model
AT1G68390 data
AT1G68390 model
AT4G12790 data
AT4G12790 model
AT4G21380 data
AT4G21380 model
AT5G17060 data
AT5G17060 model
AT2G33700 data
AT2G33700 model
AT1G67980 data
AT1G67980 model
AT4G21850 data
AT4G21850 model
AT1G62422 data
AT1G62422 model
AT5G12890 data
AT5G12890 model
AT3G03440 data
AT3G03440 model

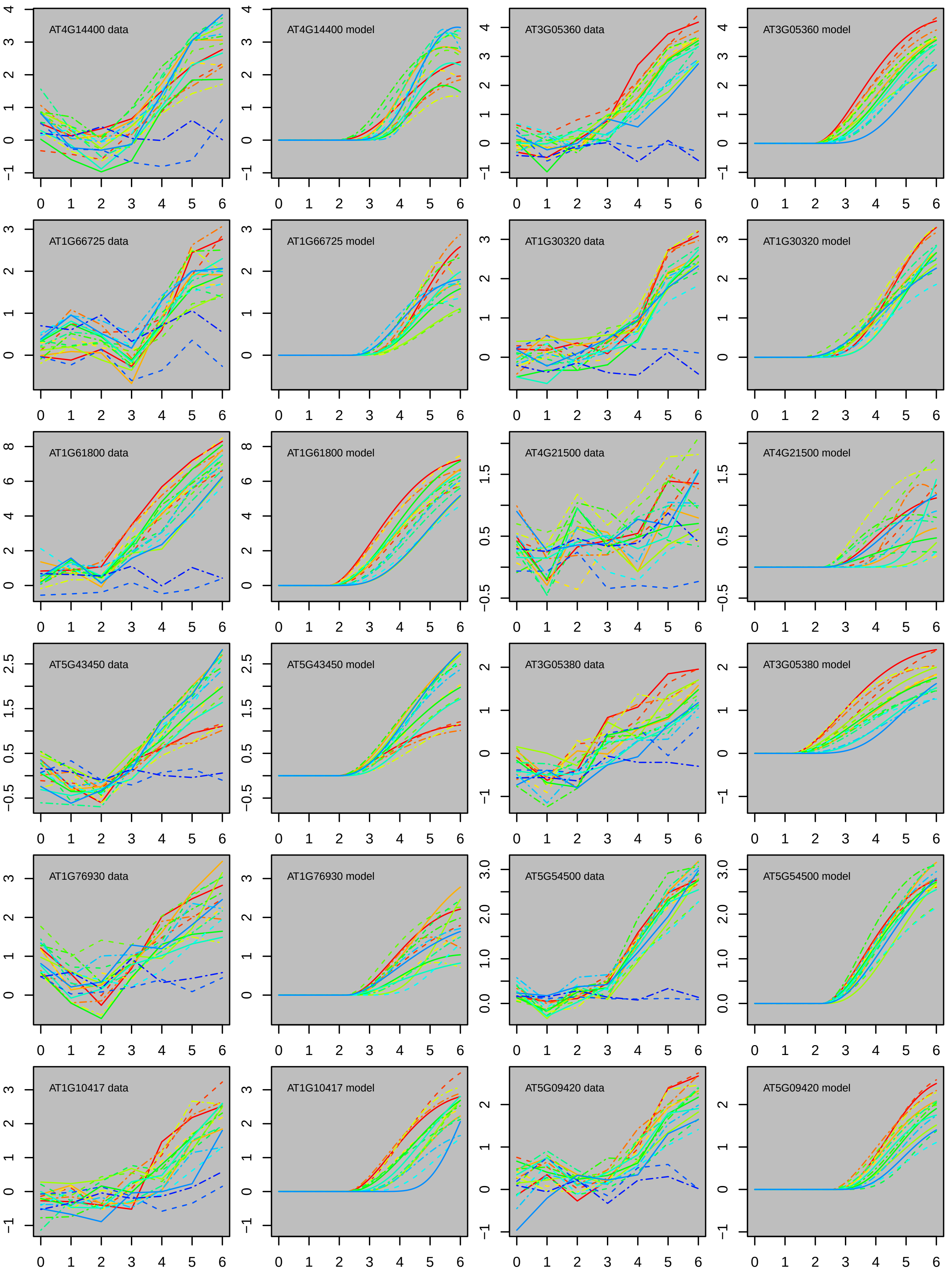

AT4G14400 data
AT4G14400 model
AT3G05360 data
AT3G05360 model
AT1G66725 data
AT1G66725 model
AT1G30320 data
AT1G30320 model
AT1G61800 data
AT1G61800 model
AT4G21500 data
AT4G21500 model
AT5G43450 data
AT5G43450 model
AT3G05380 data
AT3G05380 model
AT1G76930 data
AT1G76930 model
AT5G54500 data
AT5G54500 model
AT1G10417 data
AT1G10417 model
AT5G09420 data
AT5G09420 model

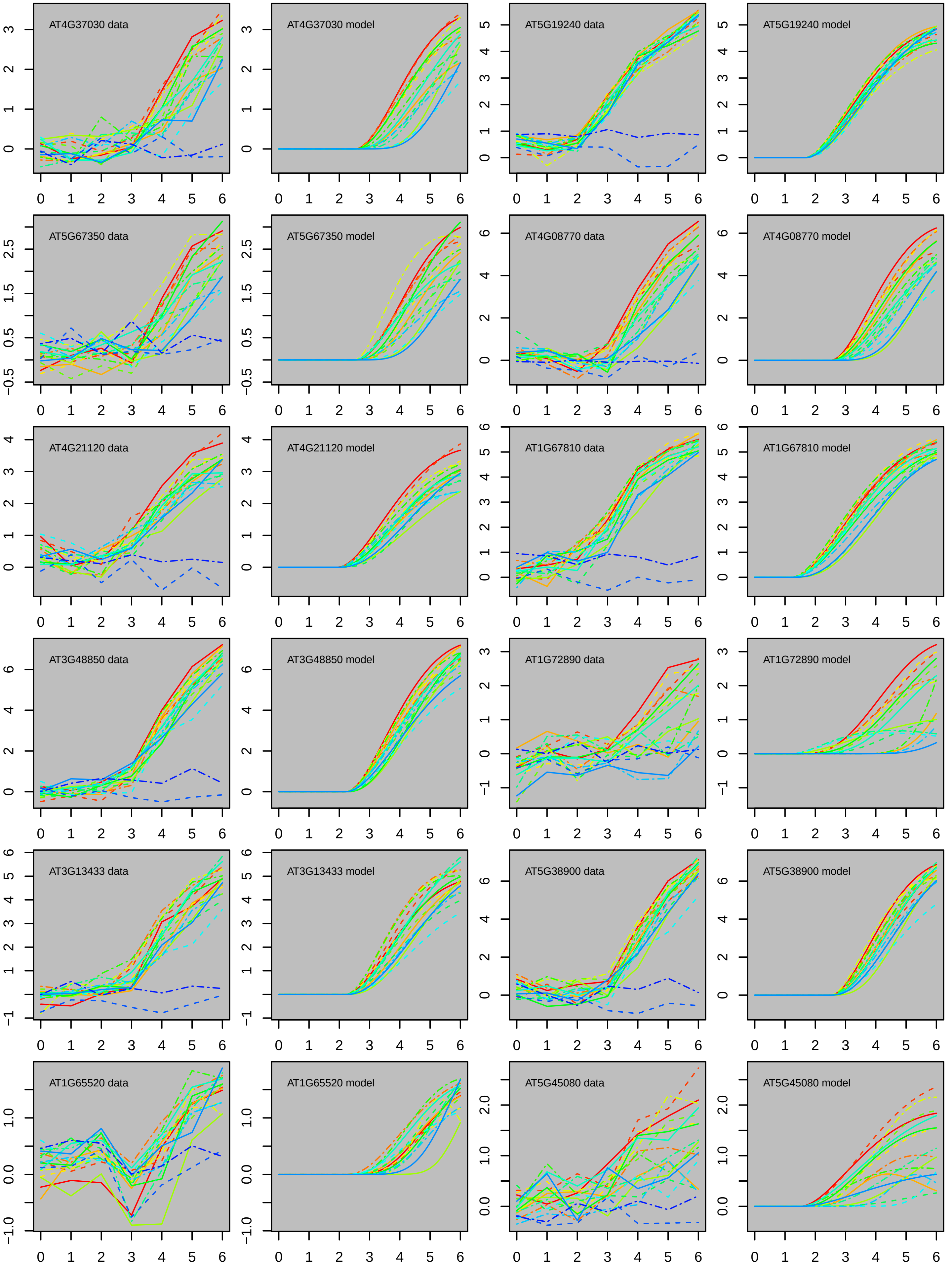

AT4G37030 data
AT4G37030 model
AT5G19240 data
AT5G19240 model
AT5G67350 data
AT5G67350 model
AT4G08770 data
AT4G08770 model
AT4G21120 data
AT4G21120 model
AT1G67810 data
AT1G67810 model
AT3G48850 data
AT3G48850 model
AT1G72890 data
AT1G72890 model
AT3G13433 data
AT3G13433 model
AT5G38900 data
AT5G38900 model
AT1G65520 data
AT1G65520 model
AT5G45080 data
AT5G45080 model

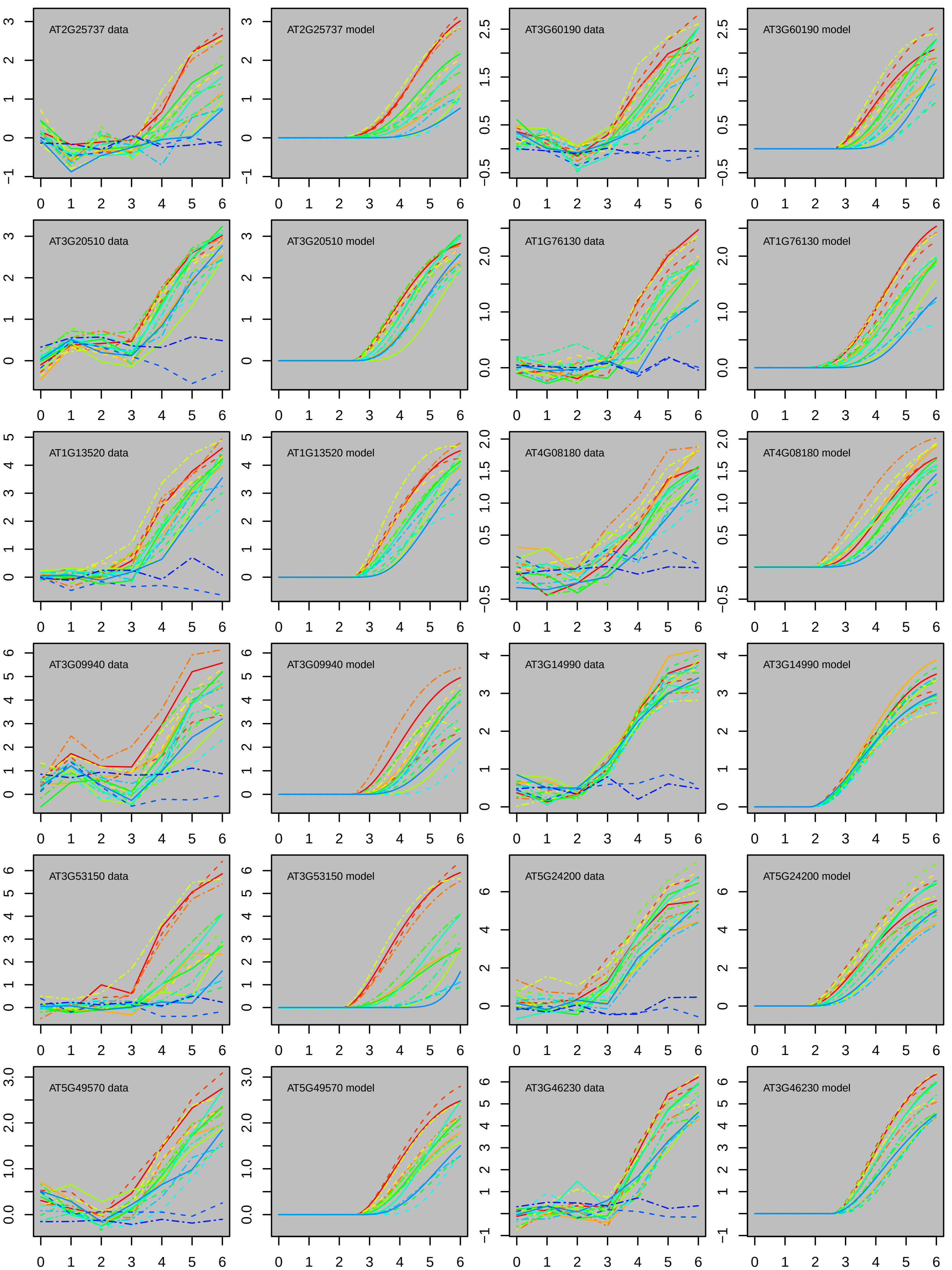

AT2G25737 data
AT2G25737 model
AT3G60190 data
AT3G60190 model
AT3G20510 data
AT3G20510 model
AT1G76130 data
AT1G76130 model
AT1G13520 data
AT1G13520 model
AT4G08180 data
AT4G08180 model
AT3G09940 data
AT3G09940 model
AT3G14990 data
AT3G14990 model
AT3G53150 data
AT3G53150 model
AT5G24200 data
AT5G24200 model
AT5G49570 data
AT5G49570 model
AT3G46230 data
AT3G46230 model

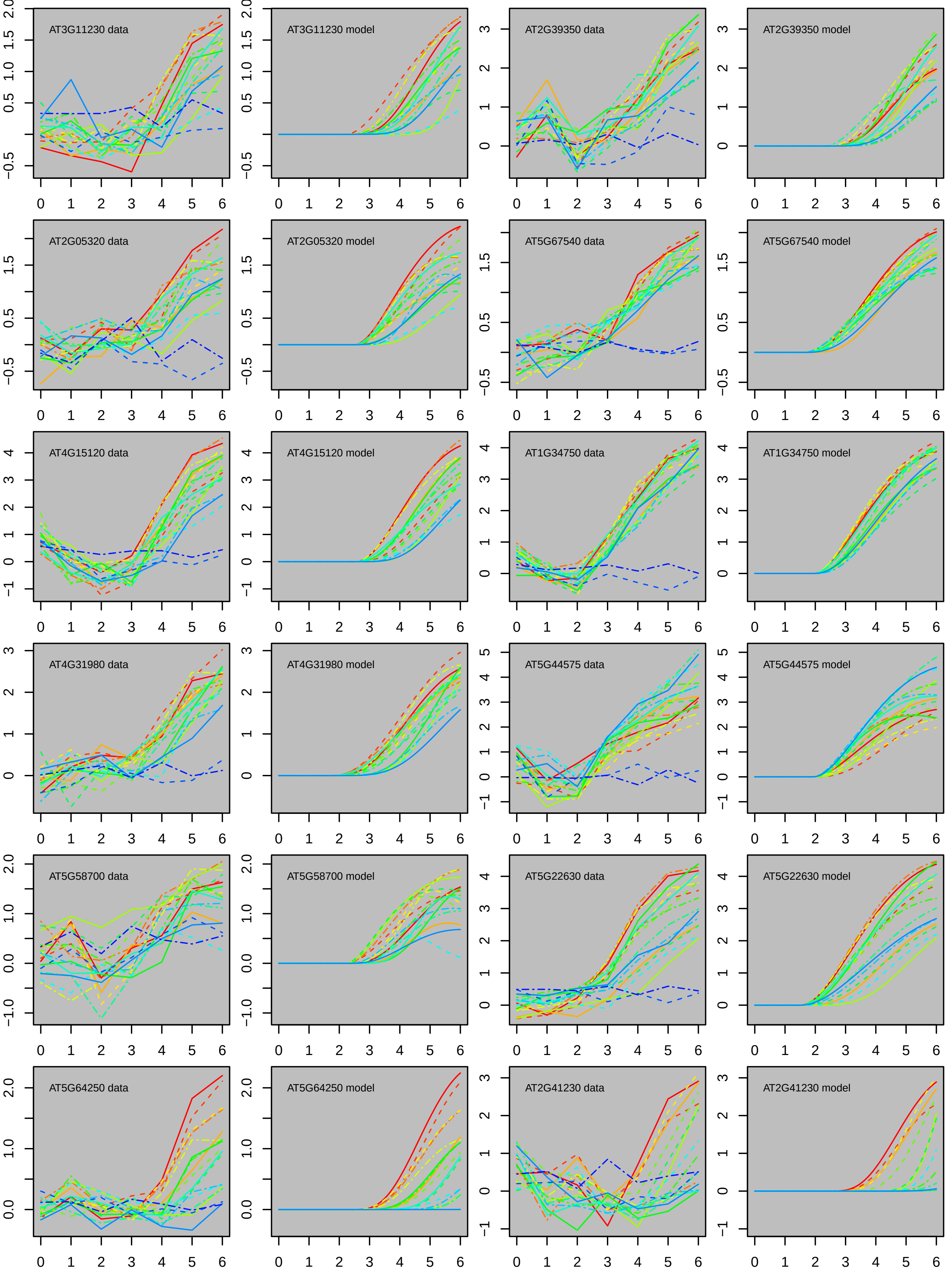
AT3G11230 data
AT3G11230 model
AT2G39350 data
AT2G39350 model
AT2G05320 data
AT2G05320 model
AT5G67540 data
AT5G67540 model
AT4G15120 data
AT4G15120 model
AT1G34750 data
AT1G34750 model
AT4G31980 data
AT4G31980 model
AT5G44575 data
AT5G44575 model
AT5G58700 data
AT5G58700 model
AT5G22630 data
AT5G22630 model
AT5G64250 data
AT5G64250 model
AT2G41230 data
AT2G41230 model

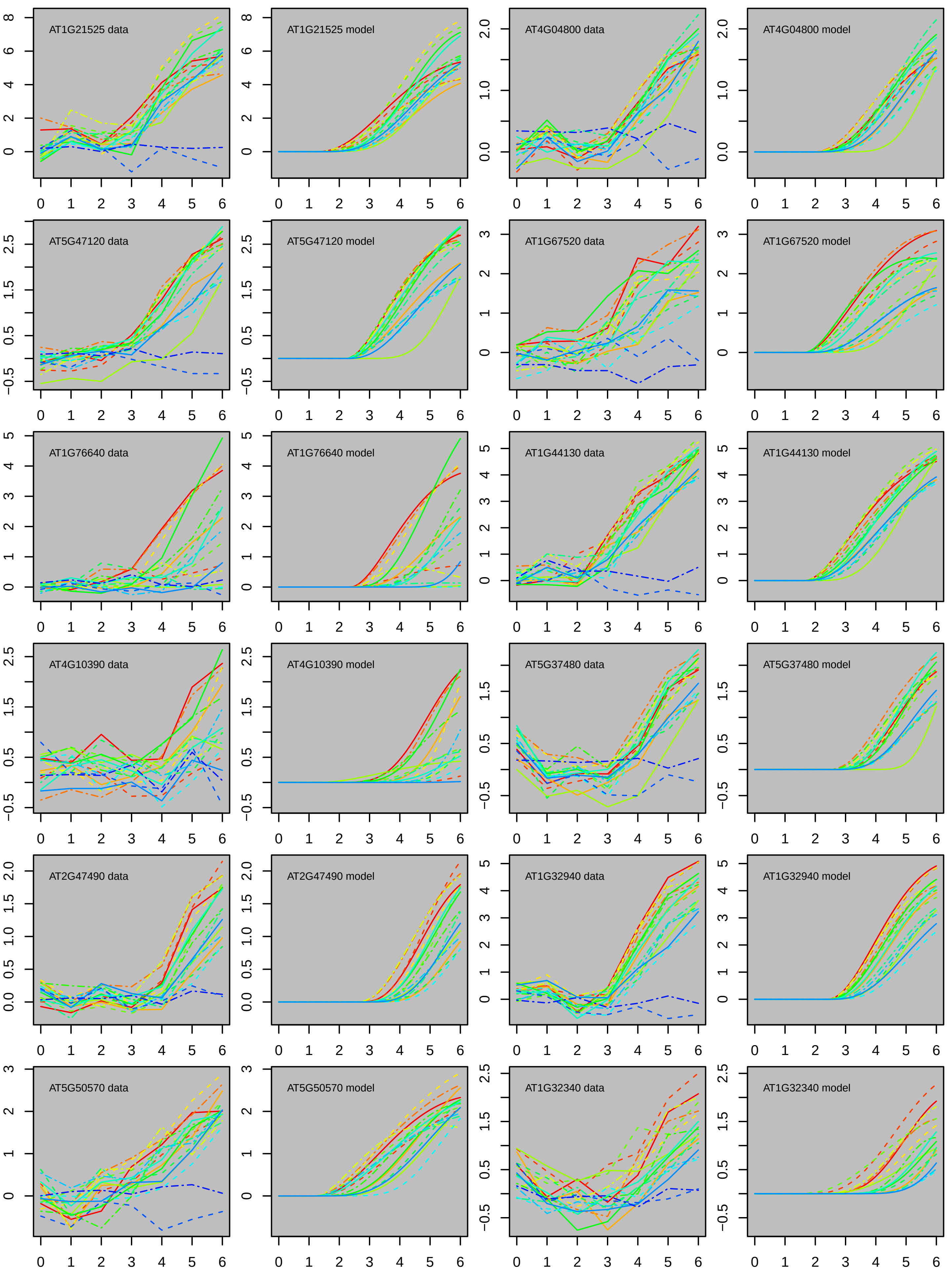
AT1G21525 data
AT1G21525 model
AT4G04800 data
AT4G04800 model
AT5G47120 data
AT5G47120 model
AT1G67520 data
AT1G67520 model
AT1G76640 data
AT1G76640 model
AT1G44130 data
AT1G44130 model
AT4G10390 data
AT4G10390 model
AT5G37480 data
AT5G37480 model
AT2G47490 data
AT2G47490 model
AT1G32940 data
AT1G32940 model
AT5G50570 data
AT5G50570 model
AT1G32340 data
AT1G32340 model

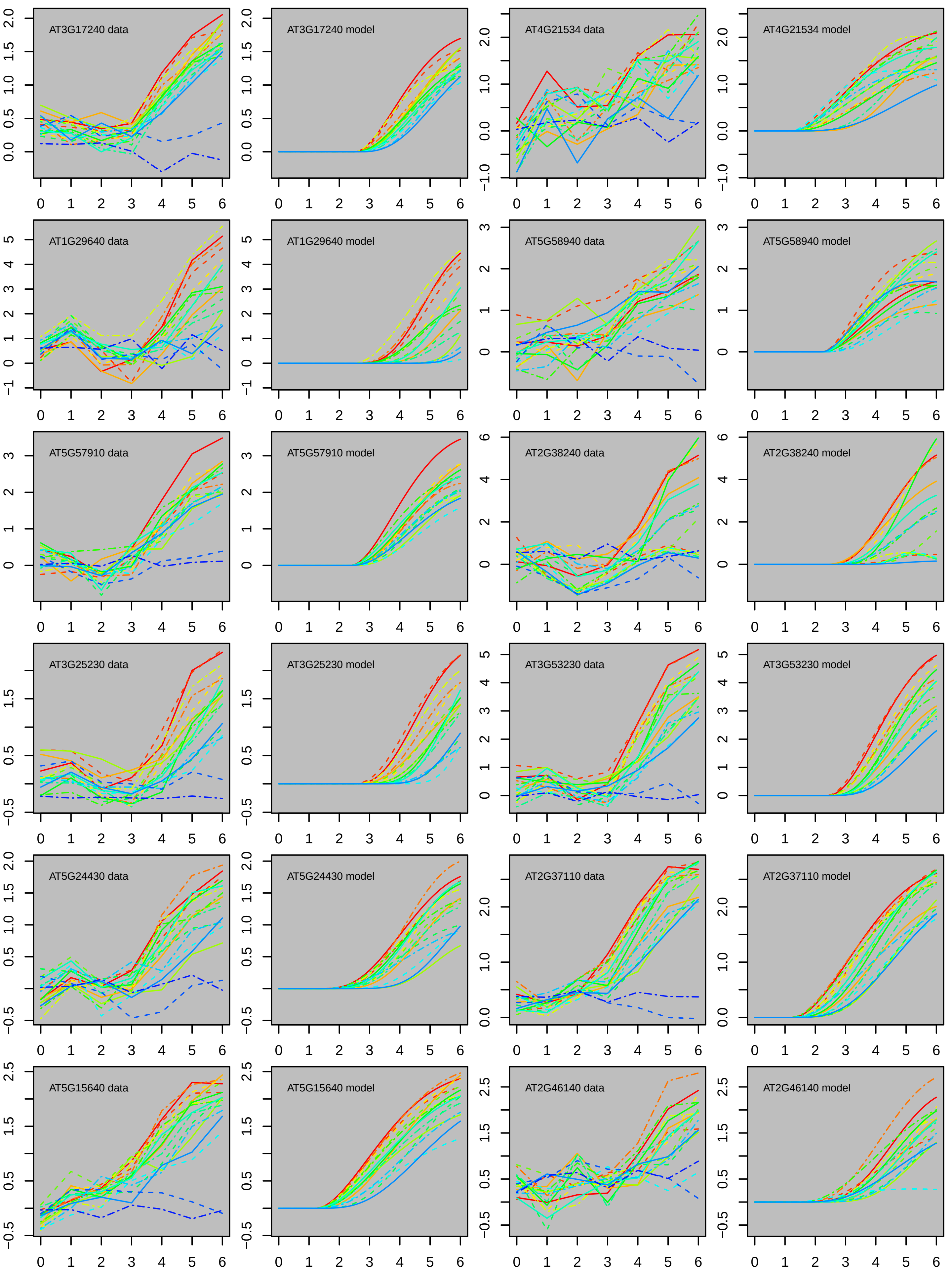

AT3G17240 data
AT3G17240 model
AT4G21534 data
AT4G21534 model
AT1G29640 data
AT1G29640 model
AT5G58940 data
AT5G58940 model
AT5G57910 data
AT5G57910 model
AT2G38240 data
AT2G38240 model
AT3G25230 data
AT3G25230 model
AT3G53230 data
AT3G53230 model
AT5G24430 data
AT5G24430 model
AT2G37110 data
AT2G37110 model
AT5G15640 data
AT5G15640 model
AT2G46140 data
AT2G46140 model

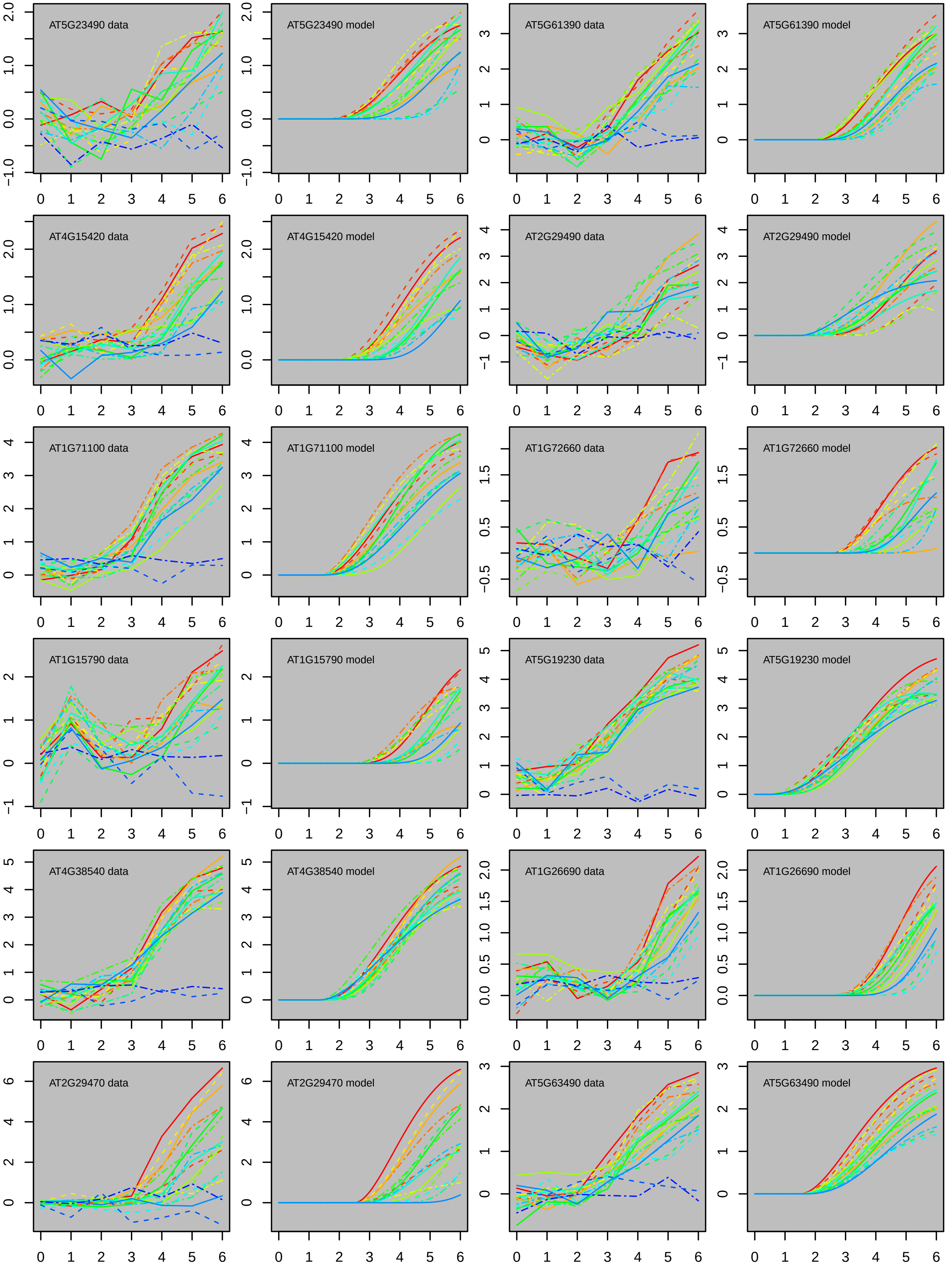
AT5G23490 data
AT5G23490 model
AT5G61390 data
AT5G61390 model
AT4G15420 data
AT4G15420 model
AT2G29490 data
AT2G29490 model
AT1G71100 data
AT1G71100 model
AT1G72660 data
AT1G72660 model
AT1G15790 data
AT1G15790 model
AT5G19230 data
AT5G19230 model
AT4G38540 data
AT4G38540 model
AT1G26690 data
AT1G26690 model
AT2G29470 data
AT2G29470 model
AT5G63490 data
AT5G63490 model

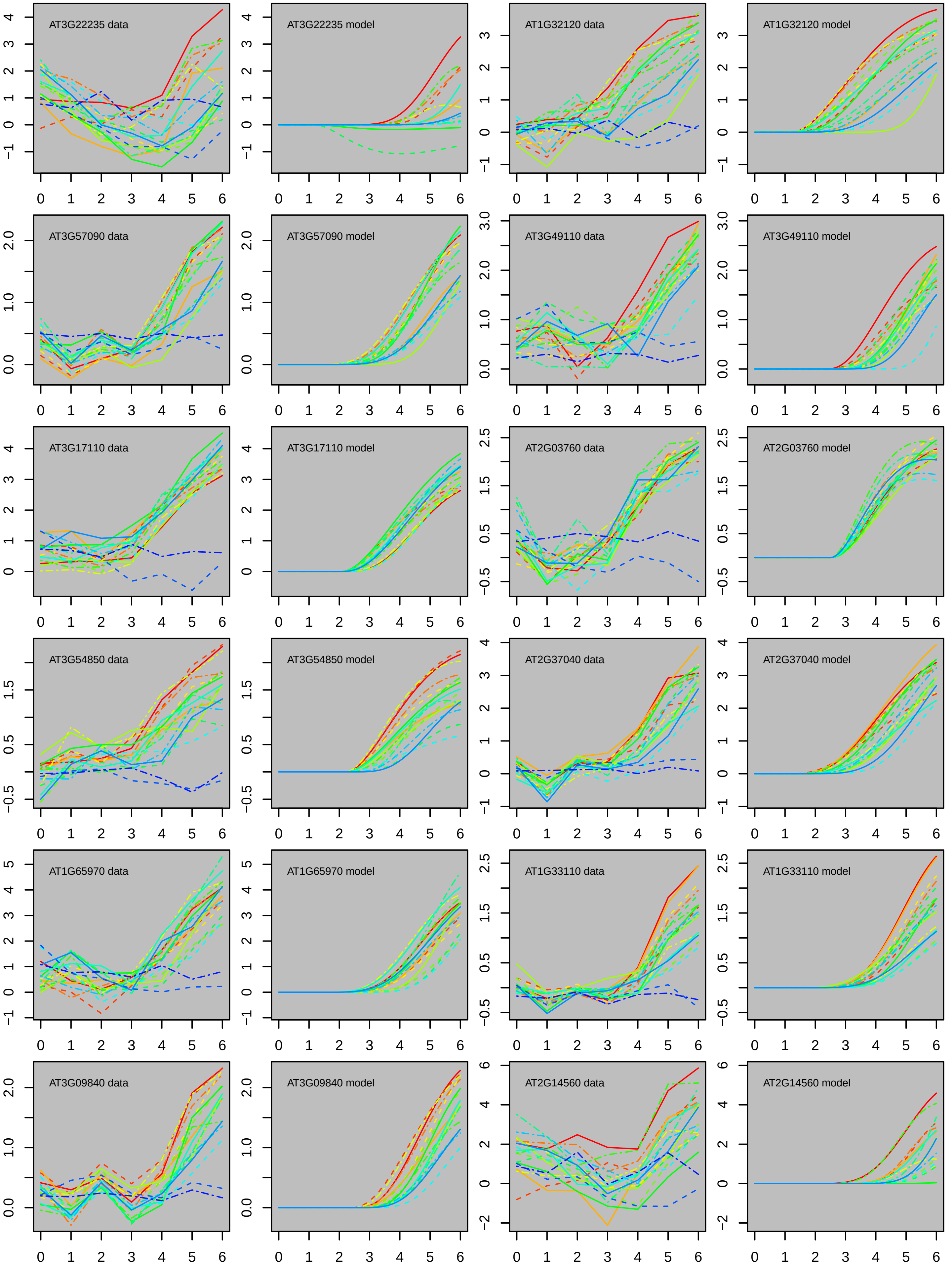
AT3G22235 data
AT3G22235 model
AT1G32120 data
AT1G32120 model
AT3G57090 data
AT3G57090 model
AT3G49110 data
AT3G49110 model
AT3G17110 data
AT3G17110 model
AT2G03760 data
AT2G03760 model
AT3G54850 data
AT3G54850 model
AT2G37040 data
AT2G37040 model
AT1G65970 data
AT1G65970 model
AT1G33110 data
AT1G33110 model
AT3G09840 data
AT3G09840 model
AT2G14560 data
AT2G14560 model

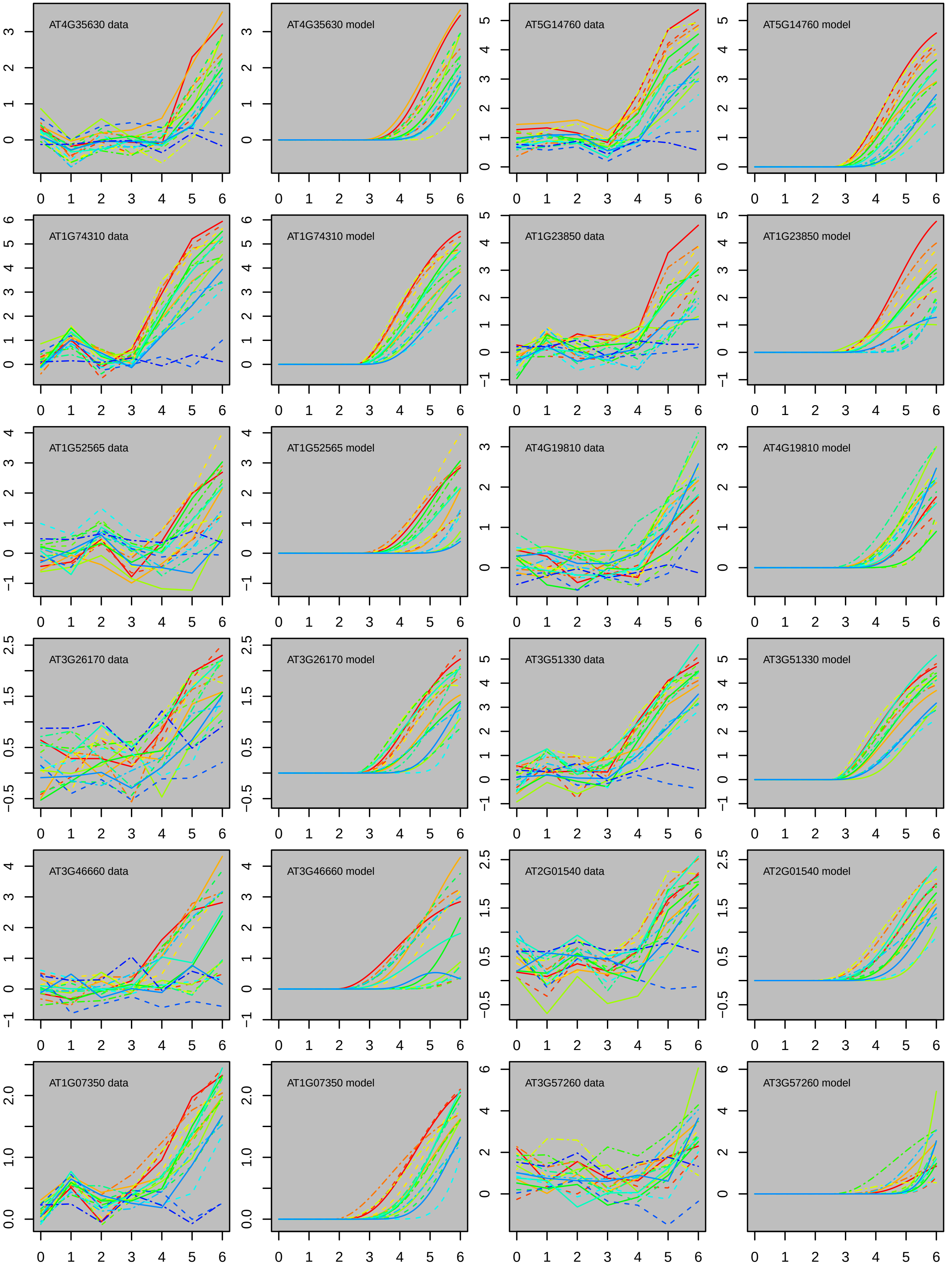

AT4G35630 data
AT4G35630 model
AT5G14760 data
AT5G14760 model
AT1G74310 data
AT1G74310 model
AT1G23850 data
AT1G23850 model
AT1G52565 data
AT1G52565 model
AT4G19810 data
AT4G19810 model
AT3G26170 data
AT3G26170 model
AT3G51330 data
AT3G51330 model
AT3G46660 data
AT3G46660 model
AT2G01540 data
AT2G01540 model
AT1G07350 data
AT1G07350 model
AT3G57260 data
AT3G57260 model

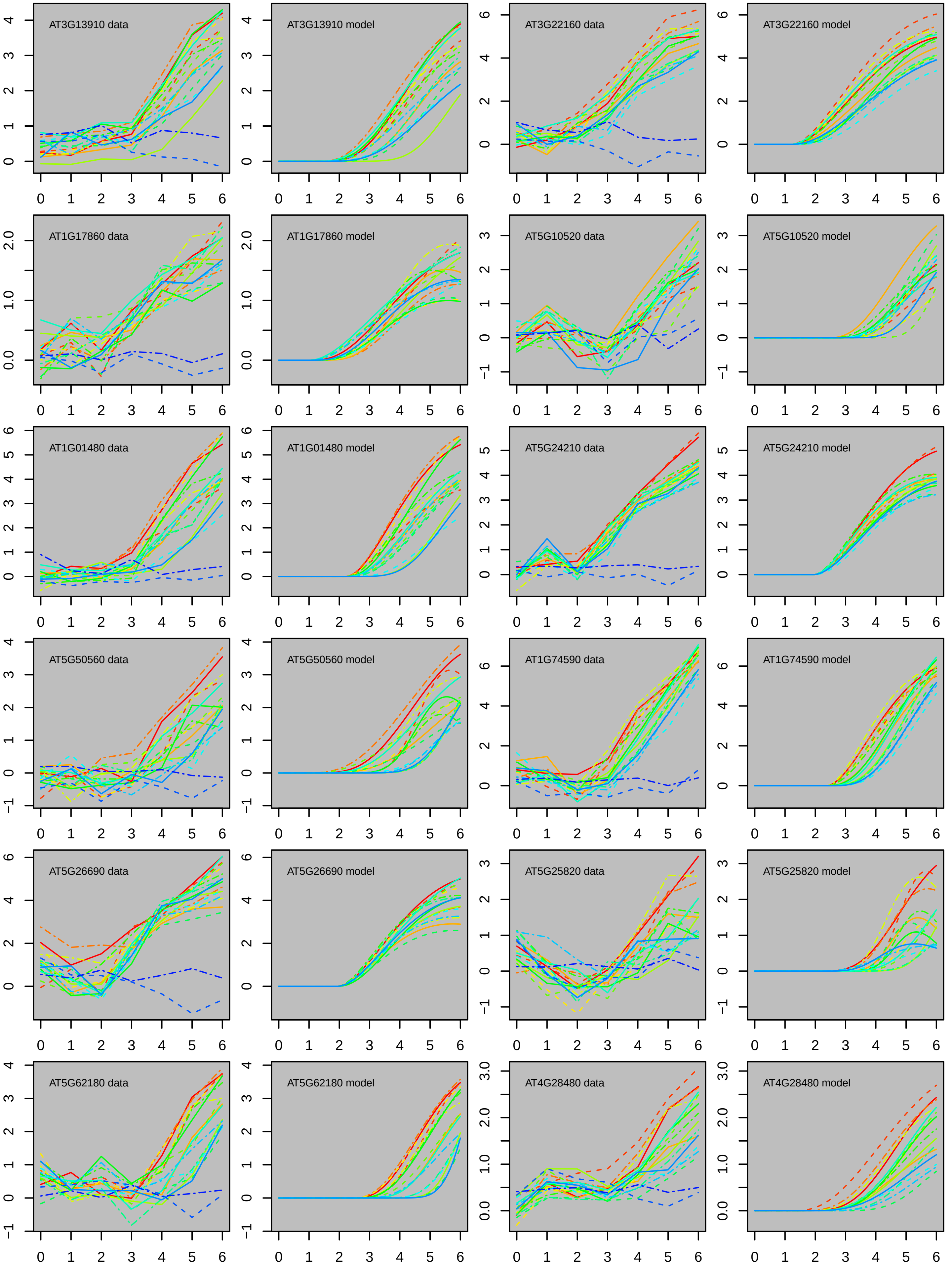

AT3G13910 data
AT3G13910 model
AT3G22160 data
AT3G22160 model
AT1G17860 data
AT1G17860 model
AT5G10520 data
AT5G10520 model
AT1G01480 data
AT1G01480 model
AT5G24210 data
AT5G24210 model
AT5G50560 data
AT5G50560 model
AT1G74590 data
AT1G74590 model
AT5G26690 data
AT5G26690 model
AT5G25820 data
AT5G25820 model
AT5G62180 data
AT5G62180 model
AT4G28480 data
AT4G28480 model

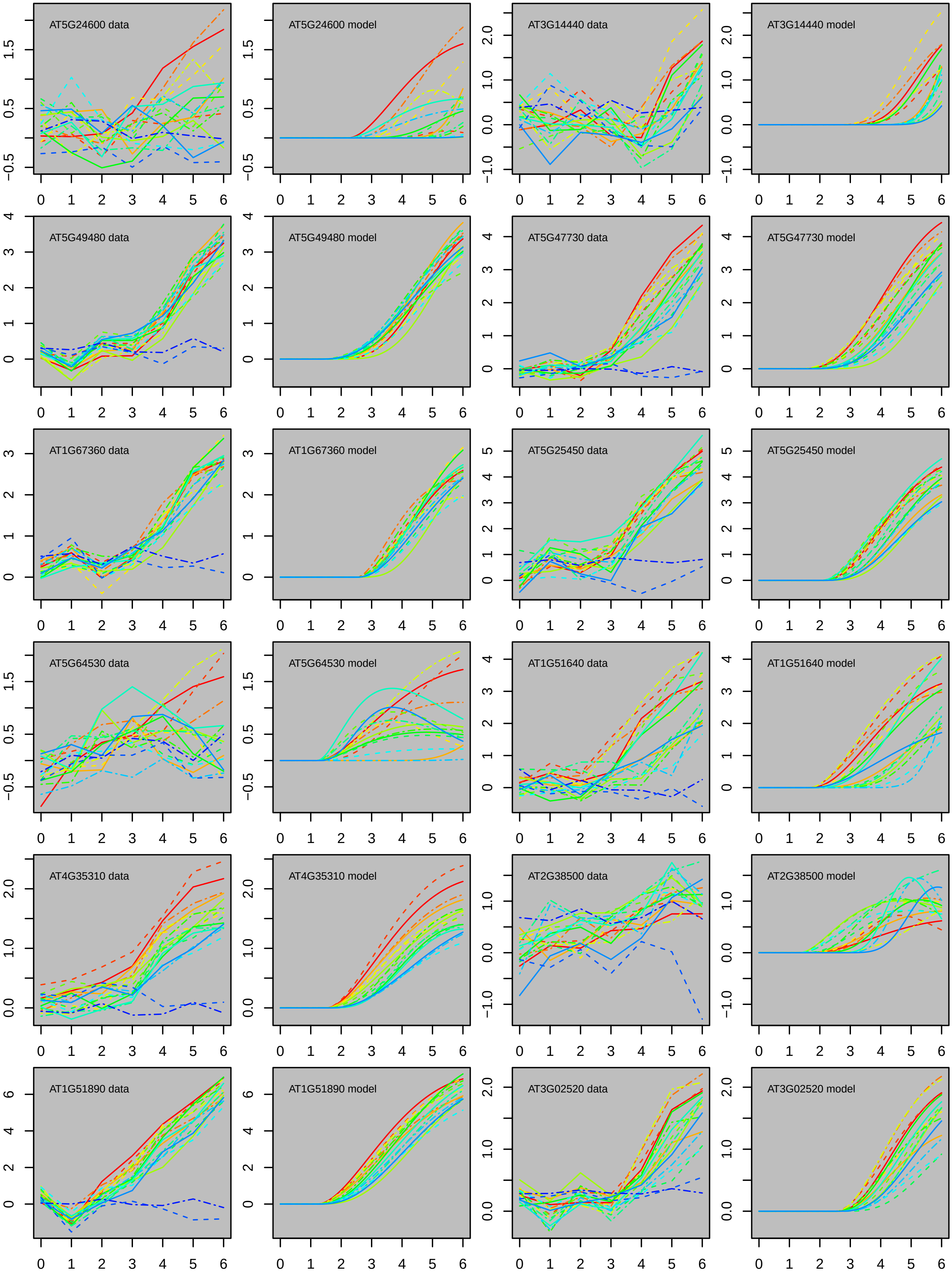

AT5G24600 data
AT5G24600 model
AT3G14440 data
AT3G14440 model
AT5G49480 data
AT5G49480 model
AT5G47730 data
AT5G47730 model
AT1G67360 data
AT1G67360 model
AT5G25450 data
AT5G25450 model
AT5G64530 data
AT5G64530 model
AT1G51640 data
AT1G51640 model
AT4G35310 data
AT4G35310 model
AT2G38500 data
AT2G38500 model
AT1G51890 data
AT1G51890 model
AT3G02520 data
AT3G02520 model

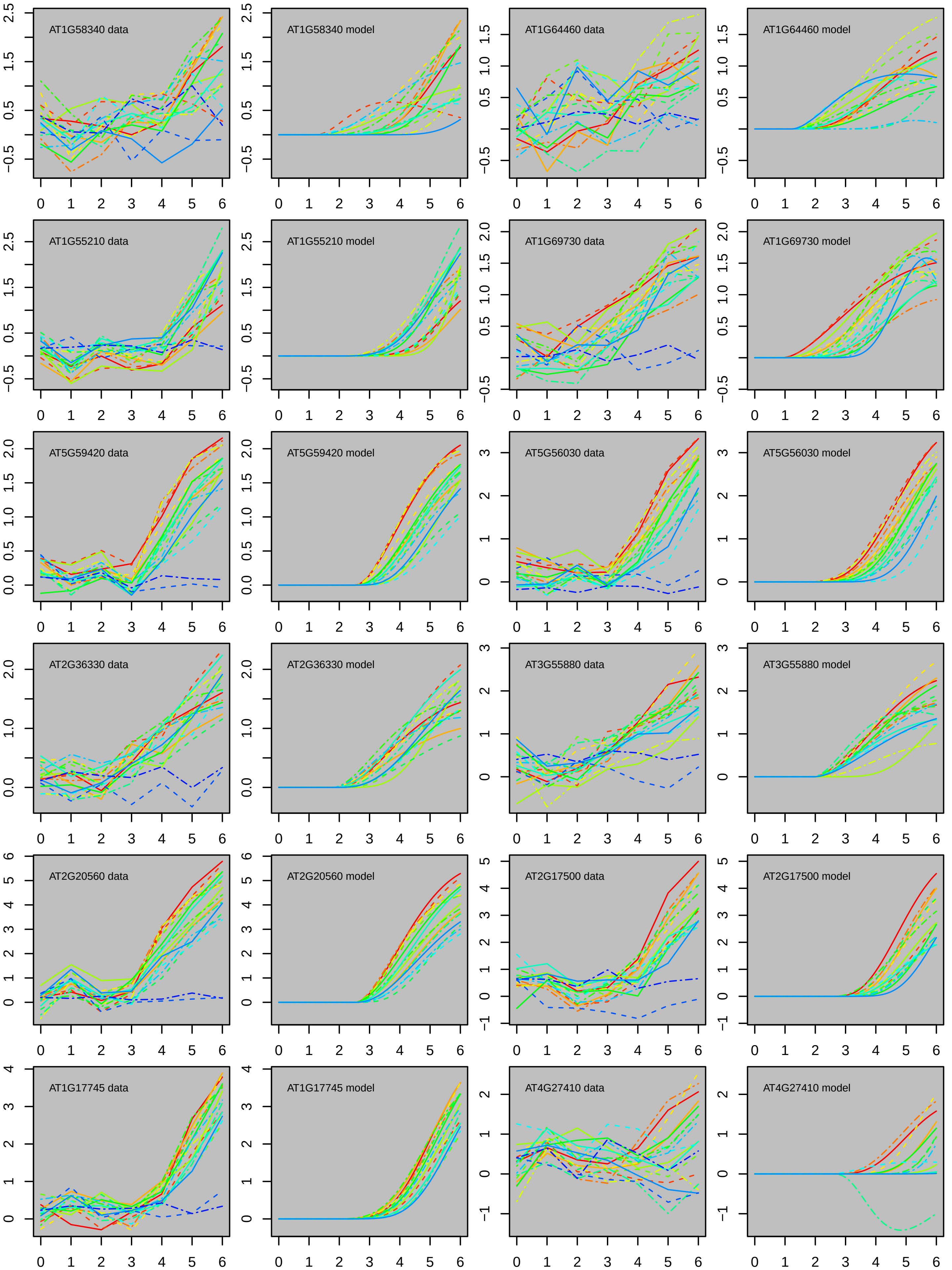
AT1G58340 data
AT1G58340 model
AT1G64460 data
AT1G64460 model
AT1G55210 data
AT1G55210 model
AT1G69730 data
AT1G69730 model
AT5G59420 data
AT5G59420 model
AT5G56030 data
AT5G56030 model
AT2G36330 data
AT2G36330 model
AT3G55880 data
AT3G55880 model
AT2G20560 data
AT2G20560 model
AT2G17500 data
AT2G17500 model
AT1G17745 data
AT1G17745 model
AT4G27410 data
AT4G27410 model

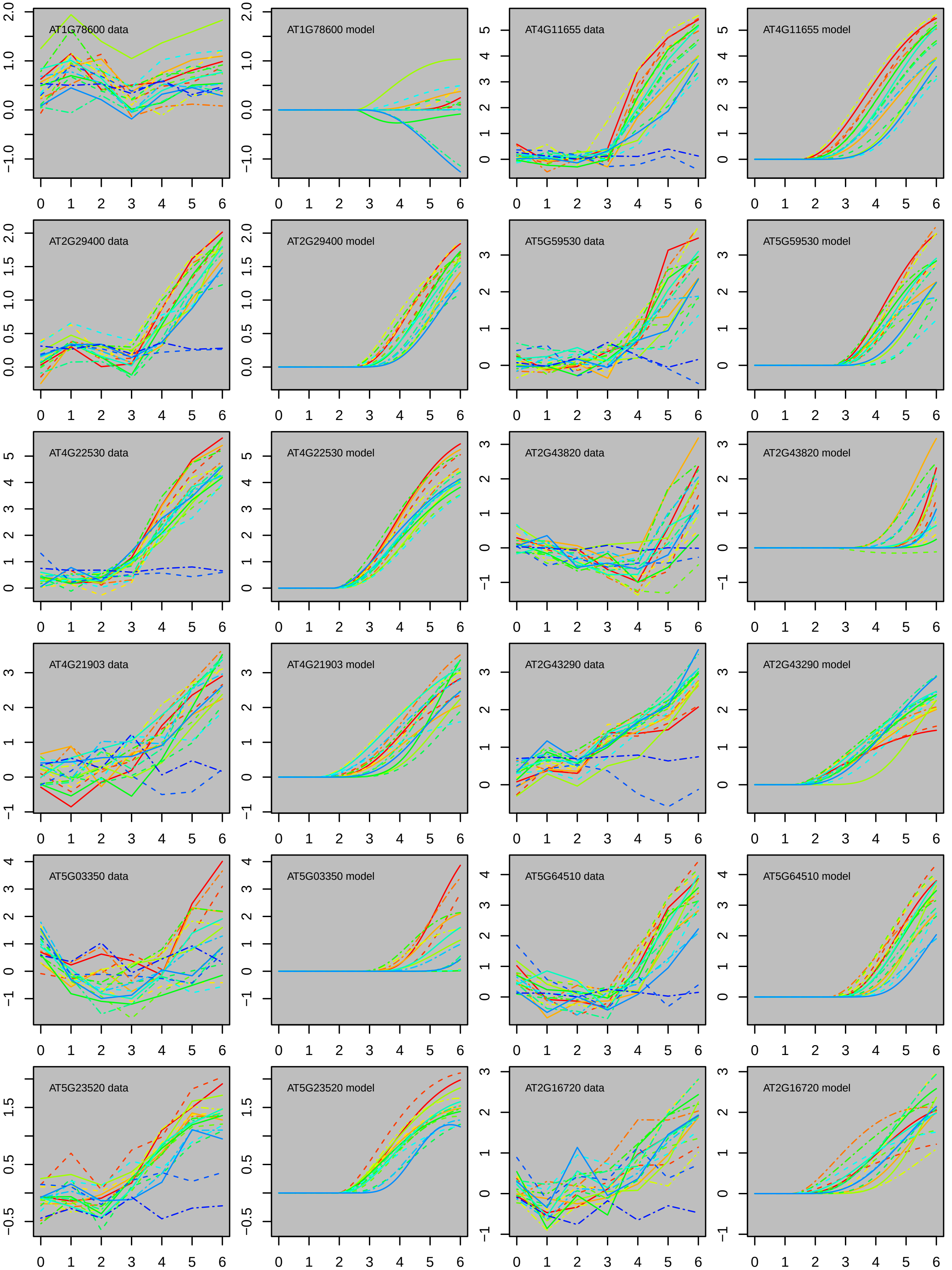

AT1G78600 data
AT1G78600 model
AT4G11655 data
AT4G11655 model
AT2G29400 data
AT2G29400 model
AT5G59530 data
AT5G59530 model
AT4G22530 data
AT4G22530 model
AT2G43820 data
AT2G43820 model
AT4G21903 data
AT4G21903 model
AT2G43290 data
AT2G43290 model
AT5G03350 data
AT5G03350 model
AT5G64510 data
AT5G64510 model
AT5G23520 data
AT5G23520 model
AT2G16720 data
AT2G16720 model

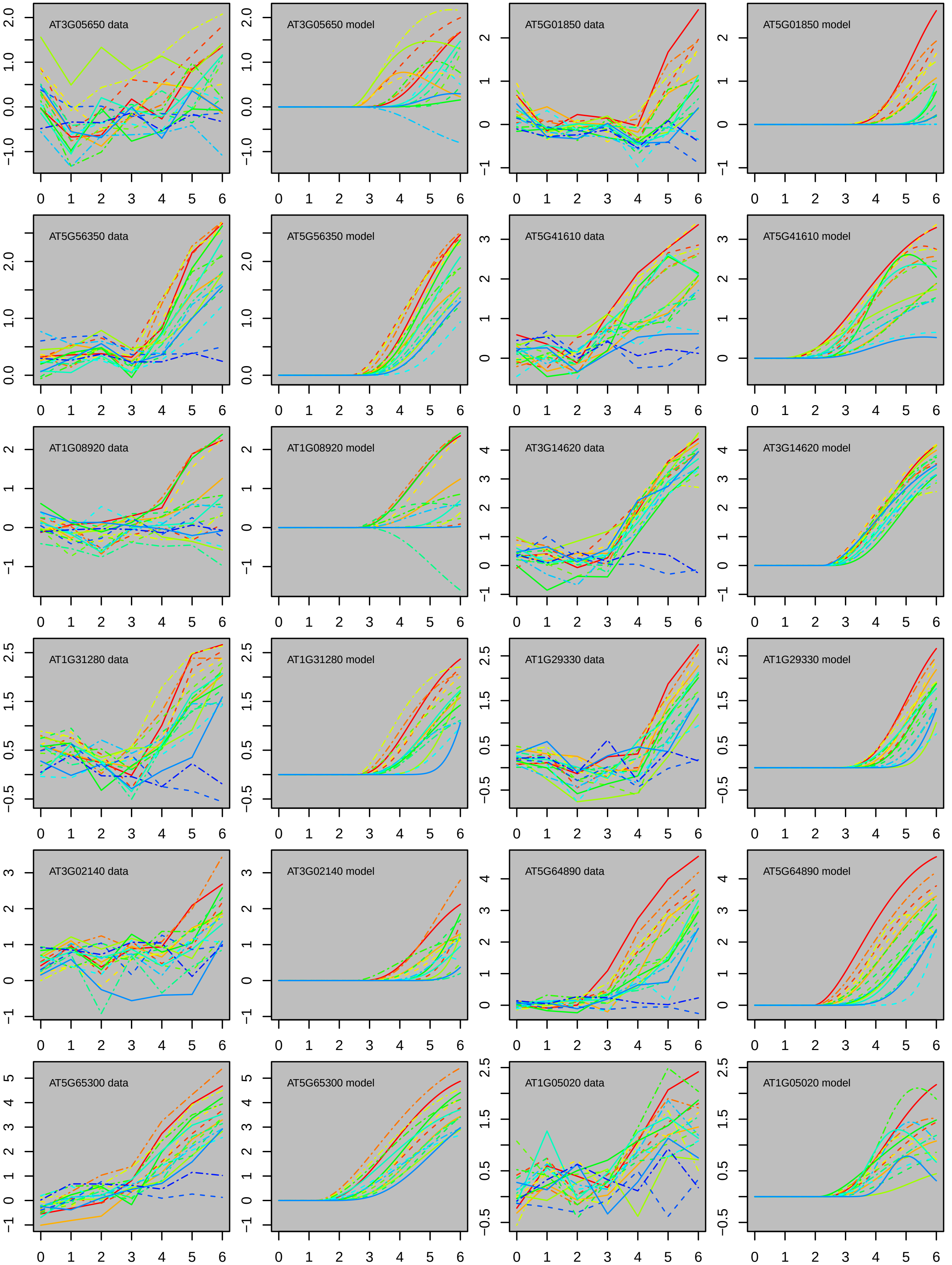

AT3G05650 data
AT3G05650 model
AT5G01850 data
AT5G01850 model
AT5G56350 data
AT5G56350 model
AT5G41610 data
AT5G41610 model
AT1G08920 data
AT1G08920 model
AT3G14620 data
AT3G14620 model
AT1G31280 data
AT1G31280 model
AT1G29330 data
AT1G29330 model
AT3G02140 data
AT3G02140 model
AT5G64890 data
AT5G64890 model
AT5G65300 data
AT5G65300 model
AT1G05020 data
AT1G05020 model

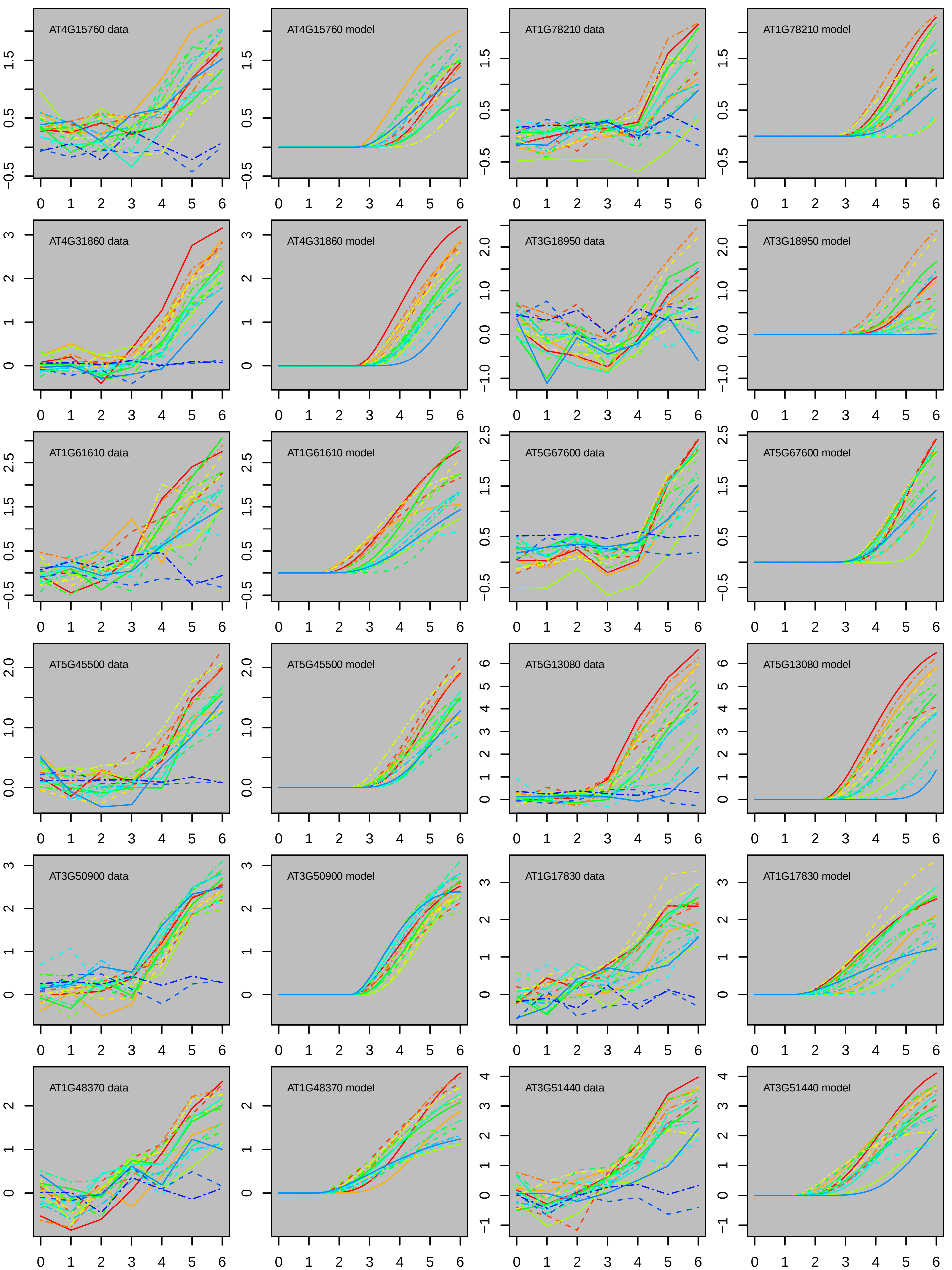

AT4G15760 data
AT4G15760 model
AT1G78210 data
AT1G78210 model
AT4G31860 data
AT4G31860 model
AT3G18950 data
AT3G18950 model
AT1G61610 data
AT1G61610 model
AT5G67600 data
AT5G67600 model
AT5G45500 data
AT5G45500 model
AT5G13080 data
AT5G13080 model
AT3G50900 data
AT3G50900 model
AT1G17830 data
AT1G17830 model
AT1G48370 data
AT1G48370 model
AT3G51440 data
AT3G51440 model

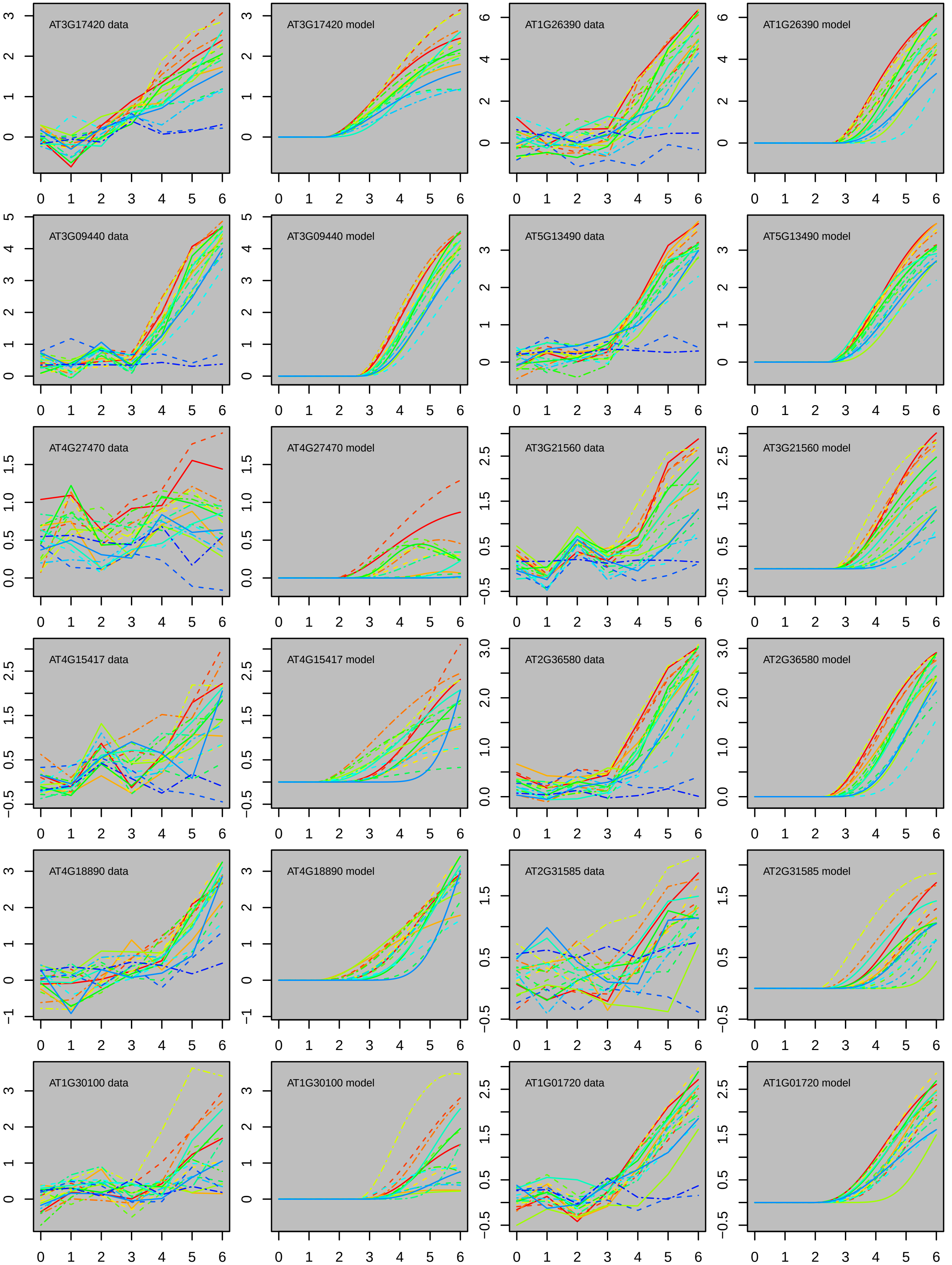
AT3G17420 data
AT3G17420 model
AT1G26390 data
AT1G26390 model
AT3G09440 data
AT3G09440 model
AT5G13490 data
AT5G13490 model
AT4G27470 data
AT4G27470 model
AT3G21560 data
AT3G21560 model
AT4G15417 data
AT4G15417 model
AT2G36580 data
AT2G36580 model
AT4G18890 data
AT4G18890 model
AT2G31585 data
AT2G31585 model
AT1G30100 data
AT1G30100 model
AT1G01720 data
AT1G01720 model

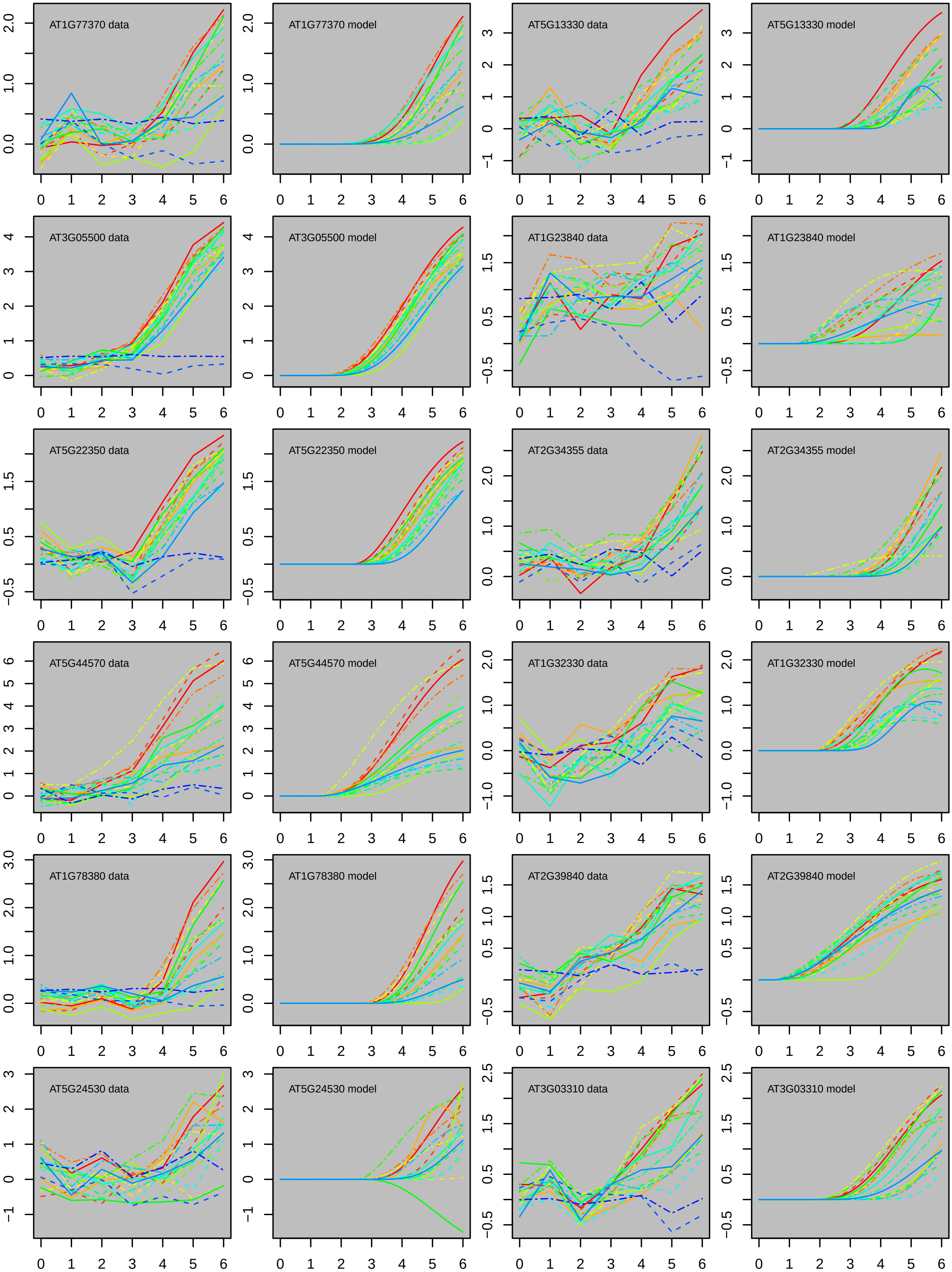
AT1G77370 data
AT1G77370 model
AT5G13330 data
AT5G13330 model
AT3G05500 data
AT3G05500 model
AT1G23840 data
AT1G23840 model
AT5G22350 data
AT5G22350 model
AT2G34355 data
AT2G34355 model
AT5G44570 data
AT5G44570 model
AT1G32330 data
AT1G32330 model
AT1G78380 data
AT1G78380 model
AT2G39840 data
AT2G39840 model
AT5G24530 data
AT5G24530 model
AT3G03310 data
AT3G03310 model

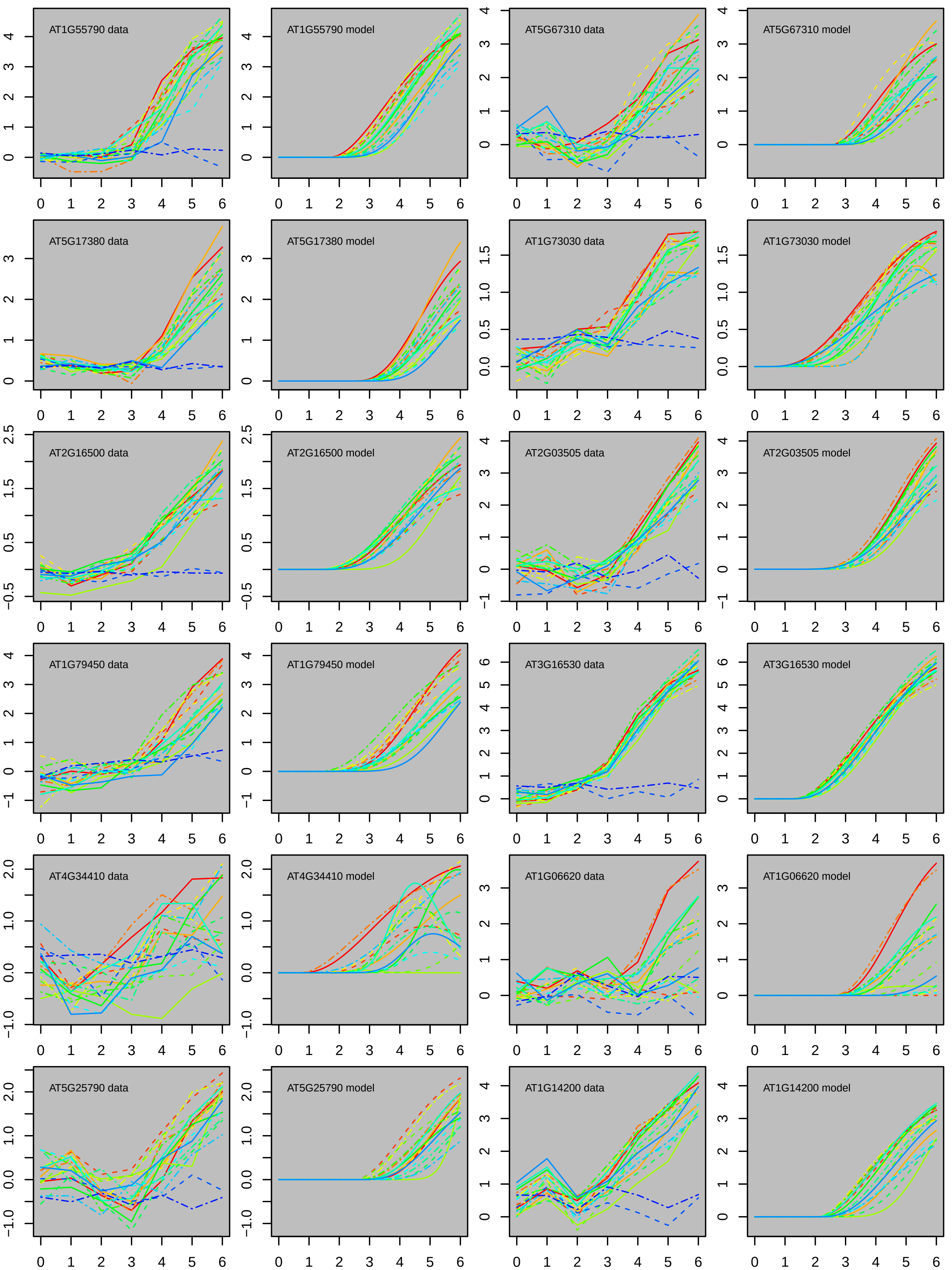

AT1G55790 data
AT1G55790 model
AT5G67310 data
AT5G67310 model
AT5G17380 data
AT5G17380 model
AT1G73030 data
AT1G73030 model
AT2G16500 data
AT2G16500 model
AT2G03505 data
AT2G03505 model
AT1G79450 data
AT1G79450 model
AT3G16530 data
AT3G16530 model
AT4G34410 data
AT4G34410 model
AT1G06620 data
AT1G06620 model
AT5G25790 data
AT5G25790 model
AT1G14200 data
AT1G14200 model

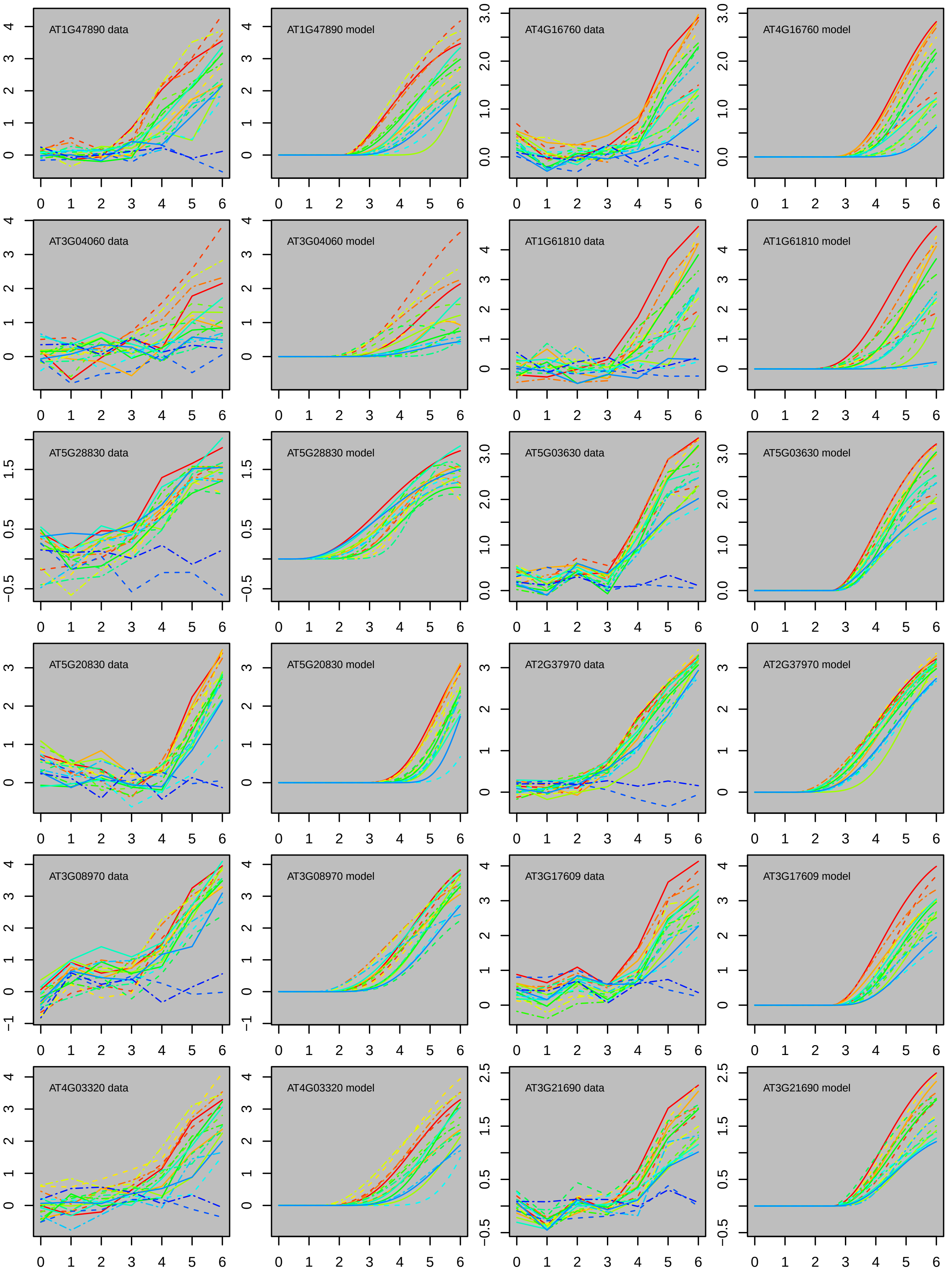
AT1G47890 data
AT1G47890 model
AT4G16760 data
AT4G16760 model
AT3G04060 data
AT3G04060 model
AT1G61810 data
AT1G61810 model
AT5G28830 data
AT5G28830 model
AT5G03630 data
AT5G03630 model
AT5G20830 data
AT5G20830 model
AT2G37970 data
AT2G37970 model
AT3G08970 data
AT3G08970 model
AT3G17609 data
AT3G17609 model
AT4G03320 data
AT4G03320 model
AT3G21690 data
AT3G21690 model

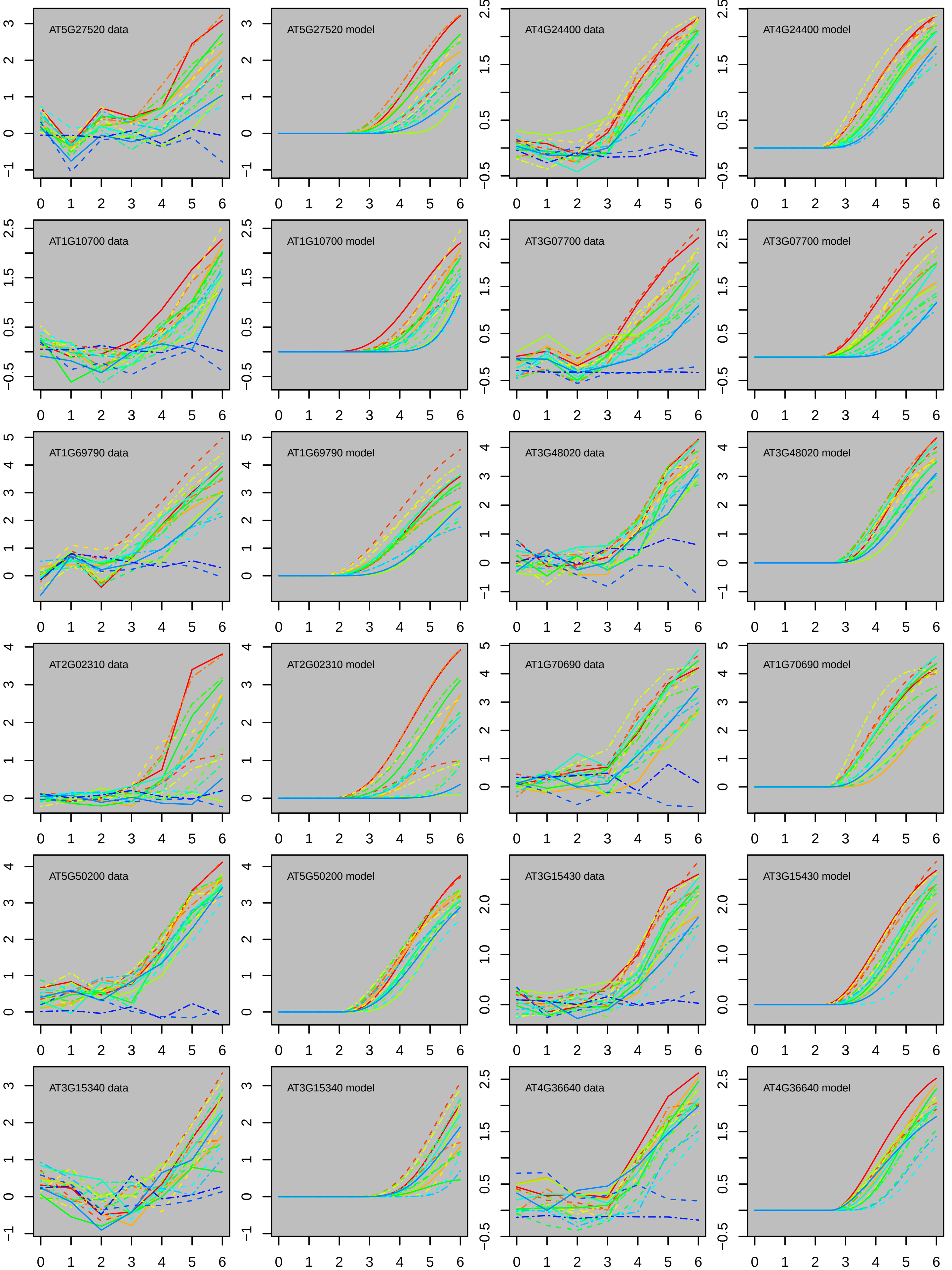
AT5G27520 data
AT5G27520 model
AT4G24400 data
AT4G24400 model
AT1G10700 data
AT1G10700 model
AT3G07700 data
AT3G07700 model
AT1G69790 data
AT1G69790 model
AT3G48020 data
AT3G48020 model
AT2G02310 data
AT2G02310 model
AT1G70690 data
AT1G70690 model
AT5G50200 data
AT5G50200 model
AT3G15430 data
AT3G15430 model
AT3G15340 data
AT3G15340 model
AT4G36640 data
AT4G36640 model

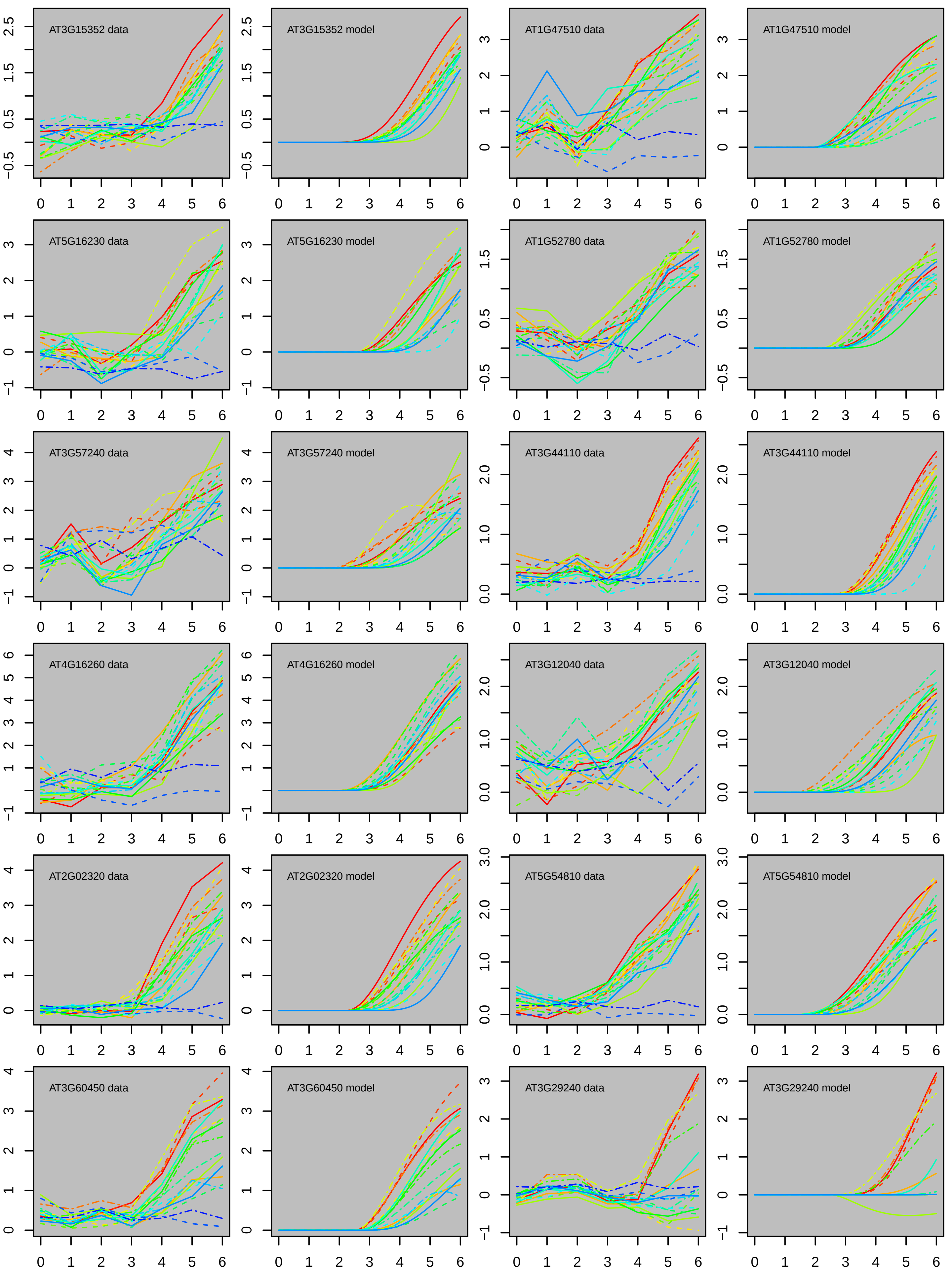

AT3G15352 data
AT3G15352 model
AT1G47510 data
AT1G47510 model
AT5G16230 data
AT5G16230 model
AT1G52780 data
AT1G52780 model
AT3G57240 data
AT3G57240 model
AT3G44110 data
AT3G44110 model
AT4G16260 data
AT4G16260 model
AT3G12040 data
AT3G12040 model
AT2G02320 data
AT2G02320 model
AT5G54810 data
AT5G54810 model
AT3G60450 data
AT3G60450 model
AT3G29240 data
AT3G29240 model

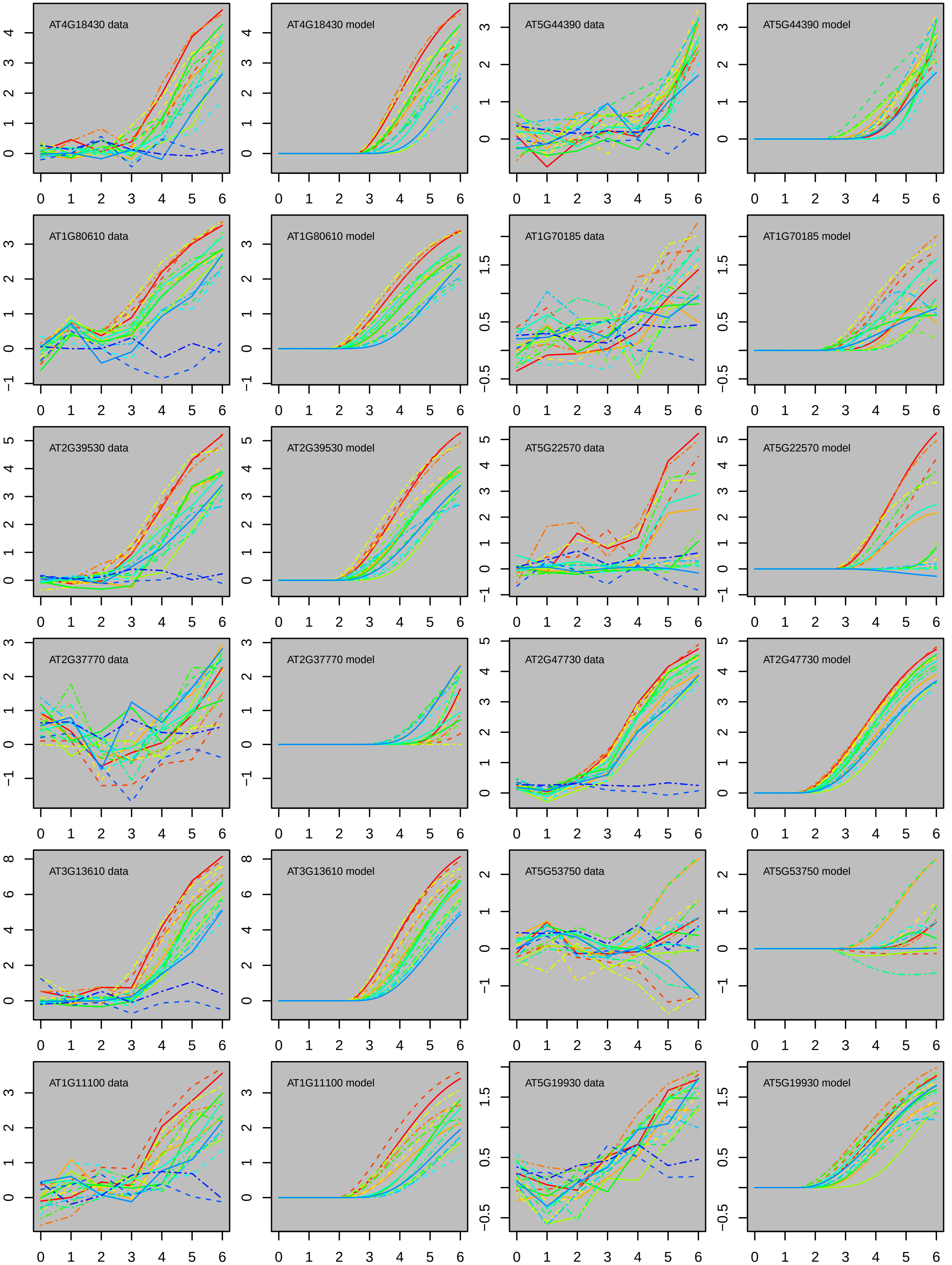
AT4G18430 data
AT4G18430 model
AT5G44390 data
AT5G44390 model
AT1G80610 data
AT1G80610 model
AT1G70185 data
AT1G70185 model
AT2G39530 data
AT2G39530 model
AT5G22570 data
AT5G22570 model
AT2G37770 data
AT2G37770 model
AT2G47730 data
AT2G47730 model
AT3G13610 data
AT3G13610 model
AT5G53750 data
AT5G53750 model
AT1G11100 data
AT1G11100 model
AT5G19930 data
AT5G19930 model

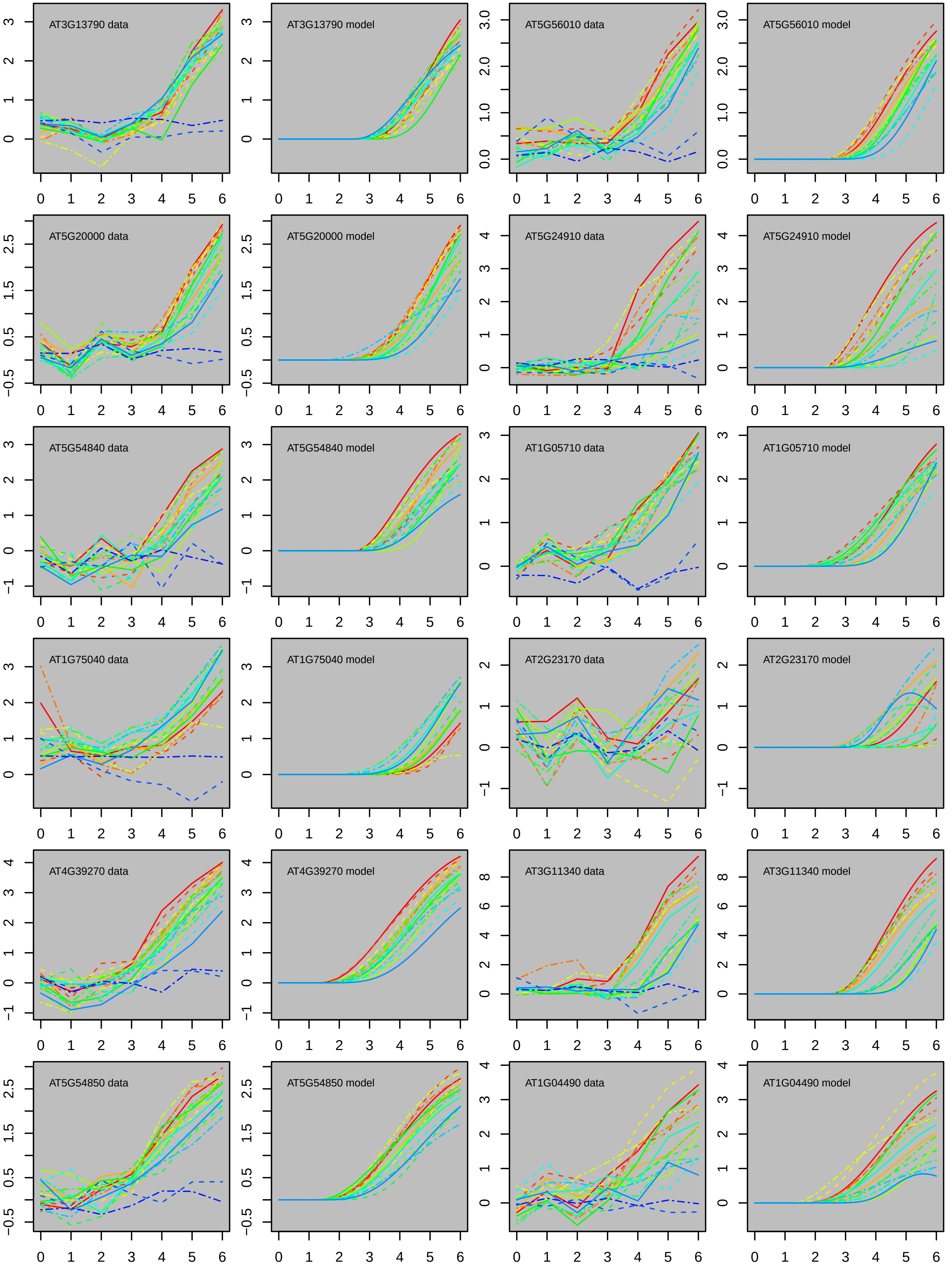
AT3G13790 data
AT3G13790 model
AT5G56010 data
AT5G56010 model
AT5G20000 data
AT5G20000 model
AT5G24910 data
AT5G24910 model
AT5G54840 data
AT5G54840 model
AT1G05710 data
AT1G05710 model
AT1G75040 data
AT1G75040 model
AT2G23170 data
AT2G23170 model
AT4G39270 data
AT4G39270 model
AT3G11340 data
AT3G11340 model
AT5G54850 data
AT5G54850 model
AT1G04490 data
AT1G04490 model

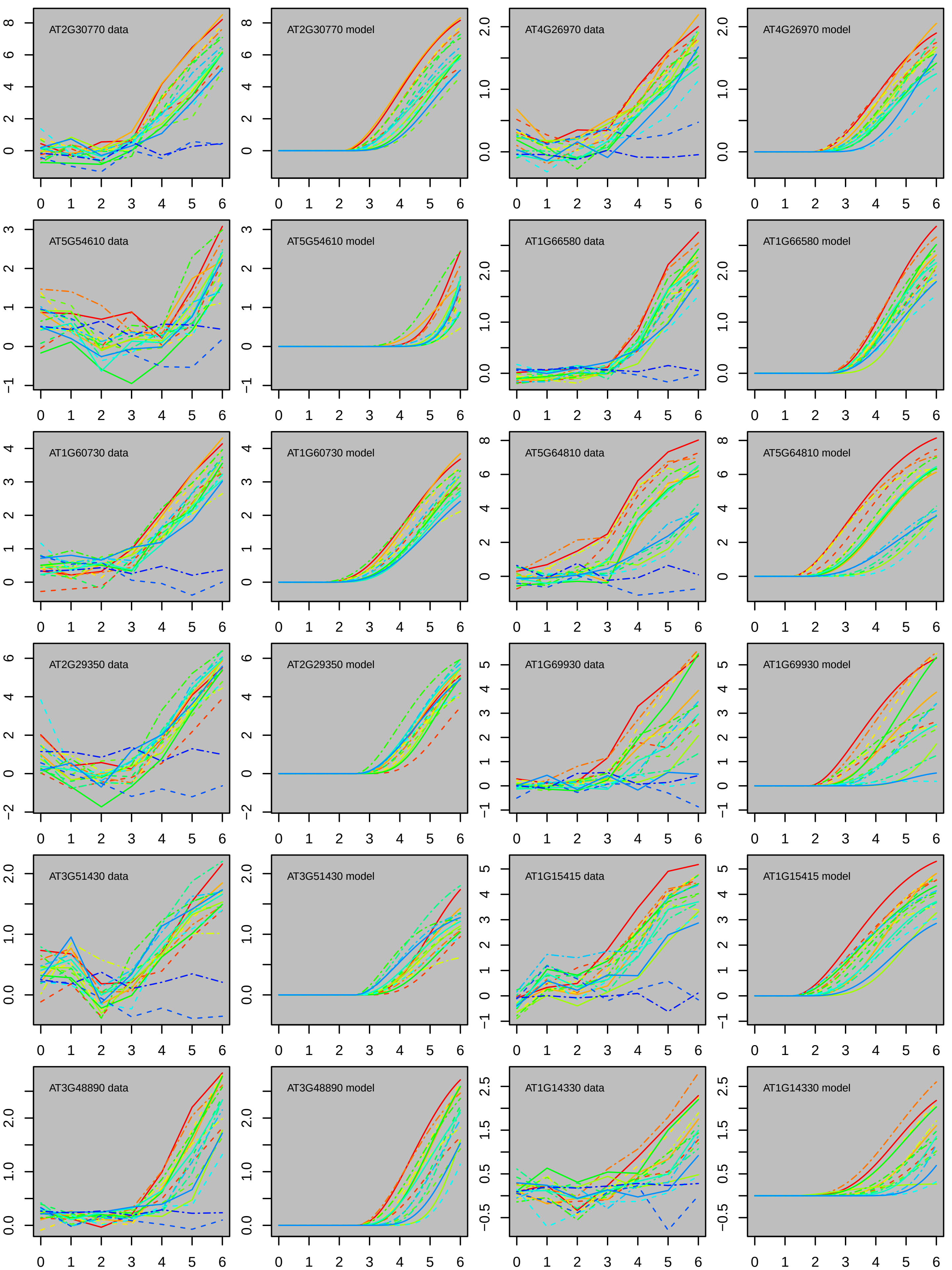

AT2G30770 data
AT2G30770 model
AT4G26970 data
AT4G26970 model
AT5G54610 data
AT5G54610 model
AT1G66580 data
AT1G66580 model
AT1G60730 data
AT1G60730 model
AT5G64810 data
AT5G64810 model
AT2G29350 data
AT2G29350 model
AT1G69930 data
AT1G69930 model
AT3G51430 data
AT3G51430 model
AT1G15415 data
AT1G15415 model
AT3G48890 data
AT3G48890 model
AT1G14330 data
AT1G14330 model

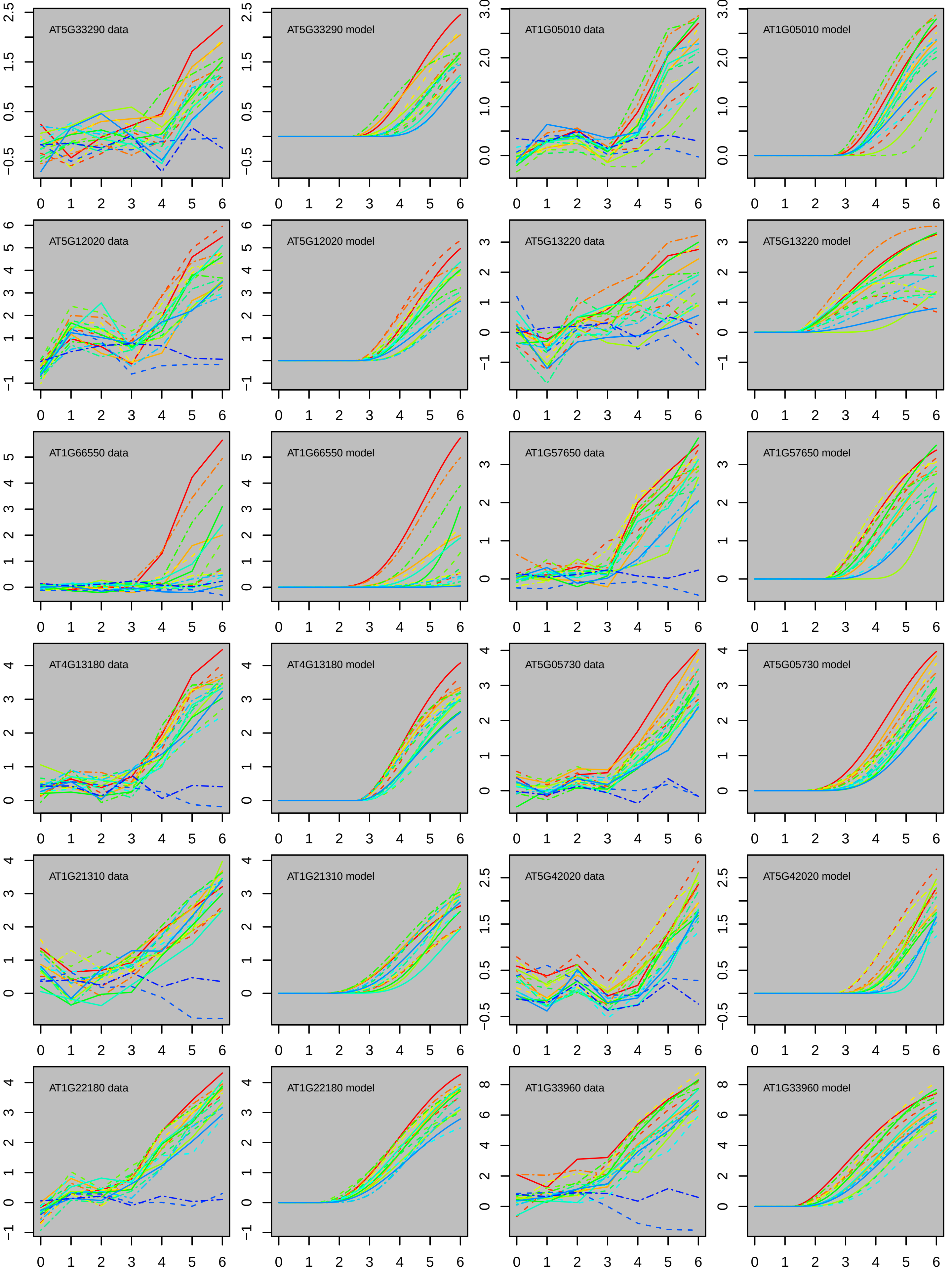
AT5G33290 data
AT5G33290 model
AT1G05010 data
AT1G05010 model
AT5G12020 data
AT5G12020 model
AT5G13220 data
AT5G13220 model
AT1G66550 data
AT1G66550 model
AT1G57650 data
AT1G57650 model
AT4G13180 data
AT4G13180 model
AT5G05730 data
AT5G05730 model
AT1G21310 data
AT1G21310 model
AT5G42020 data
AT5G42020 model
AT1G22180 data
AT1G22180 model
AT1G33960 data
AT1G33960 model

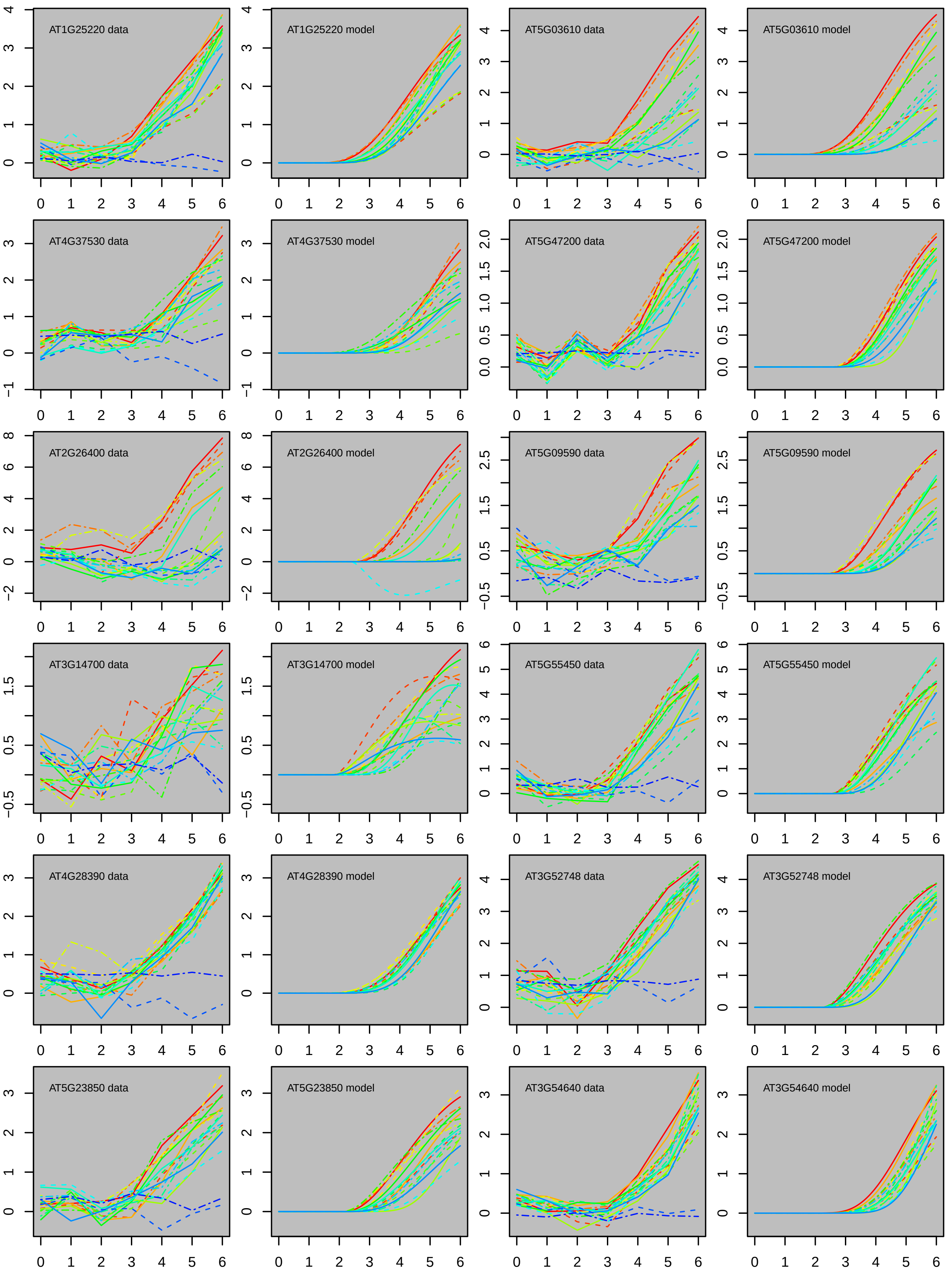
AT1G25220 data
AT1G25220 model
AT5G03610 data
AT5G03610 model
AT4G37530 data
AT4G37530 model
AT5G47200 data
AT5G47200 model
AT2G26400 data
AT2G26400 model
AT5G09590 data
AT5G09590 model
AT3G14700 data
AT3G14700 model
AT5G55450 data
AT5G55450 model
AT4G28390 data
AT4G28390 model
AT3G52748 data
AT3G52748 model
AT5G23850 data
AT5G23850 model
AT3G54640 data
AT3G54640 model

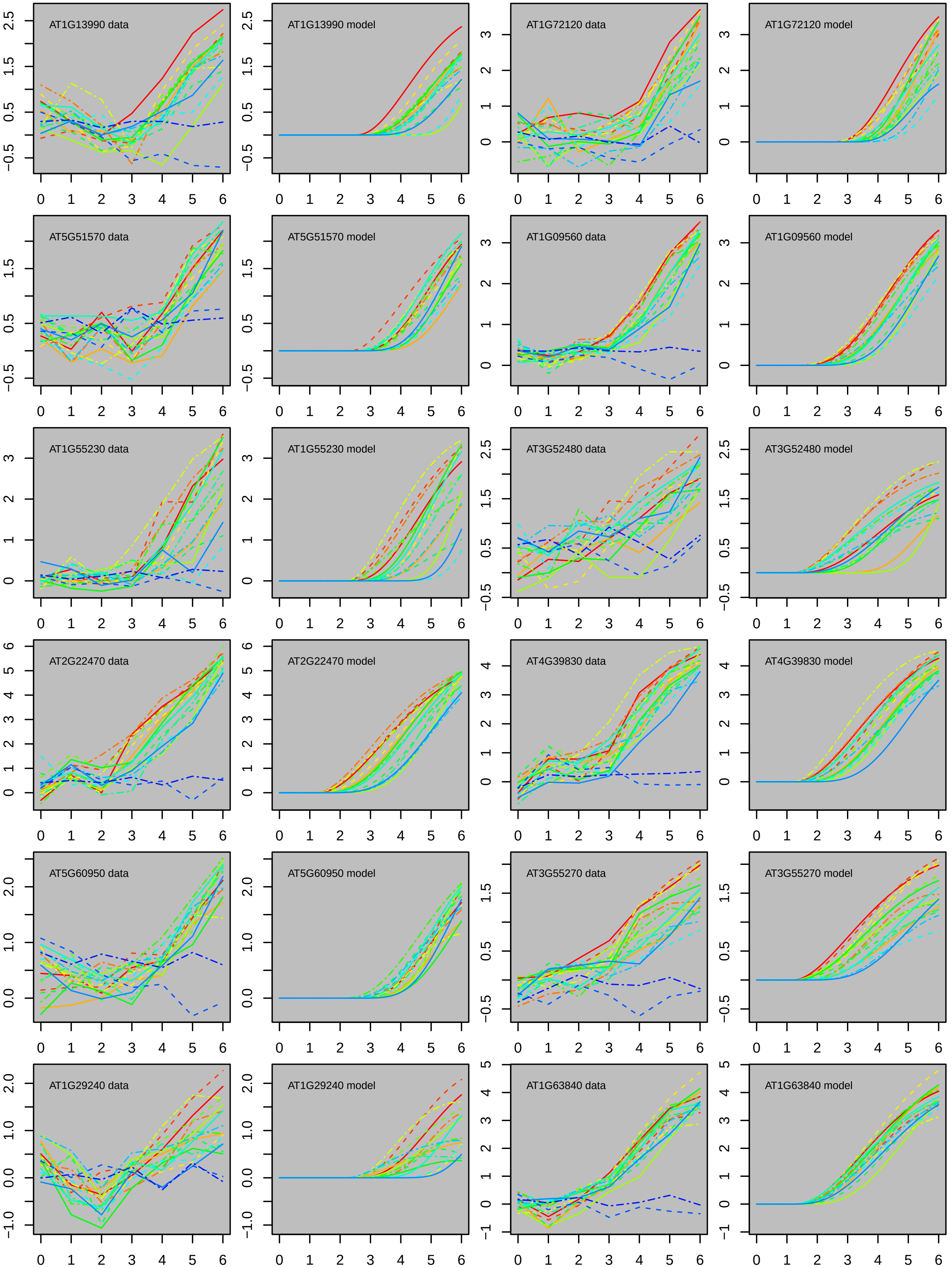
AT1G13990 data
AT1G13990 model
AT1G72120 data
AT1G72120 model
AT5G51570 data
AT5G51570 model
AT1G09560 data
AT1G09560 model
AT1G55230 data
AT1G55230 model
AT3G52480 data
AT3G52480 model
AT2G22470 data
AT2G22470 model
AT4G39830 data
AT4G39830 model
AT5G60950 data
AT5G60950 model
AT3G55270 data
AT3G55270 model
AT1G29240 data
AT1G29240 model
AT1G63840 data
AT1G63840 model

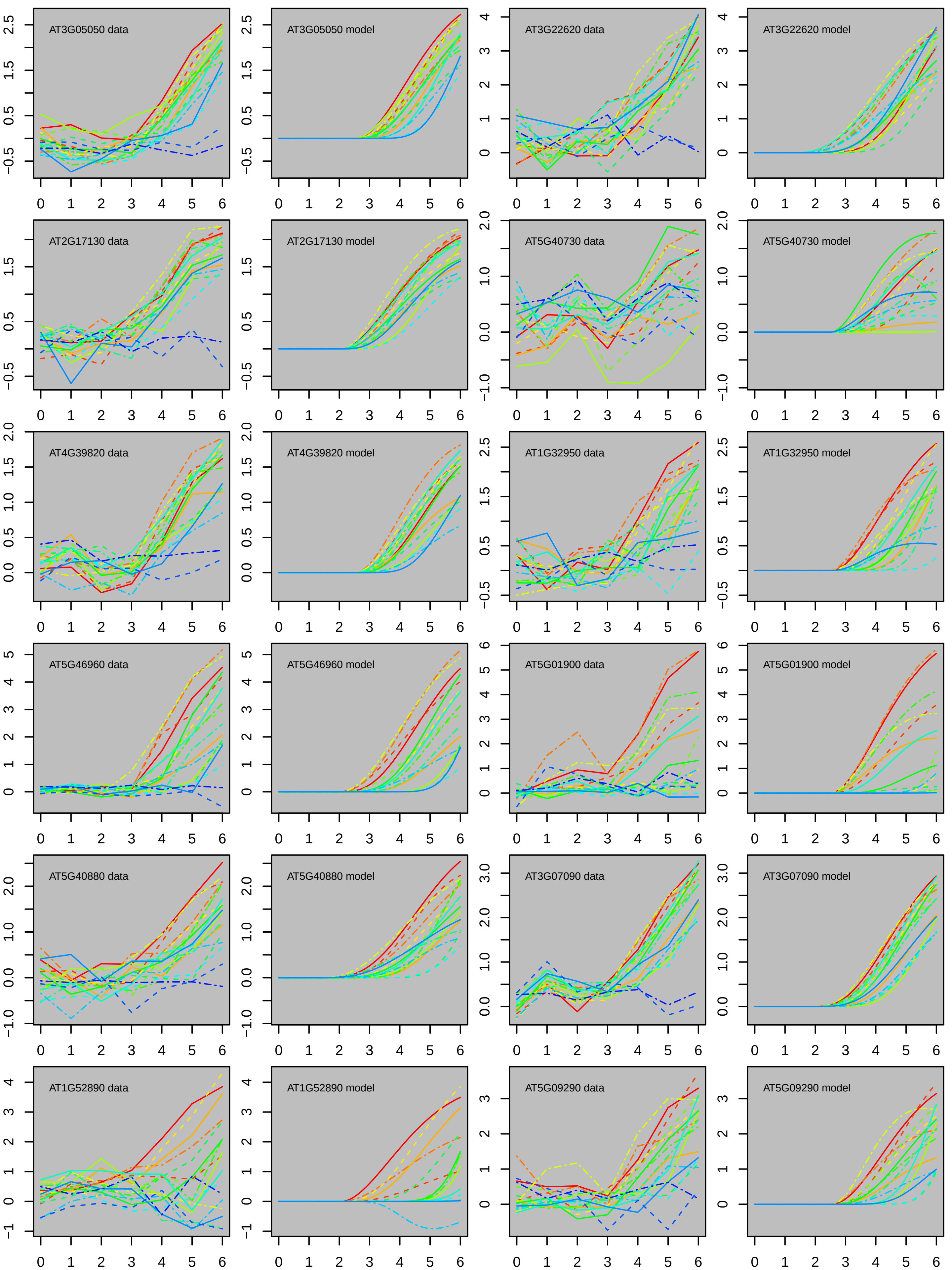

AT3G05050 data
AT3G05050 model
AT3G22620 data
AT3G22620 model
AT2G17130 data
AT2G17130 model
AT5G40730 data
AT5G40730 model
AT4G39820 data
AT4G39820 model
AT1G32950 data
AT1G32950 model
AT5G46960 data
AT5G46960 model
AT5G01900 data
AT5G01900 model
AT5G40880 data
AT5G40880 model
AT3G07090 data
AT3G07090 model
AT1G52890 data
AT1G52890 model
AT5G09290 data
AT5G09290 model

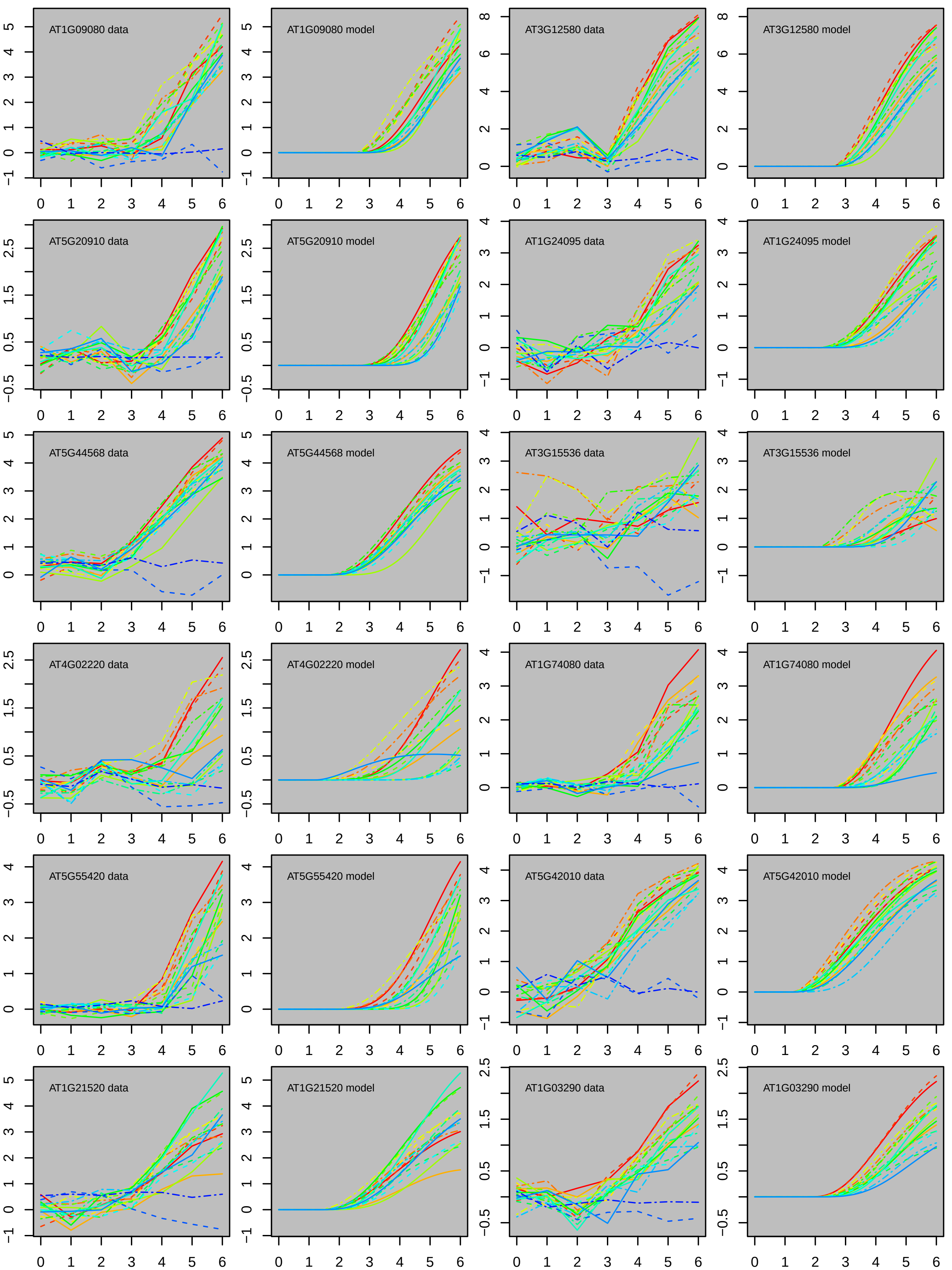

AT1G09080 data
AT1G09080 model
AT3G12580 data
AT3G12580 model
AT5G20910 data
AT5G20910 model
AT1G24095 data
AT1G24095 model
AT5G44568 data
AT5G44568 model
AT3G15536 data
AT3G15536 model
AT4G02220 data
AT4G02220 model
AT1G74080 data
AT1G74080 model
AT5G55420 data
AT5G55420 model
AT5G42010 data
AT5G42010 model
AT1G21520 data
AT1G21520 model
AT1G03290 data
AT1G03290 model

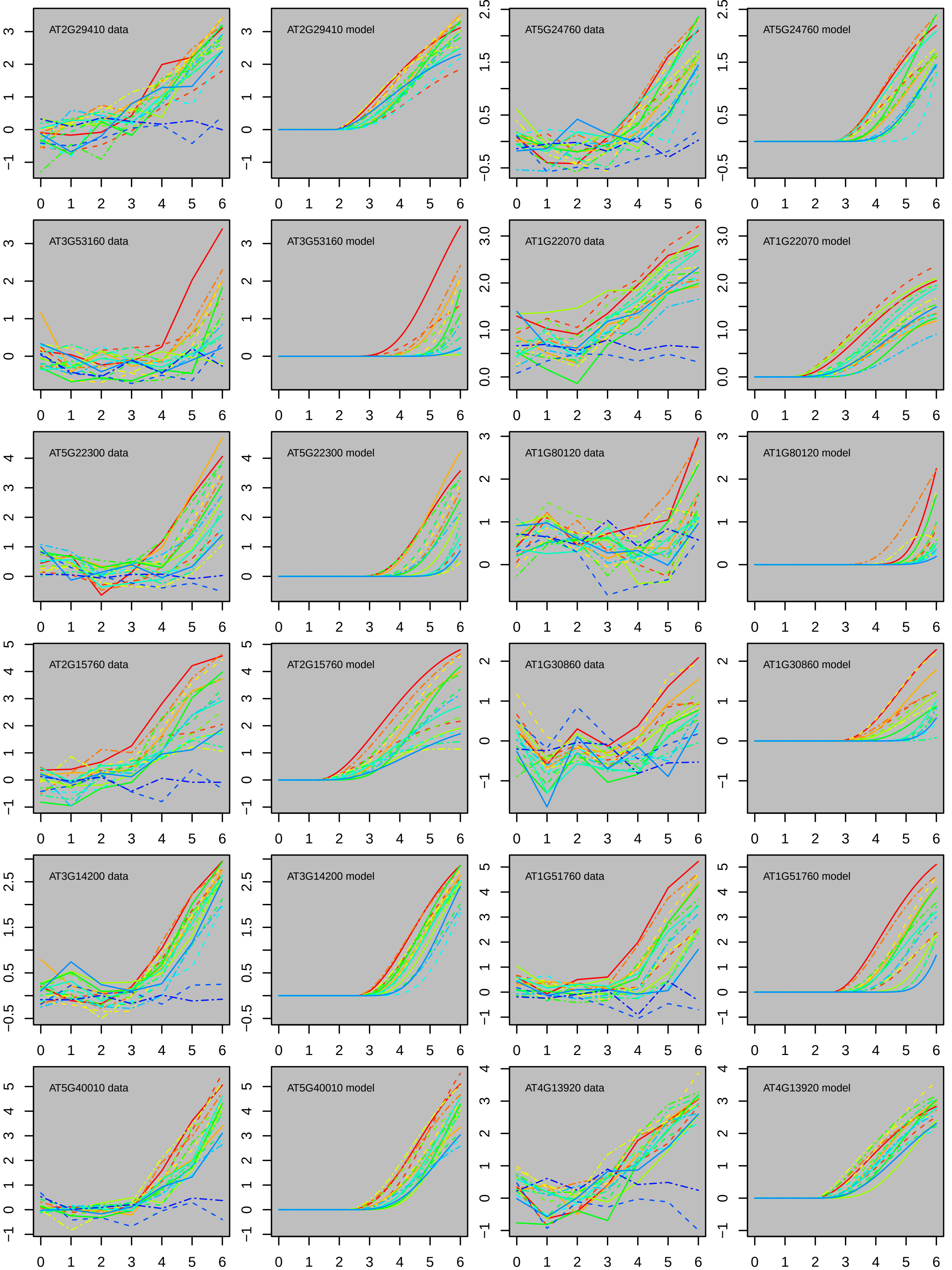

AT2G29410 data
AT2G29410 model
AT5G24760 data
AT5G24760 model
AT3G53160 data
AT3G53160 model
AT1G22070 data
AT1G22070 model
AT5G22300 data
AT5G22300 model
AT1G80120 data
AT1G80120 model
AT2G15760 data
AT2G15760 model
AT1G30860 data
AT1G30860 model
AT3G14200 data
AT3G14200 model
AT1G51760 data
AT1G51760 model
AT5G40010 data
AT5G40010 model
AT4G13920 data
AT4G13920 model

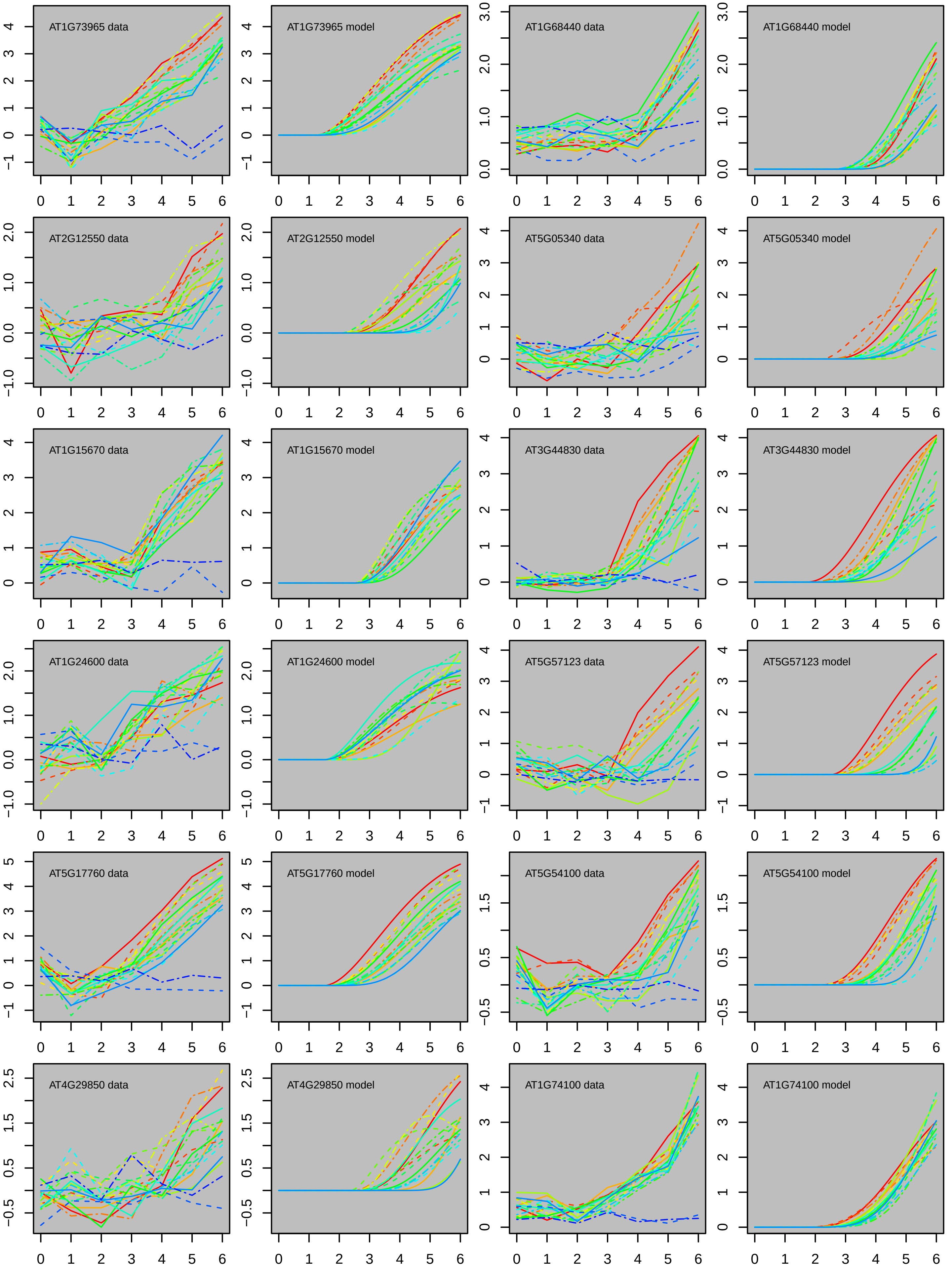
AT1G73965 data
AT1G73965 model
AT1G68440 data
AT1G68440 model
AT2G12550 data
AT2G12550 model
AT5G05340 data
AT5G05340 model
AT1G15670 data
AT1G15670 model
AT3G44830 data
AT3G44830 model
AT1G24600 data
AT1G24600 model
AT5G57123 data
AT5G57123 model
AT5G17760 data
AT5G17760 model
AT5G54100 data
AT5G54100 model
AT4G29850 data
AT4G29850 model
AT1G74100 data
AT1G74100 model

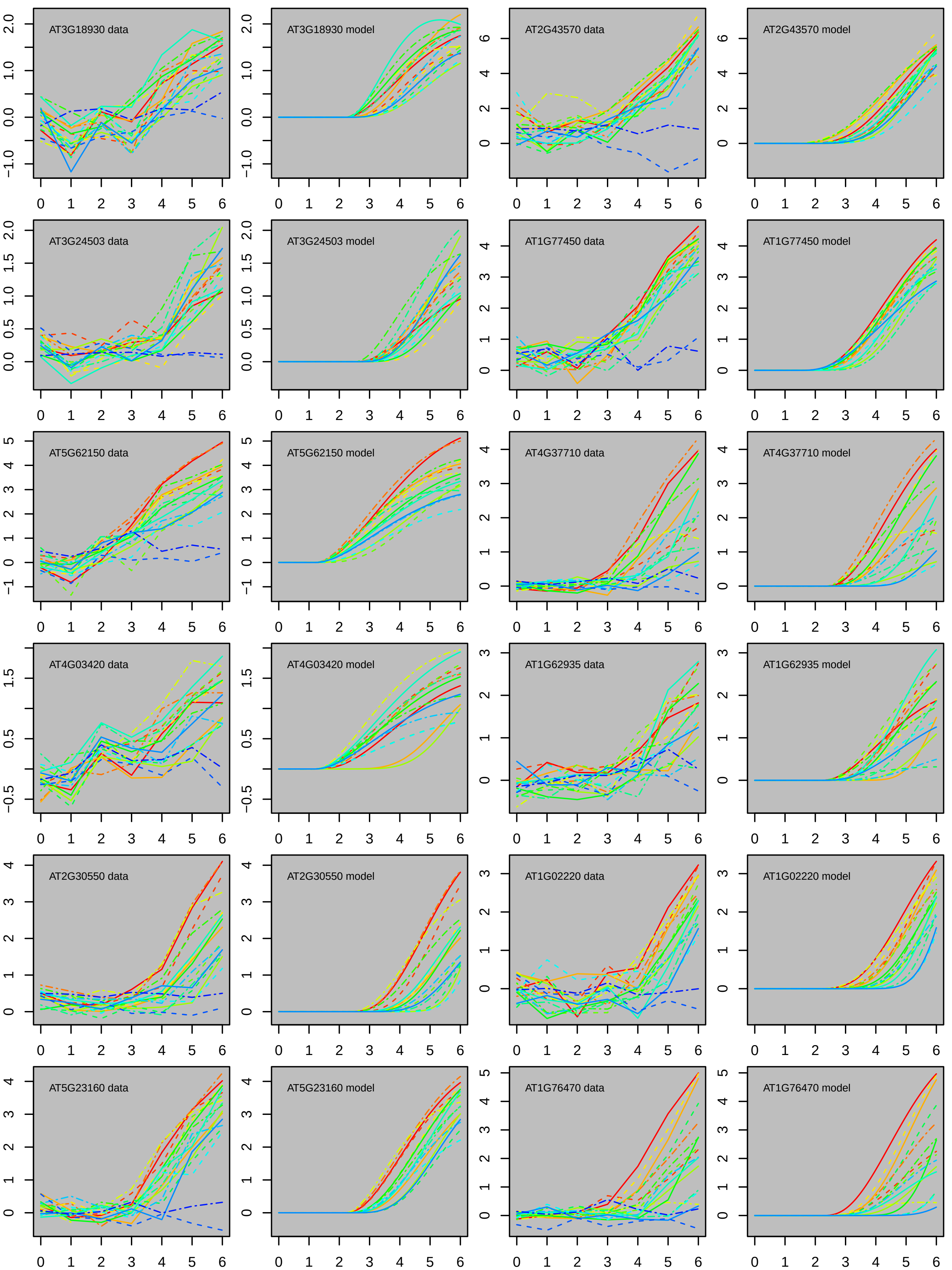

AT3G18930 data
AT3G18930 model
AT2G43570 data
AT2G43570 model
AT3G24503 data
AT3G24503 model
AT1G77450 data
AT1G77450 model
AT5G62150 data
AT5G62150 model
AT4G37710 data
AT4G37710 model
AT4G03420 data
AT4G03420 model
AT1G62935 data
AT1G62935 model
AT2G30550 data
AT2G30550 model
AT1G02220 data
AT1G02220 model
AT5G23160 data
AT5G23160 model
AT1G76470 data
AT1G76470 model
0 1 2 3 4 5 6

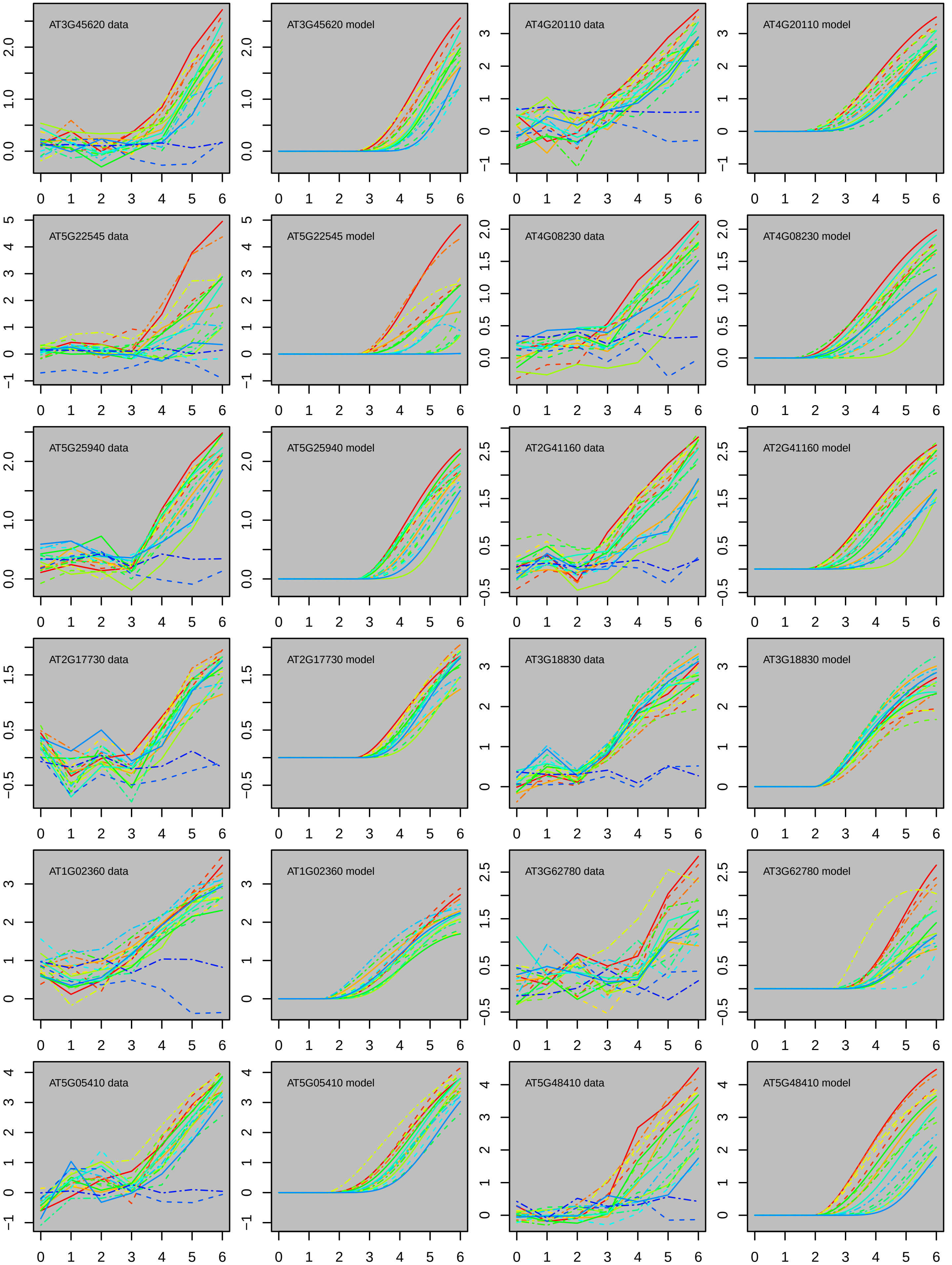
AT3G45620 data
AT3G45620 model
AT4G20110 data
AT4G20110 model
AT5G22545 data
AT5G22545 model
AT4G08230 data
AT4G08230 model
AT5G25940 data
AT5G25940 model
AT2G41160 data
AT2G41160 model
AT2G17730 data
AT2G17730 model
AT3G18830 data
AT3G18830 model
AT1G02360 data
AT1G02360 model
AT3G62780 data
AT3G62780 model
AT5G05410 data
AT5G05410 model
AT5G48410 data
AT5G48410 model

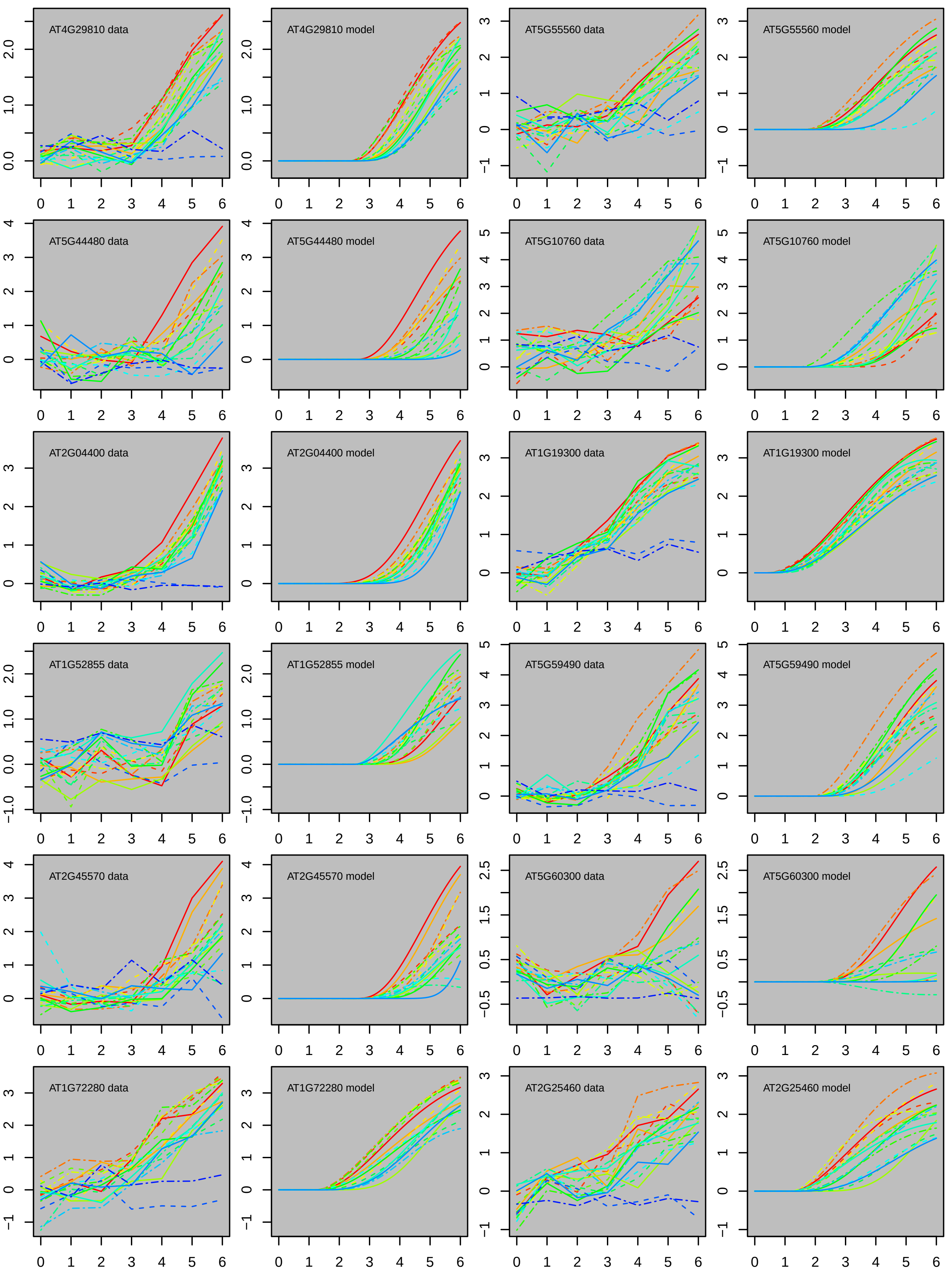
AT4G29810 data
AT4G29810 model
AT5G55560 data
AT5G55560 model
AT5G44480 data
AT5G44480 model
AT5G10760 data
AT5G10760 model
AT2G04400 data
AT2G04400 model
AT1G19300 data
AT1G19300 model
AT1G52855 data
AT1G52855 model
AT5G59490 data
AT5G59490 model
AT2G45570 data
AT2G45570 model
AT5G60300 data
AT5G60300 model
AT1G72280 data
AT1G72280 model
AT2G25460 data
AT2G25460 model

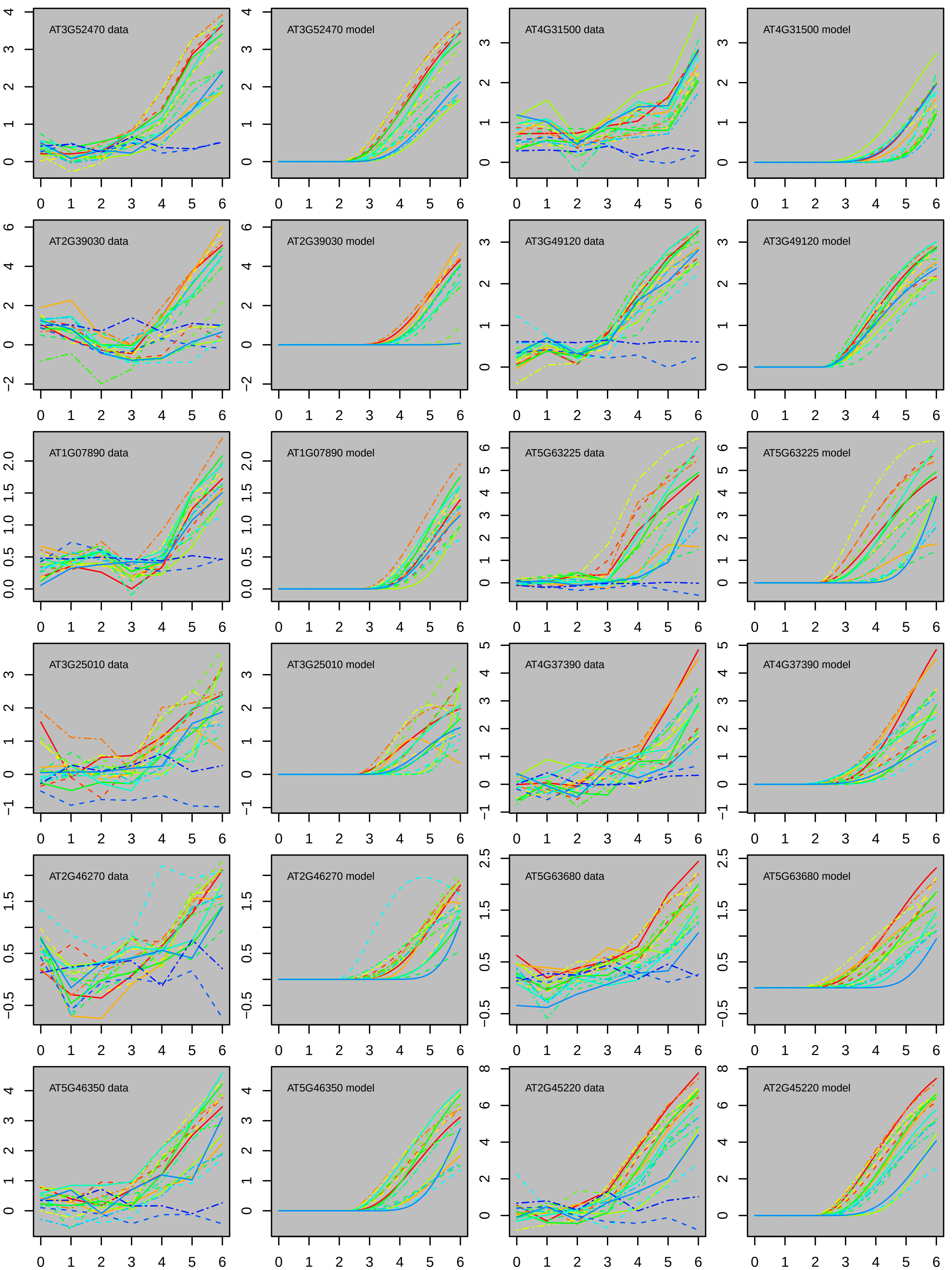
AT3G52470 data
AT3G52470 model
AT4G31500 data
AT4G31500 model
AT2G39030 data
AT2G39030 model
AT3G49120 data
AT3G49120 model
AT1G07890 data
AT1G07890 model
AT5G63225 data
AT5G63225 model
AT3G25010 data
AT3G25010 model
AT4G37390 data
AT4G37390 model
AT2G46270 data
AT2G46270 model
AT5G63680 data
AT5G63680 model
AT5G46350 data
AT5G46350 model
AT2G45220 data
AT2G45220 model

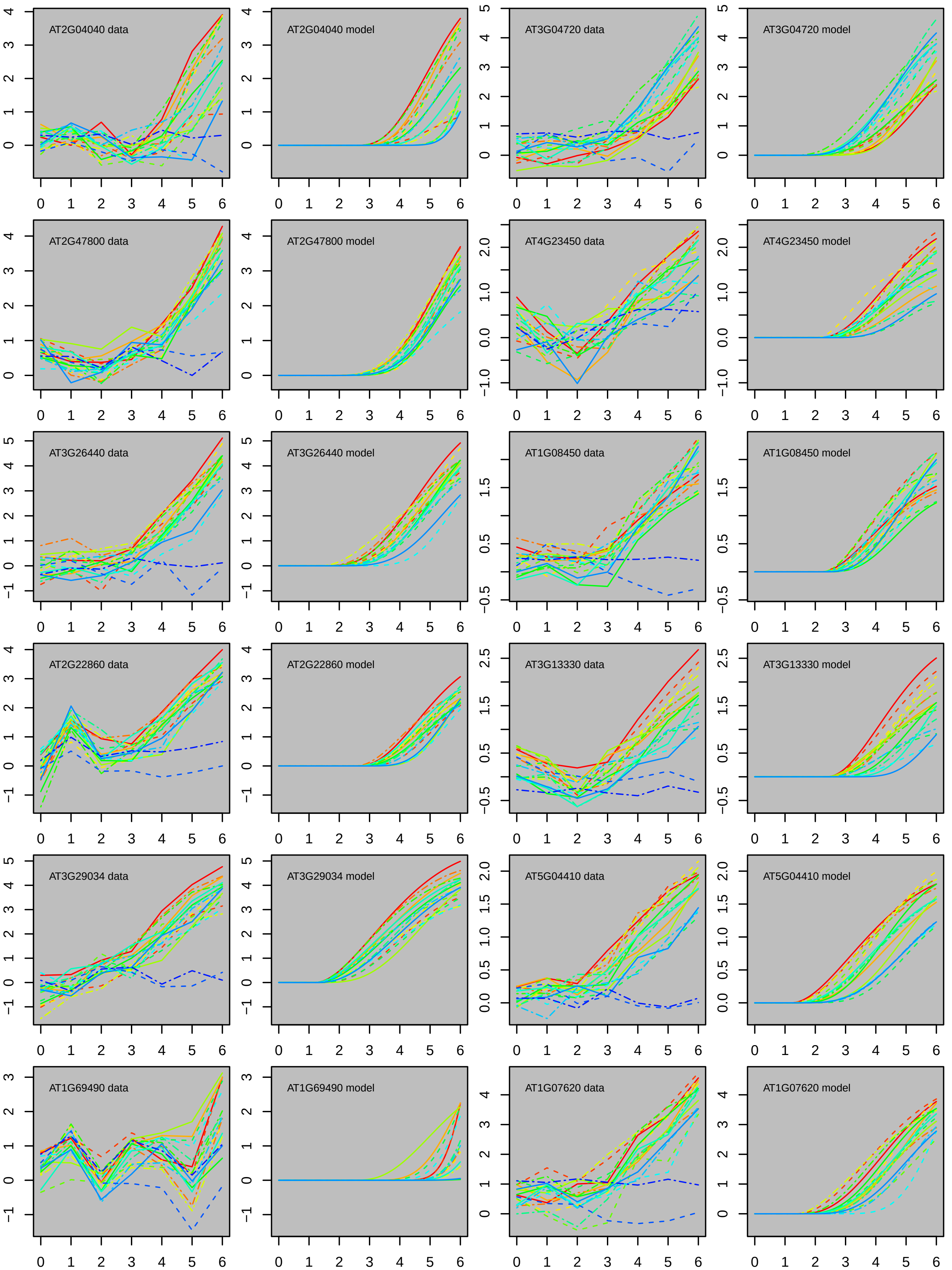
AT2G04040 data
AT2G04040 model
AT3G04720 data
AT3G04720 model
AT2G47800 data
AT2G47800 model
AT4G23450 data
AT4G23450 model
AT3G26440 data
AT3G26440 model
AT1G08450 data
AT1G08450 model
AT2G22860 data
AT2G22860 model
AT3G13330 data
AT3G13330 model
AT3G29034 data
AT3G29034 model
AT5G04410 data
AT5G04410 model
AT1G69490 data
AT1G69490 model
AT1G07620 data
AT1G07620 model

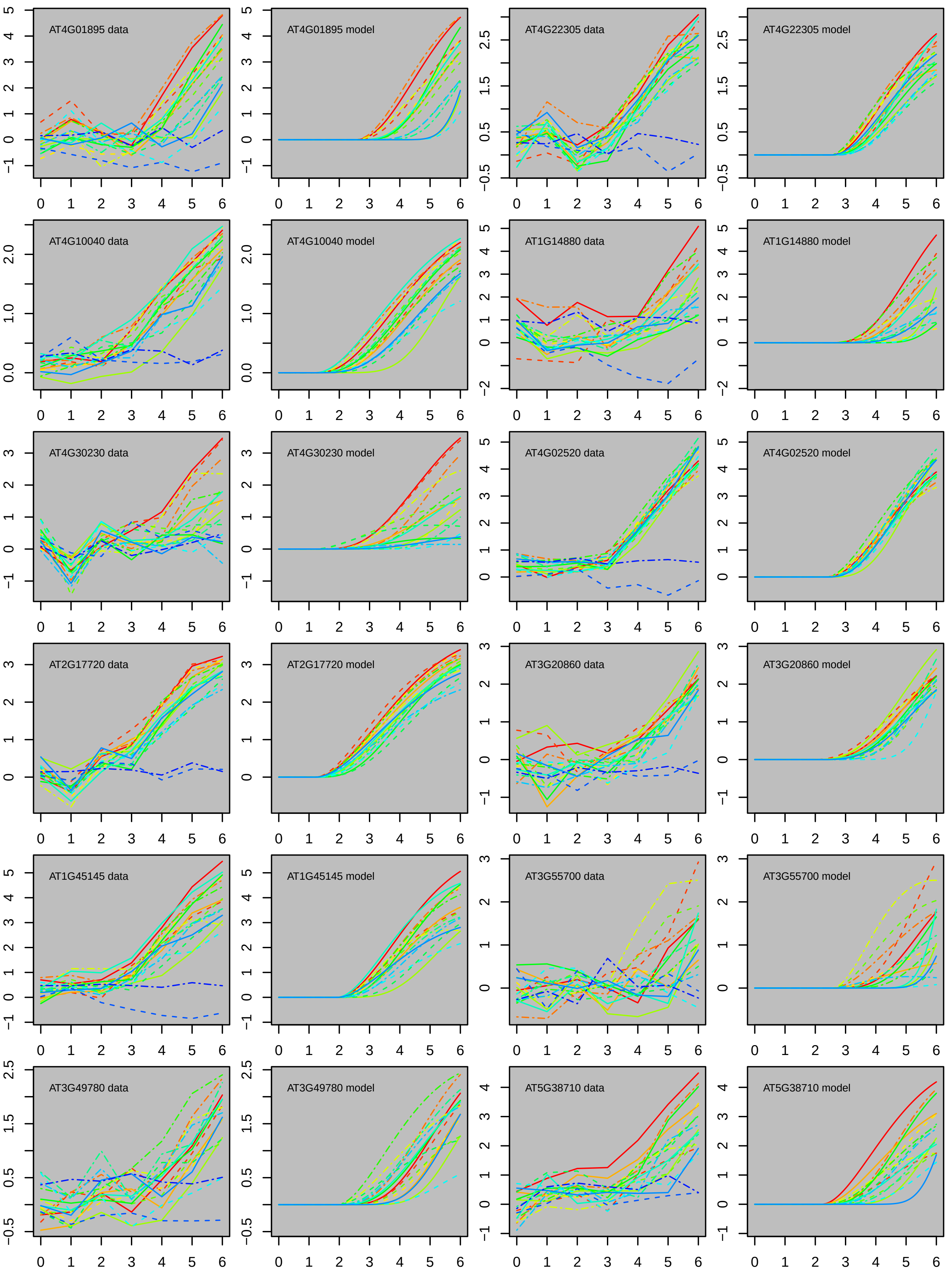

AT4G01895 data
AT4G01895 model
AT4G22305 data
AT4G22305 model
AT4G10040 data
AT4G10040 model
AT1G14880 data
AT1G14880 model
AT4G30230 data
AT4G30230 model
AT4G02520 data
AT4G02520 model
AT2G17720 data
AT2G17720 model
AT3G20860 data
AT3G20860 model
AT1G45145 data
AT1G45145 model
AT3G55700 data
AT3G55700 model
AT3G49780 data
AT3G49780 model
AT5G38710 data
AT5G38710 model

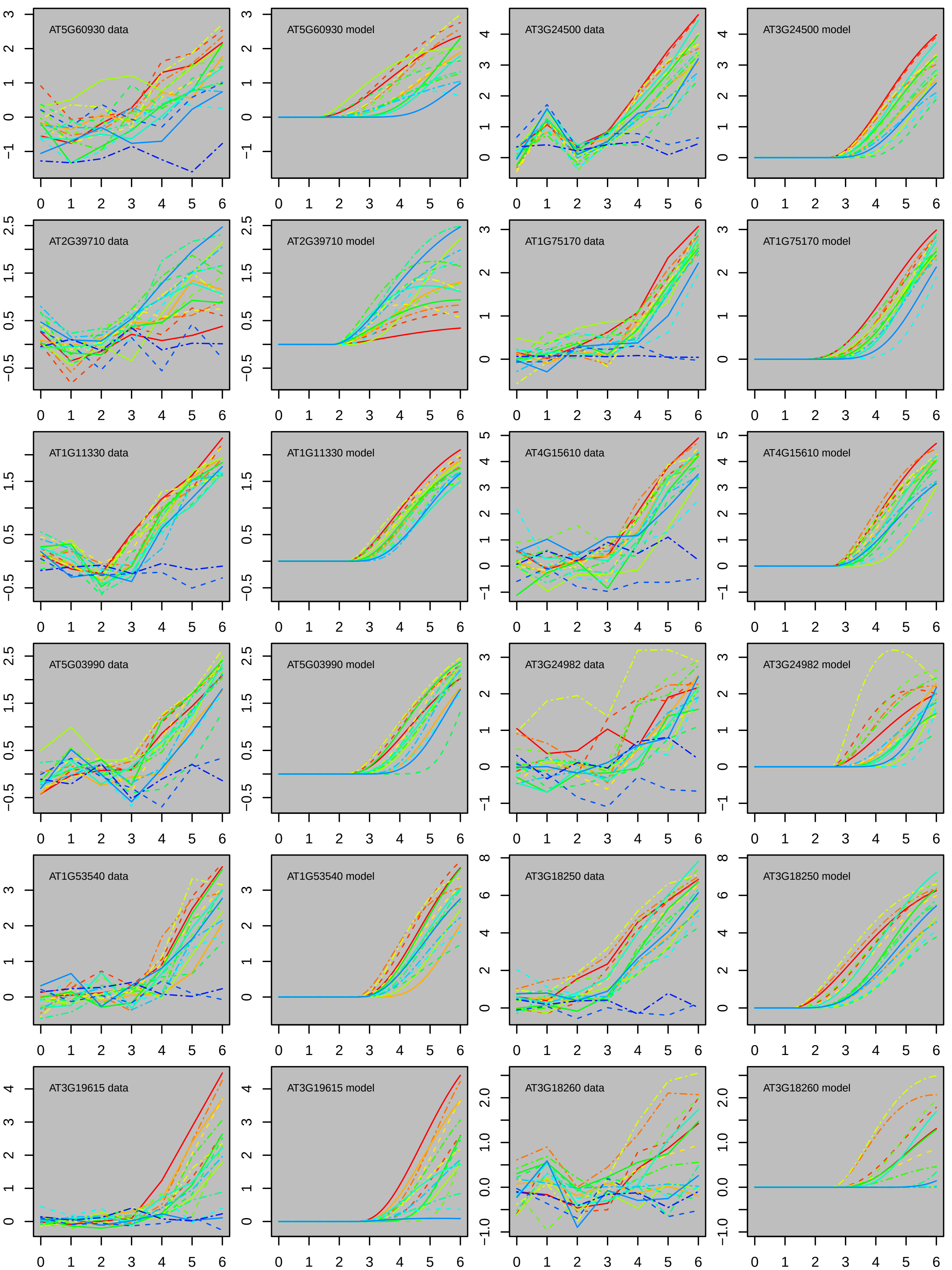
AT5G60930 data
AT5G60930 model
AT3G24500 data
AT3G24500 model
AT2G39710 data
AT2G39710 model
AT1G75170 data
AT1G75170 model
AT1G11330 data
AT1G11330 model
AT4G15610 data
AT4G15610 model
AT5G03990 data
AT5G03990 model
AT3G24982 data
AT3G24982 model
AT1G53540 data
AT1G53540 model
AT3G18250 data
AT3G18250 model
AT3G19615 data
AT3G19615 model
AT3G18260 data
AT3G18260 model

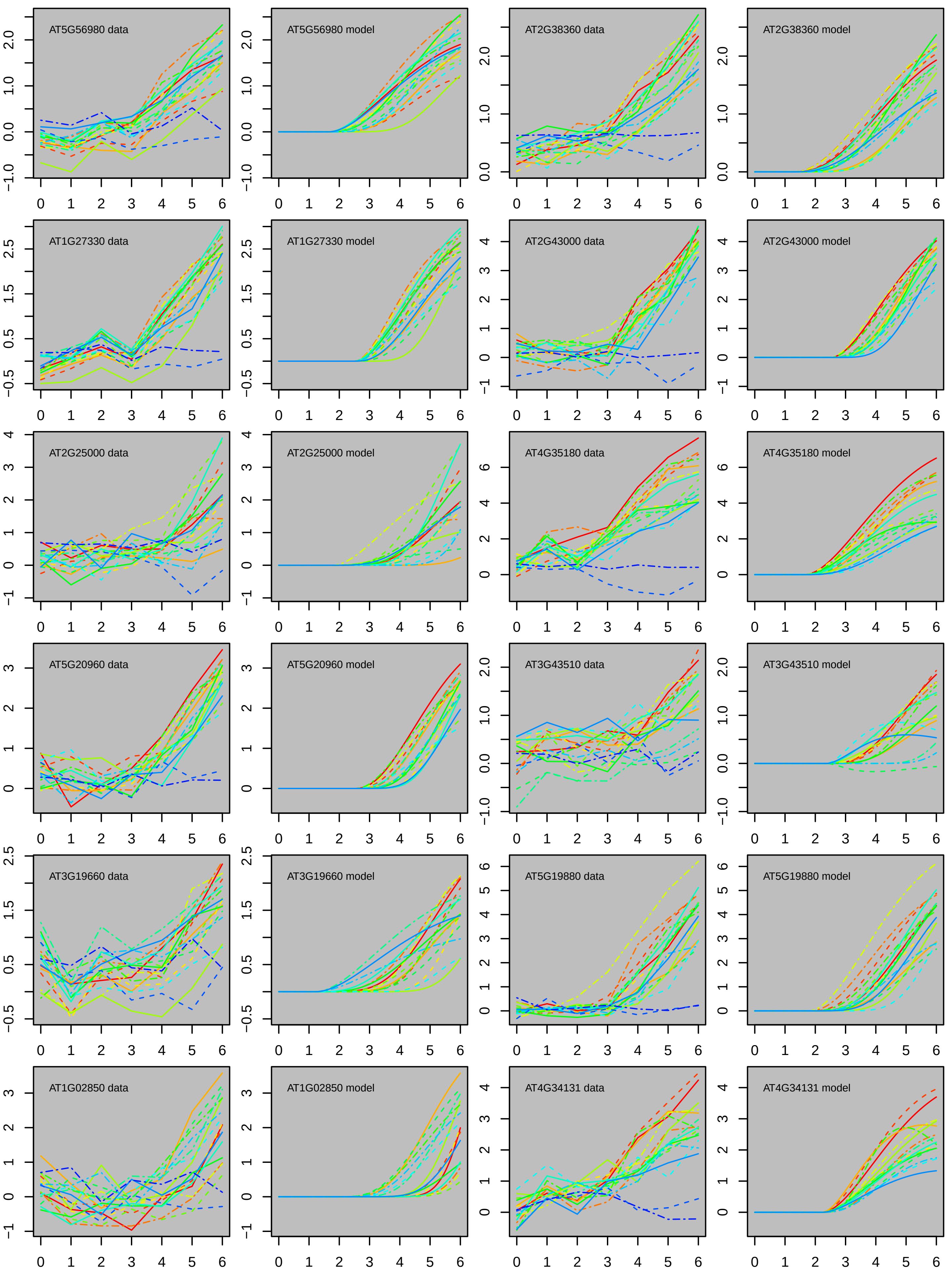
AT5G56980 data
AT5G56980 model
AT2G38360 data
AT2G38360 model
AT1G27330 data
AT1G27330 model
AT2G43000 data
AT2G43000 model
AT2G25000 data
AT2G25000 model
AT4G35180 data
AT4G35180 model
AT5G20960 data
AT5G20960 model
AT3G43510 data
AT3G43510 model
AT3G19660 data
AT3G19660 model
AT5G19880 data
AT5G19880 model
AT1G02850 data
AT1G02850 model
AT4G34131 data
AT4G34131 model

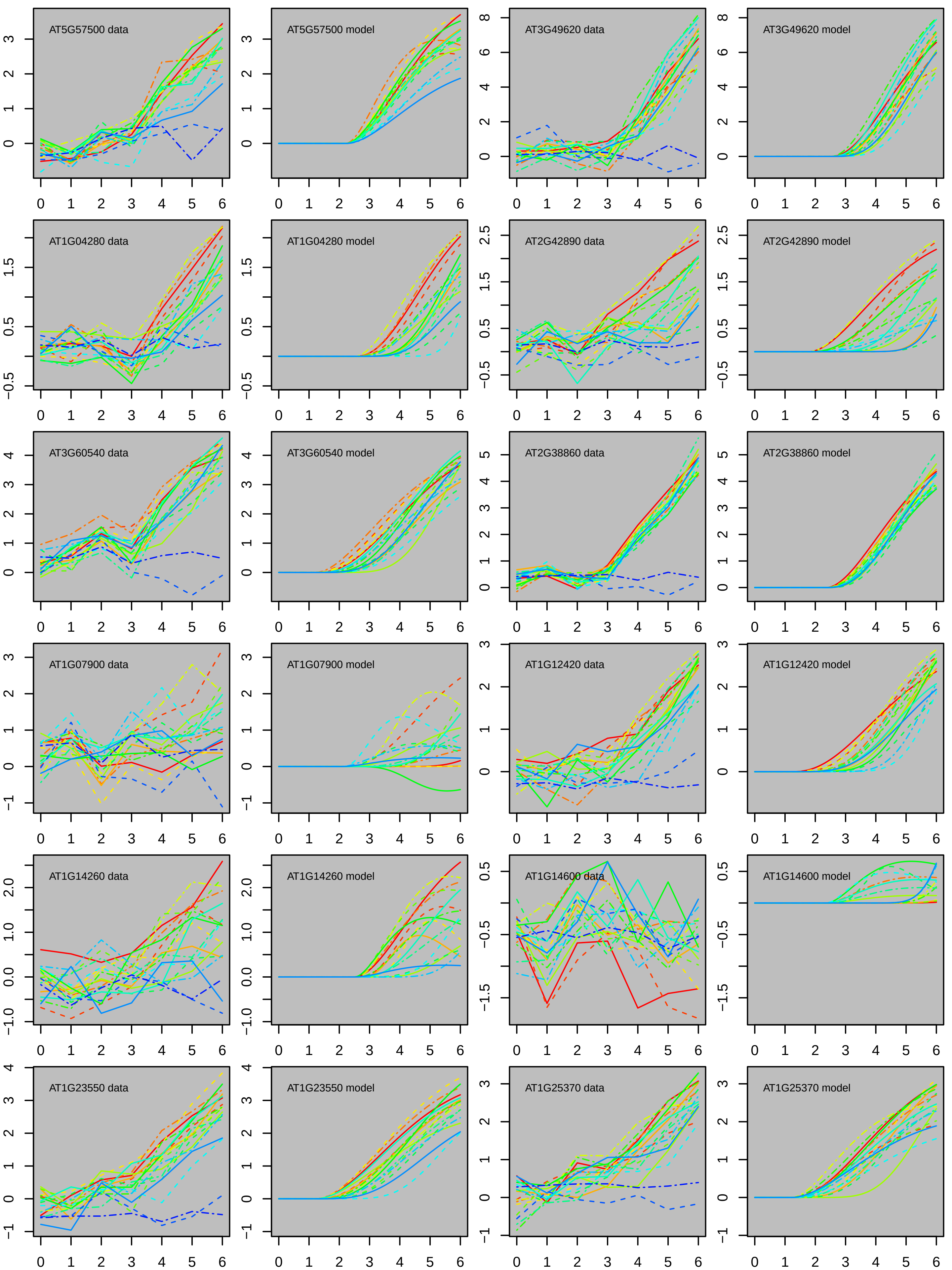

AT5G57500 data
AT5G57500 model
AT3G49620 data
AT3G49620 model
AT1G04280 data
AT1G04280 model
AT2G42890 data
AT2G42890 model
AT3G60540 data
AT3G60540 model
AT2G38860 data
AT2G38860 model
AT1G07900 data
AT1G07900 model
AT1G12420 data
AT1G12420 model
AT1G14260 data
AT1G14260 model
AT1G14600 data
AT1G14600 model
AT1G23550 data
AT1G23550 model
AT1G25370 data
AT1G25370 model
0 1 2 3 4 5 6

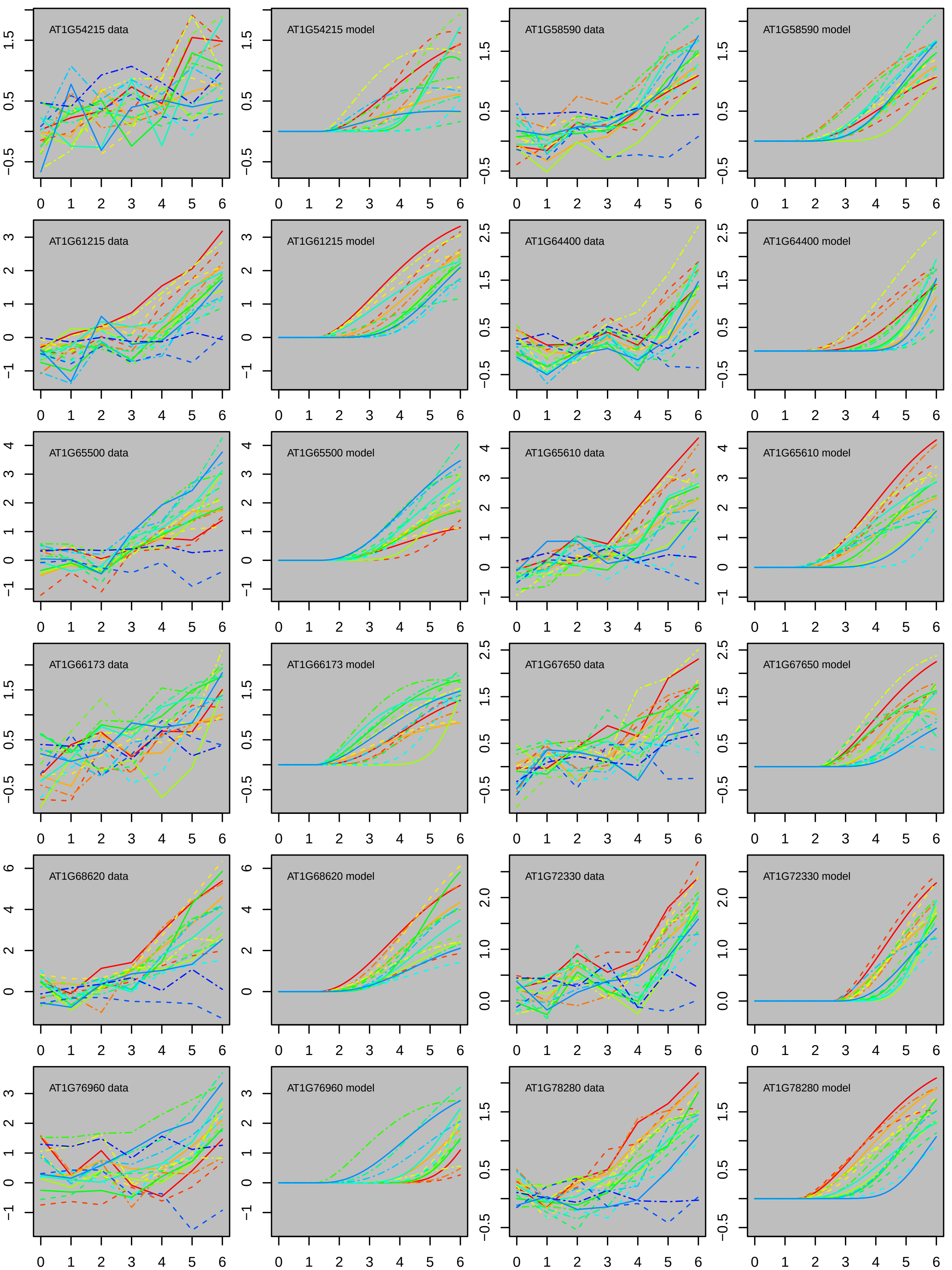
AT1G54215 data
AT1G54215 model
AT1G58590 data
AT1G58590 model
AT1G61215 data
AT1G61215 model
AT1G64400 data
AT1G64400 model
AT1G65500 data
AT1G65500 model
AT1G65610 data
AT1G65610 model
AT1G66173 data
AT1G66173 model
AT1G67650 data
AT1G67650 model
AT1G68620 data
AT1G68620 model
AT1G72330 data
AT1G72330 model
AT1G76960 data
AT1G76960 model
AT1G78280 data
AT1G78280 model

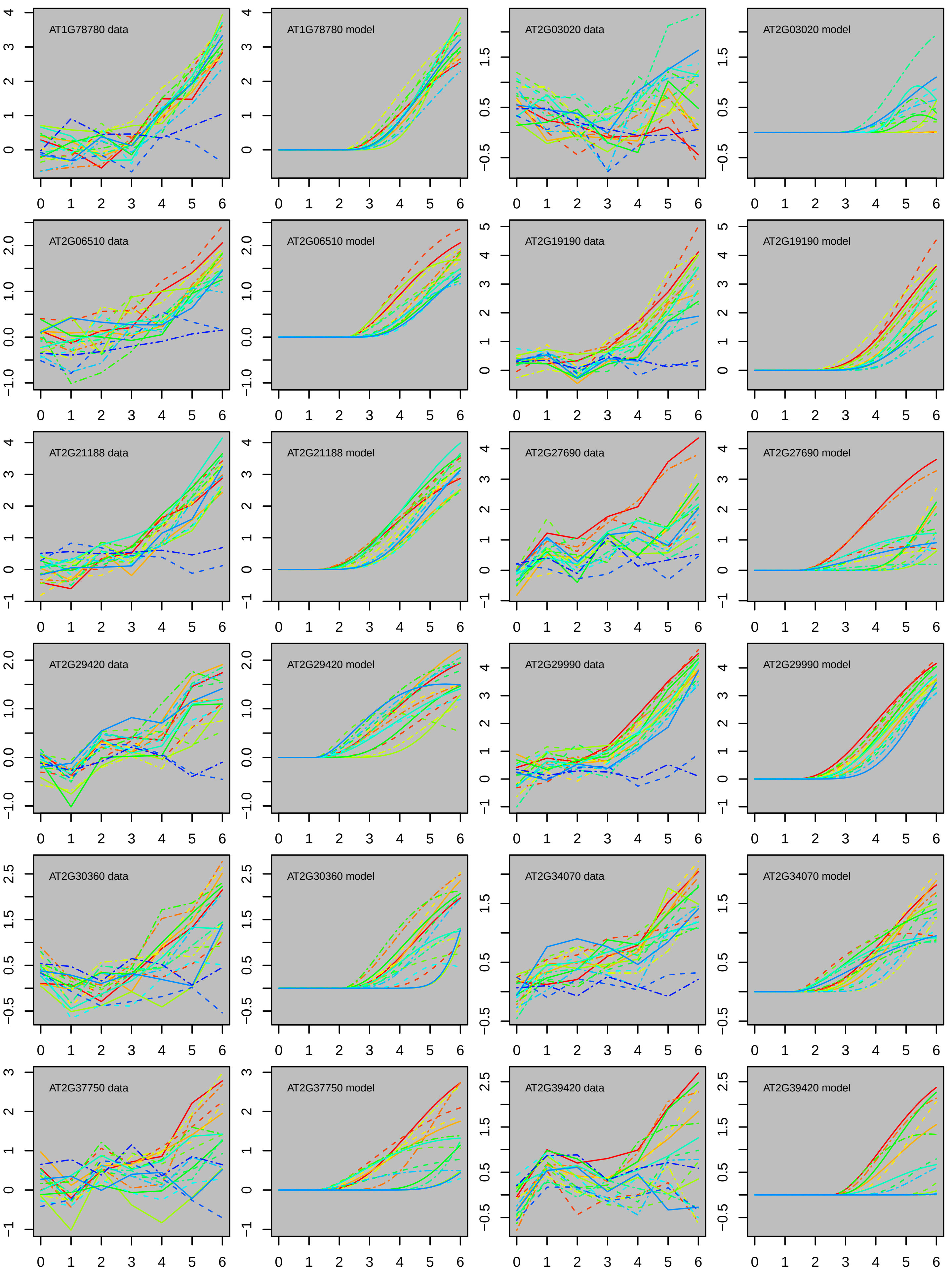

AT1G78780 data
AT1G78780 model
AT2G03020 data
AT2G03020 model
AT2G06510 data
AT2G06510 model
AT2G19190 data
AT2G19190 model
AT2G21188 data
AT2G21188 model
AT2G27690 data
AT2G27690 model
AT2G29420 data
AT2G29420 model
AT2G29990 data
AT2G29990 model
AT2G30360 data
AT2G30360 model
AT2G34070 data
AT2G34070 model
AT2G37750 data
AT2G37750 model
AT2G39420 data
AT2G39420 model

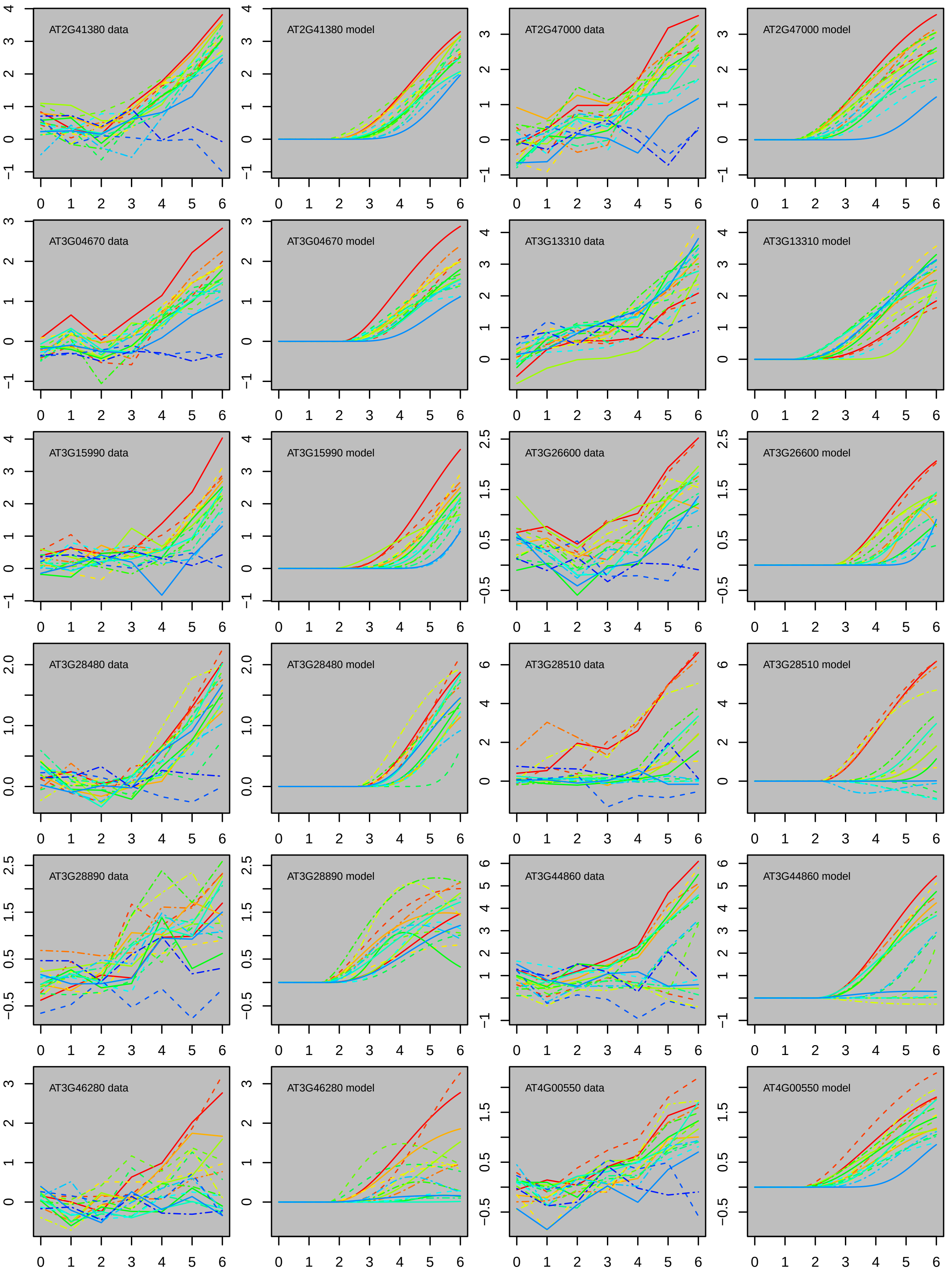
AT2G41380 data
AT2G41380 model
AT2G47000 data
AT2G47000 model
AT3G04670 data
AT3G04670 model
AT3G13310 data
AT3G13310 model
AT3G15990 data
AT3G15990 model
AT3G26600 data
AT3G26600 model
AT3G28480 data
AT3G28480 model
AT3G28510 data
AT3G28510 model
AT3G28890 data
AT3G28890 model
AT3G44860 data
AT3G44860 model
AT3G46280 data
AT3G46280 model
AT4G00550 data
AT4G00550 model

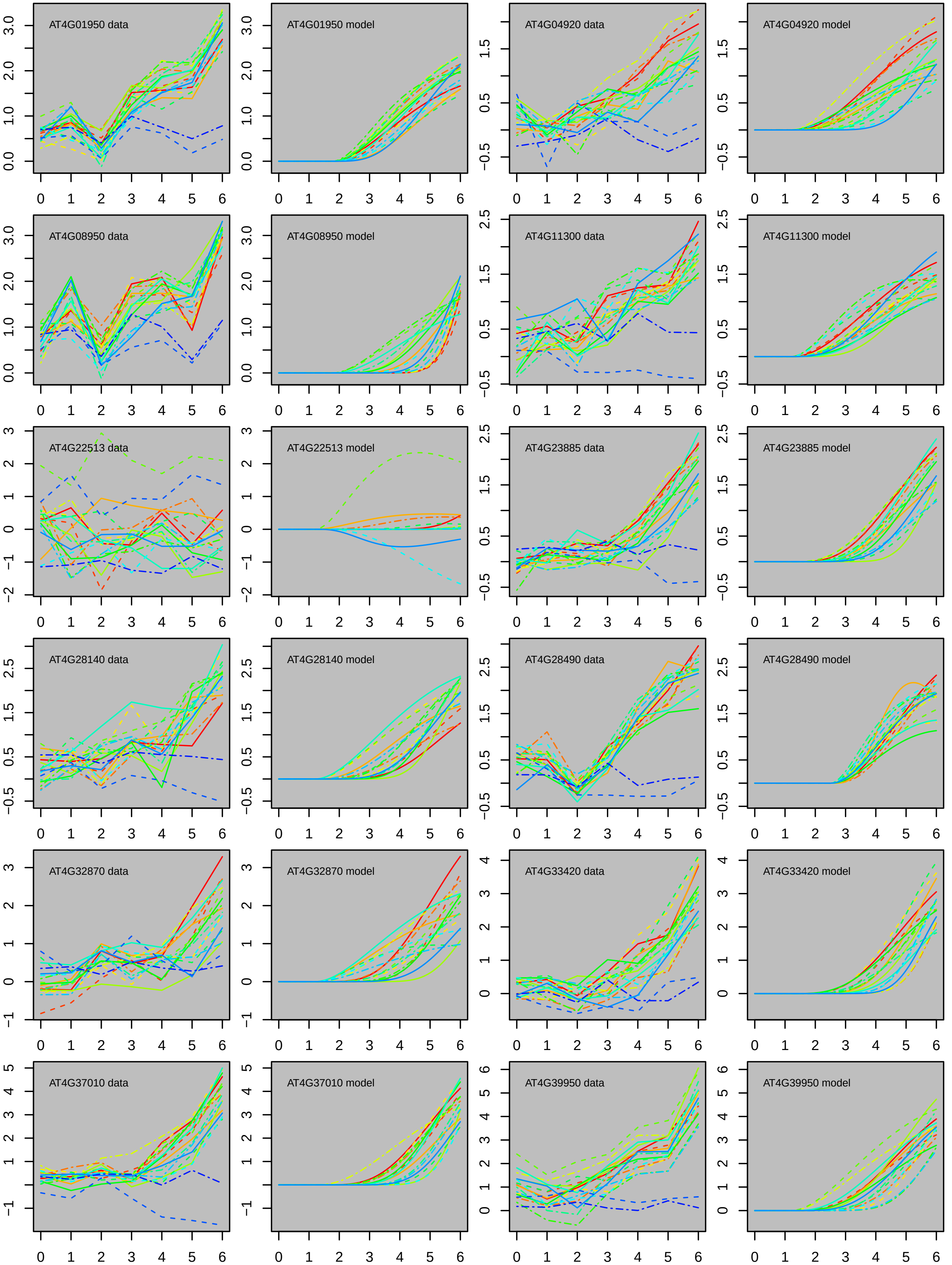

AT4G01950 data
AT4G01950 model
AT4G04920 data
AT4G04920 model
AT4G08950 data
AT4G08950 model
AT4G11300 data
AT4G11300 model
AT4G22513 data
AT4G22513 model
AT4G23885 data
AT4G23885 model
AT4G28140 data
AT4G28140 model
AT4G28490 data
AT4G28490 model
AT4G32870 data
AT4G32870 model
AT4G33420 data
AT4G33420 model
AT4G37010 data
AT4G37010 model
AT4G39950 data
AT4G39950 model

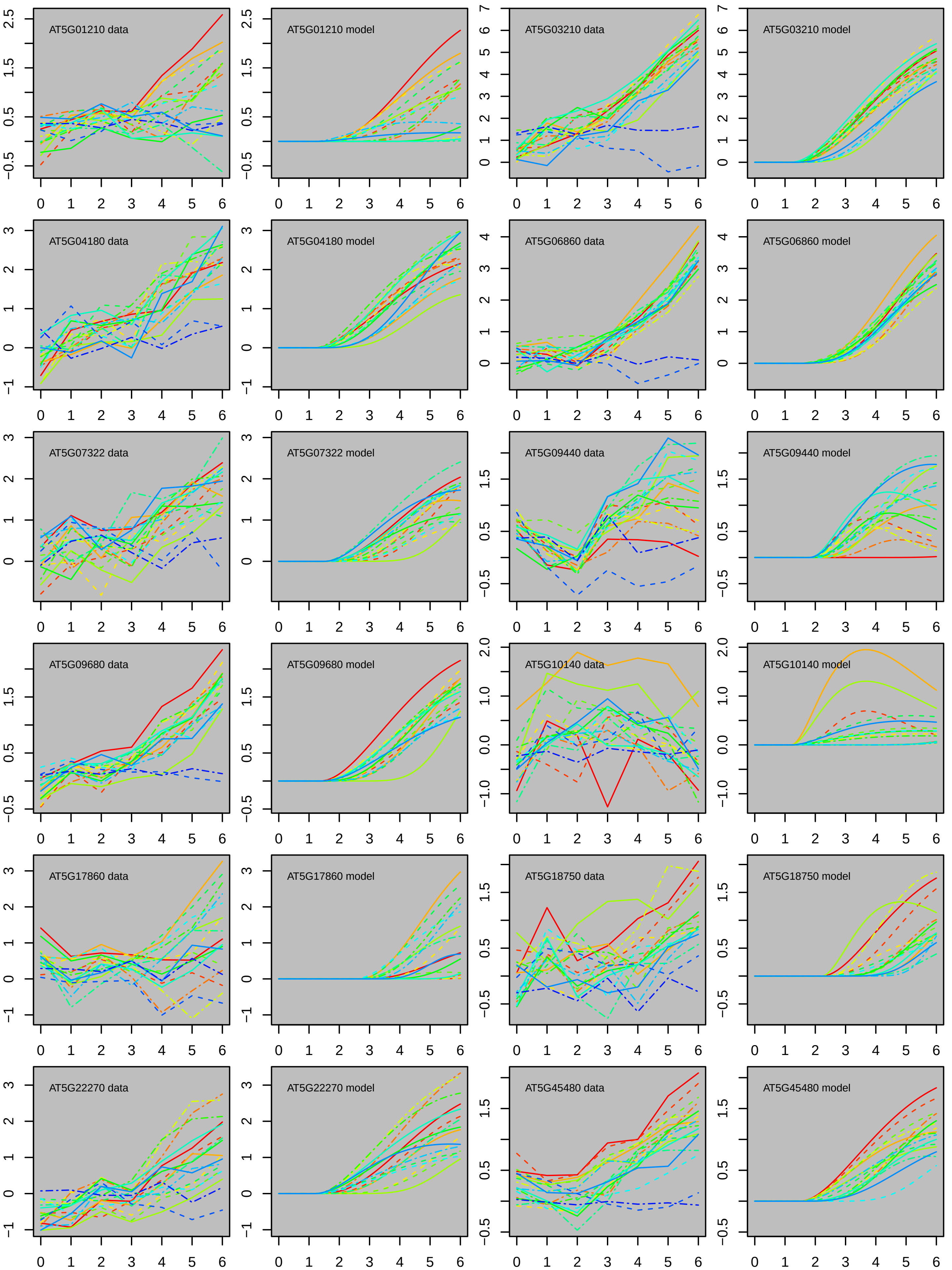

AT5G01210 data
AT5G01210 model
AT5G03210 data
AT5G03210 model
AT5G04180 data
AT5G04180 model
AT5G06860 data
AT5G06860 model
AT5G07322 data
AT5G07322 model
AT5G09440 data
AT5G09440 model
AT5G09680 data
AT5G09680 model
AT5G10140 data
AT5G10140 model
AT5G17860 data
AT5G17860 model
AT5G18750 data
AT5G18750 model
AT5G22270 data
AT5G22270 model
AT5G45480 data
AT5G45480 model

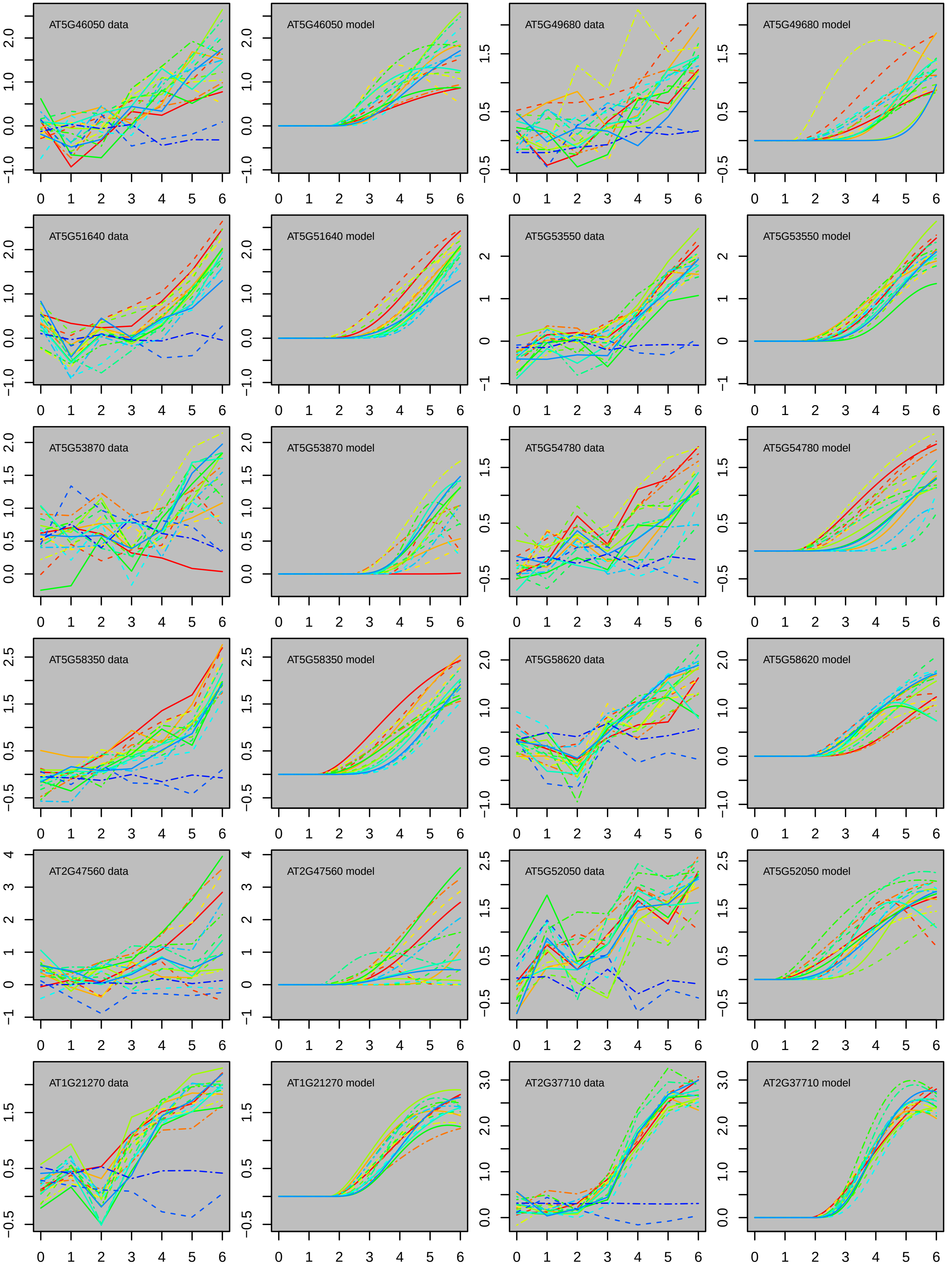
AT5G46050 data
AT5G46050 model
AT5G49680 data
AT5G49680 model
AT5G51640 data
AT5G51640 model
AT5G53550 data
AT5G53550 model
AT5G53870 data
AT5G53870 model
AT5G54780 data
AT5G54780 model
AT5G58350 data
AT5G58350 model
AT5G58620 data
AT5G58620 model
AT2G47560 data
AT2G47560 model
AT5G52050 data
AT5G52050 model
AT1G21270 data
AT1G21270 model
AT2G37710 data
AT2G37710 model

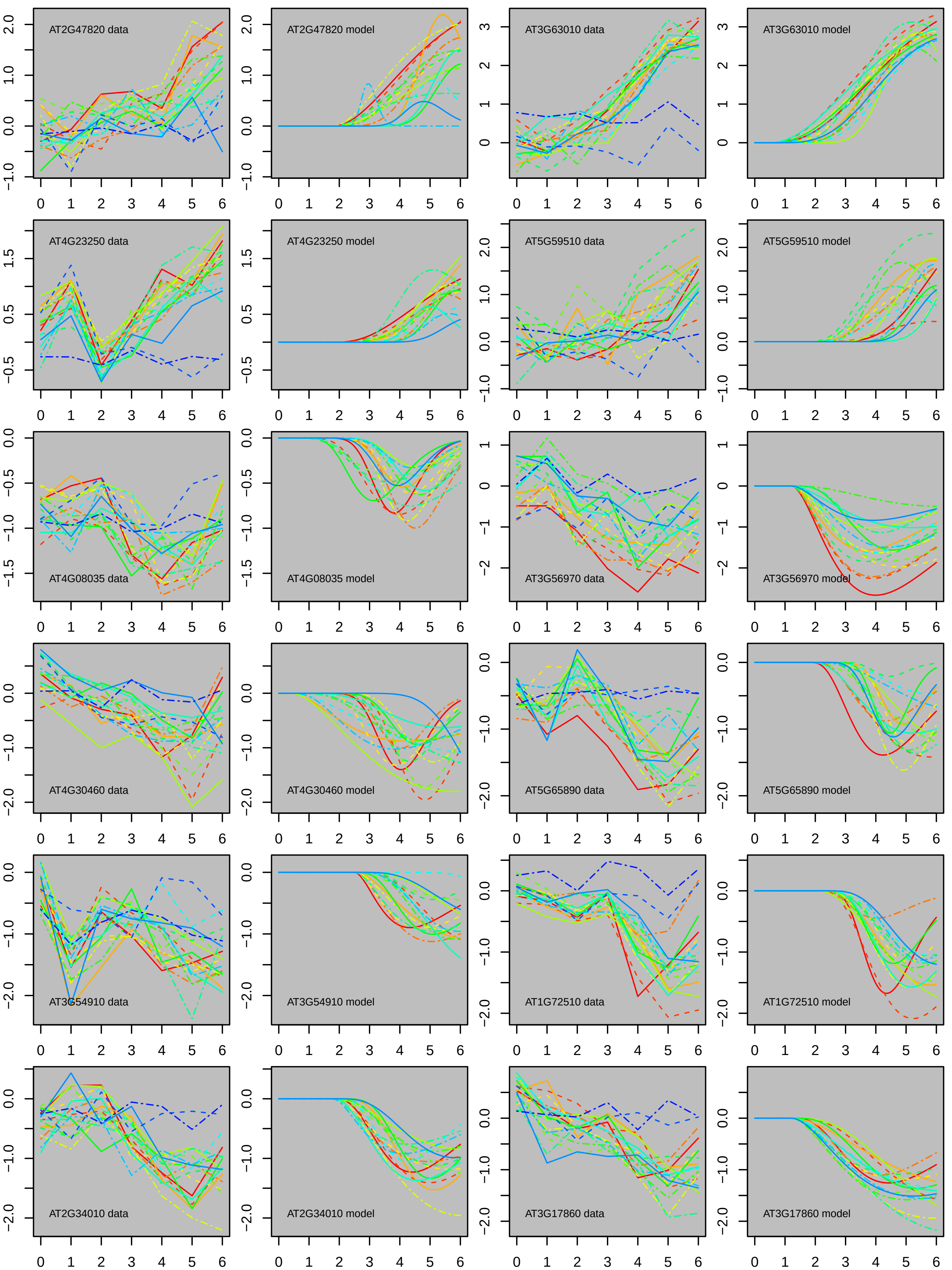
AT2G47820 data
AT2G47820 model
AT3G63010 data
AT3G63010 model
AT4G23250 data
AT4G23250 model
AT5G59510 data
AT5G59510 model
AT4G08035 data
AT4G08035 model
AT3G56970 data
AT3G56970 model
AT4G30460 data
AT4G30460 model
AT5G65890 data
AT5G65890 model
AT3G54910 data
AT3G54910 model
AT1G72510 data
AT1G72510 model
AT2G34010 data
AT2G34010 model
AT3G17860 data
AT3G17860 model

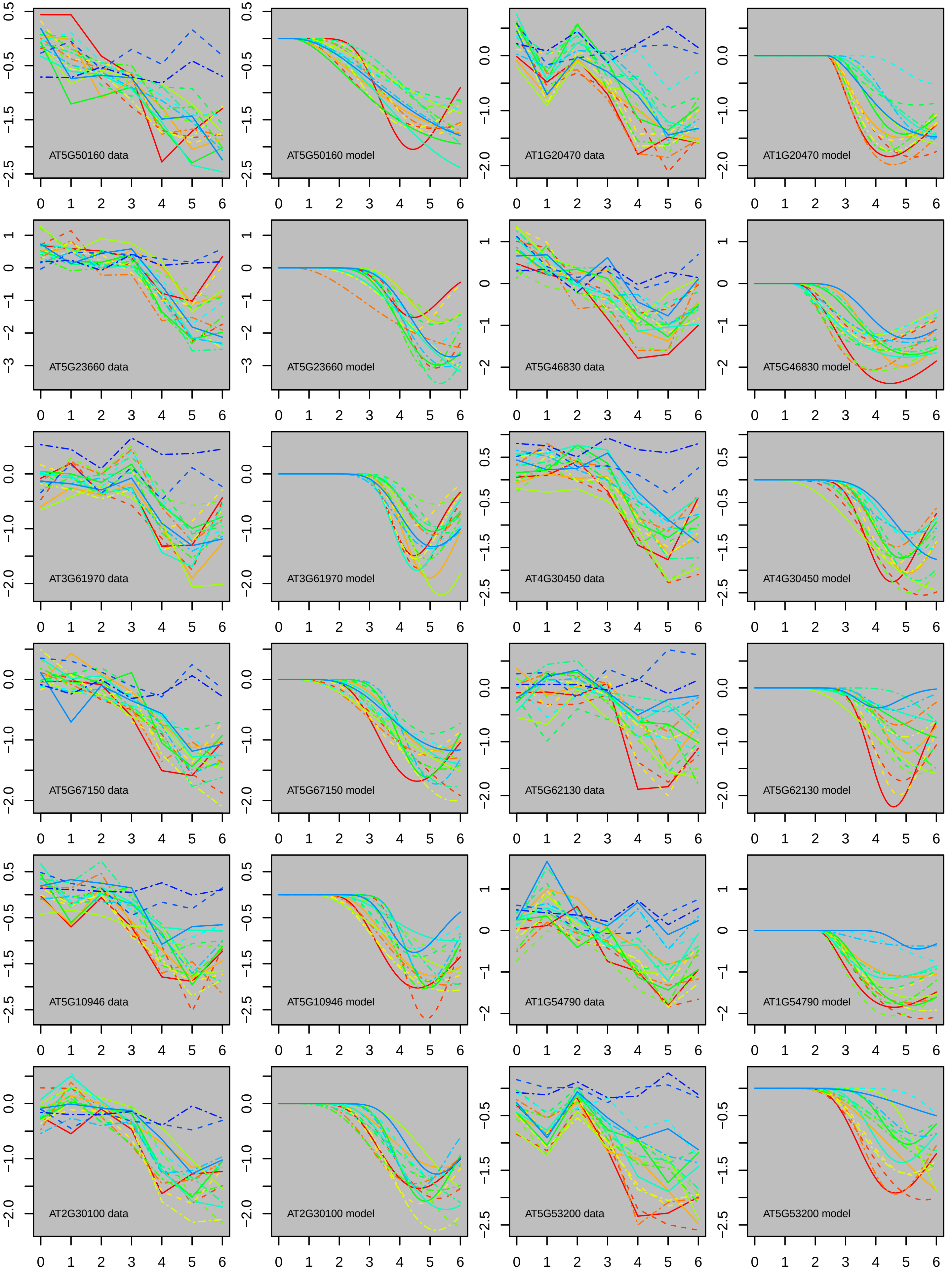

AT5G50160 data
AT5G50160 model
AT1G20470 data
AT1G20470 model
AT5G23660 data
AT5G23660 model
AT5G46830 data
AT5G46830 model
AT3G61970 data
AT3G61970 model
AT4G30450 data
AT4G30450 model
AT5G67150 data
AT5G67150 model
AT5G62130 data
AT5G62130 model
AT5G10946 data
AT5G10946 model
AT1G54790 data
AT1G54790 model
AT2G30100 data
AT2G30100 model
AT5G53200 data
AT5G53200 model

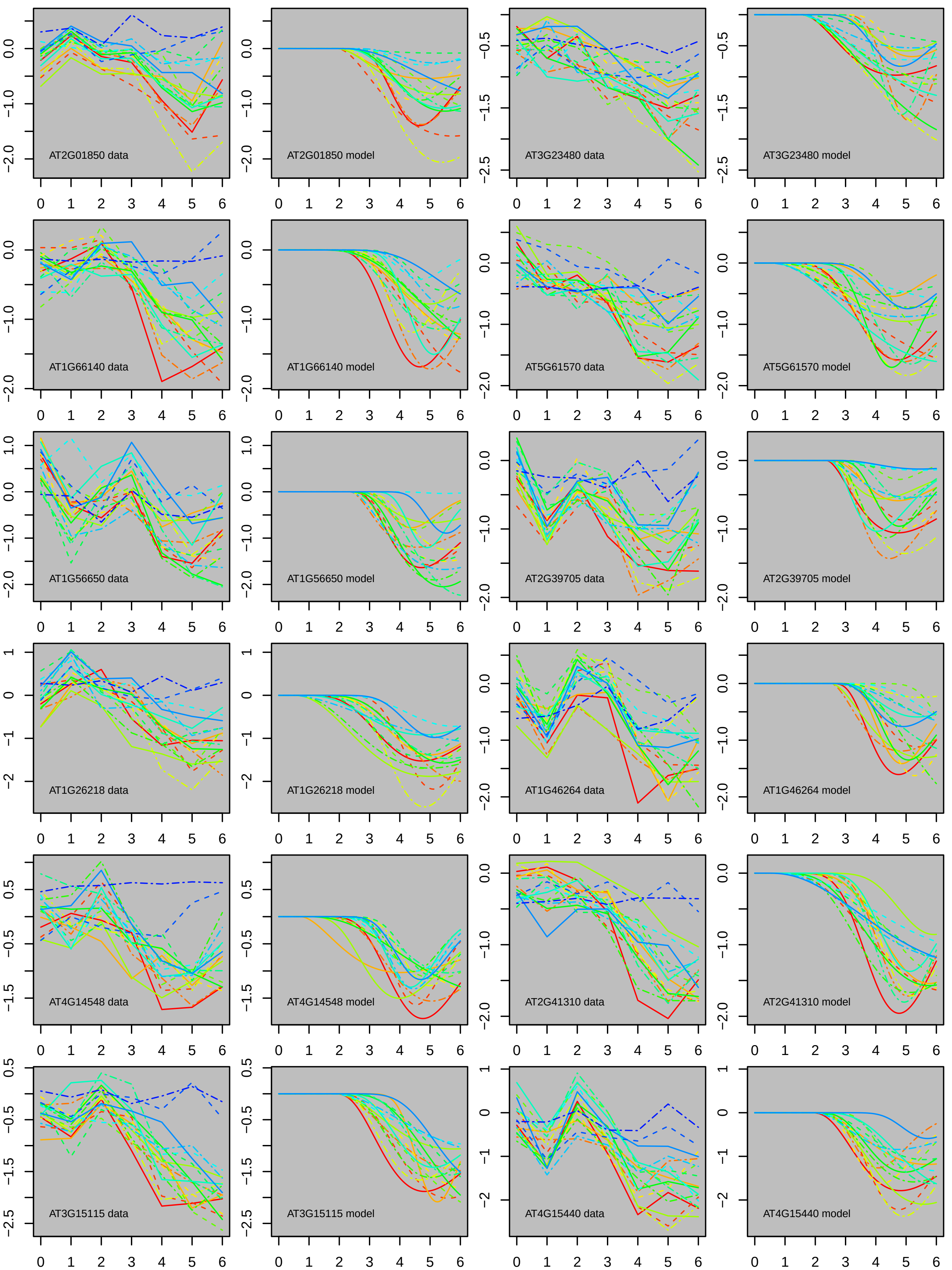

AT2G01850 data
AT2G01850 model
AT3G23480 data
AT3G23480 model
AT1G66140 data
AT1G66140 model
AT5G61570 data
AT5G61570 model
AT1G56650 data
AT1G56650 model
AT2G39705 data
AT2G39705 model
AT1G26218 data
AT1G26218 model
AT1G46264 data
AT1G46264 model
AT4G14548 data
AT4G14548 model
AT2G41310 data
AT2G41310 model
AT3G15115 data
AT3G15115 model
AT4G15440 data
AT4G15440 model

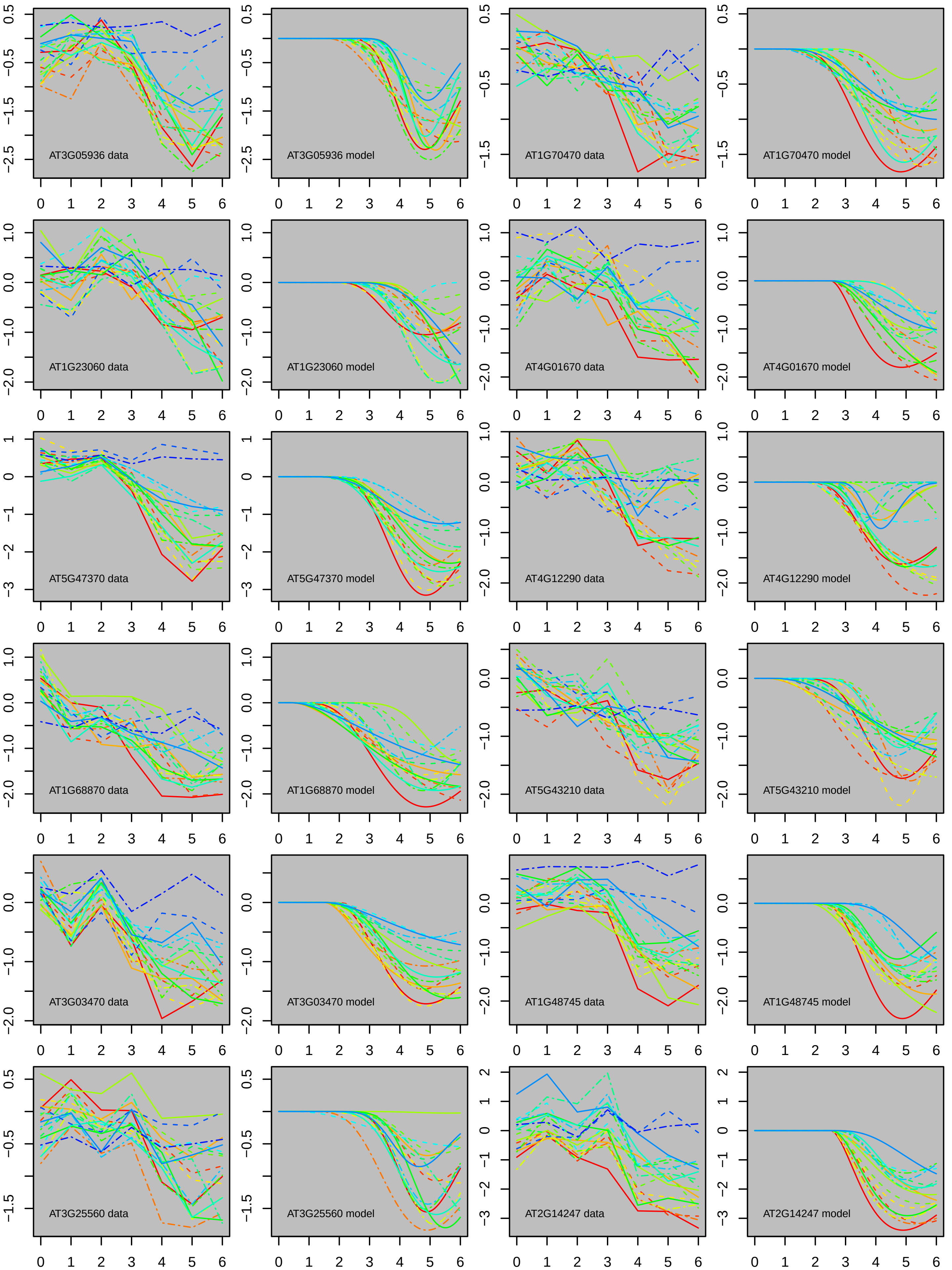
AT3G05936 data
AT3G05936 model
AT1G70470 data
AT1G70470 model
AT1G23060 data
AT1G23060 model
AT4G01670 data
AT4G01670 model
AT5G47370 data
AT5G47370 model
AT4G12290 data
AT4G12290 model
AT1G68870 data
AT1G68870 model
AT5G43210 data
AT5G43210 model
AT3G03470 data
AT3G03470 model
AT1G48745 data
AT1G48745 model
AT3G25560 data
AT3G25560 model
AT2G14247 data
AT2G14247 model

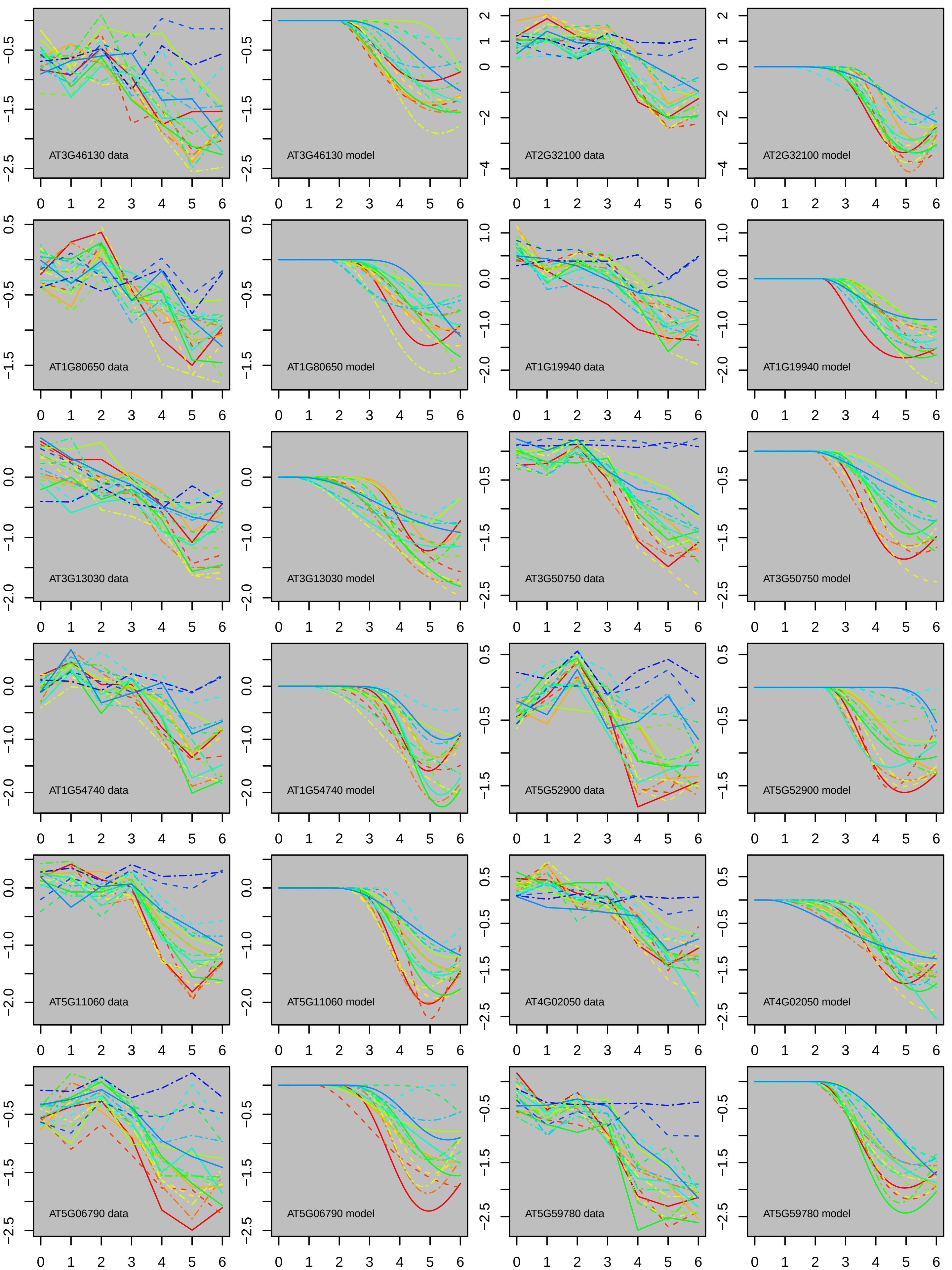
AT3G46130 data
AT3G46130 model
AT2G32100 data
AT2G32100 model
AT1G80650 data
AT1G80650 model
AT1G19940 data
AT1G19940 model
AT3G13030 data
AT3G13030 model
AT3G50750 data
AT3G50750 model
AT1G54740 data
AT1G54740 model
AT5G52900 data
AT5G52900 model
AT5G11060 data
AT5G11060 model
AT4G02050 data
AT4G02050 model
AT5G06790 data
AT5G06790 model
AT5G59780 data
AT5G59780 model

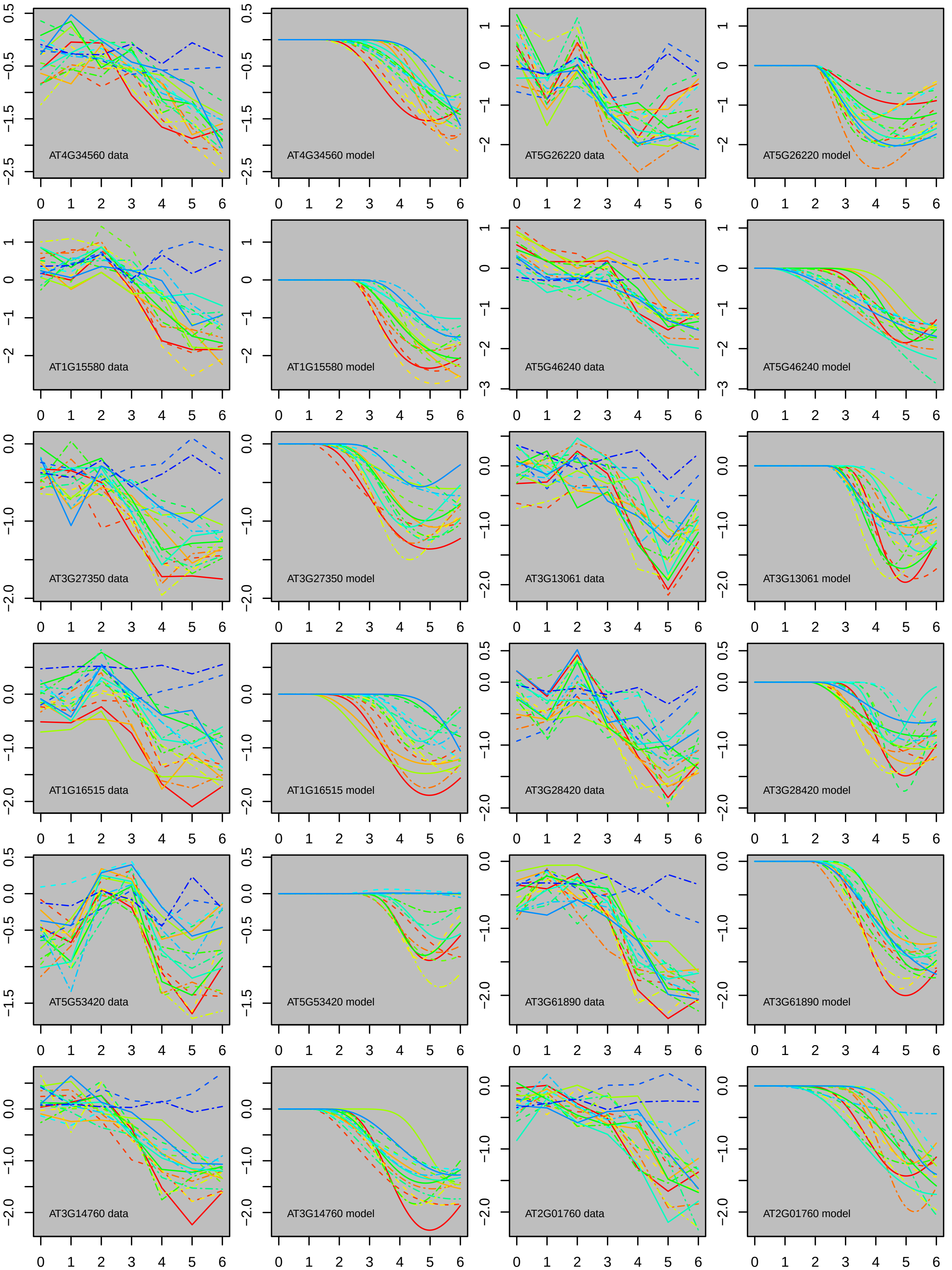

AT4G34560 data
AT4G34560 model
AT5G26220 data
AT5G26220 model
AT1G15580 data
AT1G15580 model
AT5G46240 data
AT5G46240 model
AT3G27350 data
AT3G27350 model
AT3G13061 data
AT3G13061 model
AT1G16515 data
AT1G16515 model
AT3G28420 data
AT3G28420 model
AT5G53420 data
AT5G53420 model
AT3G61890 data
AT3G61890 model
AT3G14760 data
AT3G14760 model
AT2G01760 data
AT2G01760 model

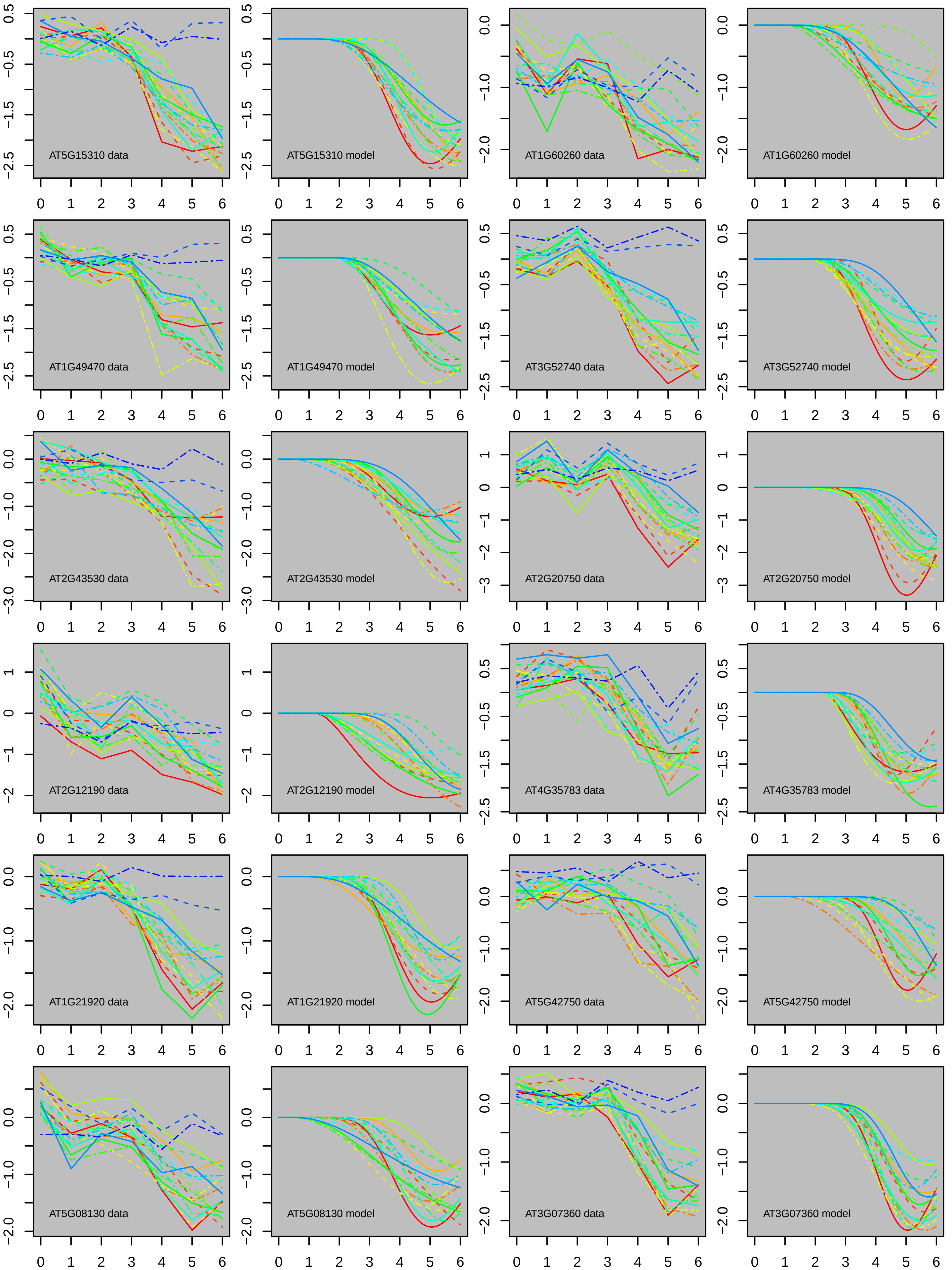

AT5G15310 data
AT5G15310 model
AT1G60260 data
AT1G60260 model
AT1G49470 data
AT1G49470 model
AT3G52740 data
AT3G52740 model
AT2G43530 data
AT2G43530 model
AT2G20750 data
AT2G20750 model
AT2G12190 data
AT2G12190 model
AT4G35783 data
AT4G35783 model
AT1G21920 data
AT1G21920 model
AT5G42750 data
AT5G42750 model
AT5G08130 data
AT5G08130 model
AT3G07360 data
AT3G07360 model

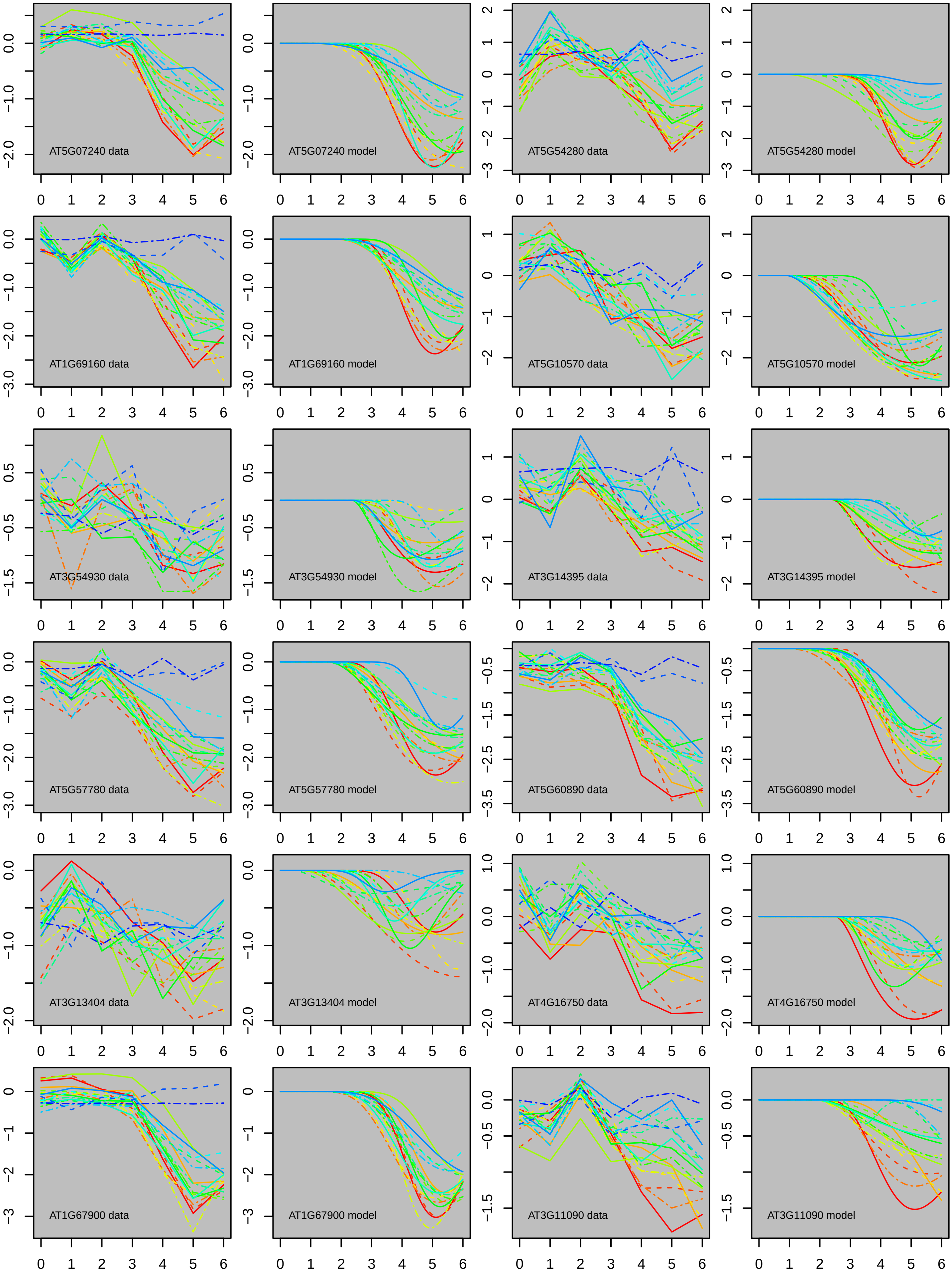

AT5G07240 data
AT5G07240 model
AT5G54280 data
AT5G54280 model
AT1G69160 data
AT1G69160 model
AT5G10570 data
AT5G10570 model
AT3G54930 data
AT3G54930 model
AT3G14395 data
AT3G14395 model
AT5G57780 data
AT5G57780 model
AT5G60890 data
AT5G60890 model
AT3G13404 data
AT3G13404 model
AT4G16750 data
AT4G16750 model
AT1G67900 data
AT1G67900 model
AT3G11090 data
AT3G11090 model

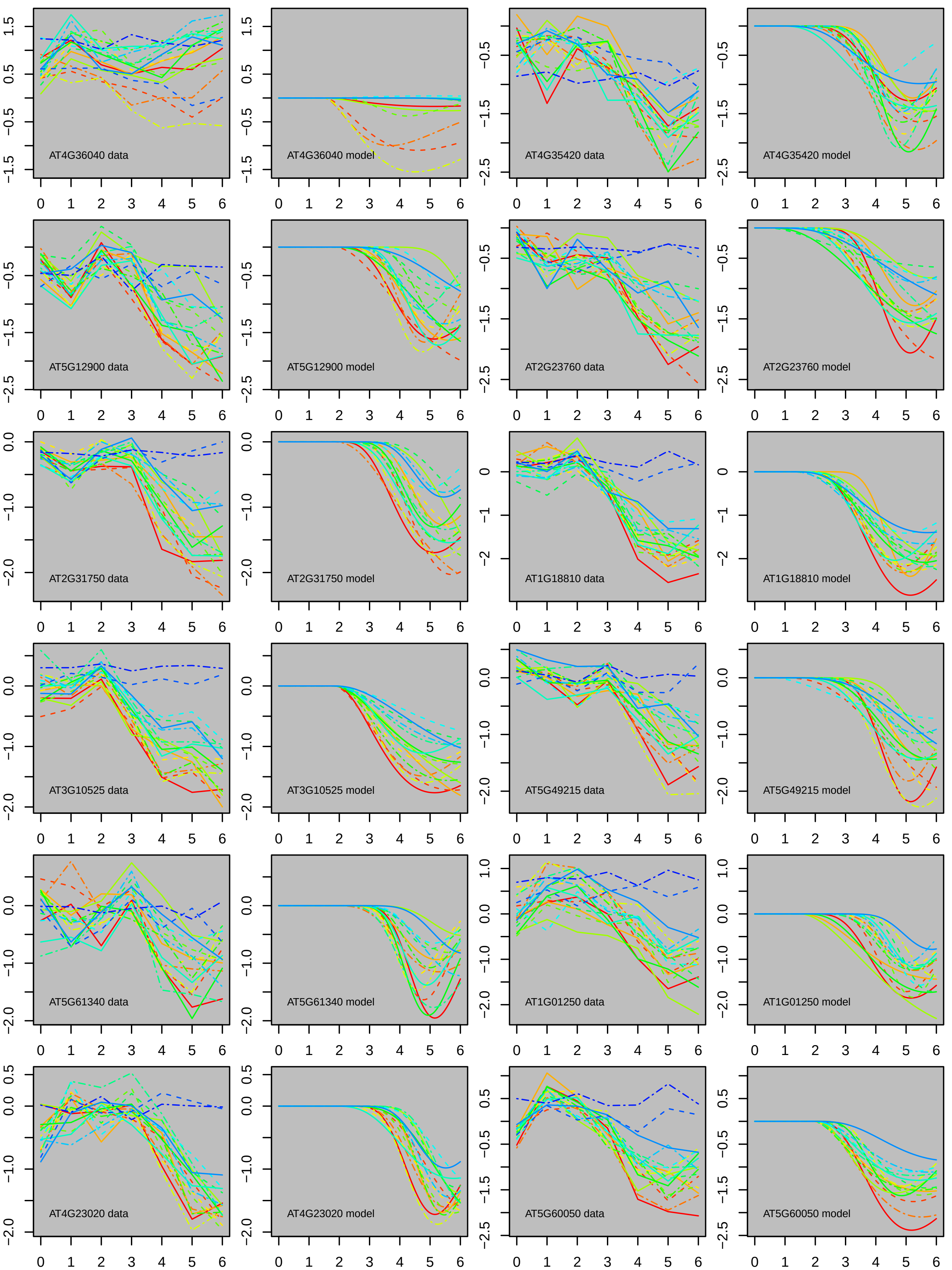

AT4G36040 data
AT4G36040 model
AT4G35420 data
AT4G35420 model
AT5G12900 data
AT5G12900 model
AT2G23760 data
AT2G23760 model
AT2G31750 data
AT2G31750 model
AT1G18810 data
AT1G18810 model
AT3G10525 data
AT3G10525 model
AT5G49215 data
AT5G49215 model
AT5G61340 data
AT5G61340 model
AT1G01250 data
AT1G01250 model
AT4G23020 data
AT4G23020 model
AT5G60050 data
AT5G60050 model

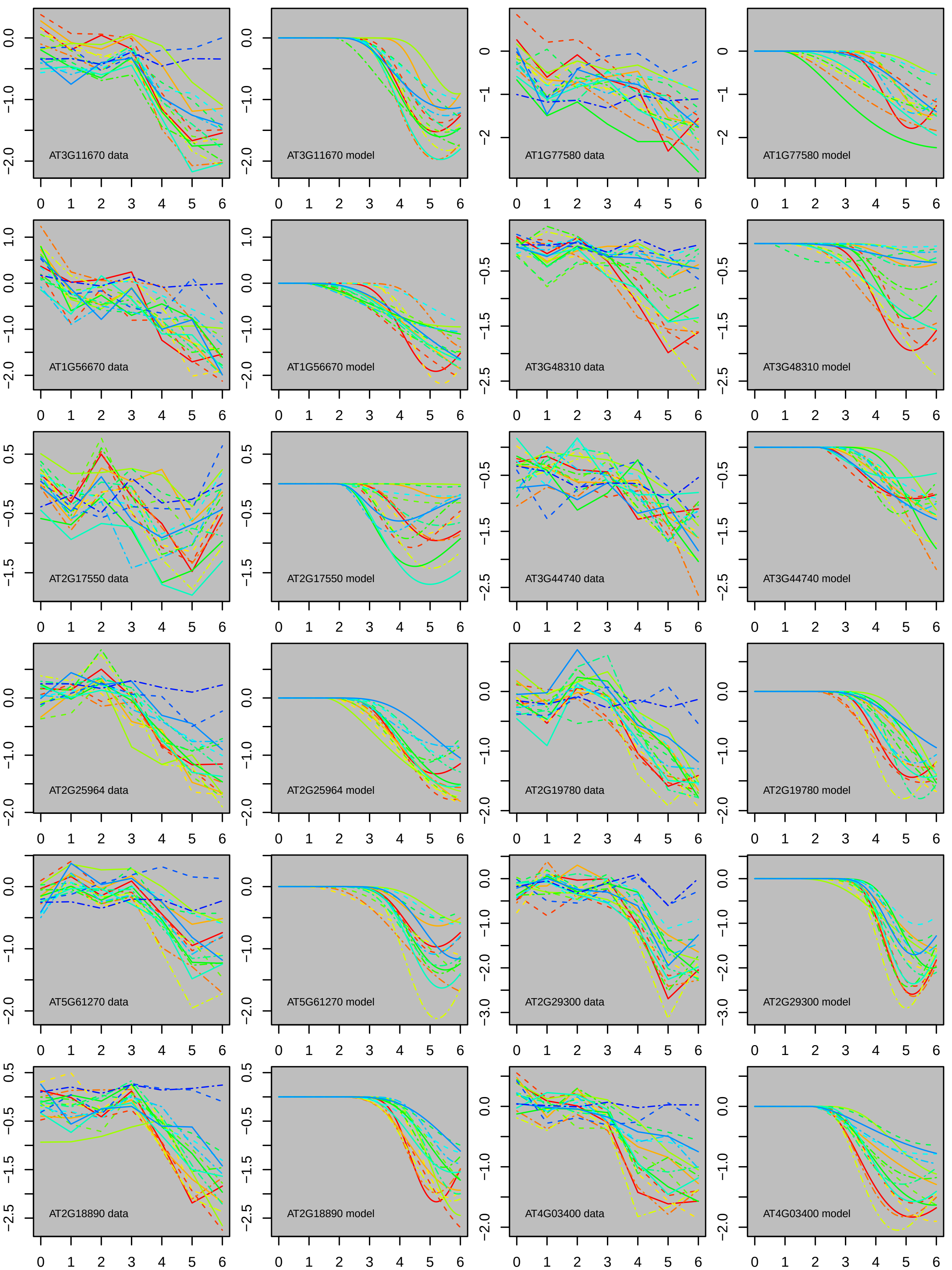

AT3G11670 data
AT3G11670 model
AT1G77580 data
AT1G77580 model
AT1G56670 data
AT1G56670 model
AT3G48310 data
AT3G48310 model
AT2G17550 data
AT2G17550 model
AT3G44740 data
AT3G44740 model
AT2G25964 data
AT2G25964 model
AT2G19780 data
AT2G19780 model
AT5G61270 data
AT5G61270 model
AT2G29300 data
AT2G29300 model
AT2G18890 data
AT2G18890 model
AT4G03400 data
AT4G03400 model

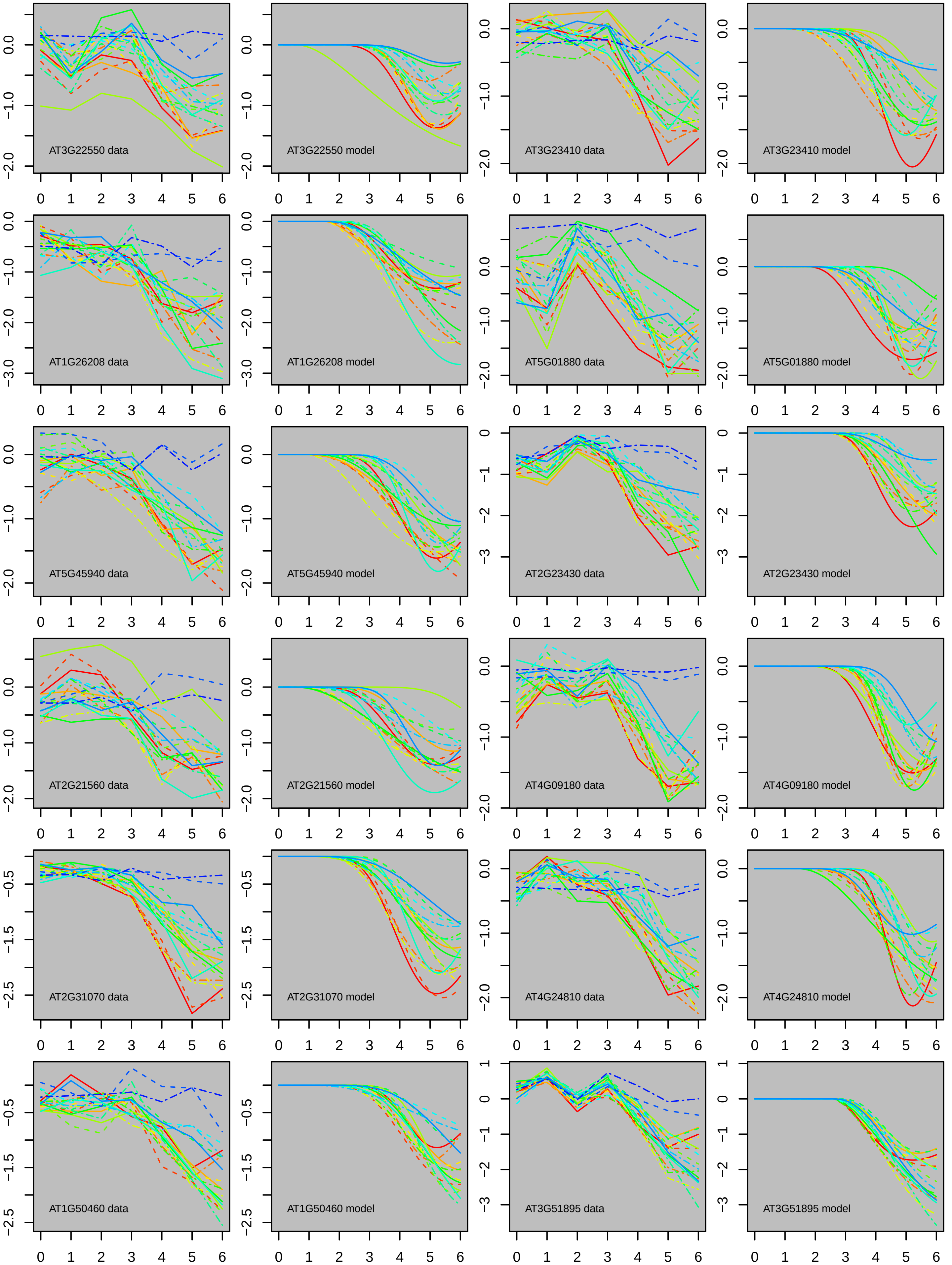
AT3G22550 data
AT3G22550 model
AT3G23410 data
AT3G23410 model
AT1G26208 data
AT1G26208 model
AT5G01880 data
AT5G01880 model
AT5G45940 data
AT5G45940 model
AT2G23430 data
AT2G23430 model
AT2G21560 data
AT2G21560 model
AT4G09180 data
AT4G09180 model
AT2G31070 data
AT2G31070 model
AT4G24810 data
AT4G24810 model
AT1G50460 data
AT1G50460 model
AT3G51895 data
AT3G51895 model

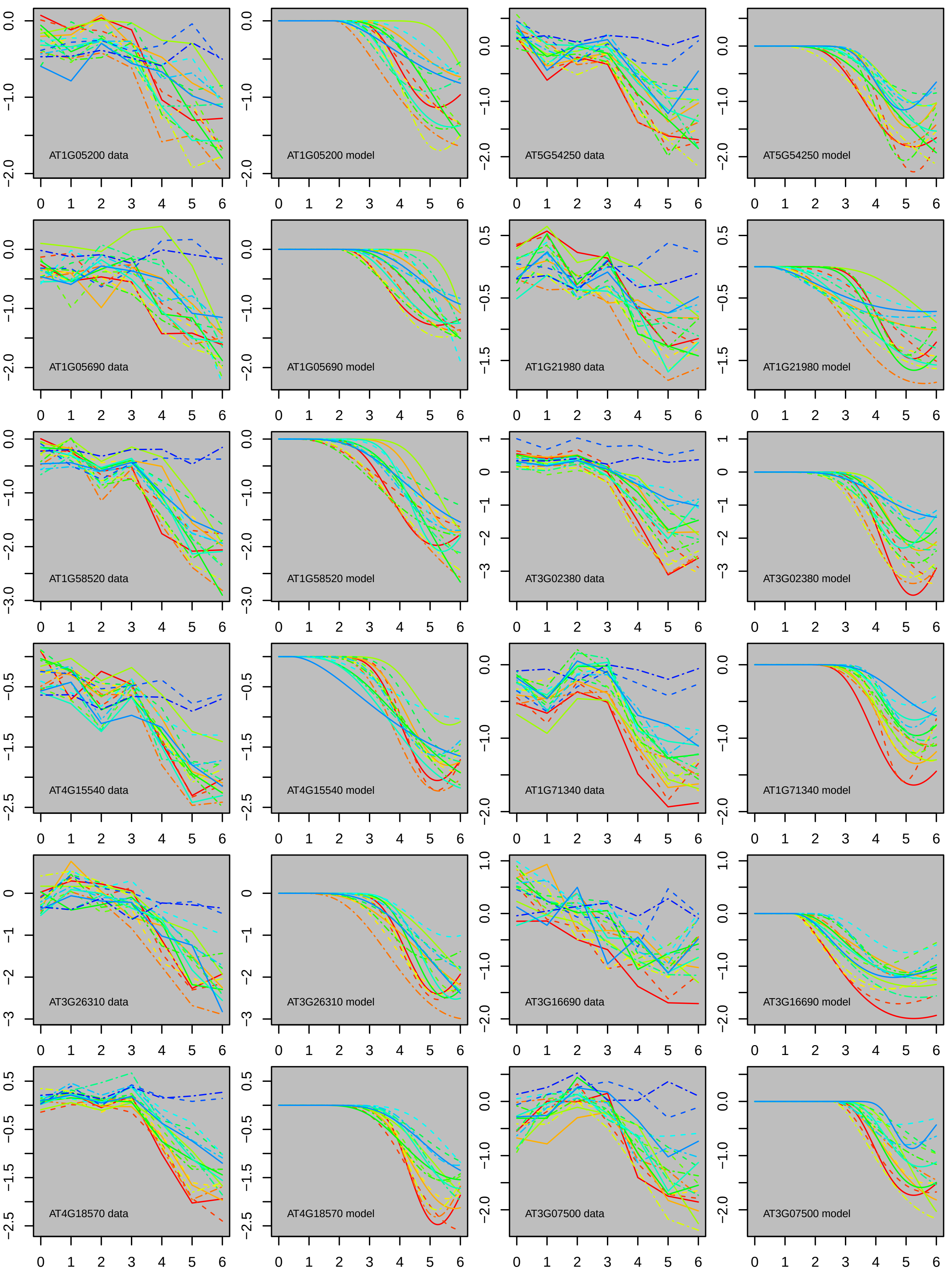
AT1G05200 data
AT1G05200 model
AT5G54250 data
AT5G54250 model
AT1G05690 data
AT1G05690 model
AT1G21980 data
AT1G21980 model
AT1G58520 data
AT1G58520 model
AT3G02380 data
AT3G02380 model
AT4G15540 data
AT4G15540 model
AT1G71340 data
AT1G71340 model
AT3G26310 data
AT3G26310 model
AT3G16690 data
AT3G16690 model
AT4G18570 data
AT4G18570 model
AT3G07500 data
AT3G07500 model

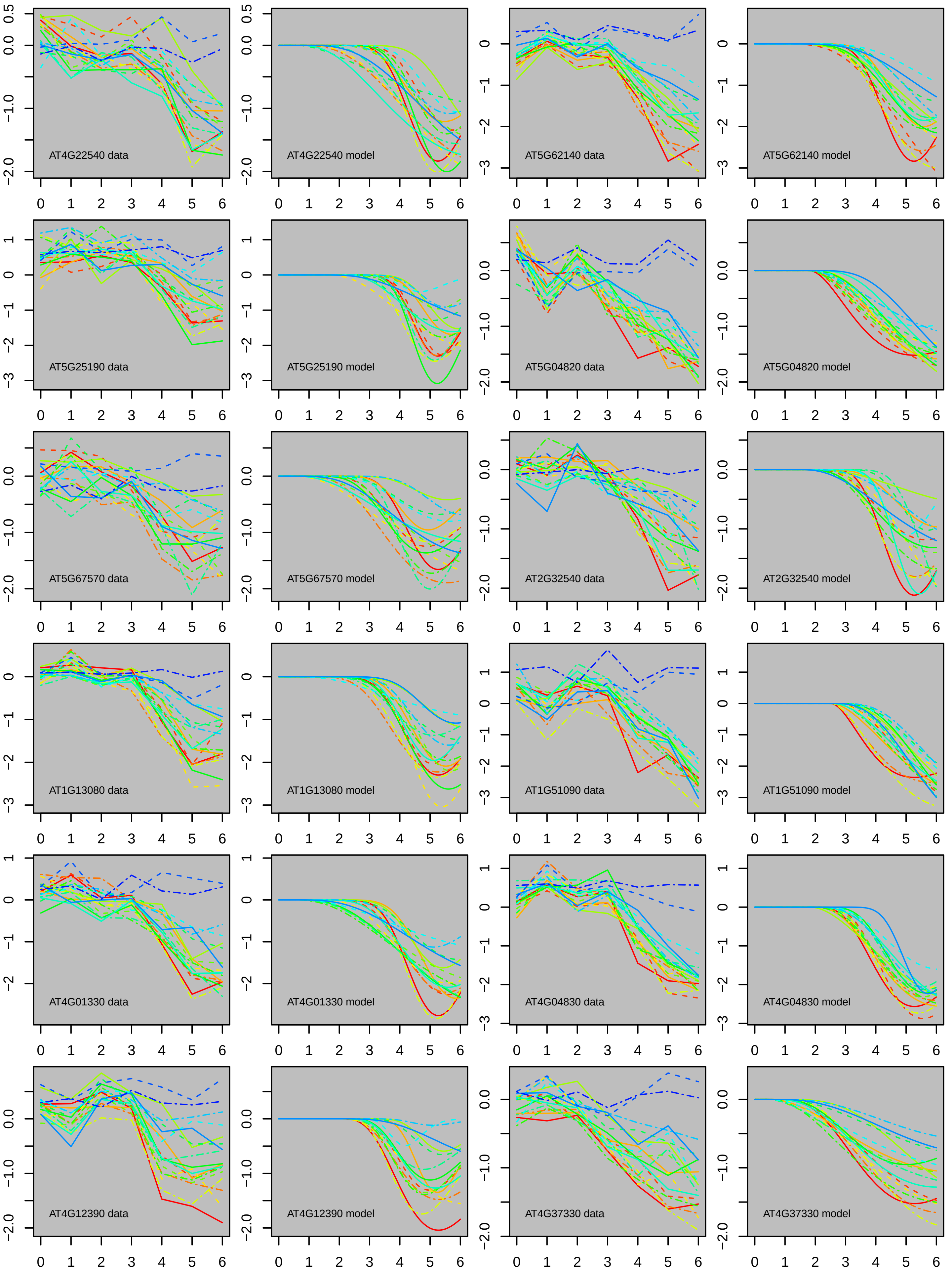
AT4G22540 data
AT4G22540 model
AT5G62140 data
AT5G62140 model
AT5G25190 data
AT5G25190 model
AT5G04820 data
AT5G04820 model
AT5G67570 data
AT5G67570 model
AT2G32540 data
AT2G32540 model
AT1G13080 data
AT1G13080 model
AT1G51090 data
AT1G51090 model
AT4G01330 data
AT4G01330 model
AT4G04830 data
AT4G04830 model
AT4G12390 data
AT4G12390 model
AT4G37330 data
AT4G37330 model

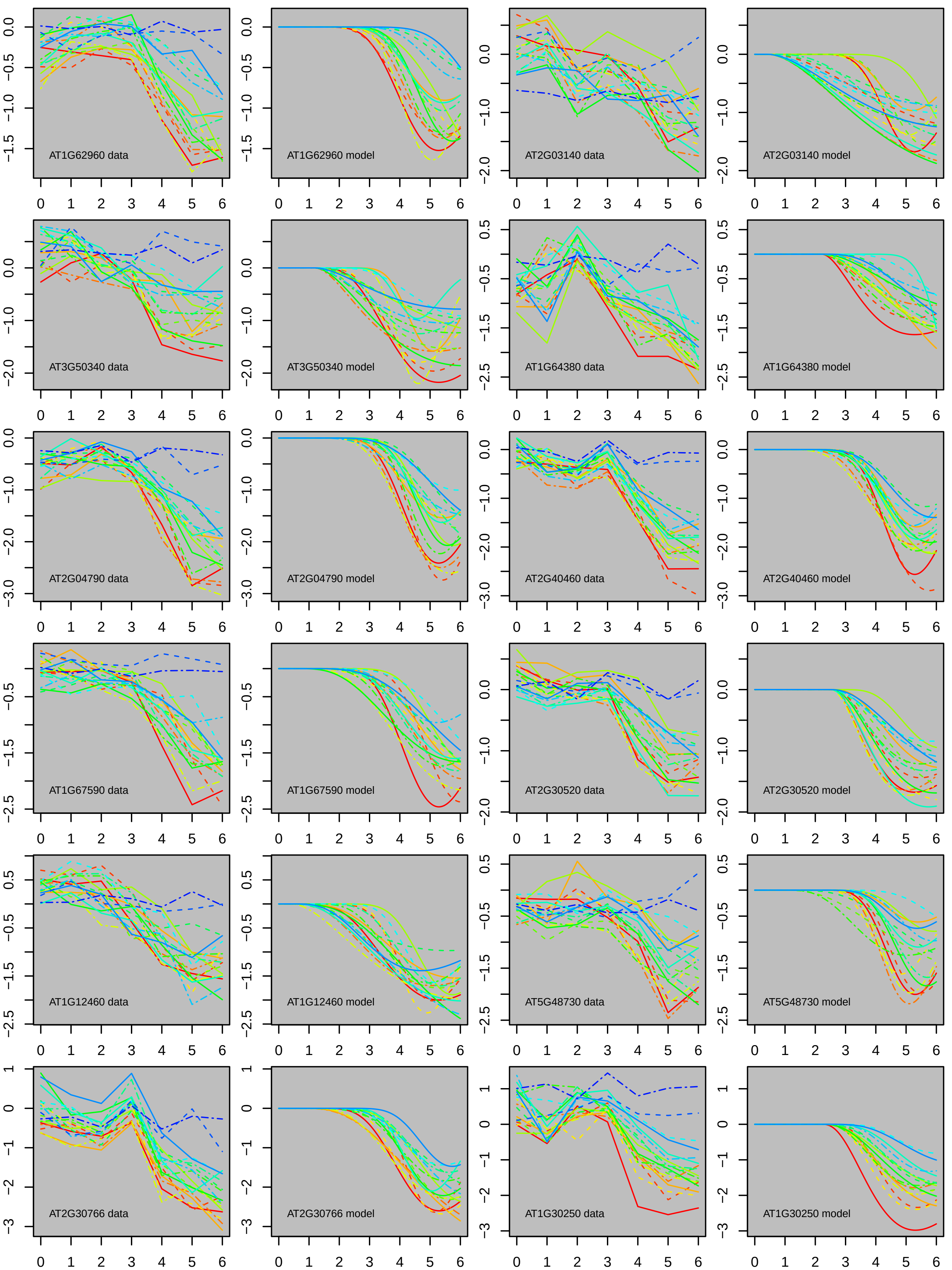

AT1G62960 data
AT1G62960 model
AT2G03140 data
AT2G03140 model
AT3G50340 data
AT3G50340 model
AT1G64380 data
AT1G64380 model
AT2G04790 data
AT2G04790 model
AT2G40460 data
AT2G40460 model
AT1G67590 data
AT1G67590 model
AT2G30520 data
AT2G30520 model
AT1G12460 data
AT1G12460 model
AT5G48730 data
AT5G48730 model
AT2G30766 data
AT2G30766 model
AT1G30250 data
AT1G30250 model

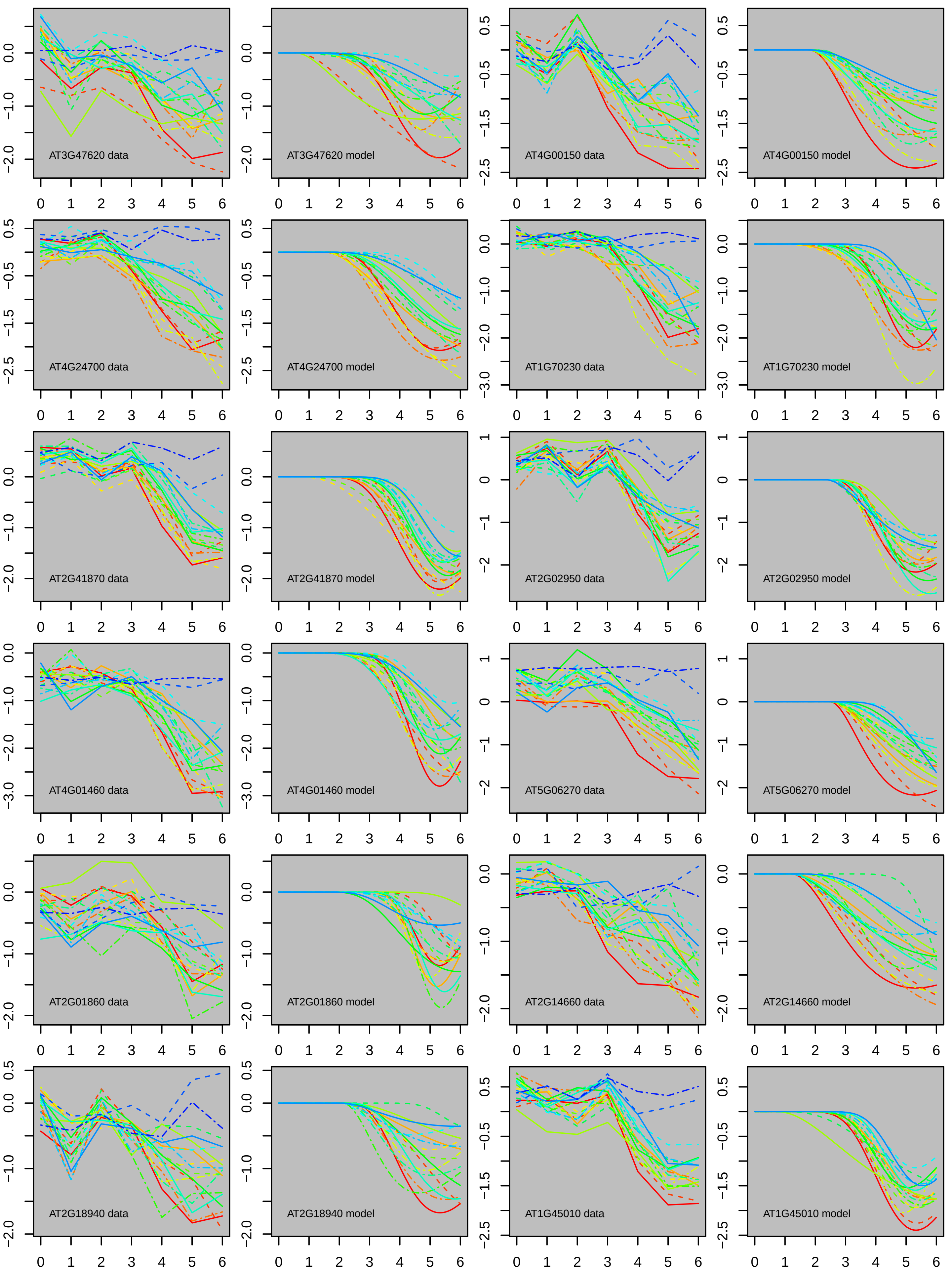

AT3G47620 data
AT3G47620 model
AT4G00150 data
AT4G00150 model
AT4G24700 data
AT4G24700 model
AT1G70230 data
AT1G70230 model
AT2G41870 data
AT2G41870 model
AT2G02950 data
AT2G02950 model
AT4G01460 data
AT4G01460 model
AT5G06270 data
AT5G06270 model
AT2G01860 data
AT2G01860 model
AT2G14660 data
AT2G14660 model
AT2G18940 data
AT2G18940 model
AT1G45010 data
AT1G45010 model

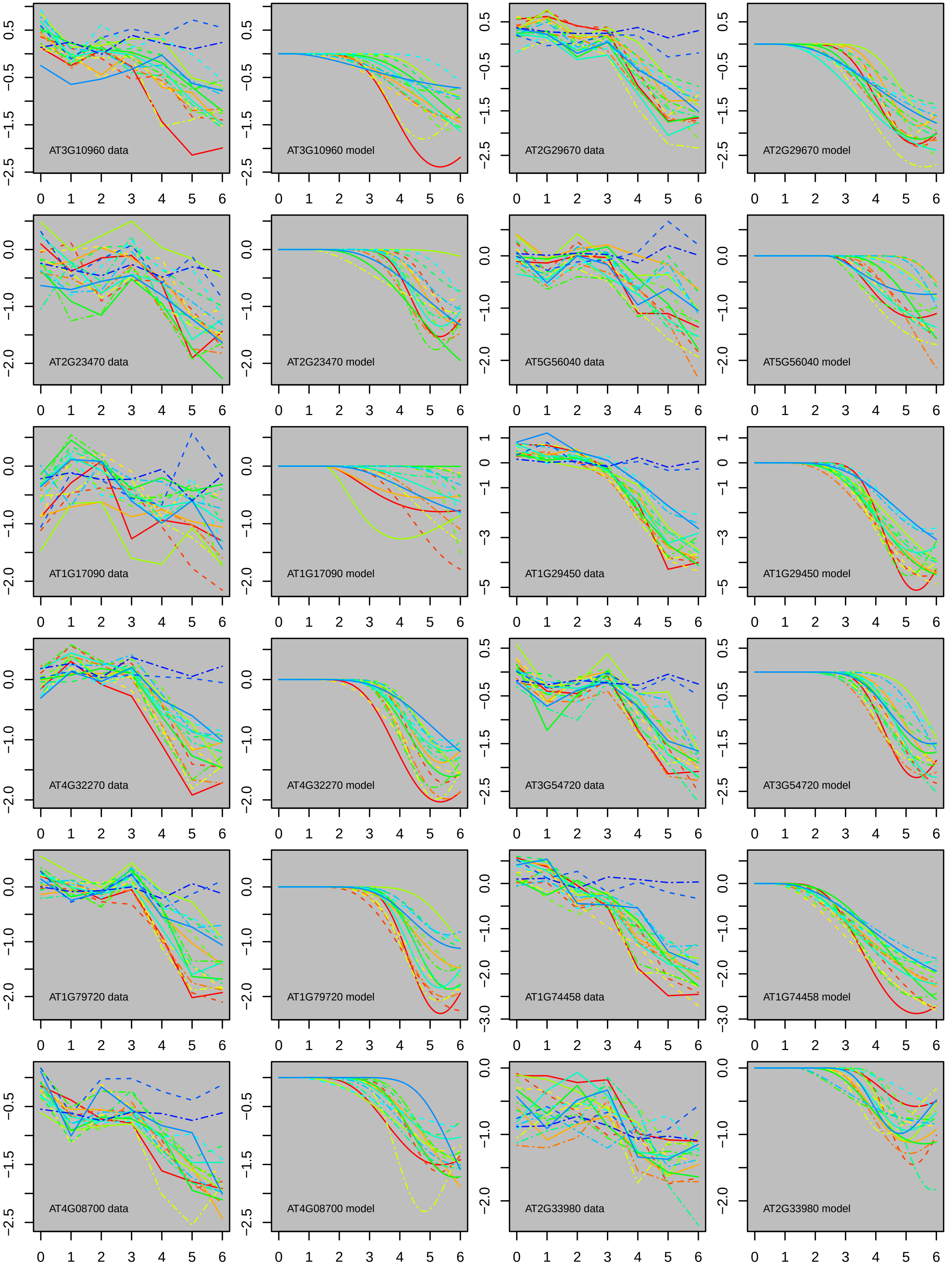
AT3G10960 data
AT3G10960 model
AT2G29670 data
AT2G29670 model
AT2G23470 data
AT2G23470 model
AT5G56040 data
AT5G56040 model
AT1G17090 data
AT1G17090 model
AT1G29450 data
AT1G29450 model
AT4G32270 data
AT4G32270 model
AT3G54720 data
AT3G54720 model
AT1G79720 data
AT1G79720 model
AT1G74458 data
AT1G74458 model
AT4G08700 data
AT4G08700 model
AT2G33980 data
AT2G33980 model

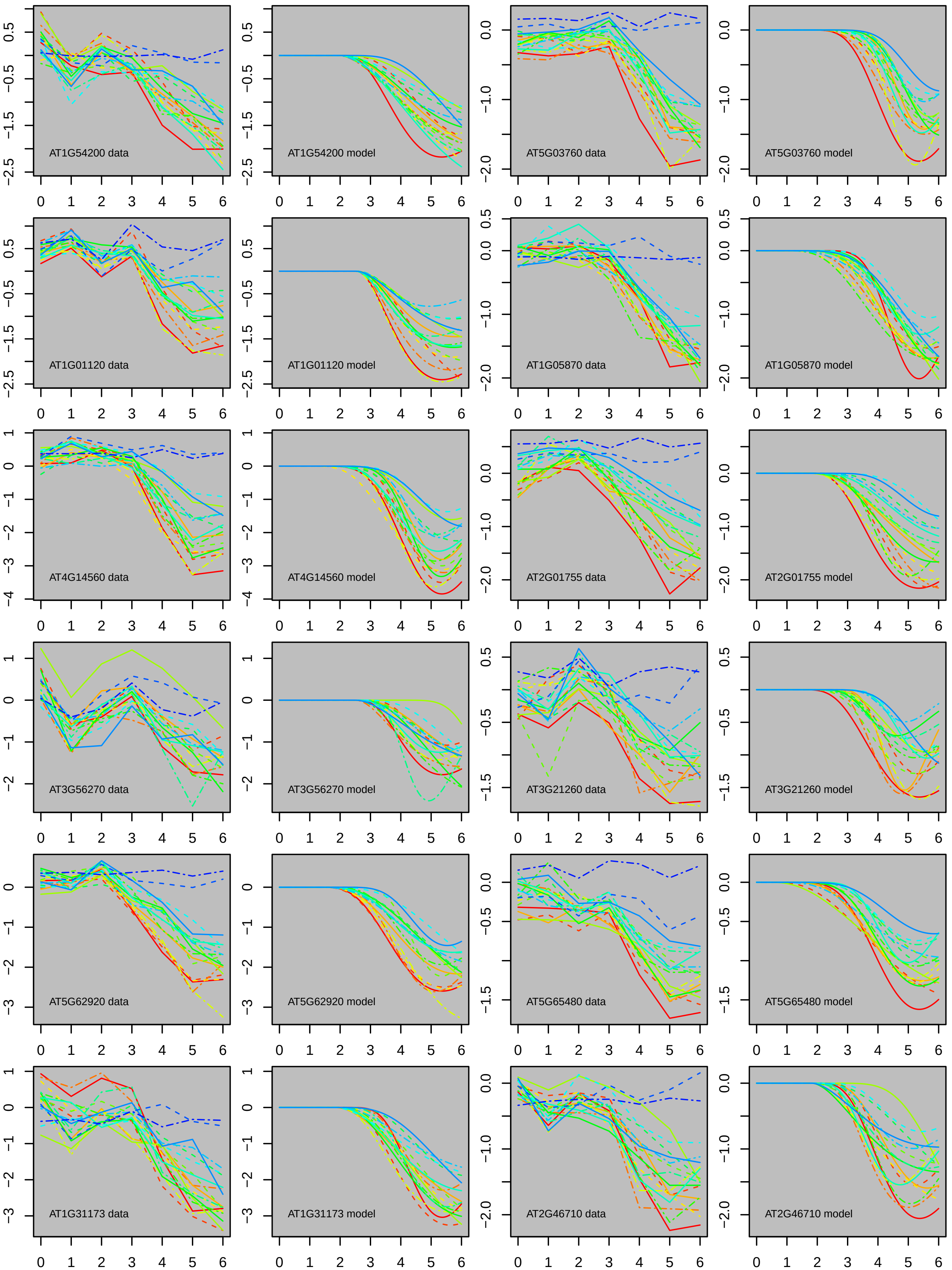
AT1G54200 data
AT1G54200 model
AT5G03760 data
AT5G03760 model
AT1G01120 data
AT1G01120 model
AT1G05870 data
AT1G05870 model
AT4G14560 data
AT4G14560 model
AT2G01755 data
AT2G01755 model
AT3G56270 data
AT3G56270 model
AT3G21260 data
AT3G21260 model
AT5G62920 data
AT5G62920 model
AT5G65480 data
AT5G65480 model
AT1G31173 data
AT1G31173 model
AT2G46710 data
AT2G46710 model

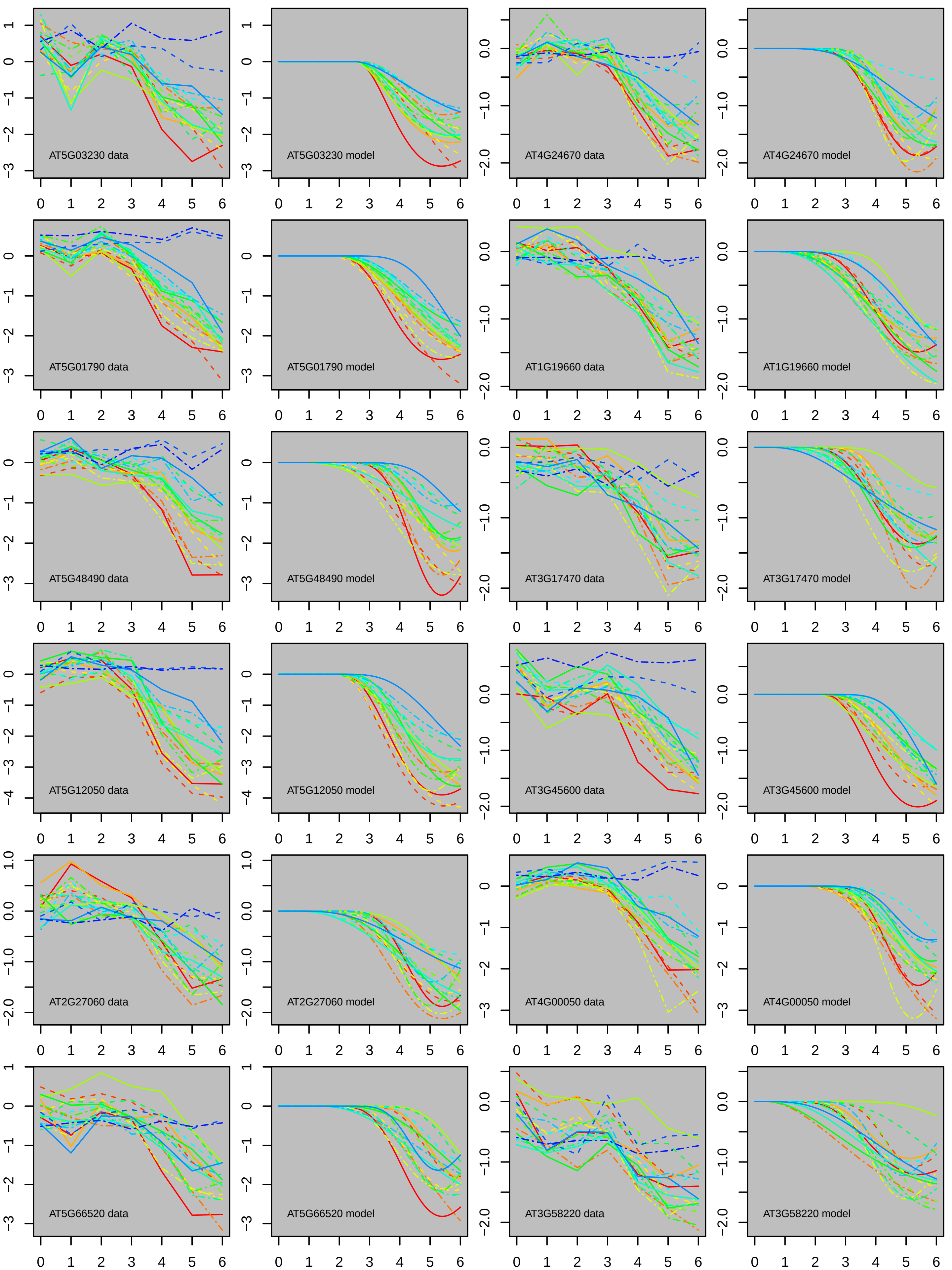
AT5G03230 data
AT5G03230 model
AT4G24670 data
AT4G24670 model
AT5G01790 data
AT5G01790 model
AT1G19660 data
AT1G19660 model
AT5G48490 data
AT5G48490 model
AT3G17470 data
AT3G17470 model
AT5G12050 data
AT5G12050 model
AT3G45600 data
AT3G45600 model
AT2G27060 data
AT2G27060 model
AT4G00050 data
AT4G00050 model
AT5G66520 data
AT5G66520 model
AT3G58220 data
AT3G58220 model

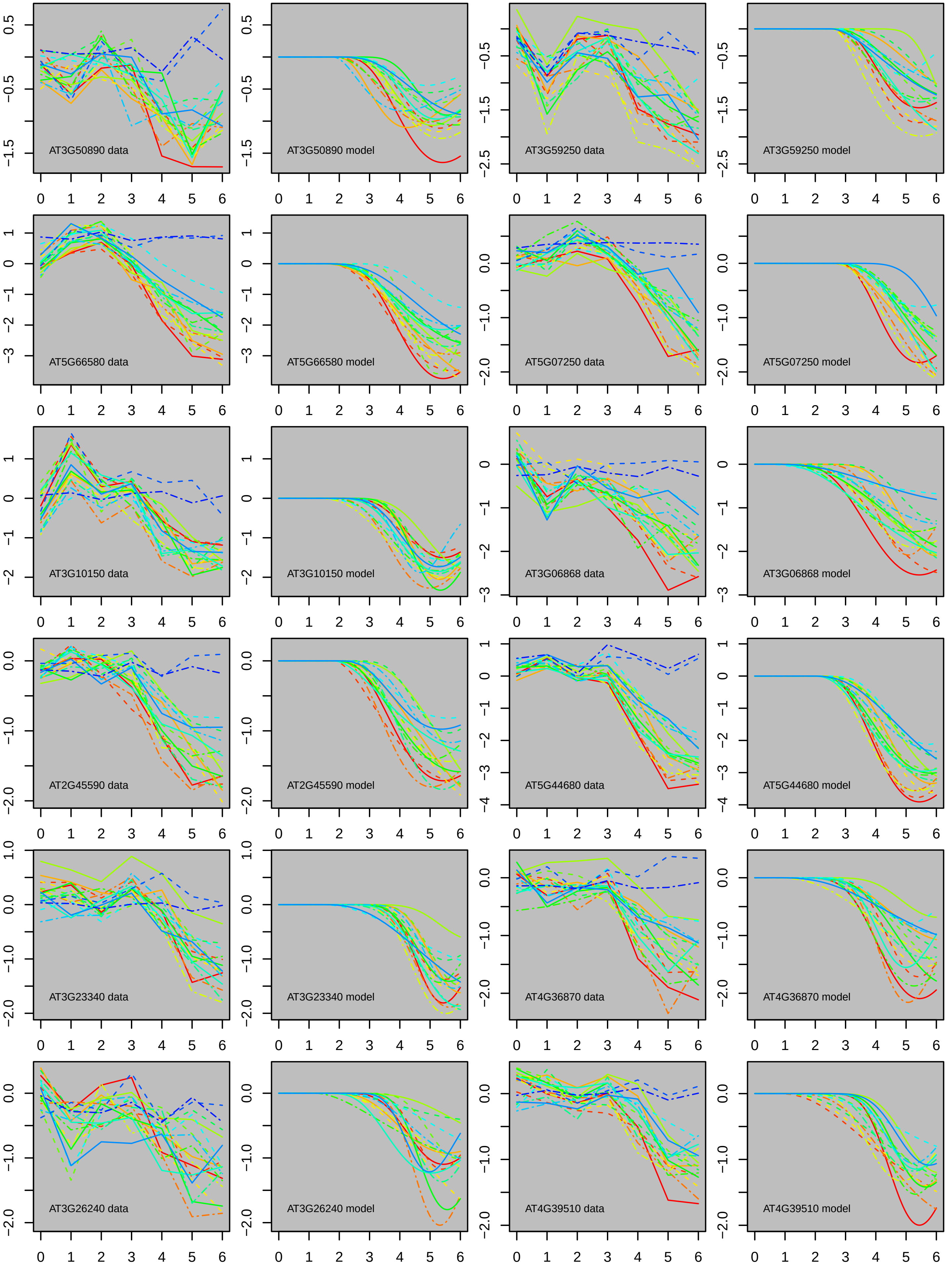
AT3G50890 data
AT3G50890 model
AT3G59250 data
AT3G59250 model
AT5G66580 data
AT5G66580 model
AT5G07250 data
AT5G07250 model
AT3G10150 data
AT3G10150 model
AT3G06868 data
AT3G06868 model
AT2G45590 data
AT2G45590 model
AT5G44680 data
AT5G44680 model
AT3G23340 data
AT3G23340 model
AT4G36870 data
AT4G36870 model
AT3G26240 data
AT3G26240 model
AT4G39510 data
AT4G39510 model

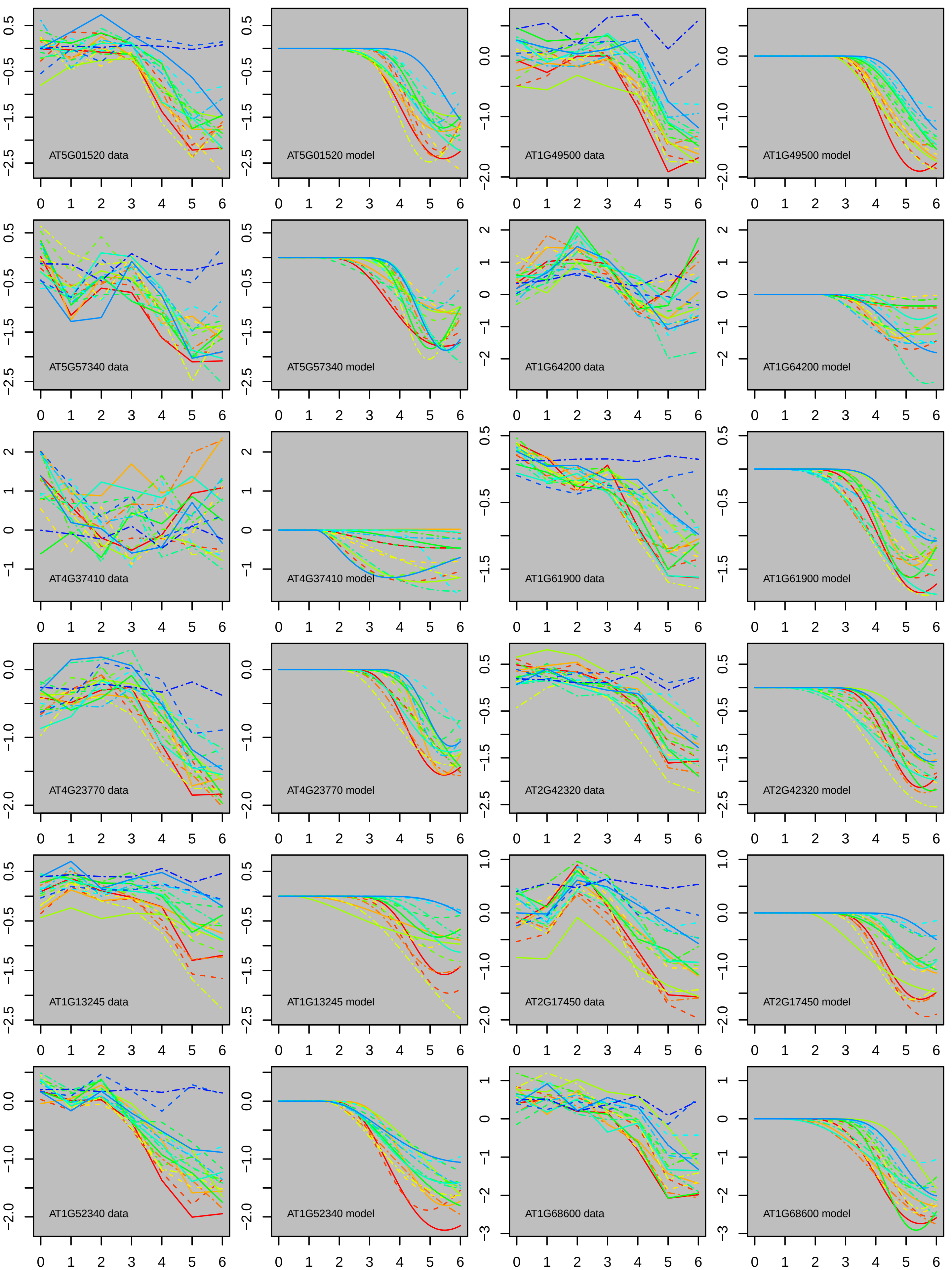

AT5G01520 data
AT5G01520 model
AT1G49500 data
AT1G49500 model
AT5G57340 data
AT5G57340 model
AT1G64200 data
AT1G64200 model
AT4G37410 data
AT4G37410 model
AT1G61900 data
AT1G61900 model
AT4G23770 data
AT4G23770 model
AT2G42320 data
AT2G42320 model
AT1G13245 data
AT1G13245 model
AT2G17450 data
AT2G17450 model
AT1G52340 data
AT1G52340 model
AT1G68600 data
AT1G68600 model

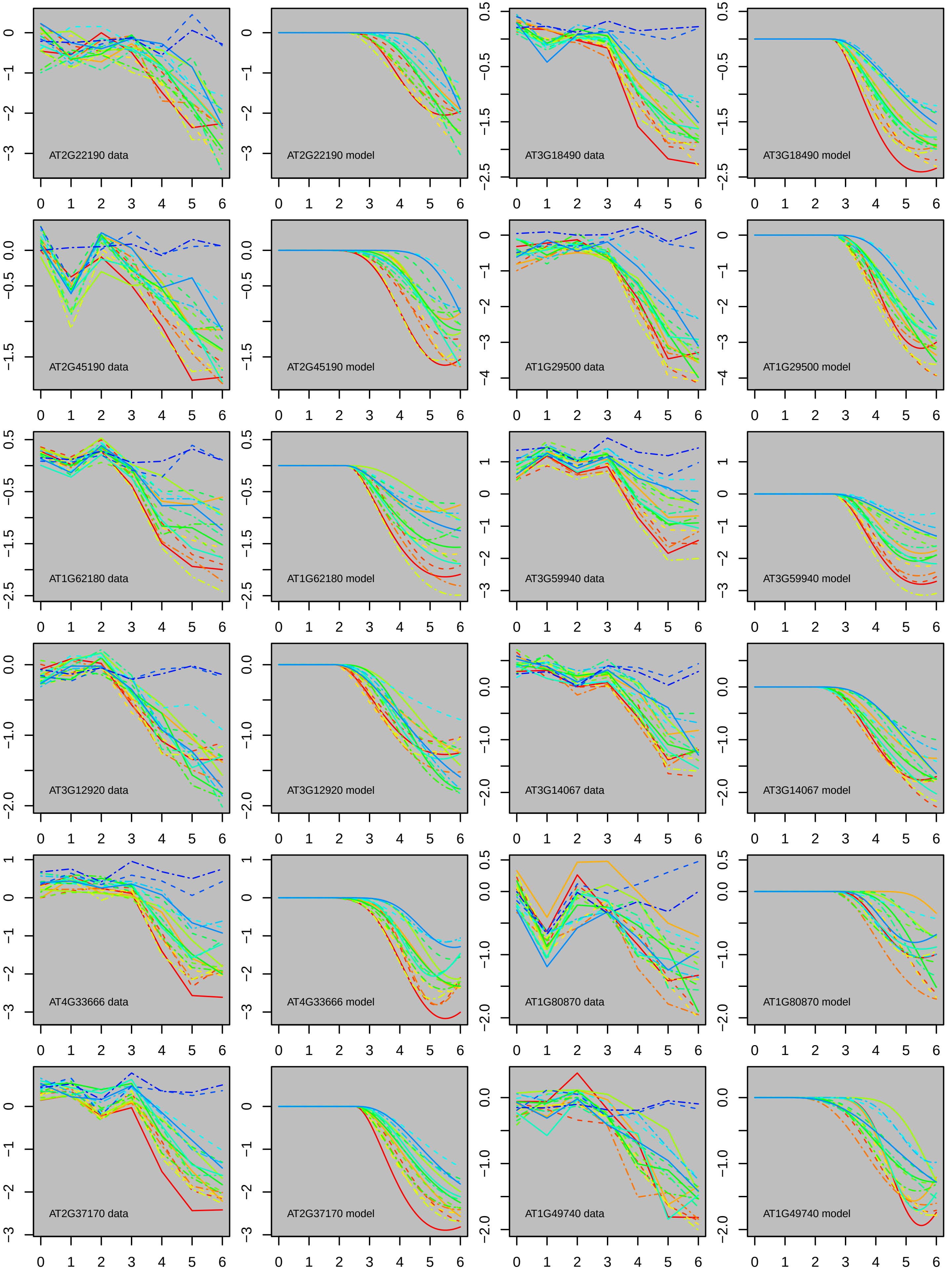

AT2G22190 data
AT2G22190 model
AT3G18490 data
AT3G18490 model
AT2G45190 data
AT2G45190 model
AT1G29500 data
AT1G29500 model
AT1G62180 data
AT1G62180 model
AT3G59940 data
AT3G59940 model
AT3G12920 data
AT3G12920 model
AT3G14067 data
AT3G14067 model
AT4G33666 data
AT4G33666 model
AT1G80870 data
AT1G80870 model
AT2G37170 data
AT2G37170 model
AT1G49740 data
AT1G49740 model

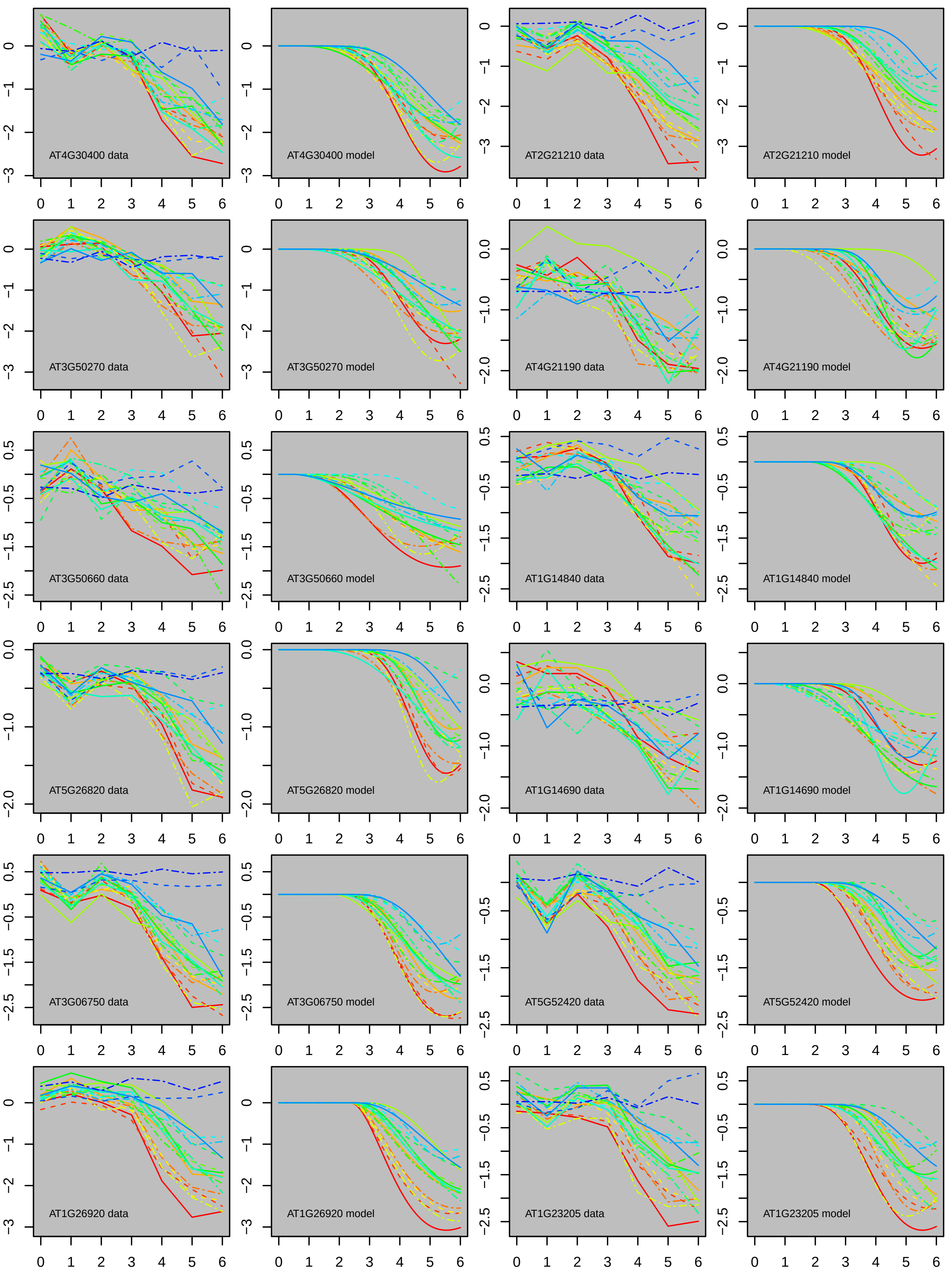
AT4G30400 data
AT4G30400 model
AT2G21210 data
AT2G21210 model
AT3G50270 data
AT3G50270 model
AT4G21190 data
AT4G21190 model
AT3G50660 data
AT3G50660 model
AT1G14840 data
AT1G14840 model
AT5G26820 data
AT5G26820 model
AT1G14690 data
AT1G14690 model
AT3G06750 data
AT3G06750 model
AT5G52420 data
AT5G52420 model
AT1G26920 data
AT1G26920 model
AT1G23205 data
AT1G23205 model

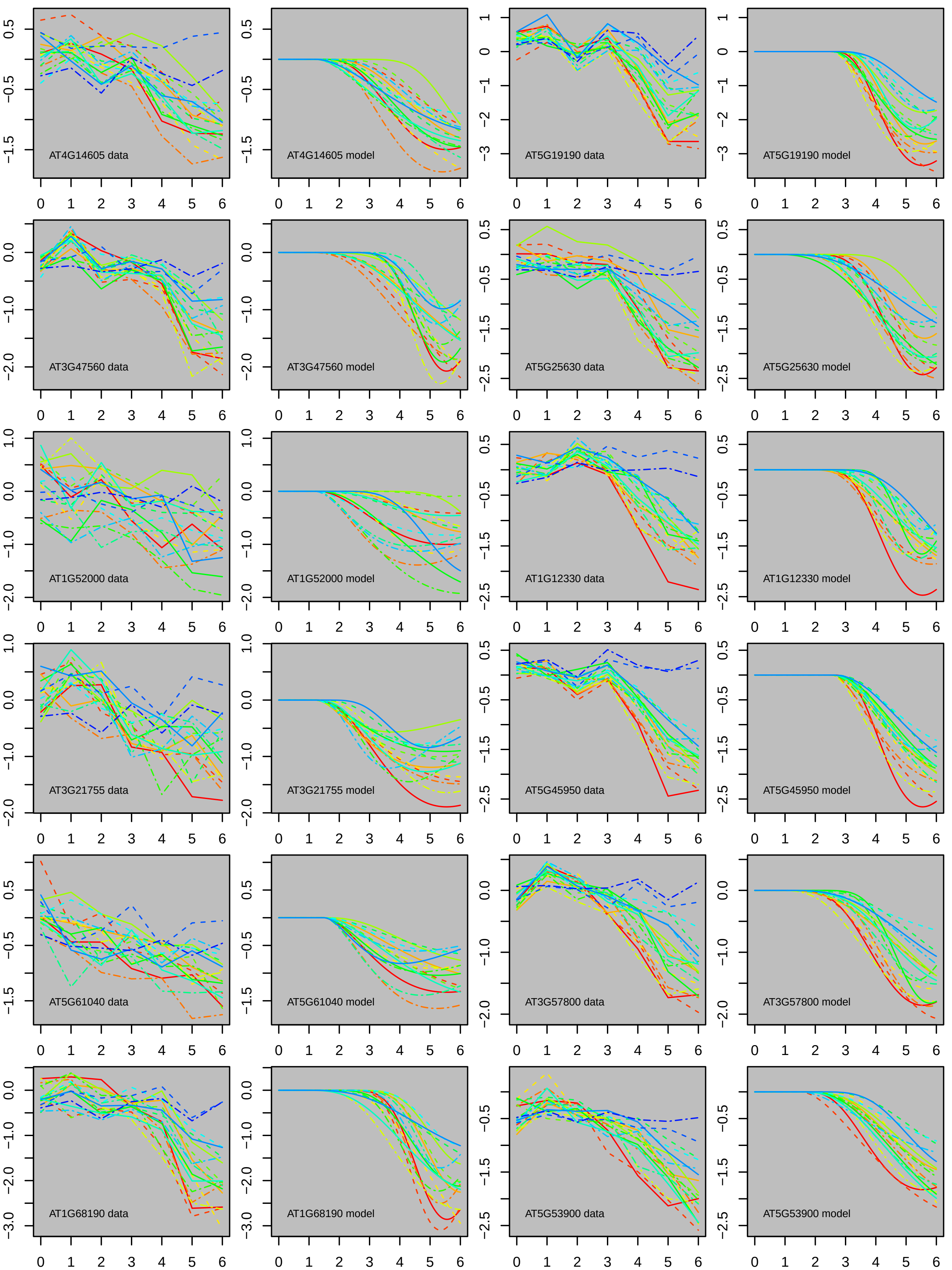
AT4G14605 data
AT4G14605 model
AT5G19190 data
AT5G19190 model
AT3G47560 data
AT3G47560 model
AT5G25630 data
AT5G25630 model
AT1G52000 data
AT1G52000 model
AT1G12330 data
AT1G12330 model
AT3G21755 data
AT3G21755 model
AT5G45950 data
AT5G45950 model
AT5G61040 data
AT5G61040 model
AT3G57800 data
AT3G57800 model
AT1G68190 data
AT1G68190 model
AT5G53900 data
AT5G53900 model

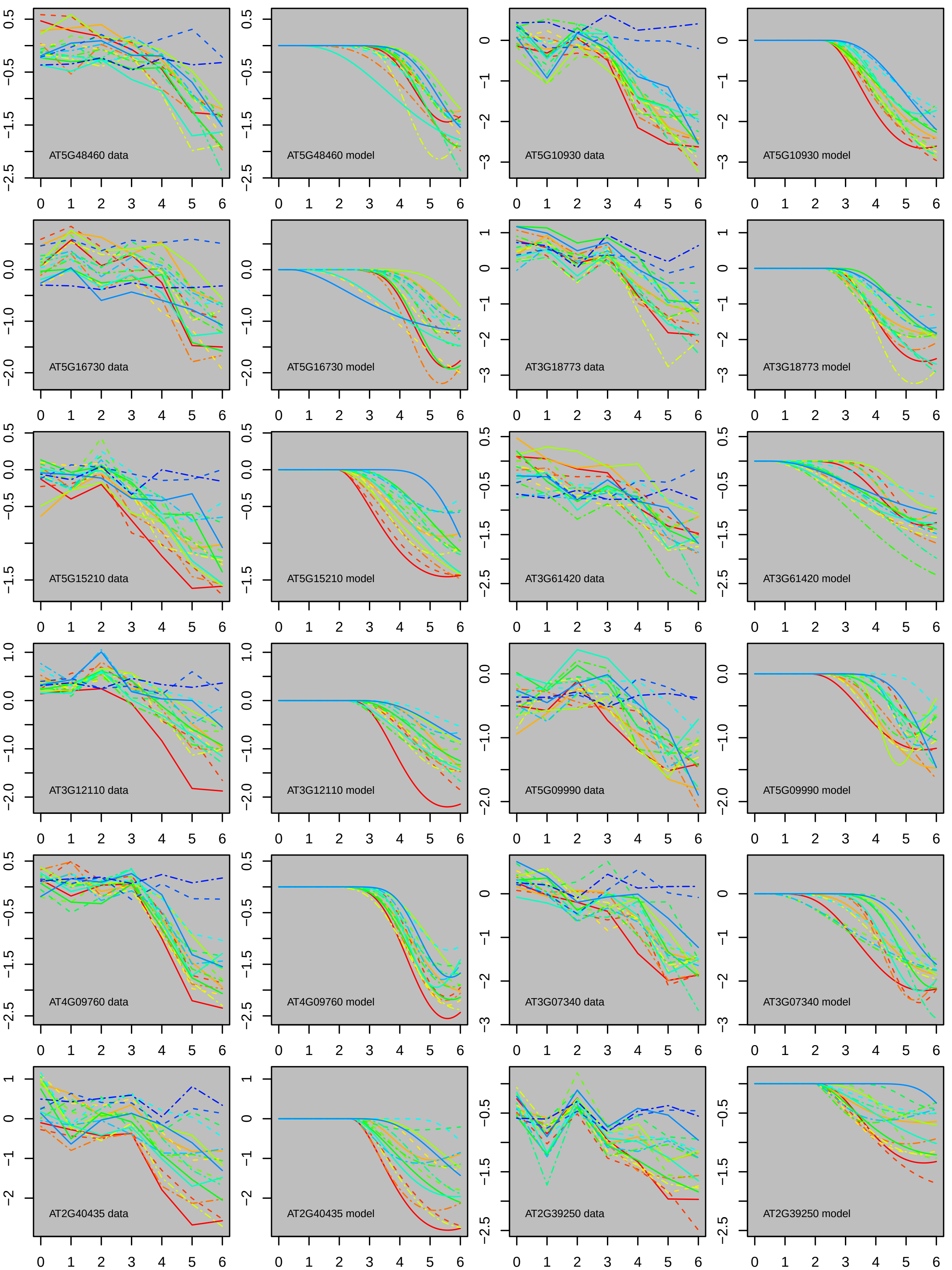
AT5G48460 data
AT5G48460 model
AT5G10930 data
AT5G10930 model
AT5G16730 data
AT5G16730 model
AT3G18773 data
AT3G18773 model
AT5G15210 data
AT5G15210 model
AT3G61420 data
AT3G61420 model
AT3G12110 data
AT3G12110 model
AT5G09990 data
AT5G09990 model
AT4G09760 data
AT4G09760 model
AT3G07340 data
AT3G07340 model
AT2G40435 data
AT2G40435 model
AT2G39250 data
AT2G39250 model

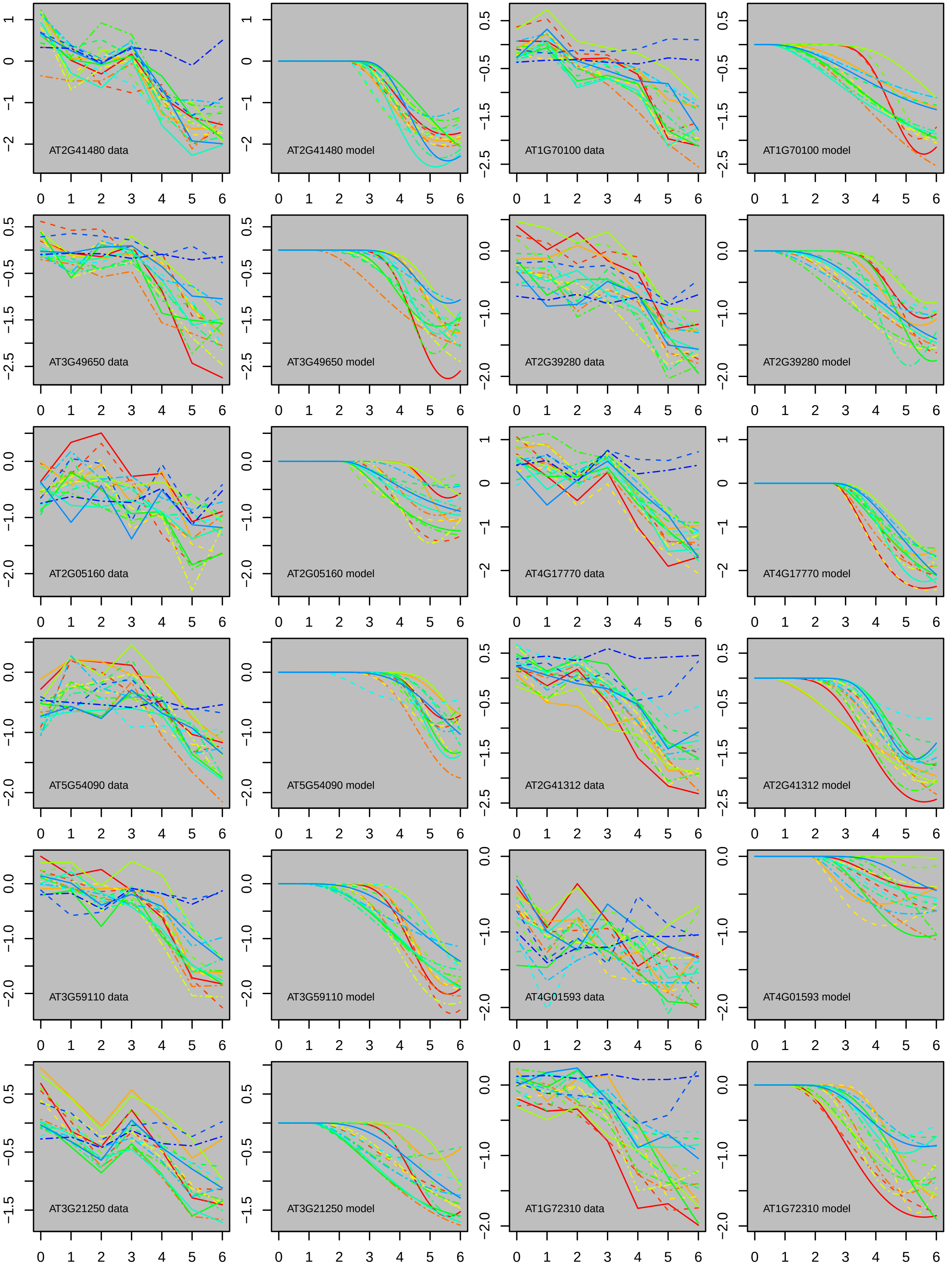
AT2G41480 data
AT2G41480 model
AT1G70100 data
AT1G70100 model
AT3G49650 data
AT3G49650 model
AT2G39280 data
AT2G39280 model
AT2G05160 data
AT2G05160 model
AT4G17770 data
AT4G17770 model
AT5G54090 data
AT5G54090 model
AT2G41312 data
AT2G41312 model
AT3G59110 data
AT3G59110 model
AT4G01593 data
AT4G01593 model
AT3G21250 data
AT3G21250 model
AT1G72310 data
AT1G72310 model

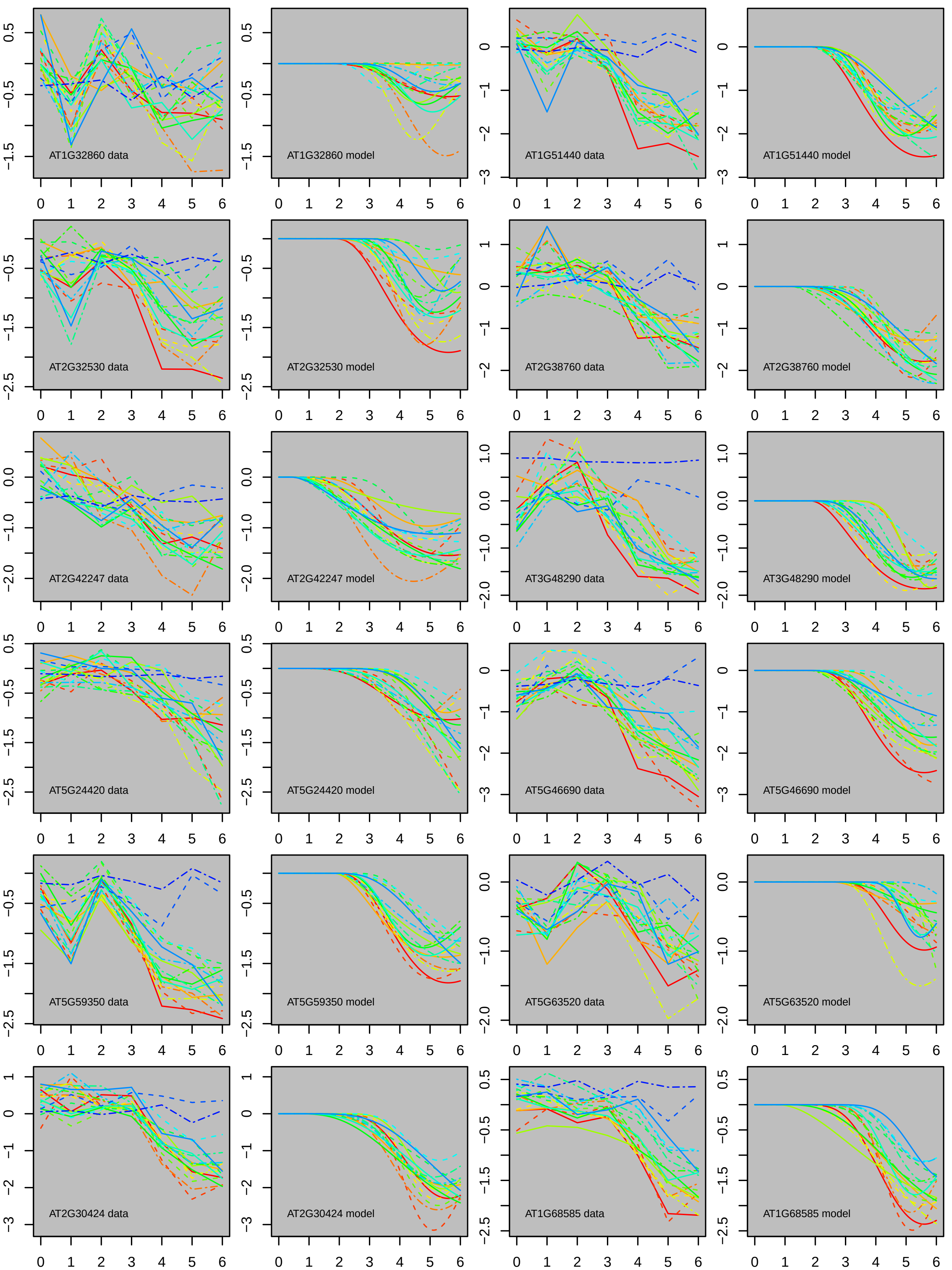

AT1G32860 data
AT1G32860 model
AT1G51440 data
AT1G51440 model
AT2G32530 data
AT2G32530 model
AT2G38760 data
AT2G38760 model
AT2G42247 data
AT2G42247 model
AT3G48290 data
AT3G48290 model
AT5G24420 data
AT5G24420 model
AT5G46690 data
AT5G46690 model
AT5G59350 data
AT5G59350 model
AT5G63520 data
AT5G63520 model
AT2G30424 data
AT2G30424 model
AT1G68585 data
AT1G68585 model

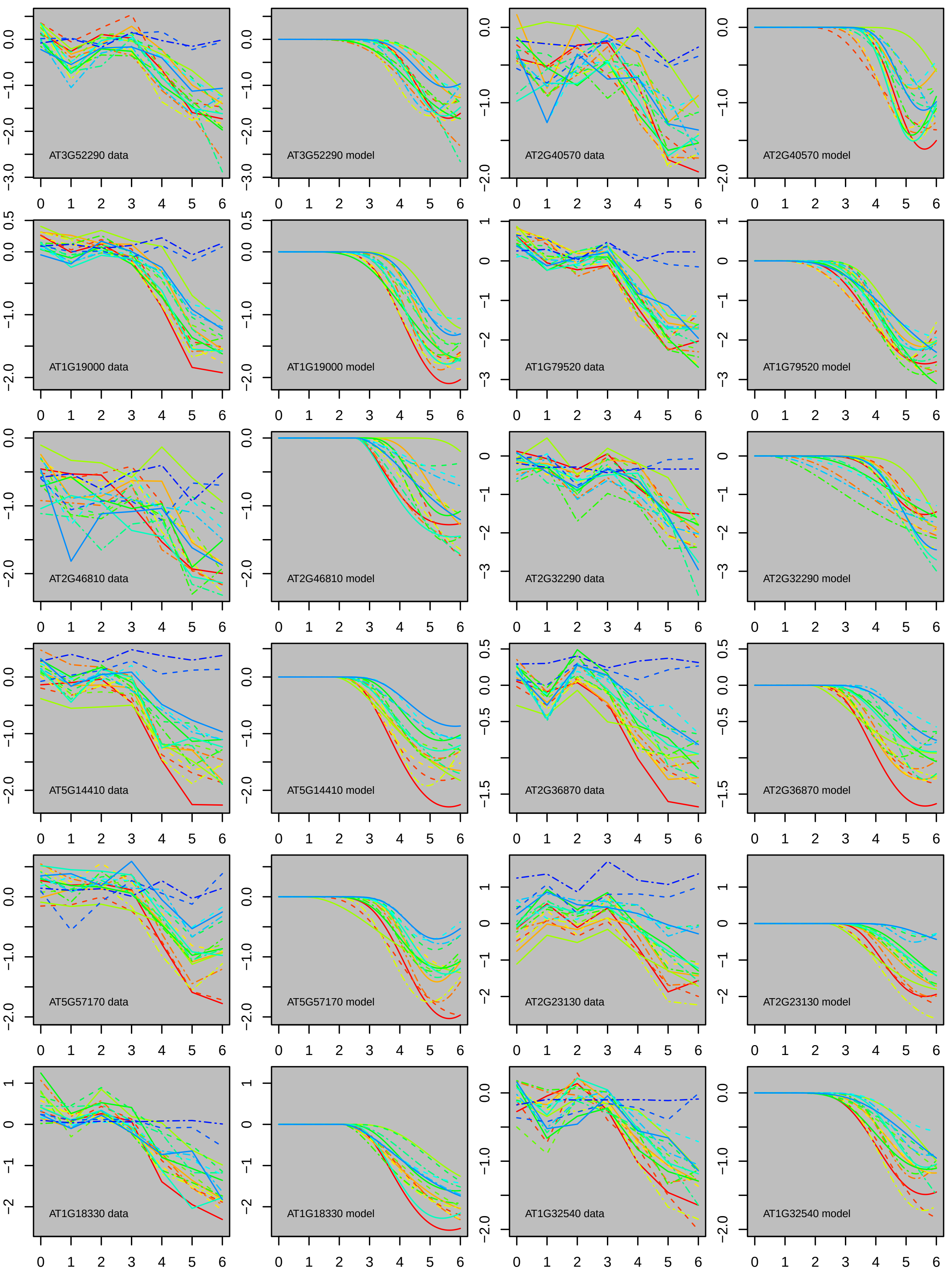

AT3G52290 data
AT3G52290 model
AT2G40570 data
AT2G40570 model
AT1G19000 data
AT1G19000 model
AT1G79520 data
AT1G79520 model
AT2G46810 data
AT2G46810 model
AT2G32290 data
AT2G32290 model
AT5G14410 data
AT5G14410 model
AT2G36870 data
AT2G36870 model
AT5G57170 data
AT5G57170 model
AT2G23130 data
AT2G23130 model
AT1G18330 data
AT1G18330 model
AT1G32540 data
AT1G32540 model

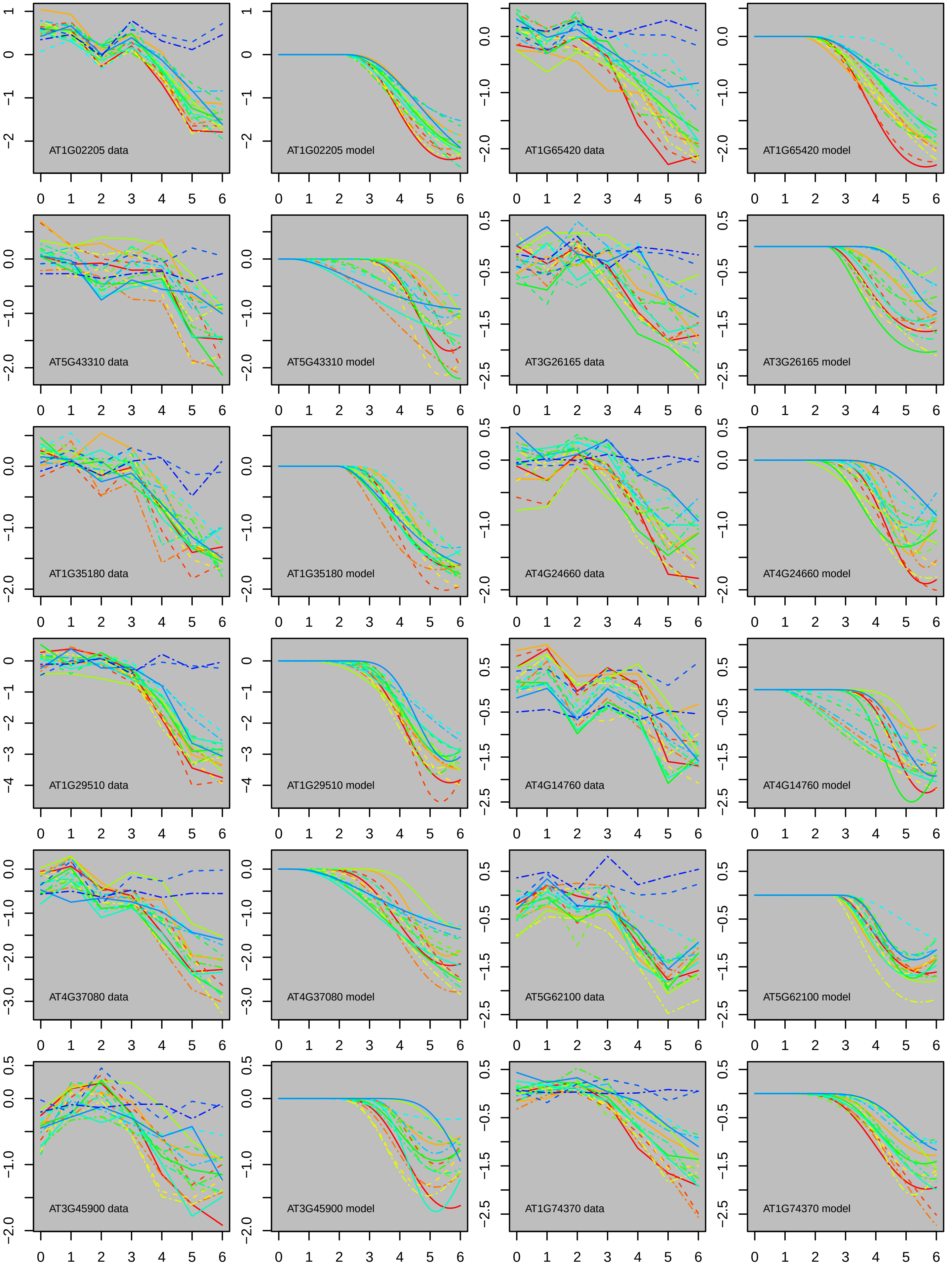
AT1G02205 data
AT1G02205 model
AT1G65420 data
AT1G65420 model
AT5G43310 data
AT5G43310 model
AT3G26165 data
AT3G26165 model
AT1G35180 data
AT1G35180 model
AT4G24660 data
AT4G24660 model
AT1G29510 data
AT1G29510 model
AT4G14760 data
AT4G14760 model
AT4G37080 data
AT4G37080 model
AT5G62100 data
AT5G62100 model
AT3G45900 data
AT3G45900 model
AT1G74370 data
AT1G74370 model

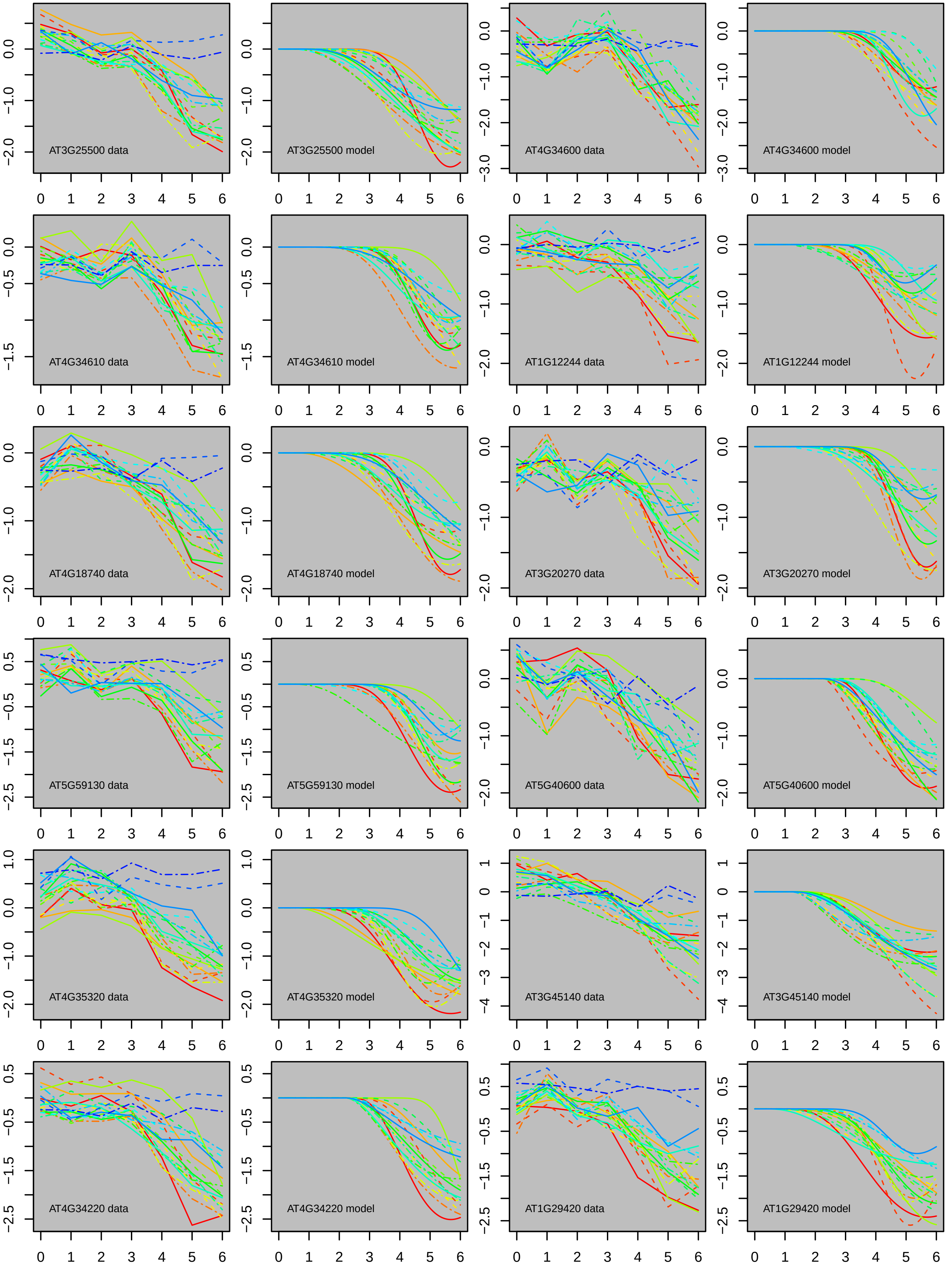
AT3G25500 data
AT3G25500 model
AT4G34600 data
AT4G34600 model
AT4G34610 data
AT4G34610 model
AT1G12244 data
AT1G12244 model
AT4G18740 data
AT4G18740 model
AT3G20270 data
AT3G20270 model
AT5G59130 data
AT5G59130 model
AT5G40600 data
AT5G40600 model
AT4G35320 data
AT4G35320 model
AT3G45140 data
AT3G45140 model
AT4G34220 data
AT4G34220 model
AT1G29420 data
AT1G29420 model

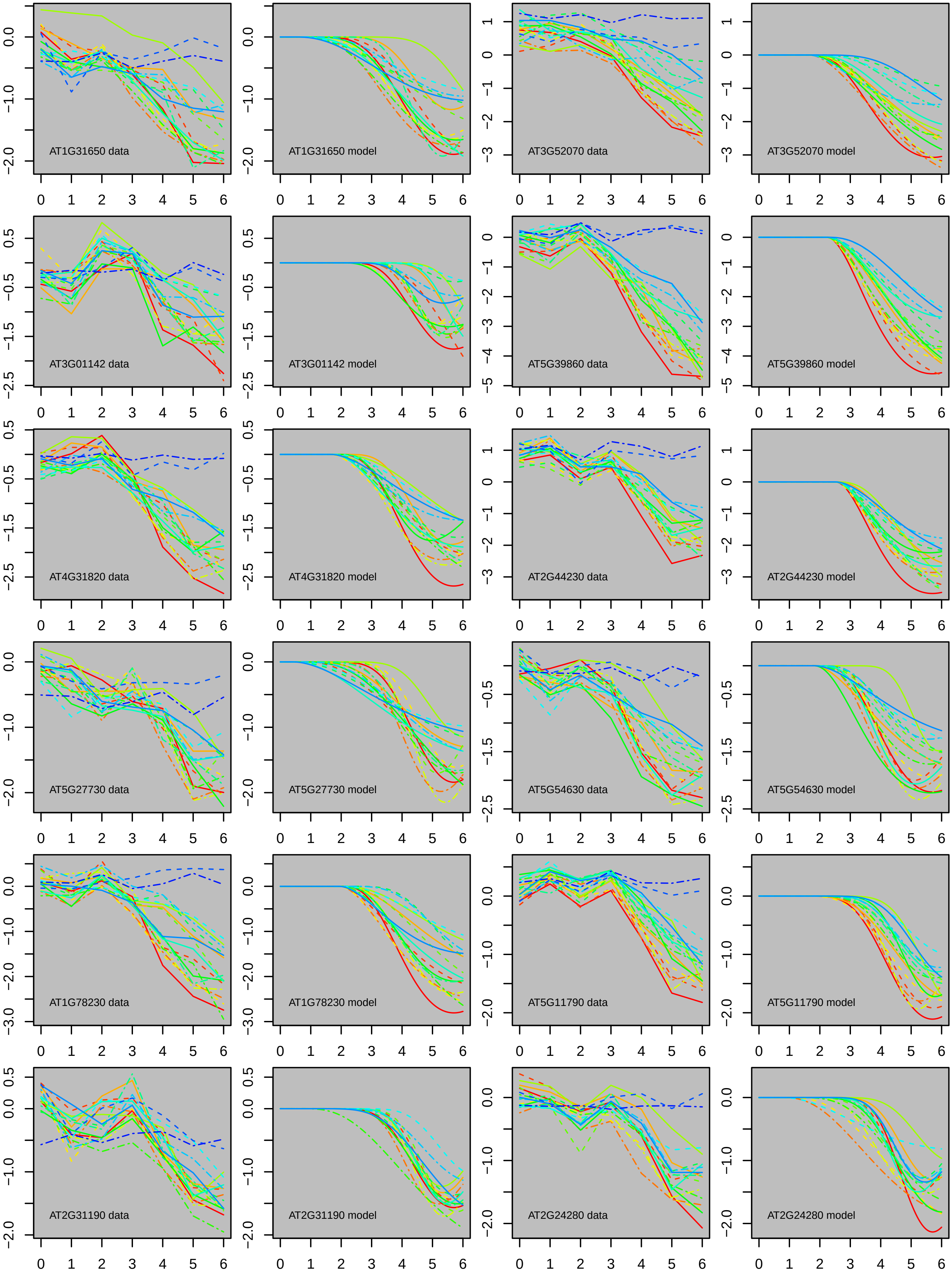
AT1G31650 data
AT1G31650 model
AT3G52070 data
AT3G52070 model
AT3G01142 data
AT3G01142 model
AT5G39860 data
AT5G39860 model
AT4G31820 data
AT4G31820 model
AT2G44230 data
AT2G44230 model
AT5G27730 data
AT5G27730 model
AT5G54630 data
AT5G54630 model
AT1G78230 data
AT1G78230 model
AT5G11790 data
AT5G11790 model
AT2G31190 data
AT2G31190 model
AT2G24280 data
AT2G24280 model

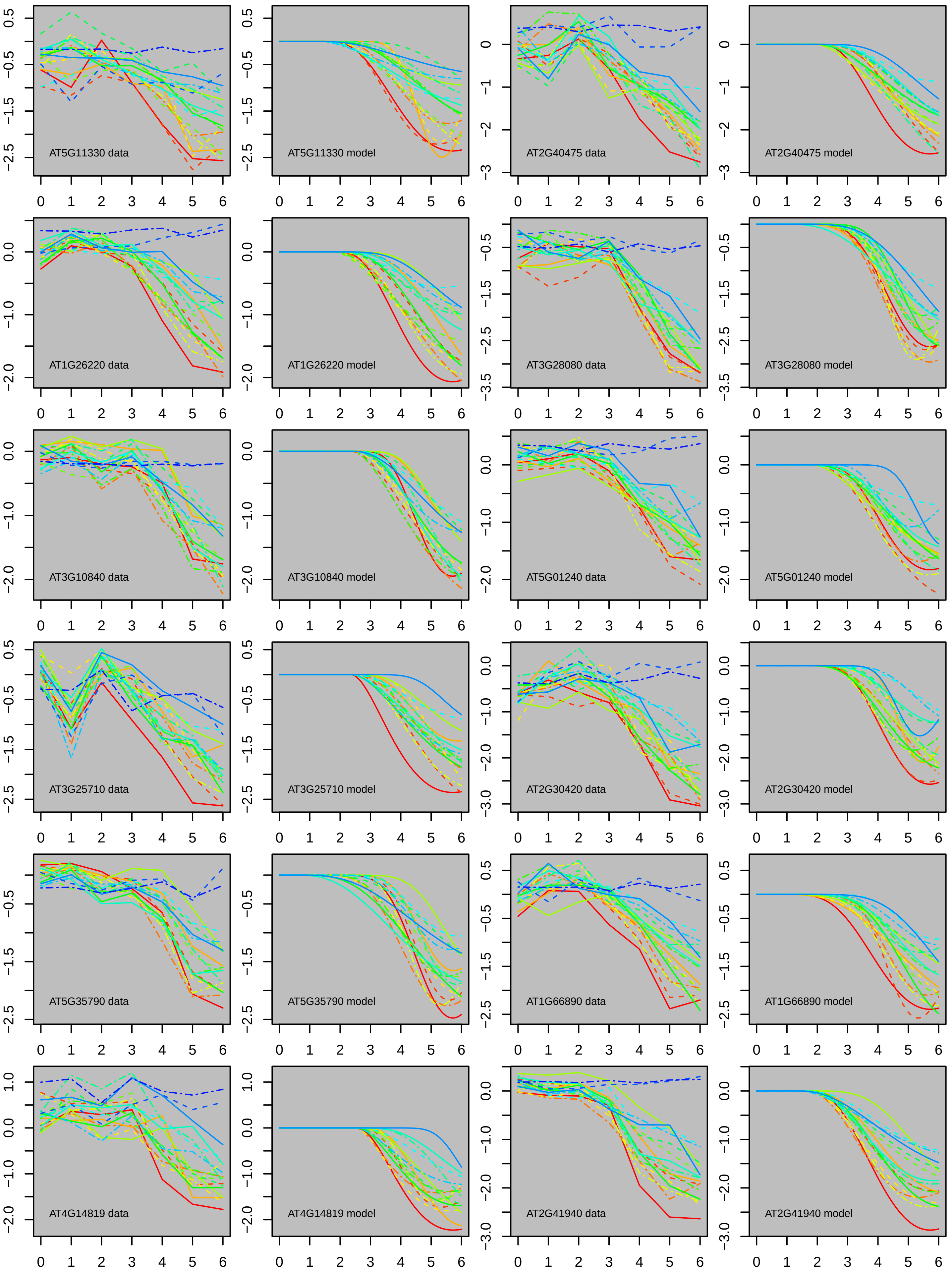

AT5G11330 data
AT5G11330 model
AT2G40475 data
AT2G40475 model
AT1G26220 data
AT1G26220 model
AT3G28080 data
AT3G28080 model
AT3G10840 data
AT3G10840 model
AT5G01240 data
AT5G01240 model
AT3G25710 data
AT3G25710 model
AT2G30420 data
AT2G30420 model
AT5G35790 data
AT5G35790 model
AT1G66890 data
AT1G66890 model
AT4G14819 data
AT4G14819 model
AT2G41940 data
AT2G41940 model

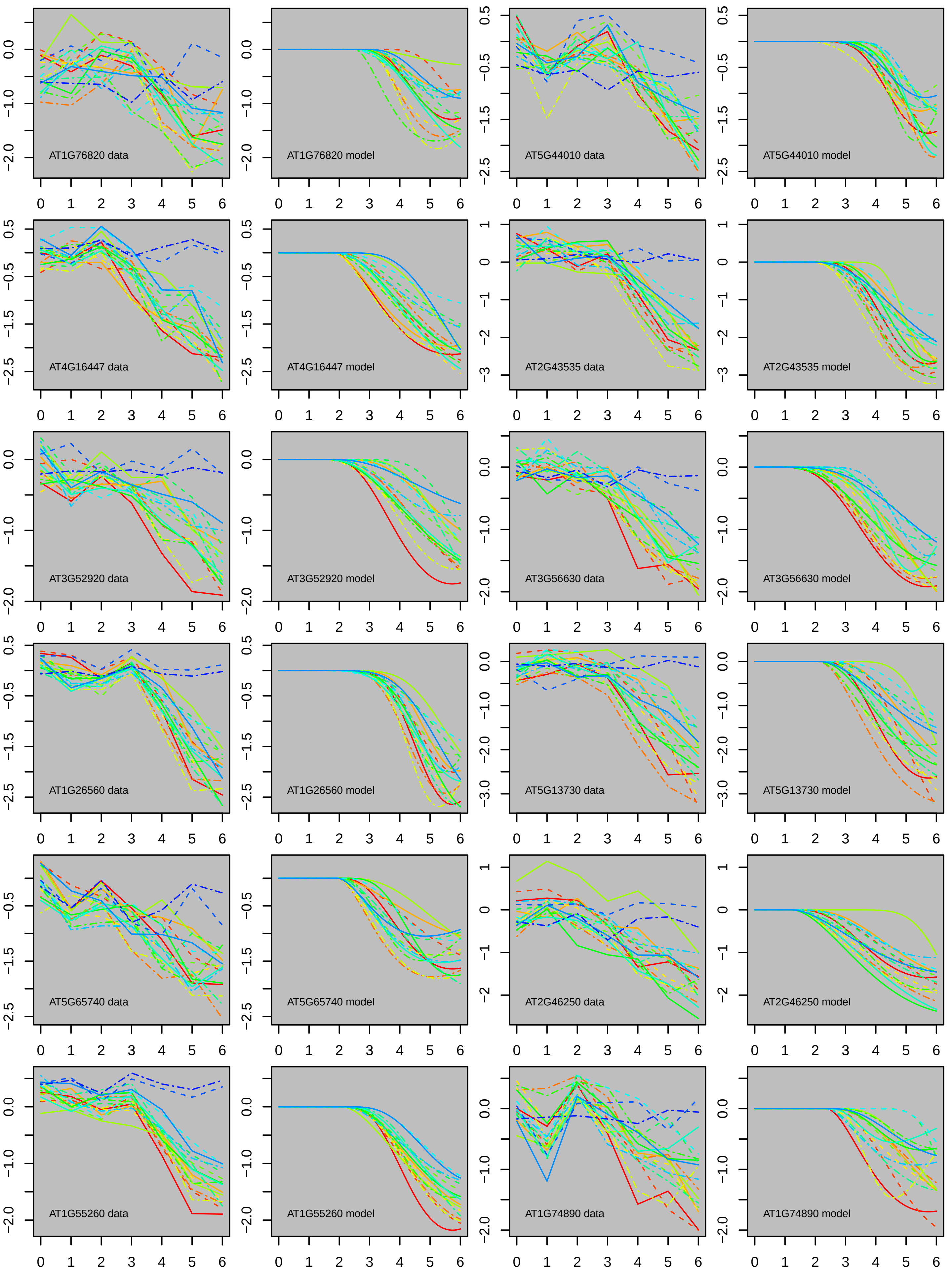
AT1G76820 data
AT1G76820 model
AT5G44010 data
AT5G44010 model
AT4G16447 data
AT4G16447 model
AT2G43535 data
AT2G43535 model
AT3G52920 data
AT3G52920 model
AT3G56630 data
AT3G56630 model
AT1G26560 data
AT1G26560 model
AT5G13730 data
AT5G13730 model
AT5G65740 data
AT5G65740 model
AT2G46250 data
AT2G46250 model
AT1G55260 data
AT1G55260 model
AT1G74890 data
AT1G74890 model

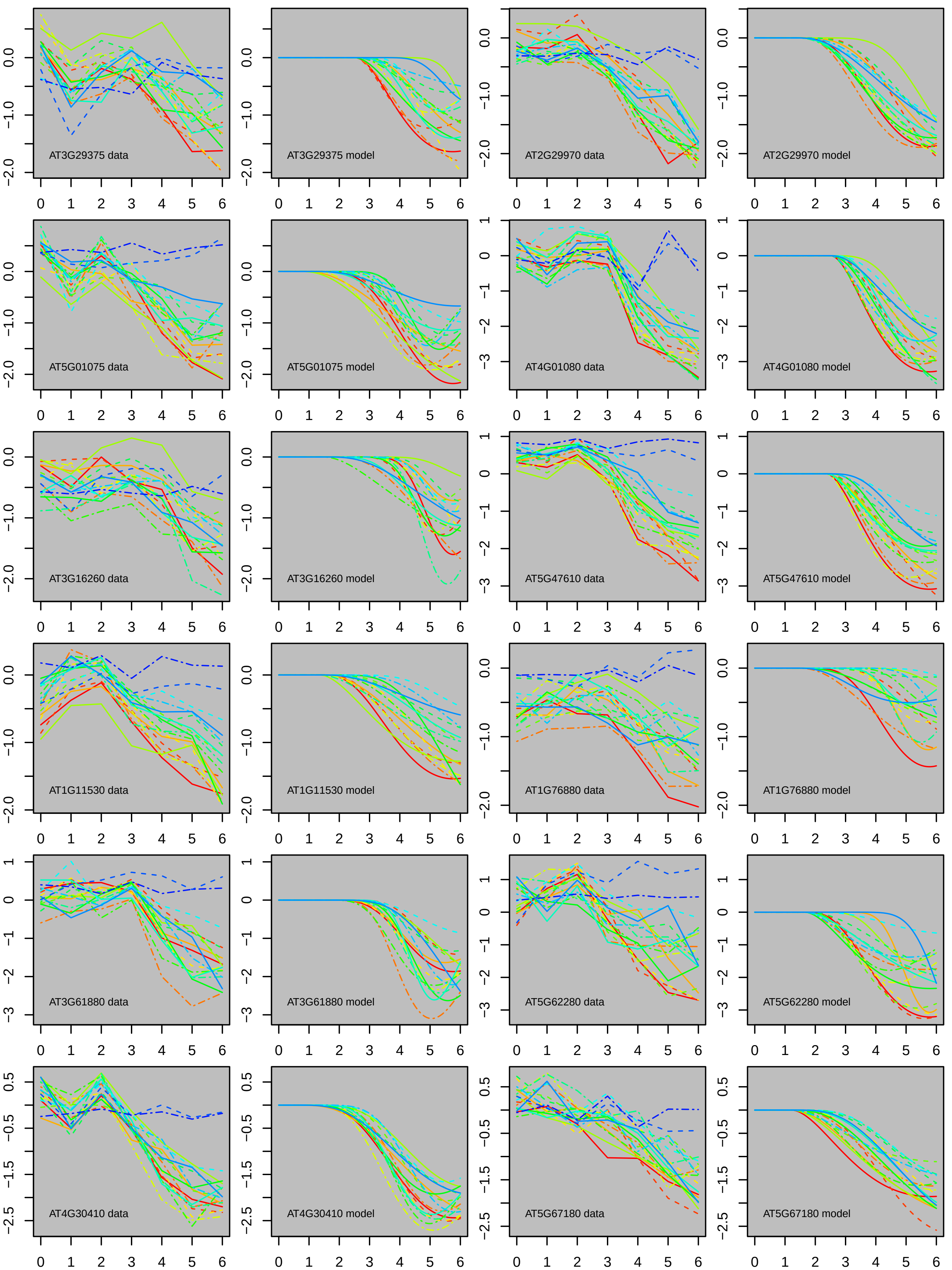
AT3G29375 data
AT3G29375 model
AT2G29970 data
AT2G29970 model
AT5G01075 data
AT5G01075 model
AT4G01080 data
AT4G01080 model
AT3G16260 data
AT3G16260 model
AT5G47610 data
AT5G47610 model
AT1G11530 data
AT1G11530 model
AT1G76880 data
AT1G76880 model
AT3G61880 data
AT3G61880 model
AT5G62280 data
AT5G62280 model
AT4G30410 data
AT4G30410 model
AT5G67180 data
AT5G67180 model

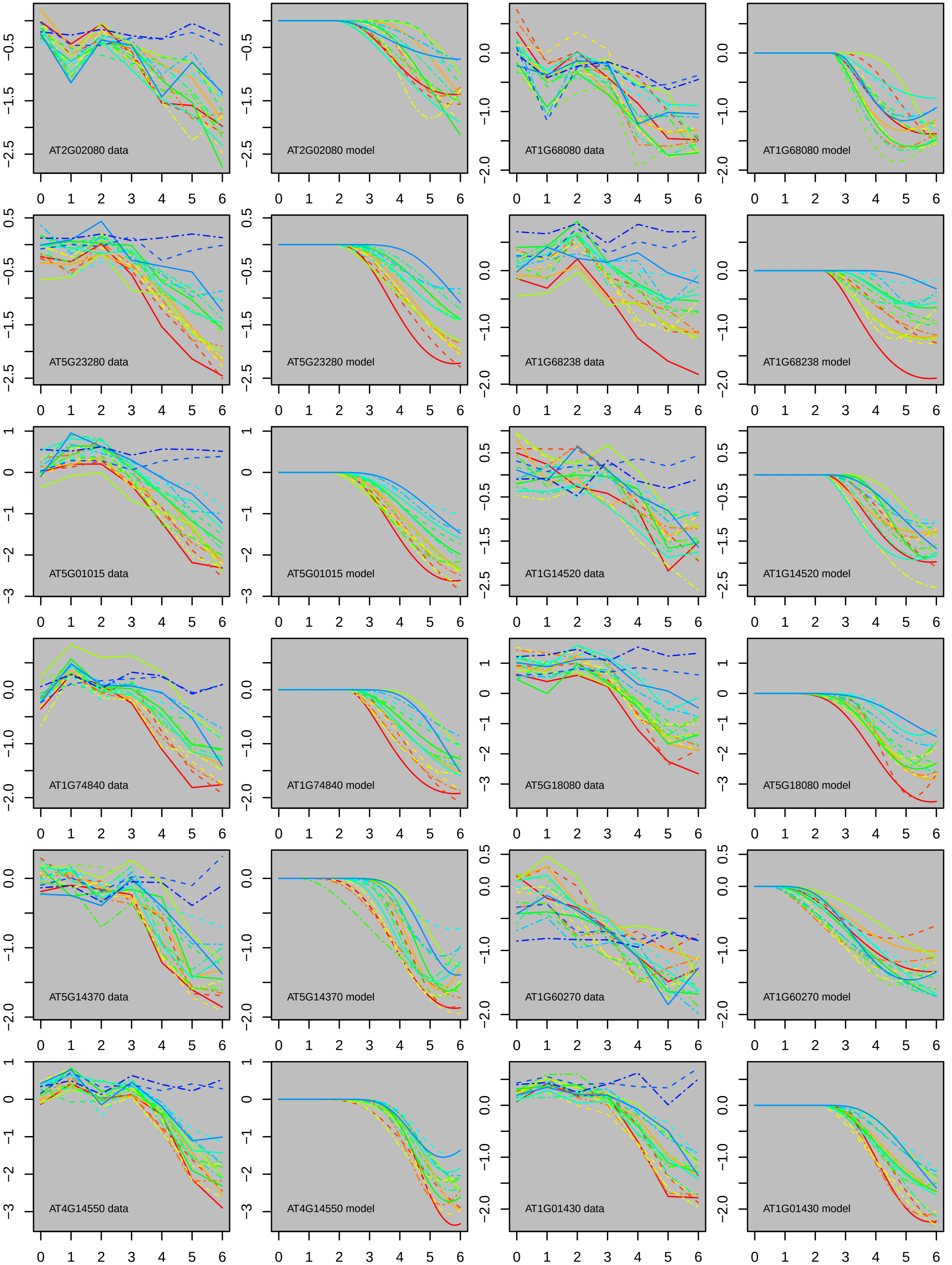

AT2G02080 data
AT2G02080 model
AT1G68080 data
AT1G68080 model
AT5G23280 data
AT5G23280 model
AT1G68238 data
AT1G68238 model
AT5G01015 data
AT5G01015 model
AT1G14520 data
AT1G14520 model
AT1G74840 data
AT1G74840 model
AT5G18080 data
AT5G18080 model
AT5G14370 data
AT5G14370 model
AT1G60270 data
AT1G60270 model
AT4G14550 data
AT4G14550 model
AT1G01430 data
AT1G01430 model

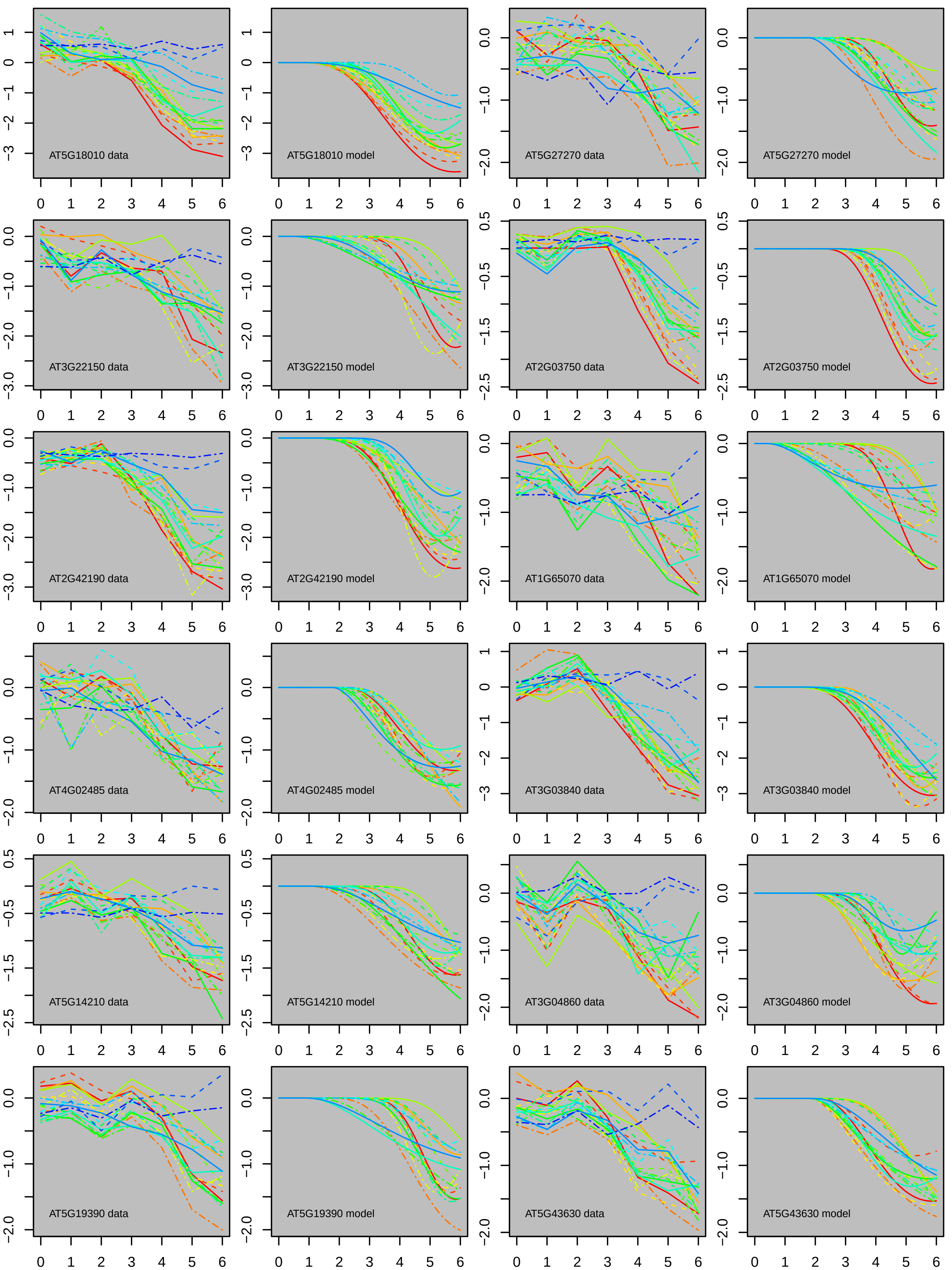

AT5G18010 data
AT5G18010 model
AT5G27270 data
AT5G27270 model
AT3G22150 data
AT3G22150 model
AT2G03750 data
AT2G03750 model
AT2G42190 data
AT2G42190 model
AT1G65070 data
AT1G65070 model
AT4G02485 data
AT4G02485 model
AT3G03840 data
AT3G03840 model
AT5G14210 data
AT5G14210 model
AT3G04860 data
AT3G04860 model
AT5G19390 data
AT5G19390 model
AT5G43630 data
AT5G43630 model

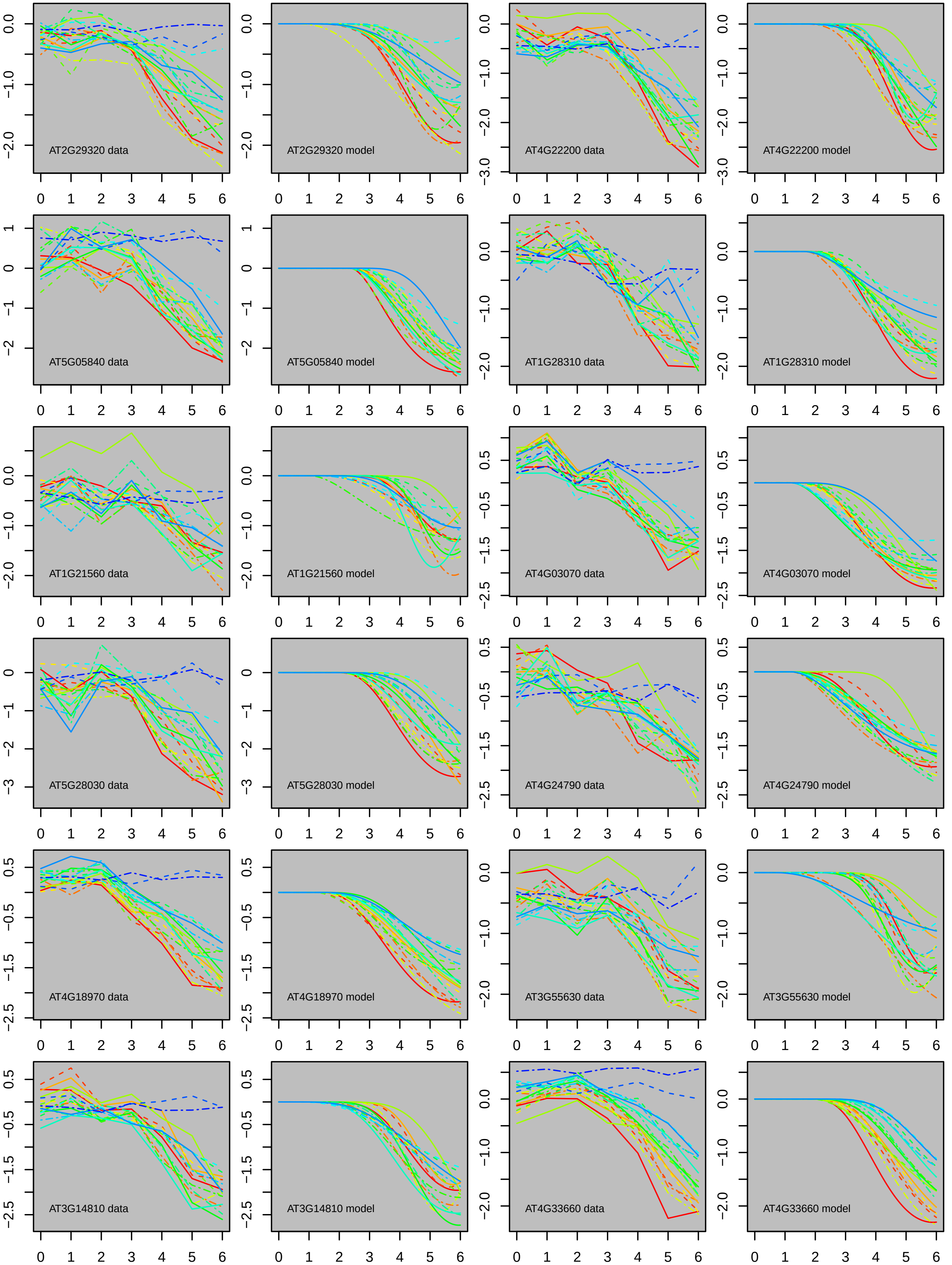

AT2G29320 data
AT2G29320 model
AT4G22200 data
AT4G22200 model
AT5G05840 data
AT5G05840 model
AT1G28310 data
AT1G28310 model
AT1G21560 data
AT1G21560 model
AT4G03070 data
AT4G03070 model
AT5G28030 data
AT5G28030 model
AT4G24790 data
AT4G24790 model
AT4G18970 data
AT4G18970 model
AT3G55630 data
AT3G55630 model
AT3G14810 data
AT3G14810 model
AT4G33660 data
AT4G33660 model

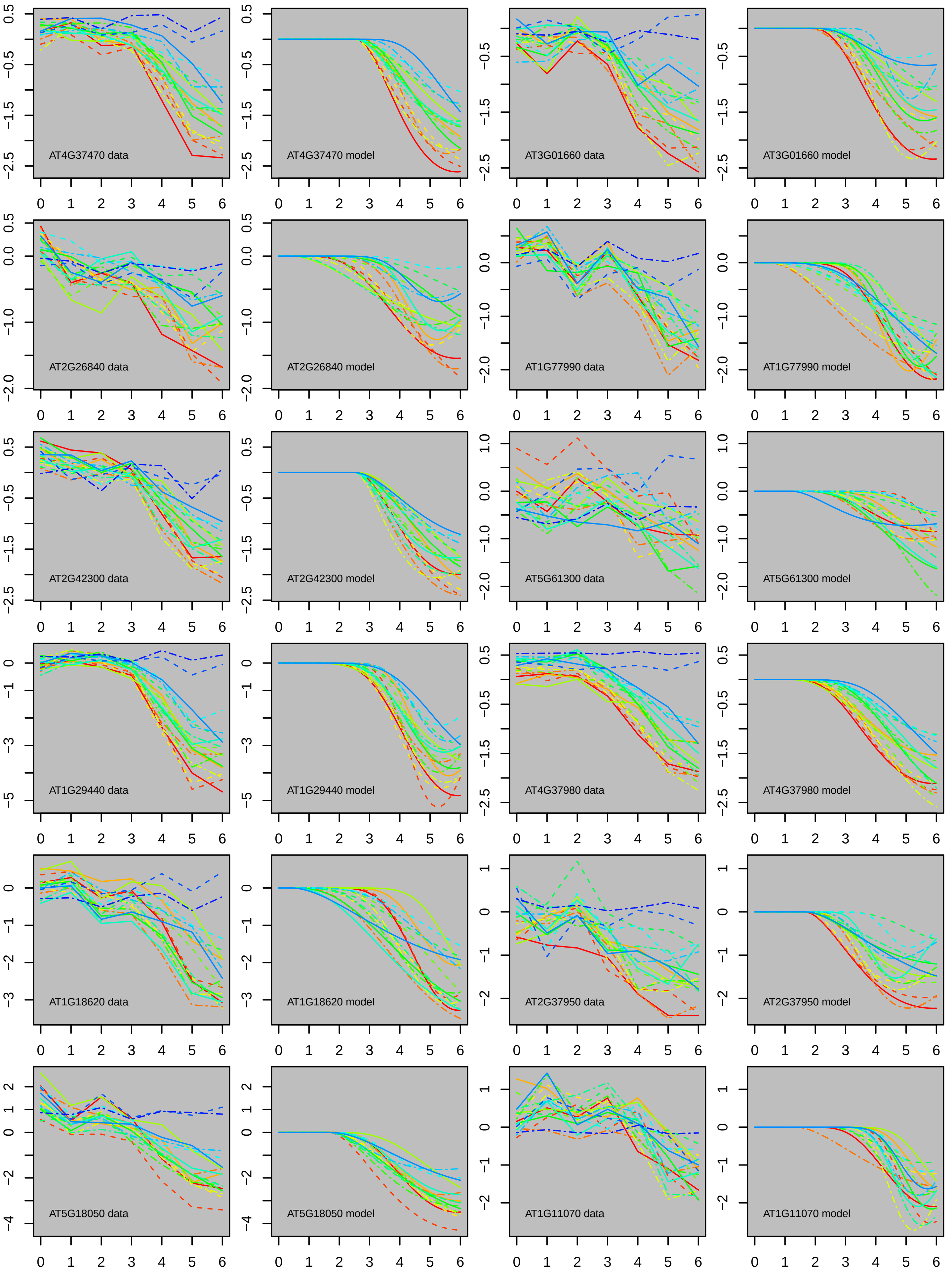

AT4G37470 data
AT4G37470 model
AT3G01660 data
AT3G01660 model
AT2G26840 data
AT2G26840 model
AT1G77990 data
AT1G77990 model
AT2G42300 data
AT2G42300 model
AT5G61300 data
AT5G61300 model
AT1G29440 data
AT1G29440 model
AT4G37980 data
AT4G37980 model
AT1G18620 data
AT1G18620 model
AT2G37950 data
AT2G37950 model
AT5G18050 data
AT5G18050 model
AT1G11070 data
AT1G11070 model

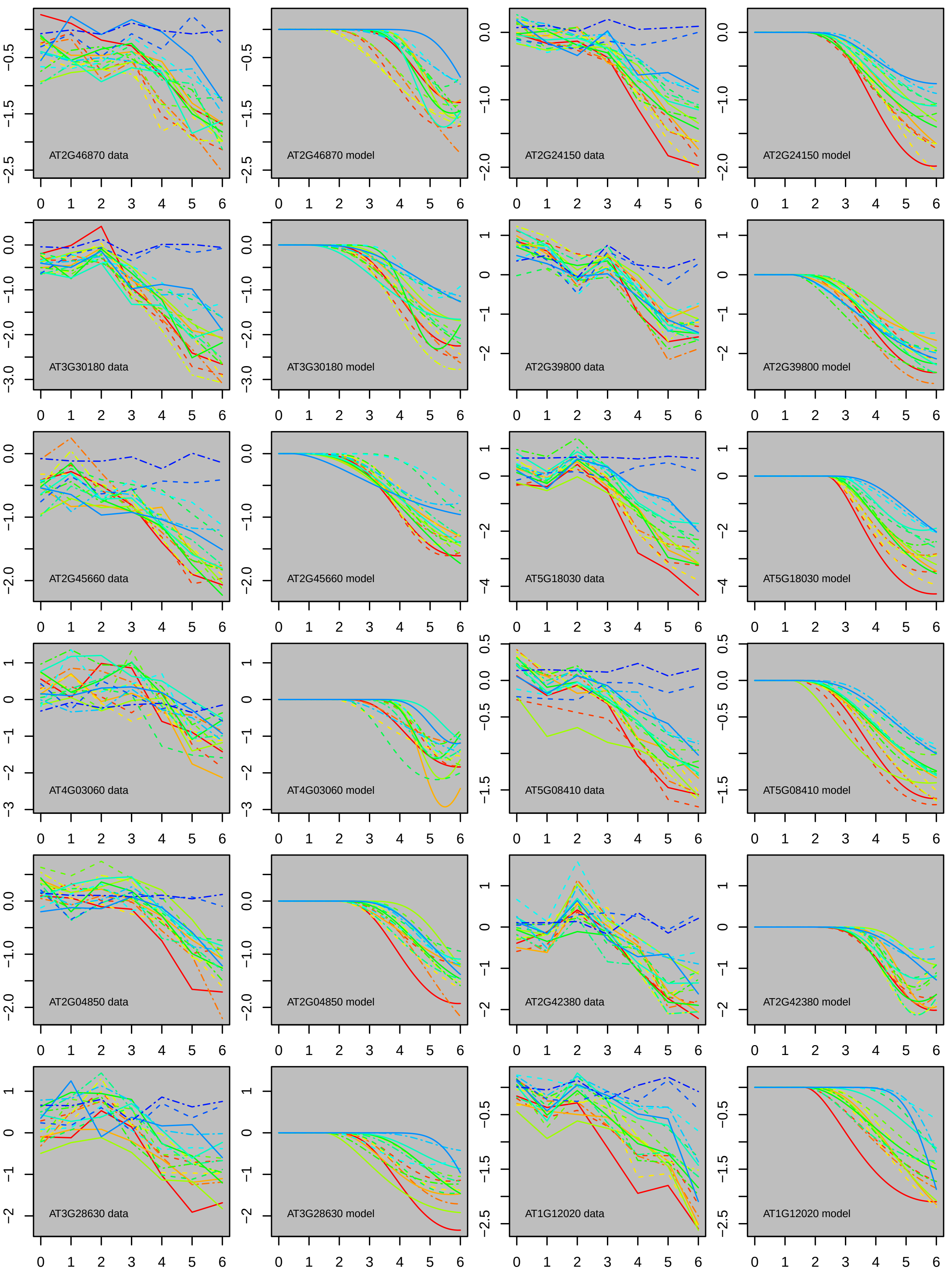
AT2G46870 data
AT2G46870 model
AT2G24150 data
AT2G24150 model
AT3G30180 data
AT3G30180 model
AT2G39800 data
AT2G39800 model
AT2G45660 data
AT2G45660 model
AT5G18030 data
AT5G18030 model
AT4G03060 data
AT4G03060 model
AT5G08410 data
AT5G08410 model
AT2G04850 data
AT2G04850 model
AT2G42380 data
AT2G42380 model
AT3G28630 data
AT3G28630 model
AT1G12020 data
AT1G12020 model

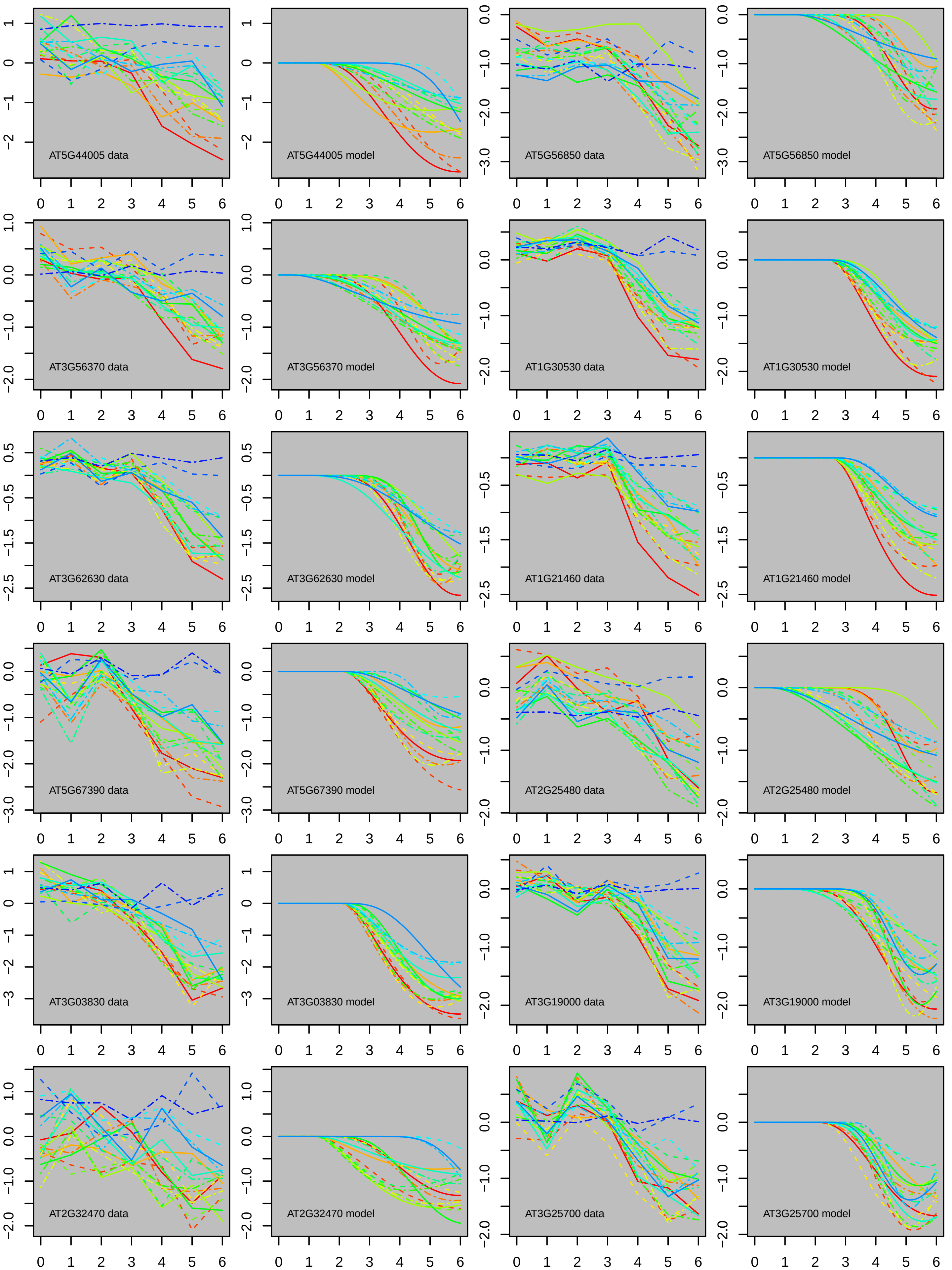
AT5G44005 data
AT5G44005 model
AT5G56850 data
AT5G56850 model
AT3G56370 data
AT3G56370 model
AT1G30530 data
AT1G30530 model
AT3G62630 data
AT3G62630 model
AT1G21460 data
AT1G21460 model
AT5G67390 data
AT5G67390 model
AT2G25480 data
AT2G25480 model
AT3G03830 data
AT3G03830 model
AT3G19000 data
AT3G19000 model
AT2G32470 data
AT2G32470 model
AT3G25700 data
AT3G25700 model

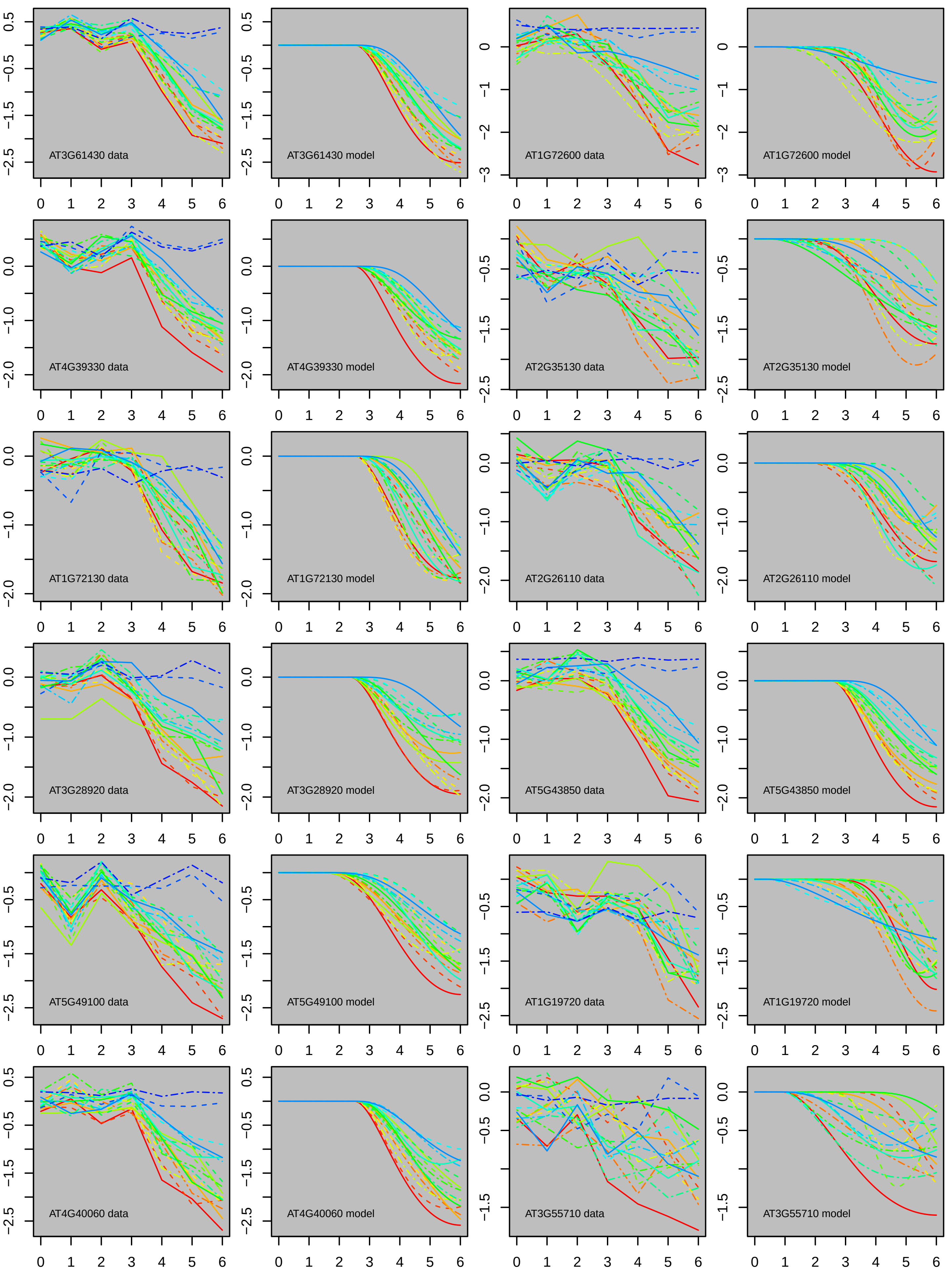
AT3G61430 data
AT3G61430 model
AT1G72600 data
AT1G72600 model
AT4G39330 data
AT4G39330 model
AT2G35130 data
AT2G35130 model
AT1G72130 data
AT1G72130 model
AT2G26110 data
AT2G26110 model
AT3G28920 data
AT3G28920 model
AT5G43850 data
AT5G43850 model
AT5G49100 data
AT5G49100 model
AT1G19720 data
AT1G19720 model
AT4G40060 data
AT4G40060 model
AT3G55710 data
AT3G55710 model

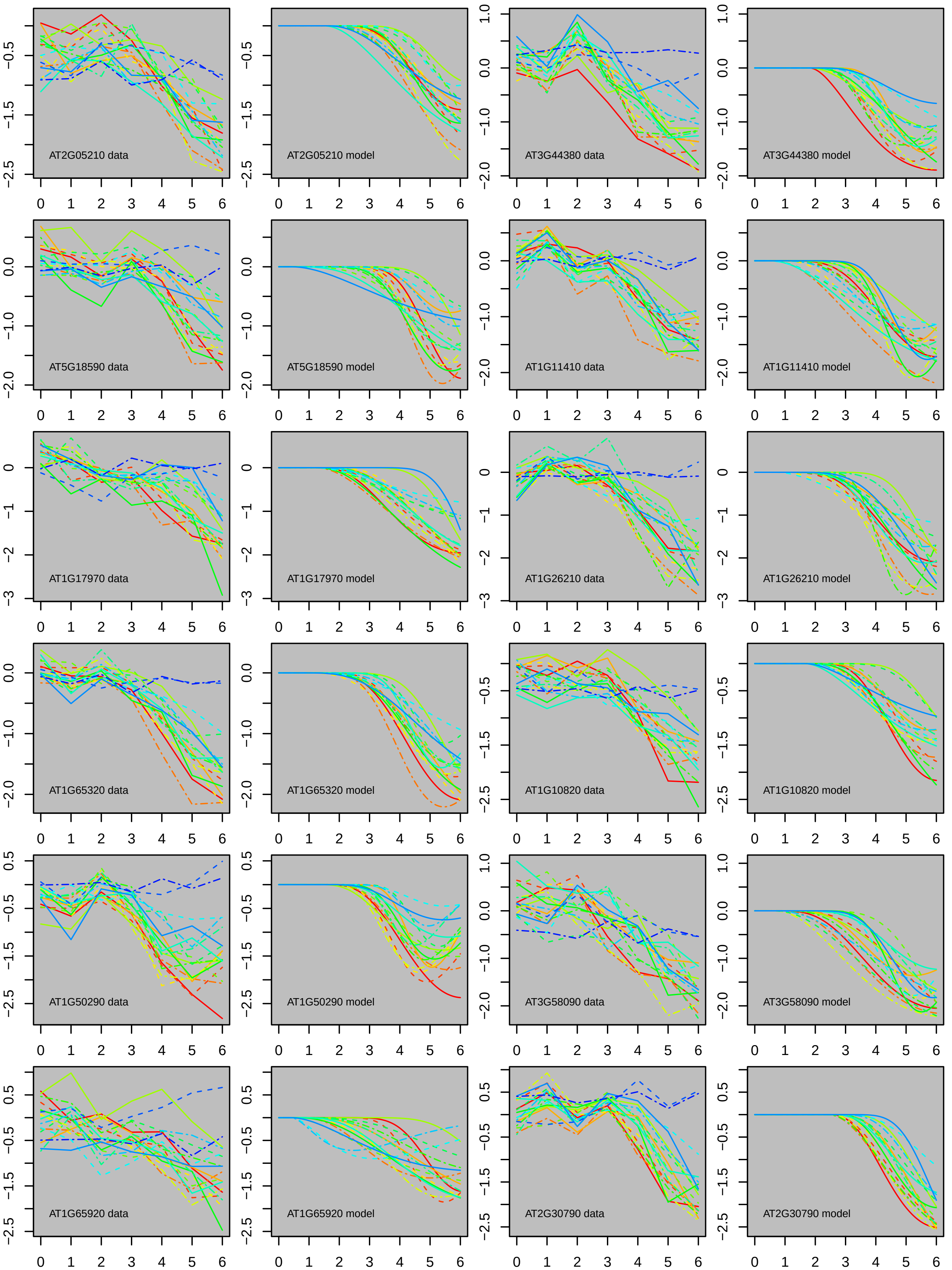
AT2G05210 data
AT2G05210 model
AT3G44380 data
AT3G44380 model
AT5G18590 data
AT5G18590 model
AT1G11410 data
AT1G11410 model
AT1G17970 data
AT1G17970 model
AT1G26210 data
AT1G26210 model
AT1G65320 data
AT1G65320 model
AT1G10820 data
AT1G10820 model
AT1G50290 data
AT1G50290 model
AT3G58090 data
AT3G58090 model
AT1G65920 data
AT1G65920 model
AT2G30790 data
AT2G30790 model

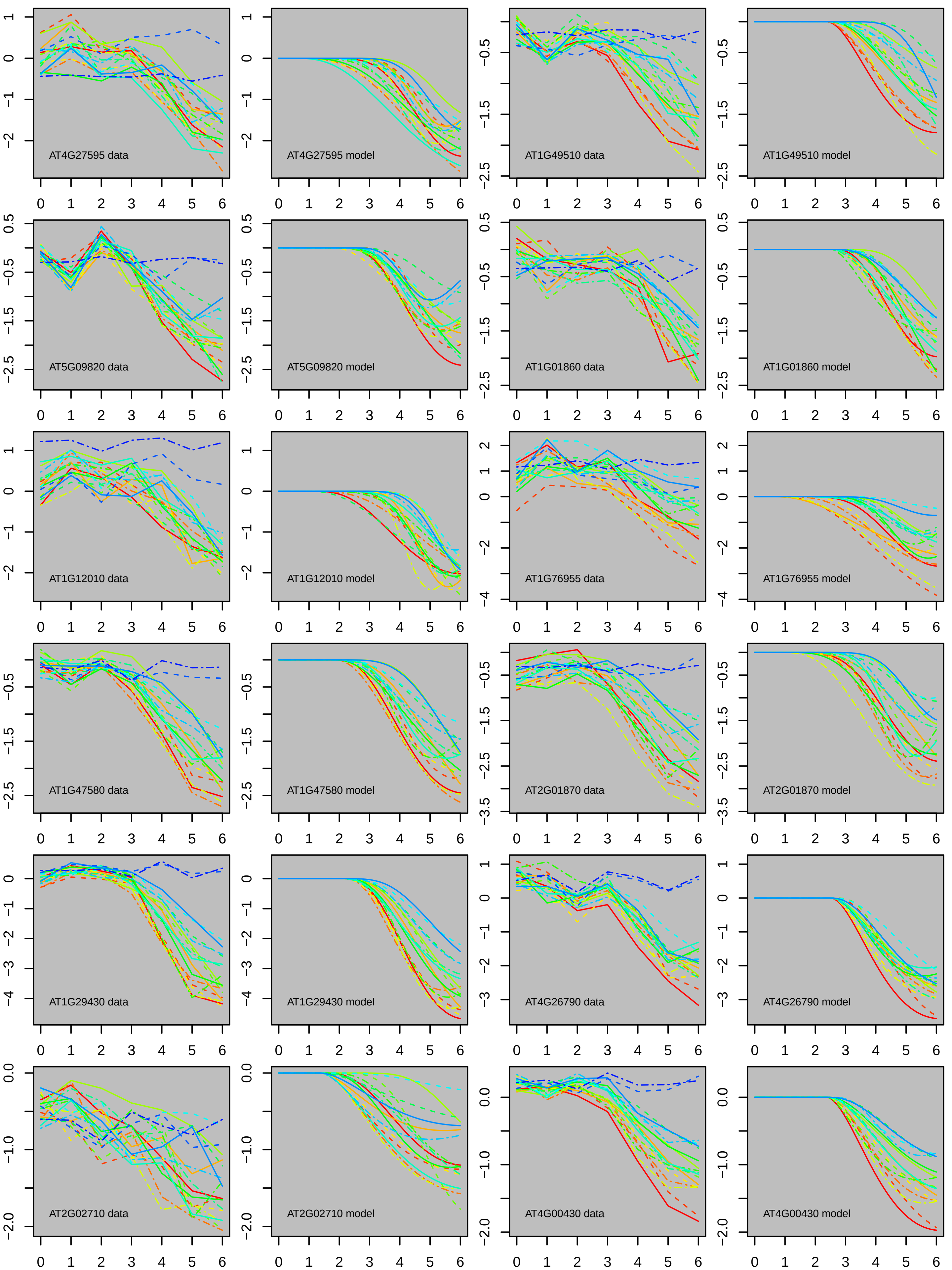

AT4G27595 data
AT4G27595 model
AT1G49510 data
AT1G49510 model
AT5G09820 data
AT5G09820 model
AT1G01860 data
AT1G01860 model
AT1G12010 data
AT1G12010 model
AT1G76955 data
AT1G76955 model
AT1G47580 data
AT1G47580 model
AT2G01870 data
AT2G01870 model
AT1G29430 data
AT1G29430 model
AT4G26790 data
AT4G26790 model
AT2G02710 data
AT2G02710 model
AT4G00430 data
AT4G00430 model

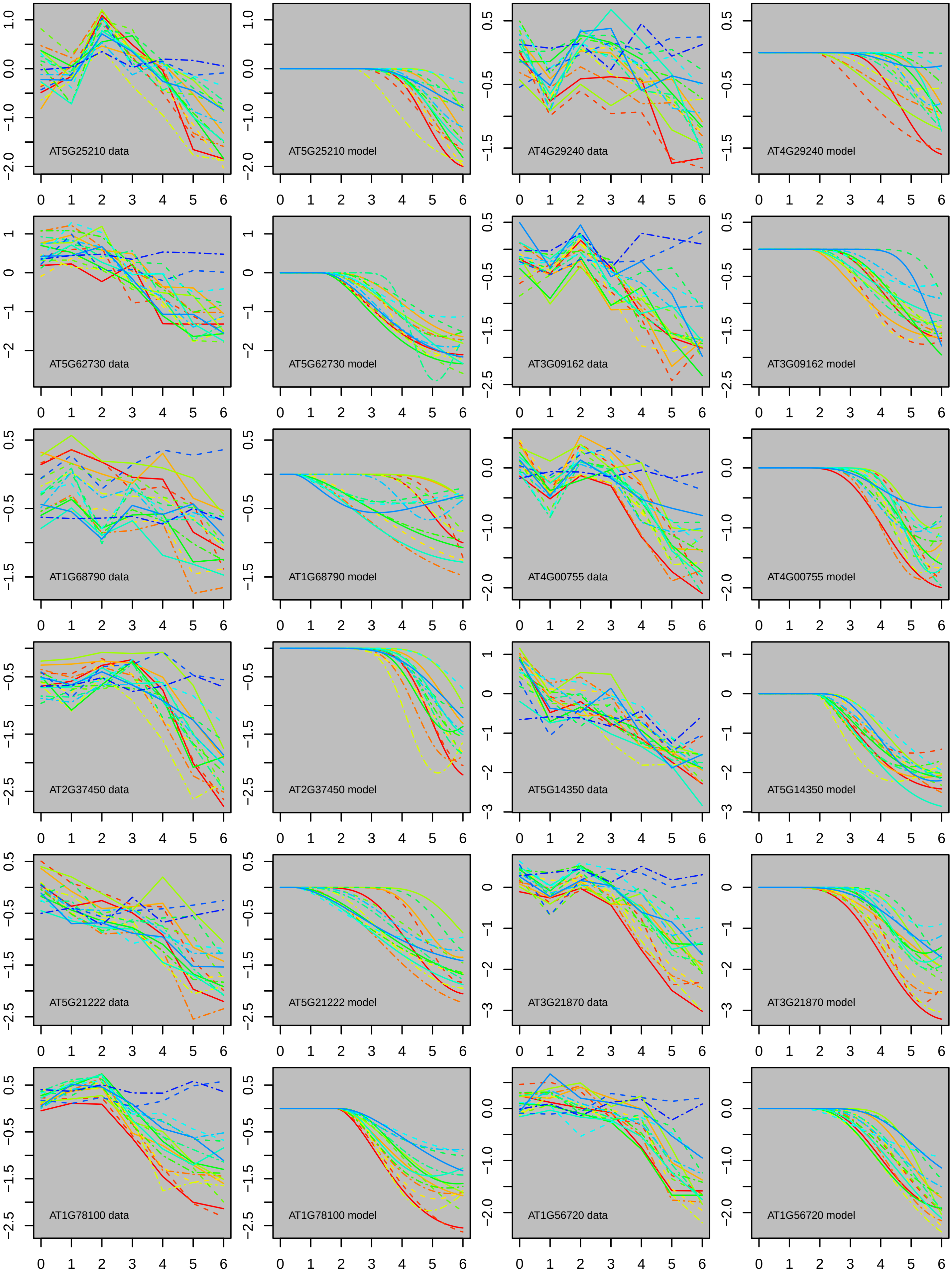
AT5G25210 data
AT5G25210 model
AT4G29240 data
AT4G29240 model
AT5G62730 data
AT5G62730 model
AT3G09162 data
AT3G09162 model
AT1G68790 data
AT1G68790 model
AT4G00755 data
AT4G00755 model
AT2G37450 data
AT2G37450 model
AT5G14350 data
AT5G14350 model
AT5G21222 data
AT5G21222 model
AT3G21870 data
AT3G21870 model
AT1G78100 data
AT1G78100 model
AT1G56720 data
AT1G56720 model

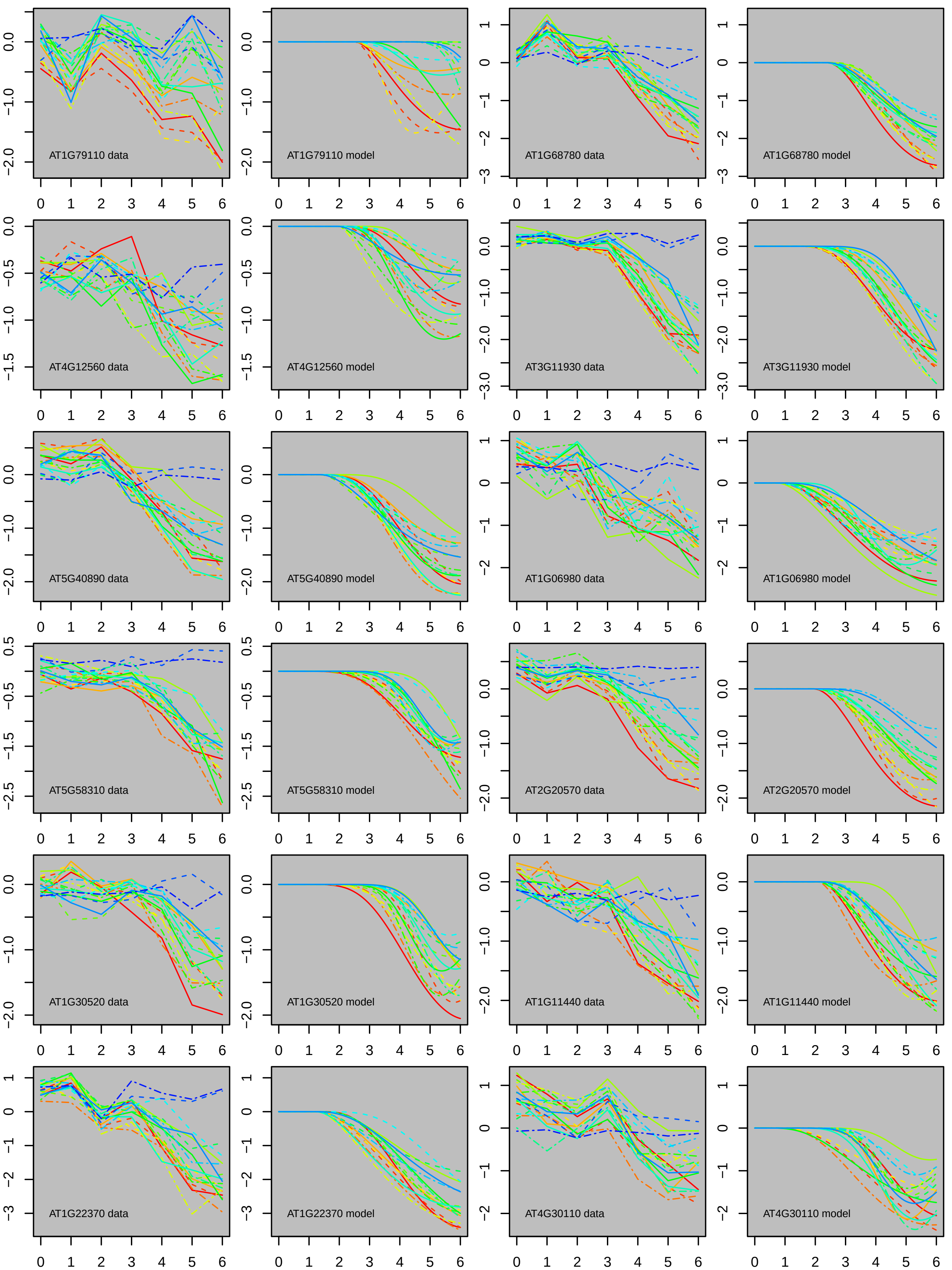

AT1G79110 data
AT1G79110 model
AT1G68780 data
AT1G68780 model
AT4G12560 data
AT4G12560 model
AT3G11930 data
AT3G11930 model
AT5G40890 data
AT5G40890 model
AT1G06980 data
AT1G06980 model
AT5G58310 data
AT5G58310 model
AT2G20570 data
AT2G20570 model
AT1G30520 data
AT1G30520 model
AT1G11440 data
AT1G11440 model
AT1G22370 data
AT1G22370 model
AT4G30110 data
AT4G30110 model

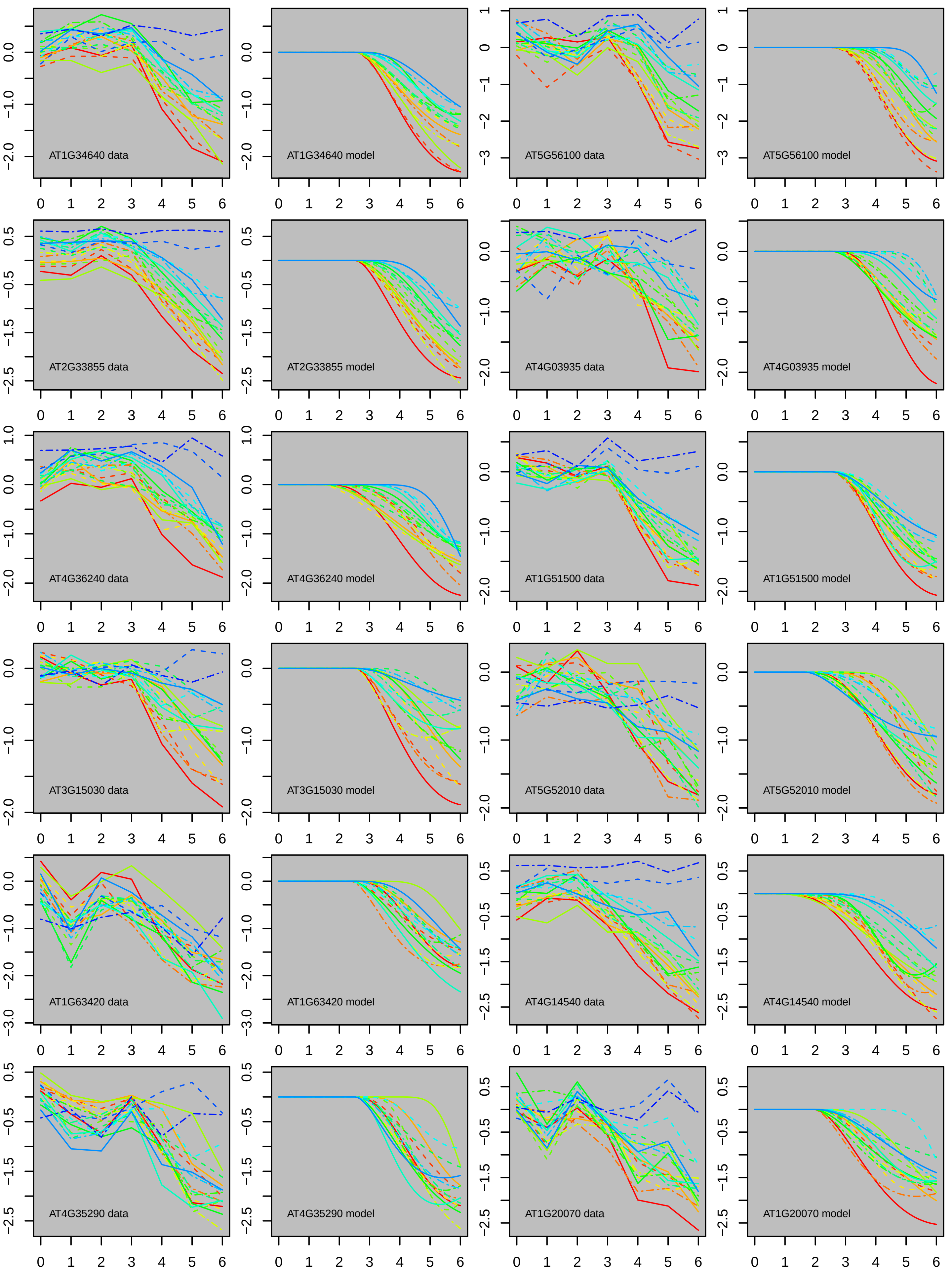

AT1G34640 data
AT1G34640 model
AT5G56100 data
AT5G56100 model
AT2G33855 data
AT2G33855 model
AT4G03935 data
AT4G03935 model
AT4G36240 data
AT4G36240 model
AT1G51500 data
AT1G51500 model
AT3G15030 data
AT3G15030 model
AT5G52010 data
AT5G52010 model
AT1G63420 data
AT1G63420 model
AT4G14540 data
AT4G14540 model
AT4G35290 data
AT4G35290 model
AT1G20070 data
AT1G20070 model

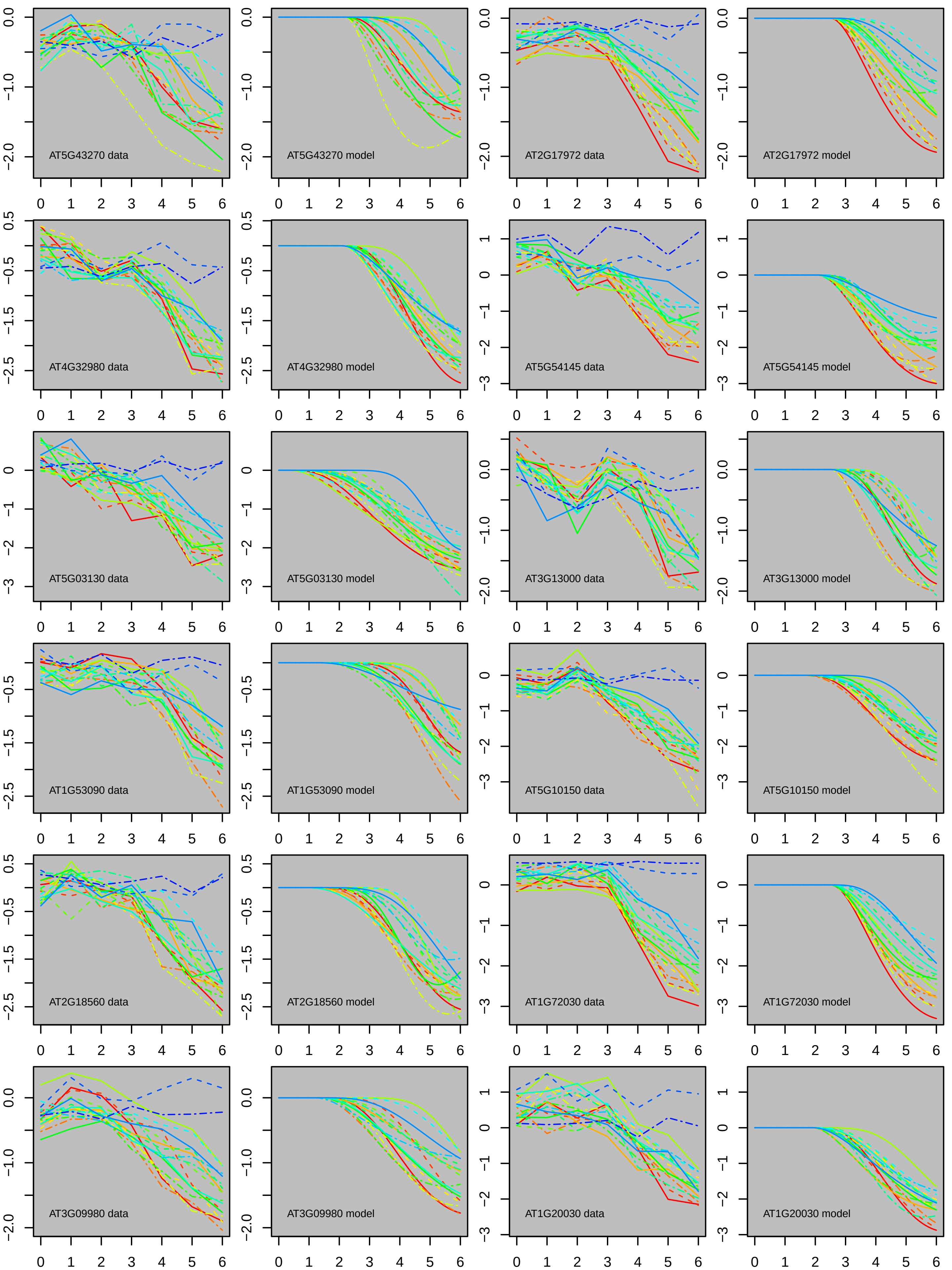
AT5G43270 data
AT5G43270 model
AT2G17972 data
AT2G17972 model
AT4G32980 data
AT4G32980 model
AT5G54145 data
AT5G54145 model
AT5G03130 data
AT5G03130 model
AT3G13000 data
AT3G13000 model
AT1G53090 data
AT1G53090 model
AT5G10150 data
AT5G10150 model
AT2G18560 data
AT2G18560 model
AT1G72030 data
AT1G72030 model
AT3G09980 data
AT3G09980 model
AT1G20030 data
AT1G20030 model

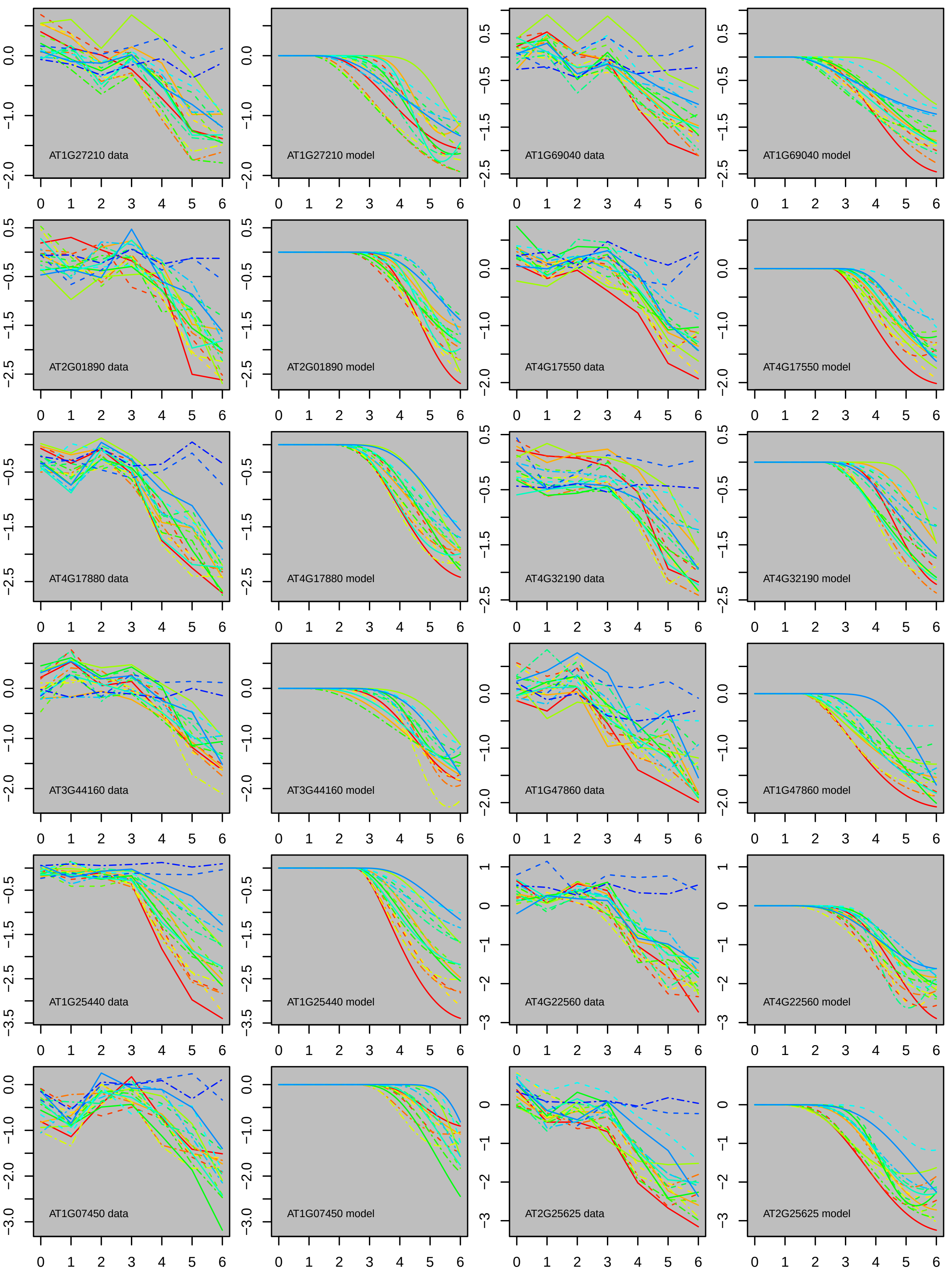

AT1G27210 data
AT1G27210 model
AT1G69040 data
AT1G69040 model
AT2G01890 data
AT2G01890 model
AT4G17550 data
AT4G17550 model
AT4G17880 data
AT4G17880 model
AT4G32190 data
AT4G32190 model
AT3G44160 data
AT3G44160 model
AT1G47860 data
AT1G47860 model
AT1G25440 data
AT1G25440 model
AT4G22560 data
AT4G22560 model
AT1G07450 data
AT1G07450 model
AT2G25625 data
AT2G25625 model

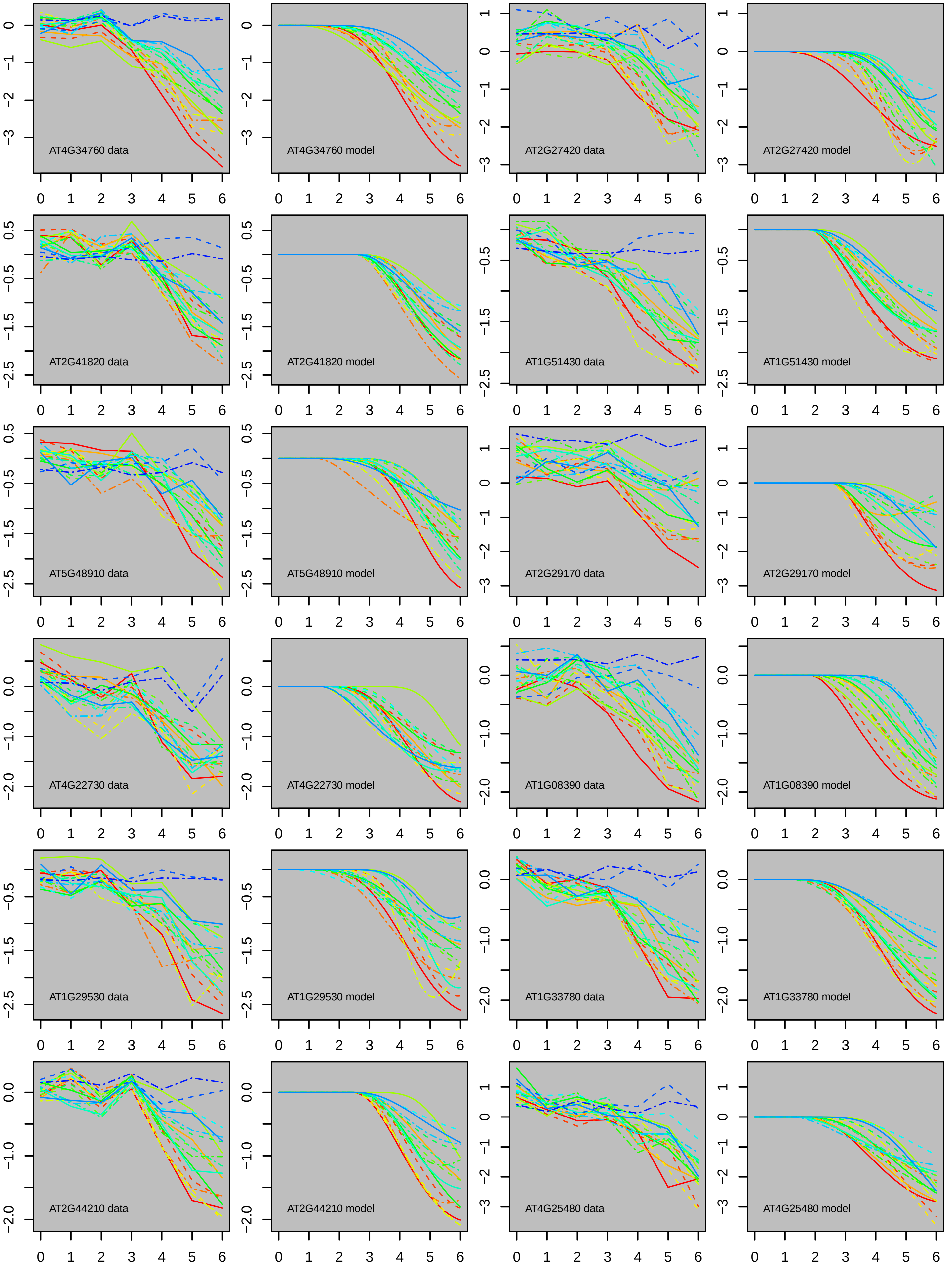

AT4G34760 data
AT4G34760 model
AT2G27420 data
AT2G27420 model
AT2G41820 data
AT2G41820 model
AT1G51430 data
AT1G51430 model
AT5G48910 data
AT5G48910 model
AT2G29170 data
AT2G29170 model
AT4G22730 data
AT4G22730 model
AT1G08390 data
AT1G08390 model
AT1G29530 data
AT1G29530 model
AT1G33780 data
AT1G33780 model
AT2G44210 data
AT2G44210 model
AT4G25480 data
AT4G25480 model

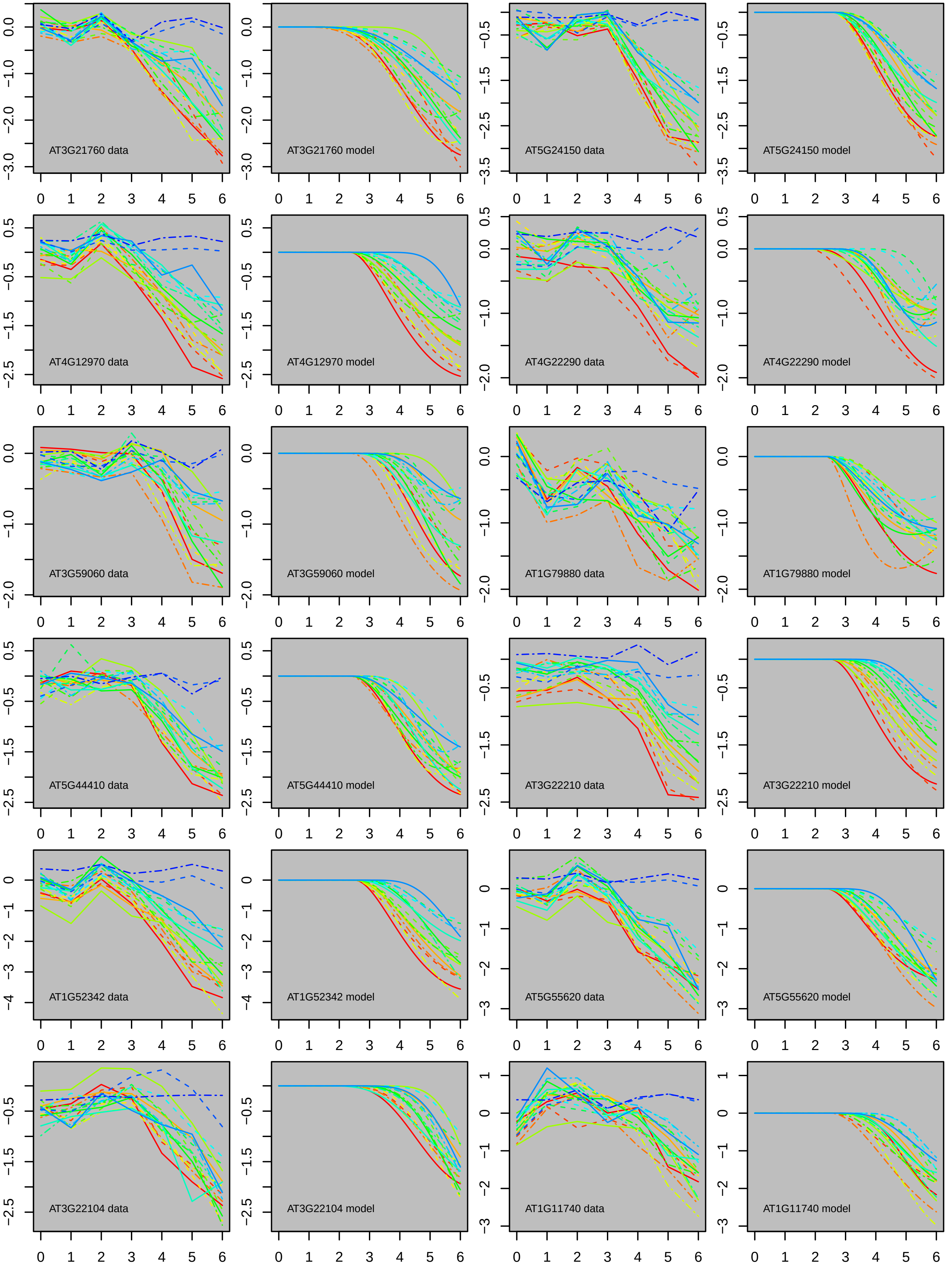
AT3G21760 data
AT3G21760 model
AT5G24150 data
AT5G24150 model
AT4G12970 data
AT4G12970 model
AT4G22290 data
AT4G22290 model
AT3G59060 data
AT3G59060 model
AT1G79880 data
AT1G79880 model
AT5G44410 data
AT5G44410 model
AT3G22210 data
AT3G22210 model
AT1G52342 data
AT1G52342 model
AT5G55620 data
AT5G55620 model
AT3G22104 data
AT3G22104 model
AT1G11740 data
AT1G11740 model

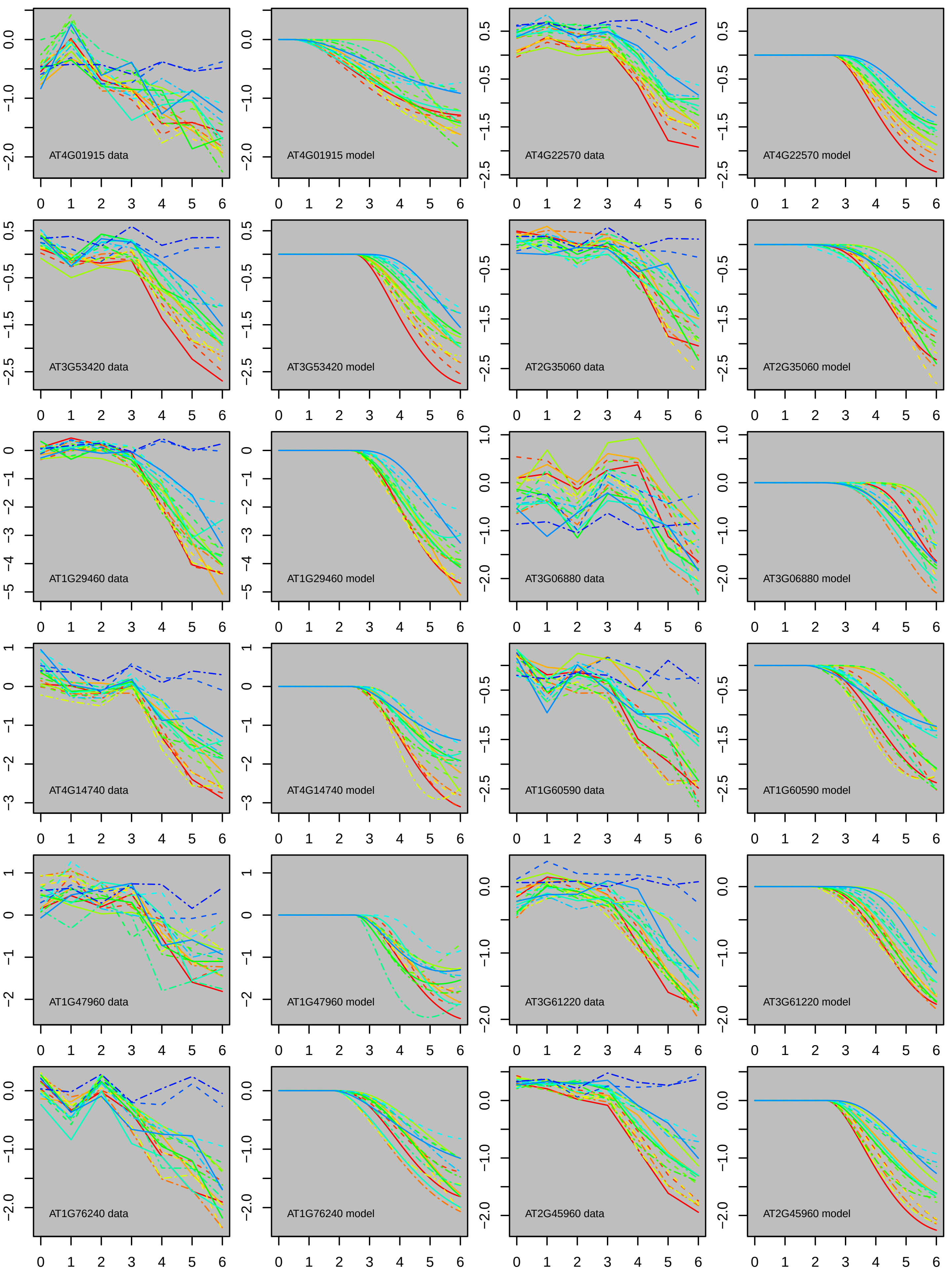
AT4G01915 data
AT4G01915 model
AT4G22570 data
AT4G22570 model
AT3G53420 data
AT3G53420 model
AT2G35060 data
AT2G35060 model
AT1G29460 data
AT1G29460 model
AT3G06880 data
AT3G06880 model
AT4G14740 data
AT4G14740 model
AT1G60590 data
AT1G60590 model
AT1G47960 data
AT1G47960 model
AT3G61220 data
AT3G61220 model
AT1G76240 data
AT1G76240 model
AT2G45960 data
AT2G45960 model

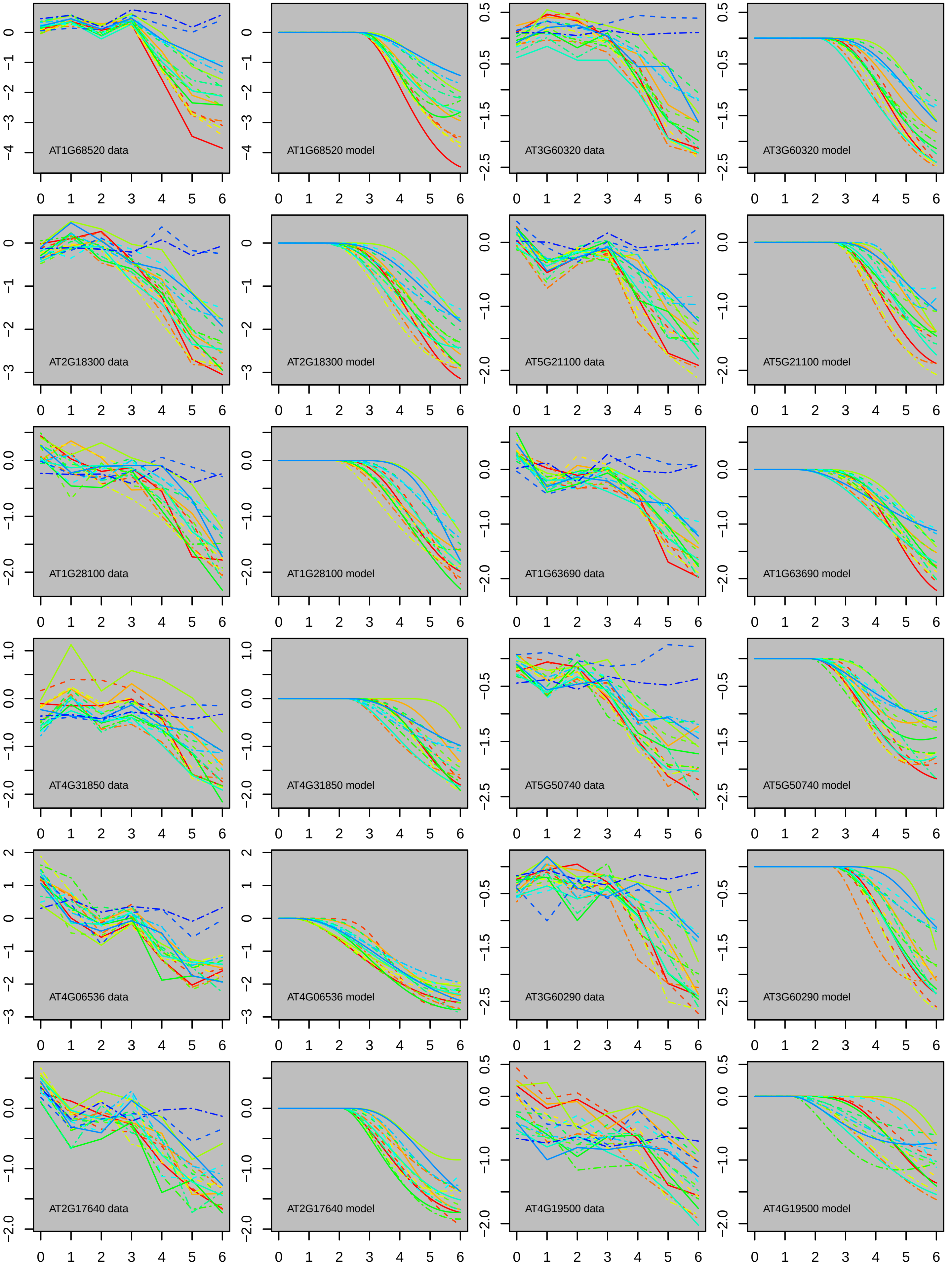

AT1G68520 data
AT1G68520 model
AT3G60320 data
AT3G60320 model
AT2G18300 data
AT2G18300 model
AT5G21100 data
AT5G21100 model
AT1G28100 data
AT1G28100 model
AT1G63690 data
AT1G63690 model
AT4G31850 data
AT4G31850 model
AT5G50740 data
AT5G50740 model
AT4G06536 data
AT4G06536 model
AT3G60290 data
AT3G60290 model
AT2G17640 data
AT2G17640 model
AT4G19500 data
AT4G19500 model

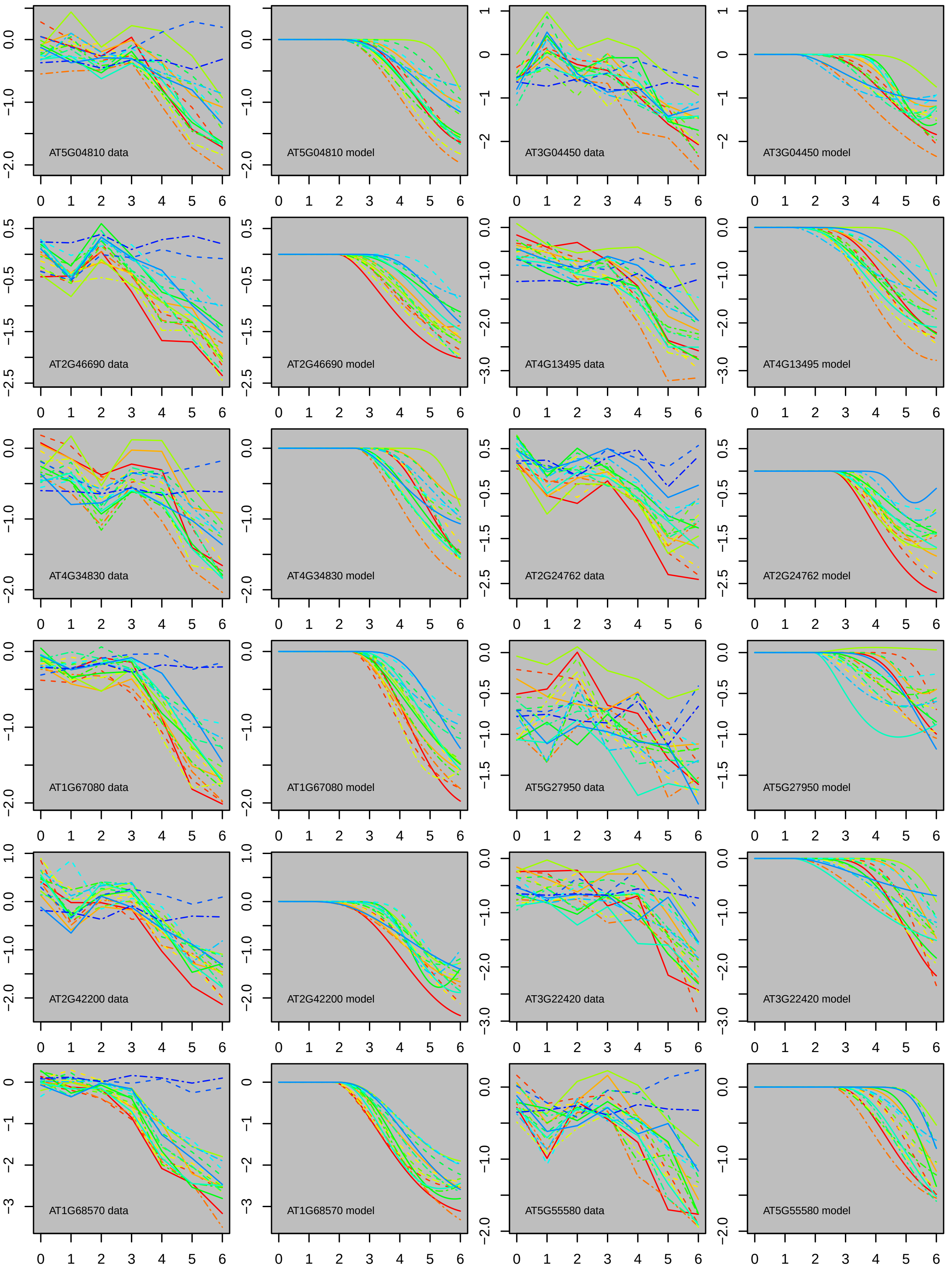
AT5G04810 data
AT5G04810 model
AT3G04450 data
AT3G04450 model
AT2G46690 data
AT2G46690 model
AT4G13495 data
AT4G13495 model
AT4G34830 data
AT4G34830 model
AT2G24762 data
AT2G24762 model
AT1G67080 data
AT1G67080 model
AT5G27950 data
AT5G27950 model
AT2G42200 data
AT2G42200 model
AT3G22420 data
AT3G22420 model
AT1G68570 data
AT1G68570 model
AT5G55580 data
AT5G55580 model

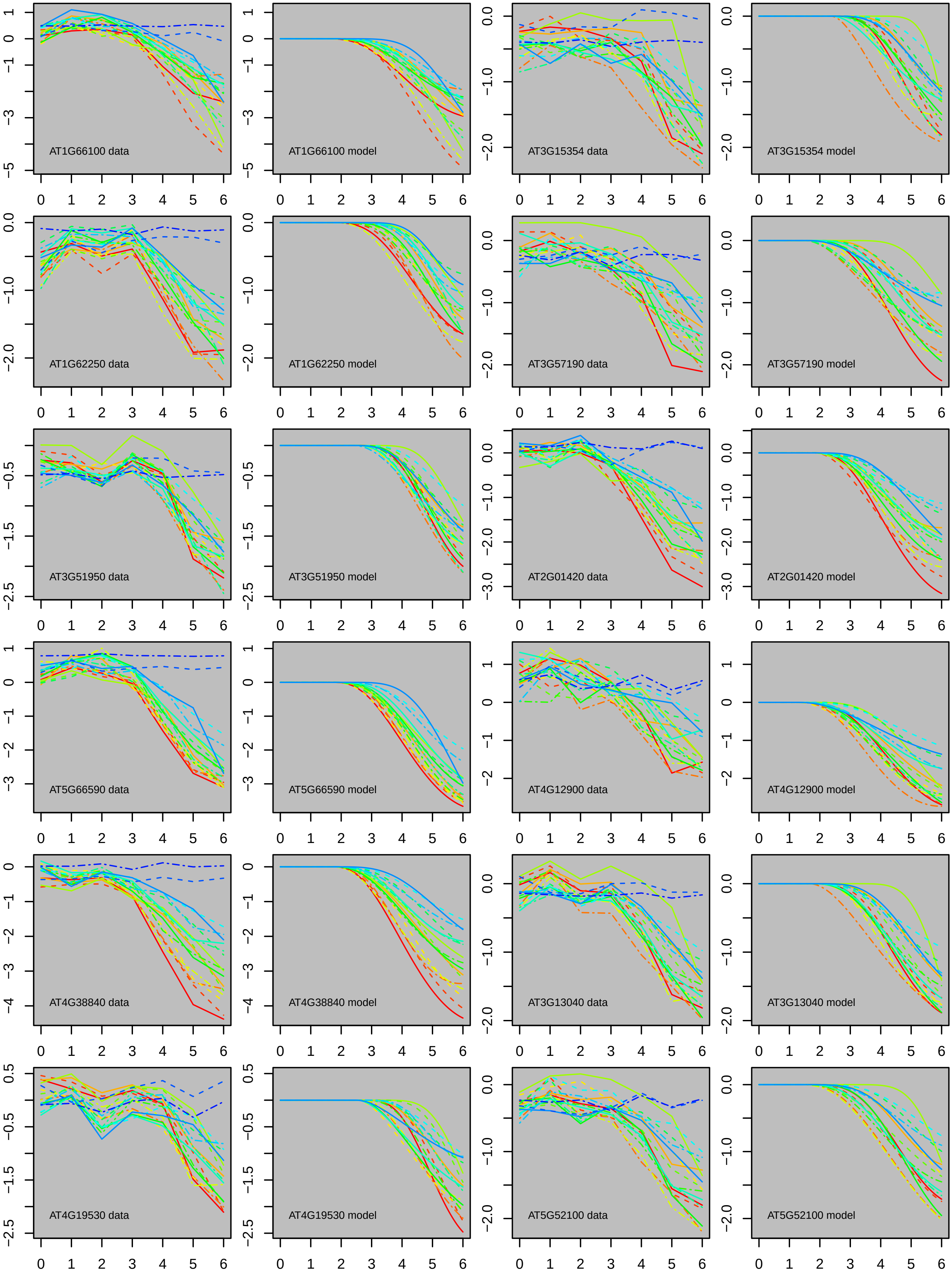

AT1G66100 data
AT1G66100 model
AT3G15354 data
AT3G15354 model
AT1G62250 data
AT1G62250 model
AT3G57190 data
AT3G57190 model
AT3G51950 data
AT3G51950 model
AT2G01420 data
AT2G01420 model
AT5G66590 data
AT5G66590 model
AT4G12900 data
AT4G12900 model
AT4G38840 data
AT4G38840 model
AT3G13040 data
AT3G13040 model
AT4G19530 data
AT4G19530 model
AT5G52100 data
AT5G52100 model

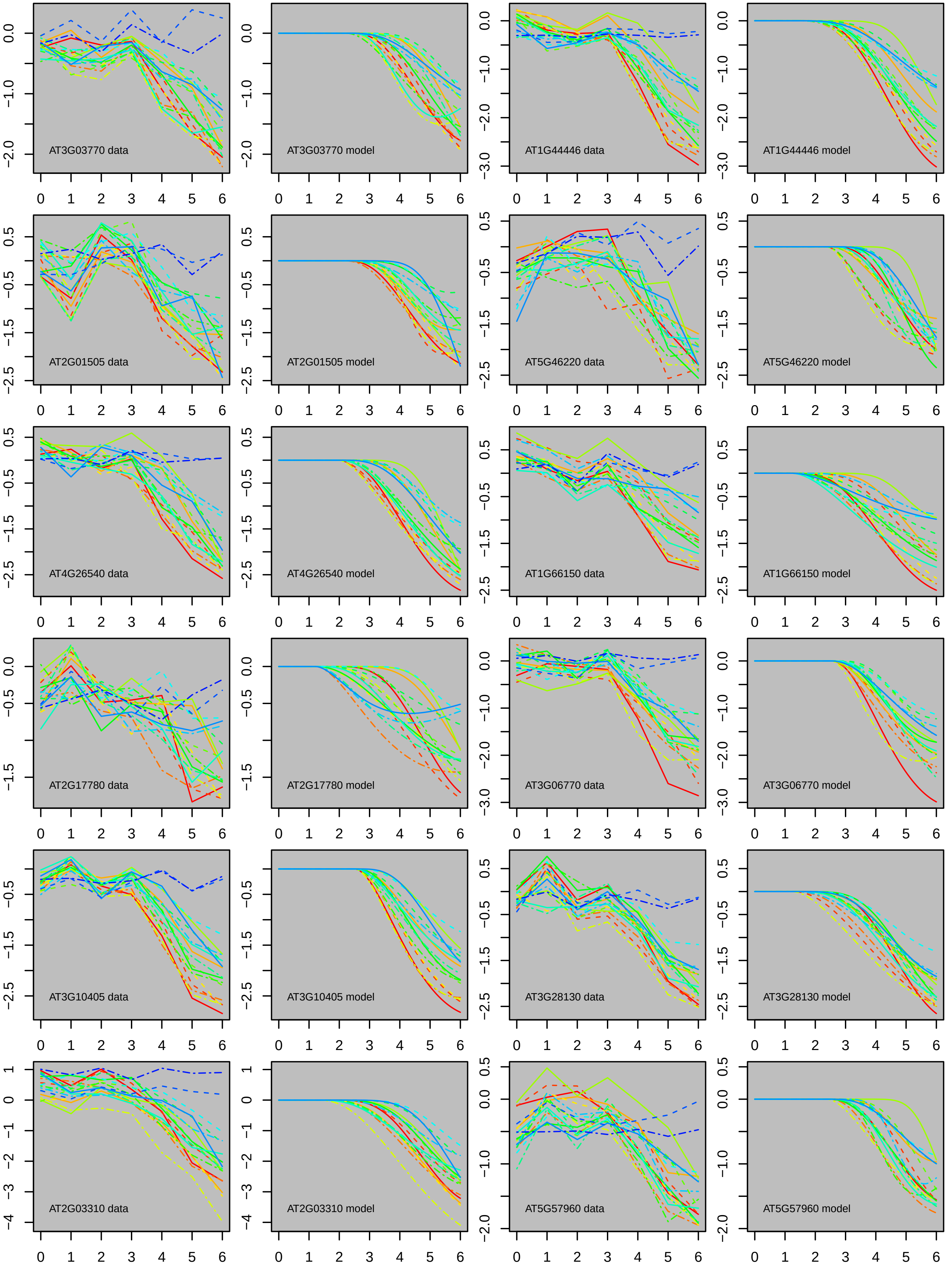
AT3G03770 data
AT3G03770 model
AT1G44446 data
AT1G44446 model
AT2G01505 data
AT2G01505 model
AT5G46220 data
AT5G46220 model
AT4G26540 data
AT4G26540 model
AT1G66150 data
AT1G66150 model
AT2G17780 data
AT2G17780 model
AT3G06770 data
AT3G06770 model
AT3G10405 data
AT3G10405 model
AT3G28130 data
AT3G28130 model
AT2G03310 data
AT2G03310 model
AT5G57960 data
AT5G57960 model

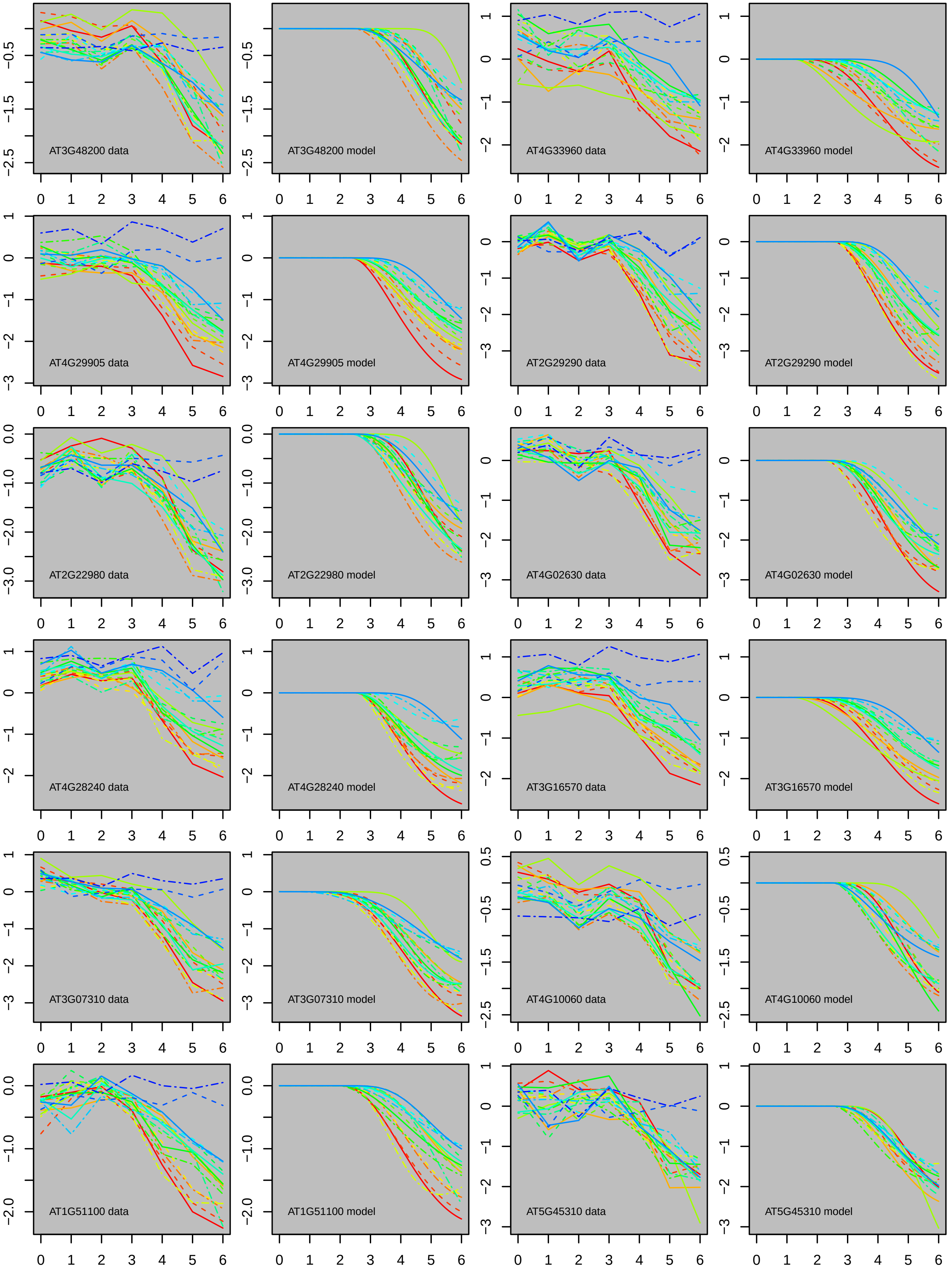

AT3G48200 data
AT3G48200 model
AT4G33960 data
AT4G33960 model
AT4G29905 data
AT4G29905 model
AT2G29290 data
AT2G29290 model
AT2G22980 data
AT2G22980 model
AT4G02630 data
AT4G02630 model
AT4G28240 data
AT4G28240 model
AT3G16570 data
AT3G16570 model
AT3G07310 data
AT3G07310 model
AT4G10060 data
AT4G10060 model
AT1G51100 data
AT1G51100 model
AT5G45310 data
AT5G45310 model

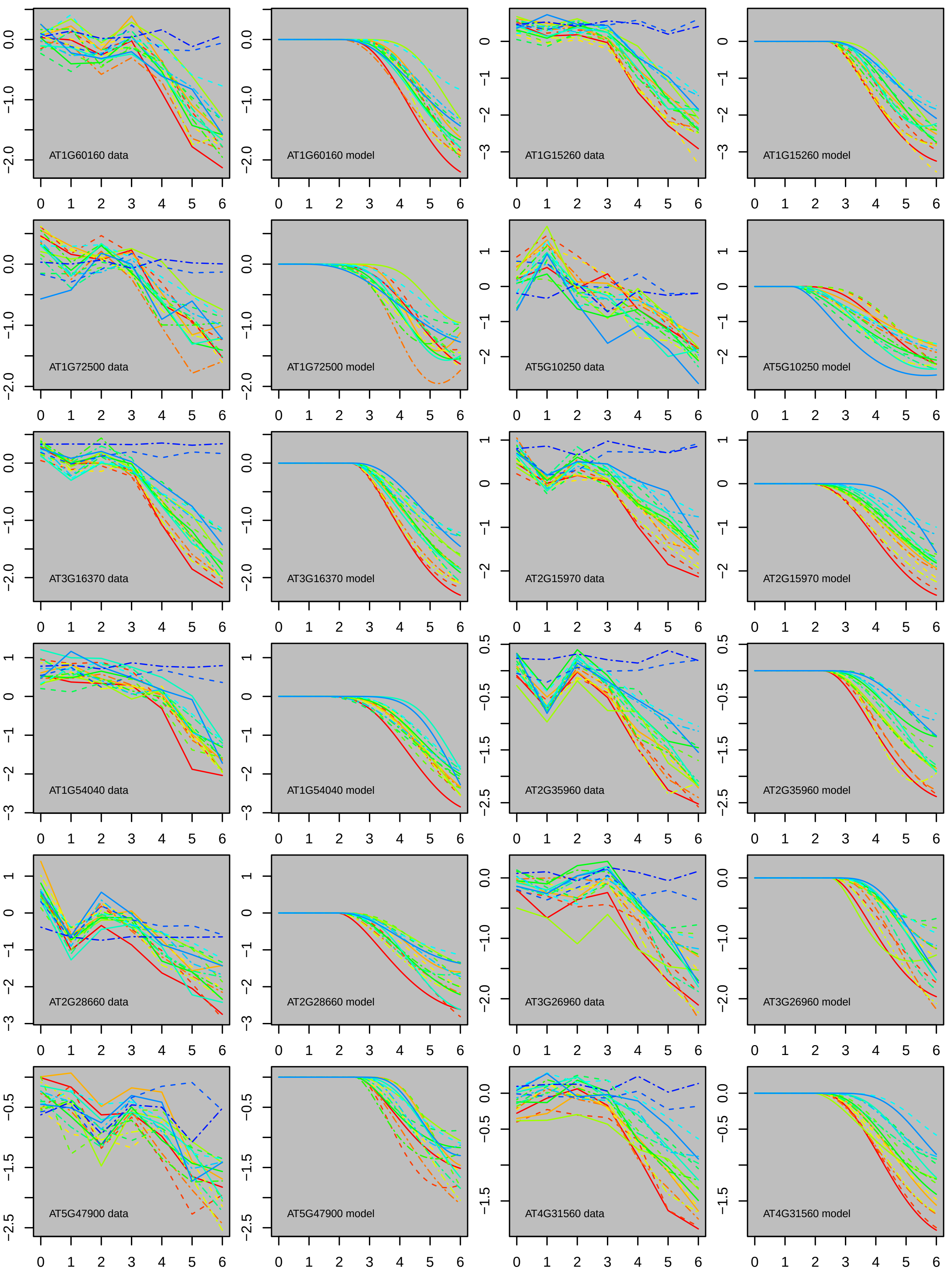

AT1G60160 data
AT1G60160 model
AT1G15260 data
AT1G15260 model
AT1G72500 data
AT1G72500 model
AT5G10250 data
AT5G10250 model
AT3G16370 data
AT3G16370 model
AT2G15970 data
AT2G15970 model
AT1G54040 data
AT1G54040 model
AT2G35960 data
AT2G35960 model
AT2G28660 data
AT2G28660 model
AT3G26960 data
AT3G26960 model
AT5G47900 data
AT5G47900 model
AT4G31560 data
AT4G31560 model

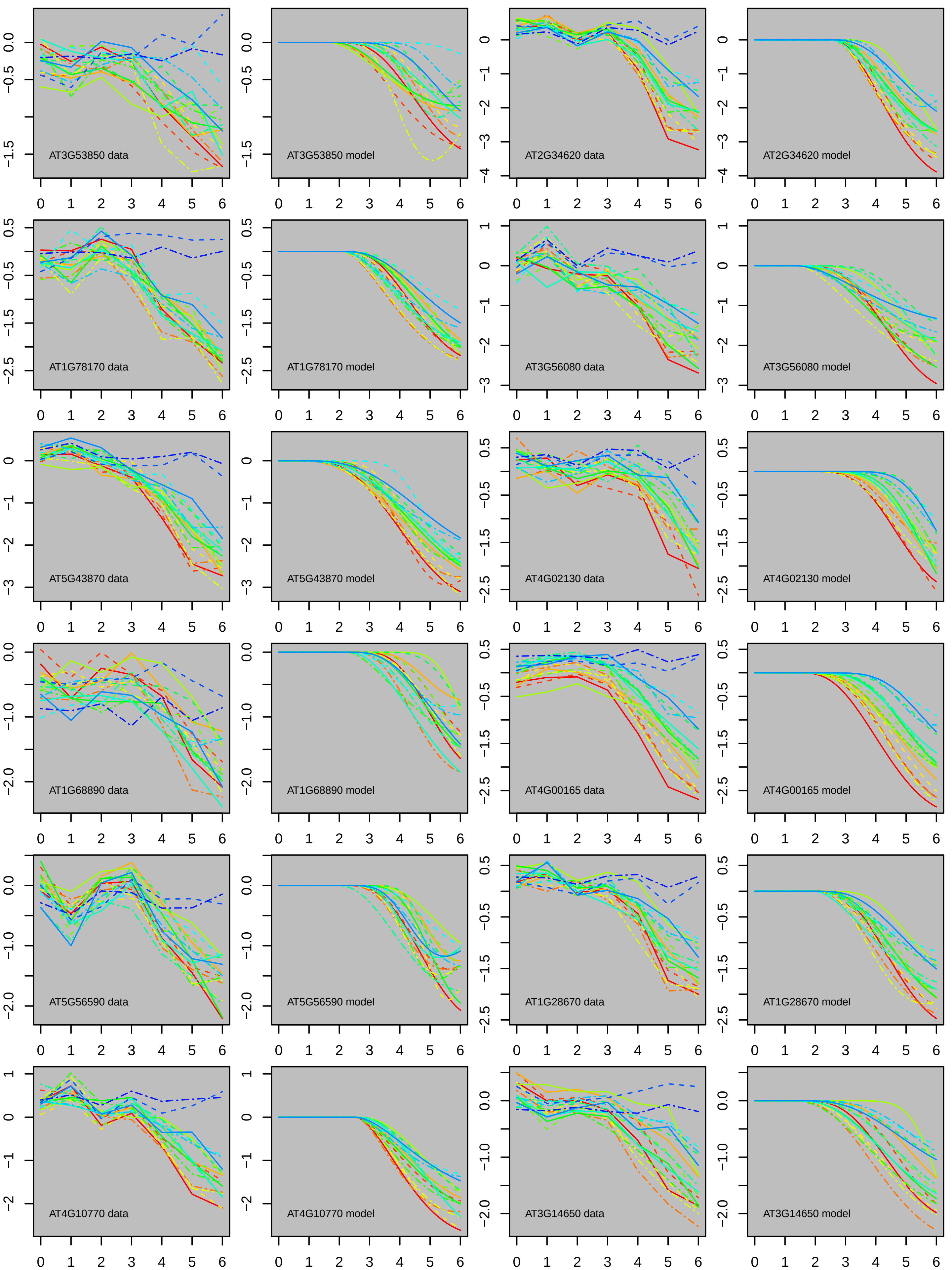
AT3G53850 data
AT3G53850 model
AT2G34620 data
AT2G34620 model
AT1G78170 data
AT1G78170 model
AT3G56080 data
AT3G56080 model
AT5G43870 data
AT5G43870 model
AT4G02130 data
AT4G02130 model
AT1G68890 data
AT1G68890 model
AT4G00165 data
AT4G00165 model
AT5G56590 data
AT5G56590 model
AT1G28670 data
AT1G28670 model
AT4G10770 data
AT4G10770 model
AT3G14650 data
AT3G14650 model

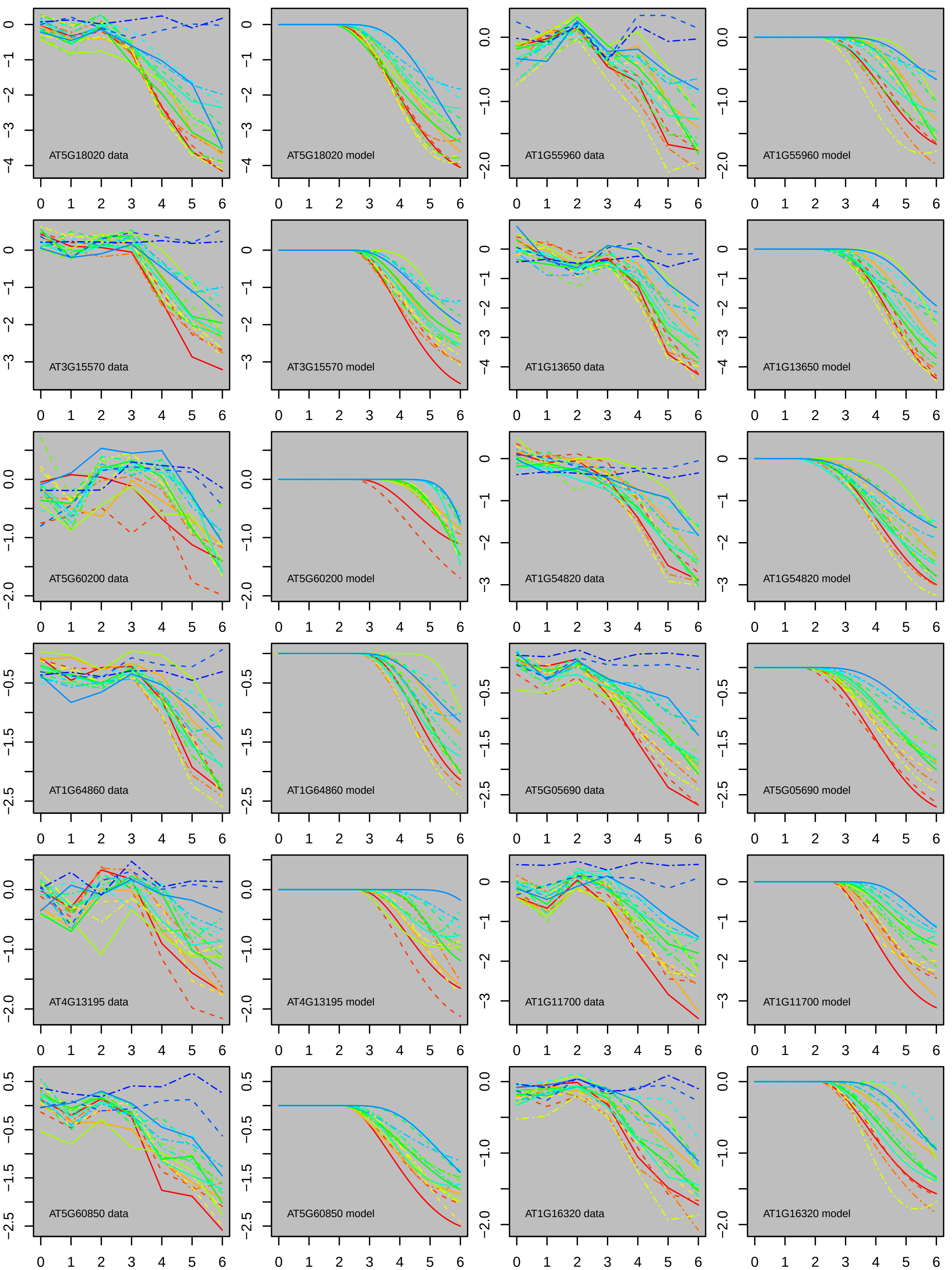

AT5G18020 data
AT5G18020 model
AT1G55960 data
AT1G55960 model
AT3G15570 data
AT3G15570 model
AT1G13650 data
AT1G13650 model
AT5G60200 data
AT5G60200 model
AT1G54820 data
AT1G54820 model
AT1G64860 data
AT1G64860 model
AT5G05690 data
AT5G05690 model
AT4G13195 data
AT4G13195 model
AT1G11700 data
AT1G11700 model
AT5G60850 data
AT5G60850 model
AT1G16320 data
AT1G16320 model
0 1 2 3 4 5 6

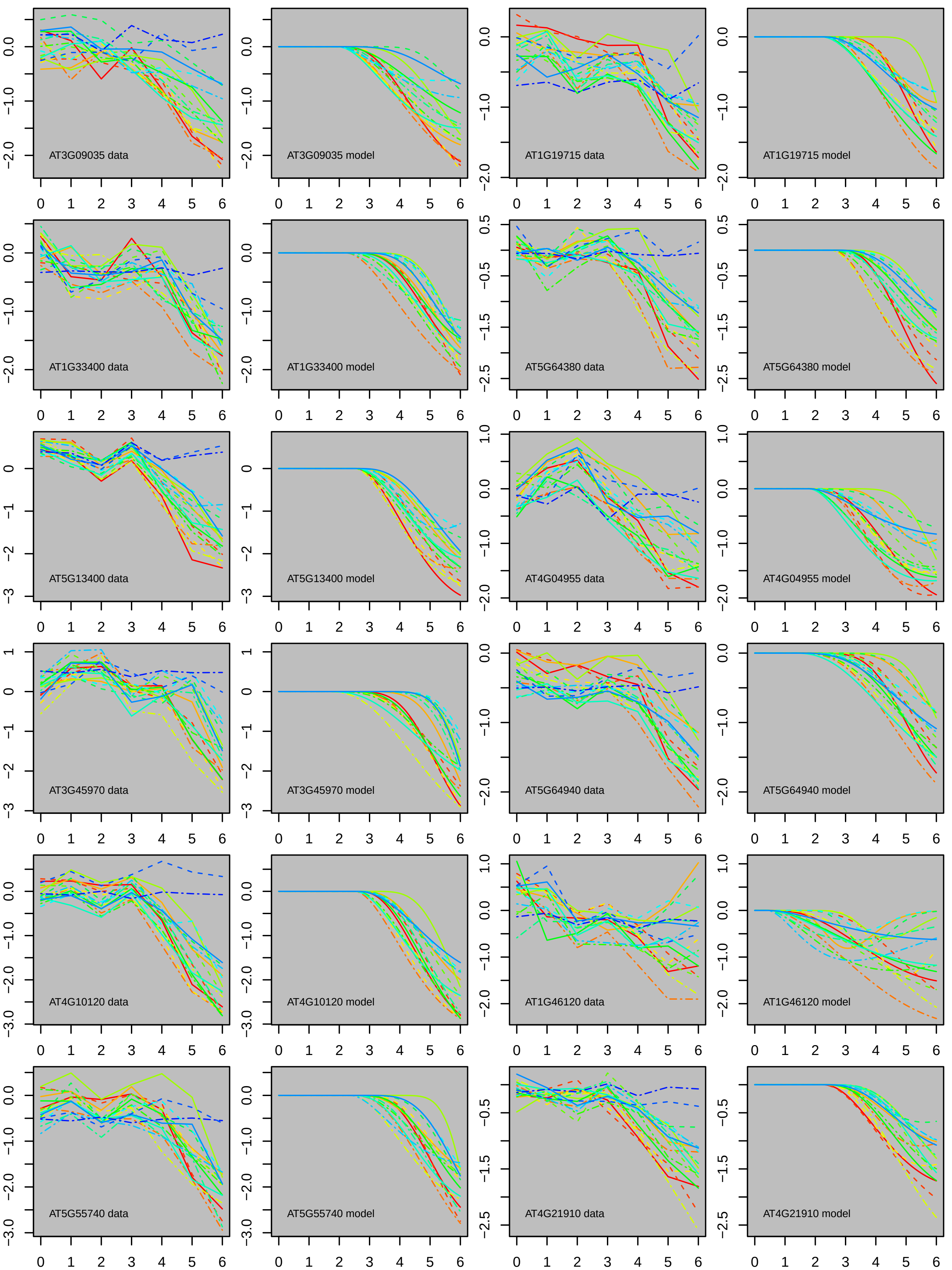
AT3G09035 data
AT3G09035 model
AT1G19715 data
AT1G19715 model
AT1G33400 data
AT1G33400 model
AT5G64380 data
AT5G64380 model
AT5G13400 data
AT5G13400 model
AT4G04955 data
AT4G04955 model
AT3G45970 data
AT3G45970 model
AT5G64940 data
AT5G64940 model
AT4G10120 data
AT4G10120 model
AT1G46120 data
AT1G46120 model
AT5G55740 data
AT5G55740 model
AT4G21910 data
AT4G21910 model

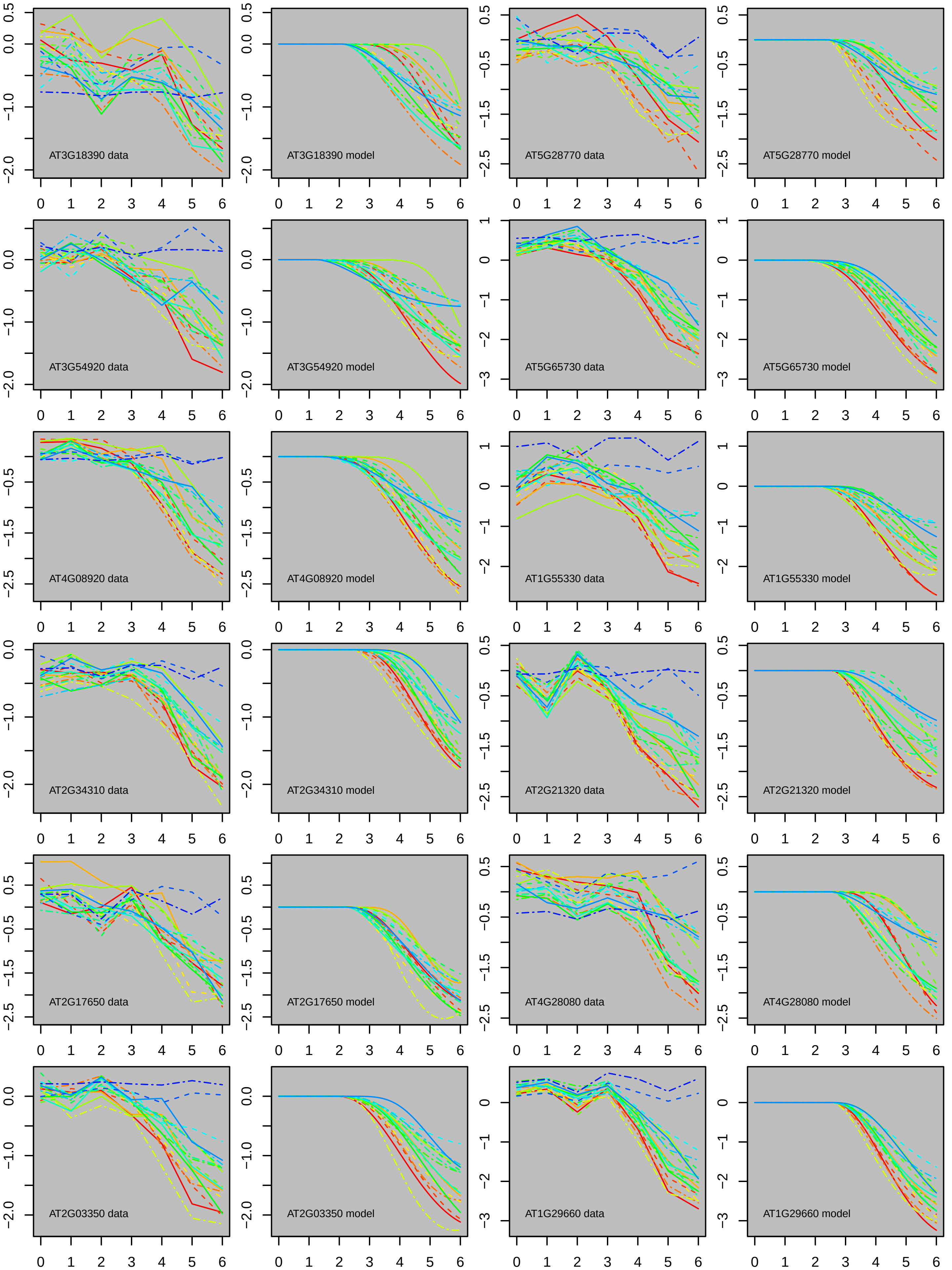
AT3G18390 data
AT3G18390 model
AT5G28770 data
AT5G28770 model
AT3G54920 data
AT3G54920 model
AT5G65730 data
AT5G65730 model
AT4G08920 data
AT4G08920 model
AT1G55330 data
AT1G55330 model
AT2G34310 data
AT2G34310 model
AT2G21320 data
AT2G21320 model
AT2G17650 data
AT2G17650 model
AT4G28080 data
AT4G28080 model
AT2G03350 data
AT2G03350 model
AT1G29660 data
AT1G29660 model

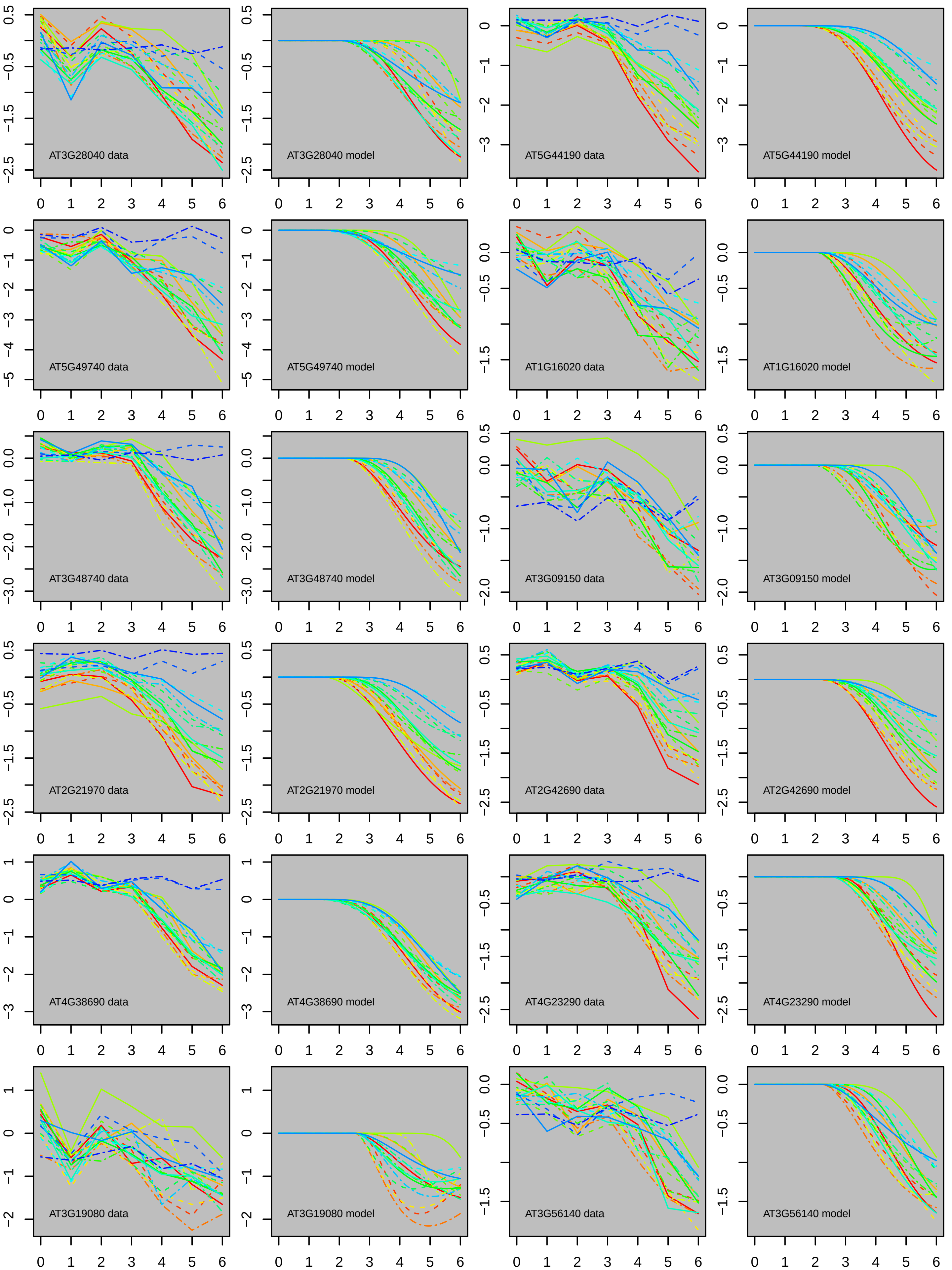
AT3G28040 data
AT3G28040 model
AT5G44190 data
AT5G44190 model
AT5G49740 data
AT5G49740 model
AT1G16020 data
AT1G16020 model
AT3G48740 data
AT3G48740 model
AT3G09150 data
AT3G09150 model
AT2G21970 data
AT2G21970 model
AT2G42690 data
AT2G42690 model
AT4G38690 data
AT4G38690 model
AT4G23290 data
AT4G23290 model
AT3G19080 data
AT3G19080 model
AT3G56140 data
AT3G56140 model

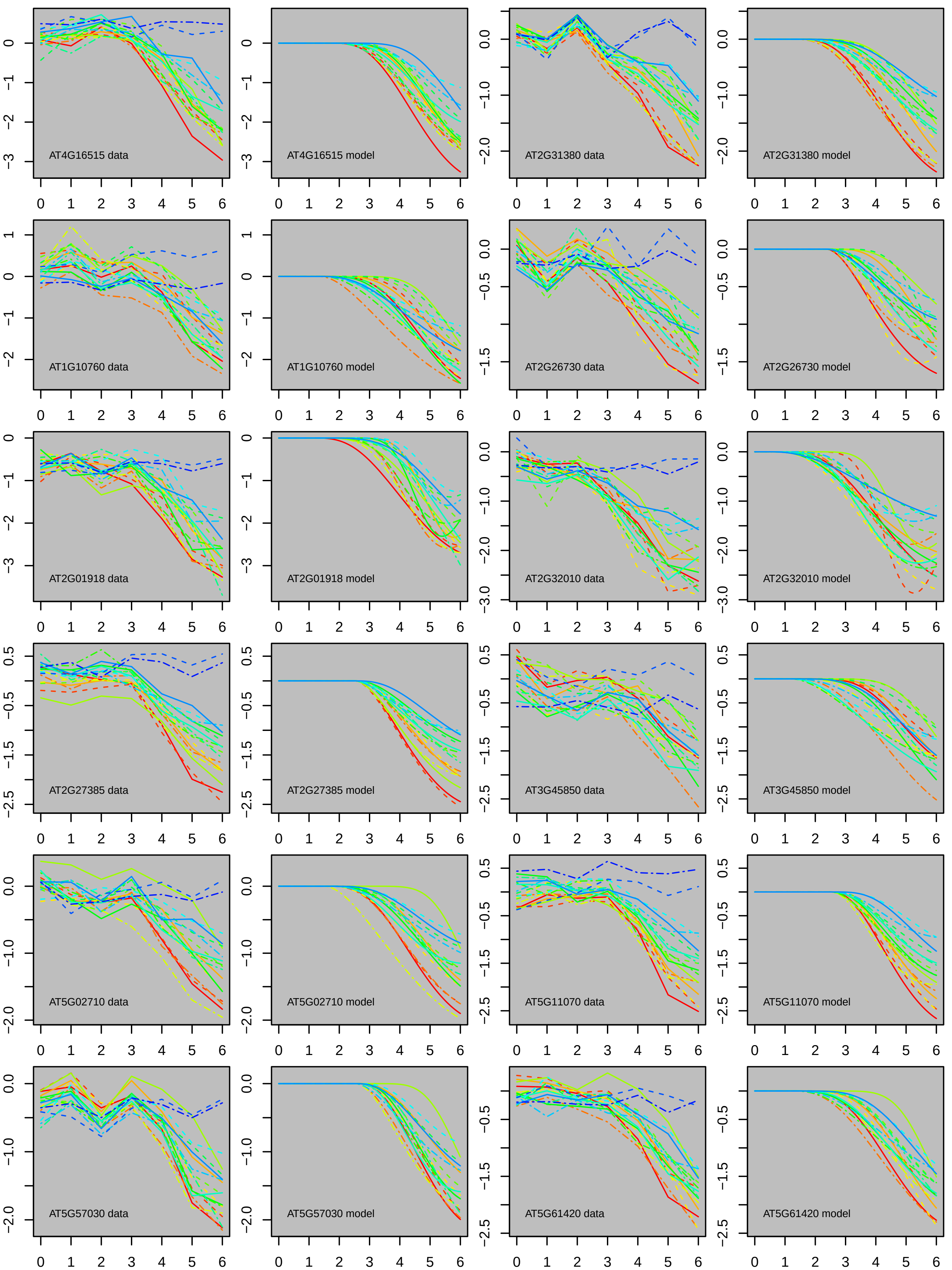
AT4G16515 data
AT4G16515 model
AT2G31380 data
AT2G31380 model
AT1G10760 data
AT1G10760 model
AT2G26730 data
AT2G26730 model
AT2G01918 data
AT2G01918 model
AT2G32010 data
AT2G32010 model
AT2G27385 data
AT2G27385 model
AT3G45850 data
AT3G45850 model
AT5G02710 data
AT5G02710 model
AT5G11070 data
AT5G11070 model
AT5G57030 data
AT5G57030 model
AT5G61420 data
AT5G61420 model

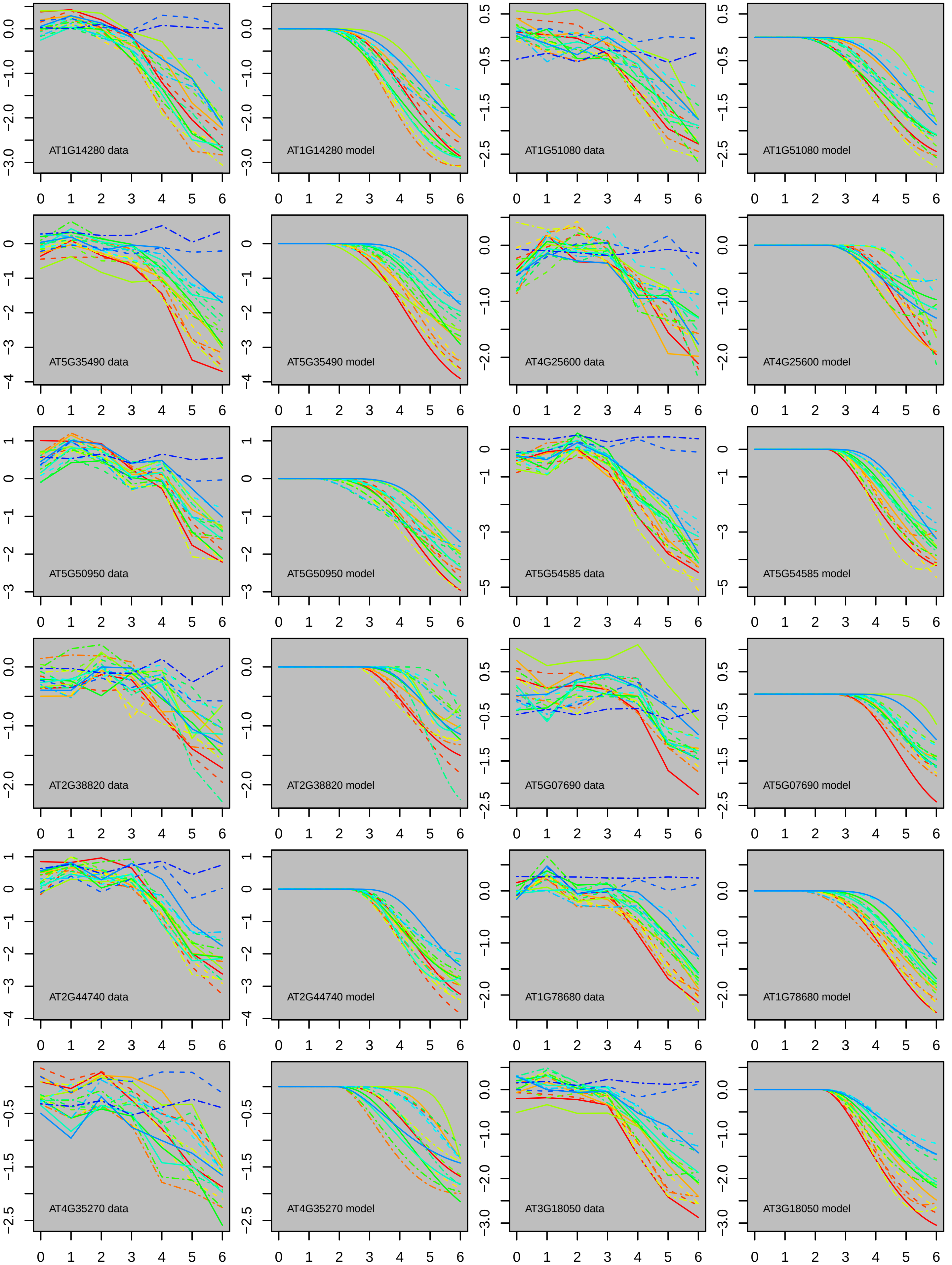

AT1G14280 data
AT1G14280 model
AT1G51080 data
AT1G51080 model
AT5G35490 data
AT5G35490 model
AT4G25600 data
AT4G25600 model
AT5G50950 data
AT5G50950 model
AT5G54585 data
AT5G54585 model
AT2G38820 data
AT2G38820 model
AT5G07690 data
AT5G07690 model
AT2G44740 data
AT2G44740 model
AT1G78680 data
AT1G78680 model
AT4G35270 data
AT4G35270 model
AT3G18050 data
AT3G18050 model

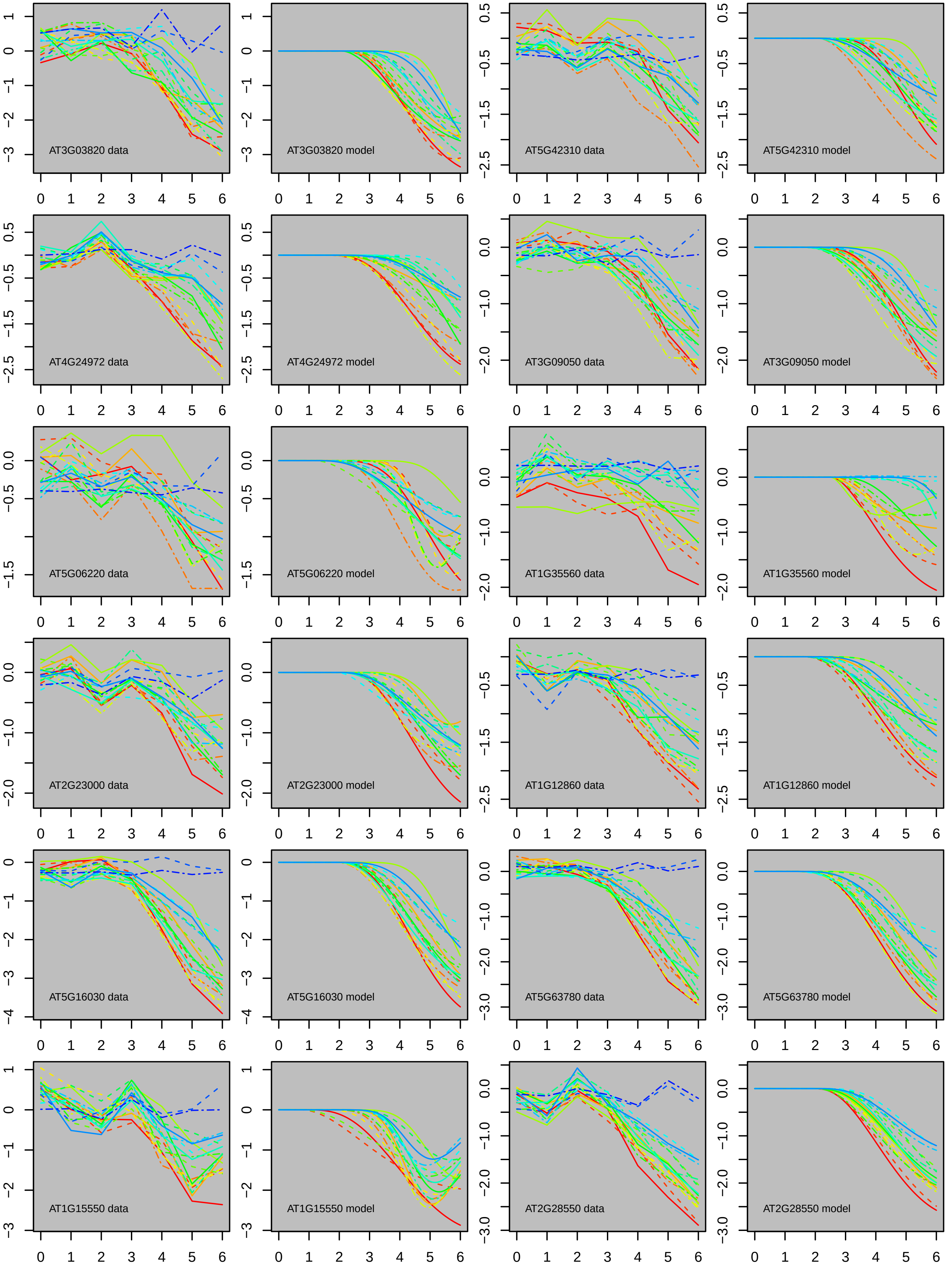
AT3G03820 data
AT3G03820 model
AT5G42310 data
AT5G42310 model
AT4G24972 data
AT4G24972 model
AT3G09050 data
AT3G09050 model
AT5G06220 data
AT5G06220 model
AT1G35560 data
AT1G35560 model
AT2G23000 data
AT2G23000 model
AT1G12860 data
AT1G12860 model
AT5G16030 data
AT5G16030 model
AT5G63780 data
AT5G63780 model
AT1G15550 data
AT1G15550 model
AT2G28550 data
AT2G28550 model

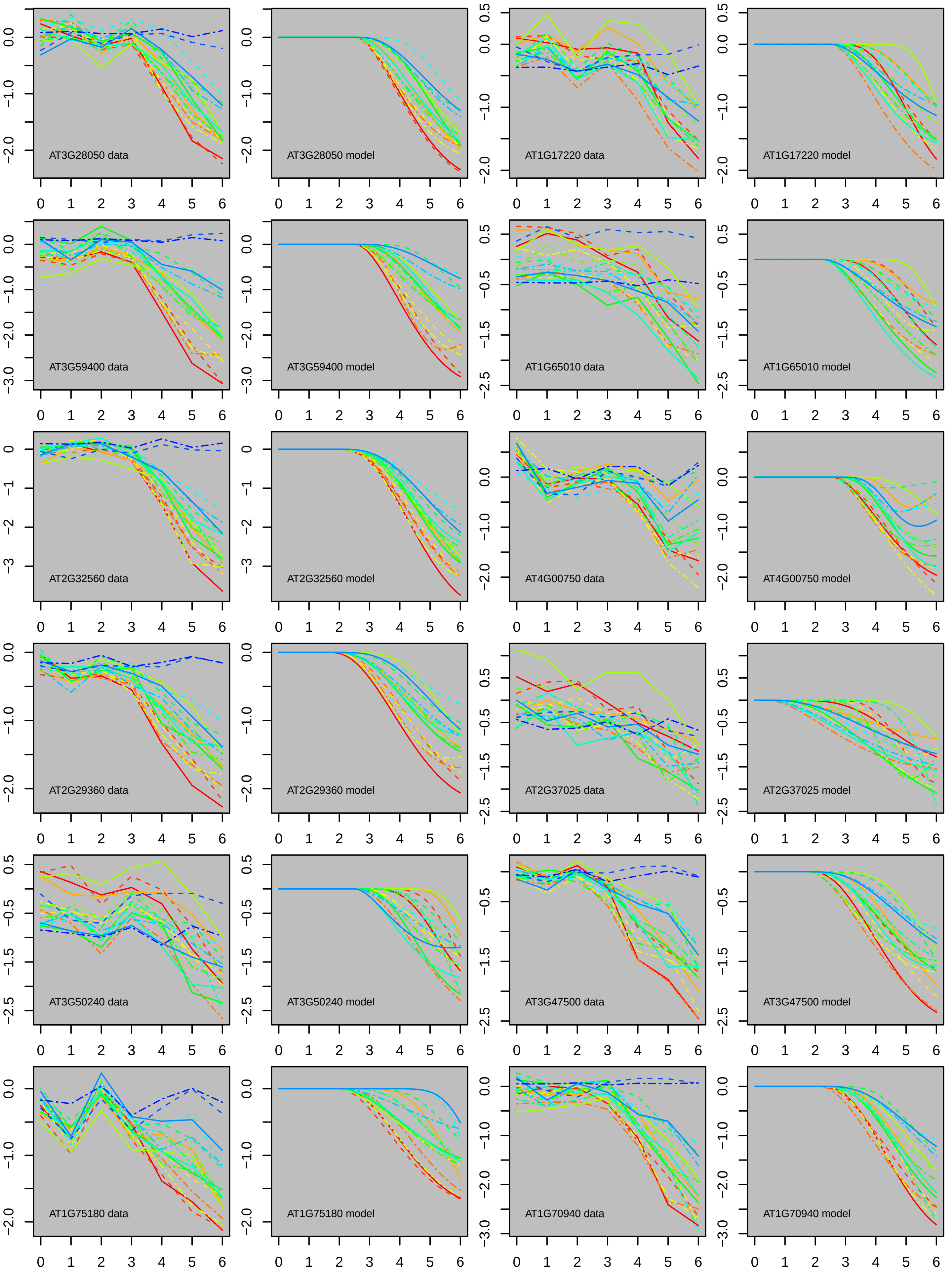

AT3G28050 data
AT3G28050 model
AT1G17220 data
AT1G17220 model
AT3G59400 data
AT3G59400 model
AT1G65010 data
AT1G65010 model
AT2G32560 data
AT2G32560 model
AT4G00750 data
AT4G00750 model
AT2G29360 data
AT2G29360 model
AT2G37025 data
AT2G37025 model
AT3G50240 data
AT3G50240 model
AT3G47500 data
AT3G47500 model
AT1G75180 data
AT1G75180 model
AT1G70940 data
AT1G70940 model

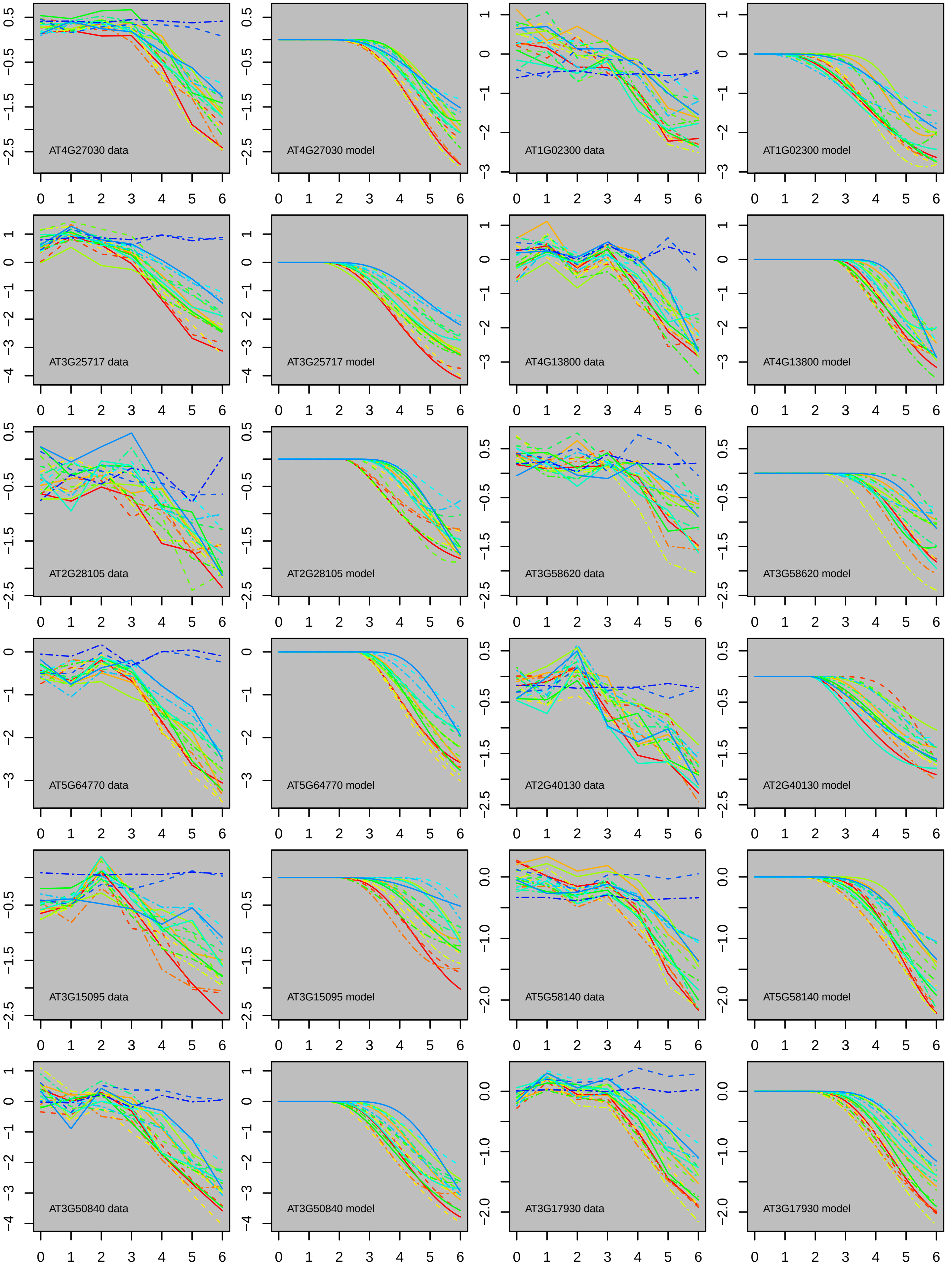
AT4G27030 data
AT4G27030 model
AT1G02300 data
AT1G02300 model
AT3G25717 data
AT3G25717 model
AT4G13800 data
AT4G13800 model
AT2G28105 data
AT2G28105 model
AT3G58620 data
AT3G58620 model
AT5G64770 data
AT5G64770 model
AT2G40130 data
AT2G40130 model
AT3G15095 data
AT3G15095 model
AT5G58140 data
AT5G58140 model
AT3G50840 data
AT3G50840 model
AT3G17930 data
AT3G17930 model

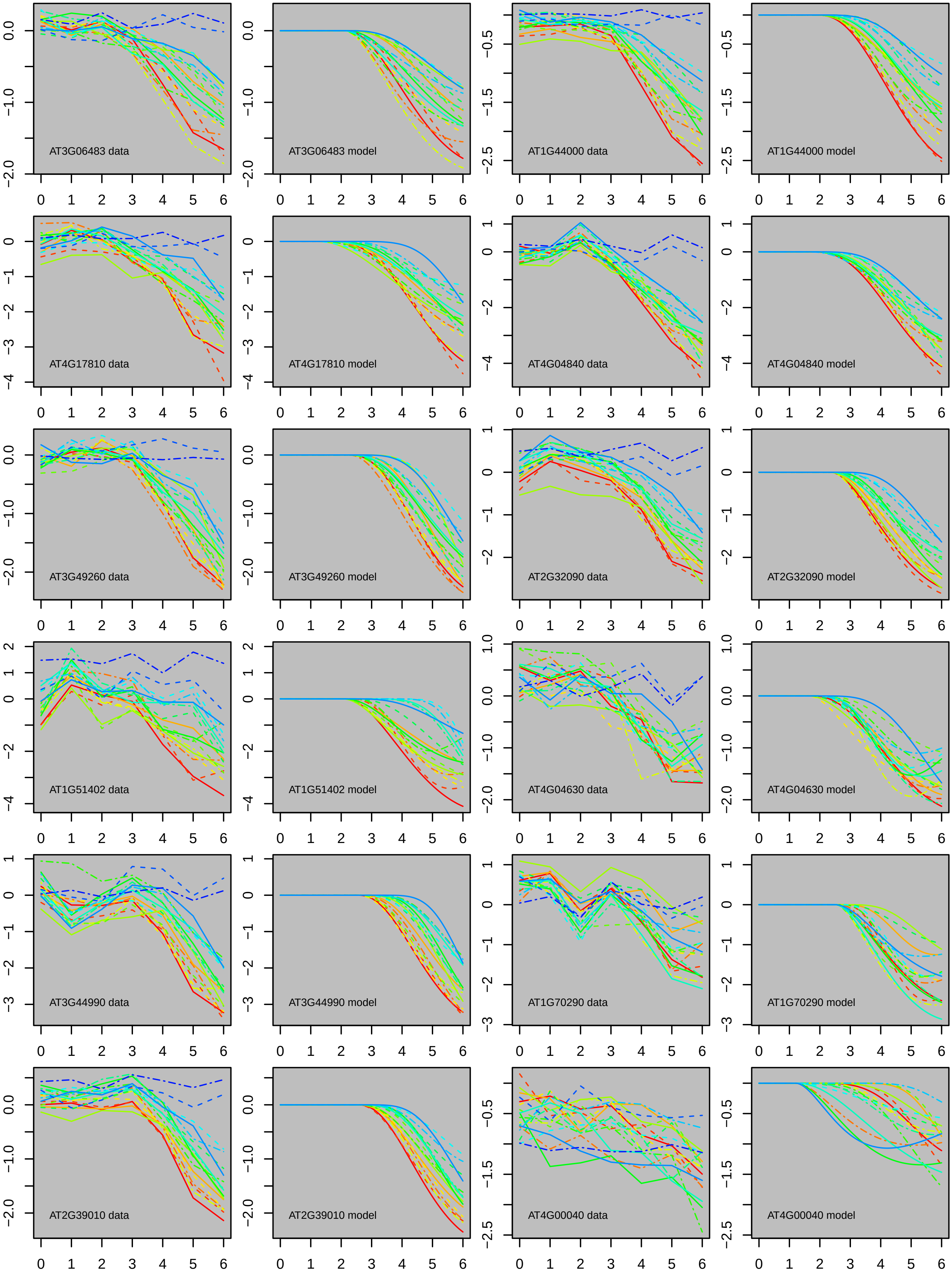
AT3G06483 data
AT3G06483 model
AT1G44000 data
AT1G44000 model
AT4G17810 data
AT4G17810 model
AT4G04840 data
AT4G04840 model
AT3G49260 data
AT3G49260 model
AT2G32090 data
AT2G32090 model
AT1G51402 data
AT1G51402 model
AT4G04630 data
AT4G04630 model
AT3G44990 data
AT3G44990 model
AT1G70290 data
AT1G70290 model
AT2G39010 data
AT2G39010 model
AT4G00040 data
AT4G00040 model

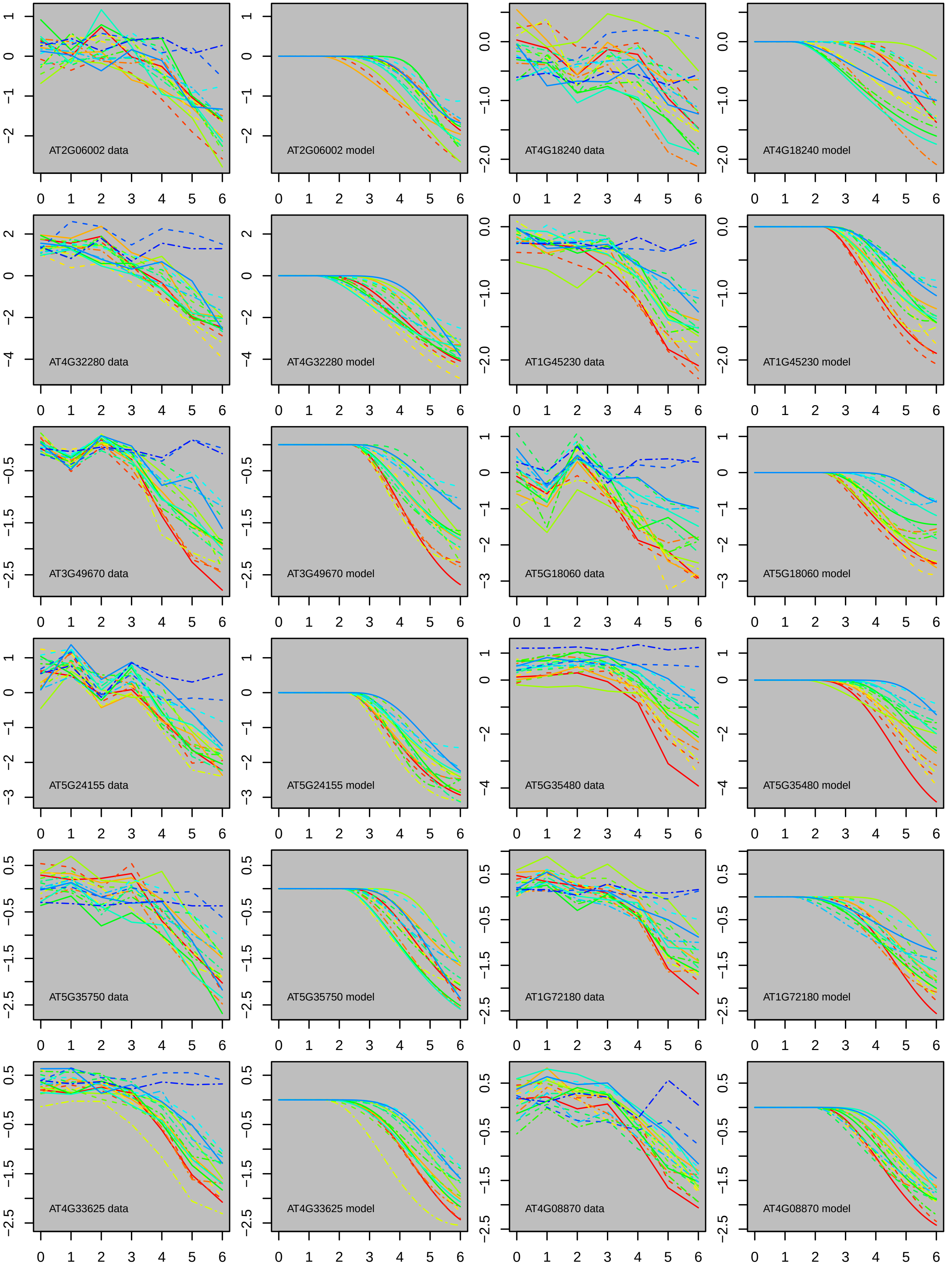
AT2G06002 data
AT2G06002 model
AT4G18240 data
AT4G18240 model
AT4G32280 data
AT4G32280 model
AT1G45230 data
AT1G45230 model
AT3G49670 data
AT3G49670 model
AT5G18060 data
AT5G18060 model
AT5G24155 data
AT5G24155 model
AT5G35480 data
AT5G35480 model
AT5G35750 data
AT5G35750 model
AT1G72180 data
AT1G72180 model
AT4G33625 data
AT4G33625 model
AT4G08870 data
AT4G08870 model

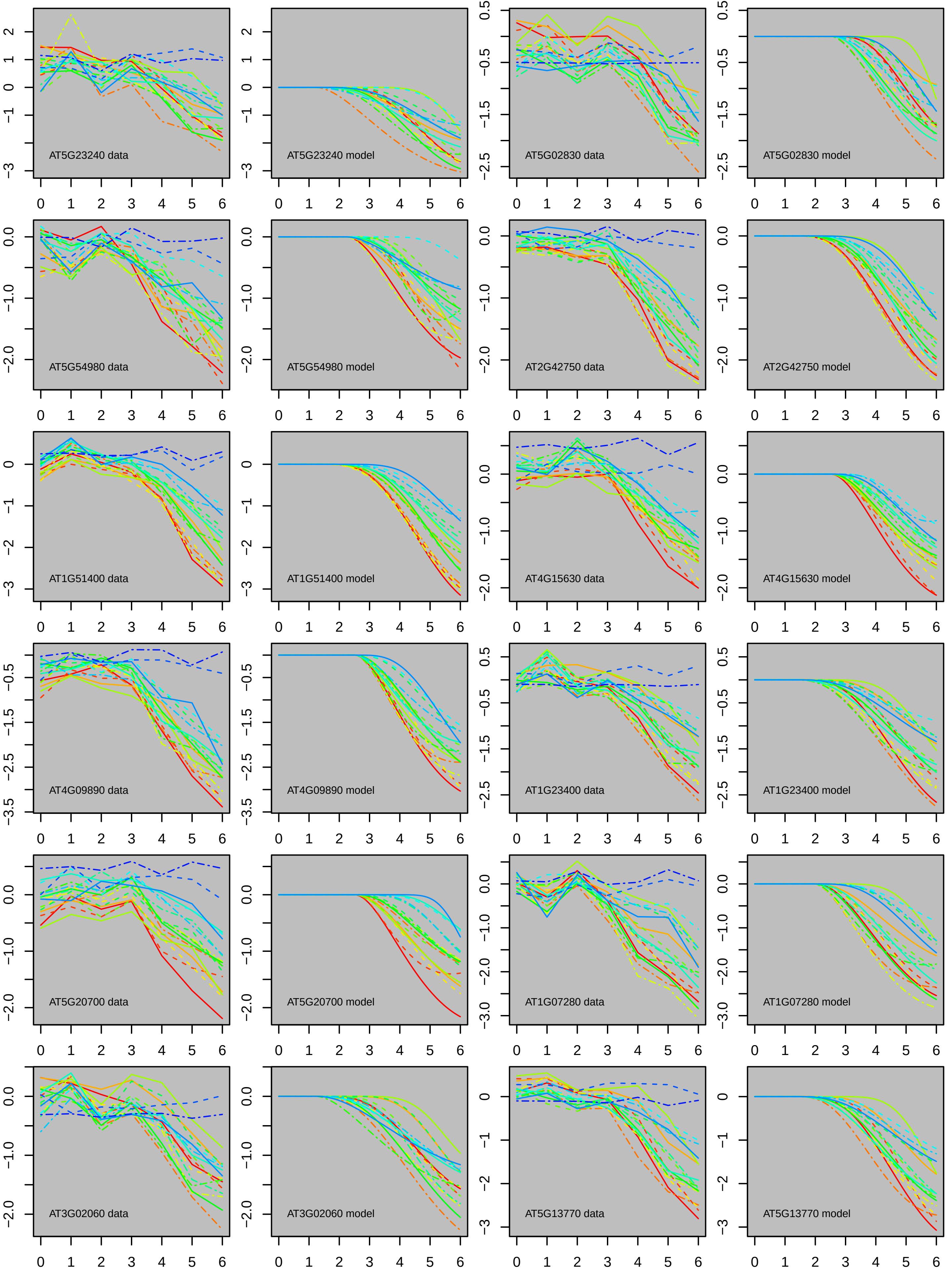
AT5G23240 data
AT5G23240 model
AT5G02830 data
AT5G02830 model
AT5G54980 data
AT5G54980 model
AT2G42750 data
AT2G42750 model
AT1G51400 data
AT1G51400 model
AT4G15630 data
AT4G15630 model
AT4G09890 data
AT4G09890 model
AT1G23400 data
AT1G23400 model
AT5G20700 data
AT5G20700 model
AT1G07280 data
AT1G07280 model
AT3G02060 data
AT3G02060 model
AT5G13770 data
AT5G13770 model

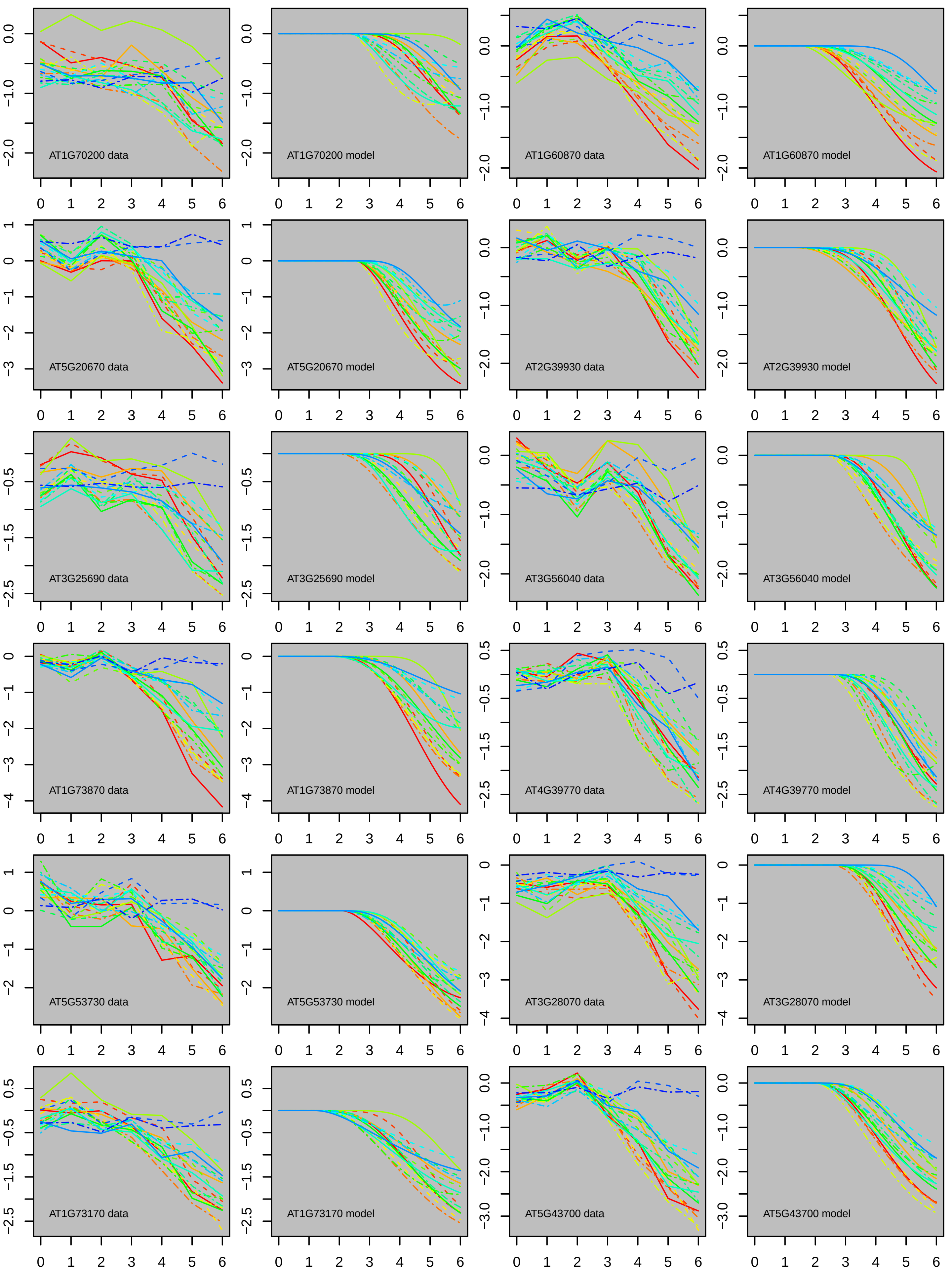
AT1G70200 data
AT1G70200 model
AT1G60870 data
AT1G60870 model
AT5G20670 data
AT5G20670 model
AT2G39930 data
AT2G39930 model
AT3G25690 data
AT3G25690 model
AT3G56040 data
AT3G56040 model
AT1G73870 data
AT1G73870 model
AT4G39770 data
AT4G39770 model
AT5G53730 data
AT5G53730 model
AT3G28070 data
AT3G28070 model
AT1G73170 data
AT1G73170 model
AT5G43700 data
AT5G43700 model

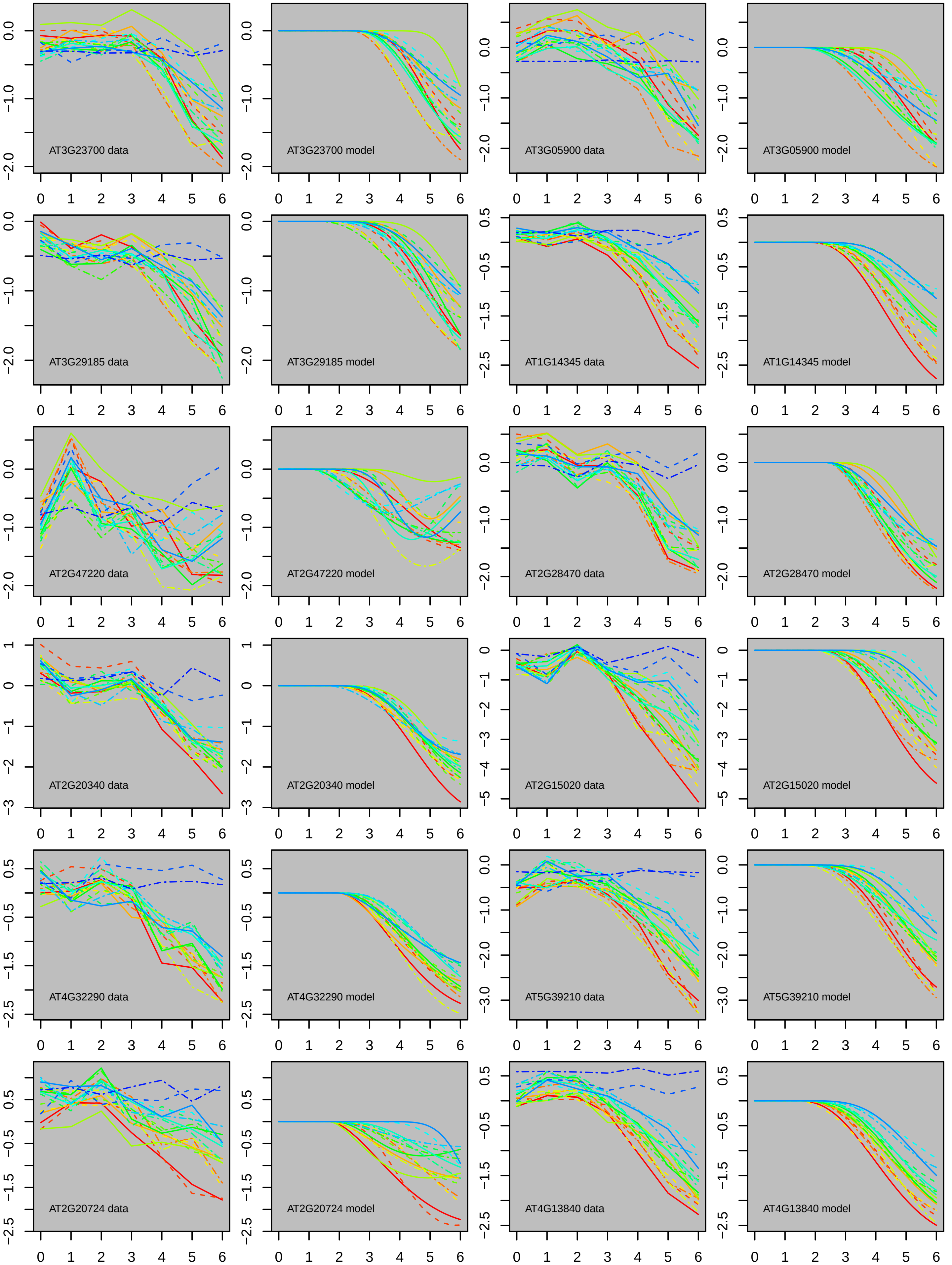

AT3G23700 data
AT3G23700 model
AT3G05900 data
AT3G05900 model
AT3G29185 data
AT3G29185 model
AT1G14345 data
AT1G14345 model
AT2G47220 data
AT2G47220 model
AT2G28470 data
AT2G28470 model
AT2G20340 data
AT2G20340 model
AT2G15020 data
AT2G15020 model
AT4G32290 data
AT4G32290 model
AT5G39210 data
AT5G39210 model
AT2G20724 data
AT2G20724 model
AT4G13840 data
AT4G13840 model

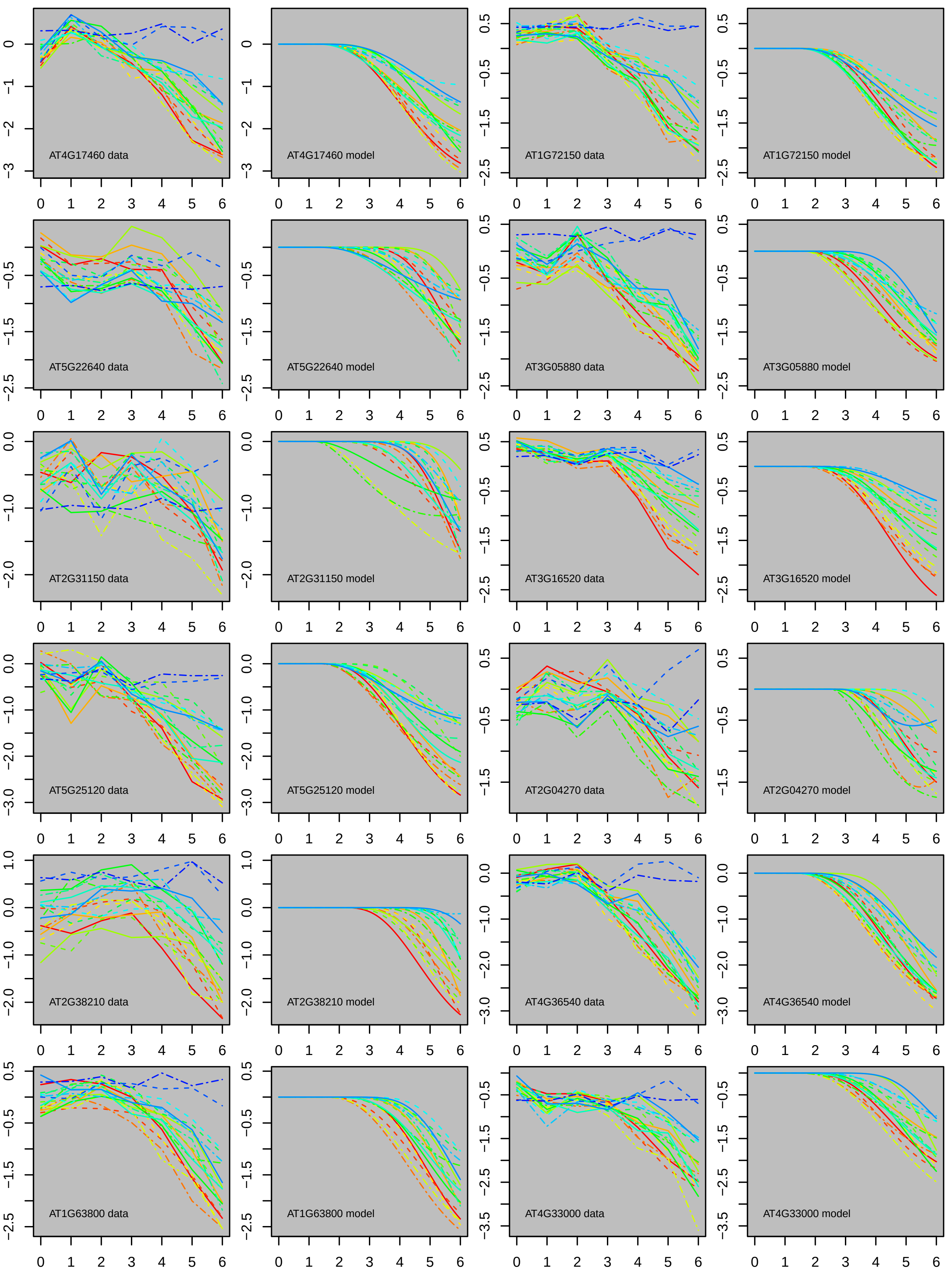

AT4G17460 data
AT4G17460 model
AT1G72150 data
AT1G72150 model
AT5G22640 data
AT5G22640 model
AT3G05880 data
AT3G05880 model
AT2G31150 data
AT2G31150 model
AT3G16520 data
AT3G16520 model
AT5G25120 data
AT5G25120 model
AT2G04270 data
AT2G04270 model
AT2G38210 data
AT2G38210 model
AT4G36540 data
AT4G36540 model
AT1G63800 data
AT1G63800 model
AT4G33000 data
AT4G33000 model

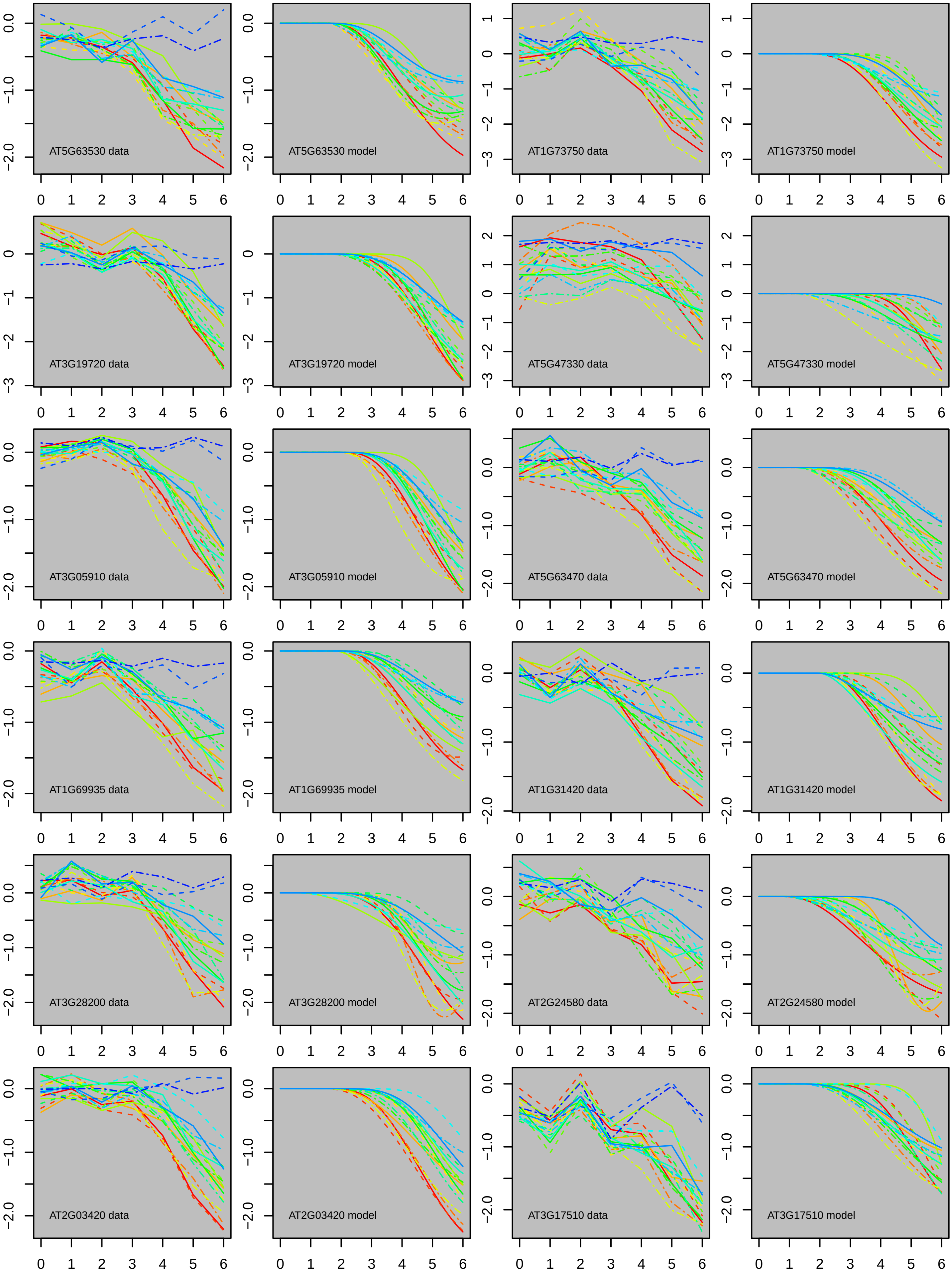

AT5G63530 data
AT5G63530 model
AT1G73750 data
AT1G73750 model
AT3G19720 data
AT3G19720 model
AT5G47330 data
AT5G47330 model
AT3G05910 data
AT3G05910 model
AT5G63470 data
AT5G63470 model
AT1G69935 data
AT1G69935 model
AT1G31420 data
AT1G31420 model
AT3G28200 data
AT3G28200 model
AT2G24580 data
AT2G24580 model
AT2G03420 data
AT2G03420 model
AT3G17510 data
AT3G17510 model

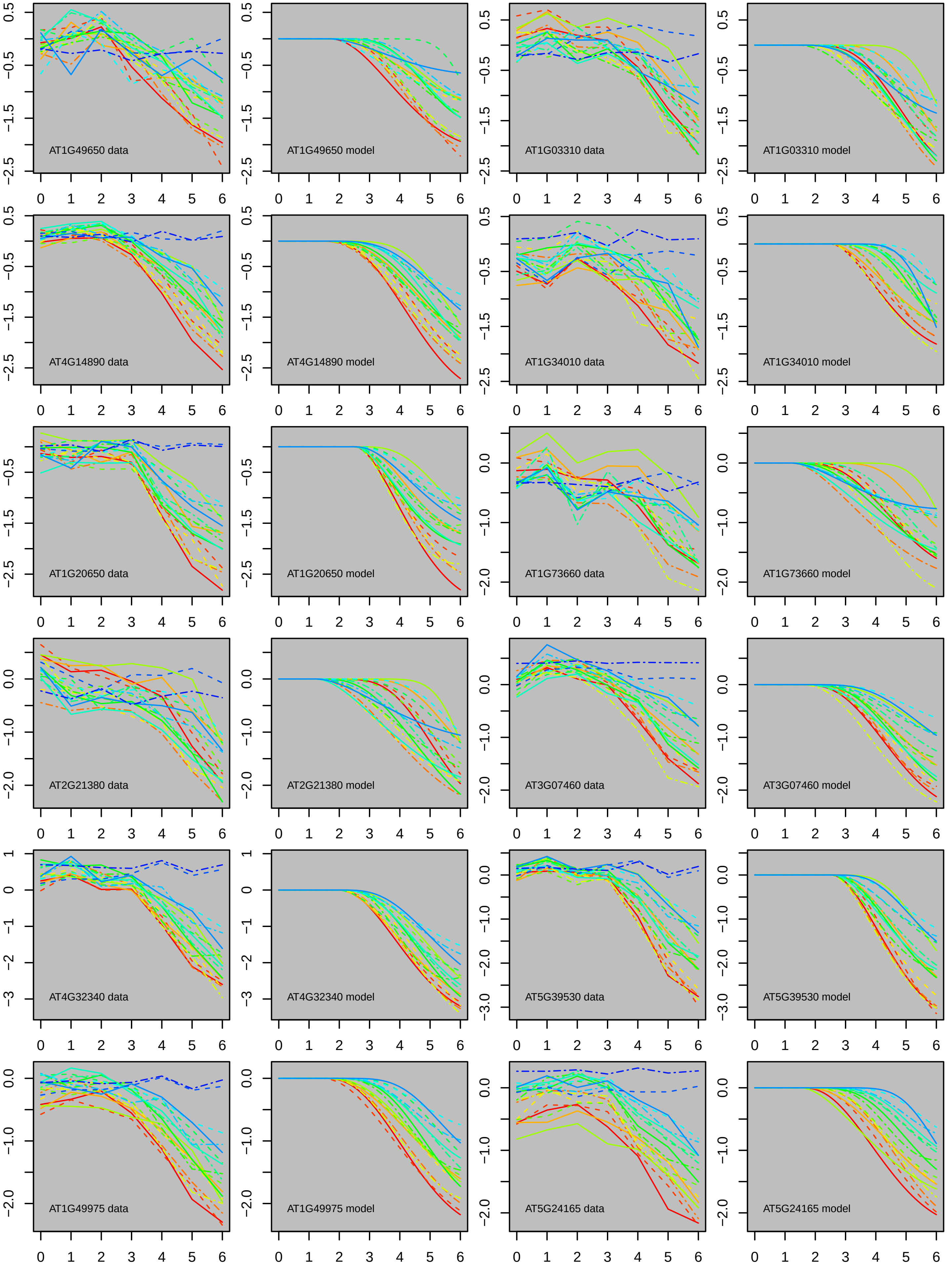
AT1G49650 data
AT1G49650 model
AT1G03310 data
AT1G03310 model
AT4G14890 data
AT4G14890 model
AT1G34010 data
AT1G34010 model
AT1G20650 data
AT1G20650 model
AT1G73660 data
AT1G73660 model
AT2G21380 data
AT2G21380 model
AT3G07460 data
AT3G07460 model
AT4G32340 data
AT4G32340 model
AT5G39530 data
AT5G39530 model
AT1G49975 data
AT1G49975 model
AT5G24165 data
AT5G24165 model

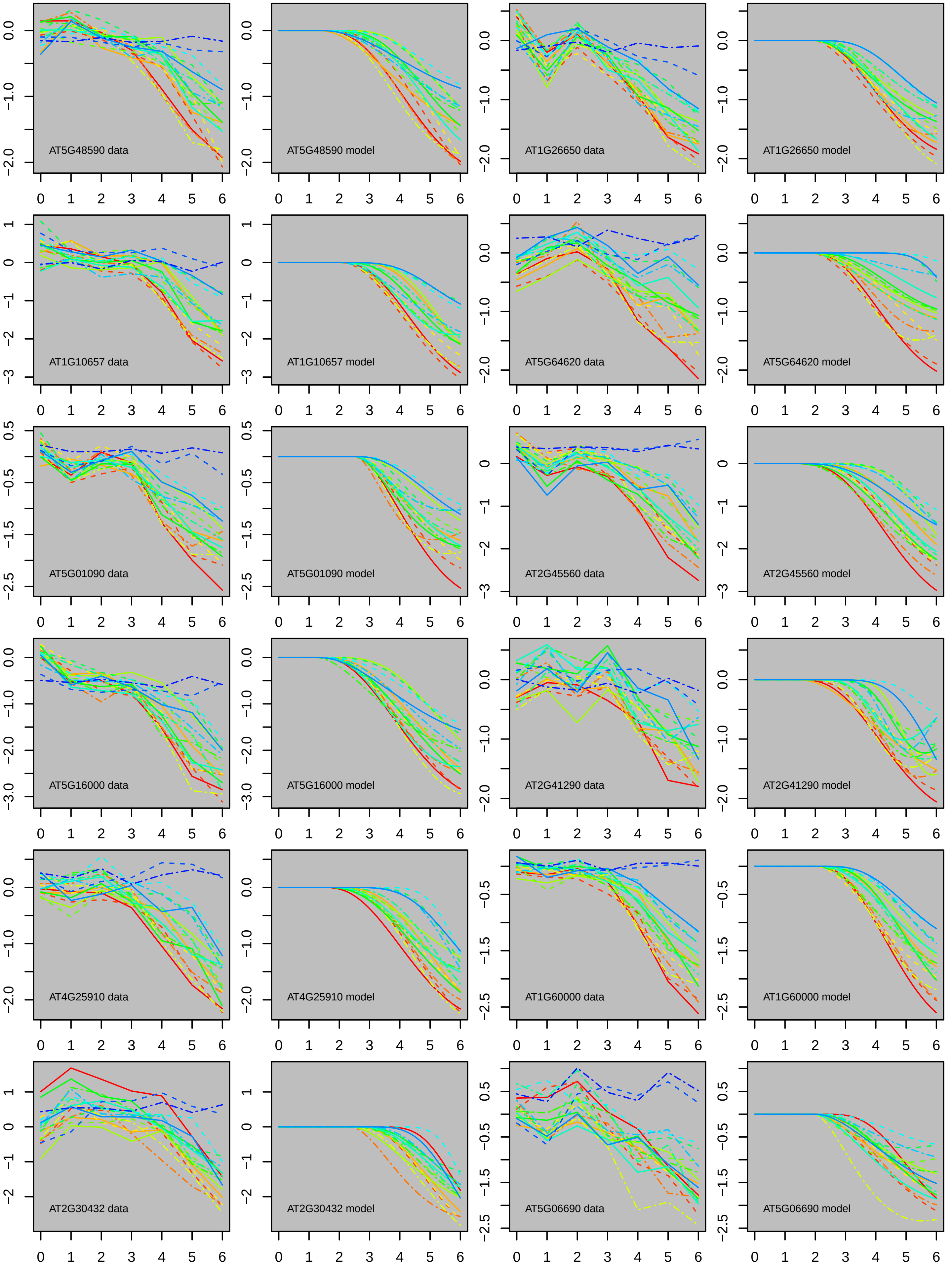

AT5G48590 data
AT5G48590 model
AT1G26650 data
AT1G26650 model
AT1G10657 data
AT1G10657 model
AT5G64620 data
AT5G64620 model
AT5G01090 data
AT5G01090 model
AT2G45560 data
AT2G45560 model
AT5G16000 data
AT5G16000 model
AT2G41290 data
AT2G41290 model
AT4G25910 data
AT4G25910 model
AT1G60000 data
AT1G60000 model
AT2G30432 data
AT2G30432 model
AT5G06690 data
AT5G06690 model

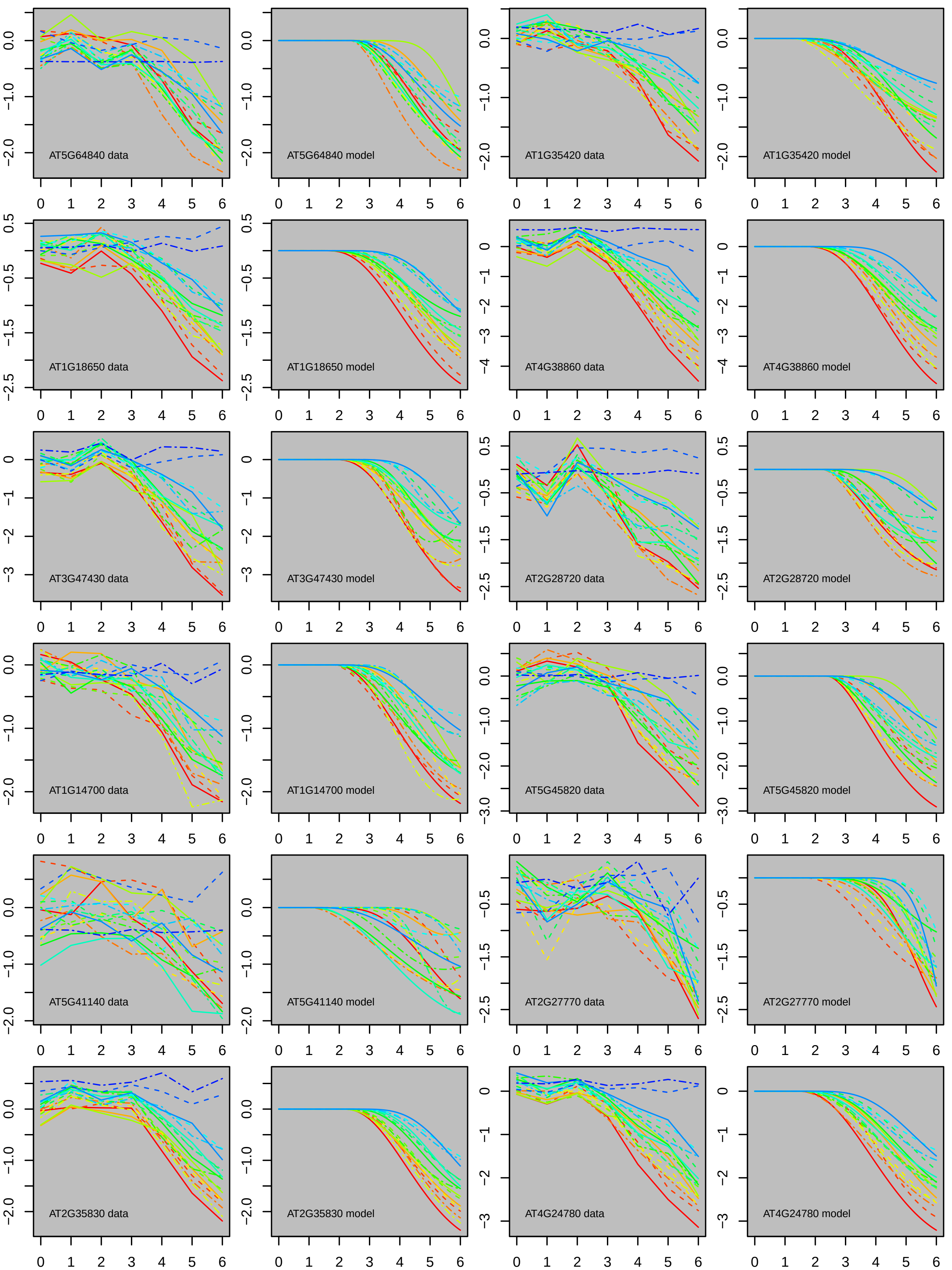
AT5G64840 data
AT5G64840 model
AT1G35420 data
AT1G35420 model
AT1G18650 data
AT1G18650 model
AT4G38860 data
AT4G38860 model
AT3G47430 data
AT3G47430 model
AT2G28720 data
AT2G28720 model
AT1G14700 data
AT1G14700 model
AT5G45820 data
AT5G45820 model
AT5G41140 data
AT5G41140 model
AT2G27770 data
AT2G27770 model
AT2G35830 data
AT2G35830 model
AT4G24780 data
AT4G24780 model

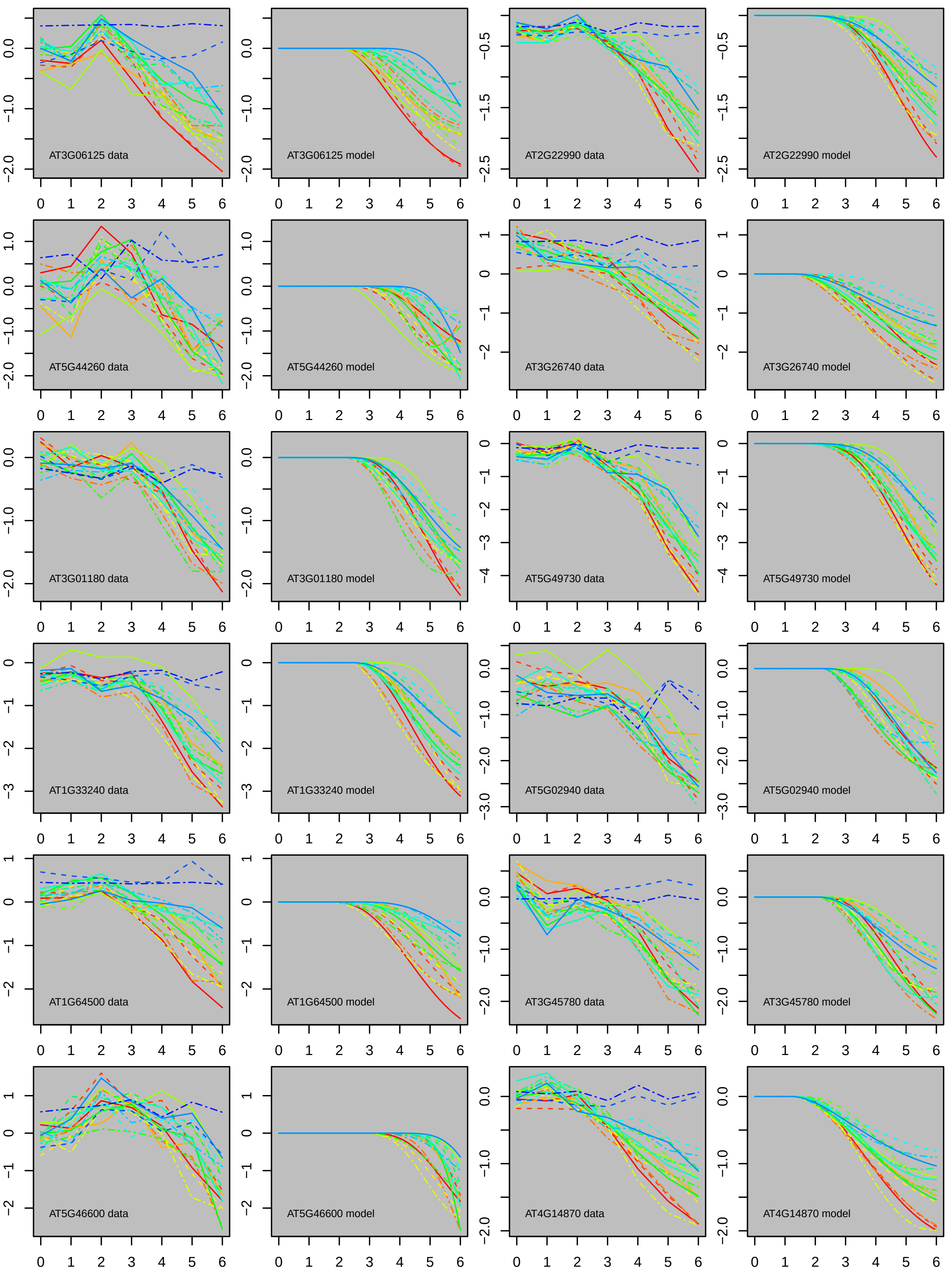
AT3G06125 data
AT3G06125 model
AT2G22990 data
AT2G22990 model
AT5G44260 data
AT5G44260 model
AT3G26740 data
AT3G26740 model
AT3G01180 data
AT3G01180 model
AT5G49730 data
AT5G49730 model
AT1G33240 data
AT1G33240 model
AT5G02940 data
AT5G02940 model
AT1G64500 data
AT1G64500 model
AT3G45780 data
AT3G45780 model
AT5G46600 data
AT5G46600 model
AT4G14870 data
AT4G14870 model

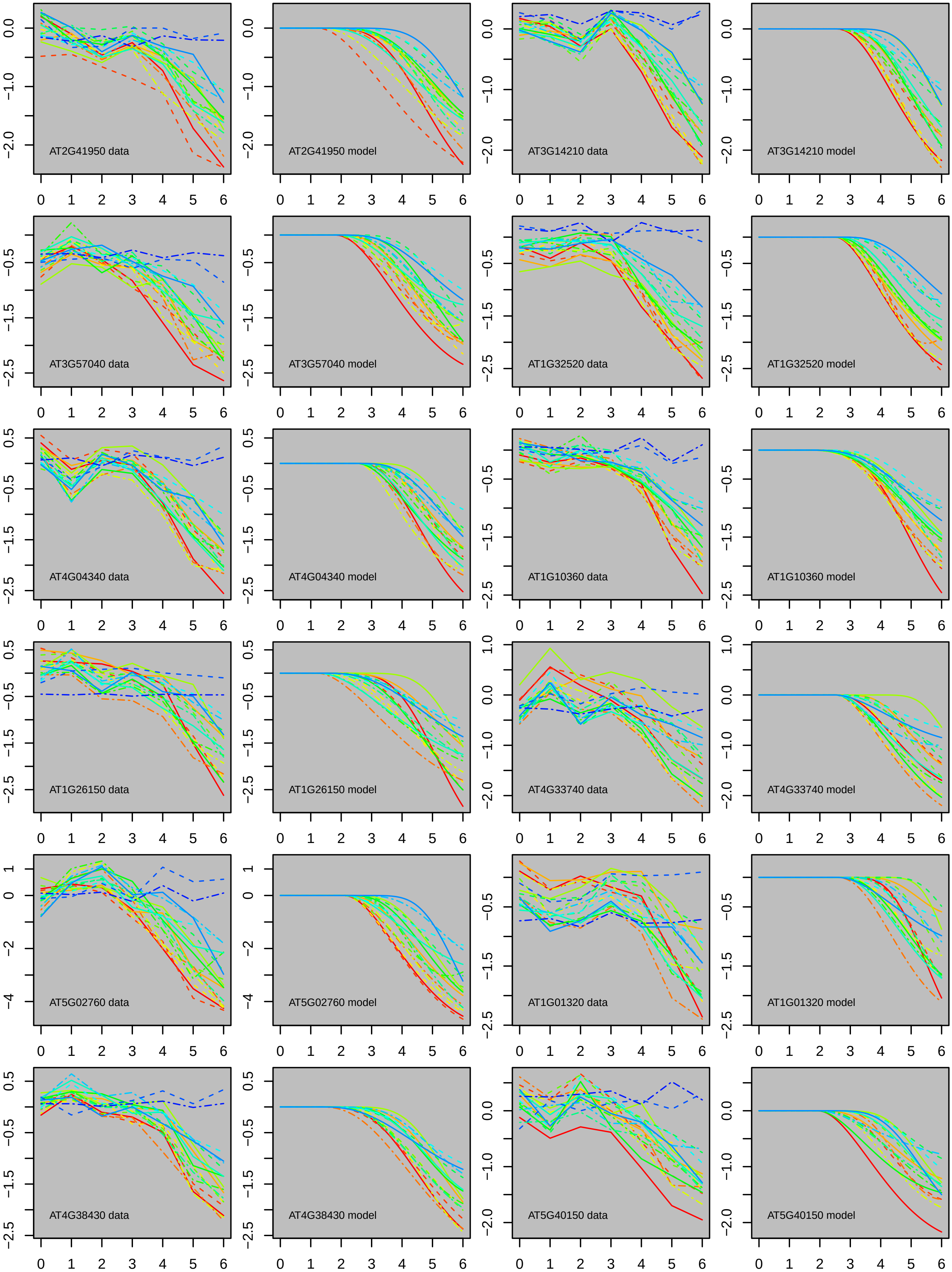
AT2G41950 data
AT2G41950 model
AT3G14210 data
AT3G14210 model
AT3G57040 data
AT3G57040 model
AT1G32520 data
AT1G32520 model
AT4G04340 data
AT4G04340 model
AT1G10360 data
AT1G10360 model
AT1G26150 data
AT1G26150 model
AT4G33740 data
AT4G33740 model
AT5G02760 data
AT5G02760 model
AT1G01320 data
AT1G01320 model
AT4G38430 data
AT4G38430 model
AT5G40150 data
AT5G40150 model

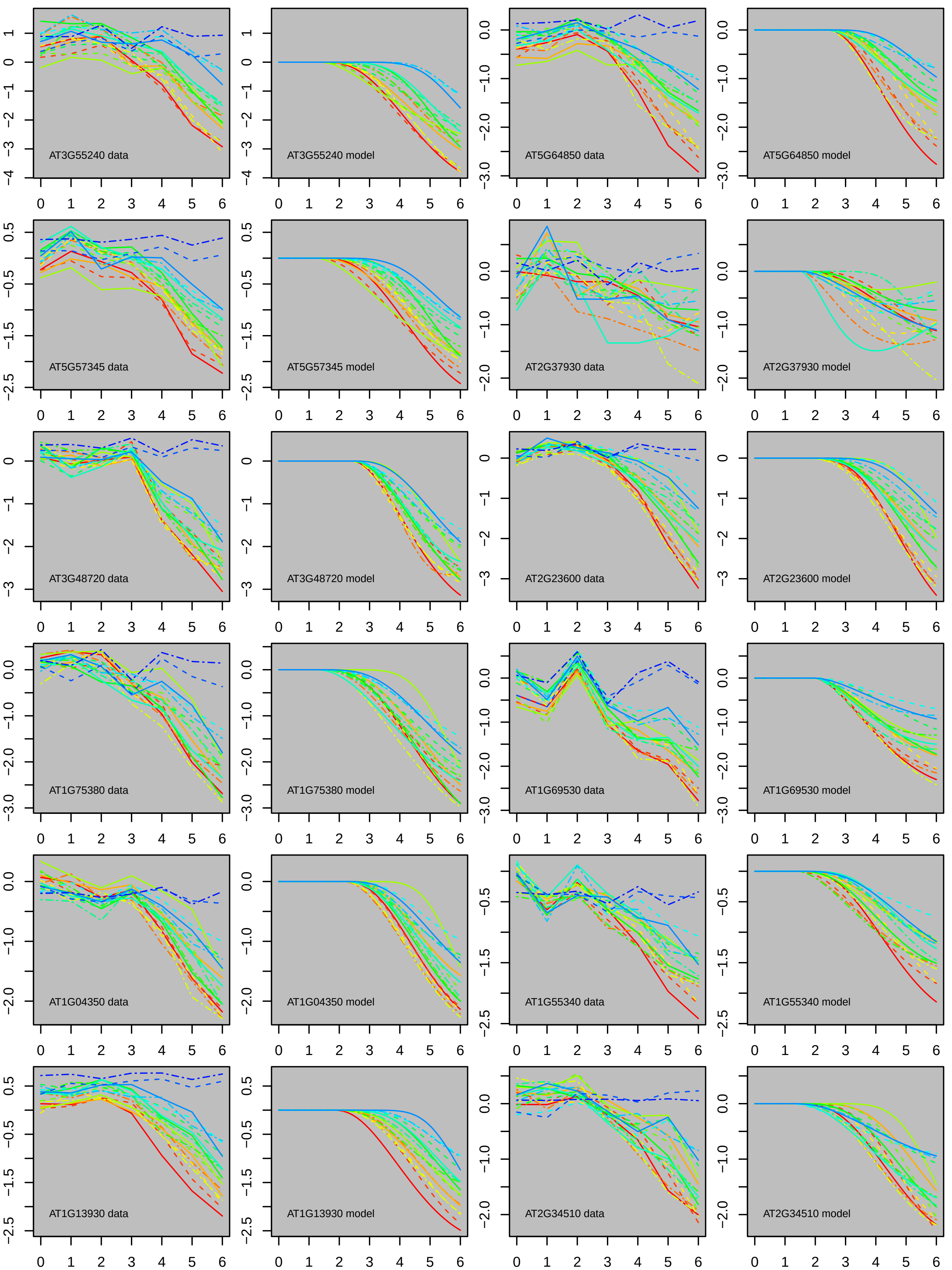
AT3G55240 data
AT3G55240 model
AT5G64850 data
AT5G64850 model
AT5G57345 data
AT5G57345 model
AT2G37930 data
AT2G37930 model
AT3G48720 data
AT3G48720 model
AT2G23600 data
AT2G23600 model
AT1G75380 data
AT1G75380 model
AT1G69530 data
AT1G69530 model
AT1G04350 data
AT1G04350 model
AT1G55340 data
AT1G55340 model
AT1G13930 data
AT1G13930 model
AT2G34510 data
AT2G34510 model

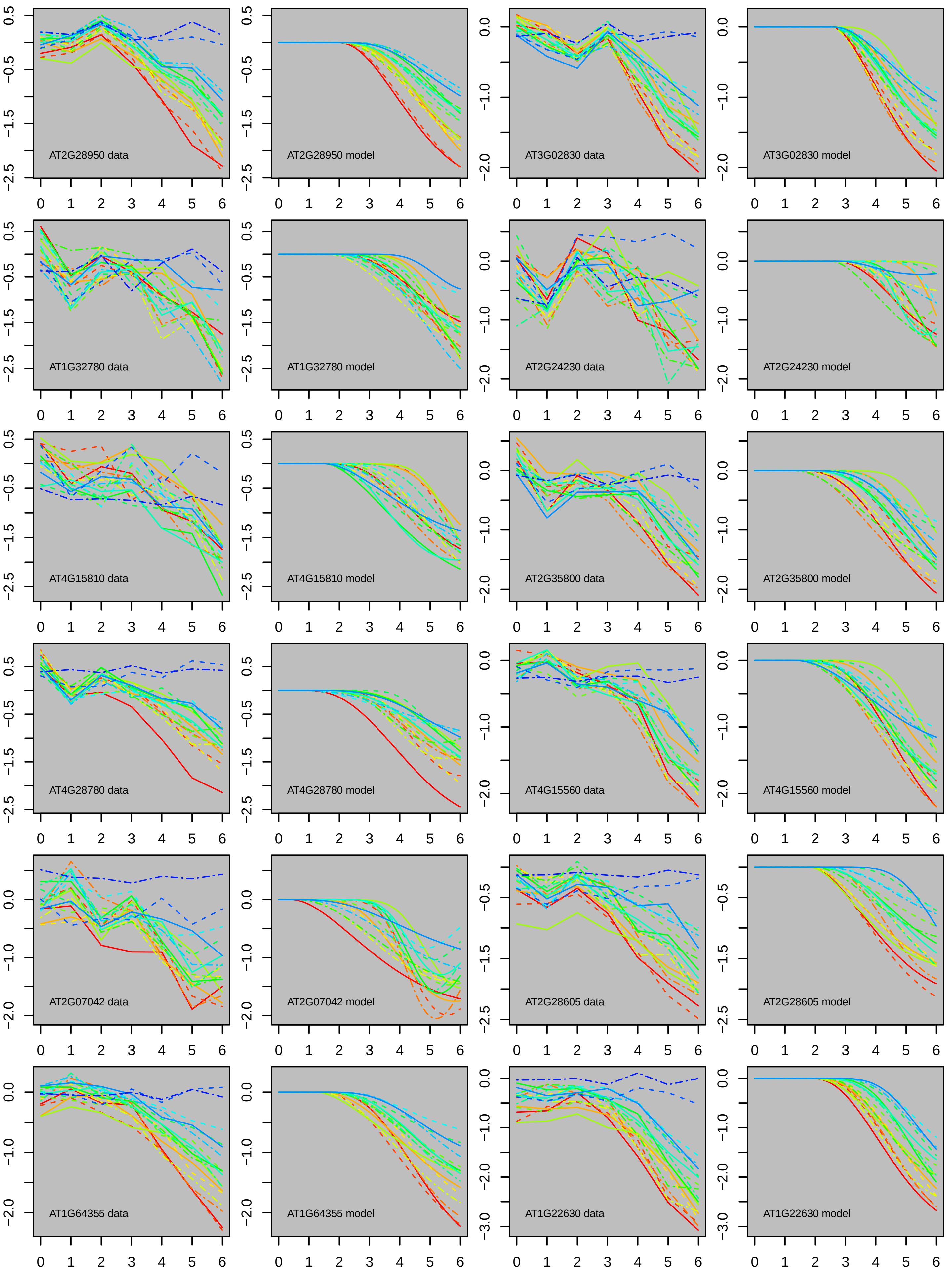
AT2G28950 data
AT2G28950 model
AT3G02830 data
AT3G02830 model
AT1G32780 data
AT1G32780 model
AT2G24230 data
AT2G24230 model
AT4G15810 data
AT4G15810 model
AT2G35800 data
AT2G35800 model
AT4G28780 data
AT4G28780 model
AT4G15560 data
AT4G15560 model
AT2G07042 data
AT2G07042 model
AT2G28605 data
AT2G28605 model
AT1G64355 data
AT1G64355 model
AT1G22630 data
AT1G22630 model

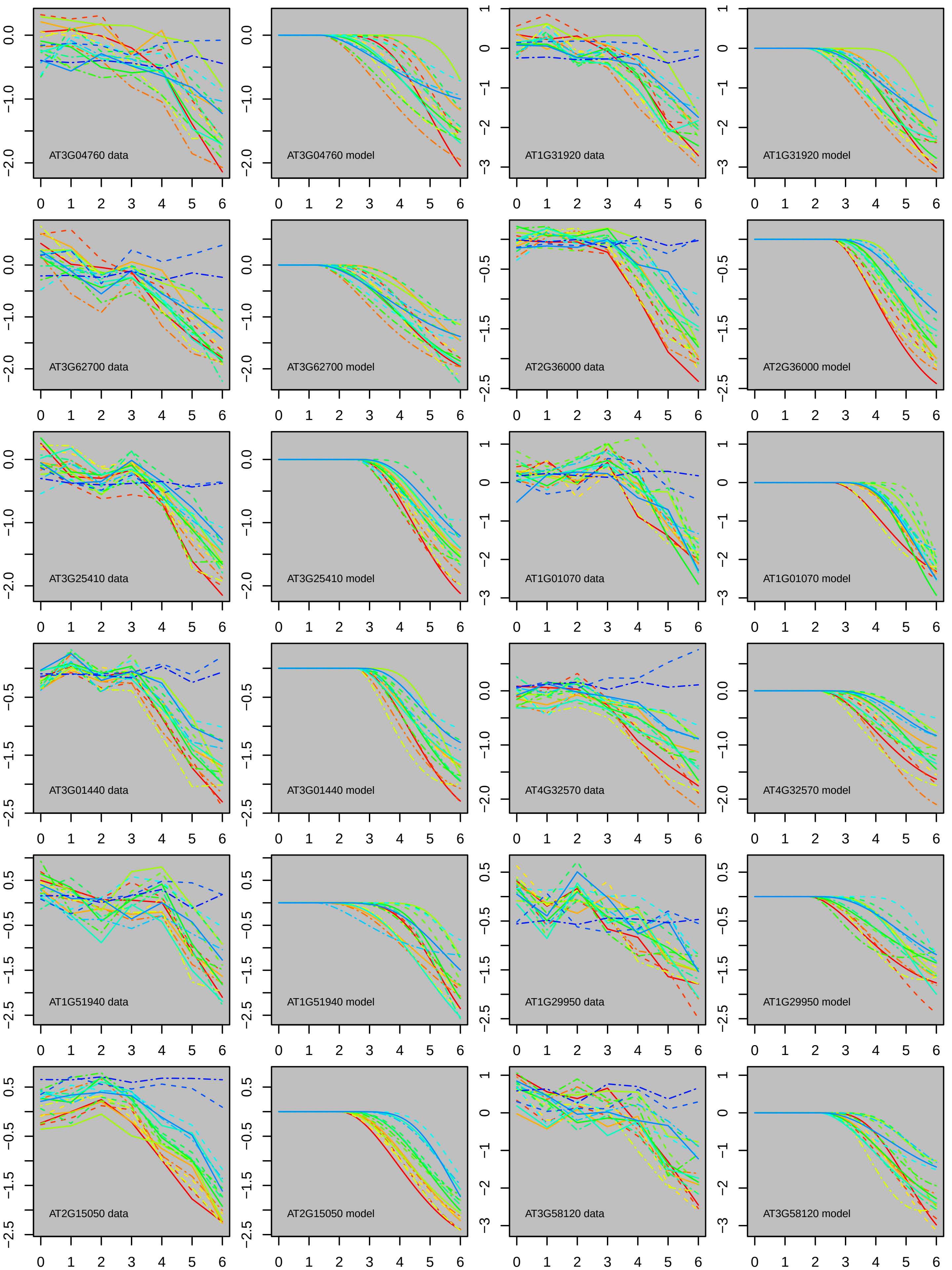

AT3G04760 data
AT3G04760 model
AT1G31920 data
AT1G31920 model
AT3G62700 data
AT3G62700 model
AT2G36000 data
AT2G36000 model
AT3G25410 data
AT3G25410 model
AT1G01070 data
AT1G01070 model
AT3G01440 data
AT3G01440 model
AT4G32570 data
AT4G32570 model
AT1G51940 data
AT1G51940 model
AT1G29950 data
AT1G29950 model
AT2G15050 data
AT2G15050 model
AT3G58120 data
AT3G58120 model

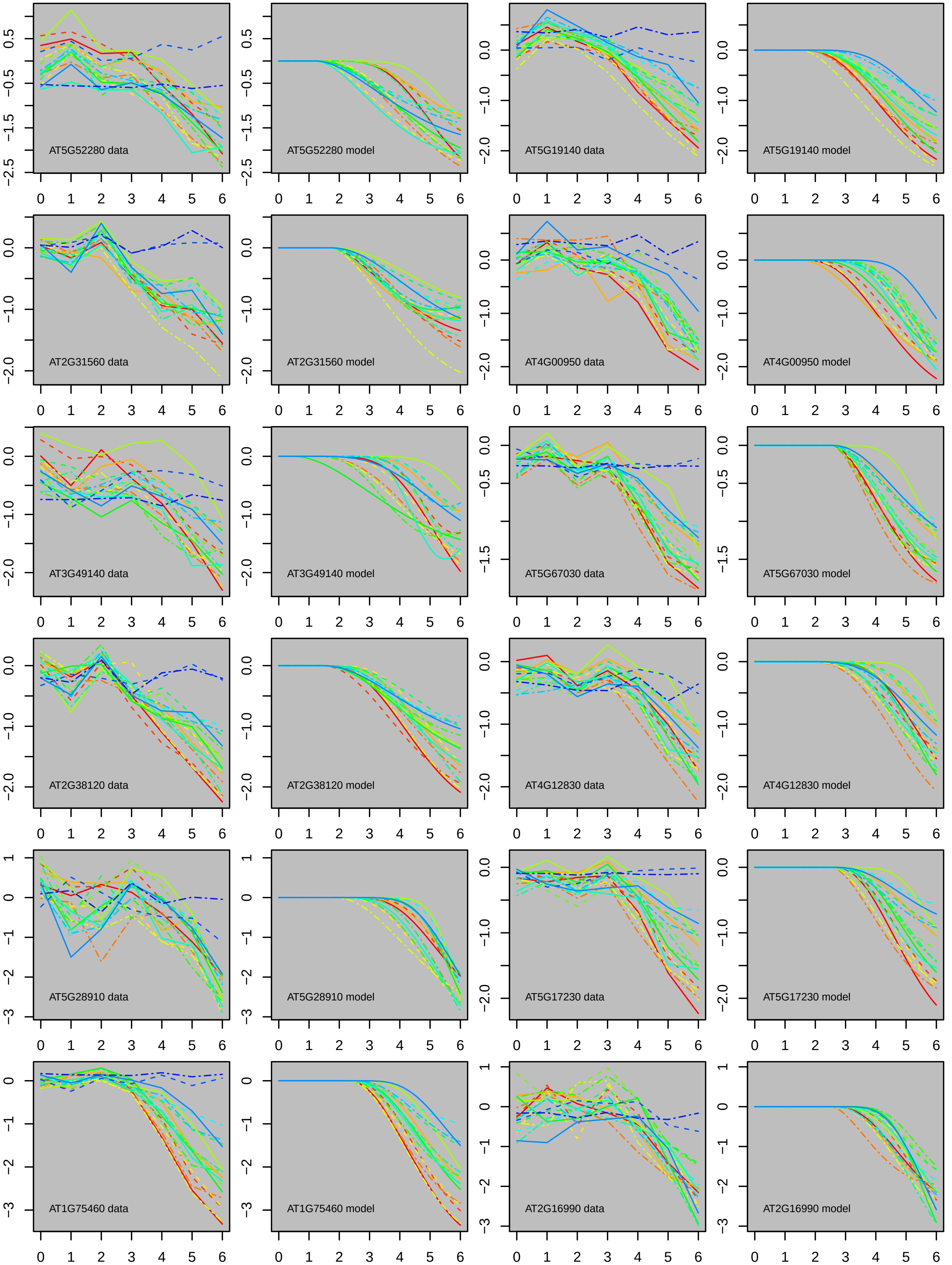
AT5G52280 data
AT5G52280 model
AT5G19140 data
AT5G19140 model
AT2G31560 data
AT2G31560 model
AT4G00950 data
AT4G00950 model
AT3G49140 data
AT3G49140 model
AT5G67030 data
AT5G67030 model
AT2G38120 data
AT2G38120 model
AT4G12830 data
AT4G12830 model
AT5G28910 data
AT5G28910 model
AT5G17230 data
AT5G17230 model
AT1G75460 data
AT1G75460 model
AT2G16990 data
AT2G16990 model

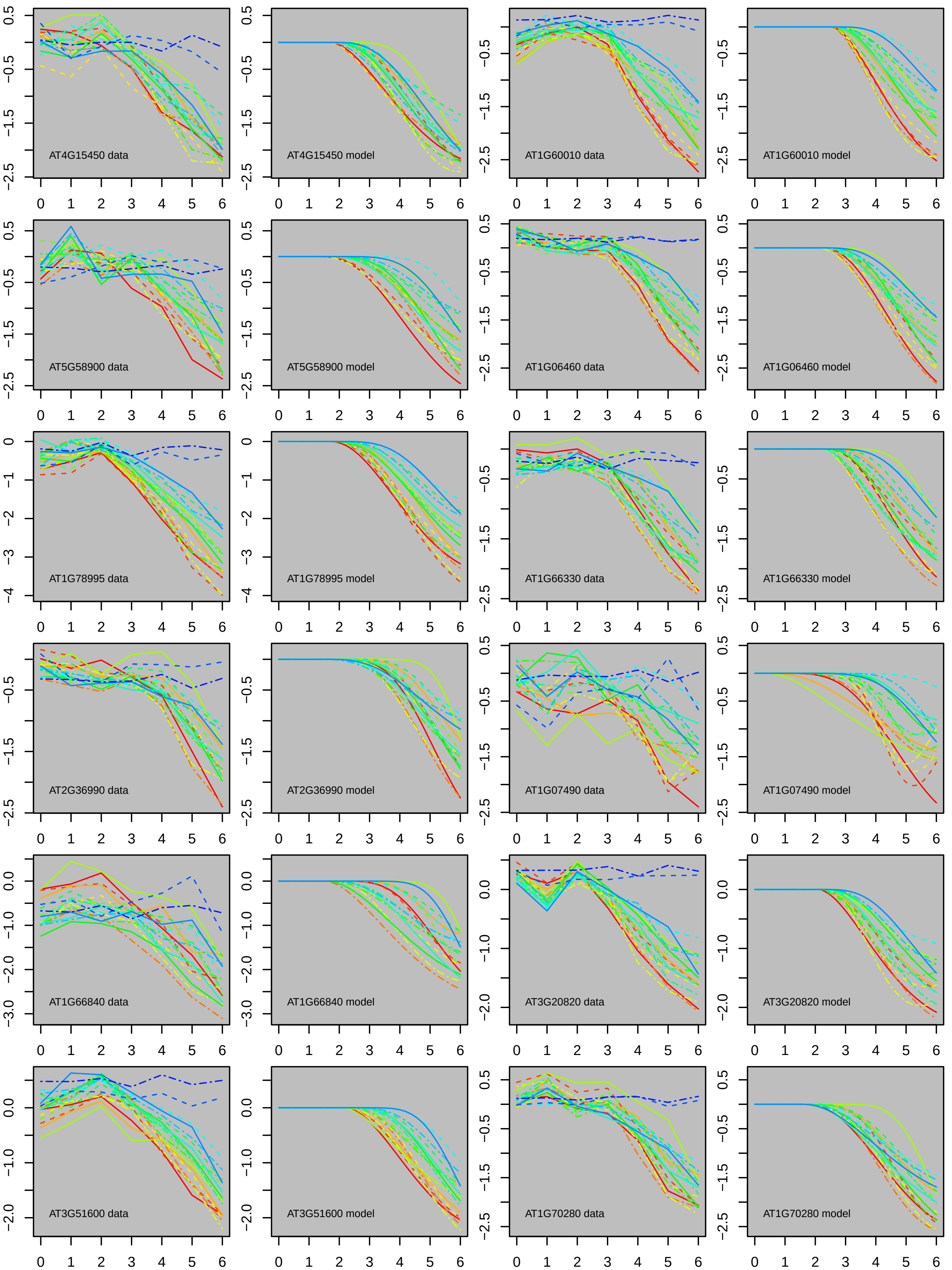
AT4G15450 data
AT4G15450 model
AT1G60010 data
AT1G60010 model
AT5G58900 data
AT5G58900 model
AT1G06460 data
AT1G06460 model
AT1G78995 data
AT1G78995 model
AT1G66330 data
AT1G66330 model
AT2G36990 data
AT2G36990 model
AT1G07490 data
AT1G07490 model
AT1G66840 data
AT1G66840 model
AT3G20820 data
AT3G20820 model
AT3G51600 data
AT3G51600 model
AT1G70280 data
AT1G70280 model

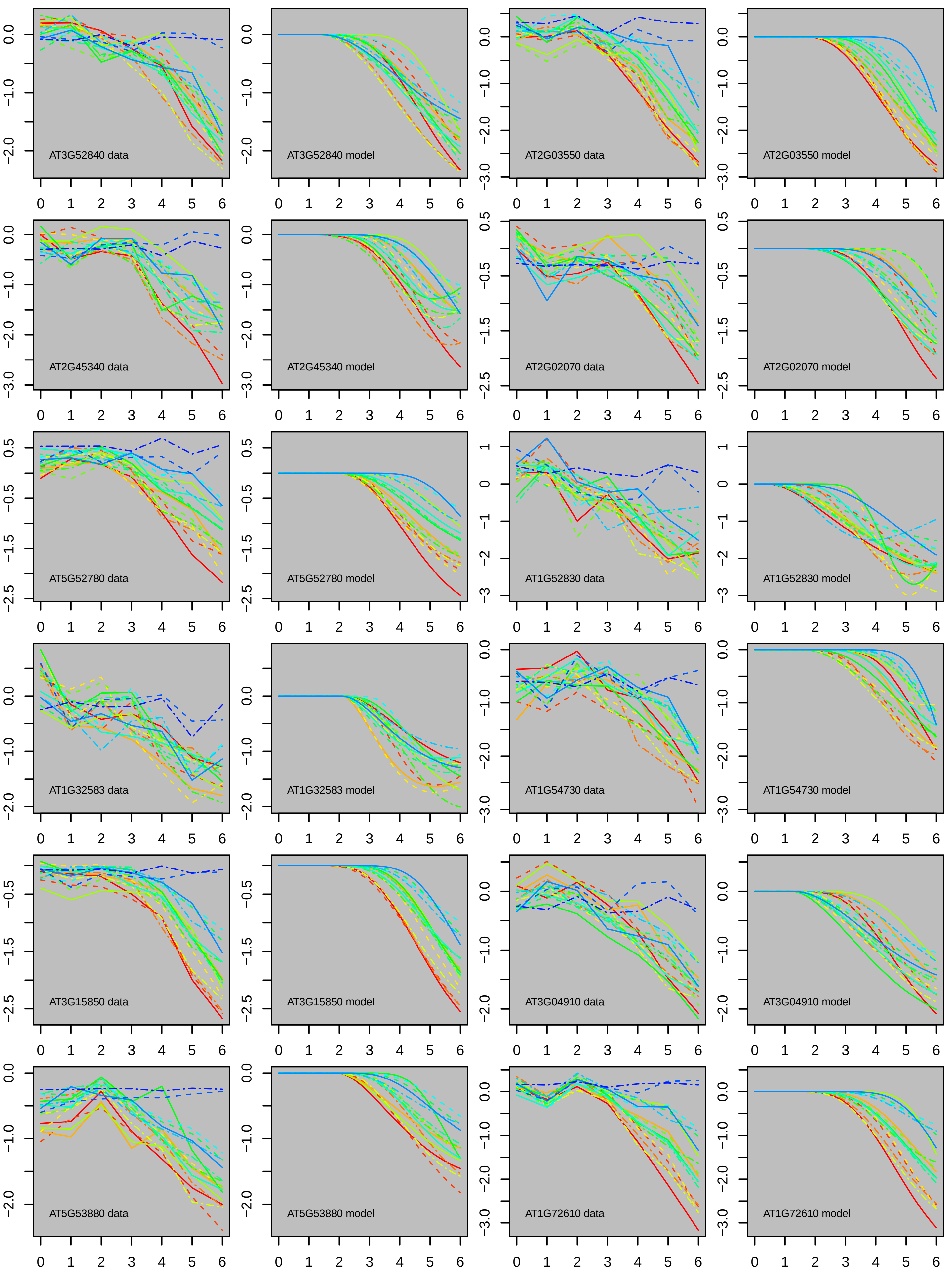
AT3G52840 data
AT3G52840 model
AT2G03550 data
AT2G03550 model
AT2G45340 data
AT2G45340 model
AT2G02070 data
AT2G02070 model
AT5G52780 data
AT5G52780 model
AT1G52830 data
AT1G52830 model
AT1G32583 data
AT1G32583 model
AT1G54730 data
AT1G54730 model
AT3G15850 data
AT3G15850 model
AT3G04910 data
AT3G04910 model
AT5G53880 data
AT5G53880 model
AT1G72610 data
AT1G72610 model

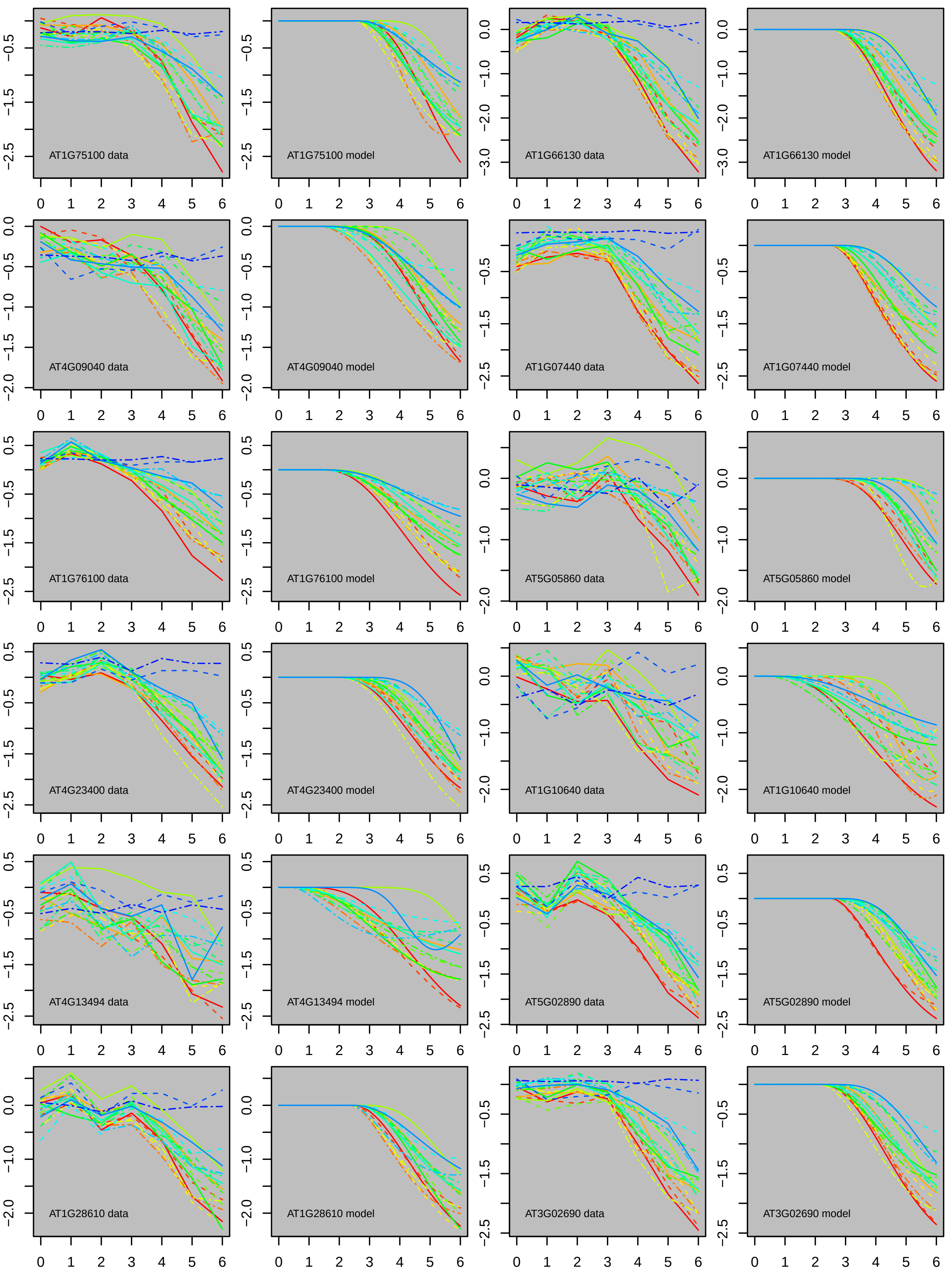
AT1G75100 data
AT1G75100 model
AT1G66130 data
AT1G66130 model
AT4G09040 data
AT4G09040 model
AT1G07440 data
AT1G07440 model
AT1G76100 data
AT1G76100 model
AT5G05860 data
AT5G05860 model
AT4G23400 data
AT4G23400 model
AT1G10640 data
AT1G10640 model
AT4G13494 data
AT4G13494 model
AT5G02890 data
AT5G02890 model
AT1G28610 data
AT1G28610 model
AT3G02690 data
AT3G02690 model

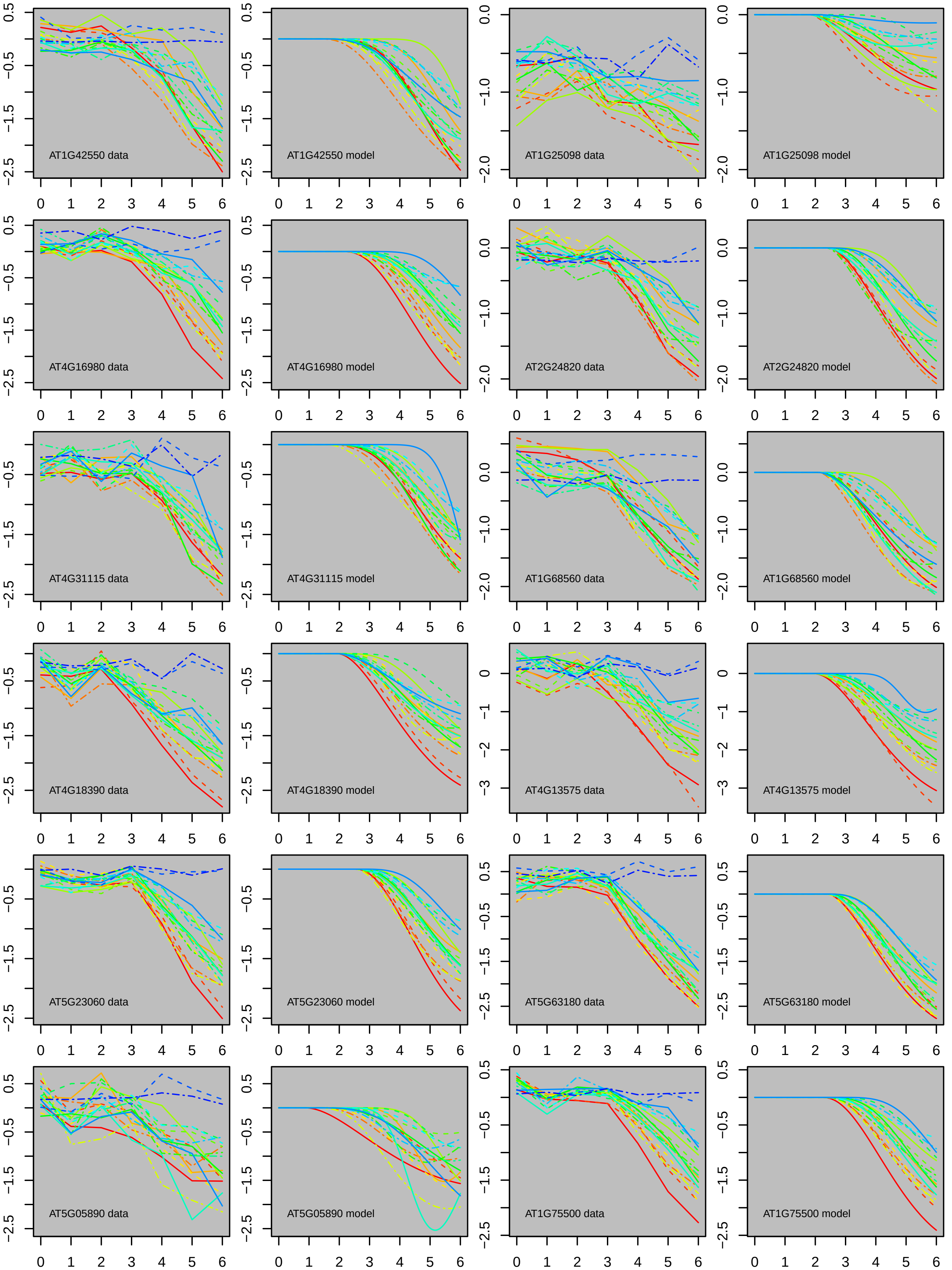
AT1G42550 data
AT1G42550 model
AT1G25098 data
AT1G25098 model
AT4G16980 data
AT4G16980 model
AT2G24820 data
AT2G24820 model
AT4G31115 data
AT4G31115 model
AT1G68560 data
AT1G68560 model
AT4G18390 data
AT4G18390 model
AT4G13575 data
AT4G13575 model
AT5G23060 data
AT5G23060 model
AT5G63180 data
AT5G63180 model
AT5G05890 data
AT5G05890 model
AT1G75500 data
AT1G75500 model

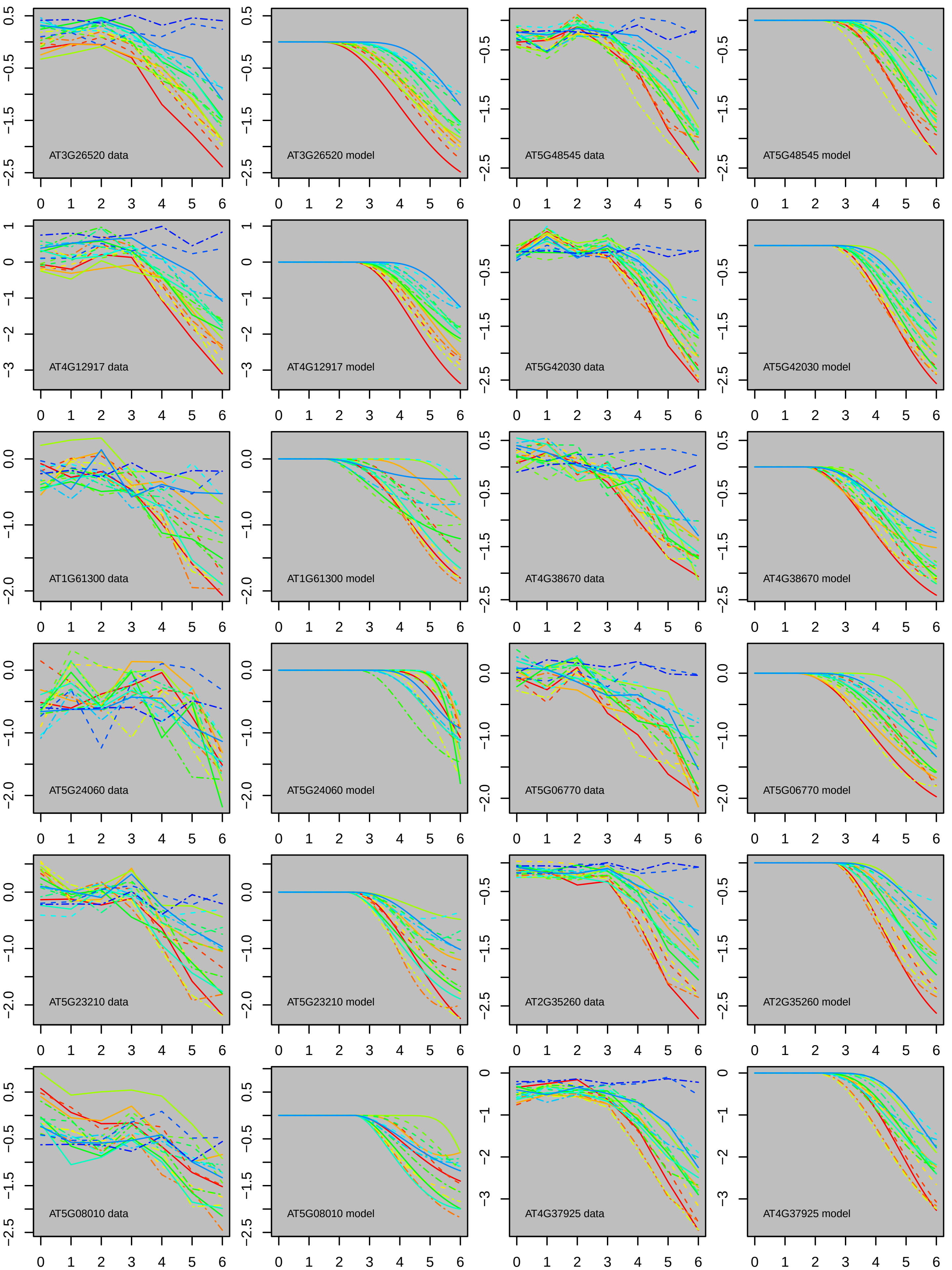
AT3G26520 data
AT3G26520 model
AT5G48545 data
AT5G48545 model
AT4G12917 data
AT4G12917 model
AT5G42030 data
AT5G42030 model
AT1G61300 data
AT1G61300 model
AT4G38670 data
AT4G38670 model
AT5G24060 data
AT5G24060 model
AT5G06770 data
AT5G06770 model
AT5G23210 data
AT5G23210 model
AT2G35260 data
AT2G35260 model
AT5G08010 data
AT5G08010 model
AT4G37925 data
AT4G37925 model

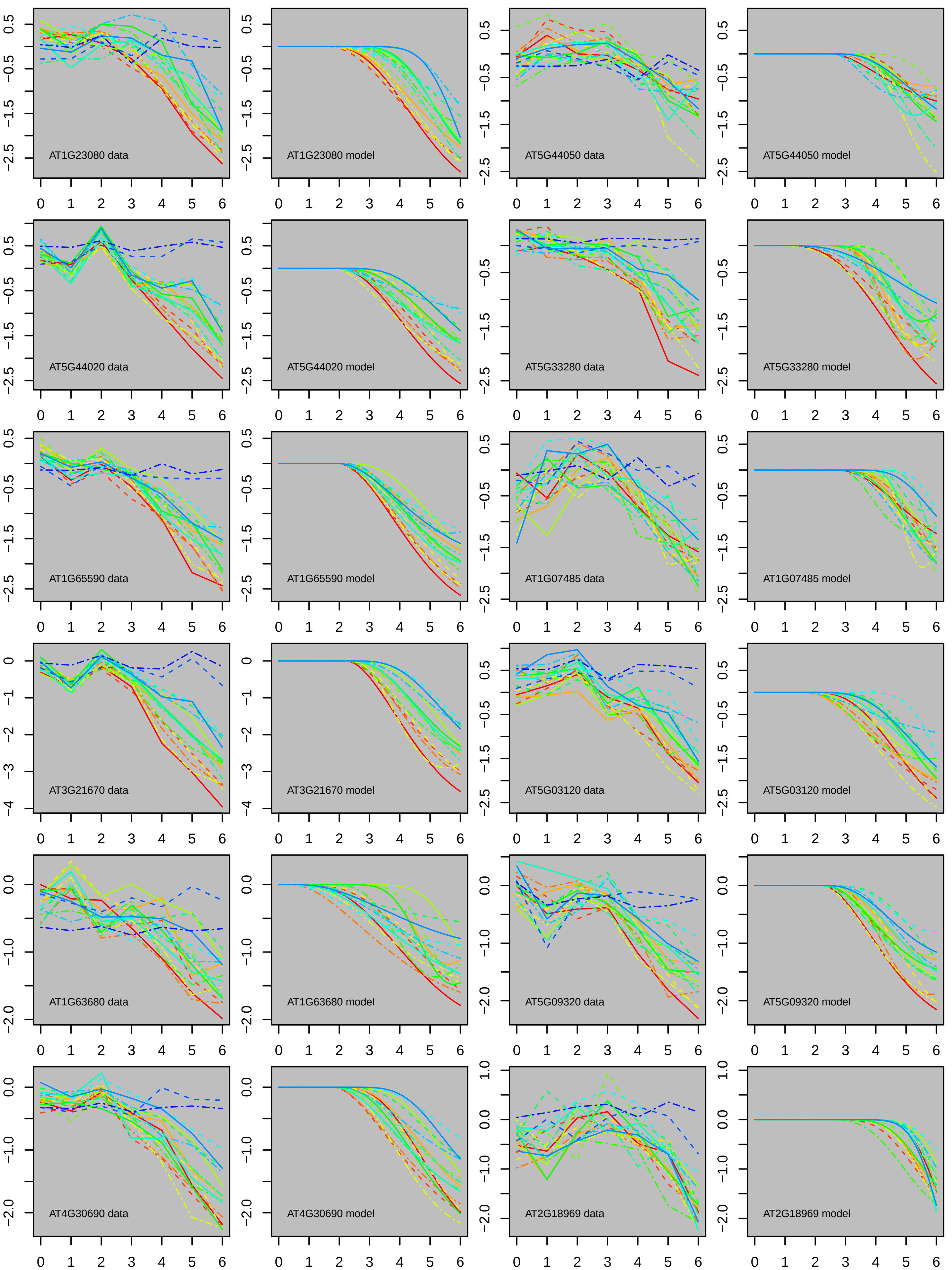

AT1G23080 data
AT1G23080 model
AT5G44050 data
AT5G44050 model
AT5G44020 data
AT5G44020 model
AT5G33280 data
AT5G33280 model
AT1G65590 data
AT1G65590 model
AT1G07485 data
AT1G07485 model
AT3G21670 data
AT3G21670 model
AT5G03120 data
AT5G03120 model
AT1G63680 data
AT1G63680 model
AT5G09320 data
AT5G09320 model
AT4G30690 data
AT4G30690 model
AT2G18969 data
AT2G18969 model

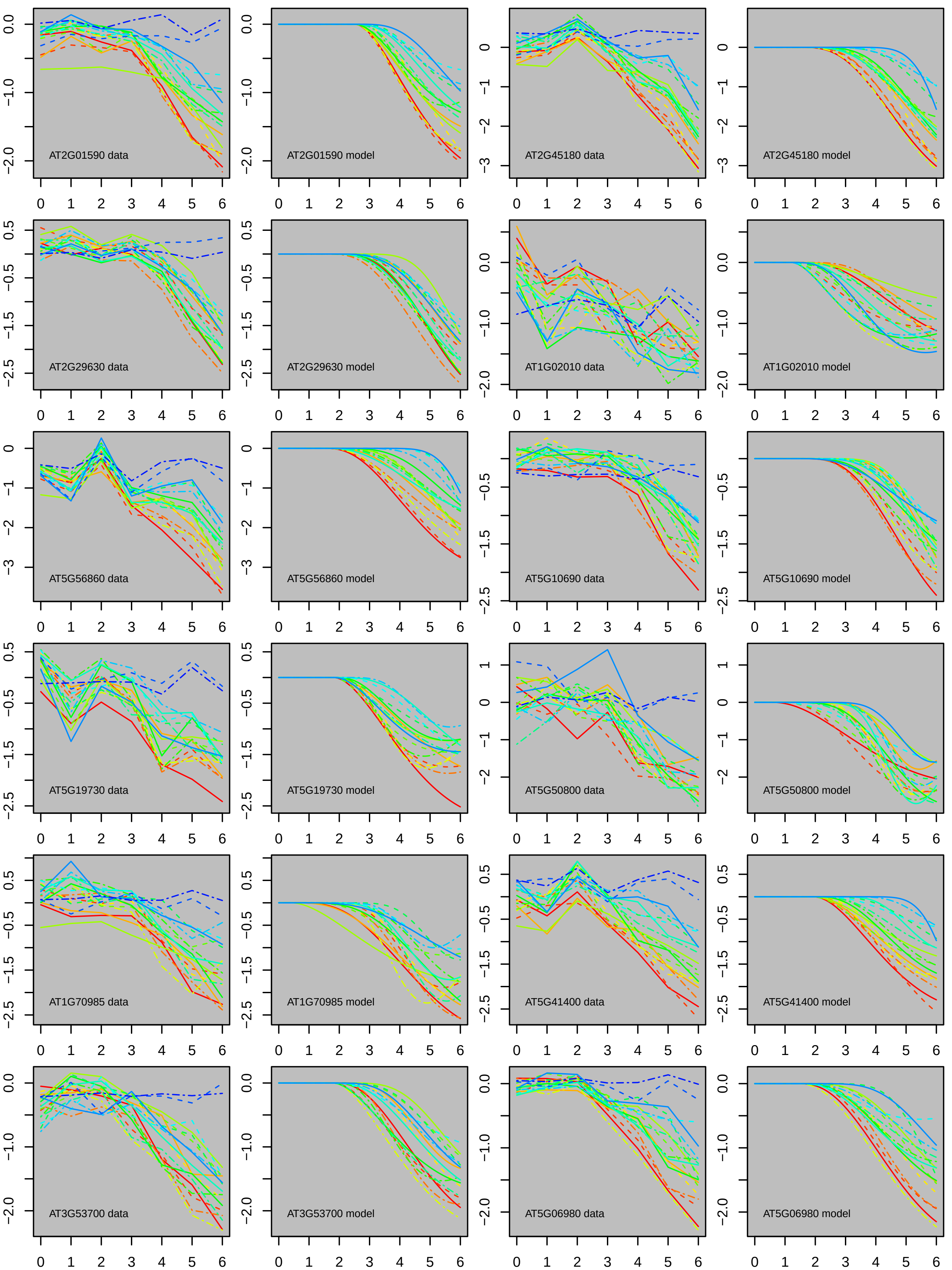

AT2G01590 data
AT2G01590 model
AT2G45180 data
AT2G45180 model
AT2G29630 data
AT2G29630 model
AT1G02010 data
AT1G02010 model
AT5G56860 data
AT5G56860 model
AT5G10690 data
AT5G10690 model
AT5G19730 data
AT5G19730 model
AT5G50800 data
AT5G50800 model
AT1G70985 data
AT1G70985 model
AT5G41400 data
AT5G41400 model
AT3G53700 data
AT3G53700 model
AT5G06980 data
AT5G06980 model

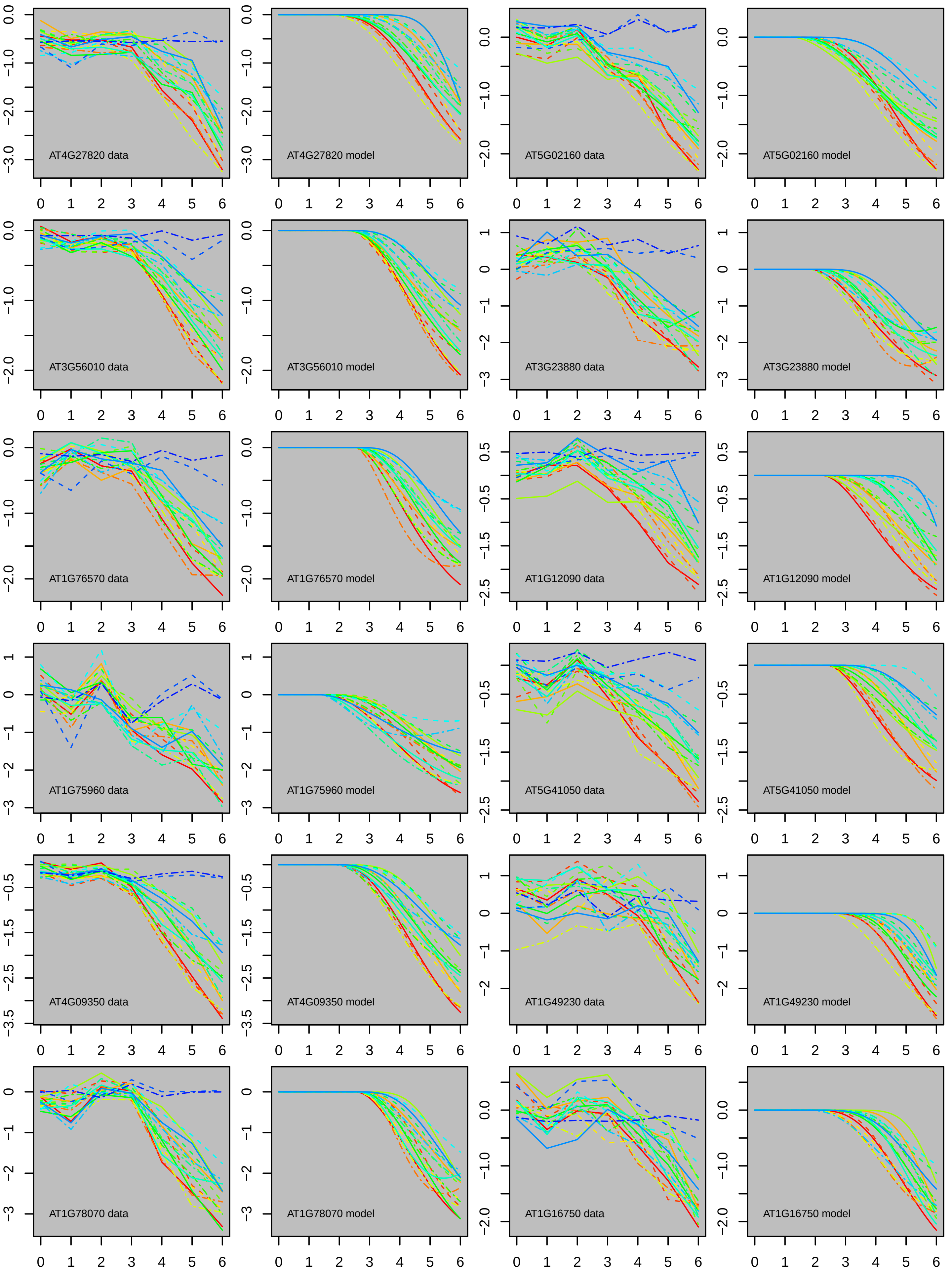

AT4G27820 data
AT4G27820 model
AT5G02160 data
AT5G02160 model
AT3G56010 data
AT3G56010 model
AT3G23880 data
AT3G23880 model
AT1G76570 data
AT1G76570 model
AT1G12090 data
AT1G12090 model
AT1G75960 data
AT1G75960 model
AT5G41050 data
AT5G41050 model
AT4G09350 data
AT4G09350 model
AT1G49230 data
AT1G49230 model
AT1G78070 data
AT1G78070 model
AT1G16750 data
AT1G16750 model

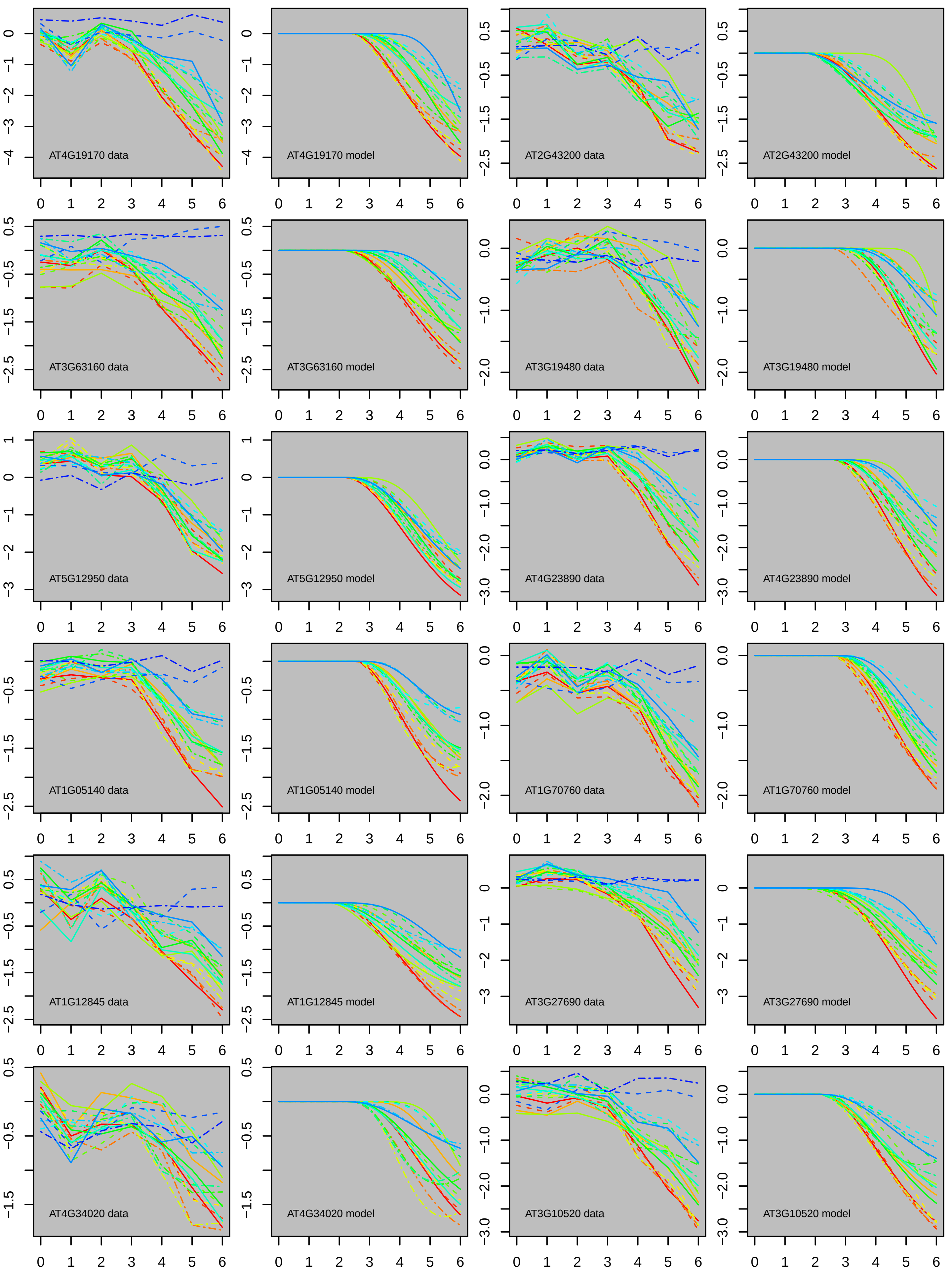
AT4G19170 data
AT4G19170 model
AT2G43200 data
AT2G43200 model
AT3G63160 data
AT3G63160 model
AT3G19480 data
AT3G19480 model
AT5G12950 data
AT5G12950 model
AT4G23890 data
AT4G23890 model
AT1G05140 data
AT1G05140 model
AT1G70760 data
AT1G70760 model
AT1G12845 data
AT1G12845 model
AT3G27690 data
AT3G27690 model
AT4G34020 data
AT4G34020 model
AT3G10520 data
AT3G10520 model

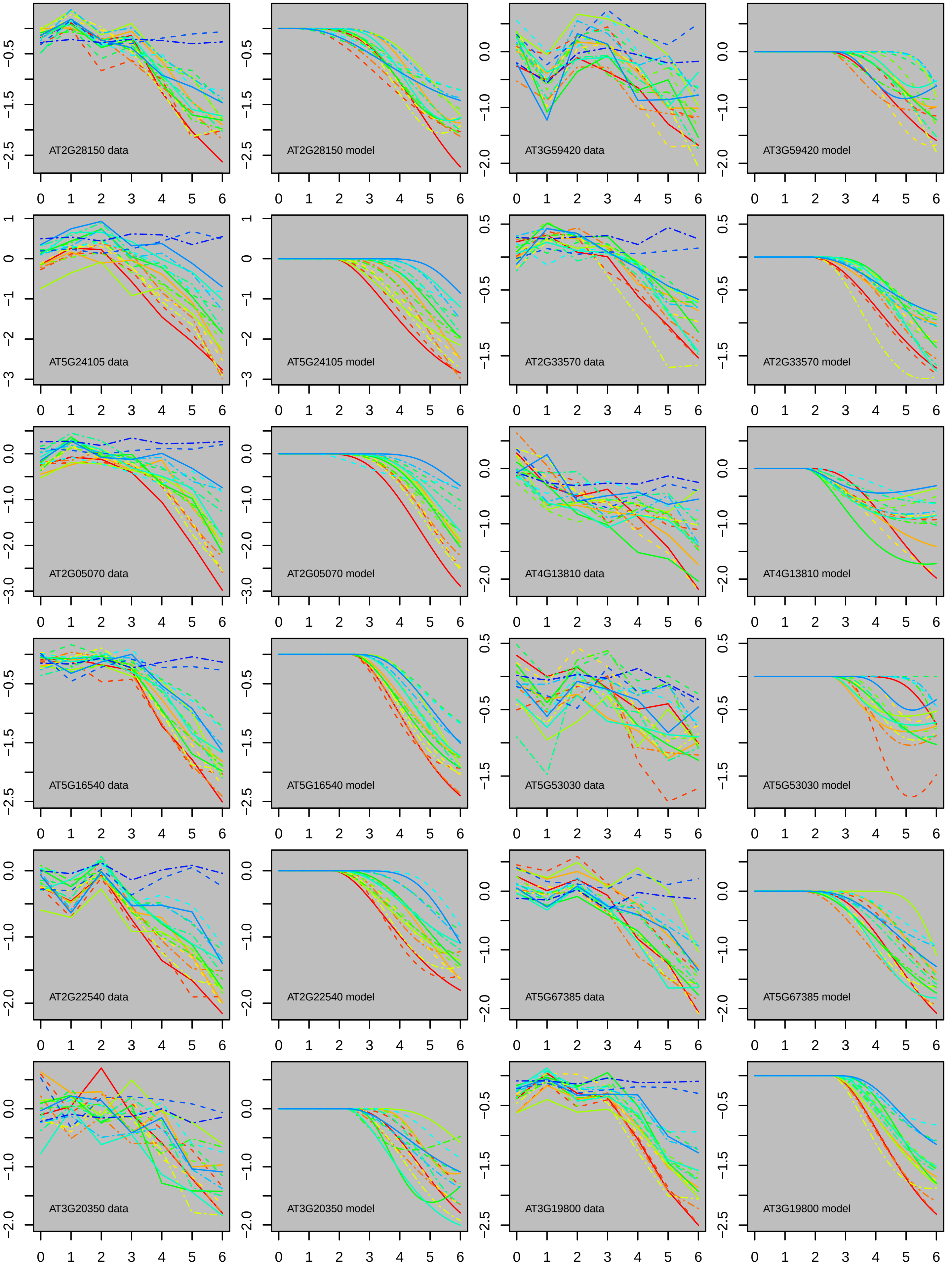
AT2G28150 data
AT2G28150 model
AT3G59420 data
AT3G59420 model
AT5G24105 data
AT5G24105 model
AT2G33570 data
AT2G33570 model
AT2G05070 data
AT2G05070 model
AT4G13810 data
AT4G13810 model
AT5G16540 data
AT5G16540 model
AT5G53030 data
AT5G53030 model
AT2G22540 data
AT2G22540 model
AT5G67385 data
AT5G67385 model
AT3G20350 data
AT3G20350 model
AT3G19800 data
AT3G19800 model

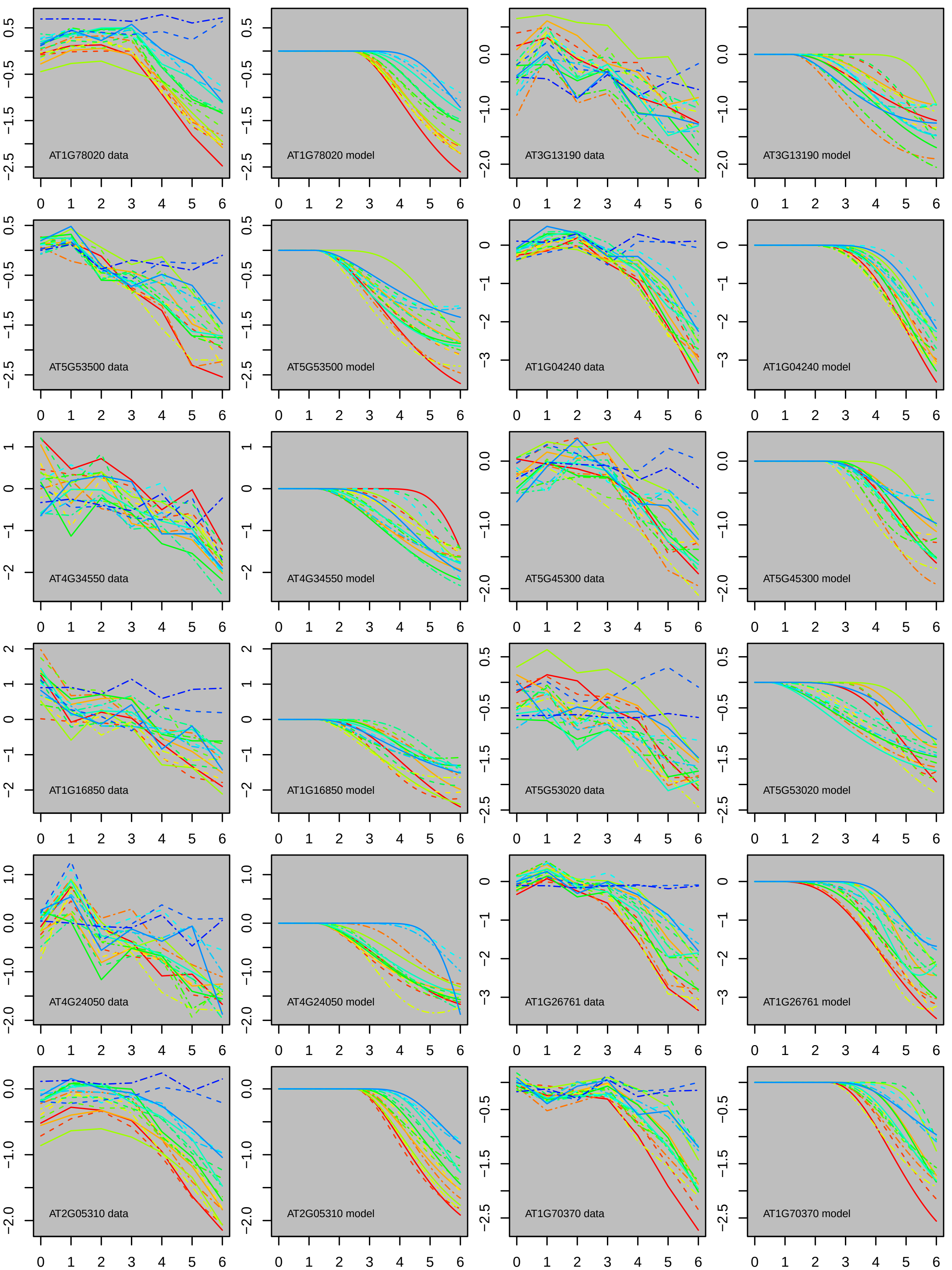
AT1G78020 data
AT1G78020 model
AT3G13190 data
AT3G13190 model
AT5G53500 data
AT5G53500 model
AT1G04240 data
AT1G04240 model
AT4G34550 data
AT4G34550 model
AT5G45300 data
AT5G45300 model
AT1G16850 data
AT1G16850 model
AT5G53020 data
AT5G53020 model
AT4G24050 data
AT4G24050 model
AT1G26761 data
AT1G26761 model
AT2G05310 data
AT2G05310 model
AT1G70370 data
AT1G70370 model

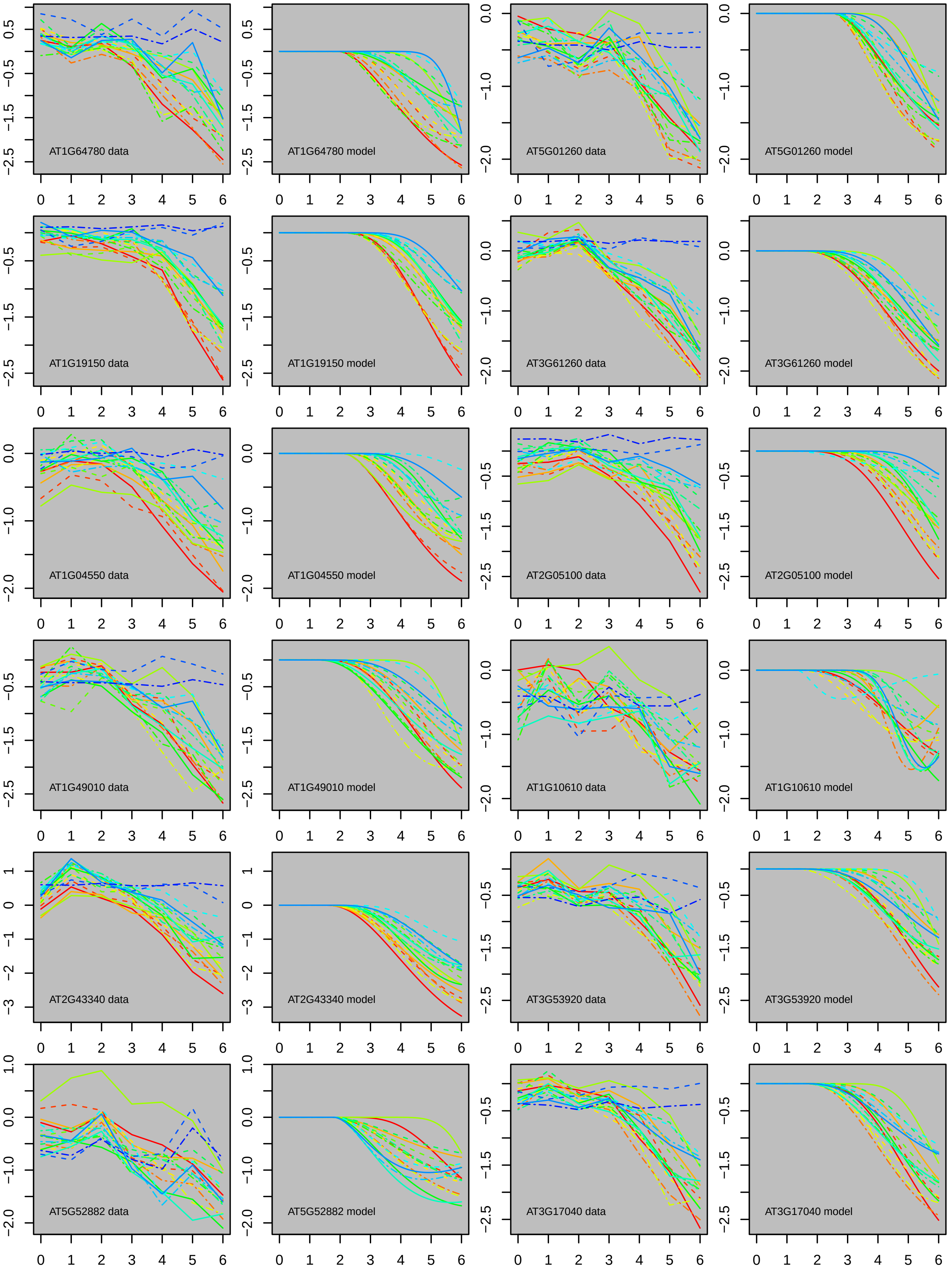

AT1G64780 data
AT1G64780 model
AT5G01260 data
AT5G01260 model
AT1G19150 data
AT1G19150 model
AT3G61260 data
AT3G61260 model
AT1G04550 data
AT1G04550 model
AT2G05100 data
AT2G05100 model
AT1G49010 data
AT1G49010 model
AT1G10610 data
AT1G10610 model
AT2G43340 data
AT2G43340 model
AT3G53920 data
AT3G53920 model
AT5G52882 data
AT5G52882 model
AT3G17040 data
AT3G17040 model

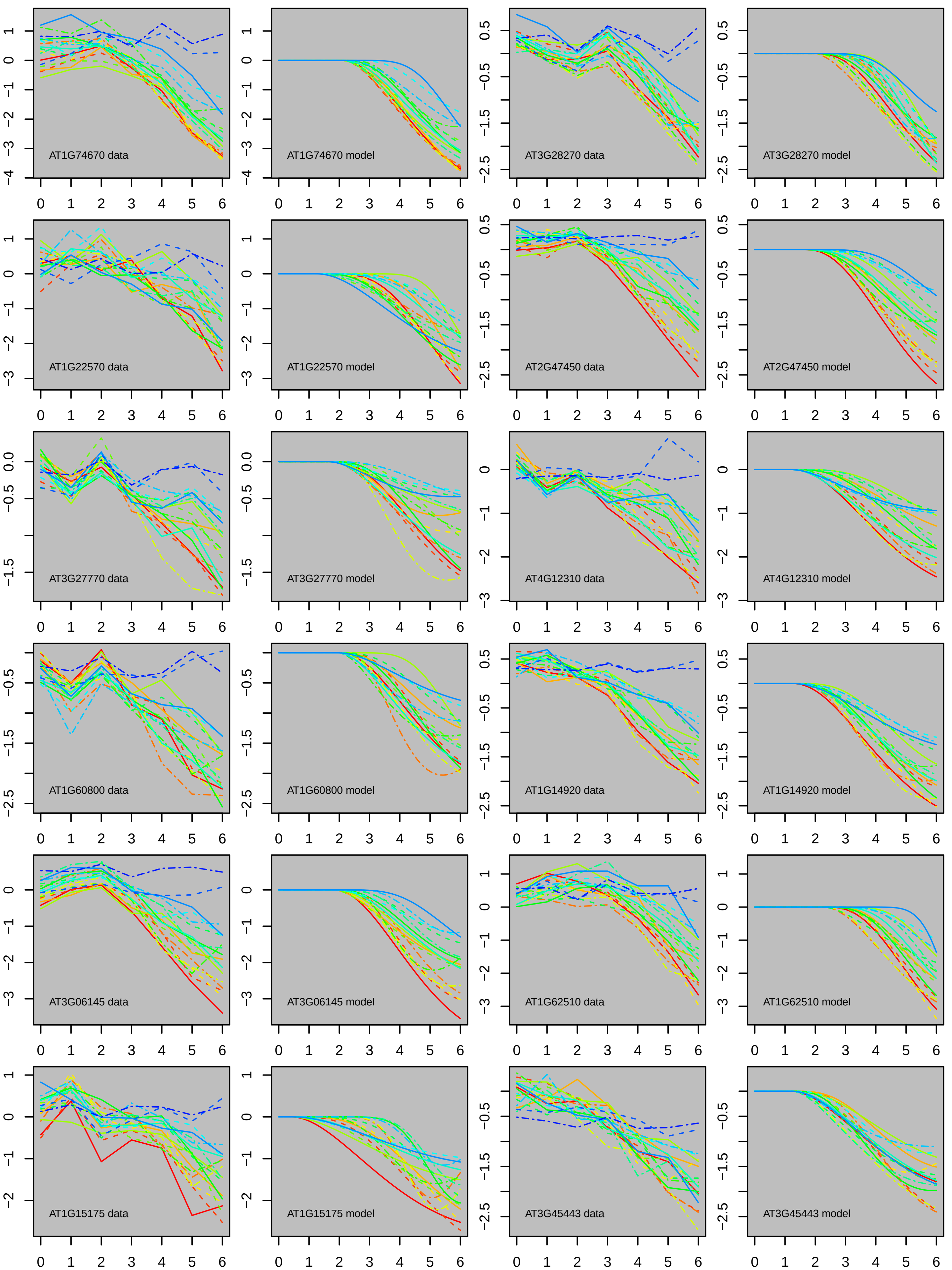
AT1G74670 data
AT1G74670 model
AT3G28270 data
AT3G28270 model
AT1G22570 data
AT1G22570 model
AT2G47450 data
AT2G47450 model
AT3G27770 data
AT3G27770 model
AT4G12310 data
AT4G12310 model
AT1G60800 data
AT1G60800 model
AT1G14920 data
AT1G14920 model
AT3G06145 data
AT3G06145 model
AT1G62510 data
AT1G62510 model
AT1G15175 data
AT1G15175 model
AT3G45443 data
AT3G45443 model

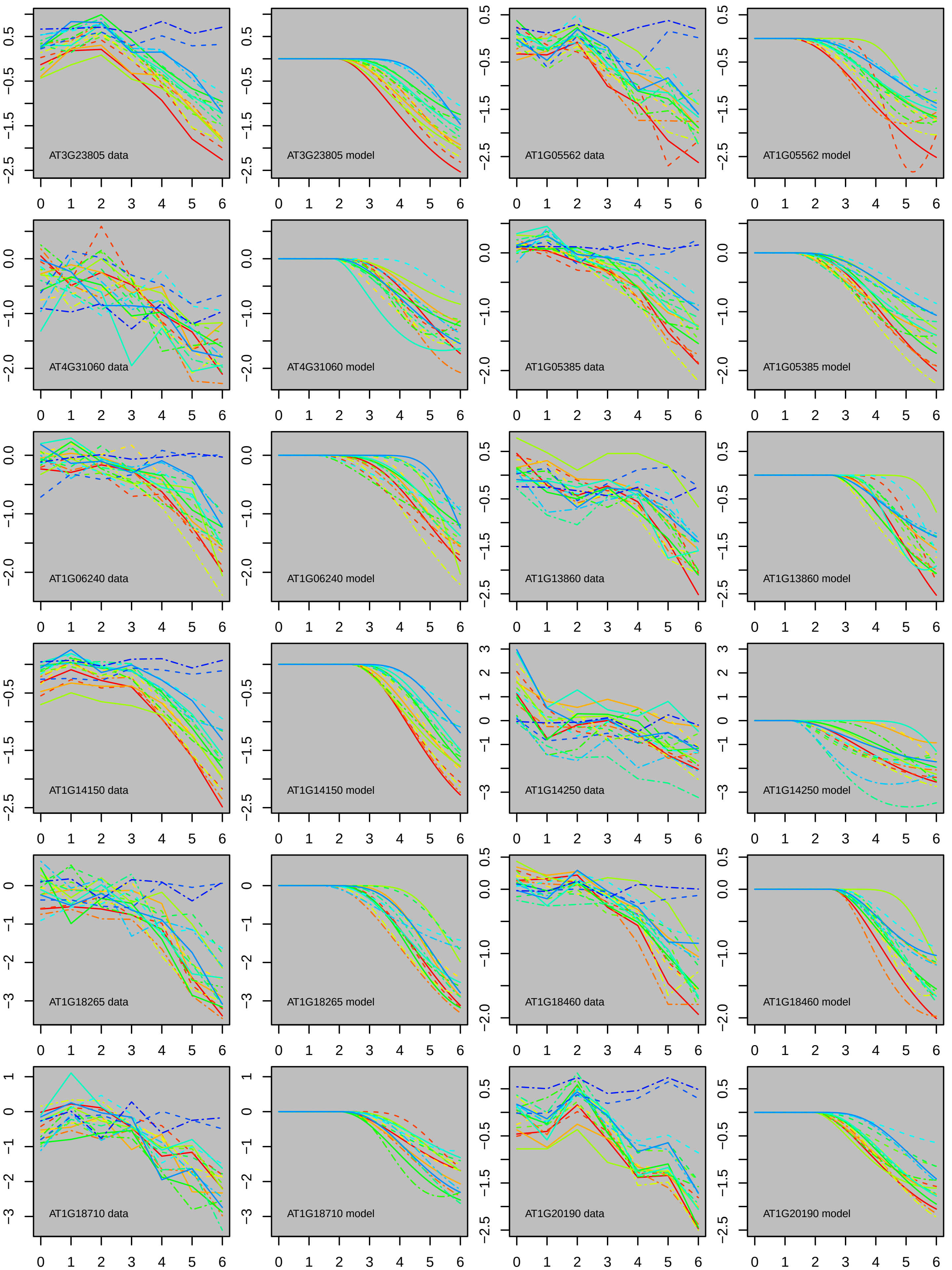
AT3G23805 data
AT3G23805 model
AT1G05562 data
AT1G05562 model
AT4G31060 data
AT4G31060 model
AT1G05385 data
AT1G05385 model
AT1G06240 data
AT1G06240 model
AT1G13860 data
AT1G13860 model
AT1G14150 data
AT1G14150 model
AT1G14250 data
AT1G14250 model
AT1G18265 data
AT1G18265 model
AT1G18460 data
AT1G18460 model
AT1G18710 data
AT1G18710 model
AT1G20190 data
AT1G20190 model

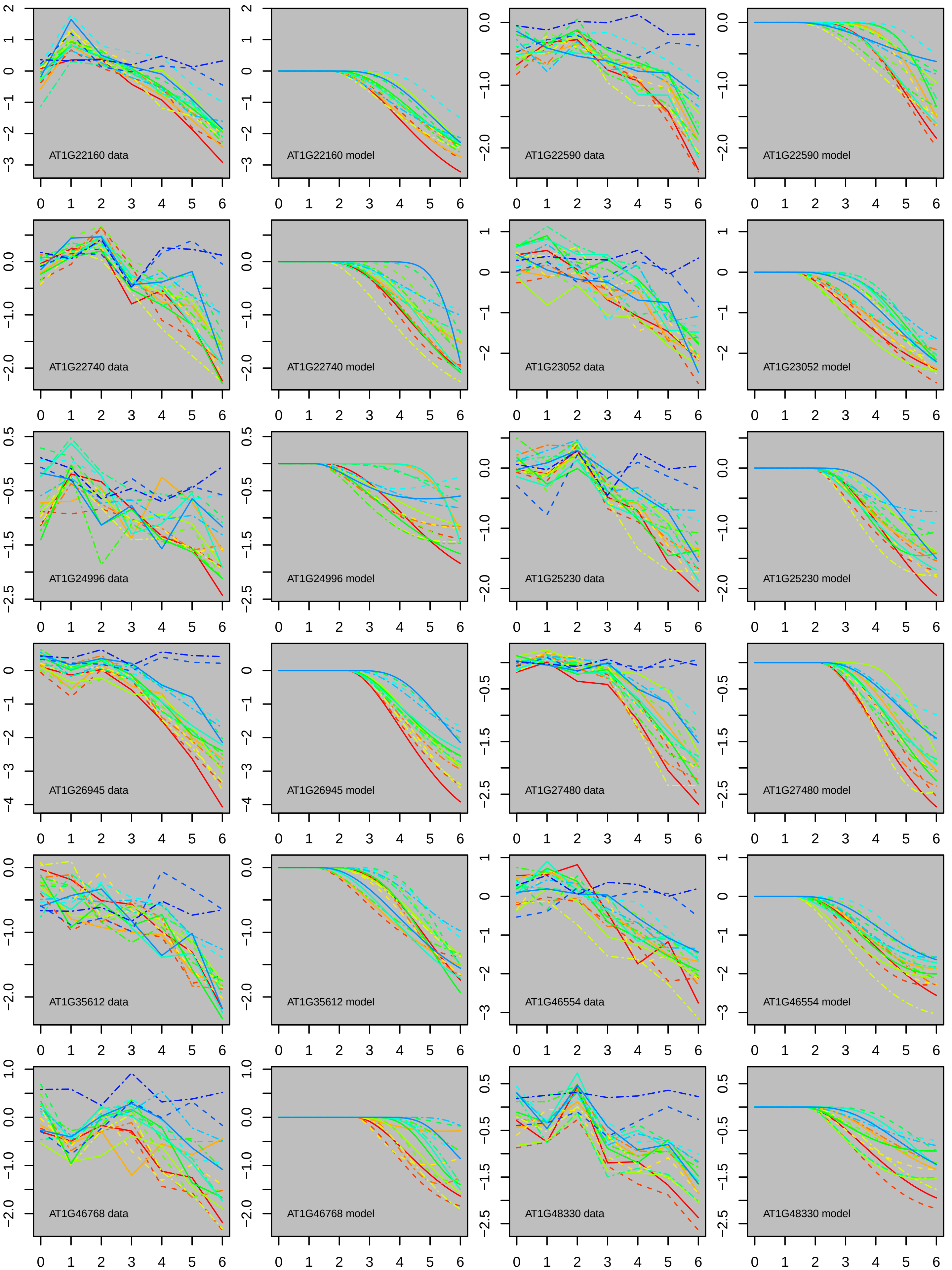
AT1G22160 data
AT1G22160 model
AT1G22590 data
AT1G22590 model
AT1G22740 data
AT1G22740 model
AT1G23052 data
AT1G23052 model
AT1G24996 data
AT1G24996 model
AT1G25230 data
AT1G25230 model
AT1G26945 data
AT1G26945 model
AT1G27480 data
AT1G27480 model
AT1G35612 data
AT1G35612 model
AT1G46554 data
AT1G46554 model
AT1G46768 data
AT1G46768 model
AT1G48330 data
AT1G48330 model

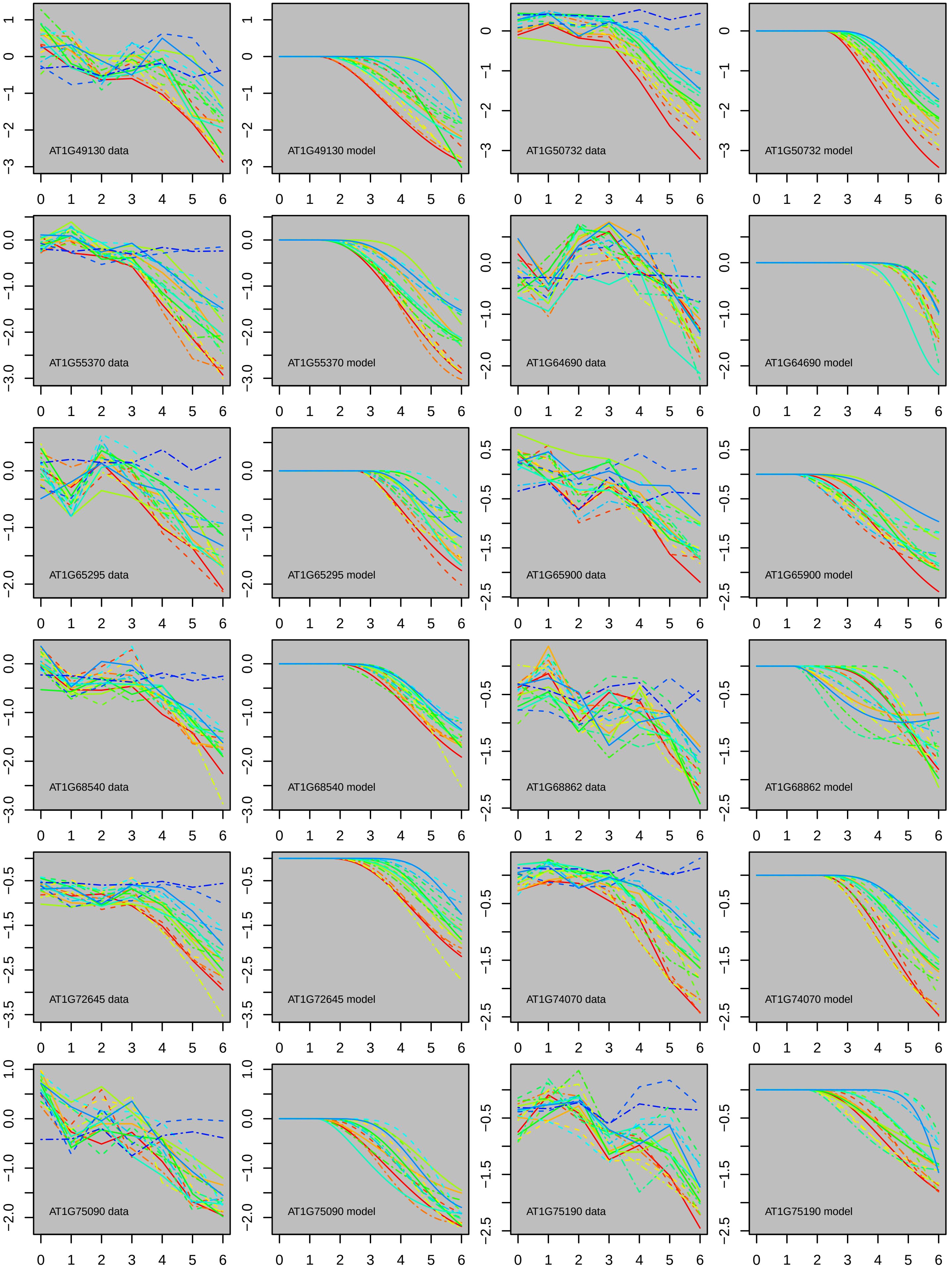
AT1G49130 data
AT1G49130 model
AT1G50732 data
AT1G50732 model
AT1G55370 data
AT1G55370 model
AT1G64690 data
AT1G64690 model
AT1G65295 data
AT1G65295 model
AT1G65900 data
AT1G65900 model
AT1G68540 data
AT1G68540 model
AT1G68862 data
AT1G68862 model
AT1G72645 data
AT1G72645 model
AT1G74070 data
AT1G74070 model
AT1G75090 data
AT1G75090 model
AT1G75190 data
AT1G75190 model

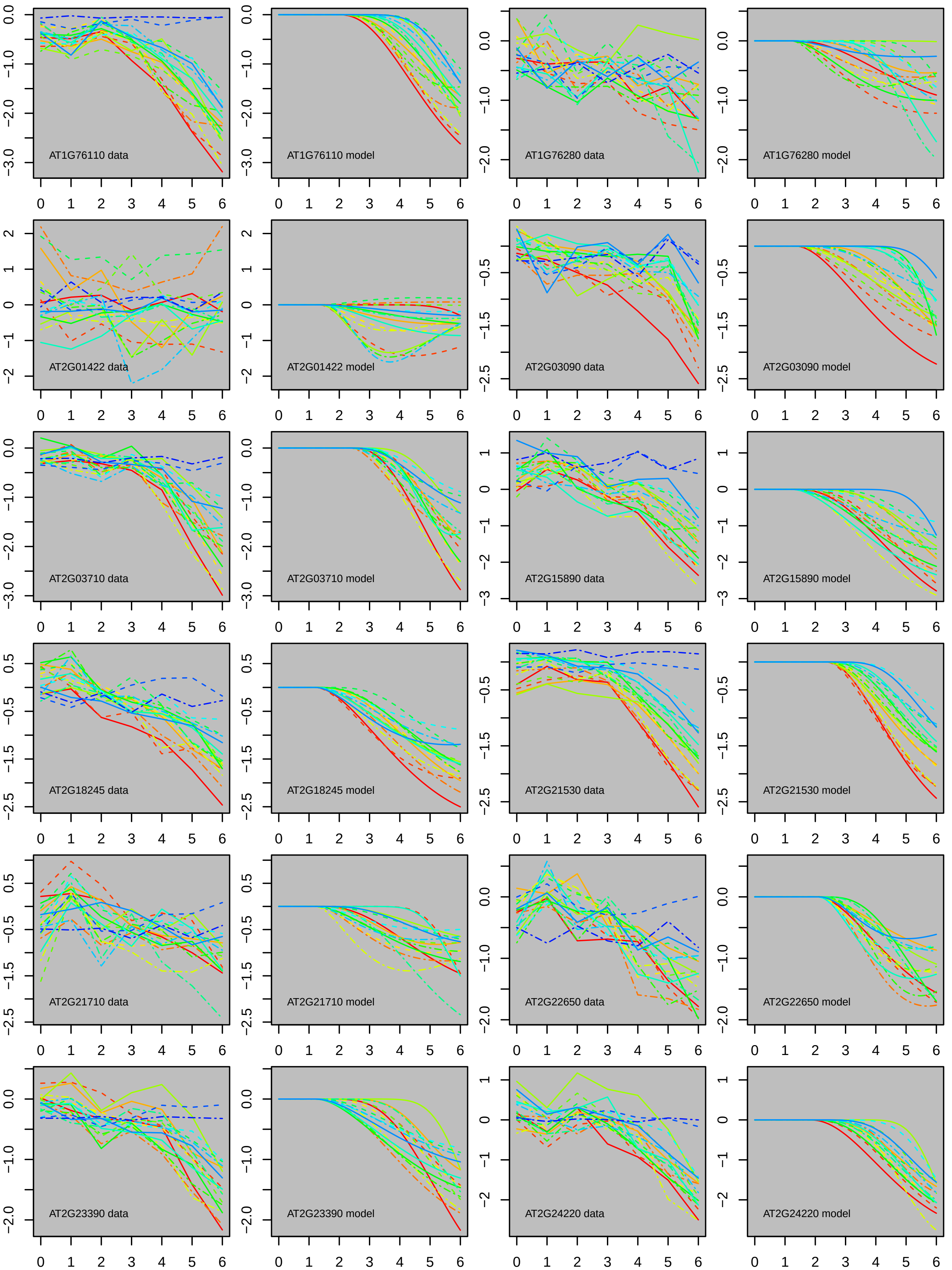
AT1G76110 data
AT1G76110 model
AT1G76280 data
AT1G76280 model
AT2G01422 data
AT2G01422 model
AT2G03090 data
AT2G03090 model
AT2G03710 data
AT2G03710 model
AT2G15890 data
AT2G15890 model
AT2G18245 data
AT2G18245 model
AT2G21530 data
AT2G21530 model
AT2G21710 data
AT2G21710 model
AT2G22650 data
AT2G22650 model
AT2G23390 data
AT2G23390 model
AT2G24220 data
AT2G24220 model

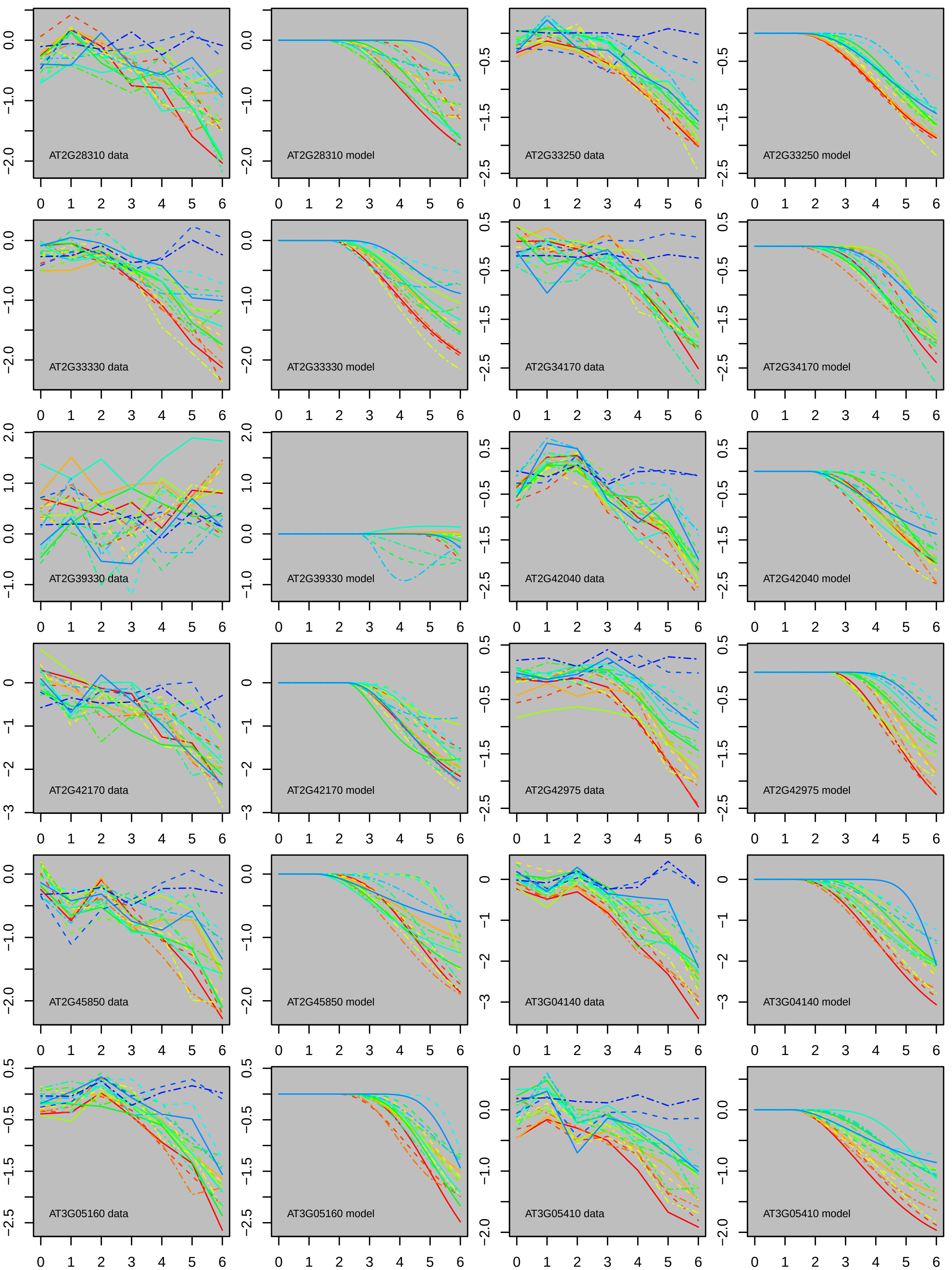
AT2G28310 data
AT2G28310 model
AT2G33250 data
AT2G33250 model
AT2G33330 data
AT2G33330 model
AT2G34170 data
AT2G34170 model
AT2G39330 data
AT2G39330 model
AT2G42040 data
AT2G42040 model
AT2G42170 data
AT2G42170 model
AT2G42975 data
AT2G42975 model
AT2G45850 data
AT2G45850 model
AT3G04140 data
AT3G04140 model
AT3G05160 data
AT3G05160 model
AT3G05410 data
AT3G05410 model

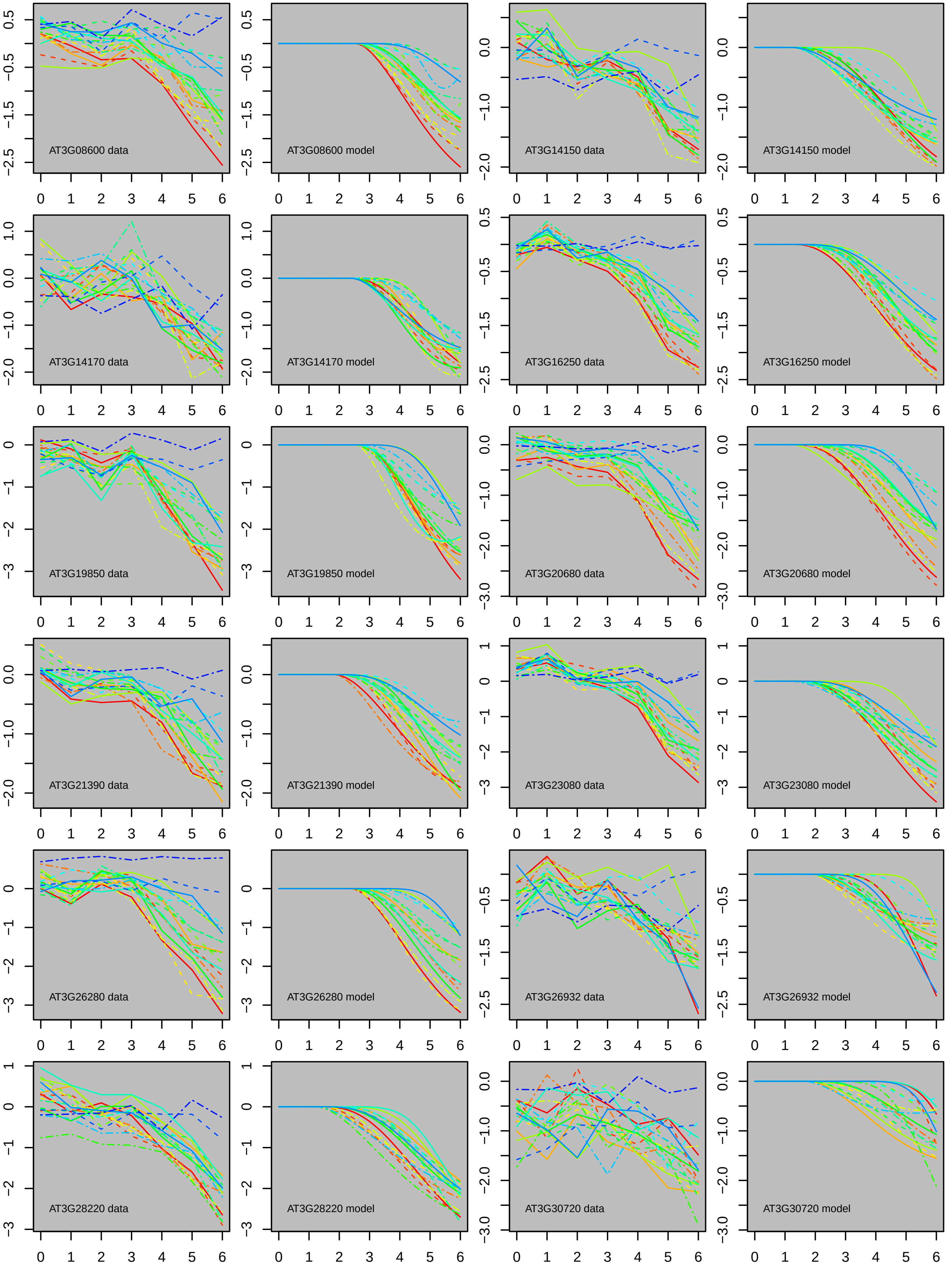
AT3G08600 data
AT3G08600 model
AT3G14150 data
AT3G14150 model
AT3G14170 data
AT3G14170 model
AT3G16250 data
AT3G16250 model
AT3G19850 data
AT3G19850 model
AT3G20680 data
AT3G20680 model
AT3G21390 data
AT3G21390 model
AT3G23080 data
AT3G23080 model
AT3G26280 data
AT3G26280 model
AT3G26932 data
AT3G26932 model
AT3G28220 data
AT3G28220 model
AT3G30720 data
AT3G30720 model

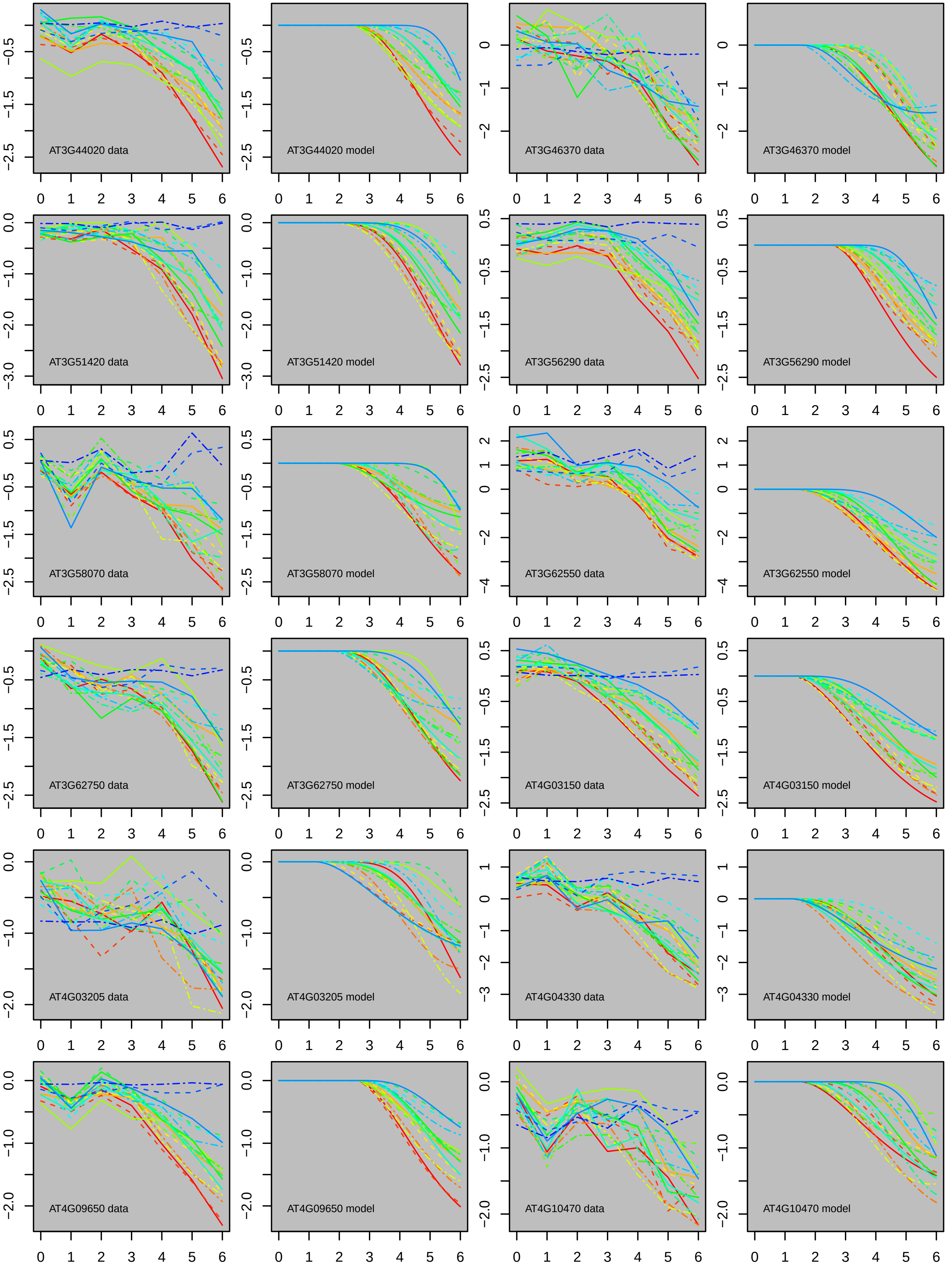
AT3G44020 data
AT3G44020 model
AT3G46370 data
AT3G46370 model
AT3G51420 data
AT3G51420 model
AT3G56290 data
AT3G56290 model
AT3G58070 data
AT3G58070 model
AT3G62550 data
AT3G62550 model
AT3G62750 data
AT3G62750 model
AT4G03150 data
AT4G03150 model
AT4G03205 data
AT4G03205 model
AT4G04330 data
AT4G04330 model
AT4G09650 data
AT4G09650 model
AT4G10470 data
AT4G10470 model

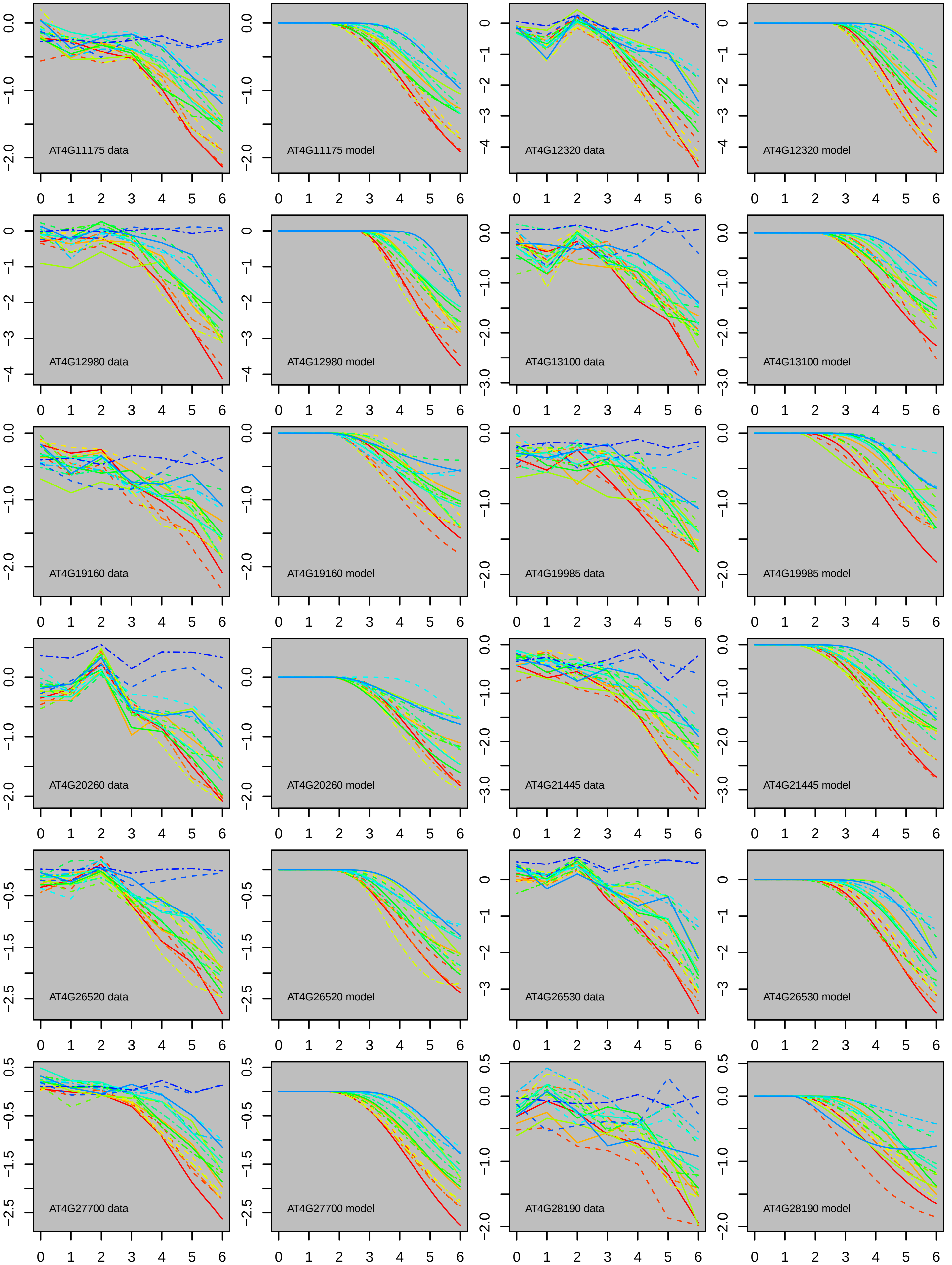

AT4G11175 data
AT4G11175 model
AT4G12320 data
AT4G12320 model
AT4G12980 data
AT4G12980 model
AT4G13100 data
AT4G13100 model
AT4G19160 data
AT4G19160 model
AT4G19985 data
AT4G19985 model
AT4G20260 data
AT4G20260 model
AT4G21445 data
AT4G21445 model
AT4G26520 data
AT4G26520 model
AT4G26530 data
AT4G26530 model
AT4G27700 data
AT4G27700 model
AT4G28190 data
AT4G28190 model

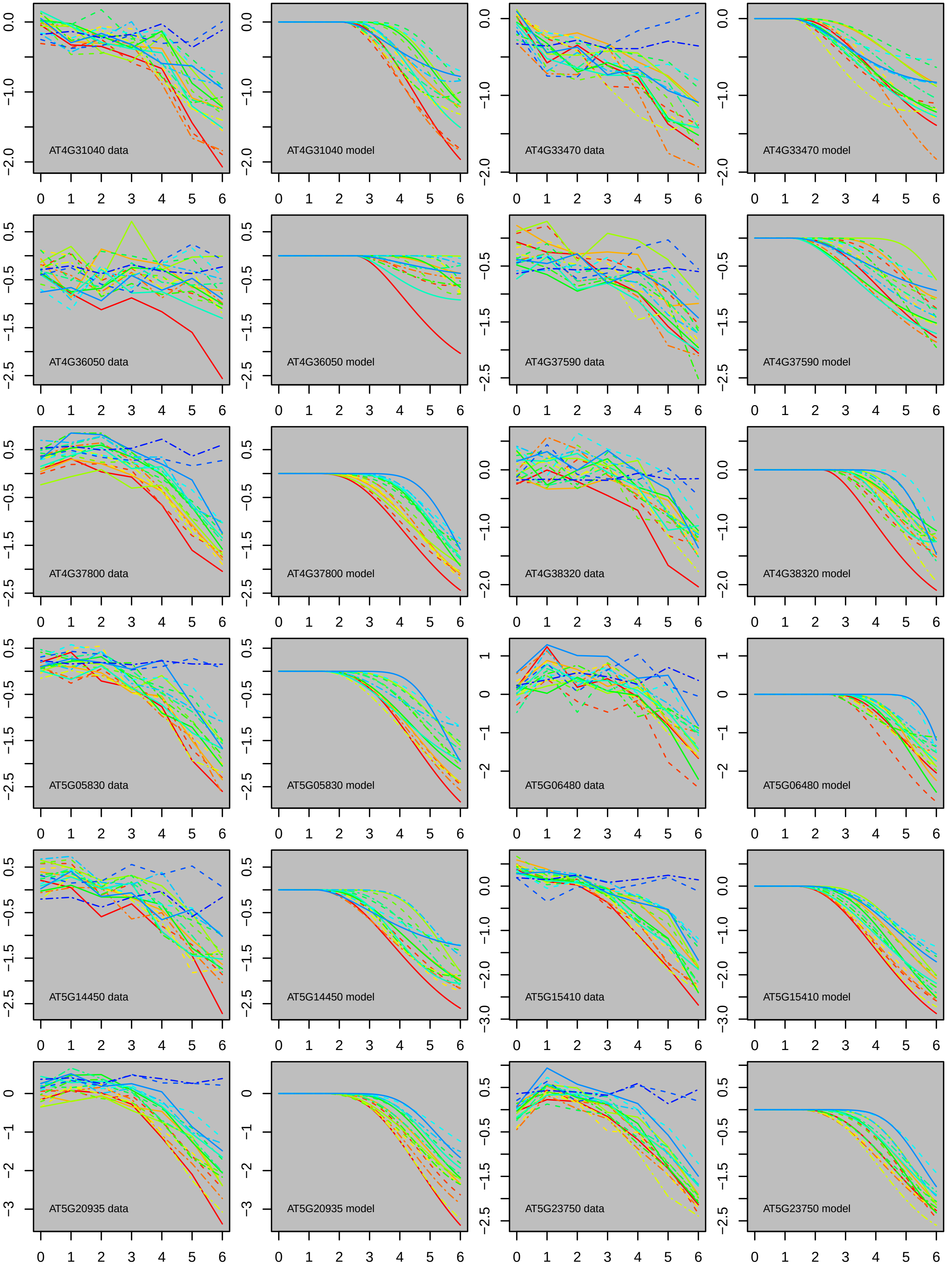

AT4G31040 data
AT4G31040 model
AT4G33470 data
AT4G33470 model
AT4G36050 data
AT4G36050 model
AT4G37590 data
AT4G37590 model
AT4G37800 data
AT4G37800 model
AT4G38320 data
AT4G38320 model
AT5G05830 data
AT5G05830 model
AT5G06480 data
AT5G06480 model
AT5G14450 data
AT5G14450 model
AT5G15410 data
AT5G15410 model
AT5G20935 data
AT5G20935 model
AT5G23750 data
AT5G23750 model

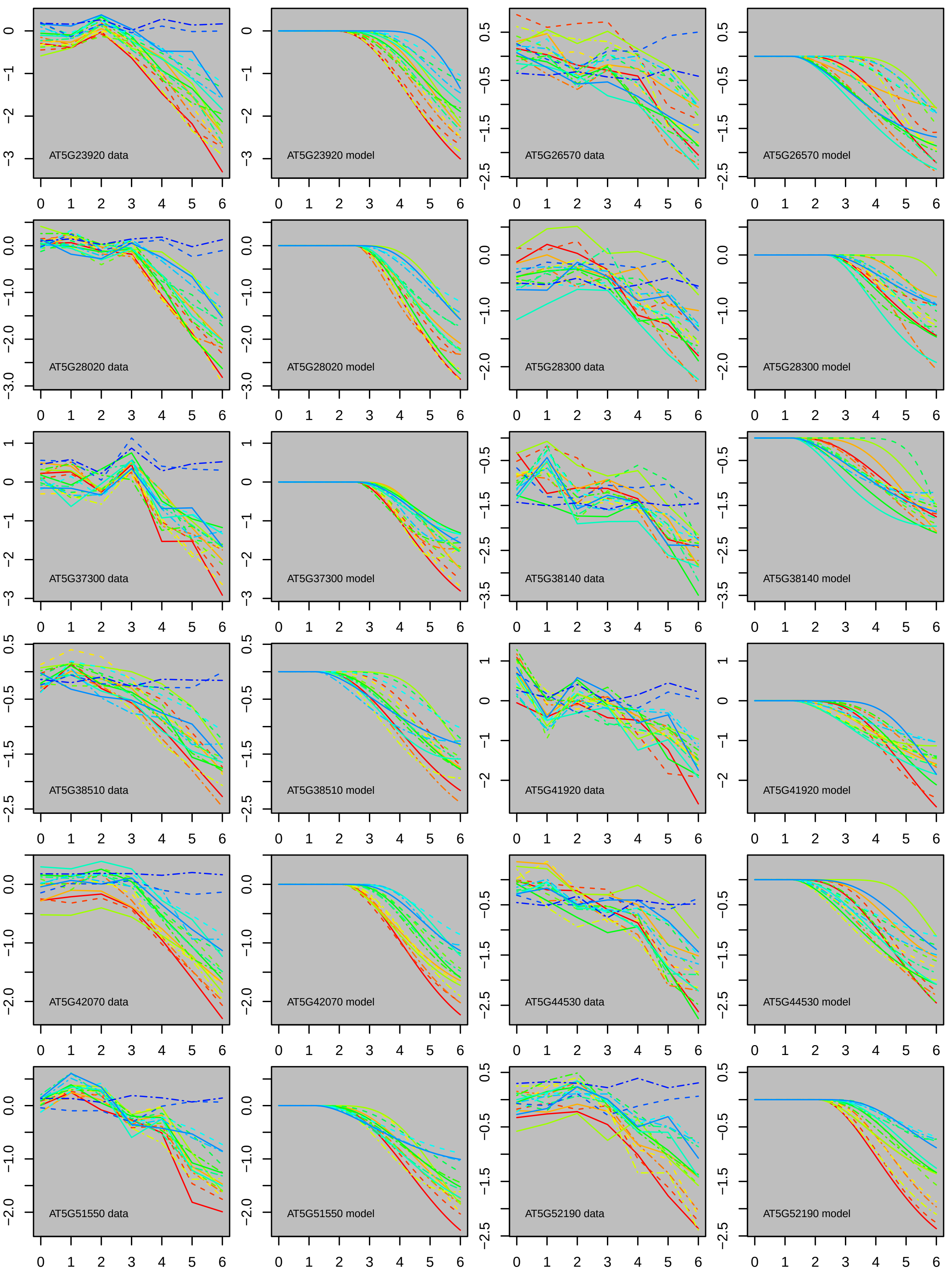
AT5G23920 data
AT5G23920 model
AT5G26570 data
AT5G26570 model
AT5G28020 data
AT5G28020 model
AT5G28300 data
AT5G28300 model
AT5G37300 data
AT5G37300 model
AT5G38140 data
AT5G38140 model
AT5G38510 data
AT5G38510 model
AT5G41920 data
AT5G41920 model
AT5G42070 data
AT5G42070 model
AT5G44530 data
AT5G44530 model
AT5G51550 data
AT5G51550 model
AT5G52190 data
AT5G52190 model

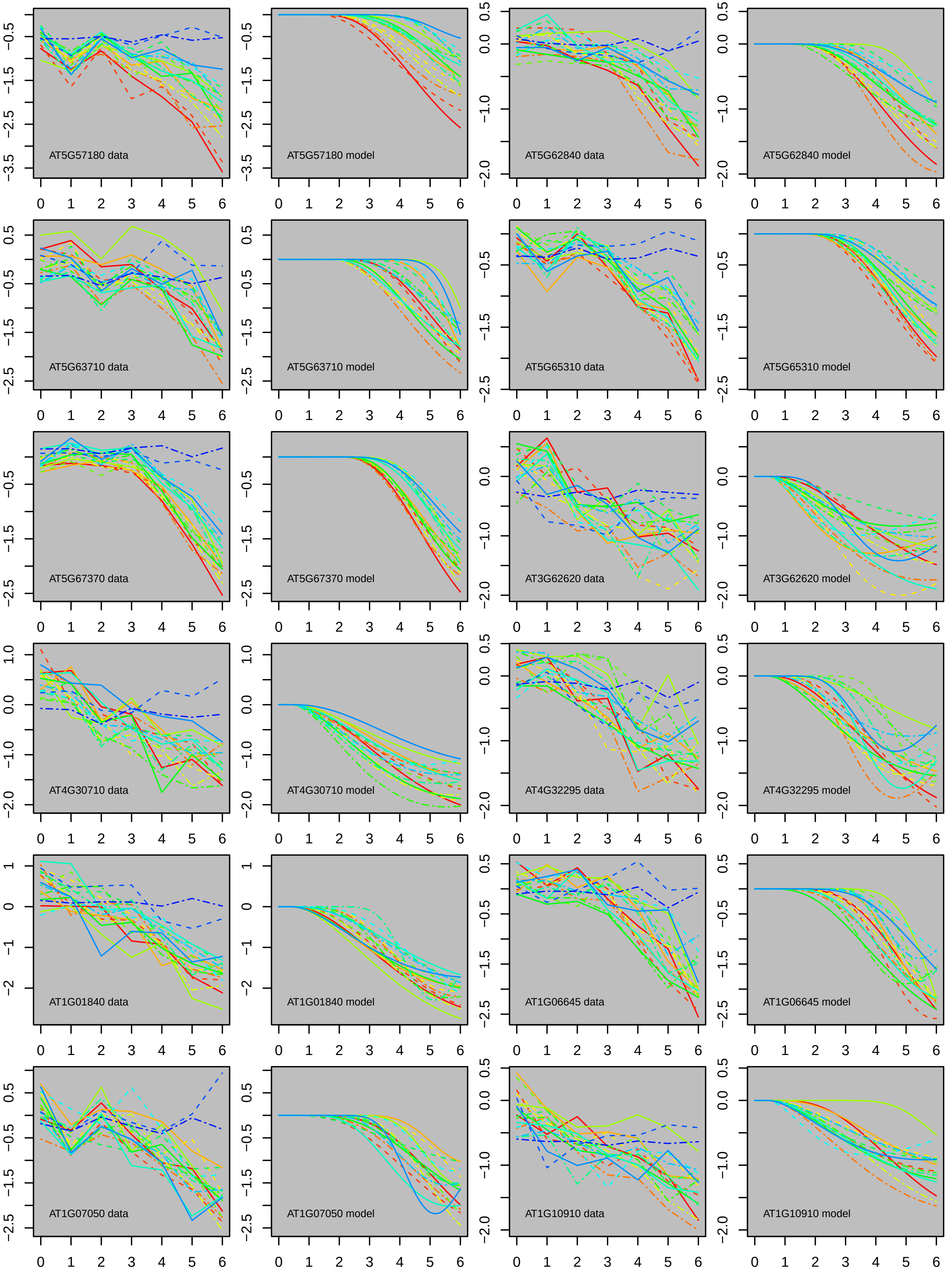

AT5G57180 data
AT5G57180 model
AT5G62840 data
AT5G62840 model
AT5G63710 data
AT5G63710 model
AT5G65310 data
AT5G65310 model
AT5G67370 data
AT5G67370 model
AT3G62620 data
AT3G62620 model
AT4G30710 data
AT4G30710 model
AT4G32295 data
AT4G32295 model
AT1G01840 data
AT1G01840 model
AT1G06645 data
AT1G06645 model
AT1G07050 data
AT1G07050 model
AT1G10910 data
AT1G10910 model

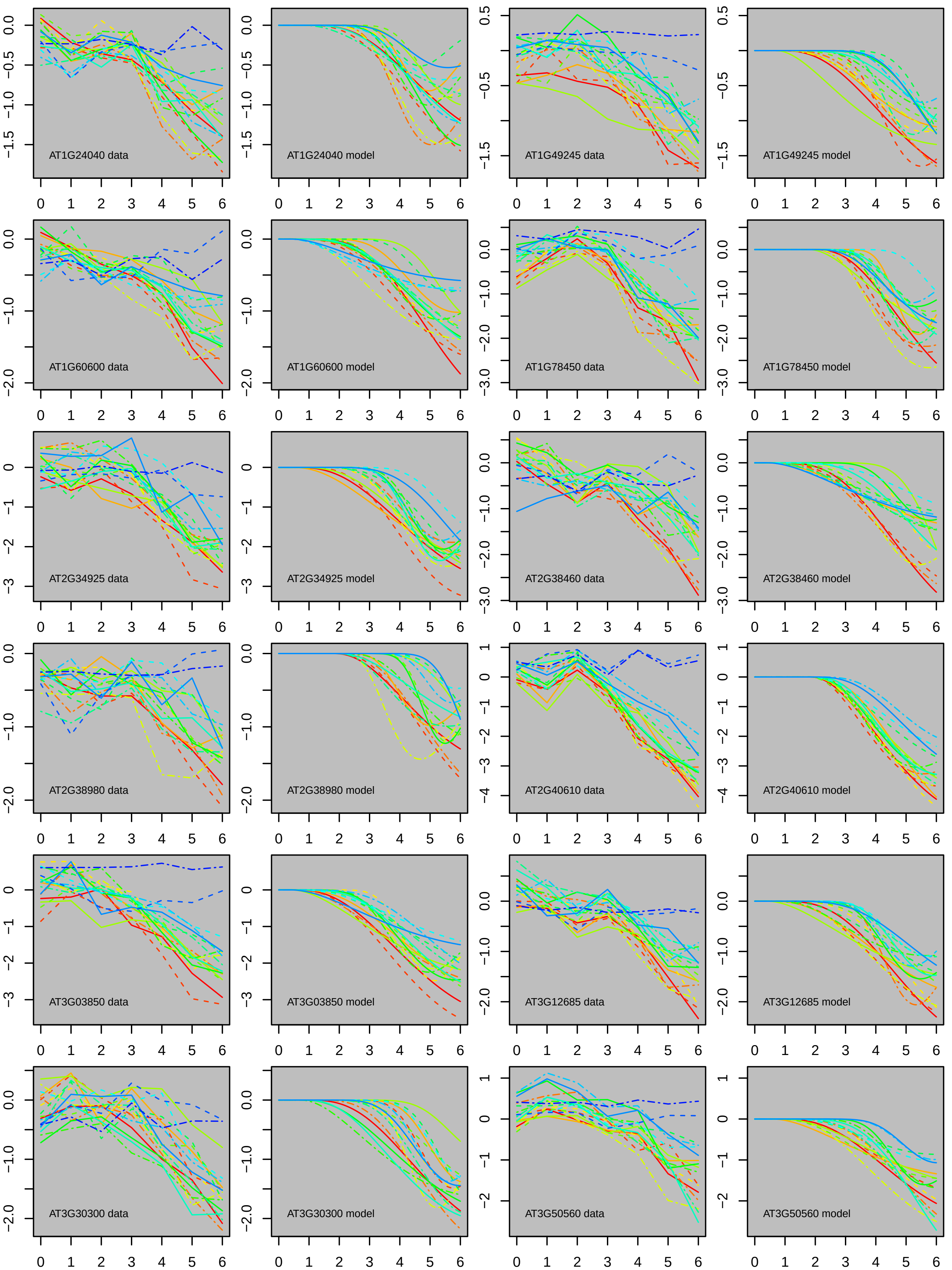
AT1G24040 data
AT1G24040 model
AT1G49245 data
AT1G49245 model
AT1G60600 data
AT1G60600 model
AT1G78450 data
AT1G78450 model
AT2G34925 data
AT2G34925 model
AT2G38460 data
AT2G38460 model
AT2G38980 data
AT2G38980 model
AT2G40610 data
AT2G40610 model
AT3G03850 data
AT3G03850 model
AT3G12685 data
AT3G12685 model
AT3G30300 data
AT3G30300 model
AT3G50560 data
AT3G50560 model

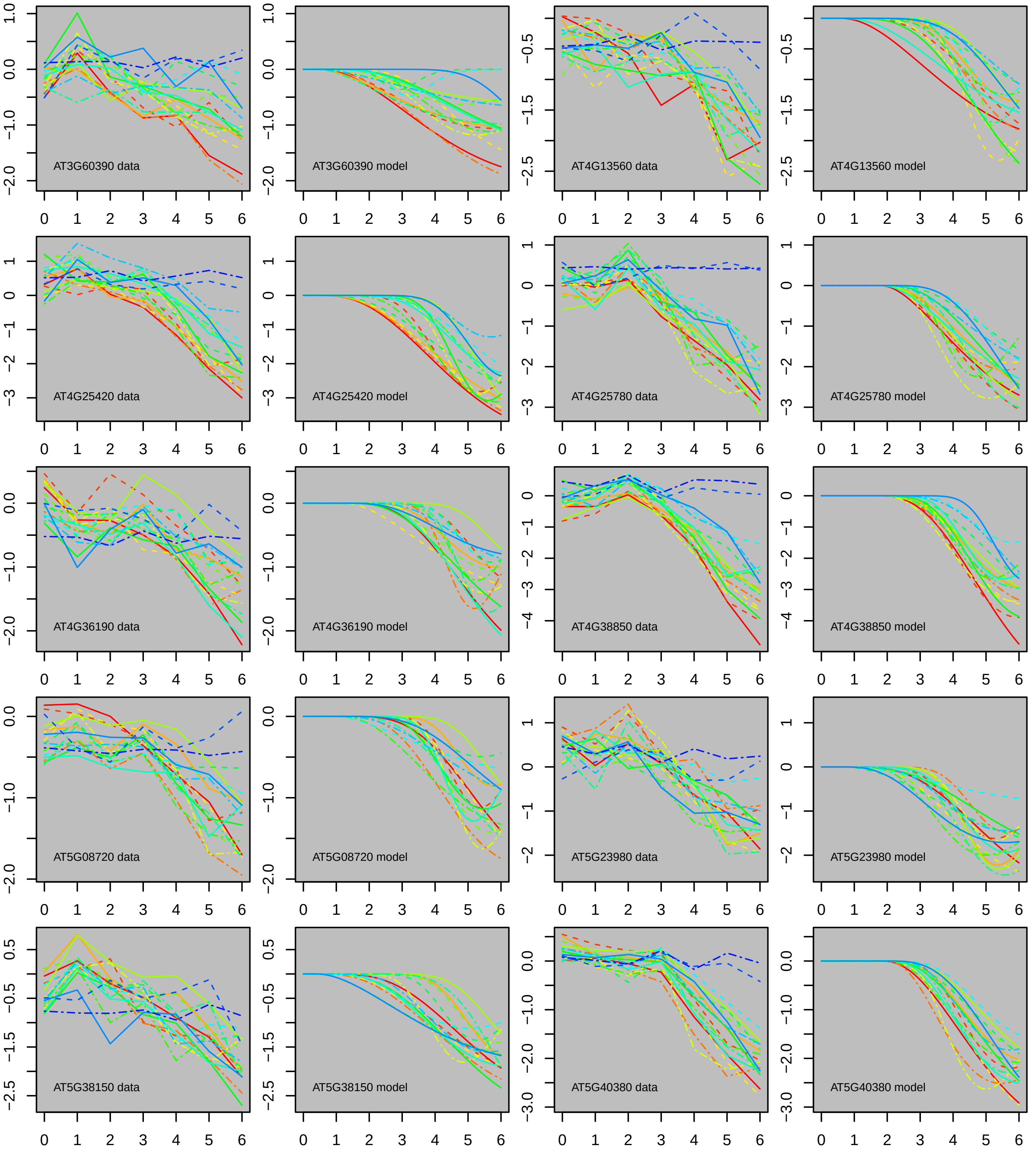

AT3G60390 data
AT3G60390 model
AT4G13560 data
AT4G13560 model
AT4G25420 data
AT4G25420 model
AT4G25780 data
AT4G25780 model
AT4G36190 data
AT4G36190 model
AT4G38850 data
AT4G38850 model
AT5G08720 data
AT5G08720 model
AT5G23980 data
AT5G23980 model
AT5G38150 data
AT5G38150 model
AT5G40380 data
AT5G40380 model
