## Supplementary material for "The kinetics and basal levels of the transcriptome response during Effector-Triggered Immunity in Arabidopsis are mainly controlled by four immune signaling sectors": Figure S5

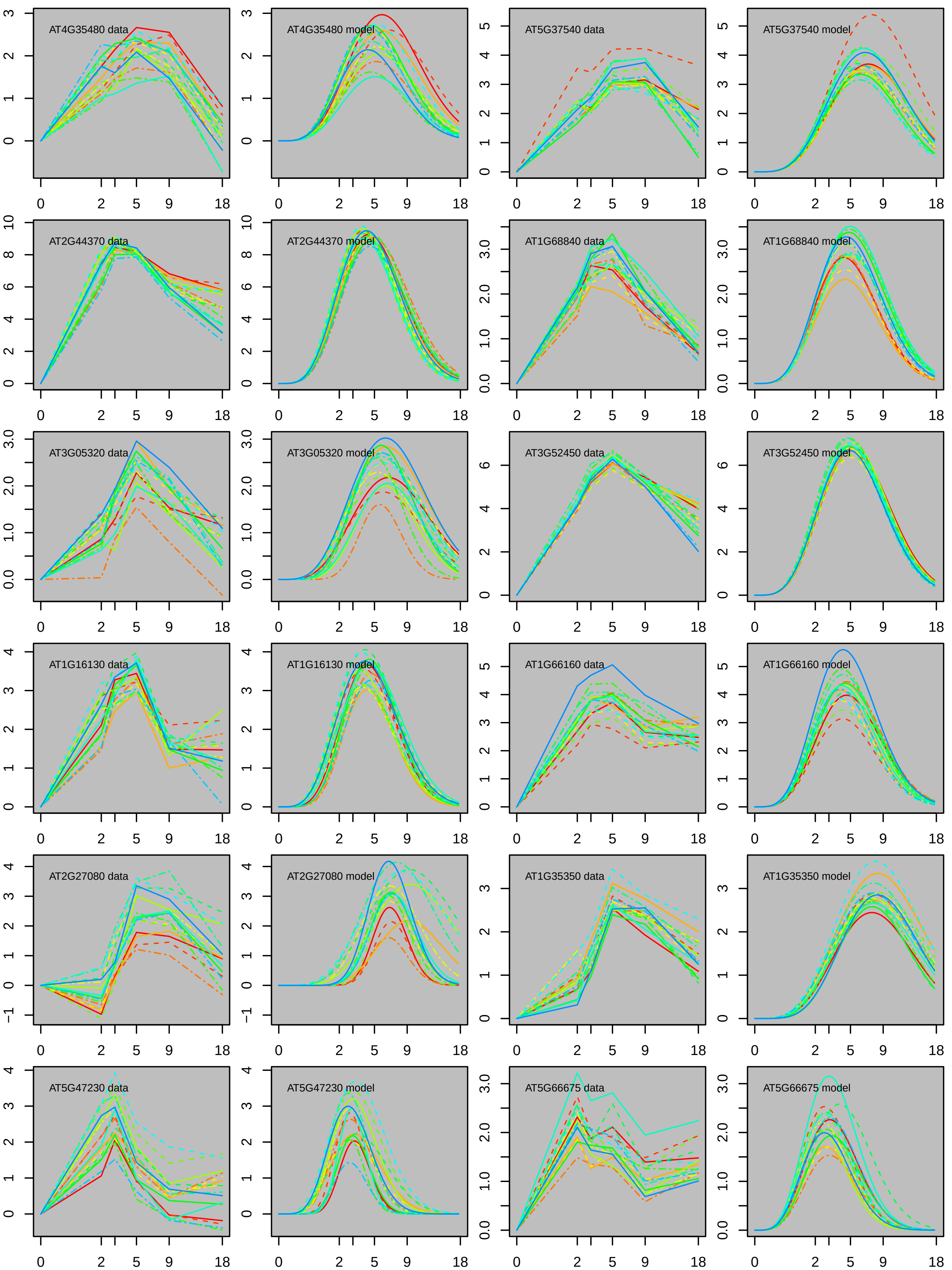
AT4G35480 data
AT4G35480 model
AT5G37540 data
AT5G37540 model
AT2G44370 data
AT2G44370 model
AT1G68840 data
AT1G68840 model
AT3G05320 data
AT3G05320 model
AT3G52450 data
AT3G52450 model
AT1G16130 data
AT1G16130 model
AT1G66160 data
AT1G66160 model
AT2G27080 data
AT2G27080 model
AT1G35350 data
AT1G35350 model
AT5G47230 data
AT5G47230 model
AT5G66675 data
AT5G66675 model
0
2
5
9
18

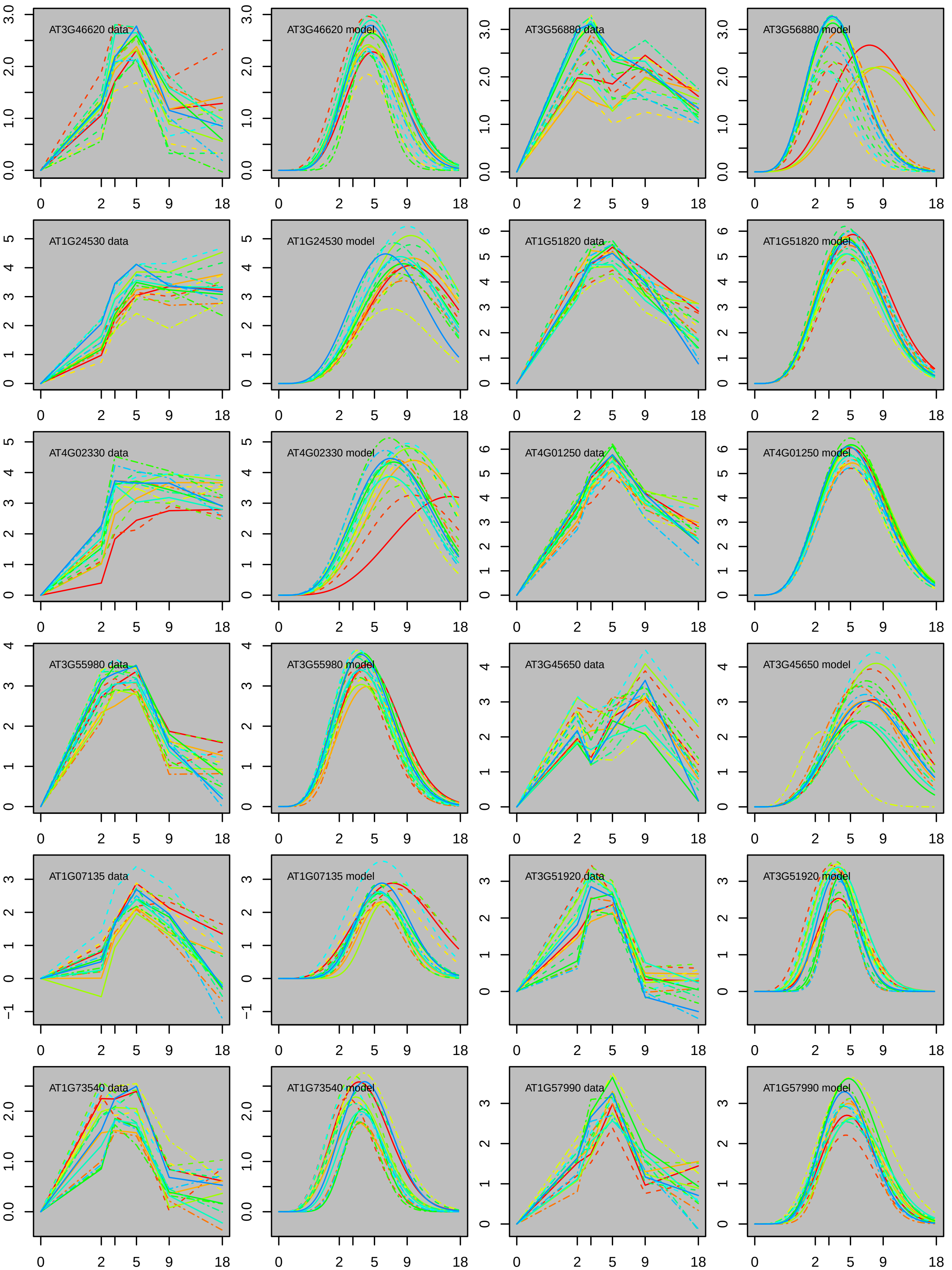
AT3G46620 data
AT3G46620 model
AT3G56880 data
AT3G56880 model
AT1G24530 data
AT1G24530 model
AT1G51820 data
AT1G51820 model
AT4G02330 data
AT4G02330 model
AT4G01250 data
AT4G01250 model
AT3G55980 data
AT3G55980 model
AT3G45650 data
AT3G45650 model
AT1G07135 data
AT1G07135 model
AT3G51920 data
AT3G51920 model
AT1G73540 data
AT1G73540 model
AT1G57990 data
AT1G57990 model
0
2
5
9
18

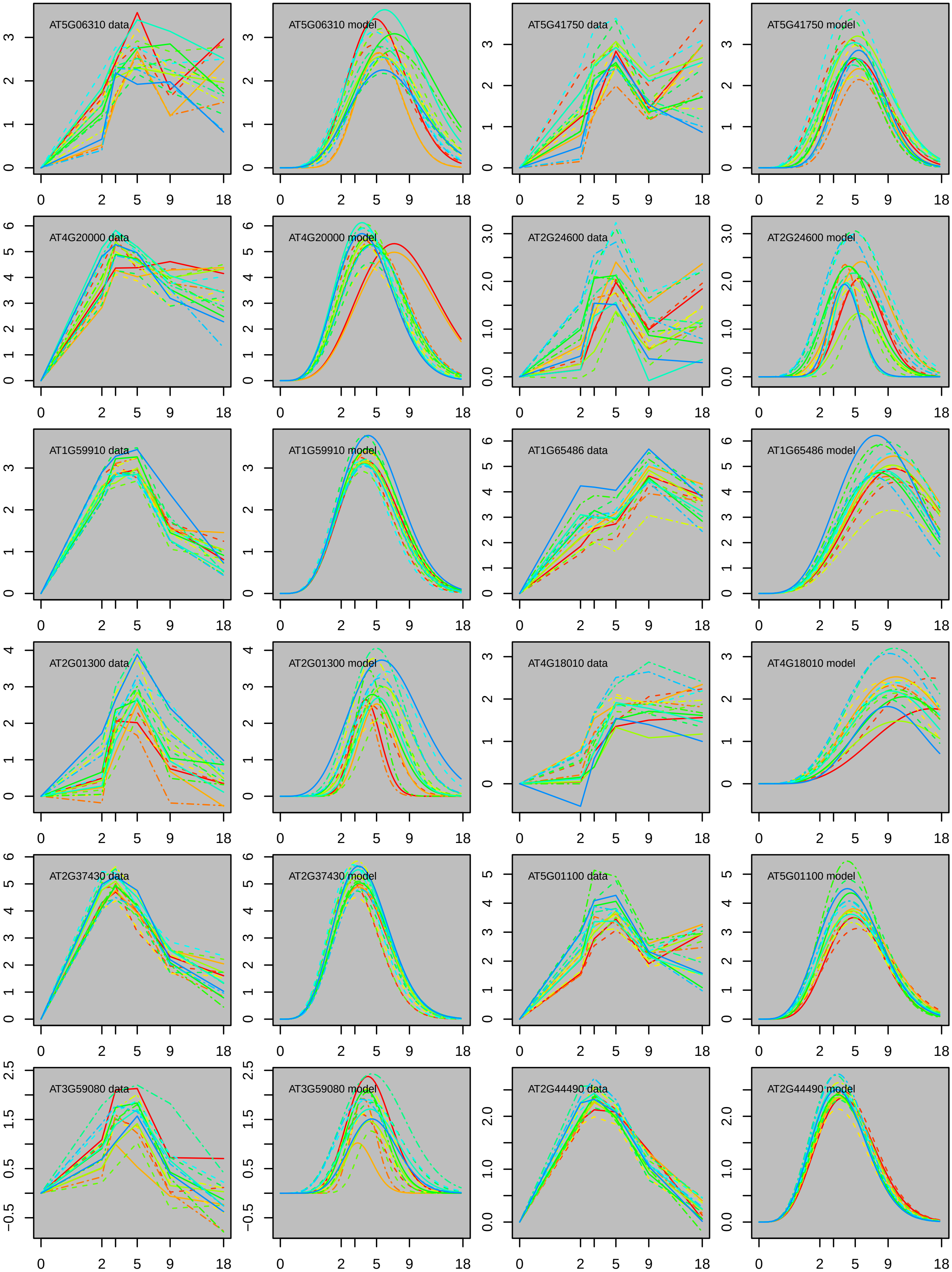
AT5G06310 data
AT5G06310 model
AT5G41750 data
AT5G41750 model
AT4G20000 data
AT4G20000 model
AT2G24600 data
AT2G24600 model
AT1G59910 data
AT1G59910 model
AT1G65486 data
AT1G65486 model
AT2G01300 data
AT2G01300 model
AT4G18010 data
AT4G18010 model
AT2G37430 data
AT2G37430 model
AT5G01100 data
AT5G01100 model
AT3G59080 data
AT3G59080 model
AT2G44490 data
AT2G44490 model
0 2 5 9 18

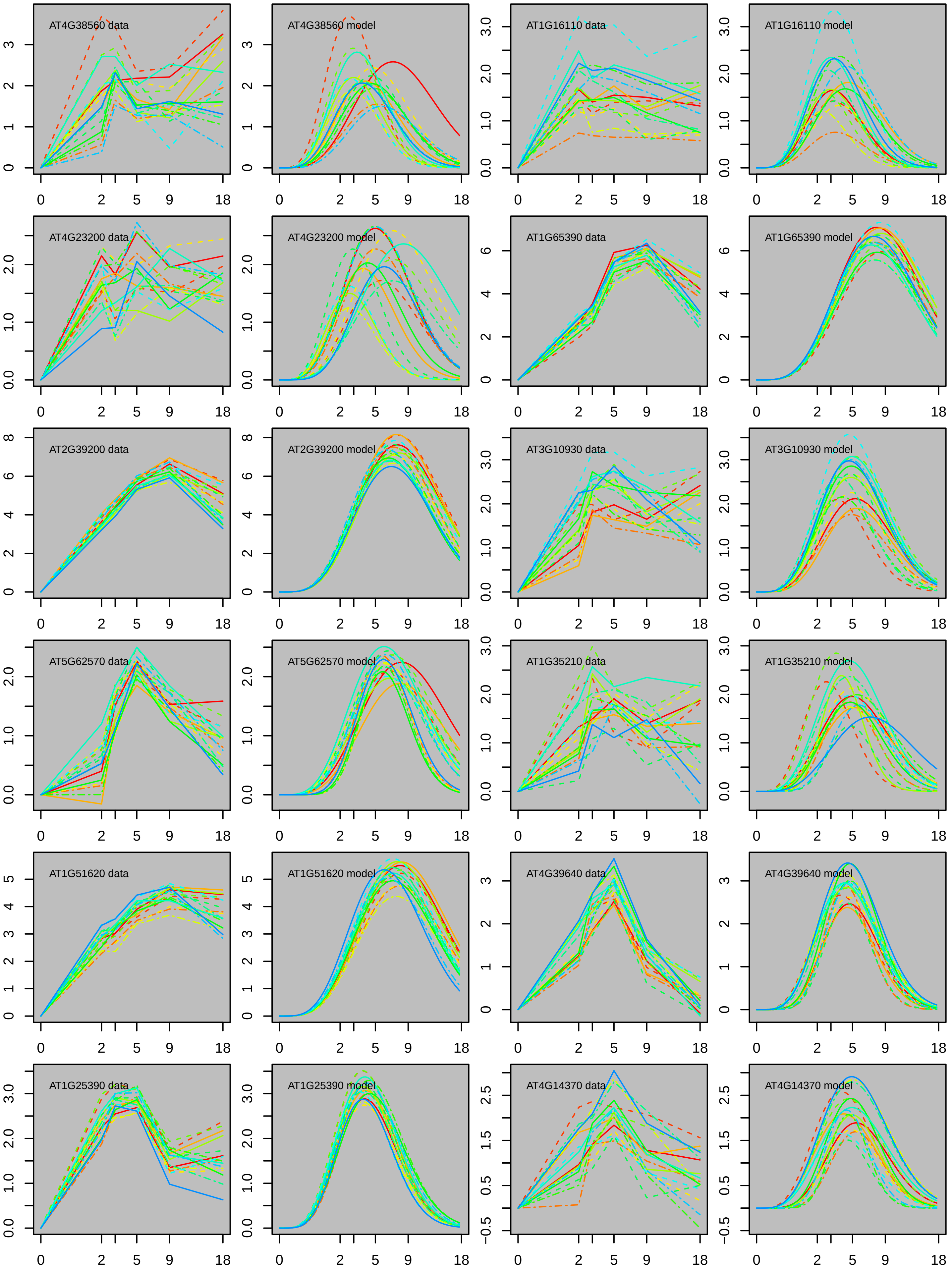
AT4G38560 data
AT4G38560 model
AT1G16110 data
AT1G16110 model
AT4G23200 data
AT4G23200 model
AT1G65390 data
AT1G65390 model
AT2G39200 data
AT2G39200 model
AT3G10930 data
AT3G10930 model
AT5G62570 data
AT5G62570 model
AT1G35210 data
AT1G35210 model
AT1G51620 data
AT1G51620 model
AT4G39640 data
AT4G39640 model
AT1G25390 data
AT1G25390 model
AT4G14370 data
AT4G14370 model
0 2 5 9 18

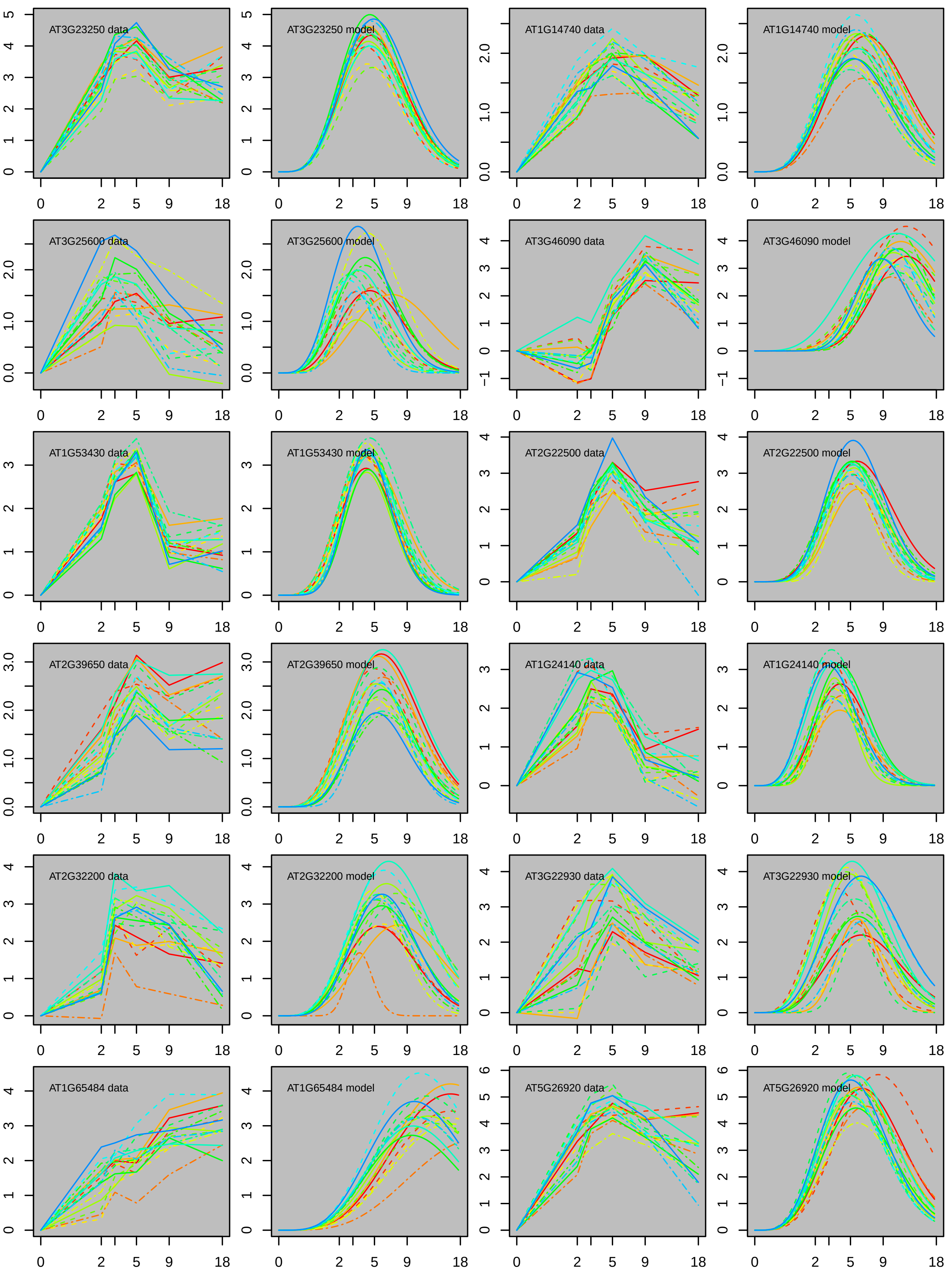

AT3G23250 data
AT3G23250 model
AT1G14740 data
AT1G14740 model
AT3G25600 data
AT3G25600 model
AT3G46090 data
AT3G46090 model
AT1G53430 data
AT1G53430 model
AT2G22500 data
AT2G22500 model
AT2G39650 data
AT2G39650 model
AT1G24140 data
AT1G24140 model
AT2G32200 data
AT2G32200 model
AT3G22930 data
AT3G22930 model
AT1G65484 data
AT1G65484 model
AT5G26920 data
AT5G26920 model

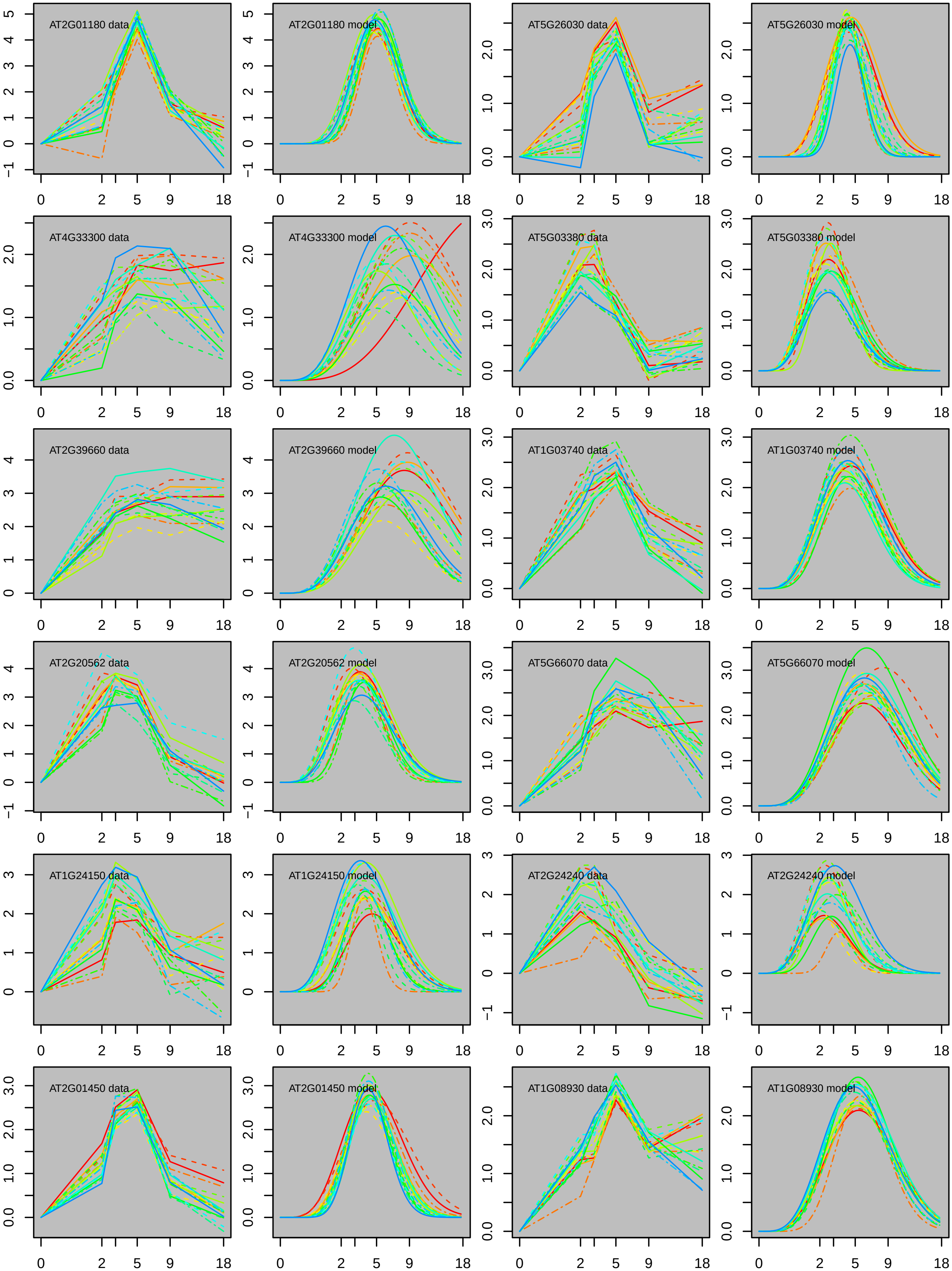
AT2G01180 data
AT2G01180 model
AT5G26030 data
AT5G26030 model
AT4G33300 data
AT4G33300 model
AT5G03380 data
AT5G03380 model
AT2G39660 data
AT2G39660 model
AT1G03740 data
AT1G03740 model
AT2G20562 data
AT2G20562 model
AT5G66070 data
AT5G66070 model
AT1G24150 data
AT1G24150 model
AT2G24240 data
AT2G24240 model
AT2G01450 data
AT2G01450 model
AT1G08930 data
AT1G08930 model
0
2
5
9
18

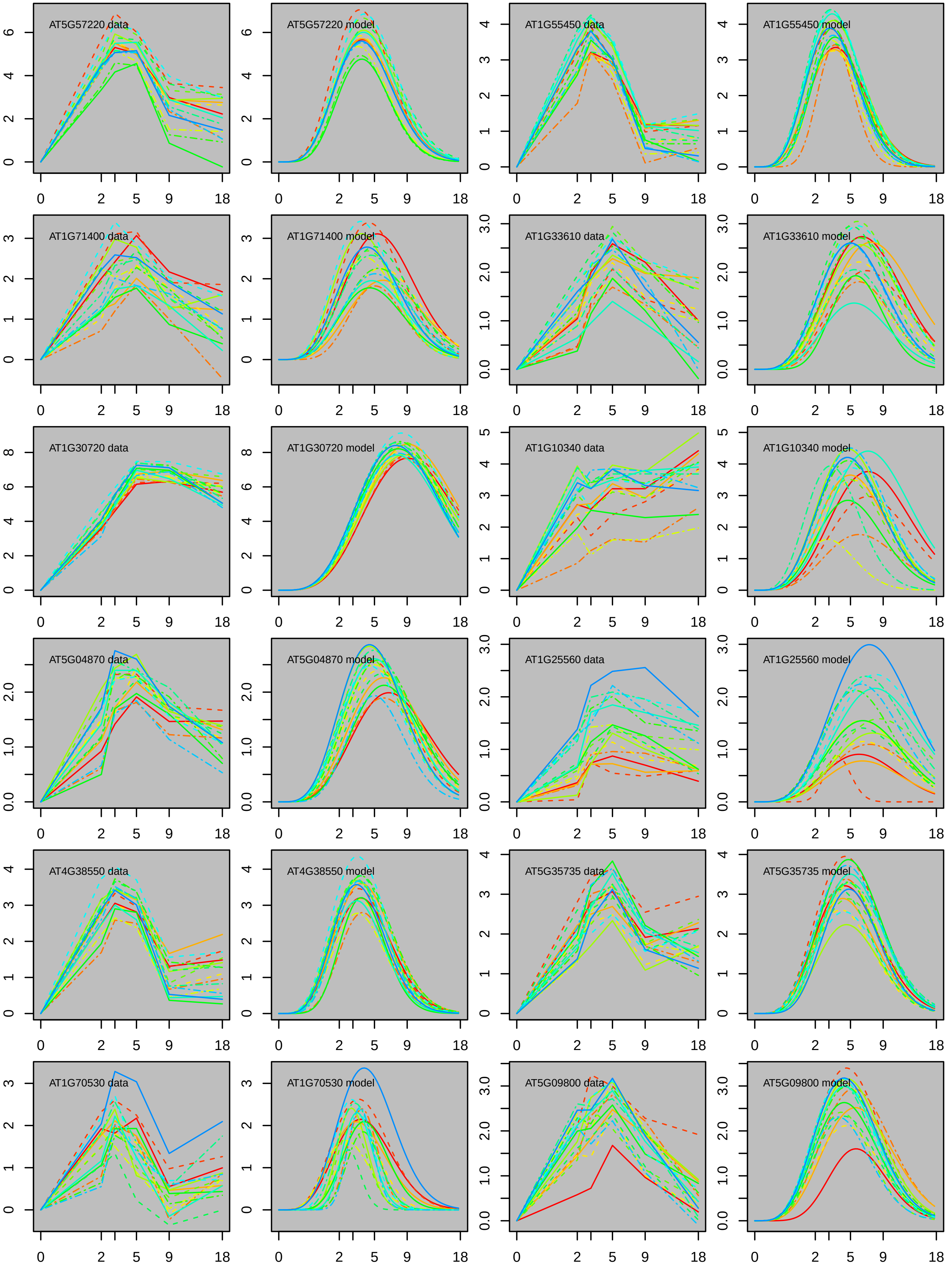
AT5G57220 data
AT5G57220 model
AT1G55450 data
AT1G55450 model
AT1G71400 data
AT1G71400 model
AT1G33610 data
AT1G33610 model
AT1G30720 data
AT1G30720 model
AT1G10340 data
AT1G10340 model
AT5G04870 data
AT5G04870 model
AT1G25560 data
AT1G25560 model
AT4G38550 data
AT4G38550 model
AT5G35735 data
AT5G35735 model
AT1G70530 data
AT1G70530 model
AT5G09800 data
AT5G09800 model

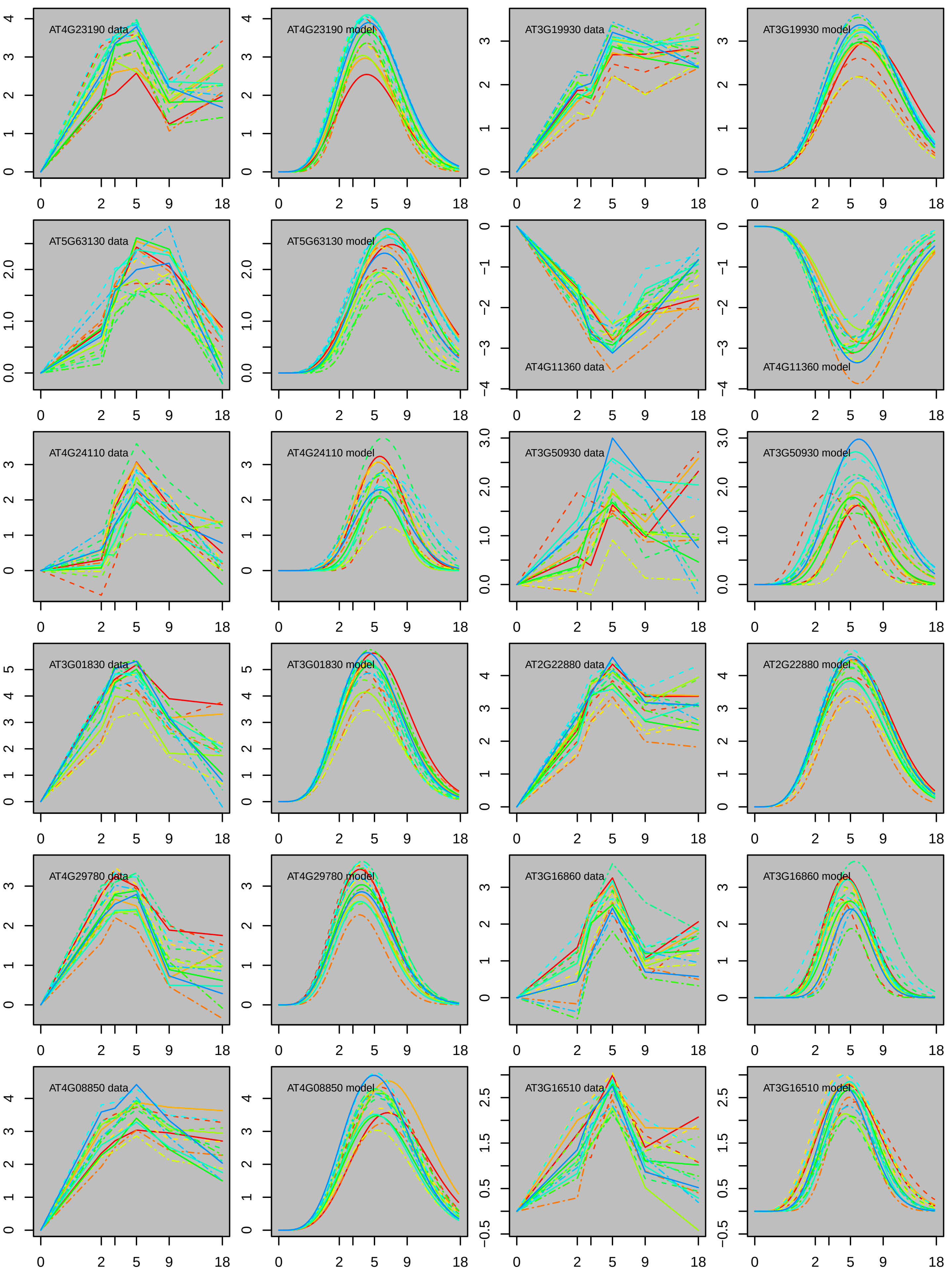

AT4G23190 data
AT4G23190 model
AT3G19930 data
AT3G19930 model
AT5G63130 data
AT5G63130 model
AT4G11360 data
AT4G11360 model
AT4G24110 data
AT4G24110 model
AT3G50930 data
AT3G50930 model
AT3G01830 data
AT3G01830 model
AT2G22880 data
AT2G22880 model
AT4G29780 data
AT4G29780 model
AT3G16860 data
AT3G16860 model
AT4G08850 data
AT4G08850 model
AT3G16510 data
AT3G16510 model

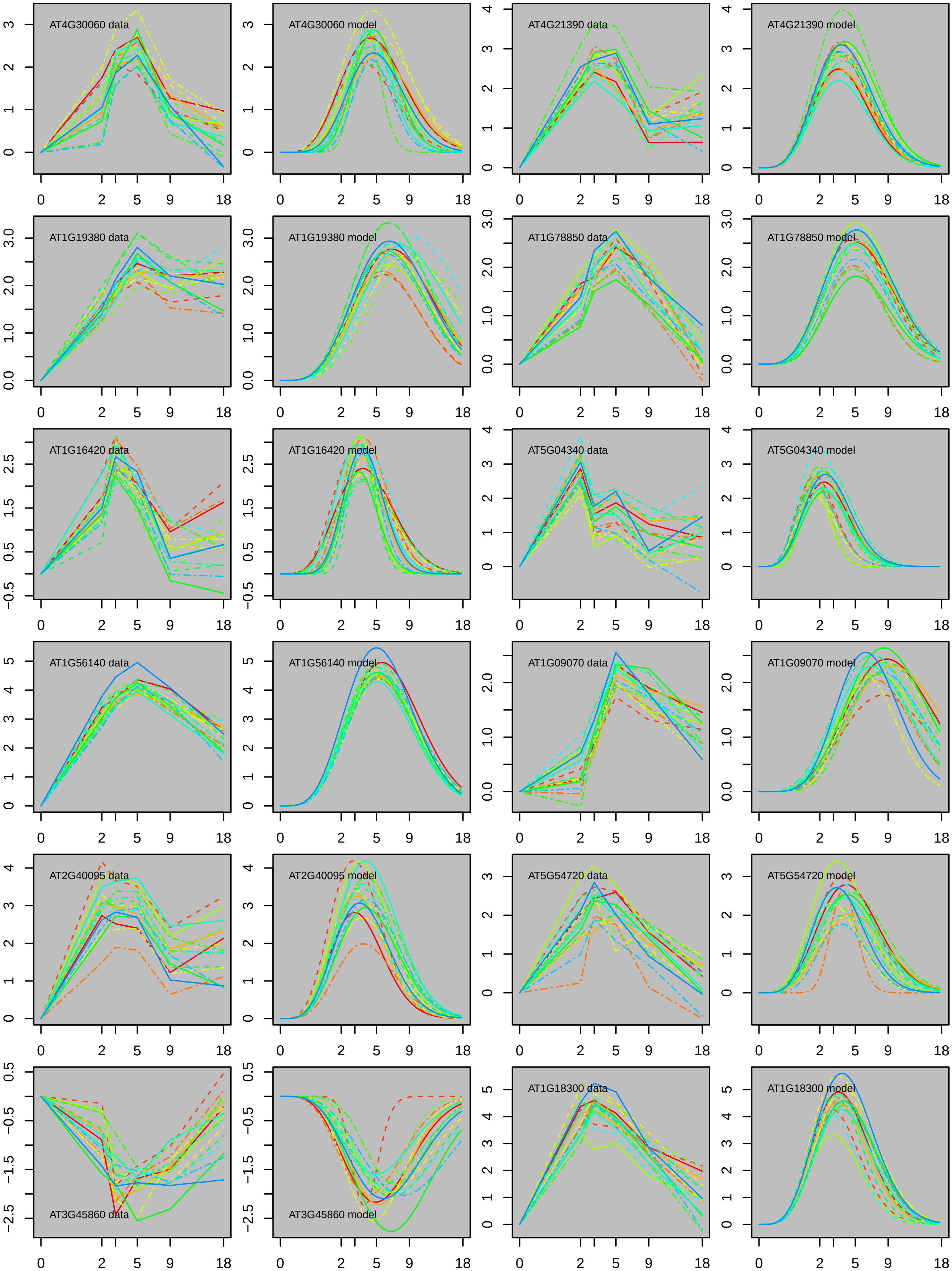
AT4G30060 data
AT4G30060 model
AT4G21390 data
AT4G21390 model
AT1G19380 data
AT1G19380 model
AT1G78850 data
AT1G78850 model
AT1G16420 data
AT1G16420 model
AT5G04340 data
AT5G04340 model
AT1G56140 data
AT1G56140 model
AT1G09070 data
AT1G09070 model
AT2G40095 data
AT2G40095 model
AT5G54720 data
AT5G54720 model
AT3G45860 data
AT3G45860 model
AT1G18300 data
AT1G18300 model
0
2
5
9
18

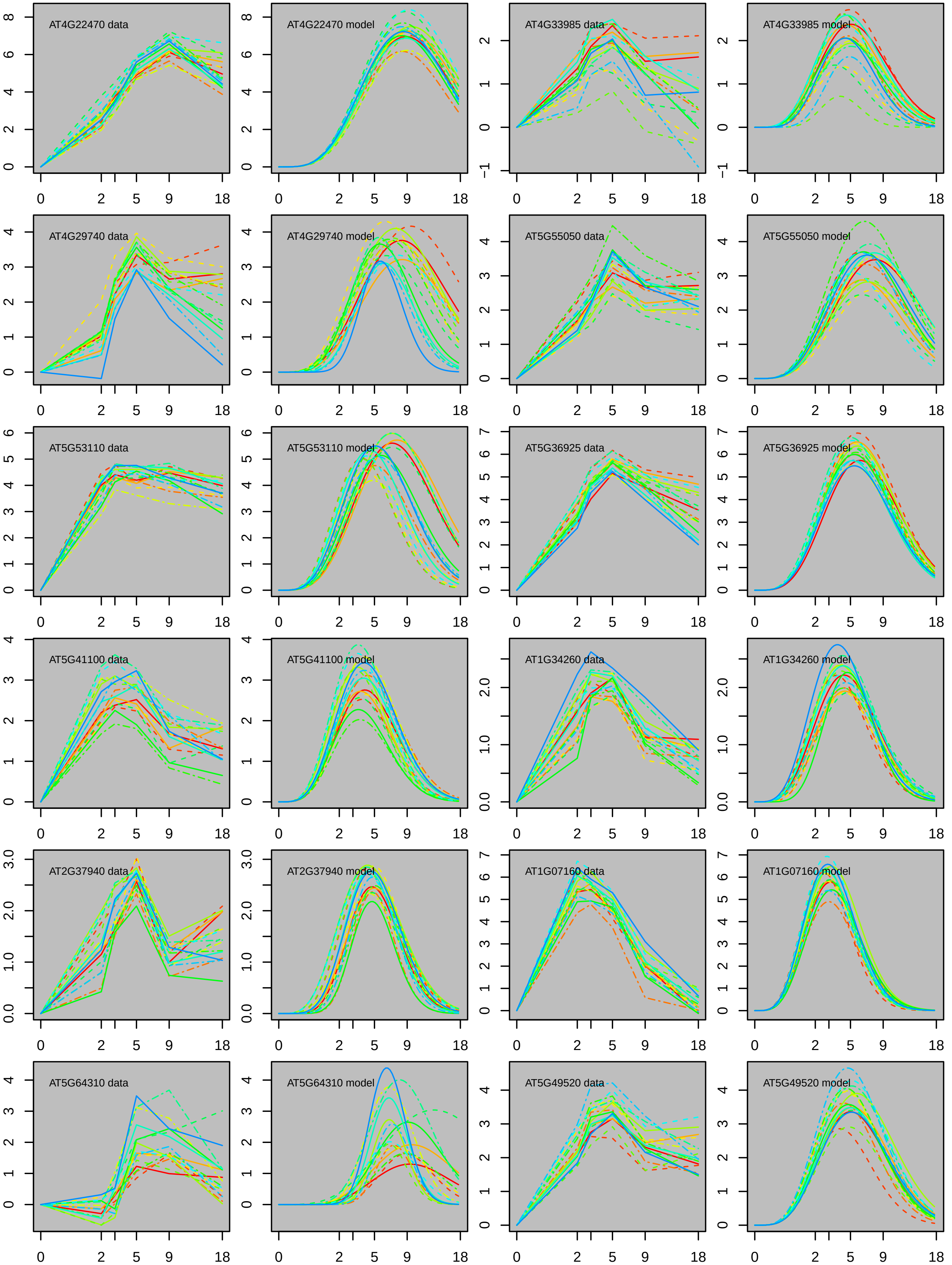
AT4G22470 data
AT4G22470 model
AT4G33985 data
AT4G33985 model
AT4G29740 data
AT4G29740 model
AT5G55050 data
AT5G55050 model
AT5G53110 data
AT5G53110 model
AT5G36925 data
AT5G36925 model
AT5G41100 data
AT5G41100 model
AT1G34260 data
AT1G34260 model
AT2G37940 data
AT2G37940 model
AT1G07160 data
AT1G07160 model
AT5G64310 data
AT5G64310 model
AT5G49520 data
AT5G49520 model
0 2 5 9 18

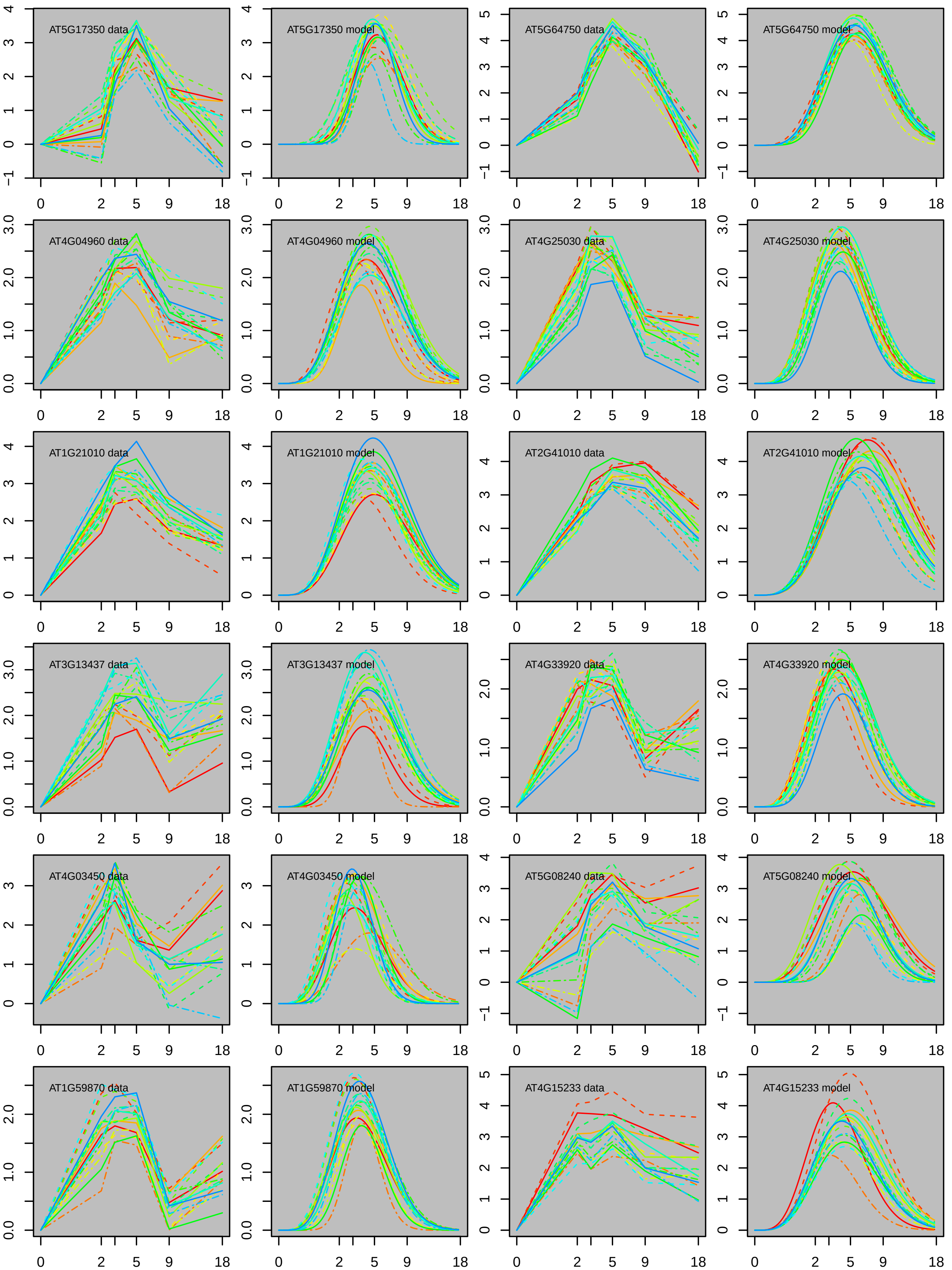
AT5G17350 data
AT5G17350 model
AT5G64750 data
AT5G64750 model
AT4G04960 data
AT4G04960 model
AT4G25030 data
AT4G25030 model
AT1G21010 data
AT1G21010 model
AT2G41010 data
AT2G41010 model
AT3G13437 data
AT3G13437 model
AT4G33920 data
AT4G33920 model
AT4G03450 data
AT4G03450 model
AT5G08240 data
AT5G08240 model
AT1G59870 data
AT1G59870 model
AT4G15233 data
AT4G15233 model
0 2 5 9 18

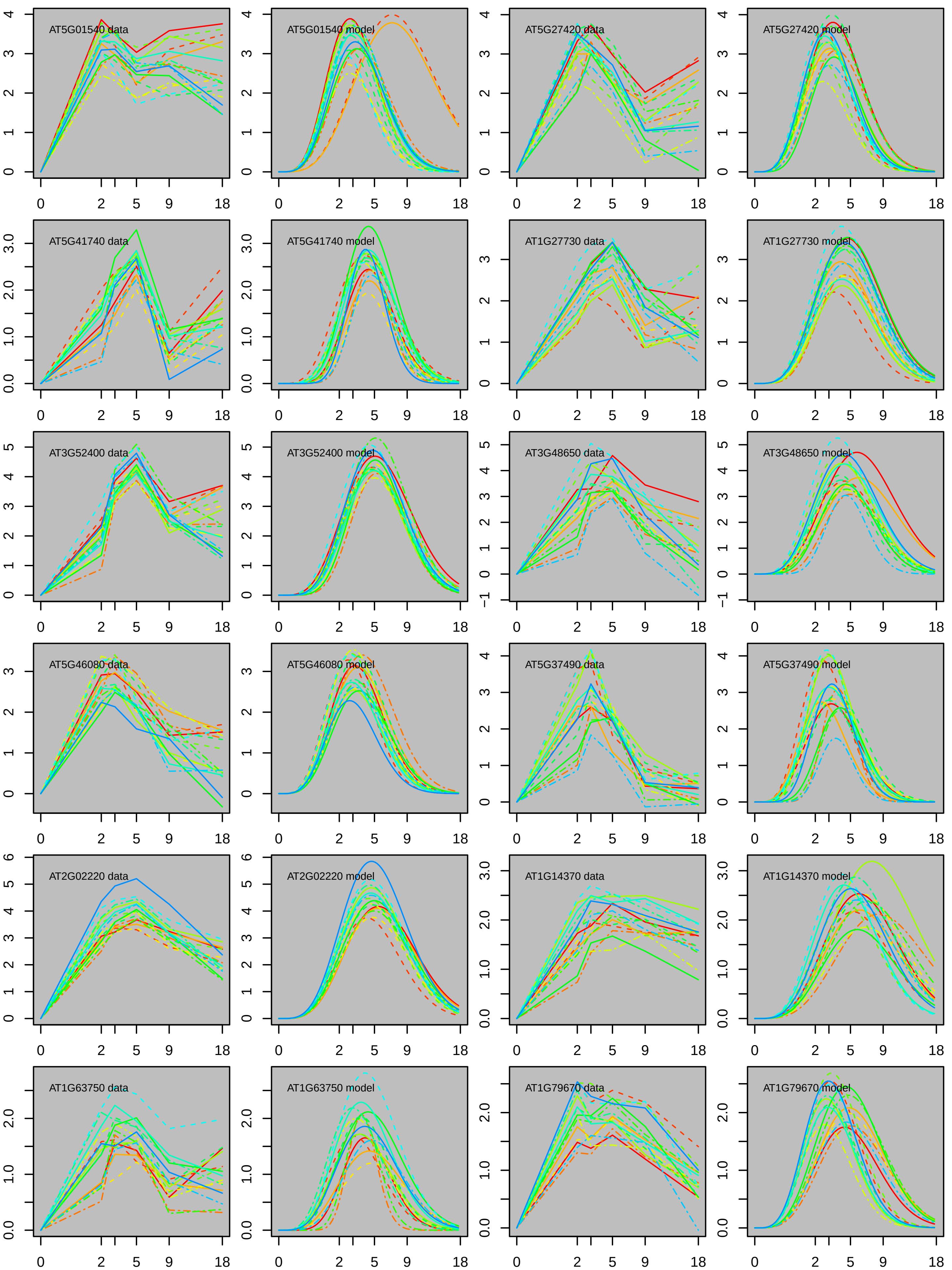

AT5G01540 data
AT5G01540 model
AT5G27420 data
AT5G27420 model
AT5G41740 data
AT5G41740 model
AT1G27730 data
AT1G27730 model
AT3G52400 data
AT3G52400 model
AT3G48650 data
AT3G48650 model
AT5G46080 data
AT5G46080 model
AT5G37490 data
AT5G37490 model
AT2G02220 data
AT2G02220 model
AT1G14370 data
AT1G14370 model
AT1G63750 data
AT1G63750 model
AT1G79670 data
AT1G79670 model

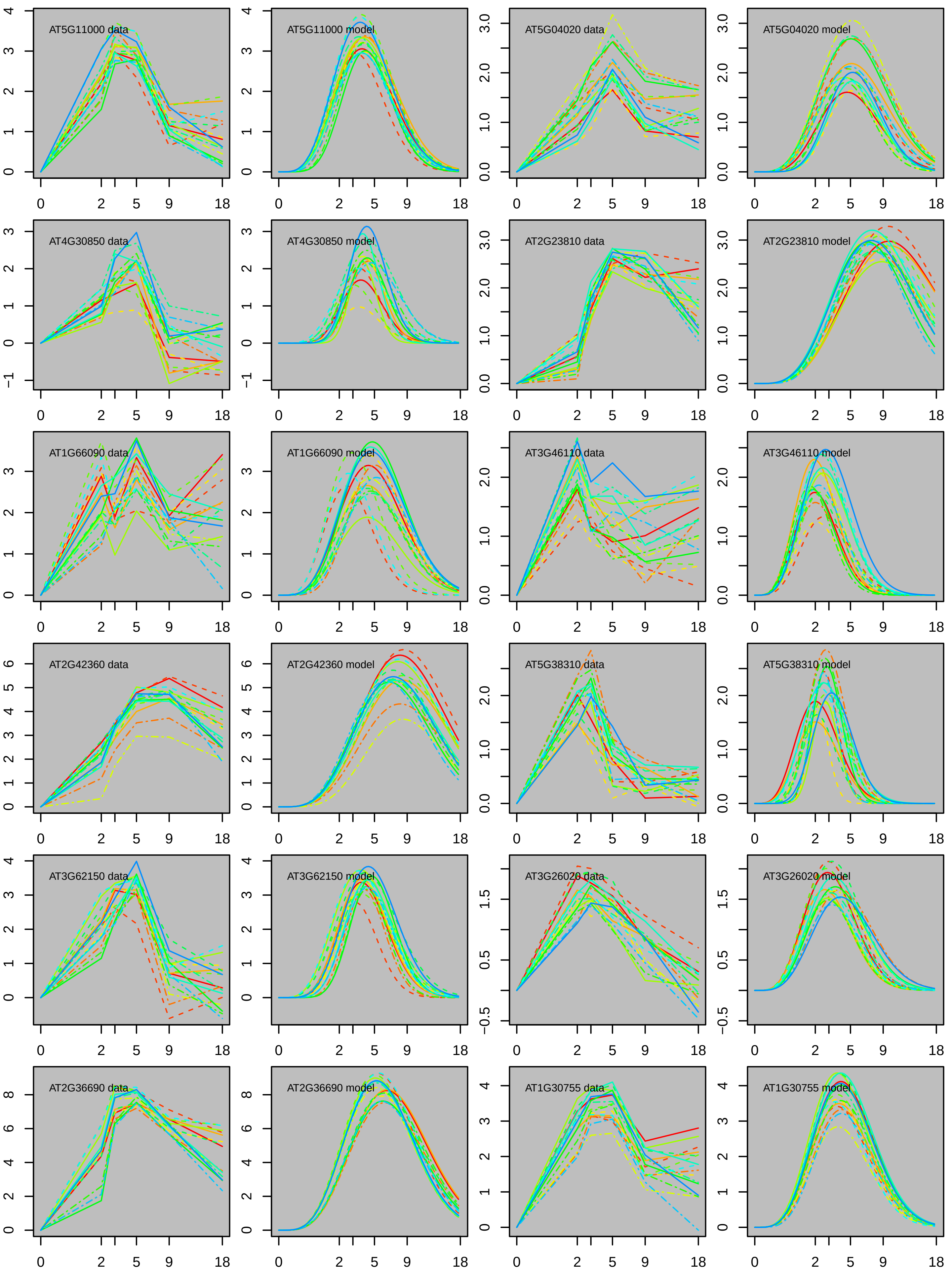
AT5G11000 data
AT5G11000 model
AT5G04020 data
AT5G04020 model
AT4G30850 data
AT4G30850 model
AT2G23810 data
AT2G23810 model
AT1G66090 data
AT1G66090 model
AT3G46110 data
AT3G46110 model
AT2G42360 data
AT2G42360 model
AT5G38310 data
AT5G38310 model
AT3G62150 data
AT3G62150 model
AT3G26020 data
AT3G26020 model
AT2G36690 data
AT2G36690 model
AT1G30755 data
AT1G30755 model

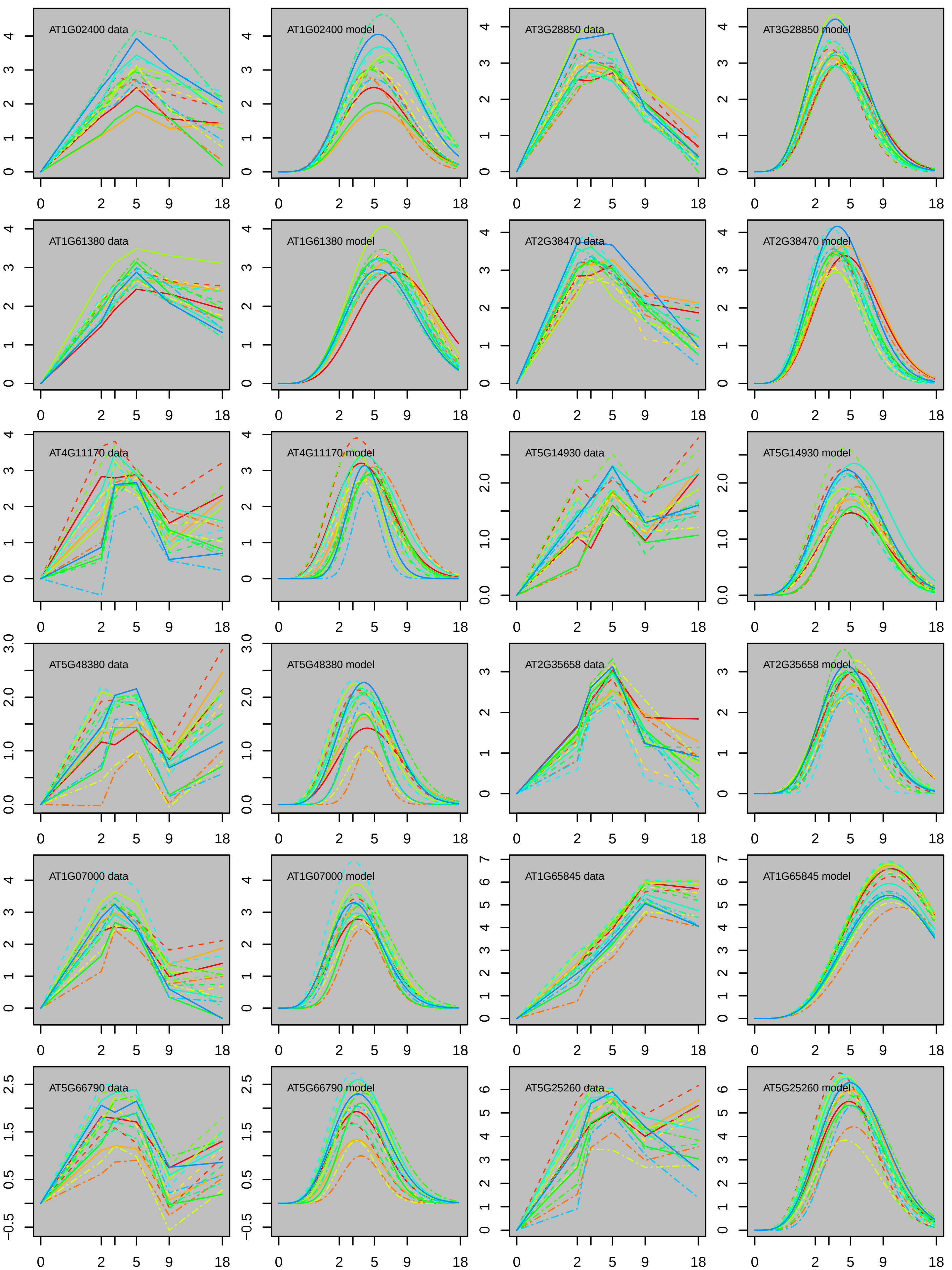
AT1G02400 data
AT1G02400 model
AT3G28850 data
AT3G28850 model
AT1G61380 data
AT1G61380 model
AT2G38470 data
AT2G38470 model
AT4G11170 data
AT4G11170 model
AT5G14930 data
AT5G14930 model
AT5G48380 data
AT5G48380 model
AT2G35658 data
AT2G35658 model
AT1G07000 data
AT1G07000 model
AT1G65845 data
AT1G65845 model
AT5G66790 data
AT5G66790 model
AT5G25260 data
AT5G25260 model
0 2 5 9 18

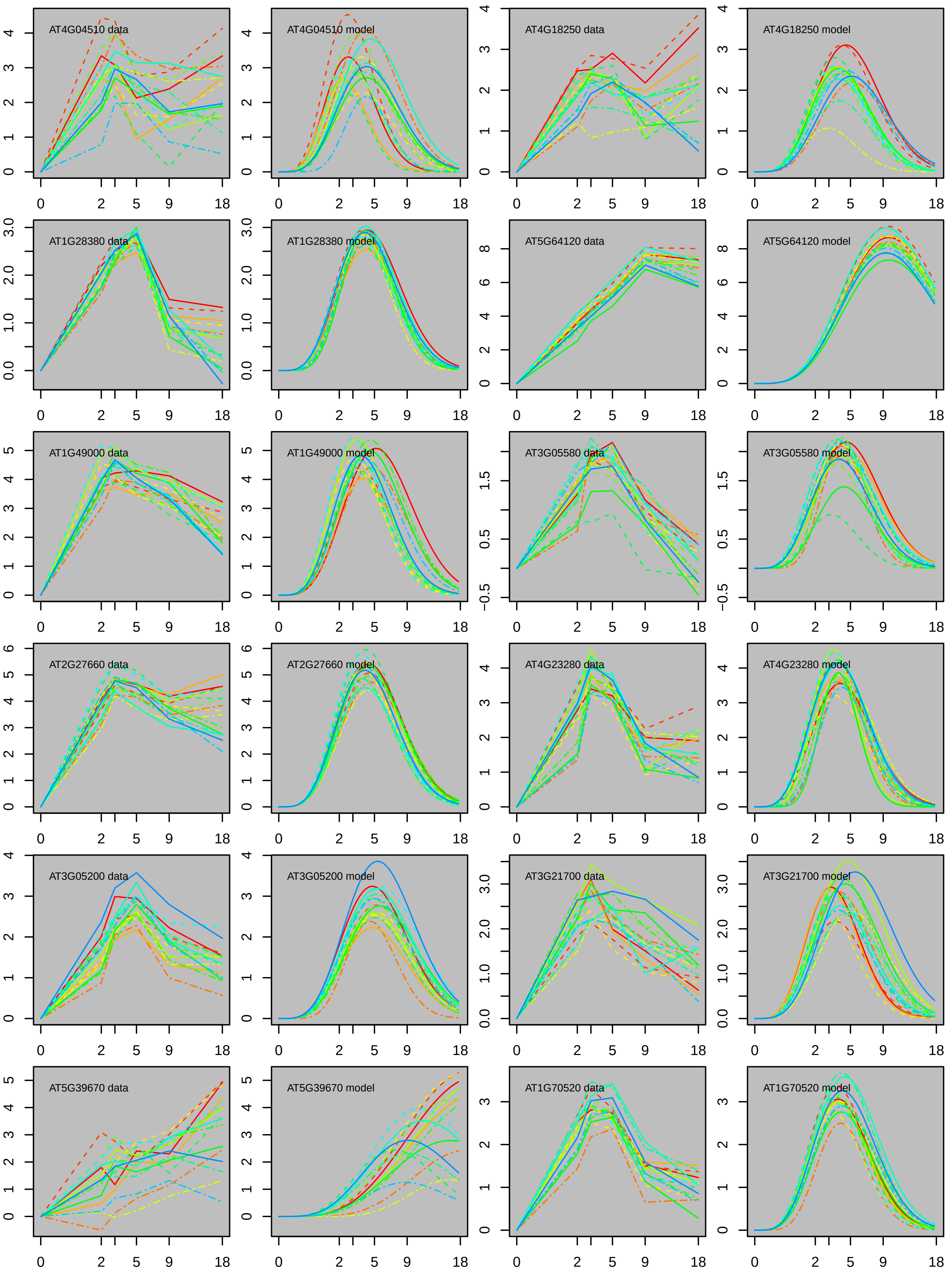

AT4G04510 data
AT4G04510 model
AT4G18250 data
AT4G18250 model
AT1G28380 data
AT1G28380 model
AT5G64120 data
AT5G64120 model
AT1G49000 data
AT1G49000 model
AT3G05580 data
AT3G05580 model
AT2G27660 data
AT2G27660 model
AT4G23280 data
AT4G23280 model
AT3G05200 data
AT3G05200 model
AT3G21700 data
AT3G21700 model
AT5G39670 data
AT5G39670 model
AT1G70520 data
AT1G70520 model

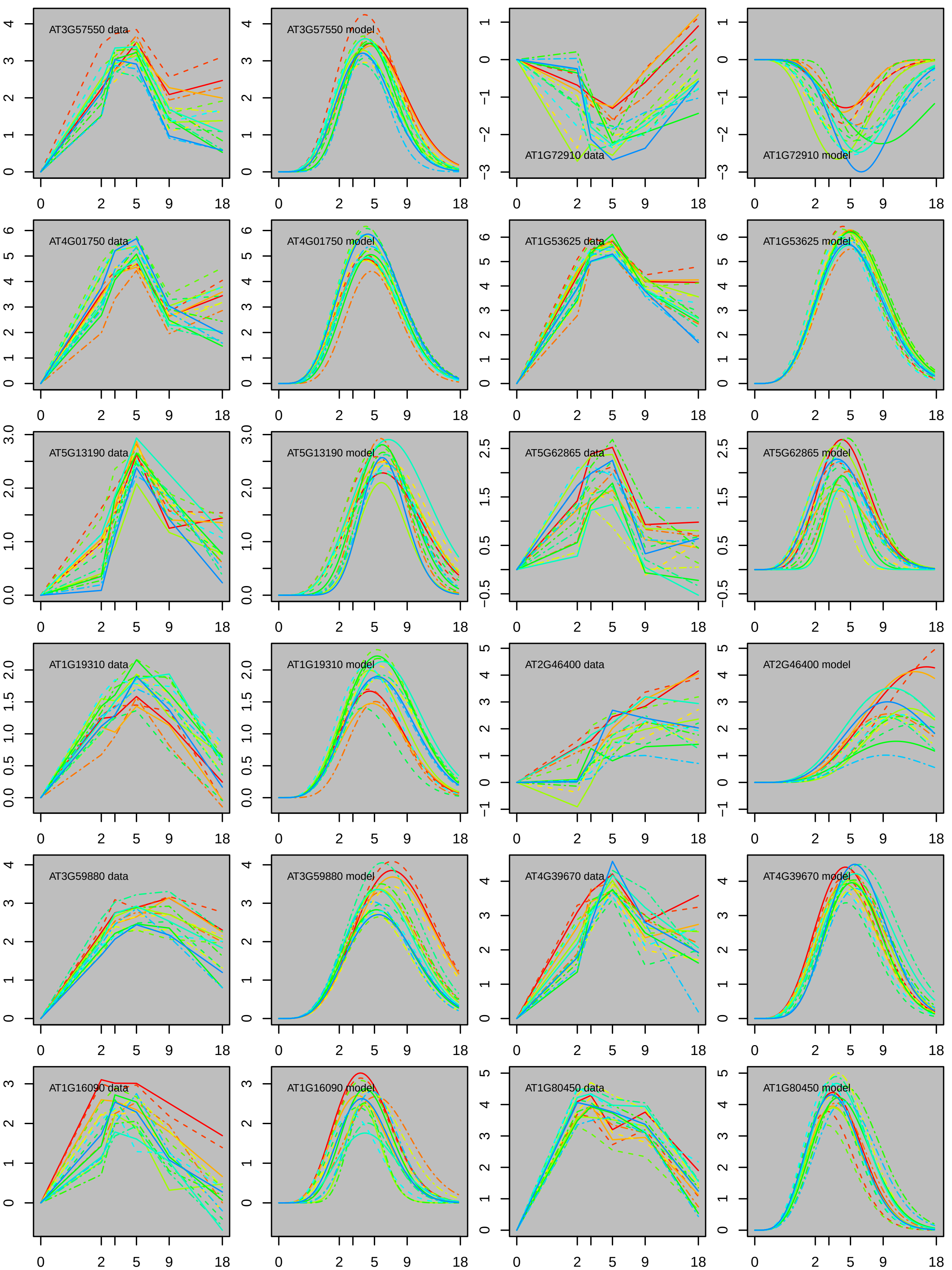
AT3G57550 data
AT3G57550 model
AT1G72910 data
AT1G72910 model
AT4G01750 data
AT4G01750 model
AT1G53625 data
AT1G53625 model
AT5G13190 data
AT5G13190 model
AT5G62865 data
AT5G62865 model
AT1G19310 data
AT1G19310 model
AT2G46400 data
AT2G46400 model
AT3G59880 data
AT3G59880 model
AT4G39670 data
AT4G39670 model
AT1G16090 data
AT1G16090 model
AT1G80450 data
AT1G80450 model

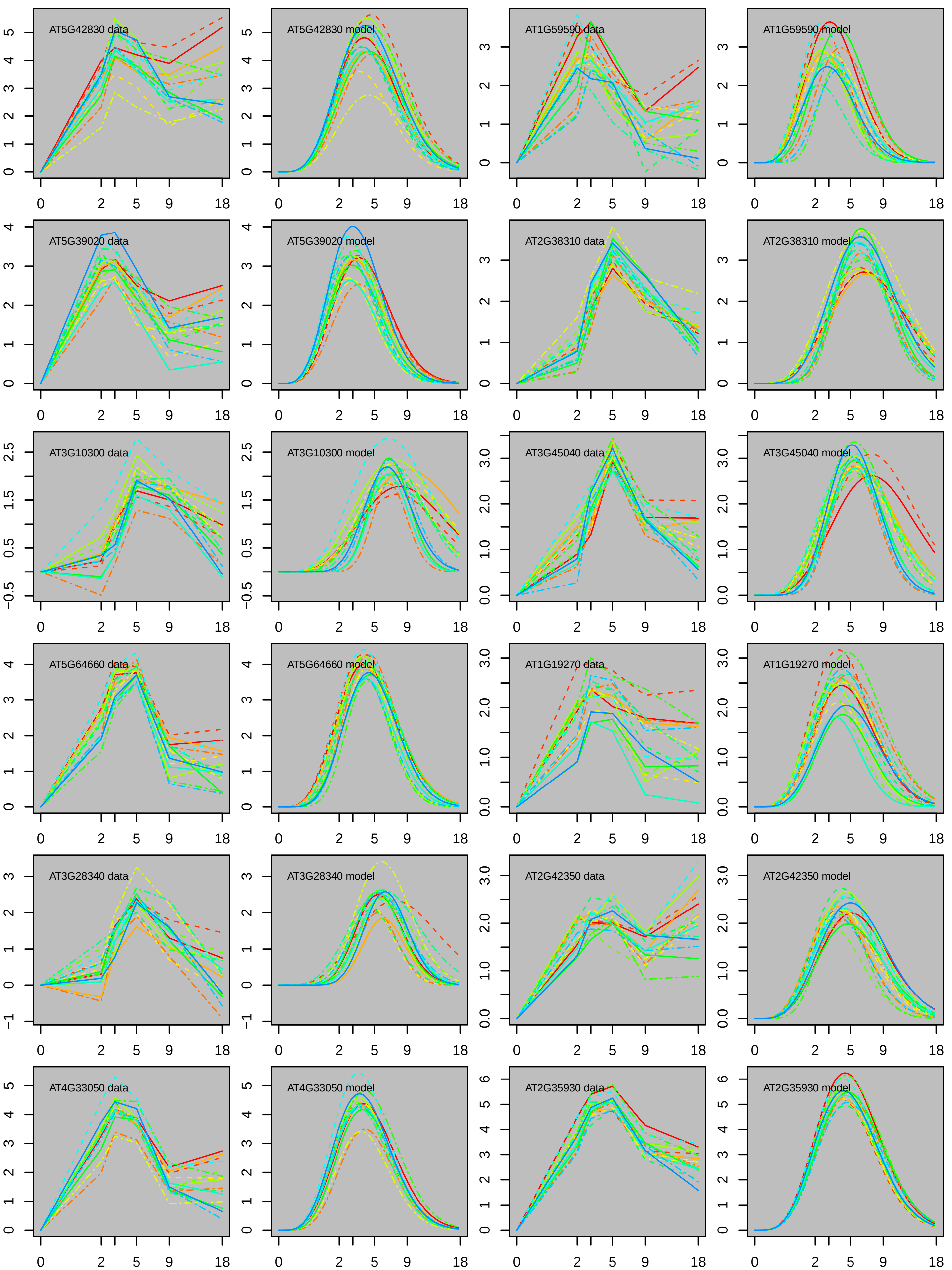

AT5G42830 data
AT5G42830 model
AT1G59590 data
AT1G59590 model
AT5G39020 data
AT5G39020 model
AT2G38310 data
AT2G38310 model
AT3G10300 data
AT3G10300 model
AT3G45040 data
AT3G45040 model
AT5G64660 data
AT5G64660 model
AT1G19270 data
AT1G19270 model
AT3G28340 data
AT3G28340 model
AT2G42350 data
AT2G42350 model
AT4G33050 data
AT4G33050 model
AT2G35930 data
AT2G35930 model

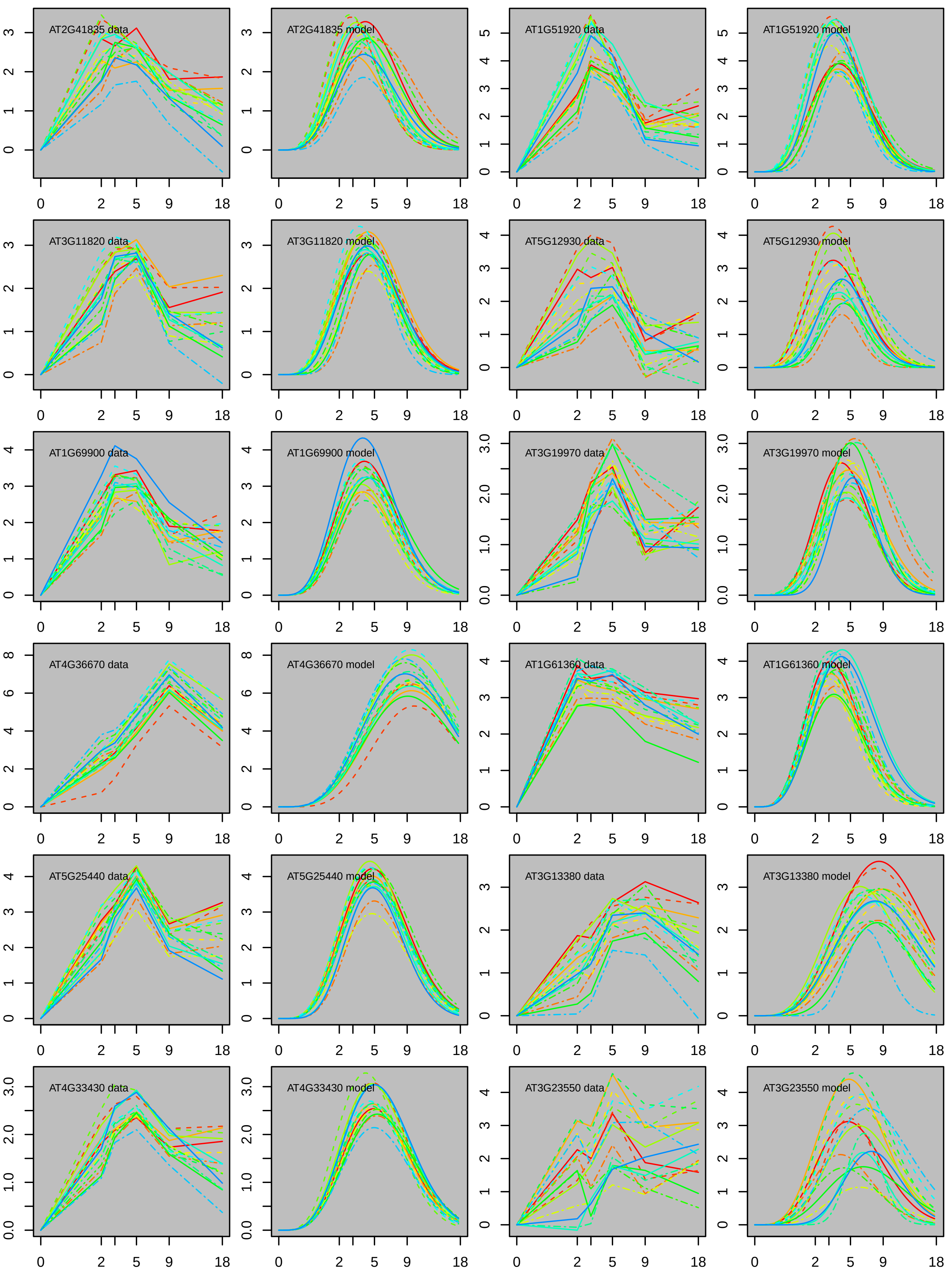

AT2G41835 data
AT2G41835 model
AT1G51920 data
AT1G51920 model
AT3G11820 data
AT3G11820 model
AT5G12930 data
AT5G12930 model
AT1G69900 data
AT1G69900 model
AT3G19970 data
AT3G19970 model
AT4G36670 data
AT4G36670 model
AT1G61360 data
AT1G61360 model
AT5G25440 data
AT5G25440 model
AT3G13380 data
AT3G13380 model
AT4G33430 data
AT4G33430 model
AT3G23550 data
AT3G23550 model
0 2 5 9 18

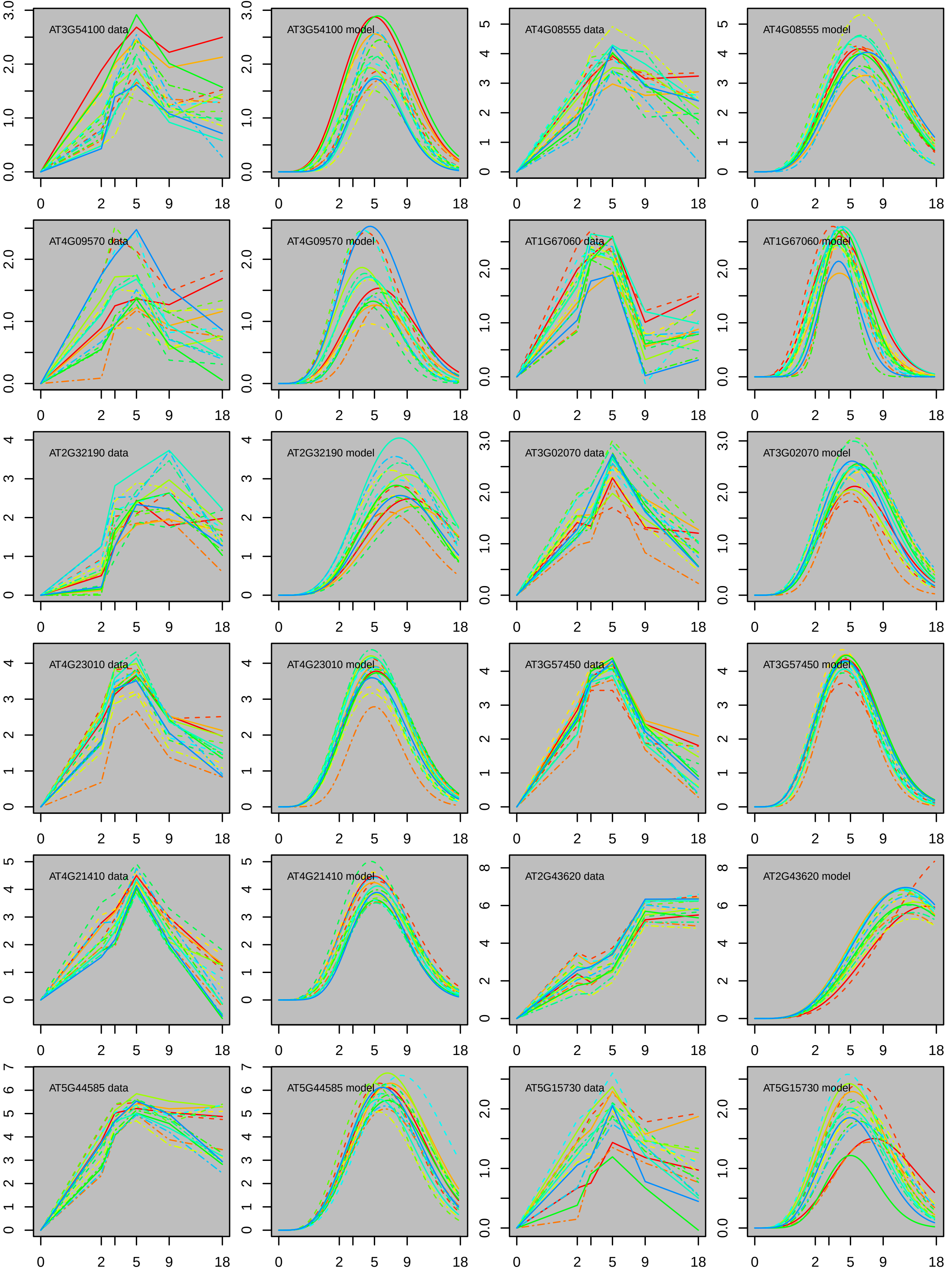

AT3G54100 data
AT3G54100 model
AT4G08555 data
AT4G08555 model
AT4G09570 data
AT4G09570 model
AT1G67060 data
AT1G67060 model
AT2G32190 data
AT2G32190 model
AT3G02070 data
AT3G02070 model
AT4G23010 data
AT4G23010 model
AT3G57450 data
AT3G57450 model
AT4G21410 data
AT4G21410 model
AT2G43620 data
AT2G43620 model
AT5G44585 data
AT5G44585 model
AT5G15730 data
AT5G15730 model

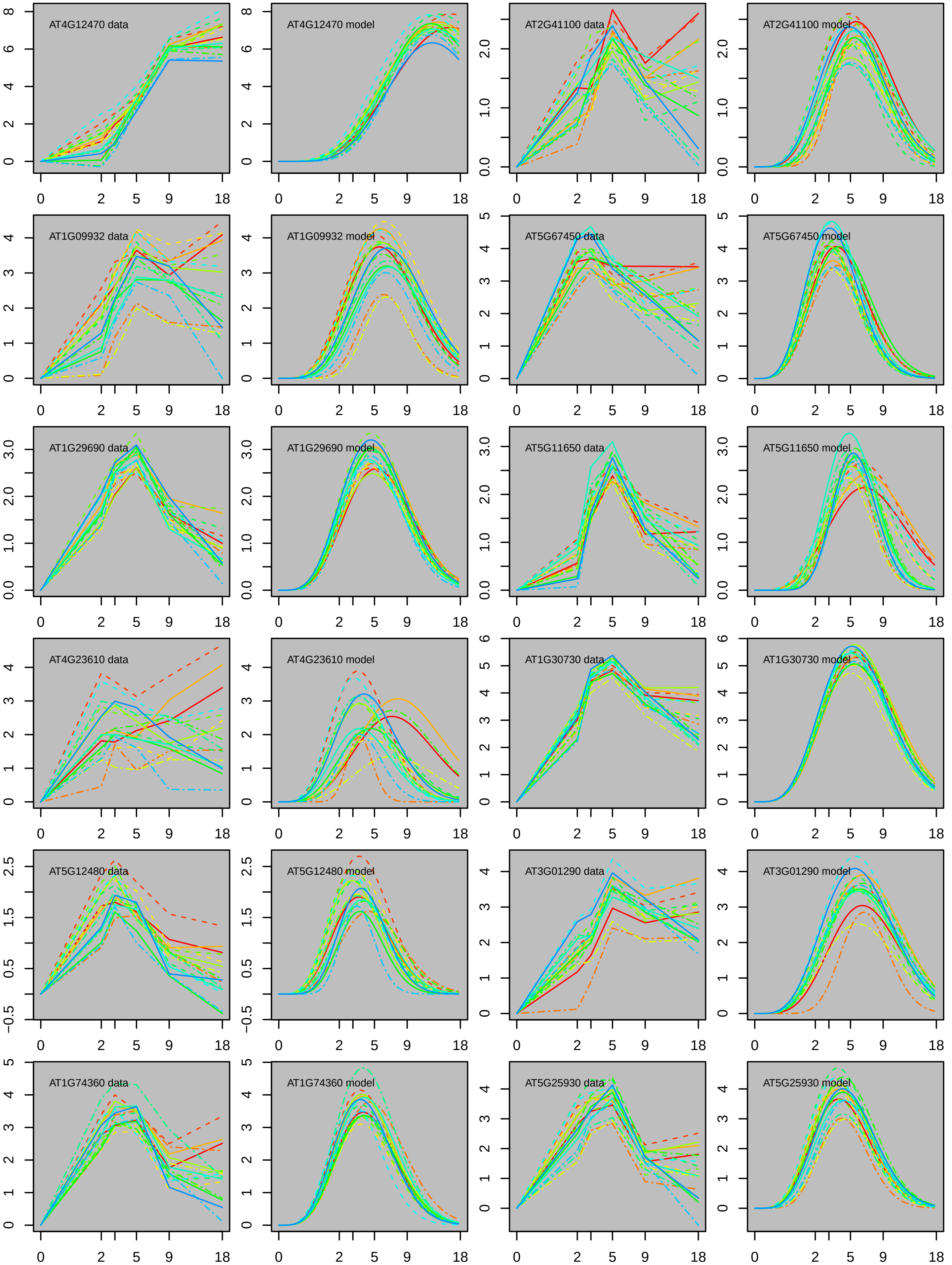
AT4G12470 data
AT4G12470 model
AT2G41100 data
AT2G41100 model
AT1G09932 data
AT1G09932 model
AT5G67450 data
AT5G67450 model
AT1G29690 data
AT1G29690 model
AT5G11650 data
AT5G11650 model
AT4G23610 data
AT4G23610 model
AT1G30730 data
AT1G30730 model
AT5G12480 data
AT5G12480 model
AT3G01290 data
AT3G01290 model
AT1G74360 data
AT1G74360 model
AT5G25930 data
AT5G25930 model

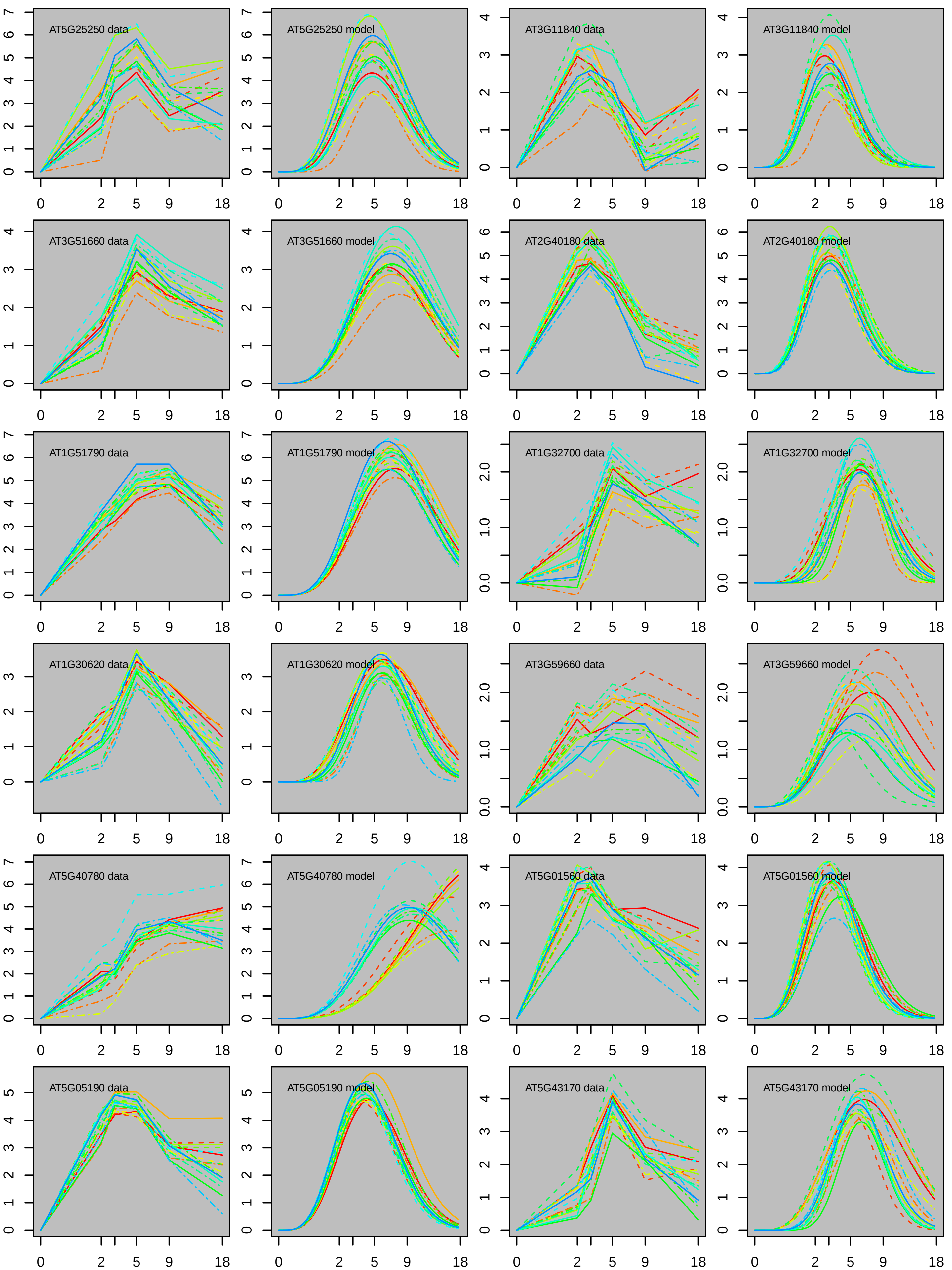

AT5G25250 data
AT5G25250 model
AT3G11840 data
AT3G11840 model
AT3G51660 data
AT3G51660 model
AT2G40180 data
AT2G40180 model
AT1G51790 data
AT1G51790 model
AT1G32700 data
AT1G32700 model
AT1G30620 data
AT1G30620 model
AT3G59660 data
AT3G59660 model
AT5G40780 data
AT5G40780 model
AT5G01560 data
AT5G01560 model
AT5G05190 data
AT5G05190 model
AT5G43170 data
AT5G43170 model

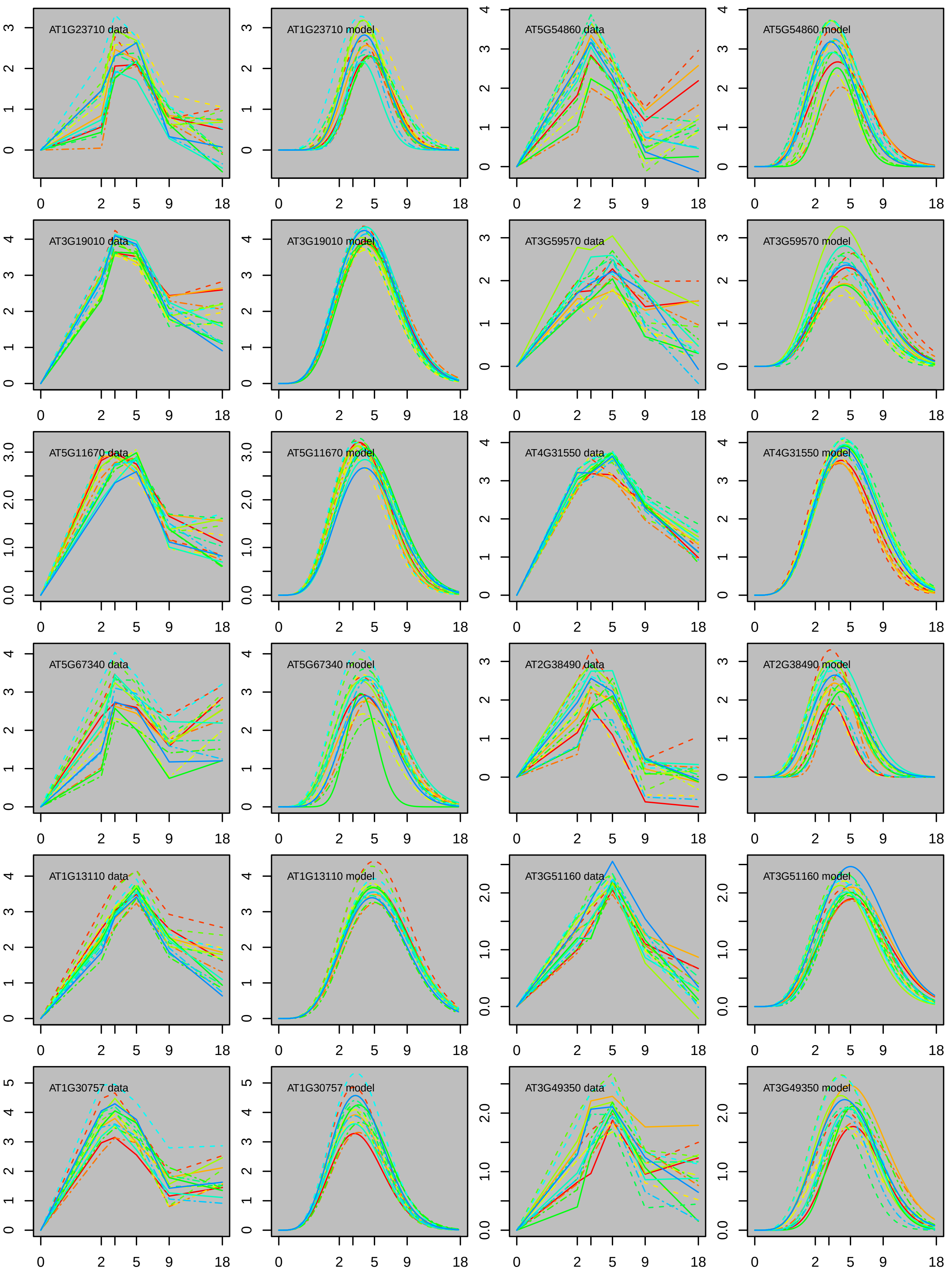
AT1G23710 data
AT1G23710 model
AT5G54860 data
AT5G54860 model
AT3G19010 data
AT3G19010 model
AT3G59570 data
AT3G59570 model
AT5G11670 data
AT5G11670 model
AT4G31550 data
AT4G31550 model
AT5G67340 data
AT5G67340 model
AT2G38490 data
AT2G38490 model
AT1G13110 data
AT1G13110 model
AT3G51160 data
AT3G51160 model
AT1G30757 data
AT1G30757 model
AT3G49350 data
AT3G49350 model

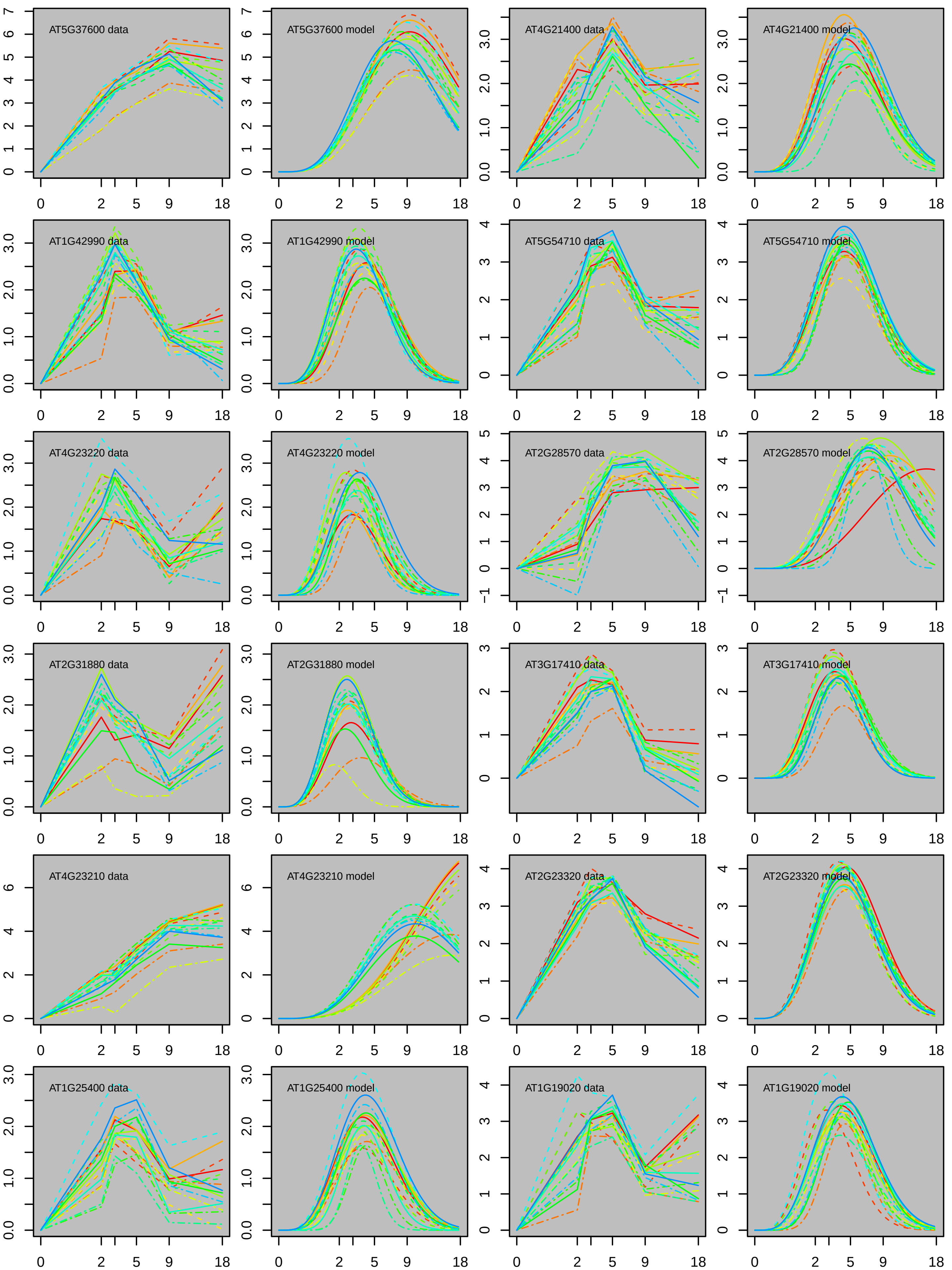
AT5G37600 data
AT5G37600 model
AT4G21400 data
AT4G21400 model
AT1G42990 data
AT1G42990 model
AT5G54710 data
AT5G54710 model
AT4G23220 data
AT4G23220 model
AT2G28570 data
AT2G28570 model
AT2G31880 data
AT2G31880 model
AT3G17410 data
AT3G17410 model
AT4G23210 data
AT4G23210 model
AT2G23320 data
AT2G23320 model
AT1G25400 data
AT1G25400 model
AT1G19020 data
AT1G19020 model

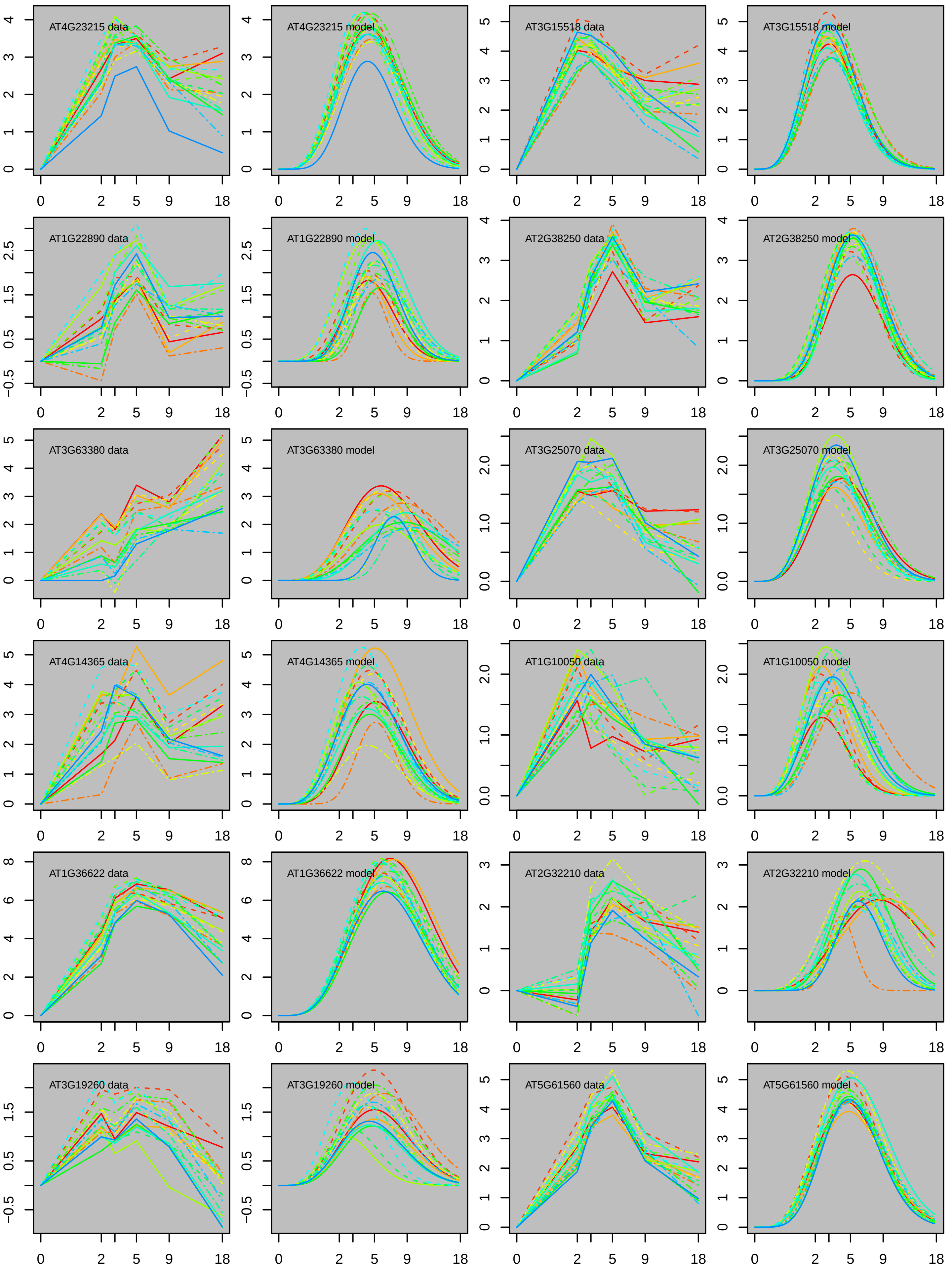
AT4G23215 data
AT4G23215 model
AT3G15518 data
AT3G15518 model
AT1G22890 data
AT1G22890 model
AT2G38250 data
AT2G38250 model
AT3G63380 data
AT3G63380 model
AT3G25070 data
AT3G25070 model
AT4G14365 data
AT4G14365 model
AT1G10050 data
AT1G10050 model
AT1G36622 data
AT1G36622 model
AT2G32210 data
AT2G32210 model
AT3G19260 data
AT3G19260 model
AT5G61560 data
AT5G61560 model
0
2
5
9
18

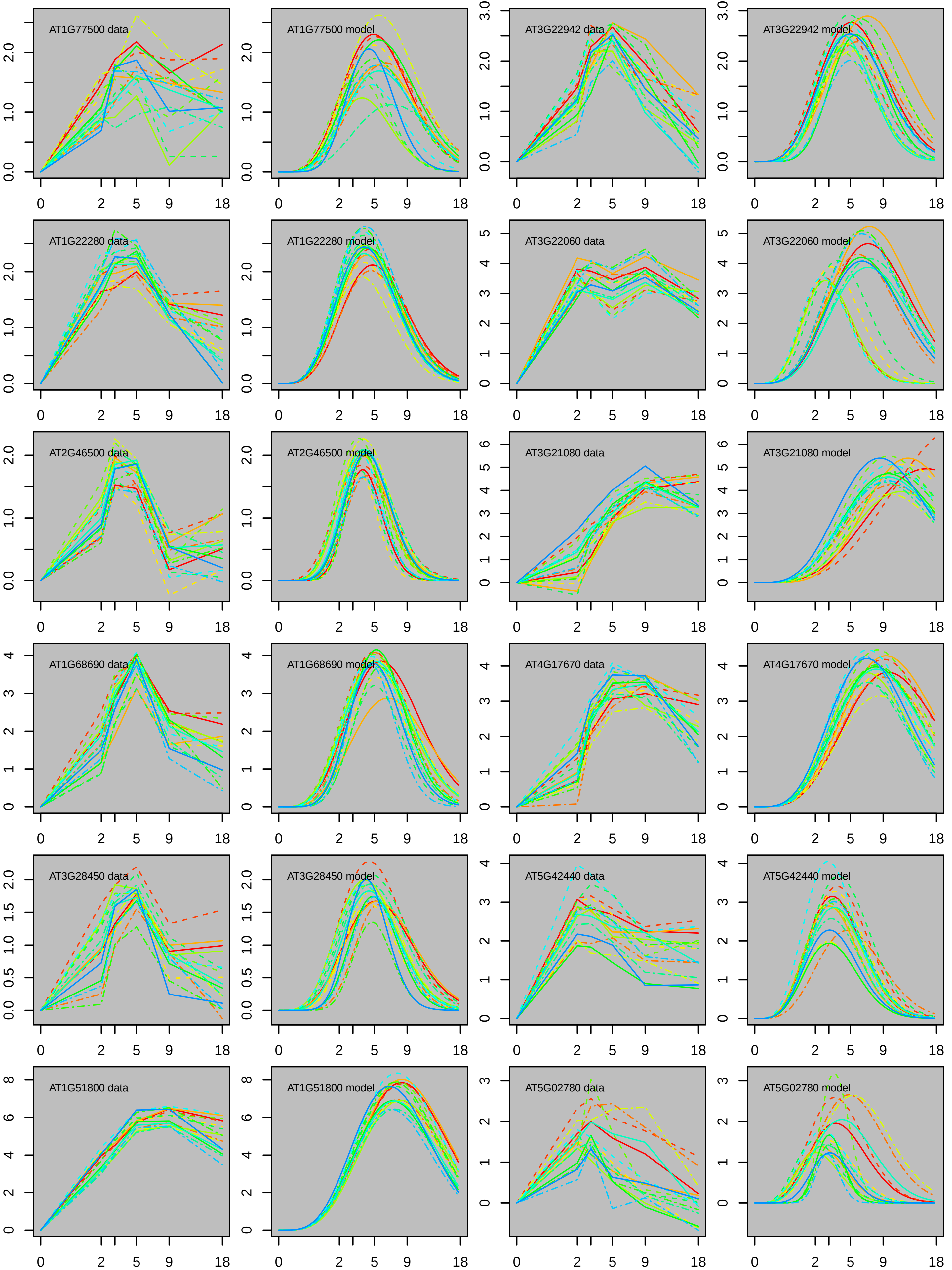
AT1G77500 data
AT1G77500 model
AT3G22942 data
AT3G22942 model
AT1G22280 data
AT1G22280 model
AT3G22060 data
AT3G22060 model
AT2G46500 data
AT2G46500 model
AT3G21080 data
AT3G21080 model
AT1G68690 data
AT1G68690 model
AT4G17670 data
AT4G17670 model
AT3G28450 data
AT3G28450 model
AT5G42440 data
AT5G42440 model
AT1G51800 data
AT1G51800 model
AT5G02780 data
AT5G02780 model
0
2
5
9
18

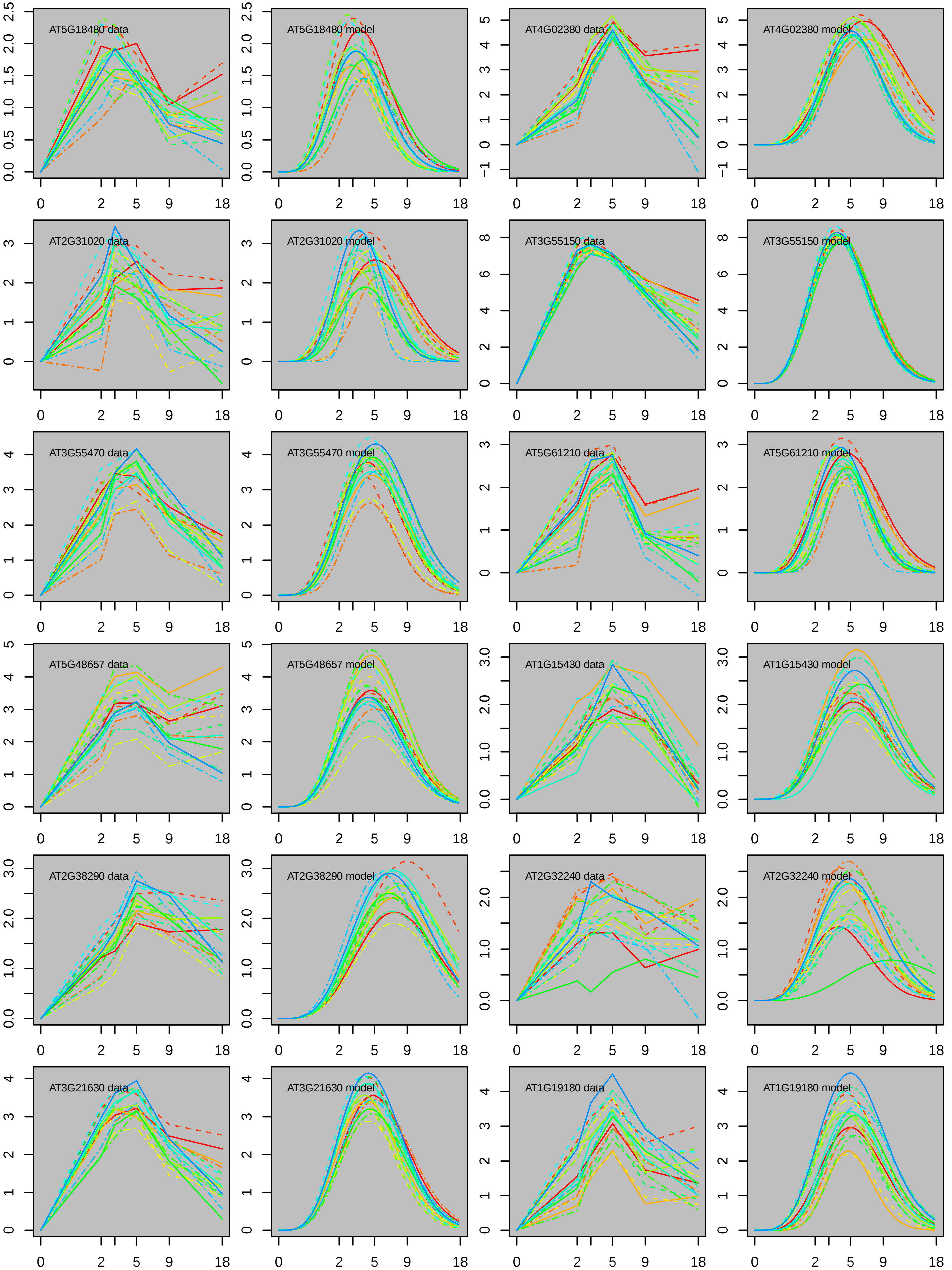

AT5G18480 data
AT5G18480 model
AT4G02380 data
AT4G02380 model
AT2G31020 data
AT2G31020 model
AT3G55150 data
AT3G55150 model
AT3G55470 data
AT3G55470 model
AT5G61210 data
AT5G61210 model
AT5G48657 data
AT5G48657 model
AT1G15430 data
AT1G15430 model
AT2G38290 data
AT2G38290 model
AT2G32240 data
AT2G32240 model
AT3G21630 data
AT3G21630 model
AT1G19180 data
AT1G19180 model
0
2
5
9
18

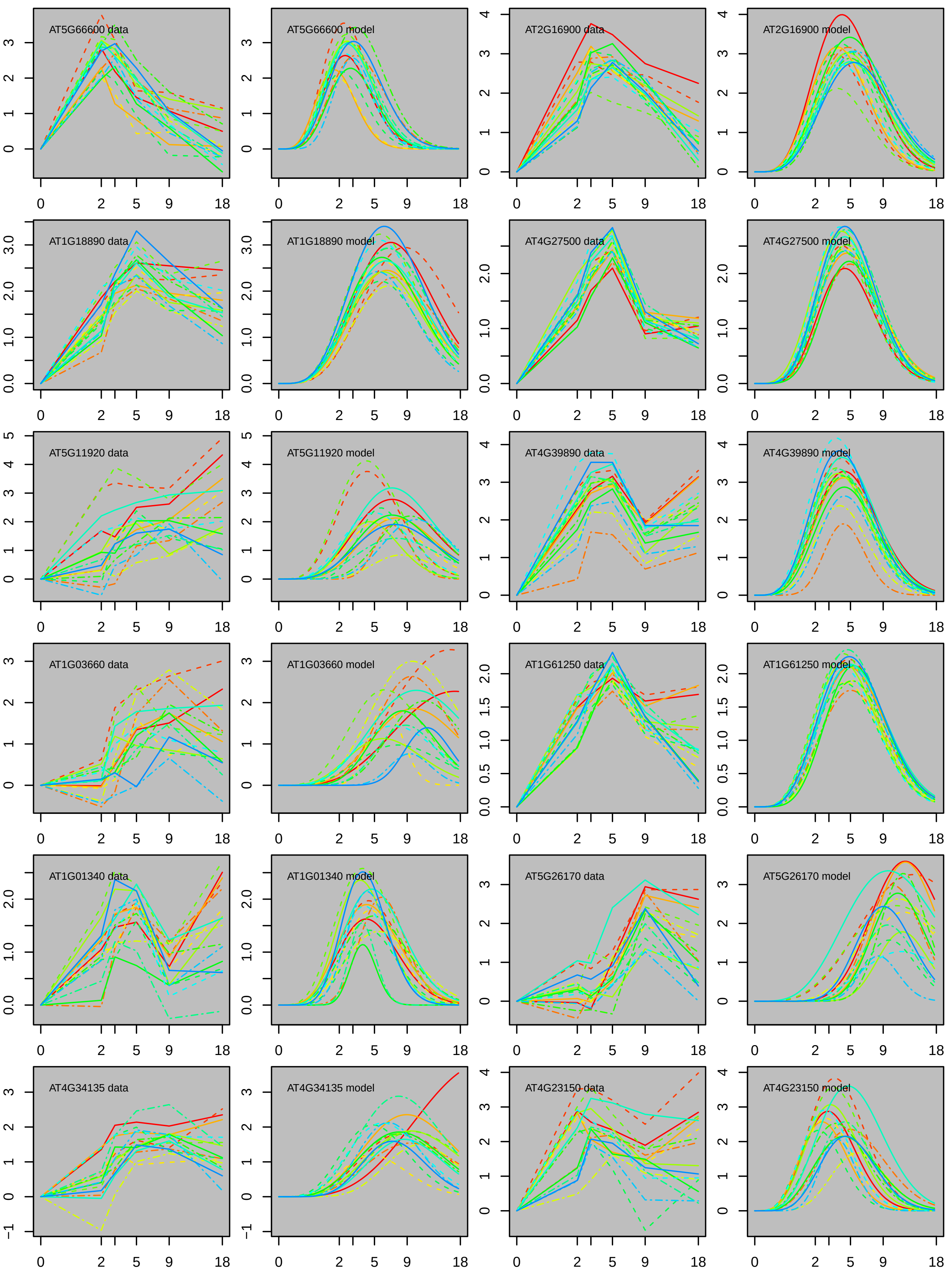
AT5G66600 data
AT5G66600 model
AT2G16900 data
AT2G16900 model
AT1G18890 data
AT1G18890 model
AT4G27500 data
AT4G27500 model
AT5G11920 data
AT5G11920 model
AT4G39890 data
AT4G39890 model
AT1G03660 data
AT1G03660 model
AT1G61250 data
AT1G61250 model
AT1G01340 data
AT1G01340 model
AT5G26170 data
AT5G26170 model
AT4G34135 data
AT4G34135 model
AT4G23150 data
AT4G23150 model

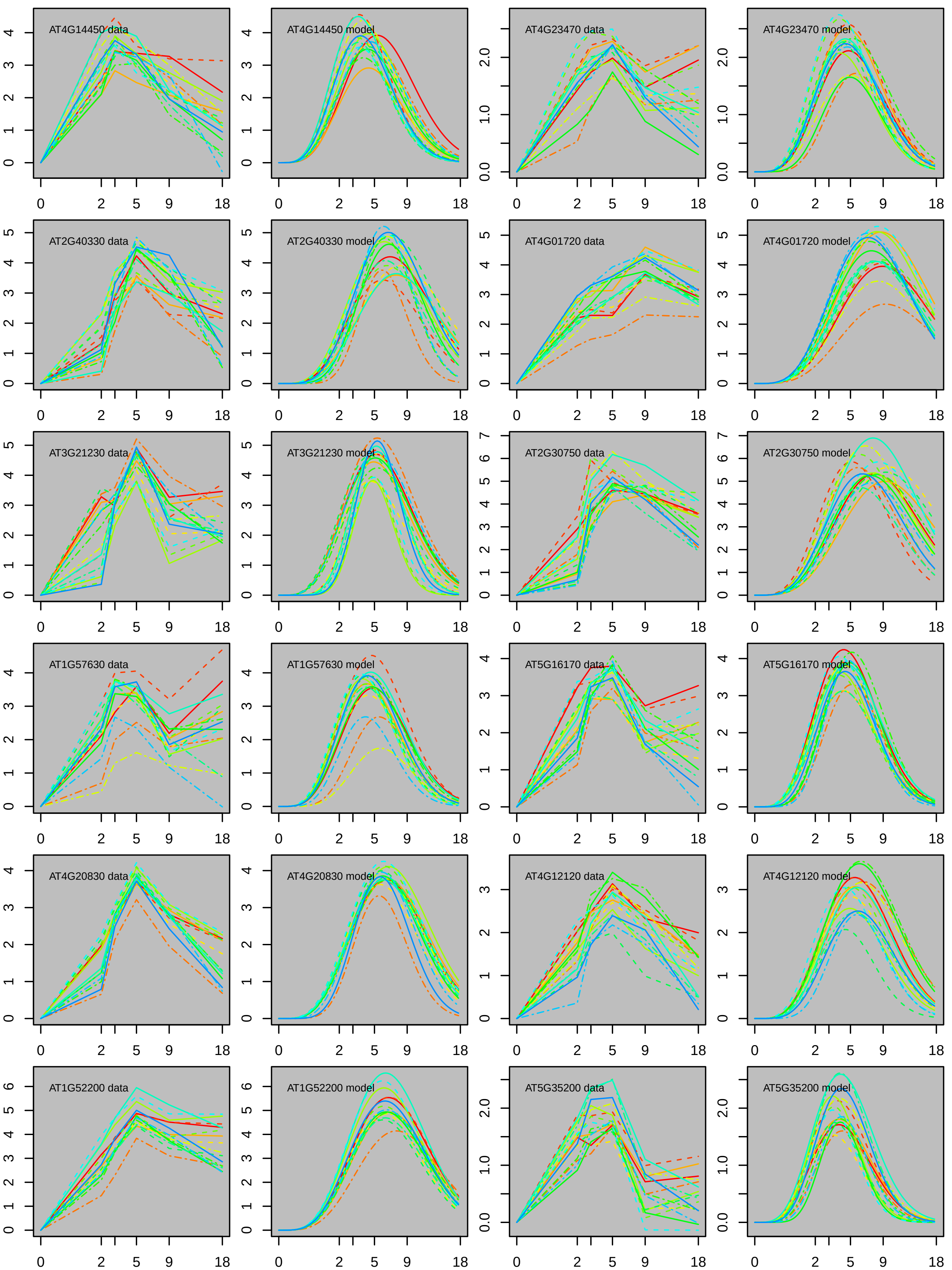

AT4G14450 data
AT4G14450 model
AT4G23470 data
AT4G23470 model
AT2G40330 data
AT2G40330 model
AT4G01720 data
AT4G01720 model
AT3G21230 data
AT3G21230 model
AT2G30750 data
AT2G30750 model
AT1G57630 data
AT1G57630 model
AT5G16170 data
AT5G16170 model
AT4G20830 data
AT4G20830 model
AT4G12120 data
AT4G12120 model
AT1G52200 data
AT1G52200 model
AT5G35200 data
AT5G35200 model
0 2 5 9 18

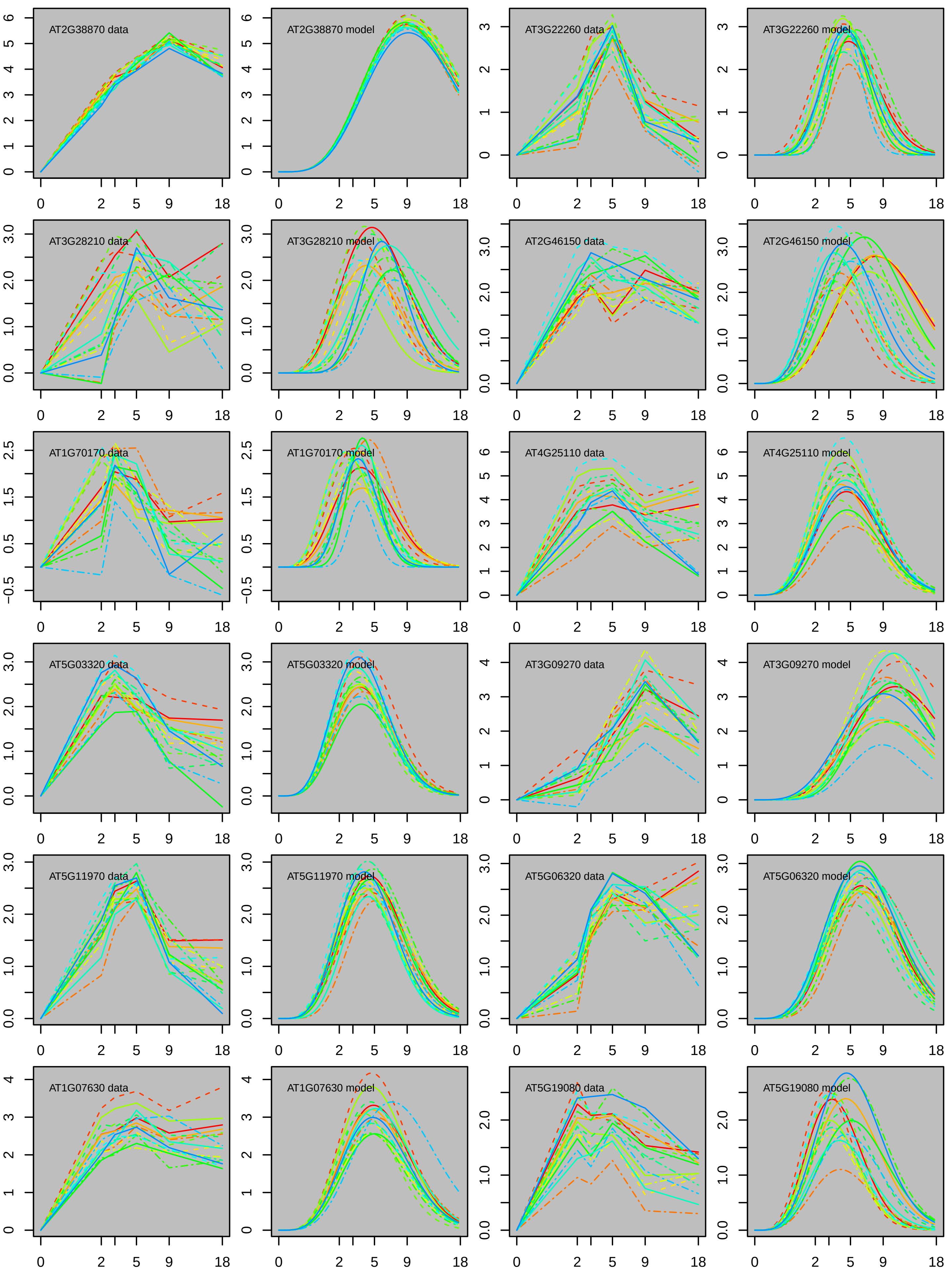

AT2G38870 data
AT2G38870 model
AT3G22260 data
AT3G22260 model
AT3G28210 data
AT3G28210 model
AT2G46150 data
AT2G46150 model
AT1G70170 data
AT1G70170 model
AT4G25110 data
AT4G25110 model
AT5G03320 data
AT5G03320 model
AT3G09270 data
AT3G09270 model
AT5G11970 data
AT5G11970 model
AT5G06320 data
AT5G06320 model
AT1G07630 data
AT1G07630 model
AT5G19080 data
AT5G19080 model

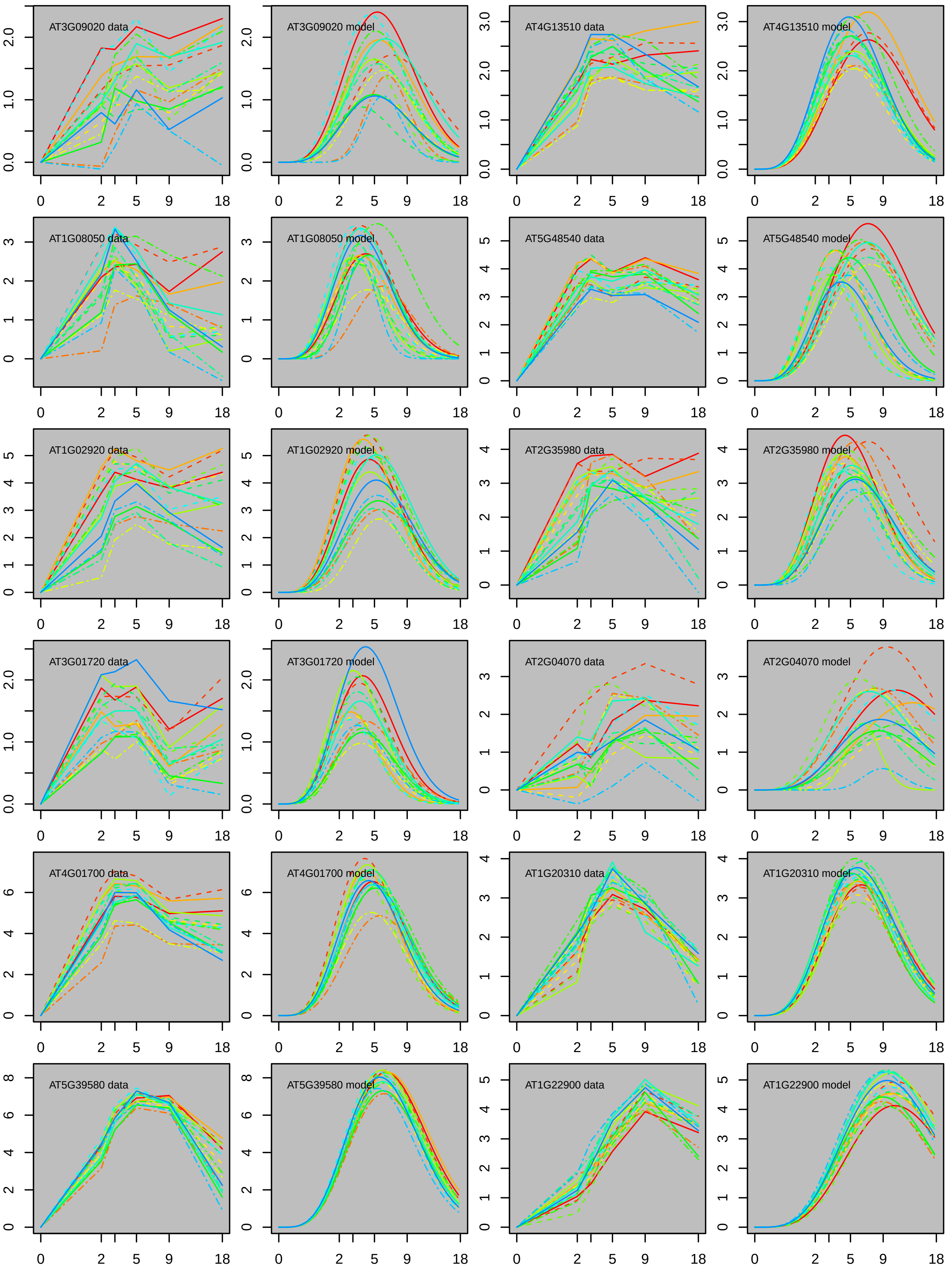
AT3G09020 data
AT3G09020 model
AT4G13510 data
AT4G13510 model
AT1G08050 data
AT1G08050 model
AT5G48540 data
AT5G48540 model
AT1G02920 data
AT1G02920 model
AT2G35980 data
AT2G35980 model
AT3G01720 data
AT3G01720 model
AT2G04070 data
AT2G04070 model
AT4G01700 data
AT4G01700 model
AT1G20310 data
AT1G20310 model
AT5G39580 data
AT5G39580 model
AT1G22900 data
AT1G22900 model
0
2
5
9
18

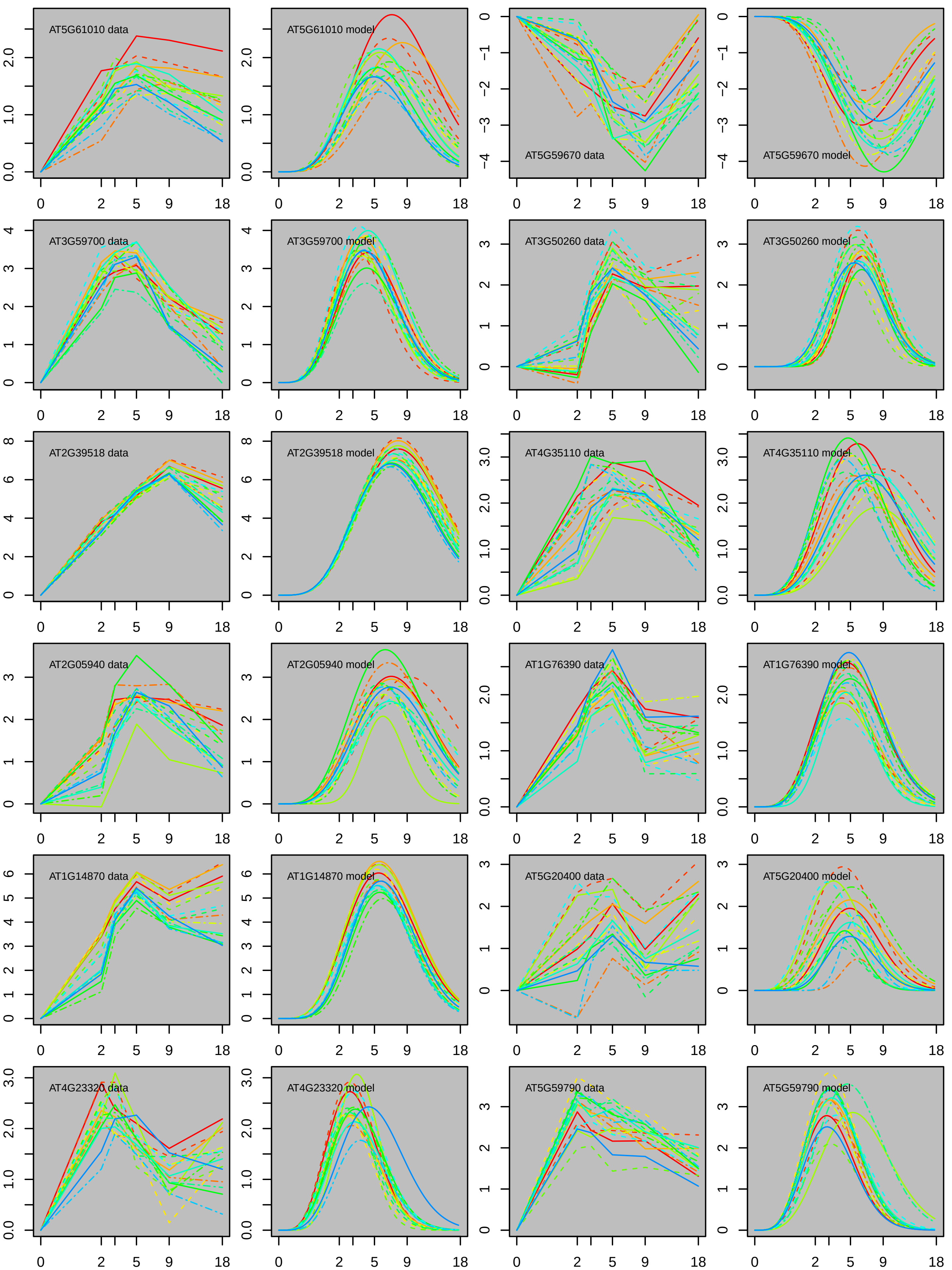
AT5G61010 data
AT5G61010 model
AT5G59670 data
AT5G59670 model
AT3G59700 data
AT3G59700 model
AT3G50260 data
AT3G50260 model
AT2G39518 data
AT2G39518 model
AT4G35110 data
AT4G35110 model
AT2G05940 data
AT2G05940 model
AT1G76390 data
AT1G76390 model
AT1G14870 data
AT1G14870 model
AT5G20400 data
AT5G20400 model
AT4G23320 data
AT4G23320 model
AT5G59790 data
AT5G59790 model

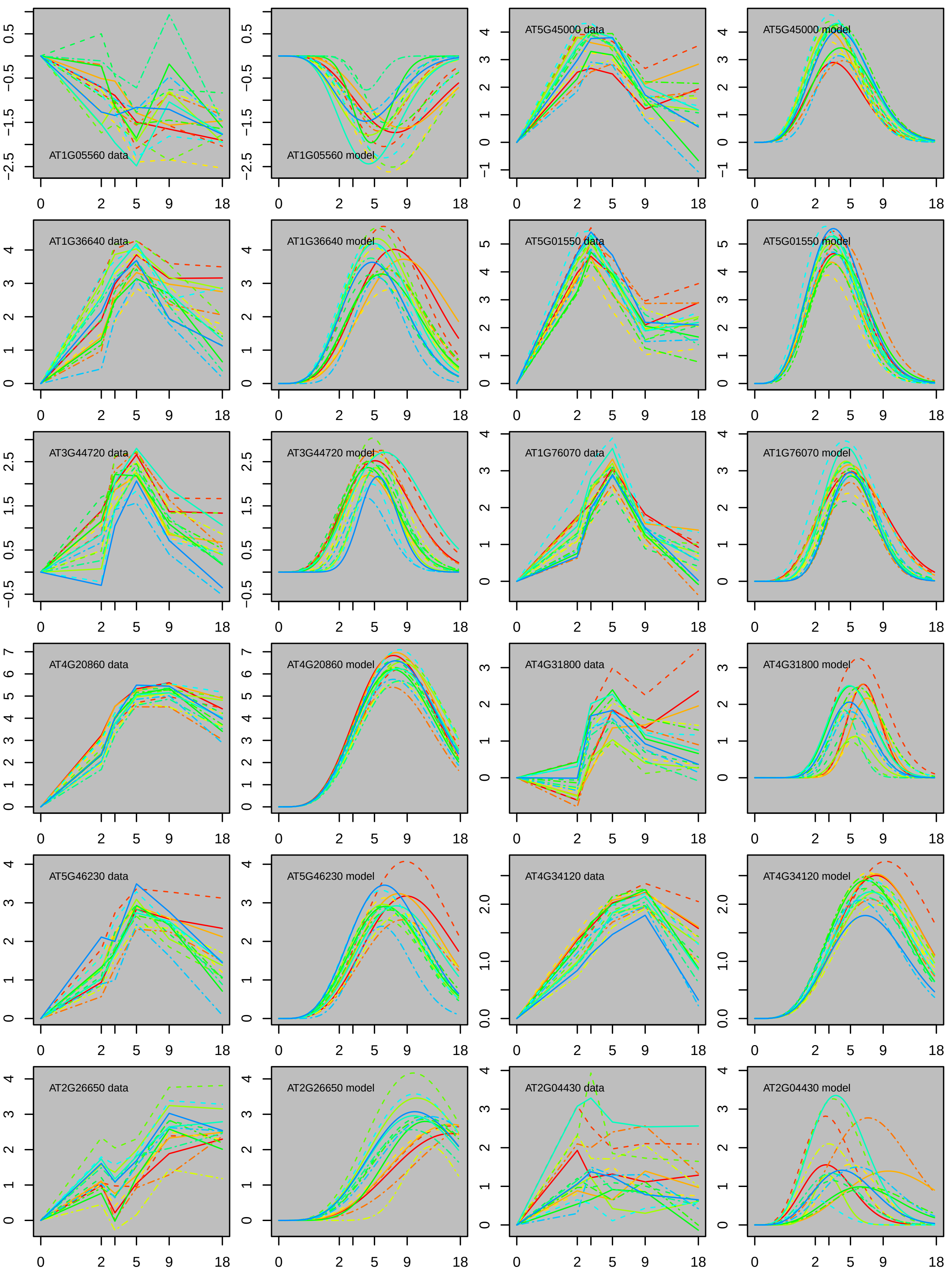
AT1G05560 data
AT1G05560 model
AT5G45000 data
AT5G45000 model
AT1G36640 data
AT1G36640 model
AT5G01550 data
AT5G01550 model
AT3G44720 data
AT3G44720 model
AT1G76070 data
AT1G76070 model
AT4G20860 data
AT4G20860 model
AT4G31800 data
AT4G31800 model
AT5G46230 data
AT5G46230 model
AT4G34120 data
AT4G34120 model
AT2G26650 data
AT2G26650 model
AT2G04430 data
AT2G04430 model

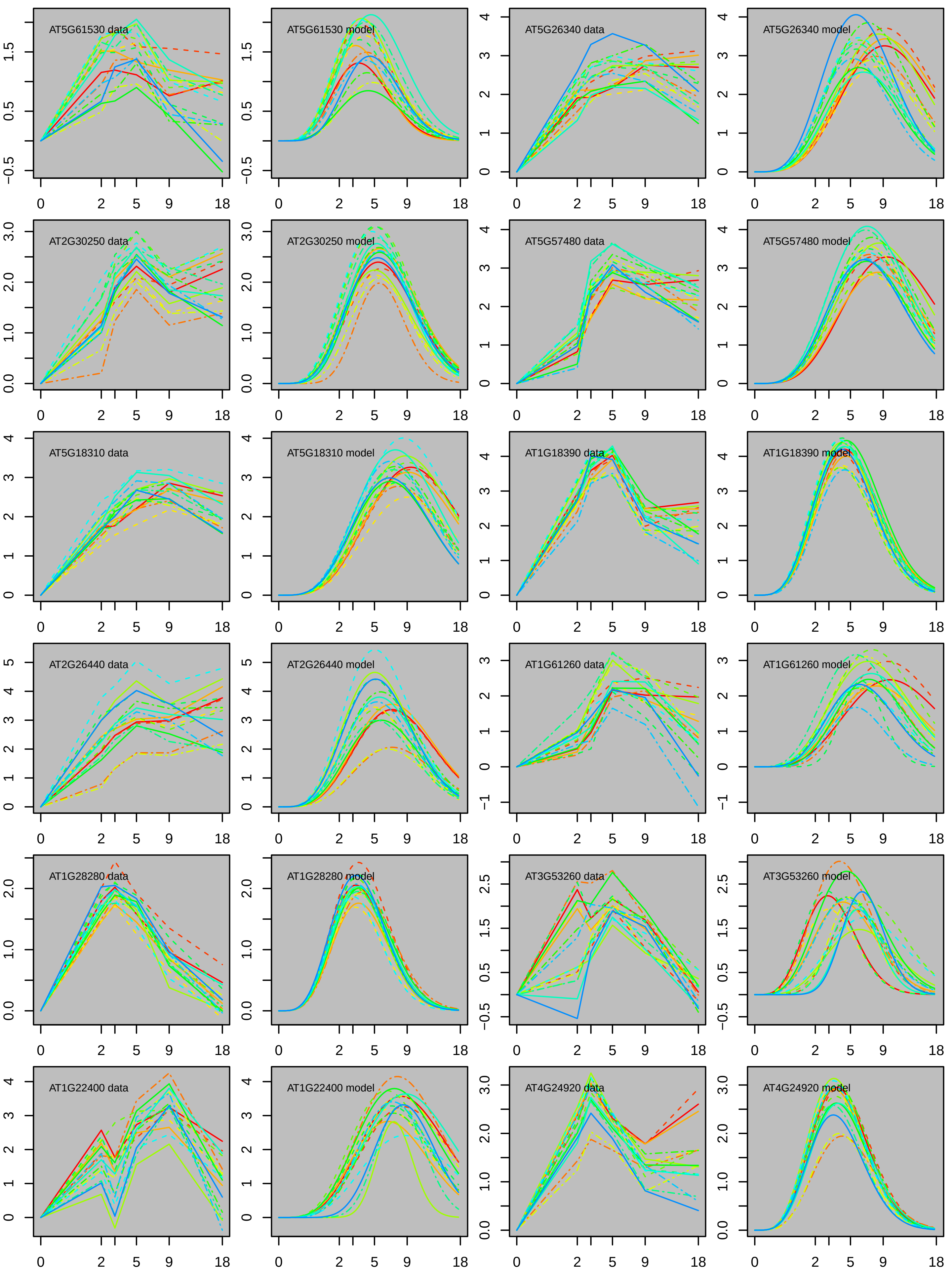
AT5G61530 data
AT5G61530 model
AT5G26340 data
AT5G26340 model
AT2G30250 data
AT2G30250 model
AT5G57480 data
AT5G57480 model
AT5G18310 data
AT5G18310 model
AT1G18390 data
AT1G18390 model
AT2G26440 data
AT2G26440 model
AT1G61260 data
AT1G61260 model
AT1G28280 data
AT1G28280 model
AT3G53260 data
AT3G53260 model
AT1G22400 data
AT1G22400 model
AT4G24920 data
AT4G24920 model
0
2
5
9
18

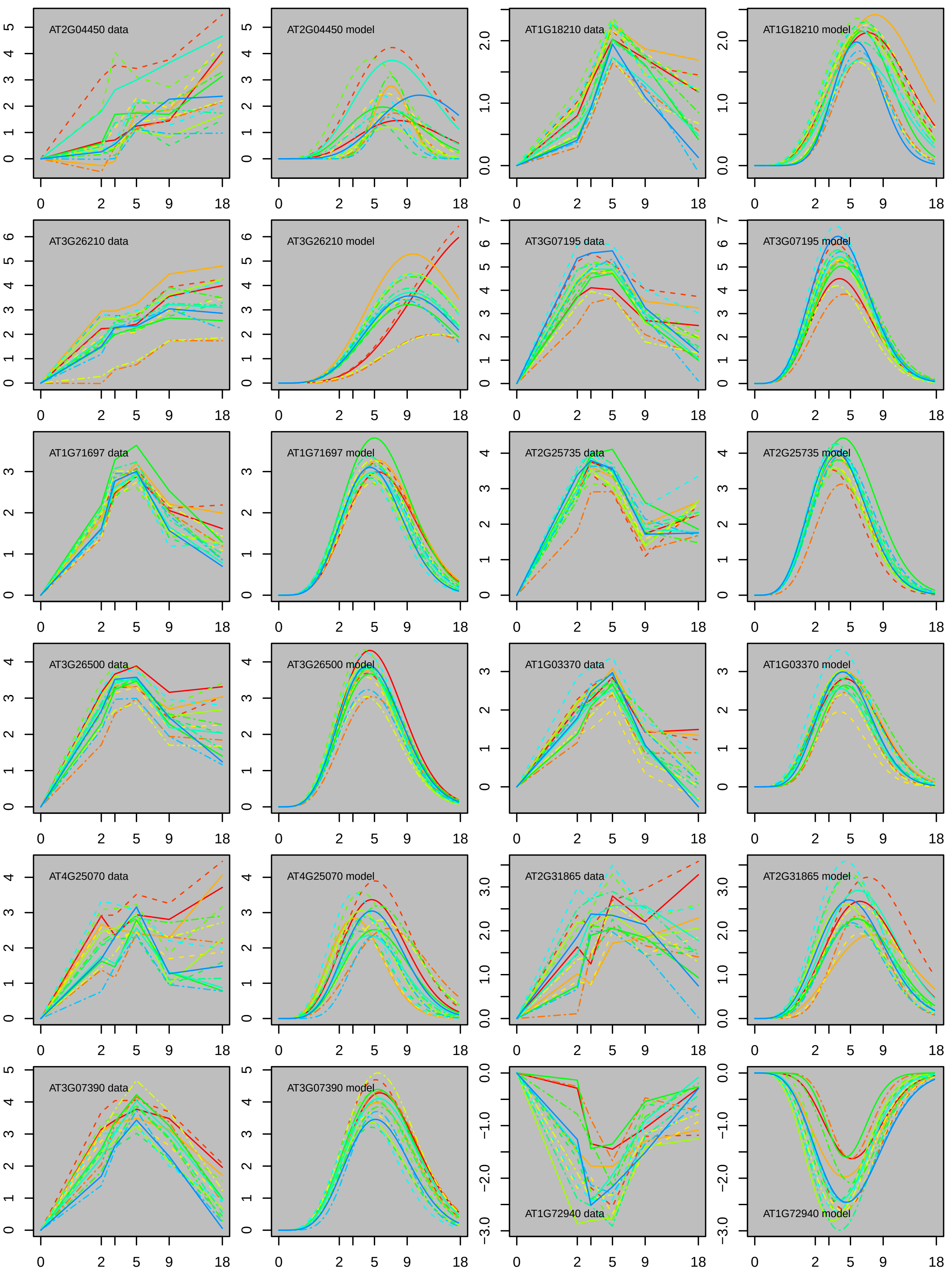

AT2G04450 data
AT2G04450 model
AT1G18210 data
AT1G18210 model
AT3G26210 data
AT3G26210 model
AT3G07195 data
AT3G07195 model
AT1G71697 data
AT1G71697 model
AT2G25735 data
AT2G25735 model
AT3G26500 data
AT3G26500 model
AT1G03370 data
AT1G03370 model
AT4G25070 data
AT4G25070 model
AT2G31865 data
AT2G31865 model
AT3G07390 data
AT3G07390 model
AT1G72940 data
AT1G72940 model
0 2 5 9 18

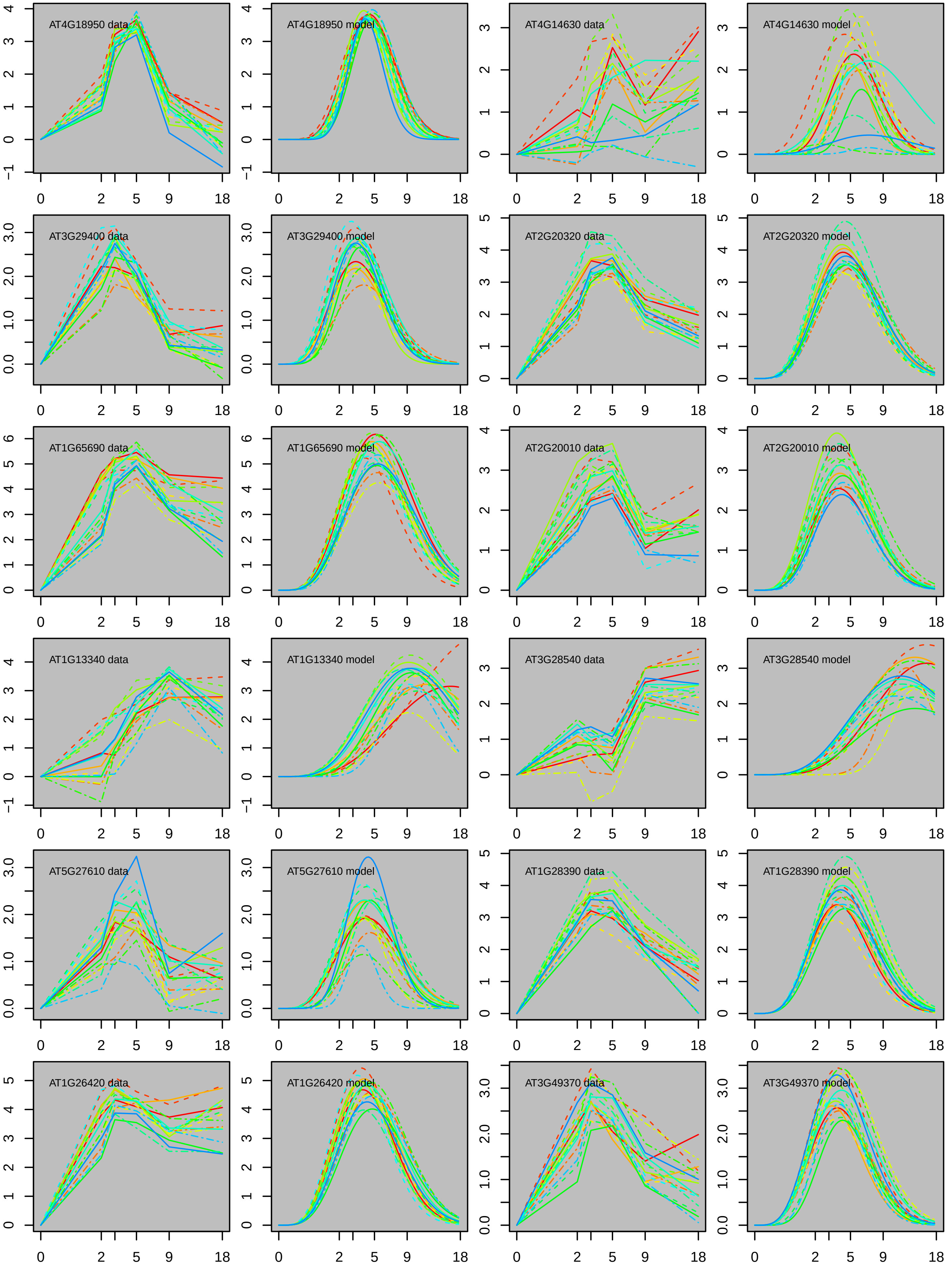
AT4G18950 data
AT4G18950 model
AT4G14630 data
AT4G14630 model
AT3G29400 data
AT3G29400 model
AT2G20320 data
AT2G20320 model
AT1G65690 data
AT1G65690 model
AT2G20010 data
AT2G20010 model
AT1G13340 data
AT1G13340 model
AT3G28540 data
AT3G28540 model
AT5G27610 data
AT5G27610 model
AT1G28390 data
AT1G28390 model
AT1G26420 data
AT1G26420 model
AT3G49370 data
AT3G49370 model
0
2
5
9
18

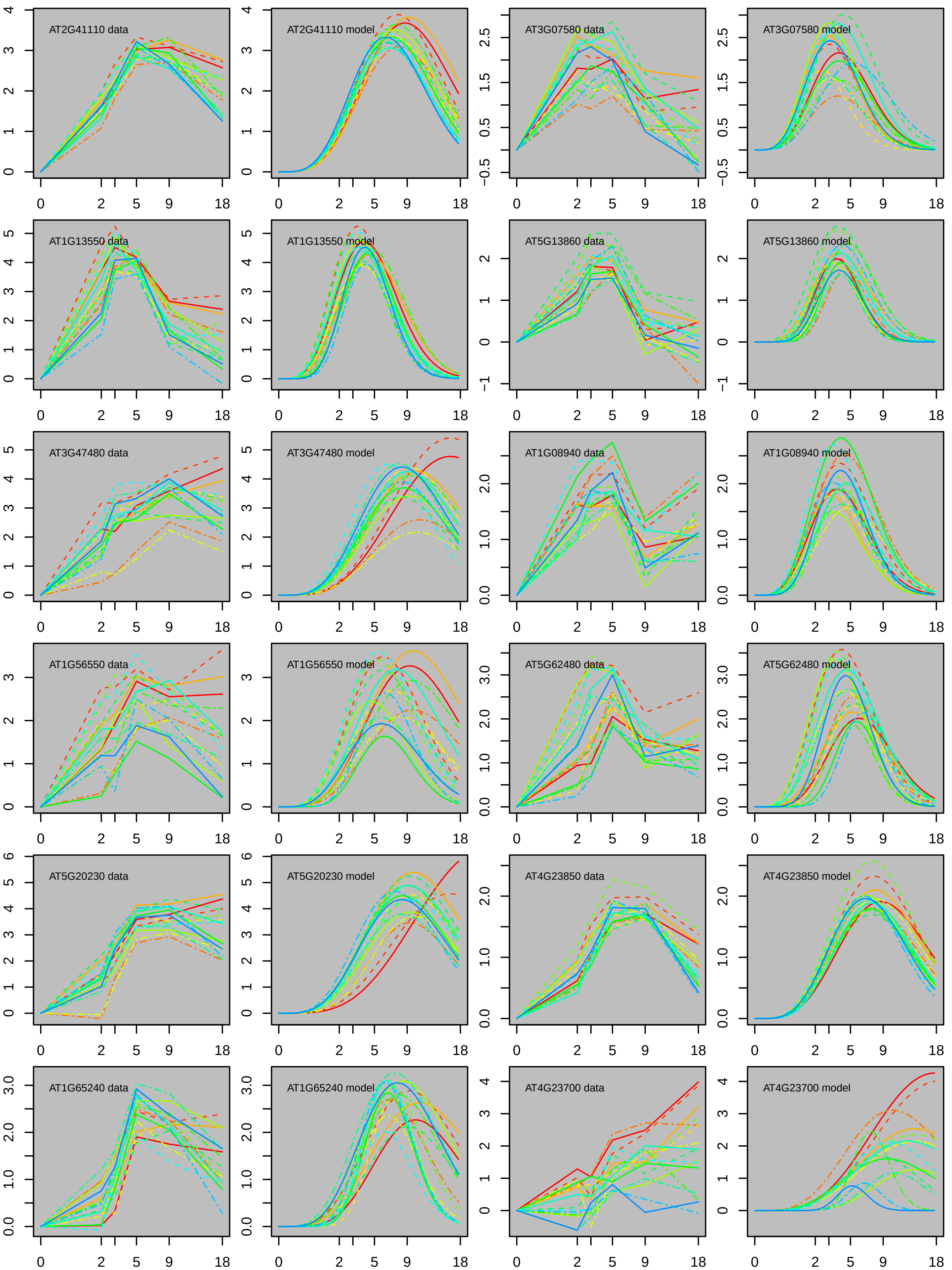
AT2G41110 data
AT2G41110 model
AT3G07580 data
AT3G07580 model
AT1G13550 data
AT1G13550 model
AT5G13860 data
AT5G13860 model
AT3G47480 data
AT3G47480 model
AT1G08940 data
AT1G08940 model
AT1G56550 data
AT1G56550 model
AT5G62480 data
AT5G62480 model
AT5G20230 data
AT5G20230 model
AT4G23850 data
AT4G23850 model
AT1G65240 data
AT1G65240 model
AT4G23700 data
AT4G23700 model

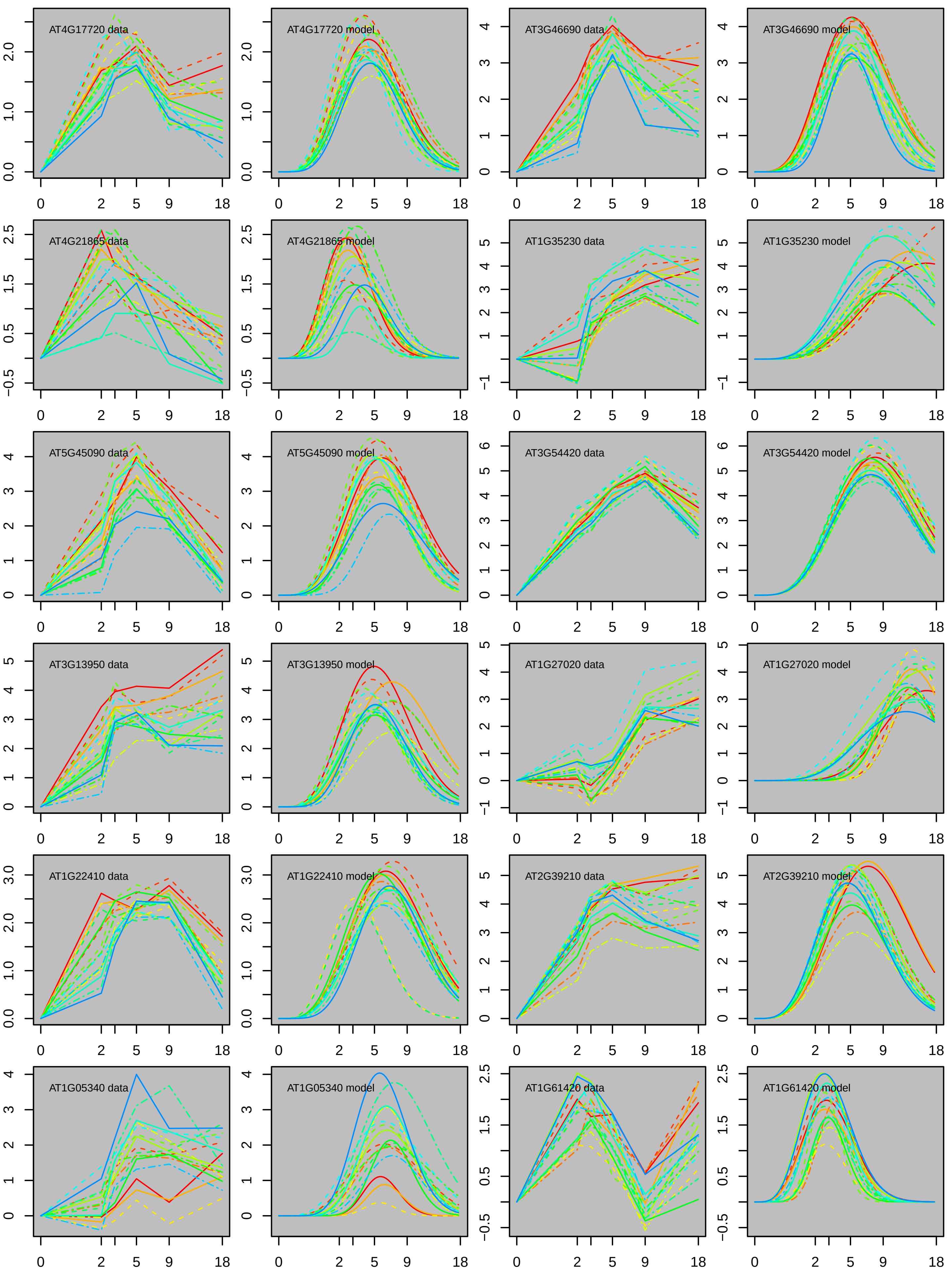

AT4G17720 data
AT4G17720 model
AT3G46690 data
AT3G46690 model
AT4G21865 data
AT4G21865 model
AT1G35230 data
AT1G35230 model
AT5G45090 data
AT5G45090 model
AT3G54420 data
AT3G54420 model
AT3G13950 data
AT3G13950 model
AT1G27020 data
AT1G27020 model
AT1G22410 data
AT1G22410 model
AT2G39210 data
AT2G39210 model
AT1G05340 data
AT1G05340 model
AT1G61420 data
AT1G61420 model

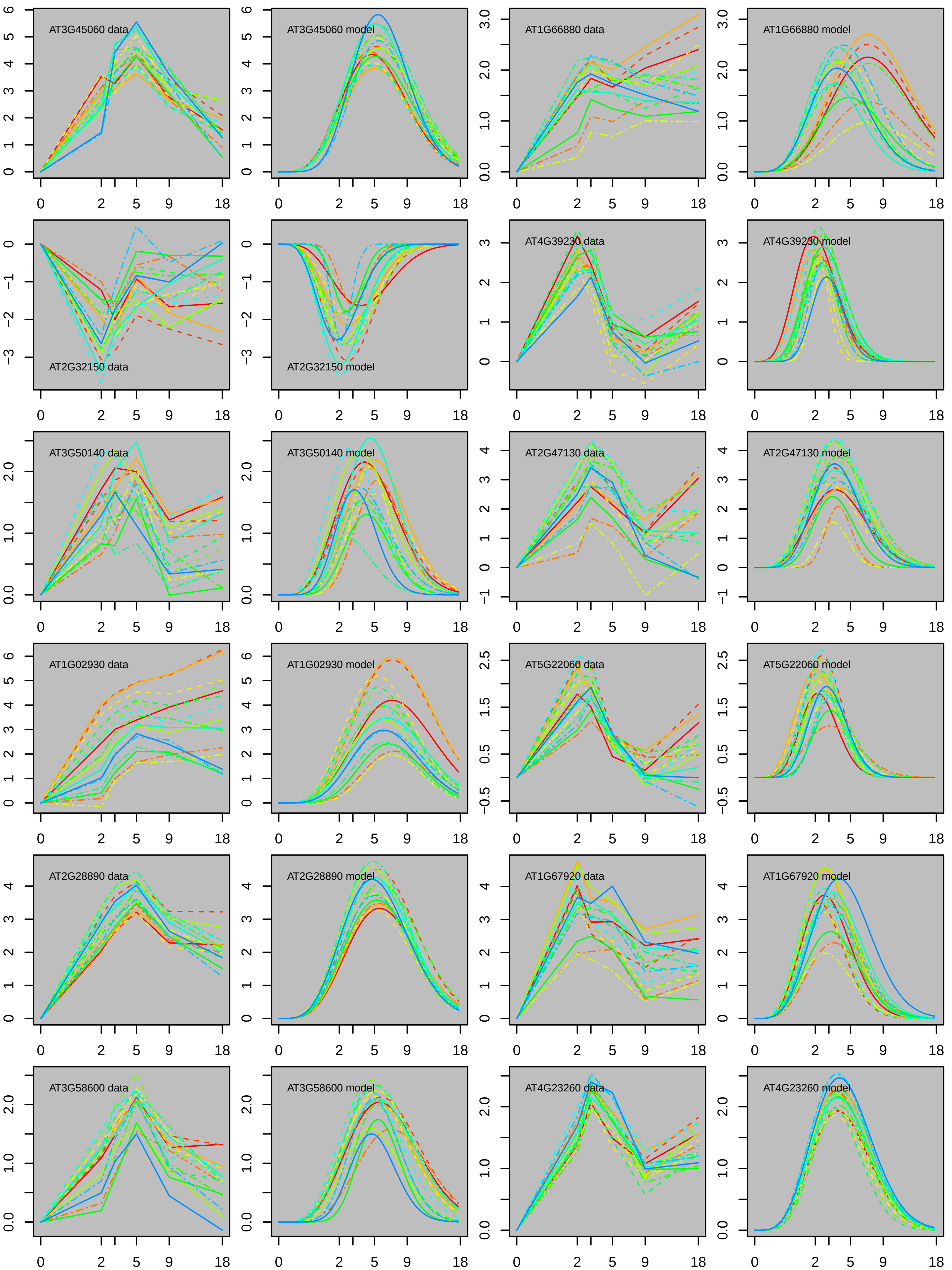
AT3G45060 data
AT3G45060 model
AT1G66880 data
AT1G66880 model
AT2G32150 data
AT2G32150 model
AT4G39230 data
AT4G39230 model
AT3G50140 data
AT3G50140 model
AT2G47130 data
AT2G47130 model
AT1G02930 data
AT1G02930 model
AT5G22060 data
AT5G22060 model
AT2G28890 data
AT2G28890 model
AT1G67920 data
AT1G67920 model
AT3G58600 data
AT3G58600 model
AT4G23260 data
AT4G23260 model

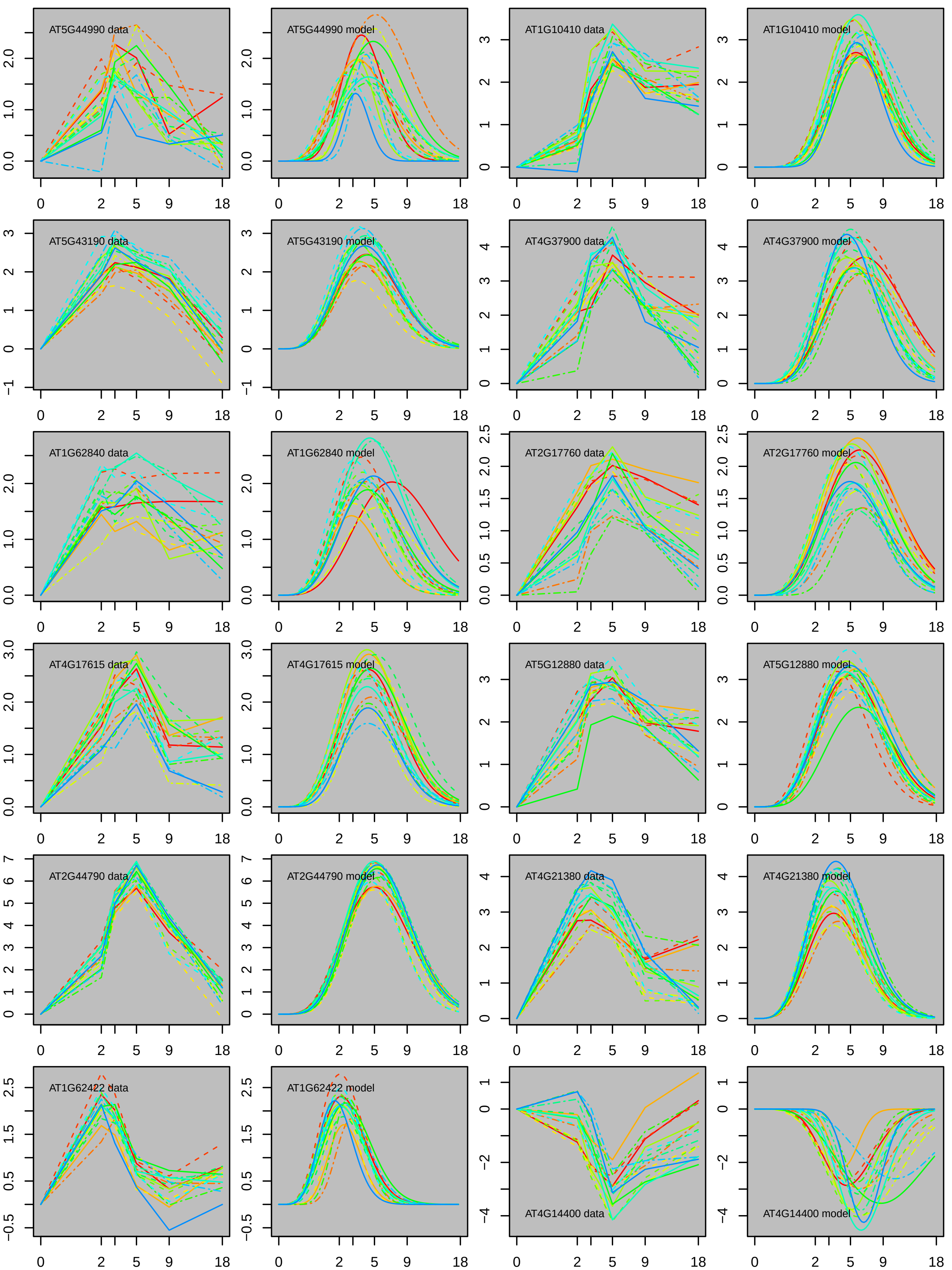
AT5G44990 data
AT5G44990 model
AT1G10410 data
AT1G10410 model
AT5G43190 data
AT5G43190 model
AT4G37900 data
AT4G37900 model
AT1G62840 data
AT1G62840 model
AT2G17760 data
AT2G17760 model
AT4G17615 data
AT4G17615 model
AT5G12880 data
AT5G12880 model
AT2G44790 data
AT2G44790 model
AT4G21380 data
AT4G21380 model
AT1G62422 data
AT1G62422 model
AT4G14400 data
AT4G14400 model
0 2 5 9 18

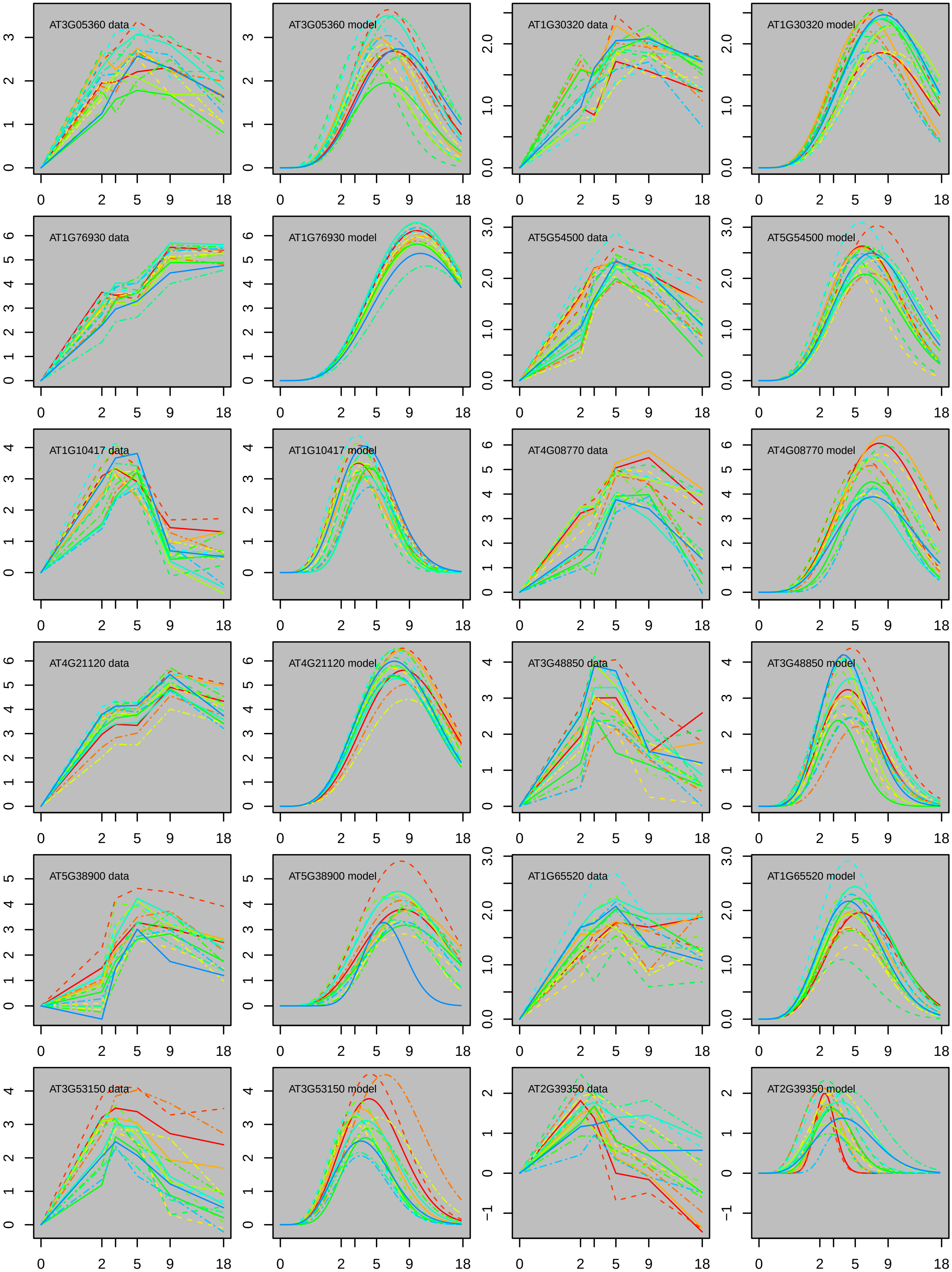
AT3G05360 data
AT3G05360 model
AT1G30320 data
AT1G30320 model
AT1G76930 data
AT1G76930 model
AT5G54500 data
AT5G54500 model
AT1G10417 data
AT1G10417 model
AT4G08770 data
AT4G08770 model
AT4G21120 data
AT4G21120 model
AT3G48850 data
AT3G48850 model
AT5G38900 data
AT5G38900 model
AT1G65520 data
AT1G65520 model
AT3G53150 data
AT3G53150 model
AT2G39350 data
AT2G39350 model

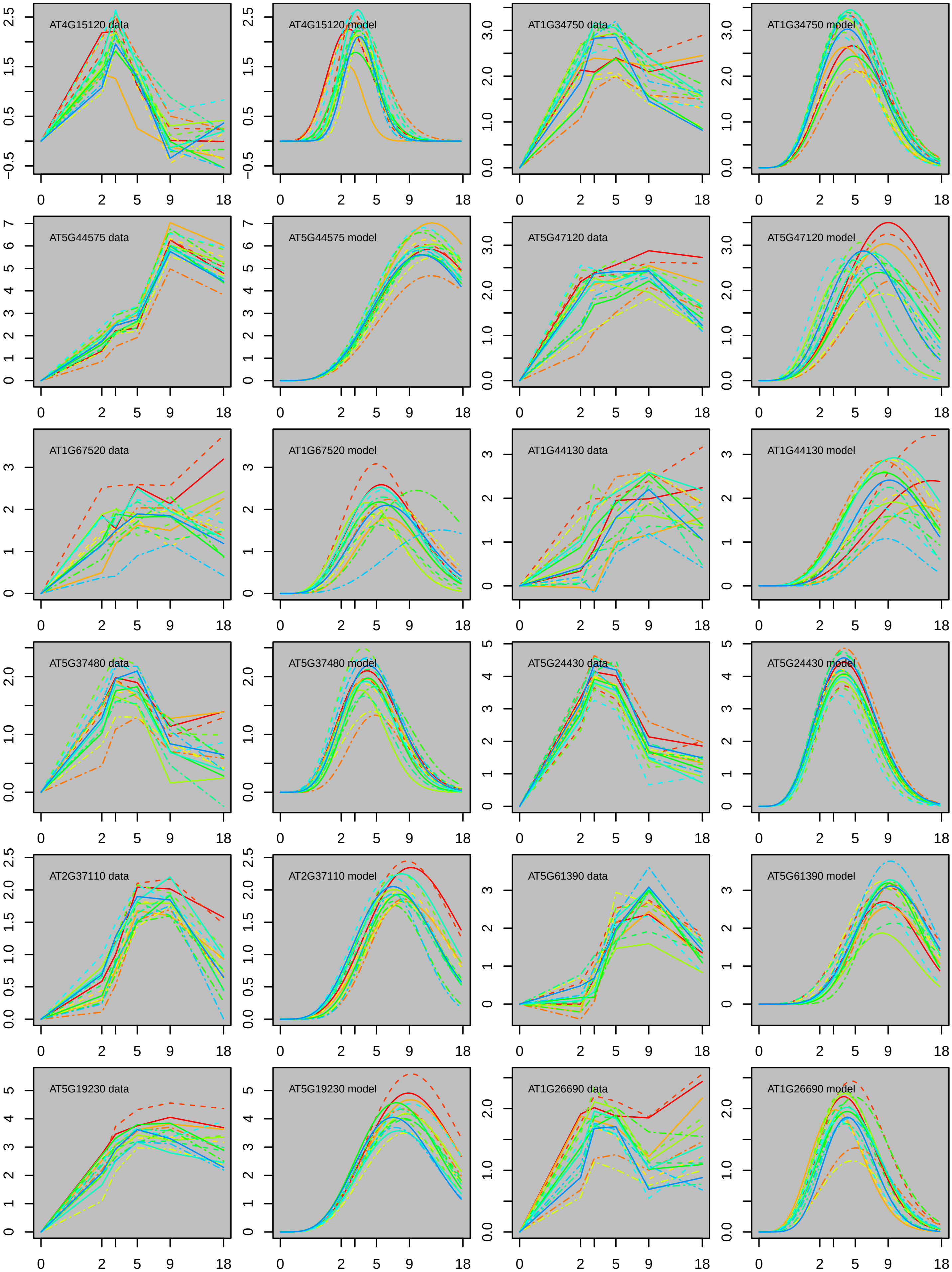

AT4G15120 data
AT4G15120 model
AT1G34750 data
AT1G34750 model
AT5G44575 data
AT5G44575 model
AT5G47120 data
AT5G47120 model
AT1G67520 data
AT1G67520 model
AT1G44130 data
AT1G44130 model
AT5G37480 data
AT5G37480 model
AT5G24430 data
AT5G24430 model
AT2G37110 data
AT2G37110 model
AT5G61390 data
AT5G61390 model
AT5G19230 data
AT5G19230 model
AT1G26690 data
AT1G26690 model
0 2 5 9 18

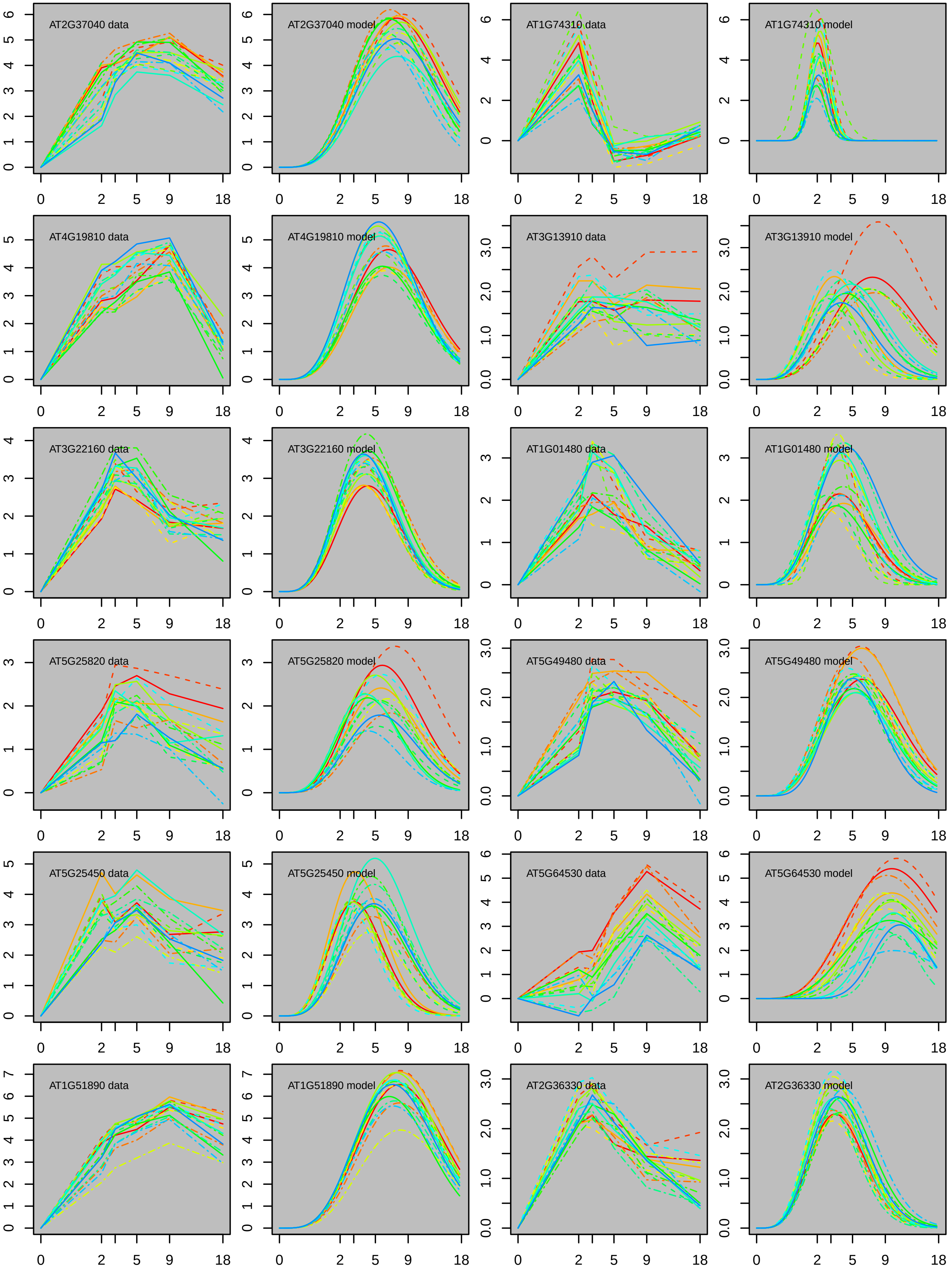
AT2G37040 data
AT2G37040 model
AT1G74310 data
AT1G74310 model
AT4G19810 data
AT4G19810 model
AT3G13910 data
AT3G13910 model
AT3G22160 data
AT3G22160 model
AT1G01480 data
AT1G01480 model
AT5G25820 data
AT5G25820 model
AT5G49480 data
AT5G49480 model
AT5G25450 data
AT5G25450 model
AT5G64530 data
AT5G64530 model
AT1G51890 data
AT1G51890 model
AT2G36330 data
AT2G36330 model
0 2 5 9 18

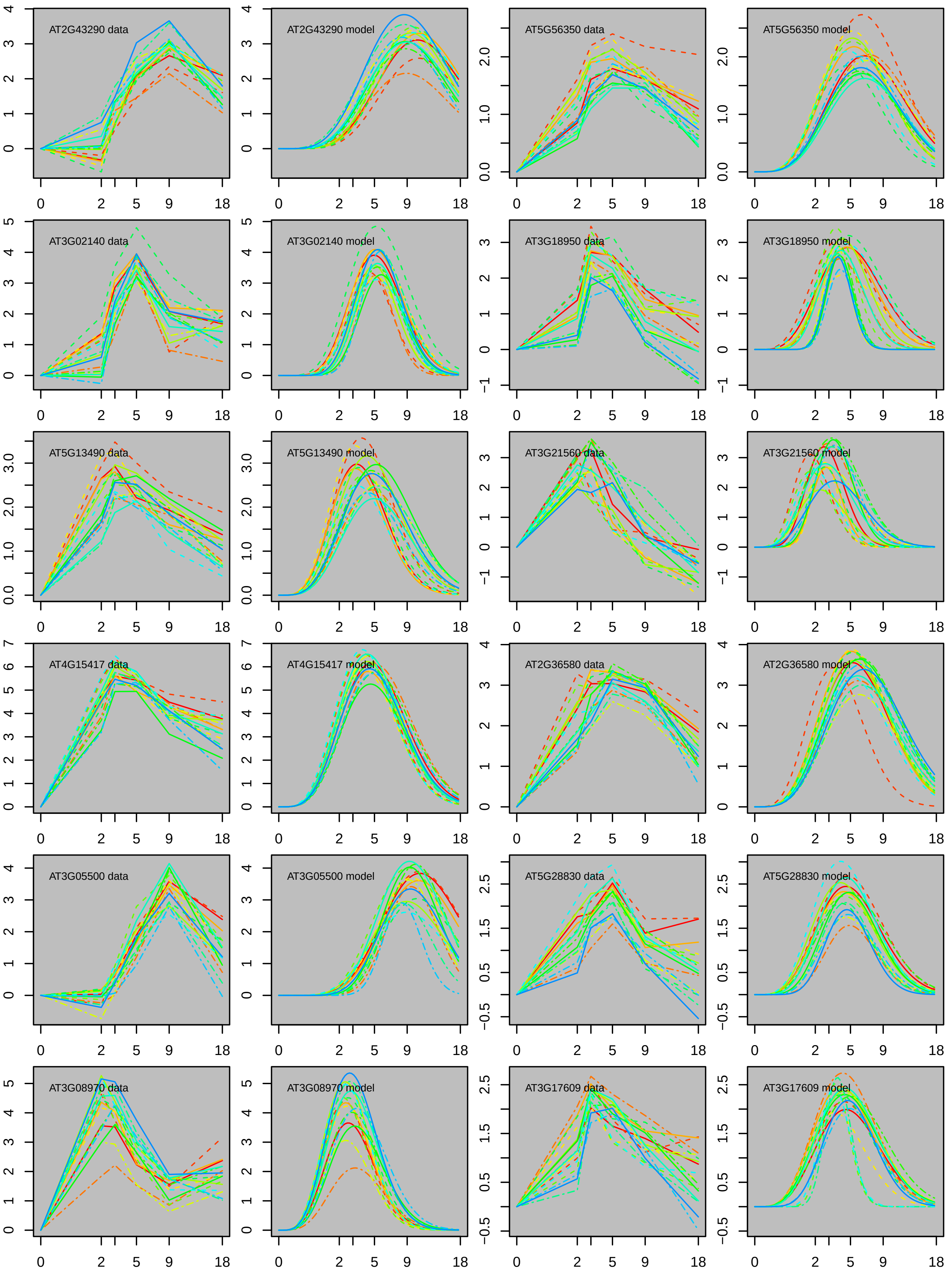

AT2G43290 data
AT2G43290 model
AT5G56350 data
AT5G56350 model
AT3G02140 data
AT3G02140 model
AT3G18950 data
AT3G18950 model
AT5G13490 data
AT5G13490 model
AT3G21560 data
AT3G21560 model
AT4G15417 data
AT4G15417 model
AT2G36580 data
AT2G36580 model
AT3G05500 data
AT3G05500 model
AT5G28830 data
AT5G28830 model
AT3G08970 data
AT3G08970 model
AT3G17609 data
AT3G17609 model

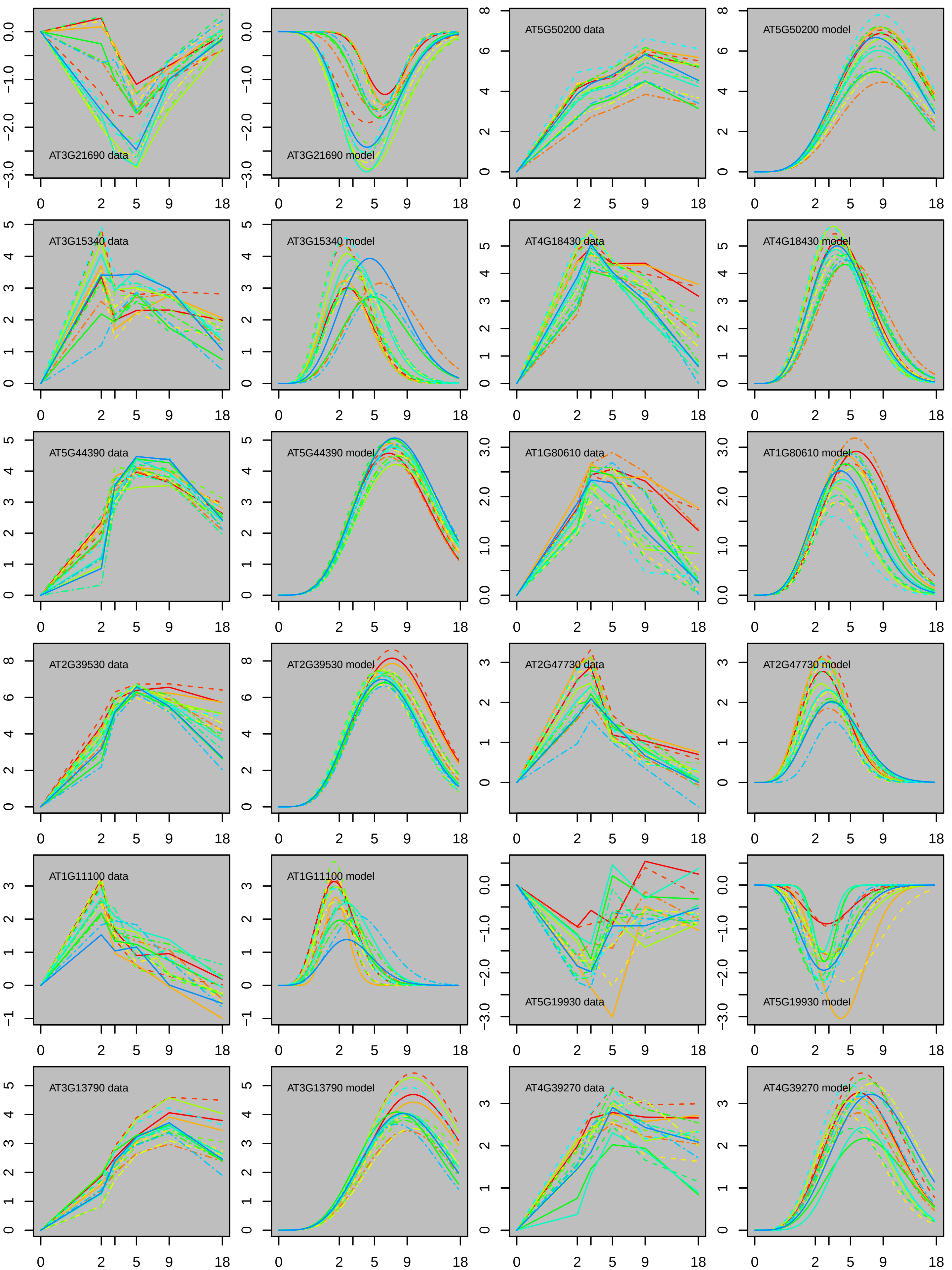

AT3G21690 data
AT3G21690 model
AT5G50200 data
AT5G50200 model
AT3G15340 data
AT3G15340 model
AT4G18430 data
AT4G18430 model
AT5G44390 data
AT5G44390 model
AT1G80610 data
AT1G80610 model
AT2G39530 data
AT2G39530 model
AT2G47730 data
AT2G47730 model
AT1G11100 data
AT1G11100 model
AT5G19930 data
AT5G19930 model
AT3G13790 data
AT3G13790 model
AT4G39270 data
AT4G39270 model
0 2 5 9 18

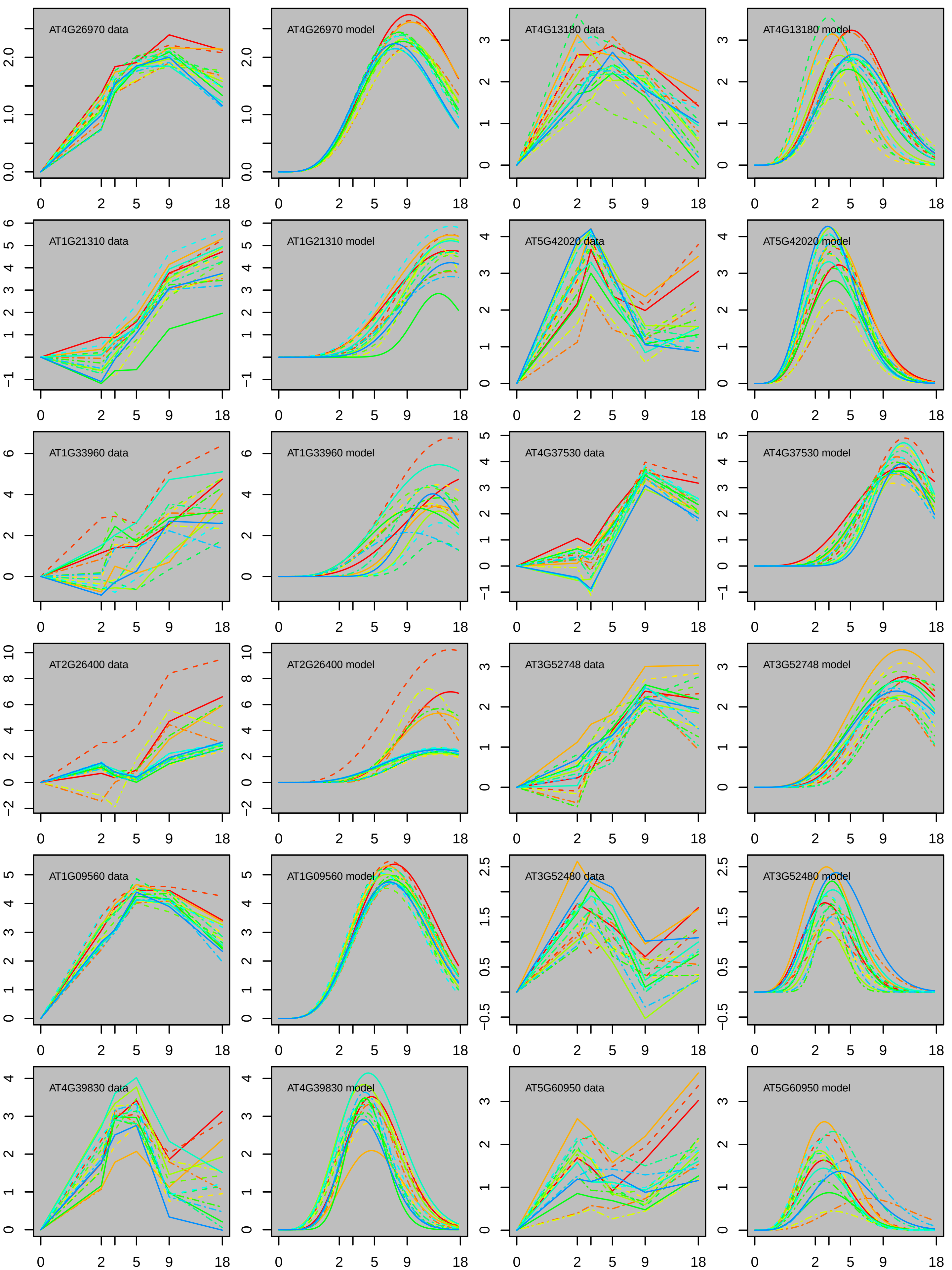
AT4G26970 data
AT4G26970 model
AT4G13180 data
AT4G13180 model
AT1G21310 data
AT1G21310 model
AT5G42020 data
AT5G42020 model
AT1G33960 data
AT1G33960 model
AT4G37530 data
AT4G37530 model
AT2G26400 data
AT2G26400 model
AT3G52748 data
AT3G52748 model
AT1G09560 data
AT1G09560 model
AT3G52480 data
AT3G52480 model
AT4G39830 data
AT4G39830 model
AT5G60950 data
AT5G60950 model
0 2 5 9 18

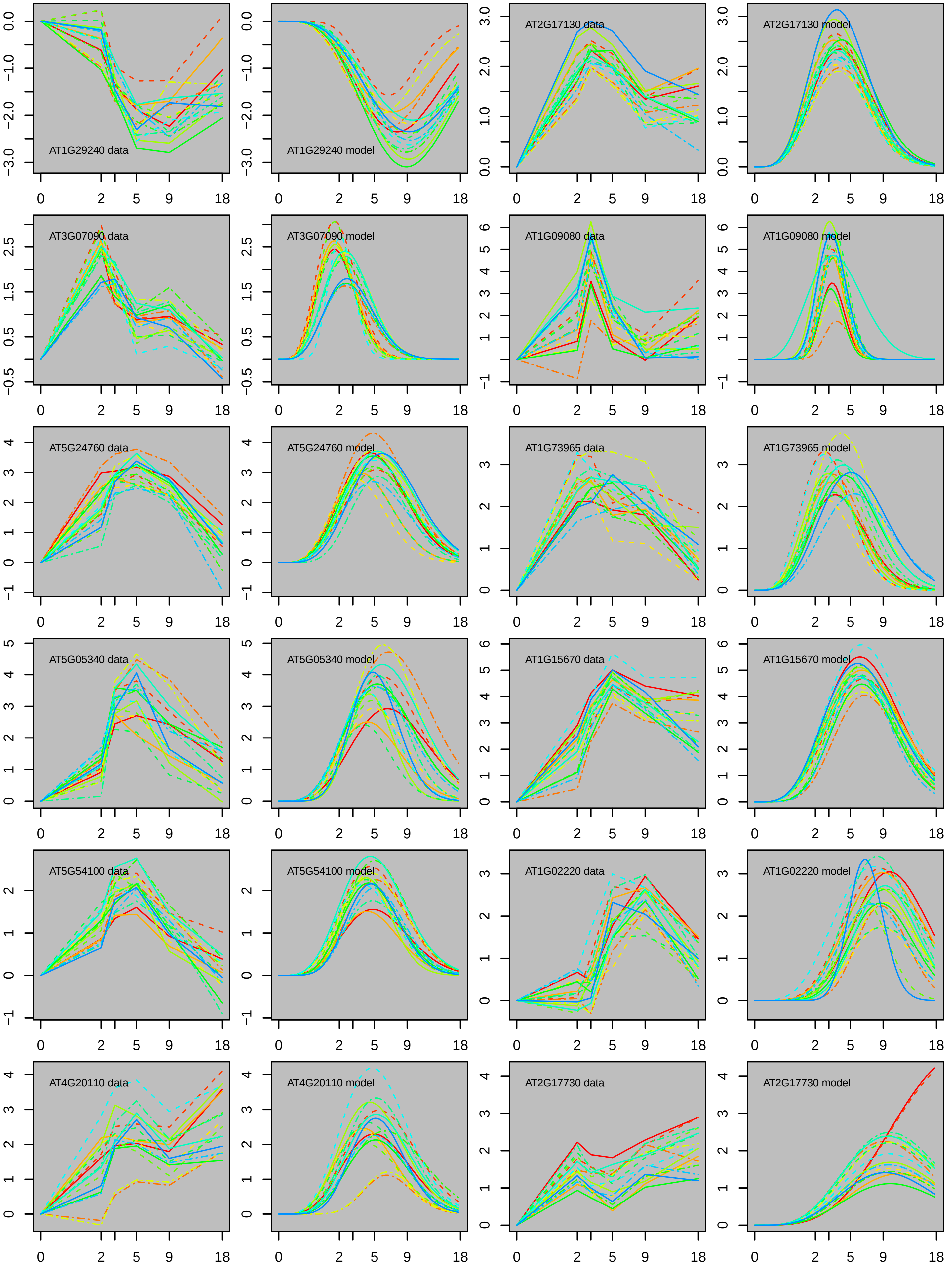
AT1G29240 data
AT1G29240 model
AT2G17130 data
AT2G17130 model
AT3G07090 data
AT3G07090 model
AT1G09080 data
AT1G09080 model
AT5G24760 data
AT5G24760 model
AT1G73965 data
AT1G73965 model
AT5G05340 data
AT5G05340 model
AT1G15670 data
AT1G15670 model
AT5G54100 data
AT5G54100 model
AT1G02220 data
AT1G02220 model
AT4G20110 data
AT4G20110 model
AT2G17730 data
AT2G17730 model

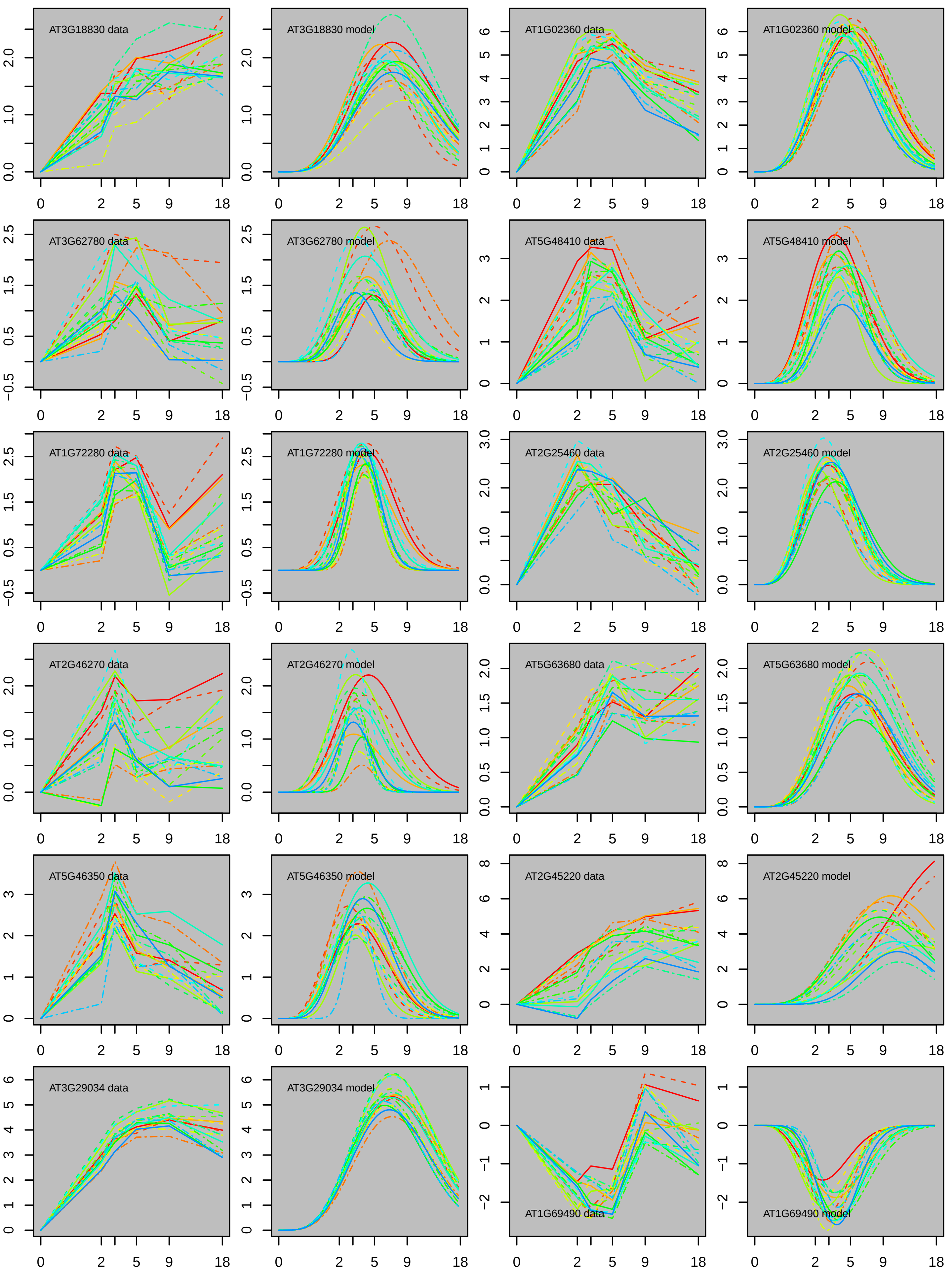
AT3G18830 data
AT3G18830 model
AT1G02360 data
AT1G02360 model
AT3G62780 data
AT3G62780 model
AT5G48410 data
AT5G48410 model
AT1G72280 data
AT1G72280 model
AT2G25460 data
AT2G25460 model
AT2G46270 data
AT2G46270 model
AT5G63680 data
AT5G63680 model
AT5G46350 data
AT5G46350 model
AT2G45220 data
AT2G45220 model
AT3G29034 data
AT3G29034 model
AT1G69490 data
AT1G69490 model
0
2
5
9
18

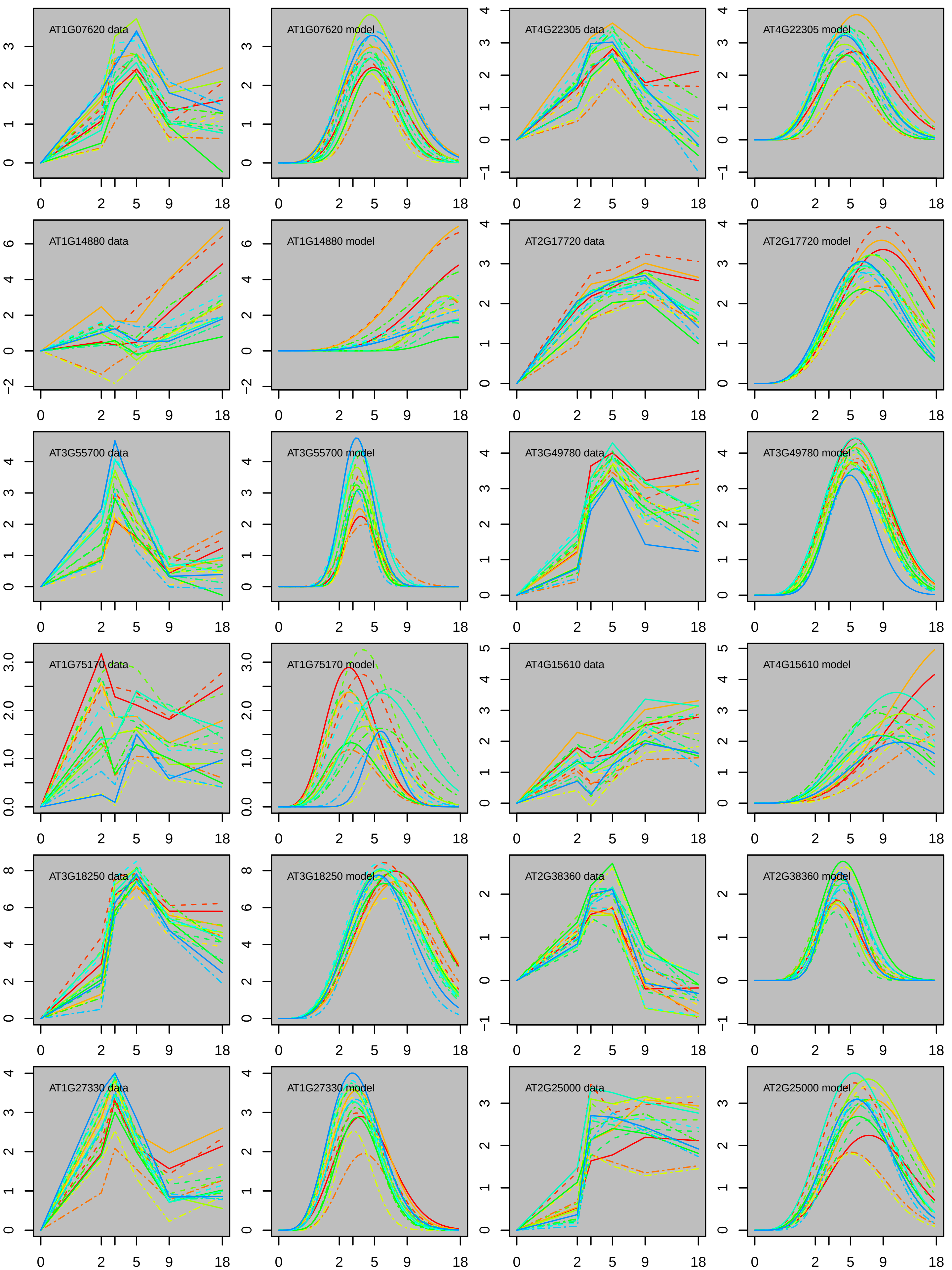
AT1G07620 data
AT1G07620 model
AT4G22305 data
AT4G22305 model
AT1G14880 data
AT1G14880 model
AT2G17720 data
AT2G17720 model
AT3G55700 data
AT3G55700 model
AT3G49780 data
AT3G49780 model
AT1G75170 data
AT1G75170 model
AT4G15610 data
AT4G15610 model
AT3G18250 data
AT3G18250 model
AT2G38360 data
AT2G38360 model
AT1G27330 data
AT1G27330 model
AT2G25000 data
AT2G25000 model

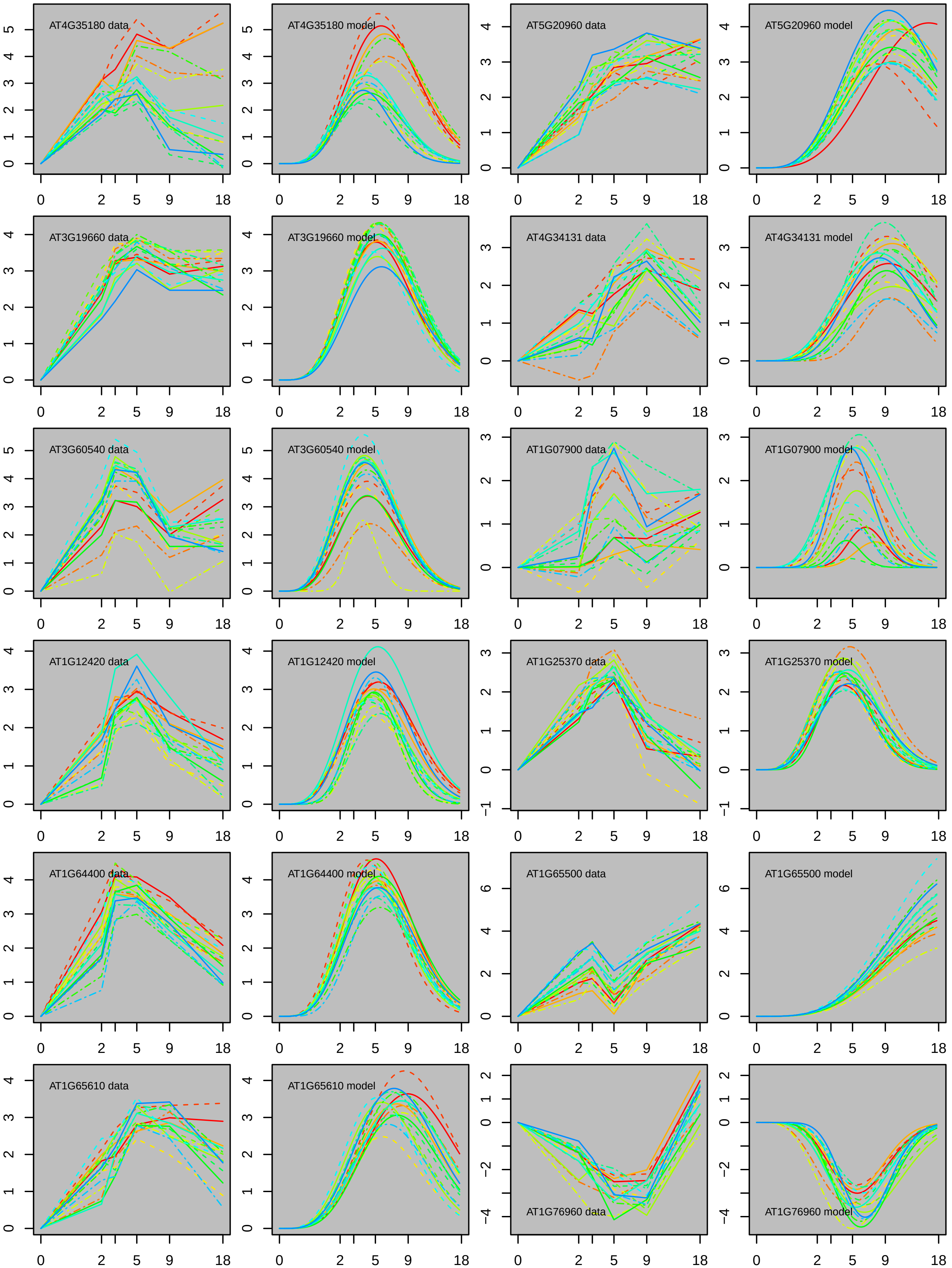
AT4G35180 data
AT4G35180 model
AT5G20960 data
AT5G20960 model
AT3G19660 data
AT3G19660 model
AT4G34131 data
AT4G34131 model
AT3G60540 data
AT3G60540 model
AT1G07900 data
AT1G07900 model
AT1G12420 data
AT1G12420 model
AT1G25370 data
AT1G25370 model
AT1G64400 data
AT1G64400 model
AT1G65500 data
AT1G65500 model
AT1G65610 data
AT1G65610 model
AT1G76960 data
AT1G76960 model
0
2
5
9
18

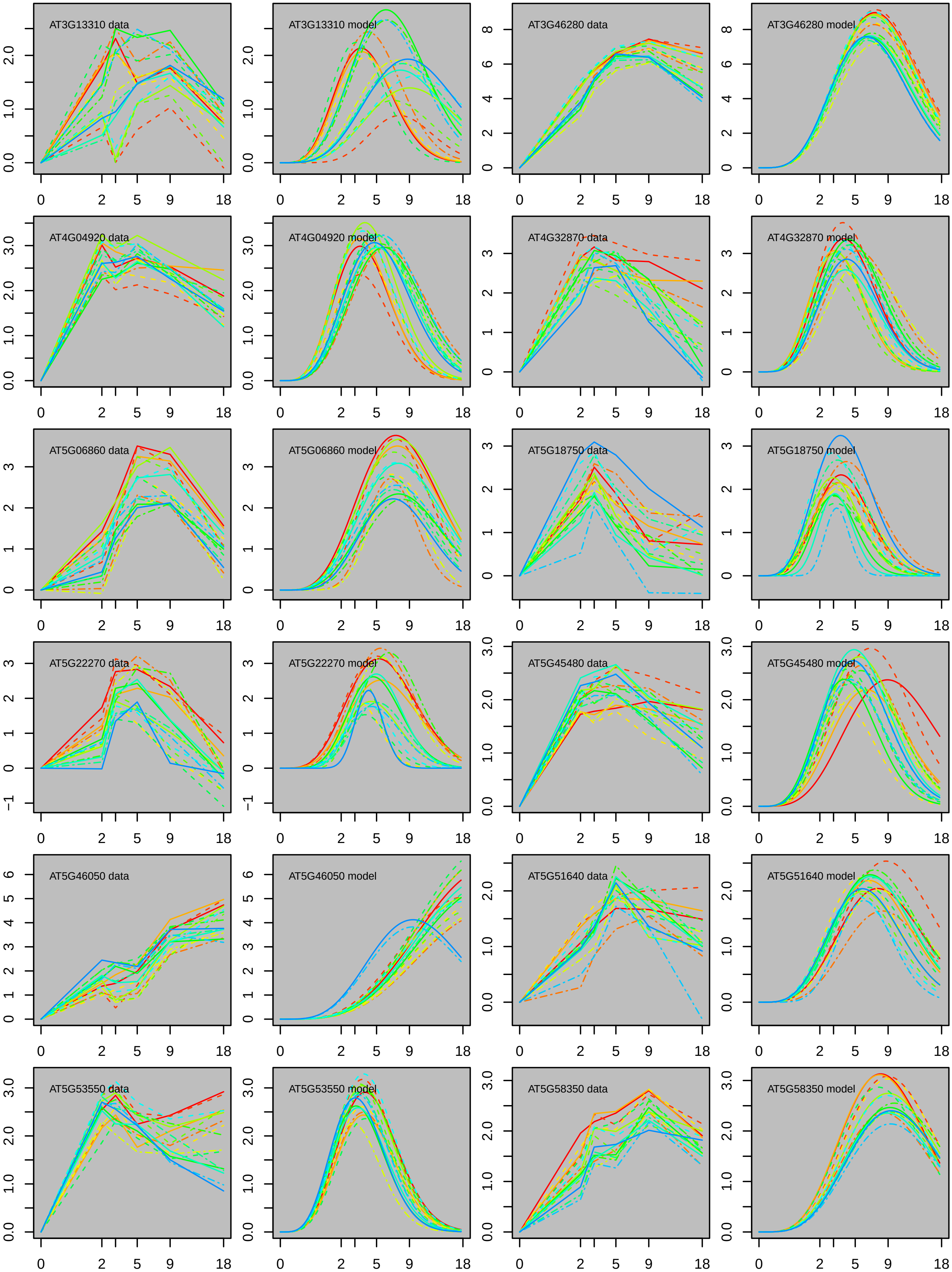
AT3G13310 data
AT3G13310 model
AT3G46280 data
AT3G46280 model
AT4G04920 data
AT4G04920 model
AT4G32870 data
AT4G32870 model
AT5G06860 data
AT5G06860 model
AT5G18750 data
AT5G18750 model
AT5G22270 data
AT5G22270 model
AT5G45480 data
AT5G45480 model
AT5G46050 data
AT5G46050 model
AT5G51640 data
AT5G51640 model
AT5G53550 data
AT5G53550 model
AT5G58350 data
AT5G58350 model
0 2 5 9 18

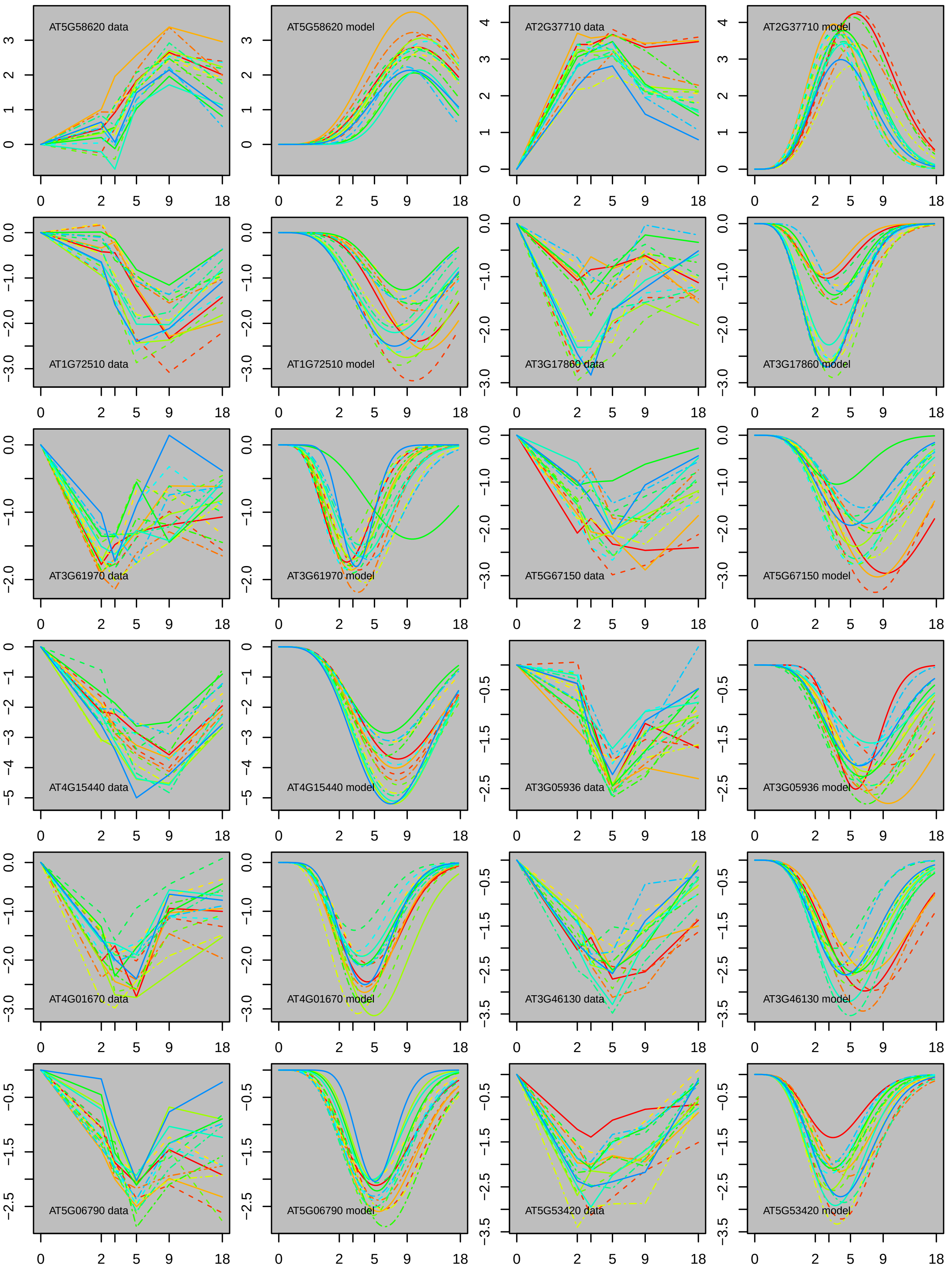
AT5G58620 data
AT5G58620 model
AT2G37710 data
AT2G37710 model
AT1G72510 data
AT1G72510 model
AT3G17860 data
AT3G17860 model
AT3G61970 data
AT3G61970 model
AT5G67150 data
AT5G67150 model
AT4G15440 data
AT4G15440 model
AT3G05936 data
AT3G05936 model
AT4G01670 data
AT4G01670 model
AT3G46130 data
AT3G46130 model
AT5G06790 data
AT5G06790 model
AT5G53420 data
AT5G53420 model

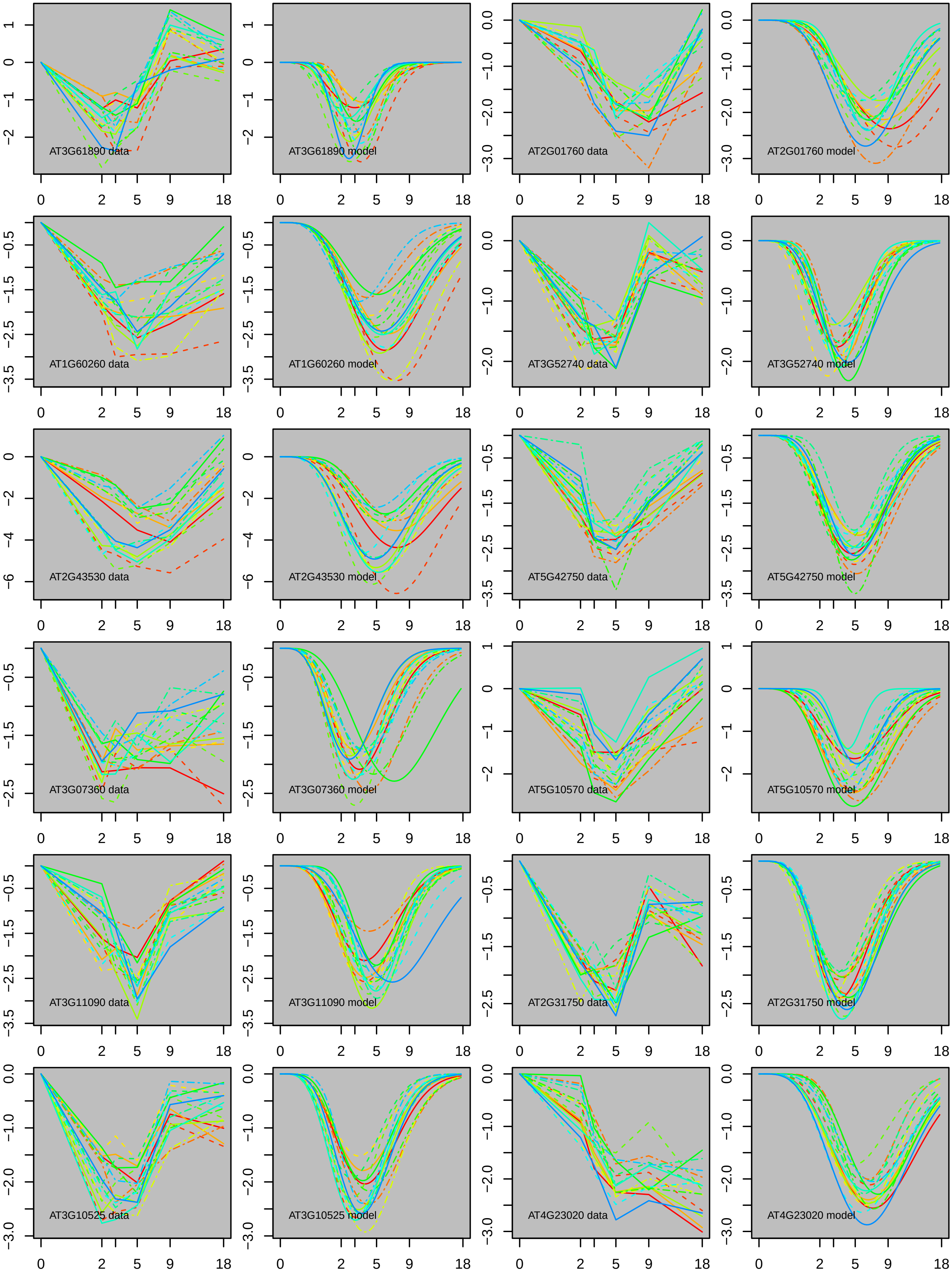

AT3G61890 data
AT3G61890 model
AT2G01760 data
AT2G01760 model
AT1G60260 data
AT1G60260 model
AT3G52740 data
AT3G52740 model
AT2G43530 data
AT2G43530 model
AT5G42750 data
AT5G42750 model
AT3G07360 data
AT3G07360 model
AT5G10570 data
AT5G10570 model
AT3G11090 data
AT3G11090 model
AT2G31750 data
AT2G31750 model
AT3G10525 data
AT3G10525 model
AT4G23020 data
AT4G23020 model
0
2
5
9
18

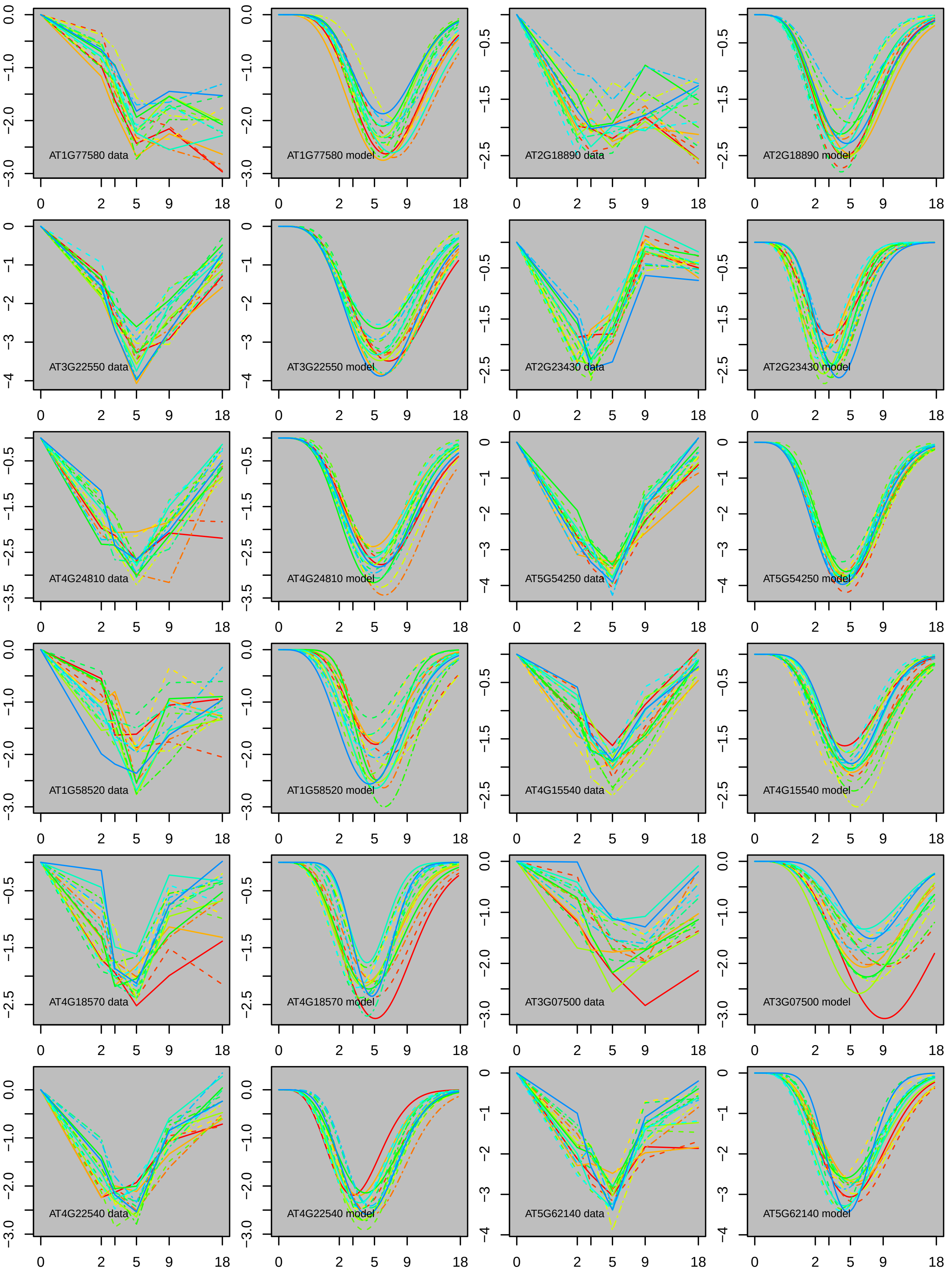

AT1G77580 data
AT1G77580 model
AT2G18890 data
AT2G18890 model
AT3G22550 data
AT3G22550 model
AT2G23430 data
AT2G23430 model
AT4G24810 data
AT4G24810 model
AT5G54250 data
AT5G54250 model
AT1G58520 data
AT1G58520 model
AT4G15540 data
AT4G15540 model
AT4G18570 data
AT4G18570 model
AT3G07500 data
AT3G07500 model
AT4G22540 data
AT4G22540 model
AT5G62140 data
AT5G62140 model

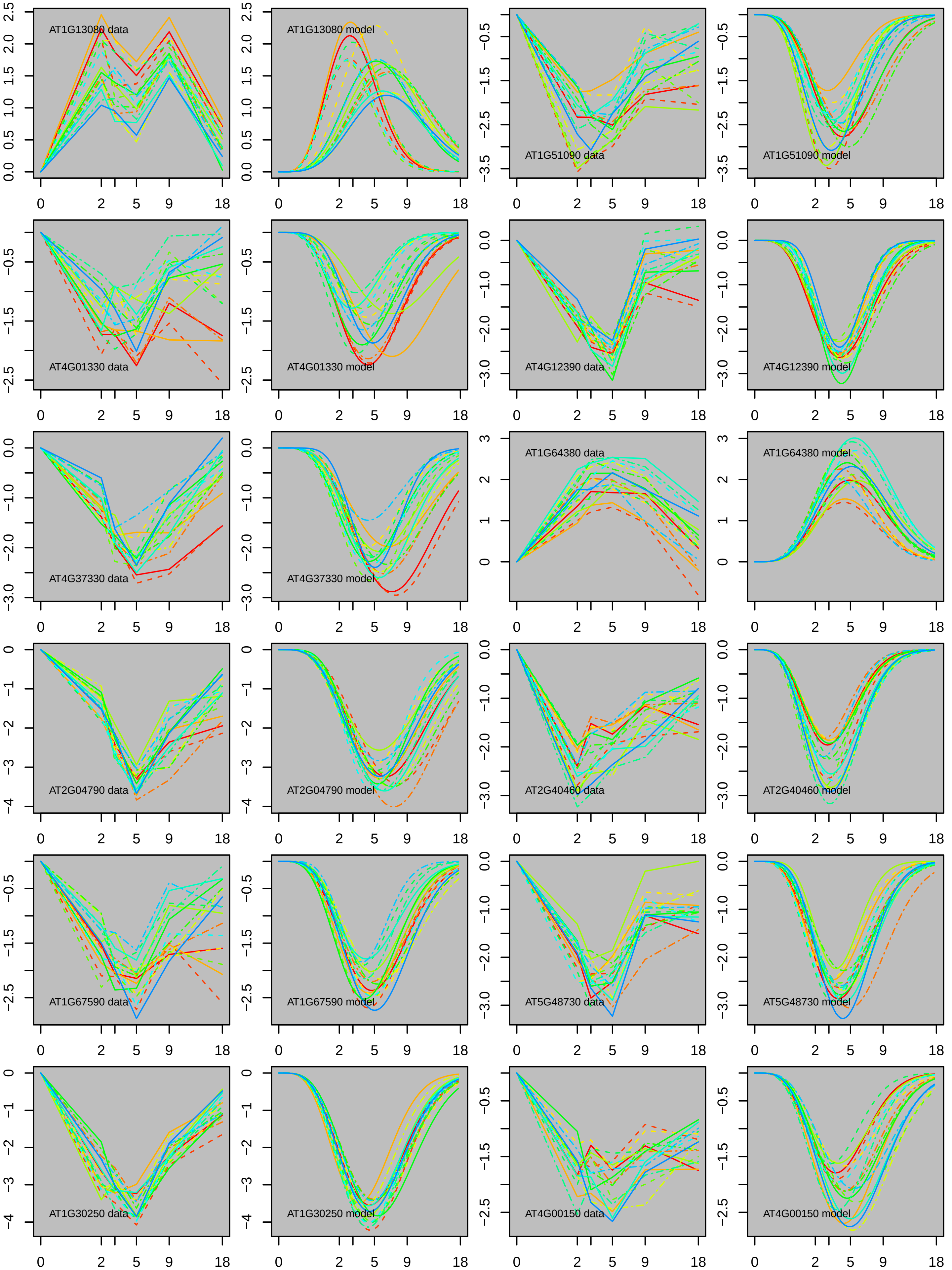

AT1G13080 data
AT1G13080 model
AT1G51090 data
AT1G51090 model
AT4G01330 data
AT4G01330 model
AT4G12390 data
AT4G12390 model
AT4G37330 data
AT4G37330 model
AT1G64380 data
AT1G64380 model
AT2G04790 data
AT2G04790 model
AT2G40460 data
AT2G40460 model
AT1G67590 data
AT1G67590 model
AT5G48730 data
AT5G48730 model
AT1G30250 data
AT1G30250 model
AT4G00150 data
AT4G00150 model

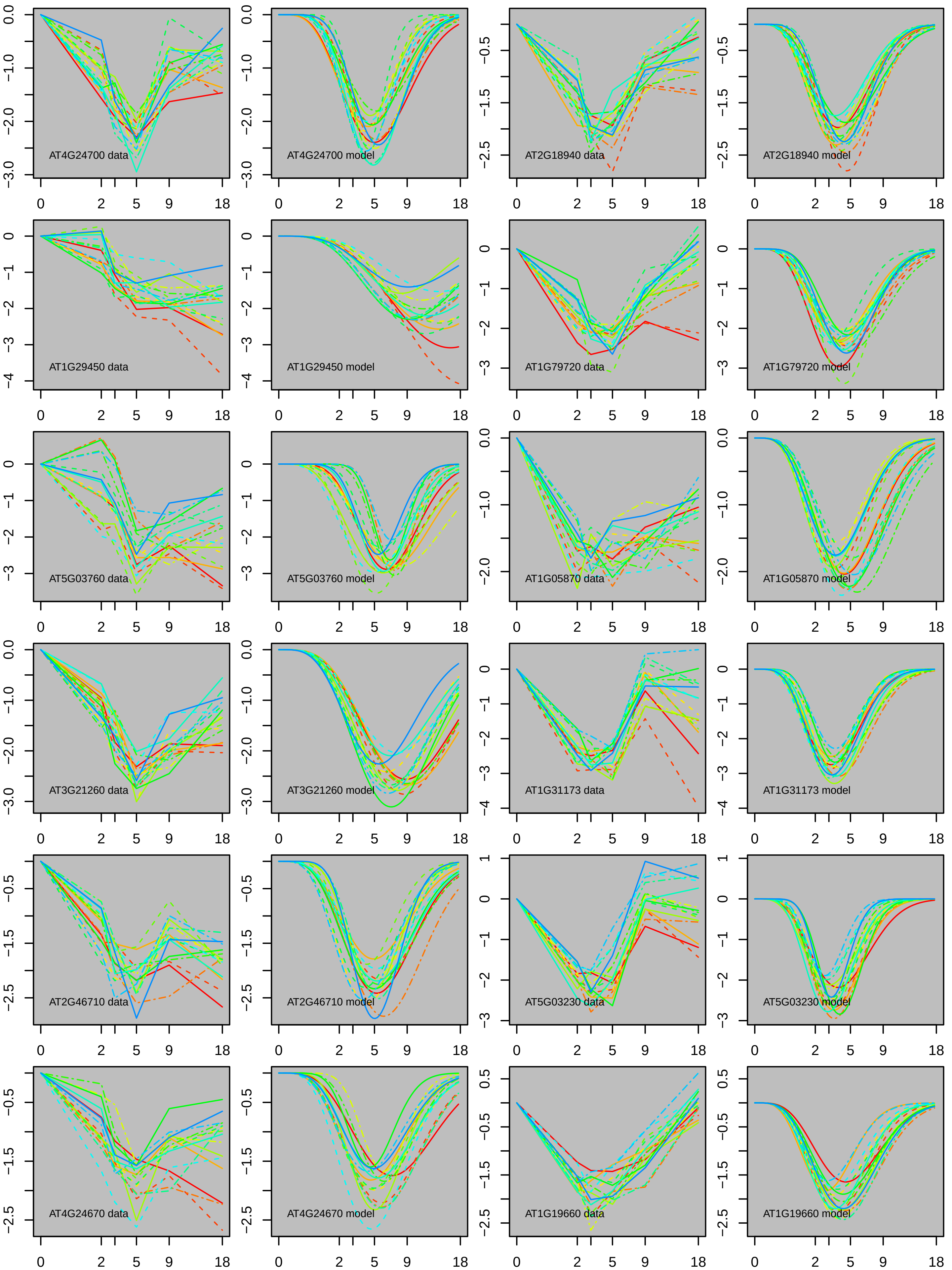
AT4G24700 data
AT4G24700 model
AT2G18940 data
AT2G18940 model
AT1G29450 data
AT1G29450 model
AT1G79720 data
AT1G79720 model
AT5G03760 data
AT5G03760 model
AT1G05870 data
AT1G05870 model
AT3G21260 data
AT3G21260 model
AT1G31173 data
AT1G31173 model
AT2G46710 data
AT2G46710 model
AT5G03230 data
AT5G03230 model
AT4G24670 data
AT4G24670 model
AT1G19660 data
AT1G19660 model

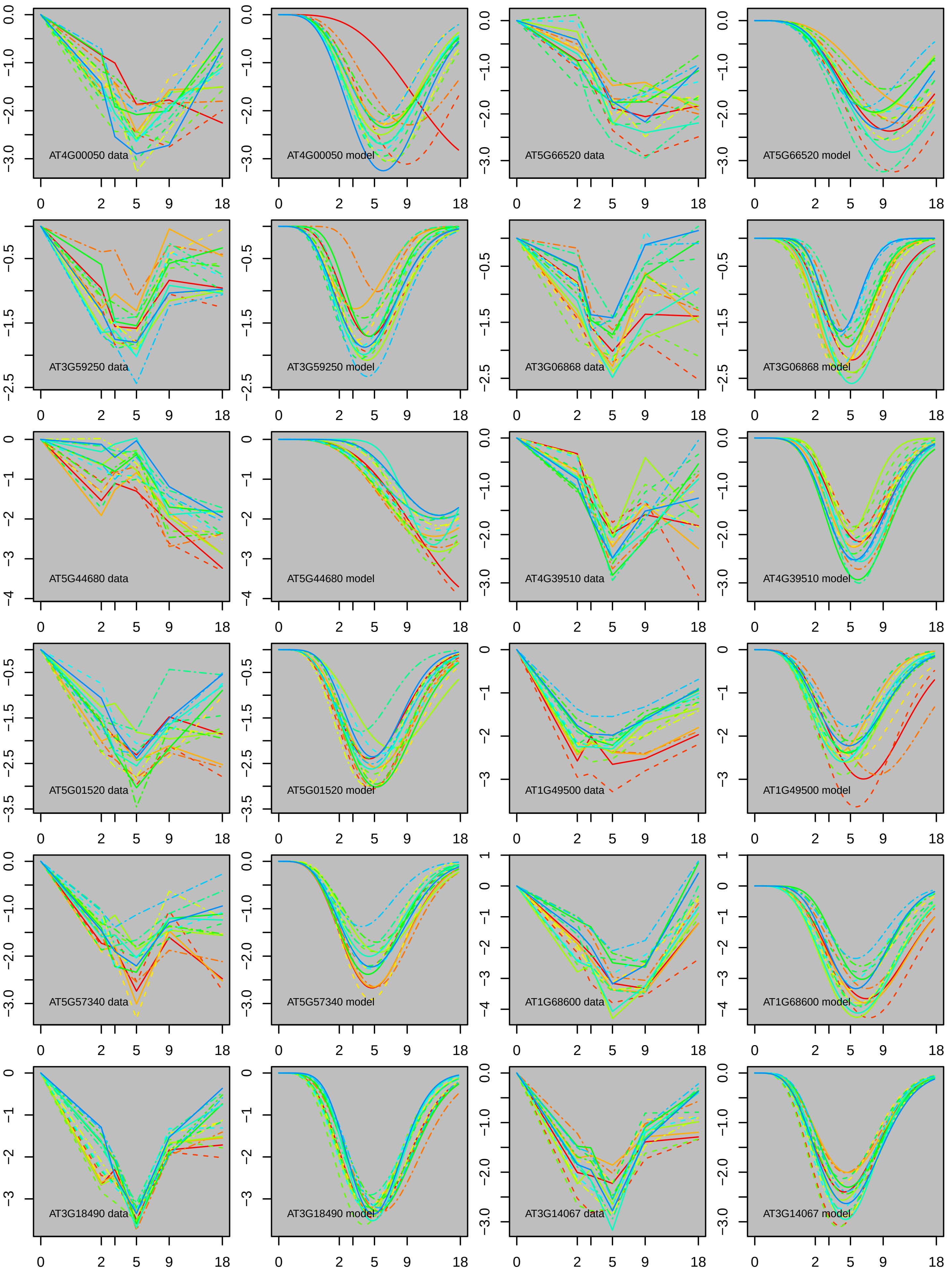
AT4G00050 data
AT4G00050 model
AT5G66520 data
AT5G66520 model
AT3G59250 data
AT3G59250 model
AT3G06868 data
AT3G06868 model
AT5G44680 data
AT5G44680 model
AT4G39510 data
AT4G39510 model
AT5G01520 data
AT5G01520 model
AT1G49500 data
AT1G49500 model
AT5G57340 data
AT5G57340 model
AT1G68600 data
AT1G68600 model
AT3G18490 data
AT3G18490 model
AT3G14067 data
AT3G14067 model

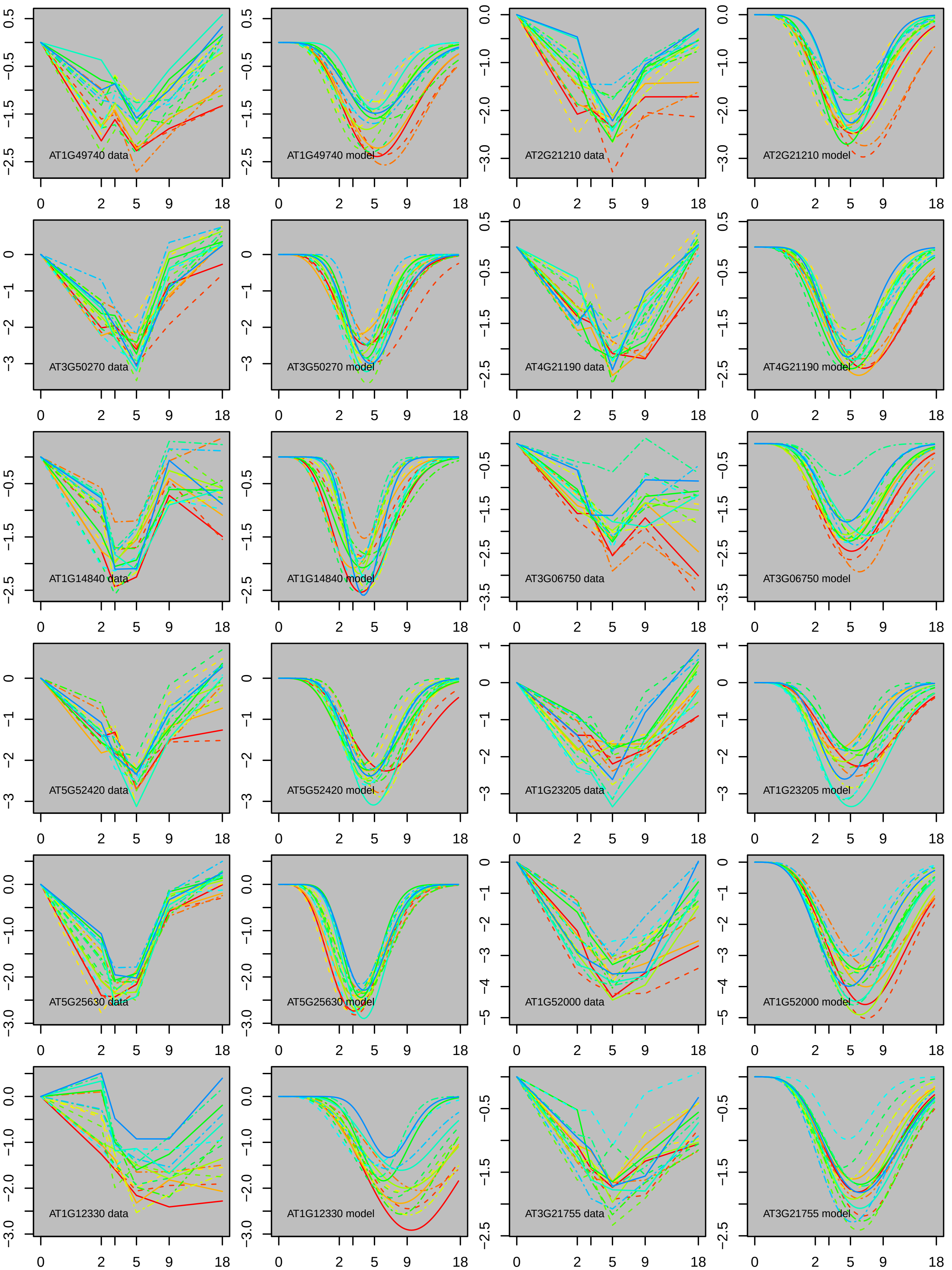
AT1G49740 data
AT1G49740 model
AT2G21210 data
AT2G21210 model
AT3G50270 data
AT3G50270 model
AT4G21190 data
AT4G21190 model
AT1G14840 data
AT1G14840 model
AT3G06750 data
AT3G06750 model
AT5G52420 data
AT5G52420 model
AT1G23205 data
AT1G23205 model
AT5G25630 data
AT5G25630 model
AT1G52000 data
AT1G52000 model
AT1G12330 data
AT1G12330 model
AT3G21755 data
AT3G21755 model
0
2
5
9
18

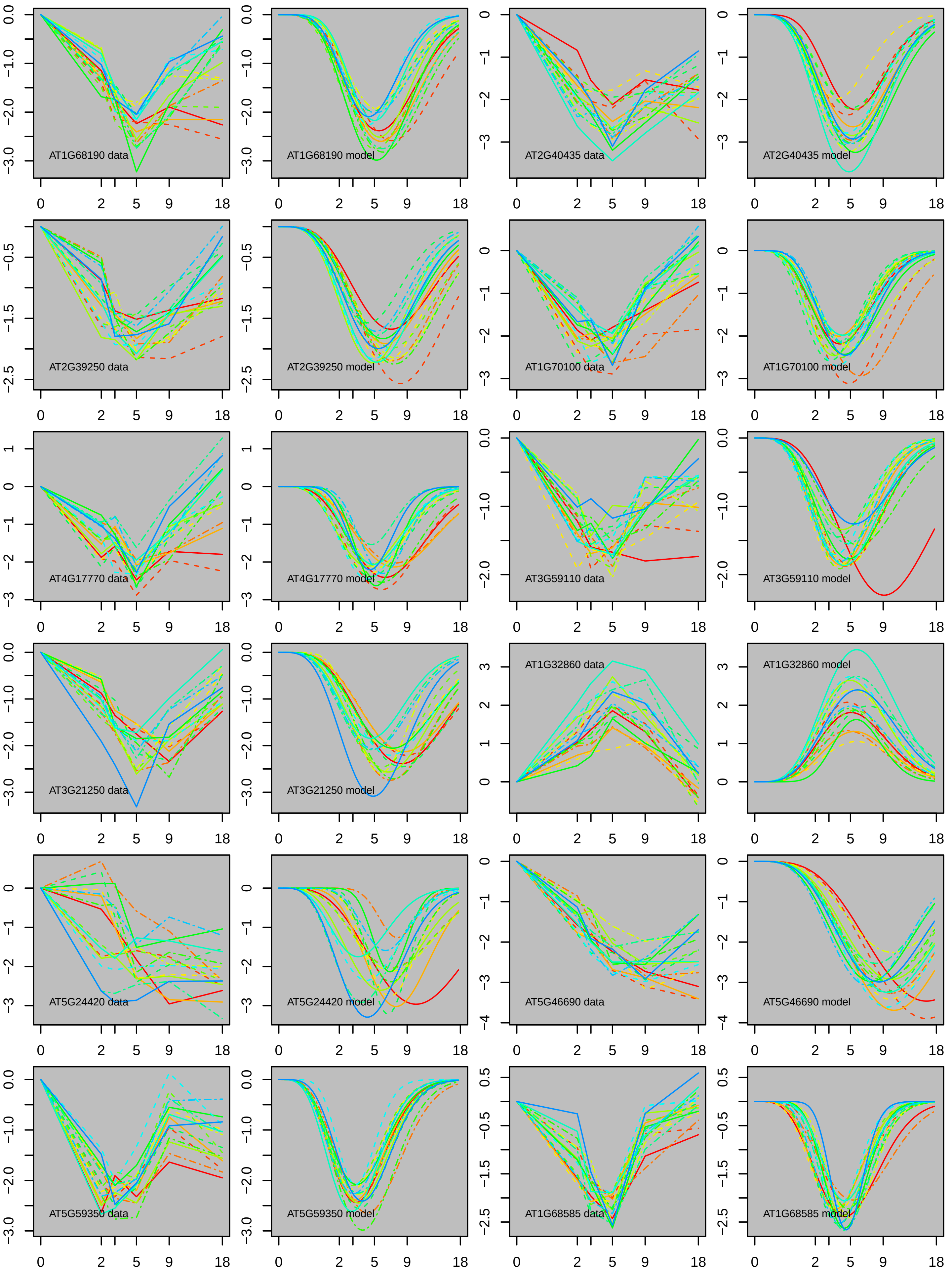

AT1G68190 data
AT1G68190 model
AT2G40435 data
AT2G40435 model
AT2G39250 data
AT2G39250 model
AT1G70100 data
AT1G70100 model
AT4G17770 data
AT4G17770 model
AT3G59110 data
AT3G59110 model
AT3G21250 data
AT3G21250 model
AT1G32860 data
AT1G32860 model
AT5G24420 data
AT5G24420 model
AT5G46690 data
AT5G46690 model
AT5G59350 data
AT5G59350 model
AT1G68585 data
AT1G68585 model

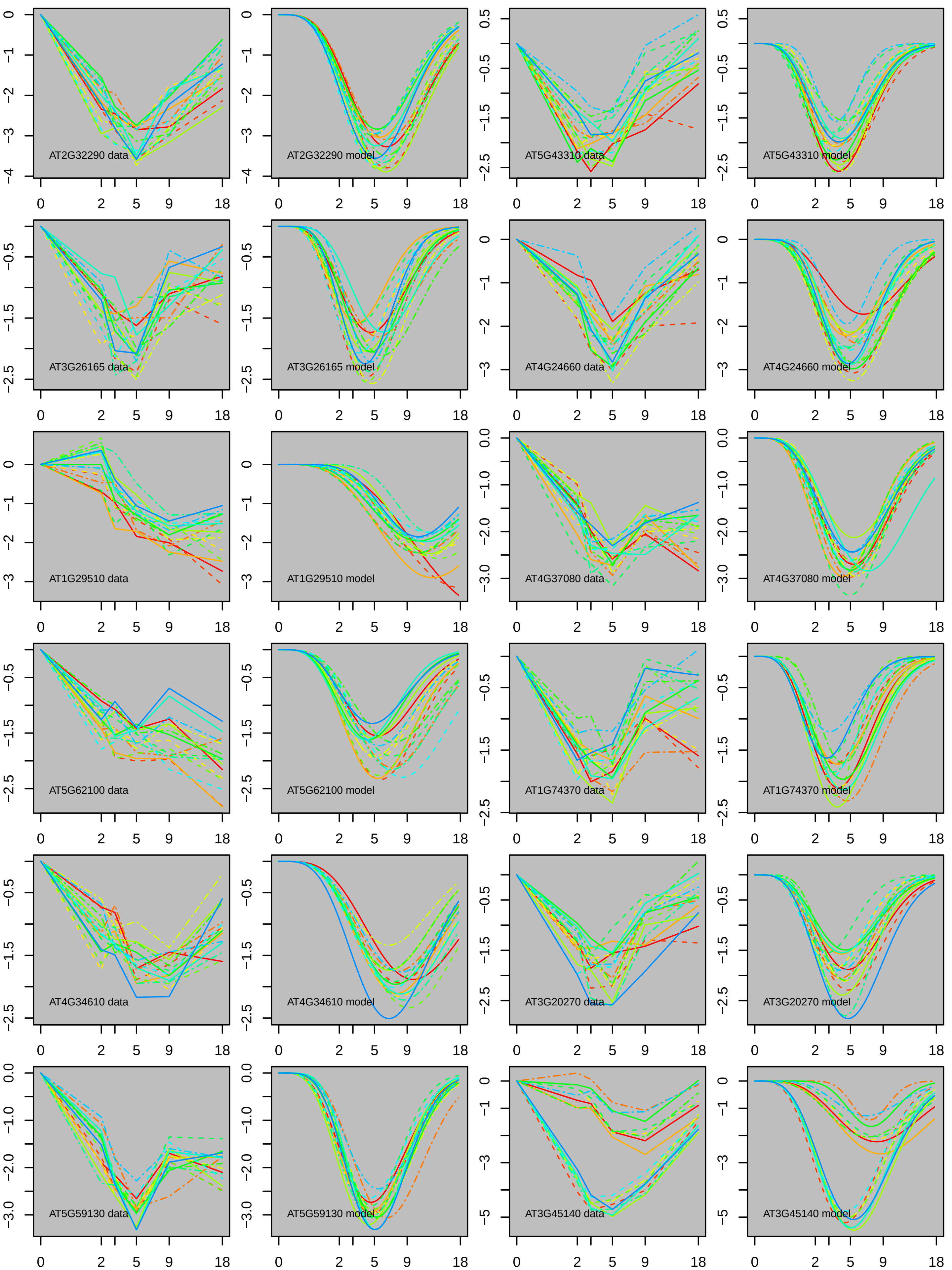
AT2G32290 data
AT2G32290 model
AT5G43310 data
AT5G43310 model
AT3G26165 data
AT3G26165 model
AT4G24660 data
AT4G24660 model
AT1G29510 data
AT1G29510 model
AT4G37080 data
AT4G37080 model
AT5G62100 data
AT5G62100 model
AT1G74370 data
AT1G74370 model
AT4G34610 data
AT4G34610 model
AT3G20270 data
AT3G20270 model
AT5G59130 data
AT5G59130 model
AT3G45140 data
AT3G45140 model

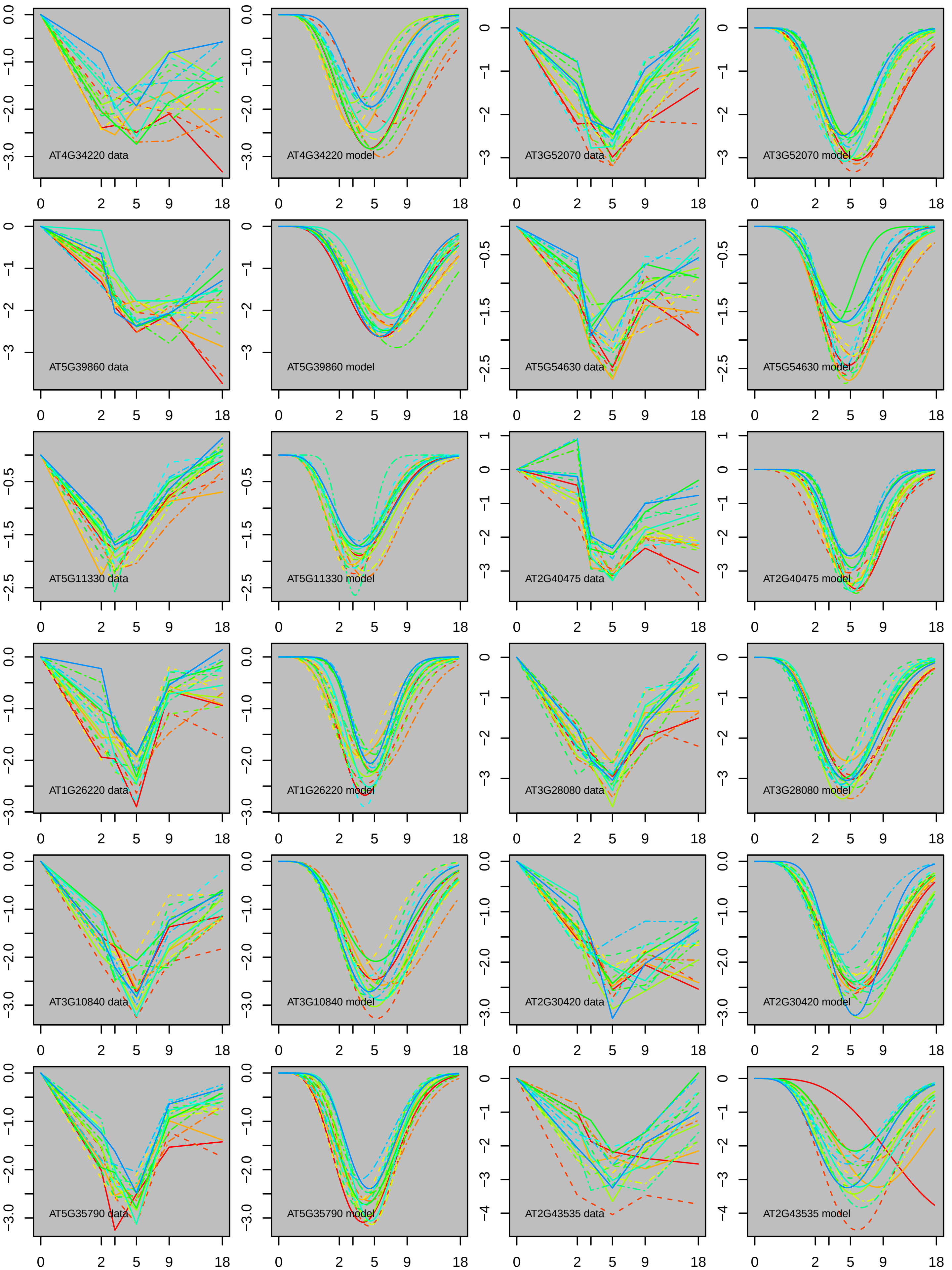

AT4G34220 data
AT4G34220 model
AT3G52070 data
AT3G52070 model
AT5G39860 data
AT5G39860 model
AT5G54630 data
AT5G54630 model
AT5G11330 data
AT5G11330 model
AT2G40475 data
AT2G40475 model
AT1G26220 data
AT1G26220 model
AT3G28080 data
AT3G28080 model
AT3G10840 data
AT3G10840 model
AT2G30420 data
AT2G30420 model
AT5G35790 data
AT5G35790 model
AT2G43535 data
AT2G43535 model
0 2 5 9 18

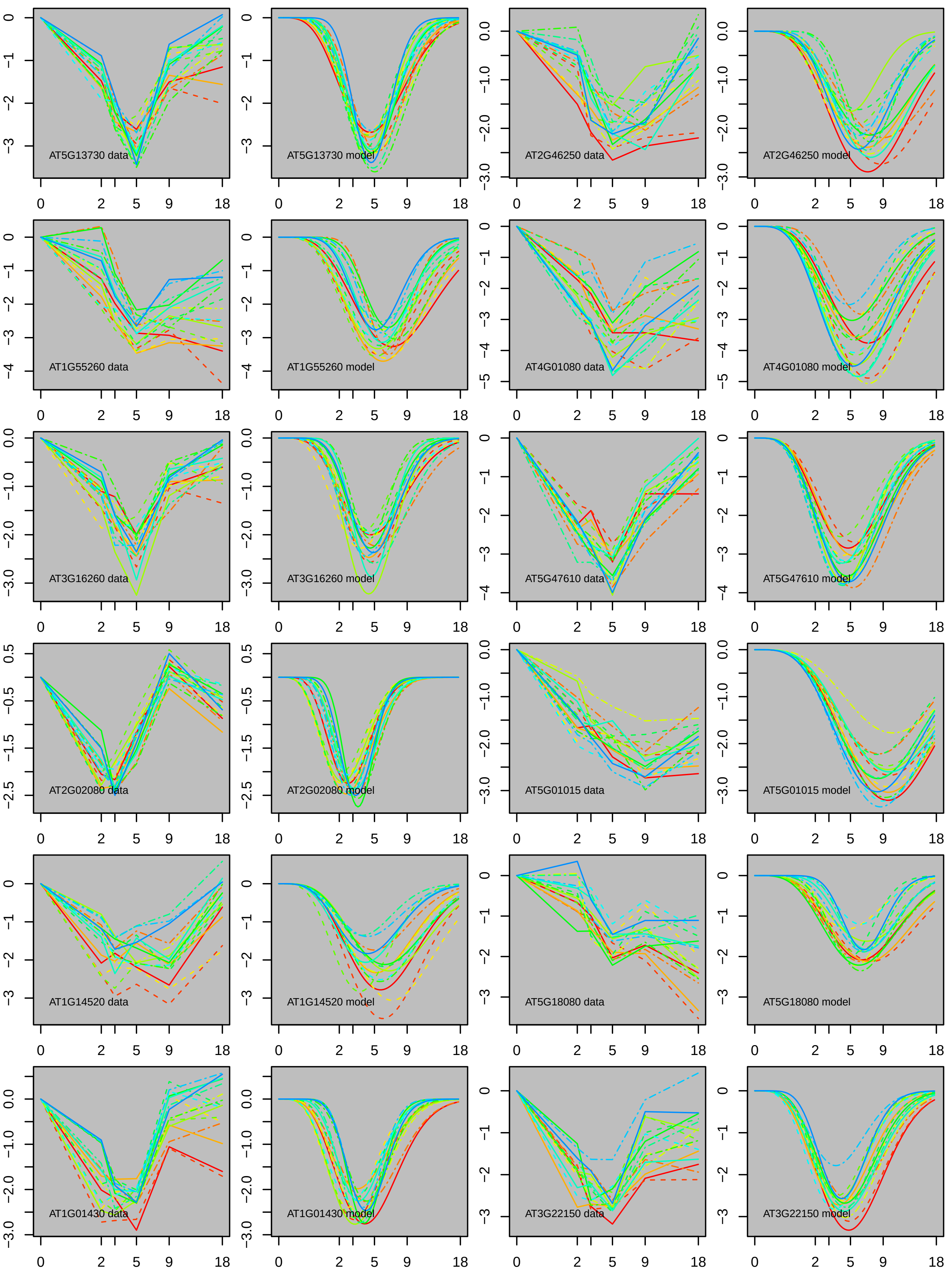
AT5G13730 data
AT5G13730 model
AT2G46250 data
AT2G46250 model
AT1G55260 data
AT1G55260 model
AT4G01080 data
AT4G01080 model
AT3G16260 data
AT3G16260 model
AT5G47610 data
AT5G47610 model
AT2G02080 data
AT2G02080 model
AT5G01015 data
AT5G01015 model
AT1G14520 data
AT1G14520 model
AT5G18080 data
AT5G18080 model
AT1G01430 data
AT1G01430 model
AT3G22150 data
AT3G22150 model
0
2
5
9
18

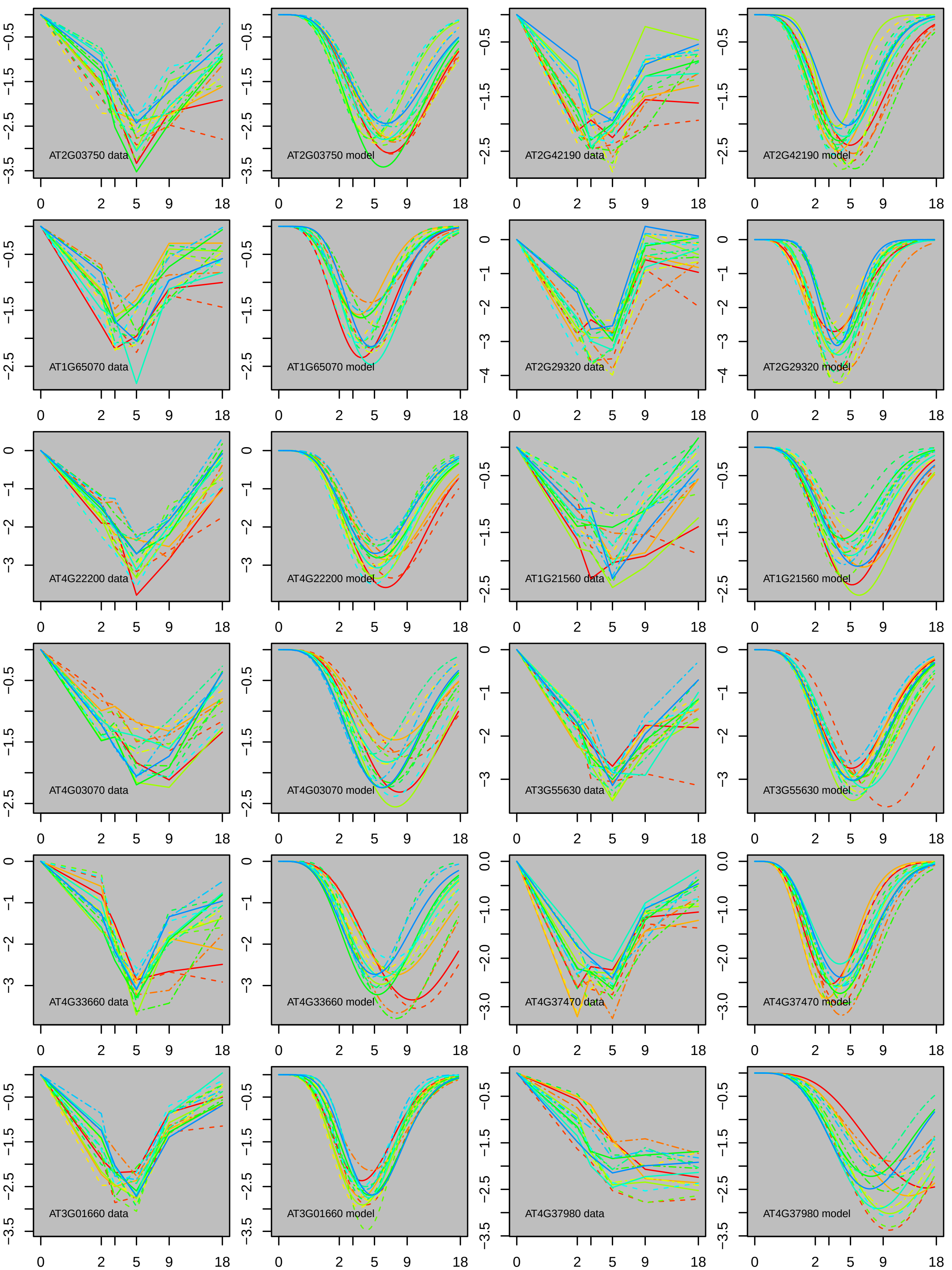
AT2G03750 data
AT2G03750 model
AT2G42190 data
AT2G42190 model
AT1G65070 data
AT1G65070 model
AT2G29320 data
AT2G29320 model
AT4G22200 data
AT4G22200 model
AT1G21560 data
AT1G21560 model
AT4G03070 data
AT4G03070 model
AT3G55630 data
AT3G55630 model
AT4G33660 data
AT4G33660 model
AT4G37470 data
AT4G37470 model
AT3G01660 data
AT3G01660 model
AT4G37980 data
AT4G37980 model
0
2
5
9
18

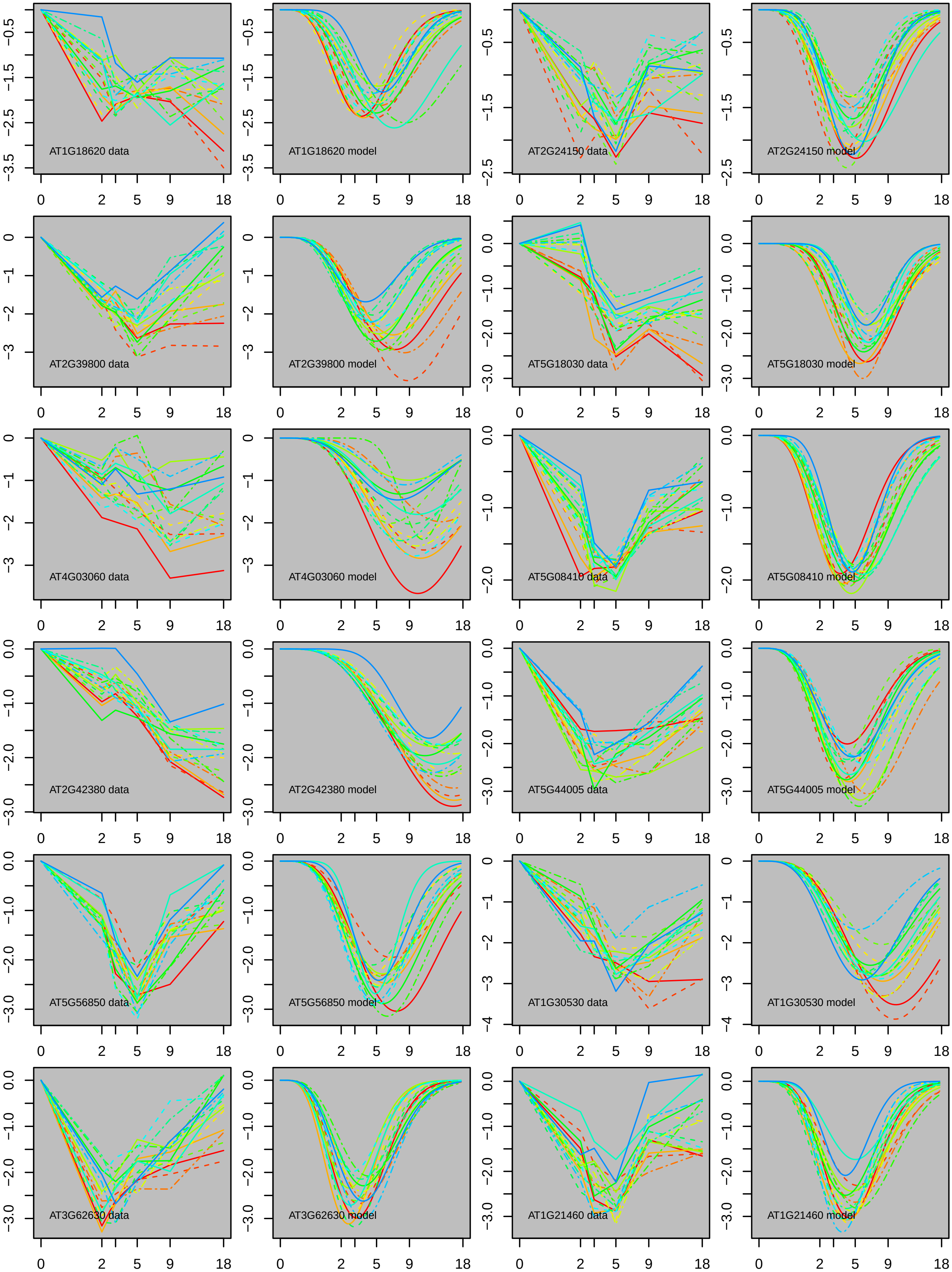

AT1G18620 data
AT1G18620 model
AT2G24150 data
AT2G24150 model
AT2G39800 data
AT2G39800 model
AT5G18030 data
AT5G18030 model
AT4G03060 data
AT4G03060 model
AT5G08410 data
AT5G08410 model
AT2G42380 data
AT2G42380 model
AT5G44005 data
AT5G44005 model
AT5G56850 data
AT5G56850 model
AT1G30530 data
AT1G30530 model
AT3G62630 data
AT3G62630 model
AT1G21460 data
AT1G21460 model
0
2
5
9
18

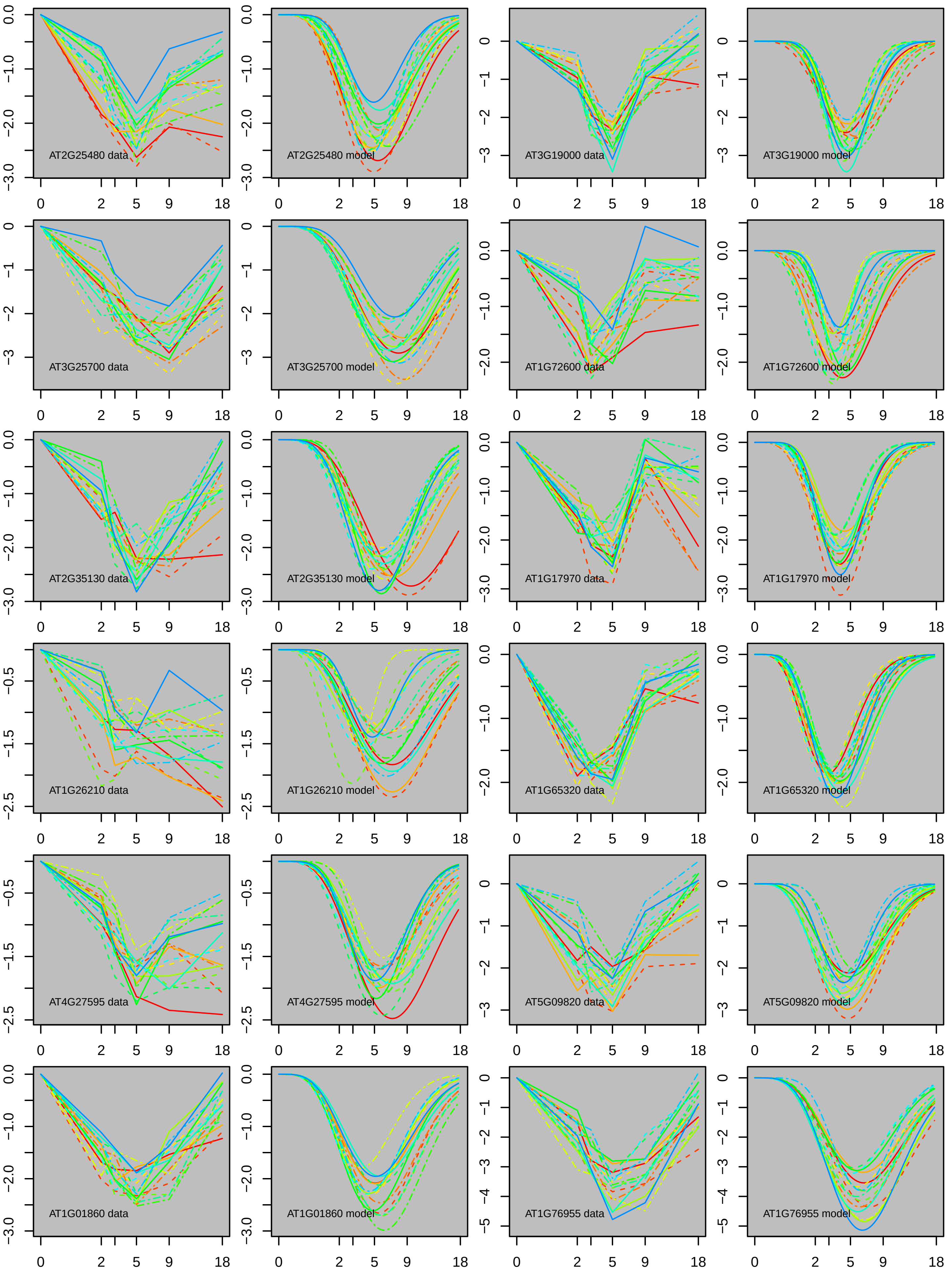
AT2G25480 data
AT2G25480 model
AT3G19000 data
AT3G19000 model
AT3G25700 data
AT3G25700 model
AT1G72600 data
AT1G72600 model
AT2G35130 data
AT2G35130 model
AT1G17970 data
AT1G17970 model
AT1G26210 data
AT1G26210 model
AT1G65320 data
AT1G65320 model
AT4G27595 data
AT4G27595 model
AT5G09820 data
AT5G09820 model
AT1G01860 data
AT1G01860 model
AT1G76955 data
AT1G76955 model

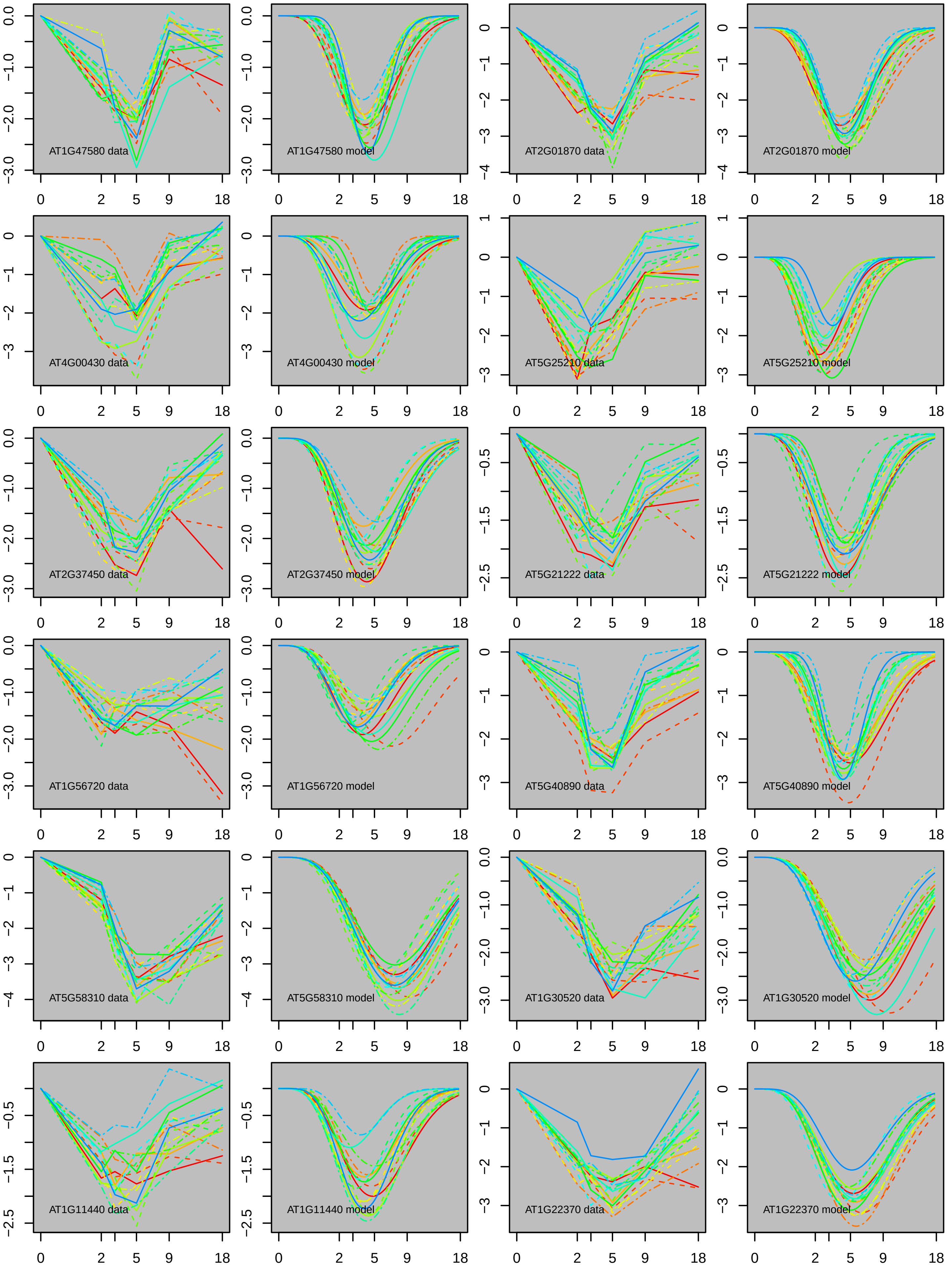
AT1G47580 data
AT1G47580 model
AT2G01870 data
AT2G01870 model
AT4G00430 data
AT4G00430 model
AT5G25210 data
AT5G25210 model
AT2G37450 data
AT2G37450 model
AT5G21222 data
AT5G21222 model
AT1G56720 data
AT1G56720 model
AT5G40890 data
AT5G40890 model
AT5G58310 data
AT5G58310 model
AT1G30520 data
AT1G30520 model
AT1G11440 data
AT1G11440 model
AT1G22370 data
AT1G22370 model
0 2 5 9 18

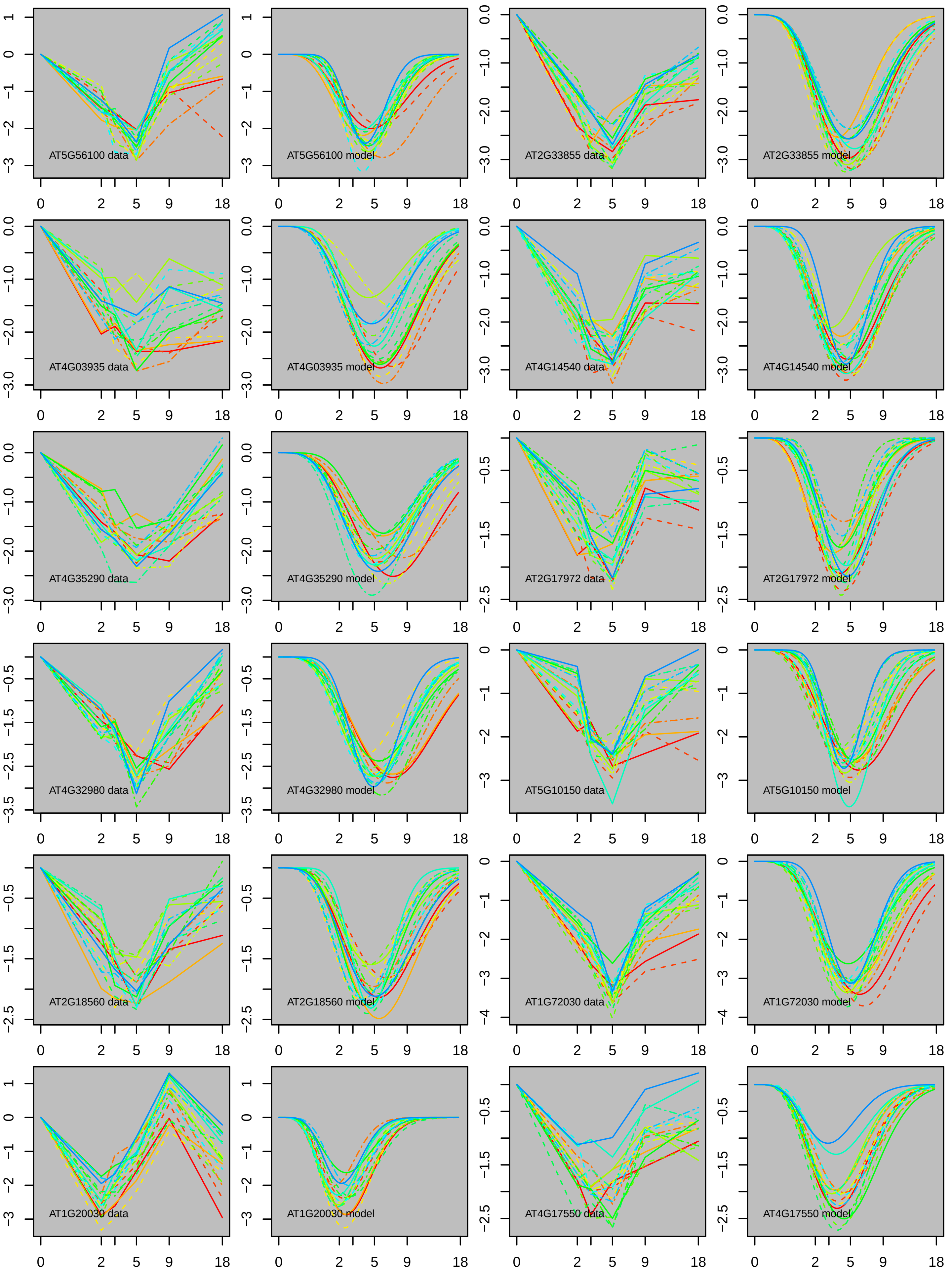

AT5G56100 data
AT5G56100 model
AT2G33855 data
AT2G33855 model
AT4G03935 data
AT4G03935 model
AT4G14540 data
AT4G14540 model
AT4G35290 data
AT4G35290 model
AT2G17972 data
AT2G17972 model
AT4G32980 data
AT4G32980 model
AT5G10150 data
AT5G10150 model
AT2G18560 data
AT2G18560 model
AT1G72030 data
AT1G72030 model
AT1G20030 data
AT1G20030 model
AT4G17550 data
AT4G17550 model

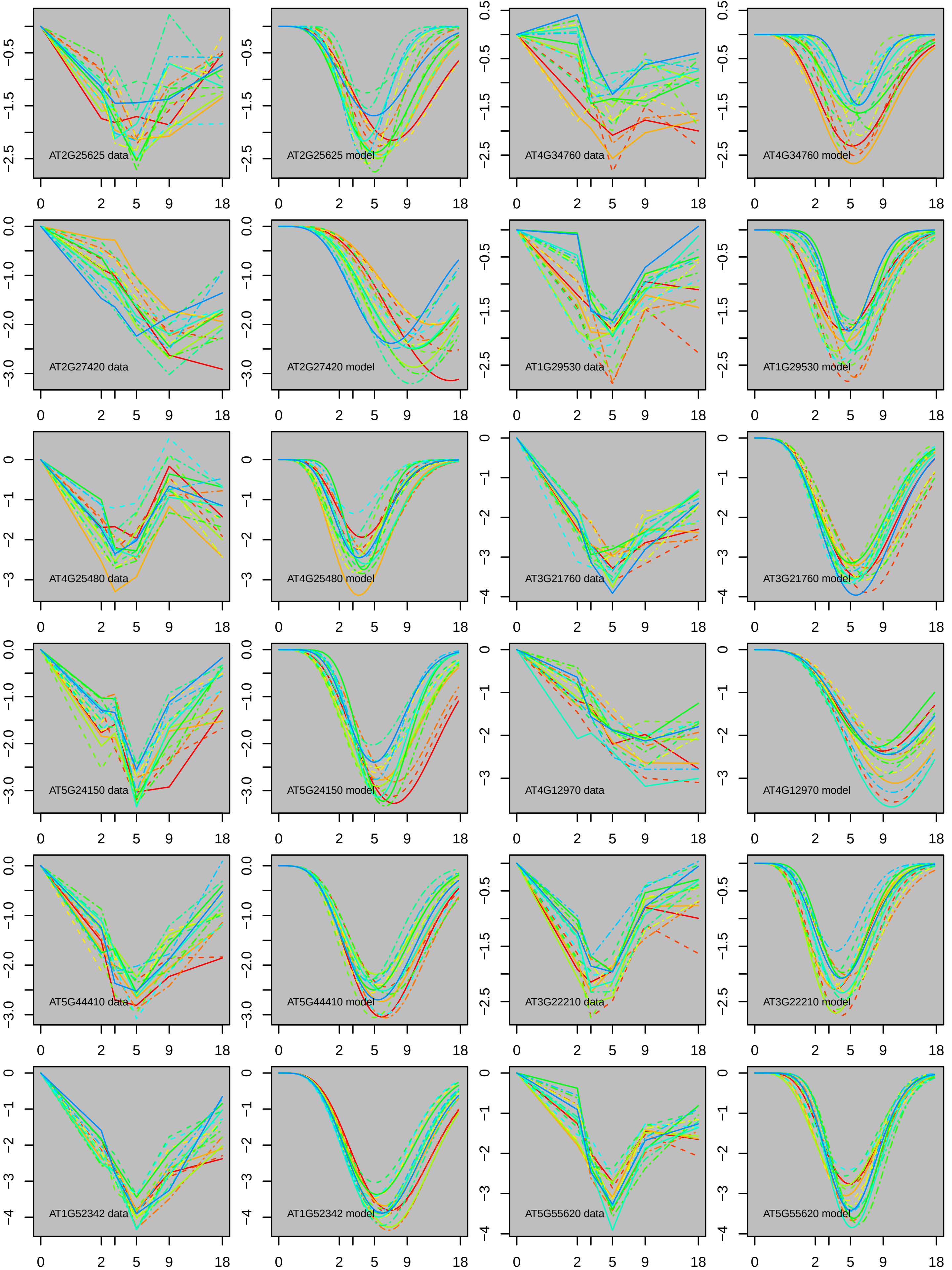
AT2G25625 data
AT2G25625 model
AT4G34760 data
AT4G34760 model
AT2G27420 data
AT2G27420 model
AT1G29530 data
AT1G29530 model
AT4G25480 data
AT4G25480 model
AT3G21760 data
AT3G21760 model
AT5G24150 data
AT5G24150 model
AT4G12970 data
AT4G12970 model
AT5G44410 data
AT5G44410 model
AT3G22210 data
AT3G22210 model
AT1G52342 data
AT1G52342 model
AT5G55620 data
AT5G55620 model

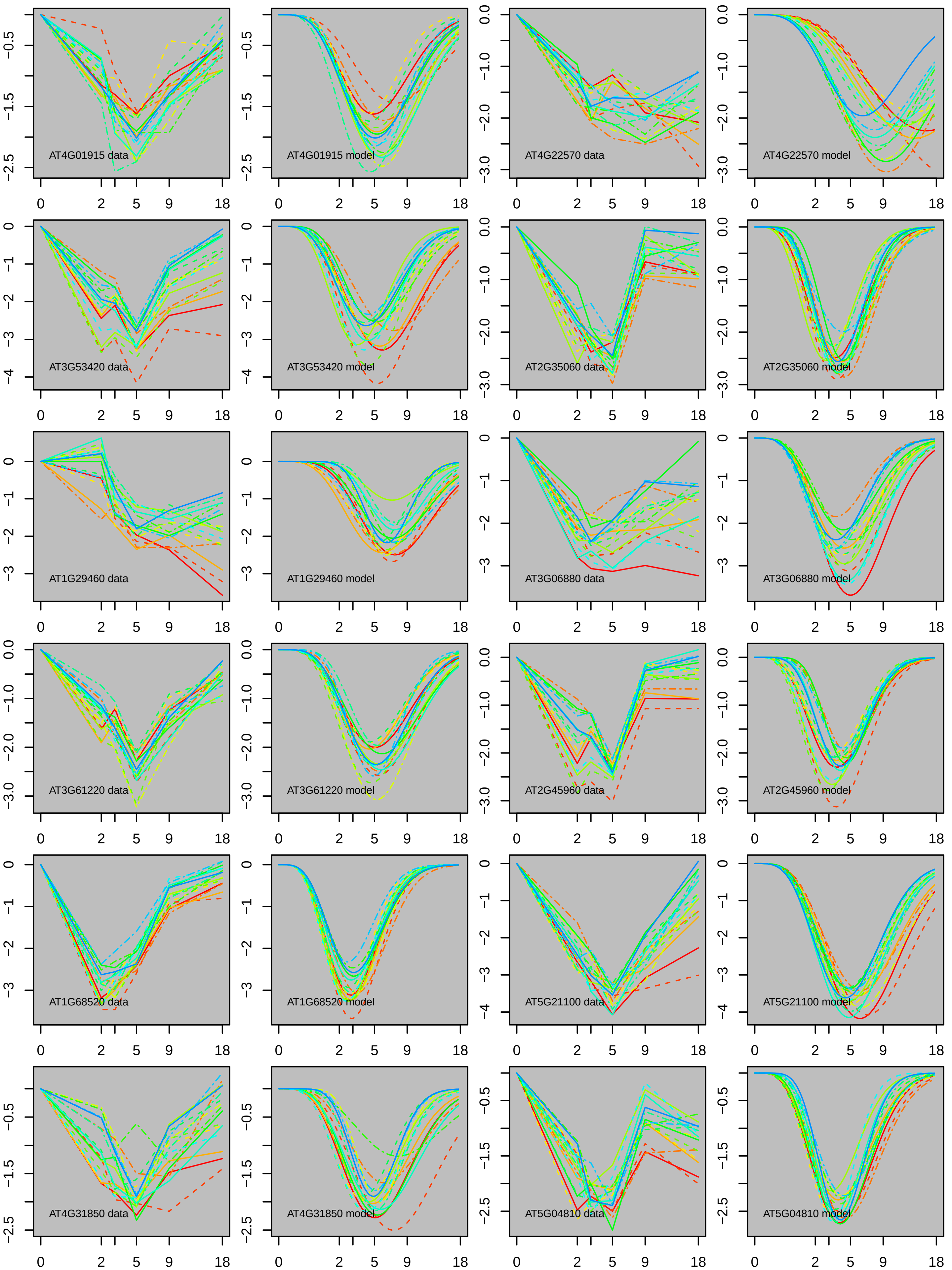
AT4G01915 data
AT4G01915 model
AT4G22570 data
AT4G22570 model
AT3G53420 data
AT3G53420 model
AT2G35060 data
AT2G35060 model
AT1G29460 data
AT1G29460 model
AT3G06880 data
AT3G06880 model
AT3G61220 data
AT3G61220 model
AT2G45960 data
AT2G45960 model
AT1G68520 data
AT1G68520 model
AT5G21100 data
AT5G21100 model
AT4G31850 data
AT4G31850 model
AT5G04810 data
AT5G04810 model

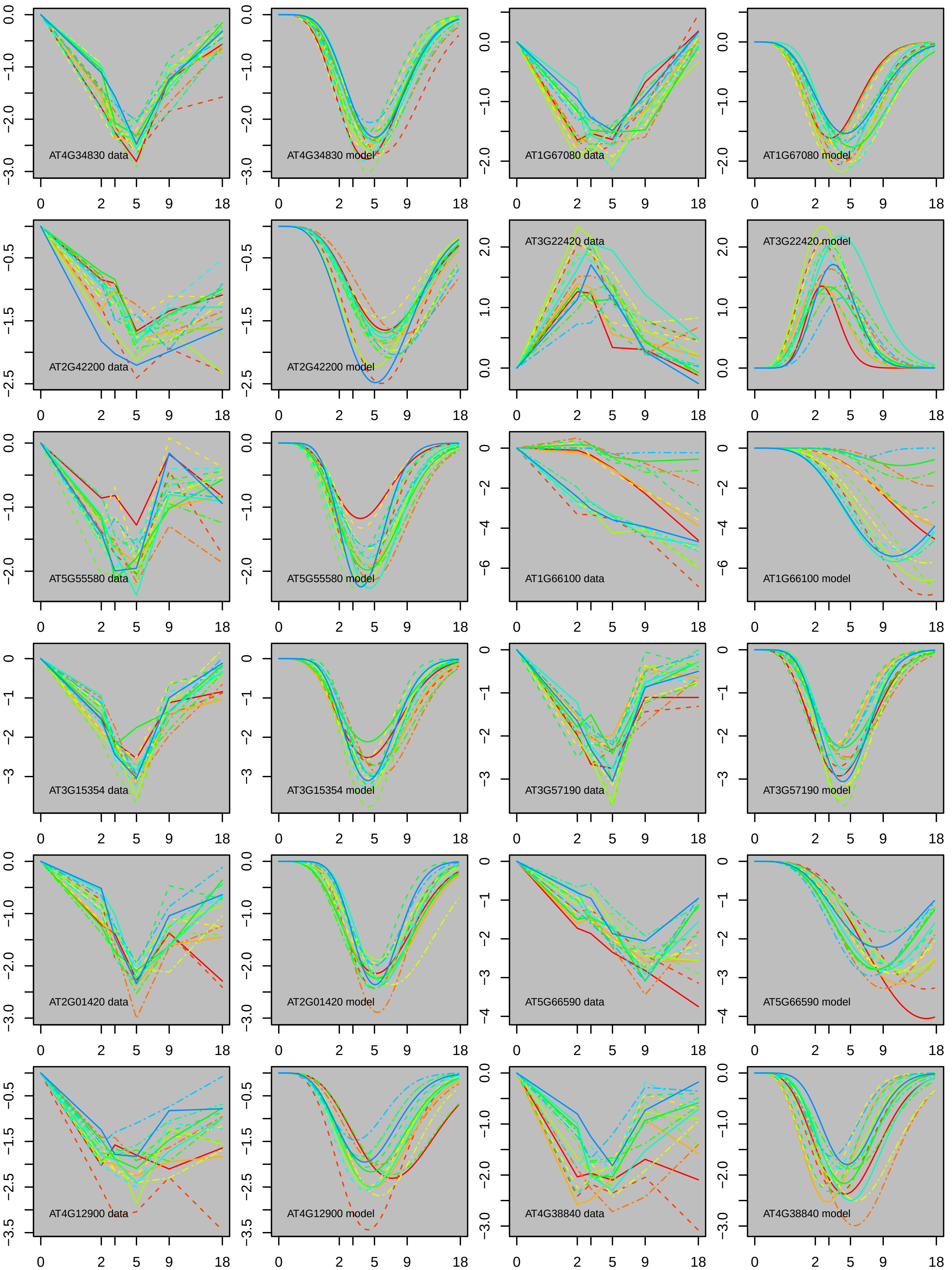

AT4G34830 data
AT4G34830 model
AT1G67080 data
AT1G67080 model
AT2G42200 data
AT2G42200 model
AT3G22420 data
AT3G22420 model
AT5G55580 data
AT5G55580 model
AT1G66100 data
AT1G66100 model
AT3G15354 data
AT3G15354 model
AT3G57190 data
AT3G57190 model
AT2G01420 data
AT2G01420 model
AT5G66590 data
AT5G66590 model
AT4G12900 data
AT4G12900 model
AT4G38840 data
AT4G38840 model

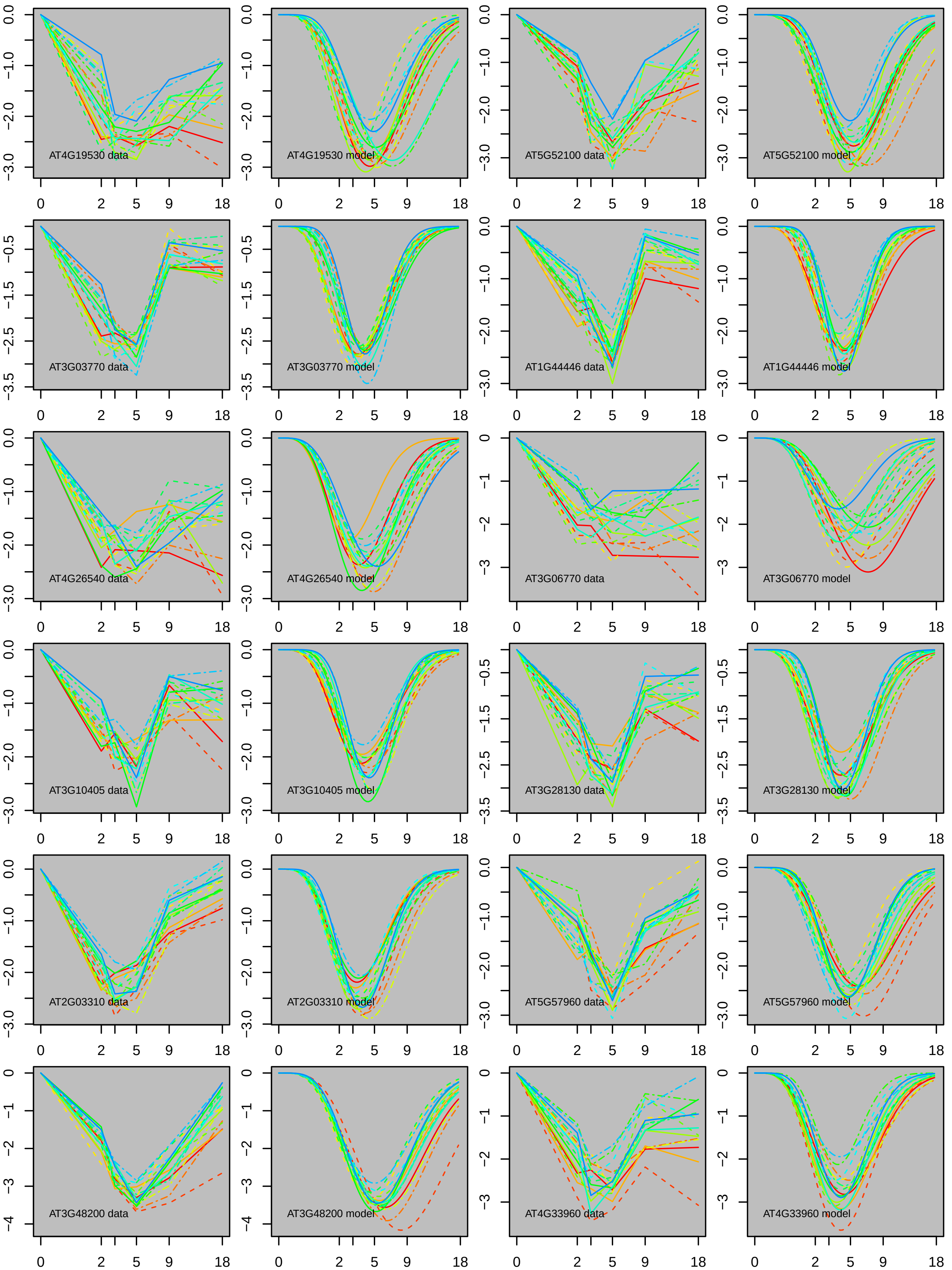

AT4G19530 data
AT4G19530 model
AT5G52100 data
AT5G52100 model
AT3G03770 data
AT3G03770 model
AT1G44446 data
AT1G44446 model
AT4G26540 data
AT4G26540 model
AT3G06770 data
AT3G06770 model
AT3G10405 data
AT3G10405 model
AT3G28130 data
AT3G28130 model
AT2G03310 data
AT2G03310 model
AT5G57960 data
AT5G57960 model
AT3G48200 data
AT3G48200 model
AT4G33960 data
AT4G33960 model
0 2 5 9 18

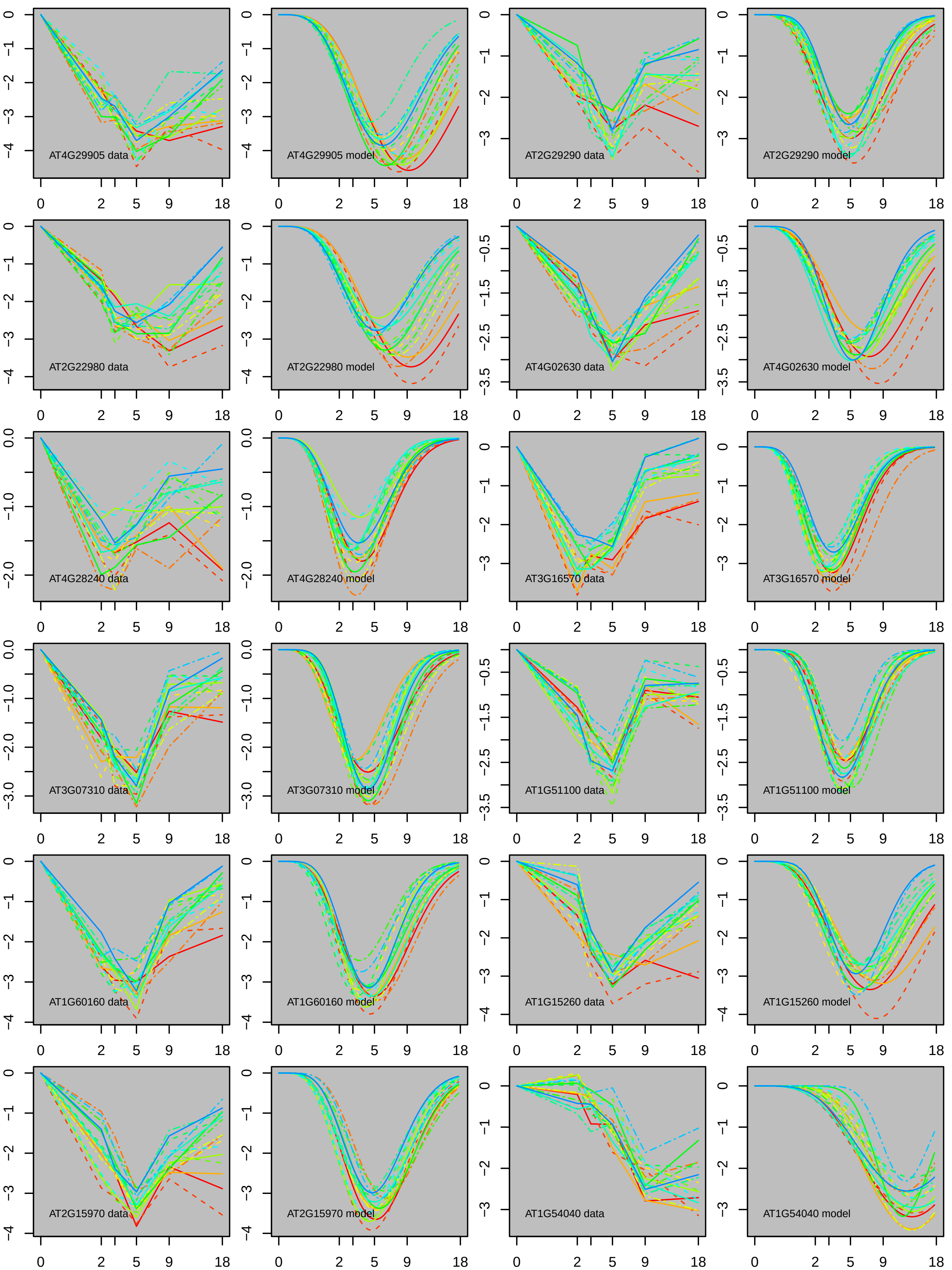
AT4G29905 data
AT4G29905 model
AT2G29290 data
AT2G29290 model
AT2G22980 data
AT2G22980 model
AT4G02630 data
AT4G02630 model
AT4G28240 data
AT4G28240 model
AT3G16570 data
AT3G16570 model
AT3G07310 data
AT3G07310 model
AT1G51100 data
AT1G51100 model
AT1G60160 data
AT1G60160 model
AT1G15260 data
AT1G15260 model
AT2G15970 data
AT2G15970 model
AT1G54040 data
AT1G54040 model

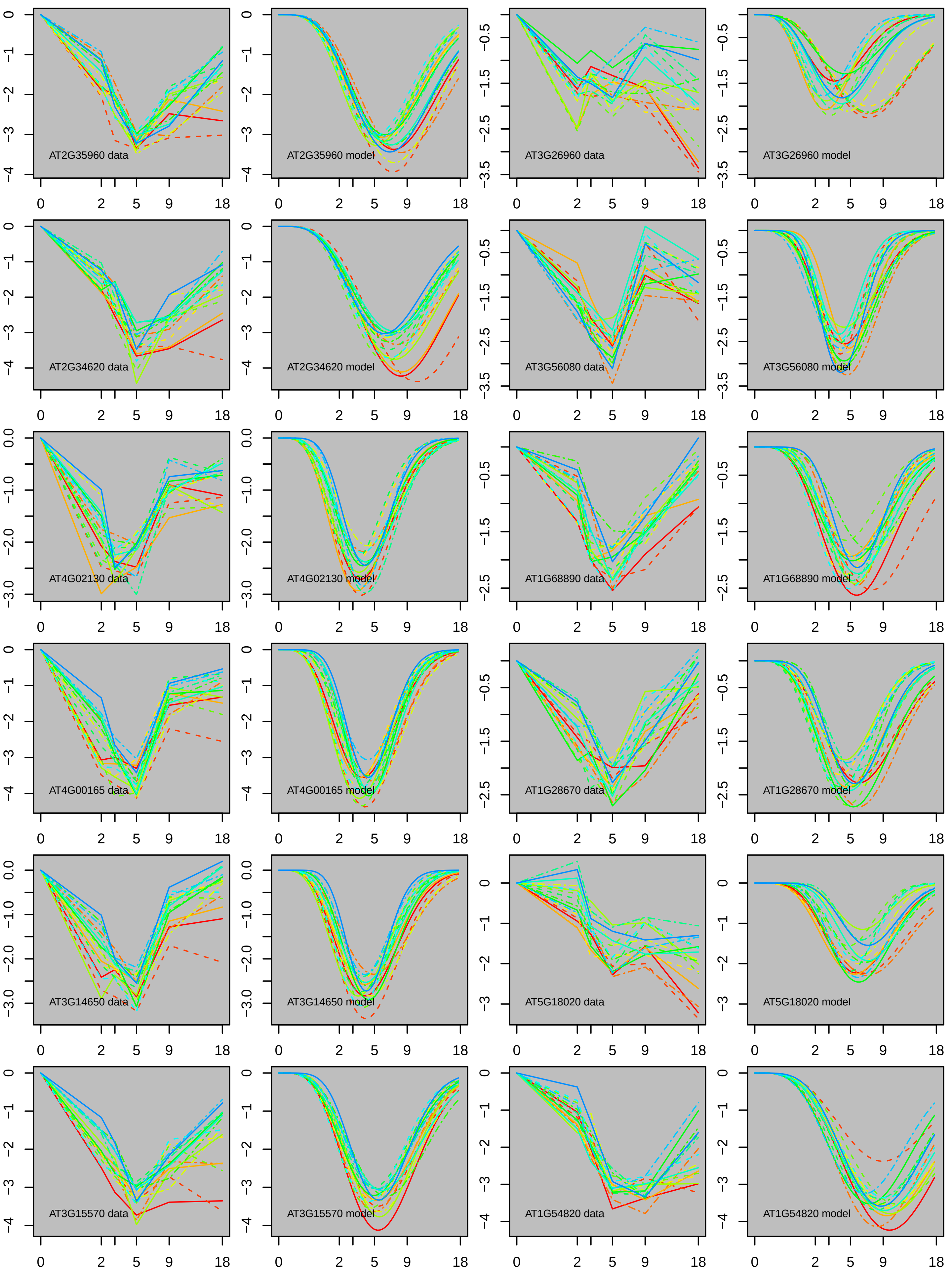
AT2G35960 data
AT2G35960 model
AT3G26960 data
AT3G26960 model
AT2G34620 data
AT2G34620 model
AT3G56080 data
AT3G56080 model
AT4G02130 data
AT4G02130 model
AT1G68890 data
AT1G68890 model
AT4G00165 data
AT4G00165 model
AT1G28670 data
AT1G28670 model
AT3G14650 data
AT3G14650 model
AT5G18020 data
AT5G18020 model
AT3G15570 data
AT3G15570 model
AT1G54820 data
AT1G54820 model

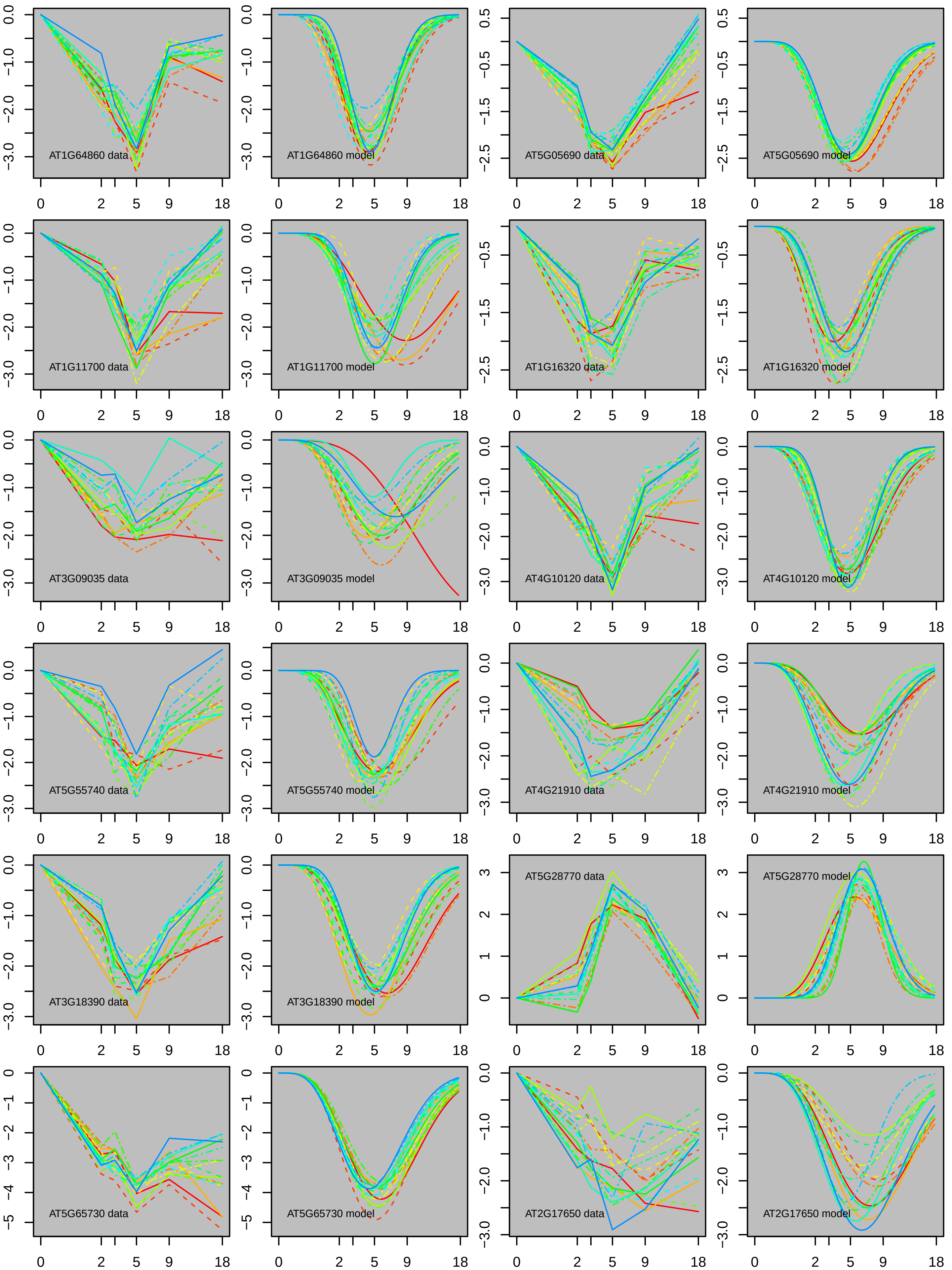
AT1G64860 data
AT1G64860 model
AT5G05690 data
AT5G05690 model
AT1G11700 data
AT1G11700 model
AT1G16320 data
AT1G16320 model
AT3G09035 data
AT3G09035 model
AT4G10120 data
AT4G10120 model
AT5G55740 data
AT5G55740 model
AT4G21910 data
AT4G21910 model
AT3G18390 data
AT3G18390 model
AT5G28770 data
AT5G28770 model
AT5G65730 data
AT5G65730 model
AT2G17650 data
AT2G17650 model
0
2
5
9
18

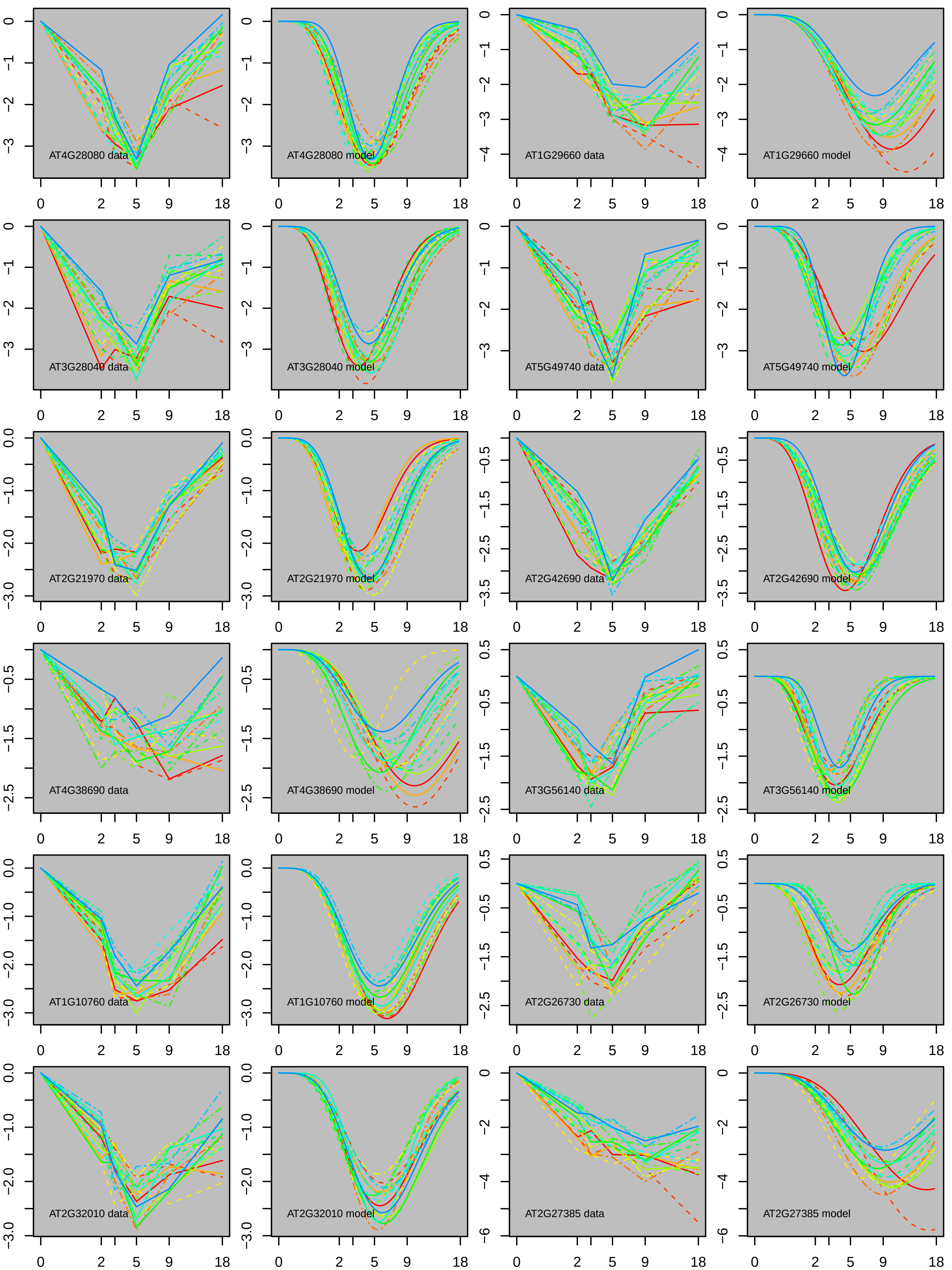
AT4G28080 data
AT4G28080 model
AT1G29660 data
AT1G29660 model
AT3G28040 data
AT3G28040 model
AT5G49740 data
AT5G49740 model
AT2G21970 data
AT2G21970 model
AT2G42690 data
AT2G42690 model
AT4G38690 data
AT4G38690 model
AT3G56140 data
AT3G56140 model
AT1G10760 data
AT1G10760 model
AT2G26730 data
AT2G26730 model
AT2G32010 data
AT2G32010 model
AT2G27385 data
AT2G27385 model

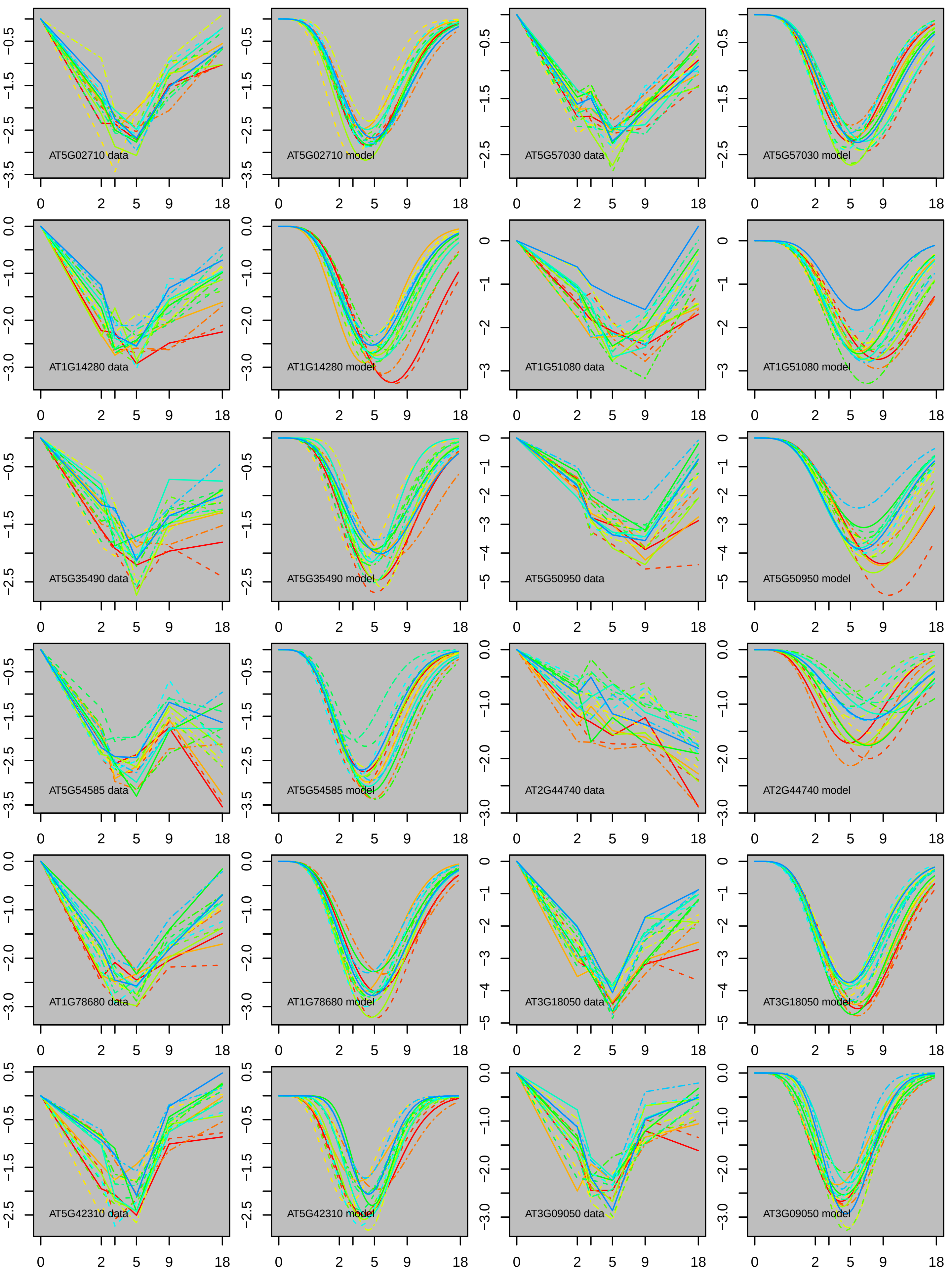
AT5G02710 data
AT5G02710 model
AT5G57030 data
AT5G57030 model
AT1G14280 data
AT1G14280 model
AT1G51080 data
AT1G51080 model
AT5G35490 data
AT5G35490 model
AT5G50950 data
AT5G50950 model
AT5G54585 data
AT5G54585 model
AT2G44740 data
AT2G44740 model
AT1G78680 data
AT1G78680 model
AT3G18050 data
AT3G18050 model
AT5G42310 data
AT5G42310 model
AT3G09050 data
AT3G09050 model

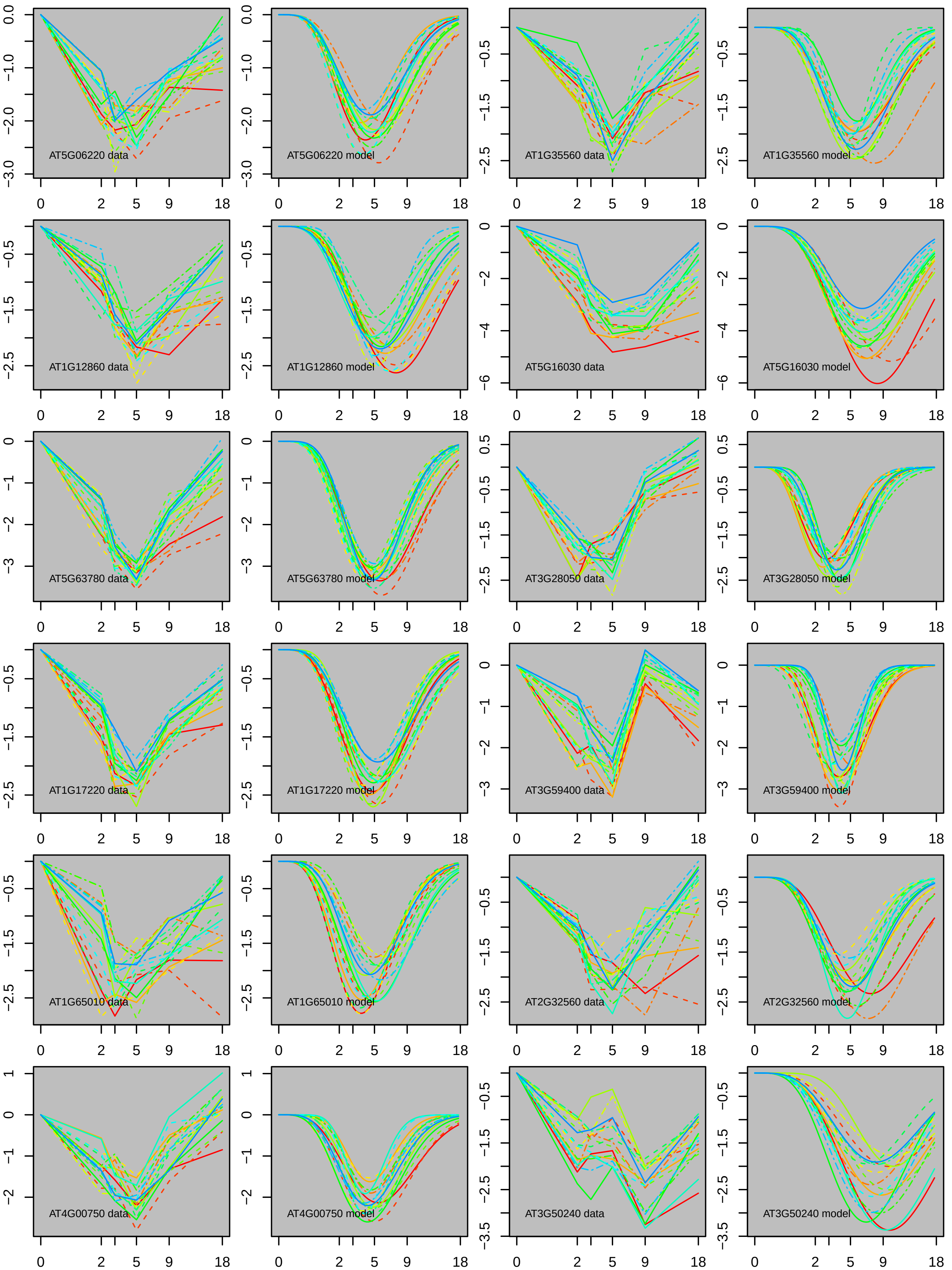

AT5G06220 data
AT5G06220 model
AT1G35560 data
AT1G35560 model
AT1G12860 data
AT1G12860 model
AT5G16030 data
AT5G16030 model
AT5G63780 data
AT5G63780 model
AT3G28050 data
AT3G28050 model
AT1G17220 data
AT1G17220 model
AT3G59400 data
AT3G59400 model
AT1G65010 data
AT1G65010 model
AT2G32560 data
AT2G32560 model
AT4G00750 data
AT4G00750 model
AT3G50240 data
AT3G50240 model
0 2 5 9 18

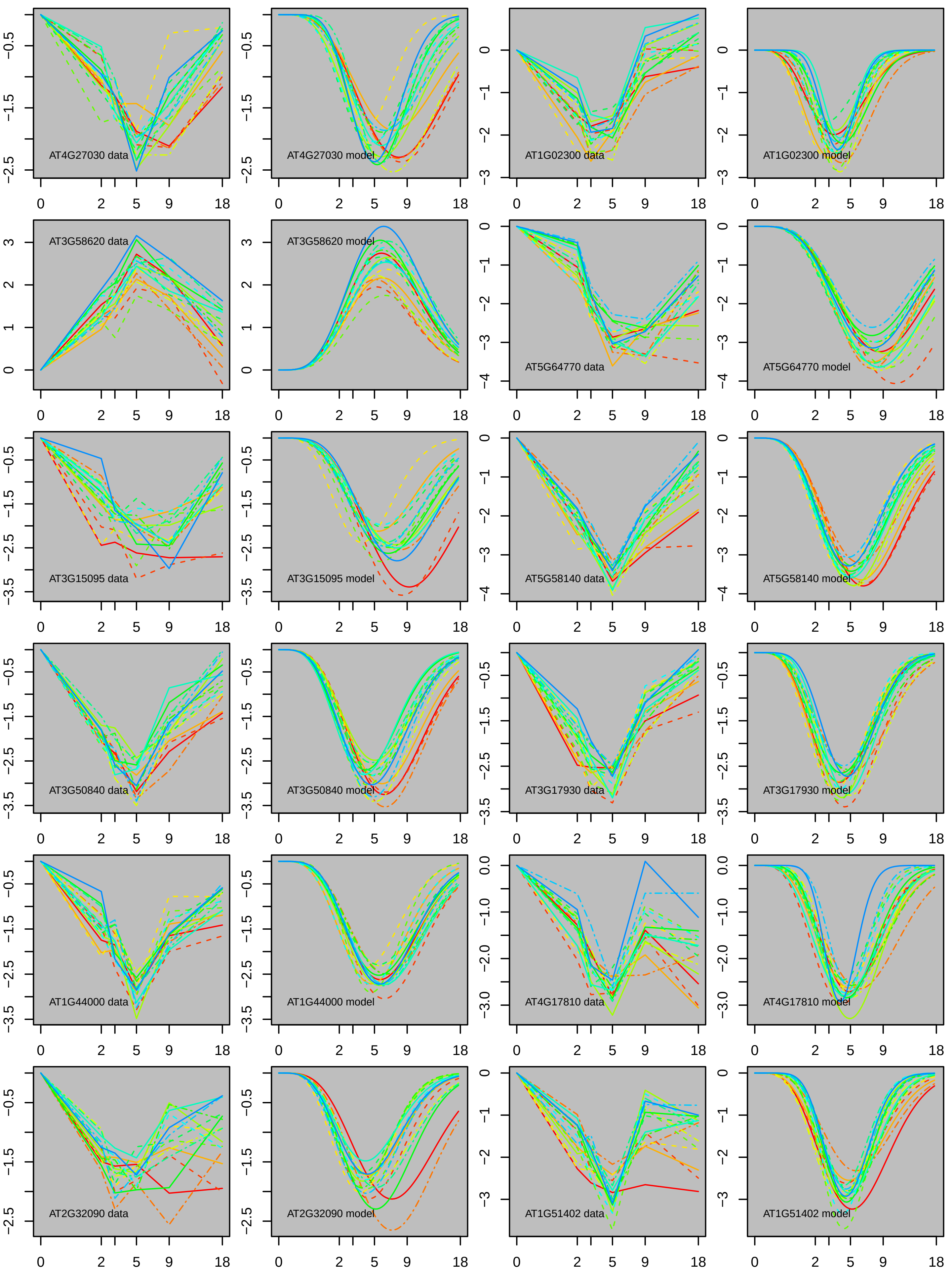
AT4G27030 data
AT4G27030 model
AT1G02300 data
AT1G02300 model
AT3G58620 data
AT3G58620 model
AT5G64770 data
AT5G64770 model
AT3G15095 data
AT3G15095 model
AT5G58140 data
AT5G58140 model
AT3G50840 data
AT3G50840 model
AT3G17930 data
AT3G17930 model
AT1G44000 data
AT1G44000 model
AT4G17810 data
AT4G17810 model
AT2G32090 data
AT2G32090 model
AT1G51402 data
AT1G51402 model

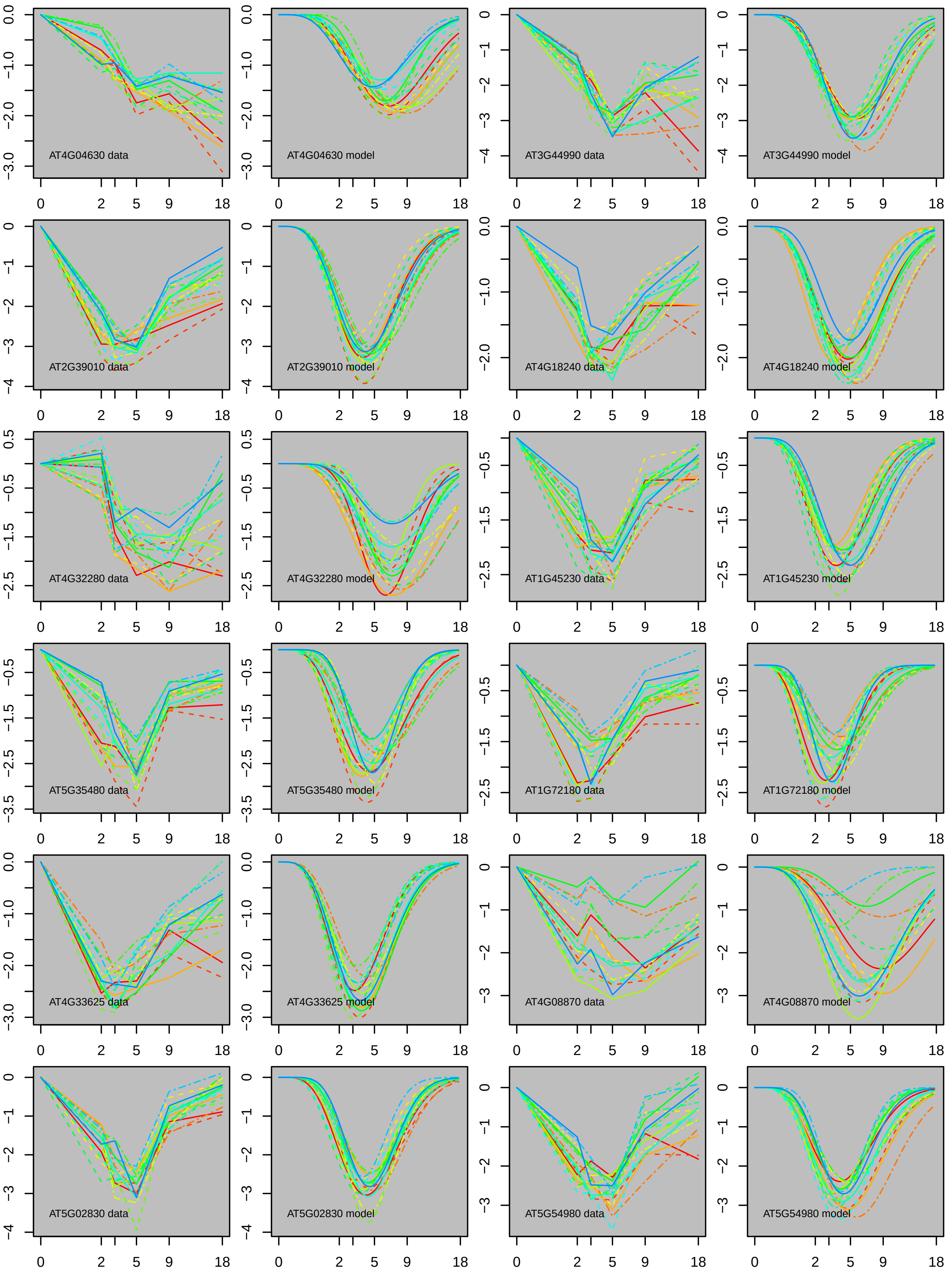
AT4G04630 data
AT4G04630 model
AT3G44990 data
AT3G44990 model
AT2G39010 data
AT2G39010 model
AT4G18240 data
AT4G18240 model
AT4G32280 data
AT4G32280 model
AT1G45230 data
AT1G45230 model
AT5G35480 data
AT5G35480 model
AT1G72180 data
AT1G72180 model
AT4G33625 data
AT4G33625 model
AT4G08870 data
AT4G08870 model
AT5G02830 data
AT5G02830 model
AT5G54980 data
AT5G54980 model

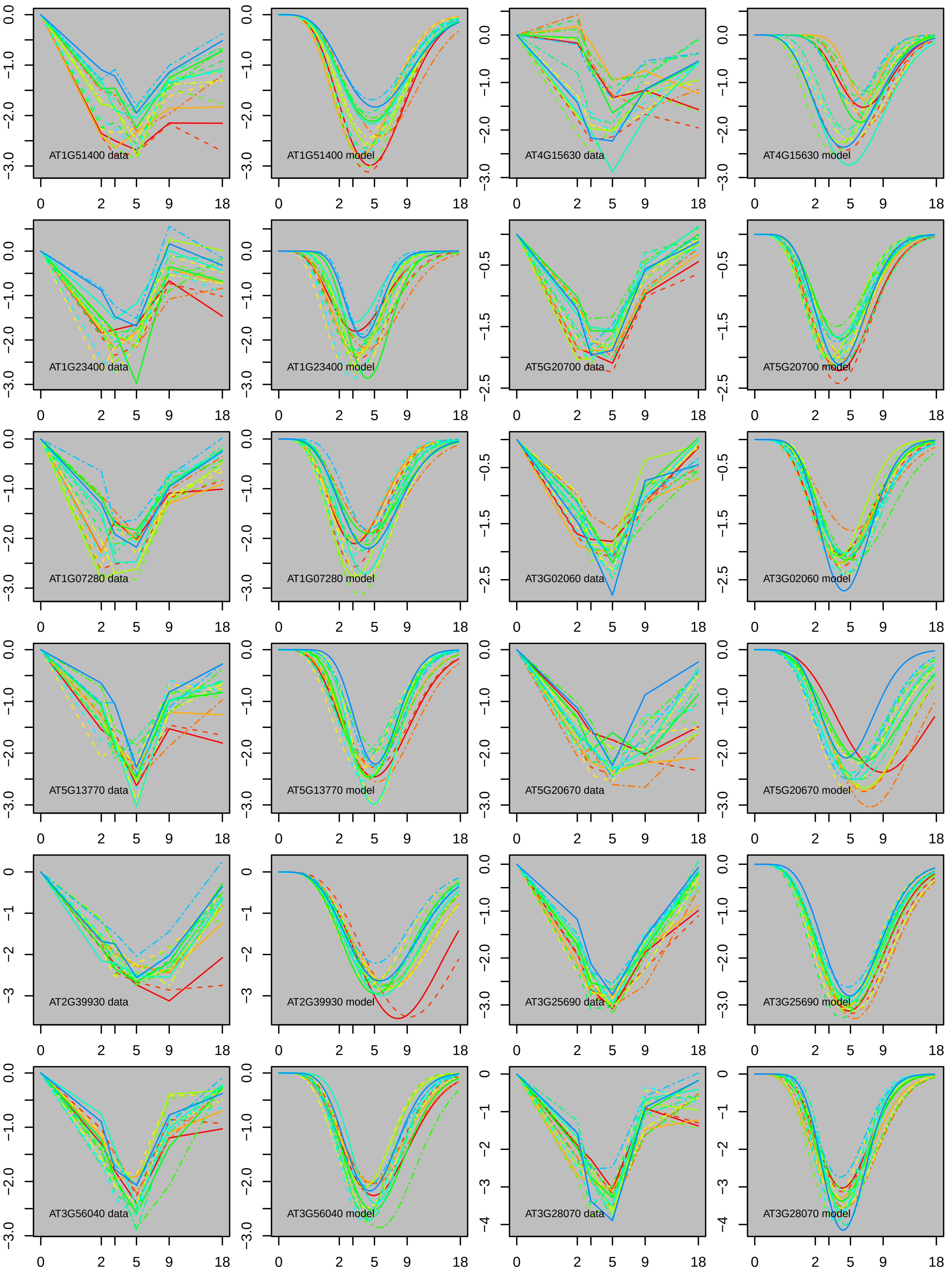
AT1G51400 data
AT1G51400 model
AT4G15630 data
AT4G15630 model
AT1G23400 data
AT1G23400 model
AT5G20700 data
AT5G20700 model
AT1G07280 data
AT1G07280 model
AT3G02060 data
AT3G02060 model
AT5G13770 data
AT5G13770 model
AT5G20670 data
AT5G20670 model
AT2G39930 data
AT2G39930 model
AT3G25690 data
AT3G25690 model
AT3G56040 data
AT3G56040 model
AT3G28070 data
AT3G28070 model
0
2
5
9
18

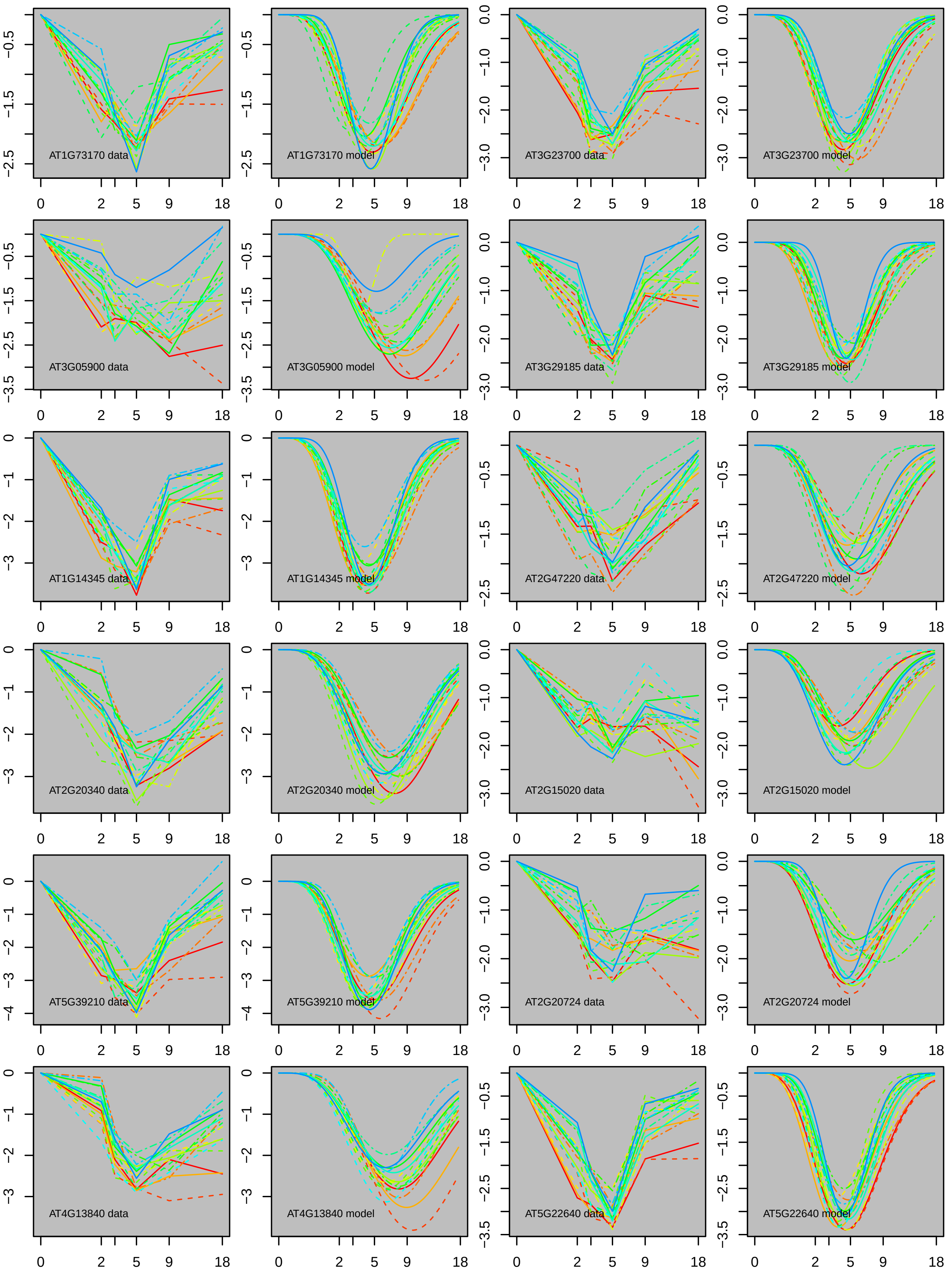
AT1G73170 data
AT1G73170 model
AT3G23700 data
AT3G23700 model
AT3G05900 data
AT3G05900 model
AT3G29185 data
AT3G29185 model
AT1G14345 data
AT1G14345 model
AT2G47220 data
AT2G47220 model
AT2G20340 data
AT2G20340 model
AT2G15020 data
AT2G15020 model
AT5G39210 data
AT5G39210 model
AT2G20724 data
AT2G20724 model
AT4G13840 data
AT4G13840 model
AT5G22640 data
AT5G22640 model

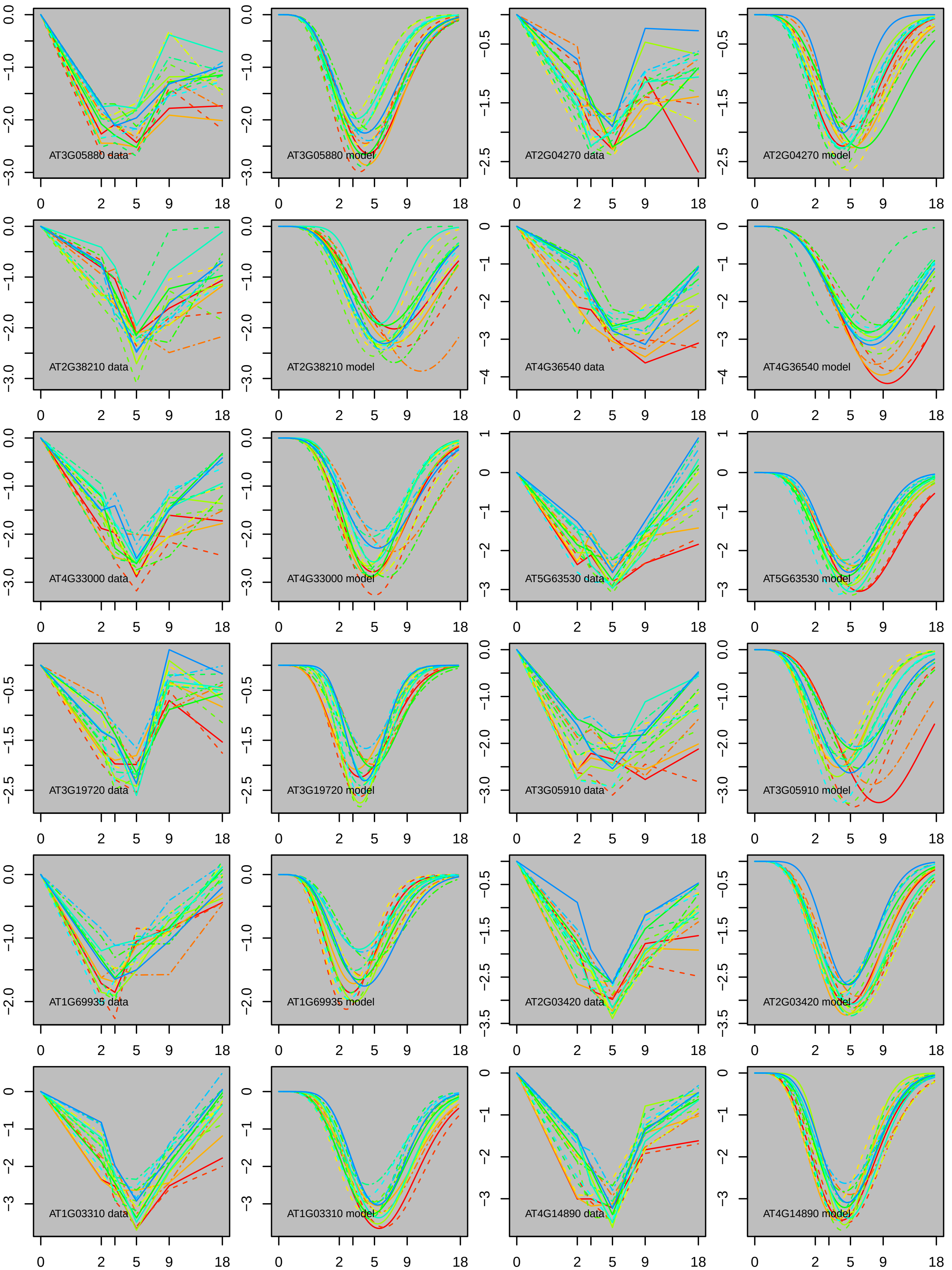
AT3G05880 data
AT3G05880 model
AT2G04270 data
AT2G04270 model
AT2G38210 data
AT2G38210 model
AT4G36540 data
AT4G36540 model
AT4G33000 data
AT4G33000 model
AT5G63530 data
AT5G63530 model
AT3G19720 data
AT3G19720 model
AT3G05910 data
AT3G05910 model
AT1G69935 data
AT1G69935 model
AT2G03420 data
AT2G03420 model
AT1G03310 data
AT1G03310 model
AT4G14890 data
AT4G14890 model

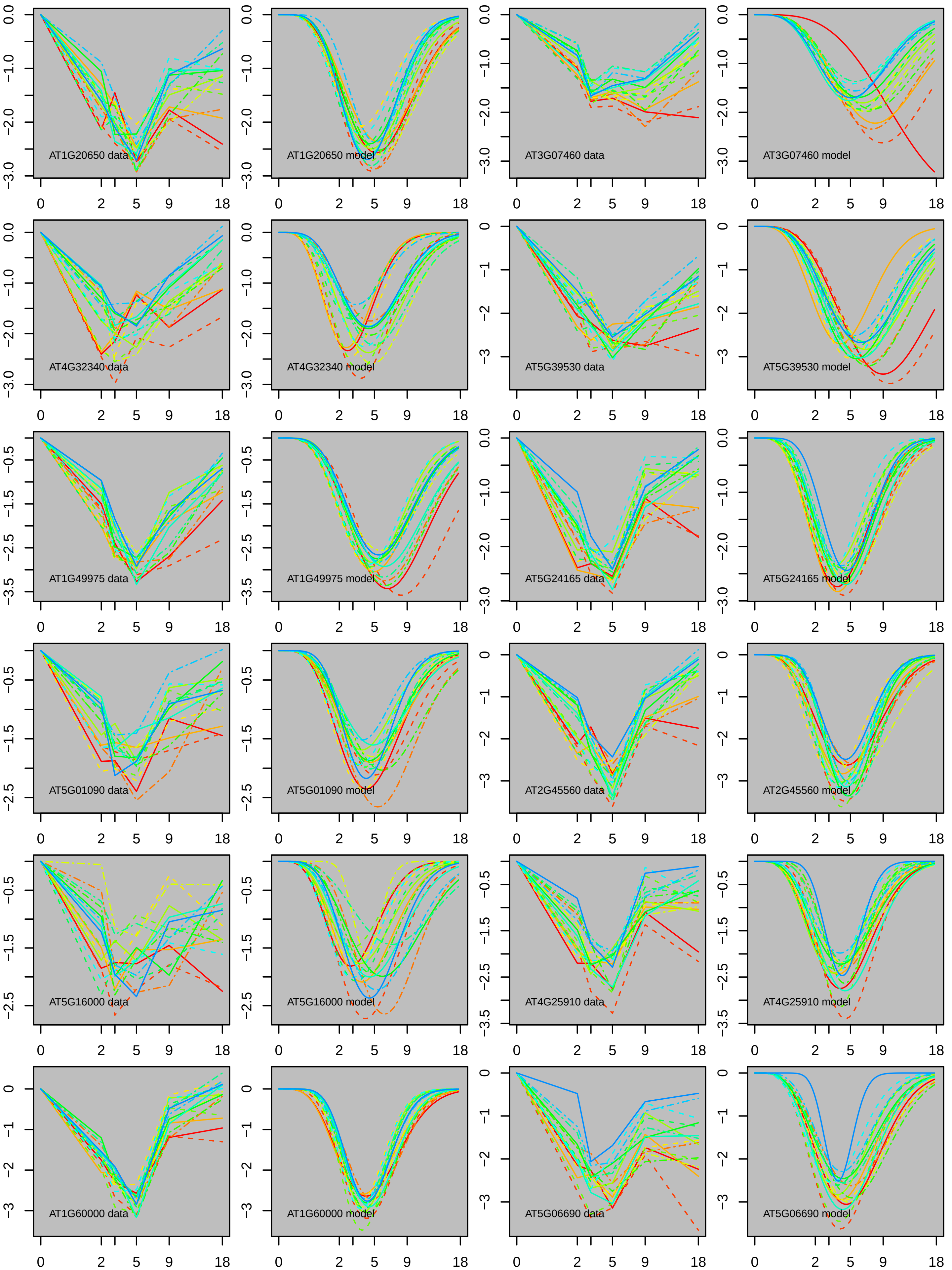
AT1G20650 data
AT1G20650 model
AT3G07460 data
AT3G07460 model
AT4G32340 data
AT4G32340 model
AT5G39530 data
AT5G39530 model
AT1G49975 data
AT1G49975 model
AT5G24165 data
AT5G24165 model
AT5G01090 data
AT5G01090 model
AT2G45560 data
AT2G45560 model
AT5G16000 data
AT5G16000 model
AT4G25910 data
AT4G25910 model
AT1G60000 data
AT1G60000 model
AT5G06690 data
AT5G06690 model
0
2
5
9
18

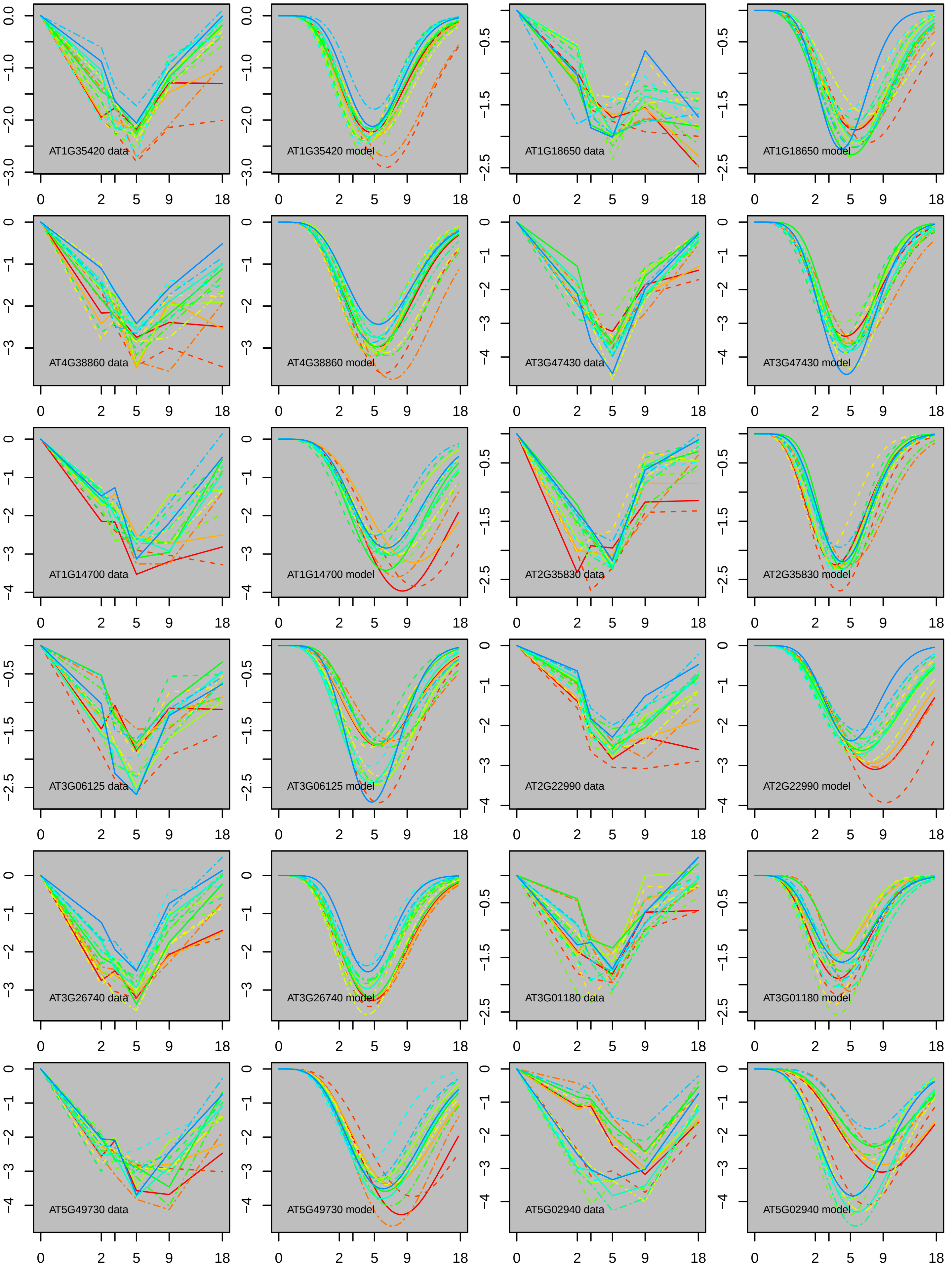
AT1G35420 data
AT1G35420 model
AT1G18650 data
AT1G18650 model
AT4G38860 data
AT4G38860 model
AT3G47430 data
AT3G47430 model
AT1G14700 data
AT1G14700 model
AT2G35830 data
AT2G35830 model
AT3G06125 data
AT3G06125 model
AT2G22990 data
AT2G22990 model
AT3G26740 data
AT3G26740 model
AT3G01180 data
AT3G01180 model
AT5G49730 data
AT5G49730 model
AT5G02940 data
AT5G02940 model

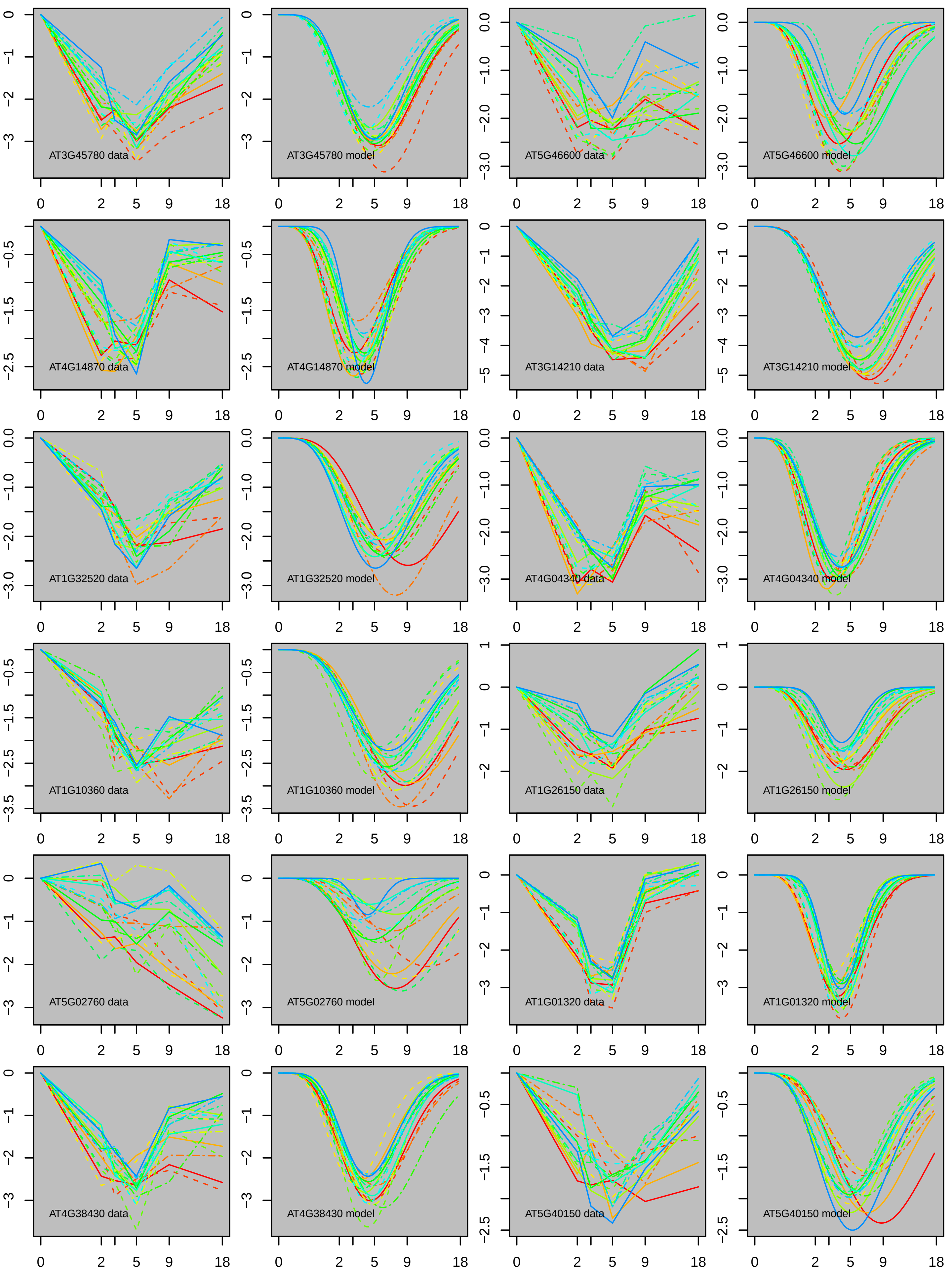
AT3G45780 data
AT3G45780 model
AT5G46600 data
AT5G46600 model
AT4G14870 data
AT4G14870 model
AT3G14210 data
AT3G14210 model
AT1G32520 data
AT1G32520 model
AT4G04340 data
AT4G04340 model
AT1G10360 data
AT1G10360 model
AT1G26150 data
AT1G26150 model
AT5G02760 data
AT5G02760 model
AT1G01320 data
AT1G01320 model
AT4G38430 data
AT4G38430 model
AT5G40150 data
AT5G40150 model

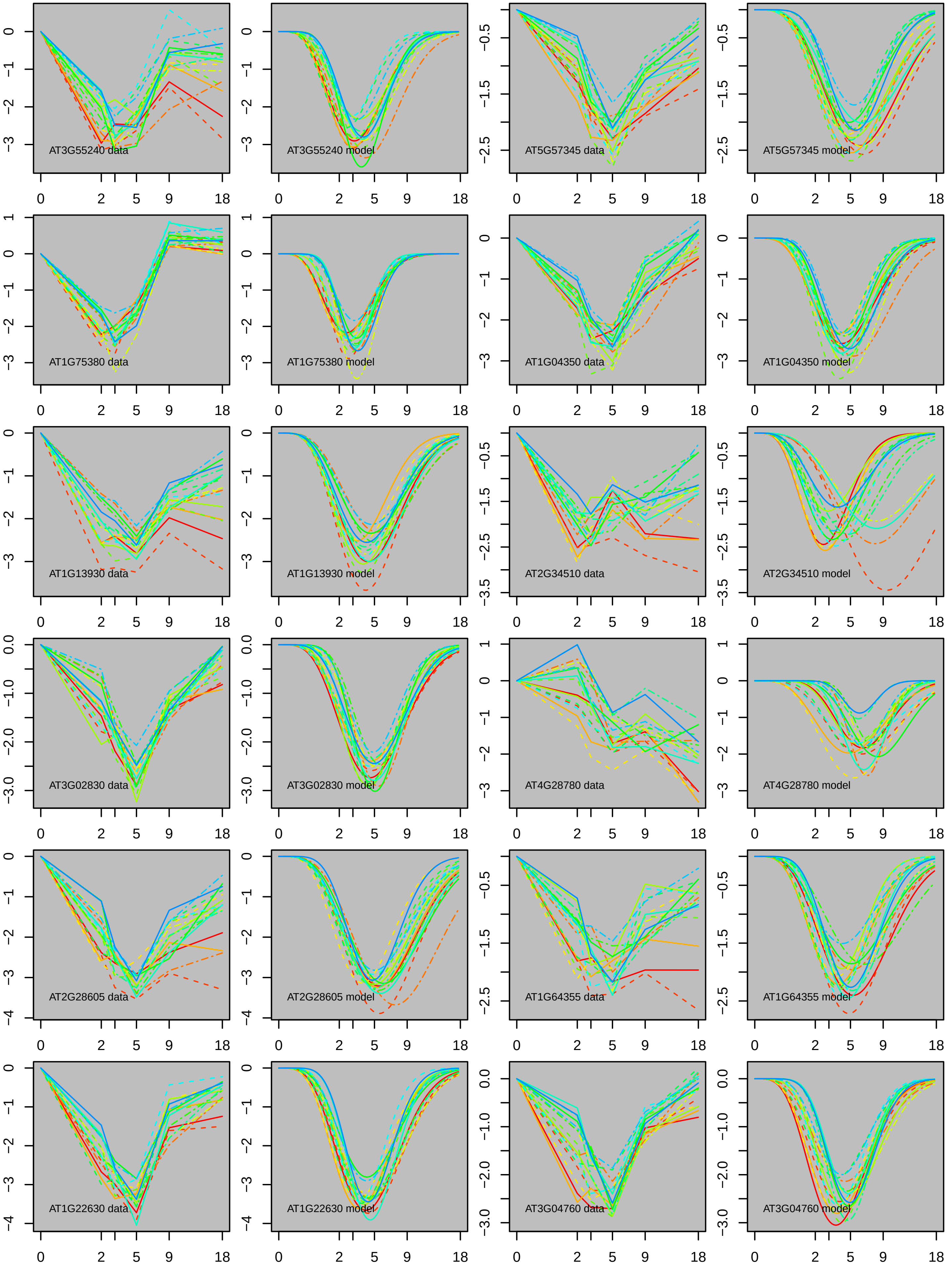
AT3G55240 data
AT3G55240 model
AT5G57345 data
AT5G57345 model
AT1G75380 data
AT1G75380 model
AT1G04350 data
AT1G04350 model
AT1G13930 data
AT1G13930 model
AT2G34510 data
AT2G34510 model
AT3G02830 data
AT3G02830 model
AT4G28780 data
AT4G28780 model
AT2G28605 data
AT2G28605 model
AT1G64355 data
AT1G64355 model
AT1G22630 data
AT1G22630 model
AT3G04760 data
AT3G04760 model

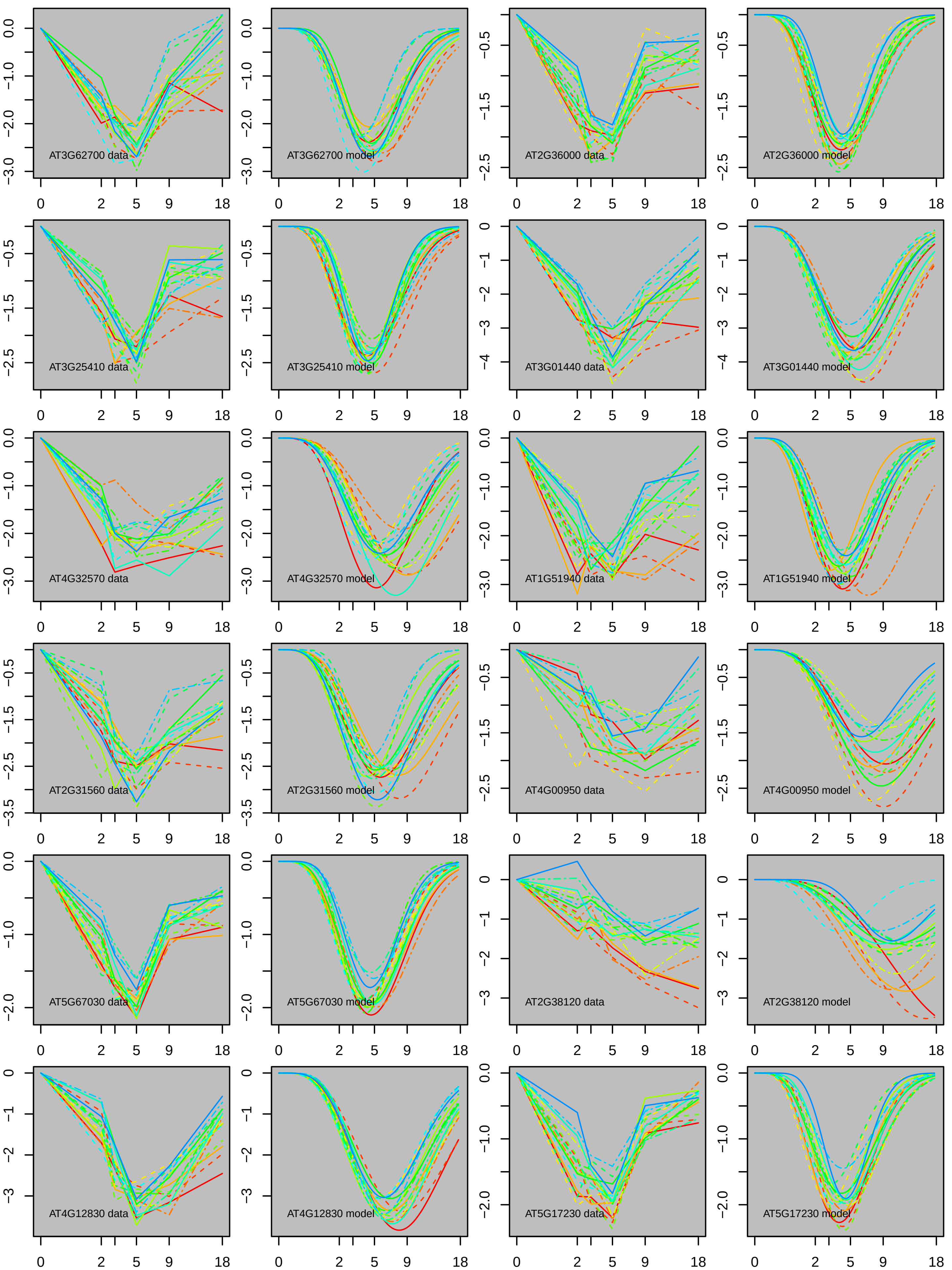
AT3G62700 data
AT3G62700 model
AT2G36000 data
AT2G36000 model
AT3G25410 data
AT3G25410 model
AT3G01440 data
AT3G01440 model
AT4G32570 data
AT4G32570 model
AT1G51940 data
AT1G51940 model
AT2G31560 data
AT2G31560 model
AT4G00950 data
AT4G00950 model
AT5G67030 data
AT5G67030 model
AT2G38120 data
AT2G38120 model
AT4G12830 data
AT4G12830 model
AT5G17230 data
AT5G17230 model

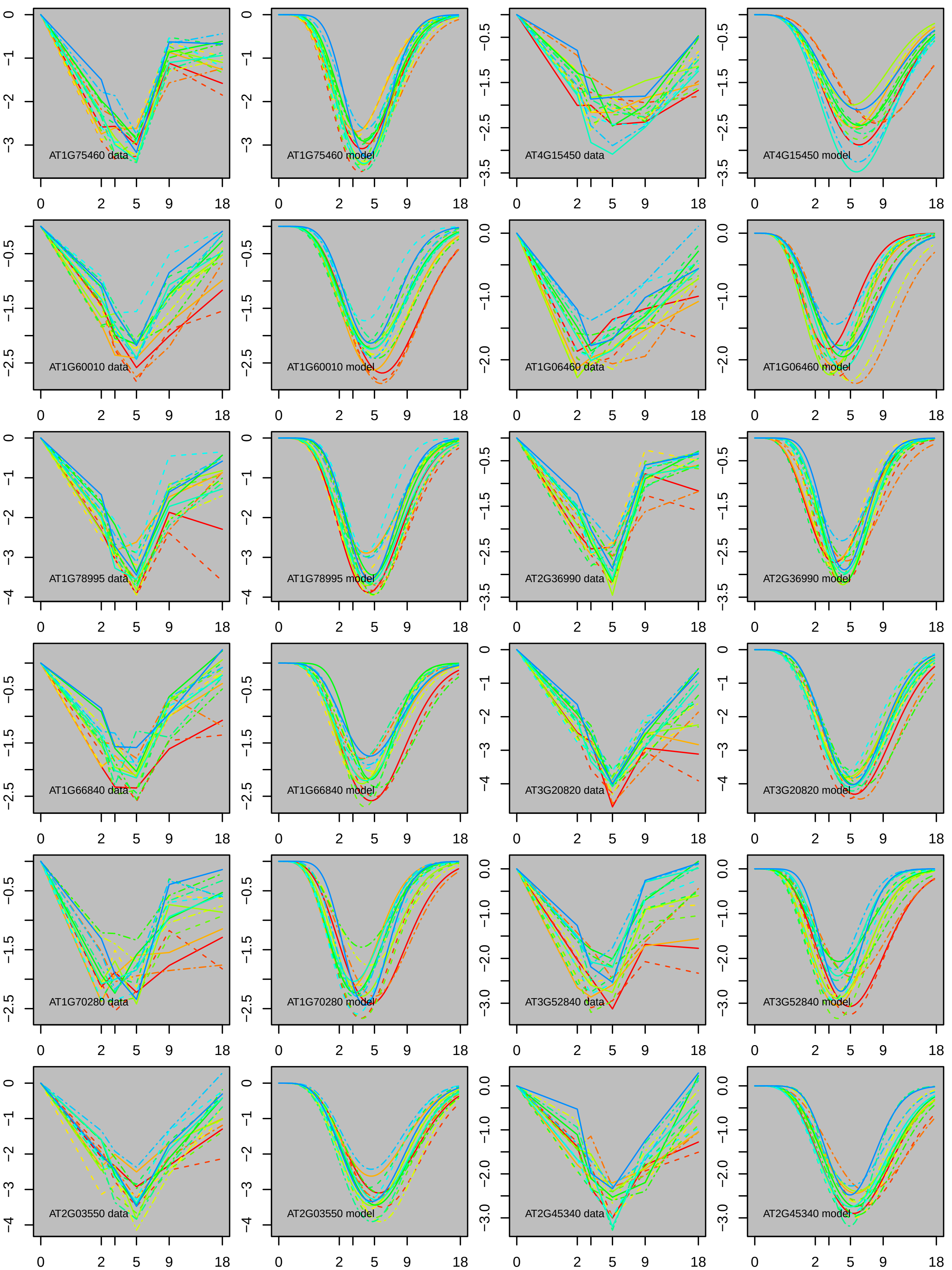
AT1G75460 data
AT1G75460 model
AT4G15450 data
AT4G15450 model
AT1G60010 data
AT1G60010 model
AT1G06460 data
AT1G06460 model
AT1G78995 data
AT1G78995 model
AT2G36990 data
AT2G36990 model
AT1G66840 data
AT1G66840 model
AT3G20820 data
AT3G20820 model
AT1G70280 data
AT1G70280 model
AT3G52840 data
AT3G52840 model
AT2G03550 data
AT2G03550 model
AT2G45340 data
AT2G45340 model

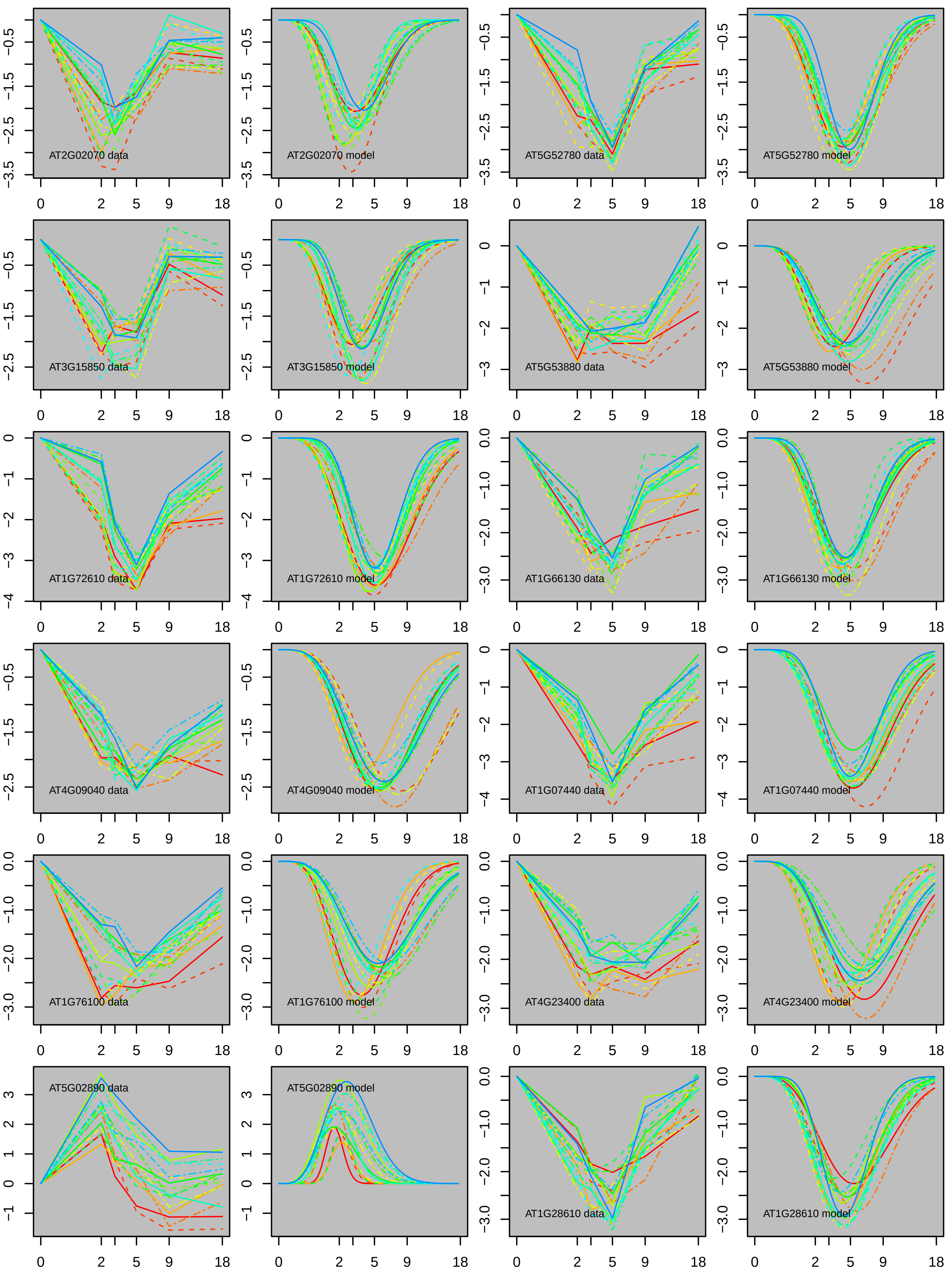

AT2G02070 data
AT2G02070 model
AT5G52780 data
AT5G52780 model
AT3G15850 data
AT3G15850 model
AT5G53880 data
AT5G53880 model
AT1G72610 data
AT1G72610 model
AT1G66130 data
AT1G66130 model
AT4G09040 data
AT4G09040 model
AT1G07440 data
AT1G07440 model
AT1G76100 data
AT1G76100 model
AT4G23400 data
AT4G23400 model
AT5G02890 data
AT5G02890 model
AT1G28610 data
AT1G28610 model

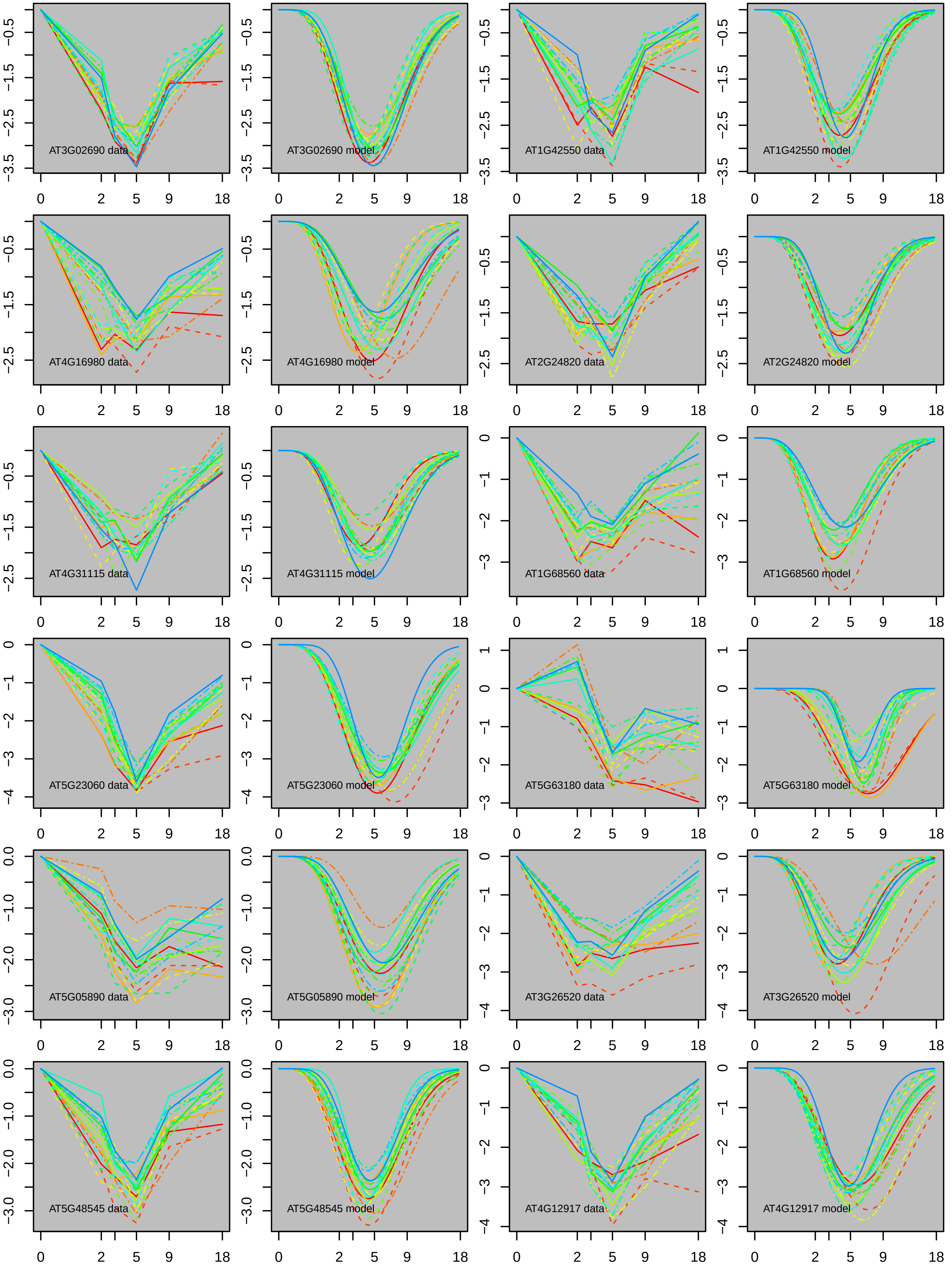

AT3G02690 data
AT3G02690 model
AT1G42550 data
AT1G42550 model
AT4G16980 data
AT4G16980 model
AT2G24820 data
AT2G24820 model
AT4G31115 data
AT4G31115 model
AT1G68560 data
AT1G68560 model
AT5G23060 data
AT5G23060 model
AT5G63180 data
AT5G63180 model
AT5G05890 data
AT5G05890 model
AT3G26520 data
AT3G26520 model
AT5G48545 data
AT5G48545 model
AT4G12917 data
AT4G12917 model

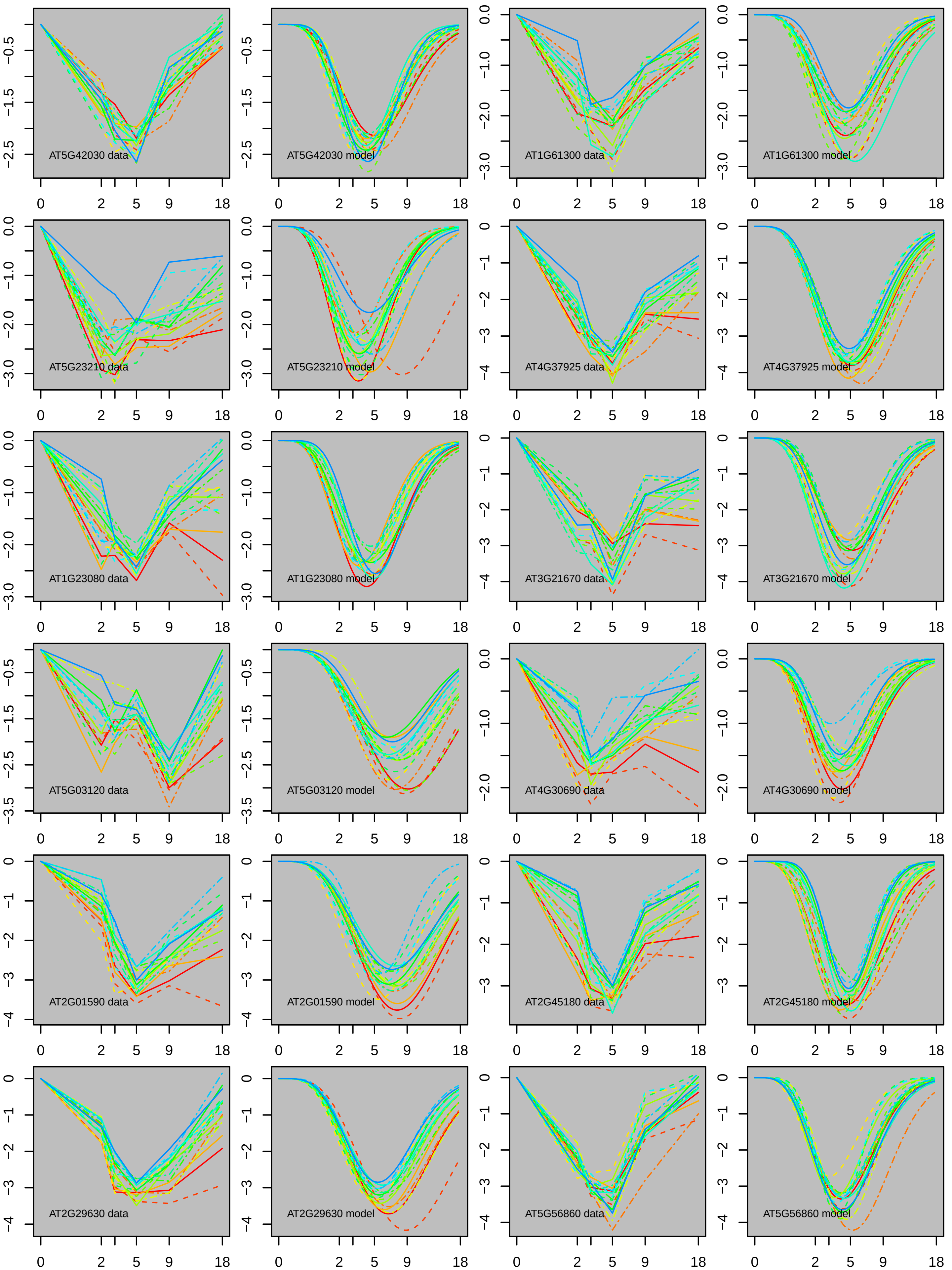
AT5G42030 data
AT5G42030 model
AT1G61300 data
AT1G61300 model
AT5G23210 data
AT5G23210 model
AT4G37925 data
AT4G37925 model
AT1G23080 data
AT1G23080 model
AT3G21670 data
AT3G21670 model
AT5G03120 data
AT5G03120 model
AT4G30690 data
AT4G30690 model
AT2G01590 data
AT2G01590 model
AT2G45180 data
AT2G45180 model
AT2G29630 data
AT2G29630 model
AT5G56860 data
AT5G56860 model

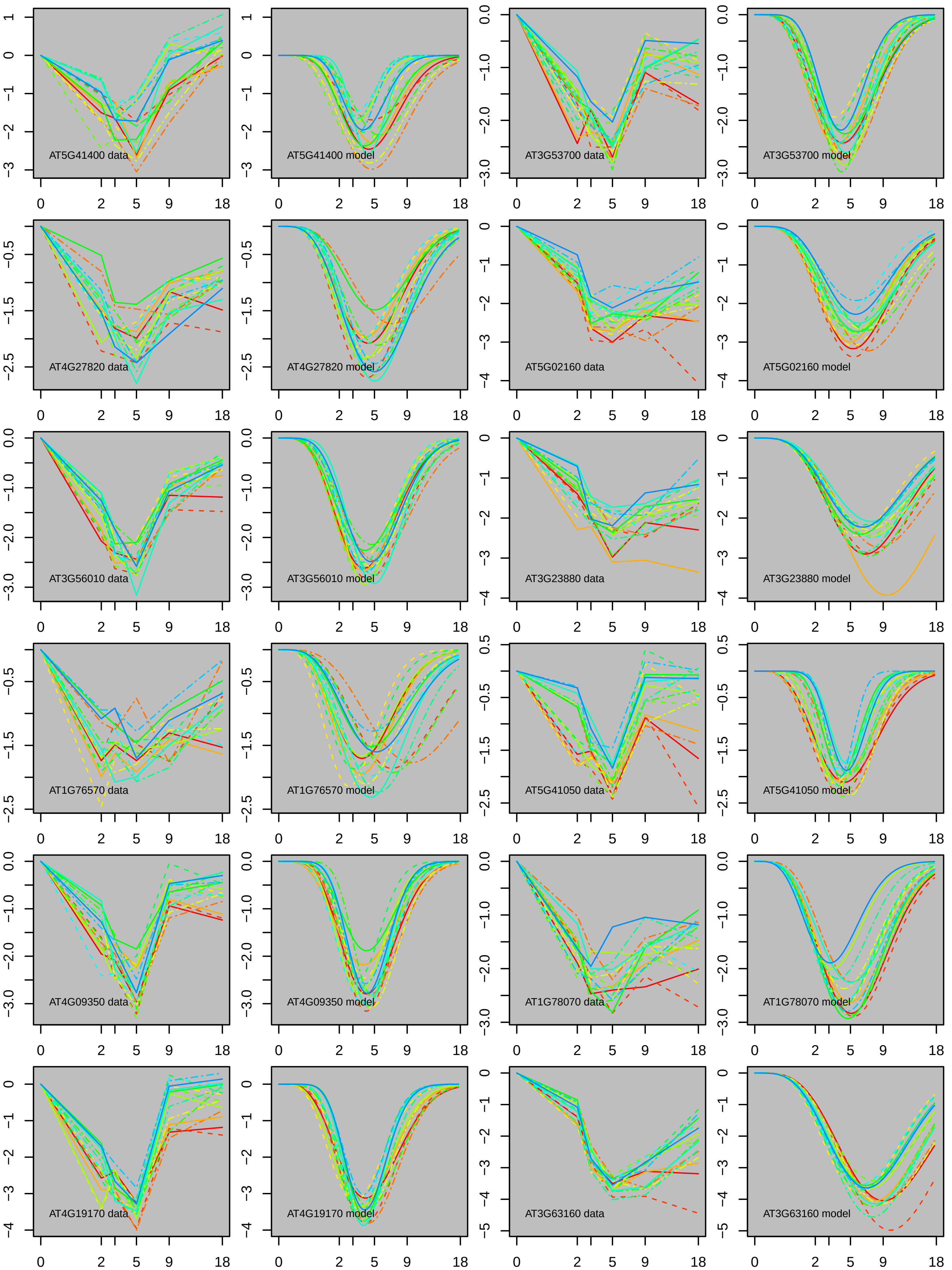
AT5G41400 data
AT5G41400 model
AT3G53700 data
AT3G53700 model
AT4G27820 data
AT4G27820 model
AT5G02160 data
AT5G02160 model
AT3G56010 data
AT3G56010 model
AT3G23880 data
AT3G23880 model
AT1G76570 data
AT1G76570 model
AT5G41050 data
AT5G41050 model
AT4G09350 data
AT4G09350 model
AT1G78070 data
AT1G78070 model
AT4G19170 data
AT4G19170 model
AT3G63160 data
AT3G63160 model
0 2 5 9 18

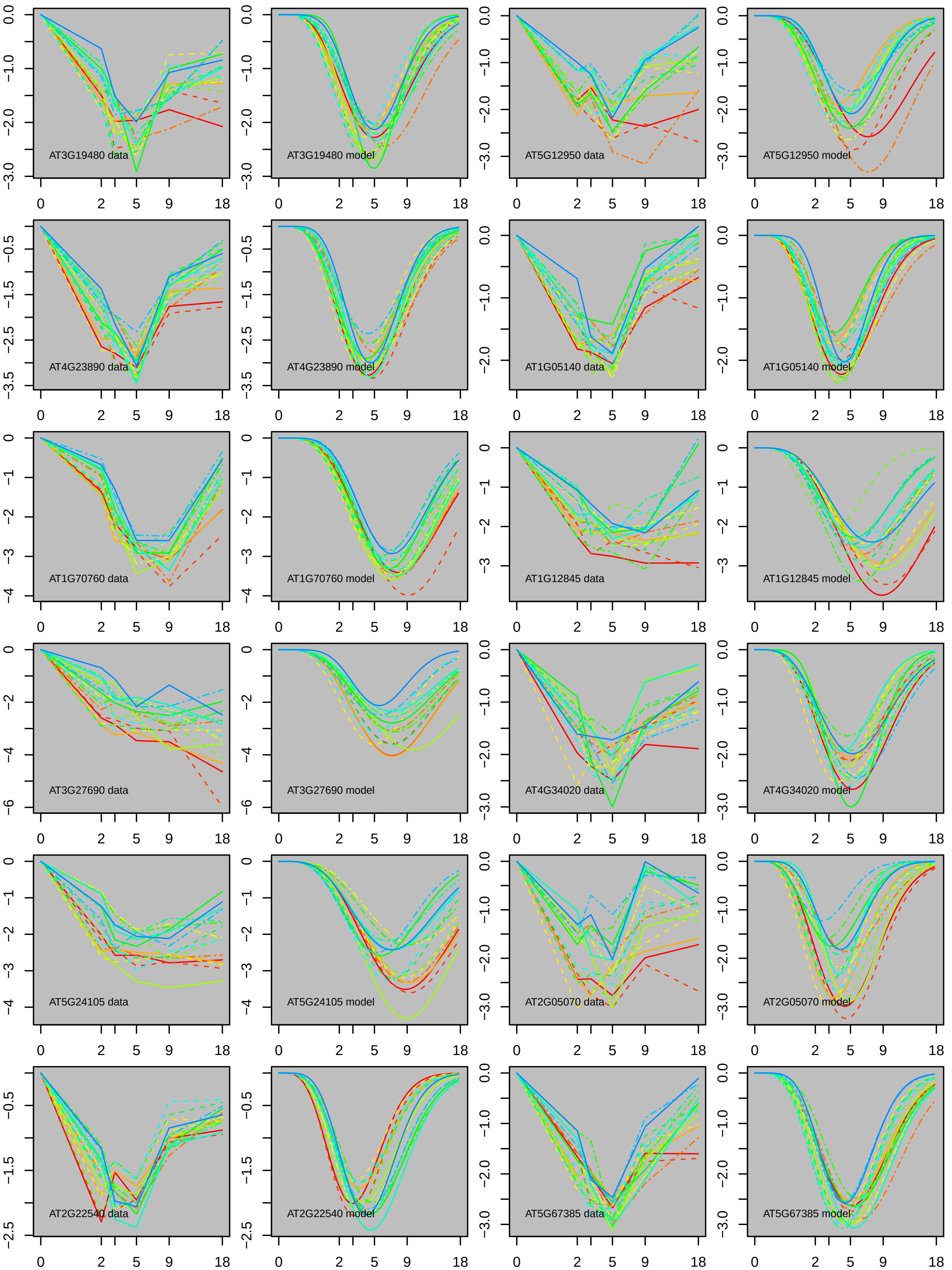

AT3G19480 data
AT3G19480 model
AT5G12950 data
AT5G12950 model
AT4G23890 data
AT4G23890 model
AT1G05140 data
AT1G05140 model
AT1G70760 data
AT1G70760 model
AT1G12845 data
AT1G12845 model
AT3G27690 data
AT3G27690 model
AT4G34020 data
AT4G34020 model
AT5G24105 data
AT5G24105 model
AT2G05070 data
AT2G05070 model
AT2G22540 data
AT2G22540 model
AT5G67385 data
AT5G67385 model

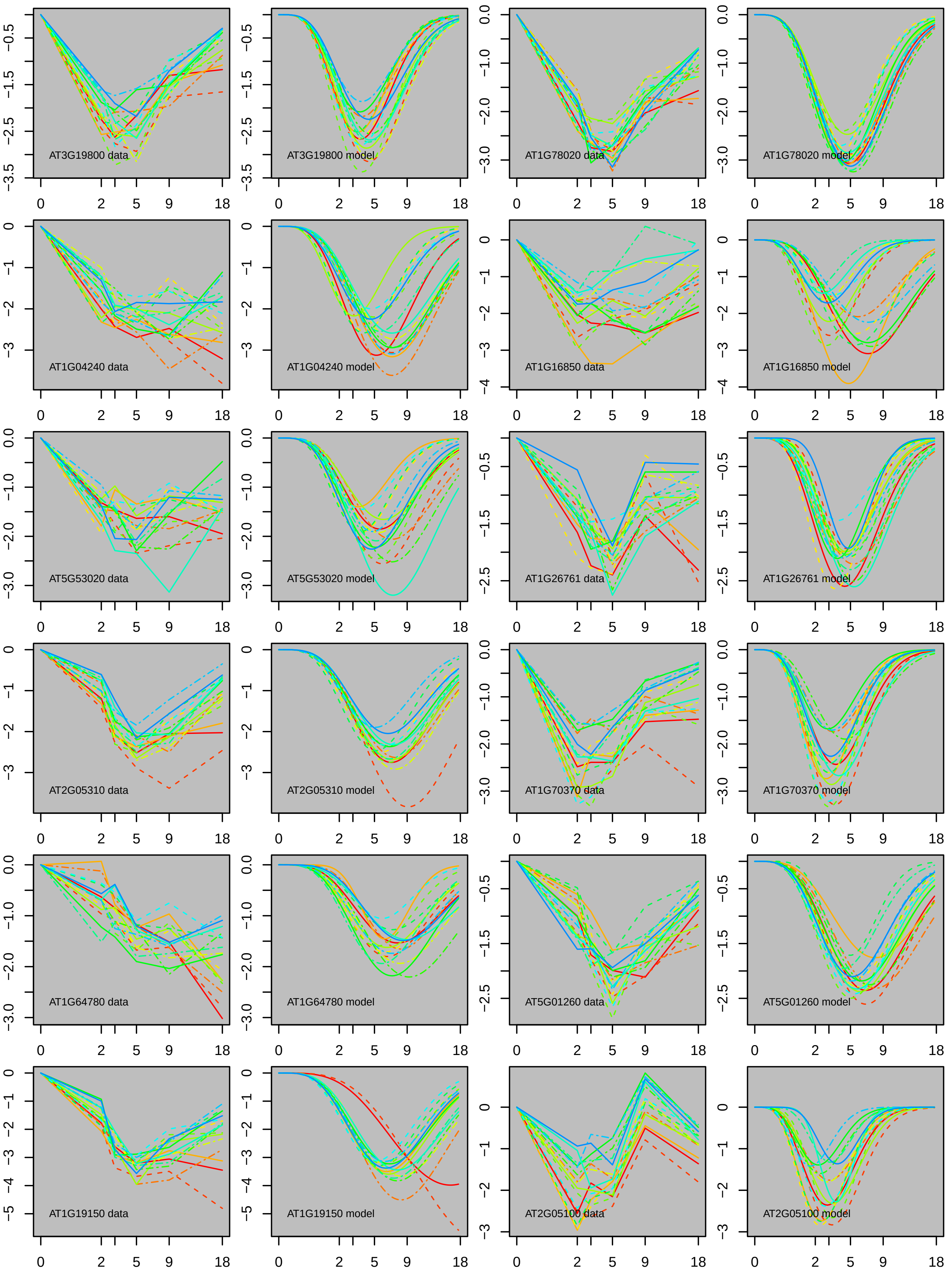
AT3G19800 data
AT3G19800 model
AT1G78020 data
AT1G78020 model
AT1G04240 data
AT1G04240 model
AT1G16850 data
AT1G16850 model
AT5G53020 data
AT5G53020 model
AT1G26761 data
AT1G26761 model
AT2G05310 data
AT2G05310 model
AT1G70370 data
AT1G70370 model
AT1G64780 data
AT1G64780 model
AT5G01260 data
AT5G01260 model
AT1G19150 data
AT1G19150 model
AT2G05100 data
AT2G05100 model

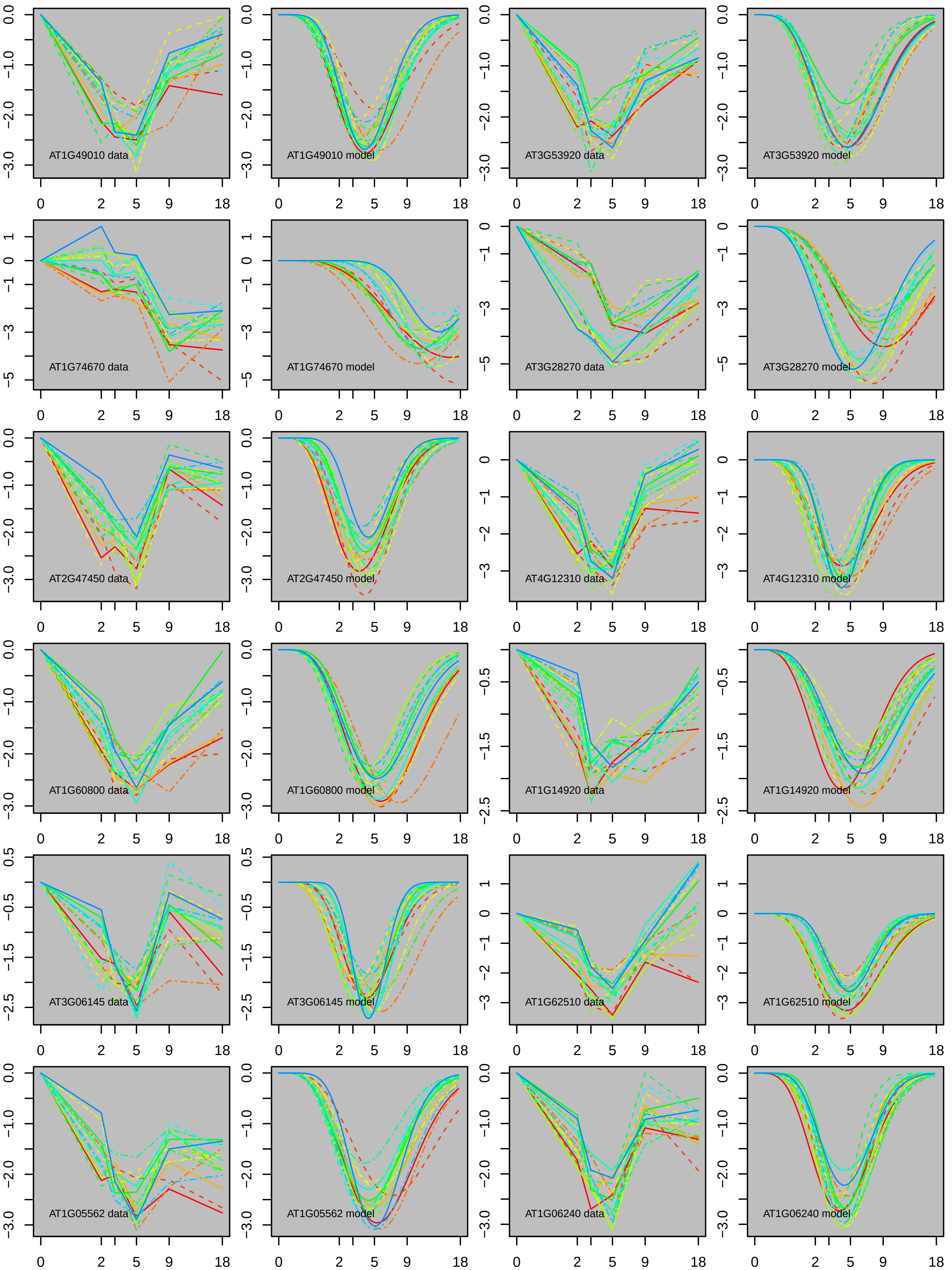

AT1G49010 data
AT1G49010 model
AT3G53920 data
AT3G53920 model
AT1G74670 data
AT1G74670 model
AT3G28270 data
AT3G28270 model
AT2G47450 data
AT2G47450 model
AT4G12310 data
AT4G12310 model
AT1G60800 data
AT1G60800 model
AT1G14920 data
AT1G14920 model
AT3G06145 data
AT3G06145 model
AT1G62510 data
AT1G62510 model
AT1G05562 data
AT1G05562 model
AT1G06240 data
AT1G06240 model
0 2 5 9 18

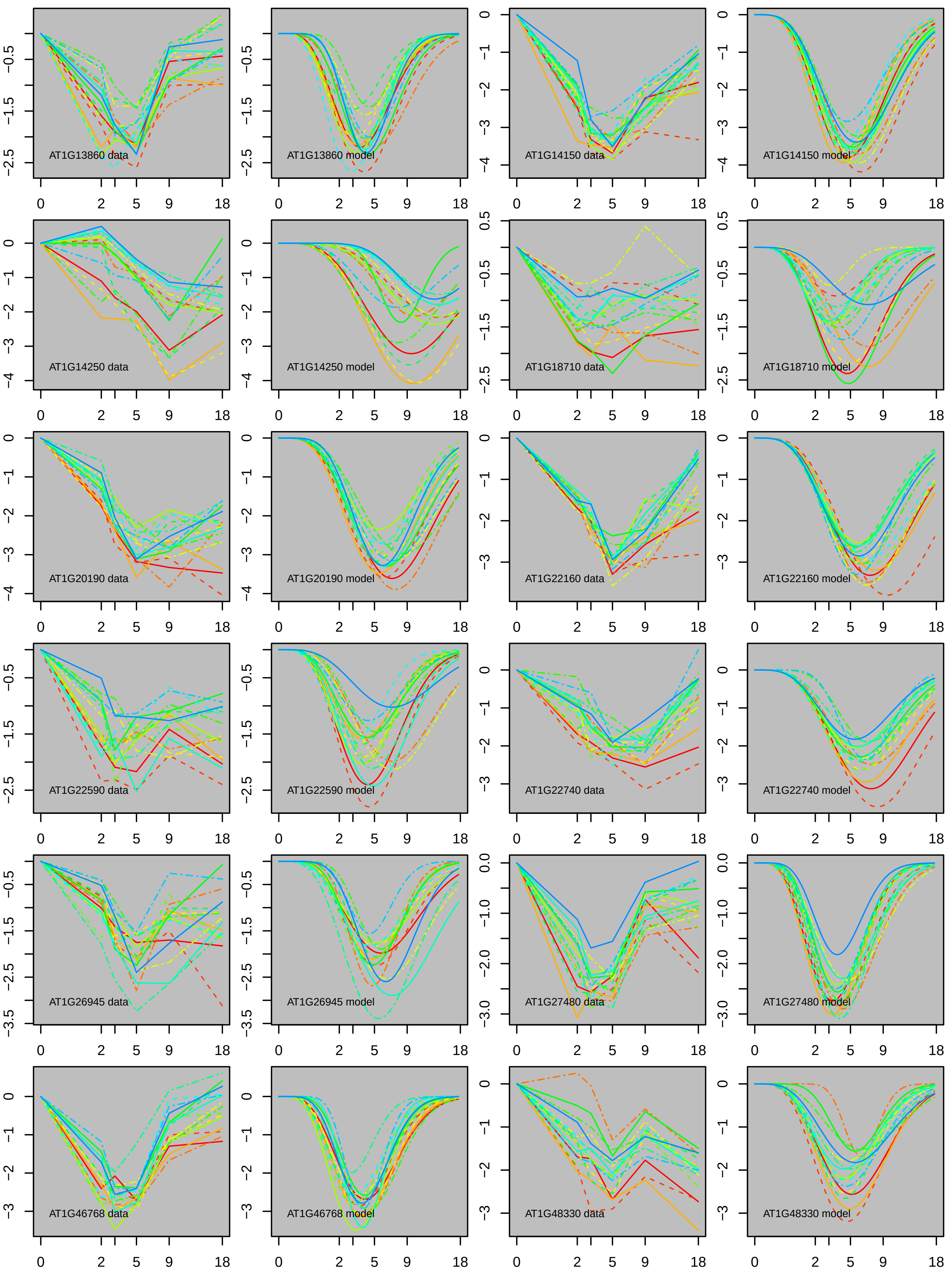
AT1G13860 data
AT1G13860 model
AT1G14150 data
AT1G14150 model
AT1G14250 data
AT1G14250 model
AT1G18710 data
AT1G18710 model
AT1G20190 data
AT1G20190 model
AT1G22160 data
AT1G22160 model
AT1G22590 data
AT1G22590 model
AT1G22740 data
AT1G22740 model
AT1G26945 data
AT1G26945 model
AT1G27480 data
AT1G27480 model
AT1G46768 data
AT1G46768 model
AT1G48330 data
AT1G48330 model

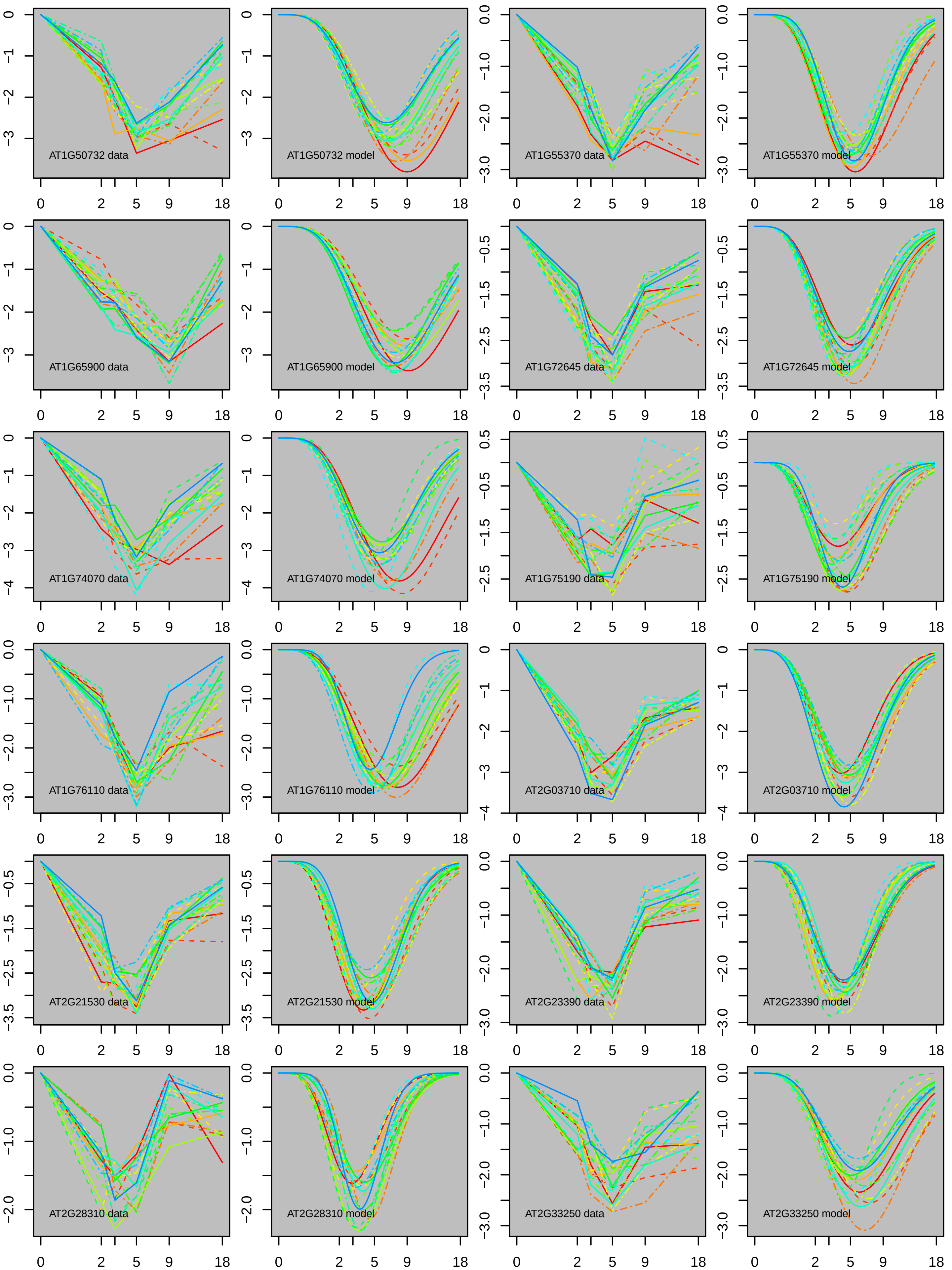
AT1G50732 data
AT1G50732 model
AT1G55370 data
AT1G55370 model
AT1G65900 data
AT1G65900 model
AT1G72645 data
AT1G72645 model
AT1G74070 data
AT1G74070 model
AT1G75190 data
AT1G75190 model
AT1G76110 data
AT1G76110 model
AT2G03710 data
AT2G03710 model
AT2G21530 data
AT2G21530 model
AT2G23390 data
AT2G23390 model
AT2G28310 data
AT2G28310 model
AT2G33250 data
AT2G33250 model
0
2
5
9
18

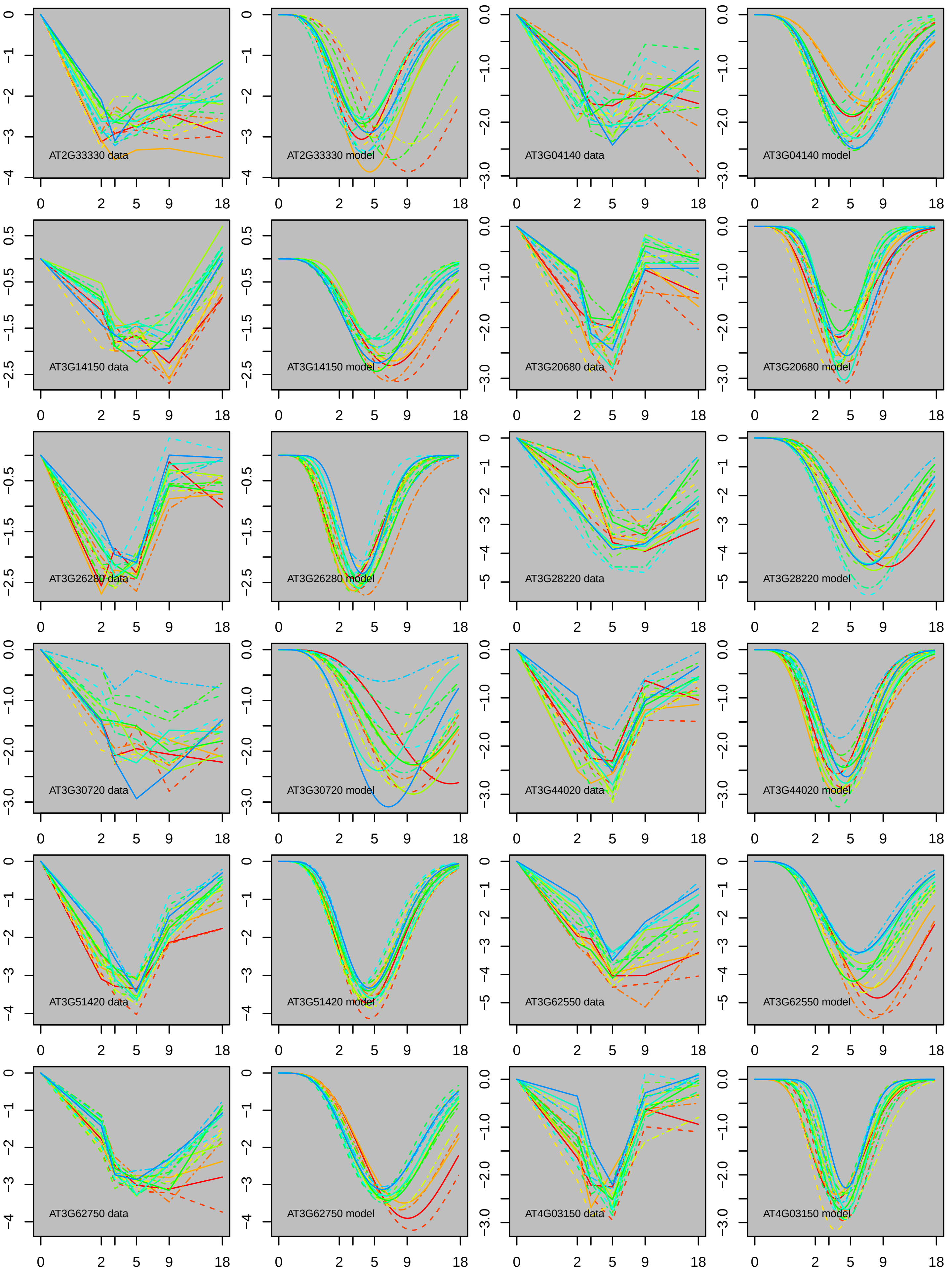
AT2G33330 data
AT2G33330 model
AT3G04140 data
AT3G04140 model
AT3G14150 data
AT3G14150 model
AT3G20680 data
AT3G20680 model
AT3G26280 data
AT3G26280 model
AT3G28220 data
AT3G28220 model
AT3G30720 data
AT3G30720 model
AT3G44020 data
AT3G44020 model
AT3G51420 data
AT3G51420 model
AT3G62550 data
AT3G62550 model
AT3G62750 data
AT3G62750 model
AT4G03150 data
AT4G03150 model

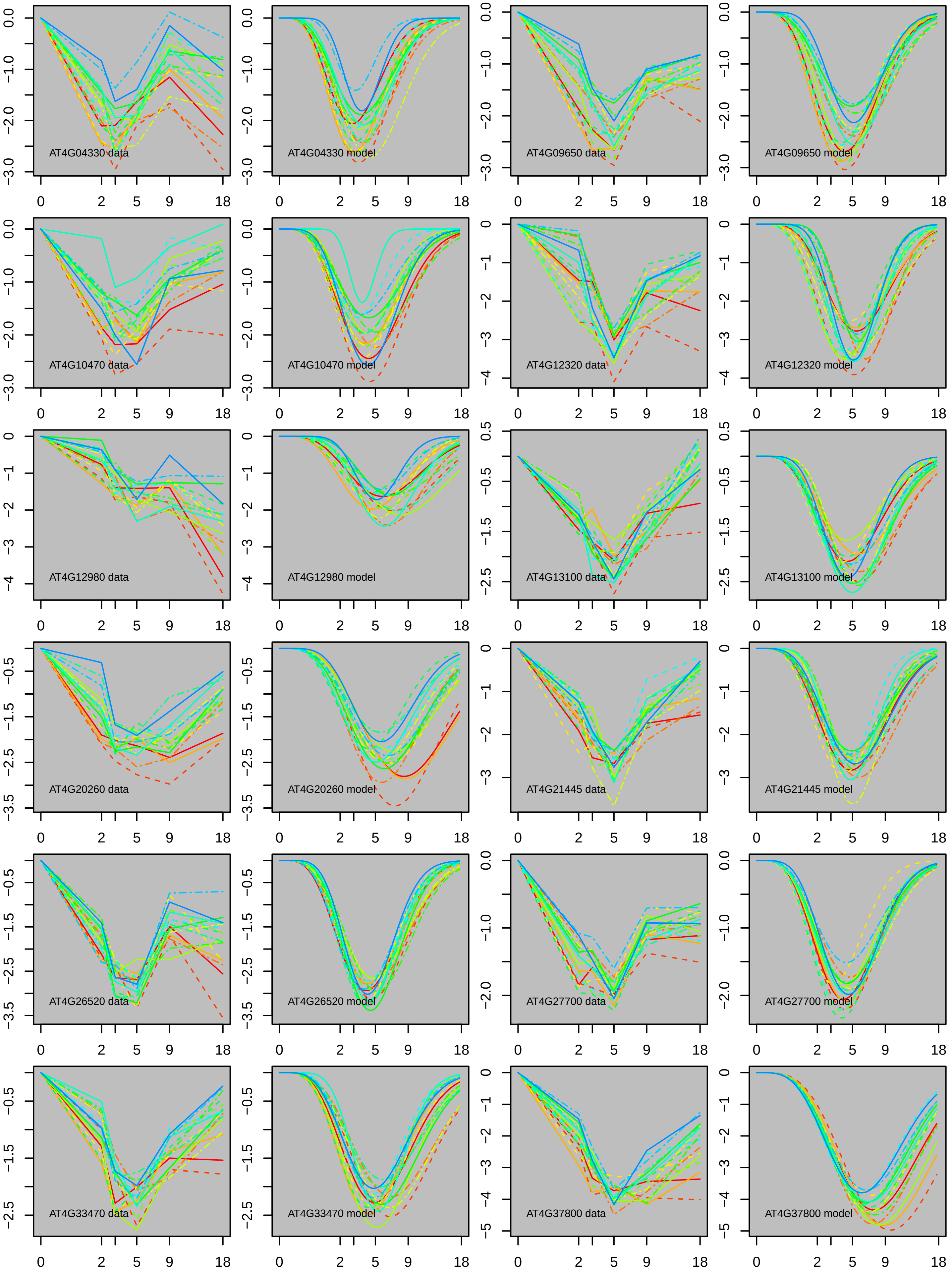

AT4G04330 data
AT4G04330 model
AT4G09650 data
AT4G09650 model
AT4G10470 data
AT4G10470 model
AT4G12320 data
AT4G12320 model
AT4G12980 data
AT4G12980 model
AT4G13100 data
AT4G13100 model
AT4G20260 data
AT4G20260 model
AT4G21445 data
AT4G21445 model
AT4G26520 data
AT4G26520 model
AT4G27700 data
AT4G27700 model
AT4G33470 data
AT4G33470 model
AT4G37800 data
AT4G37800 model

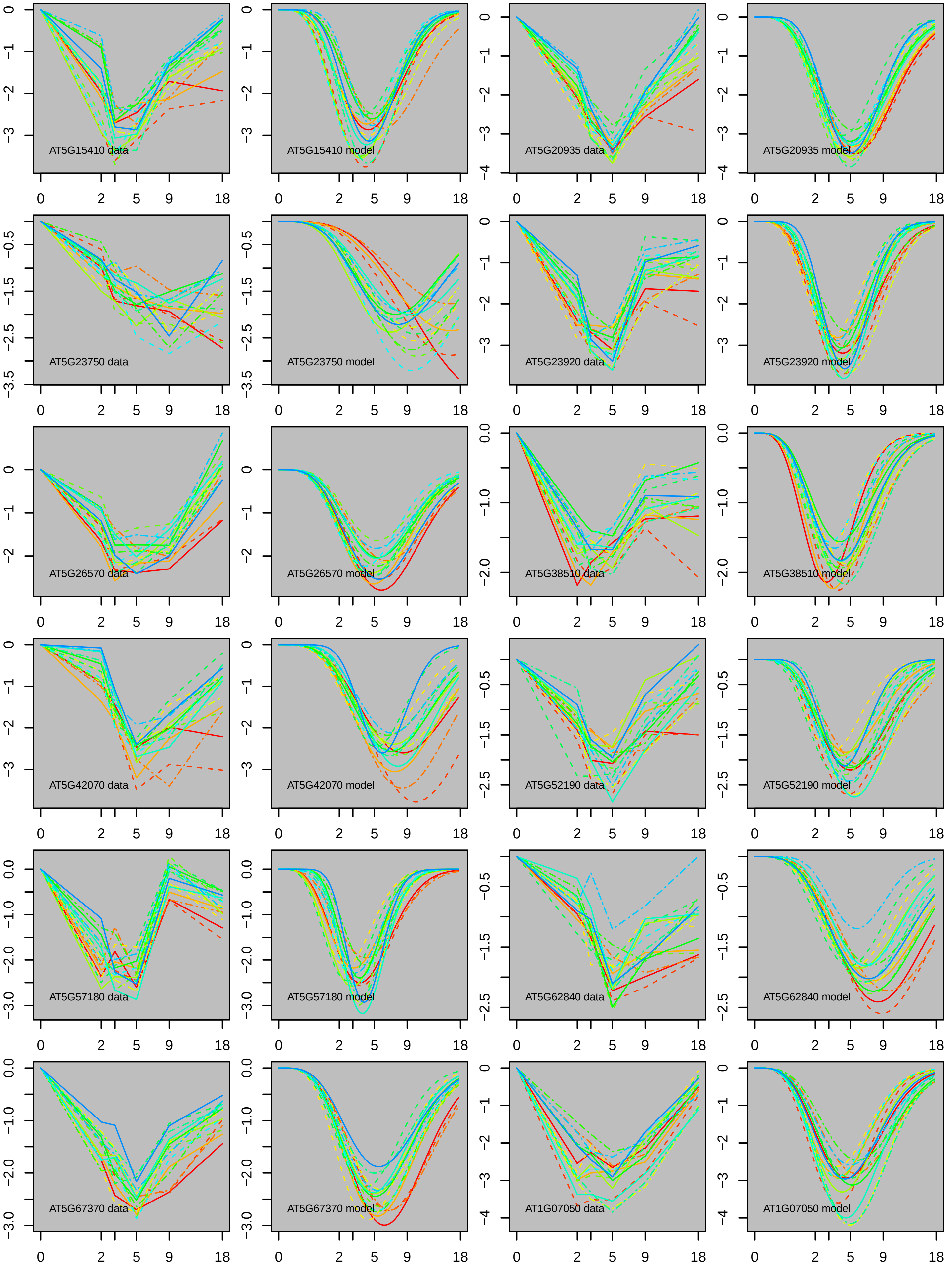
AT5G15410 data
AT5G15410 model
AT5G20935 data
AT5G20935 model
AT5G23750 data
AT5G23750 model
AT5G23920 data
AT5G23920 model
AT5G26570 data
AT5G26570 model
AT5G38510 data
AT5G38510 model
AT5G42070 data
AT5G42070 model
AT5G52190 data
AT5G52190 model
AT5G57180 data
AT5G57180 model
AT5G62840 data
AT5G62840 model
AT5G67370 data
AT5G67370 model
AT1G07050 data
AT1G07050 model
0
2
5
9
18

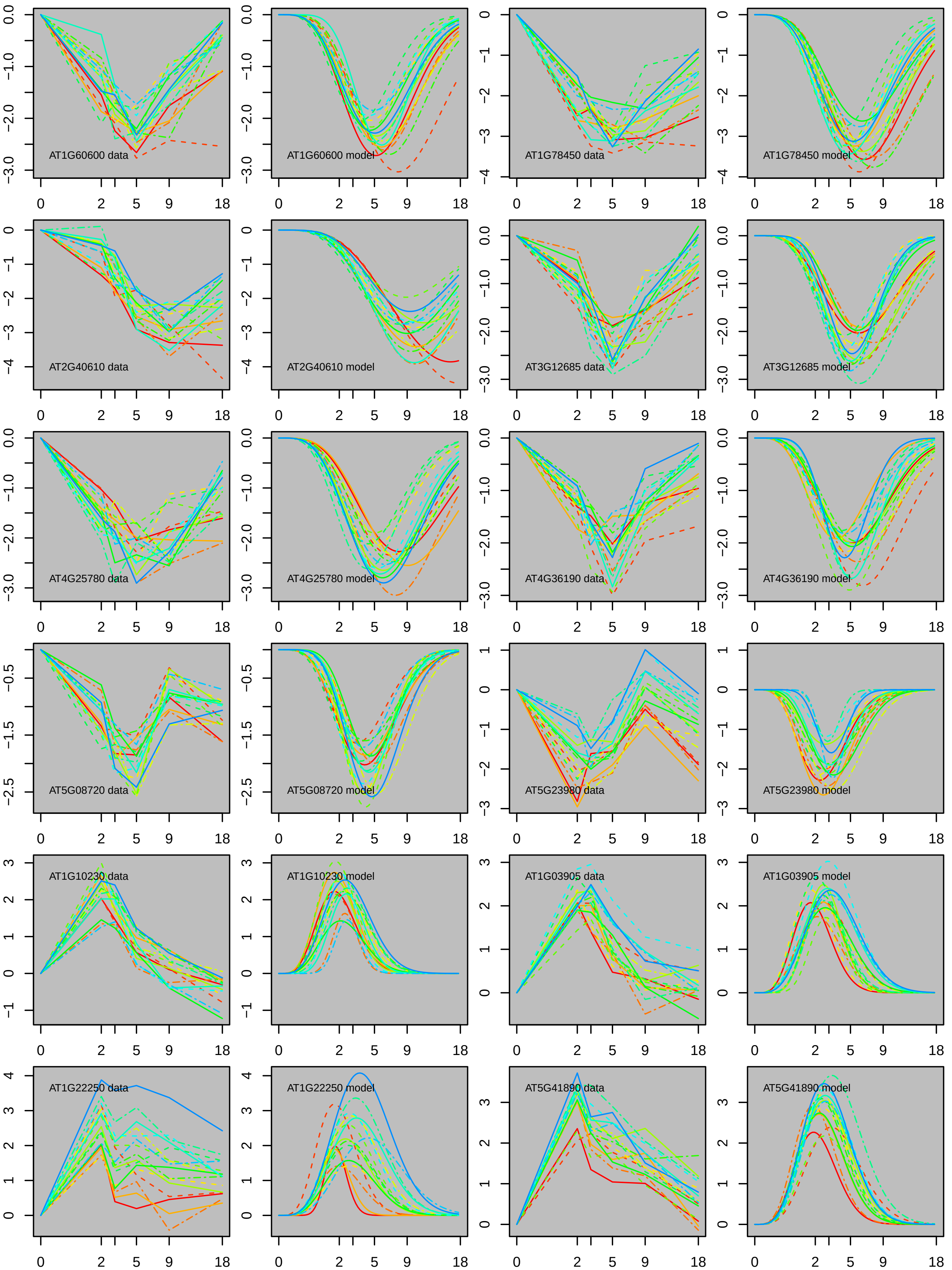
AT1G60600 data
AT1G60600 model
AT1G78450 data
AT1G78450 model
AT2G40610 data
AT2G40610 model
AT3G12685 data
AT3G12685 model
AT4G25780 data
AT4G25780 model
AT4G36190 data
AT4G36190 model
AT5G08720 data
AT5G08720 model
AT5G23980 data
AT5G23980 model
AT1G10230 data
AT1G10230 model
AT1G03905 data
AT1G03905 model
AT1G22250 data
AT1G22250 model
AT5G41890 data
AT5G41890 model
0 2 5 9 18

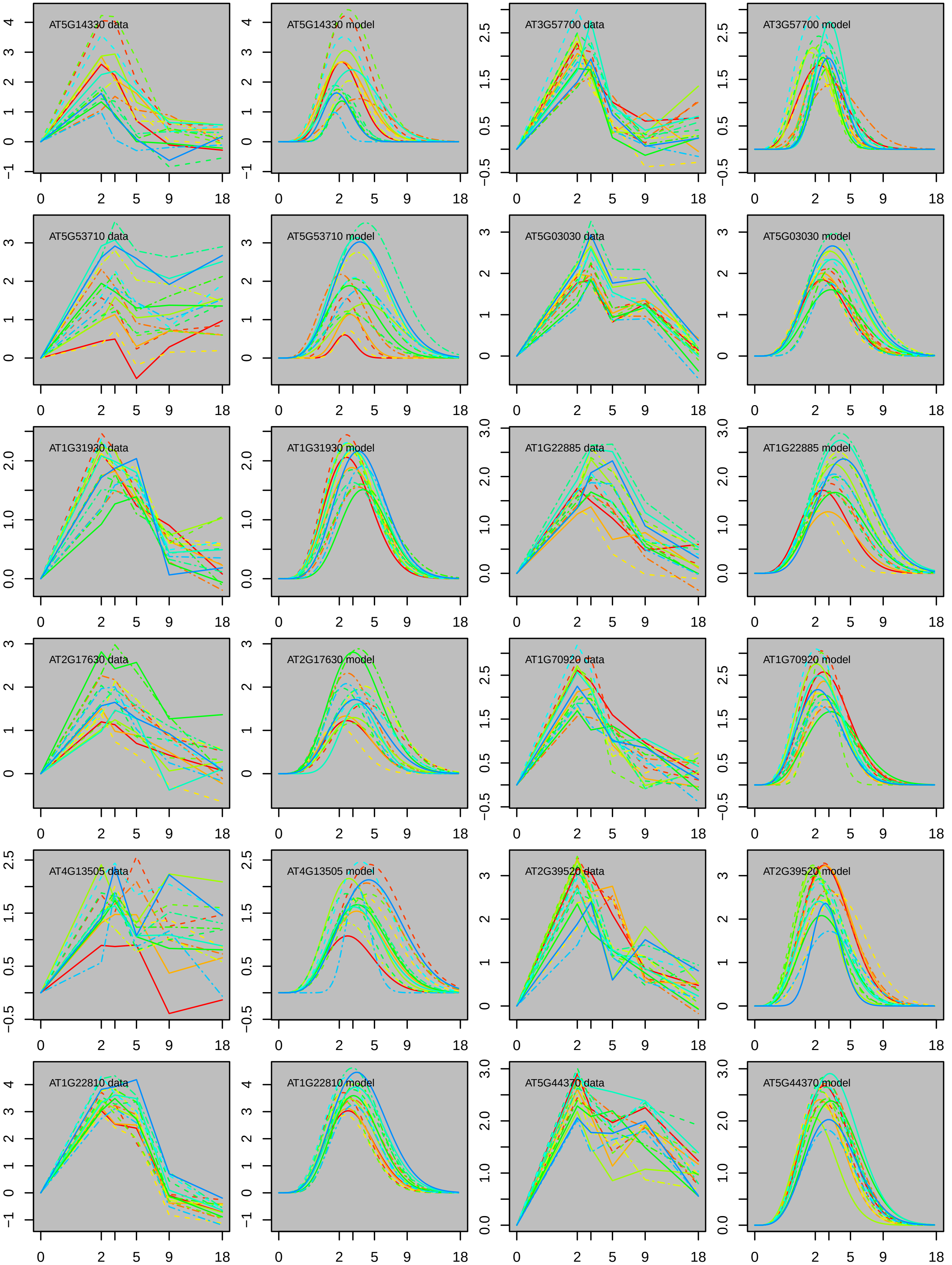
AT5G14330 data
AT5G14330 model
AT3G57700 data
AT3G57700 model
AT5G53710 data
AT5G53710 model
AT5G03030 data
AT5G03030 model
AT1G31930 data
AT1G31930 model
AT1G22885 data
AT1G22885 model
AT2G17630 data
AT2G17630 model
AT1G70920 data
AT1G70920 model
AT4G13505 data
AT4G13505 model
AT2G39520 data
AT2G39520 model
AT1G22810 data
AT1G22810 model
AT5G44370 data
AT5G44370 model
0
2
5
9
18

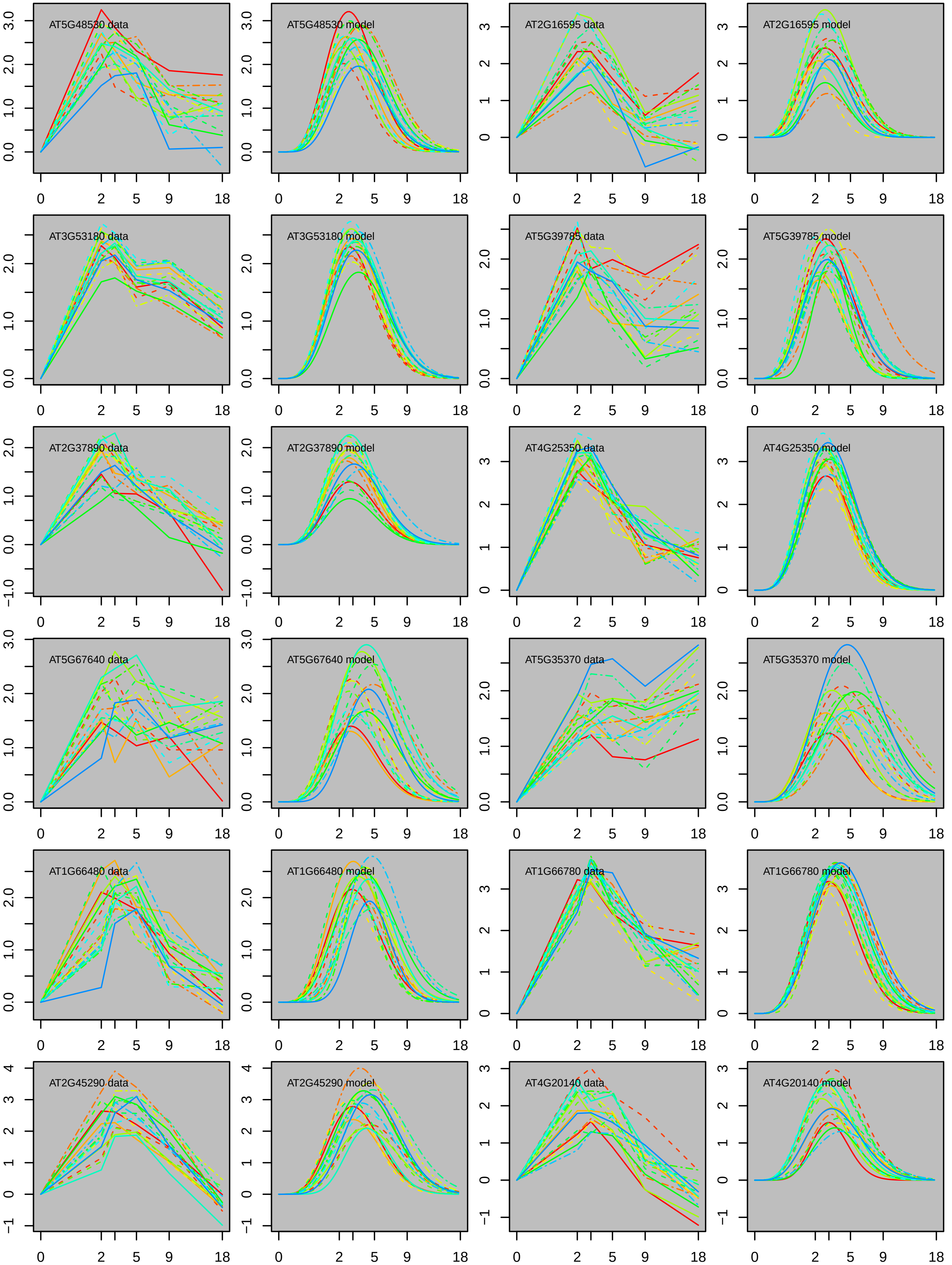
AT5G48530 data
AT5G48530 model
AT2G16595 data
AT2G16595 model
AT3G53180 data
AT3G53180 model
AT5G39785 data
AT5G39785 model
AT2G37890 data
AT2G37890 model
AT4G25350 data
AT4G25350 model
AT5G67640 data
AT5G67640 model
AT5G35370 data
AT5G35370 model
AT1G66480 data
AT1G66480 model
AT1G66780 data
AT1G66780 model
AT2G45290 data
AT2G45290 model
AT4G20140 data
AT4G20140 model
0 2 5 9 18

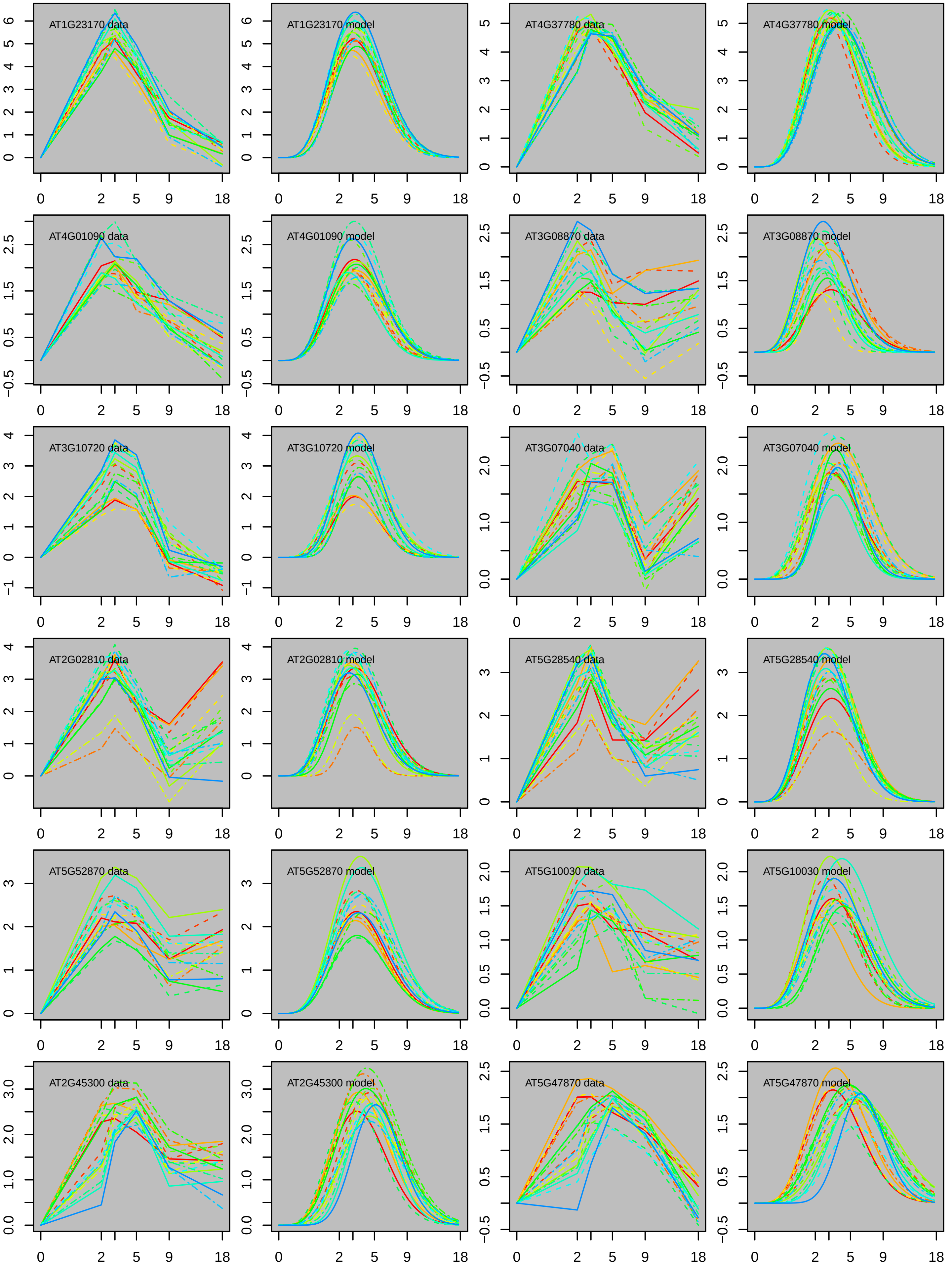
AT1G23170 data
AT1G23170 model
AT4G37780 data
AT4G37780 model
AT4G01090 data
AT4G01090 model
AT3G08870 data
AT3G08870 model
AT3G10720 data
AT3G10720 model
AT3G07040 data
AT3G07040 model
AT2G02810 data
AT2G02810 model
AT5G28540 data
AT5G28540 model
AT5G52870 data
AT5G52870 model
AT5G10030 data
AT5G10030 model
AT2G45300 data
AT2G45300 model
AT5G47870 data
AT5G47870 model

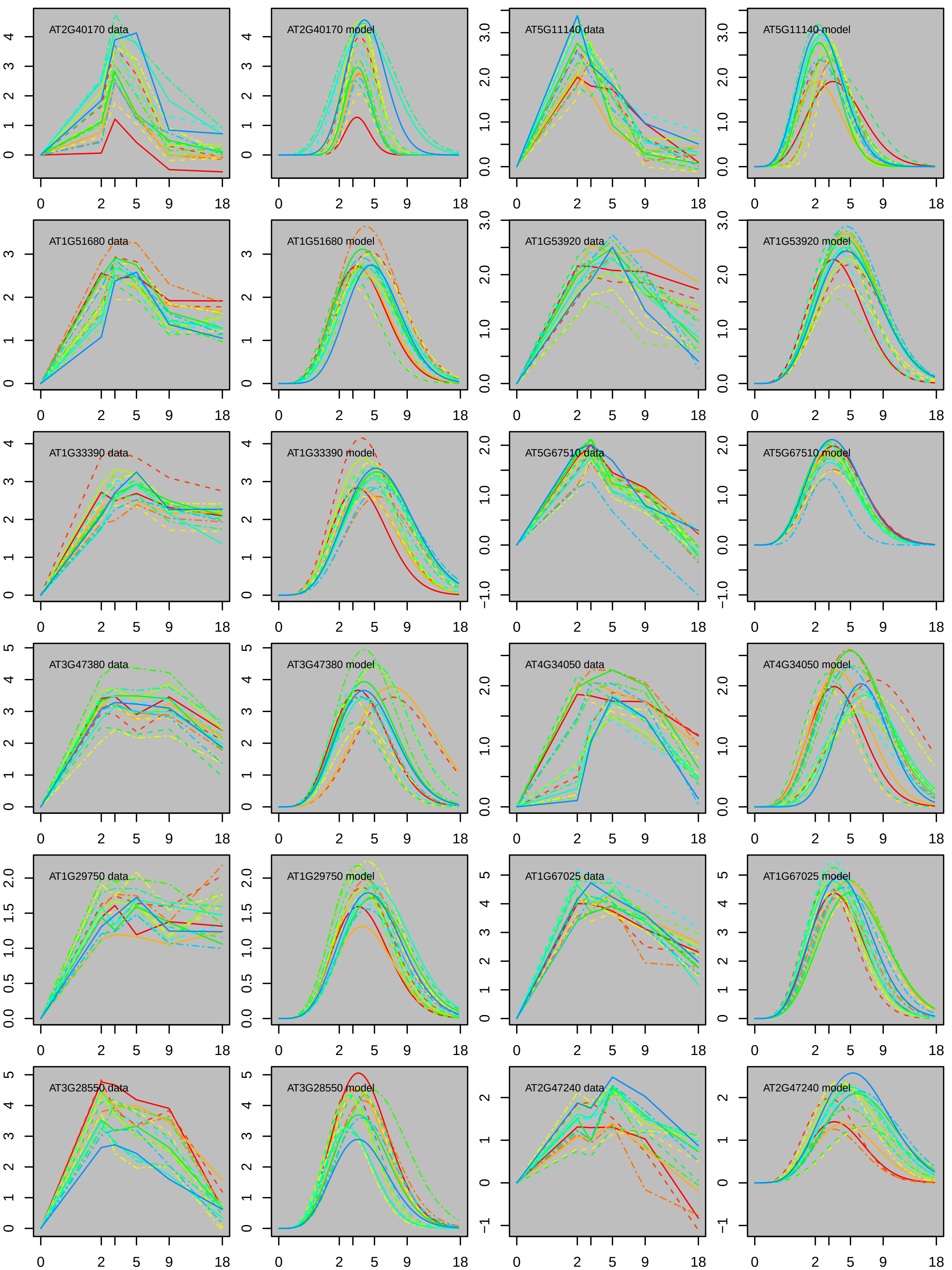

AT2G40170 data
AT2G40170 model
AT5G11140 data
AT5G11140 model
AT1G51680 data
AT1G51680 model
AT1G53920 data
AT1G53920 model
AT1G33390 data
AT1G33390 model
AT5G67510 data
AT5G67510 model
AT3G47380 data
AT3G47380 model
AT4G34050 data
AT4G34050 model
AT1G29750 data
AT1G29750 model
AT1G67025 data
AT1G67025 model
AT3G28550 data
AT3G28550 model
AT2G47240 data
AT2G47240 model
0 2 5 9 18

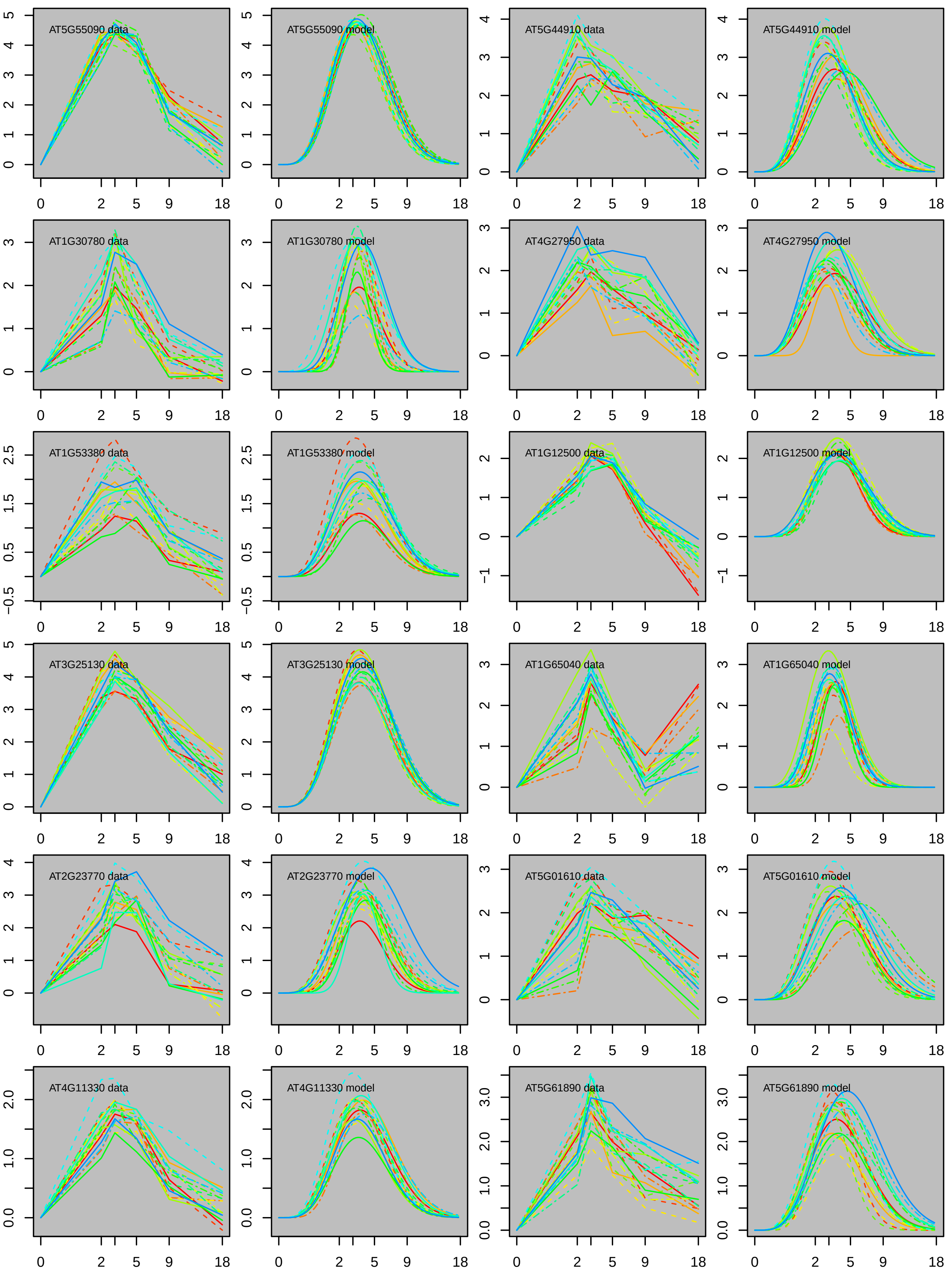
AT5G55090 data
AT5G55090 model
AT5G44910 data
AT5G44910 model
AT1G30780 data
AT1G30780 model
AT4G27950 data
AT4G27950 model
AT1G53380 data
AT1G53380 model
AT1G12500 data
AT1G12500 model
AT3G25130 data
AT3G25130 model
AT1G65040 data
AT1G65040 model
AT2G23770 data
AT2G23770 model
AT5G01610 data
AT5G01610 model
AT4G11330 data
AT4G11330 model
AT5G61890 data
AT5G61890 model
0 2 5 9 18

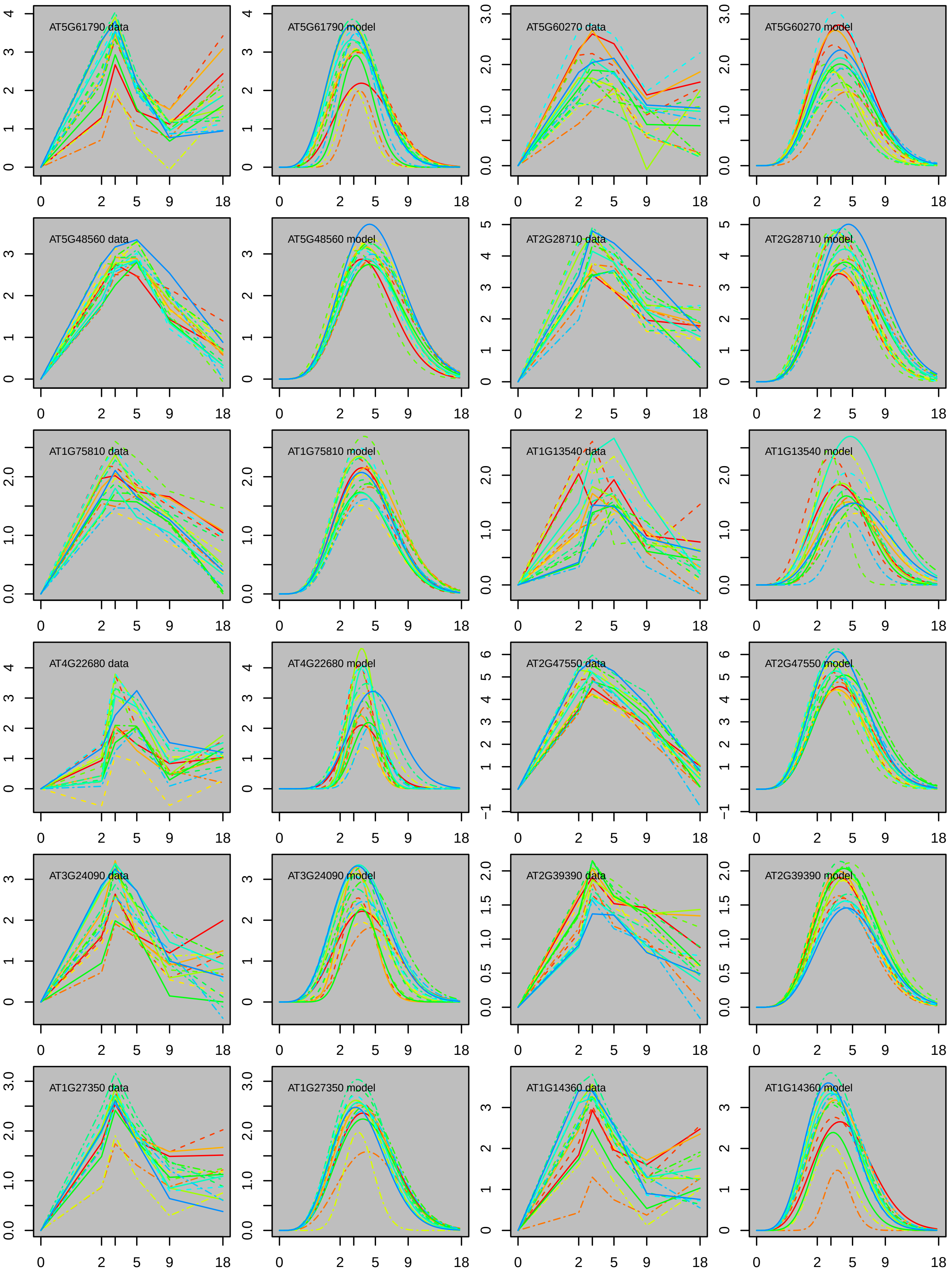

AT5G61790 data
AT5G61790 model
AT5G60270 data
AT5G60270 model
AT5G48560 data
AT5G48560 model
AT2G28710 data
AT2G28710 model
AT1G75810 data
AT1G75810 model
AT1G13540 data
AT1G13540 model
AT4G22680 data
AT4G22680 model
AT2G47550 data
AT2G47550 model
AT3G24090 data
AT3G24090 model
AT2G39390 data
AT2G39390 model
AT1G27350 data
AT1G27350 model
AT1G14360 data
AT1G14360 model

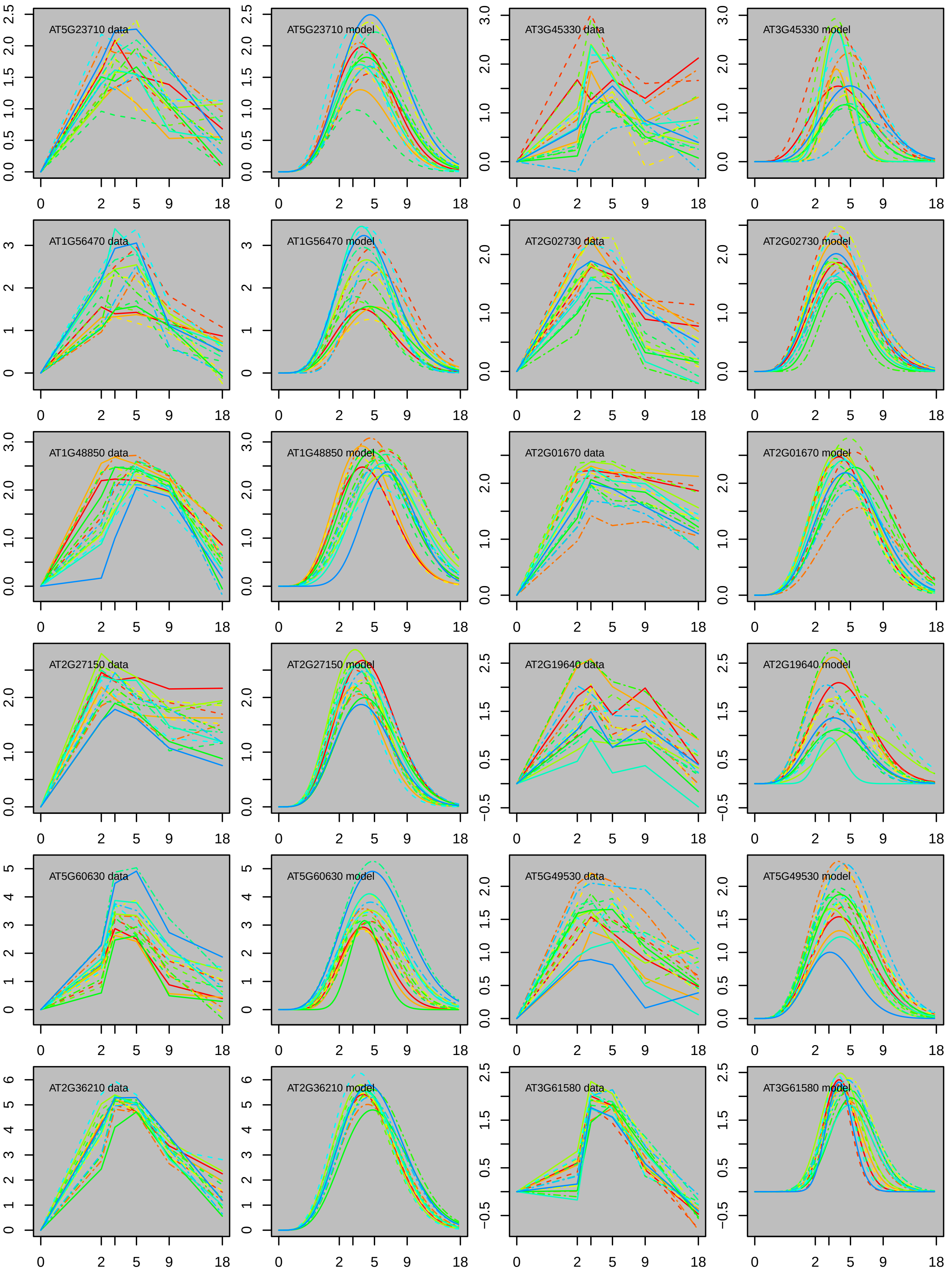
AT5G23710 data
AT5G23710 model
AT3G45330 data
AT3G45330 model
AT1G56470 data
AT1G56470 model
AT2G02730 data
AT2G02730 model
AT1G48850 data
AT1G48850 model
AT2G01670 data
AT2G01670 model
AT2G27150 data
AT2G27150 model
AT2G19640 data
AT2G19640 model
AT5G60630 data
AT5G60630 model
AT5G49530 data
AT5G49530 model
AT2G36210 data
AT2G36210 model
AT3G61580 data
AT3G61580 model
0 2 5 9 18

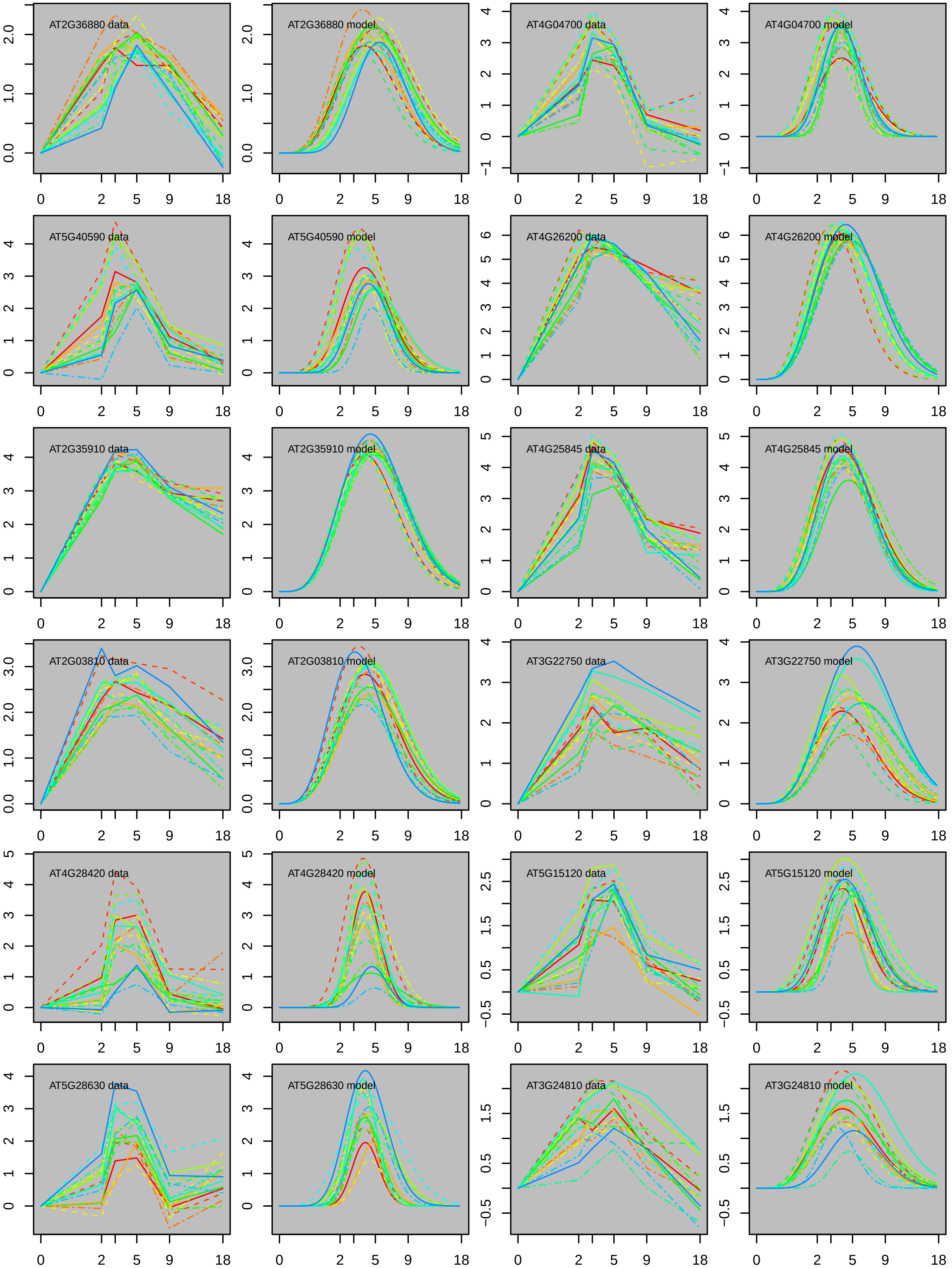
AT2G36880 data
AT2G36880 model
AT4G04700 data
AT4G04700 model
AT5G40590 data
AT5G40590 model
AT4G26200 data
AT4G26200 model
AT2G35910 data
AT2G35910 model
AT4G25845 data
AT4G25845 model
AT2G03810 data
AT2G03810 model
AT3G22750 data
AT3G22750 model
AT4G28420 data
AT4G28420 model
AT5G15120 data
AT5G15120 model
AT5G28630 data
AT5G28630 model
AT3G24810 data
AT3G24810 model

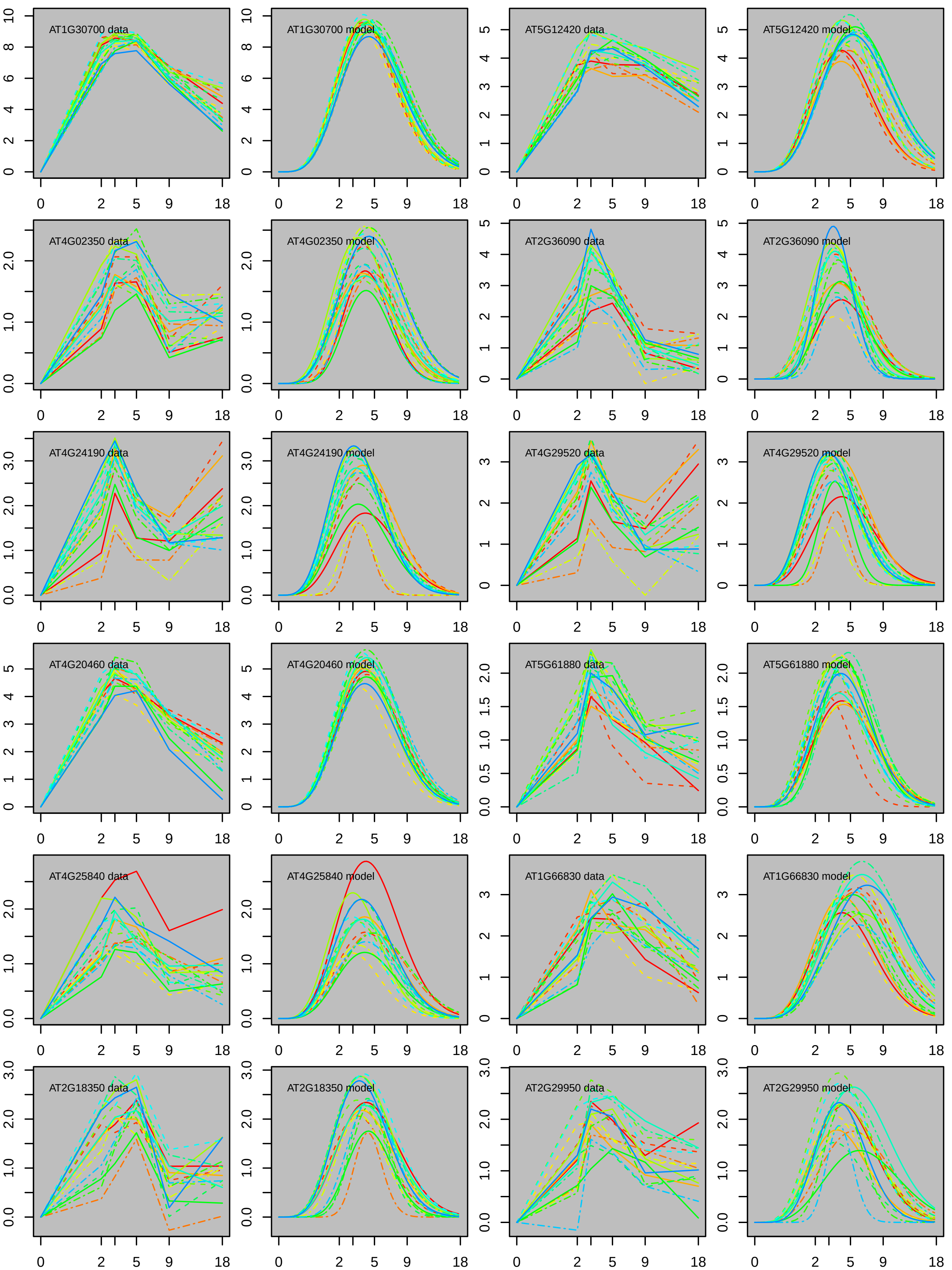
AT1G30700 data
AT1G30700 model
AT5G12420 data
AT5G12420 model
AT4G02350 data
AT4G02350 model
AT2G36090 data
AT2G36090 model
AT4G24190 data
AT4G24190 model
AT4G29520 data
AT4G29520 model
AT4G20460 data
AT4G20460 model
AT5G61880 data
AT5G61880 model
AT4G25840 data
AT4G25840 model
AT1G66830 data
AT1G66830 model
AT2G18350 data
AT2G18350 model
AT2G29950 data
AT2G29950 model

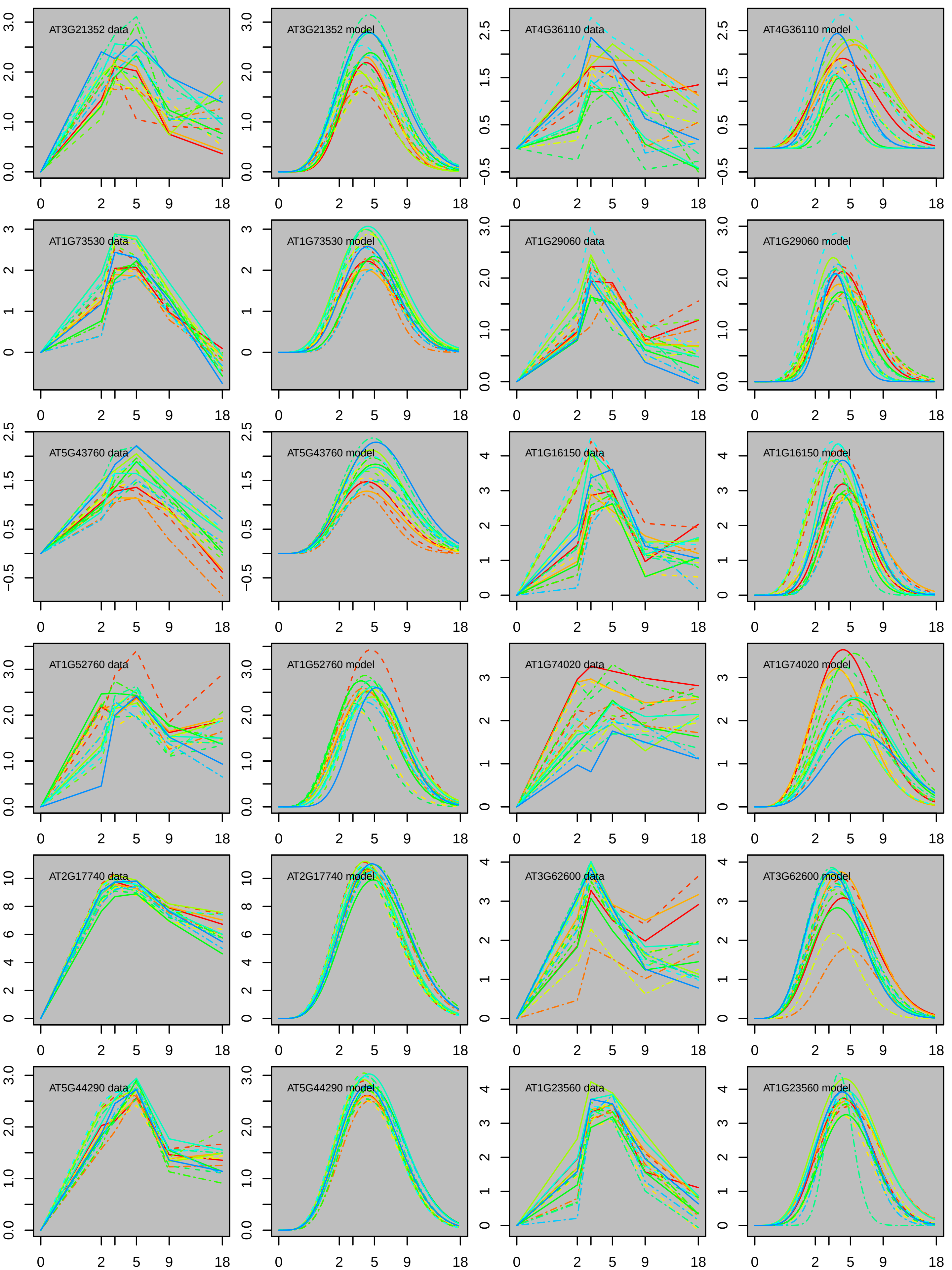

AT3G21352 data
AT3G21352 model
AT4G36110 data
AT4G36110 model
AT1G73530 data
AT1G73530 model
AT1G29060 data
AT1G29060 model
AT5G43760 data
AT5G43760 model
AT1G16150 data
AT1G16150 model
AT1G52760 data
AT1G52760 model
AT1G74020 data
AT1G74020 model
AT2G17740 data
AT2G17740 model
AT3G62600 data
AT3G62600 model
AT5G44290 data
AT5G44290 model
AT1G23560 data
AT1G23560 model
0 2 5 9 18

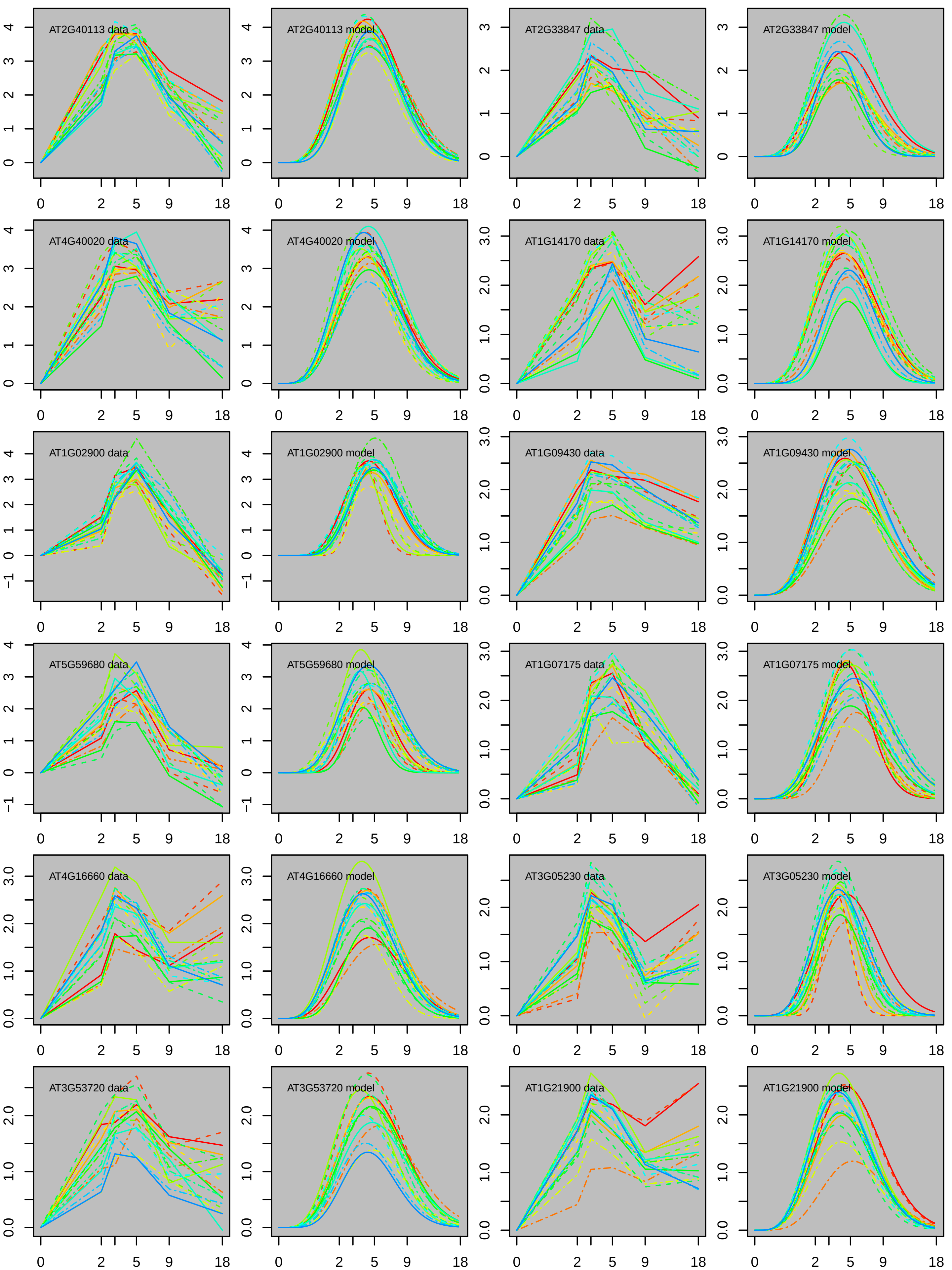
AT2G40113 data
AT2G40113 model
AT2G33847 data
AT2G33847 model
AT4G40020 data
AT4G40020 model
AT1G14170 data
AT1G14170 model
AT1G02900 data
AT1G02900 model
AT1G09430 data
AT1G09430 model
AT5G59680 data
AT5G59680 model
AT1G07175 data
AT1G07175 model
AT4G16660 data
AT4G16660 model
AT3G05230 data
AT3G05230 model
AT3G53720 data
AT3G53720 model
AT1G21900 data
AT1G21900 model
0
2
5
9
18

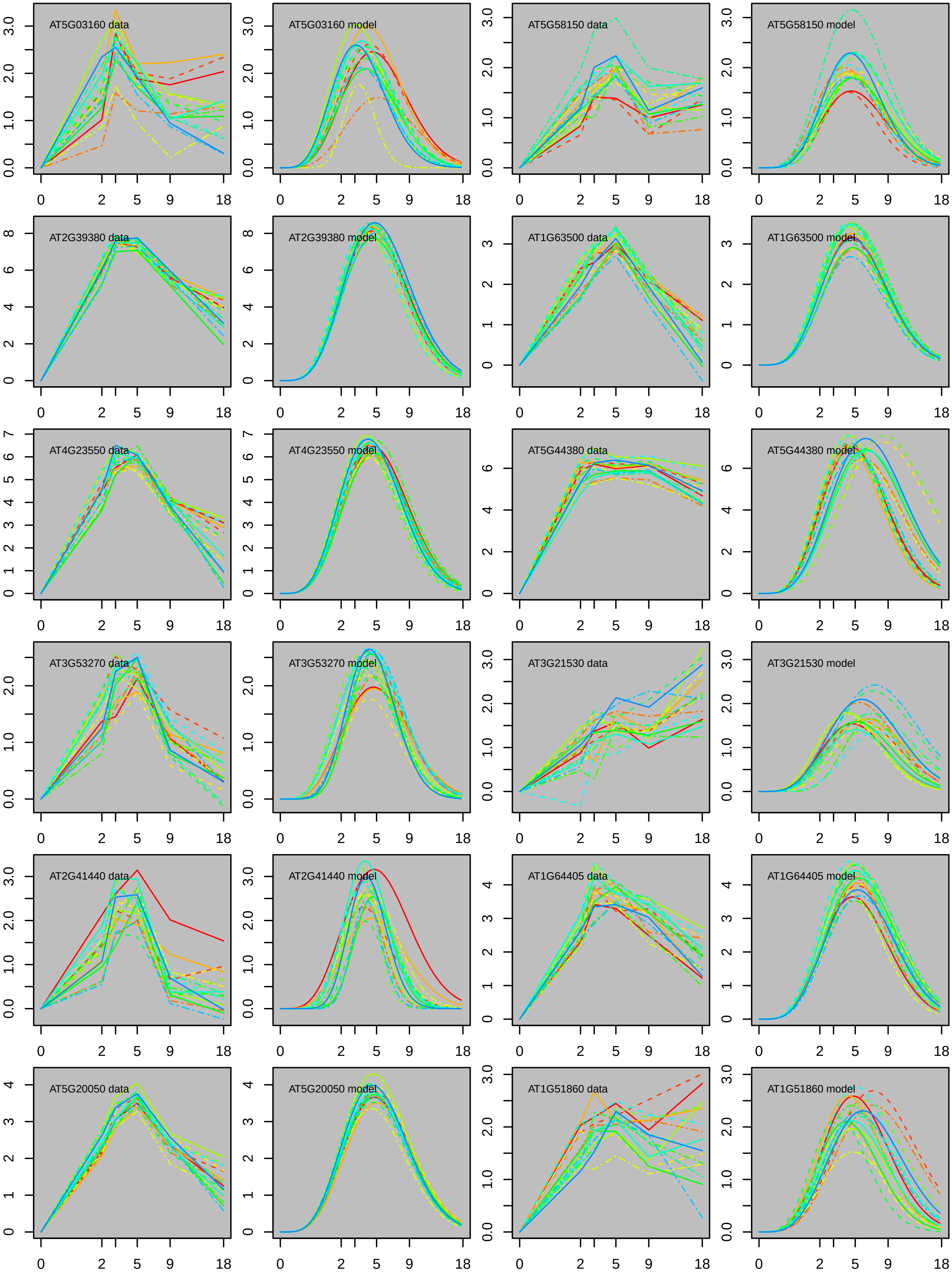
AT5G03160 data
AT5G03160 model
AT5G58150 data
AT5G58150 model
AT2G39380 data
AT2G39380 model
AT1G63500 data
AT1G63500 model
AT4G23550 data
AT4G23550 model
AT5G44380 data
AT5G44380 model
AT3G53270 data
AT3G53270 model
AT3G21530 data
AT3G21530 model
AT2G41440 data
AT2G41440 model
AT1G64405 data
AT1G64405 model
AT5G20050 data
AT5G20050 model
AT1G51860 data
AT1G51860 model
0
2
5
9
18

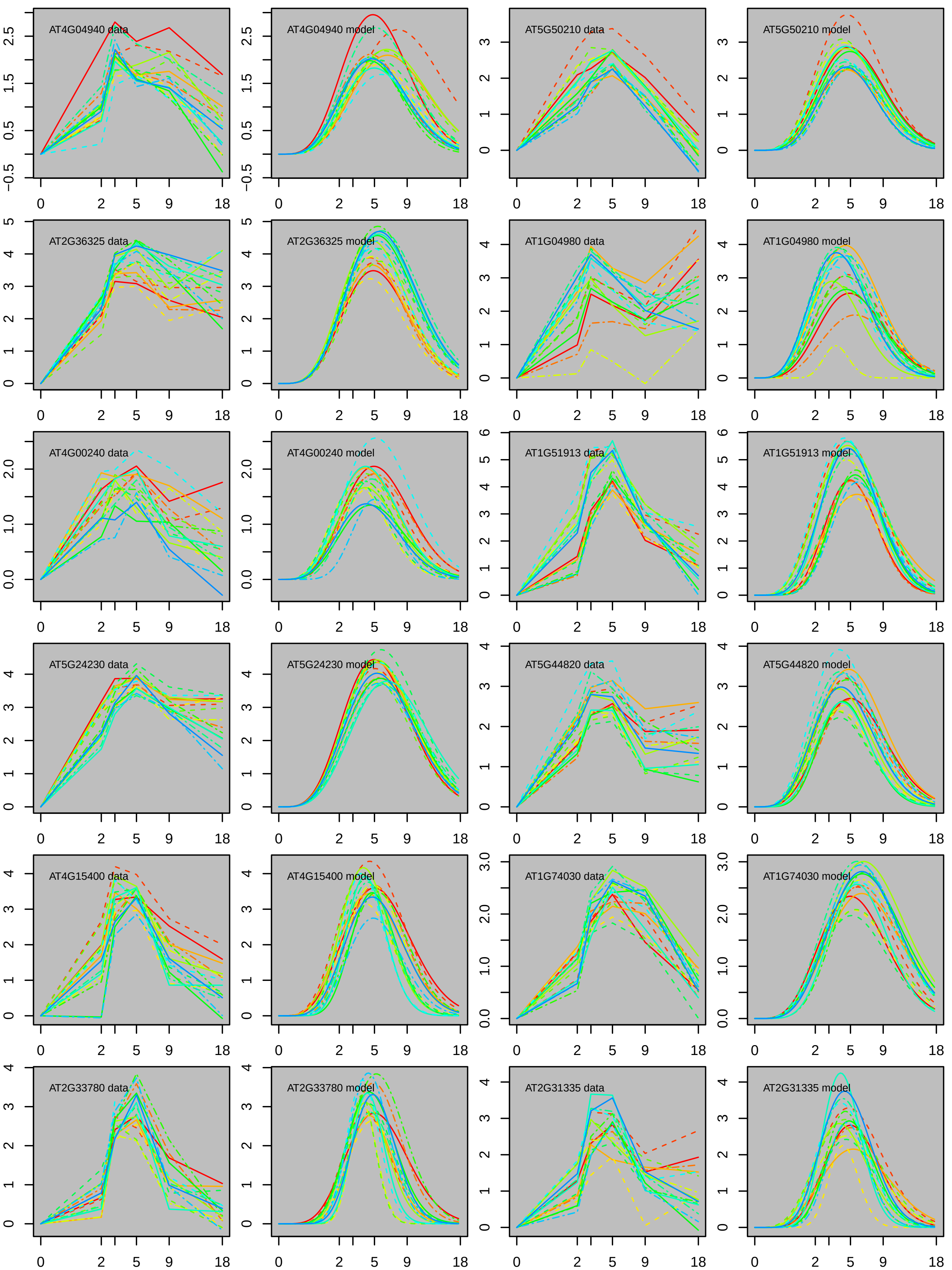
AT4G04940 data
AT4G04940 model
AT5G50210 data
AT5G50210 model
AT2G36325 data
AT2G36325 model
AT1G04980 data
AT1G04980 model
AT4G00240 data
AT4G00240 model
AT1G51913 data
AT1G51913 model
AT5G24230 data
AT5G24230 model
AT5G44820 data
AT5G44820 model
AT4G15400 data
AT4G15400 model
AT1G74030 data
AT1G74030 model
AT2G33780 data
AT2G33780 model
AT2G31335 data
AT2G31335 model

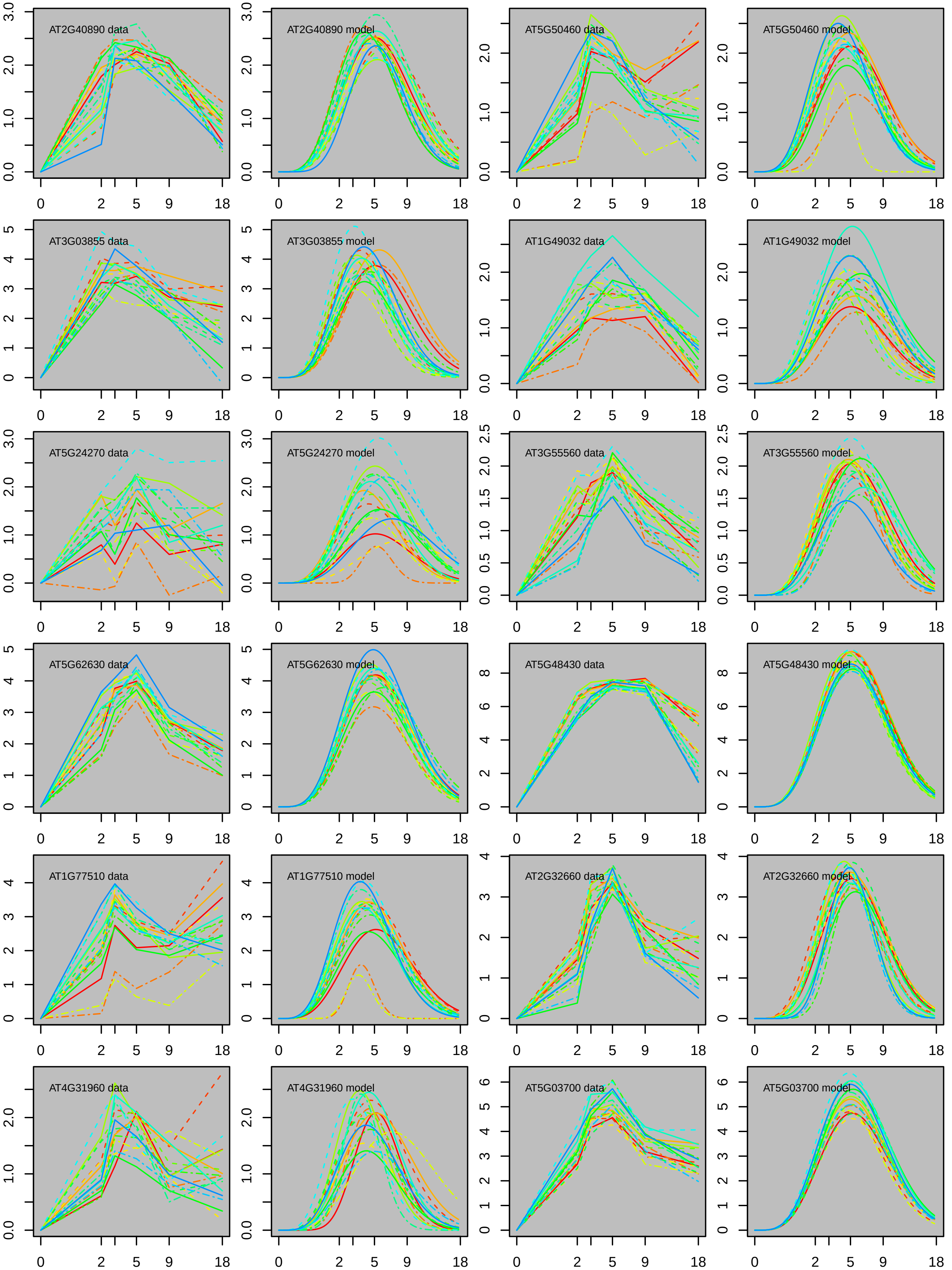
AT2G40890 data
AT2G40890 model
AT5G50460 data
AT5G50460 model
AT3G03855 data
AT3G03855 model
AT1G49032 data
AT1G49032 model
AT5G24270 data
AT5G24270 model
AT3G55560 data
AT3G55560 model
AT5G62630 data
AT5G62630 model
AT5G48430 data
AT5G48430 model
AT1G77510 data
AT1G77510 model
AT2G32660 data
AT2G32660 model
AT4G31960 data
AT4G31960 model
AT5G03700 data
AT5G03700 model
0
2
5
9
18

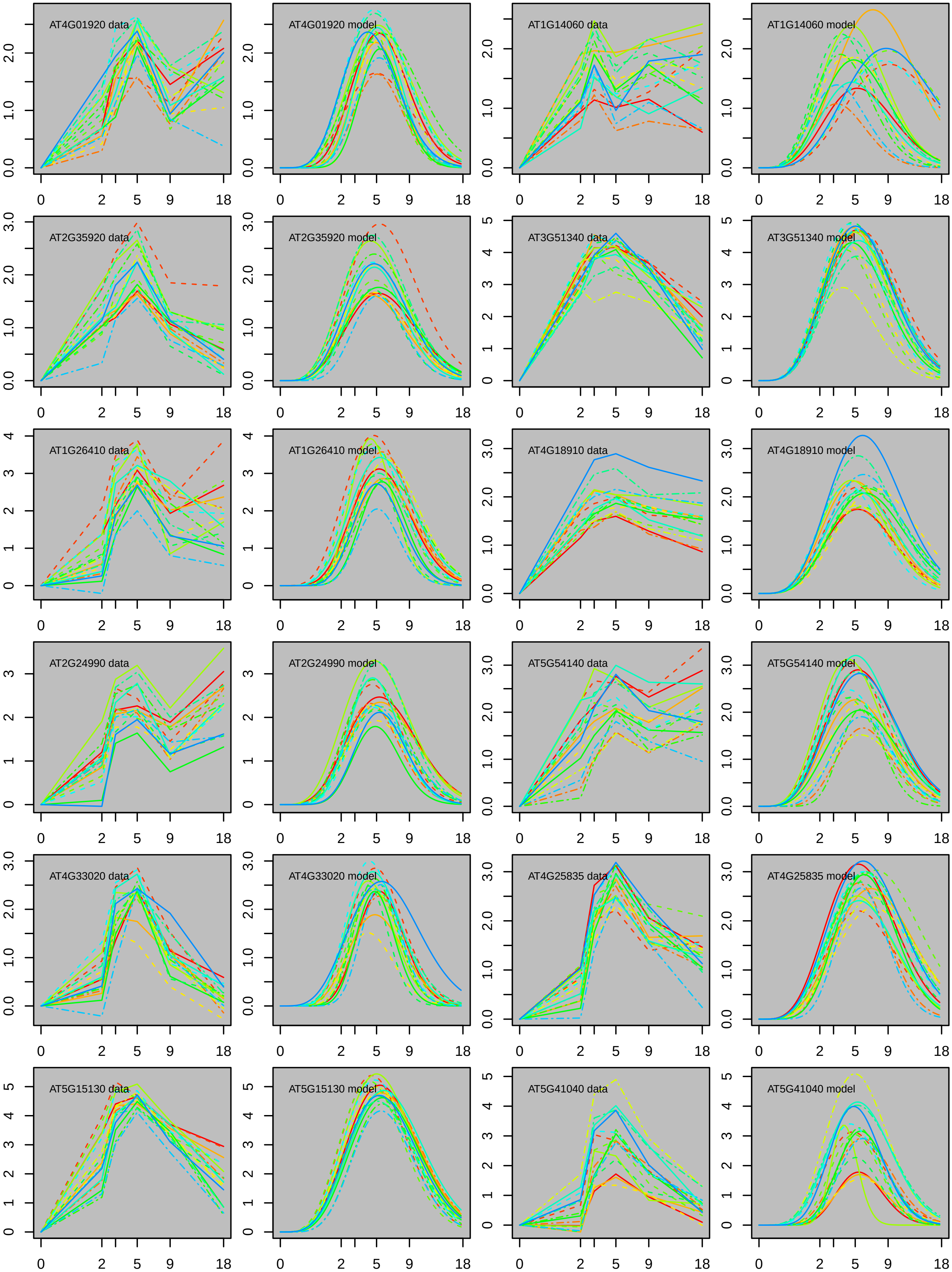

AT4G01920 data
AT4G01920 model
AT1G14060 data
AT1G14060 model
AT2G35920 data
AT2G35920 model
AT3G51340 data
AT3G51340 model
AT1G26410 data
AT1G26410 model
AT4G18910 data
AT4G18910 model
AT2G24990 data
AT2G24990 model
AT5G54140 data
AT5G54140 model
AT4G33020 data
AT4G33020 model
AT4G25835 data
AT4G25835 model
AT5G15130 data
AT5G15130 model
AT5G41040 data
AT5G41040 model
0 2 5 9 18

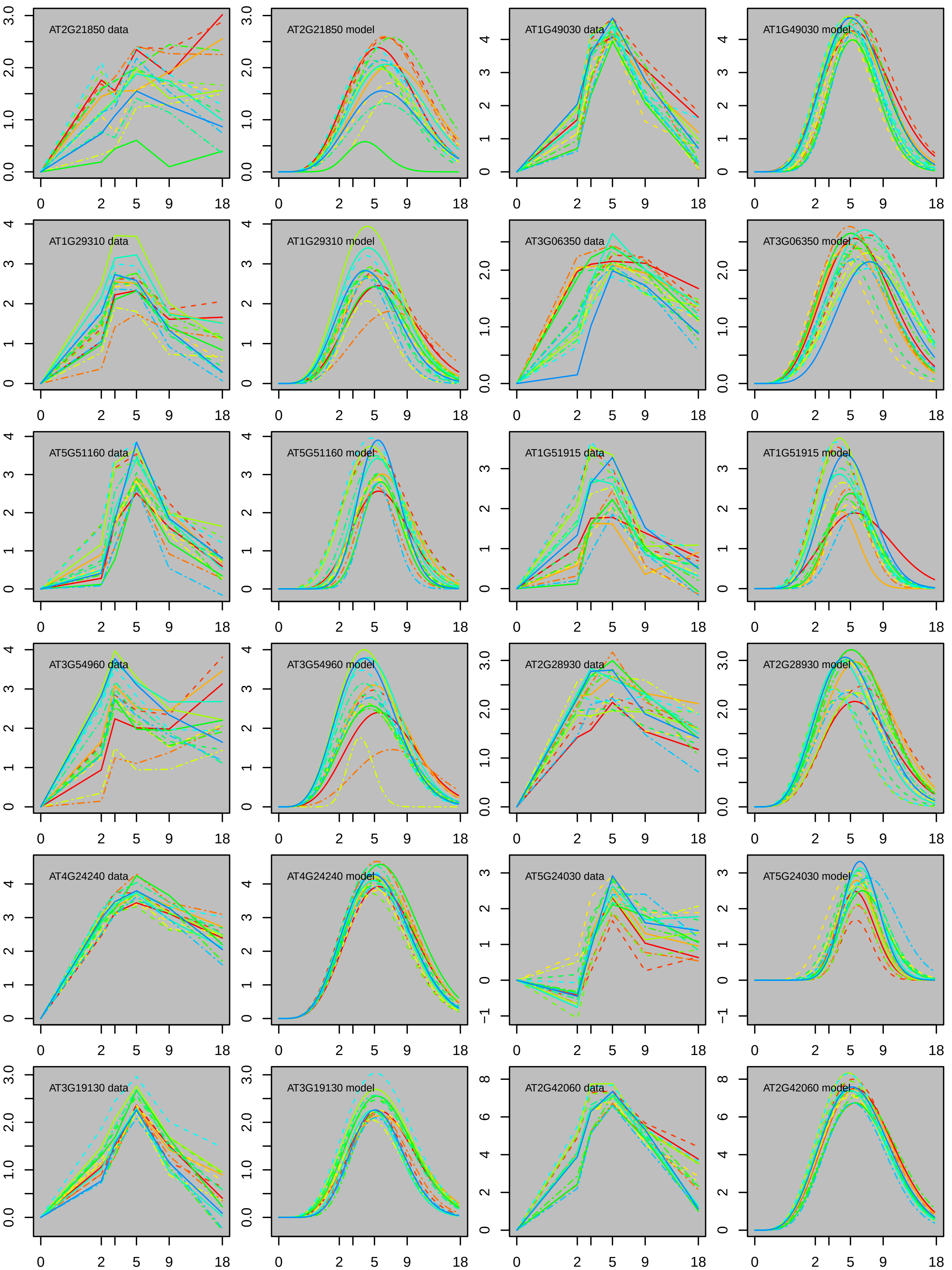

AT2G21850 data
AT2G21850 model
AT1G49030 data
AT1G49030 model
AT1G29310 data
AT1G29310 model
AT3G06350 data
AT3G06350 model
AT5G51160 data
AT5G51160 model
AT1G51915 data
AT1G51915 model
AT3G54960 data
AT3G54960 model
AT2G28930 data
AT2G28930 model
AT4G24240 data
AT4G24240 model
AT5G24030 data
AT5G24030 model
AT3G19130 data
AT3G19130 model
AT2G42060 data
AT2G42060 model
0
2
5
9
18

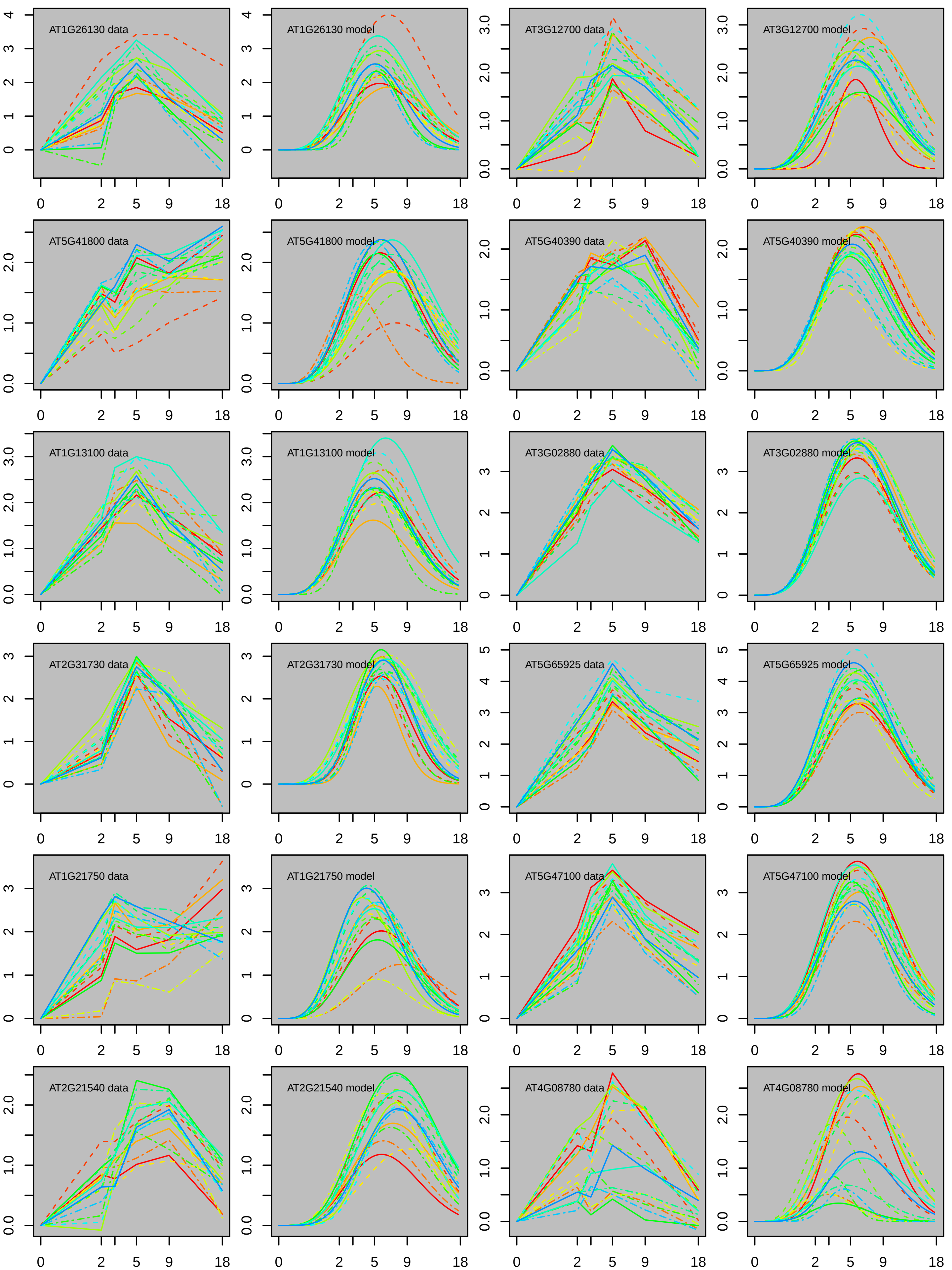
AT1G26130 data
AT1G26130 model
AT3G12700 data
AT3G12700 model
AT5G41800 data
AT5G41800 model
AT5G40390 data
AT5G40390 model
AT1G13100 data
AT1G13100 model
AT3G02880 data
AT3G02880 model
AT2G31730 data
AT2G31730 model
AT5G65925 data
AT5G65925 model
AT1G21750 data
AT1G21750 model
AT5G47100 data
AT5G47100 model
AT2G21540 data
AT2G21540 model
AT4G08780 data
AT4G08780 model
0 2 5 9 18

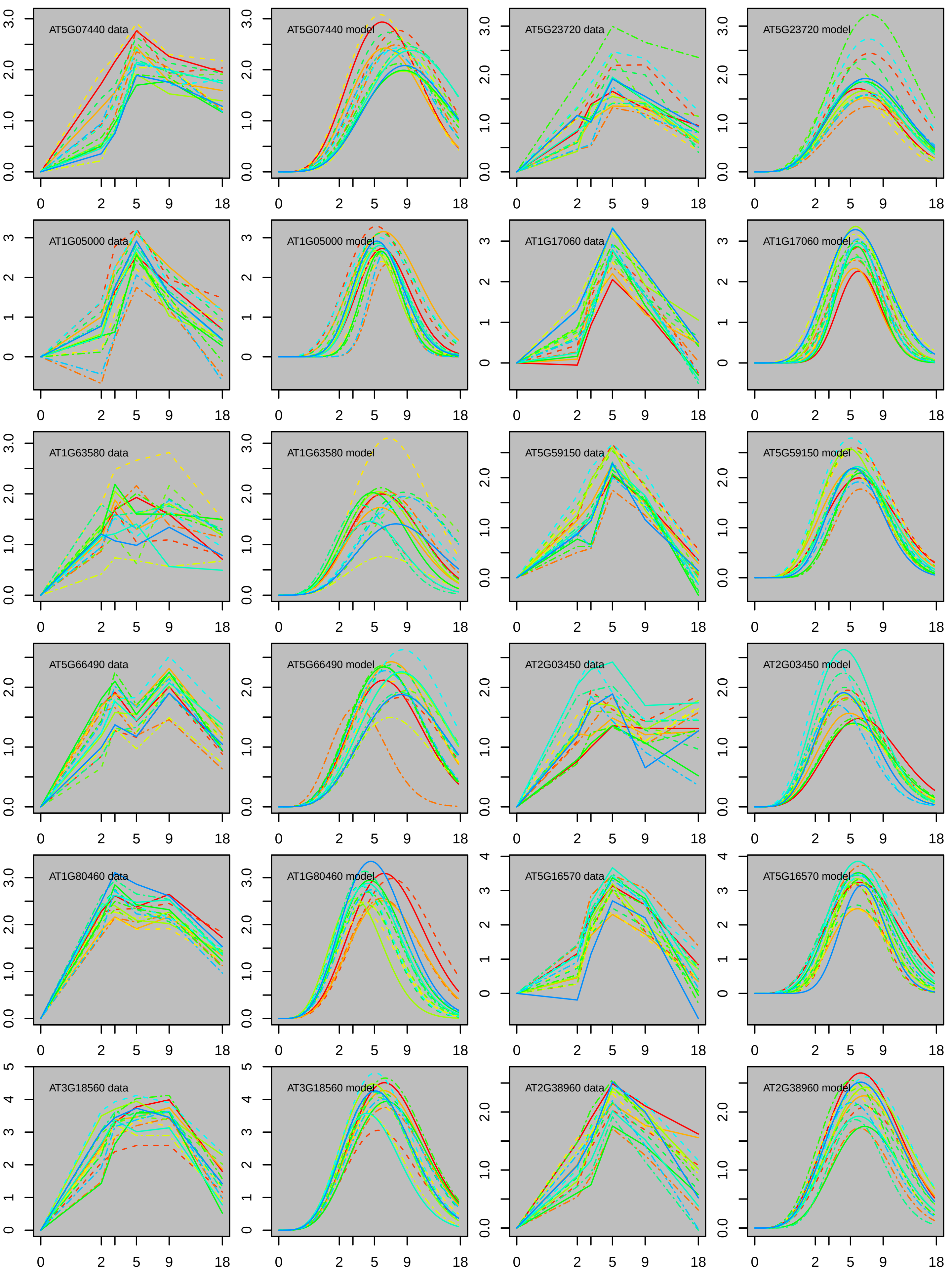

AT5G07440 data
AT5G07440 model
AT5G23720 data
AT5G23720 model
AT1G05000 data
AT1G05000 model
AT1G17060 data
AT1G17060 model
AT1G63580 data
AT1G63580 model
AT5G59150 data
AT5G59150 model
AT5G66490 data
AT5G66490 model
AT2G03450 data
AT2G03450 model
AT1G80460 data
AT1G80460 model
AT5G16570 data
AT5G16570 model
AT3G18560 data
AT3G18560 model
AT2G38960 data
AT2G38960 model
0 2 5 9 18

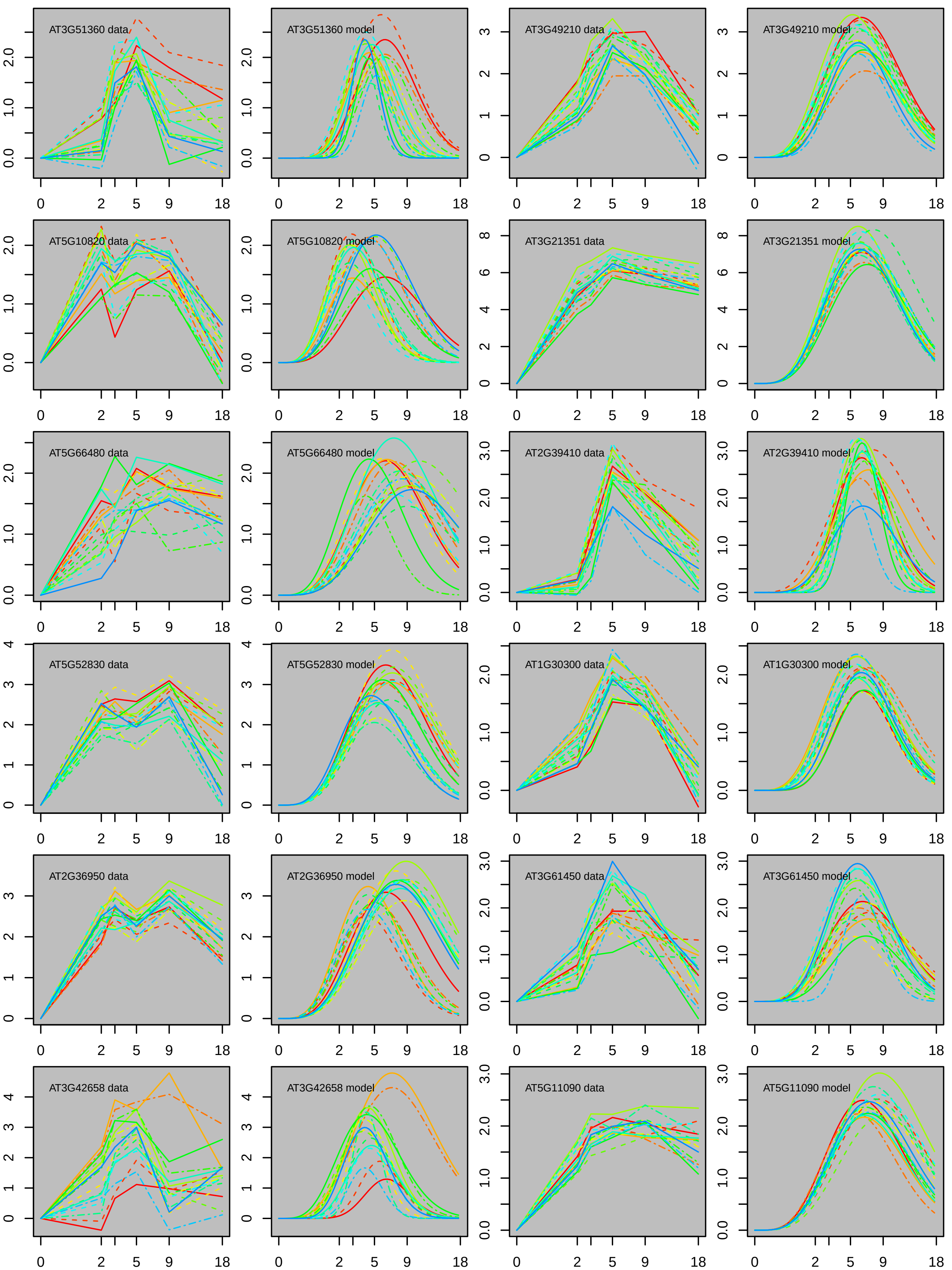

AT3G51360 data
AT3G51360 model
AT3G49210 data
AT3G49210 model
AT5G10820 data
AT5G10820 model
AT3G21351 data
AT3G21351 model
AT5G66480 data
AT5G66480 model
AT2G39410 data
AT2G39410 model
AT5G52830 data
AT5G52830 model
AT1G30300 data
AT1G30300 model
AT2G36950 data
AT2G36950 model
AT3G61450 data
AT3G61450 model
AT3G42658 data
AT3G42658 model
AT5G11090 data
AT5G11090 model

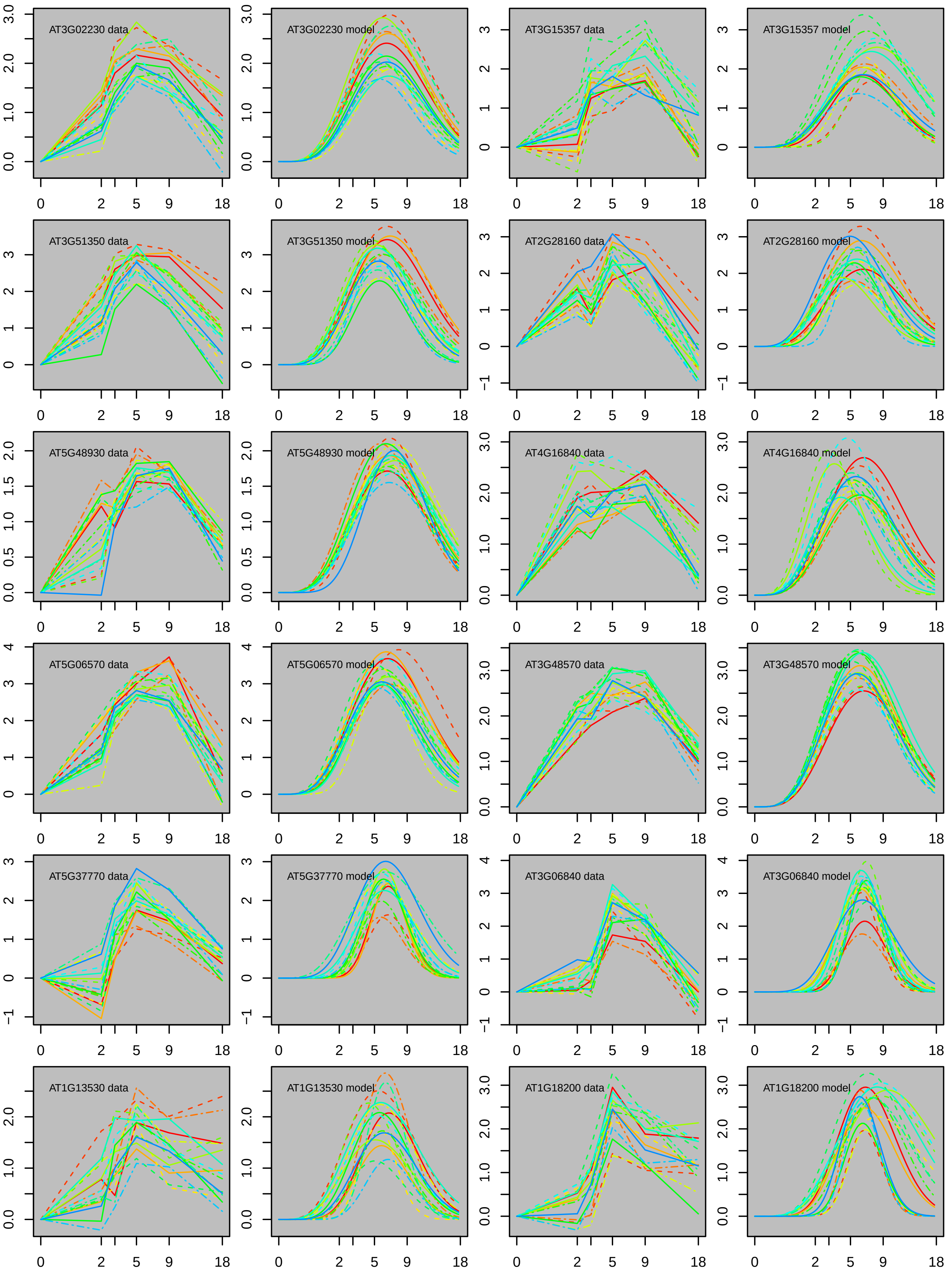
AT3G02230 data
AT3G02230 model
AT3G15357 data
AT3G15357 model
AT3G51350 data
AT3G51350 model
AT2G28160 data
AT2G28160 model
AT5G48930 data
AT5G48930 model
AT4G16840 data
AT4G16840 model
AT5G06570 data
AT5G06570 model
AT3G48570 data
AT3G48570 model
AT5G37770 data
AT5G37770 model
AT3G06840 data
AT3G06840 model
AT1G13530 data
AT1G13530 model
AT1G18200 data
AT1G18200 model

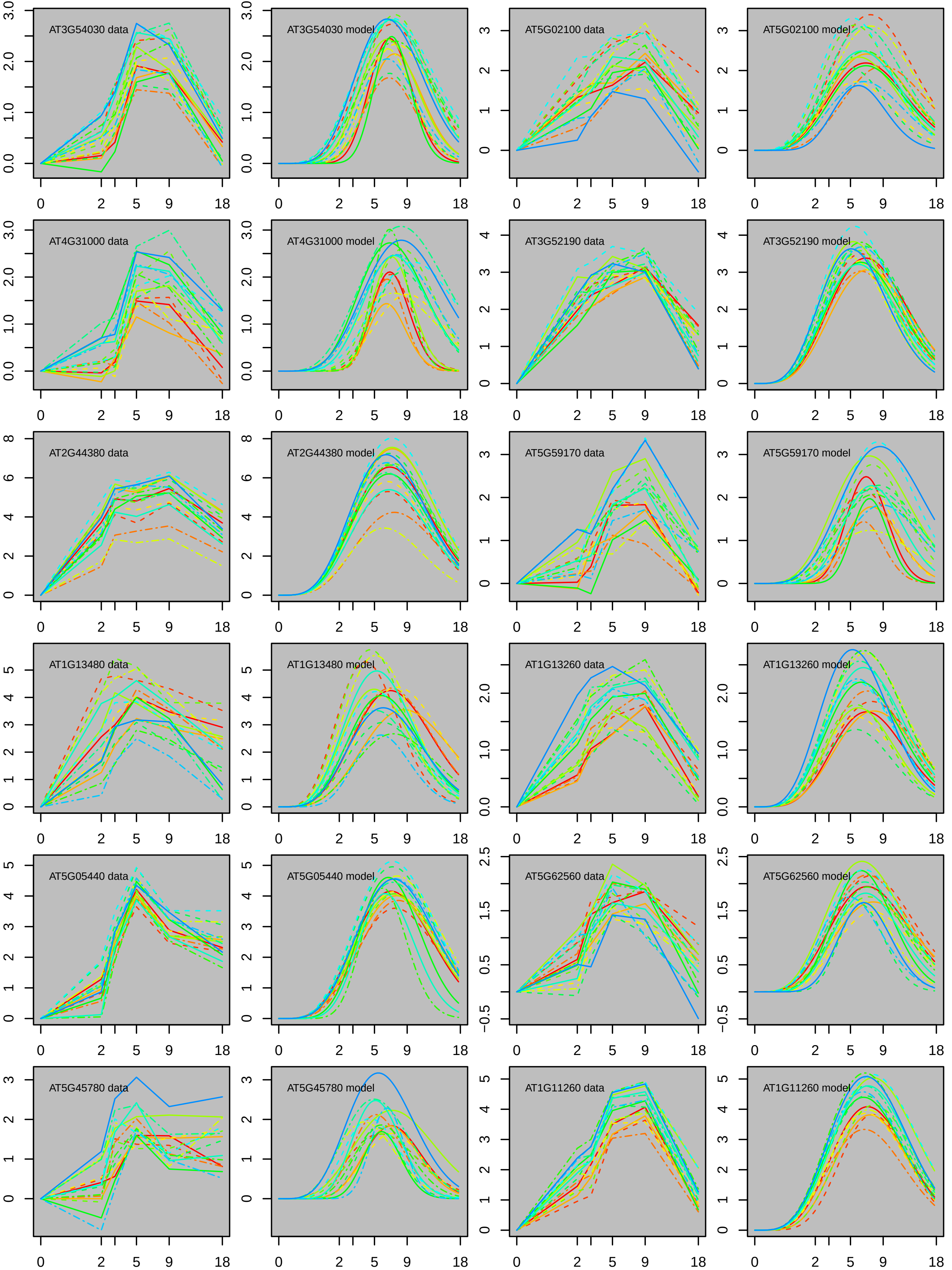
AT3G54030 data
AT3G54030 model
AT5G02100 data
AT5G02100 model
AT4G31000 data
AT4G31000 model
AT3G52190 data
AT3G52190 model
AT2G44380 data
AT2G44380 model
AT5G59170 data
AT5G59170 model
AT1G13480 data
AT1G13480 model
AT1G13260 data
AT1G13260 model
AT5G05440 data
AT5G05440 model
AT5G62560 data
AT5G62560 model
AT5G45780 data
AT5G45780 model
AT1G11260 data
AT1G11260 model
0 2 5 9 18

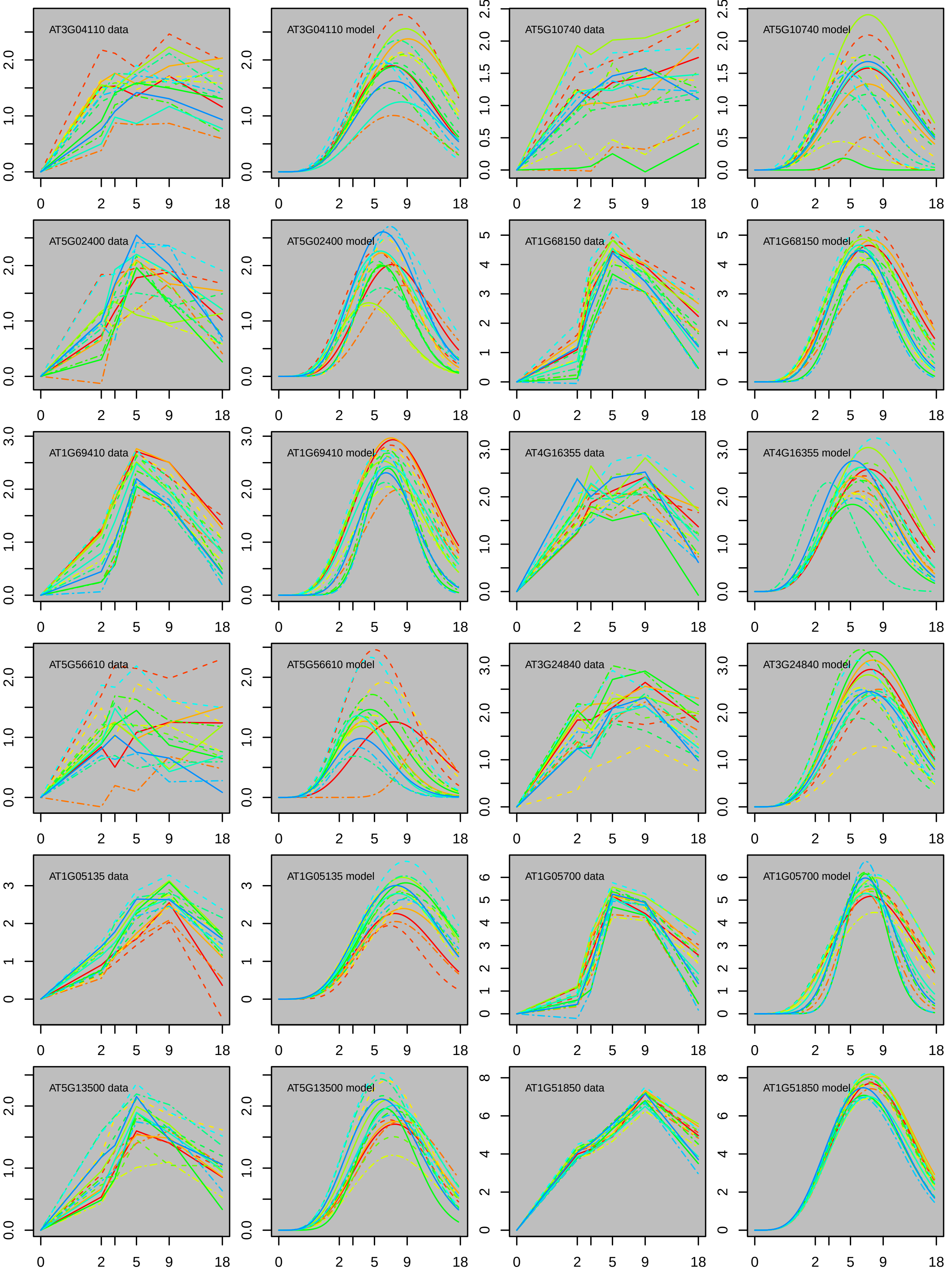

AT3G04110 data
AT3G04110 model
AT5G10740 data
AT5G10740 model
AT5G02400 data
AT5G02400 model
AT1G68150 data
AT1G68150 model
AT1G69410 data
AT1G69410 model
AT4G16355 data
AT4G16355 model
AT5G56610 data
AT5G56610 model
AT3G24840 data
AT3G24840 model
AT1G05135 data
AT1G05135 model
AT1G05700 data
AT1G05700 model
AT5G13500 data
AT5G13500 model
AT1G51850 data
AT1G51850 model

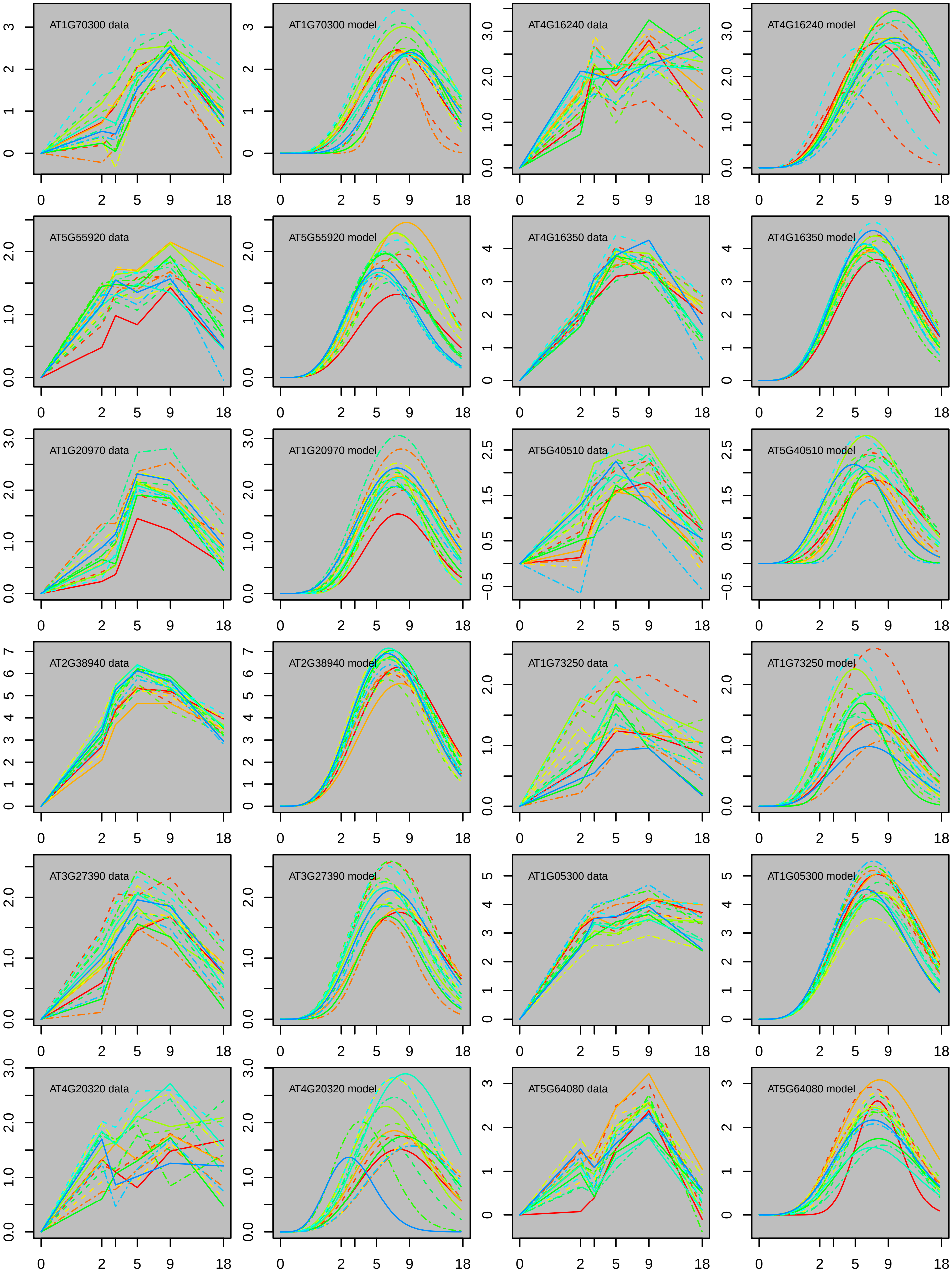
AT1G70300 data
AT1G70300 model
AT4G16240 data
AT4G16240 model
AT5G55920 data
AT5G55920 model
AT4G16350 data
AT4G16350 model
AT1G20970 data
AT1G20970 model
AT5G40510 data
AT5G40510 model
AT2G38940 data
AT2G38940 model
AT1G73250 data
AT1G73250 model
AT3G27390 data
AT3G27390 model
AT1G05300 data
AT1G05300 model
AT4G20320 data
AT4G20320 model
AT5G64080 data
AT5G64080 model
0 2 5 9 18

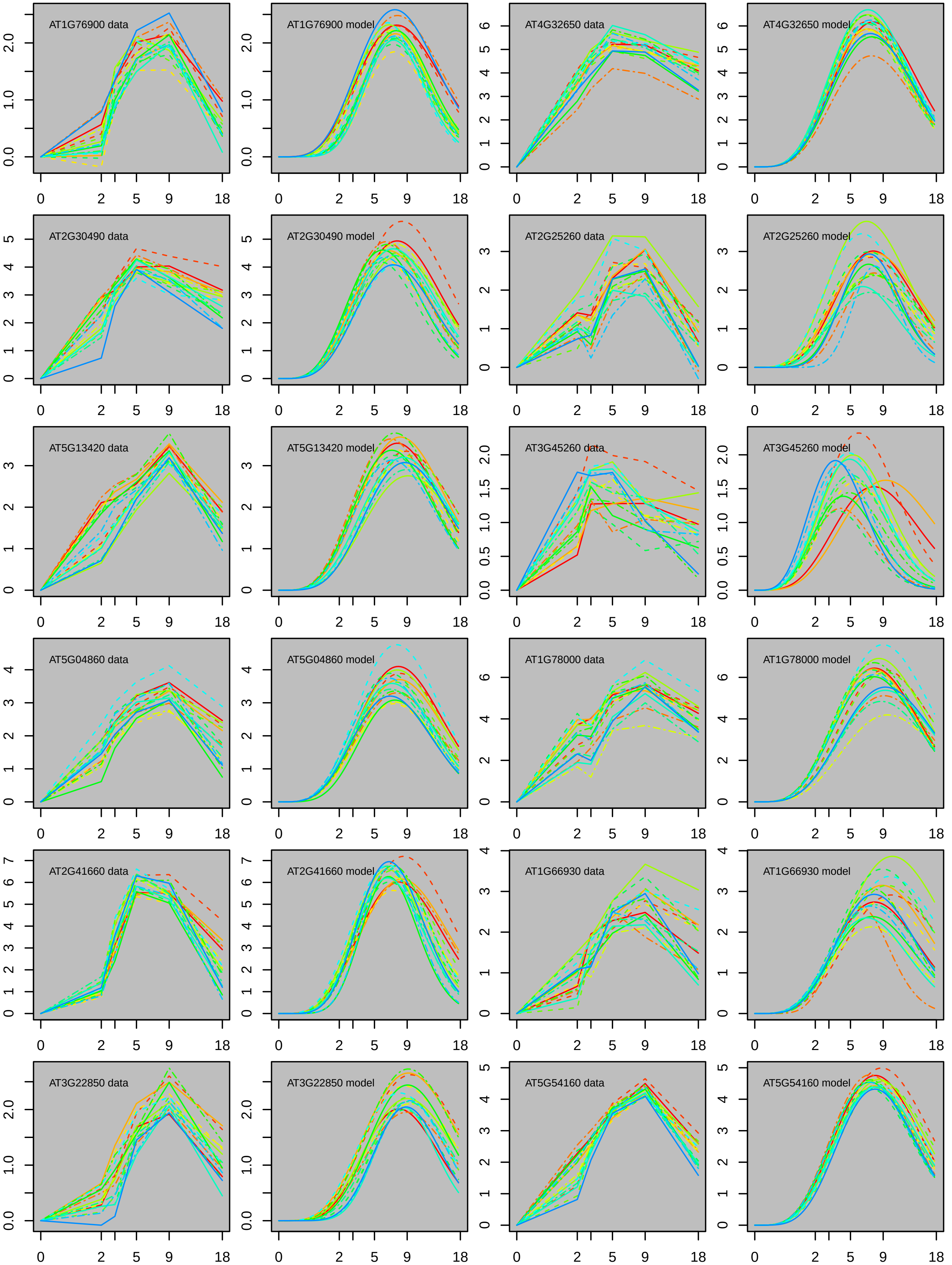
AT1G76900 data
AT1G76900 model
AT4G32650 data
AT4G32650 model
AT2G30490 data
AT2G30490 model
AT2G25260 data
AT2G25260 model
AT5G13420 data
AT5G13420 model
AT3G45260 data
AT3G45260 model
AT5G04860 data
AT5G04860 model
AT1G78000 data
AT1G78000 model
AT2G41660 data
AT2G41660 model
AT1G66930 data
AT1G66930 model
AT3G22850 data
AT3G22850 model
AT5G54160 data
AT5G54160 model

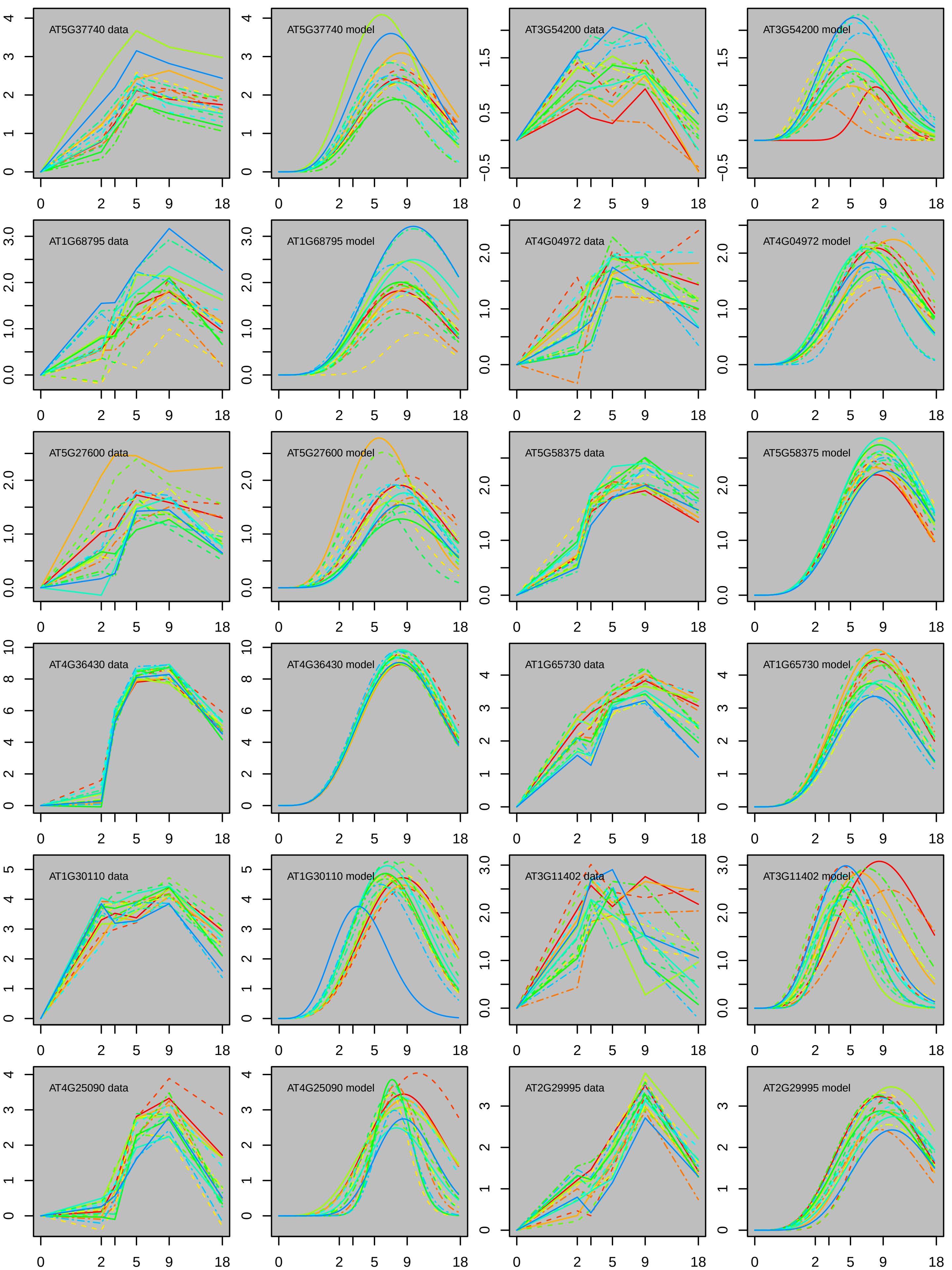
AT5G37740 data
AT5G37740 model
AT3G54200 data
AT3G54200 model
AT1G68795 data
AT1G68795 model
AT4G04972 data
AT4G04972 model
AT5G27600 data
AT5G27600 model
AT5G58375 data
AT5G58375 model
AT4G36430 data
AT4G36430 model
AT1G65730 data
AT1G65730 model
AT1G30110 data
AT1G30110 model
AT3G11402 data
AT3G11402 model
AT4G25090 data
AT4G25090 model
AT2G29995 data
AT2G29995 model

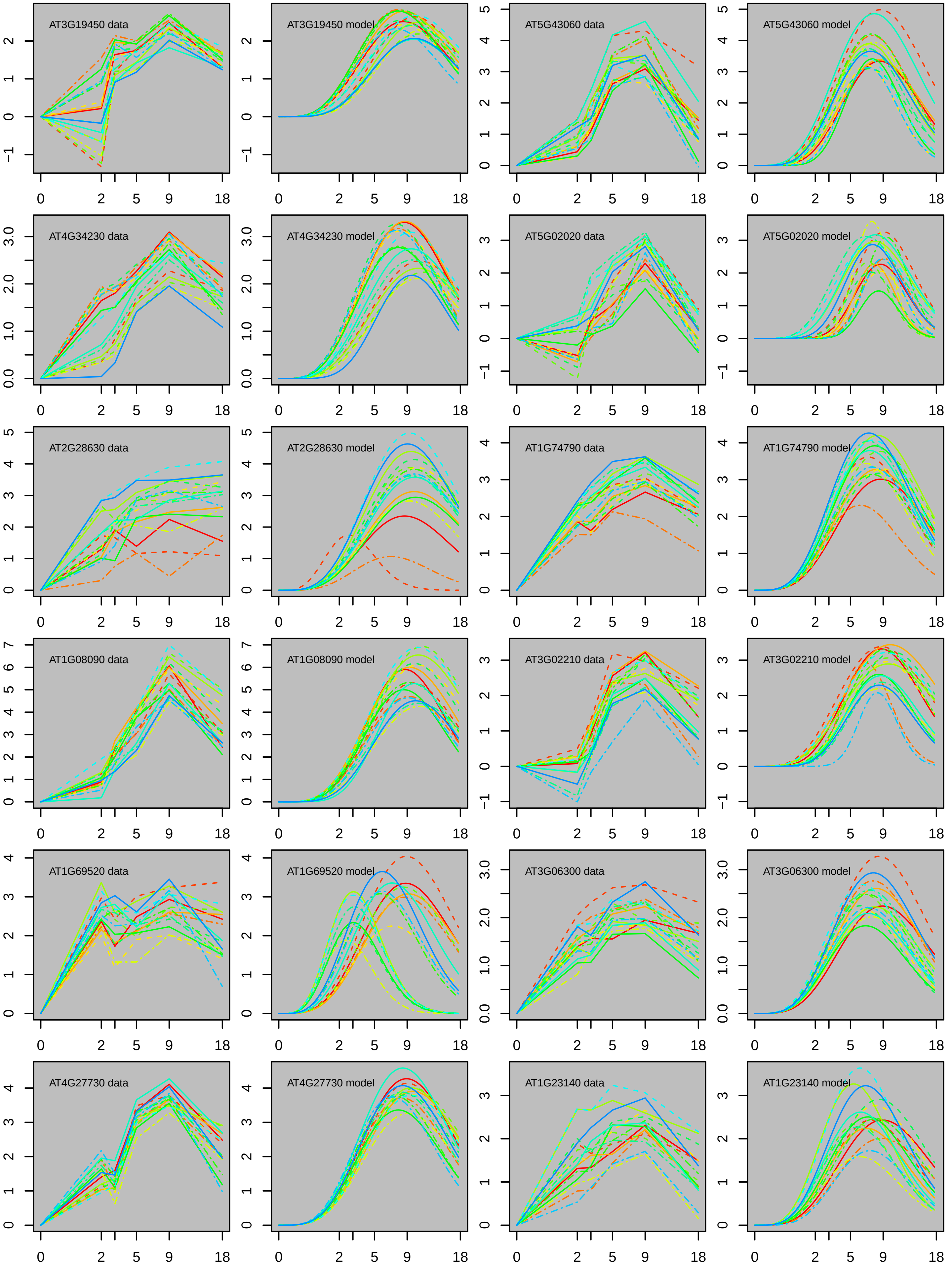
AT3G19450 data
AT3G19450 model
AT5G43060 data
AT5G43060 model
AT4G34230 data
AT4G34230 model
AT5G02020 data
AT5G02020 model
AT2G28630 data
AT2G28630 model
AT1G74790 data
AT1G74790 model
AT1G08090 data
AT1G08090 model
AT3G02210 data
AT3G02210 model
AT1G69520 data
AT1G69520 model
AT3G06300 data
AT3G06300 model
AT4G27730 data
AT4G27730 model
AT1G23140 data
AT1G23140 model
0 2 5 9 18

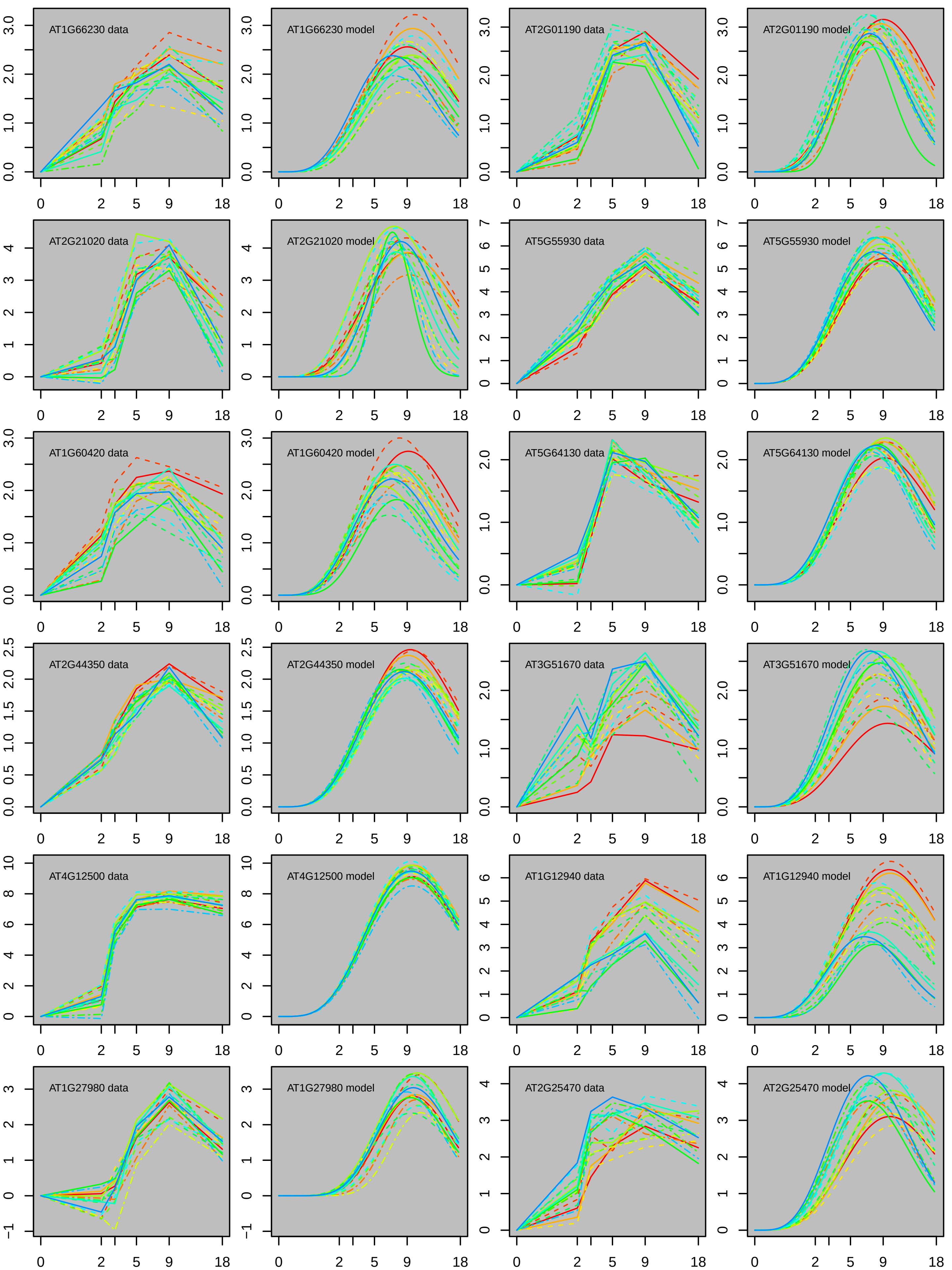

AT1G66230 data
AT1G66230 model
AT2G01190 data
AT2G01190 model
AT2G21020 data
AT2G21020 model
AT5G55930 data
AT5G55930 model
AT1G60420 data
AT1G60420 model
AT5G64130 data
AT5G64130 model
AT2G44350 data
AT2G44350 model
AT3G51670 data
AT3G51670 model
AT4G12500 data
AT4G12500 model
AT1G12940 data
AT1G12940 model
AT1G27980 data
AT1G27980 model
AT2G25470 data
AT2G25470 model
0 2 5 9 18

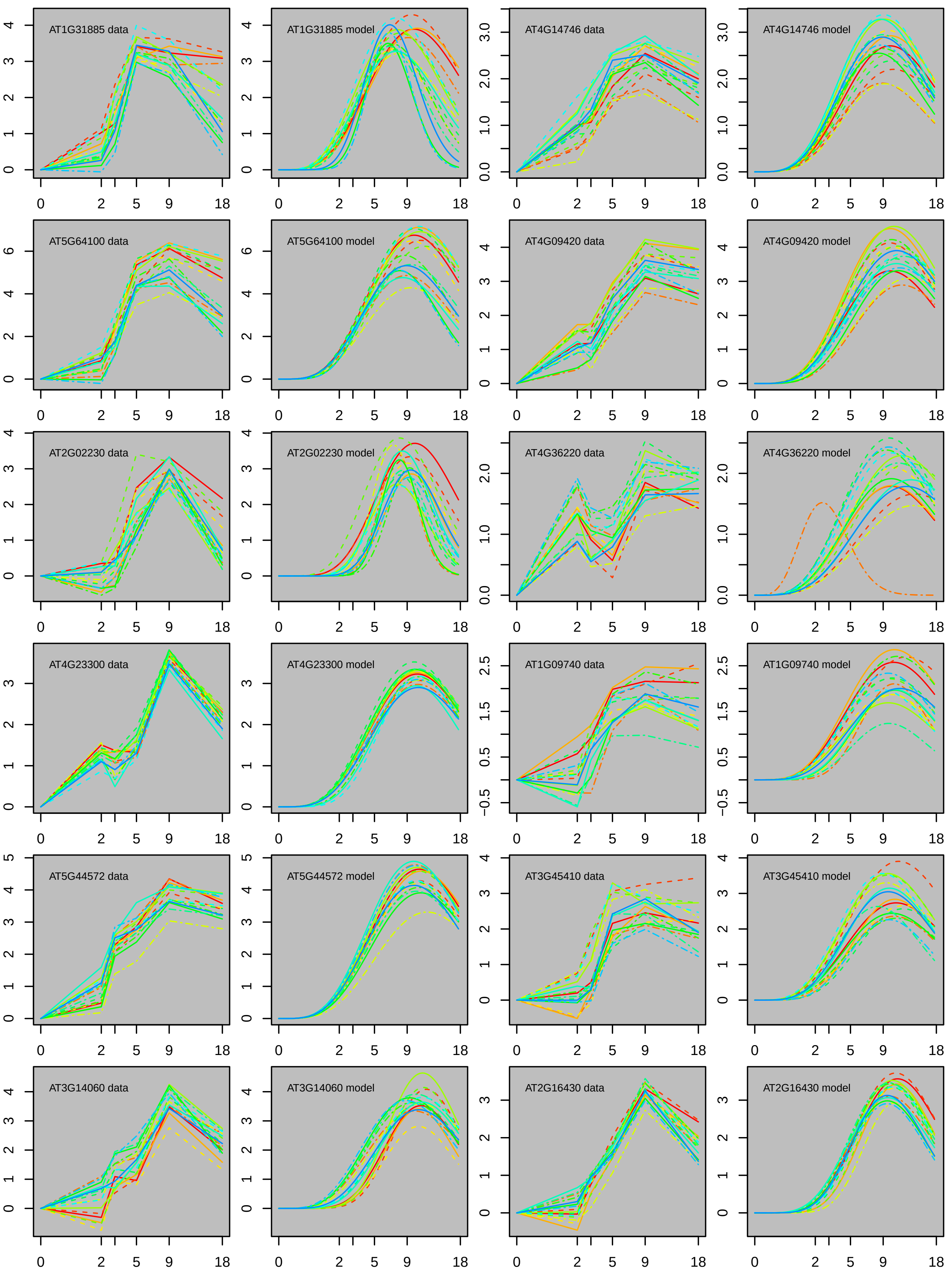

AT1G31885 data
AT1G31885 model
AT4G14746 data
AT4G14746 model
AT5G64100 data
AT5G64100 model
AT4G09420 data
AT4G09420 model
AT2G02230 data
AT2G02230 model
AT4G36220 data
AT4G36220 model
AT4G23300 data
AT4G23300 model
AT1G09740 data
AT1G09740 model
AT5G44572 data
AT5G44572 model
AT3G45410 data
AT3G45410 model
AT3G14060 data
AT3G14060 model
AT2G16430 data
AT2G16430 model
0
2
5
9
18

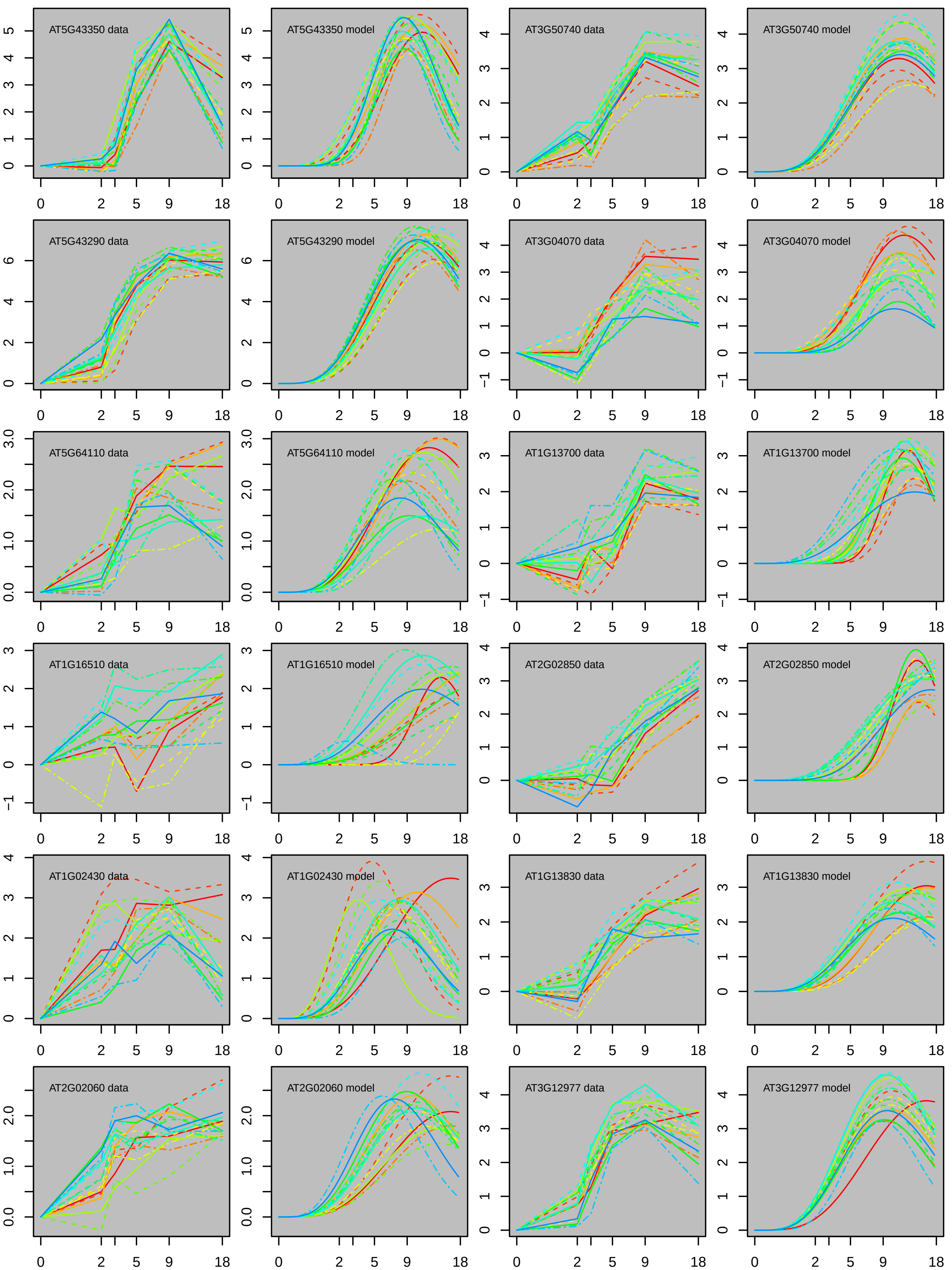

AT5G43350 data
AT5G43350 model
AT3G50740 data
AT3G50740 model
AT5G43290 data
AT5G43290 model
AT3G04070 data
AT3G04070 model
AT5G64110 data
AT5G64110 model
AT1G13700 data
AT1G13700 model
AT1G16510 data
AT1G16510 model
AT2G02850 data
AT2G02850 model
AT1G02430 data
AT1G02430 model
AT1G13830 data
AT1G13830 model
AT2G02060 data
AT2G02060 model
AT3G12977 data
AT3G12977 model

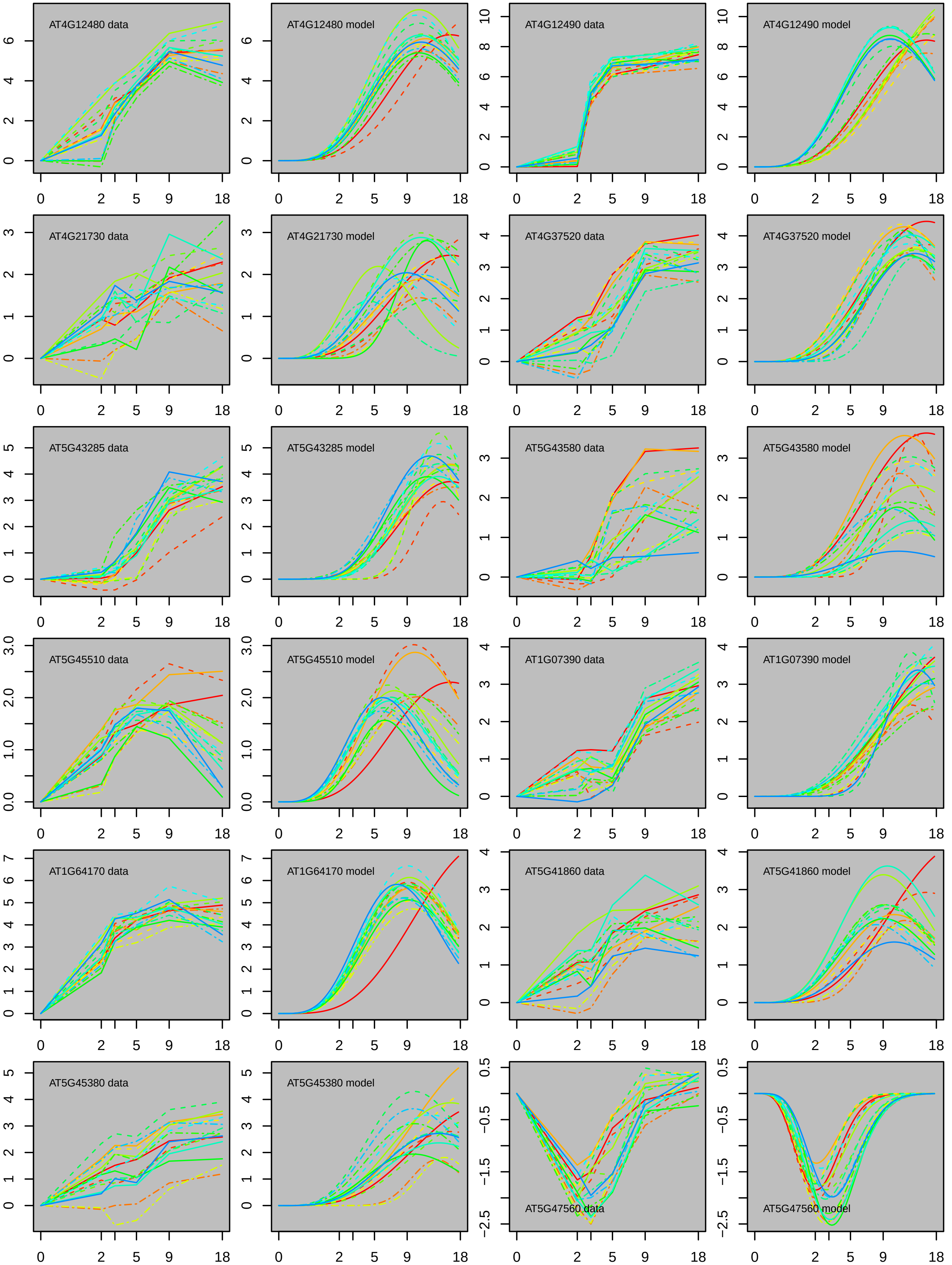
AT4G12480 data
AT4G12480 model
AT4G12490 data
AT4G12490 model
AT4G21730 data
AT4G21730 model
AT4G37520 data
AT4G37520 model
AT5G43285 data
AT5G43285 model
AT5G43580 data
AT5G43580 model
AT5G45510 data
AT5G45510 model
AT1G07390 data
AT1G07390 model
AT1G64170 data
AT1G64170 model
AT5G41860 data
AT5G41860 model
AT5G45380 data
AT5G45380 model
AT5G47560 data
AT5G47560 model
0 2 5 9 18

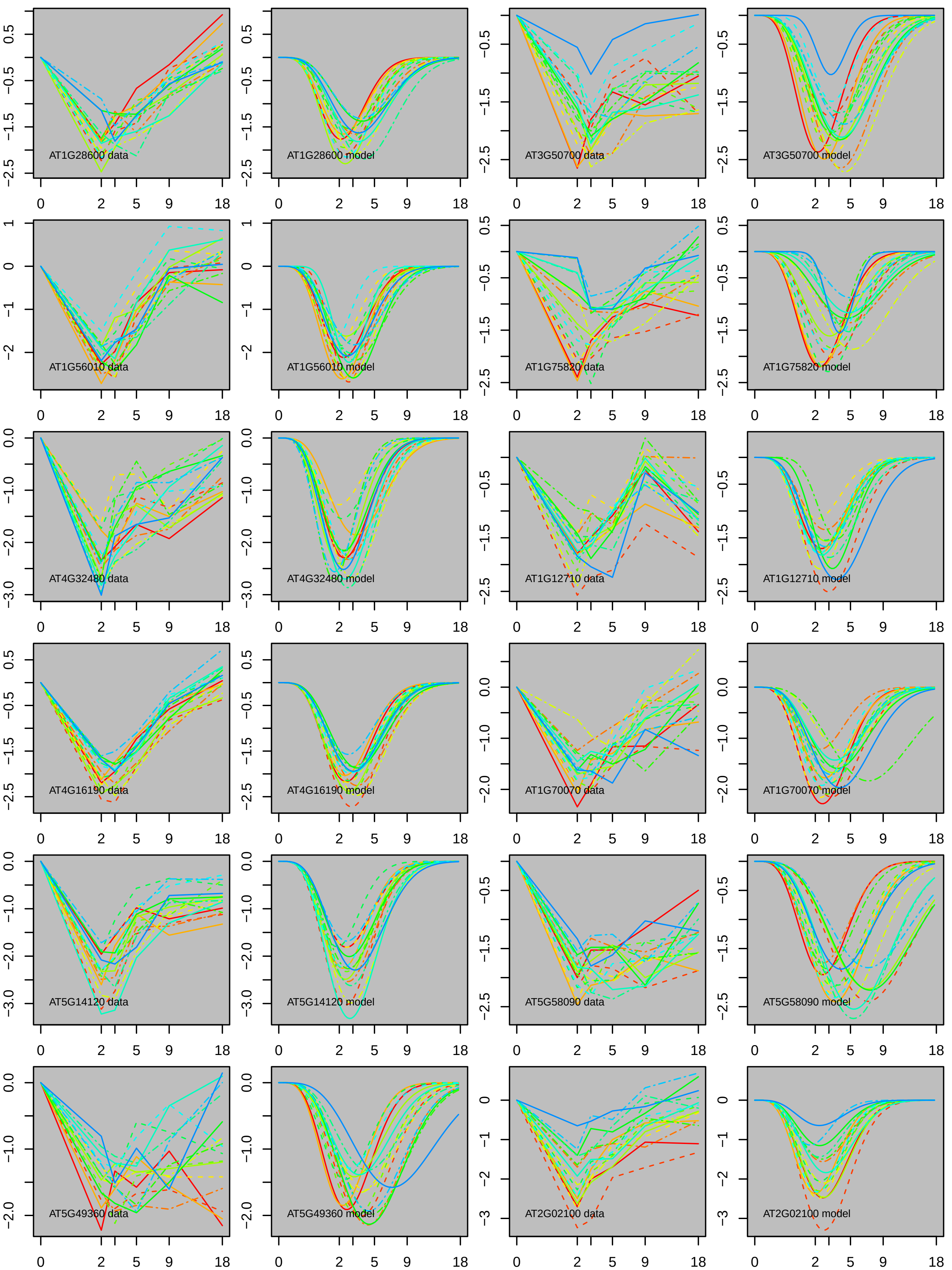

AT1G28600 data
AT1G28600 model
AT3G50700 data
AT3G50700 model
AT1G56010 data
AT1G56010 model
AT1G75820 data
AT1G75820 model
AT4G32480 data
AT4G32480 model
AT1G12710 data
AT1G12710 model
AT4G16190 data
AT4G16190 model
AT1G70070 data
AT1G70070 model
AT5G14120 data
AT5G14120 model
AT5G58090 data
AT5G58090 model
AT5G49360 data
AT5G49360 model
AT2G02100 data
AT2G02100 model

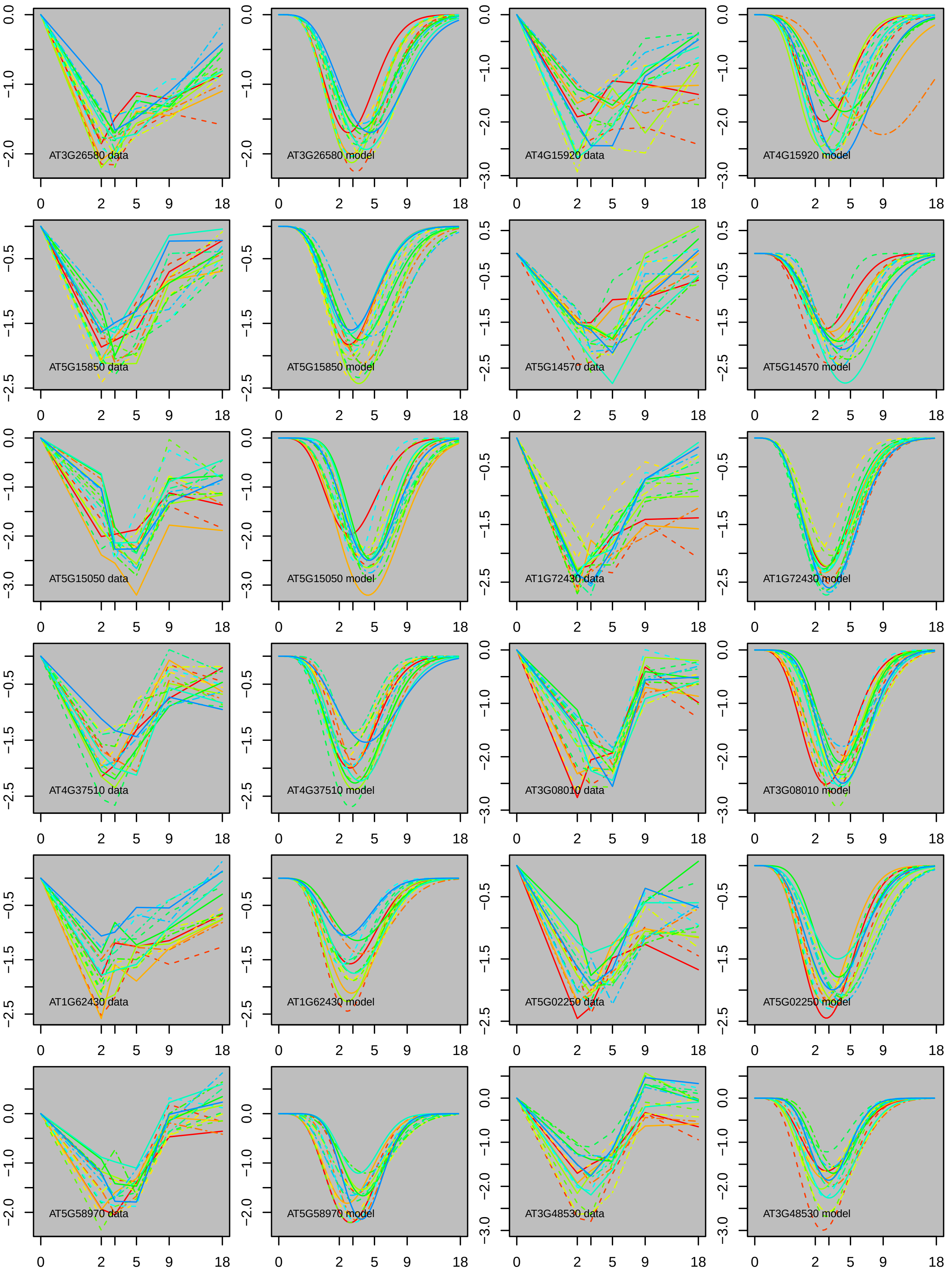

AT3G26580 data
AT3G26580 model
AT4G15920 data
AT4G15920 model
AT5G15850 data
AT5G15850 model
AT5G14570 data
AT5G14570 model
AT5G15050 data
AT5G15050 model
AT1G72430 data
AT1G72430 model
AT4G37510 data
AT4G37510 model
AT3G08010 data
AT3G08010 model
AT1G62430 data
AT1G62430 model
AT5G02250 data
AT5G02250 model
AT5G58970 data
AT5G58970 model
AT3G48530 data
AT3G48530 model

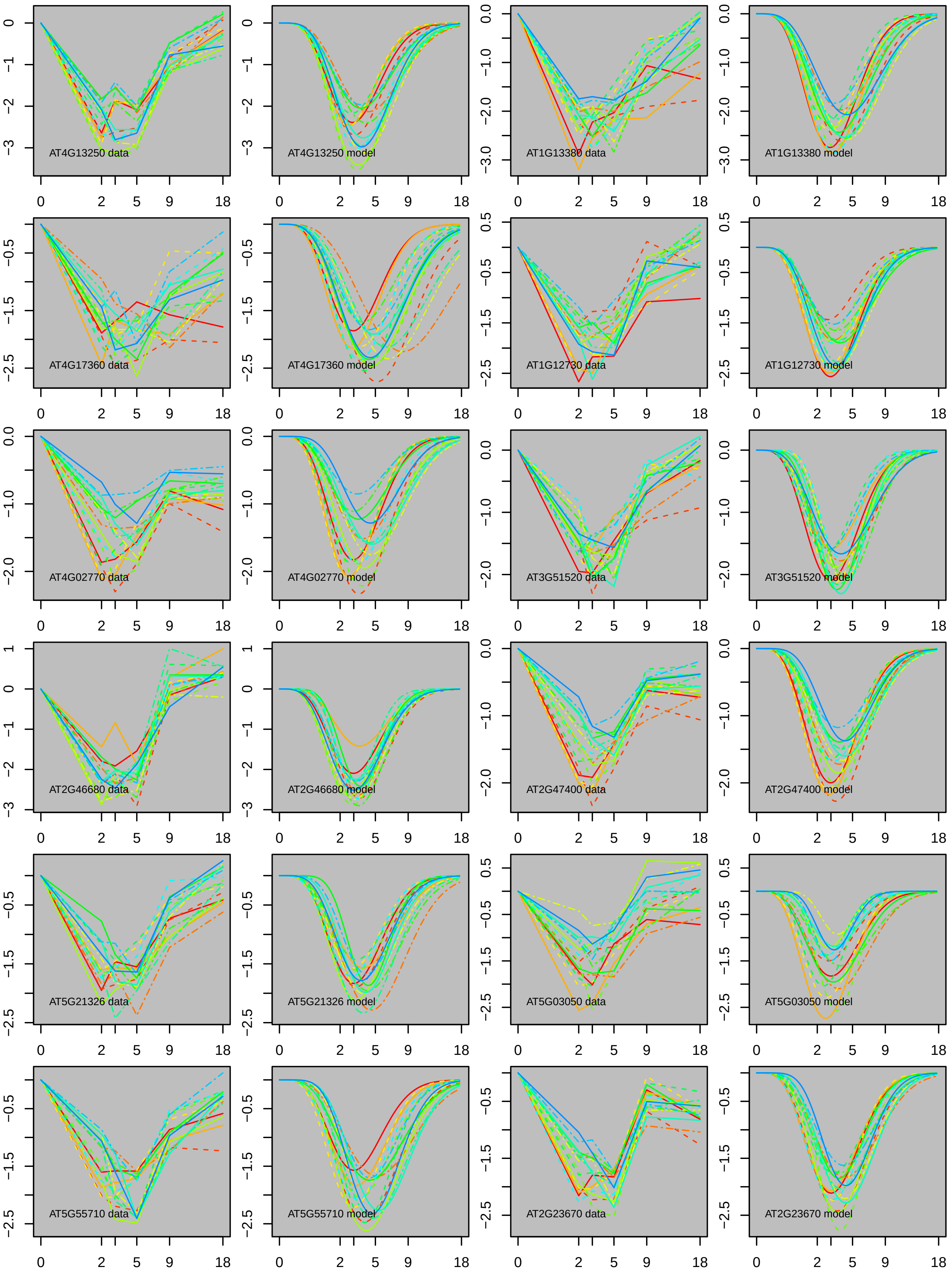

AT4G13250 data
AT4G13250 model
AT1G13380 data
AT1G13380 model
AT4G17360 data
AT4G17360 model
AT1G12730 data
AT1G12730 model
AT4G02770 data
AT4G02770 model
AT3G51520 data
AT3G51520 model
AT2G46680 data
AT2G46680 model
AT2G47400 data
AT2G47400 model
AT5G21326 data
AT5G21326 model
AT5G03050 data
AT5G03050 model
AT5G55710 data
AT5G55710 model
AT2G23670 data
AT2G23670 model

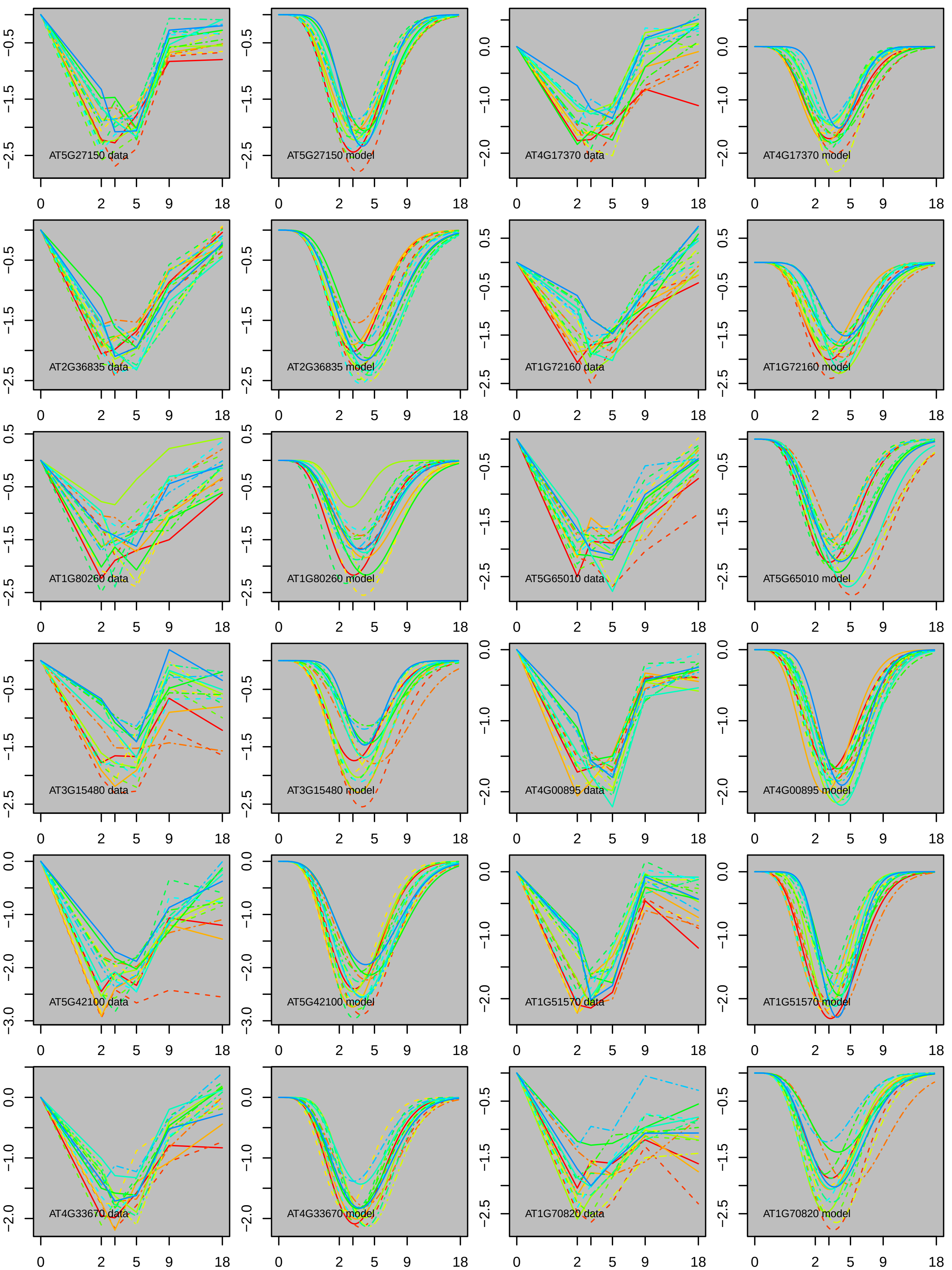
AT5G27150 data
AT5G27150 model
AT4G17370 data
AT4G17370 model
AT2G36835 data
AT2G36835 model
AT1G72160 data
AT1G72160 model
AT1G80260 data
AT1G80260 model
AT5G65010 data
AT5G65010 model
AT3G15480 data
AT3G15480 model
AT4G00895 data
AT4G00895 model
AT5G42100 data
AT5G42100 model
AT1G51570 data
AT1G51570 model
AT4G33670 data
AT4G33670 model
AT1G70820 data
AT1G70820 model
0
2
5
9
18

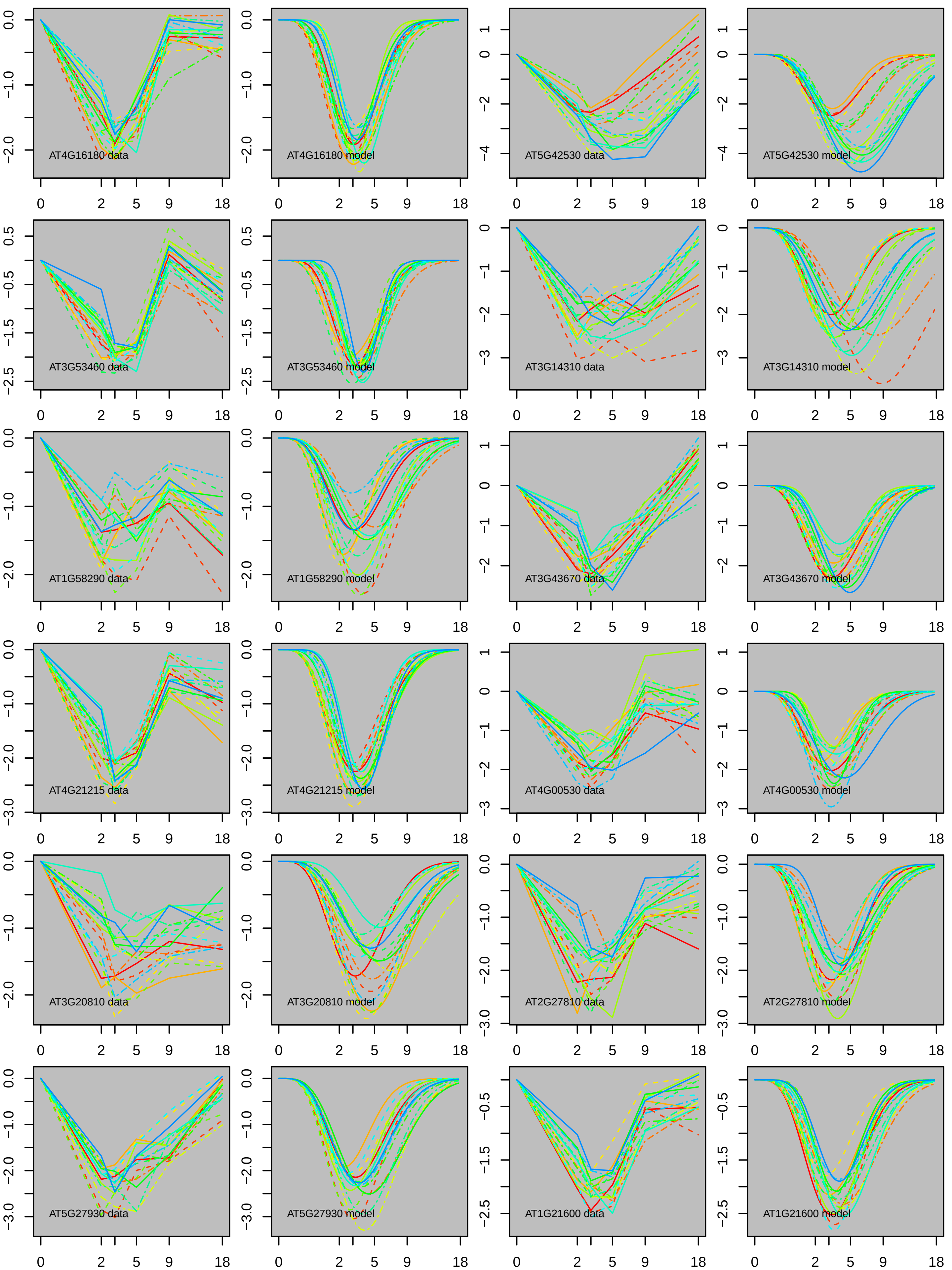

AT4G16180 data
AT4G16180 model
AT5G42530 data
AT5G42530 model
AT3G53460 data
AT3G53460 model
AT3G14310 data
AT3G14310 model
AT1G58290 data
AT1G58290 model
AT3G43670 data
AT3G43670 model
AT4G21215 data
AT4G21215 model
AT4G00530 data
AT4G00530 model
AT3G20810 data
AT3G20810 model
AT2G27810 data
AT2G27810 model
AT5G27930 data
AT5G27930 model
AT1G21600 data
AT1G21600 model

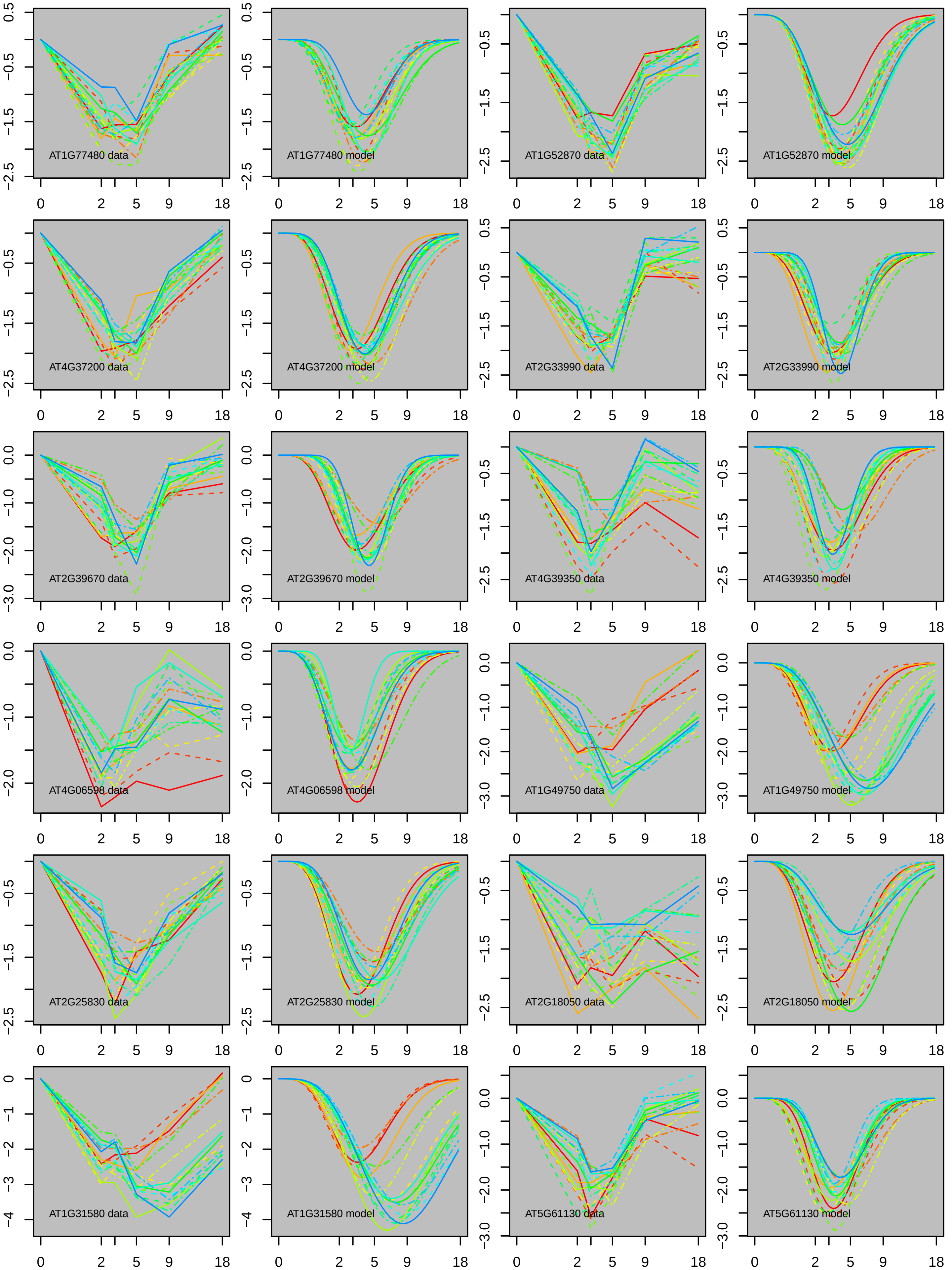

AT1G77480 data
AT1G77480 model
AT1G52870 data
AT1G52870 model
AT4G37200 data
AT4G37200 model
AT2G33990 data
AT2G33990 model
AT2G39670 data
AT2G39670 model
AT4G39350 data
AT4G39350 model
AT4G06598 data
AT4G06598 model
AT1G49750 data
AT1G49750 model
AT2G25830 data
AT2G25830 model
AT2G18050 data
AT2G18050 model
AT1G31580 data
AT1G31580 model
AT5G61130 data
AT5G61130 model

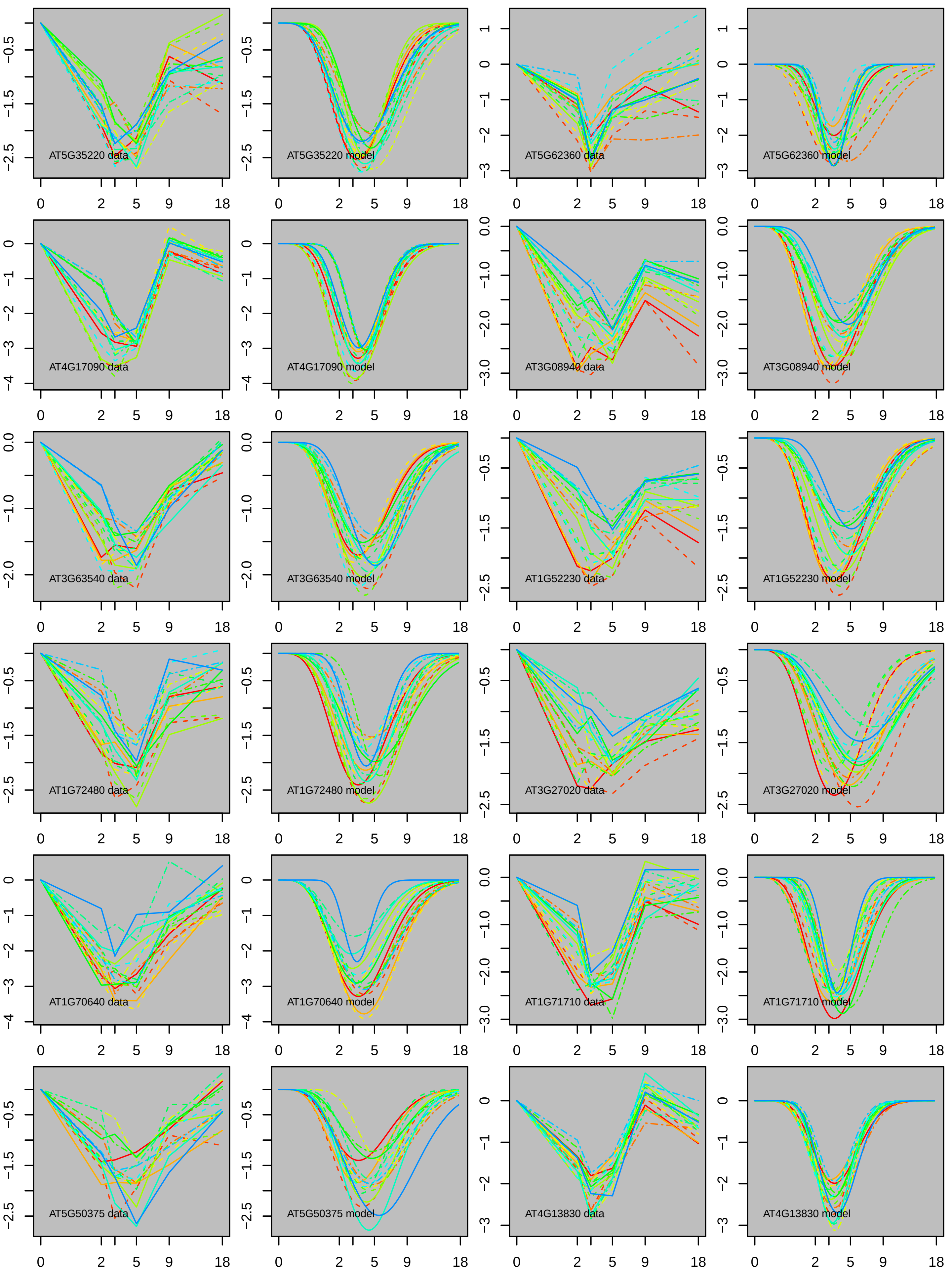

AT5G35220 data
AT5G35220 model
AT5G62360 data
AT5G62360 model
AT4G17090 data
AT4G17090 model
AT3G08940 data
AT3G08940 model
AT3G63540 data
AT3G63540 model
AT1G52230 data
AT1G52230 model
AT1G72480 data
AT1G72480 model
AT3G27020 data
AT3G27020 model
AT1G70640 data
AT1G70640 model
AT1G71710 data
AT1G71710 model
AT5G50375 data
AT5G50375 model
AT4G13830 data
AT4G13830 model

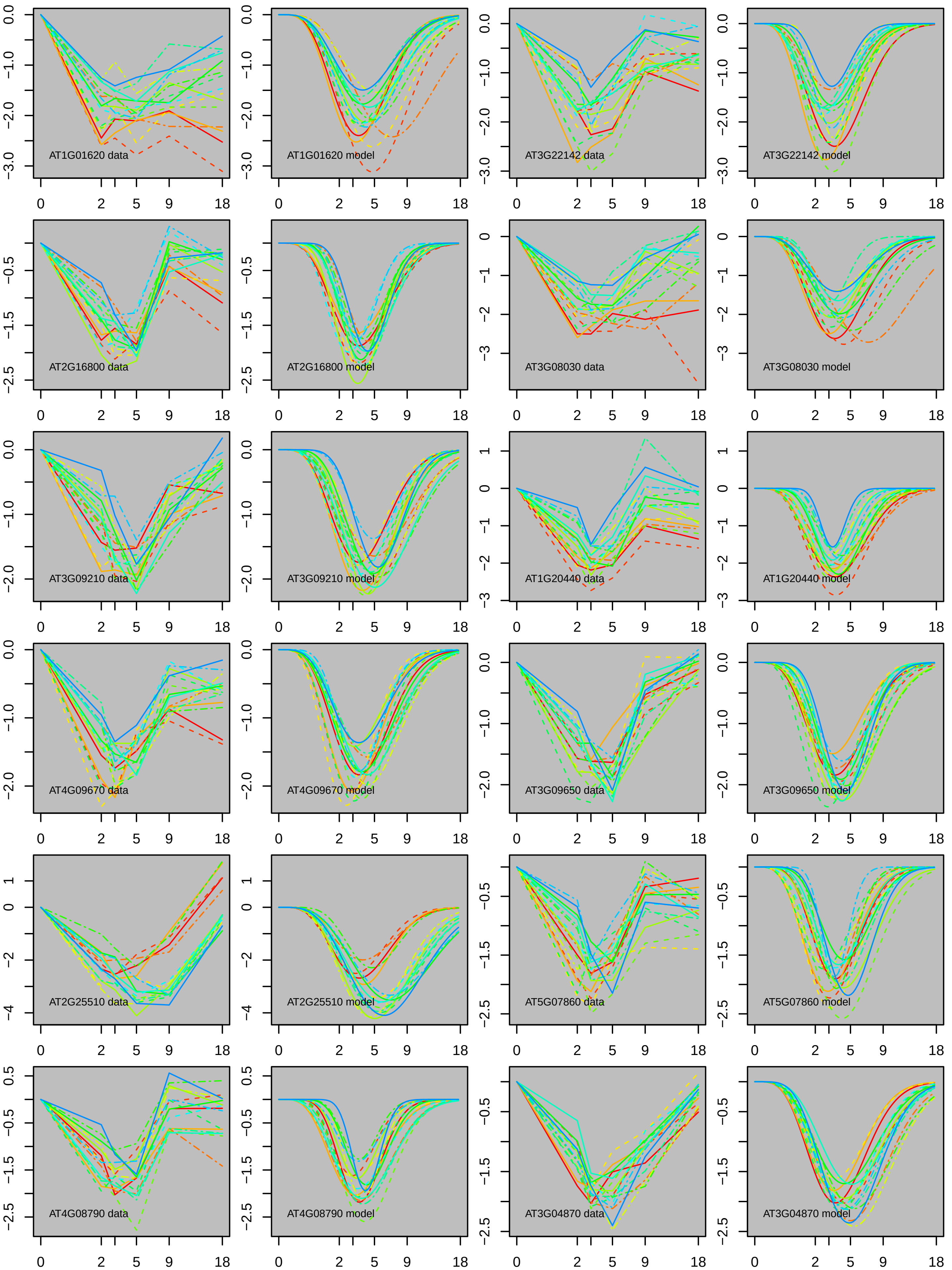
AT1G01620 data
AT1G01620 model
AT3G22142 data
AT3G22142 model
AT2G16800 data
AT2G16800 model
AT3G08030 data
AT3G08030 model
AT3G09210 data
AT3G09210 model
AT1G20440 data
AT1G20440 model
AT4G09670 data
AT4G09670 model
AT3G09650 data
AT3G09650 model
AT2G25510 data
AT2G25510 model
AT5G07860 data
AT5G07860 model
AT4G08790 data
AT4G08790 model
AT3G04870 data
AT3G04870 model

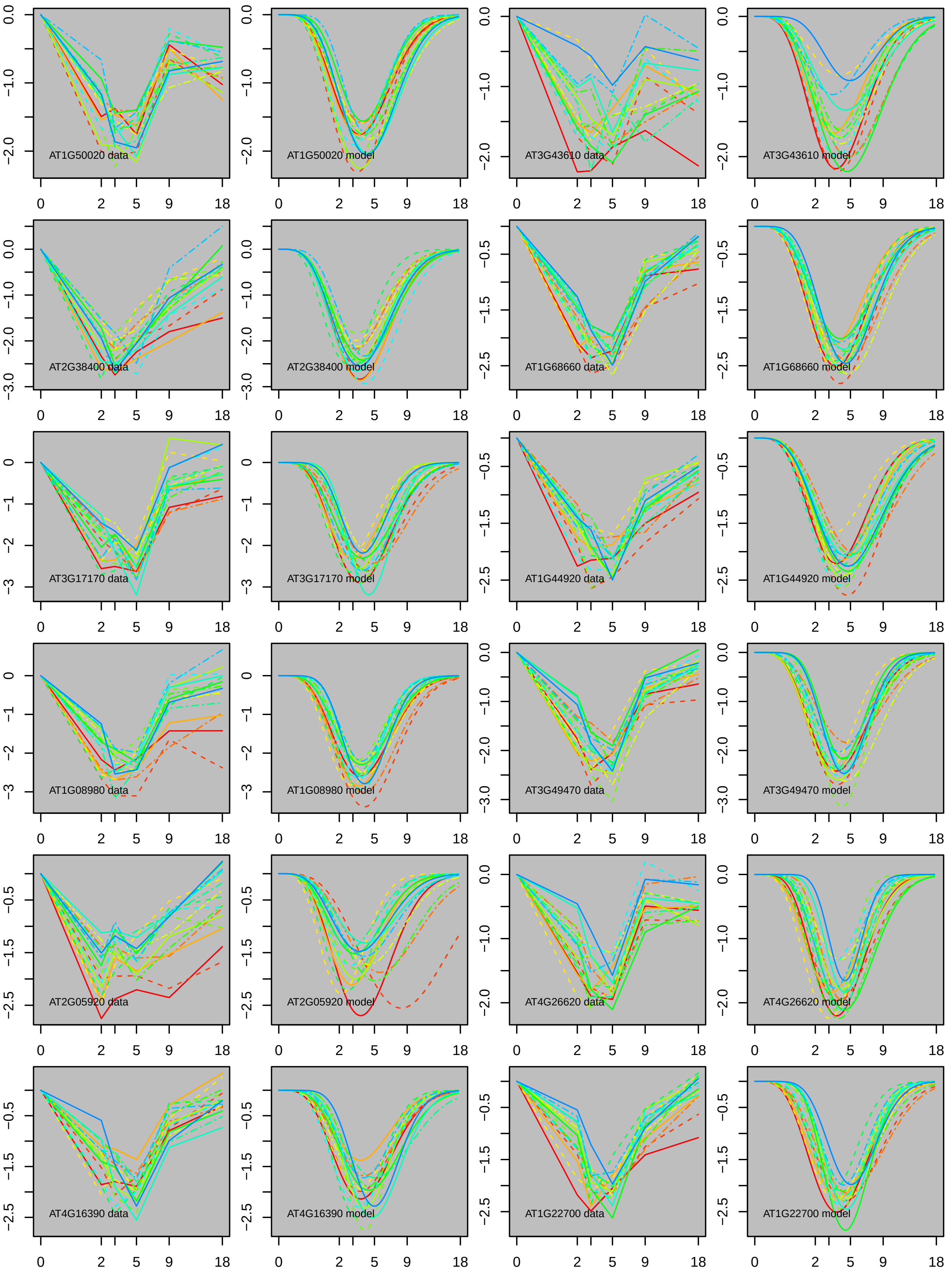

AT1G50020 data
AT1G50020 model
AT3G43610 data
AT3G43610 model
AT2G38400 data
AT2G38400 model
AT1G68660 data
AT1G68660 model
AT3G17170 data
AT3G17170 model
AT1G44920 data
AT1G44920 model
AT1G08980 data
AT1G08980 model
AT3G49470 data
AT3G49470 model
AT2G05920 data
AT2G05920 model
AT4G26620 data
AT4G26620 model
AT4G16390 data
AT4G16390 model
AT1G22700 data
AT1G22700 model

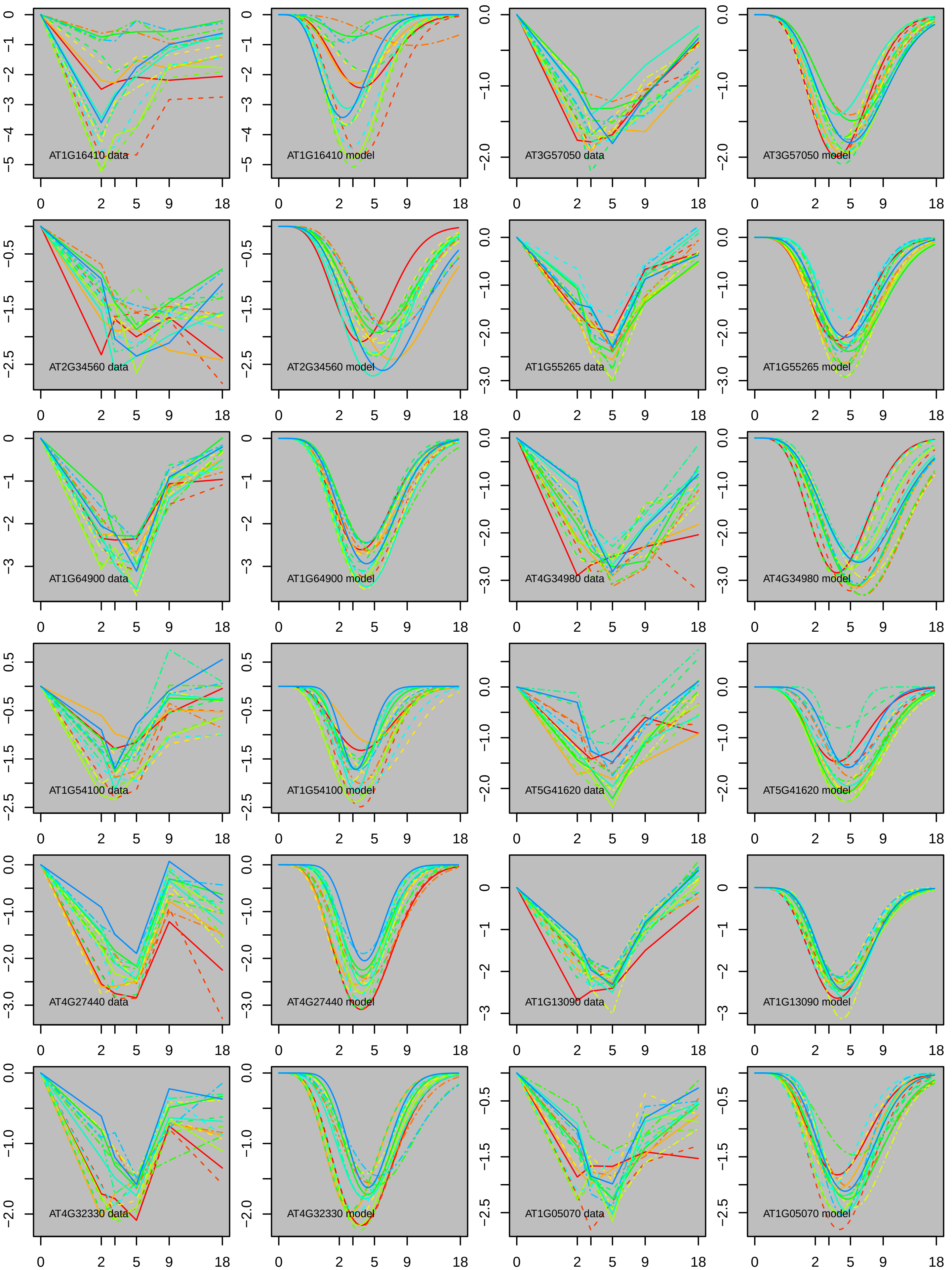
AT1G16410 data
AT1G16410 model
AT3G57050 data
AT3G57050 model
AT2G34560 data
AT2G34560 model
AT1G55265 data
AT1G55265 model
AT1G64900 data
AT1G64900 model
AT4G34980 data
AT4G34980 model
AT1G54100 data
AT1G54100 model
AT5G41620 data
AT5G41620 model
AT4G27440 data
AT4G27440 model
AT1G13090 data
AT1G13090 model
AT4G32330 data
AT4G32330 model
AT1G05070 data
AT1G05070 model

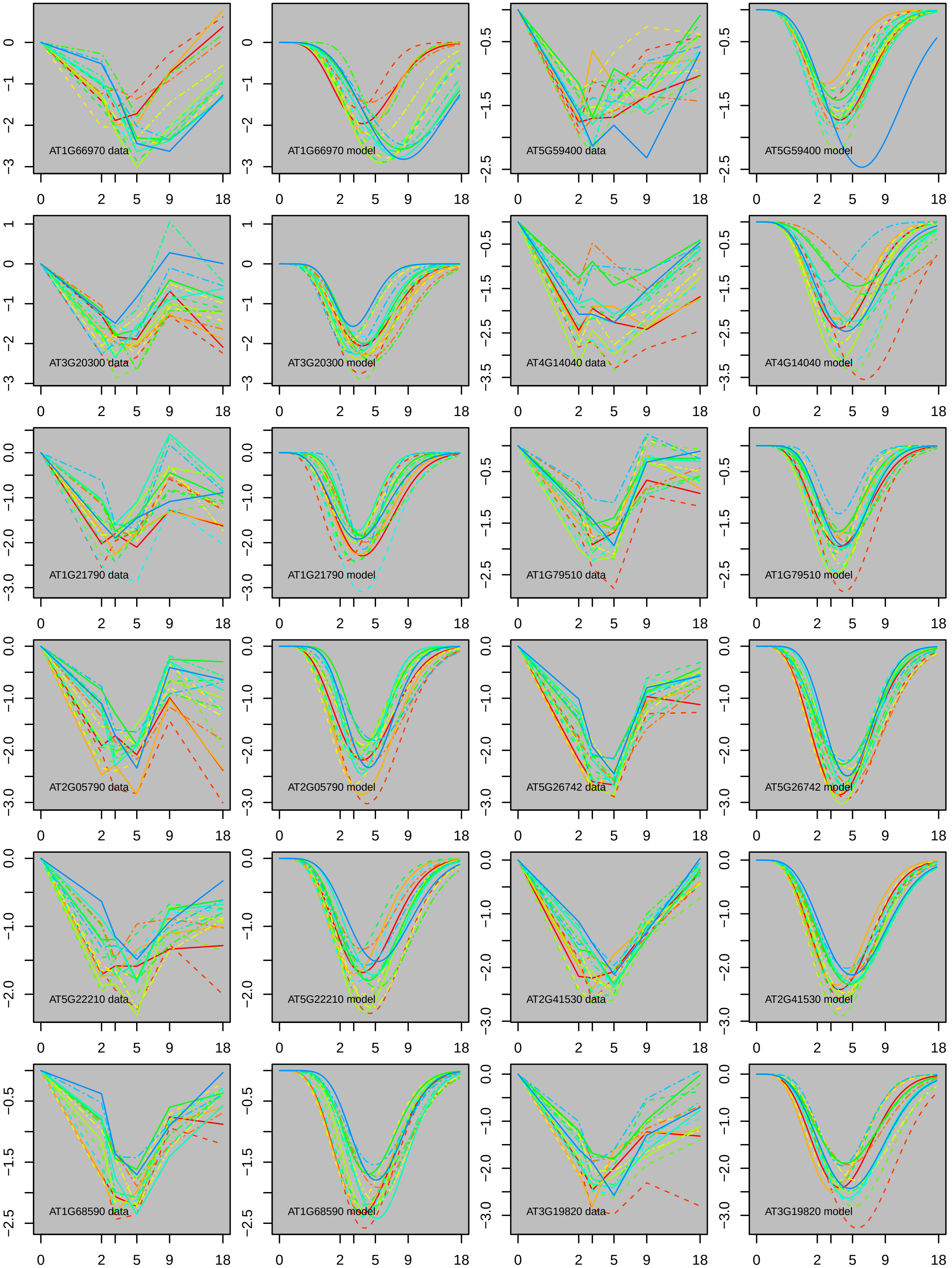
AT1G66970 data
AT1G66970 model
AT5G59400 data
AT5G59400 model
AT3G20300 data
AT3G20300 model
AT4G14040 data
AT4G14040 model
AT1G21790 data
AT1G21790 model
AT1G79510 data
AT1G79510 model
AT2G05790 data
AT2G05790 model
AT5G26742 data
AT5G26742 model
AT5G22210 data
AT5G22210 model
AT2G41530 data
AT2G41530 model
AT1G68590 data
AT1G68590 model
AT3G19820 data
AT3G19820 model
0
2
5
9
18

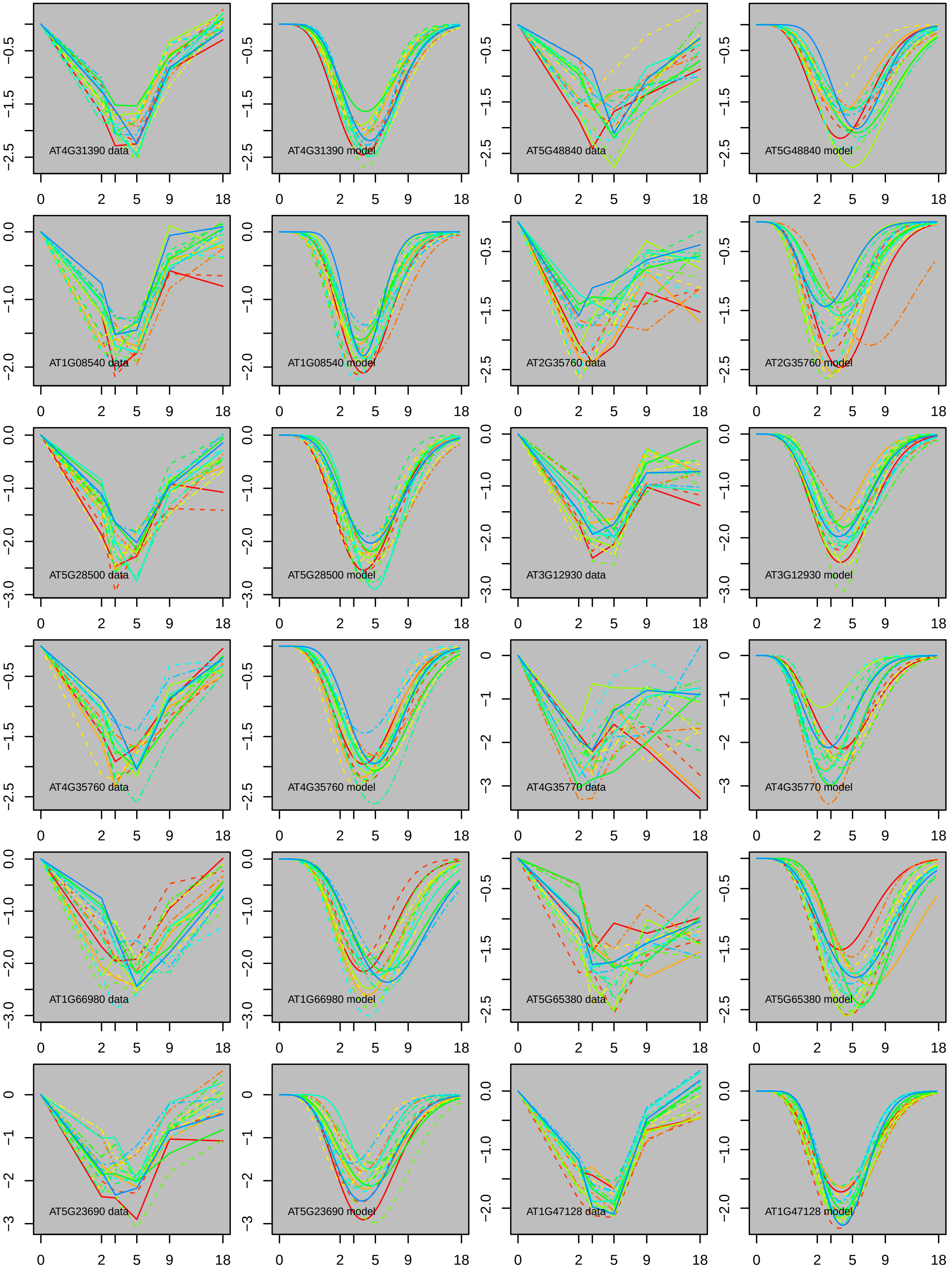
AT4G31390 data
AT4G31390 model
AT5G48840 data
AT5G48840 model
AT1G08540 data
AT1G08540 model
AT2G35760 data
AT2G35760 model
AT5G28500 data
AT5G28500 model
AT3G12930 data
AT3G12930 model
AT4G35760 data
AT4G35760 model
AT4G35770 data
AT4G35770 model
AT1G66980 data
AT1G66980 model
AT5G65380 data
AT5G65380 model
AT5G23690 data
AT5G23690 model
AT1G47128 data
AT1G47128 model

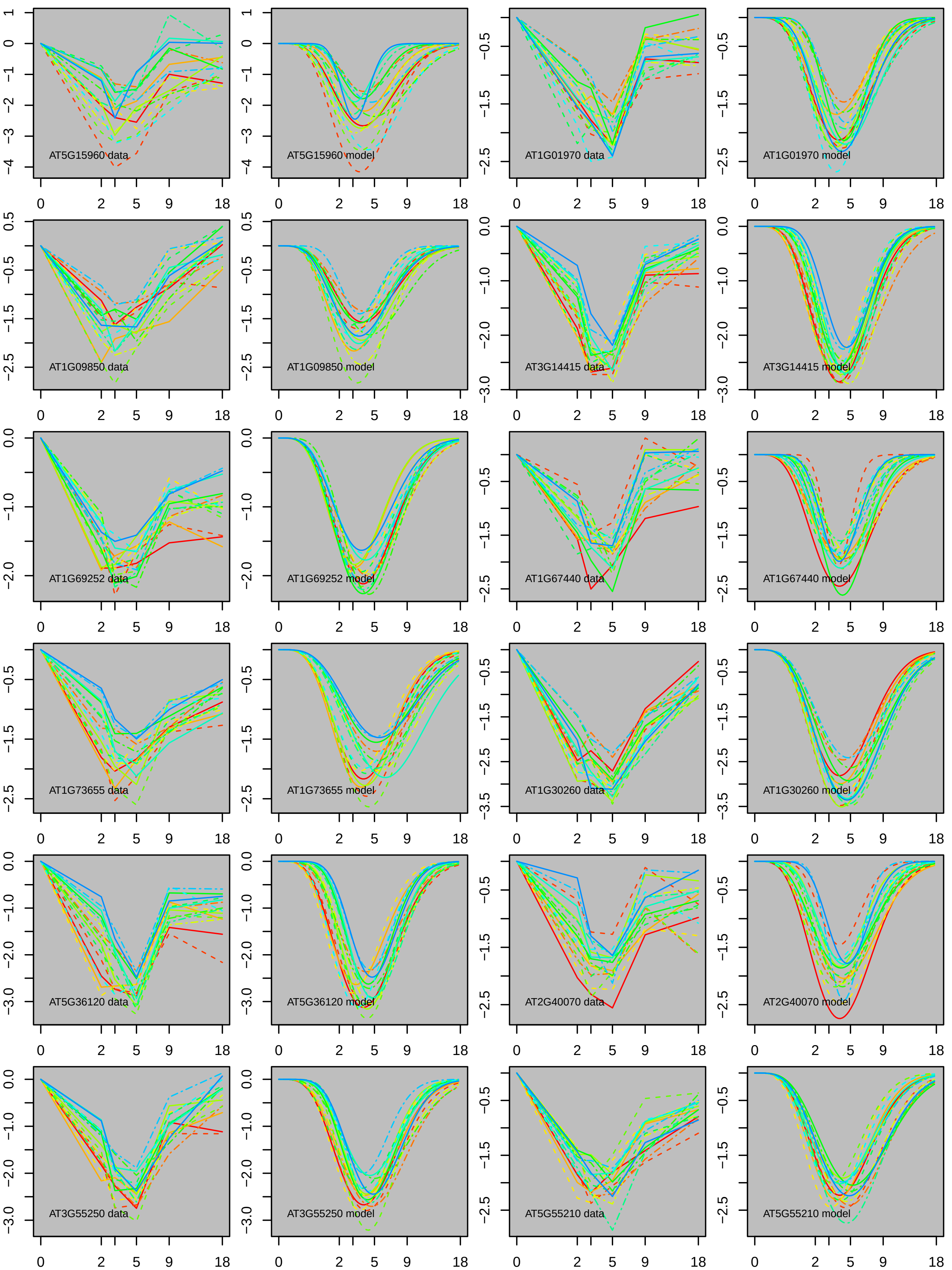

AT5G15960 data
AT5G15960 model
AT1G01970 data
AT1G01970 model
AT1G09850 data
AT1G09850 model
AT3G14415 data
AT3G14415 model
AT1G69252 data
AT1G69252 model
AT1G67440 data
AT1G67440 model
AT1G73655 data
AT1G73655 model
AT1G30260 data
AT1G30260 model
AT5G36120 data
AT5G36120 model
AT2G40070 data
AT2G40070 model
AT3G55250 data
AT3G55250 model
AT5G55210 data
AT5G55210 model

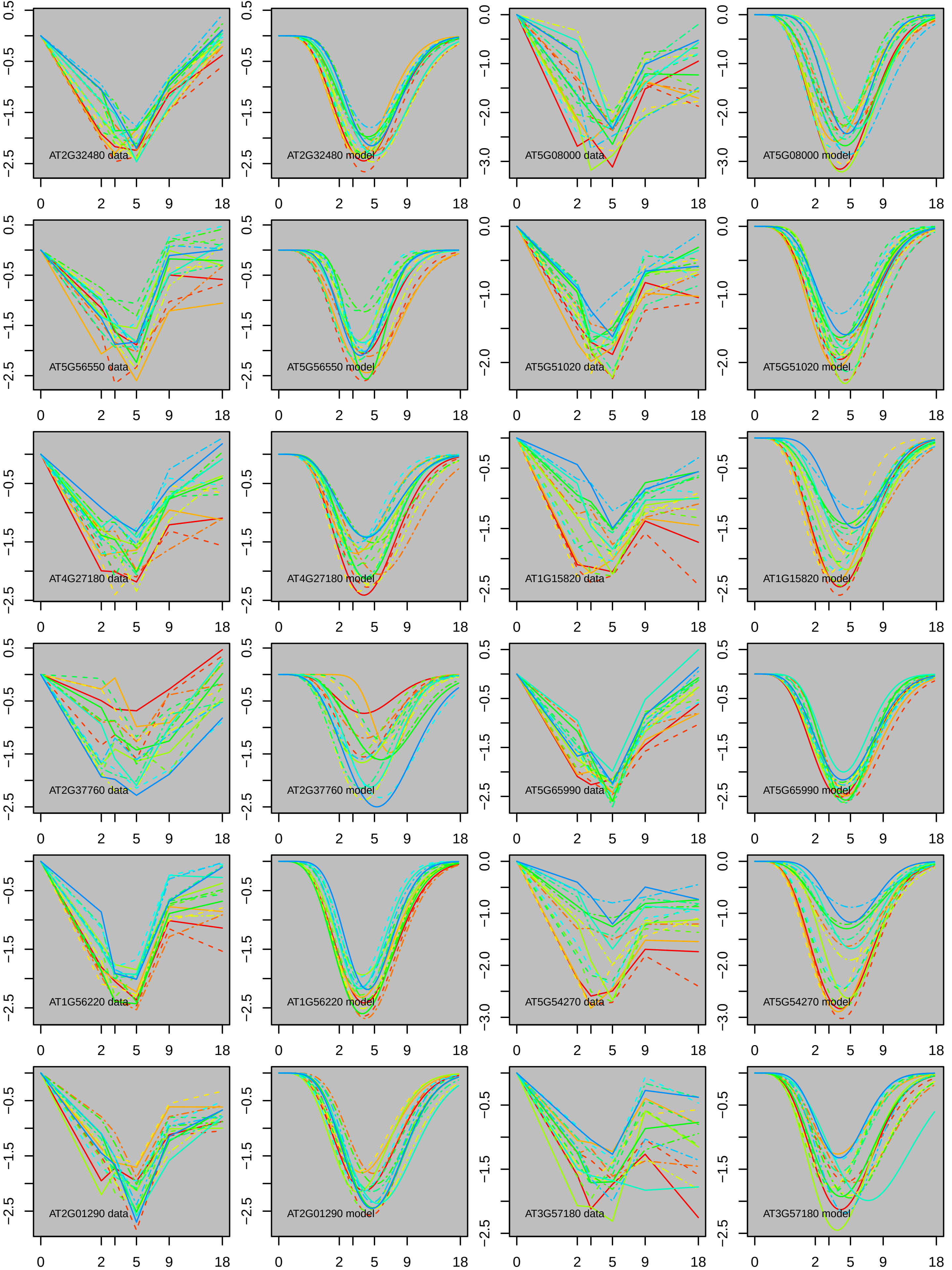
AT2G32480 data
AT2G32480 model
AT5G08000 data
AT5G08000 model
AT5G56550 data
AT5G56550 model
AT5G51020 data
AT5G51020 model
AT4G27180 data
AT4G27180 model
AT1G15820 data
AT1G15820 model
AT2G37760 data
AT2G37760 model
AT5G65990 data
AT5G65990 model
AT1G56220 data
AT1G56220 model
AT5G54270 data
AT5G54270 model
AT2G01290 data
AT2G01290 model
AT3G57180 data
AT3G57180 model

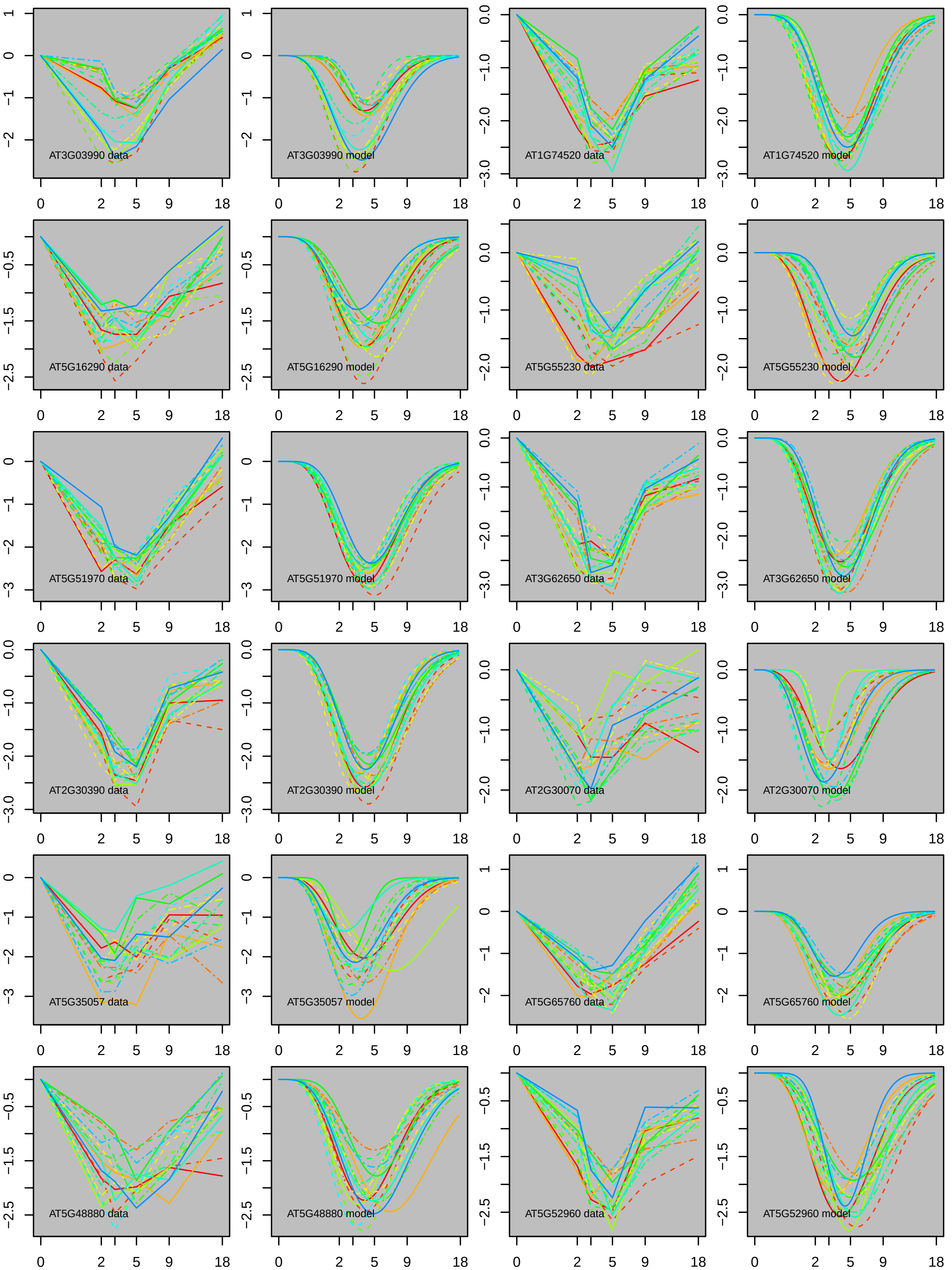
AT3G03990 data
AT3G03990 model
AT1G74520 data
AT1G74520 model
AT5G16290 data
AT5G16290 model
AT5G55230 data
AT5G55230 model
AT5G51970 data
AT5G51970 model
AT3G62650 data
AT3G62650 model
AT2G30390 data
AT2G30390 model
AT2G30070 data
AT2G30070 model
AT5G35057 data
AT5G35057 model
AT5G65760 data
AT5G65760 model
AT5G48880 data
AT5G48880 model
AT5G52960 data
AT5G52960 model

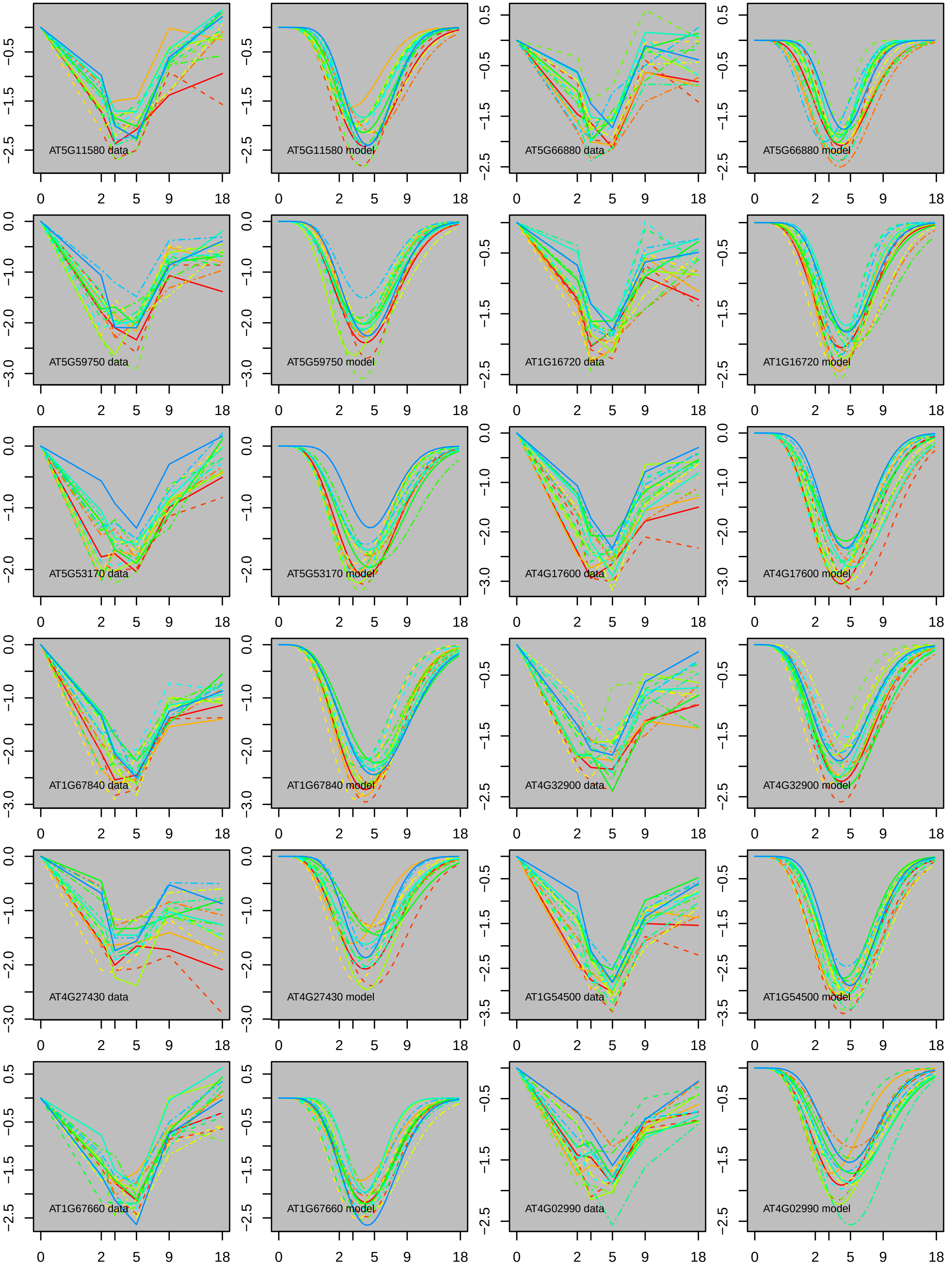

AT5G11580 data
AT5G11580 model
AT5G66880 data
AT5G66880 model
AT5G59750 data
AT5G59750 model
AT1G16720 data
AT1G16720 model
AT5G53170 data
AT5G53170 model
AT4G17600 data
AT4G17600 model
AT1G67840 data
AT1G67840 model
AT4G32900 data
AT4G32900 model
AT4G27430 data
AT4G27430 model
AT1G54500 data
AT1G54500 model
AT1G67660 data
AT1G67660 model
AT4G02990 data
AT4G02990 model

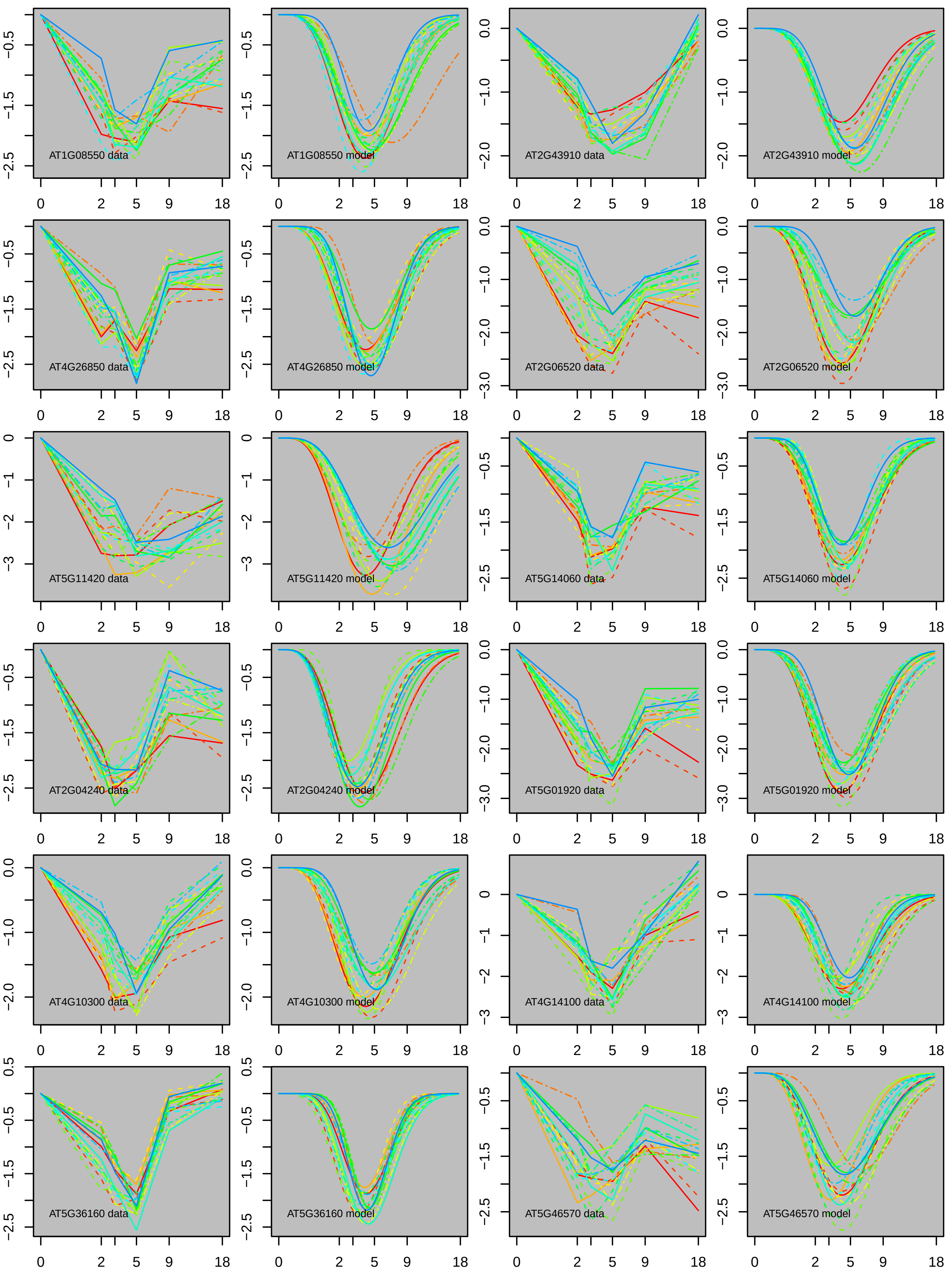
AT1G08550 data
AT1G08550 model
AT2G43910 data
AT2G43910 model
AT4G26850 data
AT4G26850 model
AT2G06520 data
AT2G06520 model
AT5G11420 data
AT5G11420 model
AT5G14060 data
AT5G14060 model
AT2G04240 data
AT2G04240 model
AT5G01920 data
AT5G01920 model
AT4G10300 data
AT4G10300 model
AT4G14100 data
AT4G14100 model
AT5G36160 data
AT5G36160 model
AT5G46570 data
AT5G46570 model
0 2 5 9 18

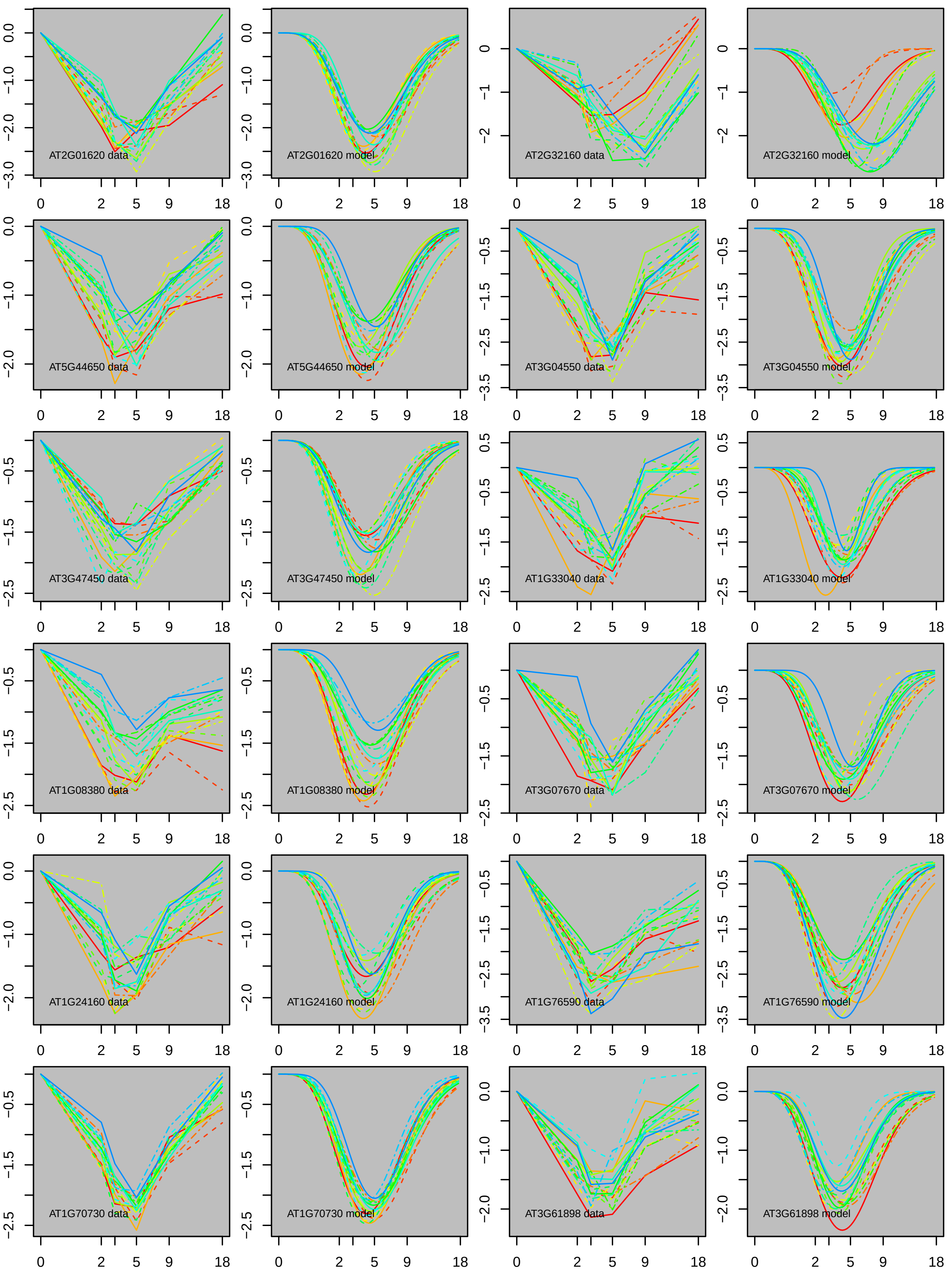
AT2G01620 data
AT2G01620 model
AT2G32160 data
AT2G32160 model
AT5G44650 data
AT5G44650 model
AT3G04550 data
AT3G04550 model
AT3G47450 data
AT3G47450 model
AT1G33040 data
AT1G33040 model
AT1G08380 data
AT1G08380 model
AT3G07670 data
AT3G07670 model
AT1G24160 data
AT1G24160 model
AT1G76590 data
AT1G76590 model
AT1G70730 data
AT1G70730 model
AT3G61898 data
AT3G61898 model

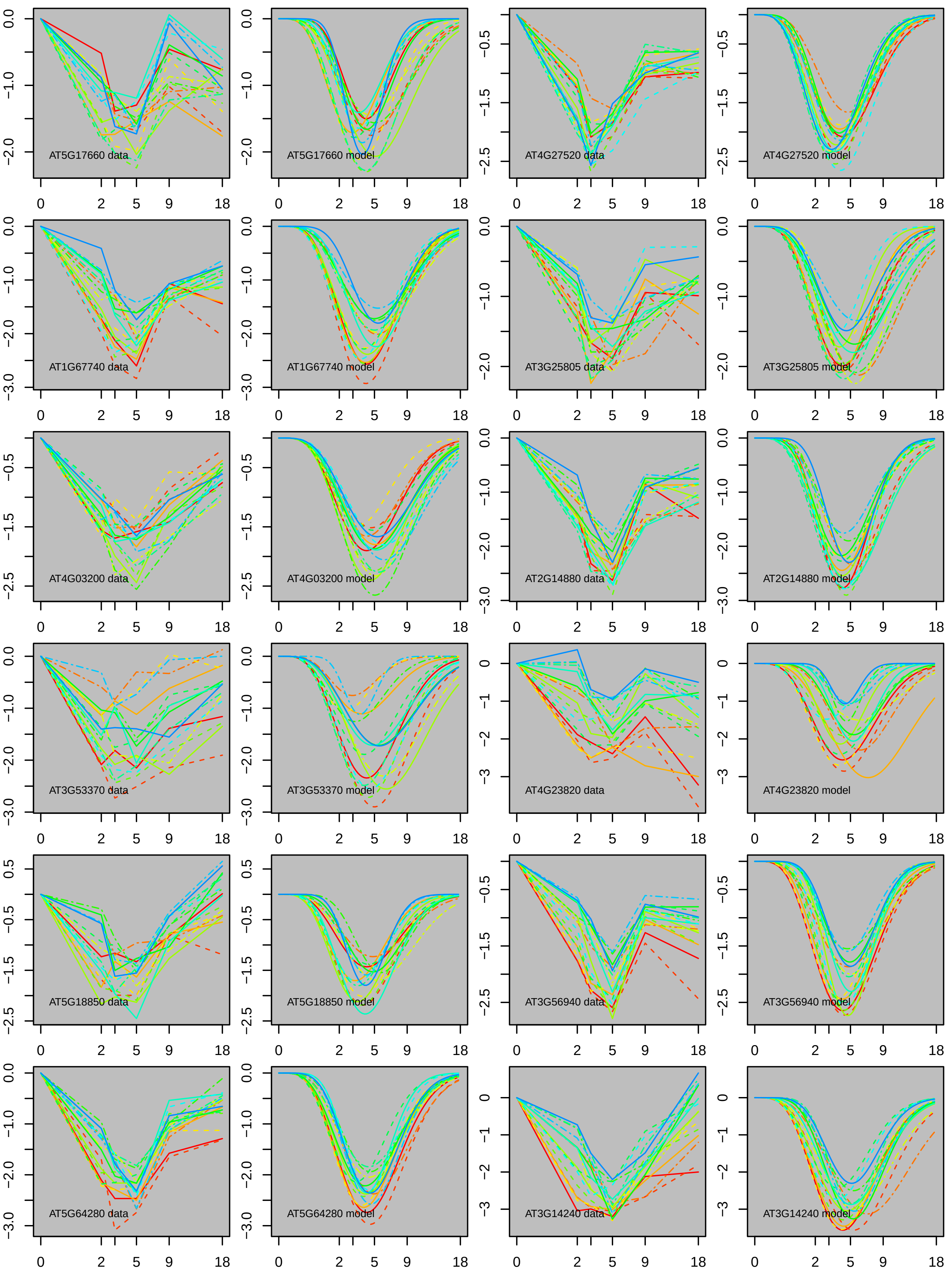

AT5G17660 data
AT5G17660 model
AT4G27520 data
AT4G27520 model
AT1G67740 data
AT1G67740 model
AT3G25805 data
AT3G25805 model
AT4G03200 data
AT4G03200 model
AT2G14880 data
AT2G14880 model
AT3G53370 data
AT3G53370 model
AT4G23820 data
AT4G23820 model
AT5G18850 data
AT5G18850 model
AT3G56940 data
AT3G56940 model
AT5G64280 data
AT5G64280 model
AT3G14240 data
AT3G14240 model

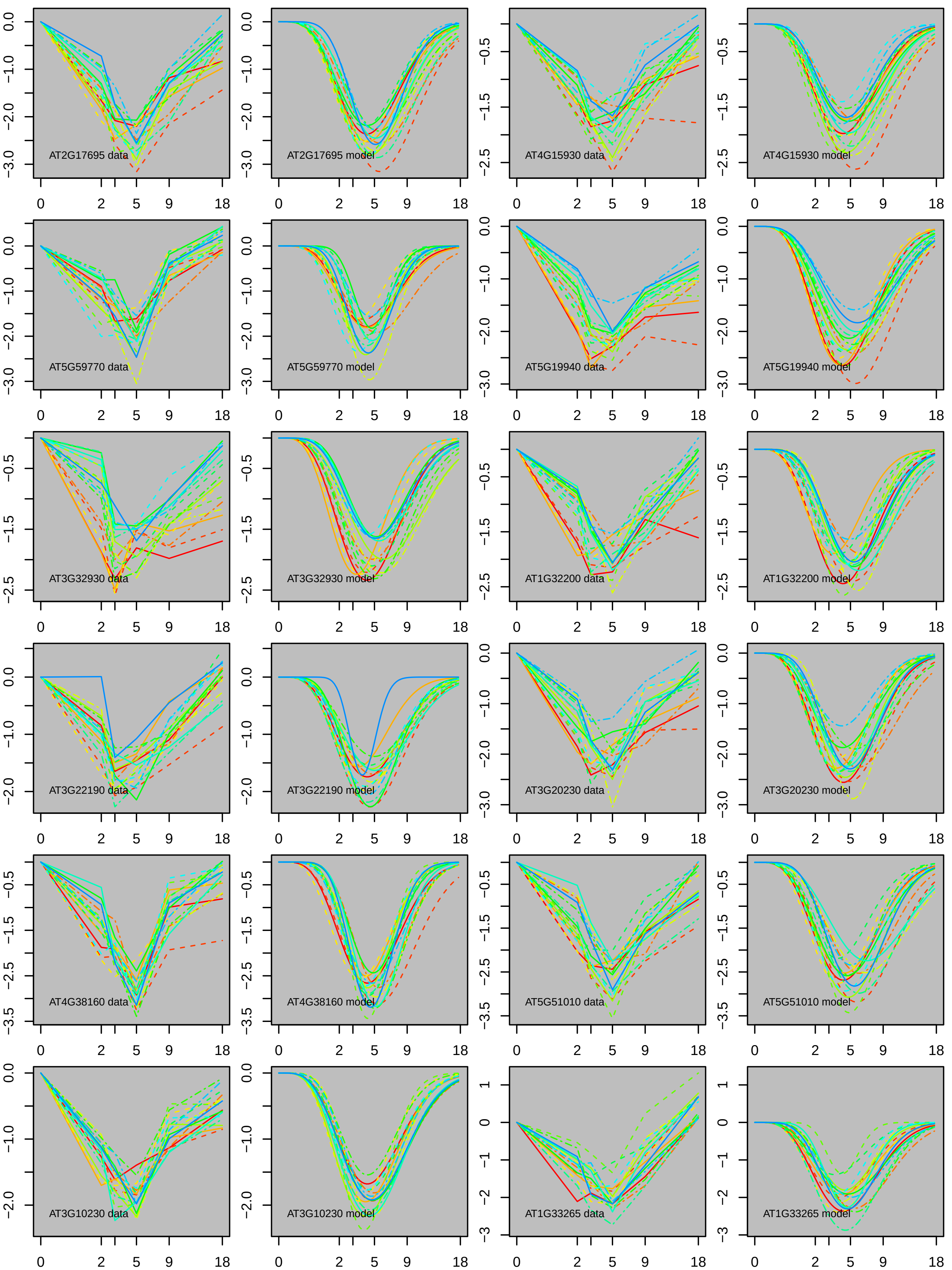
AT2G17695 data
AT2G17695 model
AT4G15930 data
AT4G15930 model
AT5G59770 data
AT5G59770 model
AT5G19940 data
AT5G19940 model
AT3G32930 data
AT3G32930 model
AT1G32200 data
AT1G32200 model
AT3G22190 data
AT3G22190 model
AT3G20230 data
AT3G20230 model
AT4G38160 data
AT4G38160 model
AT5G51010 data
AT5G51010 model
AT3G10230 data
AT3G10230 model
AT1G33265 data
AT1G33265 model

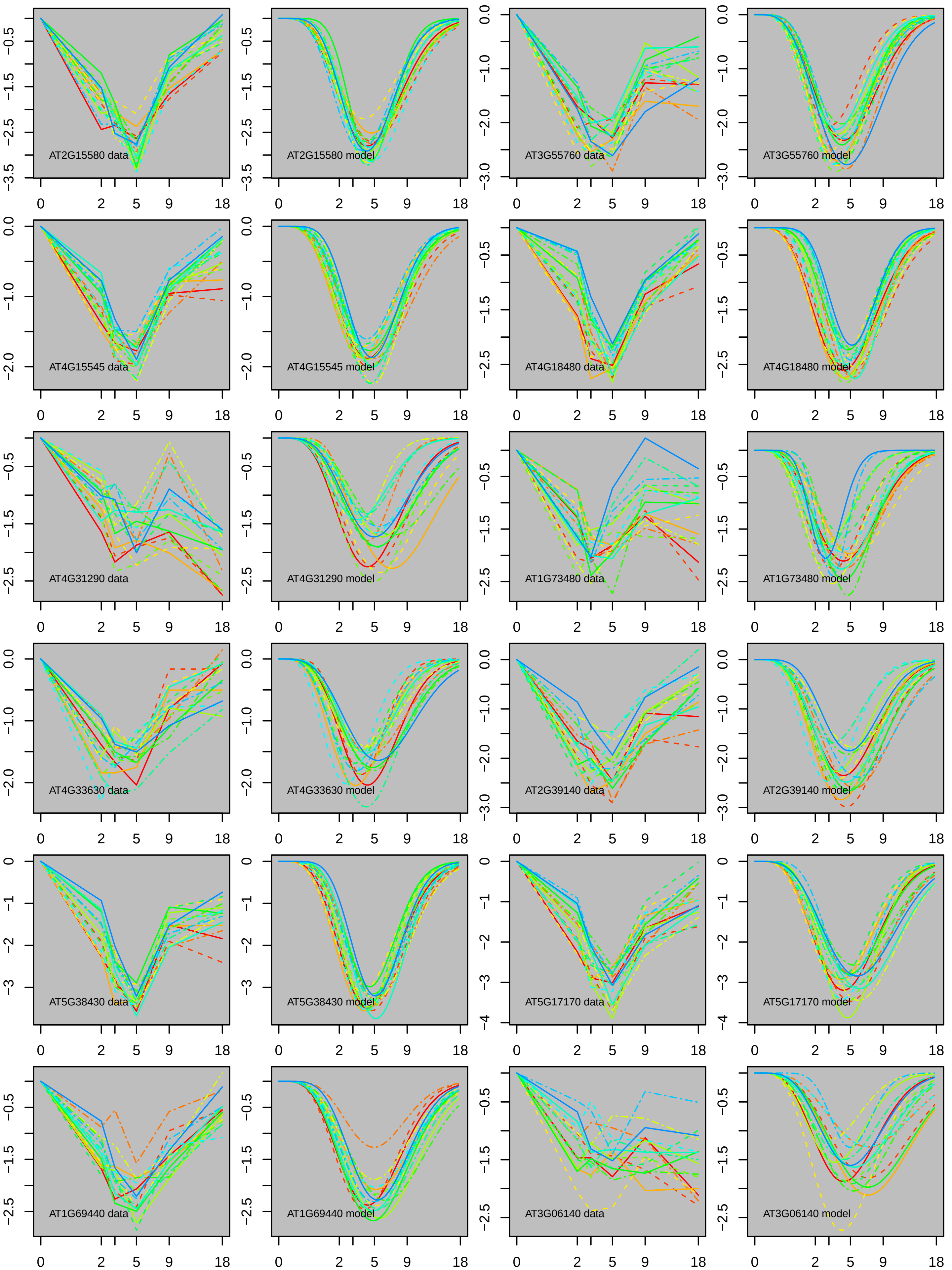
AT2G15580 data
AT2G15580 model
AT3G55760 data
AT3G55760 model
AT4G15545 data
AT4G15545 model
AT4G18480 data
AT4G18480 model
AT4G31290 data
AT4G31290 model
AT1G73480 data
AT1G73480 model
AT4G33630 data
AT4G33630 model
AT2G39140 data
AT2G39140 model
AT5G38430 data
AT5G38430 model
AT5G17170 data
AT5G17170 model
AT1G69440 data
AT1G69440 model
AT3G06140 data
AT3G06140 model

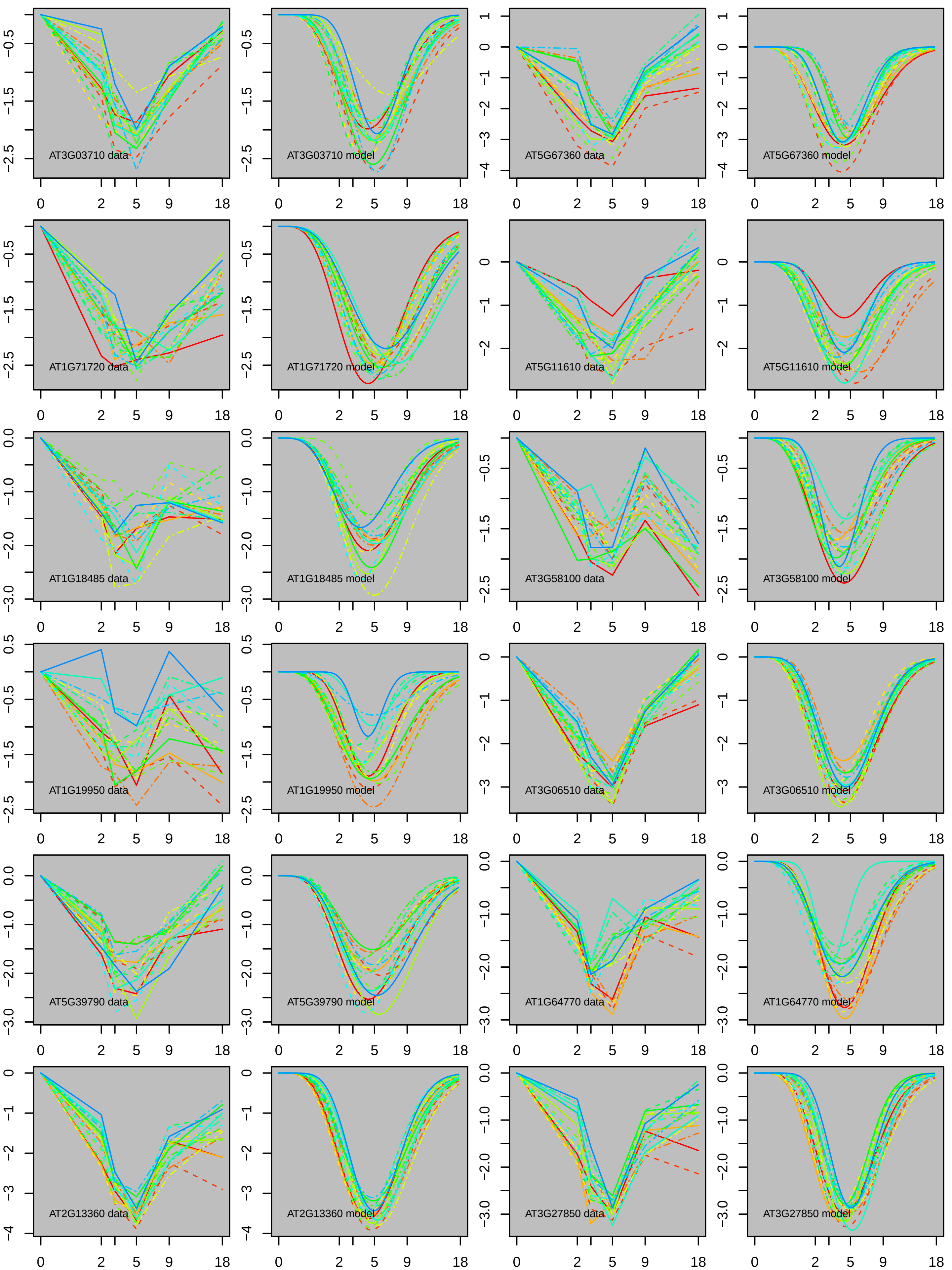
AT3G03710 data
AT3G03710 model
AT5G67360 data
AT5G67360 model
AT1G71720 data
AT1G71720 model
AT5G11610 data
AT5G11610 model
AT1G18485 data
AT1G18485 model
AT3G58100 data
AT3G58100 model
AT1G19950 data
AT1G19950 model
AT3G06510 data
AT3G06510 model
AT5G39790 data
AT5G39790 model
AT1G64770 data
AT1G64770 model
AT2G13360 data
AT2G13360 model
AT3G27850 data
AT3G27850 model

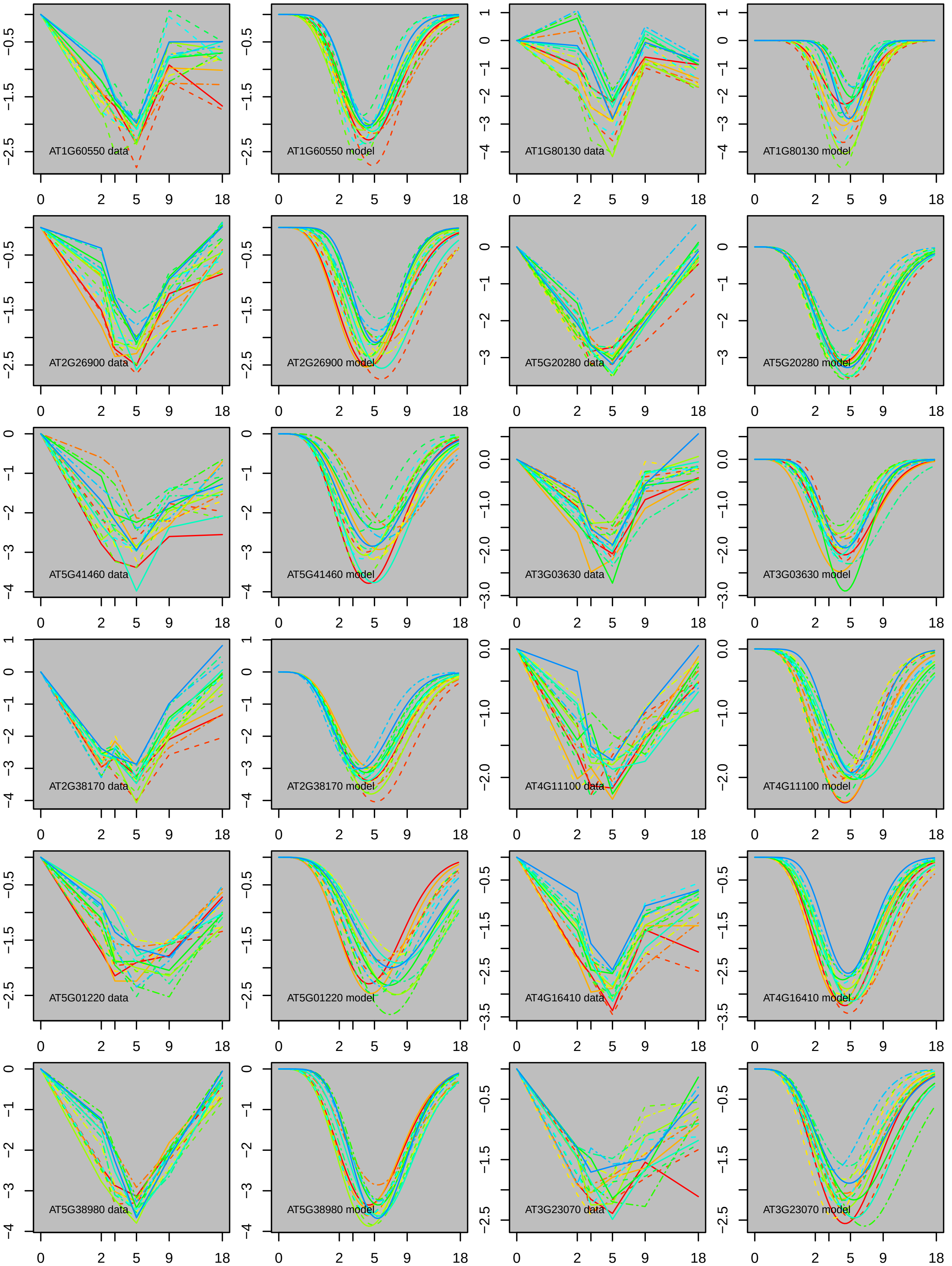
AT1G60550 data
AT1G60550 model
AT1G80130 data
AT1G80130 model
AT2G26900 data
AT2G26900 model
AT5G20280 data
AT5G20280 model
AT5G41460 data
AT5G41460 model
AT3G03630 data
AT3G03630 model
AT2G38170 data
AT2G38170 model
AT4G11100 data
AT4G11100 model
AT5G01220 data
AT5G01220 model
AT4G16410 data
AT4G16410 model
AT5G38980 data
AT5G38980 model
AT3G23070 data
AT3G23070 model

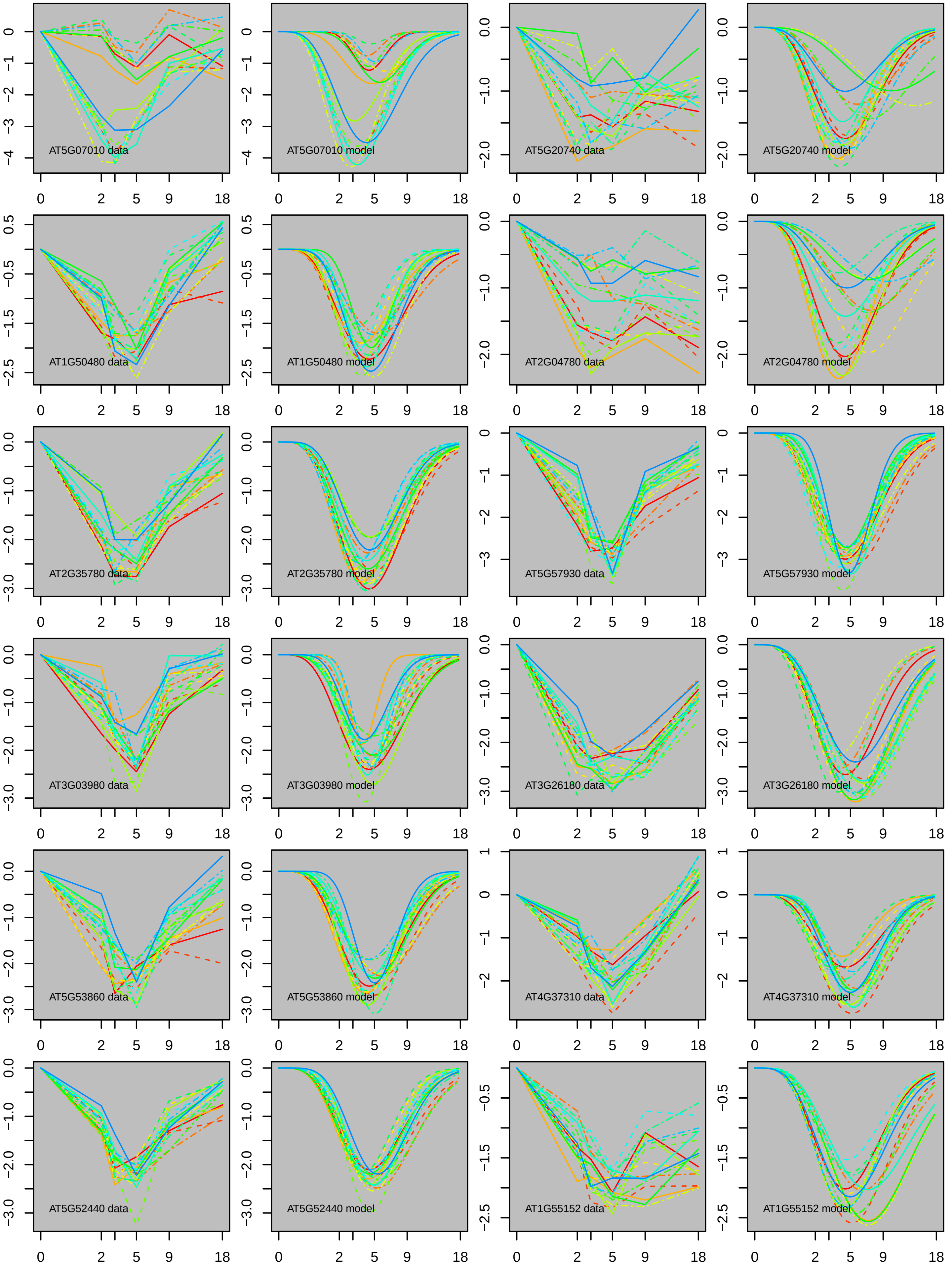

AT5G07010 data
AT5G07010 model
AT5G20740 data
AT5G20740 model
AT1G50480 data
AT1G50480 model
AT2G04780 data
AT2G04780 model
AT2G35780 data
AT2G35780 model
AT5G57930 data
AT5G57930 model
AT3G03980 data
AT3G03980 model
AT3G26180 data
AT3G26180 model
AT5G53860 data
AT5G53860 model
AT4G37310 data
AT4G37310 model
AT5G52440 data
AT5G52440 model
AT1G55152 data
AT1G55152 model

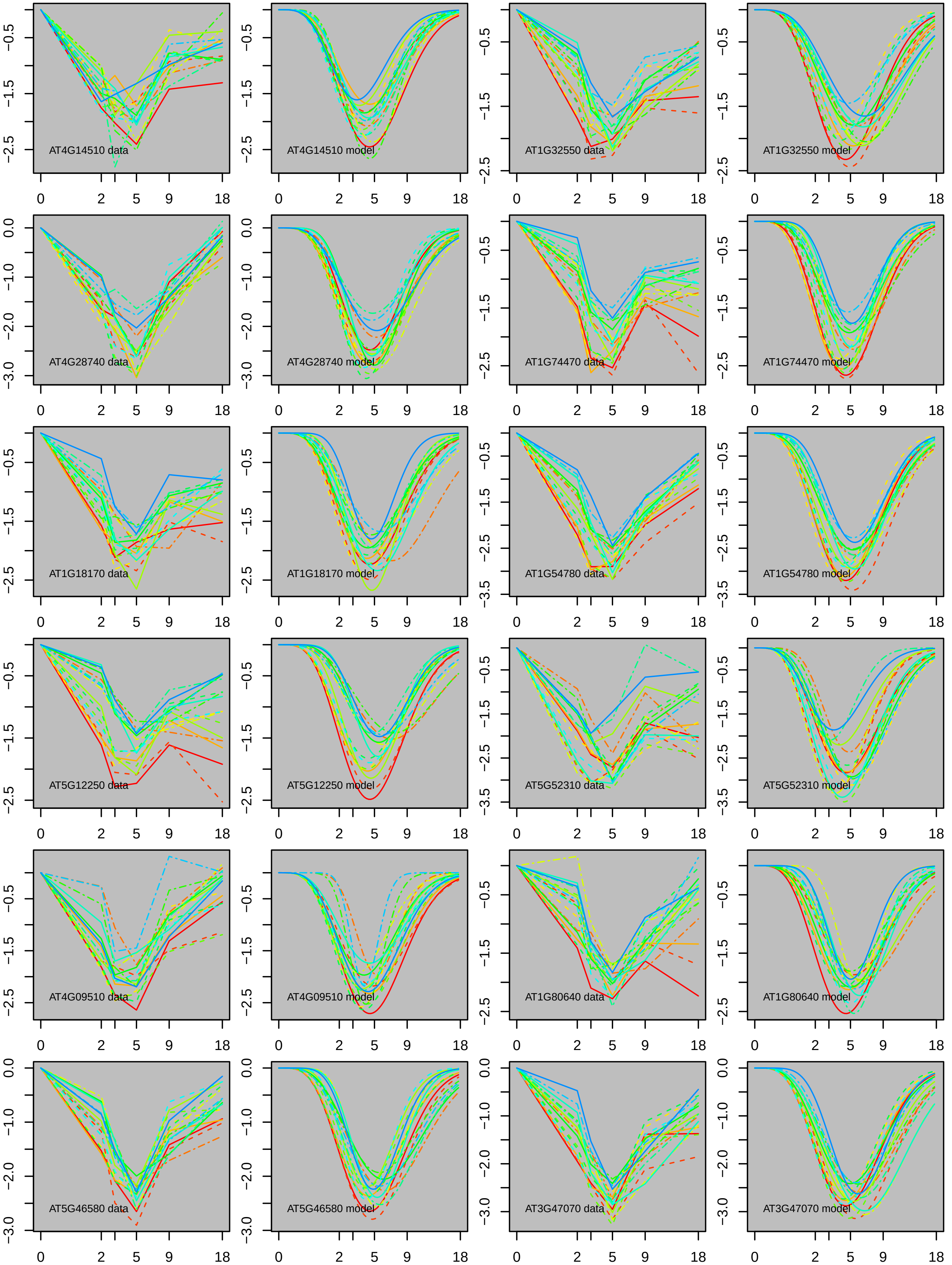

AT4G14510 data
AT4G14510 model
AT1G32550 data
AT1G32550 model
AT4G28740 data
AT4G28740 model
AT1G74470 data
AT1G74470 model
AT1G18170 data
AT1G18170 model
AT1G54780 data
AT1G54780 model
AT5G12250 data
AT5G12250 model
AT5G52310 data
AT5G52310 model
AT4G09510 data
AT4G09510 model
AT1G80640 data
AT1G80640 model
AT5G46580 data
AT5G46580 model
AT3G47070 data
AT3G47070 model
0 2 5 9 18

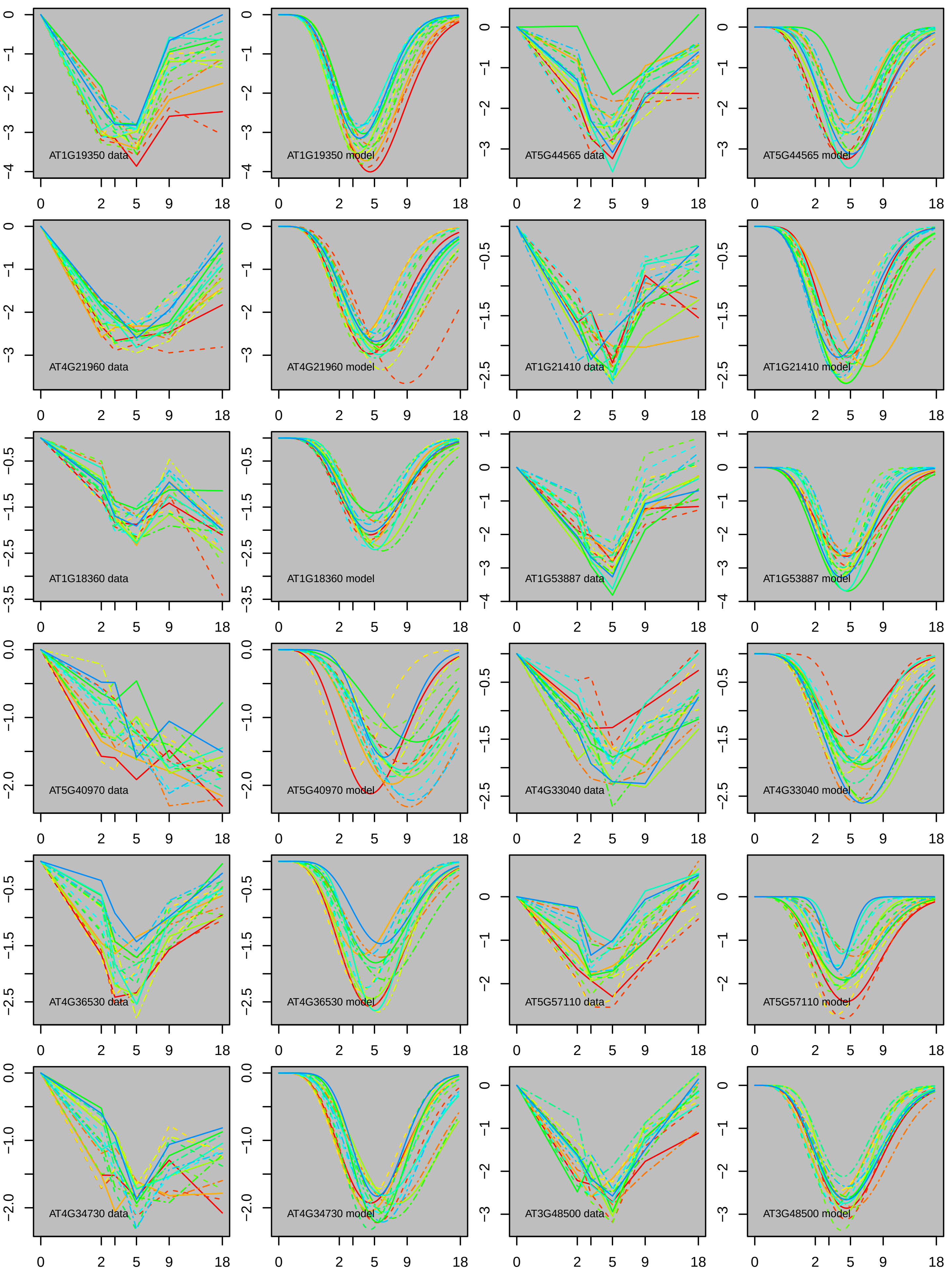
AT1G19350 data
AT1G19350 model
AT5G44565 data
AT5G44565 model
AT4G21960 data
AT4G21960 model
AT1G21410 data
AT1G21410 model
AT1G18360 data
AT1G18360 model
AT1G53887 data
AT1G53887 model
AT5G40970 data
AT5G40970 model
AT4G33040 data
AT4G33040 model
AT4G36530 data
AT4G36530 model
AT5G57110 data
AT5G57110 model
AT4G34730 data
AT4G34730 model
AT3G48500 data
AT3G48500 model

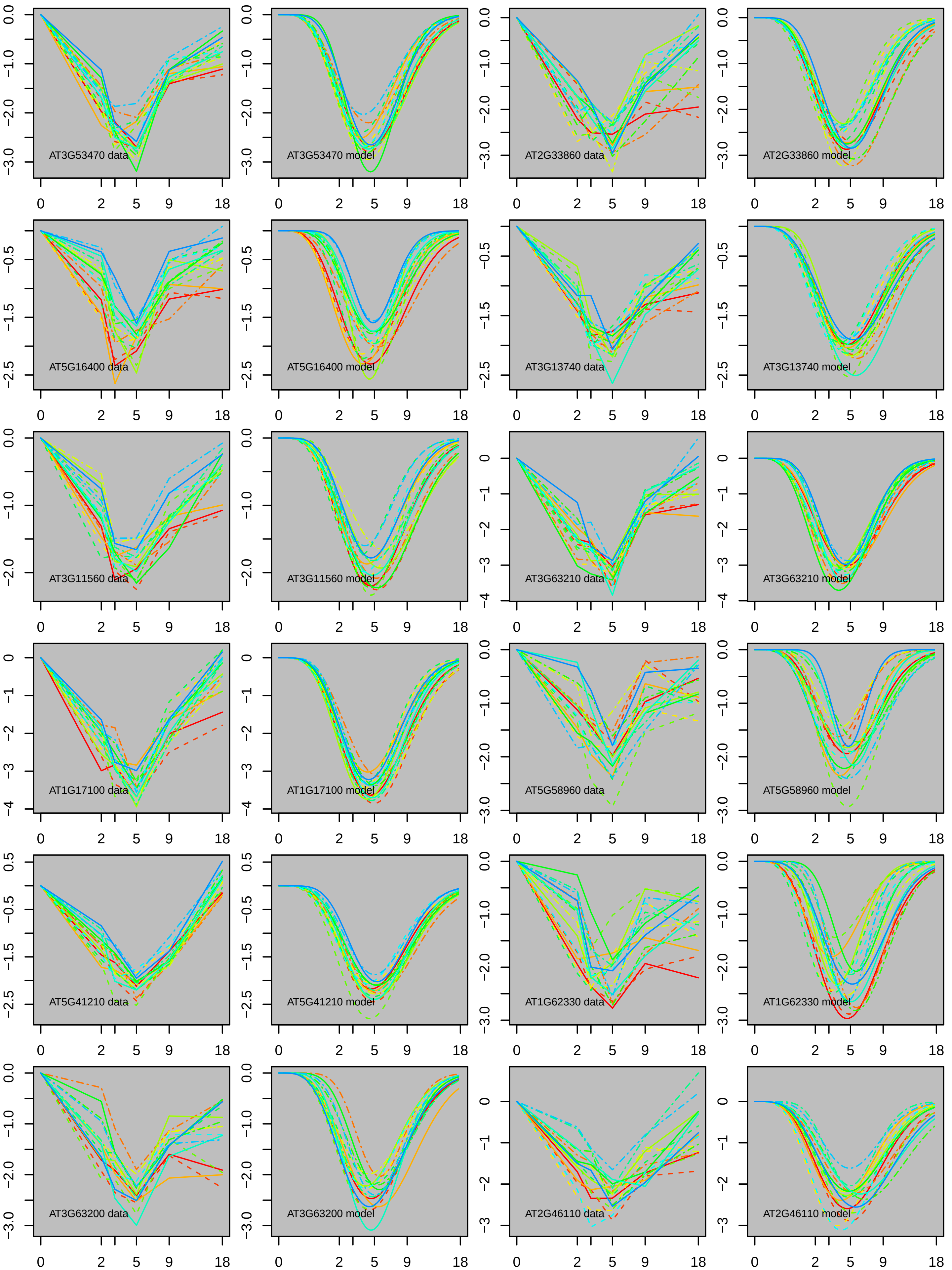

AT3G53470 data
AT3G53470 model
AT2G33860 data
AT2G33860 model
AT5G16400 data
AT5G16400 model
AT3G13740 data
AT3G13740 model
AT3G11560 data
AT3G11560 model
AT3G63210 data
AT3G63210 model
AT1G17100 data
AT1G17100 model
AT5G58960 data
AT5G58960 model
AT5G41210 data
AT5G41210 model
AT1G62330 data
AT1G62330 model
AT3G63200 data
AT3G63200 model
AT2G46110 data
AT2G46110 model

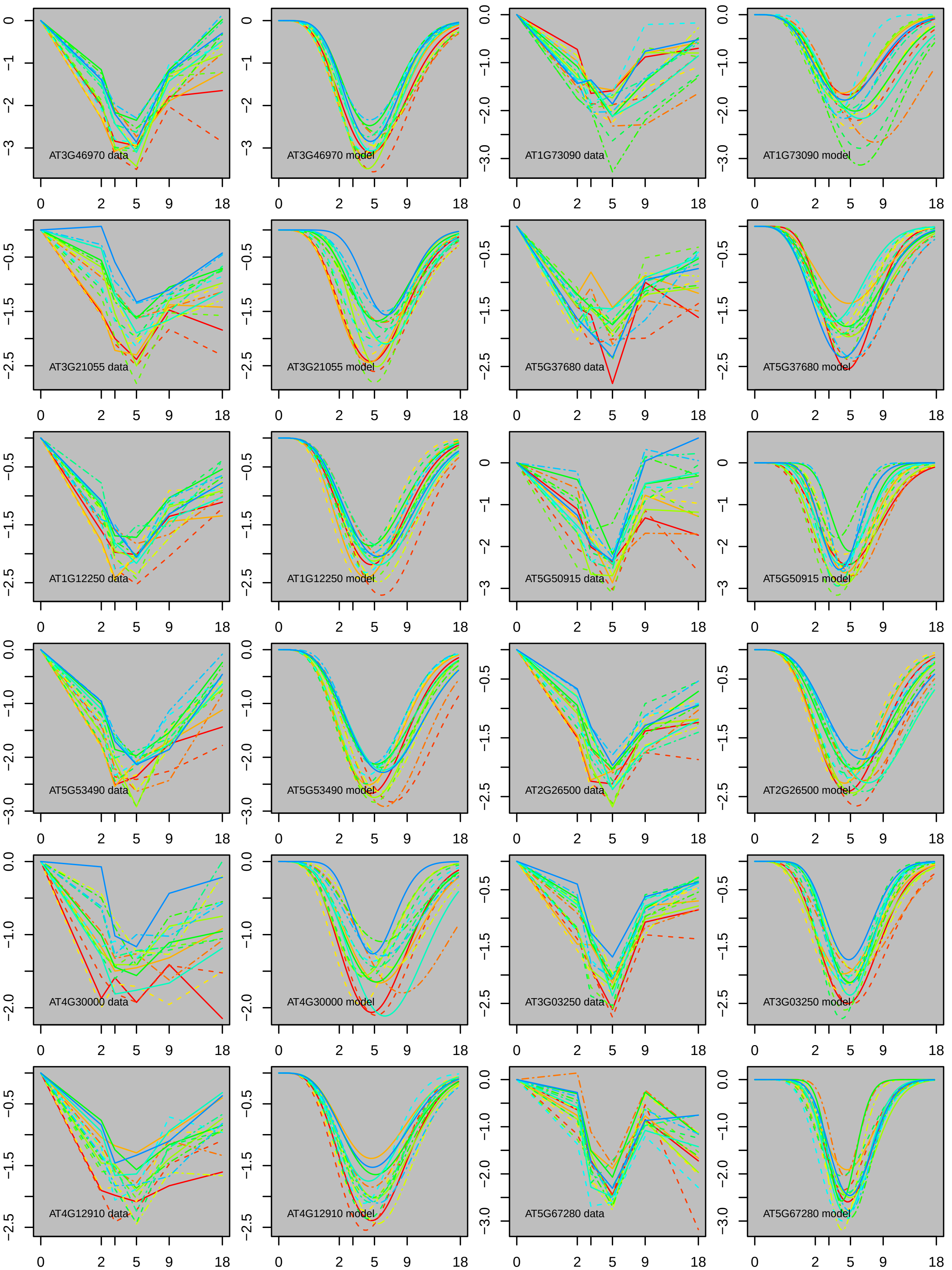

AT3G46970 data
AT3G46970 model
AT1G73090 data
AT1G73090 model
AT3G21055 data
AT3G21055 model
AT5G37680 data
AT5G37680 model
AT1G12250 data
AT1G12250 model
AT5G50915 data
AT5G50915 model
AT5G53490 data
AT5G53490 model
AT2G26500 data
AT2G26500 model
AT4G30000 data
AT4G30000 model
AT3G03250 data
AT3G03250 model
AT4G12910 data
AT4G12910 model
AT5G67280 data
AT5G67280 model

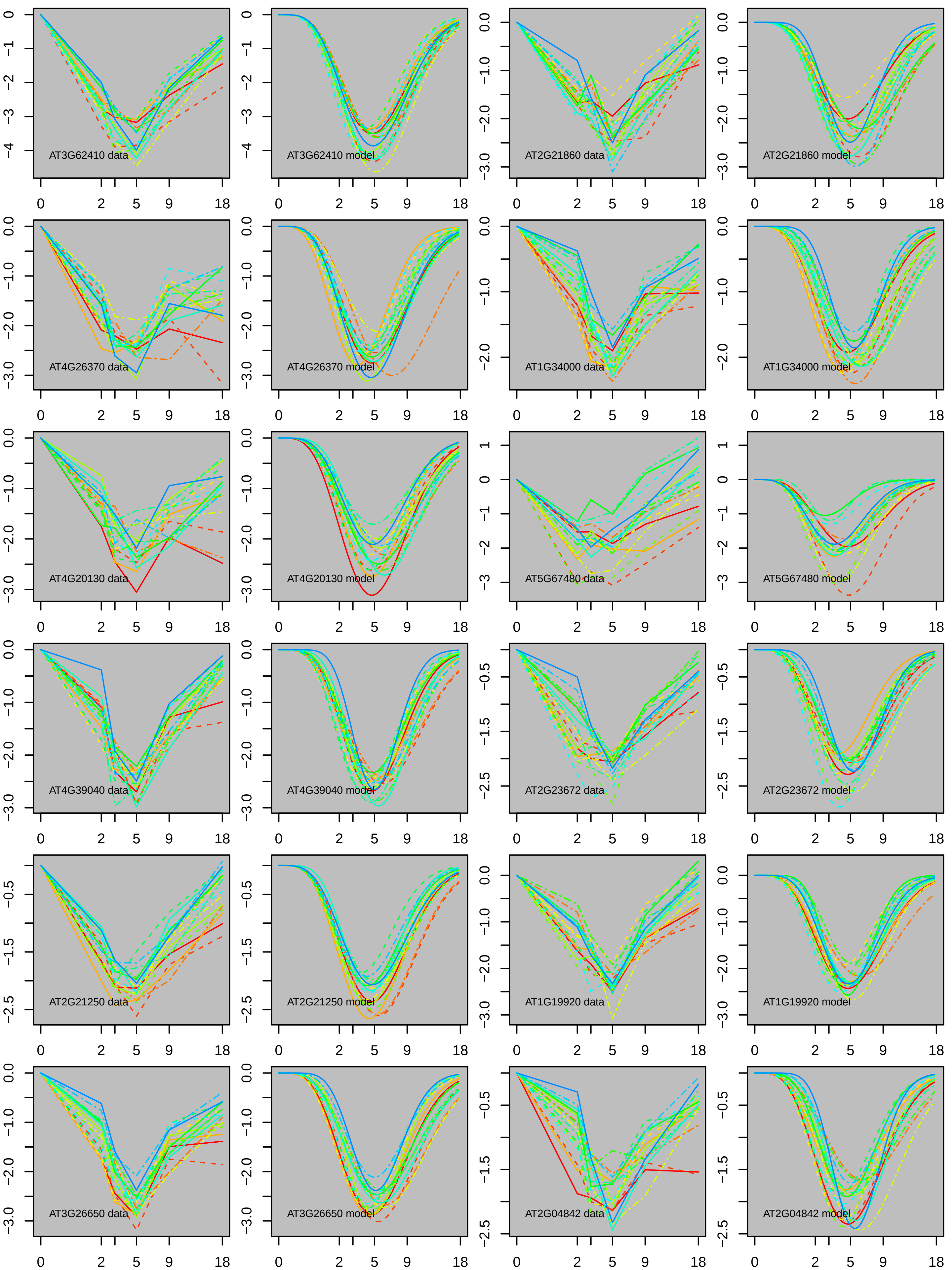
AT3G62410 data
AT3G62410 model
AT2G21860 data
AT2G21860 model
AT4G26370 data
AT4G26370 model
AT1G34000 data
AT1G34000 model
AT4G20130 data
AT4G20130 model
AT5G67480 data
AT5G67480 model
AT4G39040 data
AT4G39040 model
AT2G23672 data
AT2G23672 model
AT2G21250 data
AT2G21250 model
AT1G19920 data
AT1G19920 model
AT3G26650 data
AT3G26650 model
AT2G04842 data
AT2G04842 model
0 2 5 9 18

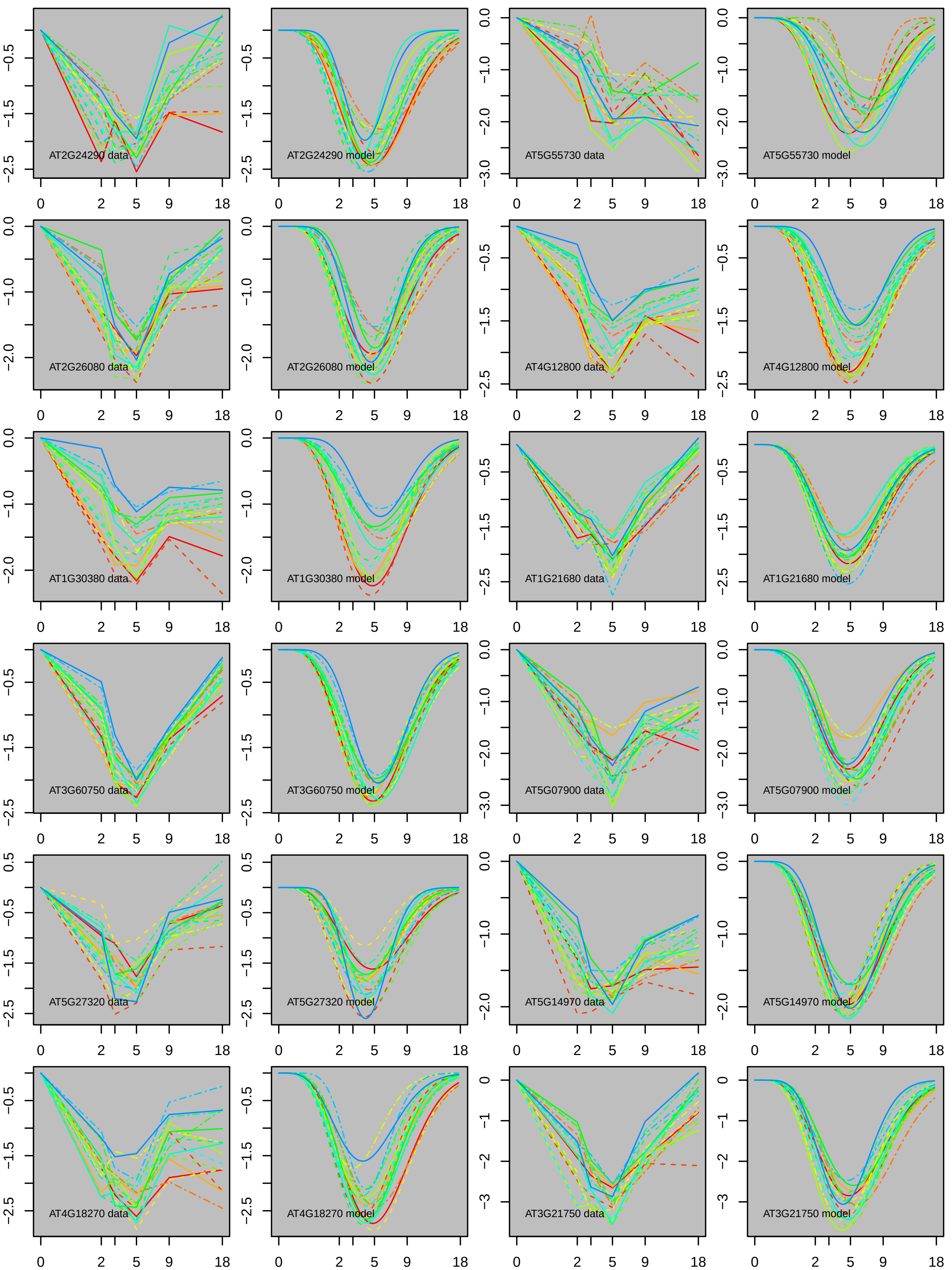
AT2G24290 data
AT2G24290 model
AT5G55730 data
AT5G55730 model
AT2G26080 data
AT2G26080 model
AT4G12800 data
AT4G12800 model
AT1G30380 data
AT1G30380 model
AT1G21680 data
AT1G21680 model
AT3G60750 data
AT3G60750 model
AT5G07900 data
AT5G07900 model
AT5G27320 data
AT5G27320 model
AT5G14970 data
AT5G14970 model
AT4G18270 data
AT4G18270 model
AT3G21750 data
AT3G21750 model
0 2 5 9 18

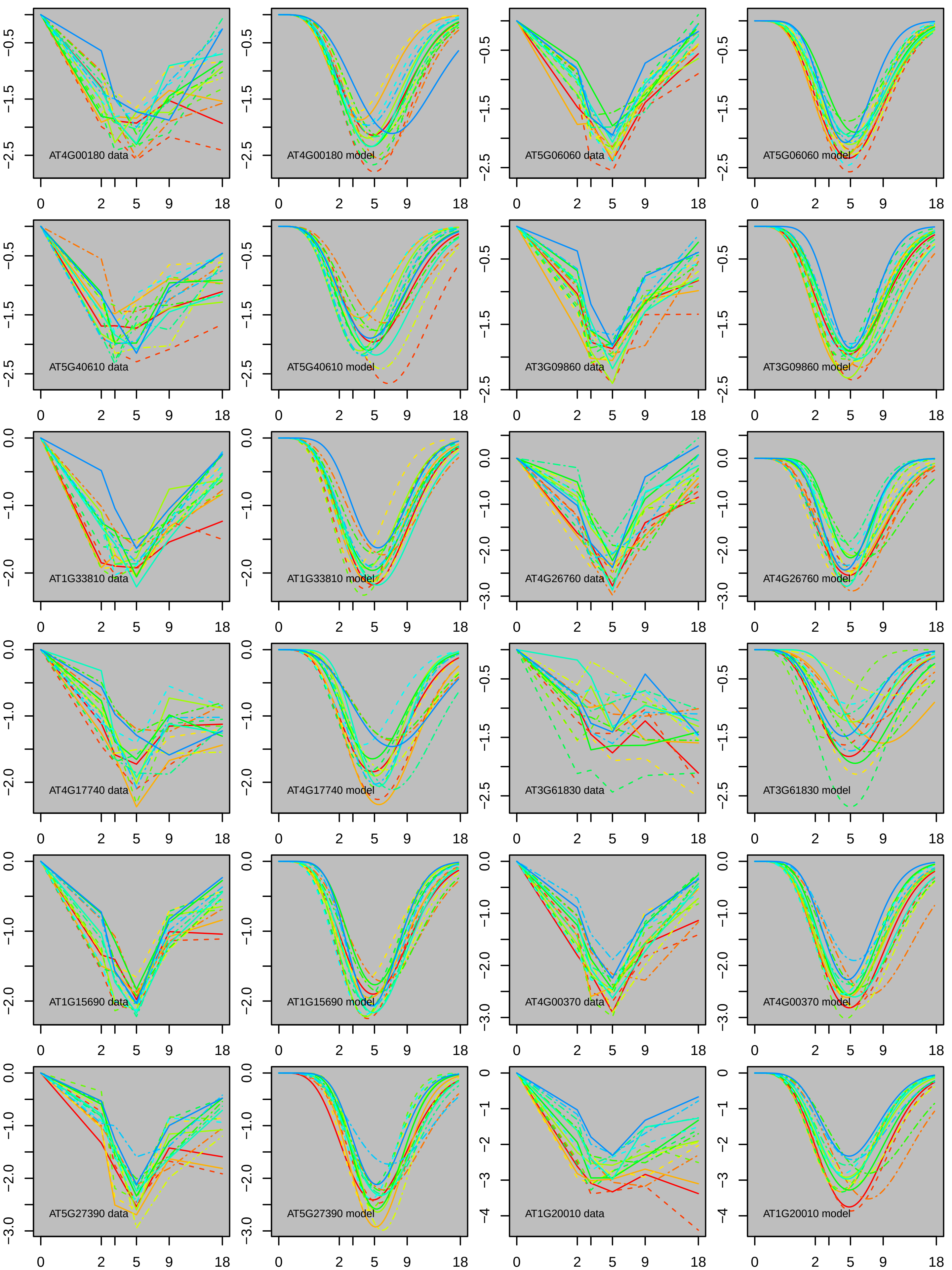
AT4G00180 data
AT4G00180 model
AT5G06060 data
AT5G06060 model
AT5G40610 data
AT5G40610 model
AT3G09860 data
AT3G09860 model
AT1G33810 data
AT1G33810 model
AT4G26760 data
AT4G26760 model
AT4G17740 data
AT4G17740 model
AT3G61830 data
AT3G61830 model
AT1G15690 data
AT1G15690 model
AT4G00370 data
AT4G00370 model
AT5G27390 data
AT5G27390 model
AT1G20010 data
AT1G20010 model

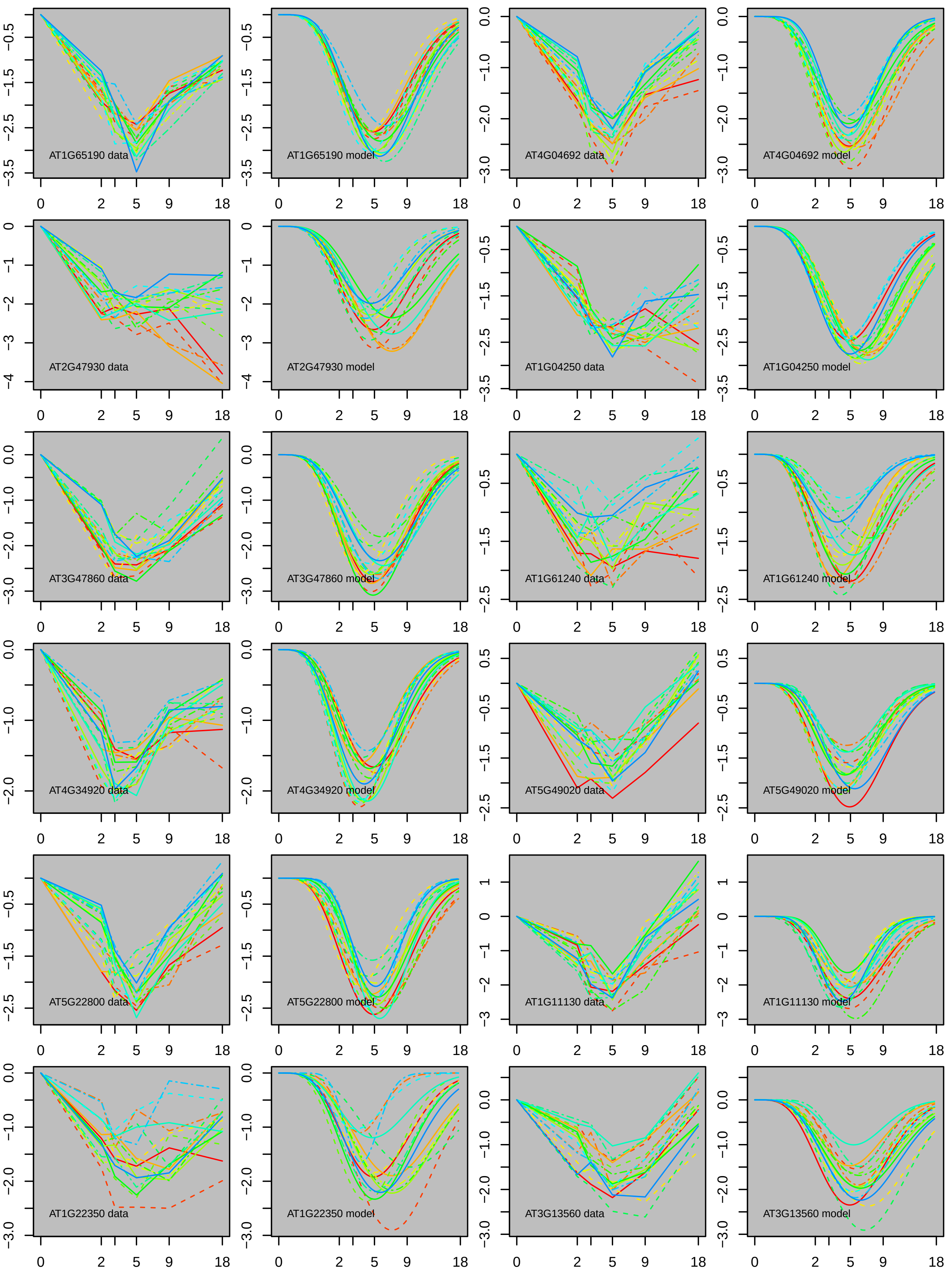
AT1G65190 data
AT1G65190 model
AT4G04692 data
AT4G04692 model
AT2G47930 data
AT2G47930 model
AT1G04250 data
AT1G04250 model
AT3G47860 data
AT3G47860 model
AT1G61240 data
AT1G61240 model
AT4G34920 data
AT4G34920 model
AT5G49020 data
AT5G49020 model
AT5G22800 data
AT5G22800 model
AT1G11130 data
AT1G11130 model
AT1G22350 data
AT1G22350 model
AT3G13560 data
AT3G13560 model

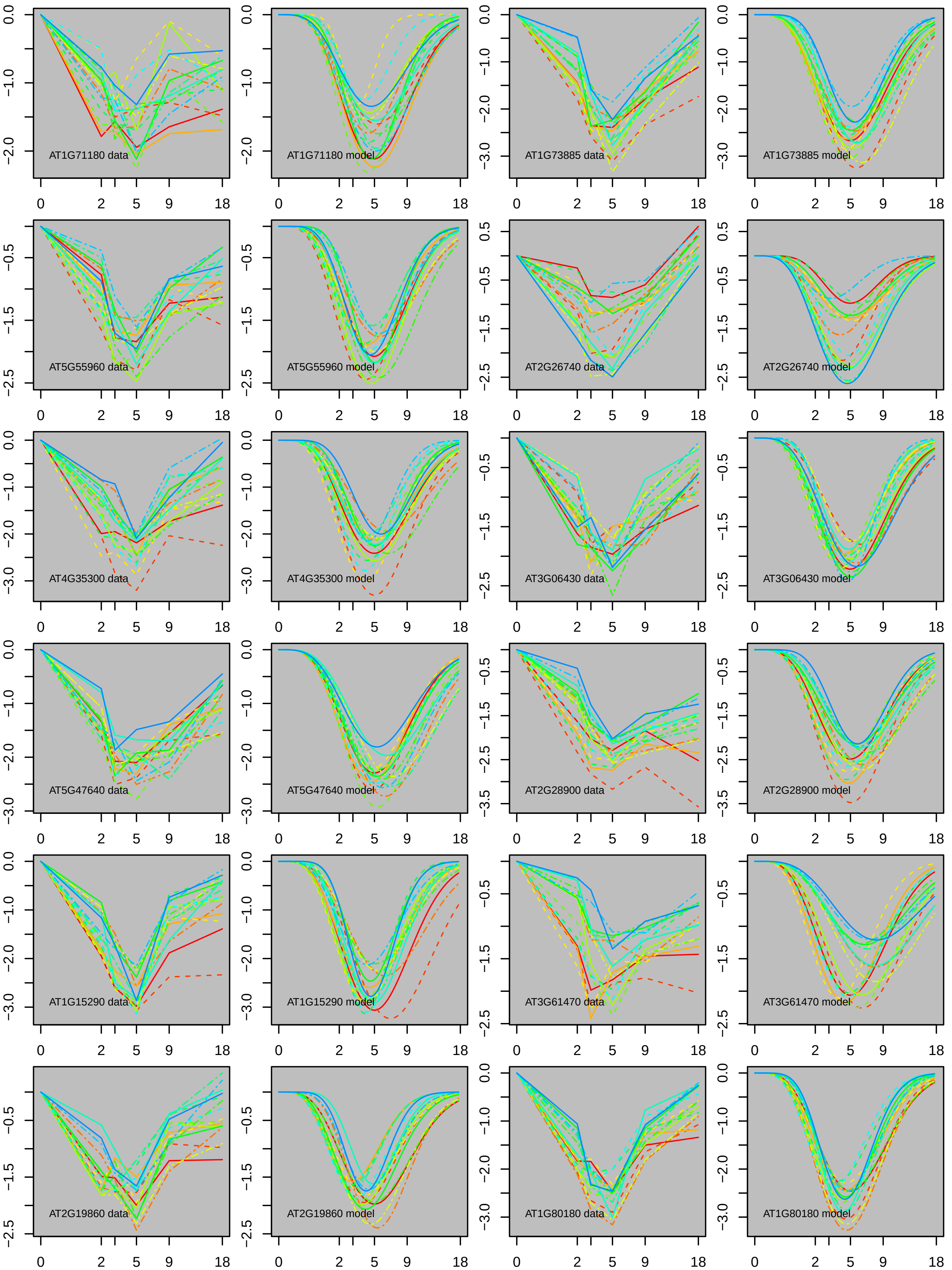
AT1G71180 data
AT1G71180 model
AT1G73885 data
AT1G73885 model
AT5G55960 data
AT5G55960 model
AT2G26740 data
AT2G26740 model
AT4G35300 data
AT4G35300 model
AT3G06430 data
AT3G06430 model
AT5G47640 data
AT5G47640 model
AT2G28900 data
AT2G28900 model
AT1G15290 data
AT1G15290 model
AT3G61470 data
AT3G61470 model
AT2G19860 data
AT2G19860 model
AT1G80180 data
AT1G80180 model
0 2 5 9 18

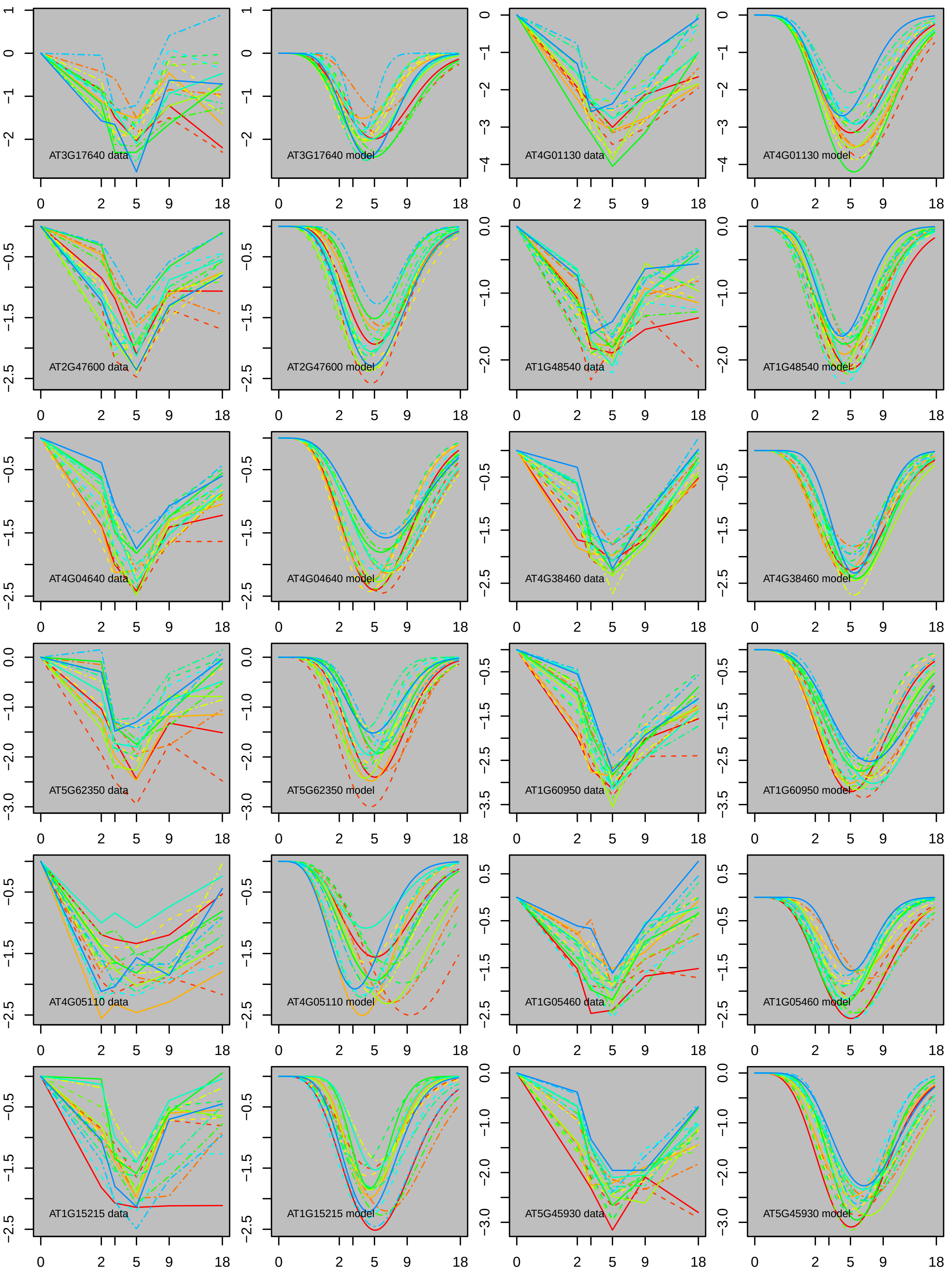
AT3G17640 data
AT3G17640 model
AT4G01130 data
AT4G01130 model
AT2G47600 data
AT2G47600 model
AT1G48540 data
AT1G48540 model
AT4G04640 data
AT4G04640 model
AT4G38460 data
AT4G38460 model
AT5G62350 data
AT5G62350 model
AT1G60950 data
AT1G60950 model
AT4G05110 data
AT4G05110 model
AT1G05460 data
AT1G05460 model
AT1G15215 data
AT1G15215 model
AT5G45930 data
AT5G45930 model

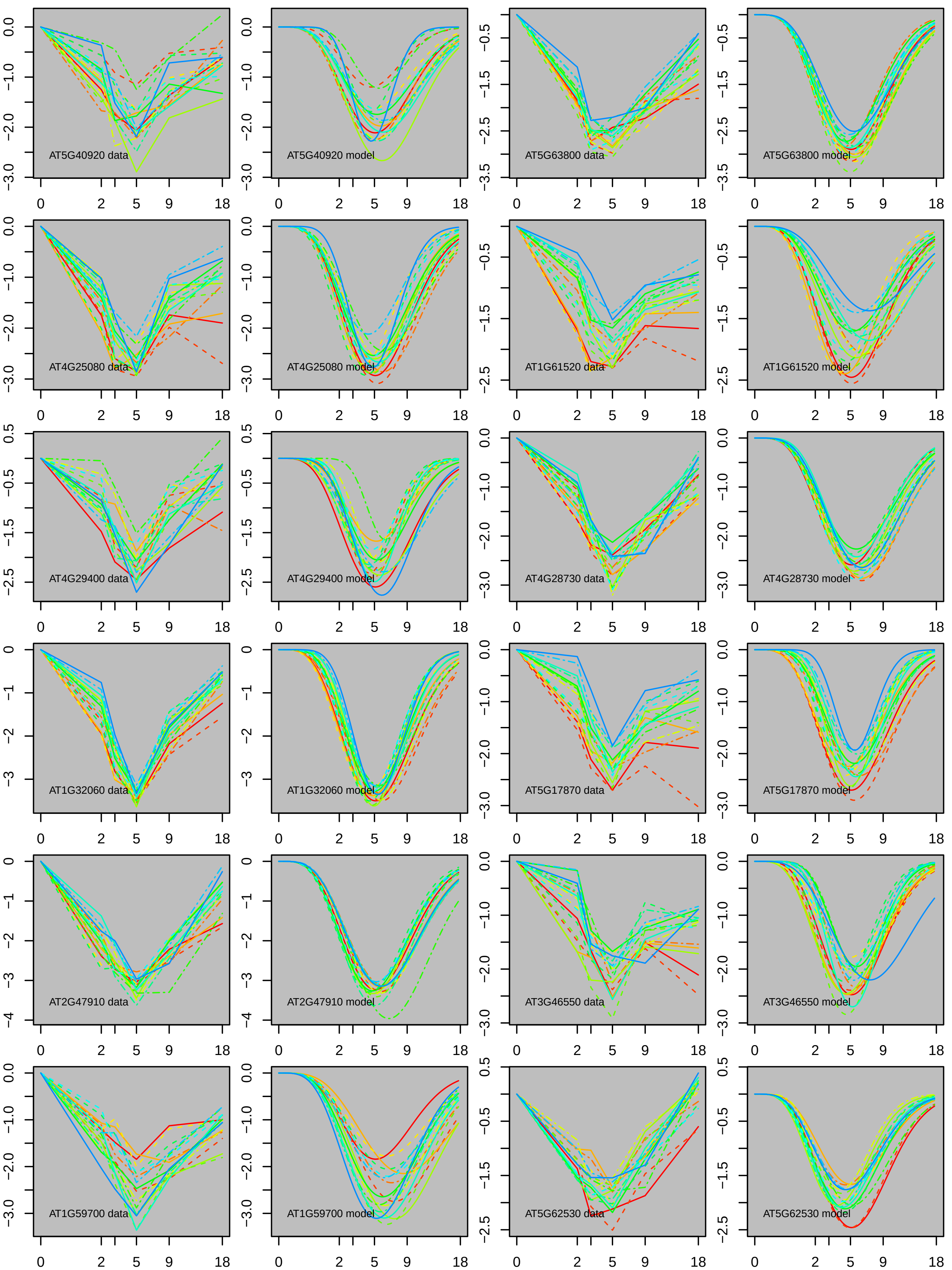

AT5G40920 data
AT5G40920 model
AT5G63800 data
AT5G63800 model
AT4G25080 data
AT4G25080 model
AT1G61520 data
AT1G61520 model
AT4G29400 data
AT4G29400 model
AT4G28730 data
AT4G28730 model
AT1G32060 data
AT1G32060 model
AT5G17870 data
AT5G17870 model
AT2G47910 data
AT2G47910 model
AT3G46550 data
AT3G46550 model
AT1G59700 data
AT1G59700 model
AT5G62530 data
AT5G62530 model

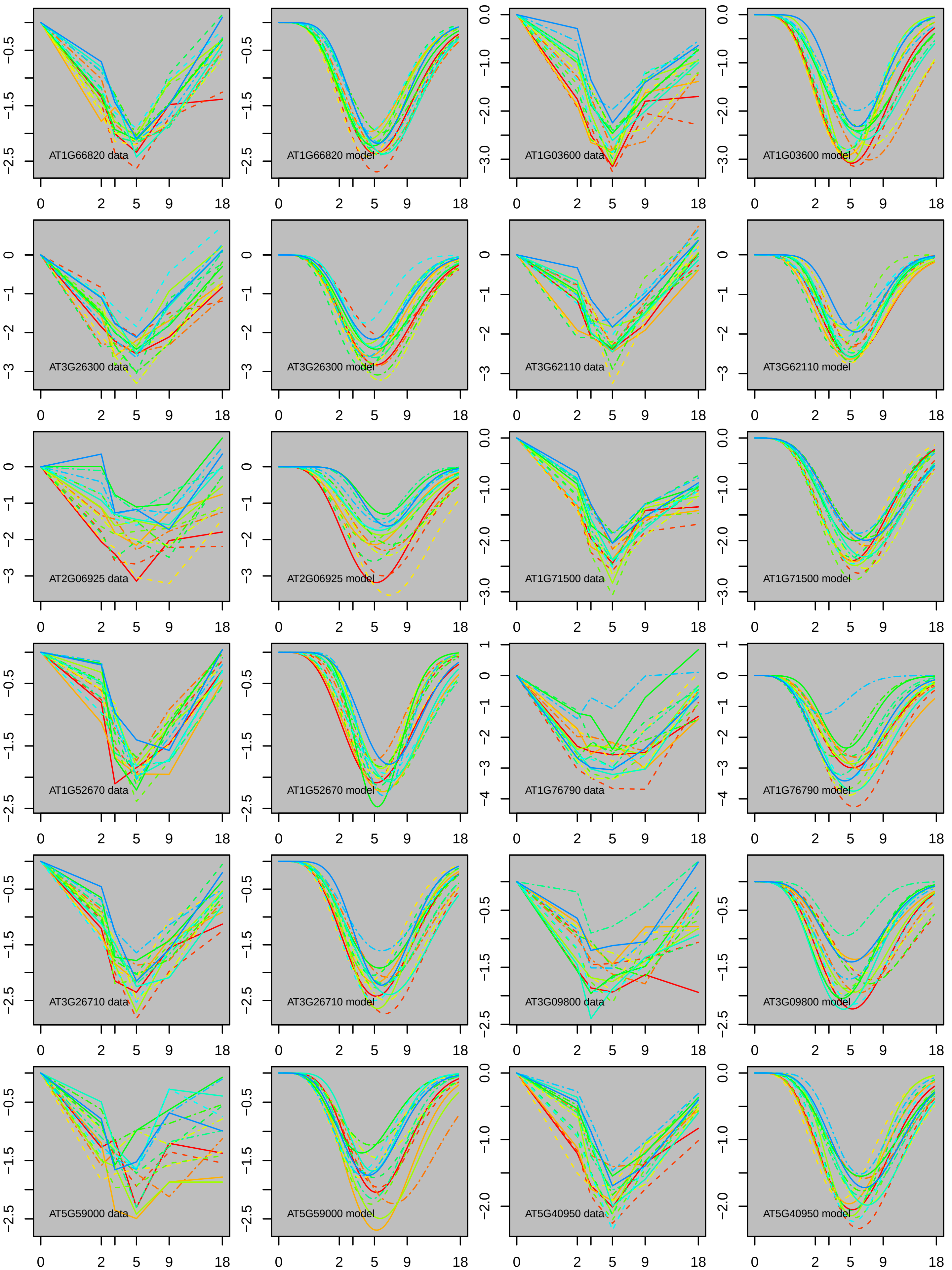
AT1G66820 data
AT1G66820 model
AT1G03600 data
AT1G03600 model
AT3G26300 data
AT3G26300 model
AT3G62110 data
AT3G62110 model
AT2G06925 data
AT2G06925 model
AT1G71500 data
AT1G71500 model
AT1G52670 data
AT1G52670 model
AT1G76790 data
AT1G76790 model
AT3G26710 data
AT3G26710 model
AT3G09800 data
AT3G09800 model
AT5G59000 data
AT5G59000 model
AT5G40950 data
AT5G40950 model

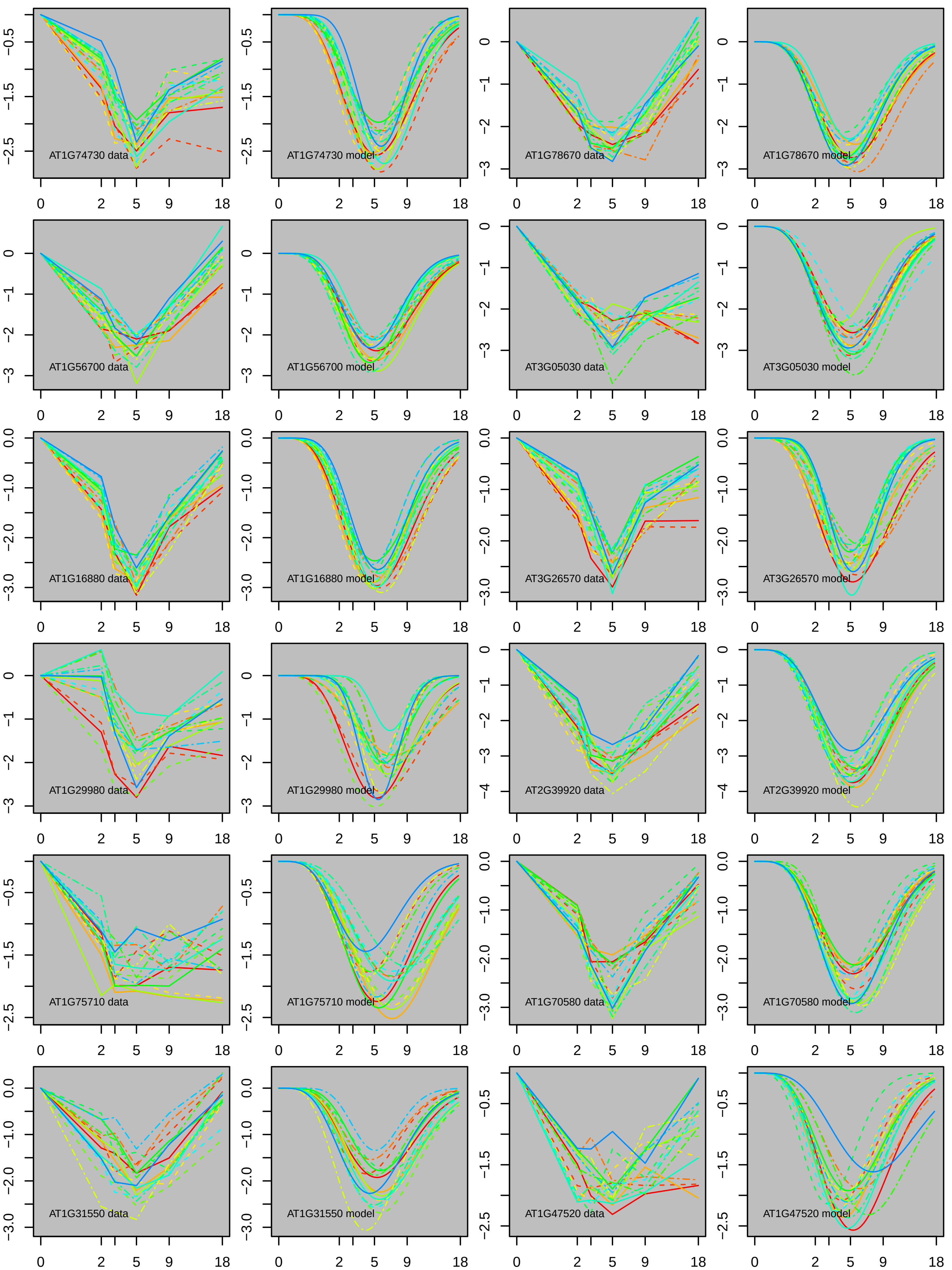
AT1G74730 data
AT1G74730 model
AT1G78670 data
AT1G78670 model
AT1G56700 data
AT1G56700 model
AT3G05030 data
AT3G05030 model
AT1G16880 data
AT1G16880 model
AT3G26570 data
AT3G26570 model
AT1G29980 data
AT1G29980 model
AT2G39920 data
AT2G39920 model
AT1G75710 data
AT1G75710 model
AT1G70580 data
AT1G70580 model
AT1G31550 data
AT1G31550 model
AT1G47520 data
AT1G47520 model

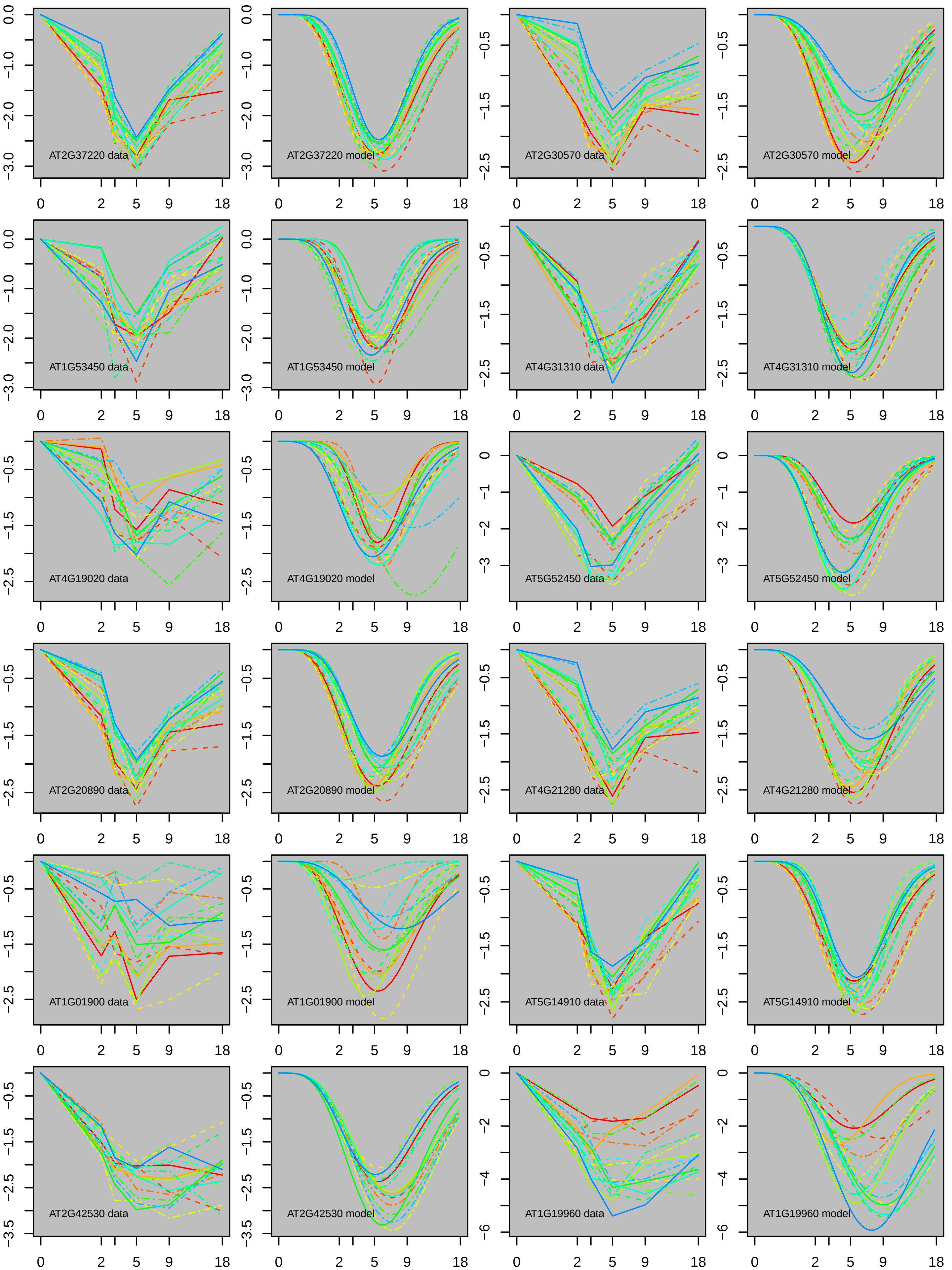
AT2G37220 data
AT2G37220 model
AT2G30570 data
AT2G30570 model
AT1G53450 data
AT1G53450 model
AT4G31310 data
AT4G31310 model
AT4G19020 data
AT4G19020 model
AT5G52450 data
AT5G52450 model
AT2G20890 data
AT2G20890 model
AT4G21280 data
AT4G21280 model
AT1G01900 data
AT1G01900 model
AT5G14910 data
AT5G14910 model
AT2G42530 data
AT2G42530 model
AT1G19960 data
AT1G19960 model

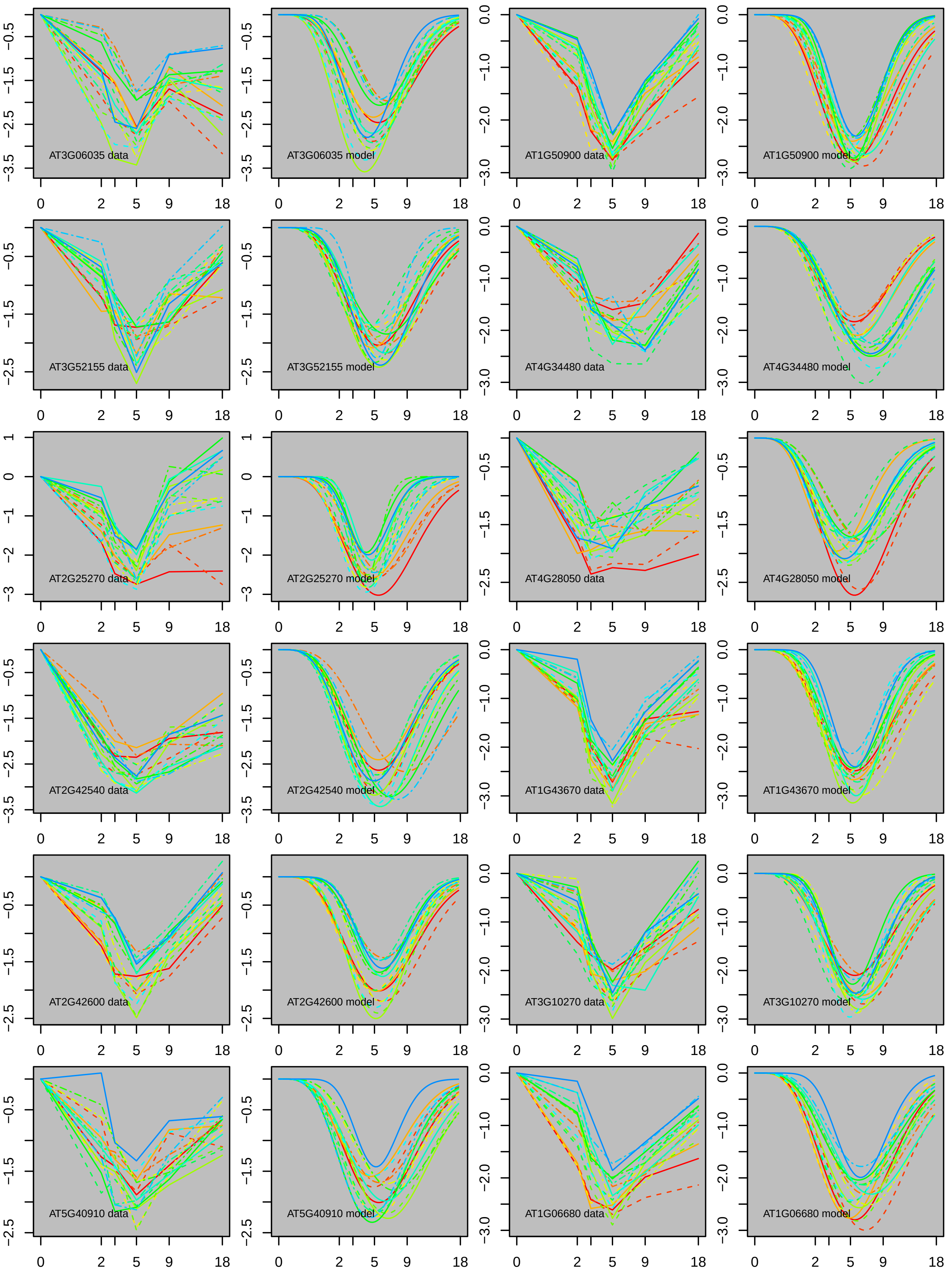
AT3G06035 data
AT3G06035 model
AT1G50900 data
AT1G50900 model
AT3G52155 data
AT3G52155 model
AT4G34480 data
AT4G34480 model
AT2G25270 data
AT2G25270 model
AT4G28050 data
AT4G28050 model
AT2G42540 data
AT2G42540 model
AT1G43670 data
AT1G43670 model
AT2G42600 data
AT2G42600 model
AT3G10270 data
AT3G10270 model
AT5G40910 data
AT5G40910 model
AT1G06680 data
AT1G06680 model

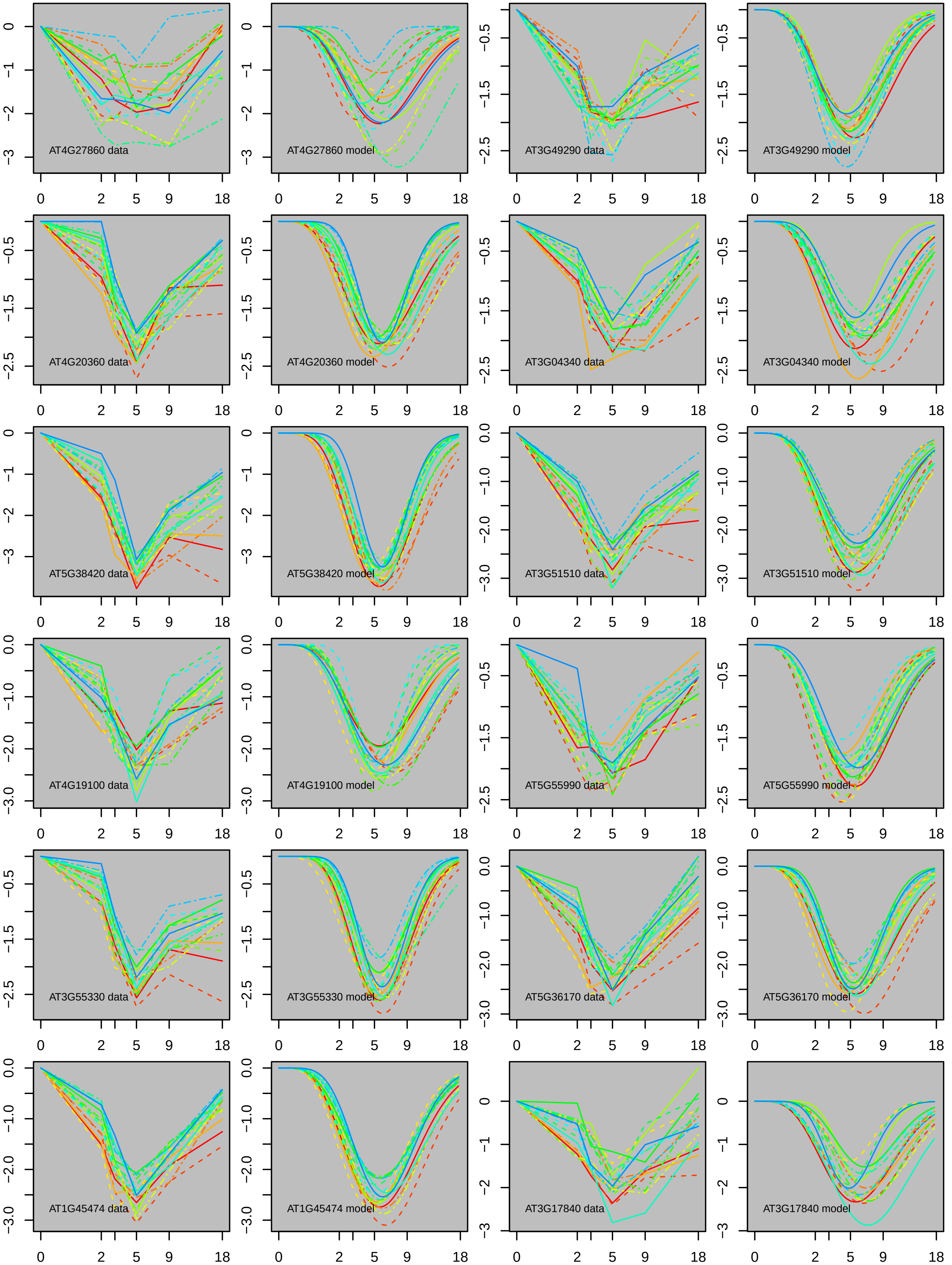
AT4G27860 data
AT4G27860 model
AT3G49290 data
AT3G49290 model
AT4G20360 data
AT4G20360 model
AT3G04340 data
AT3G04340 model
AT5G38420 data
AT5G38420 model
AT3G51510 data
AT3G51510 model
AT4G19100 data
AT4G19100 model
AT5G55990 data
AT5G55990 model
AT3G55330 data
AT3G55330 model
AT5G36170 data
AT5G36170 model
AT1G45474 data
AT1G45474 model
AT3G17840 data
AT3G17840 model

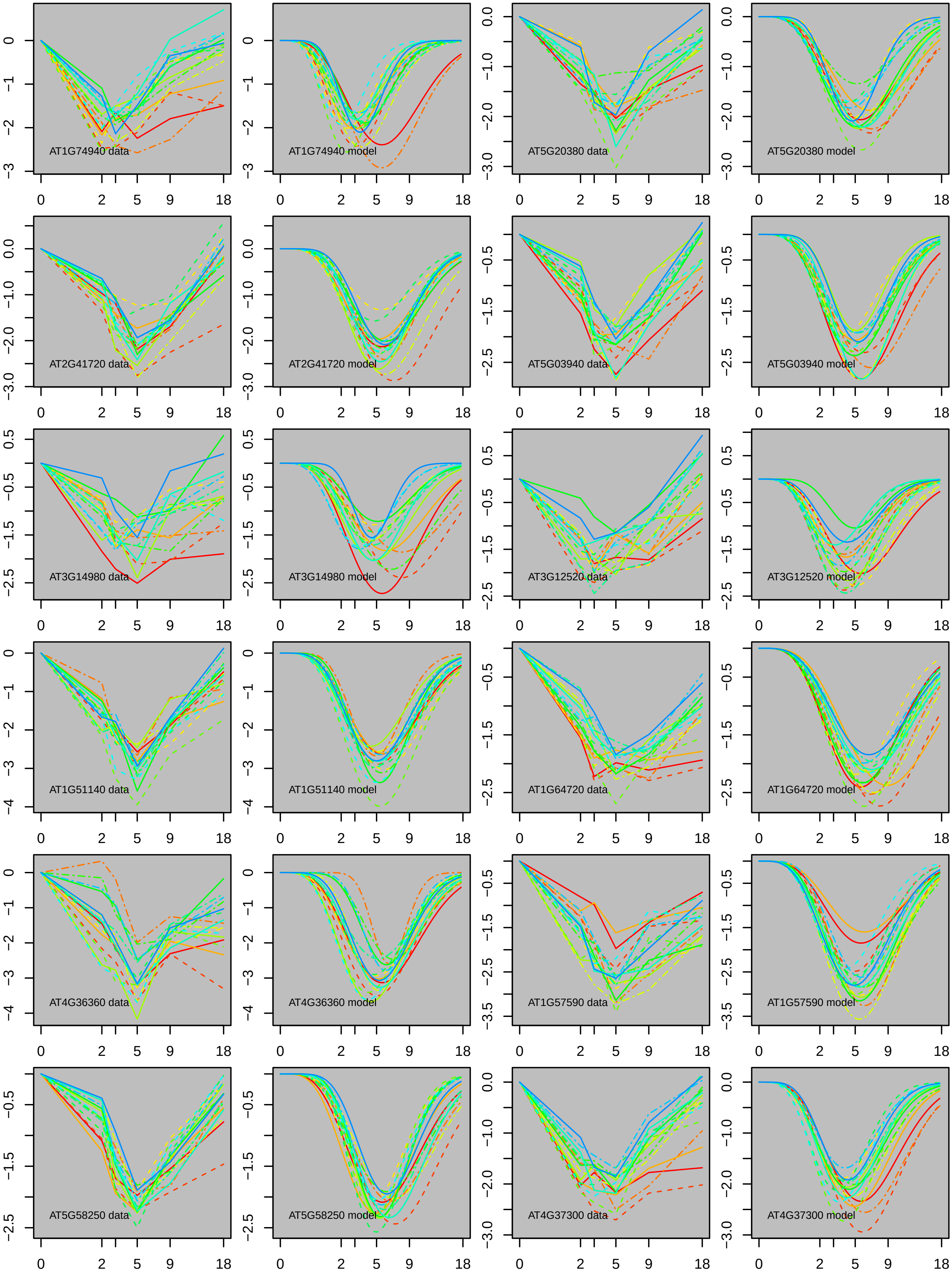
AT1G74940 data
AT1G74940 model
AT5G20380 data
AT5G20380 model
AT2G41720 data
AT2G41720 model
AT5G03940 data
AT5G03940 model
AT3G14980 data
AT3G14980 model
AT3G12520 data
AT3G12520 model
AT1G51140 data
AT1G51140 model
AT1G64720 data
AT1G64720 model
AT4G36360 data
AT4G36360 model
AT1G57590 data
AT1G57590 model
AT5G58250 data
AT5G58250 model
AT4G37300 data
AT4G37300 model
0 2 5 9 18

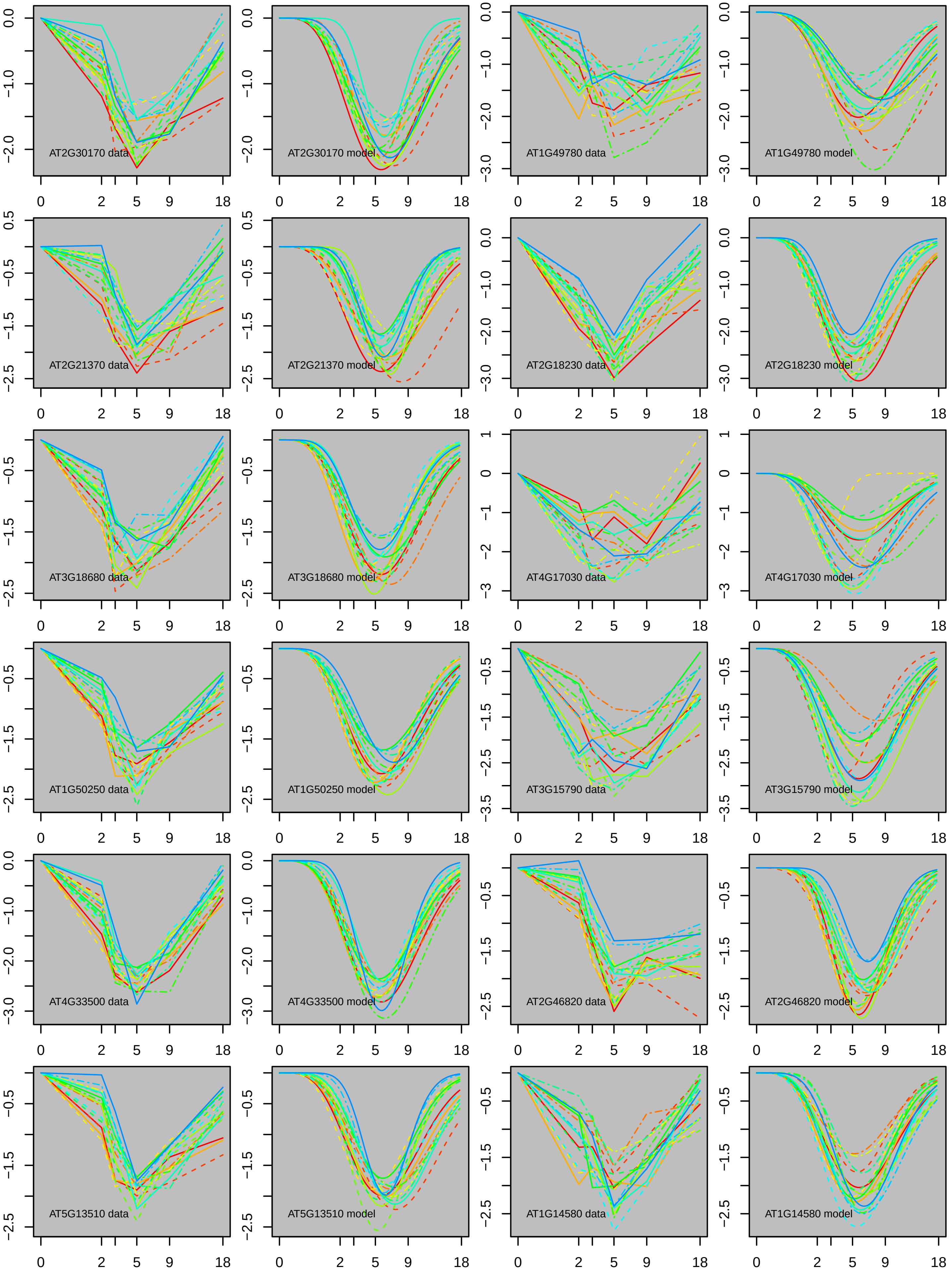

AT2G30170 data
AT2G30170 model
AT1G49780 data
AT1G49780 model
AT2G21370 data
AT2G21370 model
AT2G18230 data
AT2G18230 model
AT3G18680 data
AT3G18680 model
AT4G17030 data
AT4G17030 model
AT1G50250 data
AT1G50250 model
AT3G15790 data
AT3G15790 model
AT4G33500 data
AT4G33500 model
AT2G46820 data
AT2G46820 model
AT5G13510 data
AT5G13510 model
AT1G14580 data
AT1G14580 model

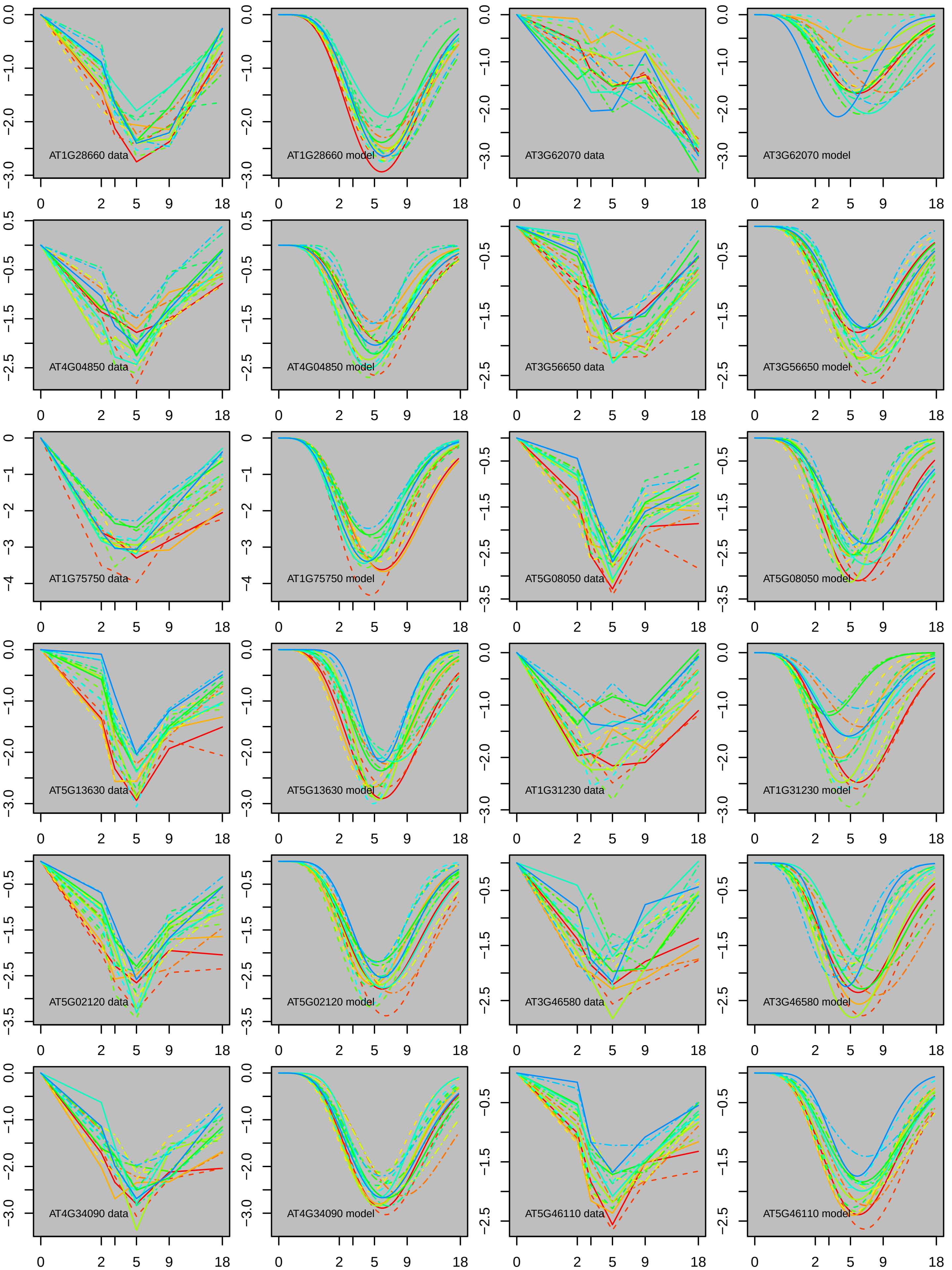
AT1G28660 data
AT1G28660 model
AT3G62070 data
AT3G62070 model
AT4G04850 data
AT4G04850 model
AT3G56650 data
AT3G56650 model
AT1G75750 data
AT1G75750 model
AT5G08050 data
AT5G08050 model
AT5G13630 data
AT5G13630 model
AT1G31230 data
AT1G31230 model
AT5G02120 data
AT5G02120 model
AT3G46580 data
AT3G46580 model
AT4G34090 data
AT4G34090 model
AT5G46110 data
AT5G46110 model
0 2 5 9 18

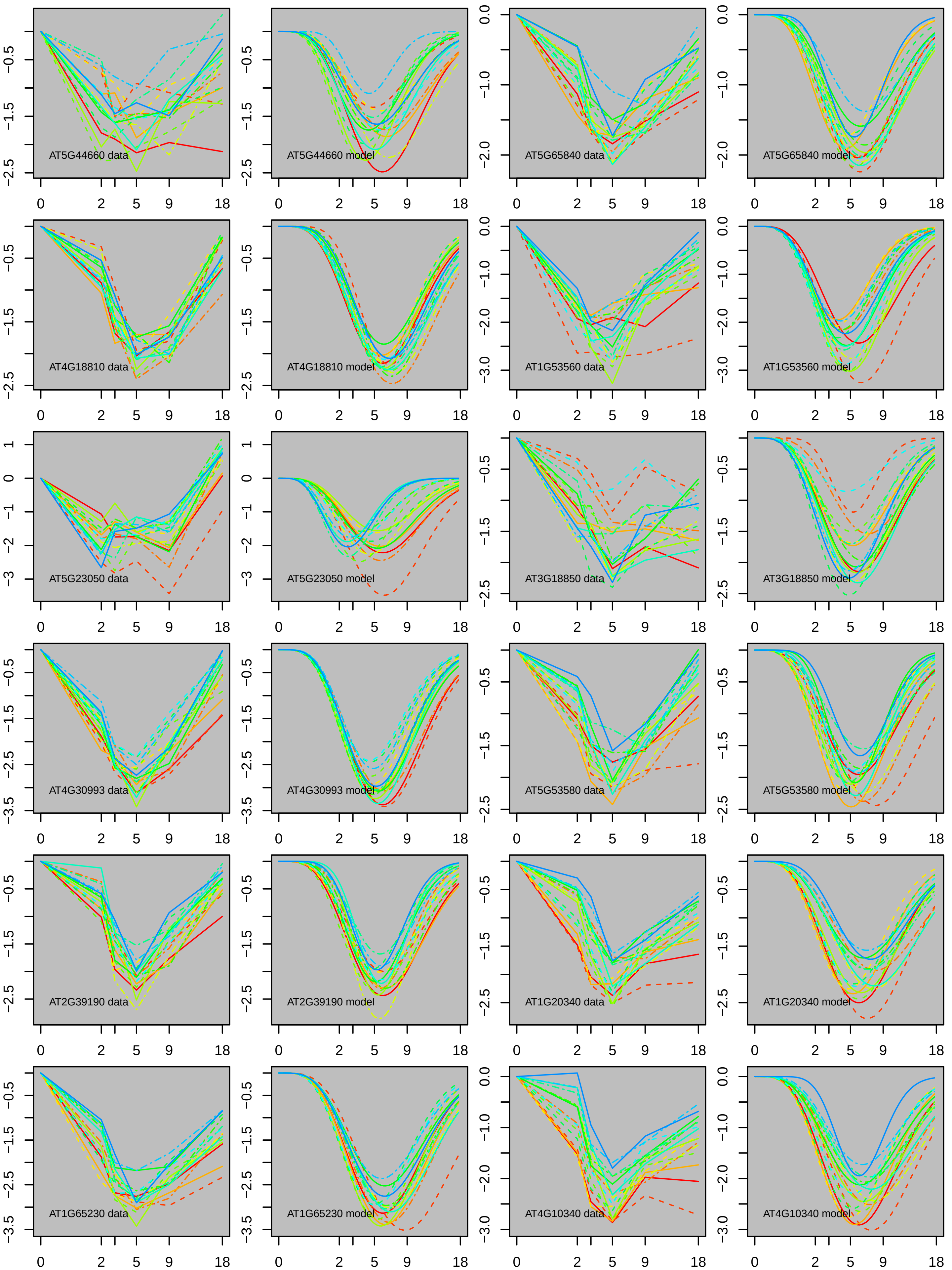

AT5G44660 data
AT5G44660 model
AT5G65840 data
AT5G65840 model
AT4G18810 data
AT4G18810 model
AT1G53560 data
AT1G53560 model
AT5G23050 data
AT5G23050 model
AT3G18850 data
AT3G18850 model
AT4G30993 data
AT4G30993 model
AT5G53580 data
AT5G53580 model
AT2G39190 data
AT2G39190 model
AT1G20340 data
AT1G20340 model
AT1G65230 data
AT1G65230 model
AT4G10340 data
AT4G10340 model

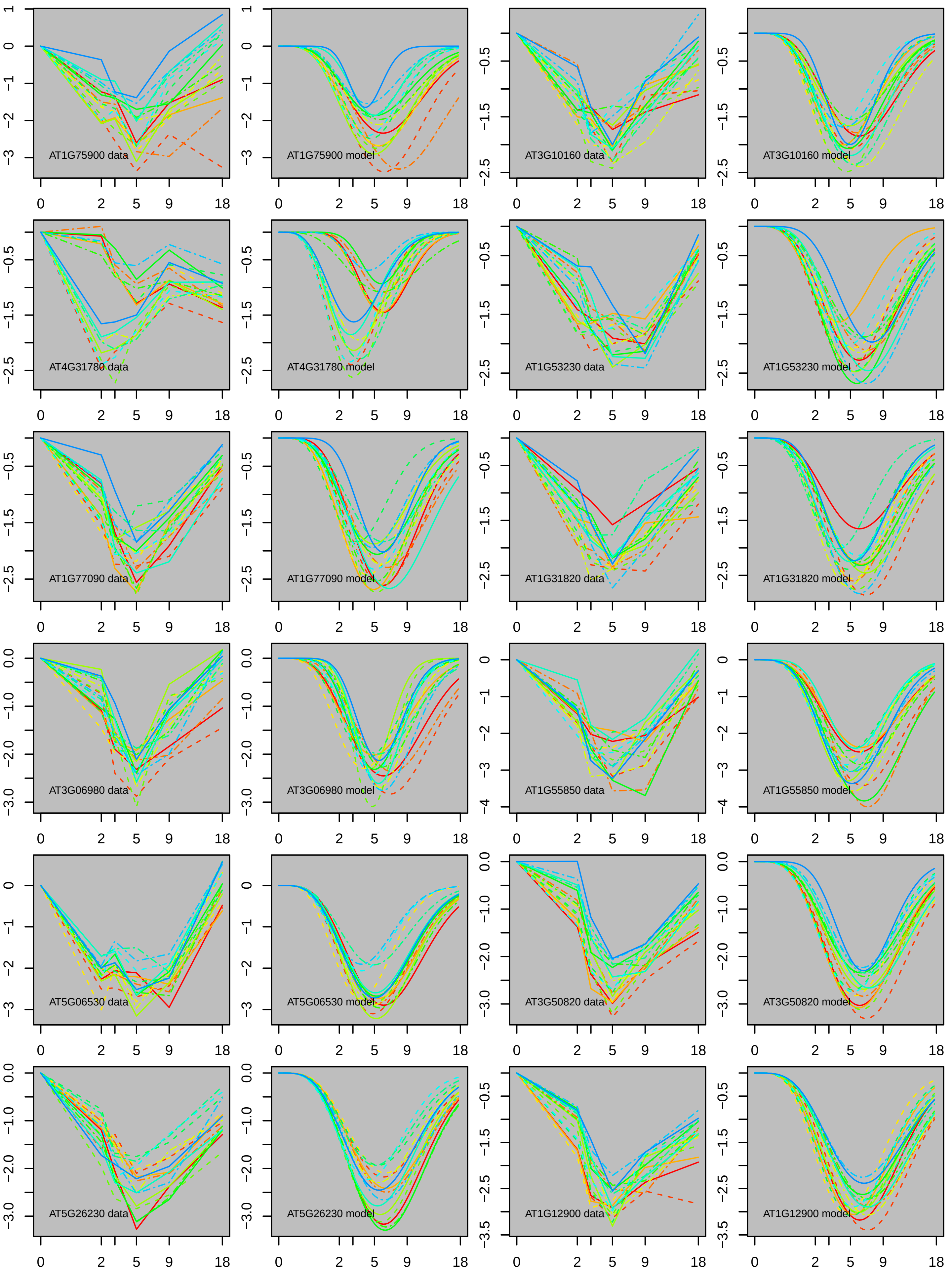
AT1G75900 data
AT1G75900 model
AT3G10160 data
AT3G10160 model
AT4G31780 data
AT4G31780 model
AT1G53230 data
AT1G53230 model
AT1G77090 data
AT1G77090 model
AT1G31820 data
AT1G31820 model
AT3G06980 data
AT3G06980 model
AT1G55850 data
AT1G55850 model
AT5G06530 data
AT5G06530 model
AT3G50820 data
AT3G50820 model
AT5G26230 data
AT5G26230 model
AT1G12900 data
AT1G12900 model

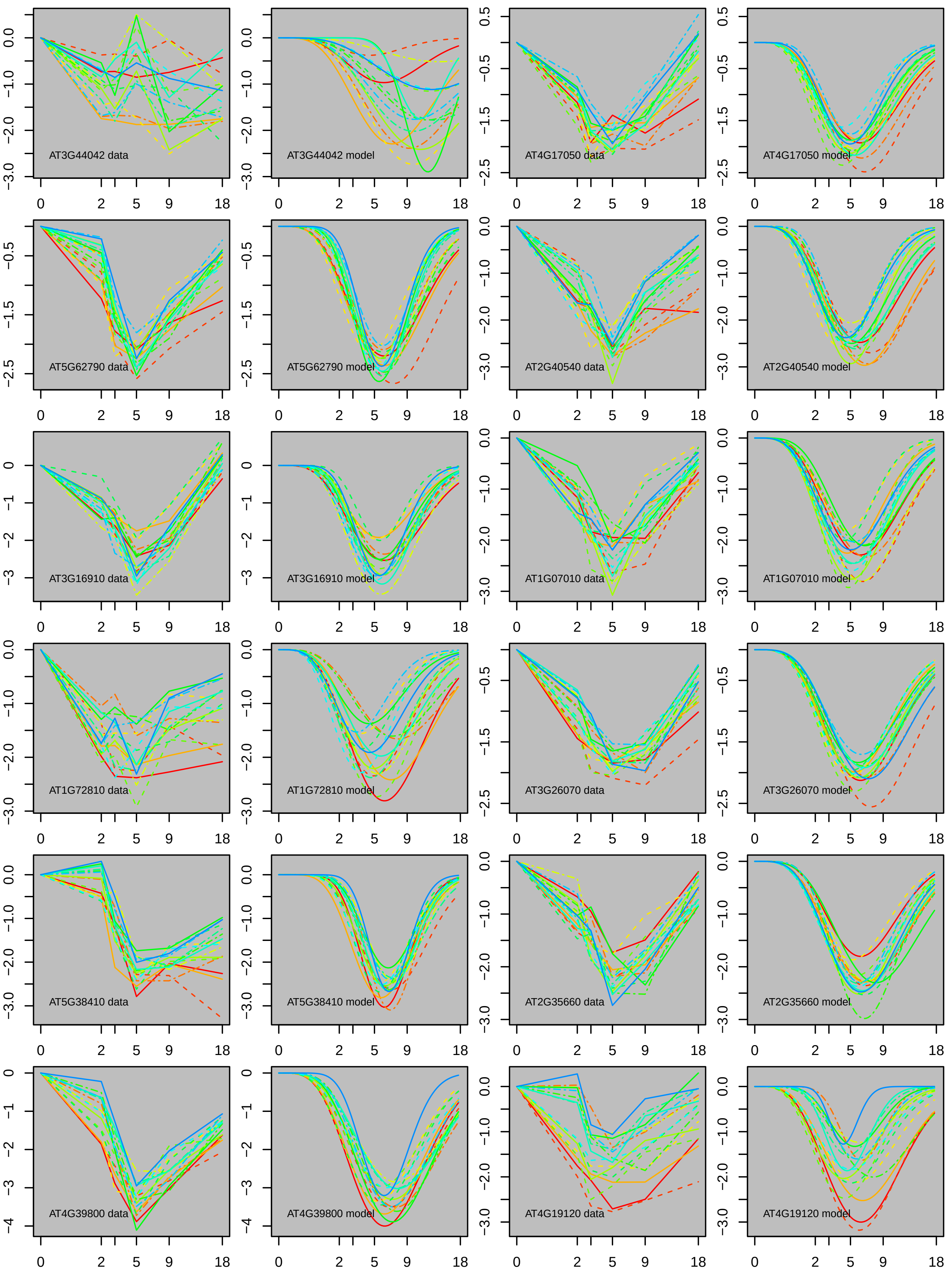
AT3G44042 data
AT3G44042 model
AT4G17050 data
AT4G17050 model
AT5G62790 data
AT5G62790 model
AT2G40540 data
AT2G40540 model
AT3G16910 data
AT3G16910 model
AT1G07010 data
AT1G07010 model
AT1G72810 data
AT1G72810 model
AT3G26070 data
AT3G26070 model
AT5G38410 data
AT5G38410 model
AT2G35660 data
AT2G35660 model
AT4G39800 data
AT4G39800 model
AT4G19120 data
AT4G19120 model

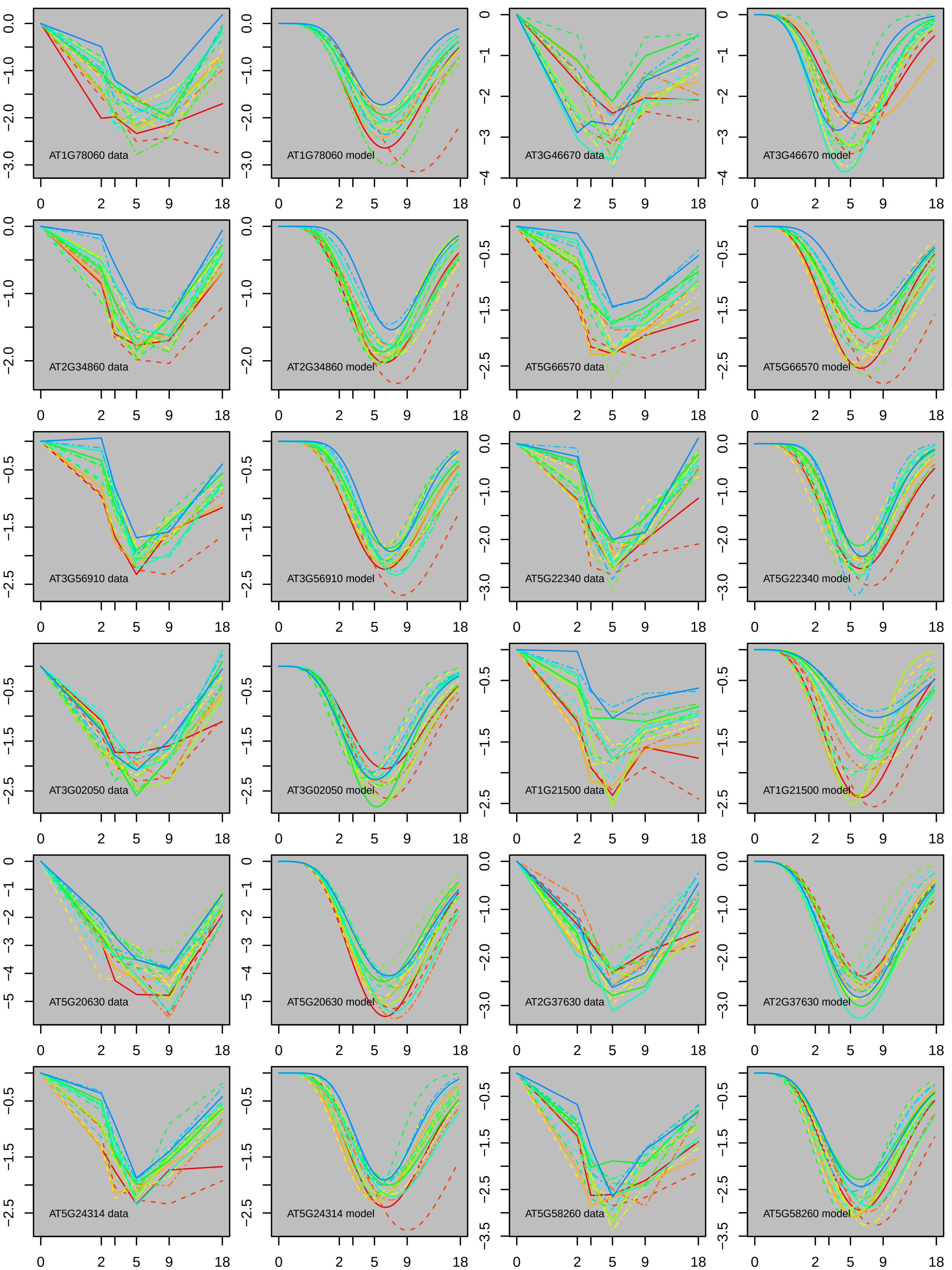
AT1G78060 data
AT1G78060 model
AT3G46670 data
AT3G46670 model
AT2G34860 data
AT2G34860 model
AT5G66570 data
AT5G66570 model
AT3G56910 data
AT3G56910 model
AT5G22340 data
AT5G22340 model
AT3G02050 data
AT3G02050 model
AT1G21500 data
AT1G21500 model
AT5G20630 data
AT5G20630 model
AT2G37630 data
AT2G37630 model
AT5G24314 data
AT5G24314 model
AT5G58260 data
AT5G58260 model
0
2
5
9
18

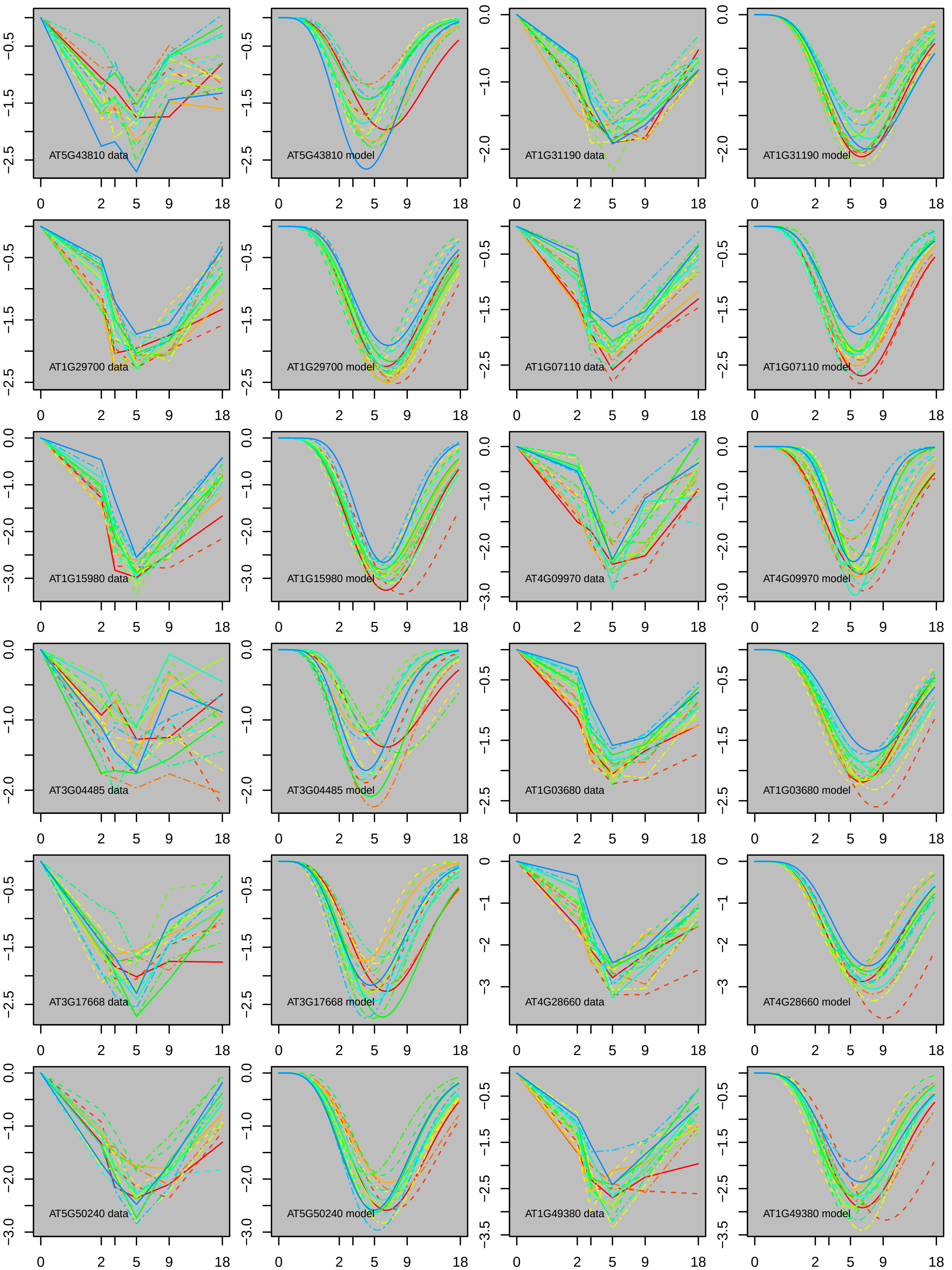
AT5G43810 data
AT5G43810 model
AT1G31190 data
AT1G31190 model
AT1G29700 data
AT1G29700 model
AT1G07110 data
AT1G07110 model
AT1G15980 data
AT1G15980 model
AT4G09970 data
AT4G09970 model
AT3G04485 data
AT3G04485 model
AT1G03680 data
AT1G03680 model
AT3G17668 data
AT3G17668 model
AT4G28660 data
AT4G28660 model
AT5G50240 data
AT5G50240 model
AT1G49380 data
AT1G49380 model

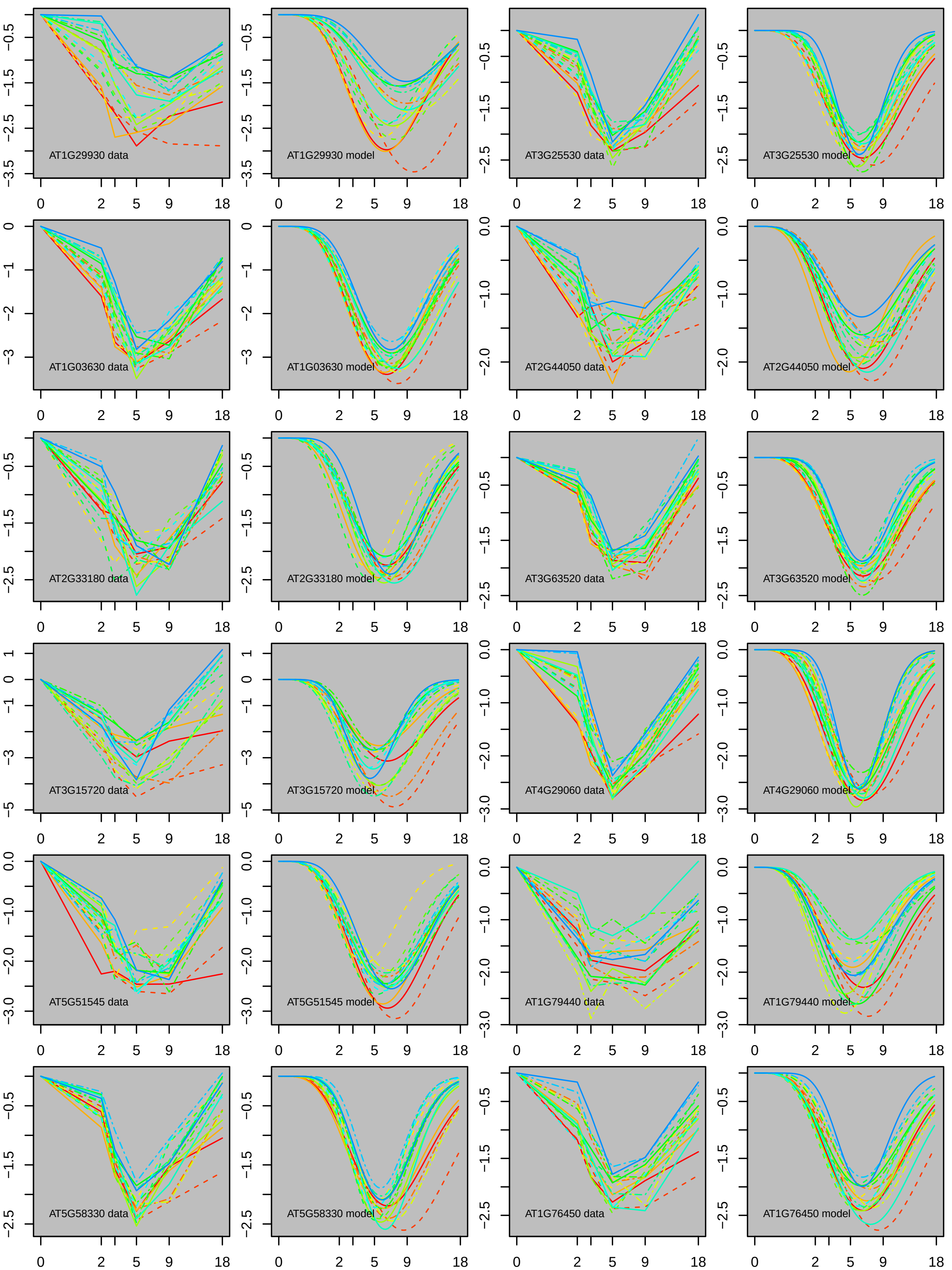
AT1G29930 data
AT1G29930 model
AT3G25530 data
AT3G25530 model
AT1G03630 data
AT1G03630 model
AT2G44050 data
AT2G44050 model
AT2G33180 data
AT2G33180 model
AT3G63520 data
AT3G63520 model
AT3G15720 data
AT3G15720 model
AT4G29060 data
AT4G29060 model
AT5G51545 data
AT5G51545 model
AT1G79440 data
AT1G79440 model
AT5G58330 data
AT5G58330 model
AT1G76450 data
AT1G76450 model

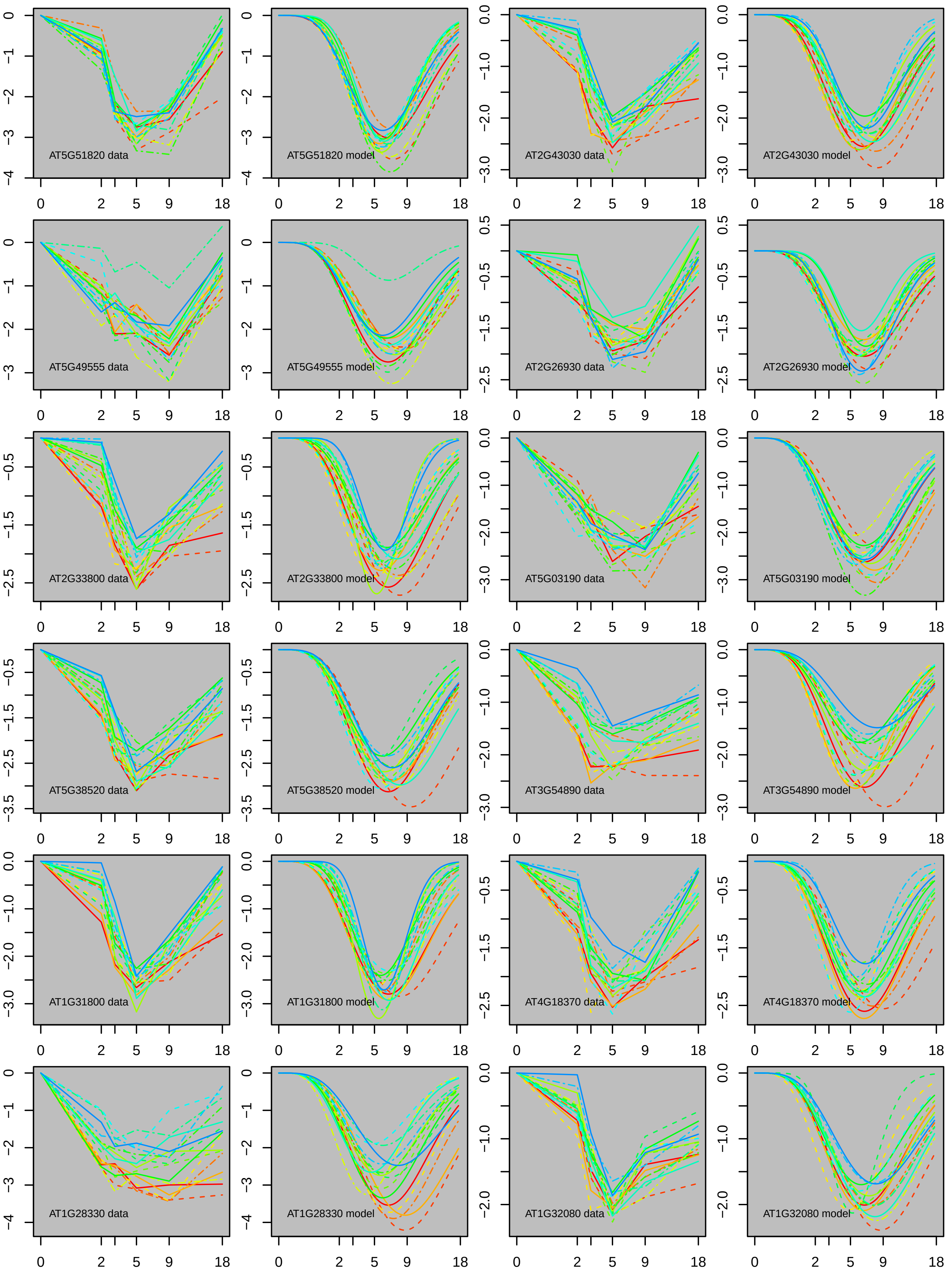

AT5G51820 data
AT5G51820 model
AT2G43030 data
AT2G43030 model
AT5G49555 data
AT5G49555 model
AT2G26930 data
AT2G26930 model
AT2G33800 data
AT2G33800 model
AT5G03190 data
AT5G03190 model
AT5G38520 data
AT5G38520 model
AT3G54890 data
AT3G54890 model
AT1G31800 data
AT1G31800 model
AT4G18370 data
AT4G18370 model
AT1G28330 data
AT1G28330 model
AT1G32080 data
AT1G32080 model

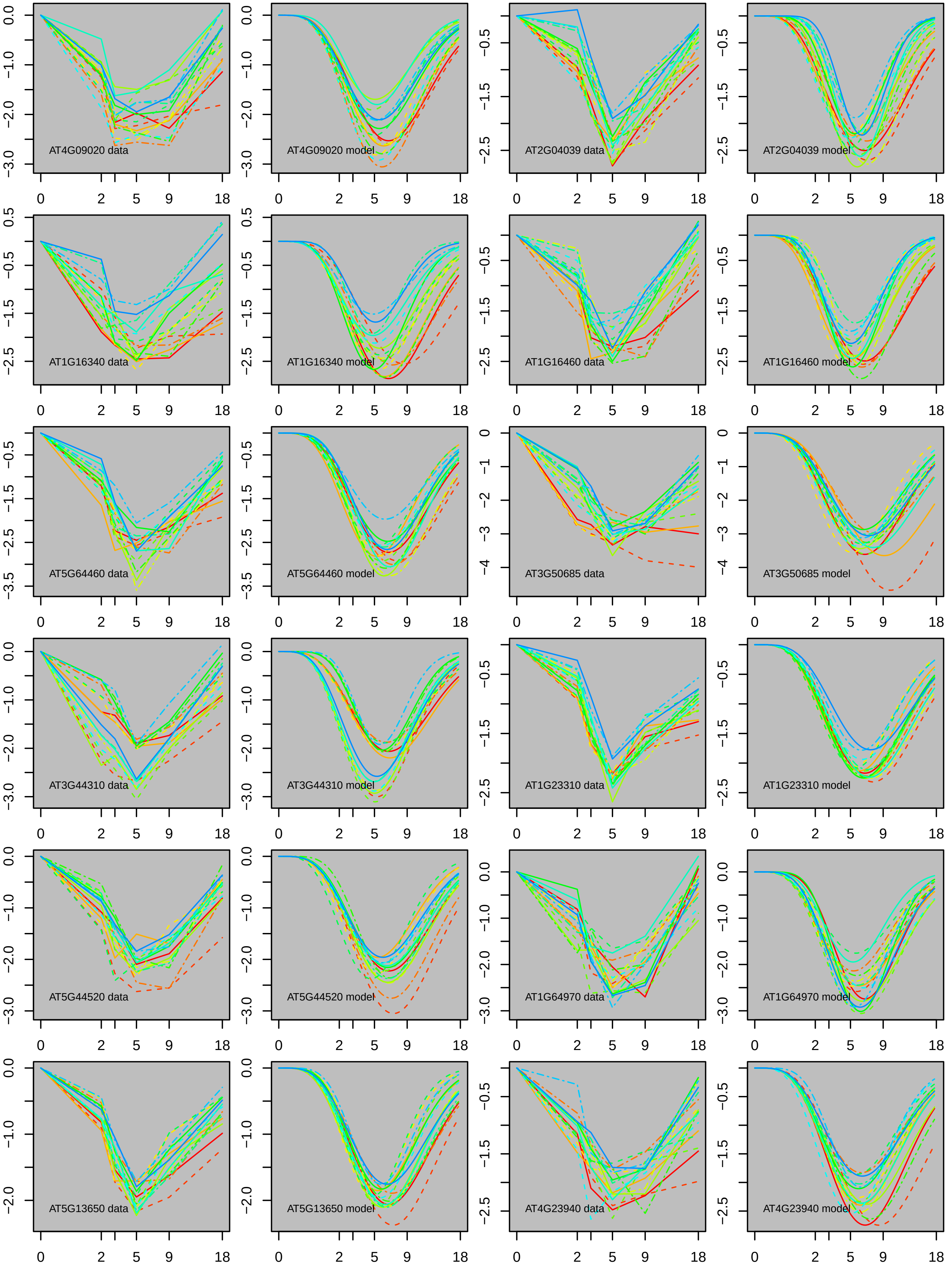
AT4G09020 data
AT4G09020 model
AT2G04039 data
AT2G04039 model
AT1G16340 data
AT1G16340 model
AT1G16460 data
AT1G16460 model
AT5G64460 data
AT5G64460 model
AT3G50685 data
AT3G50685 model
AT3G44310 data
AT3G44310 model
AT1G23310 data
AT1G23310 model
AT5G44520 data
AT5G44520 model
AT1G64970 data
AT1G64970 model
AT5G13650 data
AT5G13650 model
AT4G23940 data
AT4G23940 model

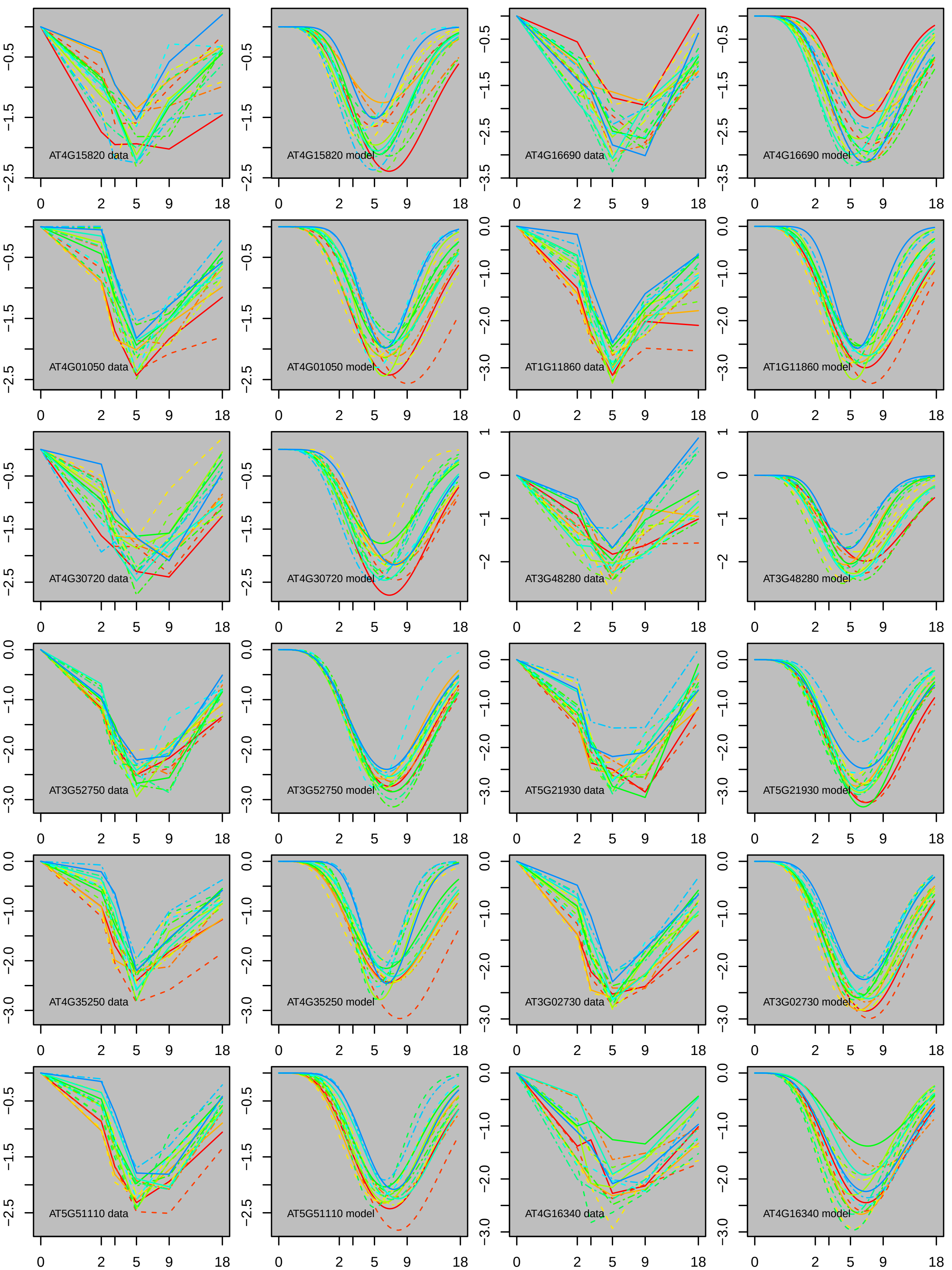
AT4G15820 data
AT4G15820 model
AT4G16690 data
AT4G16690 model
AT4G01050 data
AT4G01050 model
AT1G11860 data
AT1G11860 model
AT4G30720 data
AT4G30720 model
AT3G48280 data
AT3G48280 model
AT3G52750 data
AT3G52750 model
AT5G21930 data
AT5G21930 model
AT4G35250 data
AT4G35250 model
AT3G02730 data
AT3G02730 model
AT5G51110 data
AT5G51110 model
AT4G16340 data
AT4G16340 model
0
2
5
9
18

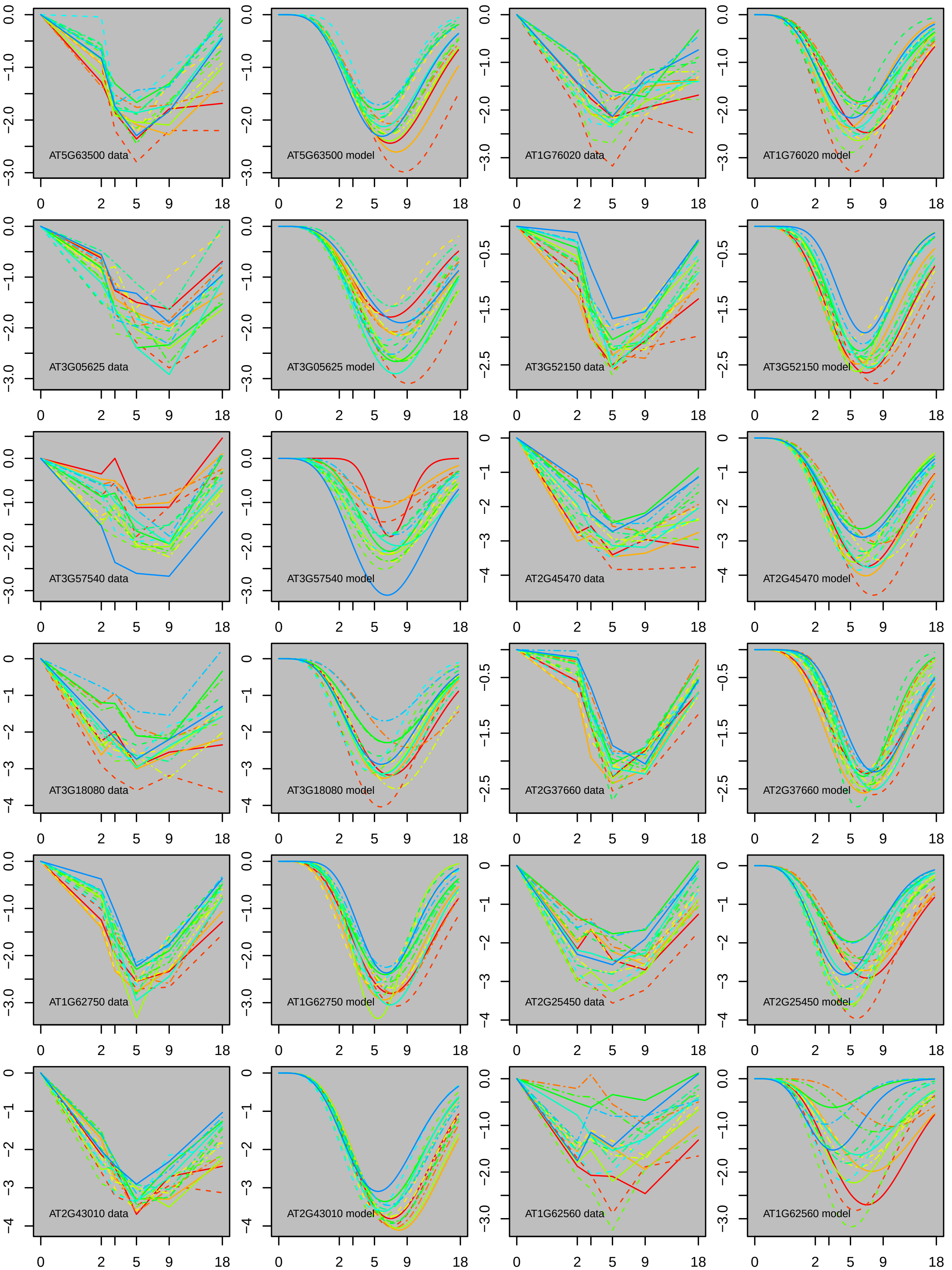
AT5G63500 data
AT5G63500 model
AT1G76020 data
AT1G76020 model
AT3G05625 data
AT3G05625 model
AT3G52150 data
AT3G52150 model
AT3G57540 data
AT3G57540 model
AT2G45470 data
AT2G45470 model
AT3G18080 data
AT3G18080 model
AT2G37660 data
AT2G37660 model
AT1G62750 data
AT1G62750 model
AT2G25450 data
AT2G25450 model
AT2G43010 data
AT2G43010 model
AT1G62560 data
AT1G62560 model

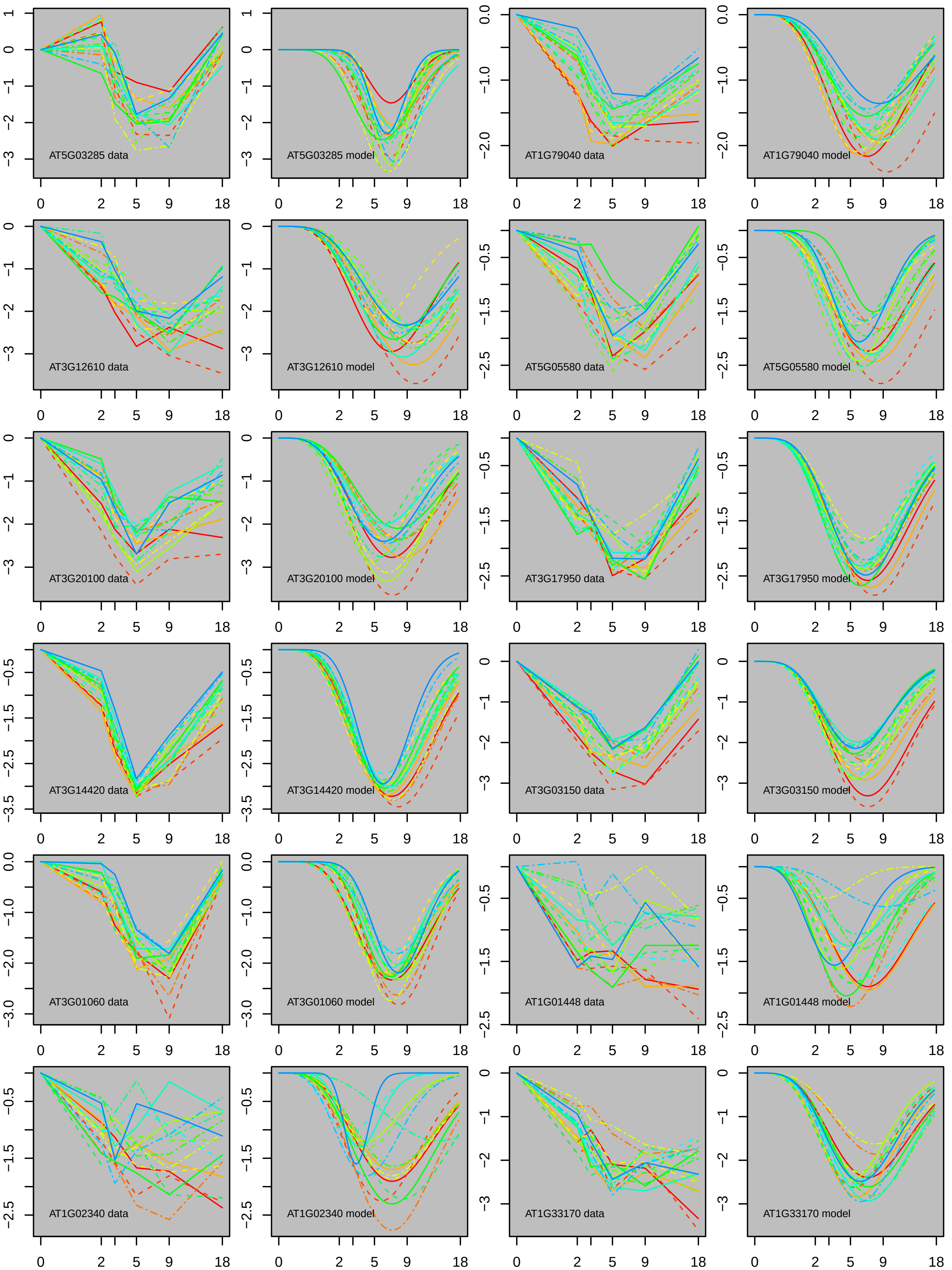
AT5G03285 data
AT5G03285 model
AT1G79040 data
AT1G79040 model
AT3G12610 data
AT3G12610 model
AT5G05580 data
AT5G05580 model
AT3G20100 data
AT3G20100 model
AT3G17950 data
AT3G17950 model
AT3G14420 data
AT3G14420 model
AT3G03150 data
AT3G03150 model
AT3G01060 data
AT3G01060 model
AT1G01448 data
AT1G01448 model
AT1G02340 data
AT1G02340 model
AT1G33170 data
AT1G33170 model
0
2
5
9
18

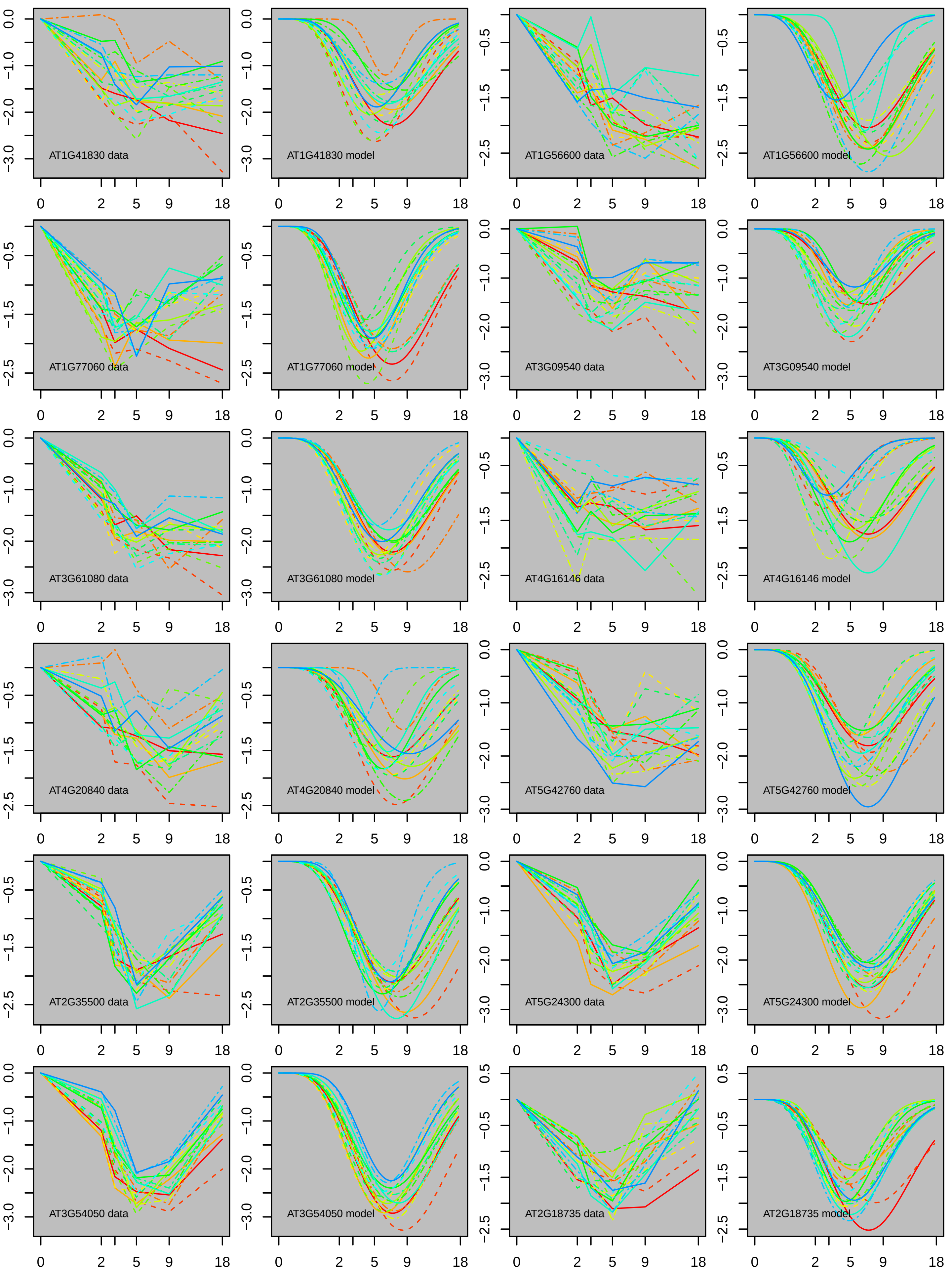
AT1G41830 data
AT1G41830 model
AT1G56600 data
AT1G56600 model
AT1G77060 data
AT1G77060 model
AT3G09540 data
AT3G09540 model
AT3G61080 data
AT3G61080 model
AT4G16146 data
AT4G16146 model
AT4G20840 data
AT4G20840 model
AT5G42760 data
AT5G42760 model
AT2G35500 data
AT2G35500 model
AT5G24300 data
AT5G24300 model
AT3G54050 data
AT3G54050 model
AT2G18735 data
AT2G18735 model

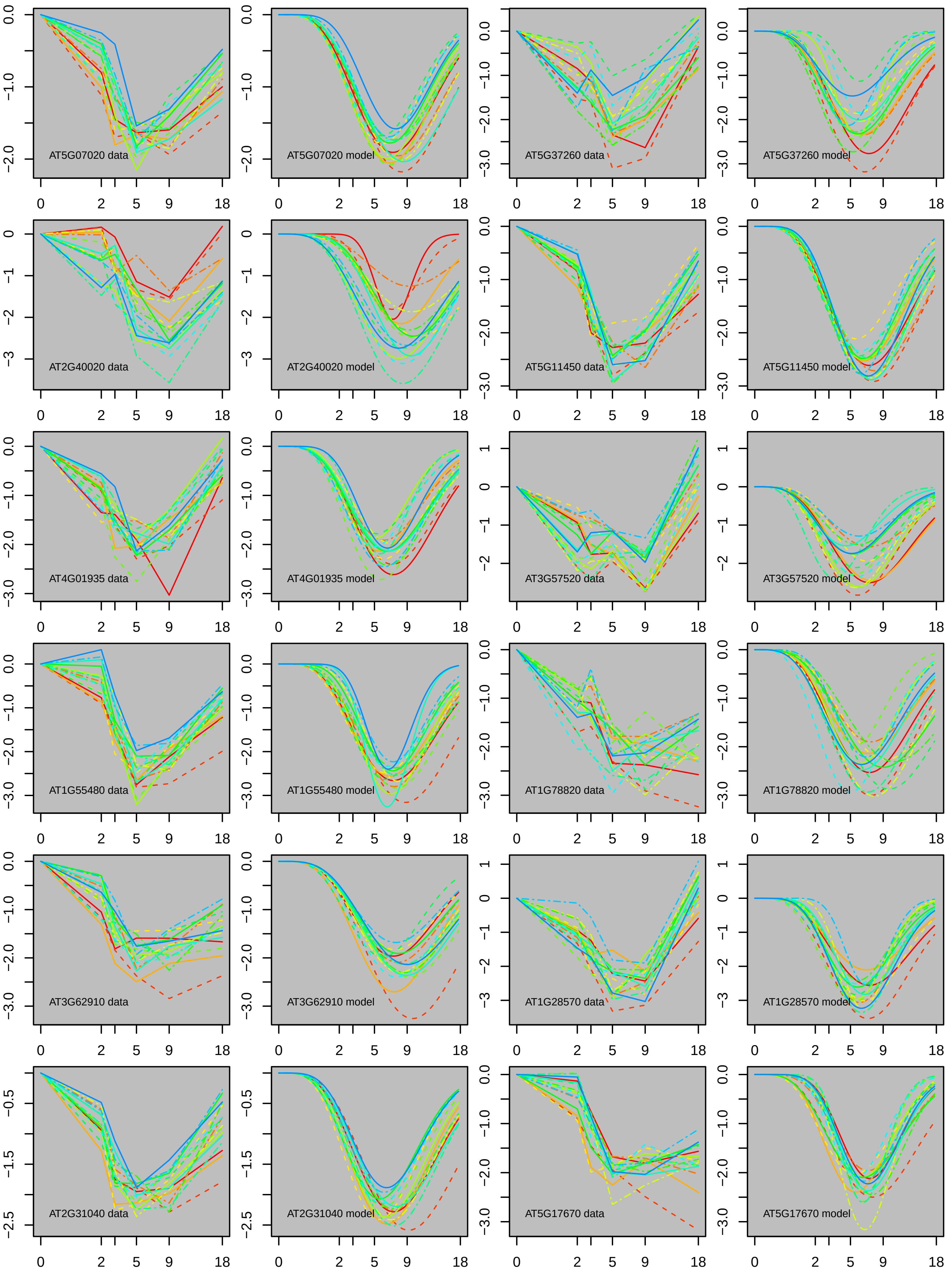
AT5G07020 data
AT5G07020 model
AT5G37260 data
AT5G37260 model
AT2G40020 data
AT2G40020 model
AT5G11450 data
AT5G11450 model
AT4G01935 data
AT4G01935 model
AT3G57520 data
AT3G57520 model
AT1G55480 data
AT1G55480 model
AT1G78820 data
AT1G78820 model
AT3G62910 data
AT3G62910 model
AT1G28570 data
AT1G28570 model
AT2G31040 data
AT2G31040 model
AT5G17670 data
AT5G17670 model

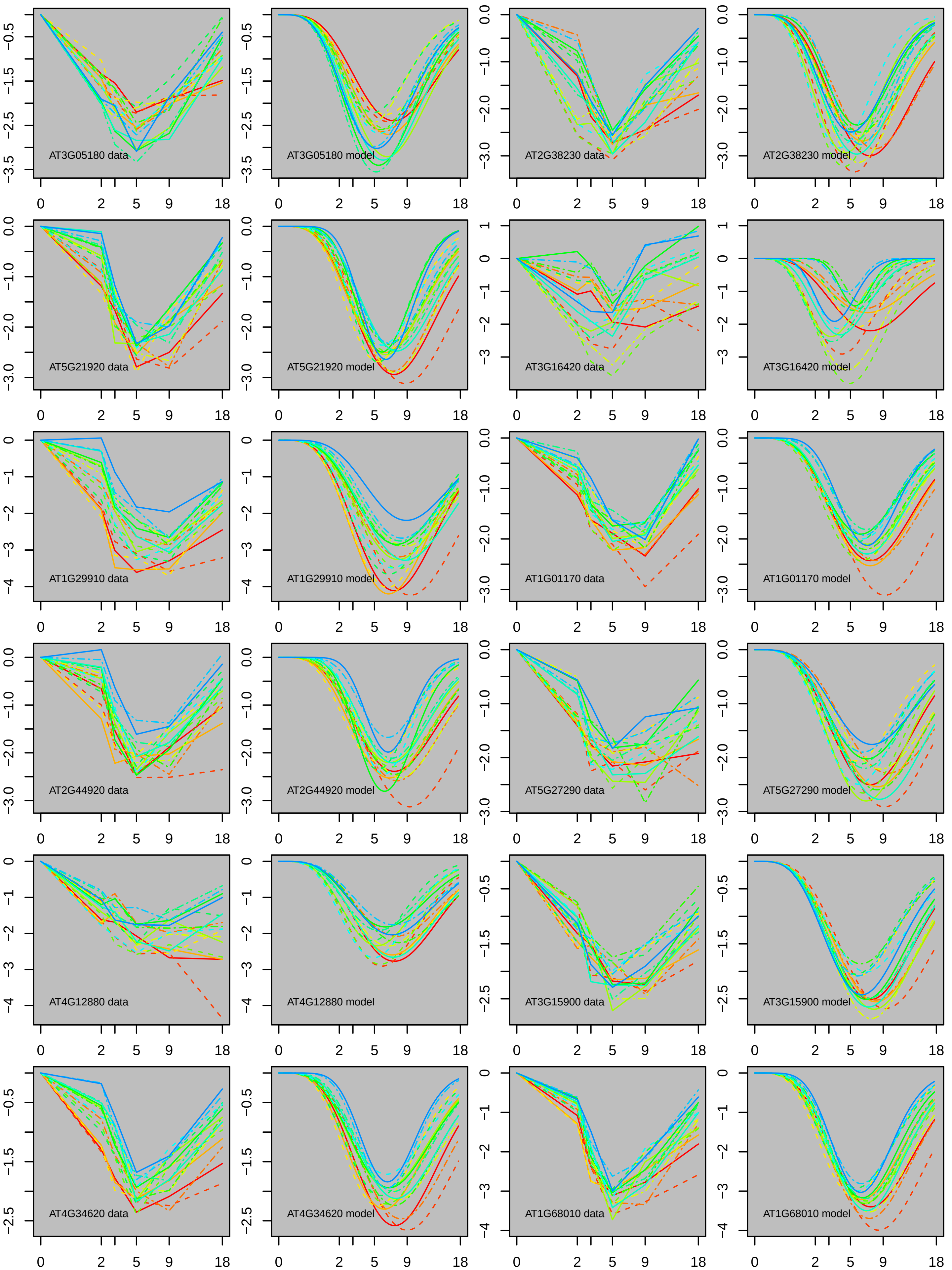
AT3G05180 data
AT3G05180 model
AT2G38230 data
AT2G38230 model
AT5G21920 data
AT5G21920 model
AT3G16420 data
AT3G16420 model
AT1G29910 data
AT1G29910 model
AT1G01170 data
AT1G01170 model
AT2G44920 data
AT2G44920 model
AT5G27290 data
AT5G27290 model
AT4G12880 data
AT4G12880 model
AT3G15900 data
AT3G15900 model
AT4G34620 data
AT4G34620 model
AT1G68010 data
AT1G68010 model
0
2
5
9
18

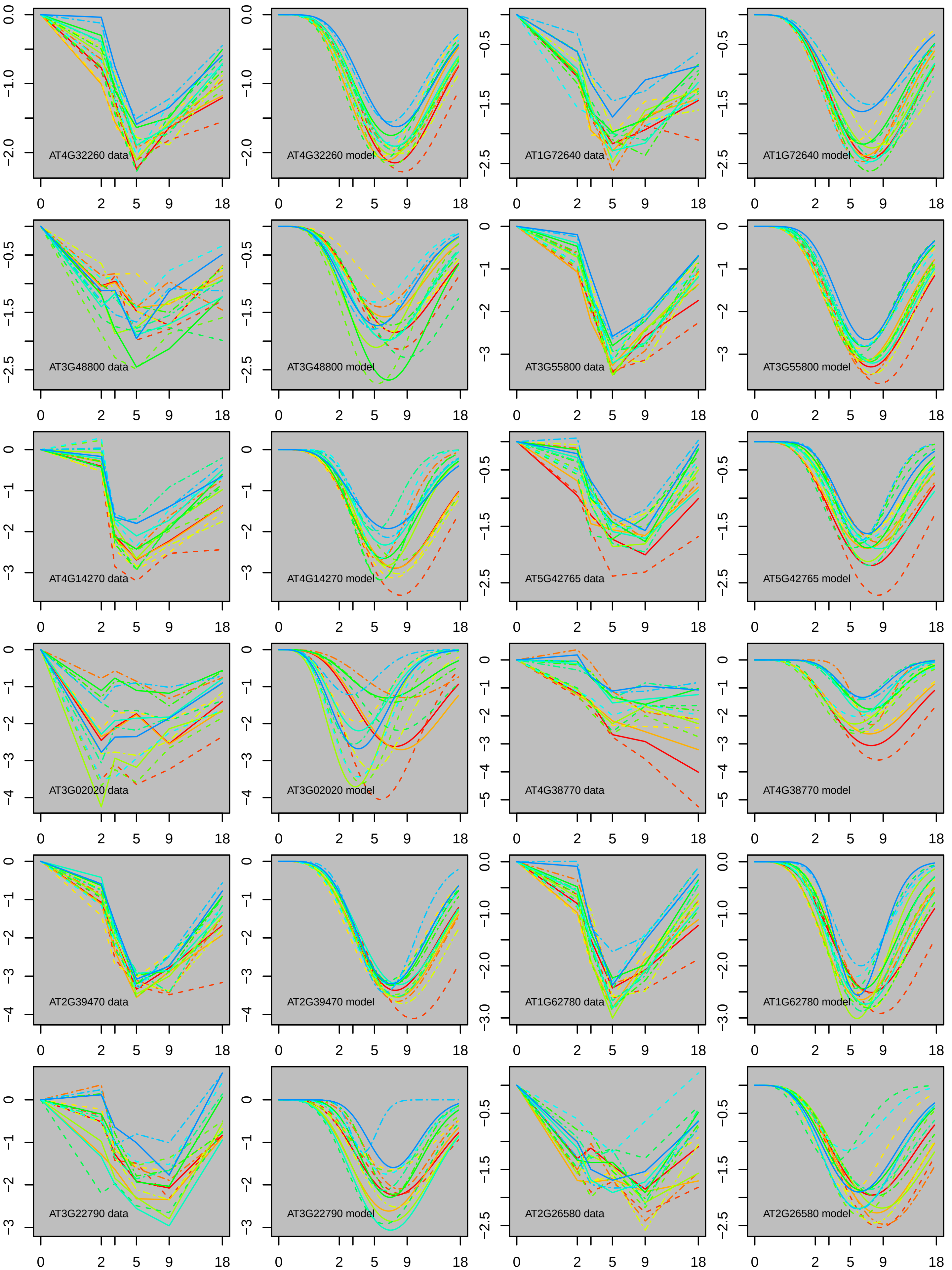

AT4G32260 data
AT4G32260 model
AT1G72640 data
AT1G72640 model
AT3G48800 data
AT3G48800 model
AT3G55800 data
AT3G55800 model
AT4G14270 data
AT4G14270 model
AT5G42765 data
AT5G42765 model
AT3G02020 data
AT3G02020 model
AT4G38770 data
AT4G38770 model
AT2G39470 data
AT2G39470 model
AT1G62780 data
AT1G62780 model
AT3G22790 data
AT3G22790 model
AT2G26580 data
AT2G26580 model
0 2 5 9 18

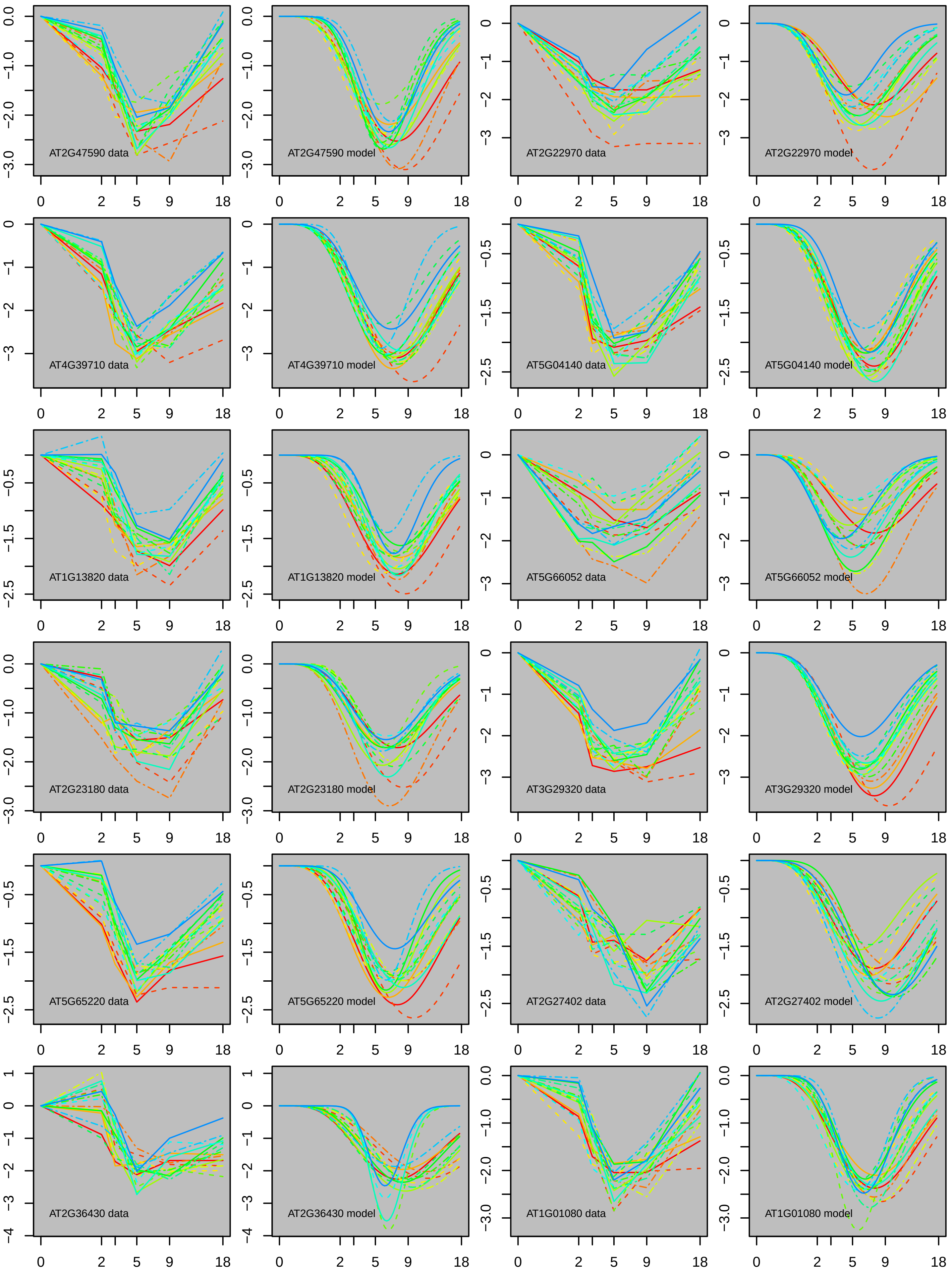
AT2G47590 data
AT2G47590 model
AT2G22970 data
AT2G22970 model
AT4G39710 data
AT4G39710 model
AT5G04140 data
AT5G04140 model
AT1G13820 data
AT1G13820 model
AT5G66052 data
AT5G66052 model
AT2G23180 data
AT2G23180 model
AT3G29320 data
AT3G29320 model
AT5G65220 data
AT5G65220 model
AT2G27402 data
AT2G27402 model
AT2G36430 data
AT2G36430 model
AT1G01080 data
AT1G01080 model

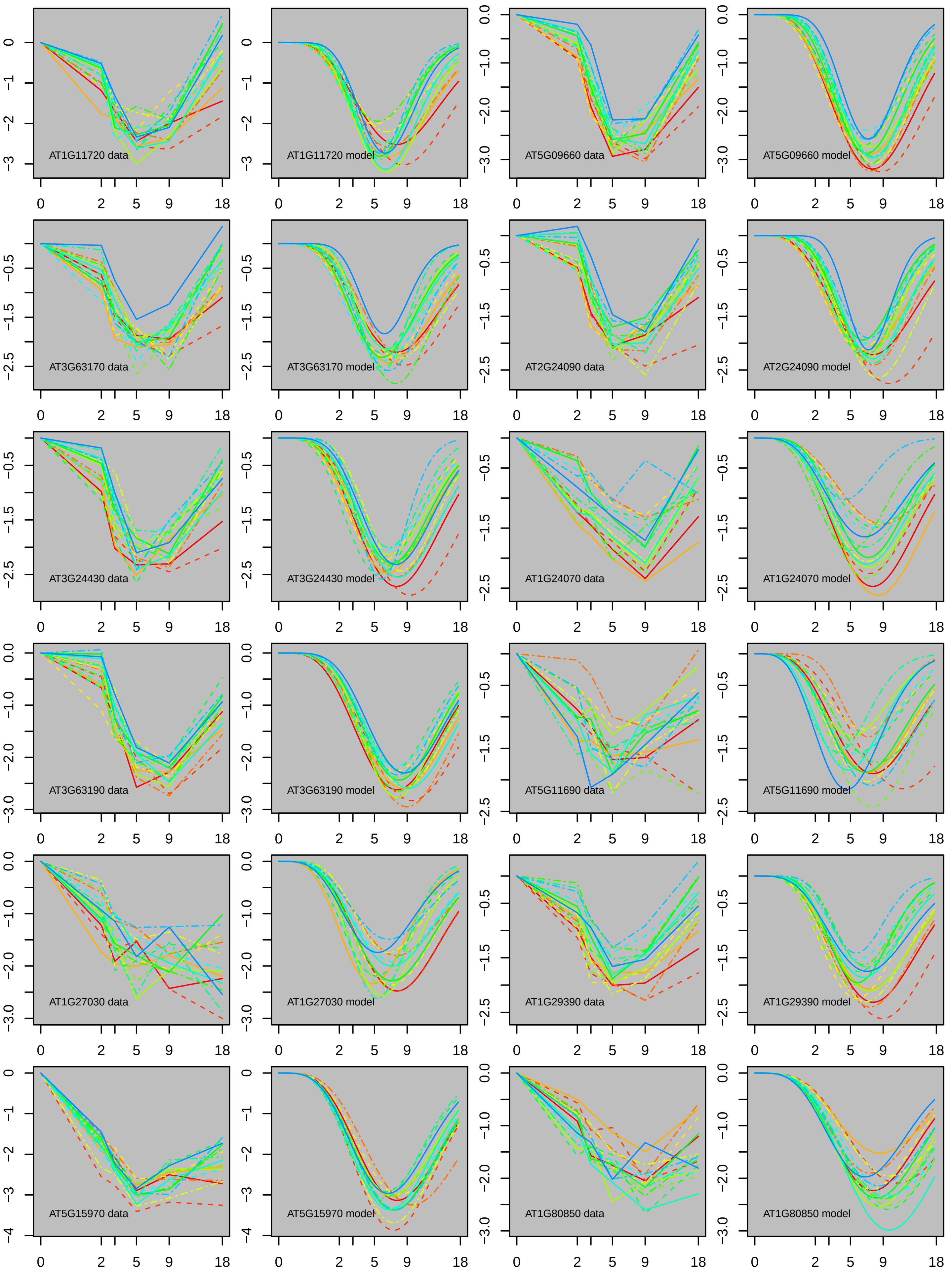

AT1G11720 data
AT1G11720 model
AT5G09660 data
AT5G09660 model
AT3G63170 data
AT3G63170 model
AT2G24090 data
AT2G24090 model
AT3G24430 data
AT3G24430 model
AT1G24070 data
AT1G24070 model
AT3G63190 data
AT3G63190 model
AT5G11690 data
AT5G11690 model
AT1G27030 data
AT1G27030 model
AT1G29390 data
AT1G29390 model
AT5G15970 data
AT5G15970 model
AT1G80850 data
AT1G80850 model

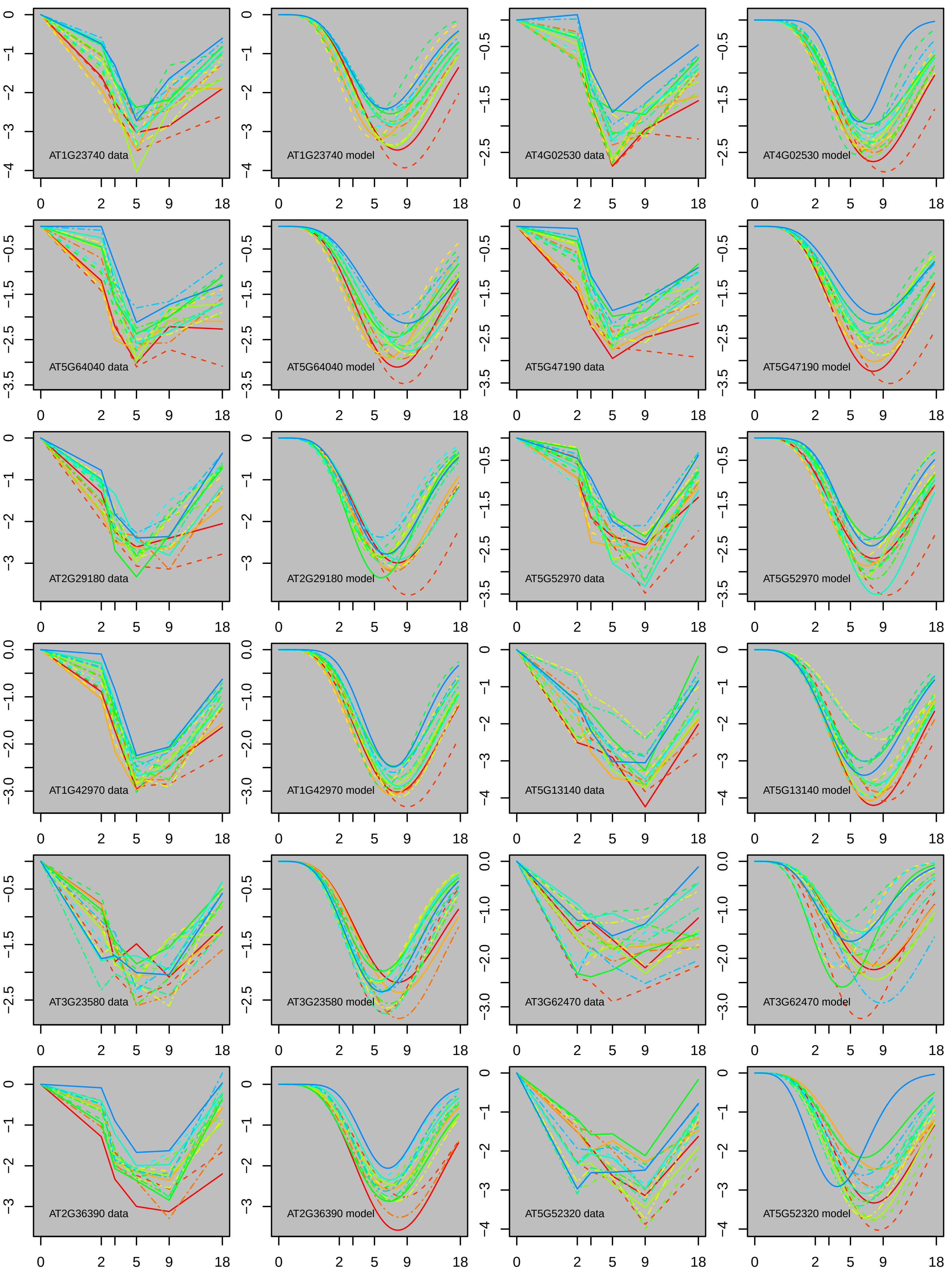
AT1G23740 data
AT1G23740 model
AT4G02530 data
AT4G02530 model
AT5G64040 data
AT5G64040 model
AT5G47190 data
AT5G47190 model
AT2G29180 data
AT2G29180 model
AT5G52970 data
AT5G52970 model
AT1G42970 data
AT1G42970 model
AT5G13140 data
AT5G13140 model
AT3G23580 data
AT3G23580 model
AT3G62470 data
AT3G62470 model
AT2G36390 data
AT2G36390 model
AT5G52320 data
AT5G52320 model

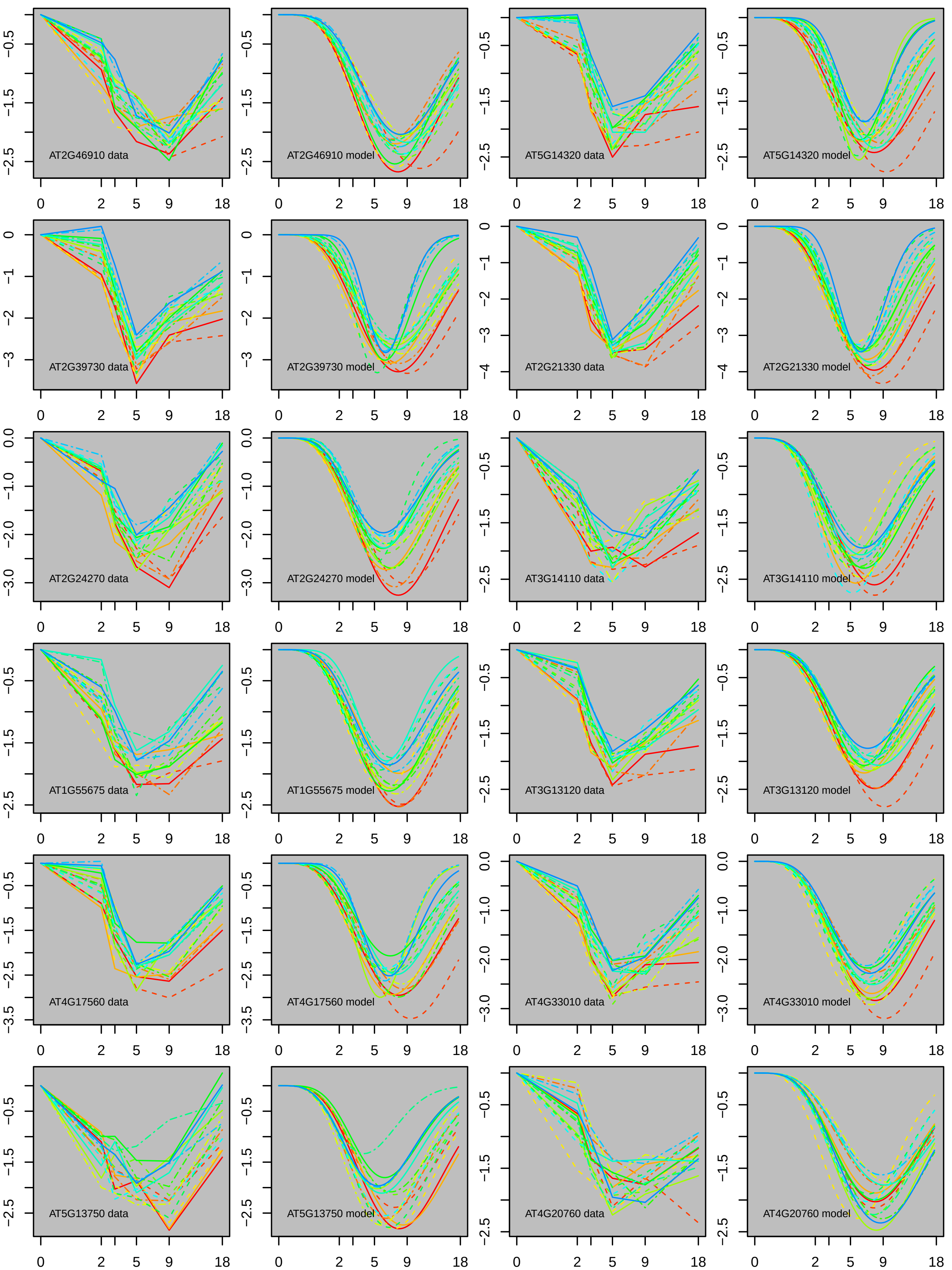
AT2G46910 data
AT2G46910 model
AT5G14320 data
AT5G14320 model
AT2G39730 data
AT2G39730 model
AT2G21330 data
AT2G21330 model
AT2G24270 data
AT2G24270 model
AT3G14110 data
AT3G14110 model
AT1G55675 data
AT1G55675 model
AT3G13120 data
AT3G13120 model
AT4G17560 data
AT4G17560 model
AT4G33010 data
AT4G33010 model
AT5G13750 data
AT5G13750 model
AT4G20760 data
AT4G20760 model
0
2
5
9
18

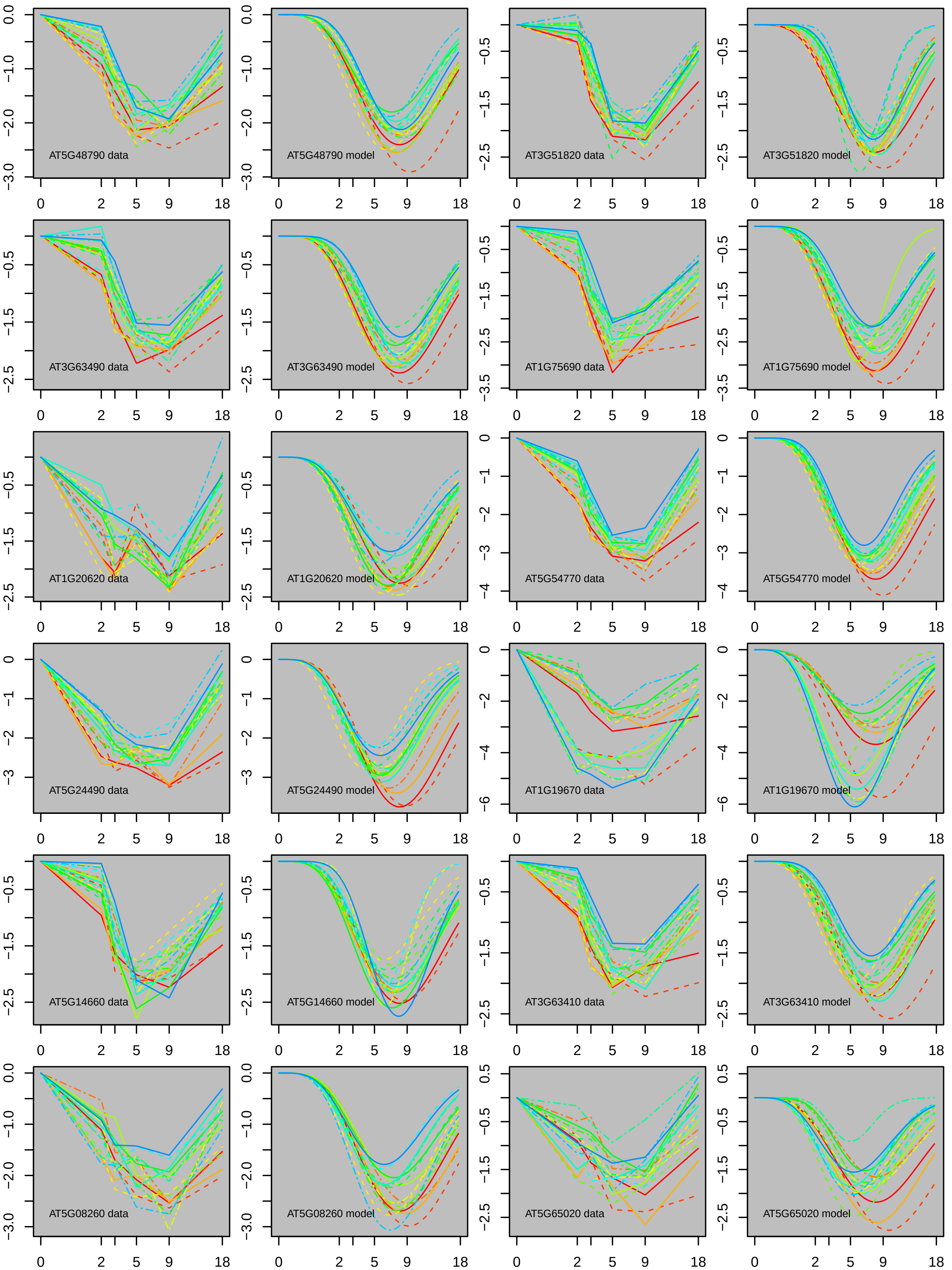
AT5G48790 data
AT5G48790 model
AT3G51820 data
AT3G51820 model
AT3G63490 data
AT3G63490 model
AT1G75690 data
AT1G75690 model
AT1G20620 data
AT1G20620 model
AT5G54770 data
AT5G54770 model
AT5G24490 data
AT5G24490 model
AT1G19670 data
AT1G19670 model
AT5G14660 data
AT5G14660 model
AT3G63410 data
AT3G63410 model
AT5G08260 data
AT5G08260 model
AT5G65020 data
AT5G65020 model

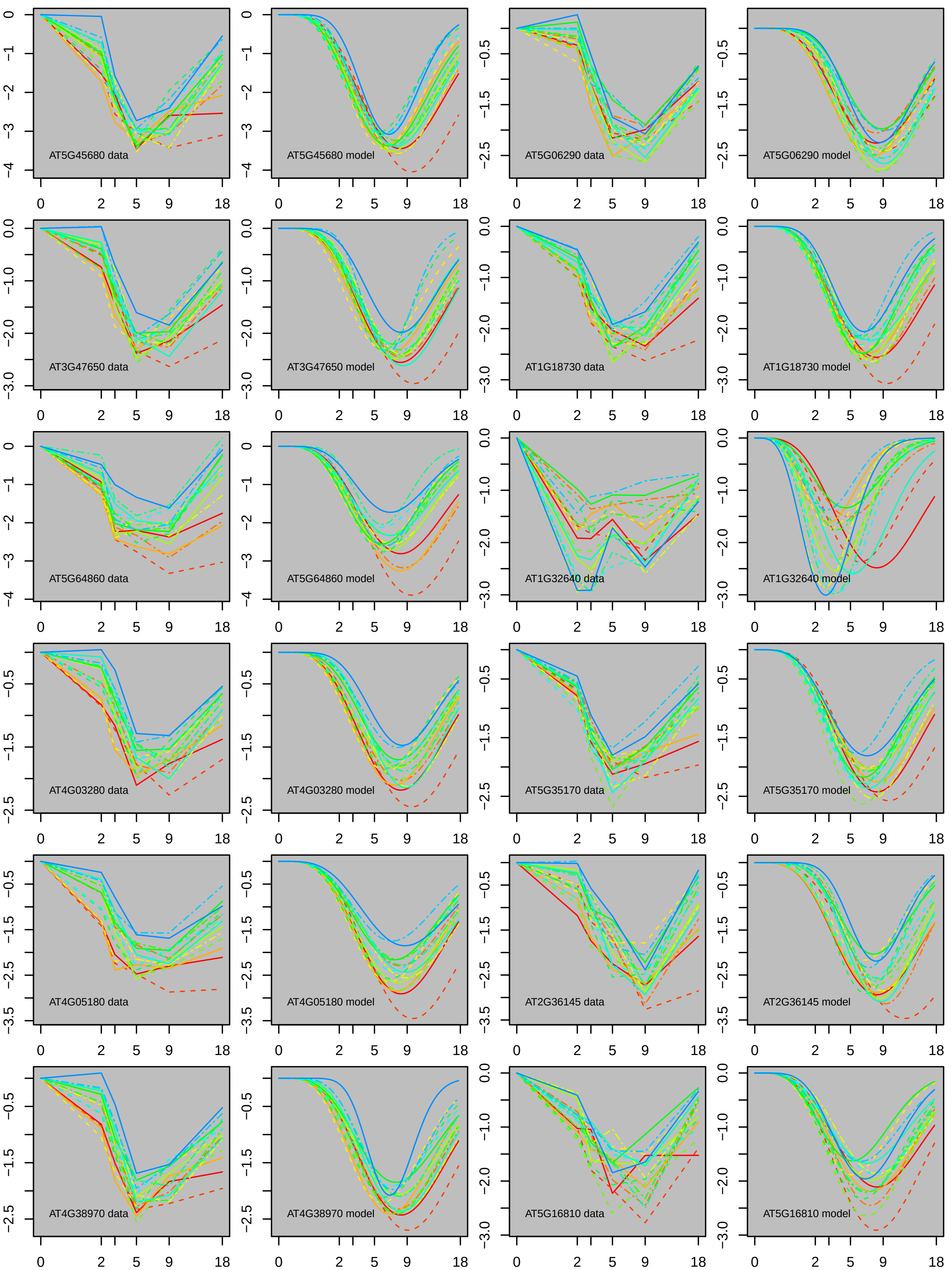
AT5G45680 data
AT5G45680 model
AT5G06290 data
AT5G06290 model
AT3G47650 data
AT3G47650 model
AT1G18730 data
AT1G18730 model
AT5G64860 data
AT5G64860 model
AT1G32640 data
AT1G32640 model
AT4G03280 data
AT4G03280 model
AT5G35170 data
AT5G35170 model
AT4G05180 data
AT4G05180 model
AT2G36145 data
AT2G36145 model
AT4G38970 data
AT4G38970 model
AT5G16810 data
AT5G16810 model

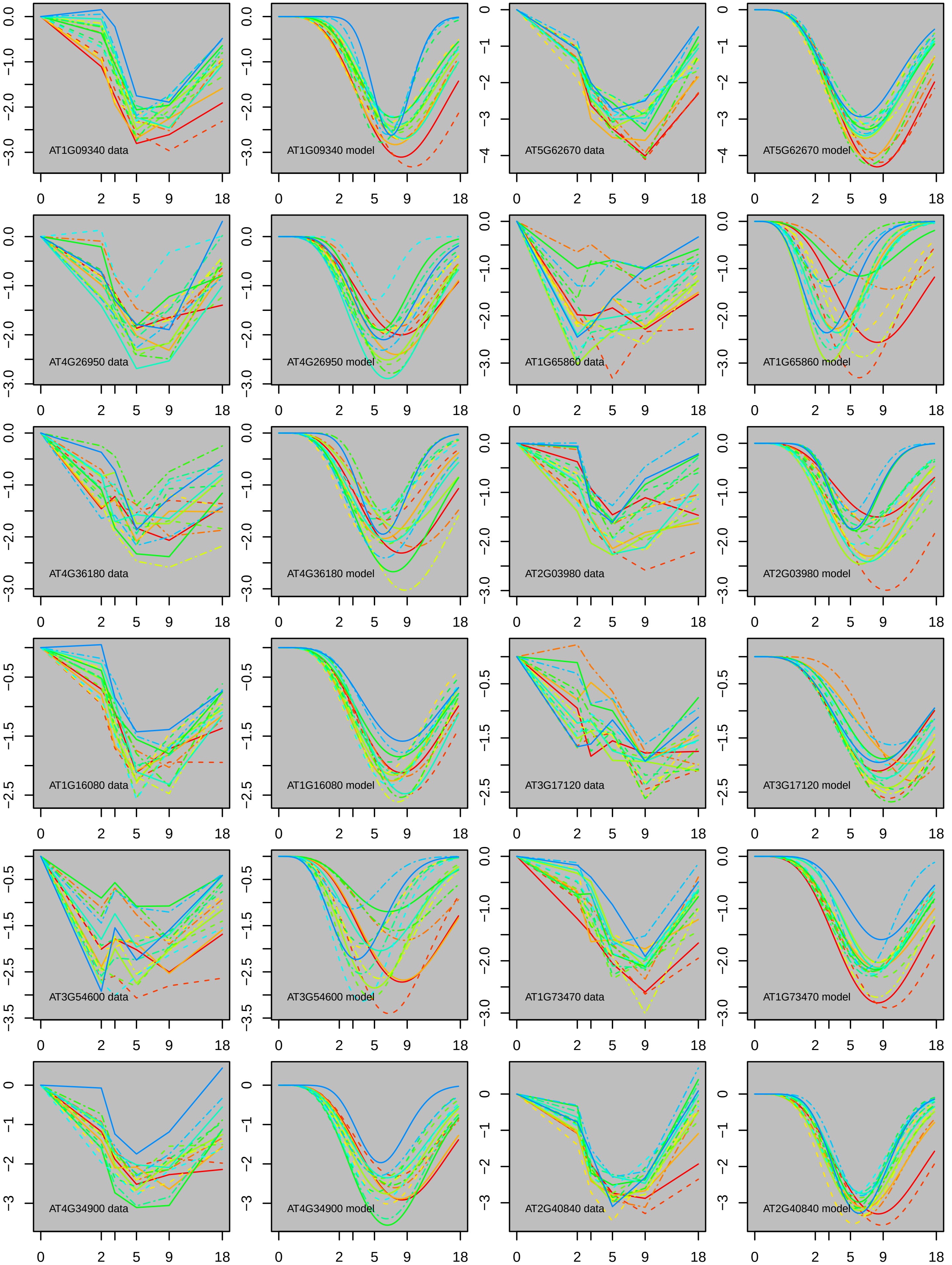
AT1G09340 data
AT1G09340 model
AT5G62670 data
AT5G62670 model
AT4G26950 data
AT4G26950 model
AT1G65860 data
AT1G65860 model
AT4G36180 data
AT4G36180 model
AT2G03980 data
AT2G03980 model
AT1G16080 data
AT1G16080 model
AT3G17120 data
AT3G17120 model
AT3G54600 data
AT3G54600 model
AT1G73470 data
AT1G73470 model
AT4G34900 data
AT4G34900 model
AT2G40840 data
AT2G40840 model
0 2 5 9 18

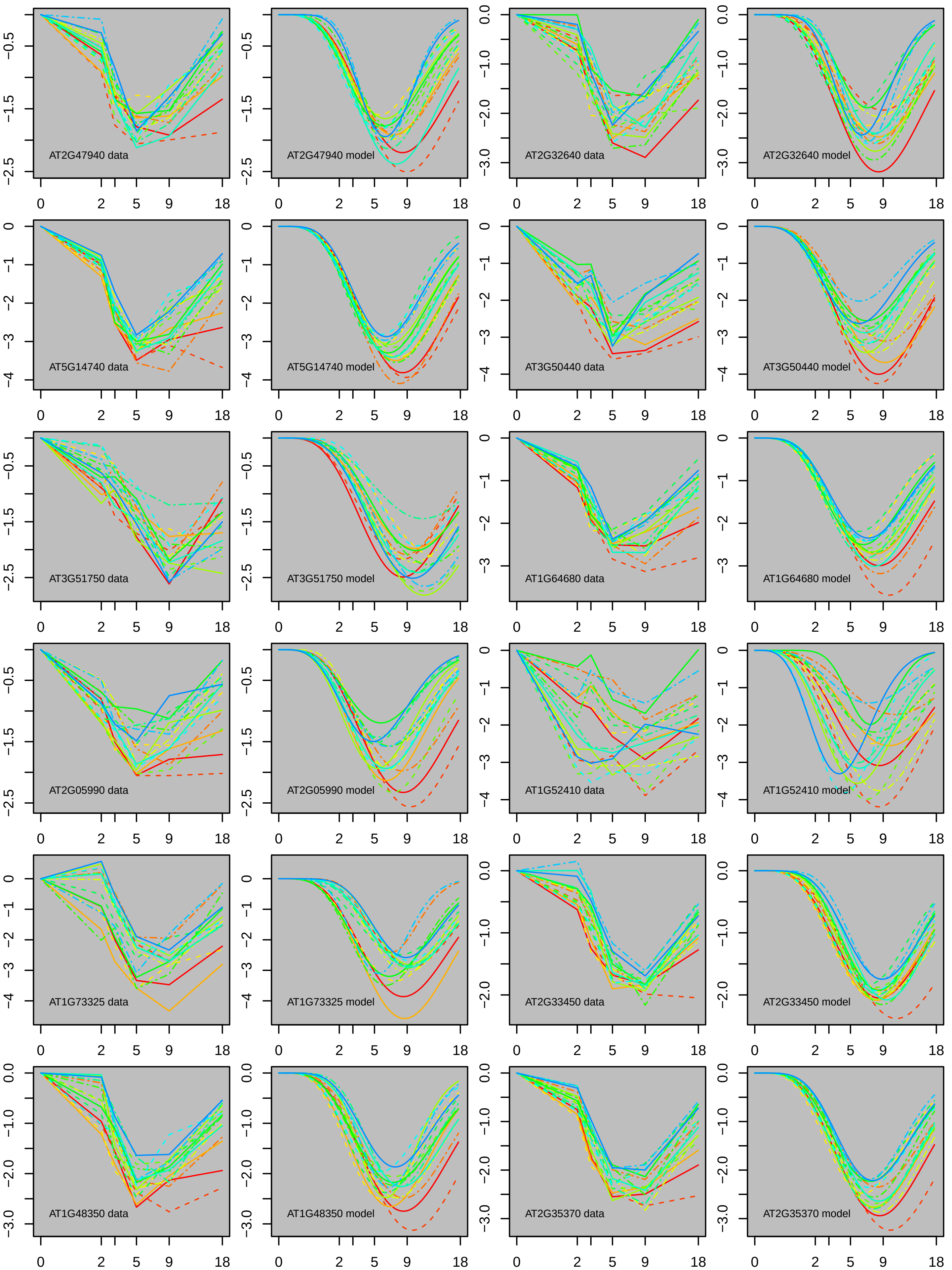
AT2G47940 data
AT2G47940 model
AT2G32640 data
AT2G32640 model
AT5G14740 data
AT5G14740 model
AT3G50440 data
AT3G50440 model
AT3G51750 data
AT3G51750 model
AT1G64680 data
AT1G64680 model
AT2G05990 data
AT2G05990 model
AT1G52410 data
AT1G52410 model
AT1G73325 data
AT1G73325 model
AT2G33450 data
AT2G33450 model
AT1G48350 data
AT1G48350 model
AT2G35370 data
AT2G35370 model

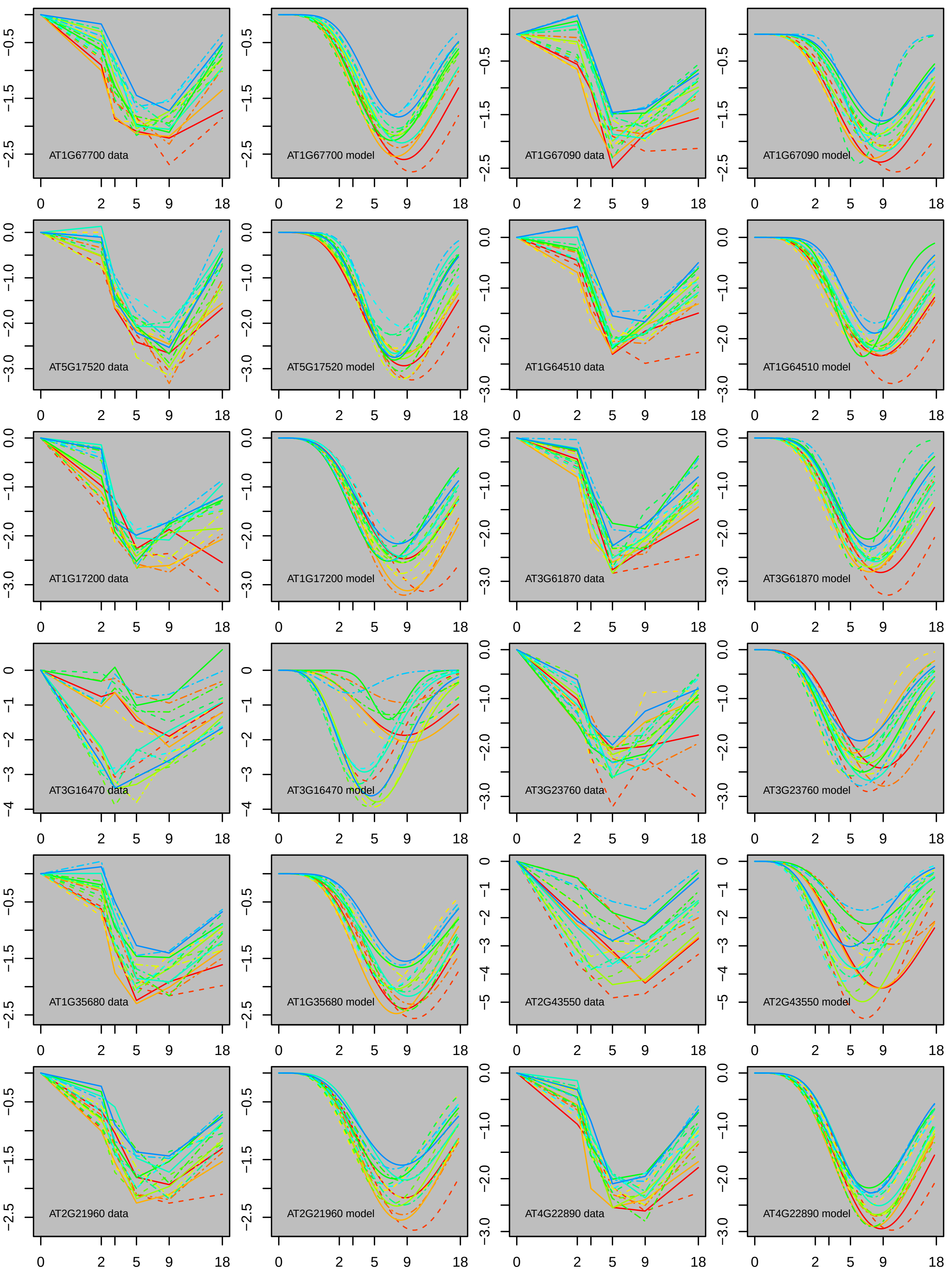
AT1G67700 data
AT1G67700 model
AT1G67090 data
AT1G67090 model
AT5G17520 data
AT5G17520 model
AT1G64510 data
AT1G64510 model
AT1G17200 data
AT1G17200 model
AT3G61870 data
AT3G61870 model
AT3G16470 data
AT3G16470 model
AT3G23760 data
AT3G23760 model
AT1G35680 data
AT1G35680 model
AT2G43550 data
AT2G43550 model
AT2G21960 data
AT2G21960 model
AT4G22890 data
AT4G22890 model

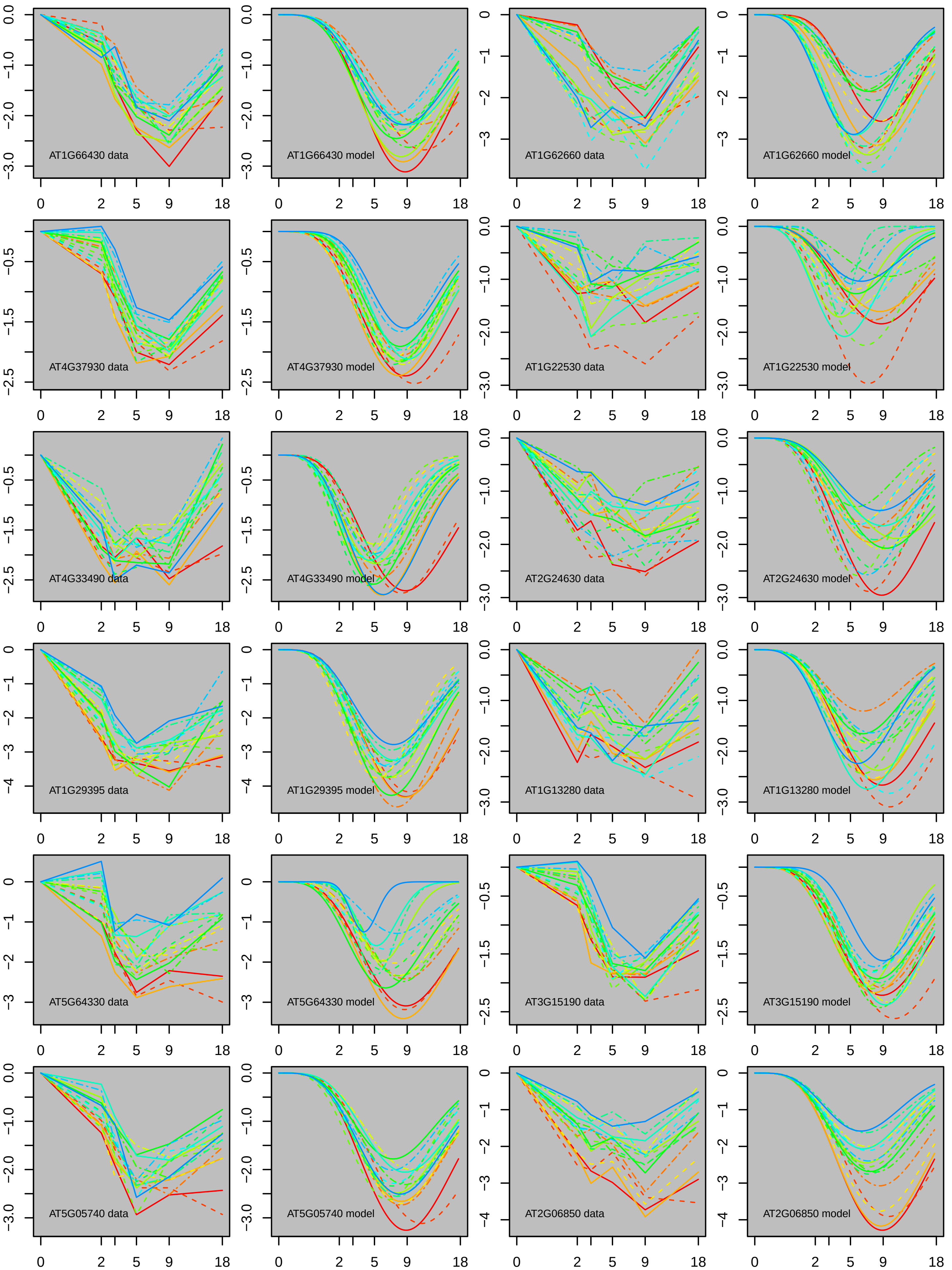
AT1G66430 data
AT1G66430 model
AT1G62660 data
AT1G62660 model
AT4G37930 data
AT4G37930 model
AT1G22530 data
AT1G22530 model
AT4G33490 data
AT4G33490 model
AT2G24630 data
AT2G24630 model
AT1G29395 data
AT1G29395 model
AT1G13280 data
AT1G13280 model
AT5G64330 data
AT5G64330 model
AT3G15190 data
AT3G15190 model
AT5G05740 data
AT5G05740 model
AT2G06850 data
AT2G06850 model

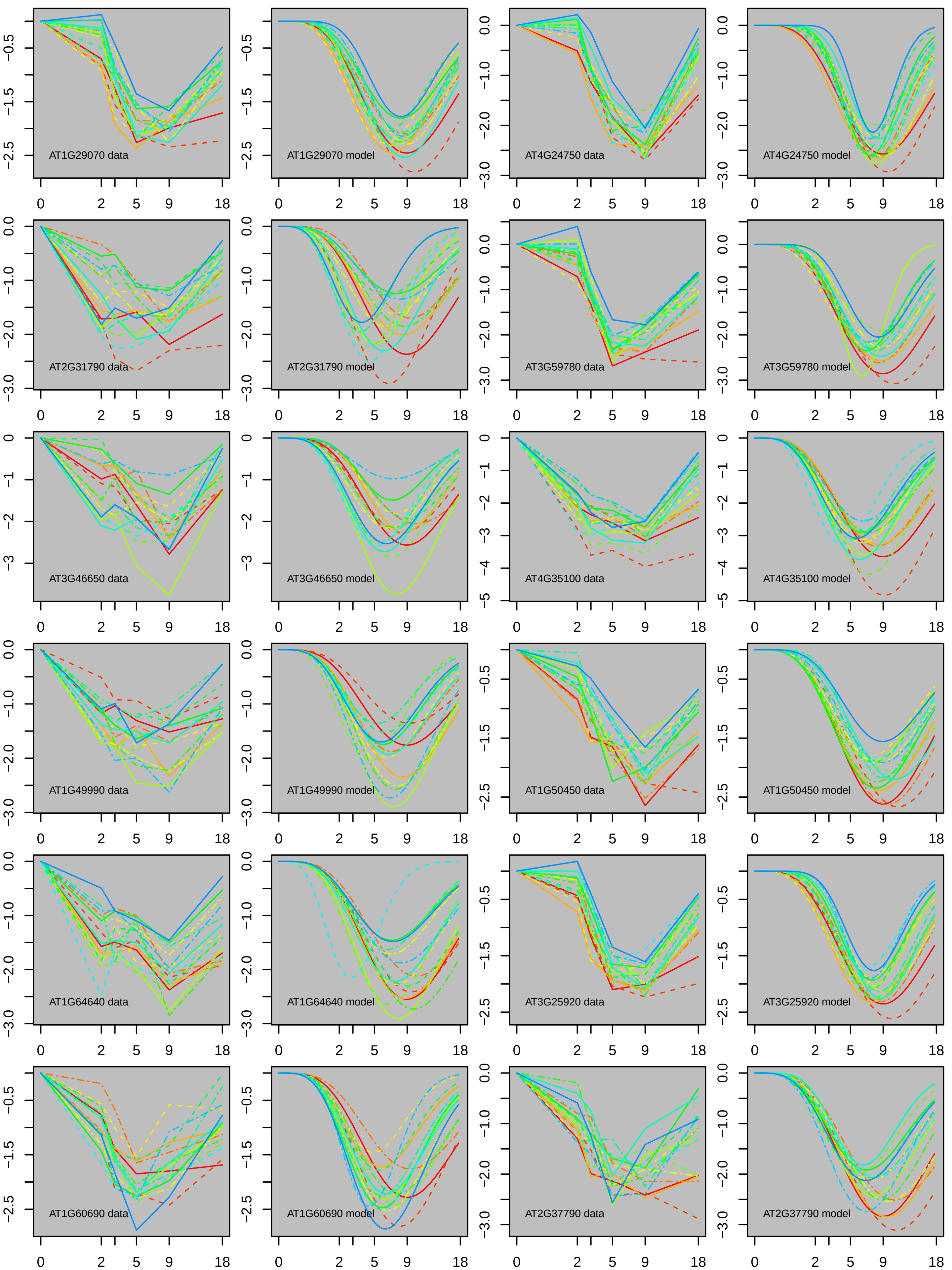
AT1G29070 data
AT1G29070 model
AT4G24750 data
AT4G24750 model
AT2G31790 data
AT2G31790 model
AT3G59780 data
AT3G59780 model
AT3G46650 data
AT3G46650 model
AT4G35100 data
AT4G35100 model
AT1G49990 data
AT1G49990 model
AT1G50450 data
AT1G50450 model
AT1G64640 data
AT1G64640 model
AT3G25920 data
AT3G25920 model
AT1G60690 data
AT1G60690 model
AT2G37790 data
AT2G37790 model
0 2 5 9 18

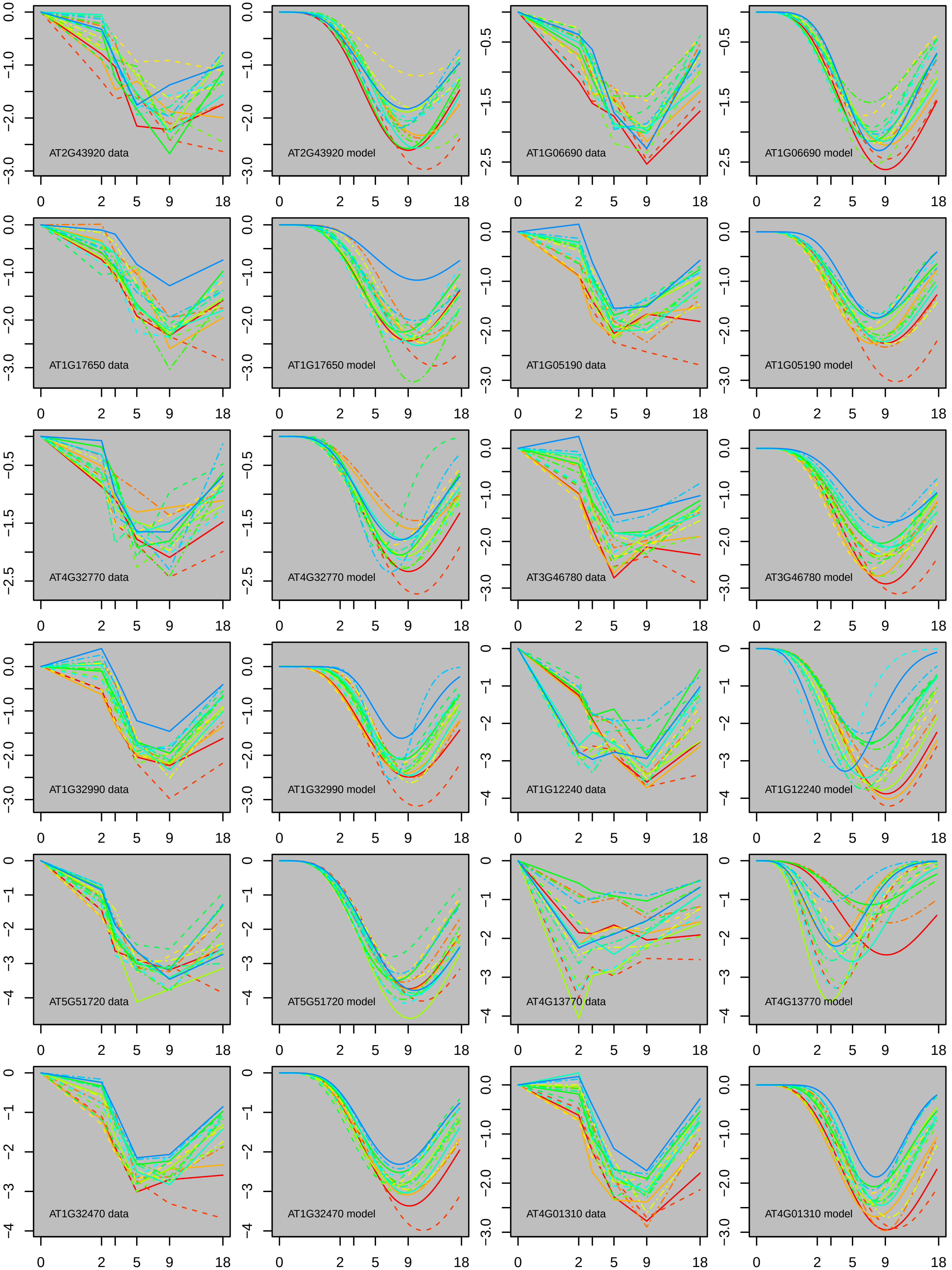
AT2G43920 data
AT2G43920 model
AT1G06690 data
AT1G06690 model
AT1G17650 data
AT1G17650 model
AT1G05190 data
AT1G05190 model
AT4G32770 data
AT4G32770 model
AT3G46780 data
AT3G46780 model
AT1G32990 data
AT1G32990 model
AT1G12240 data
AT1G12240 model
AT5G51720 data
AT5G51720 model
AT4G13770 data
AT4G13770 model
AT1G32470 data
AT1G32470 model
AT4G01310 data
AT4G01310 model

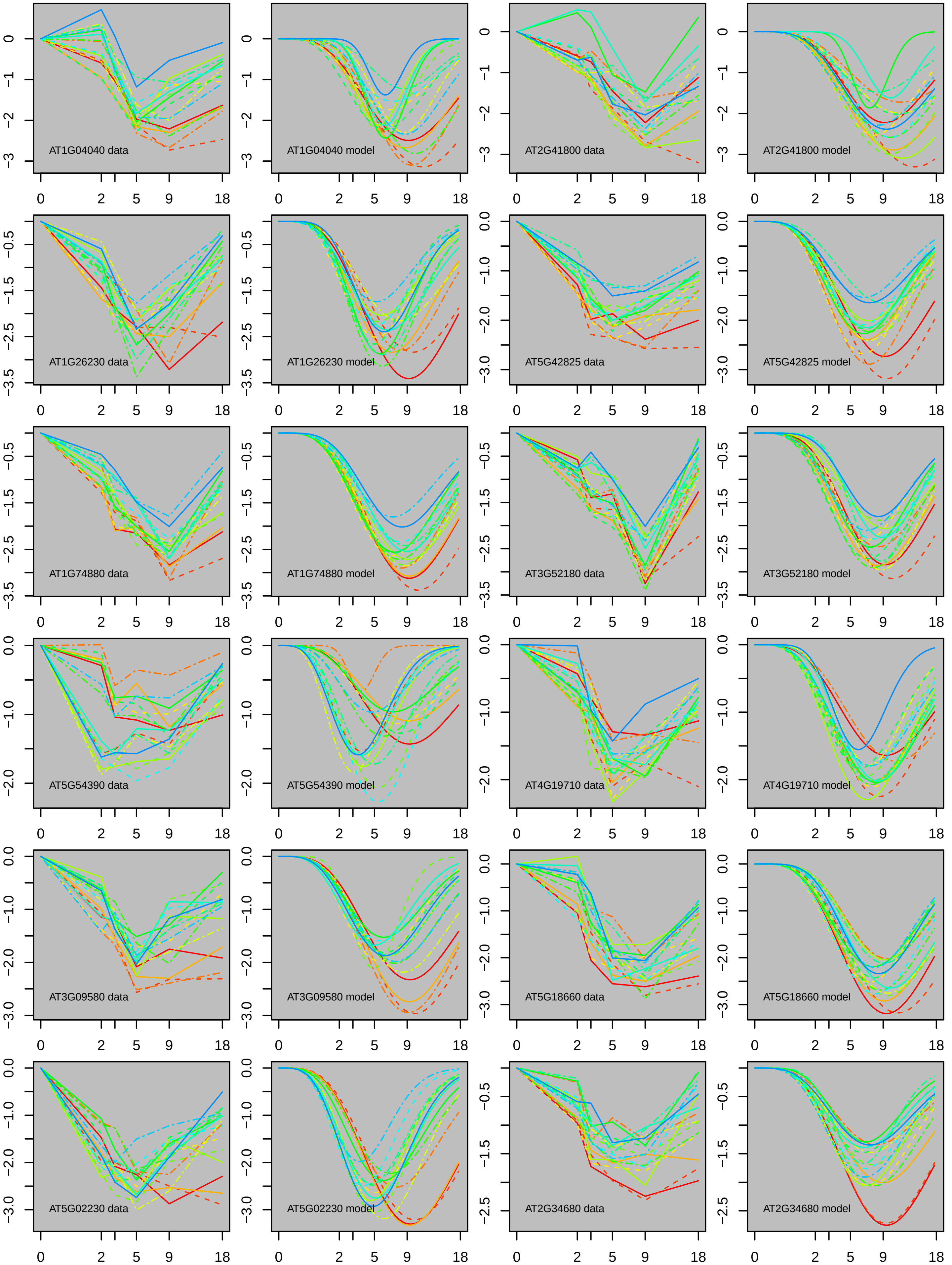
AT1G04040 data
AT1G04040 model
AT2G41800 data
AT2G41800 model
AT1G26230 data
AT1G26230 model
AT5G42825 data
AT5G42825 model
AT1G74880 data
AT1G74880 model
AT3G52180 data
AT3G52180 model
AT5G54390 data
AT5G54390 model
AT4G19710 data
AT4G19710 model
AT3G09580 data
AT3G09580 model
AT5G18660 data
AT5G18660 model
AT5G02230 data
AT5G02230 model
AT2G34680 data
AT2G34680 model

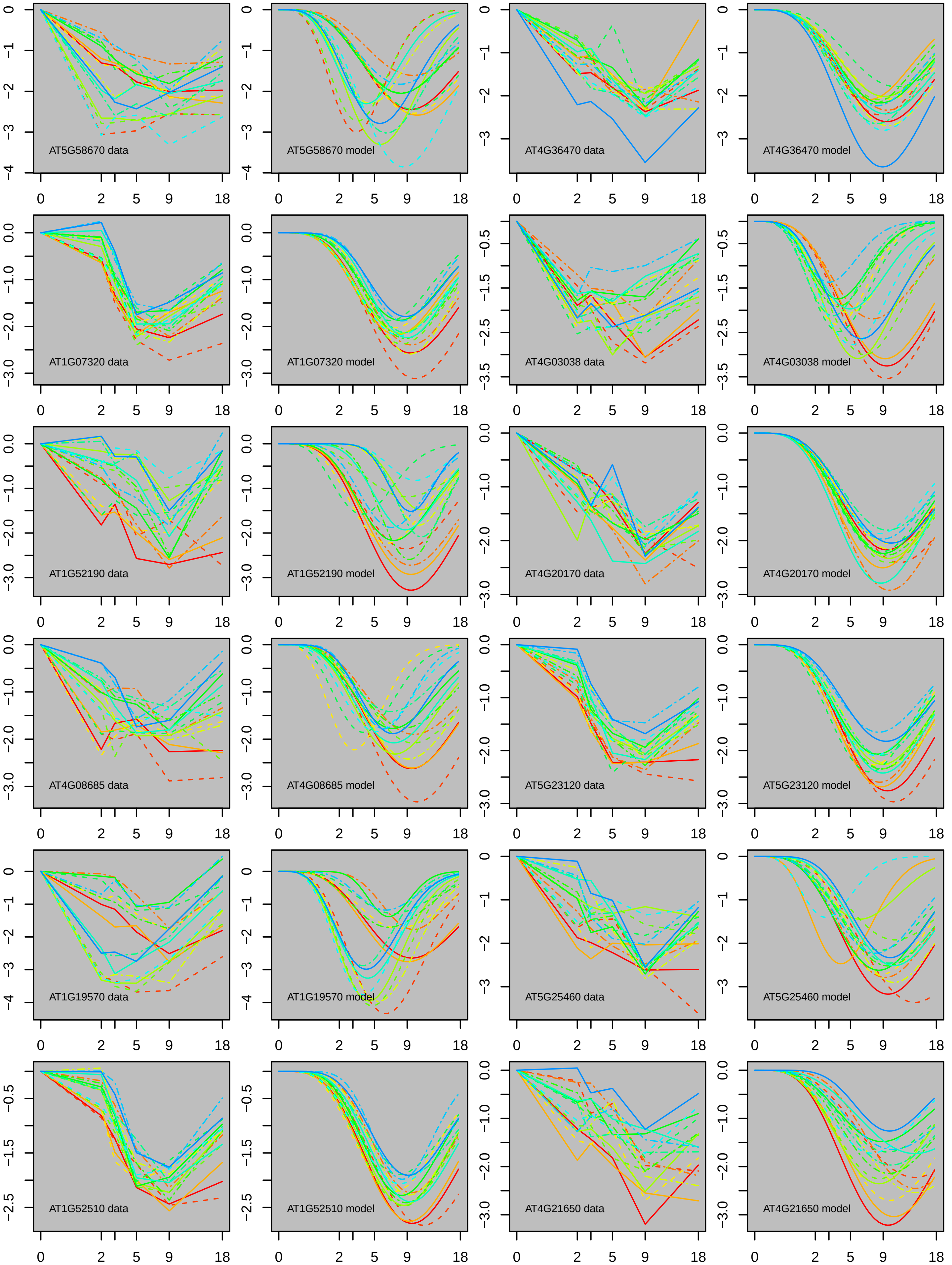
AT5G58670 data
AT5G58670 model
AT4G36470 data
AT4G36470 model
AT1G07320 data
AT1G07320 model
AT4G03038 data
AT4G03038 model
AT1G52190 data
AT1G52190 model
AT4G20170 data
AT4G20170 model
AT4G08685 data
AT4G08685 model
AT5G23120 data
AT5G23120 model
AT1G19570 data
AT1G19570 model
AT5G25460 data
AT5G25460 model
AT1G52510 data
AT1G52510 model
AT4G21650 data
AT4G21650 model
0
2
5
9
18

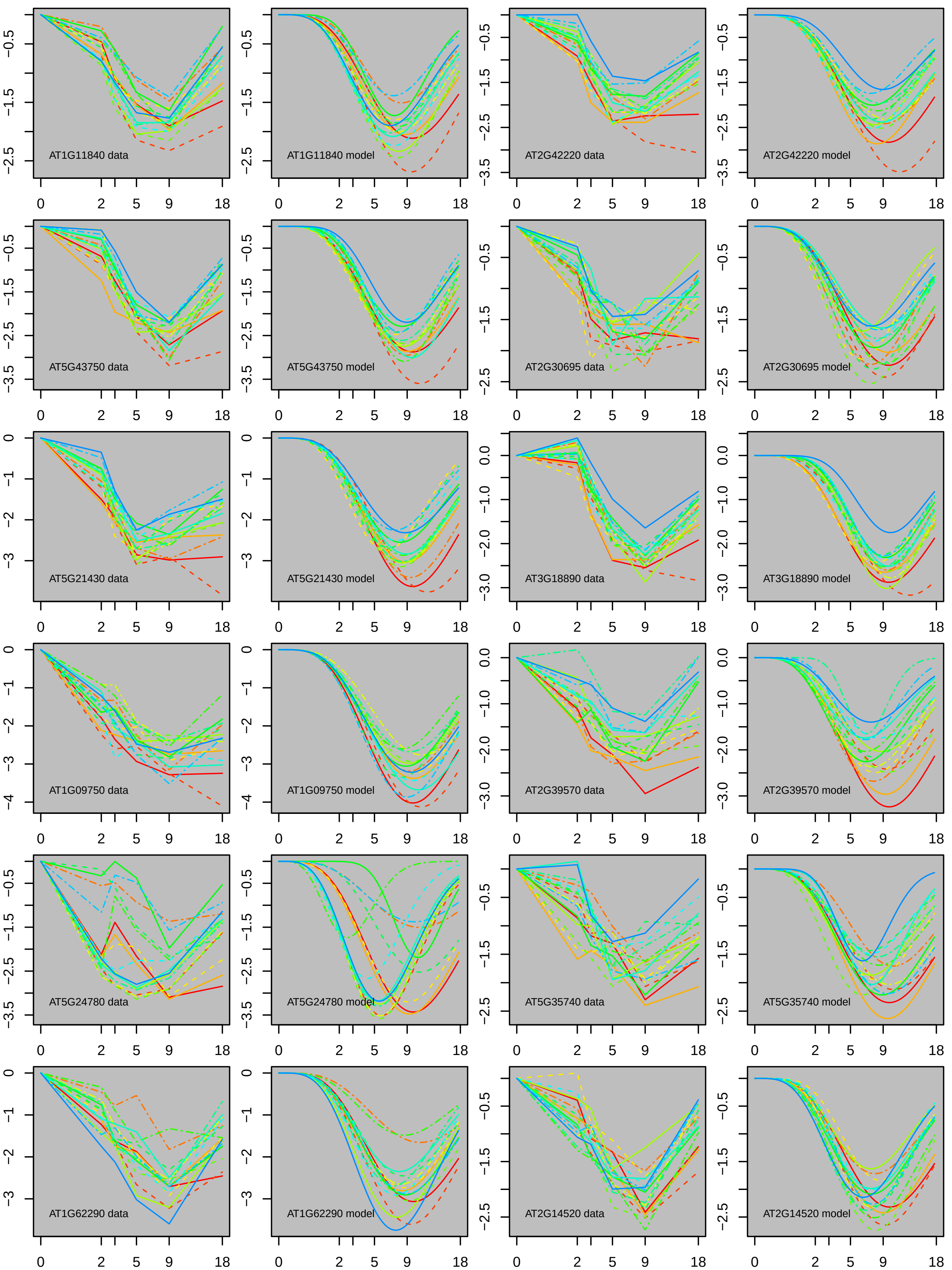

AT1G11840 data
AT1G11840 model
AT2G42220 data
AT2G42220 model
AT5G43750 data
AT5G43750 model
AT2G30695 data
AT2G30695 model
AT5G21430 data
AT5G21430 model
AT3G18890 data
AT3G18890 model
AT1G09750 data
AT1G09750 model
AT2G39570 data
AT2G39570 model
AT5G24780 data
AT5G24780 model
AT5G35740 data
AT5G35740 model
AT1G62290 data
AT1G62290 model
AT2G14520 data
AT2G14520 model

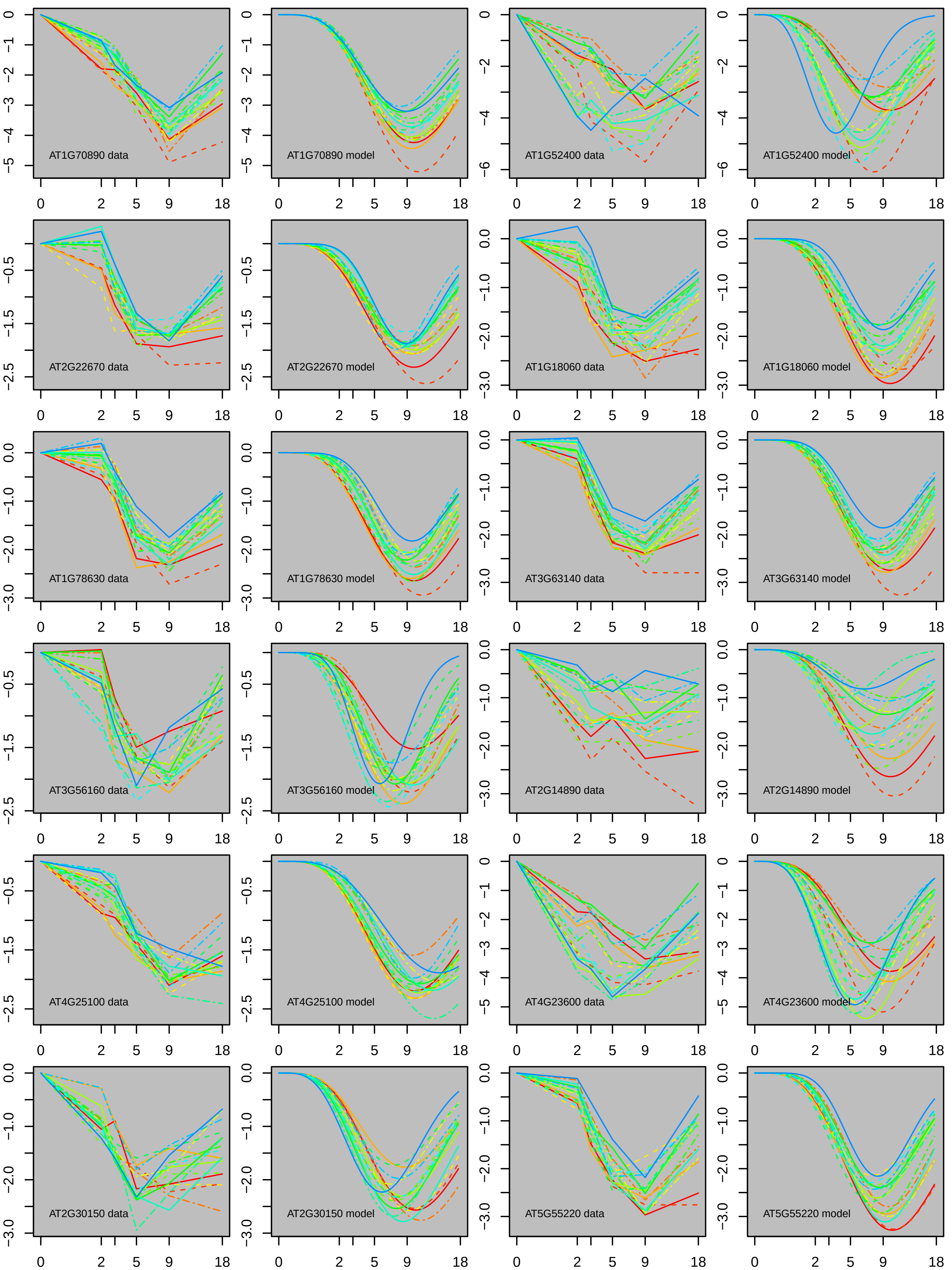
AT1G70890 data
AT1G70890 model
AT1G52400 data
AT1G52400 model
AT2G22670 data
AT2G22670 model
AT1G18060 data
AT1G18060 model
AT1G78630 data
AT1G78630 model
AT3G63140 data
AT3G63140 model
AT3G56160 data
AT3G56160 model
AT2G14890 data
AT2G14890 model
AT4G25100 data
AT4G25100 model
AT4G23600 data
AT4G23600 model
AT2G30150 data
AT2G30150 model
AT5G55220 data
AT5G55220 model

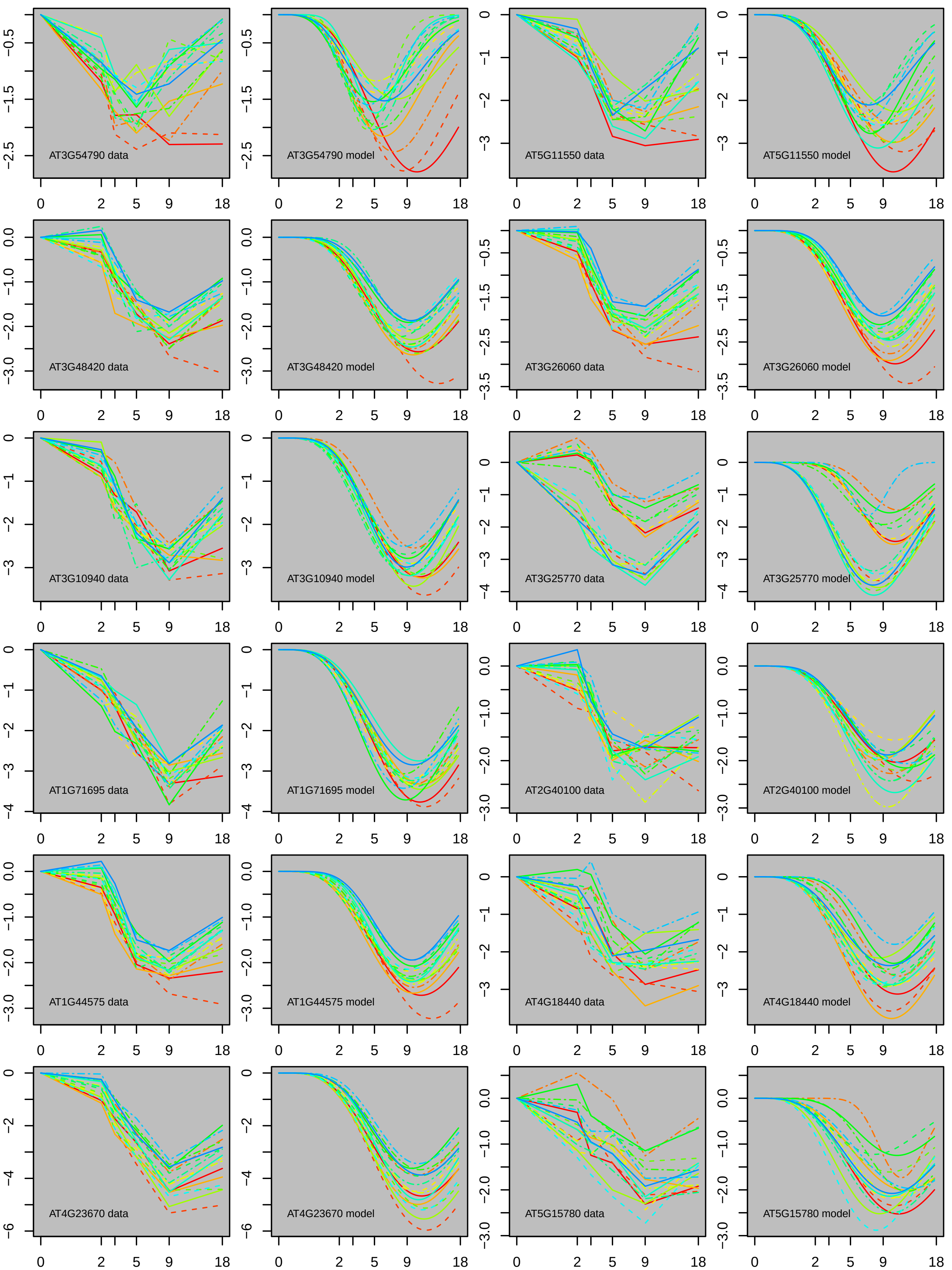
AT3G54790 data
AT3G54790 model
AT5G11550 data
AT5G11550 model
AT3G48420 data
AT3G48420 model
AT3G26060 data
AT3G26060 model
AT3G10940 data
AT3G10940 model
AT3G25770 data
AT3G25770 model
AT1G71695 data
AT1G71695 model
AT2G40100 data
AT2G40100 model
AT1G44575 data
AT1G44575 model
AT4G18440 data
AT4G18440 model
AT4G23670 data
AT4G23670 model
AT5G15780 data
AT5G15780 model

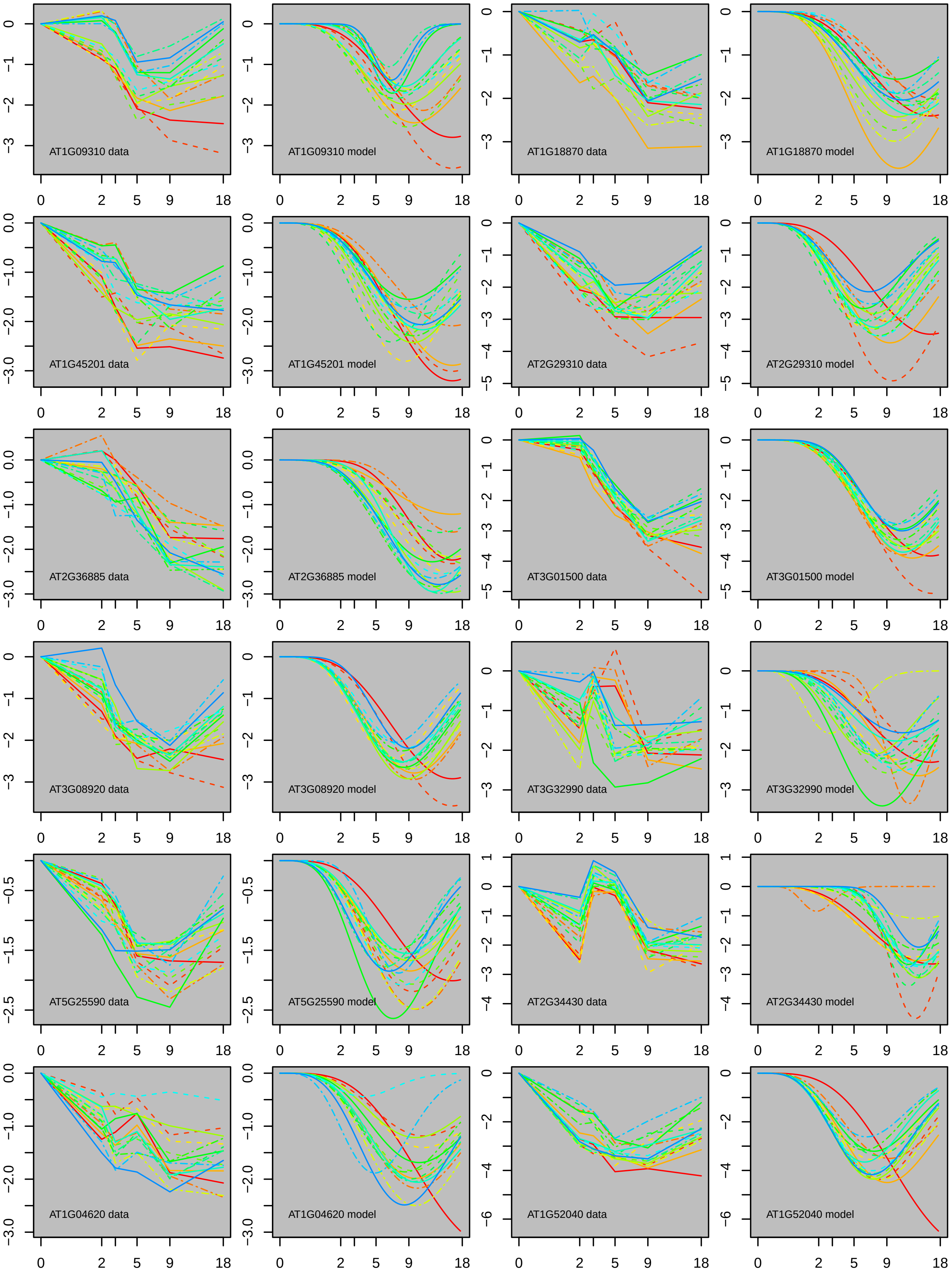
AT1G09310 data
AT1G09310 model
AT1G18870 data
AT1G18870 model
AT1G45201 data
AT1G45201 model
AT2G29310 data
AT2G29310 model
AT2G36885 data
AT2G36885 model
AT3G01500 data
AT3G01500 model
AT3G08920 data
AT3G08920 model
AT3G32990 data
AT3G32990 model
AT5G25590 data
AT5G25590 model
AT2G34430 data
AT2G34430 model
AT1G04620 data
AT1G04620 model
AT1G52040 data
AT1G52040 model

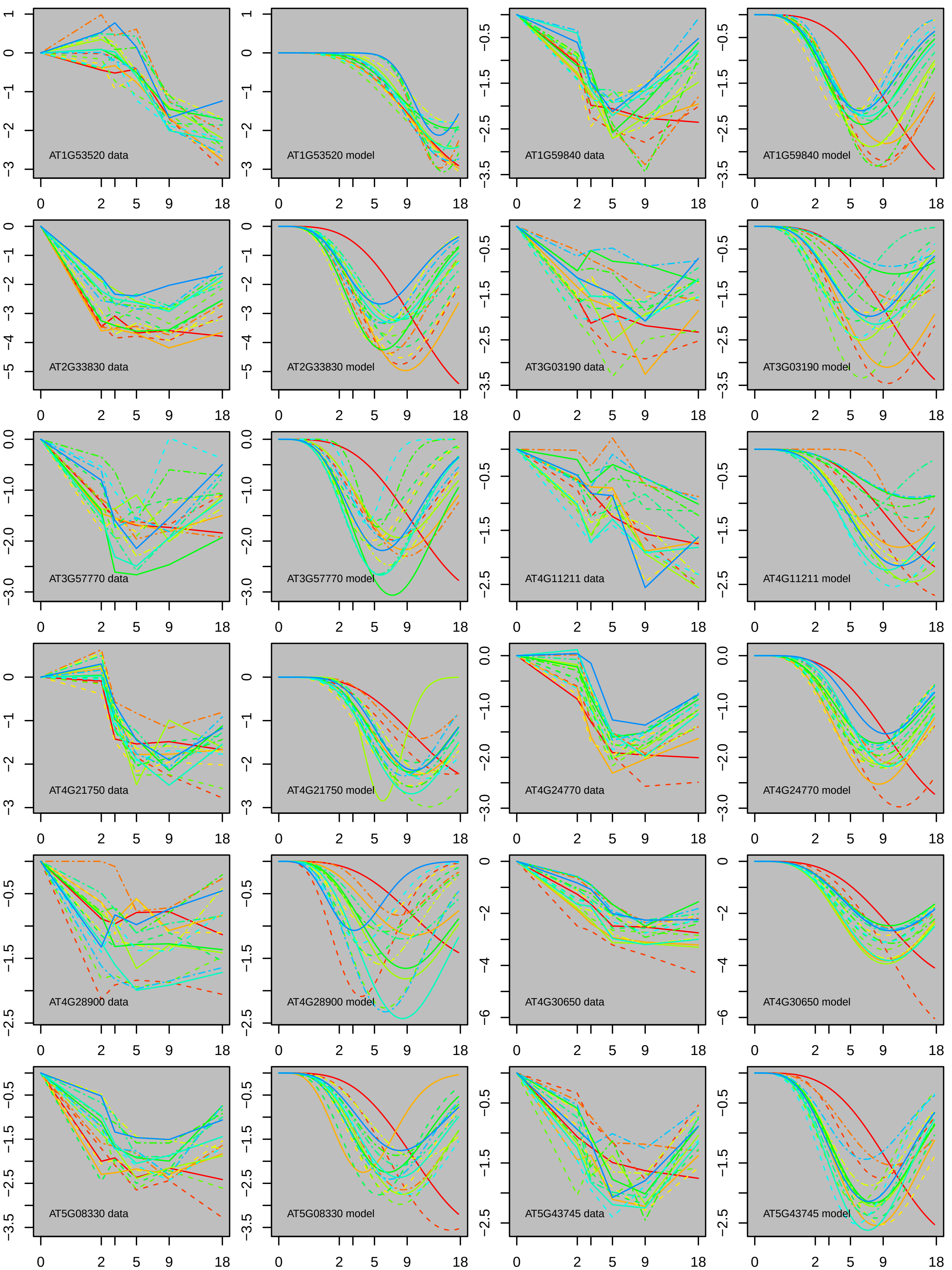
AT1G53520 data
AT1G53520 model
AT1G59840 data
AT1G59840 model
AT2G33830 data
AT2G33830 model
AT3G03190 data
AT3G03190 model
AT3G57770 data
AT3G57770 model
AT4G11211 data
AT4G11211 model
AT4G21750 data
AT4G21750 model
AT4G24770 data
AT4G24770 model
AT4G28900 data
AT4G28900 model
AT4G30650 data
AT4G30650 model
AT5G08330 data
AT5G08330 model
AT5G43745 data
AT5G43745 model
